# Footprint-seq: a simple method to quantitatively map *in vitro* protein-DNA interactions on a genome-wide scale at high spatial resolution

**DOI:** 10.64898/2026.01.16.700004

**Authors:** Archer J. Wang, Anne M. Stringer, Matthew Adams, Patricia M. Aquino, Bradley N. Jenner, James E. Galagan, David C. Grainger, Joseph T. Wade

## Abstract

Protein–DNA interactions underlie essential cellular processes. The position and/or strength of protein interactions with DNA have historically been studied for short DNA regions using *in vitro* assays. More recently, microarray- and sequencing-based assays have been developed to map protein-DNA interactions on a genome-wide scale. Here, we introduce Footprint-seq, a simple, sequencing-based method that maps and quantifies protein–DNA interactions *in vitro* with high resolution on a genome-wide scale. Footprint-seq measures local protection from transposition in the presence of a DNA-binding protein. Using the well-studied *Escherichia coli* DNA-binding protein, CRP, as a test case, we show that Footprint-seq detects CRP binding sites with high resolution across plasmid or genomic DNA. Moreover, we show that Footprint-seq enables inference of dissociation constants for hundreds of genomic sites. Comparison to an alternative method shows that Footprint-seq has both higher sensitivity and higher spatial resolution. Thus, Footprint-seq provides a rapid, accessible, and quantitative method for mapping protein–DNA interactions in bacteria and other systems.

## INTRODUCTION

Protein-DNA interactions play critical roles in many cellular processes, including replication, transcription, and DNA repair. There are numerous methods for measuring the affinity of protein-DNA interactions and/or their position on DNA, both *in vivo* and *in vitro*. Most *in vitro* methods are targeted to individual protein-DNA interactions of interest. For example, the Electromobility Shift Assay (EMSA) can be used to measure the affinity of an interaction between a protein and a specific DNA sequence (1). The assay relies upon the altered mobility of DNA fragments on a non-denaturing gel when bound by a protein. DNase I footprinting assays can be used to measure the position of an interaction between a protein and a specific DNA sequence (2). The assay relies upon DNase I cleavage throughout the DNA fragment of interest, but not in regions that are protected by a bound protein. For these and other targeted *in vitro* assays, DNA regions of interest are typically limited in size, often <100 bp.

The advent of microarrays and next-generation DNA sequencing led to the development of genome-scale methods for interrogating protein-DNA interactions *in vitro*. Most of these methods rely on the capture of sheared genomic DNA by the protein of interest. For example, in DIP-chip (DNA immunoprecipitation with microarray detection) (3), DAP-seq (DNA affinity purification sequencing), IDAP-seq (*in vitro* DNA affinity purification sequencing), *in vitro* ChIP-seq (Chromatin immunoprecipitation followed by sequencing) (4), Affinity-seq (5), genomic SELEX (Systematic evolution of ligands by exponential enrichment) (6), and GHT-SELEX (genomic high-throughput SELEX) (7), genomic DNA fragments are bound by a protein of interest, which is then immunoprecipitated or affinity purified, enriching for protein-bound DNA regions that are then quantified using microarrays or DNA sequencing. Nitro-seq uses a similar approach, but isolates protein-DNA complexes by filtration with nitrocellulose (8). Because these approaches all use sheared genomic DNA, the binding site location is inferred from the position with the highest overall enrichment across overlapping DNA fragments, somewhat limiting the spatial resolution for weaker sites. An alternative approach for mapping protein-DNA interactions *in vitro* is the use of PBMs (protein-binding microarrays), in which a protein is bound to DNA on a microarray, with protein occupancy quantified by the addition of a fluorescently tagged antibody specific to the DNA-binding protein (9). PBMs have been used predominantly to measure protein binding to artificial DNA sequences, but the microarrays can be designed to use genomic DNA sequences (10).

Here, we develop a new method, Footprint-seq, for interrogating protein-DNA interactions *in vitro*. Footprint-seq is a simple, rapid method that identifies the position and affinity of protein interactions with DNA. Footprint-seq can be performed with DNA of any size, and is ideally suited to bacterial genomes. Using *Escherichia coli* CRP (cyclic-AMP receptor protein), a well-characterized bacterial transcription factor, we show that Footprint-seq has high spatial resolution, is highly sensitive, and can be used to infer *K*_d_ values for individual sites on a genome-wide scale.

## RESULTS

### Footprint-seq: a sequencing-based method to map and quantify protein-DNA interactions in vitro with high spatial resolution

We sought to develop a method to measure protein-DNA interactions *in vitro* on a genomic scale and with high spatial resolution. We leveraged the “Assay for Transposase-Accessible Chromatin using sequencing” (ATAC-seq) method that is widely used to map chromatin accessibility in eukaryotes (11). ATAC-seq relies on “tagmentation”, wherein a hyperactive Tn5 transposase is used to introduce short sequence tags into DNA (12). The positions of Tn5 integration are determined and quantified by PCR coupled with deep sequencing. Nucleosome-associated regions have reduced levels of transposition due to their lower accessibility. Thus, ATAC-seq can map nucleosome density across whole genomes. ATAC-seq captures the binding of all nuclear proteins to DNA. We reasoned that a similar approach could be used to map the association of specific proteins with DNA *in vitro*, a method we call “Footprint-seq” (Figure 1) due to its similarity to the widely used DNA footprinting approaches that map the association of DNA-binding proteins with short DNA regions.

**Figure 1.**
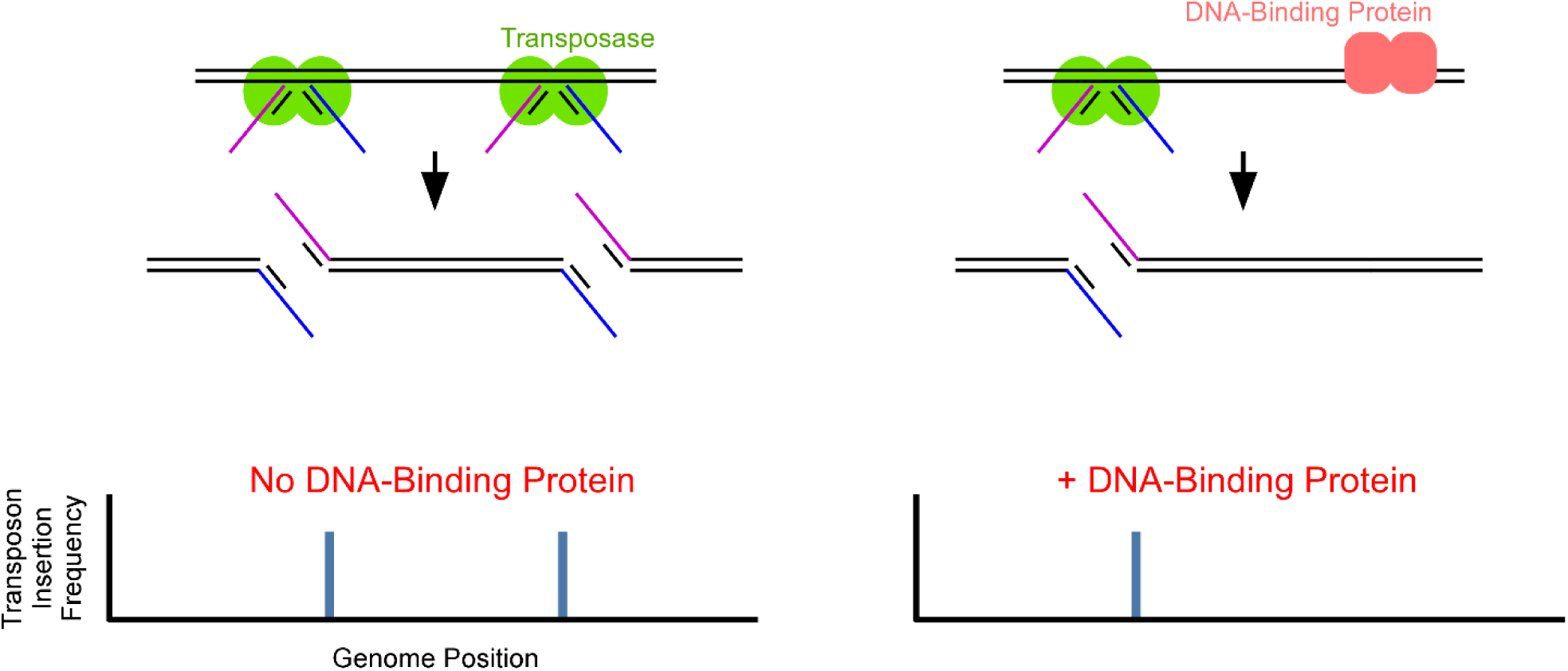
Schematic of the Footprint-seq method. DNA is incubated with transposon-bound transposase in a tagmentation reaction. PCR-amplification and sequencing of the tagmented DNA identifies sites of transposon insertion. Addition of a DNA-binding protein can reduce the transposon insertion frequency at positions close to the binding site.

### Footprint-seq identifies binding sites for E. coli CRP on a plasmid

To establish Footprint-seq, we selected *E. coli* CRP, since its DNA sequence preference is well-characterized, and many high-affinity CRP sites in the *E. coli* genome have been previously described. We first determined whether tagmentation occurs under conditions commonly used to study CRP-DNA interactions *in vitro*. We used the Nextera Tn5 tagmentation system that is widely used to generate Illumina sequencing libraries from genomic DNA (13). We targeted a plasmid DNA template that includes a promoter region with two strong CRP sites (14). Tagmentation was less efficient in buffer conditions optimized for CRP-DNA interactions than buffer conditions optimized for Tn5 transposon activity; nonetheless, we detected sufficient tagmentation by PCR with an extended incubation of DNA with the transposase-transposon complex. We next incubated plasmid DNA with a range of CRP concentrations prior to tagmentation. We sequenced the corresponding PCR products and quantified transposition at positions across the plasmid for all samples. Transposition frequencies across the plasmid were highly correlated between replicates (Figure S1); hence, we combined replicate datasets for further analysis. For most positions across the plasmid, we observed essentially identical patterns of transposition regardless of the CRP concentration used. However, in the presence of CRP, we observed reduced transposition at each of the two CRP binding sites, i.e., a transposon footprint (Figure 2A). Moreover, the extent of footprinting correlated with the CRP concentration (Figure 2A). The size of the footprint around each of the CRP sites was ∼10 bp larger than that observed previously for DNase I footprinting (14). At higher concentrations of CRP, we observed footprinting of a third site (Figure 2B). Visual inspection of the plasmid sequence at this third site revealed a close match to the CRP consensus site.

**Figure 2.**
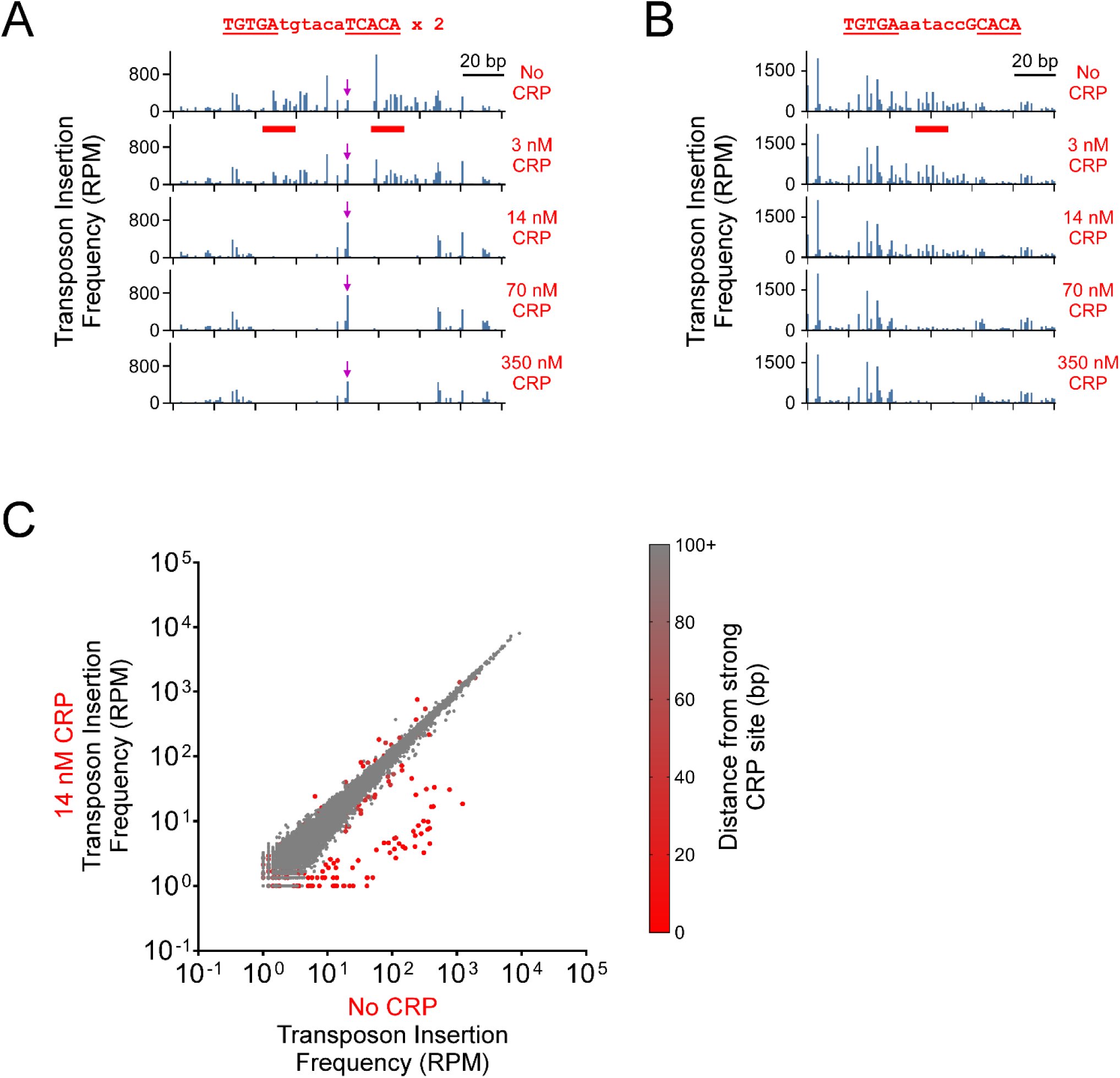
Benchmarking Footprint-seq with CRP using a plasmid. **(A)** Transposon insertion frequencies for a 160 bp region of a plasmid, centered on two high-affinity CRP sites (red horizontal bars). The DNA sequence of the two identical sites is indicated above the graphs. Data are shown for DNA incubated with 0, 3, 14, 70, or 350 nM CRP. The red arrow indicates a hypersensitive site that has increased transposition with increasing concentrations of CRP. **(B)** Transposon insertion frequencies for a 120 bp region of the same plasmid, highlighting an additional CRP site (red bar). **(C)** Scatter-plot comparing transposon insertion frequencies at every position on the 9,010 bp plasmid. Each data point represents a different position on the plasmid. Transposon insertion frequencies are plotted for a sample with no CRP added (*x*-axis) and a sample incubated with 14 nM CRP (*y*-axis). Plasmid positions within 100 bp of the edge of either of the two high-affinity CRP sites (i.e., the edge of the two TGTGAtgtacaTCACA sequences) are colored red, with the intensity of red indicating the distance to the CRP sites.

We compared the extent of footprinting at all positions across the plasmid with 14 nM CRP (Figure 2C). Footprinting by CRP was observed predominantly at positions near the two CRP sites. We also observed evidence of hypersensitive positions flanking the CRP binding sequence, i.e., positions with increased transposition in the presence of CRP (purple arrows in Figure 2A).

### Genome-wide Footprint-seq of CRP

To determine the genome-wide binding profile of CRP, we performed Footprint-seq using *E. coli* genomic DNA. We matched the concentration of genomic DNA to that used for plasmid DNA, and we used the same concentrations of CRP as used with the plasmid. Transposition frequencies across the chromosome were highly correlated between replicates (Figure S2); hence, we combined replicate datasets for most further analysis. We determined the relative extent of CRP-dependent footprinting in regions containing CRP sites described by the RegulonDB database as (i) “confirmed” (66 sites), (ii) supported by “strong” evidence (114 sites), or (iii) supported by “weak” evidence (123 sites) (15). Note that CRP sites upstream of *lacZ*, *mhpR*/*mhpA*, *galE*, and *mlc* could not be analyzed because they were in regions deleted in the strain used to generate genomic DNA. Examples of CRP-bound sites are shown in Figure 3. The site upstream of *focA* is classified by RegulonDB as “confirmed”, and the site upstream of *glnA* is classified as being supported by “weak” evidence. Similar plots for all regions tested are shown in Figures S3, S4, and S5.

**Figure 3.**
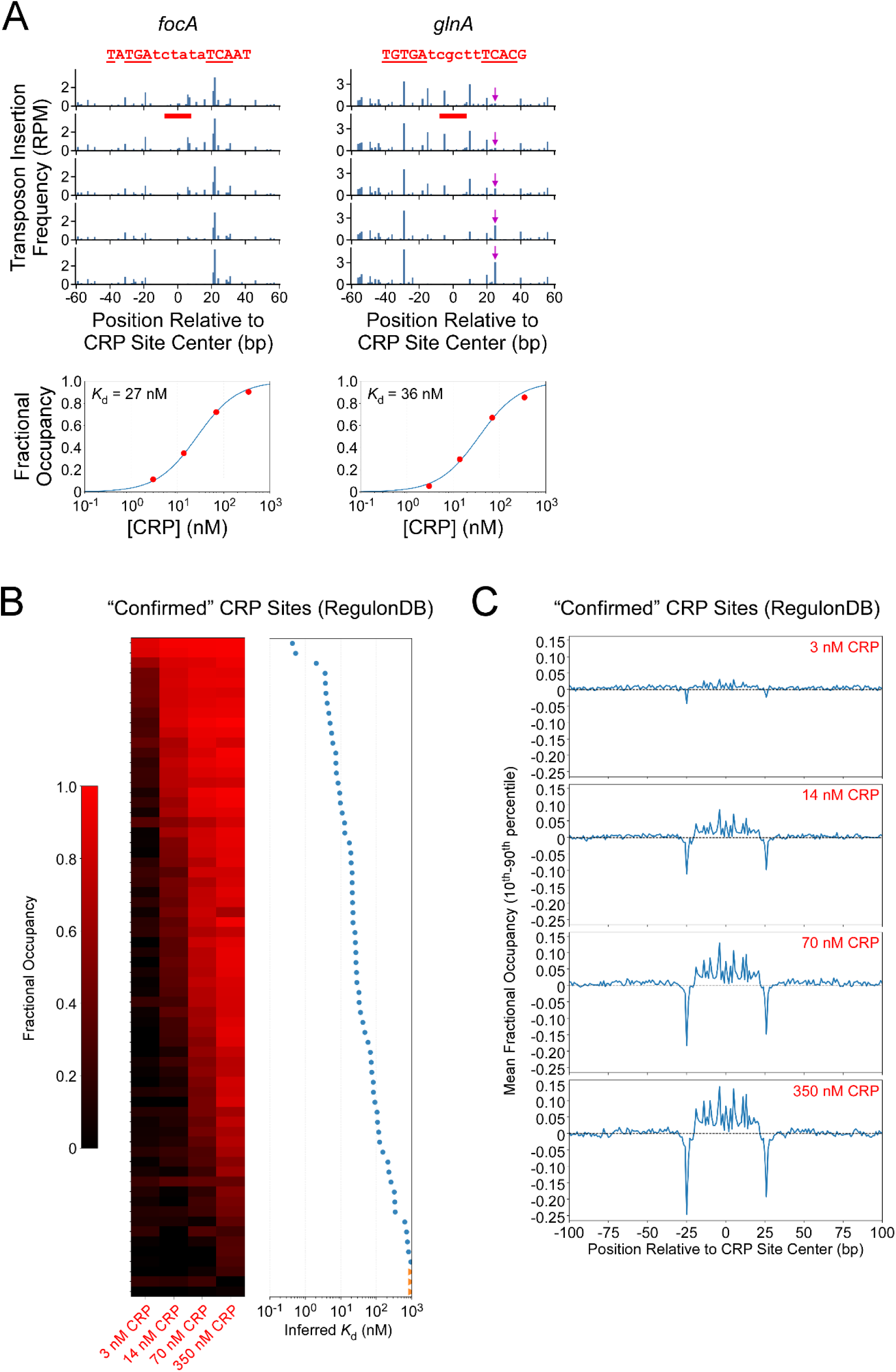
Footprint-seq data for previously identified chromosomal CRP sites. **(A)** Upper panels: transposon insertion frequencies for two chromosomal regions of 120 bp, centered on previously described CRP sites (red horizontal bars). DNA sequences of the CRP sites are indicated above the graphs. Purple arrows indicate a hypersensitive site. Lower panels: Fractional occupancy of the indicated CRP sites (0 = no occupancy, 1 = fully occupied) plotted as a function of CRP concentration. A modeled binding curve was used to estimate the *K*_d_ for each site. **(B)** Heatmap showing fractional occupancy for all CRP sites classified by RegulonDB as “confirmed” across all CRP concentrations tested. The color scale indicates the degree of occupancy. Sites are ordered by estimated *K*_d_, with *K*_d_ values plotted next to the heatmap; data points in orange indicate an estimated *K*_d_ >1 μM. **(C)** Mean difference (10^th^ to 90^th^ percentile values) in coverage in experimental versus control samples for the 200 bp regions surrounding CRP sites classified by RegulonDB as “confirmed”. Values >0 indicate footprinting by CRP, whereas values <0 indicate hypersensitivity. Position 0 is the underlined base in TGTGAnnn<u>n</u>nnTCACA.

The extent of footprinting varied between DNA sites, but overall correlated well with CRP concentration (Figure 3A-C, S3-S7): at 3 nM CRP, little footprinting was observed, whereas at 350 nM CRP, most sites were strongly footprinted. By comparing the extent of footprinting (the reduction in transposition frequency compared to the “no protein” control) by different concentrations of CRP in a 30 bp window centered on each CRP site, we modeled binding curves that allowed us to estimate *K*_d_ values for each site (Figure 3A, B, S3-S7).

In addition to observing a reduction in transposition frequency due to CRP footprinting, we saw hypersensitive positions (e.g., purple arrow in Figure 3A) similar to those seen for the plasmid sites. Comparison of transposition frequencies across all the tested CRP sites revealed a symmetric pattern of protection from positions 19 bp upstream to 20 bp downstream of the center of the TGTGA–N_6_–TCACA motif, flanked by a pair of prominent, precisely positioned hypersensitive positions 25 bp upstream and 26 bp downstream of the center of the TGTGA–N_6_–TCACA motif (Figure 3C) [we consider the center of a site to be the underlined base, with respect to an optimal CRP half-site: TGTGTNNN**<u>N</u>**]. We conclude that Footprint-seq can (i) map protein-DNA interactions with high resolution, and (ii) quantitatively measure occupancy across sites.

### De novo identification of CRP sites using Footprint-seq

To identify CRP sites across the *E. coli* genome directly from Footprint-seq data, we searched for 30 bp regions with significantly lower transposition density with 350 nM CRP than with no CRP, taking into account variability between biological replicates. Differences in transposition density between samples with/without protein could be due to footprinting by CRP (i.e., a true positive) or random variation between experiments (i.e., a false positive). Differences in transposition density due to random variation should occur equally in both directions, i.e., increased or decreased density at 350 nM CRP. We selected a *p*-value threshold of 1e^-5^, which identified no regions with higher transposition density at 350 nM CRP (red arrow on the *y*-axis in Figure 4A). Thus, we identified 1,217 putative CRP-bound regions. We note that this is a highly stringent threshold, with an estimated false discovery rate close to 0. Moreover, it is likely that some regions with higher transposon density at 350 nM CRP than with no CRP are not false positives, but rather represent hypersensitive sites flanking CRP-bound regions. Hence, we expect false negative sites, i.e., genuine CRP-bound regions that failed to meet the stringent *p*-value threshold.

**Figure 4.**
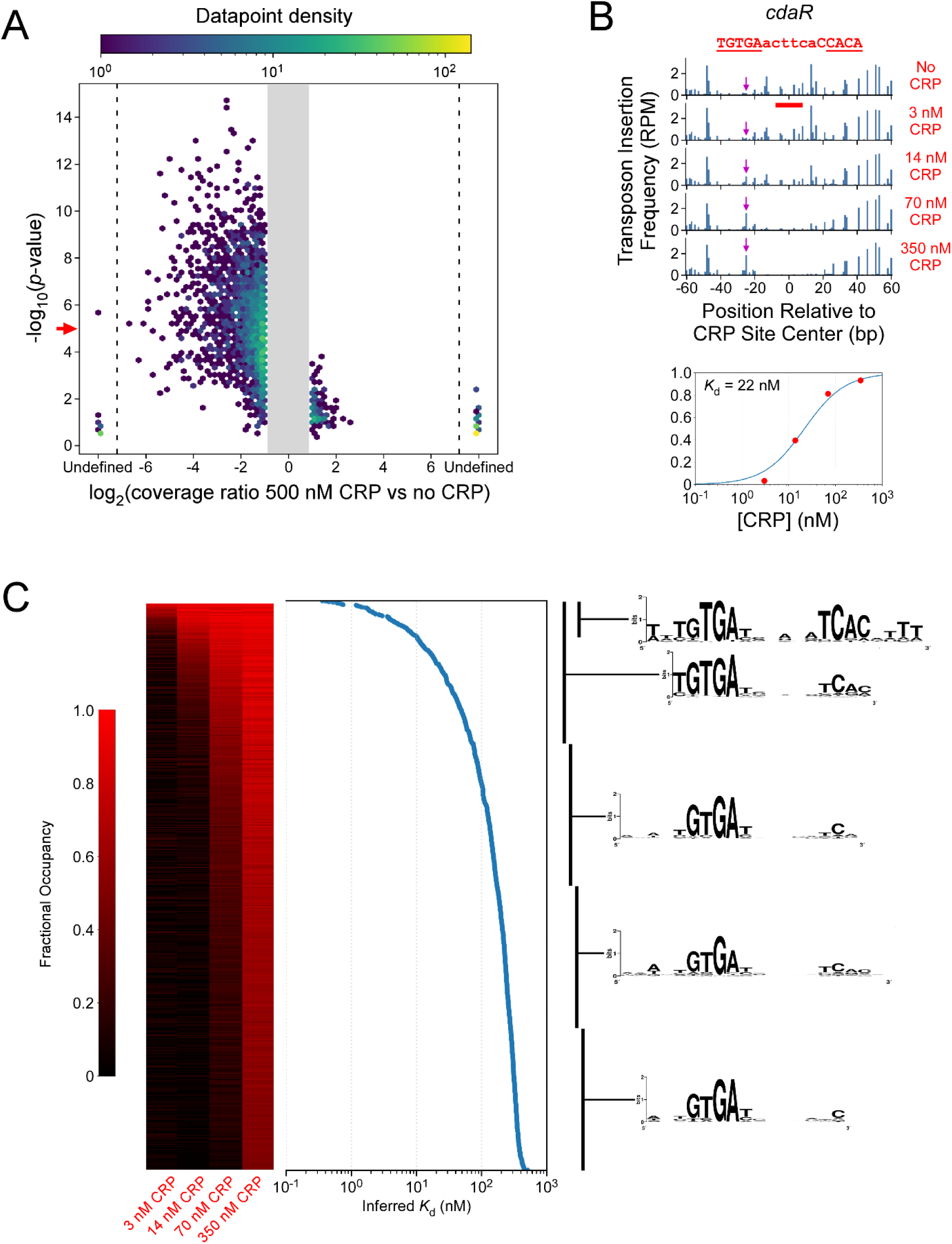
*De novo* identification of CRP binding sites genome-wide. **(A)** Volcano hex-plot showing the ratio in transposon insertion frequency +/− CRP (350 nM) for 30 bp regions across the chromosome (*x*-axis) plotted against the probability of the observed difference in transposon insertion frequency occurring due to random noise (not corrected for multiple-hypothesis testing). The color indicates the density of data points. Values are not plotted for ratios between 0.5 and 2. “Undefined” values on the *x*-axis indicate regions with zero coverage in either sample. The red arrow on the *y*-axis indicates the *p*-value cut-off (10^-5^) used to select the top candidate regions. **(B)** Upper panel: transposon insertion frequencies for a chromosomal region of 120 bp, centered on the newly discovered CRP site upstream of *cdaR*. The inferred CRP site is inidicated by a red horizontal bar, and the DNA sequence of the CRP site is indicated above the graph. Purples arrow indicate a hypersensitive site. Lower panel: Fractional occupancy of the indicated CRP site (0 = no occupancy, 1 = fully occupied) plotted as a function of CRP concentration. A modeled binding curve was used to estimate the *K*_d_. **(C)** Heatmap showing fractional occupancy for all *de novo* identified CRP sites across all CRP concentrations tested. The color scale indicates the degree of occupancy. Sites are ordered by estimated *K*_d_, with values plotted next to the heatmap. Enriched sequence motifs are indicated for sites with an estimated *K*_d_ < 10 nM (“< 10 nM”), and each of the first (“Q1”), second (“Q2”), third (“Q3”), and fourth (“Q4”) quartiles based on estimated *K*_d_.

The 1,217 putative CRP-bound regions we identified have inferred *K*_d_ values ranging from 0.4 to 510 nM. An example of a novel CRP site, upstream of *cdaR*, is shown in Figure 4B. We searched for enriched sequence motifs within selected groups of putative CRP-bound regions. First, we searched within the 77 regions with an inferred *K*_d_ of <10 nM (Table 1). These sites are all associated with an instance of a sequence motif that closely matches the consensus CRP binding site. Moreover, enrichment of A/T-rich sequence in the 3-4 bp flanking the TGTGA/TCACA half-sites suggests that the identity of those positions influences CRP affinity, perhaps by affecting the ability of the DNA to be bent. We next stratified the 1,217 putative CRP-bound regions into four quartiles based on inferred *K*_d_. We searched within each quartile group for enriched sequence motifs. For the top three quartile groups, we identified a match to an enriched DNA sequence motif in every sequence; the enriched DNA sequence motif closely resembles the known CRP consensus site (Figure 4C). For the bottom quartile of regions, we identified a match to an enriched DNA sequence motif in all but two regions (Figure 4C). Across all quartiles, the center of the predicted binding site was within 10 bp of the center of the match to the enriched sequence motif in 92% of cases, highlighting the high resolution of Footprint-seq. As *K*_d_ values increased (i.e., weaker CRP binding), the information content of the motif decreased. Specifically, the outside position of each half-site (underlined bases: <u>T</u>GTGA/TCAC<u>A</u>) appears to be the least important for CRP binding, but is required for high-affinity binding. Moreover, the weakest sites rely primarily on a single half-site.

**Table 1.**
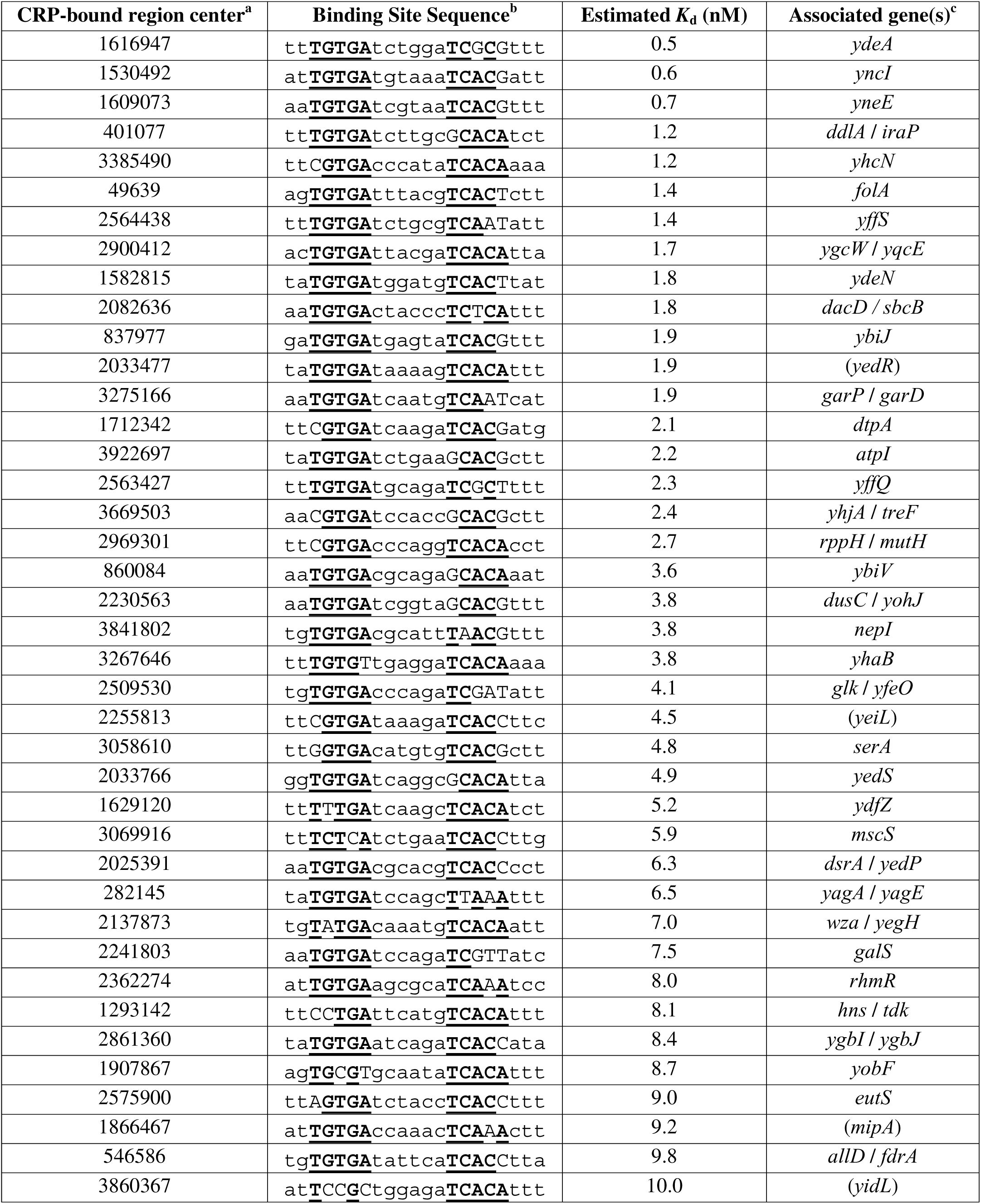
Novel CRP sites with inferred *K*_d_ values below 10 nM.

Of the 308 CRP sites that have been previously described or predicted, 178 are within 100 bp of one of the 1,217 CRP-bound regions we identified, with a median distance of 5 bp from the CRP site center inferred from Footprint-seq data alone (i.e., without searching for motif matches). We also identified many high-affinity CRP sites that have not been previously described. Of the 77 CRP-bound regions with inferred *K*_d_ values of <10 nM, 37 are >500 bp from a previously described site, and 33 of these 37 novel sites are intergenic, strongly suggesting most or all are regulatory (Table 1). Overall, our data indicate that Footprint-seq can identify sites of protein-DNA interaction with high sensitivity and high resolution across a wide range of affinities.

### Validation of CRP binding sites identified using Footprint-seq

We used electromobility shift assays (EMSAs) to independently determine the *K*_d_ of CRP for two sites identified using Footprint-seq: a previously described CRP site in the intergenic region between *gcd* and *hpt* (16), and a novel site in the intergenic region upstream of *cdaR* (Figure 4B). The *K*_d_ values for these sites inferred from Footprint-seq data are 0.44 nM and 22 nM, respectively. We used EMSAs to compare the binding of CRP to the two sites across a range of CRP concentrations from 0.5 to 128 nM (Figure 5). In both cases, the EMSA data are consistent with the *K*_d_ values inferred from Footprint-seq data, suggesting that Footprint-seq *K*_d_ measurements accurately reflect the affinity of the analyzed DNA sites.

**Figure 5.**
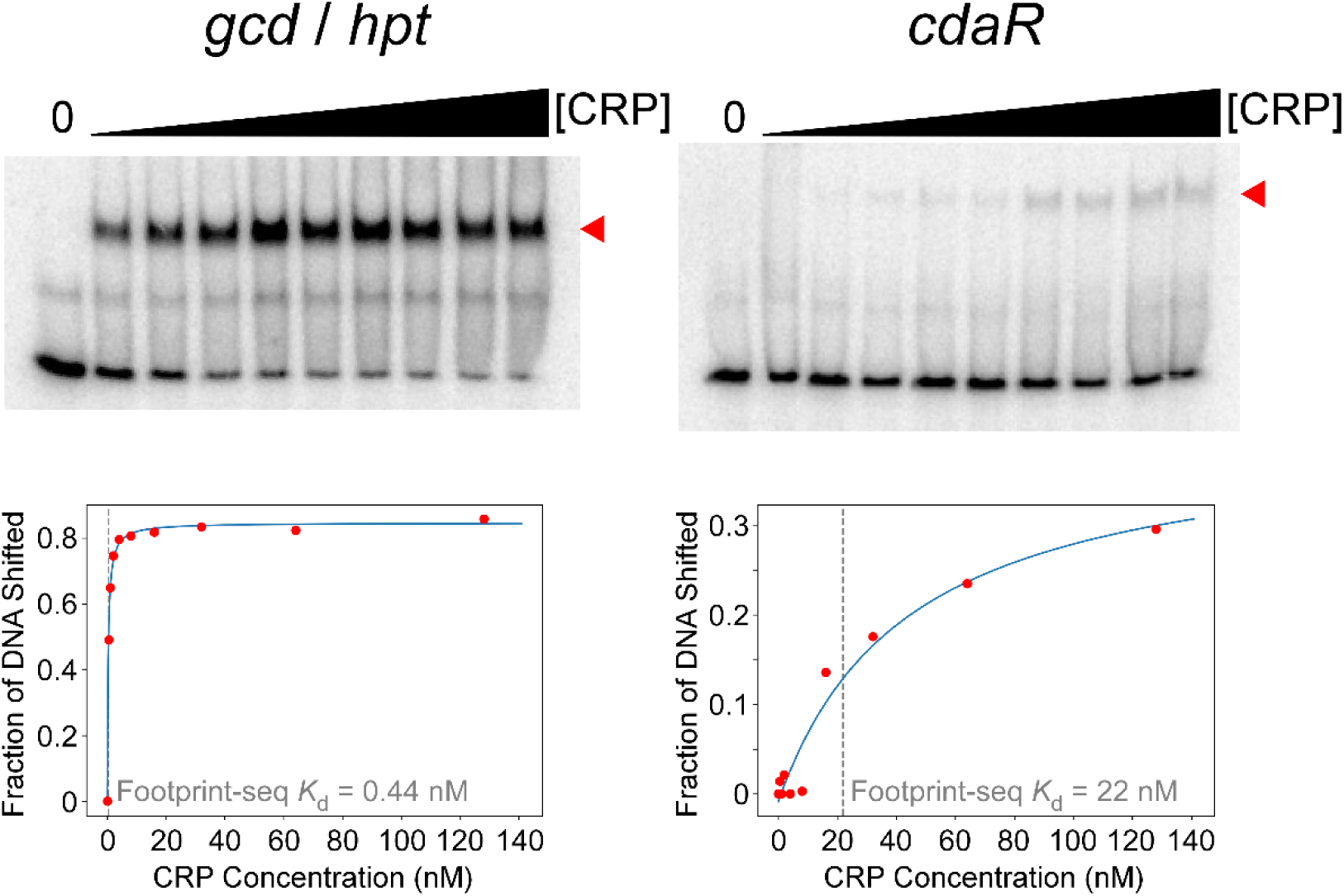
Electromobility shift assays support *K*_d_ estimates from Footprint-seq. Electromobility shift assasy for two putative CRP sites identified by Footprint-seq. The CRP site upstream of *gcd* and *hpt* is classified by RegulonDB as “confirmed”, and has an inferred *K*_d_ from Footprint-seq of 0.44 nM. The CRP site upstream of *cdaR* has not been described previously, and has an inferred *K*_d_ from Footprint-seq of 22 nM. Gels show reactions incubated with 0, 0.5, 1, 2, 4, 8, 16, 32, 64, and 128 nM CRP (left to right). Red triangles indicate the position of the shifted bands. Graphs below each gel show quantification of the proportion of shifted DNA; gray vertical lines indicate the *K*_d_ values inferred from Footprint-seq data.

### Comparison of Footprint-seq to in vitro ChIP-seq

We previously used *in vitro* ChIP-seq to map the binding of five transcription factors across the *E. coli* genome, combining those data together with *in vivo* ChIP-seq data to train BoltzNet, a neural network that accurately predicts transcription factor binding energy from DNA sequence alone (4). The *in vitro* ChIP-seq method involves mixing of purified DNA-binding protein with sheared genomic DNA, followed by immunoprecipitation of the protein, library amplification, and sequencing. To benchmark Footprint-seq against an established method, we used *in vitro* ChIP-seq to map the binding of CRP across the *E. coli* genome, identifying 396 enriched regions. We searched these regions for enriched sequence motifs using MEME (17), identifying 254 “high-confidence” CRP sites (Table S3; see Methods for details). Three of these putative CRP sites are in genomic regions missing in the strain we isolated genomic DNA from for Footprint-seq. We analyzed the remaining 251 high-confidence sites using our Footprint-seq data. In 232 cases (92%), we identified a CRP site with a *K*_d_ of <500 nM (Figure 6; Figure S10). Thus, Footprint-seq detects almost all high-confidence CRP sites identified using *in vitro* ChIP-seq.

**Figure 6.**
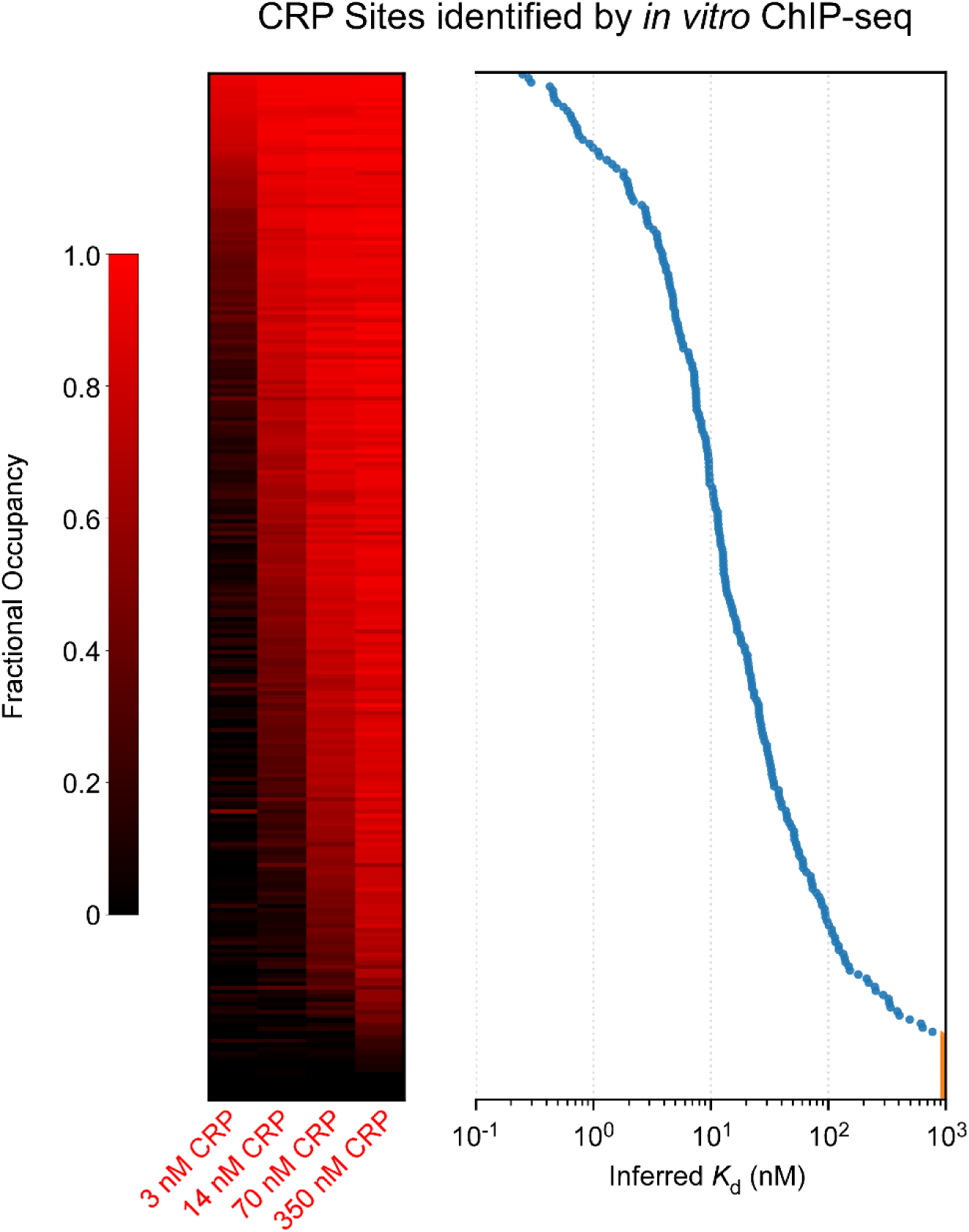
Comparison of Footprint-seq to *in vitro* ChIP-seq. Heatmap showing Footprint-seq fractional occupancy across all CRP concentrations tested, for “high-confidence” CRP sites identified by *in vitro* ChIP-seq. The color scale indicates the degree of occupancy. Sites are ordered by estimated *K*_d_, with *K*_d_ values plotted next to the heatmap; data points in orange indicate an estimated *K*_d_ >1 μM.

We next compared the spatial resolution of Footprint-seq and *in vitro* ChIP-seq. For *in vitro* ChIP-seq, we compared the peak position to the center of the associated CRP site in cases where we found a match to the CRP site motif. The median distance between the peak position and the site center is 11 bp. For Footprint-seq, we searched for enriched sequence motifs in the top 396 sites (the same number of sites as there are enriched regions from *in vitro* ChIP-seq) identified in our *de novo* search. We found an associated CRP site for every CRP-bound region identified by Footprint-seq; the median distance between the center of the CRP-bound region and the center of the site is 4 bp. Thus, Footprint-seq has higher spatial resolution than *in vitro* ChIP-seq.

## DISCUSSION

We have established Footprint-seq as a method that determines the position and strength of protein-DNA interactions *in vitro*, on a genome-wide scale. By comparison with *in vitro* ChIP-seq, we conclude that Footprint-seq offers advantages over methods for genome-scale mapping of protein-DNA interactions that rely on enrichment of fragmented DNA. First, the resolution of Footprint-seq is higher than that of *in vitro* ChIP-seq; although *in vitro* ChIP-seq has high resolution (median of 11 bp between the inferred and actual site center), the resolution of Footprint-seq is considerably higher (median of 4 bp between the inferred and actual site center). For methods that use fragmented genomic DNA, a single binding site can be found in many overlapping DNA fragments. Hence, the binding site location must be inferred from the position with the highest overall enrichment across overlapping DNA fragments, similar to analysis of ChIP-seq data. This can limit the resolution, especially for weaker binding sites. By contrast, when using Footprint-seq, protection from transposition is tightly bounded around the binding site (Figure 3C). Second, the sensitivity of Footprint-seq is higher than that of *in vitro* ChIP-seq. We identified ∼3 times as many putative CRP sites using Footprint-seq than we did using *in vitro* ChIP-seq, and the extremely high association of even the weakest of these sites with sequences resembling a CRP site provides high confidence in the calls. Third, because Footprint-seq leverages tagmentation, the entire method is technically simple, with only ∼1 hour of hands-on time. Fourth, hypersensitive sites can provide additional position information, although we do not know if proteins other than CRP will cause hypersensitive sites. We speculate that hypersensitive sites are due either to distortions in the DNA caused by CRP binding, or physical interactions between CRP and the transposase.

There are also weaknesses of Footprint-seq relative to other methods for mapping protein-DNA interactions genome-wide *in vitro*. First, Footprint-seq relies largely on the loss of signal, meaning that genomic regions with inherently low transposition frequencies will be more difficult to interrogate. This weakness can be effectively mitigated by increasing sequencing depth (we aim for 40 million reads per sample for *E. coli* genomic DNA). Second, because *de novo* identification of protein-bound DNA sites relies on searching for regions of reduced transposition frequency, it could be complicated by the presence of hypersensitive positions. In the case of CRP, we relied on the fact that the hypersensitive positions are at the edge of the region with altered transposition frequency relative to the DNA site. Third, Footprint-seq requires that protein-DNA interactions can occur under conditions that are conducive to tagmentation. We found that extending the time for transposition mitigated this problem for CRP.

We applied Footprint-seq to a single plasmid and to genomic DNA. It could be used to study the binding of proteins to any similarly sized genome for which DNA can be isolated, providing a simple method to study the regulons of DNA-binding transcription factors in species that are not genetically tractable. It could also be applied to plasmids or PCR products, either individually or in pools. Thus, Footprint-seq could functionally replace an EMSA, while simultaneously providing high-resolution positional information.

## MATERIALS AND METHODS

### Source of plasmid and *E. coli* chromosomal DNA

The plasmid used in this study was ML1, which contains a pair of strong CRP sites (14). *E. coli* genomic DNA for Footprint-seq was isolated from strain RR1023, which carries a derivative of the F factor, pOX38 (18). *E. coli* genomic DNA for *in vitro* ChIP-seq was isolated from strain MG1655 (19).

### Expression and purification of CRP

CRP for Footprint-seq and electromobility shift assays was purified as described previously (20). For *in vitro* ChIP-seq, CRP with a C-terminal His_6_-tag was purified by overexpression from a T7 promoter with a *lac* operator. Expression and purification were performed using a QIAexpress kit (Qiagen) following the manufacturer’s protocol.

### Footprint-seq

The CRP Footprint-seq reactions were assembled in 1X binding buffer (40 mM HEPES, pH 8.0, 5 mM MgCl_2_, 100 mM potassium glutamate), 50 ng DNA (plasmid or genomic; genomic DNA was isolated using the Qiagen Blood and Tissue kit, following the manufacturer’s protocol), 0.2 mM cAMP, and CRP at final concentrations of 3 nM, 14 nM, 70 nM, or 350 nM, supplied in CRP stock buffer (10 mM sodium phosphate, pH 6.8, 0.1 mM EDTA, 0.2 M NaCl, 50% [w/v] glycerol). A no-CRP control received the same volume of CRP stock buffer without protein. Reactions were incubated at 37 °C for 10 minutes, followed by addition of 5 μL TDE1 Nextera transposase and incubation for 30 minutes at 37 °C. DNA was purified using a Qiagen MinElute kit and eluted in 10 μL of elution buffer. Footprint-seq libraries were amplified using the Illumina Nextera DNA Library Prep Kit, following the manufacturer’s 2016 instructions. The kit-supplied custom Nextera PCR primers were used with the following cycling parameters: 72 °C for 5 minutes and 98 °C for 30 seconds, followed by 6 cycles of 98 °C for 10 seconds, 63 °C for 30 seconds, and 72 °C for 1 minute. Sequencing was performed using an Illumina NextSeq Instrument at the Wadsworth Center Applied Genomics Technologies Core Facility.

### Footprint-seq analysis

#### Calculating normalized transposition frequency values

Sequence reads from .fastq files for Footprint-seq experiments with chromosomal DNA were aligned to the *E. coli* MG1655 genome sequence (GenBank ID U00096.3) using Rockhopper (version 2.03) (21). The resulting .sam files were used to generate .gff files that list transposon insertion frequency at each associated genomic position. For .fastq files for Footprint-seq experiments with plasmid DNA, we used Rockhopper (version 2.03) to align sequence reads to a reference genome consisting of two copies of the plasmid followed by the *Mycobacterium tuberculosis* H37Rv chromosome sequence. The position of sequence reads aligning to the second copy of the plasmid was adjusted to match the first copy of the plasmid, and sequence reads aligning to non-plasmid sequence were discarded. To generate .gff files, we determined the sequence at the transposon insertion site (5’ end of each sequence read) and shifted reads on the plus strand by +4 bp, and reads on the minus strand by −5 bp, to account for the 9 bp sequence duplication that occurs during Tn5 transposition (22). Sequence read coverage in .gff files was normalized to reads per million (RPM). Most analyses presented here use .gff files that represent an average (mean) of two replicate datasets.

#### Calculating Fractional Occupancy Values

The transposon insertion frequency values in a 30 bp window were summed for the control (no CRP) sample and for an experimental sample. “Fractional Occupancy” is defined as the ratio of the sum from the experimental sample to that from the control sample, subtracted from 1. Values >1 were set to 1. By this definition, if the transposon insertion frequency values are the same in the control and experimental samples, the Fractional Occupancy is 0, whereas if there is a complete inhibition of transposition in the experimental sample, the Fractional Occupancy is 1.

#### Meta-analysis of transposon insertion frequency differences around selected sites

Graphs showing meta-analysis of transposon insertion frequency differences between control and experimental samples were plotted by calculating the absolute difference in transposon insertion frequency between a control and experimental sample in a 200 bp window around a defined site position. Values at each position were rank-ordered, and the mean of the 10^th^ to the 90^th^ percentile values was calculated. These mean values were plotted as a function of position relative to the site centers.

#### Estimating K_d_ values

To estimate *K*_d_ values for individual sites, we fitted a non-cooperative Langmuir binding model to Fractional Occupancy values across a range of CRP concentrations tested. The binding model assumes the formula *F* = *x* / (*K*_d_ + *x*), where *F* is the Fractional Occupancy and *x* is CRP concentration. The model assumes a single binding site with non-cooperative CRP binding, and a dynamic range of Fractional Occupancy from 0 to 1. Before model-fitting, we inferred an approximate *K*_d_ as a starting value by searching for neighboring protein concentrations for which Fractional Occupancy for the lower concentration was <0.5 and Fractional Occupancy for the higher concentration was >0.5. In such a situation, the starting *K*_d_ estimate was determined by interpolating the two data points and determining the concentration for which Fractional Occupancy was 0.5. If none of the Fractional Occupancy exceeded 0.5, the geometric mean of protein concentrations tested was used as the starting value for modeling. A model was fitted using the scipy.optimize.least_squares function (23).

#### De novo Identification of CRP binding sites from Footprint-seq data

We compared transposition frequencies between two replicates of a control dataset (i.e., no CRP) and two replicates of an experimental dataset that used 350 nM CRP, in sliding 30 bp windows across the genome, with a 1 bp step size. At each genomic position, we determined the total transposition frequency within the 30 bp window centered on that position for all four datasets. We summed values for replicates and determined the ratio of the sums. If the ratio was >2 or <0.5, we fit a mixed-effects model using the Python module *statsmodel* (24), and determined a *p*-value for the condition effect, i.e., the difference in transposition frequency between control and experimental samples, taking into account variability between replicates. To collapse overlapping windows into a single genome coordinate, windows were ranked in ascending order of *p*-value. Beginning with the lowest *p*-value, a window was retained only if its midpoint was more than 60 bp from every previously retained midpoint. Thus, selected sites represent local *p*-value minima, with all sites separated by at least 61 bp. Only sites with a *p*-value below 1e^-5^ were reported as putative CRP sites.

### Electrombilty shift assays

DNA fragments were generated by PCR using E. coli MG1655 genomic DNA as template and oligonucleotides Bio-TTGCCATAAAGAGACACCAT and AAGACACGCAGTATACCAG (*gcd*/*hpt* fragment), and Bio-CACCATCTACCTGTTAATGC and GCATAAGGAAGTACGTAACG (*cdaR* fragment). After purification, fragments were end-labelled with γ^32^P-ATP (Hartmann Analytic) and T4 polynucleotide kinase (New England Biolabs). Binding reactions were set up with 0.1 nM radiolabeled DNA and CRP (0–128 nM). Reaction buffer contained 125 μM cAMP, 40 mM Tris-acetate (pH 7.9), 150 mM potassium glutamate, 4 mM magnesium glutamate, 100 μM EDTA, 10 μM DTT, 10 mM potassium phosphate (pH 7.9), and 100 μg/mL recombinant albumin (New England Biolabs). Reactions were incubated for 30 min at 37 °C, before protein–DNA complexes were resolved by electrophoresis on a 5% (w/v) non-denaturing polyacrylamide gel (Protogel) in 0.5X TBE buffer, run at 160 V at room temperature. Gels were then dried and exposed to a phosphor screen (BioRad), which was scanned using an Amersham Typhoon biomolecular imager. Band intensities were quantified using ImageJ. Full gel images are shown in Figure S9.

Best-fit curves for EMSA data were generated by nonlinear least-squares fitting of a one-site, non-cooperative binding model to the fraction of DNA shifted over a range of CRP concentrations. The fraction shifted was calculated as *S*/(*S* + *U*), where *S* and *U* are the background-corrected intensities of the shifted and unshifted DNA bands, respectively. The fitted model was *Y = Bottom + (Top − Bottom) × x/(K*_d_ *+ x)*, where *Y* is the fraction shifted, *x* is the protein concentration, and *Bottom* and *Top* are fitted parameters representing the baseline and maximum observed fraction shifted, respectively. Initial parameter estimates were obtained by setting *Bottom* and *Top* to the minimum and maximum observed fractions shifted, respectively, and estimating an initial *K*_d_ as the protein concentration whose observed fraction shifted was closest to the midpoint between *Bottom* and *Top*. Nonlinear least-squares fitting was performed using the scipy.optimize.curve_fit function in SciPy (23).

### Comparison of Footprint-seq to i*n vitro* ChIP-seq

*In vitro* ChIP-seq and associated peak-calling was performed as described previously (4), using 50 ng (42 nM final concentration) or 500 ng (420 nM final concentration) of CRP. “High-confidence” sites (Table S3) were identified by extracting 101 bp regions surrounding peak positions, searching for enriched sequence motifs using MEME (17), and selecting regions with a match to a motif with the known CRP consensus. Analysis of Footprint-seq data was performed using the position of the center of the sequence matching the CRP binding site motif.

## DATA AVAILABILITY

Raw and processed Footprint-seq data are available at EBI ArrayExpress using accession number E-MTAB-16502. Raw *in vitro* ChIP-seq data are available at the NIH Short Read Archive using BioProject ID PRJNA1514151. All Python code is available at https://github.com/wade-lab/Footprint-seq_analysis.

## ACKNOWLEDGEMENTS

We thank the Wadsworth Center Protein Purification Core Facility, the Wadsworth Center Applied Genomic Technologies Core Facility, the Wadsworth Center Media and Glassware Core Facility, and the Boston University Microarray and Sequencing Resource Core Facility for technical support. We thank Keith Derbyshire, Jon Paczkowski, and Tung Le for helpful discussions.

