## Supplementary figures and images for "Footprint-seq: a simple method to quantitatively map *in vitro* protein-DNA interactions on a genome-wide scale at high spatial resolution"

### Figures S1, S2, S6, S7, S9

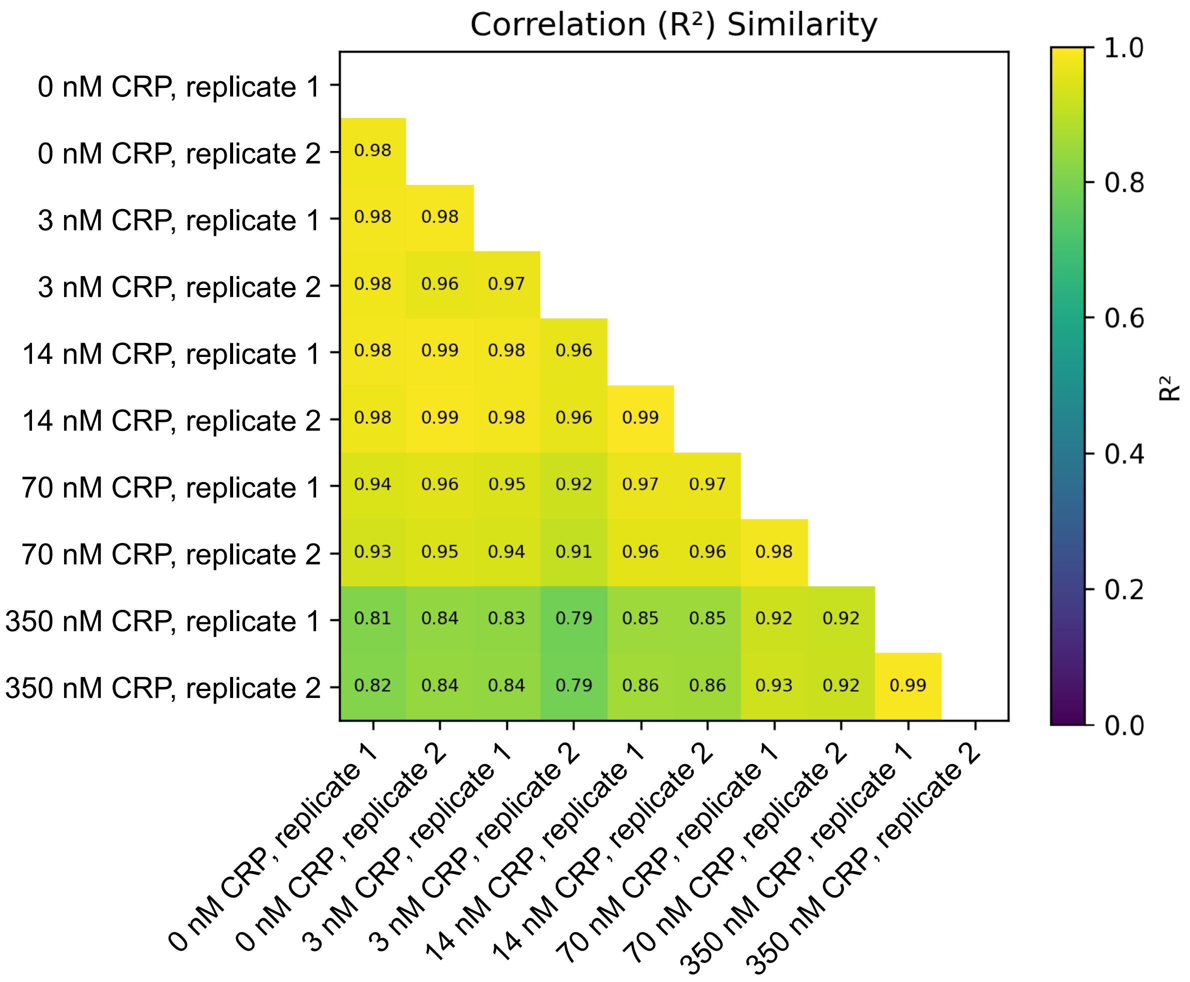

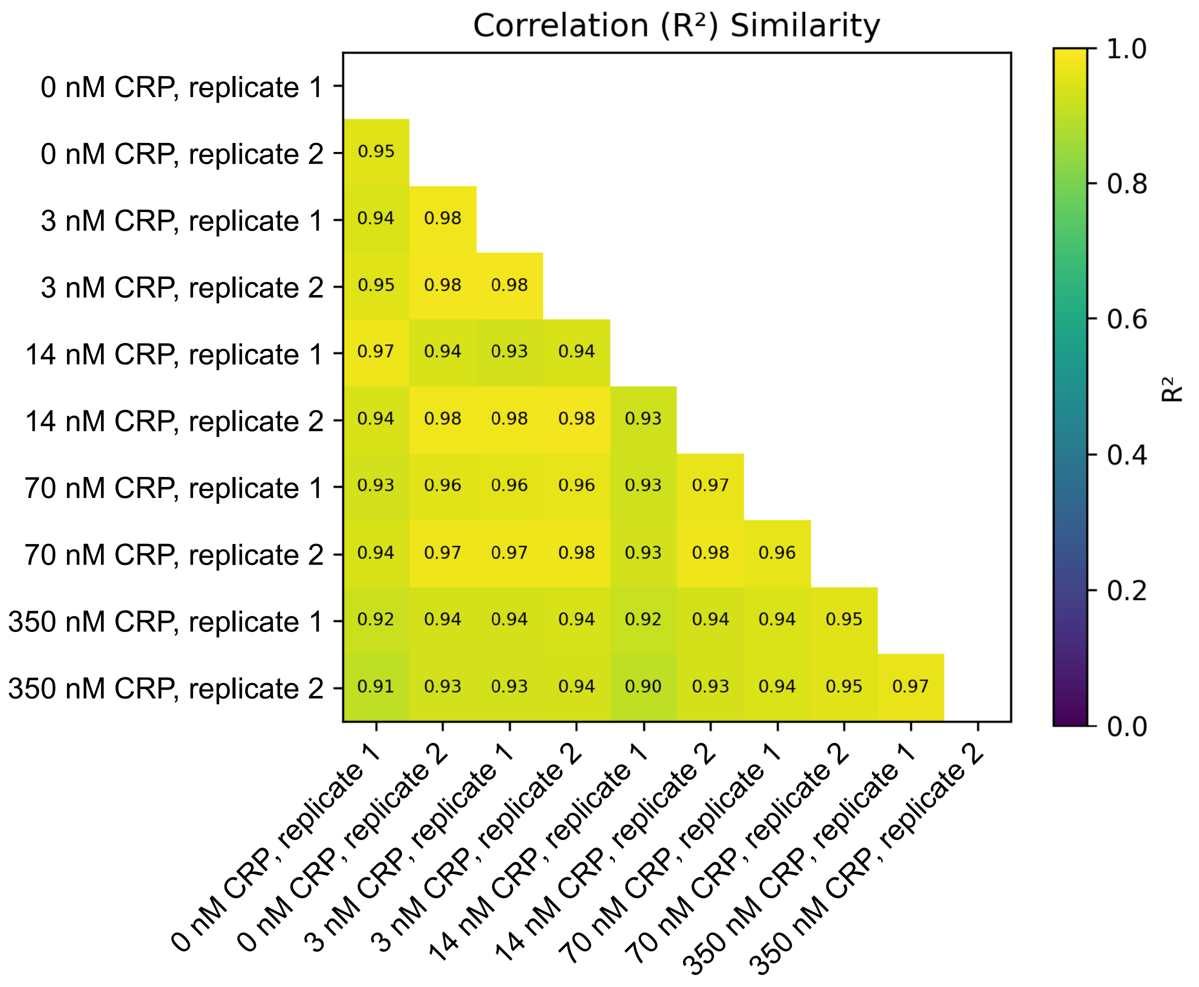

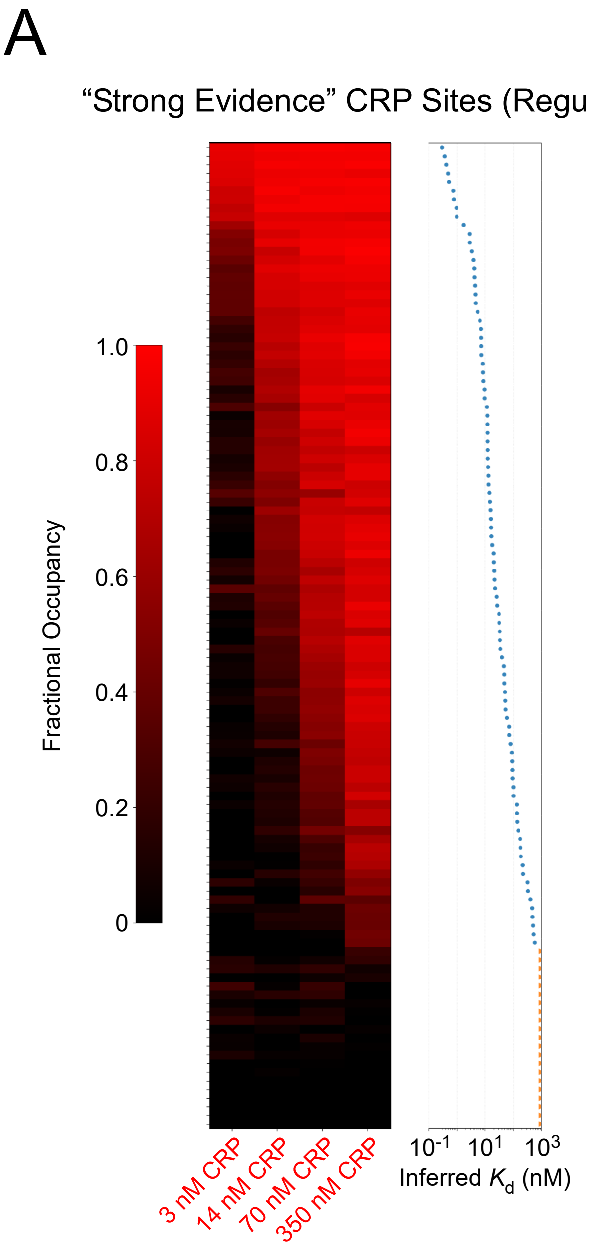

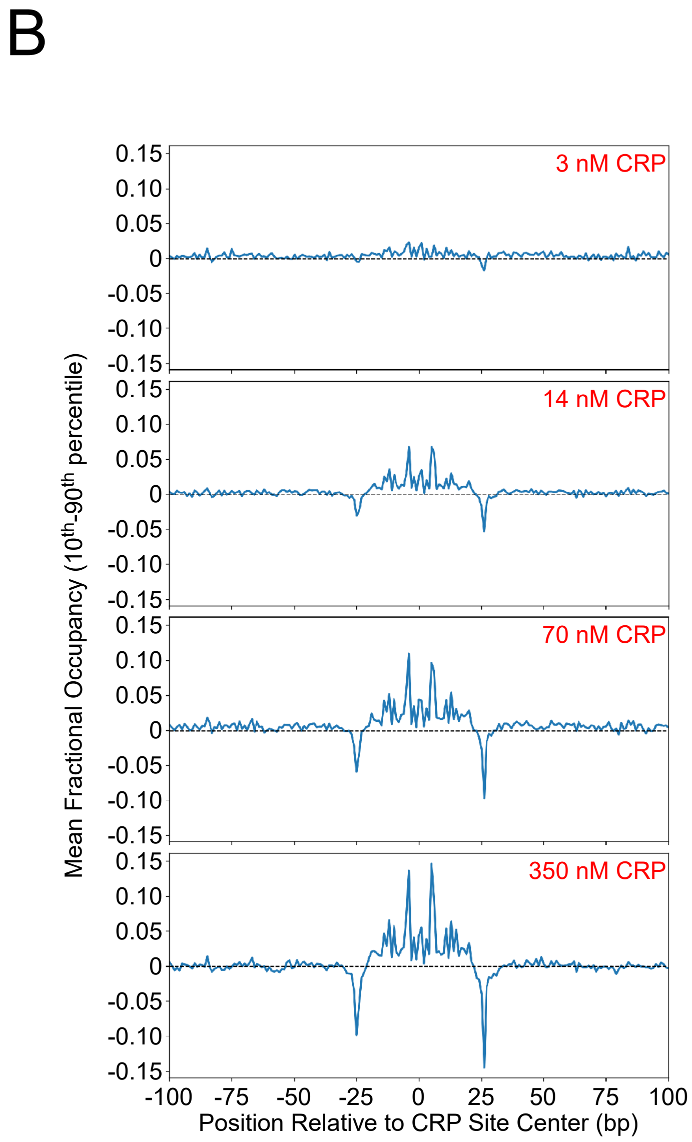

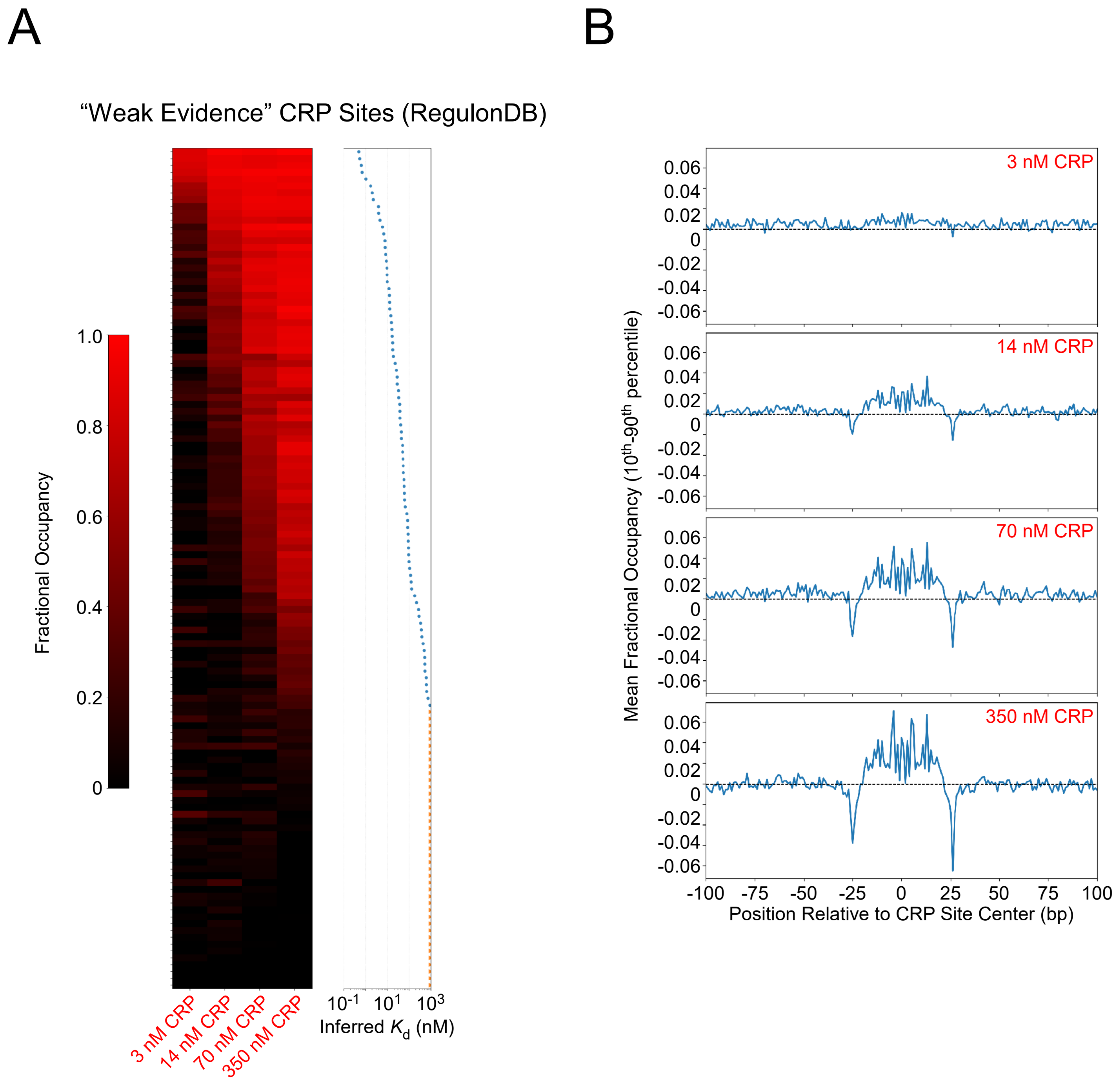

#### Figure S9

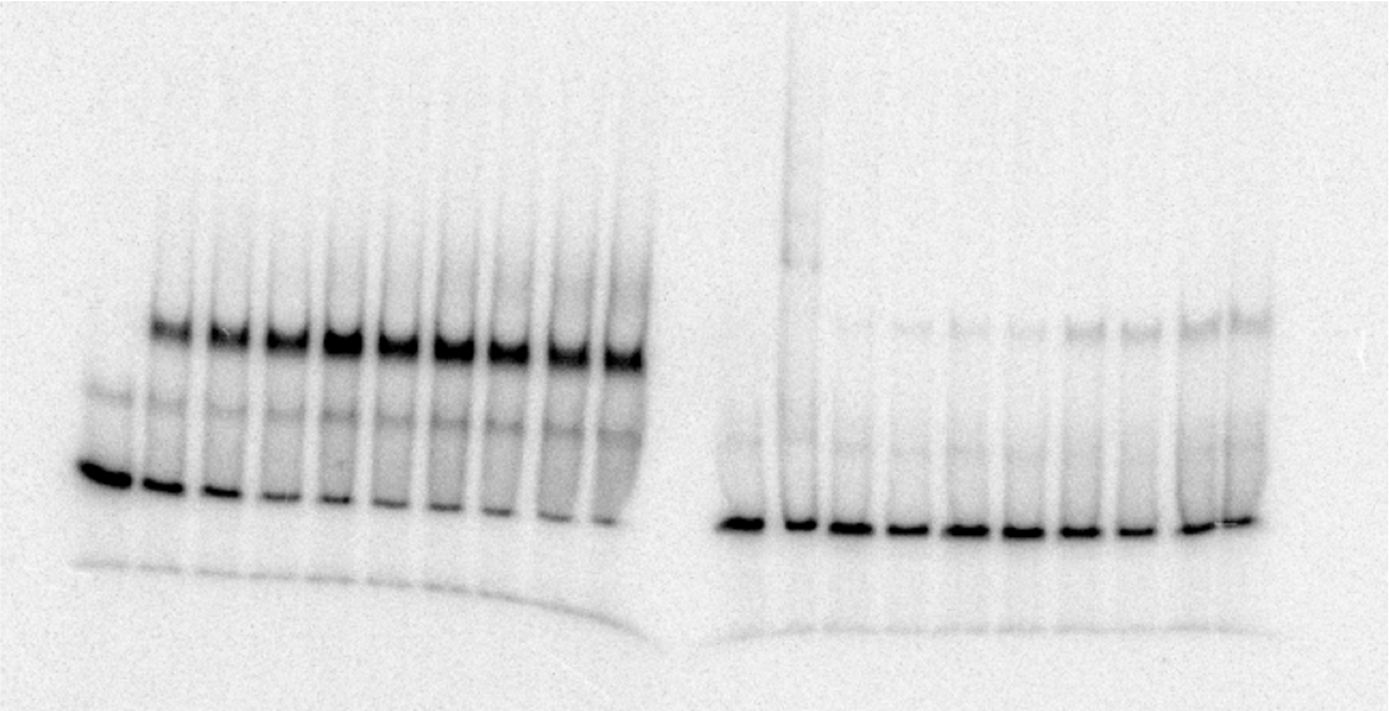
