## Supplementary material for "Footprint-seq: a simple method to quantitatively map *in vitro* protein-DNA interactions on a genome-wide scale at high spatial resolution": Figure S3

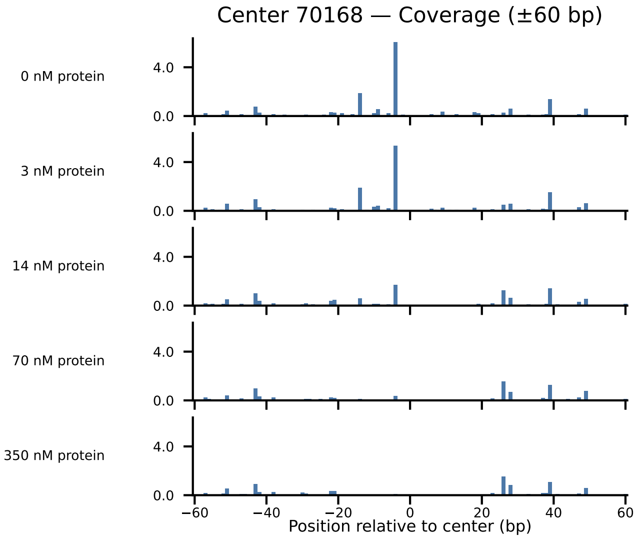
Center 70168 — Coverage (±60 bp)
0 nM protein
3 nM protein
14 nM protein
70 nM protein
350 nM protein
4.0
0.0
−60
−40
−20
0
20
40
60
Position relative to center (bp)

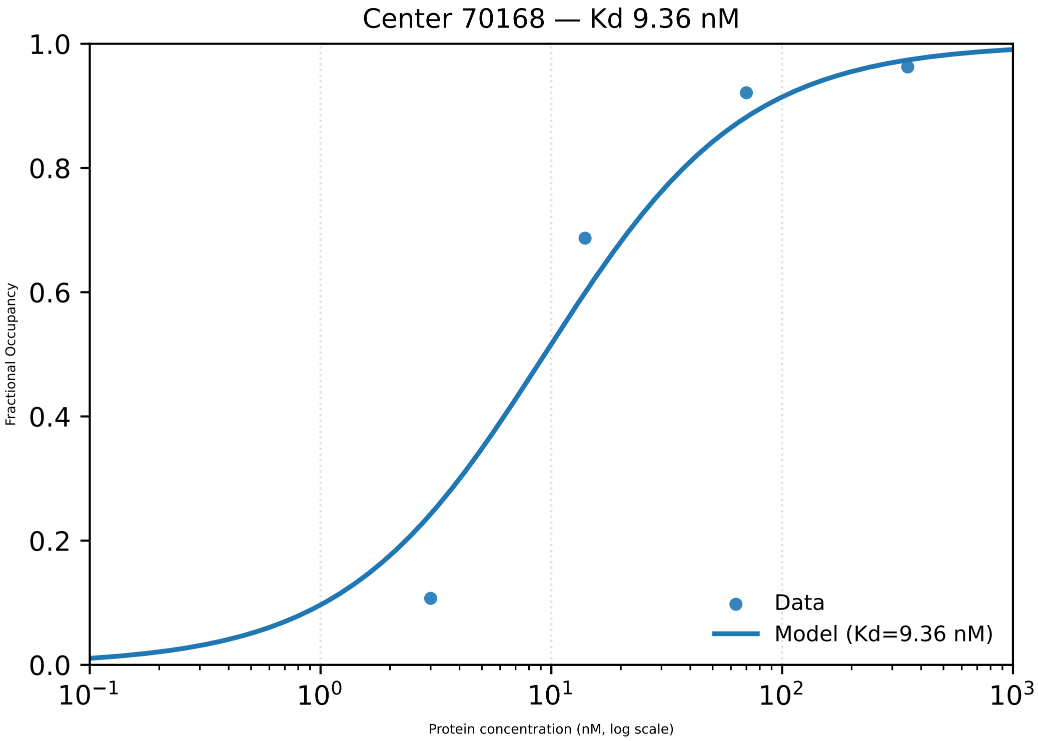

Center 70168 — Kd 9.36 nM
1.0
0.8
0.6
0.4
0.2
0.0
Fractional Occupancy
$10^{-1}$
$10^{0}$
$10^{1}$
$10^{2}$
$10^{3}$
Protein concentration (nM, log scale)
Data
Model (Kd=9.36 nM)

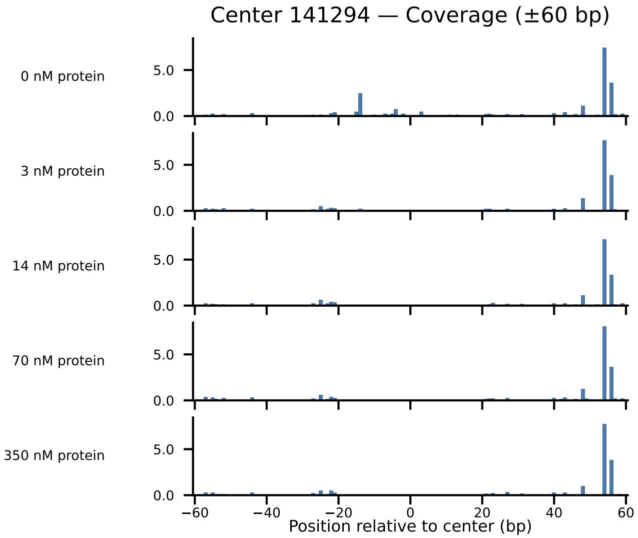

Center 141294 — Coverage (±60 bp)
0 nM protein
3 nM protein
14 nM protein
70 nM protein
350 nM protein
5.0
0.0
−60
−40
−20
0
20
40
60
Position relative to center (bp)

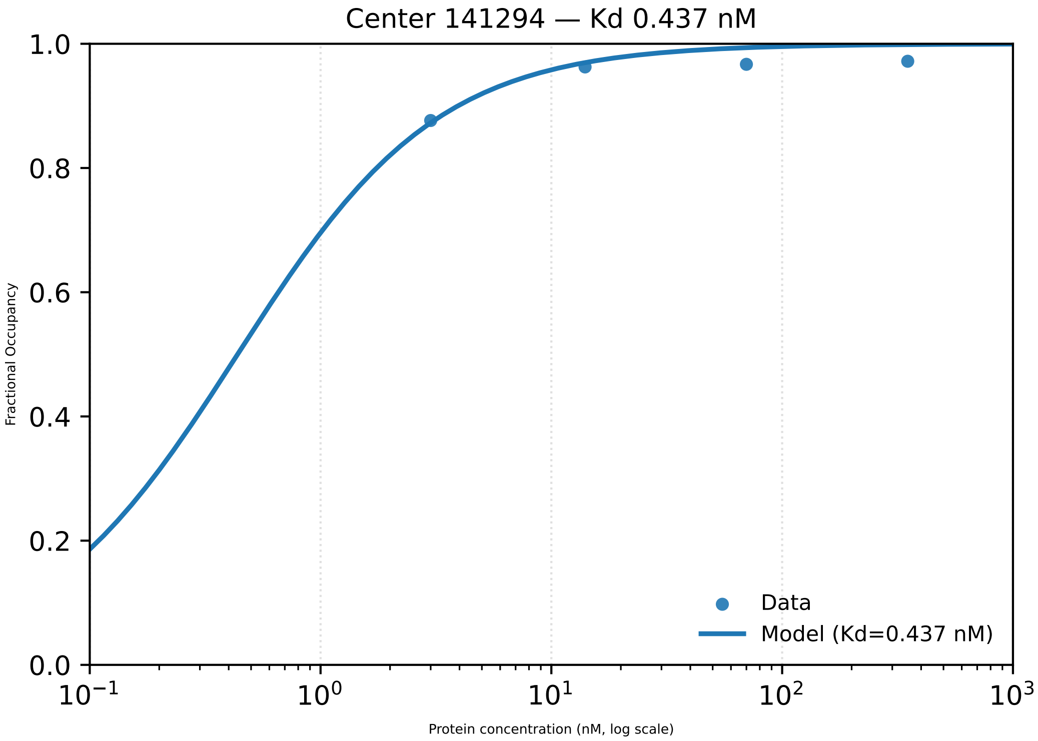

Center 141294 — Kd 0.437 nM
Fractional Occupancy
1.0
0.8
0.6
0.4
0.2
0.0
$10^{-1}$
$10^{0}$
$10^{1}$
$10^{2}$
$10^{3}$
Protein concentration (nM, log scale)
Data
Model (Kd=0.437 nM)

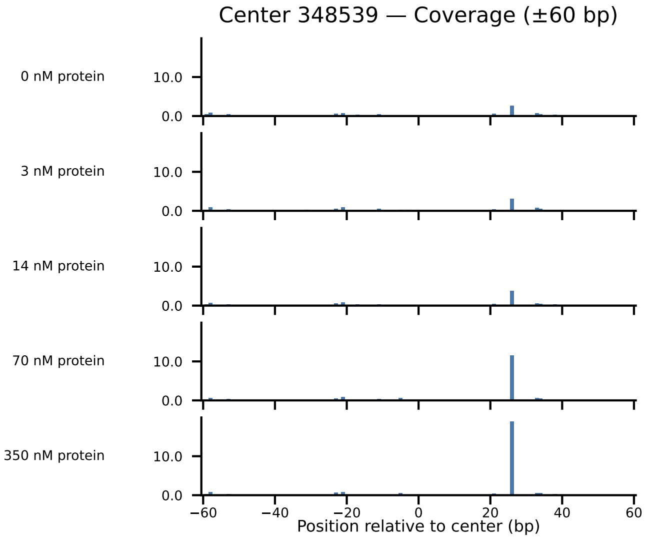

Center 348539 — Coverage (±60 bp)
0 nM protein
3 nM protein
14 nM protein
70 nM protein
350 nM protein
10.0
0.0
−60
−40
−20
0
20
40
60
Position relative to center (bp)

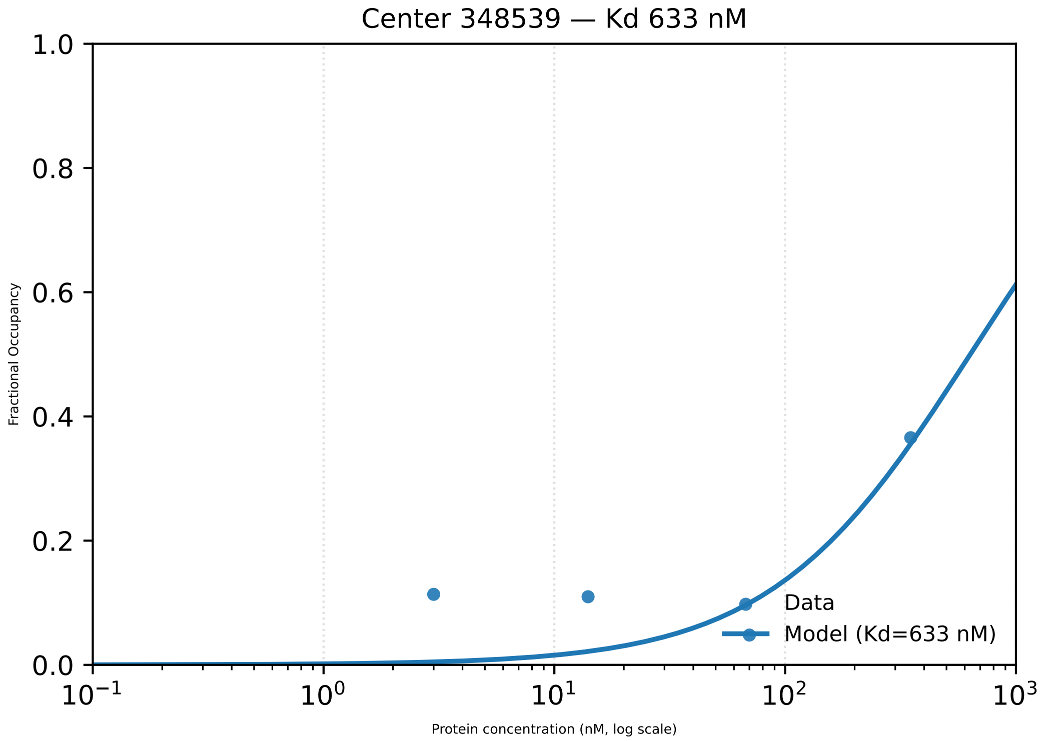

Center 348539 — Kd 633 nM
1.0
0.8
0.6
0.4
0.2
0.0
Fractional Occupancy
$10^{-1}$
$10^{0}$
$10^{1}$
$10^{2}$
$10^{3}$
Protein concentration (nM, log scale)
Data
Model (Kd=633 nM)

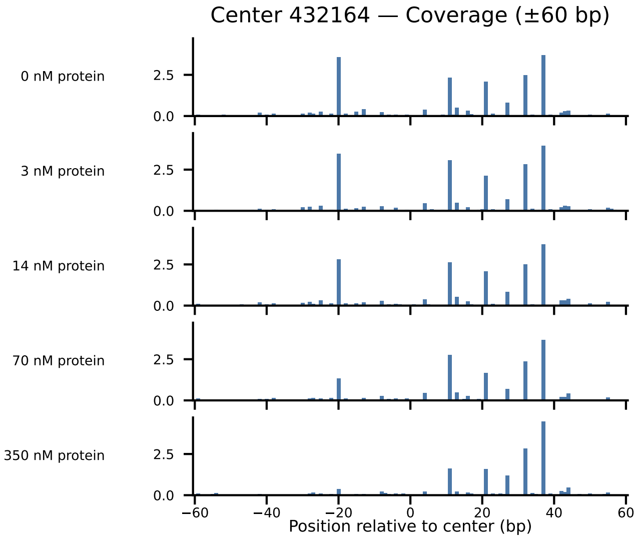
Center 432164 — Coverage (±60 bp)
0 nM protein
3 nM protein
14 nM protein
70 nM protein
350 nM protein
2.5
0.0
−60
−40
−20
0
20
40
60
Position relative to center (bp)

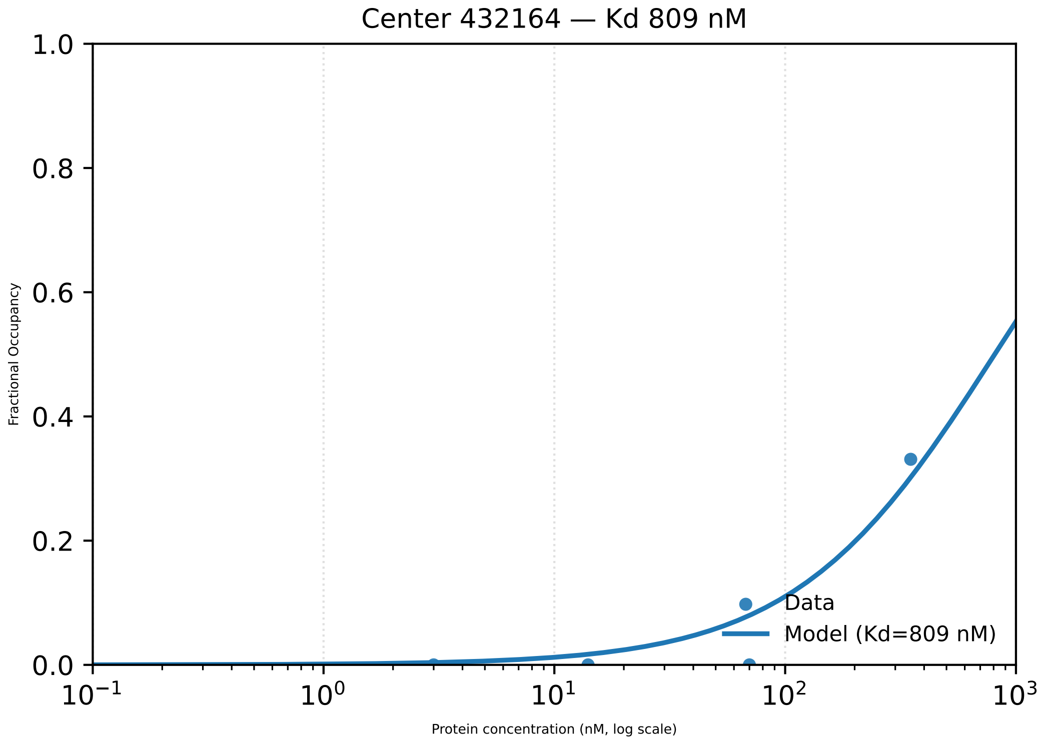
Center 432164 — Kd 809 nM
Fractional Occupancy
1.0
0.8
0.6
0.4
0.2
0.0
$10^{-1}$
$10^{0}$
$10^{1}$
$10^{2}$
$10^{3}$
Protein concentration (nM, log scale)
Data
Model (Kd=809 nM)

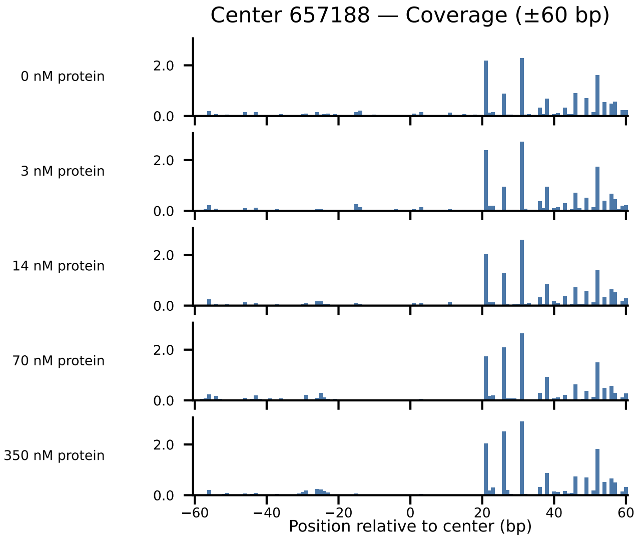

Center 657188 — Coverage (±60 bp)
0 nM protein
3 nM protein
14 nM protein
70 nM protein
350 nM protein
2.0
0.0
−60
−40
−20
0
20
40
60
Position relative to center (bp)

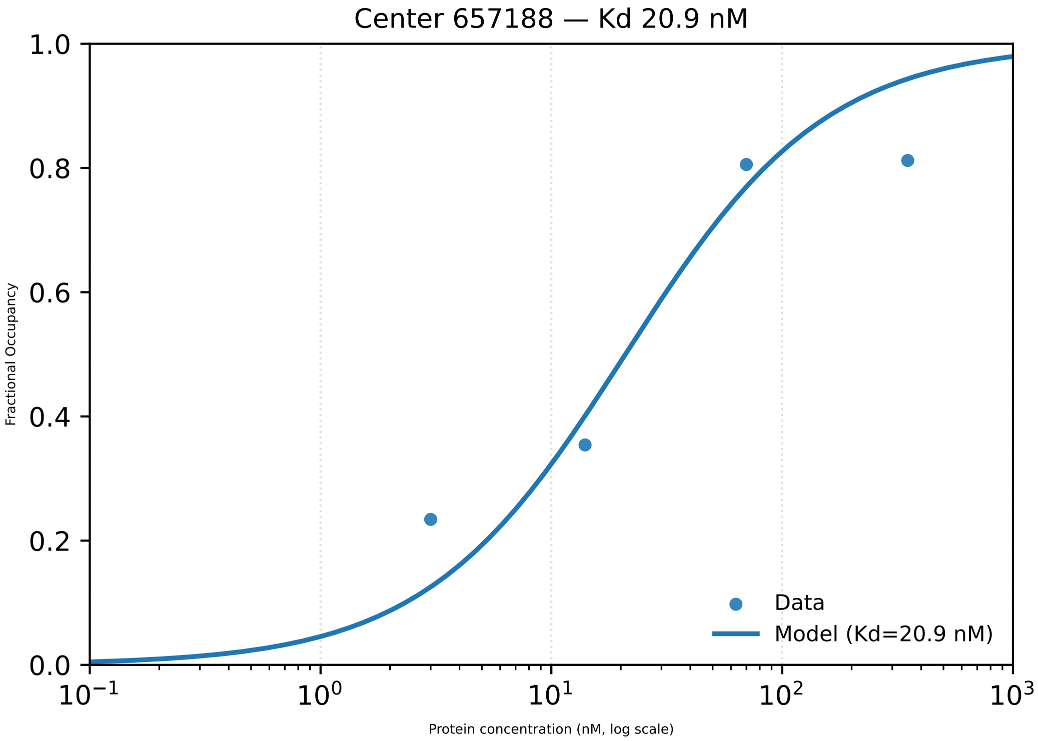

Center 657188 — Kd 20.9 nM
Fractional Occupancy
1.0
0.8
0.6
0.4
0.2
0.0
$10^{-1}$
$10^{0}$
$10^{1}$
$10^{2}$
$10^{3}$
Protein concentration (nM, log scale)
Data
Model (Kd=20.9 nM)

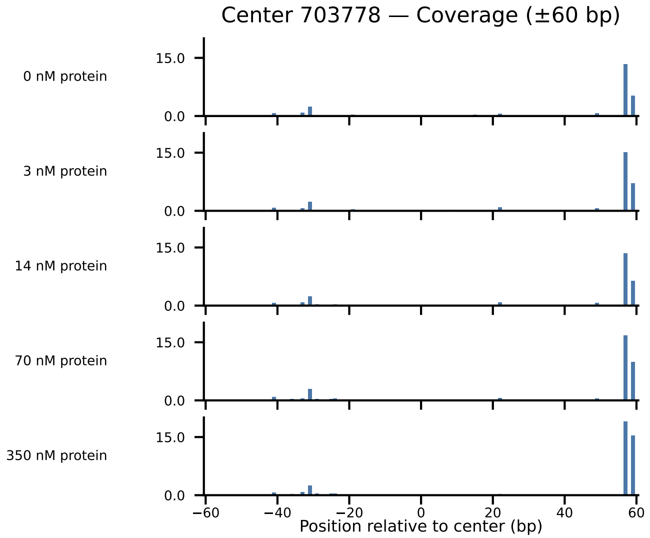

Center 703778 — Coverage (±60 bp)
0 nM protein
3 nM protein
14 nM protein
70 nM protein
350 nM protein
15.0
0.0
−60
−40
−20
0
20
40
60
Position relative to center (bp)

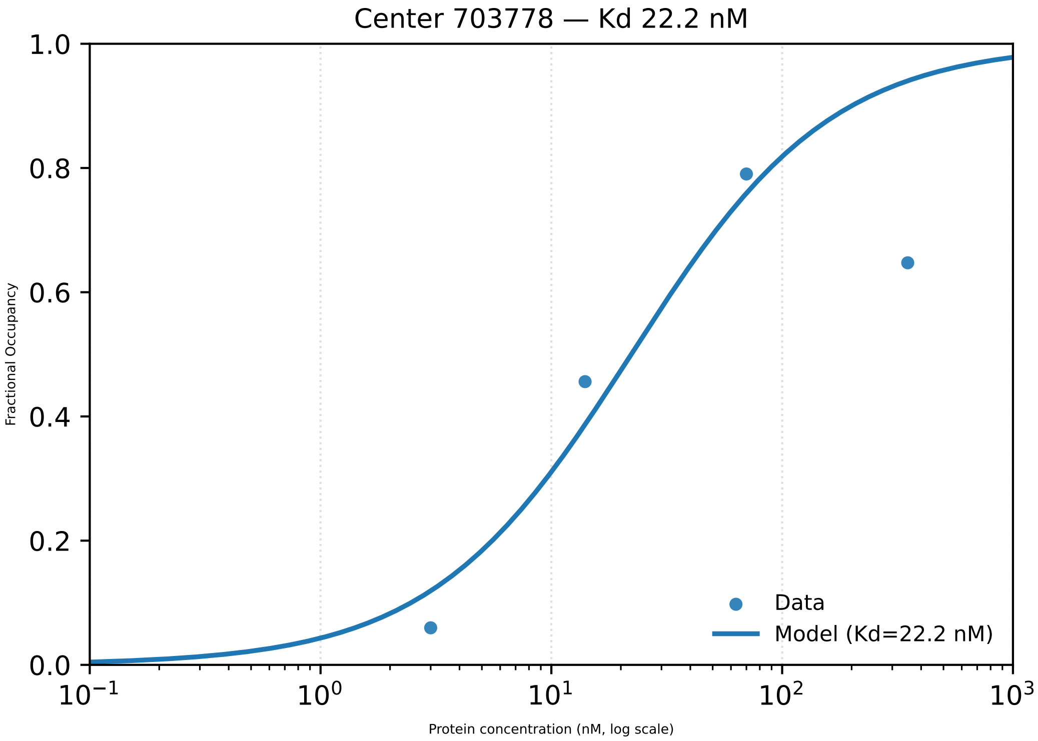

Center 703778 — Kd 22.2 nM
1.0
0.8
0.6
0.4
0.2
0.0
Fractional Occupancy
Data
Model (Kd=22.2 nM)
$10^{-1}$
$10^{0}$
$10^{1}$
$10^{2}$
$10^{3}$
Protein concentration (nM, log scale)

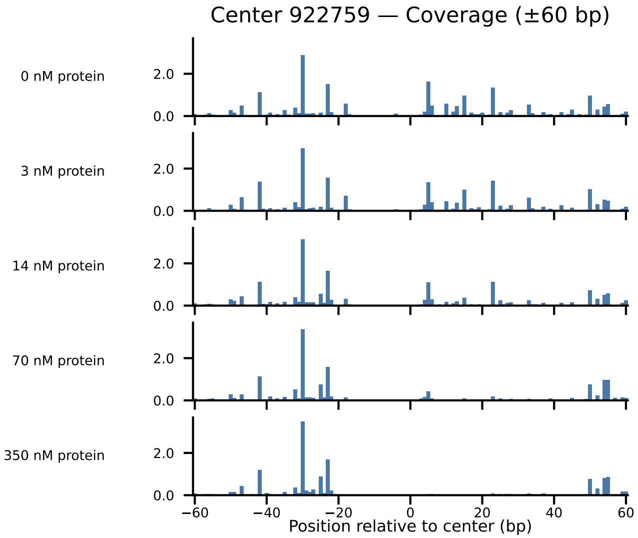

Center 922759 — Coverage (±60 bp)
0 nM protein
3 nM protein
14 nM protein
70 nM protein
350 nM protein
0.0
2.0
−60
−40
−20
0
20
40
60
Position relative to center (bp)

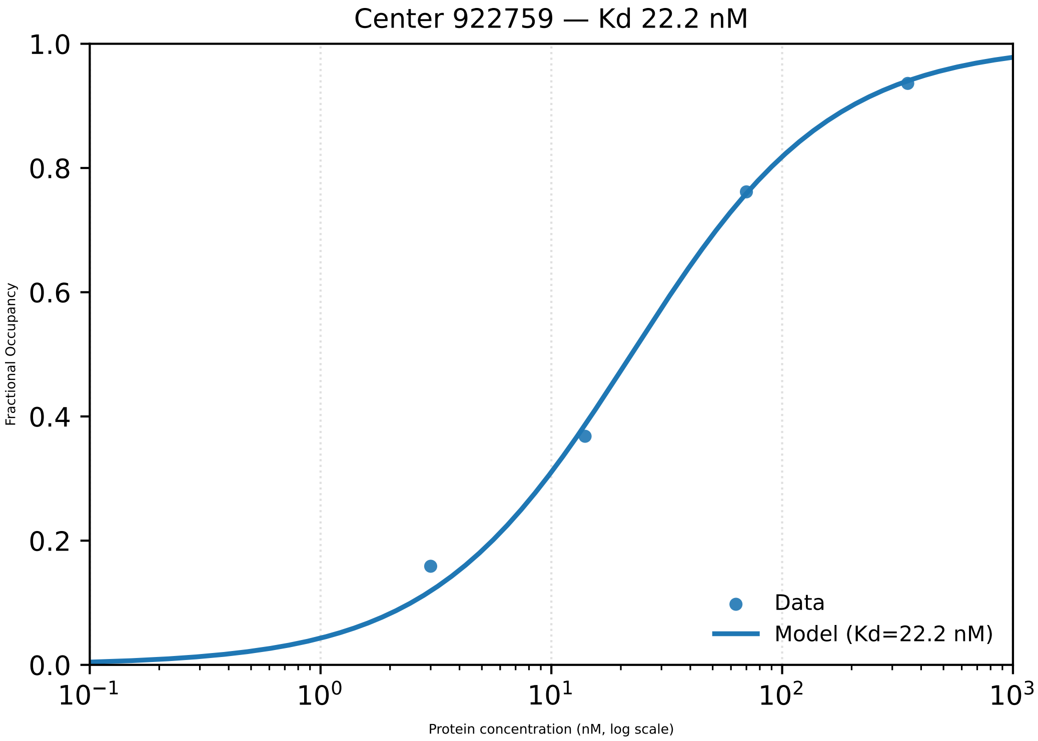

Center 922759 — Kd 22.2 nM
1.0
0.8
0.6
0.4
0.2
0.0
Fractional Occupancy
$10^{-1}$
$10^{0}$
$10^{1}$
$10^{2}$
$10^{3}$
Protein concentration (nM, log scale)
Data
Model (Kd=22.2 nM)

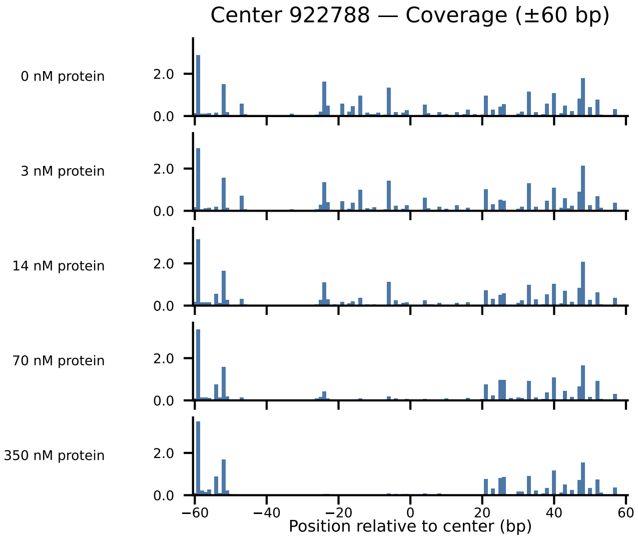
Center 922788 — Coverage (±60 bp)
0 nM protein
3 nM protein
14 nM protein
70 nM protein
350 nM protein
2.0
0.0
−60
−40
−20
0
20
40
60
Position relative to center (bp)

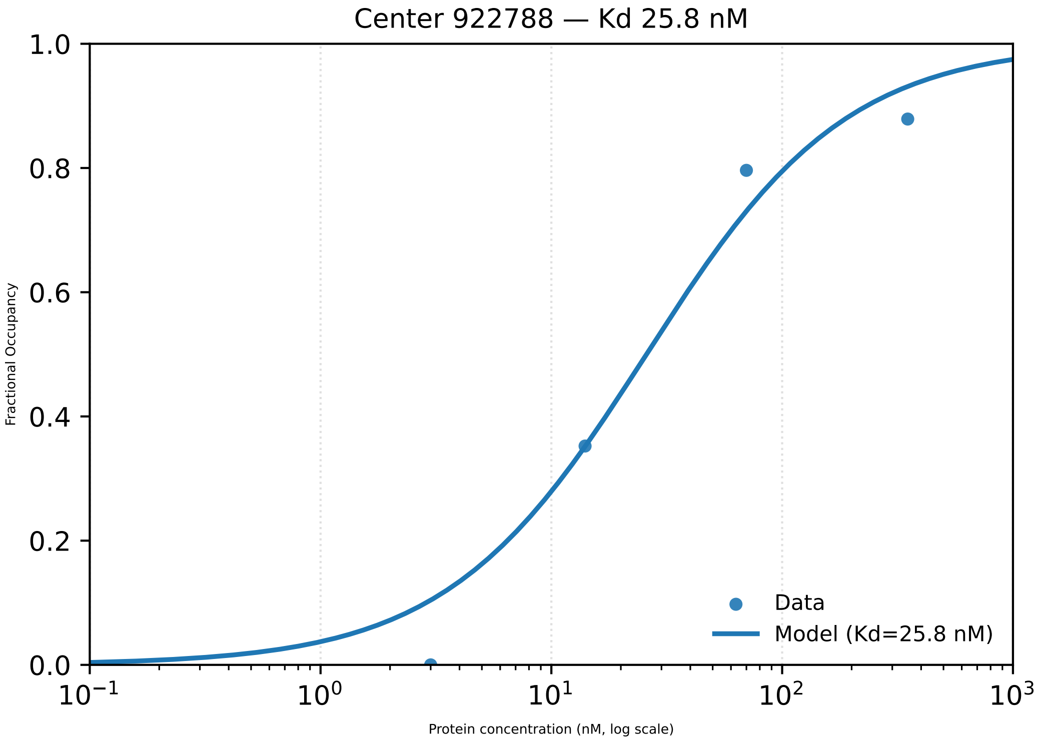

Center 922788 — Kd 25.8 nM
Fractional Occupancy
1.0
0.8
0.6
0.4
0.2
0.0
$10^{-1}$
$10^{0}$
$10^{1}$
$10^{2}$
$10^{3}$
Protein concentration (nM, log scale)
Data
Model (Kd=25.8 nM)

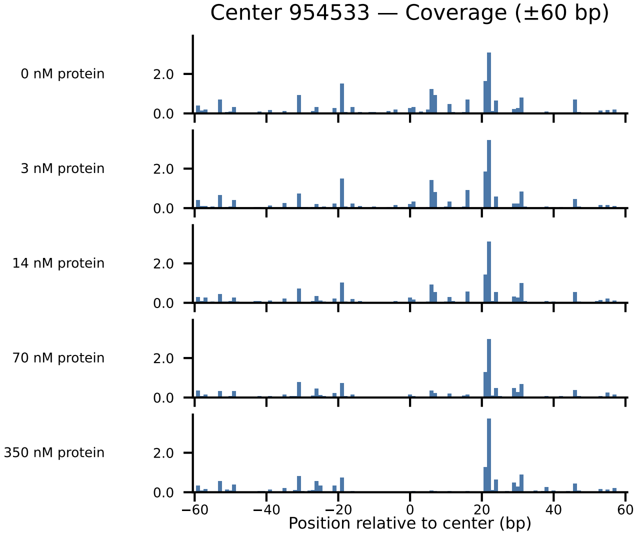

Center 954533 — Coverage (±60 bp)
0 nM protein
3 nM protein
14 nM protein
70 nM protein
350 nM protein
0.0
2.0
−60
−40
−20
0
20
40
60
Position relative to center (bp)

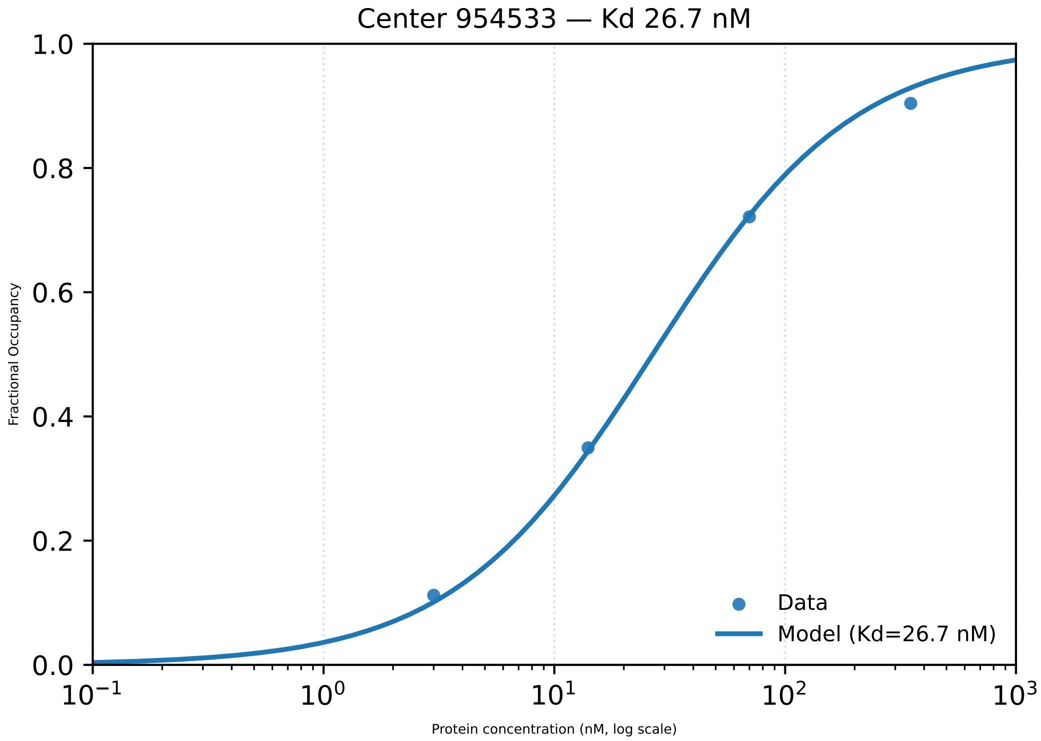

Center 954533 — Kd 26.7 nM
1.0
0.8
0.6
0.4
0.2
0.0
Fractional Occupancy
$10^{-1}$
$10^{0}$
$10^{1}$
$10^{2}$
$10^{3}$
Protein concentration (nM, log scale)
Data
Model (Kd=26.7 nM)

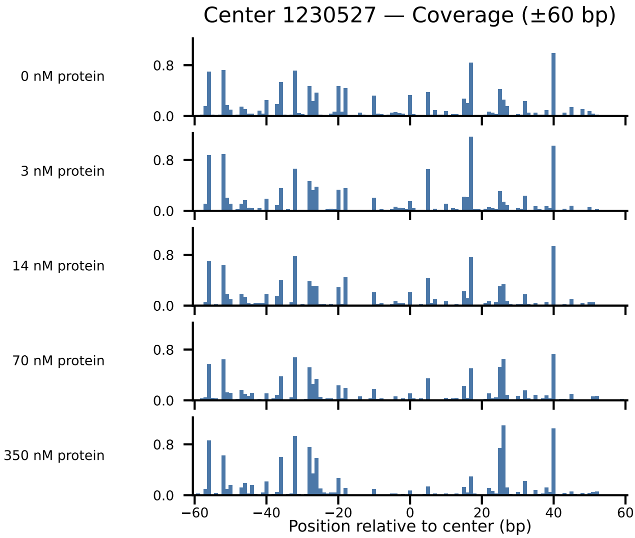

Center 1230527 — Coverage (±60 bp)
0 nM protein
3 nM protein
14 nM protein
70 nM protein
350 nM protein
0.8
0.0
−60
−40
−20
0
20
40
60
Position relative to center (bp)

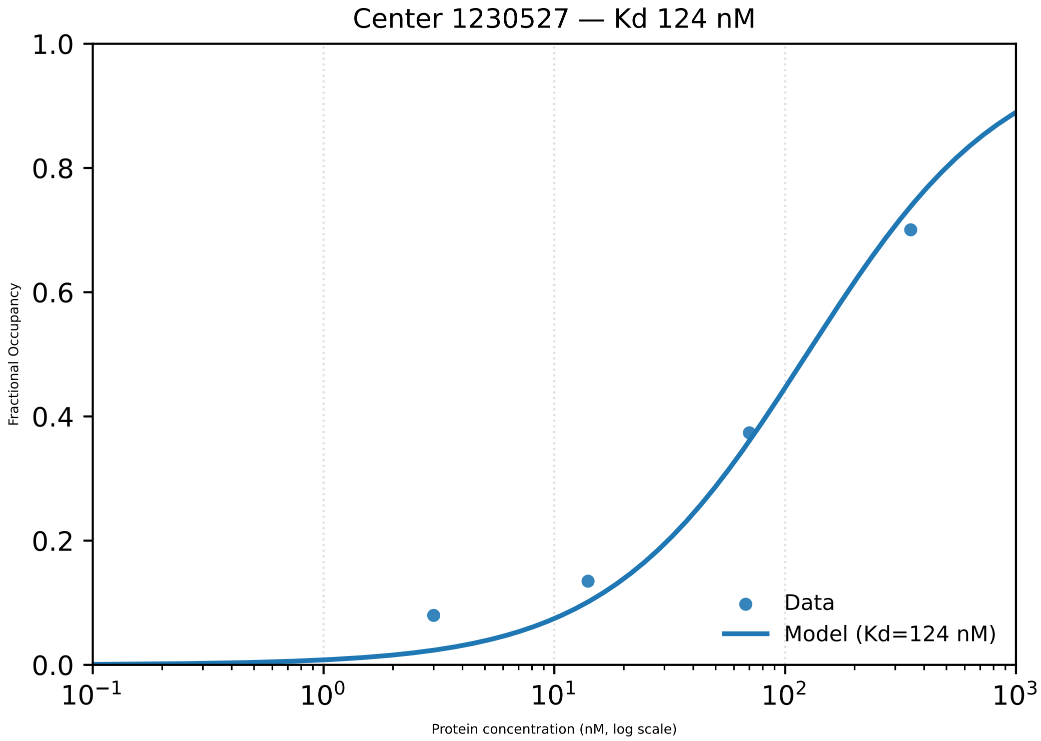

Center 1230527 — Kd 124 nM
Fractional Occupancy
1.0
0.8
0.6
0.4
0.2
0.0
$10^{-1}$
$10^{0}$
$10^{1}$
$10^{2}$
$10^{3}$
Protein concentration (nM, log scale)
Data
Model (Kd=124 nM)

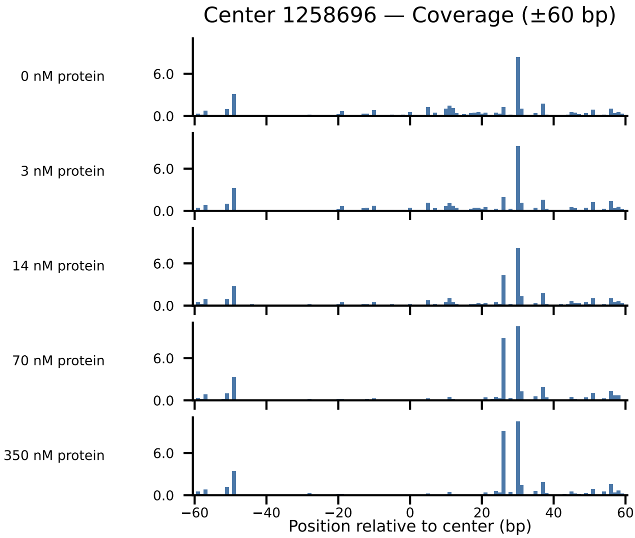

Center 1258696 — Coverage (±60 bp)
0 nM protein
3 nM protein
14 nM protein
70 nM protein
350 nM protein
6.0
0.0
−60
−40
−20
0
20
40
60
Position relative to center (bp)

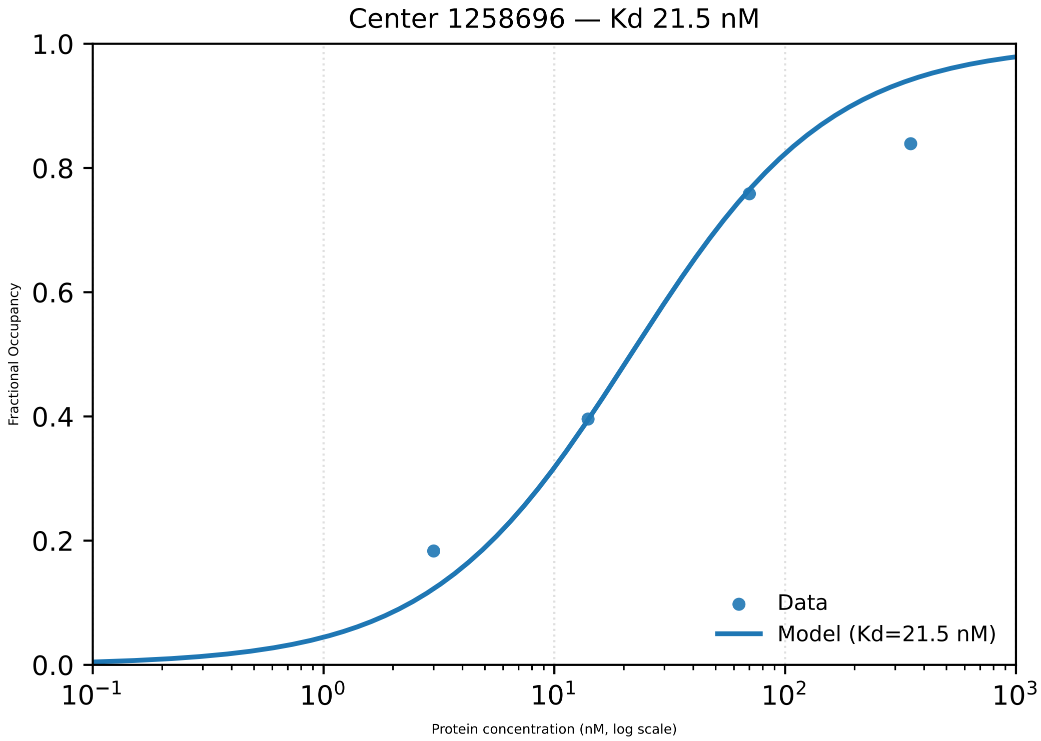

Center 1258696 — Kd 21.5 nM
1.0
0.8
0.6
0.4
0.2
0.0
Fractional Occupancy
$10^{-1}$
$10^{0}$
$10^{1}$
$10^{2}$
$10^{3}$
Protein concentration (nM, log scale)
Data
Model (Kd=21.5 nM)

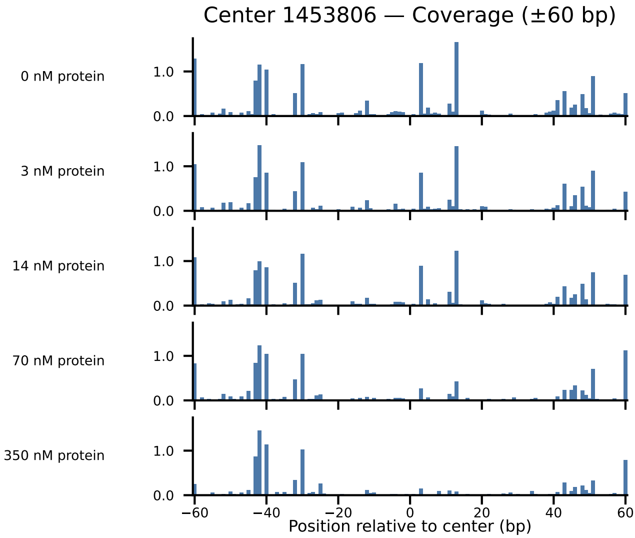

Center 1453806 — Coverage (±60 bp)
0 nM protein
3 nM protein
14 nM protein
70 nM protein
350 nM protein
1.0
0.0
−60
−40
−20
0
20
40
60
Position relative to center (bp)

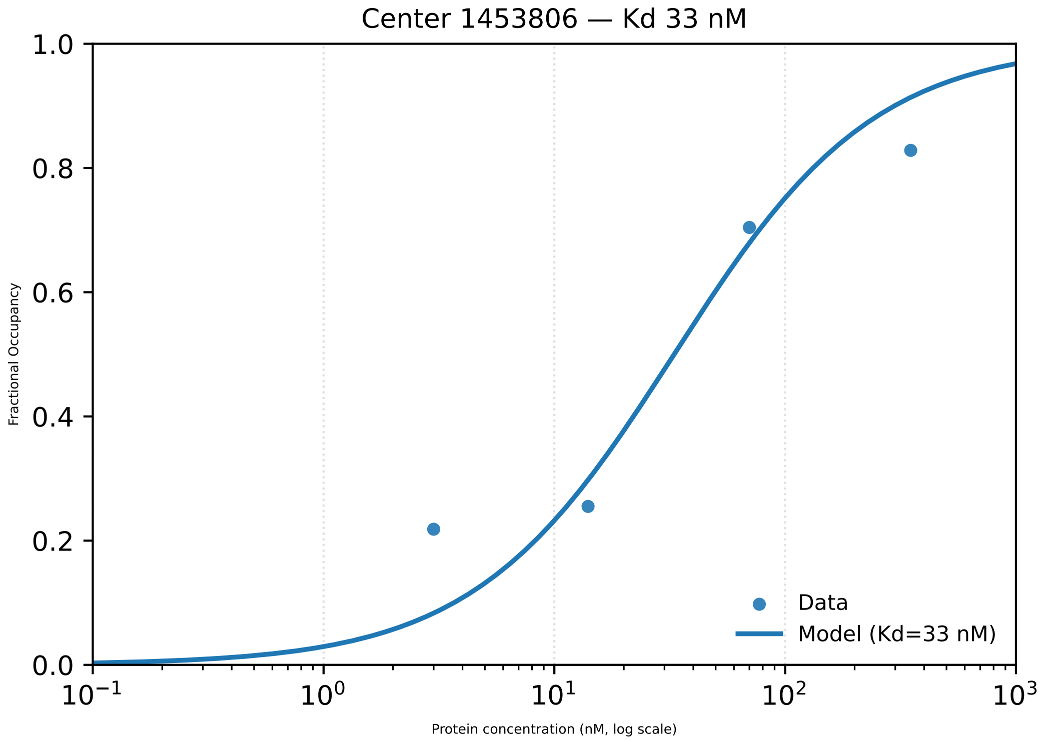

Center 1453806 — Kd 33 nM
1.0
0.8
0.6
0.4
0.2
0.0
Fractional Occupancy
$10^{-1}$
$10^{0}$
$10^{1}$
$10^{2}$
$10^{3}$
Protein concentration (nM, log scale)
Data
Model (Kd=33 nM)

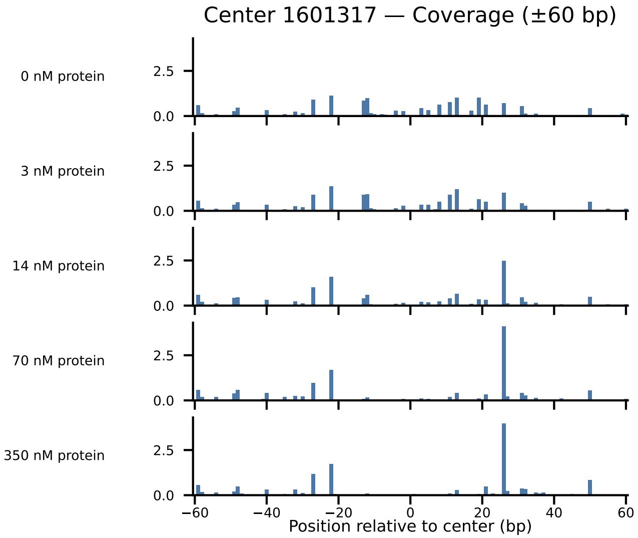

Center 1601317 — Coverage (±60 bp)
0 nM protein
3 nM protein
14 nM protein
70 nM protein
350 nM protein
2.5
0.0
−60
−40
−20
0
20
40
60
Position relative to center (bp)

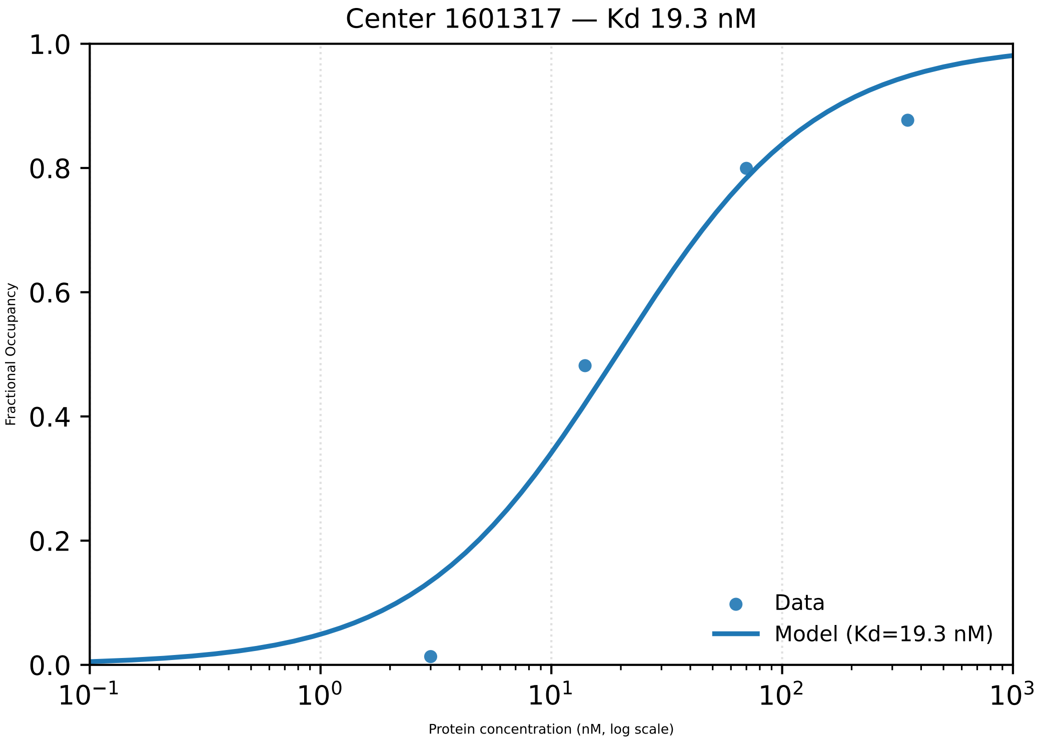

Center 1601317 — Kd 19.3 nM
Fractional Occupancy
Protein concentration (nM, log scale)
1.0
0.8
0.6
0.4
0.2
0.0
$10^{-1}$
$10^{0}$
$10^{1}$
$10^{2}$
$10^{3}$
Data
Model (Kd=19.3 nM)

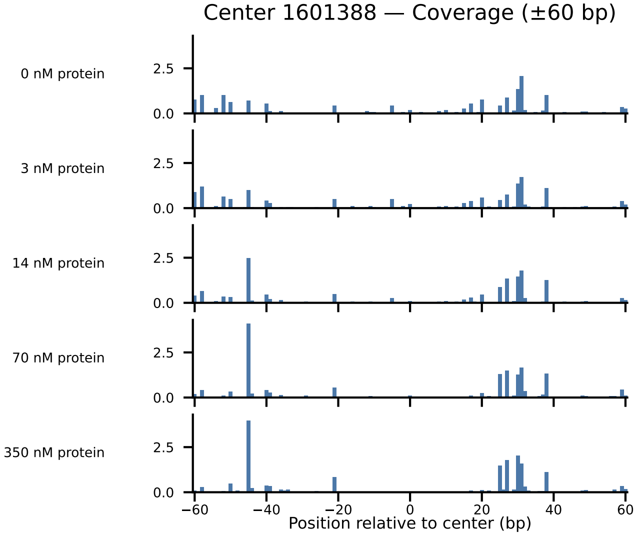

Center 1601388 — Coverage (±60 bp)
0 nM protein
3 nM protein
14 nM protein
70 nM protein
350 nM protein
2.5
0.0
−60
−40
−20
0
20
40
60
Position relative to center (bp)

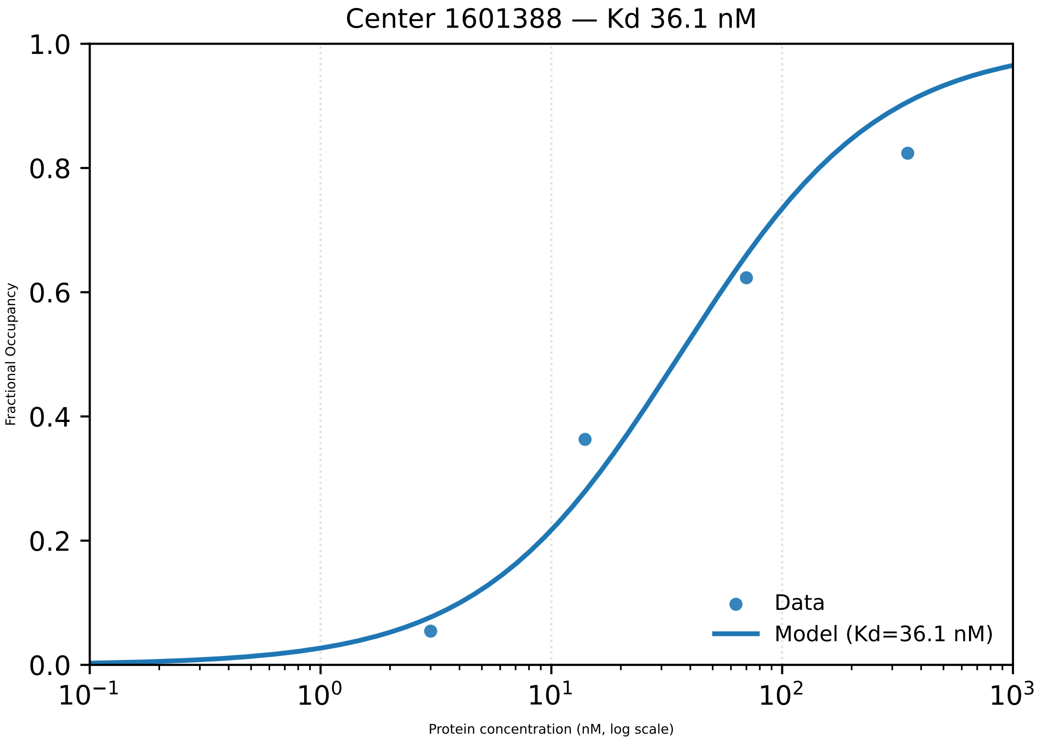

Center 1601388 — Kd 36.1 nM
1.0
0.8
0.6
0.4
0.2
0.0
Fractional Occupancy
Data
Model (Kd=36.1 nM)
$10^{-1}$
$10^{0}$
$10^{1}$
$10^{2}$
$10^{3}$
Protein concentration (nM, log scale)

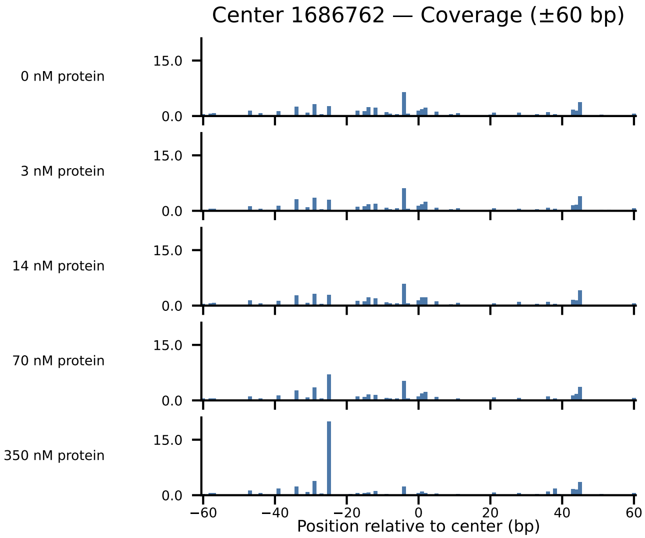

Center 1686762 — Coverage (±60 bp)
0 nM protein
3 nM protein
14 nM protein
70 nM protein
350 nM protein
15.0
0.0
−60
−40
−20
0
20
40
60
Position relative to center (bp)

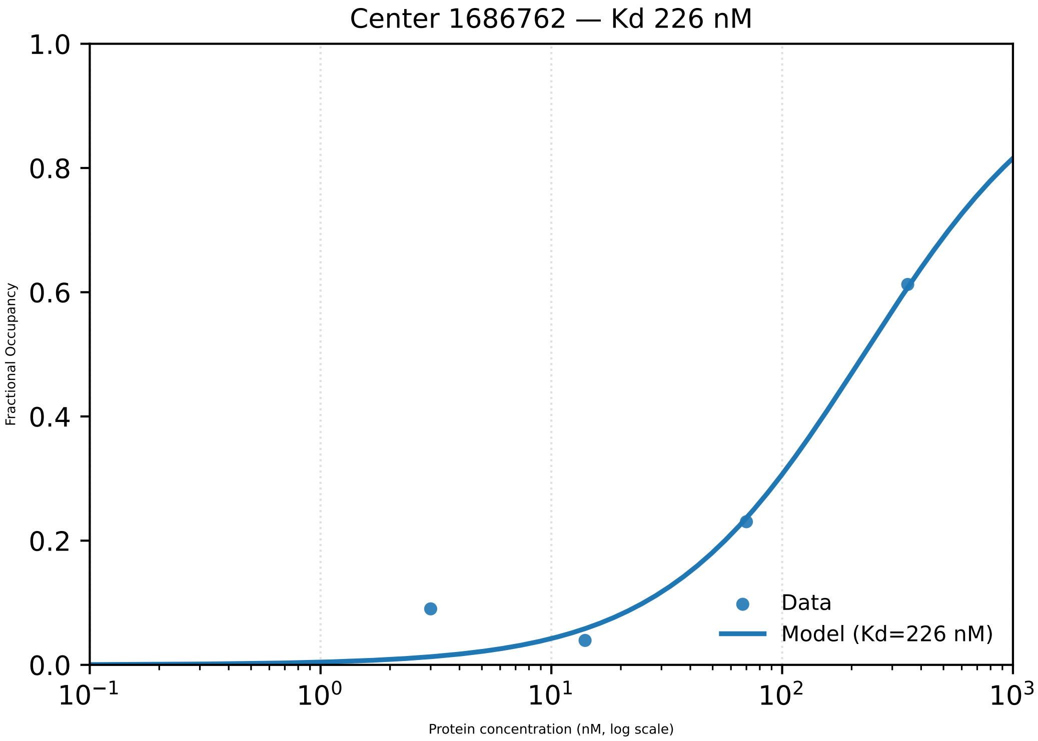

Center 1686762 — Kd 226 nM
1.0
0.8
0.6
0.4
0.2
0.0
Fractional Occupancy
$10^{-1}$
$10^{0}$
$10^{1}$
$10^{2}$
$10^{3}$
Protein concentration (nM, log scale)
Data
Model (Kd=226 nM)

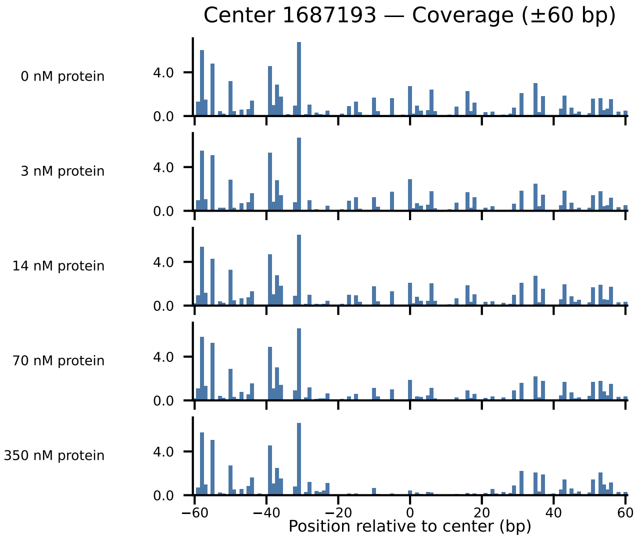

Center 1687193 — Coverage (±60 bp)
0 nM protein
3 nM protein
14 nM protein
70 nM protein
350 nM protein
4.0
0.0
−60
−40
−20
0
20
40
60
Position relative to center (bp)

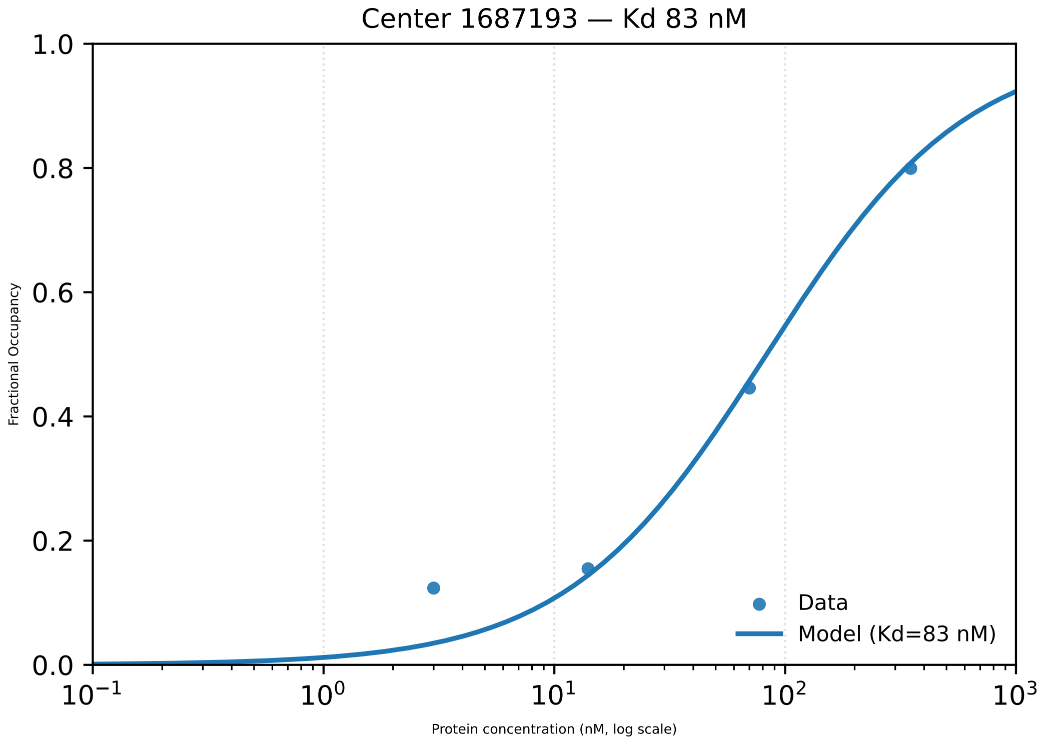

Center 1687193 — Kd 83 nM
1.0
0.8
0.6
0.4
0.2
0.0
Fractional Occupancy
$10^{-1}$
$10^{0}$
$10^{1}$
$10^{2}$
$10^{3}$
Protein concentration (nM, log scale)
Data
Model (Kd=83 nM)

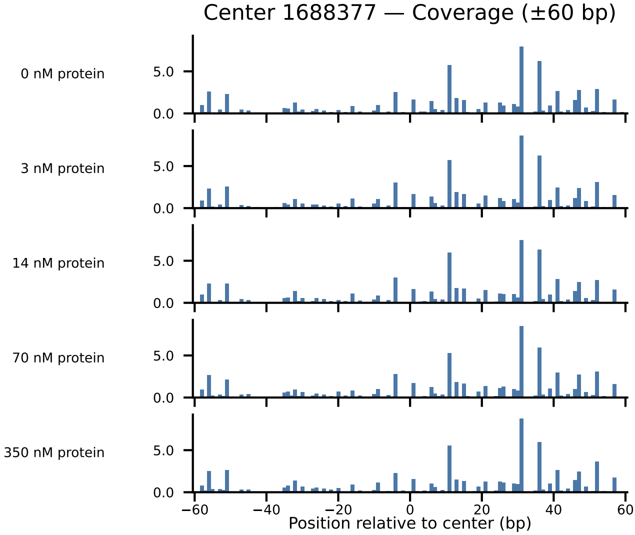
Center 1688377 — Coverage (±60 bp)
0 nM protein
3 nM protein
14 nM protein
70 nM protein
350 nM protein
5.0
0.0
−60
−40
−20
0
20
40
60
Position relative to center (bp)

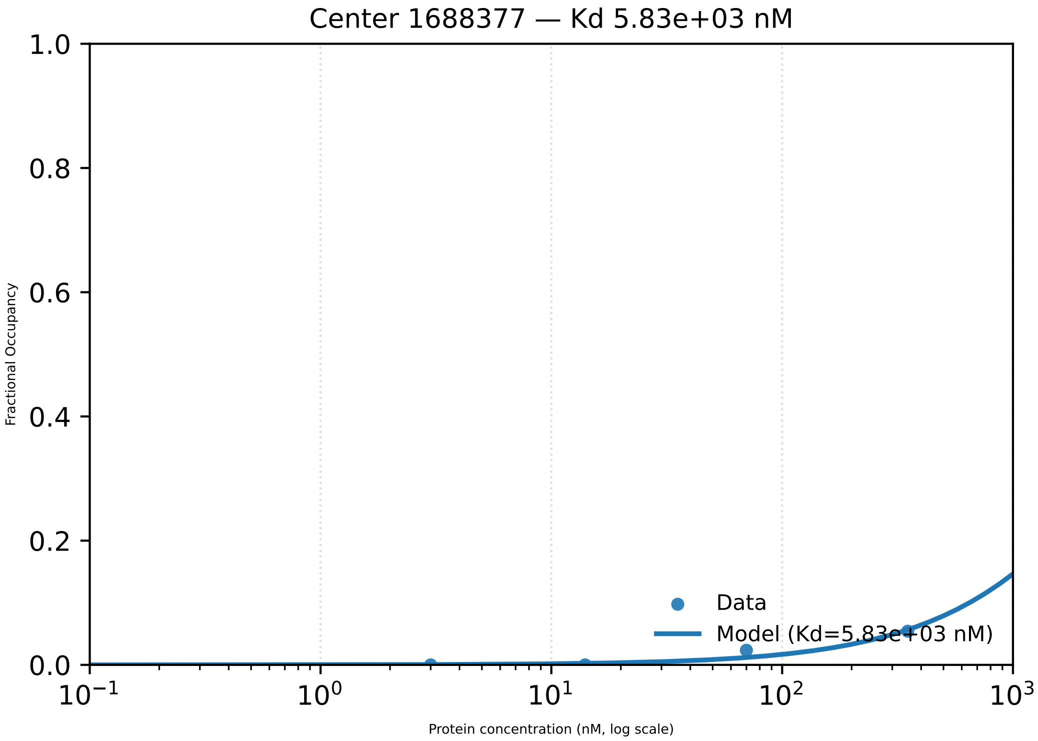

Center 1688377 — Kd 5.83e+03 nM
1.0
0.8
0.6
0.4
0.2
0.0
Fractional Occupancy
Data
Model (Kd=5.83e+03 nM)
$10^{-1}$
$10^{0}$
$10^{1}$
$10^{2}$
$10^{3}$
Protein concentration (nM, log scale)

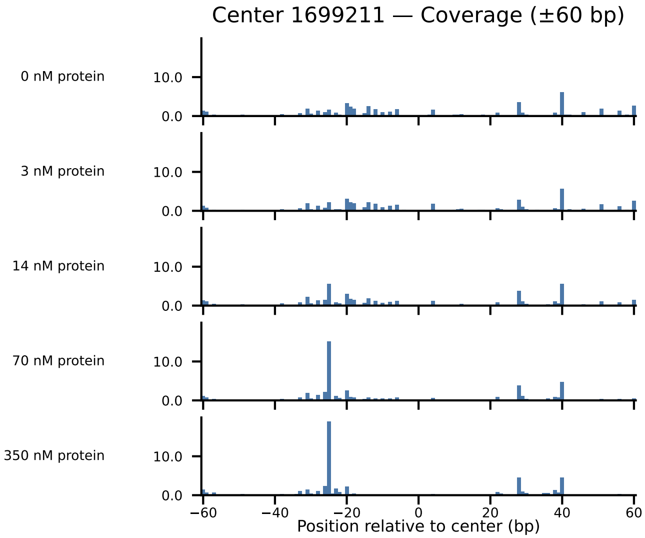

Center 1699211 — Coverage (±60 bp)
0 nM protein
3 nM protein
14 nM protein
70 nM protein
350 nM protein
10.0
0.0
−60
−40
−20
0
20
40
60
Position relative to center (bp)

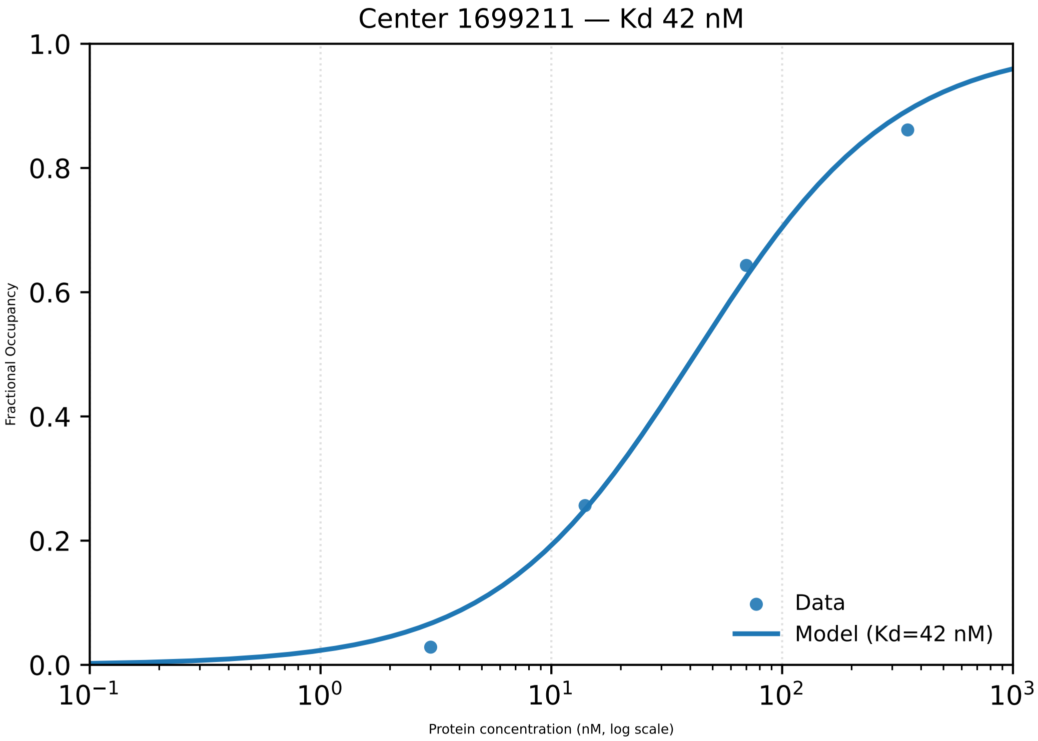
Center 1699211 — Kd 42 nM
Fractional Occupancy
1.0
0.8
0.6
0.4
0.2
0.0
$10^{-1}$
$10^{0}$
$10^{1}$
$10^{2}$
$10^{3}$
Protein concentration (nM, log scale)
Data
Model (Kd=42 nM)

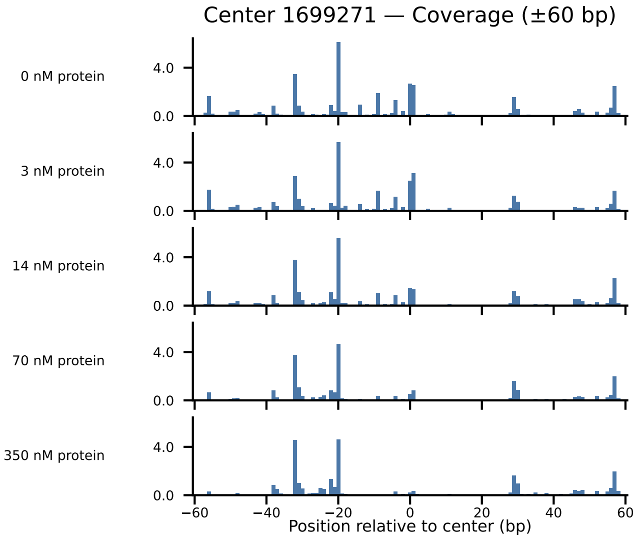

Center 1699271 — Coverage (±60 bp)
0 nM protein
3 nM protein
14 nM protein
70 nM protein
350 nM protein
4.0
0.0
−60
−40
−20
0
20
40
60
Position relative to center (bp)

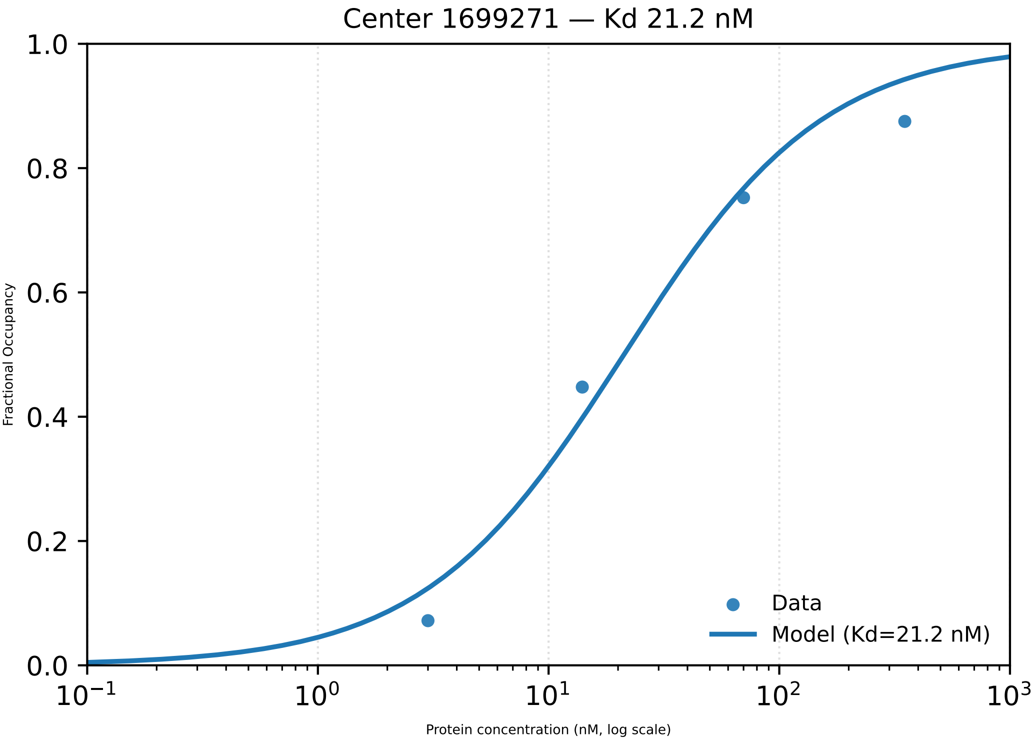

Center 1699271 — Kd 21.2 nM
Fractional Occupancy
1.0
0.8
0.6
0.4
0.2
0.0
$10^{-1}$
$10^{0}$
$10^{1}$
$10^{2}$
$10^{3}$
Protein concentration (nM, log scale)
Data
Model (Kd=21.2 nM)

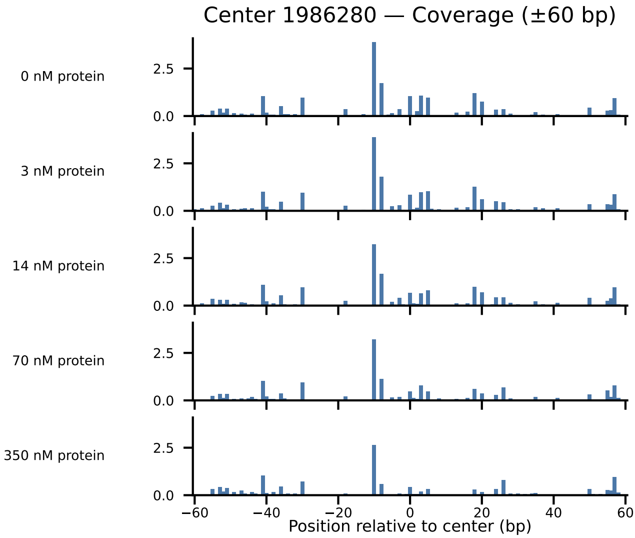

Center 1986280 — Coverage (±60 bp)
0 nM protein
3 nM protein
14 nM protein
70 nM protein
350 nM protein
2.5
0.0
−60
−40
−20
0
20
40
60
Position relative to center (bp)

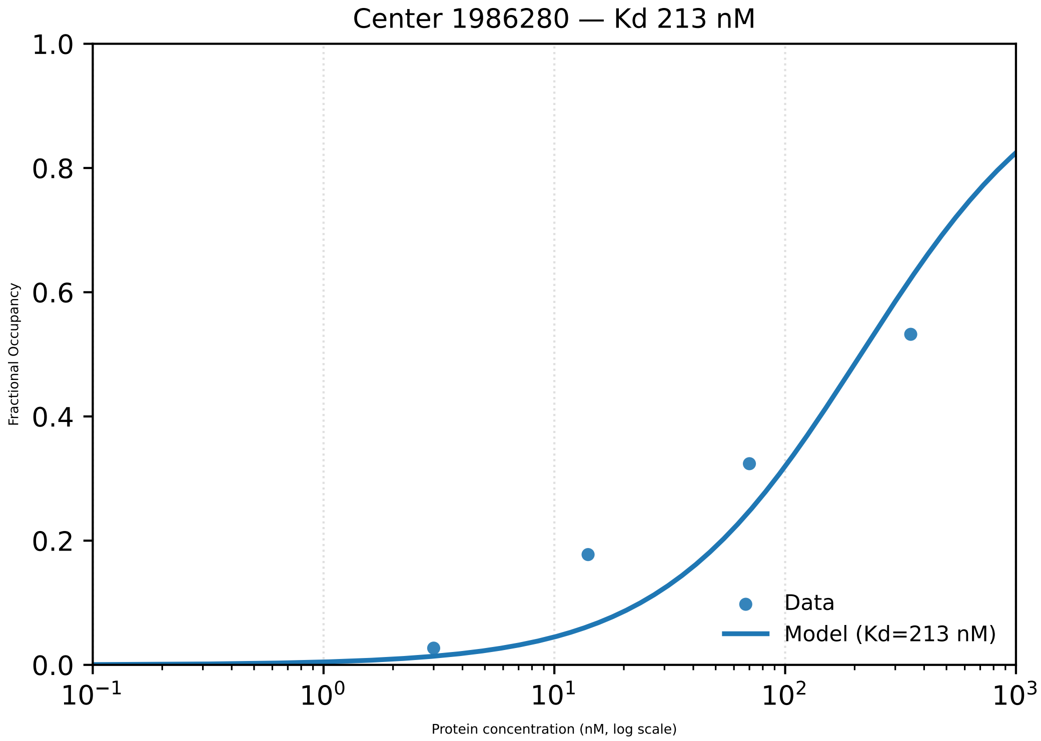

Center 1986280 — Kd 213 nM
Fractional Occupancy
1.0
0.8
0.6
0.4
0.2
0.0
$10^{-1}$
$10^{0}$
$10^{1}$
$10^{2}$
$10^{3}$
Protein concentration (nM, log scale)
Data
Model (Kd=213 nM)

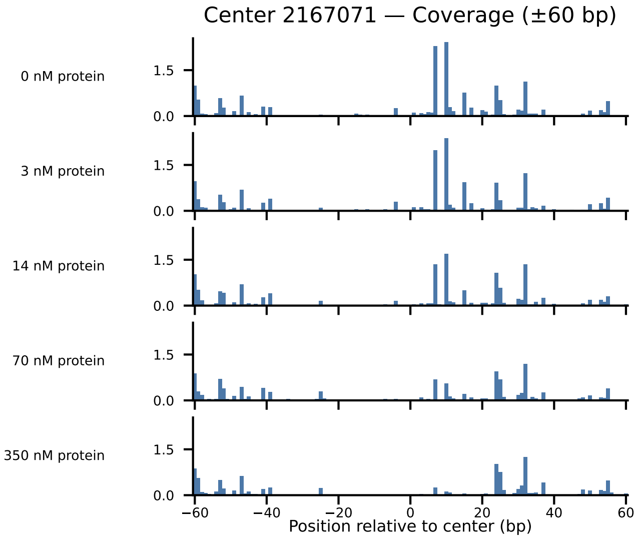

Center 2167071 — Coverage (±60 bp)
0 nM protein
3 nM protein
14 nM protein
70 nM protein
350 nM protein
1.5
0.0
−60
−40
−20
0
20
40
60
Position relative to center (bp)

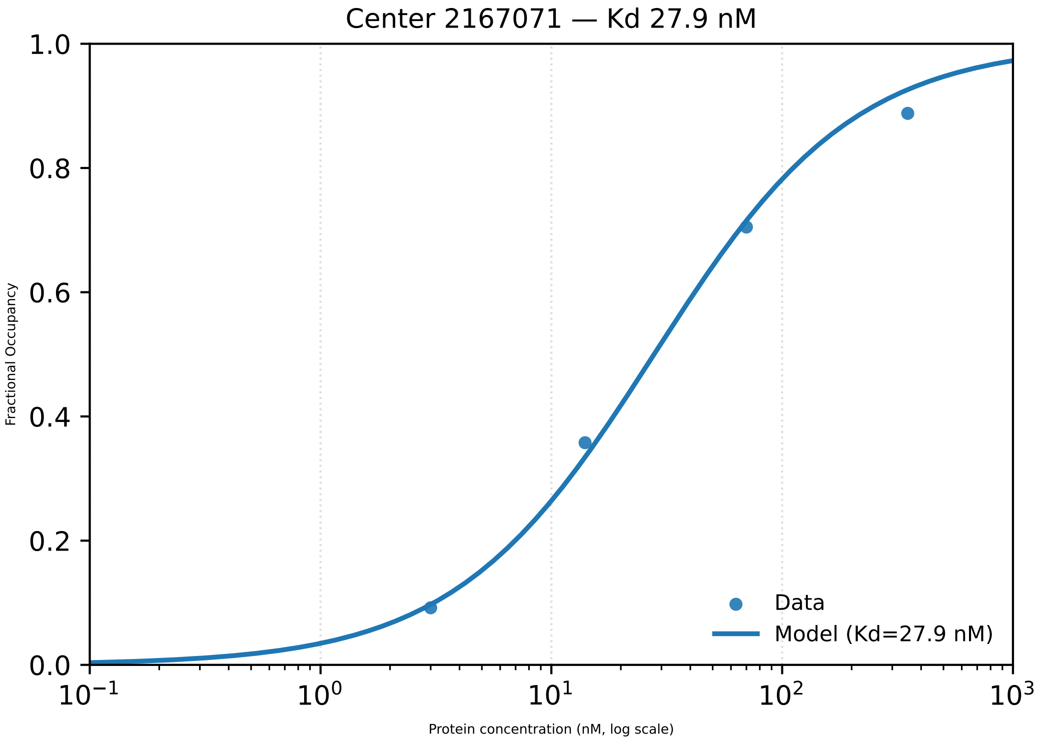
Center 2167071 — Kd 27.9 nM
1.0
0.8
0.6
0.4
0.2
0.0
Fractional Occupancy
$10^{-1}$
$10^{0}$
$10^{1}$
$10^{2}$
$10^{3}$
Protein concentration (nM, log scale)
Data
Model (Kd=27.9 nM)

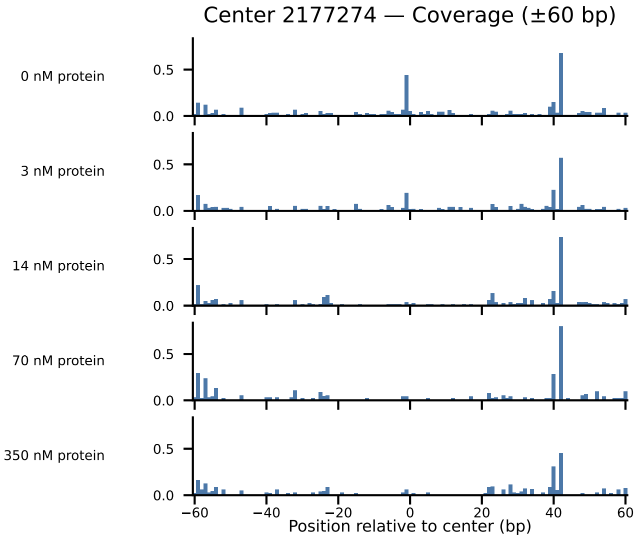

Center 2177274 — Coverage (±60 bp)
0 nM protein
3 nM protein
14 nM protein
70 nM protein
350 nM protein
0.5
0.0
−60
−40
−20
0
20
40
60
Position relative to center (bp)

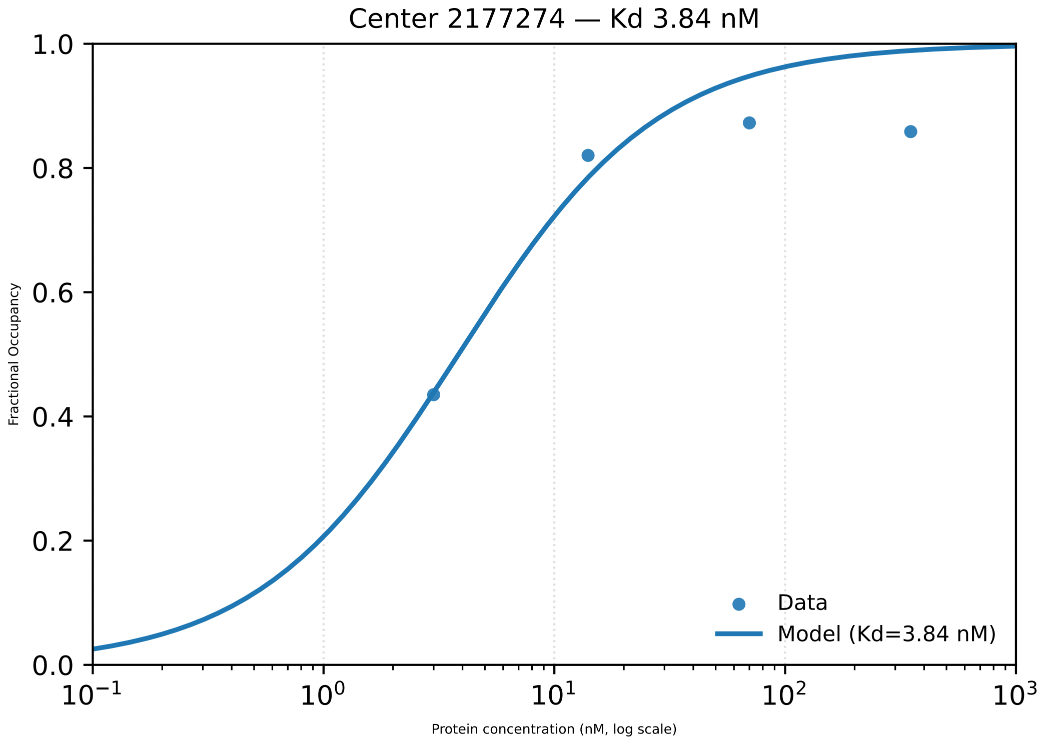

Center 2177274 — Kd 3.84 nM
1.0
0.8
0.6
0.4
0.2
0.0
Fractional Occupancy
$10^{-1}$
$10^{0}$
$10^{1}$
$10^{2}$
$10^{3}$
Protein concentration (nM, log scale)
Data
Model (Kd=3.84 nM)

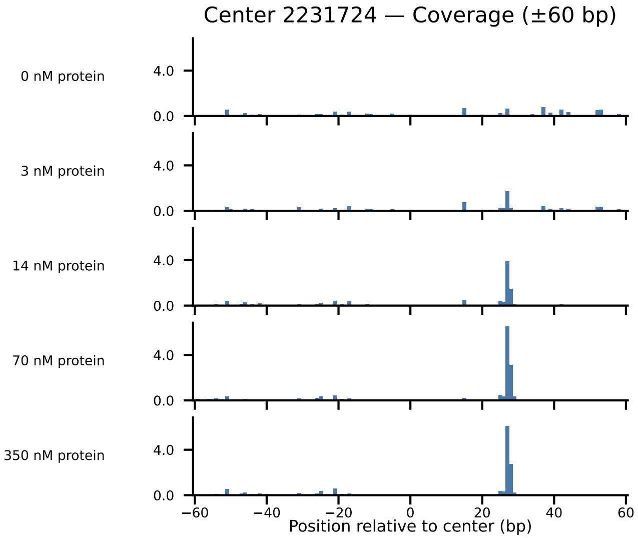
Center 2231724 — Coverage (±60 bp)
0 nM protein
3 nM protein
14 nM protein
70 nM protein
350 nM protein
4.0
0.0
−60
−40
−20
0
20
40
60
Position relative to center (bp)

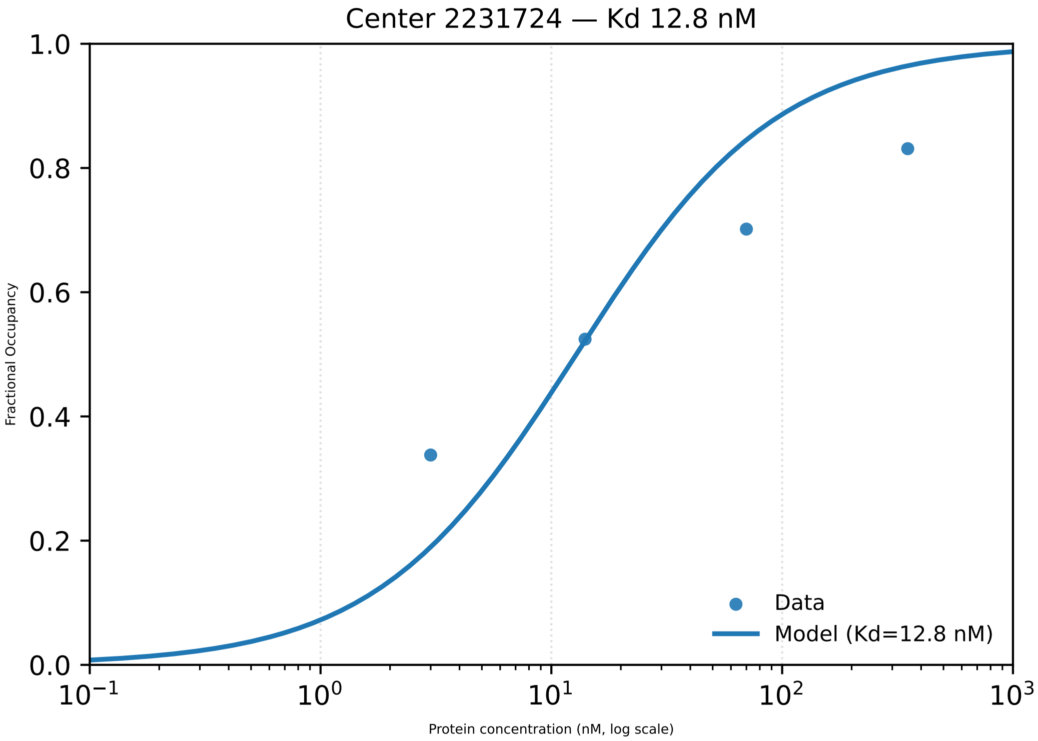

Center 2231724 — Kd 12.8 nM
1.0
0.8
0.6
0.4
0.2
0.0
Fractional Occupancy
$10^{-1}$
$10^{0}$
$10^{1}$
$10^{2}$
$10^{3}$
Protein concentration (nM, log scale)
Data
Model (Kd=12.8 nM)

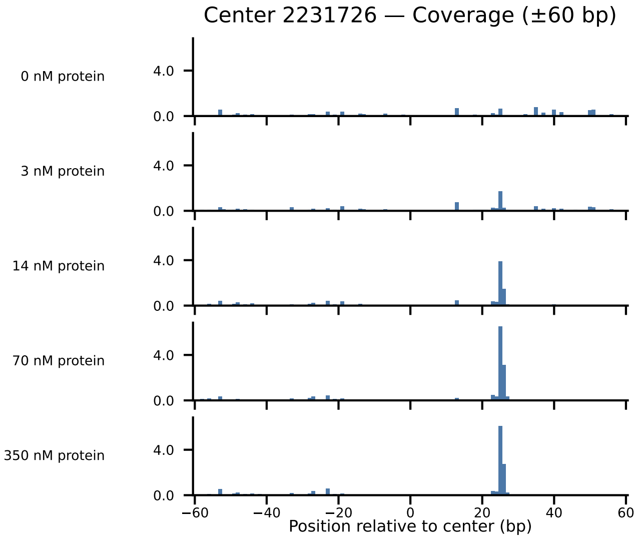
Center 2231726 — Coverage (±60 bp)
0 nM protein
3 nM protein
14 nM protein
70 nM protein
350 nM protein
4.0
0.0
−60
−40
−20
0
20
40
60
Position relative to center (bp)

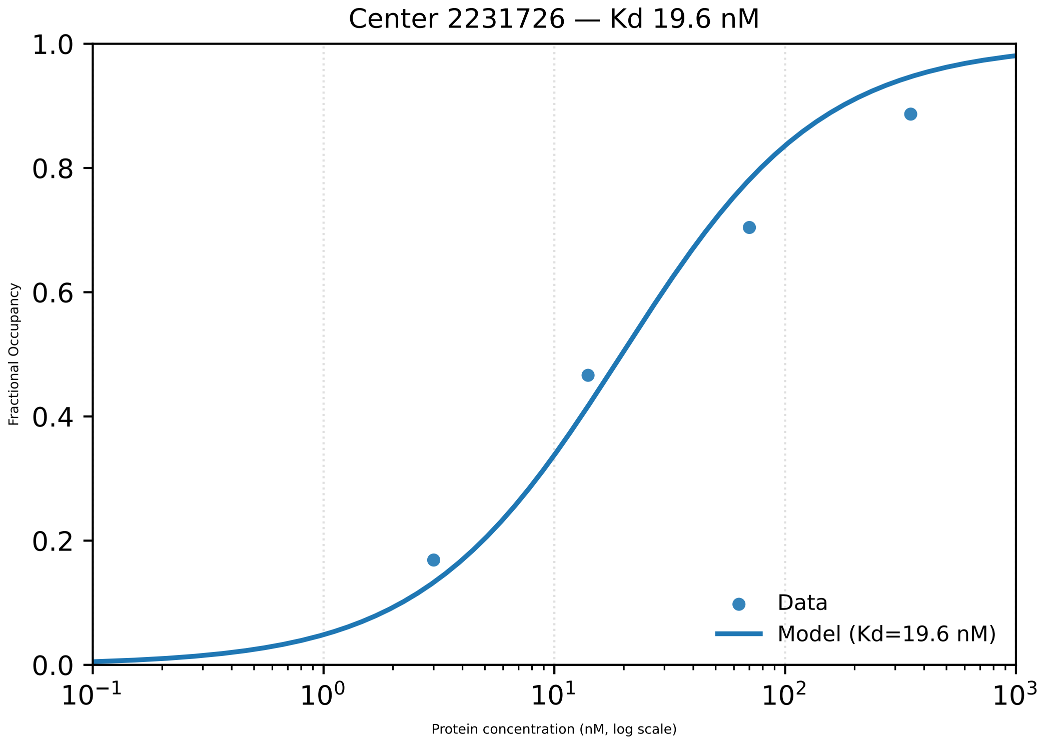

Center 2231726 — Kd 19.6 nM
1.0
0.8
0.6
0.4
0.2
0.0
Fractional Occupancy
$10^{-1}$
$10^{0}$
$10^{1}$
$10^{2}$
$10^{3}$
Protein concentration (nM, log scale)
Data
Model (Kd=19.6 nM)

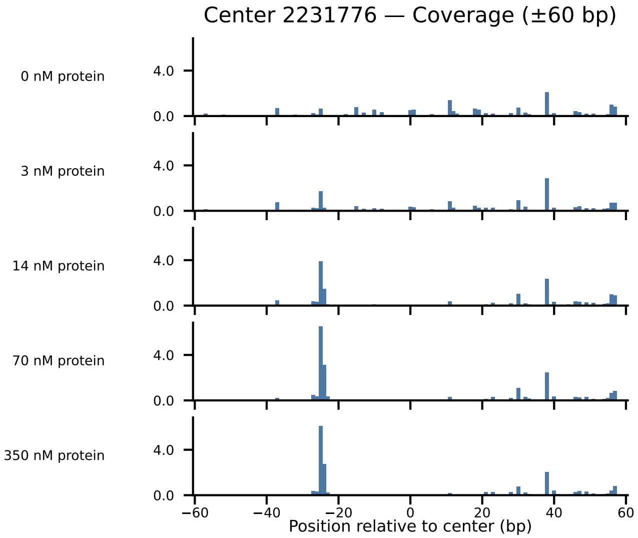

Center 2231776 — Coverage (±60 bp)
0 nM protein
3 nM protein
14 nM protein
70 nM protein
350 nM protein
4.0
0.0
−60
−40
−20
0
20
40
60
Position relative to center (bp)

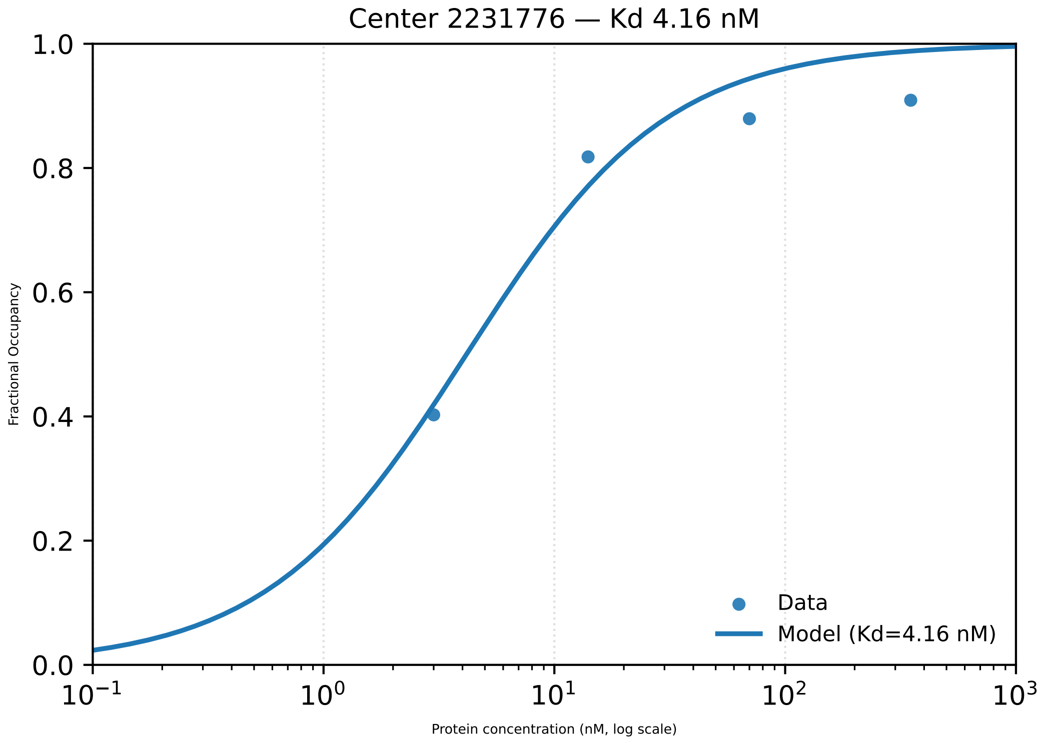

Center 2231776 — Kd 4.16 nM
1.0
0.8
0.6
0.4
0.2
0.0
Fractional Occupancy
$10^{-1}$
$10^{0}$
$10^{1}$
$10^{2}$
$10^{3}$
Protein concentration (nM, log scale)
Data
Model (Kd=4.16 nM)

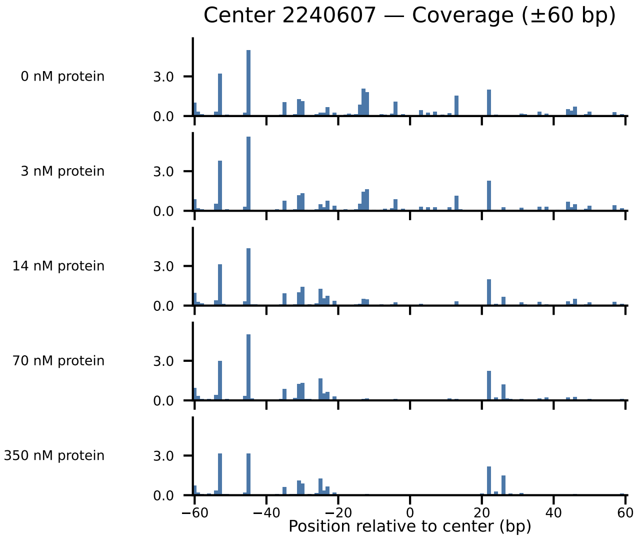

Center 2240607 — Coverage (±60 bp)
0 nM protein
3 nM protein
14 nM protein
70 nM protein
350 nM protein
3.0
0.0
−60
−40
−20
0
20
40
60
Position relative to center (bp)

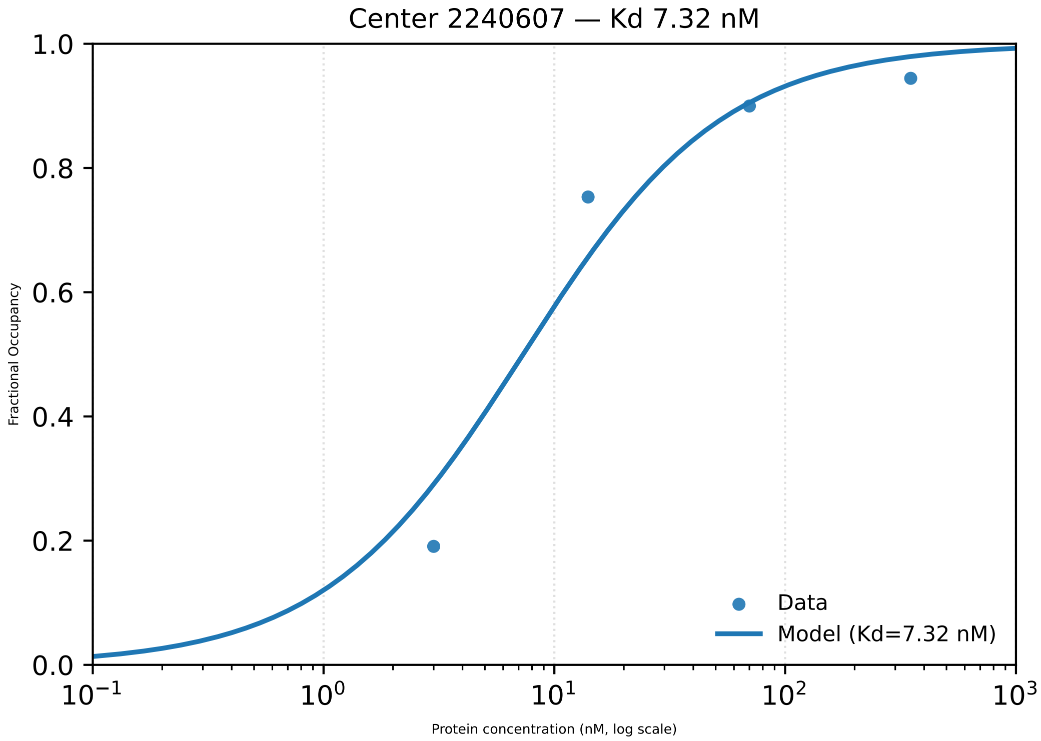

Center 2240607 — Kd 7.32 nM
Fractional Occupancy
1.0
0.8
0.6
0.4
0.2
0.0
$10^{-1}$
$10^{0}$
$10^{1}$
$10^{2}$
$10^{3}$
Protein concentration (nM, log scale)
Data
Model (Kd=7.32 nM)

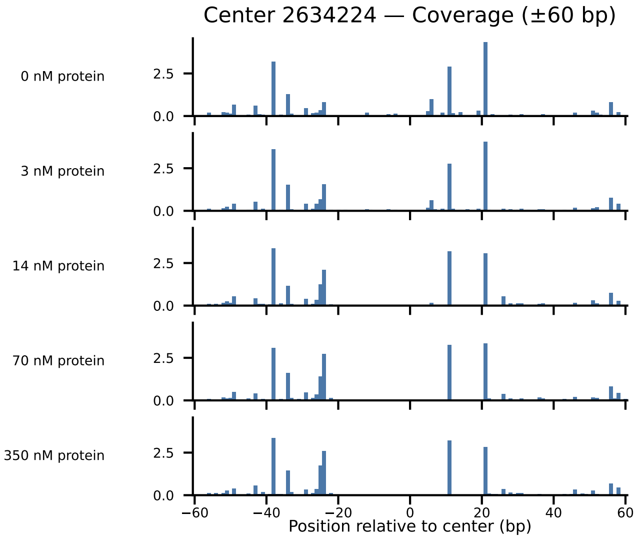

Center 2634224 — Coverage (±60 bp)
0 nM protein
3 nM protein
14 nM protein
70 nM protein
350 nM protein
2.5
0.0
−60
−40
−20
0
20
40
60
Position relative to center (bp)

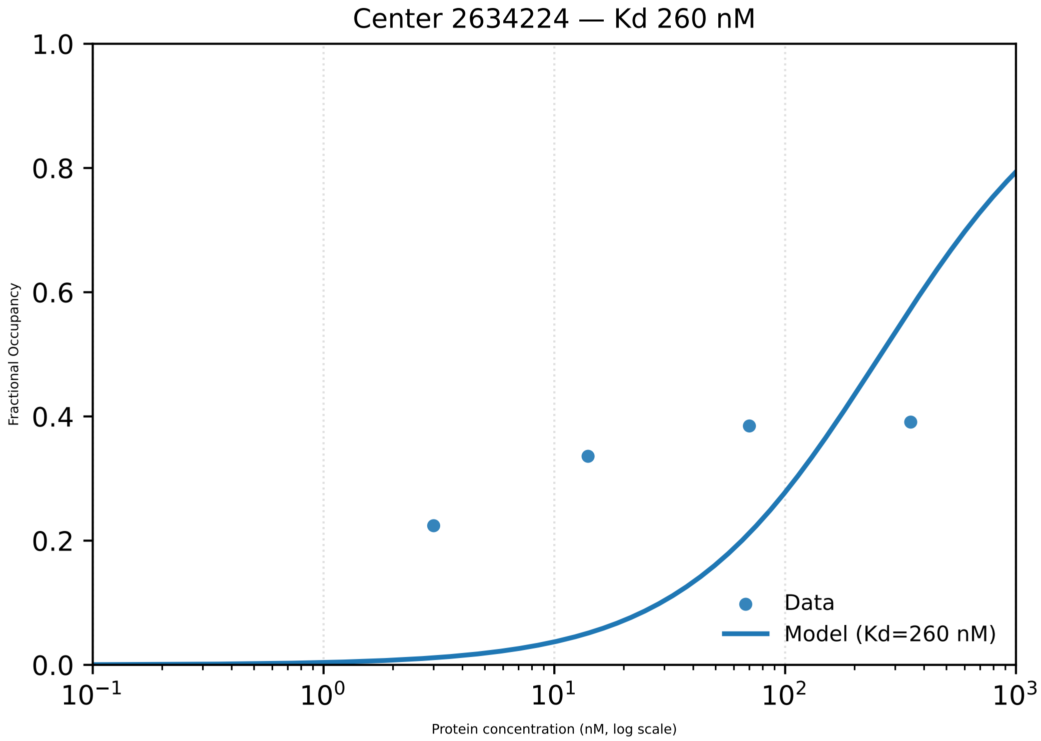

Center 2634224 — Kd 260 nM
Fractional Occupancy
Protein concentration (nM, log scale)
1.0
0.8
0.6
0.4
0.2
0.0
$10^{-1}$
$10^{0}$
$10^{1}$
$10^{2}$
$10^{3}$
Data
Model (Kd=260 nM)

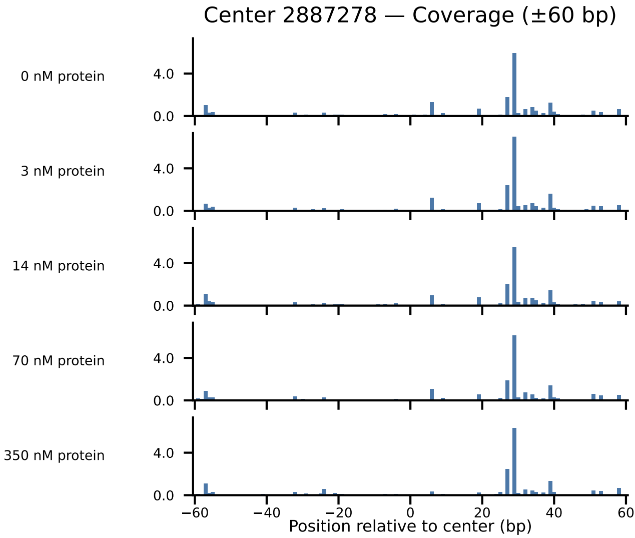
Center 2887278 — Coverage (±60 bp)
0 nM protein
3 nM protein
14 nM protein
70 nM protein
350 nM protein
4.0
0.0
−60
−40
−20
0
20
40
60
Position relative to center (bp)

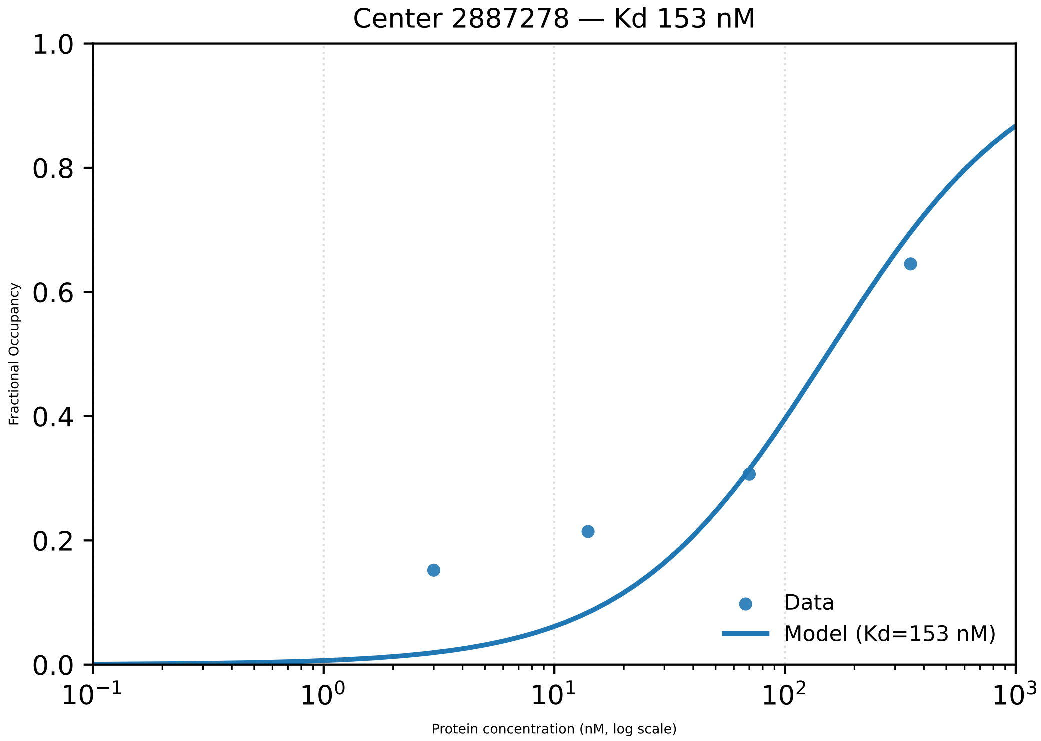

Center 2887278 — Kd 153 nM
1.0
0.8
0.6
0.4
0.2
0.0
Fractional Occupancy
$10^{-1}$
$10^{0}$
$10^{1}$
$10^{2}$
$10^{3}$
Protein concentration (nM, log scale)
Data
Model (Kd=153 nM)

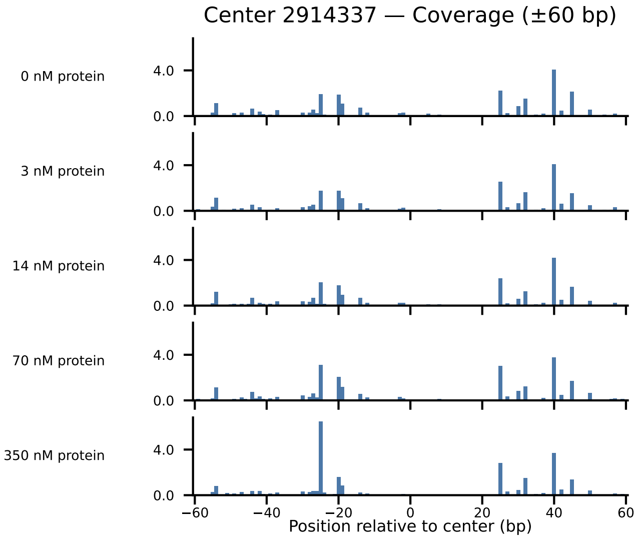

Center 2914337 — Coverage (±60 bp)
0 nM protein
3 nM protein
14 nM protein
70 nM protein
350 nM protein
4.0
0.0
−60
−40
−20
0
20
40
60
Position relative to center (bp)

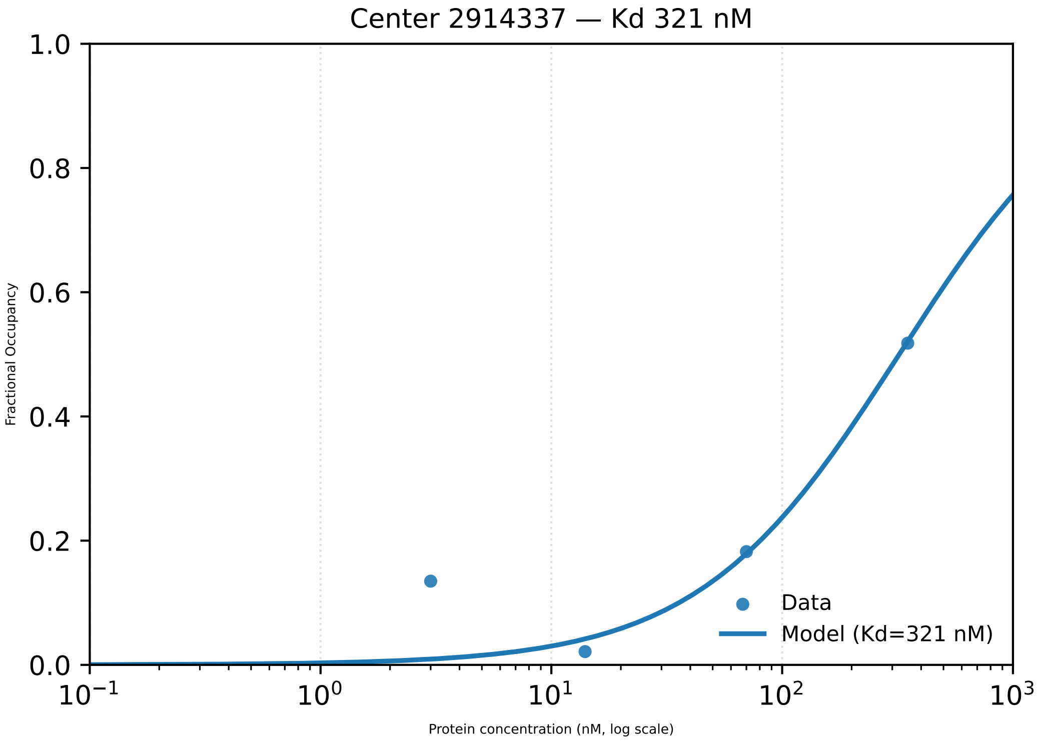

Center 2914337 — Kd 321 nM
1.0
0.8
0.6
0.4
0.2
0.0
Fractional Occupancy
Data
Model (Kd=321 nM)
$10^{-1}$
$10^{0}$
$10^{1}$
$10^{2}$
$10^{3}$
Protein concentration (nM, log scale)

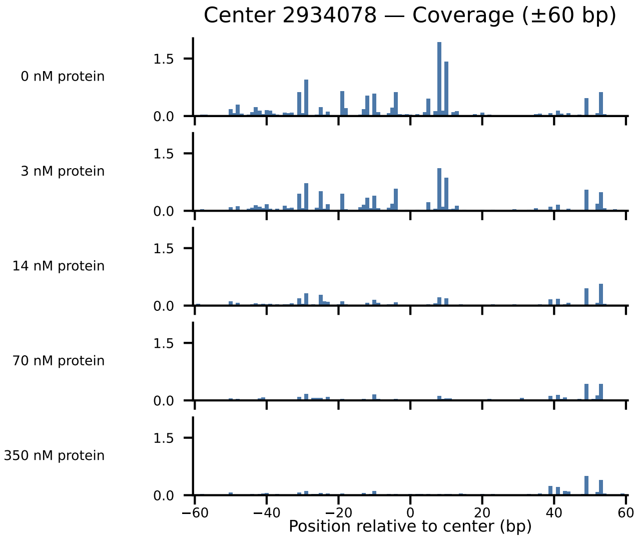

Center 2934078 — Coverage (±60 bp)
0 nM protein
3 nM protein
14 nM protein
70 nM protein
350 nM protein
1.5
0.0
−60
−40
−20
0
20
40
60
Position relative to center (bp)

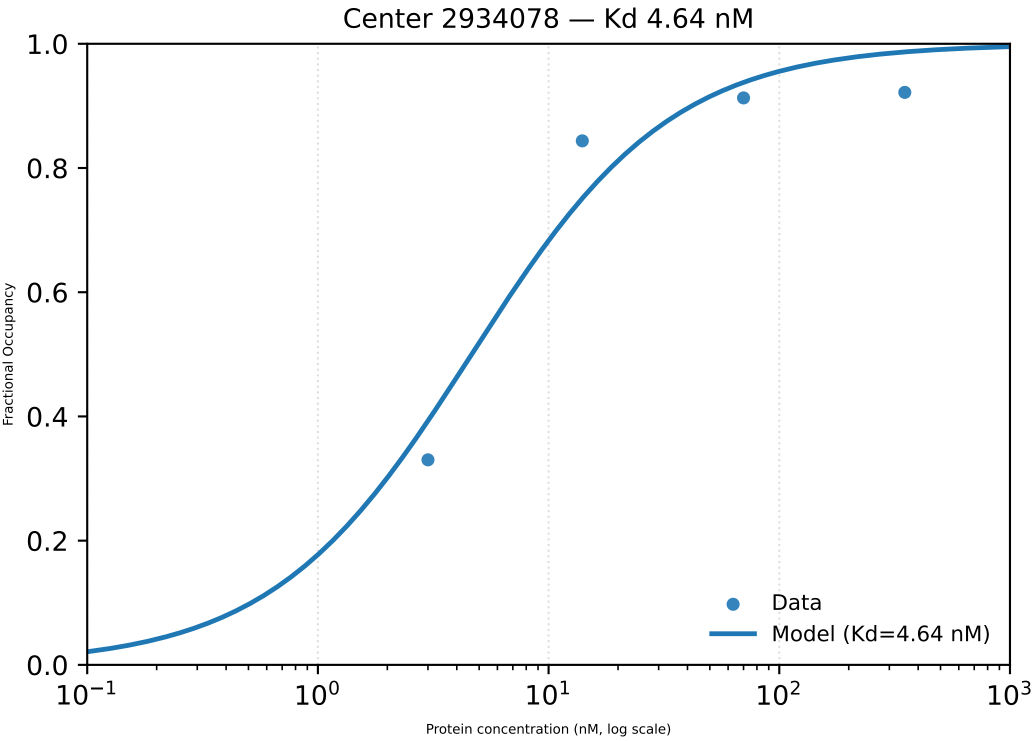

Center 2934078 — Kd 4.64 nM
1.0
0.8
0.6
0.4
0.2
0.0
Fractional Occupancy
$10^{-1}$
$10^{0}$
$10^{1}$
$10^{2}$
$10^{3}$
Protein concentration (nM, log scale)
Data
Model (Kd=4.64 nM)

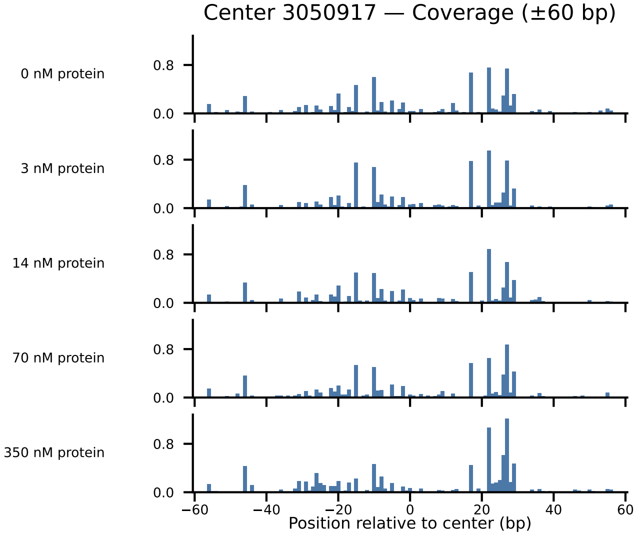

Center 3050917 — Coverage (±60 bp)
0 nM protein
3 nM protein
14 nM protein
70 nM protein
350 nM protein
0.8
0.0
−60
−40
−20
0
20
40
60
Position relative to center (bp)

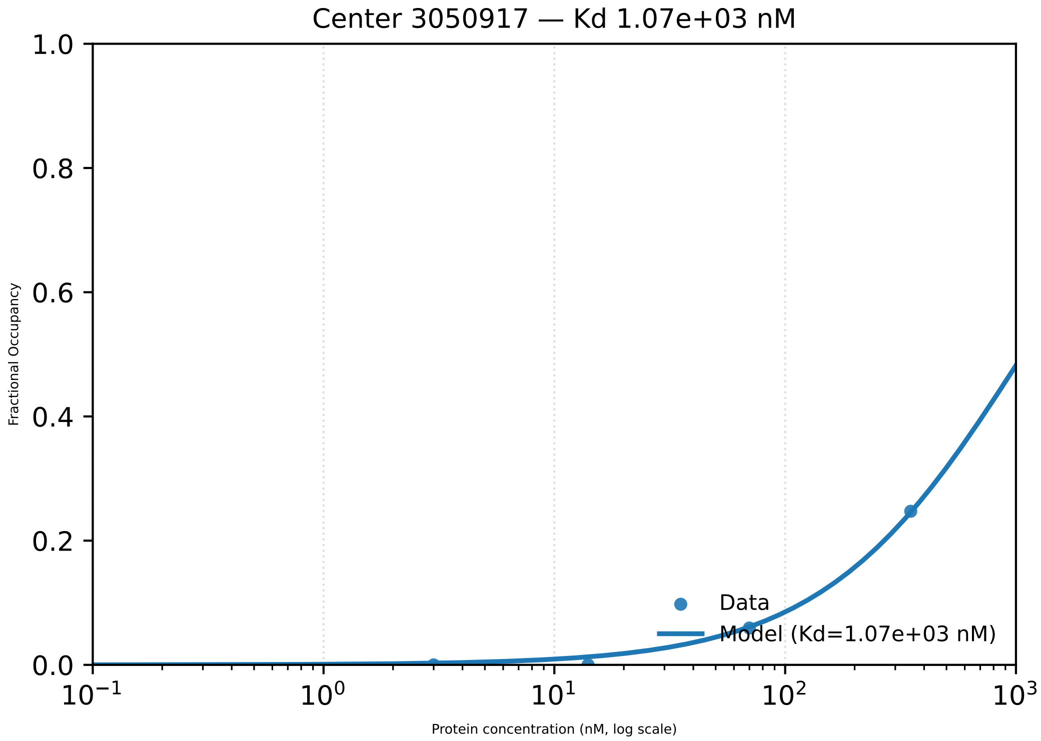

Center 3050917 — Kd 1.07e+03 nM
1.0
0.8
0.6
0.4
0.2
0.0
Fractional Occupancy
Data
Model (Kd=1.07e+03 nM)
$10^{-1}$
$10^{0}$
$10^{1}$
$10^{2}$
$10^{3}$
Protein concentration (nM, log scale)

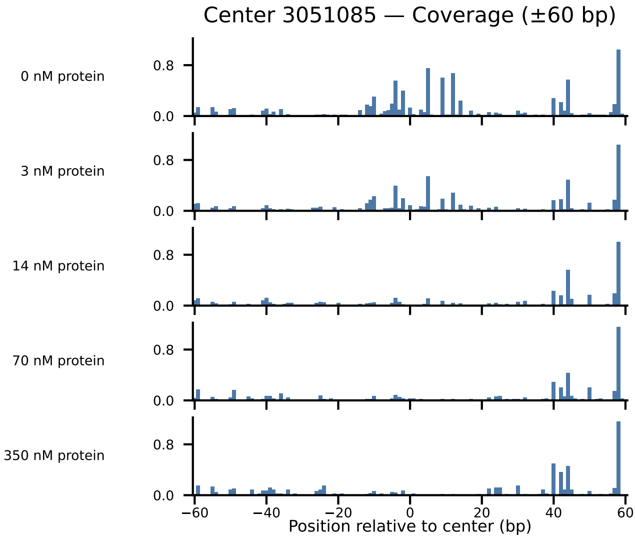

Center 3051085 — Coverage (±60 bp)
0 nM protein
3 nM protein
14 nM protein
70 nM protein
350 nM protein
0.8
0.0
−60
−40
−20
0
20
40
60
Position relative to center (bp)

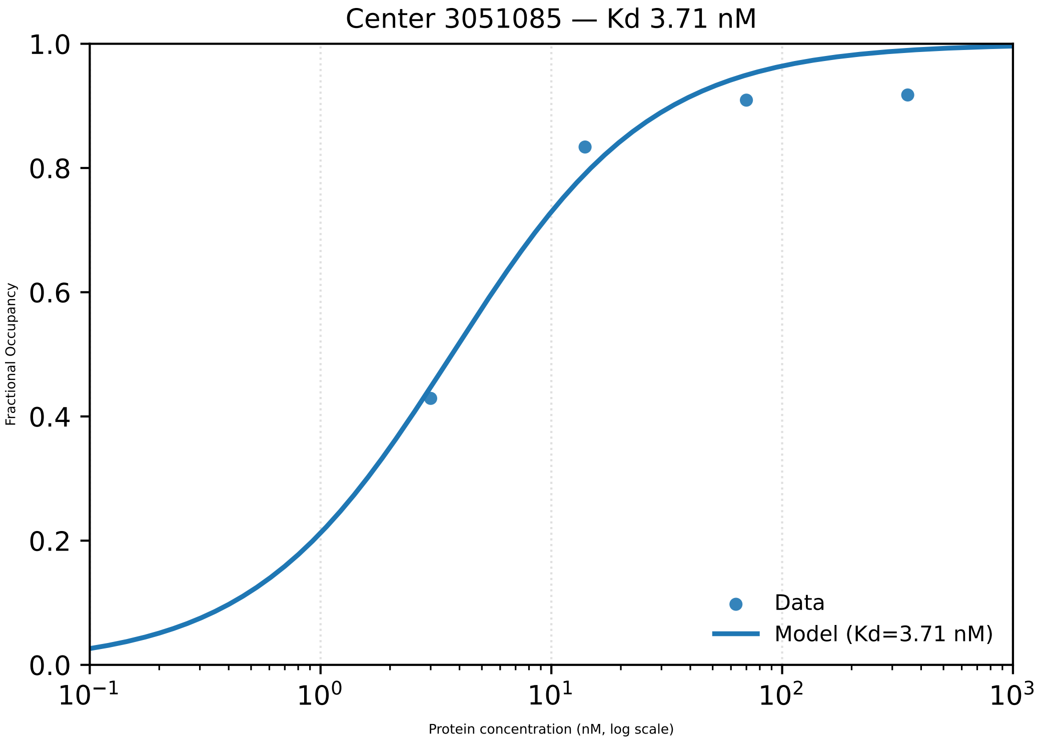

Center 3051085 — Kd 3.71 nM
Fractional Occupancy
1.0
0.8
0.6
0.4
0.2
0.0
$10^{-1}$
$10^{0}$
$10^{1}$
$10^{2}$
$10^{3}$
Protein concentration (nM, log scale)
Data
Model (Kd=3.71 nM)

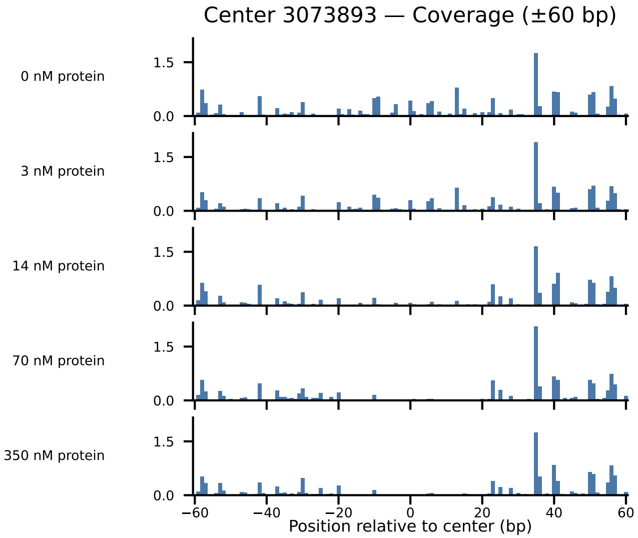
Center 3073893 — Coverage (±60 bp)
0 nM protein
3 nM protein
14 nM protein
70 nM protein
350 nM protein
1.5
0.0
−60
−40
−20
0
20
40
60
Position relative to center (bp)

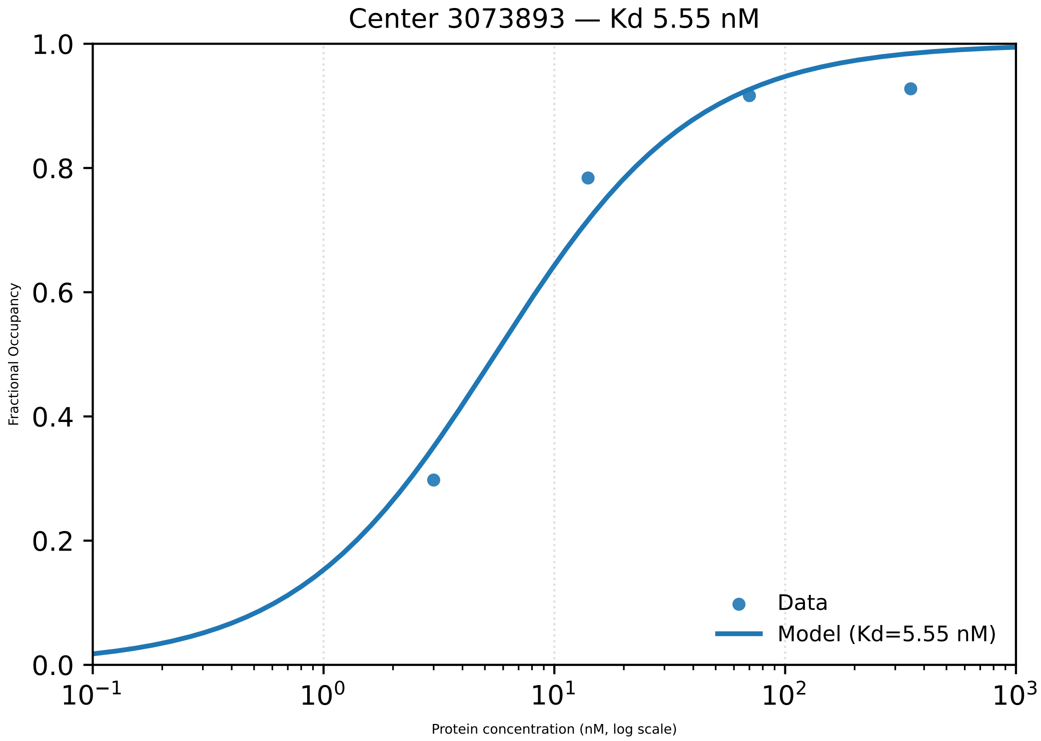

Center 3073893 — Kd 5.55 nM
Fractional Occupancy
Protein concentration (nM, log scale)
Data
Model (Kd=5.55 nM)

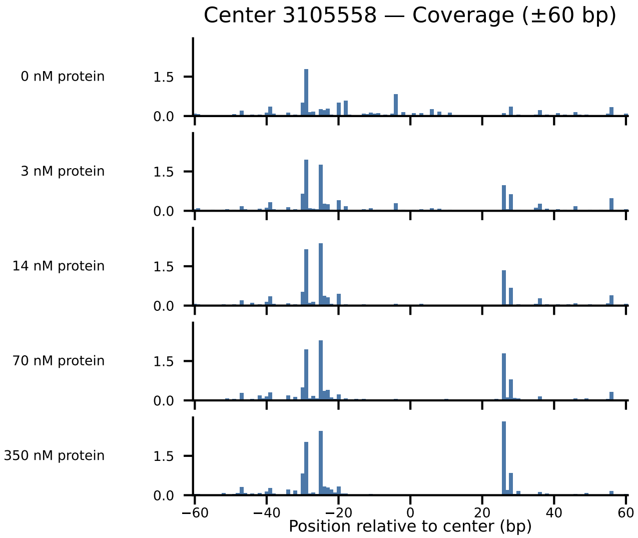

Center 3105558 — Coverage (±60 bp)
0 nM protein
3 nM protein
14 nM protein
70 nM protein
350 nM protein
1.5
0.0
−60
−40
−20
0
20
40
60
Position relative to center (bp)

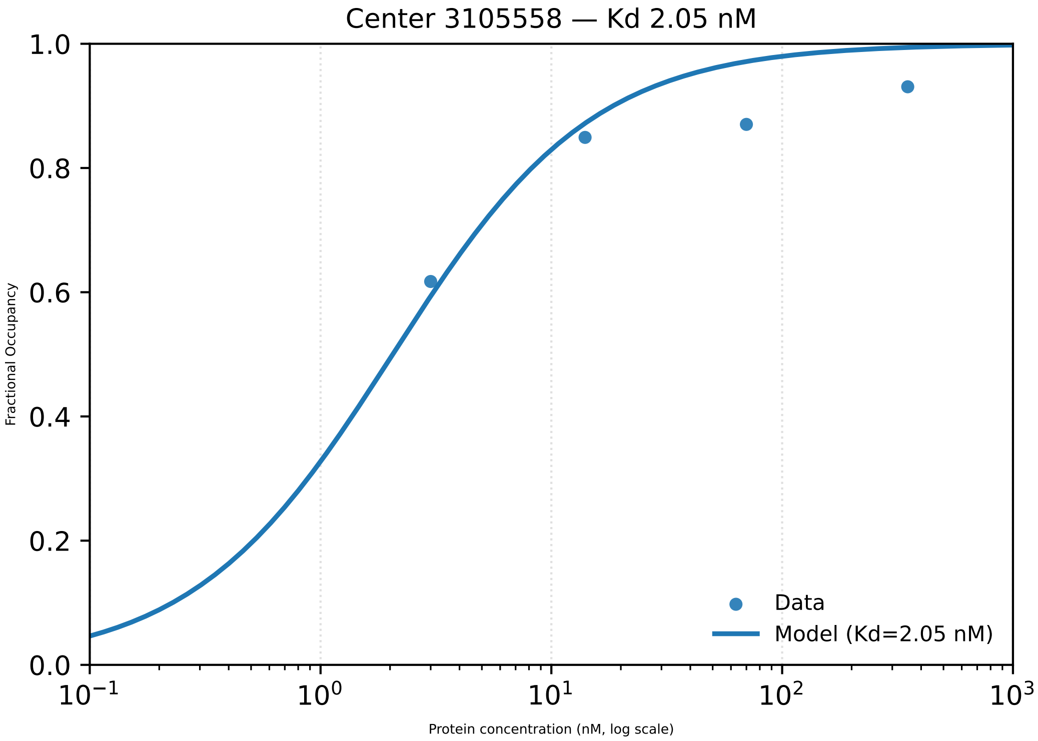
Center 3105558 — Kd 2.05 nM
Fractional Occupancy
1.0
0.8
0.6
0.4
0.2
0.0
$10^{-1}$
$10^{0}$
$10^{1}$
$10^{2}$
$10^{3}$
Protein concentration (nM, log scale)
Data
Model (Kd=2.05 nM)

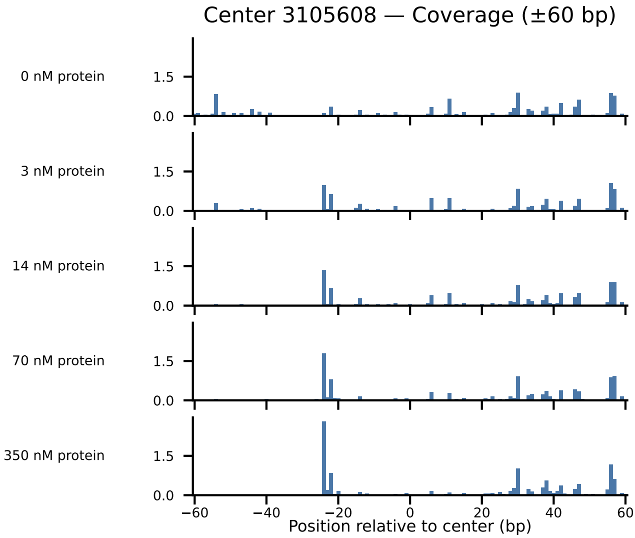

Center 3105608 — Coverage (±60 bp)
0 nM protein
3 nM protein
14 nM protein
70 nM protein
350 nM protein
1.5
0.0
−60
−40
−20
0
20
40
60
Position relative to center (bp)

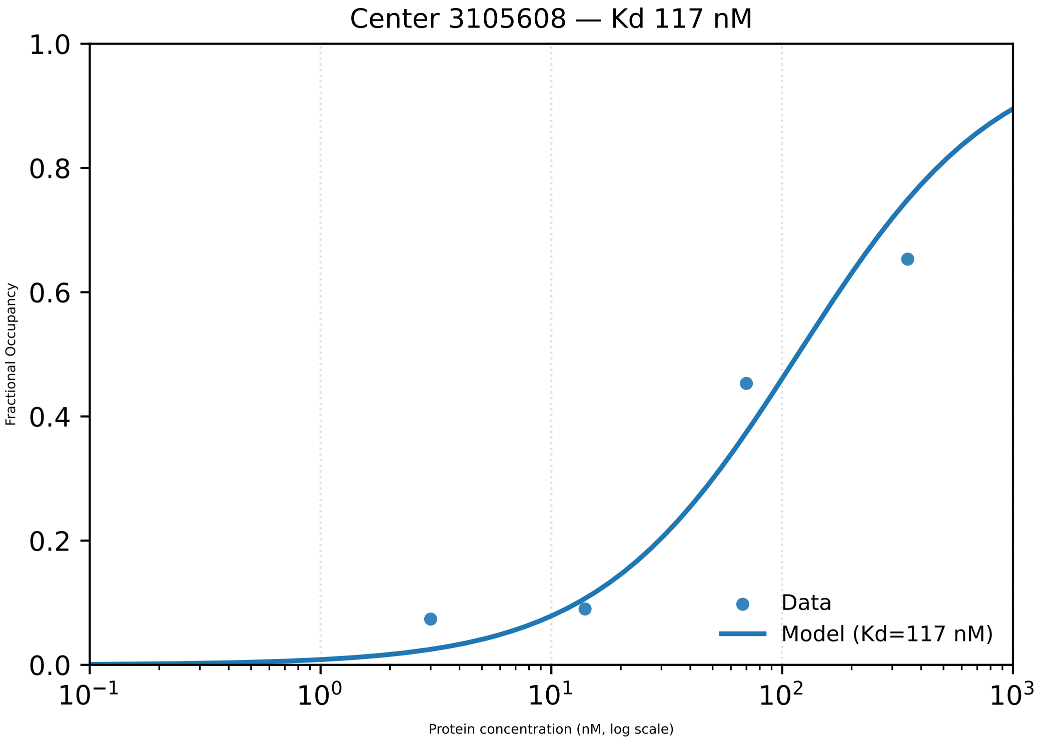

Center 3105608 — Kd 117 nM
1.0
0.8
0.6
0.4
0.2
0.0
Fractional Occupancy
$10^{-1}$
$10^{0}$
$10^{1}$
$10^{2}$
$10^{3}$
Protein concentration (nM, log scale)
Data
Model (Kd=117 nM)

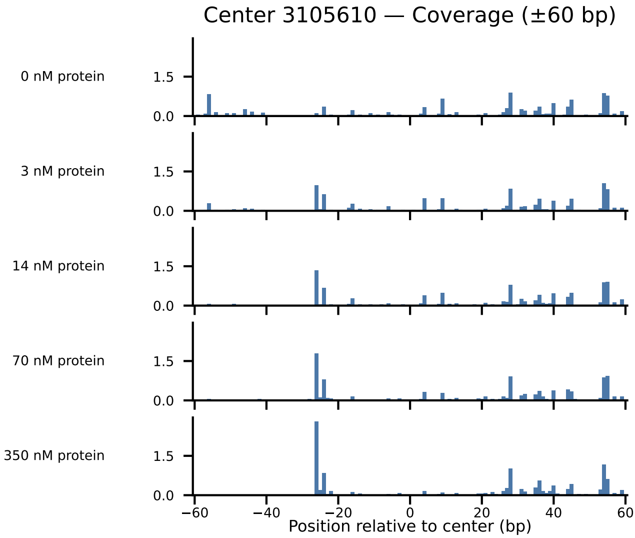

Center 3105610 — Coverage (±60 bp)
0 nM protein
3 nM protein
14 nM protein
70 nM protein
350 nM protein
1.5
0.0
−60
−40
−20
0
20
40
60
Position relative to center (bp)

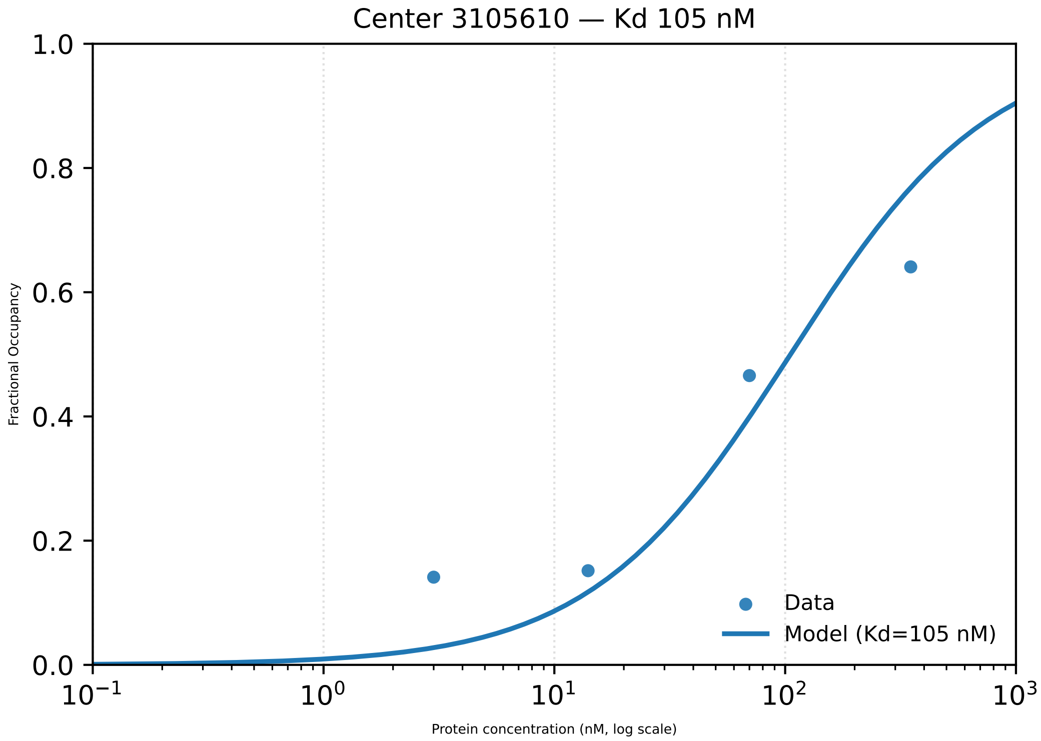

Center 3105610 — Kd 105 nM
1.0
0.8
0.6
0.4
0.2
0.0
Fractional Occupancy
$10^{-1}$
$10^{0}$
$10^{1}$
$10^{2}$
$10^{3}$
Protein concentration (nM, log scale)
Data
Model (Kd=105 nM)

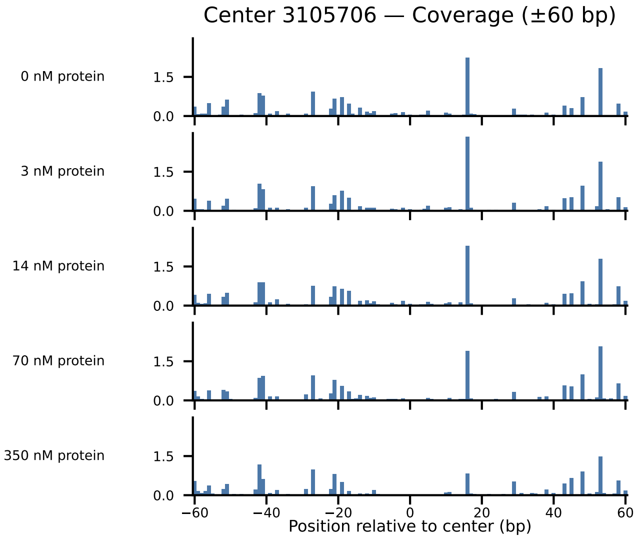

Center 3105706 — Coverage (±60 bp)
0 nM protein
3 nM protein
14 nM protein
70 nM protein
350 nM protein
1.5
0.0
−60
−40
−20
0
20
40
60
Position relative to center (bp)

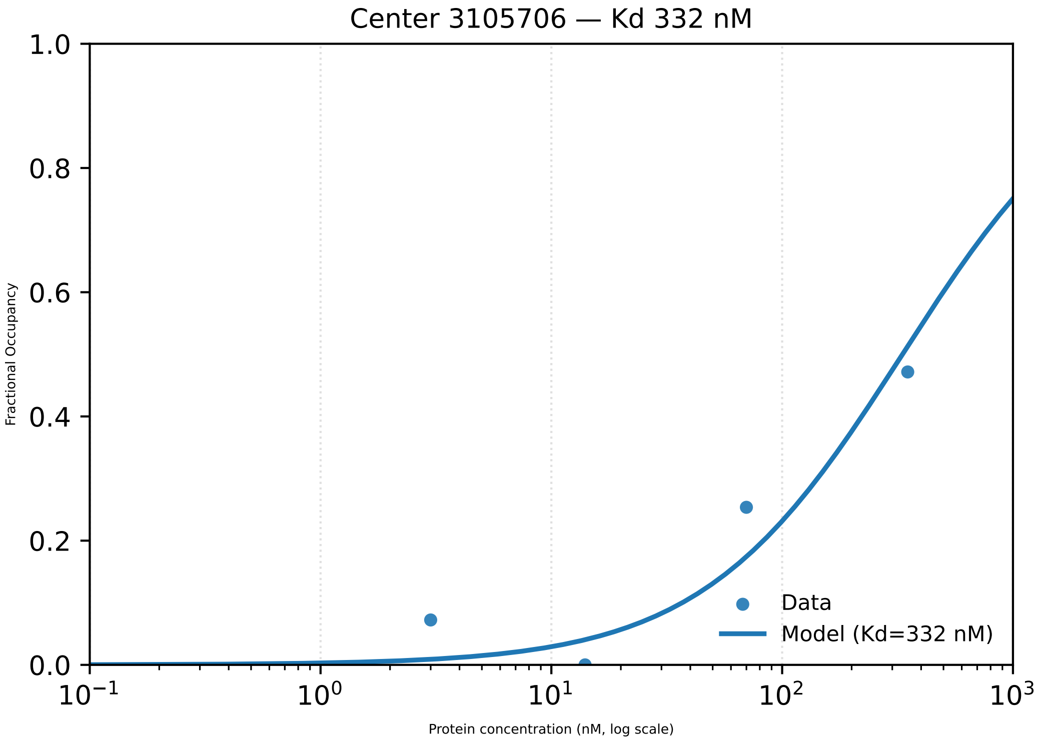

Center 3105706 — Kd 332 nM
1.0
0.8
0.6
0.4
0.2
0.0
Fractional Occupancy
$10^{-1}$
$10^{0}$
$10^{1}$
$10^{2}$
$10^{3}$
Protein concentration (nM, log scale)
Data
Model (Kd=332 nM)

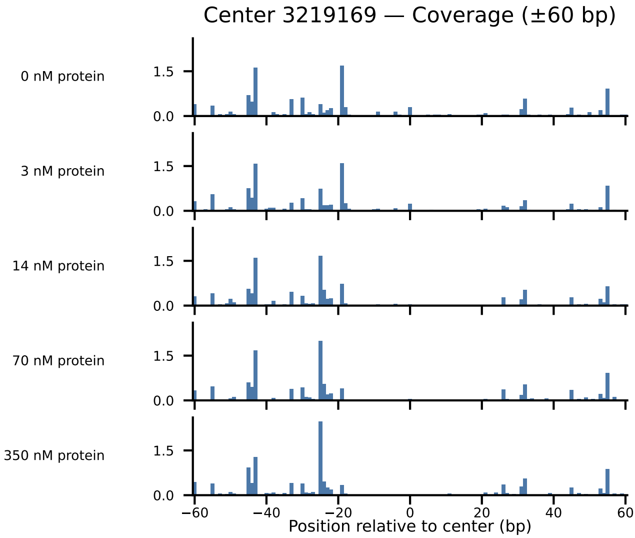

Center 3219169 — Coverage (±60 bp)
0 nM protein
3 nM protein
14 nM protein
70 nM protein
350 nM protein
1.5
0.0
−60
−40
−20
0
20
40
60
Position relative to center (bp)

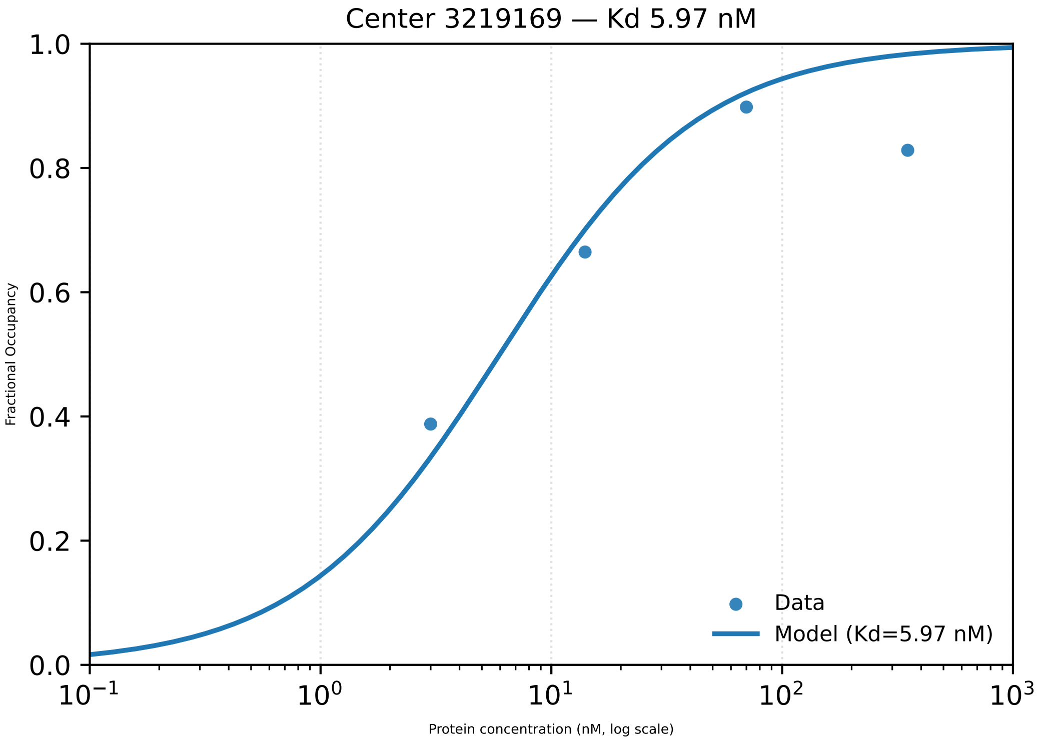

Center 3219169 — Kd 5.97 nM
1.0
0.8
0.6
0.4
0.2
0.0
Fractional Occupancy
$10^{-1}$
$10^{0}$
$10^{1}$
$10^{2}$
$10^{3}$
Protein concentration (nM, log scale)
Data
Model (Kd=5.97 nM)

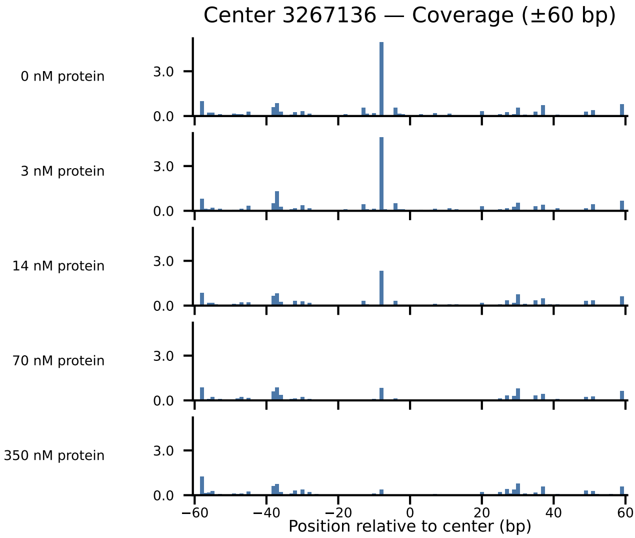

Center 3267136 — Coverage (±60 bp)
0 nM protein
3 nM protein
14 nM protein
70 nM protein
350 nM protein
3.0
0.0
−60
−40
−20
0
20
40
60
Position relative to center (bp)

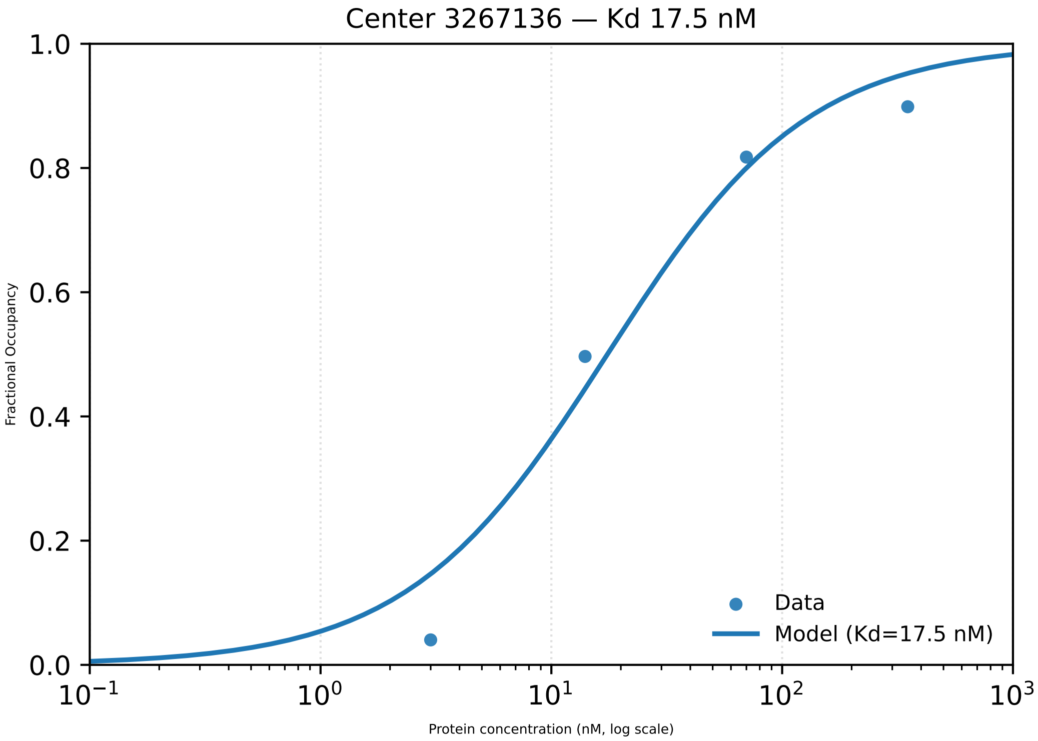
Center 3267136 — Kd 17.5 nM
Fractional Occupancy
1.0
0.8
0.6
0.4
0.2
0.0
$10^{-1}$
$10^{0}$
$10^{1}$
$10^{2}$
$10^{3}$
Protein concentration (nM, log scale)
Data
Model (Kd=17.5 nM)

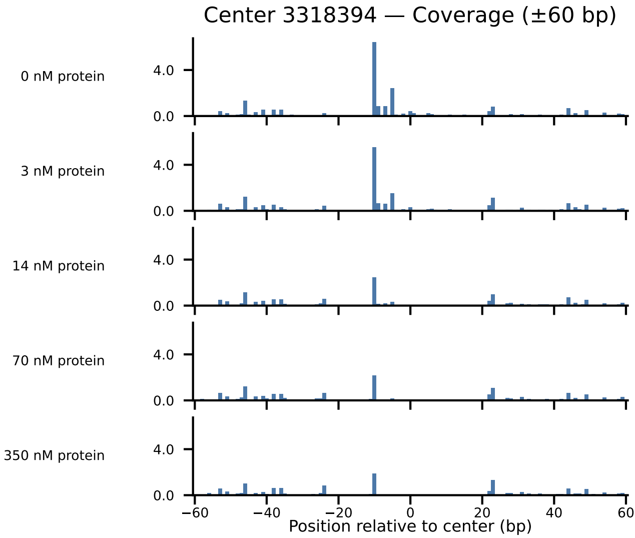
Center 3318394 — Coverage (±60 bp)
0 nM protein
3 nM protein
14 nM protein
70 nM protein
350 nM protein
4.0
0.0
−60
−40
−20
0
20
40
60
Position relative to center (bp)

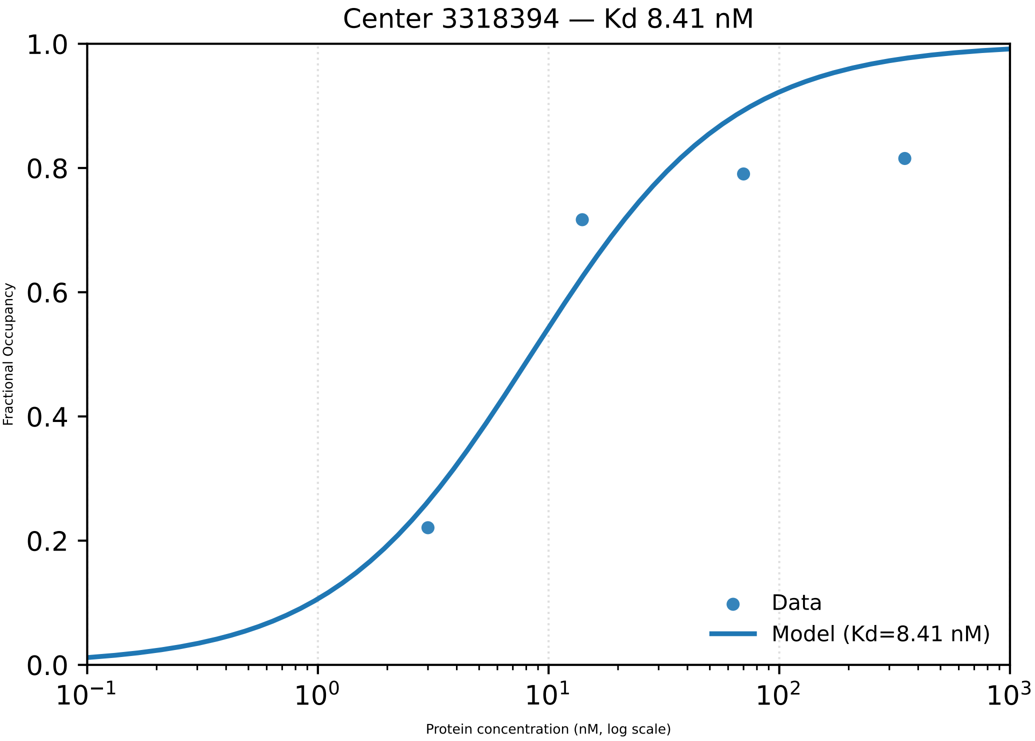

Center 3318394 — Kd 8.41 nM
Fractional Occupancy
1.0
0.8
0.6
0.4
0.2
0.0
$10^{-1}$
$10^{0}$
$10^{1}$
$10^{2}$
$10^{3}$
Protein concentration (nM, log scale)
Data
Model (Kd=8.41 nM)

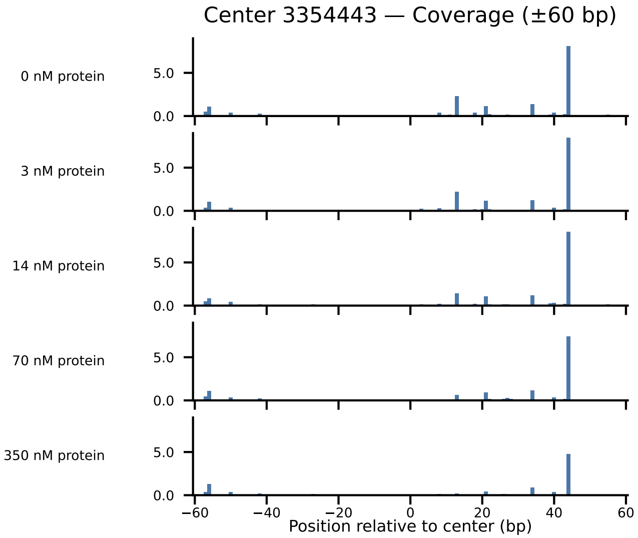

Center 3354443 — Coverage (±60 bp)
0 nM protein
3 nM protein
14 nM protein
70 nM protein
350 nM protein
5.0
0.0
−60
−40
−20
0
20
40
60
Position relative to center (bp)

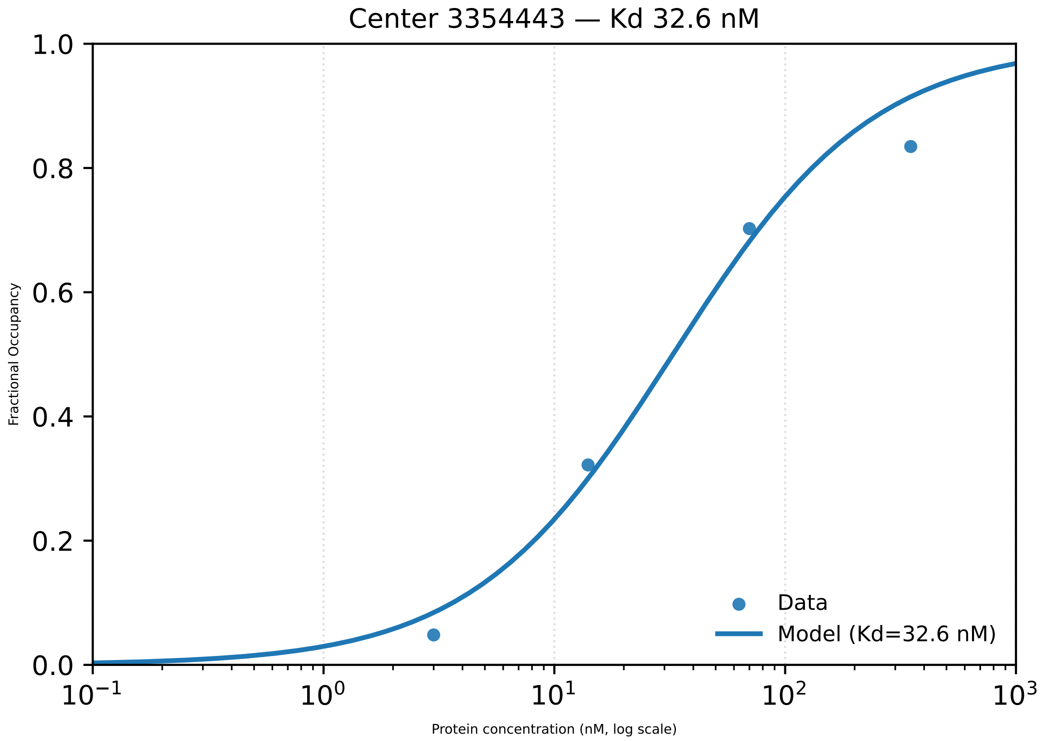

Center 3354443 — Kd 32.6 nM
1.0
0.8
0.6
0.4
0.2
0.0
Fractional Occupancy
$10^{-1}$
$10^{0}$
$10^{1}$
$10^{2}$
$10^{3}$
Protein concentration (nM, log scale)
Data
Model (Kd=32.6 nM)

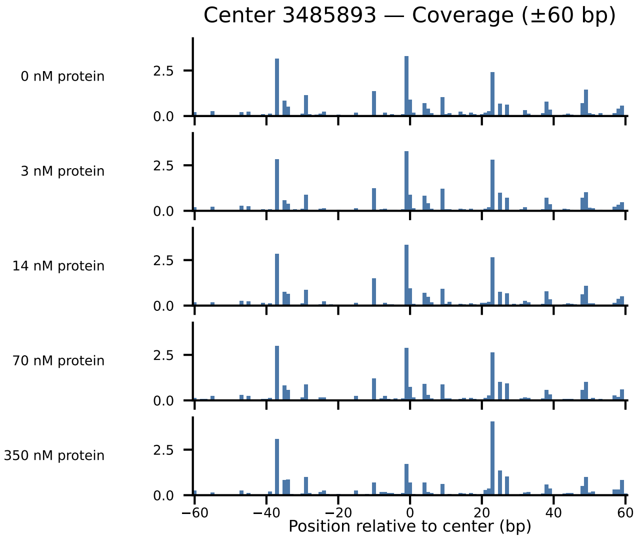
Center 3485893 — Coverage (±60 bp)
0 nM protein
3 nM protein
14 nM protein
70 nM protein
350 nM protein
2.5
0.0
−60
−40
−20
0
20
40
60
Position relative to center (bp)

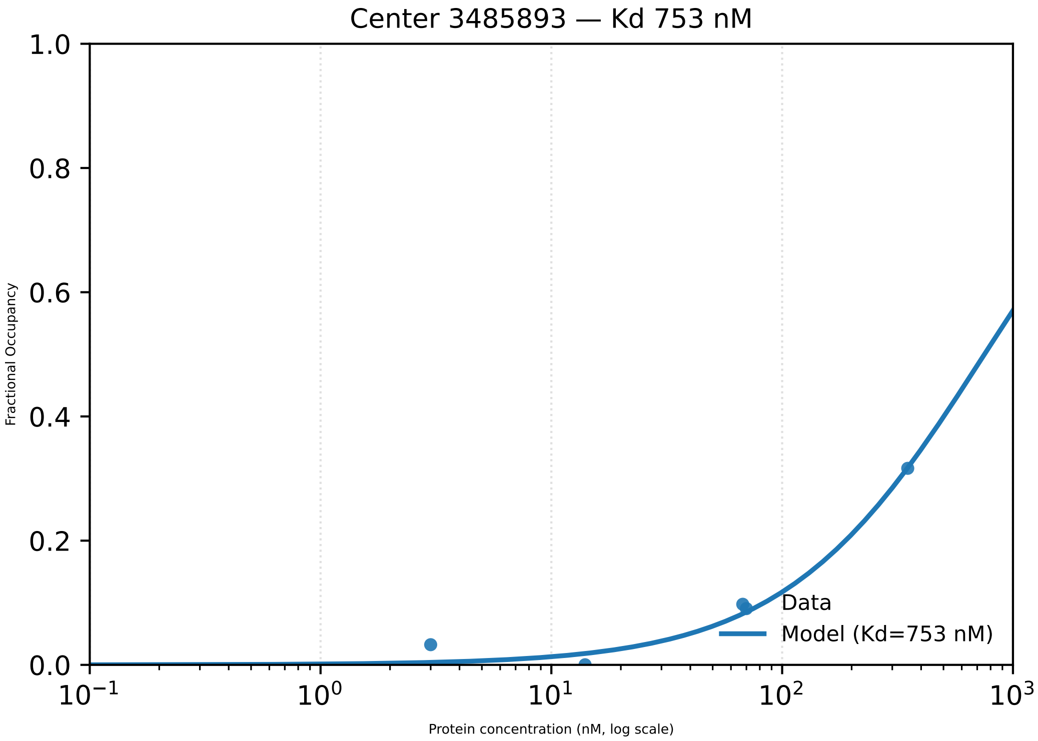

Center 3485893 — Kd 753 nM
1.0
0.8
0.6
0.4
0.2
0.0
Fractional Occupancy
$10^{-1}$
$10^{0}$
$10^{1}$
$10^{2}$
$10^{3}$
Protein concentration (nM, log scale)
Data
Model (Kd=753 nM)

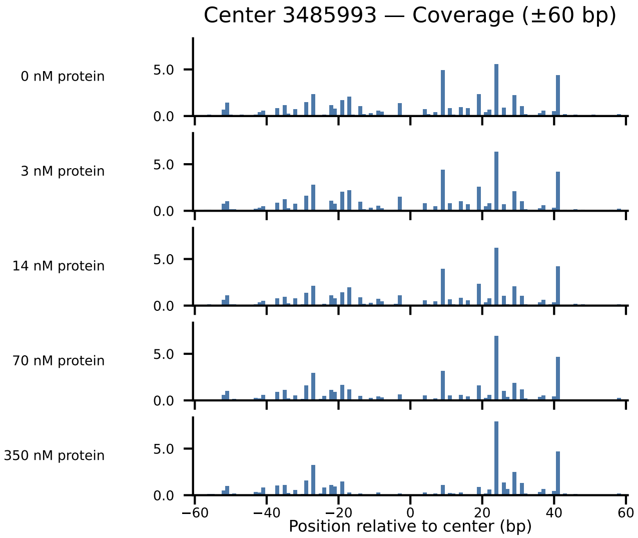

Center 3485993 — Coverage (±60 bp)
0 nM protein
3 nM protein
14 nM protein
70 nM protein
350 nM protein
5.0
0.0
−60
−40
−20
0
20
40
60
Position relative to center (bp)

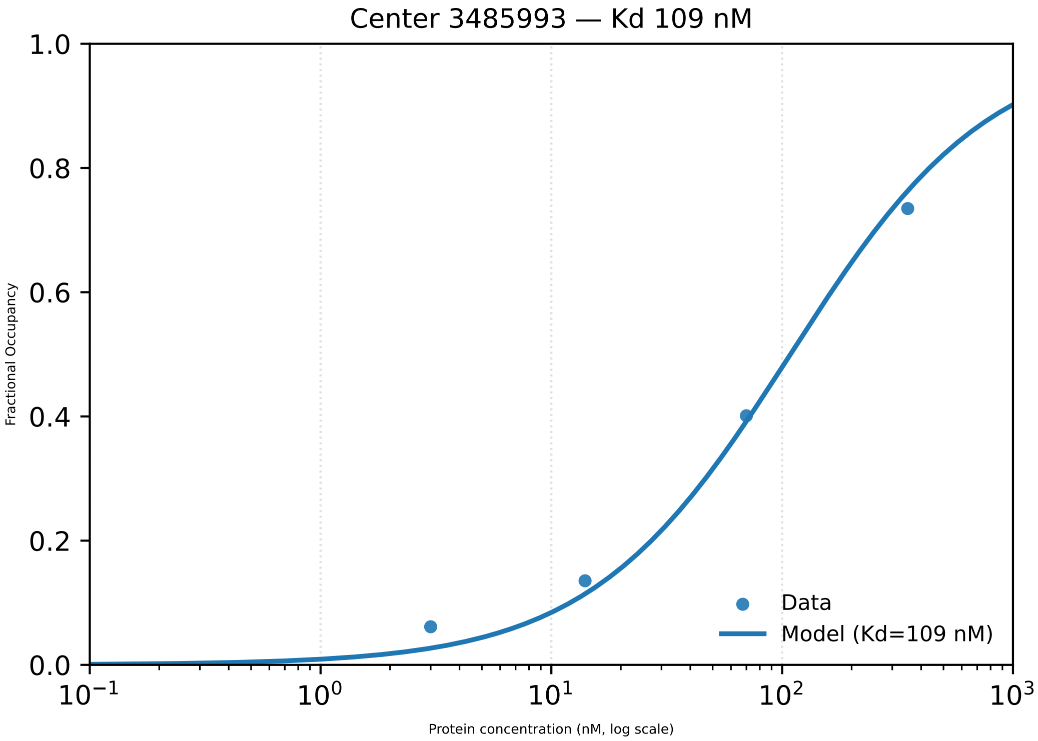

Center 3485993 — Kd 109 nM
1.0
0.8
0.6
0.4
0.2
0.0
Fractional Occupancy
$10^{-1}$
$10^{0}$
$10^{1}$
$10^{2}$
$10^{3}$
Protein concentration (nM, log scale)
Data
Model (Kd=109 nM)

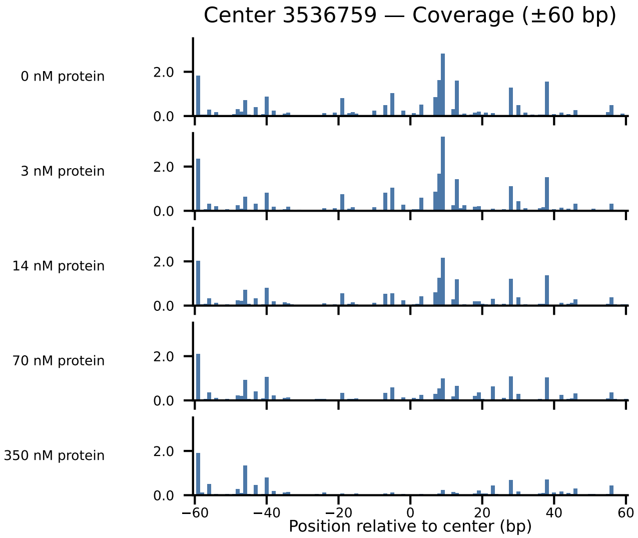

Center 3536759 — Coverage (±60 bp)
0 nM protein
3 nM protein
14 nM protein
70 nM protein
350 nM protein
2.0
0.0
−60
−40
−20
0
20
40
60
Position relative to center (bp)

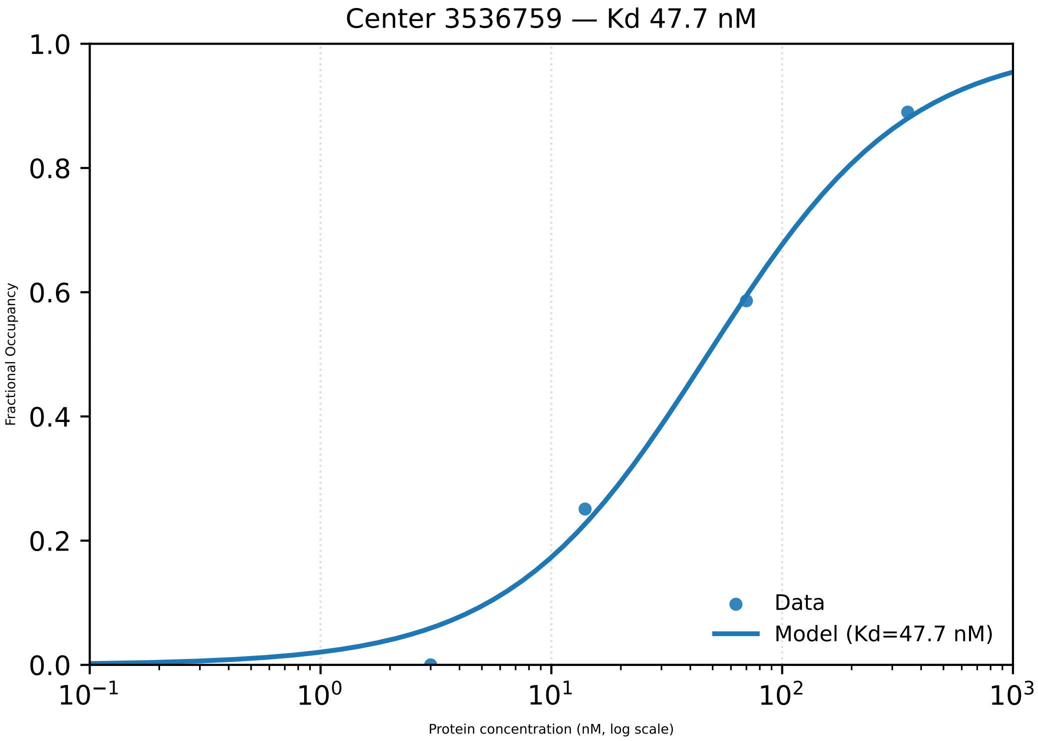

Center 3536759 — Kd 47.7 nM
1.0
0.8
0.6
0.4
0.2
0.0
Fractional Occupancy
$10^{-1}$
$10^{0}$
$10^{1}$
$10^{2}$
$10^{3}$
Protein concentration (nM, log scale)
Data
Model (Kd=47.7 nM)

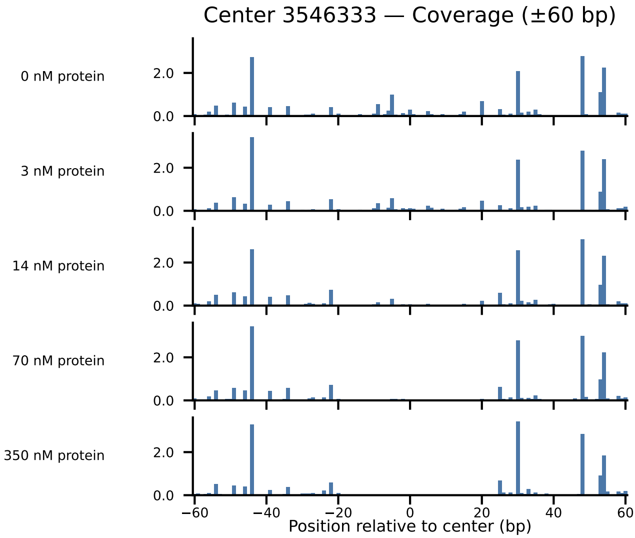

Center 3546333 — Coverage (±60 bp)
0 nM protein
3 nM protein
14 nM protein
70 nM protein
350 nM protein
2.0
0.0
−60
−40
−20
0
20
40
60
Position relative to center (bp)

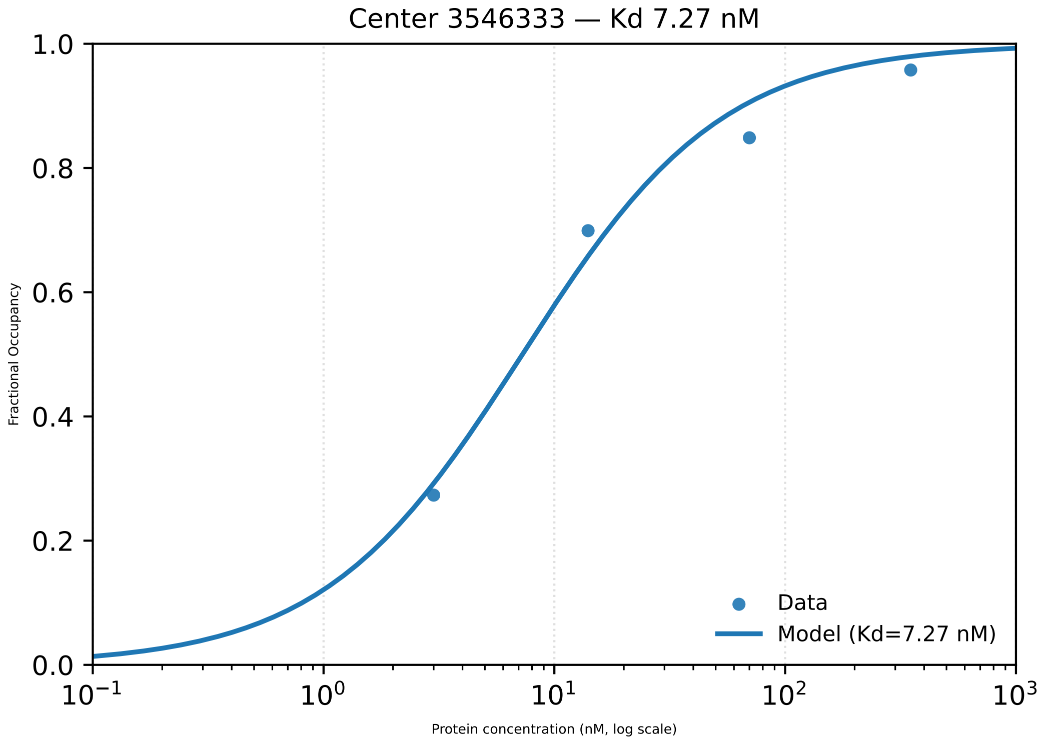

Center 3546333 — Kd 7.27 nM
Fractional Occupancy
1.0
0.8
0.6
0.4
0.2
0.0
$10^{-1}$
$10^{0}$
$10^{1}$
$10^{2}$
$10^{3}$
Protein concentration (nM, log scale)
Data
Model (Kd=7.27 nM)

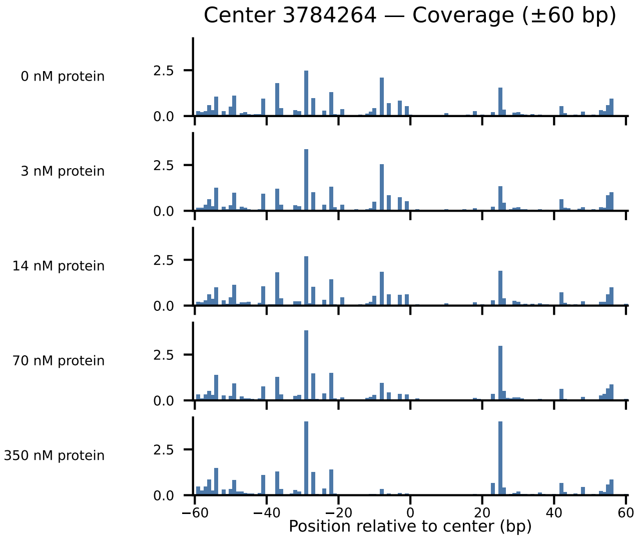

Center 3784264 — Coverage (±60 bp)
0 nM protein
3 nM protein
14 nM protein
70 nM protein
350 nM protein
2.5
0.0
−60
−40
−20
0
20
40
60
Position relative to center (bp)

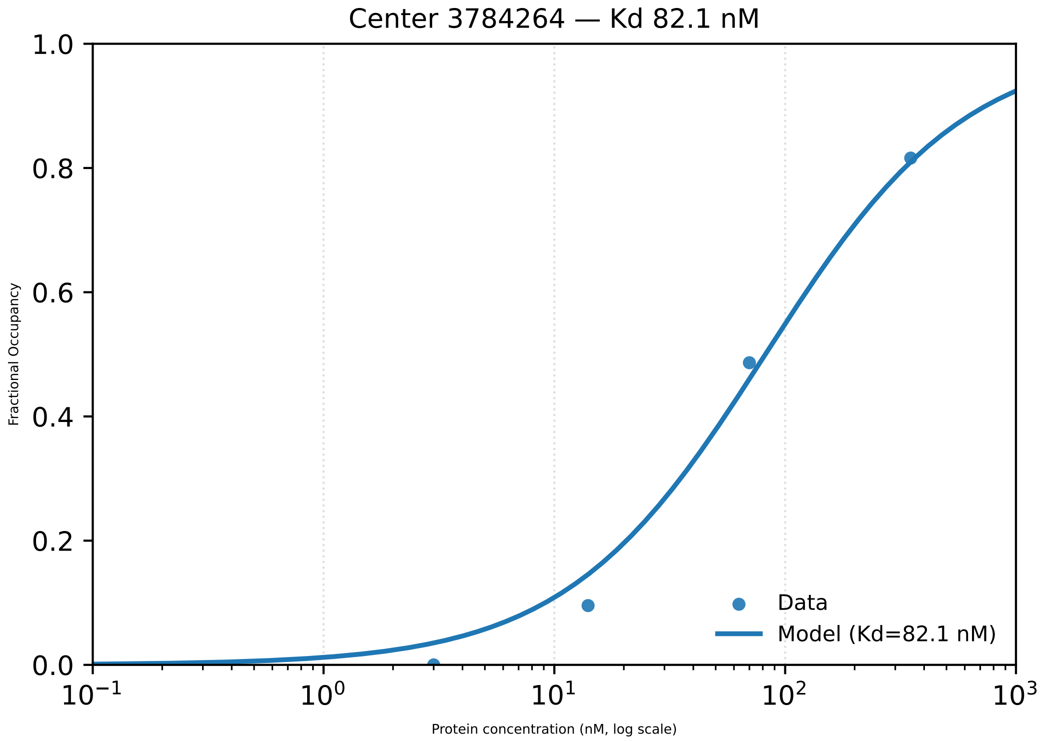

Center 3784264 — Kd 82.1 nM
1.0
0.8
0.6
0.4
0.2
0.0
Fractional Occupancy
$10^{-1}$
$10^{0}$
$10^{1}$
$10^{2}$
$10^{3}$
Protein concentration (nM, log scale)
Data
Model (Kd=82.1 nM)

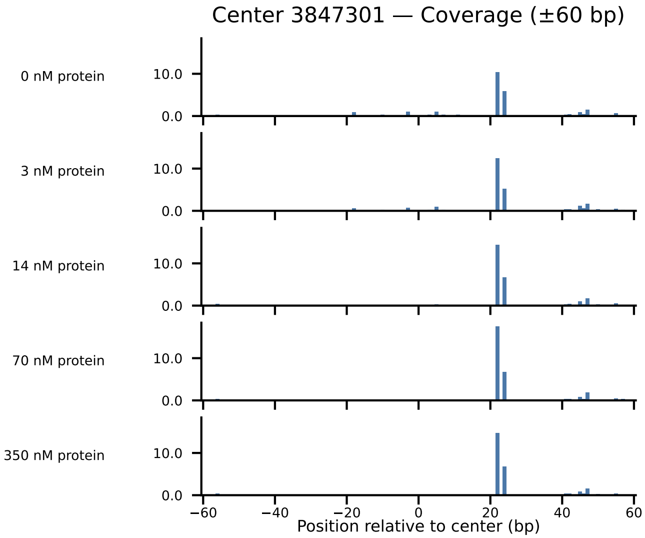

Center 3847301 — Coverage (±60 bp)
0 nM protein
3 nM protein
14 nM protein
70 nM protein
350 nM protein
10.0
0.0
−60
−40
−20
0
20
40
60
Position relative to center (bp)

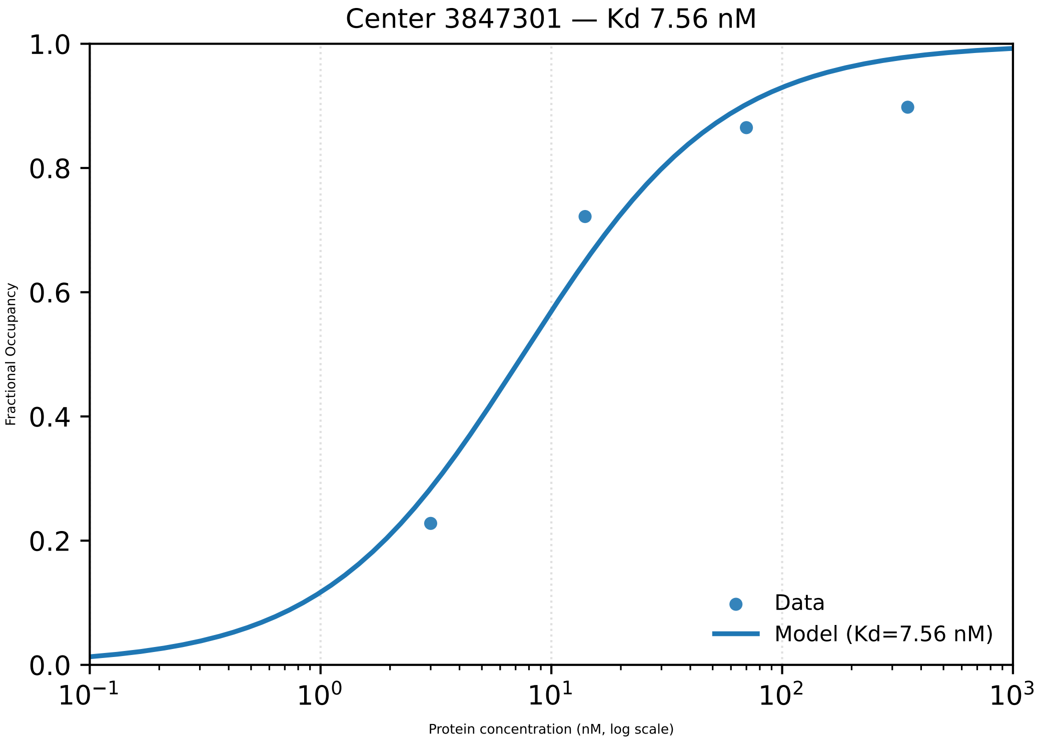

Center 3847301 — Kd 7.56 nM
1.0
0.8
0.6
0.4
0.2
0.0
Fractional Occupancy
Data
Model (Kd=7.56 nM)
$10^{-1}$
$10^{0}$
$10^{1}$
$10^{2}$
$10^{3}$
Protein concentration (nM, log scale)

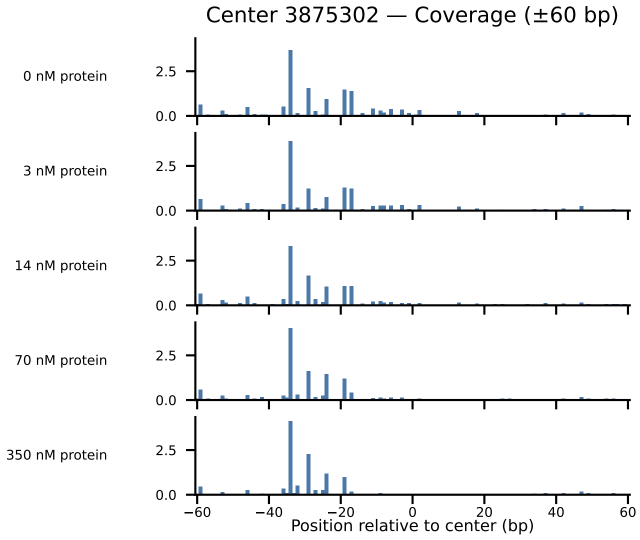
Center 3875302 — Coverage (±60 bp)
0 nM protein
3 nM protein
14 nM protein
70 nM protein
350 nM protein
2.5
0.0
−60
−40
−20
0
20
40
60
Position relative to center (bp)

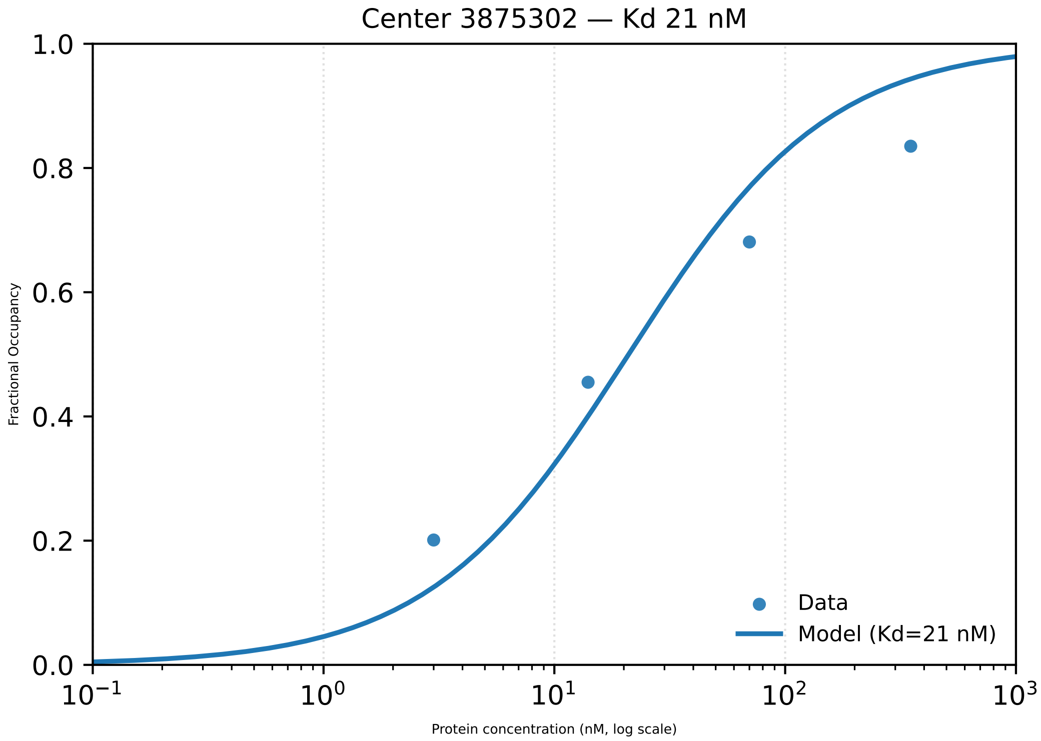

Center 3875302 — Kd 21 nM
Fractional Occupancy
1.0
0.8
0.6
0.4
0.2
0.0
$10^{-1}$
$10^{0}$
$10^{1}$
$10^{2}$
$10^{3}$
Protein concentration (nM, log scale)
Data
Model (Kd=21 nM)

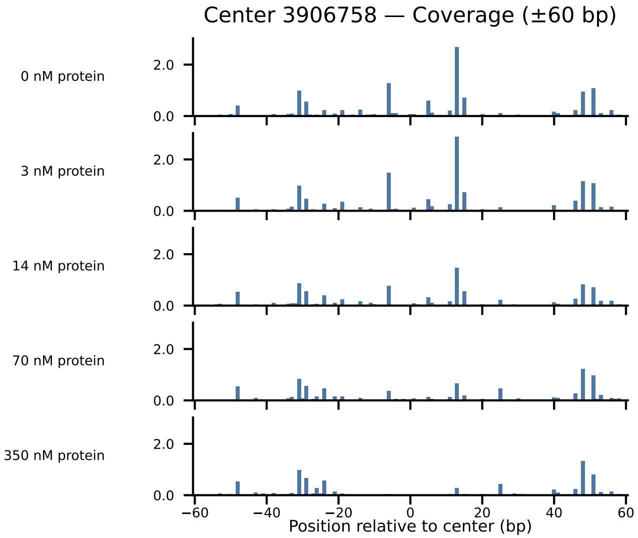

Center 3906758 — Coverage (±60 bp)
0 nM protein
3 nM protein
14 nM protein
70 nM protein
350 nM protein
2.0
0.0
−60
−40
−20
0
20
40
60
Position relative to center (bp)

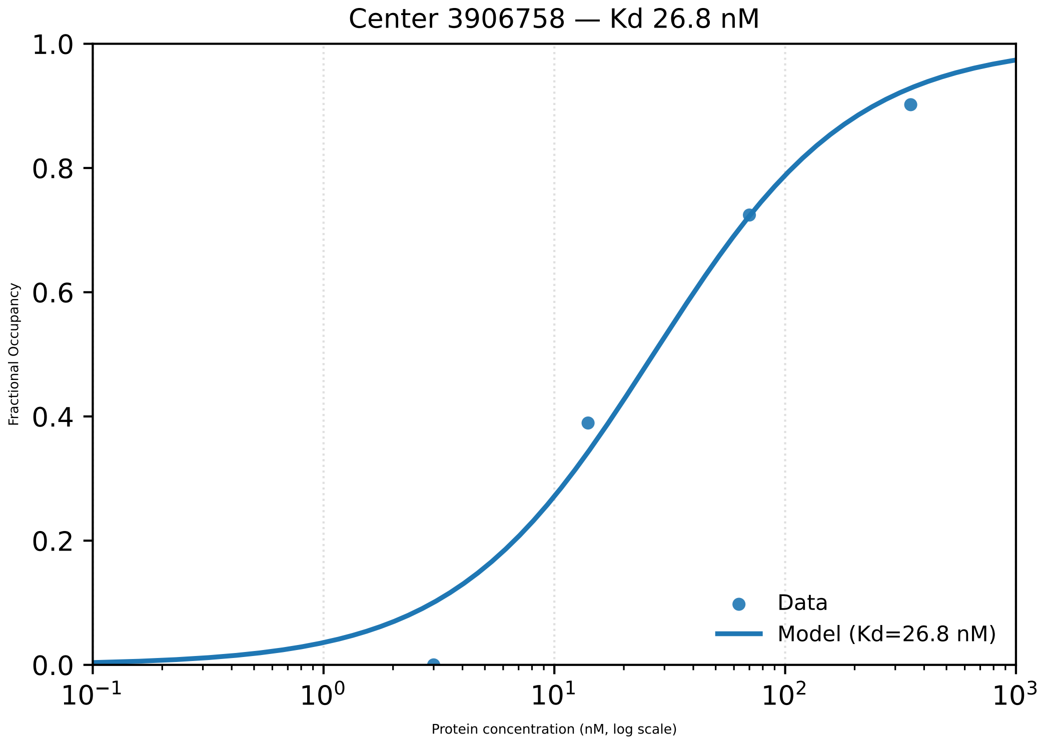

Center 3906758 — Kd 26.8 nM
1.0
0.8
0.6
0.4
0.2
0.0
Fractional Occupancy
$10^{-1}$
$10^{0}$
$10^{1}$
$10^{2}$
$10^{3}$
Protein concentration (nM, log scale)
Data
Model (Kd=26.8 nM)

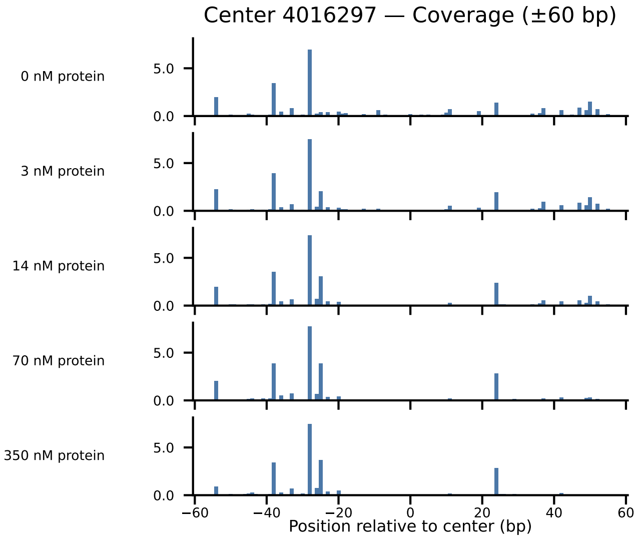
Center 4016297 — Coverage (±60 bp)
0 nM protein
3 nM protein
14 nM protein
70 nM protein
350 nM protein
5.0
0.0
−60
−40
−20
0
20
40
60
Position relative to center (bp)

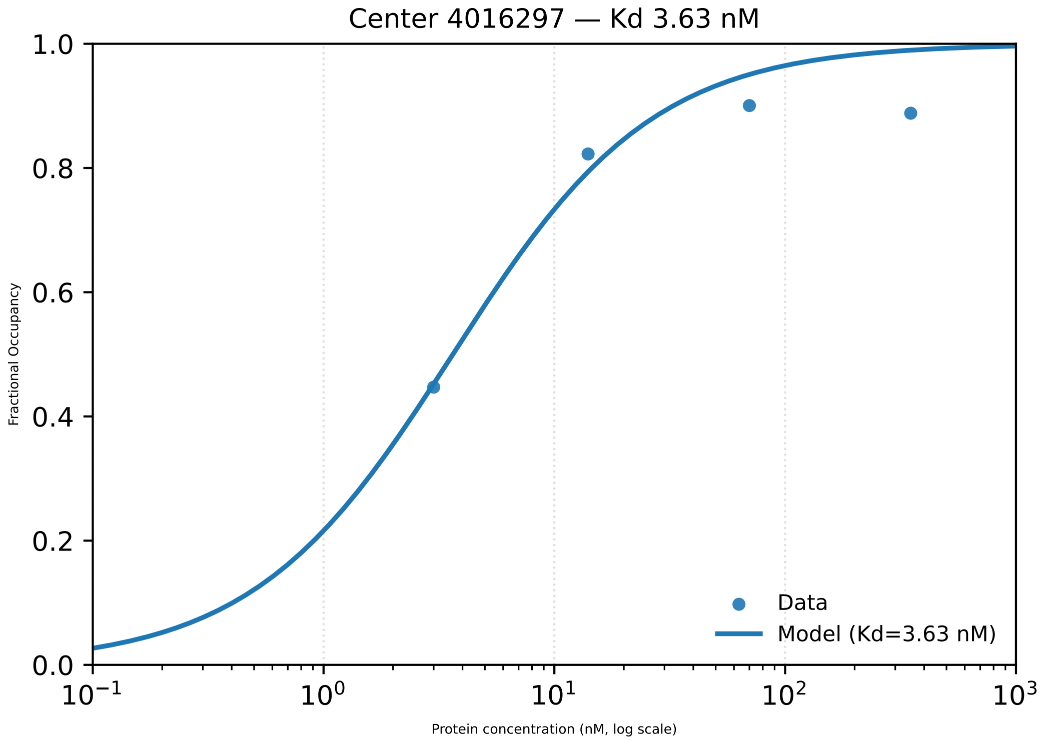

Center 4016297 — Kd 3.63 nM
1.0
0.8
0.6
0.4
0.2
0.0
Fractional Occupancy
Data
Model (Kd=3.63 nM)
$10^{-1}$
$10^{0}$
$10^{1}$
$10^{2}$
$10^{3}$
Protein concentration (nM, log scale)

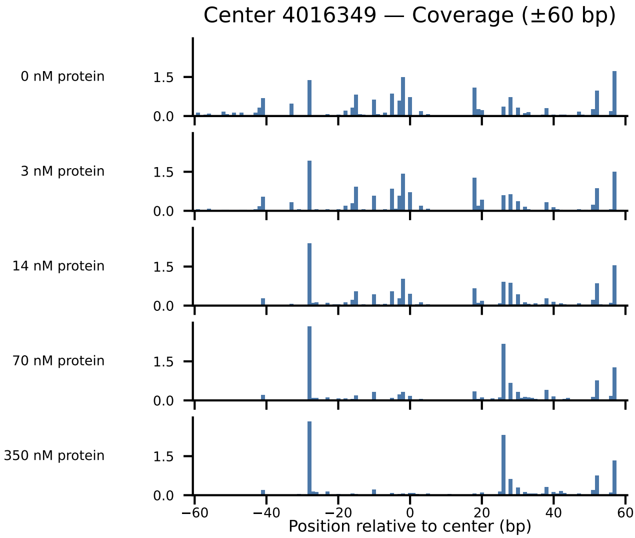

Center 4016349 — Coverage (±60 bp)
0 nM protein
3 nM protein
14 nM protein
70 nM protein
350 nM protein
1.5
0.0
−60
−40
−20
0
20
40
60
Position relative to center (bp)

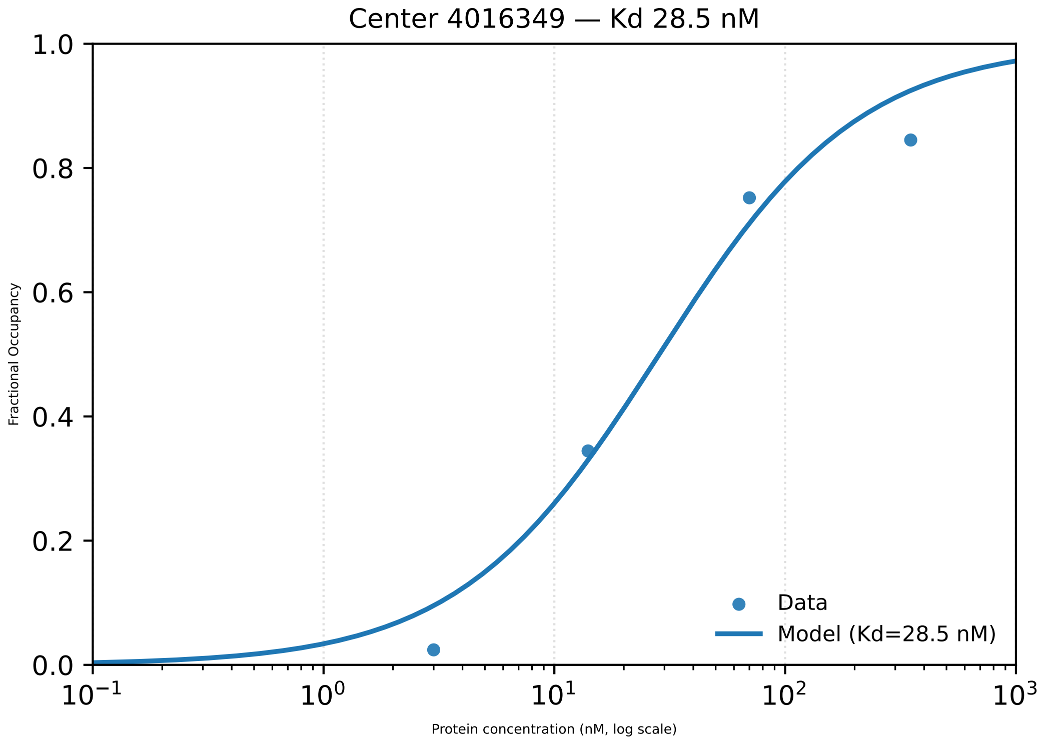

Center 4016349 — Kd 28.5 nM
1.0
0.8
0.6
0.4
0.2
0.0
Fractional Occupancy
Data
Model (Kd=28.5 nM)
$10^{-1}$
$10^{0}$
$10^{1}$
$10^{2}$
$10^{3}$
Protein concentration (nM, log scale)

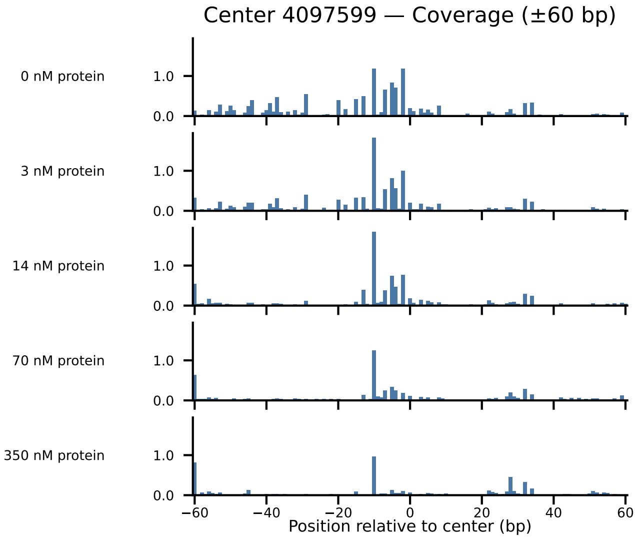
Center 4097599 — Coverage (±60 bp)
0 nM protein
3 nM protein
14 nM protein
70 nM protein
350 nM protein
1.0
0.0
−60
−40
−20
0
20
40
60
Position relative to center (bp)

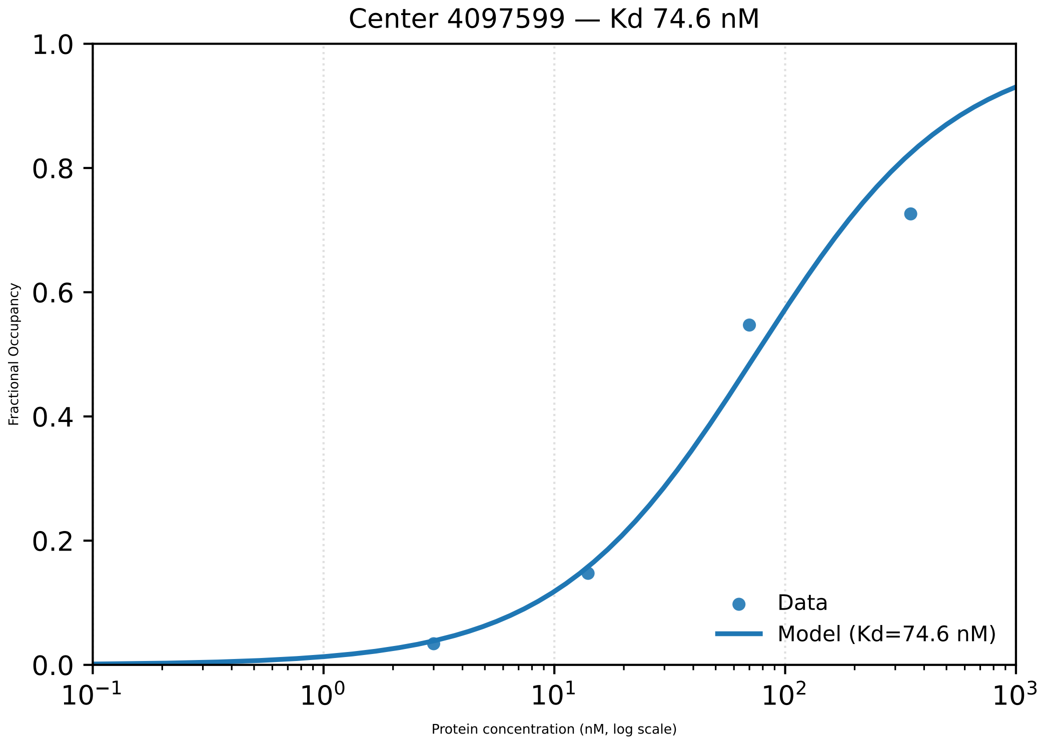

Center 4097599 — Kd 74.6 nM
1.0
0.8
0.6
0.4
0.2
0.0
Fractional Occupancy
$10^{-1}$
$10^{0}$
$10^{1}$
$10^{2}$
$10^{3}$
Protein concentration (nM, log scale)
Data
Model (Kd=74.6 nM)

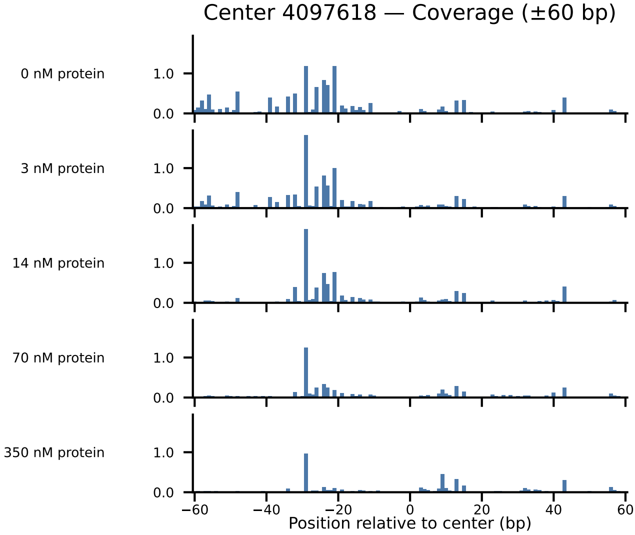

Center 4097618 — Coverage (±60 bp)
0 nM protein
3 nM protein
14 nM protein
70 nM protein
350 nM protein
1.0
0.0
−60
−40
−20
0
20
40
60
Position relative to center (bp)

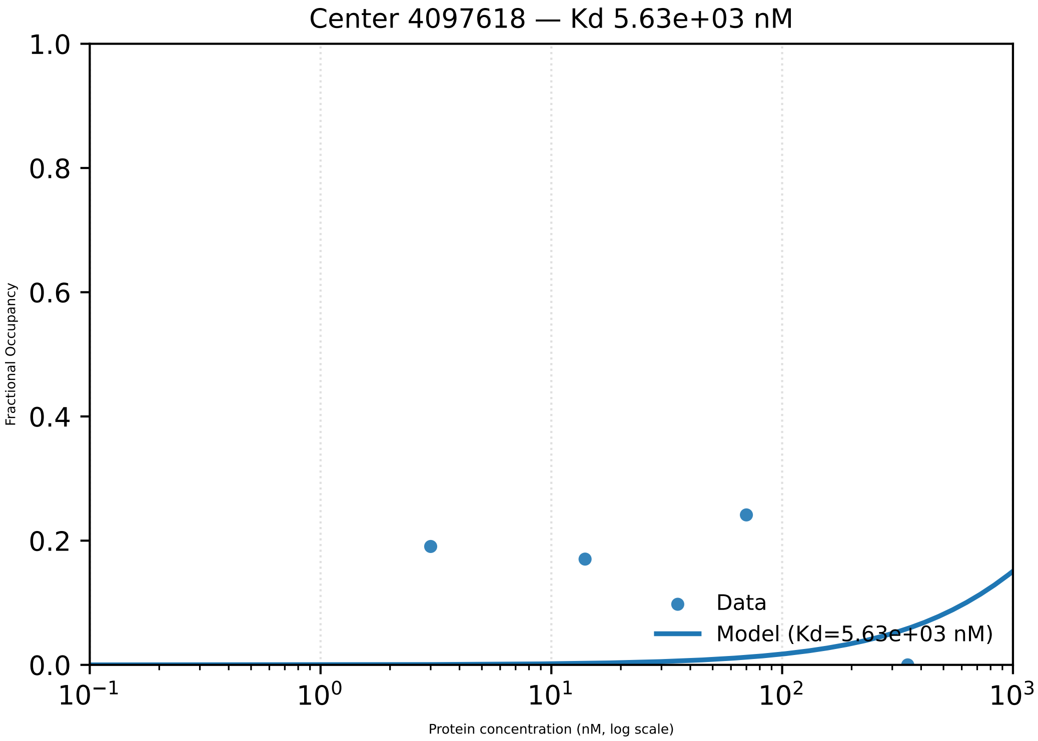

Center 4097618 — Kd 5.63e+03 nM
1.0
0.8
0.6
0.4
0.2
0.0
Fractional Occupancy
$10^{-1}$
$10^{0}$
$10^{1}$
$10^{2}$
$10^{3}$
Protein concentration (nM, log scale)
Data
Model (Kd=5.63e+03 nM)

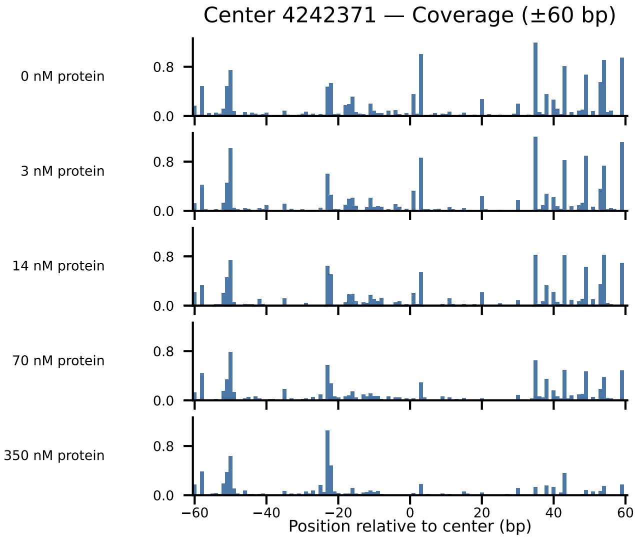

Center 4242371 — Coverage (±60 bp)
0 nM protein
3 nM protein
14 nM protein
70 nM protein
350 nM protein
0.8
0.0
−60
−40
−20
0
20
40
60
Position relative to center (bp)

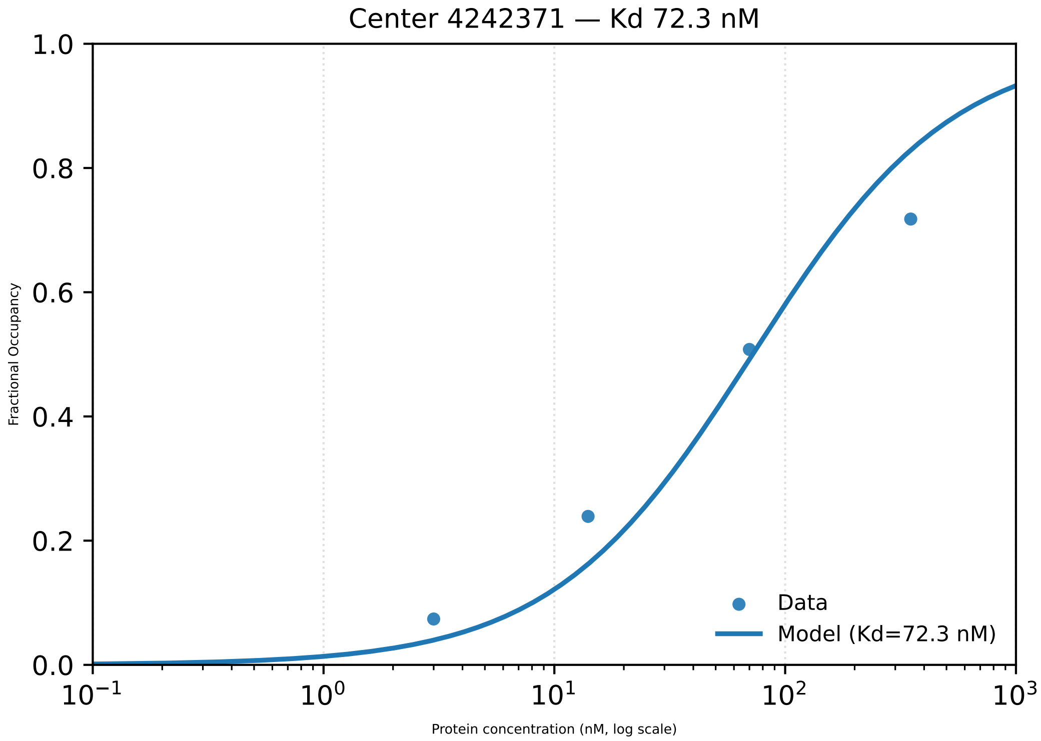

Center 4242371 — Kd 72.3 nM
1.0
0.8
0.6
0.4
0.2
0.0
Fractional Occupancy
$10^{-1}$
$10^{0}$
$10^{1}$
$10^{2}$
$10^{3}$
Protein concentration (nM, log scale)
Data
Model (Kd=72.3 nM)

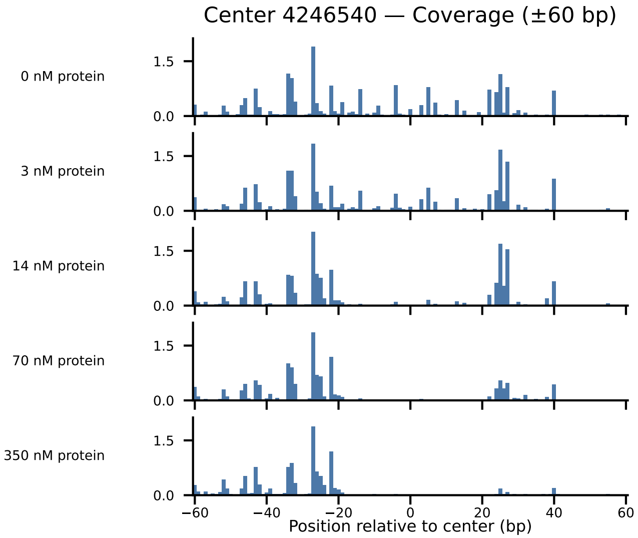
Center 4246540 — Coverage (±60 bp)
0 nM protein
3 nM protein
14 nM protein
70 nM protein
350 nM protein
1.5
0.0
−60
−40
−20
0
20
40
60
Position relative to center (bp)

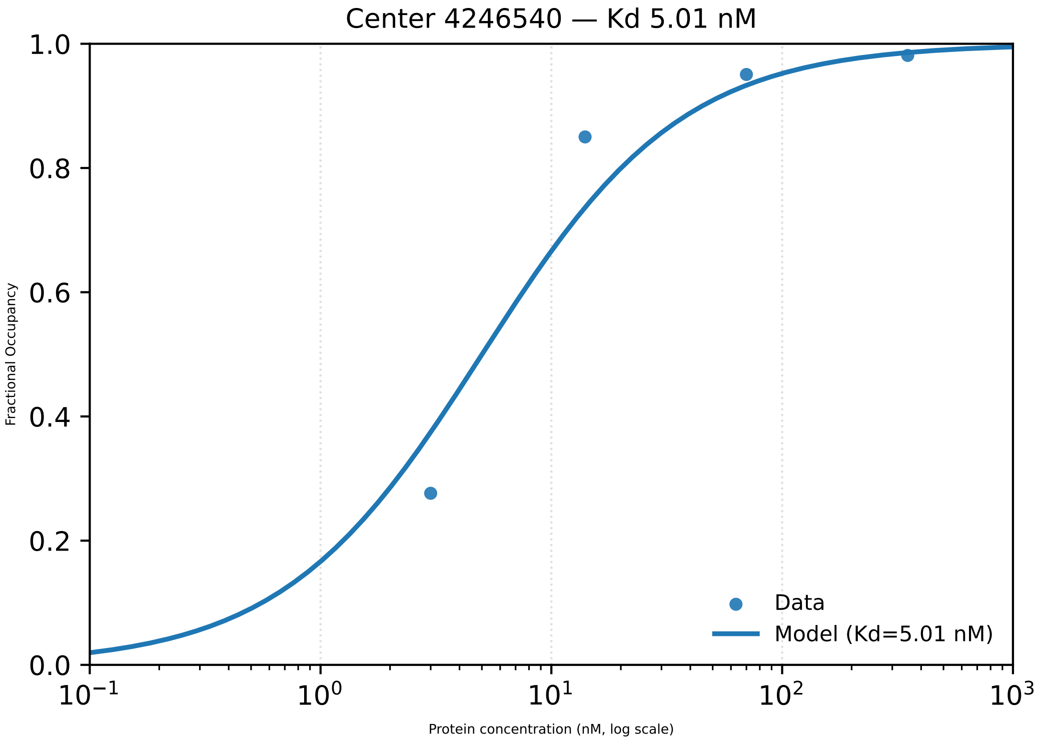

Center 4246540 — Kd 5.01 nM
Fractional Occupancy
1.0
0.8
0.6
0.4
0.2
0.0
$10^{-1}$
$10^{0}$
$10^{1}$
$10^{2}$
$10^{3}$
Protein concentration (nM, log scale)
Data
Model (Kd=5.01 nM)

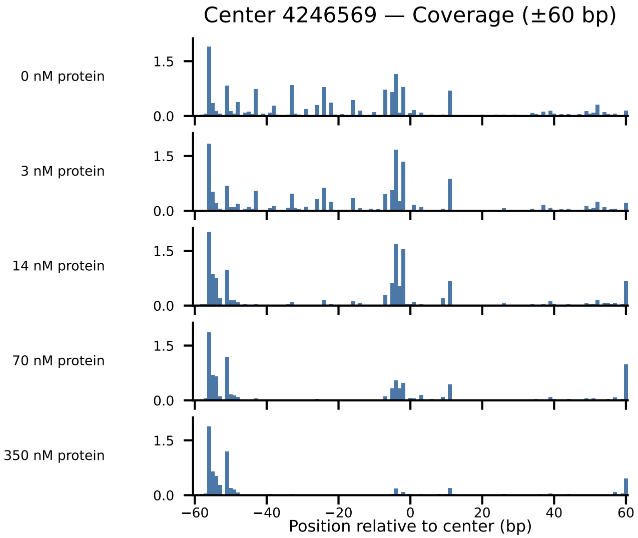
Center 4246569 — Coverage (±60 bp)
0 nM protein
3 nM protein
14 nM protein
70 nM protein
350 nM protein
1.5
0.0
−60
−40
−20
0
20
40
60
Position relative to center (bp)

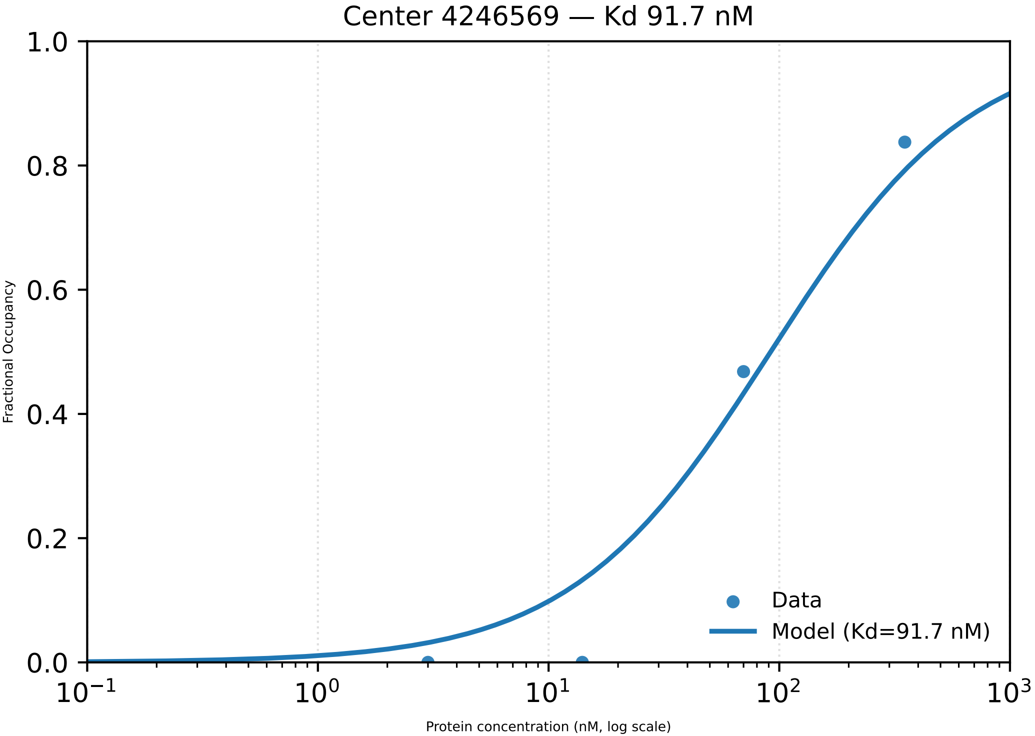

Center 4246569 — Kd 91.7 nM
1.0
0.8
0.6
0.4
0.2
0.0
Fractional Occupancy
Data
Model (Kd=91.7 nM)
$10^{-1}$
$10^{0}$
$10^{1}$
$10^{2}$
$10^{3}$
Protein concentration (nM, log scale)

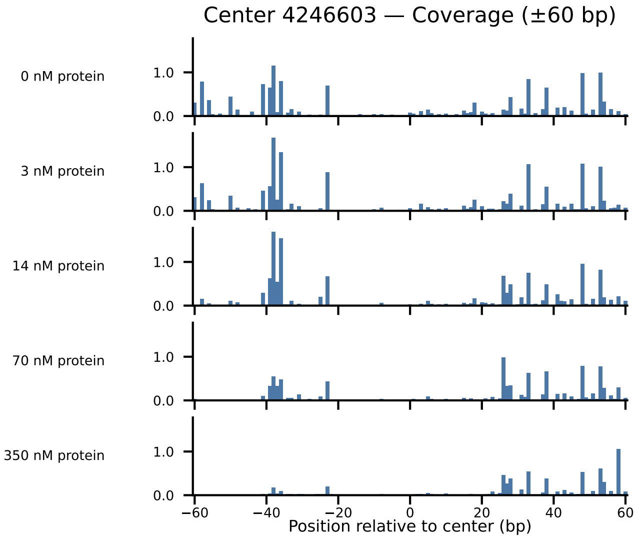

Center 4246603 — Coverage (±60 bp)
0 nM protein
3 nM protein
14 nM protein
70 nM protein
350 nM protein
1.0
0.0
−60
−40
−20
0
20
40
60
Position relative to center (bp)

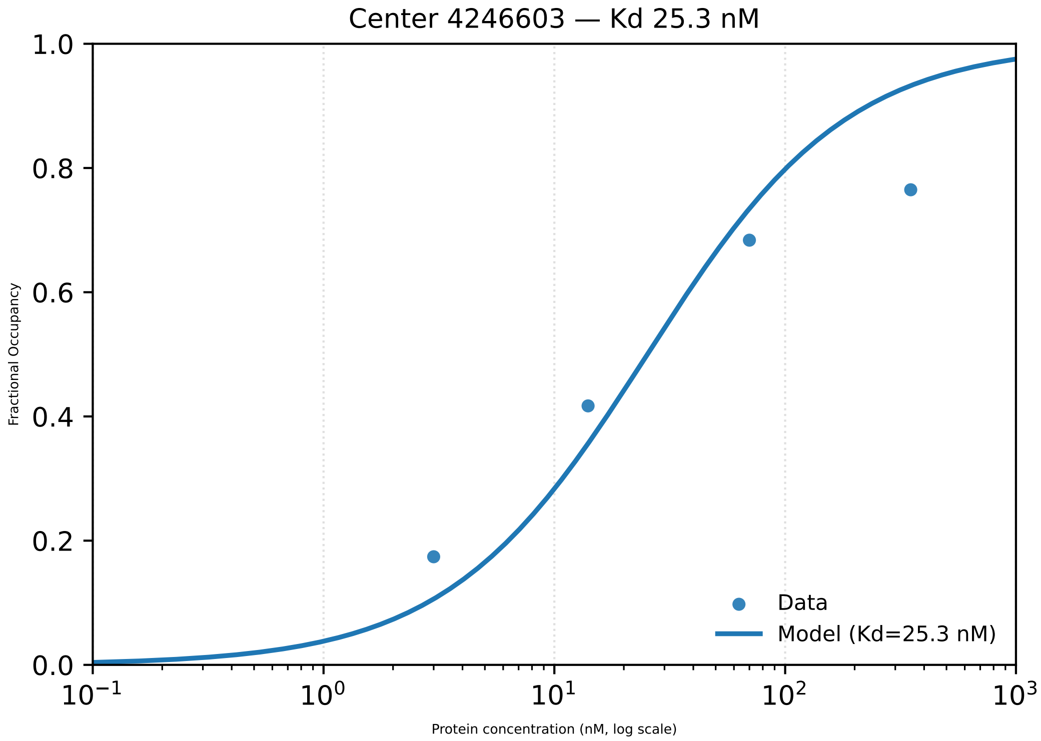

Center 4246603 — Kd 25.3 nM
1.0
0.8
0.6
0.4
0.2
0.0
Fractional Occupancy
$10^{-1}$
$10^{0}$
$10^{1}$
$10^{2}$
$10^{3}$
Protein concentration (nM, log scale)
Data
Model (Kd=25.3 nM)

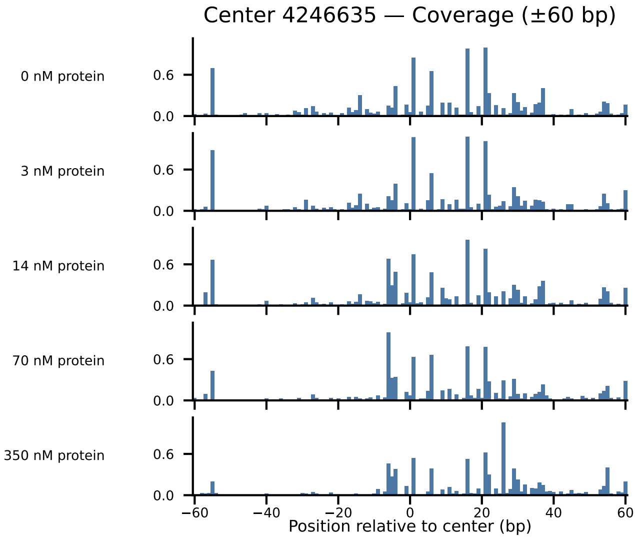

Center 4246635 — Coverage (±60 bp)
0 nM protein
3 nM protein
14 nM protein
70 nM protein
350 nM protein
0.6
0.0
−60
−40
−20
0
20
40
60
Position relative to center (bp)

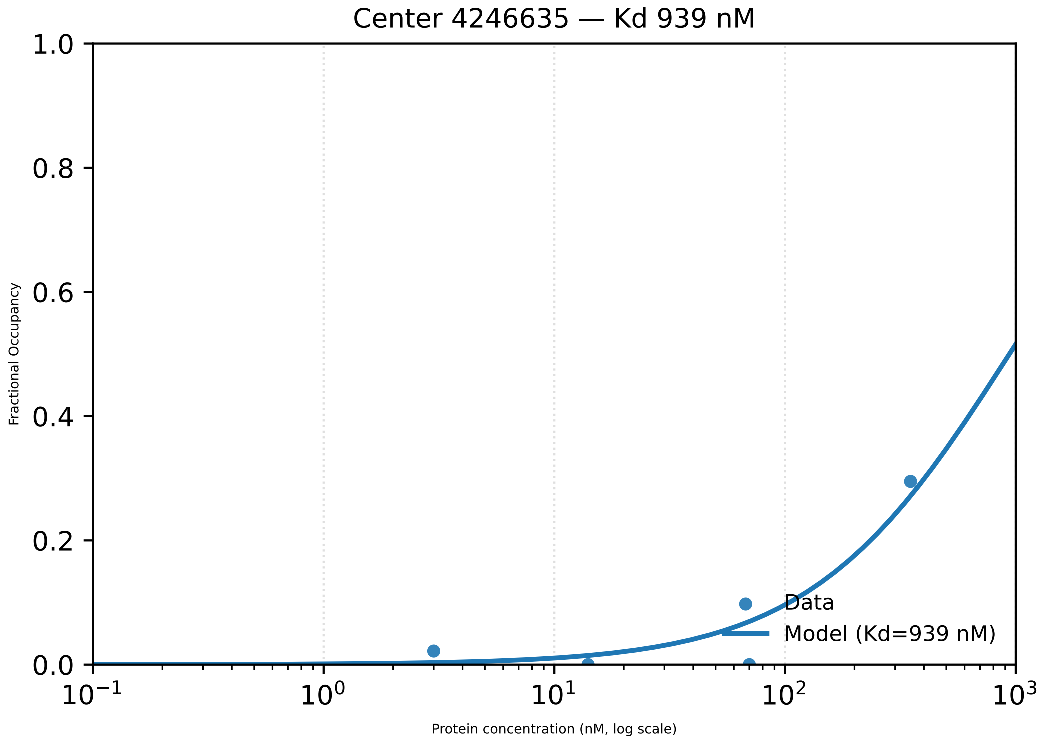
Center 4246635 — Kd 939 nM
1.0
0.8
0.6
0.4
0.2
0.0
Fractional Occupancy
$10^{-1}$
$10^{0}$
$10^{1}$
$10^{2}$
$10^{3}$
Protein concentration (nM, log scale)
Data
Model (Kd=939 nM)

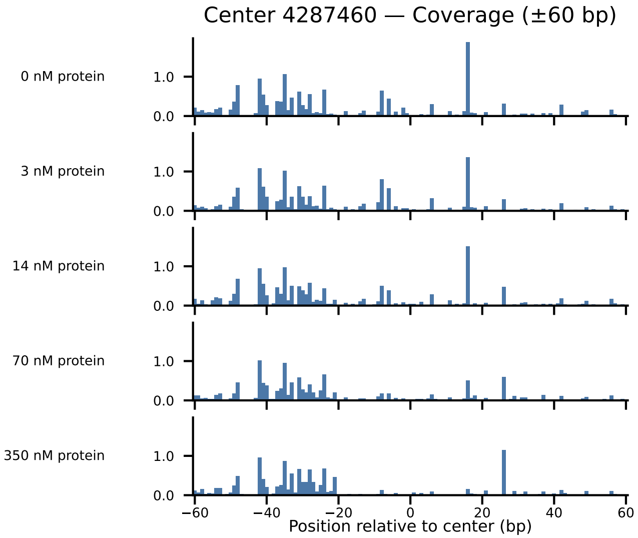

Center 4287460 — Coverage (±60 bp)
0 nM protein
3 nM protein
14 nM protein
70 nM protein
350 nM protein
1.0
0.0
−60
−40
−20
0
20
40
60
Position relative to center (bp)

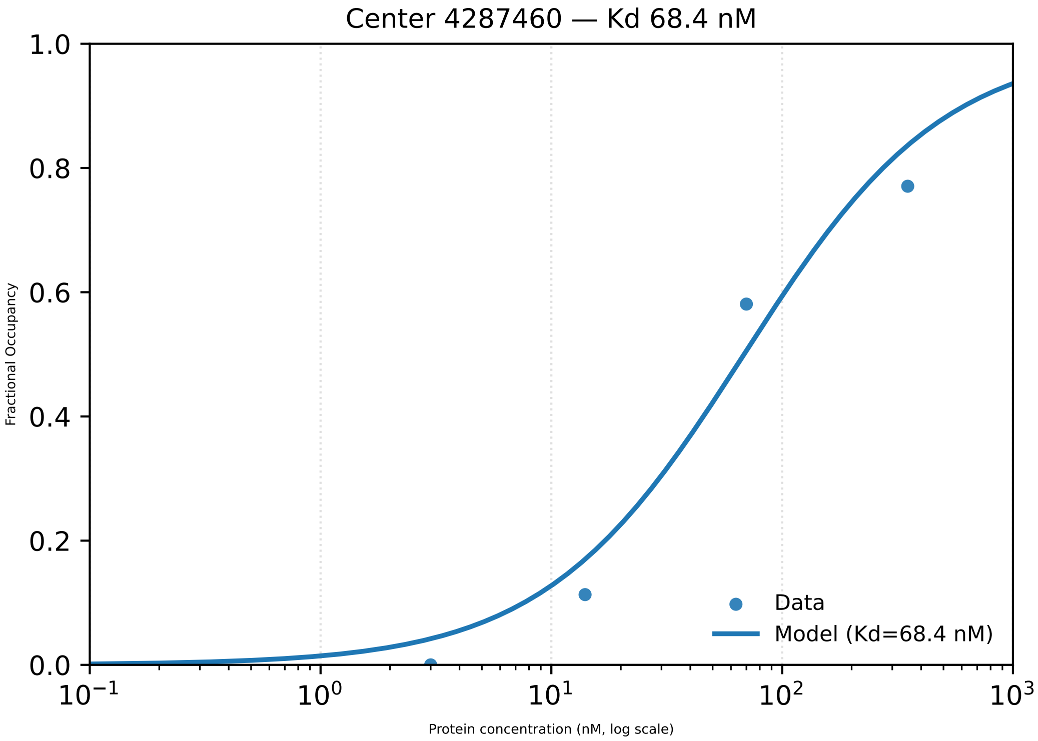

Center 4287460 — Kd 68.4 nM
Fractional Occupancy
1.0
0.8
0.6
0.4
0.2
0.0
$10^{-1}$
$10^{0}$
$10^{1}$
$10^{2}$
$10^{3}$
Protein concentration (nM, log scale)
Data
Model (Kd=68.4 nM)

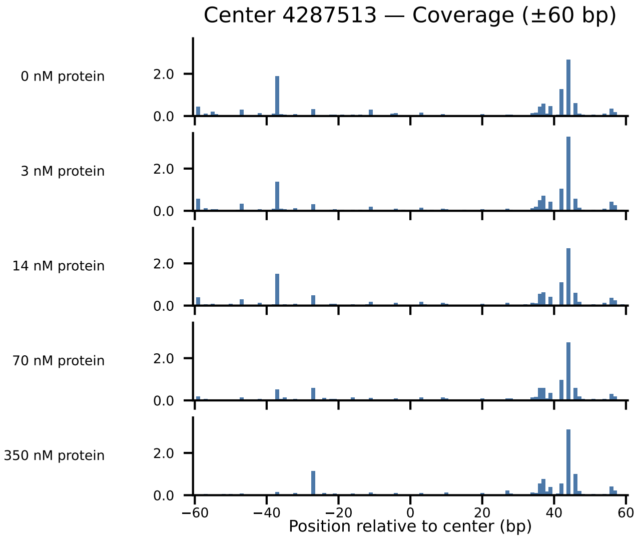
Center 4287513 — Coverage (±60 bp)
0 nM protein
3 nM protein
14 nM protein
70 nM protein
350 nM protein
2.0
0.0
−60
−40
−20
0
20
40
60
Position relative to center (bp)

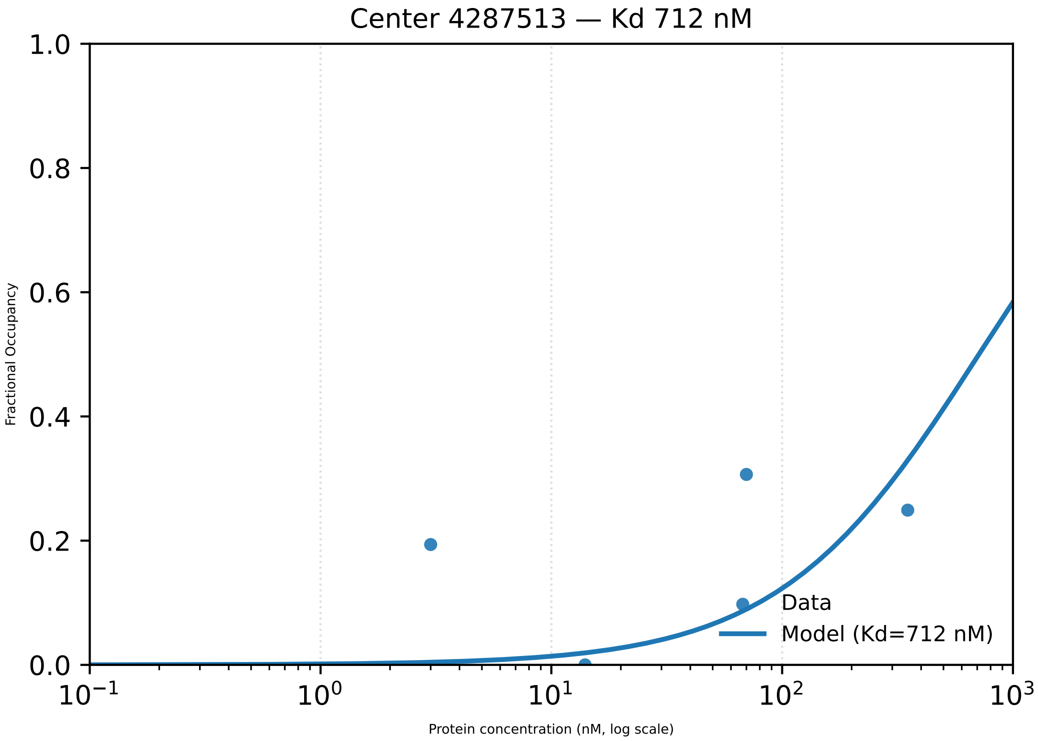

Center 4287513 — Kd 712 nM
Fractional Occupancy
Protein concentration (nM, log scale)
Data
Model (Kd=712 nM)
1.0
0.8
0.6
0.4
0.2
0.0
$10^{-1}$
$10^{0}$
$10^{1}$
$10^{2}$
$10^{3}$

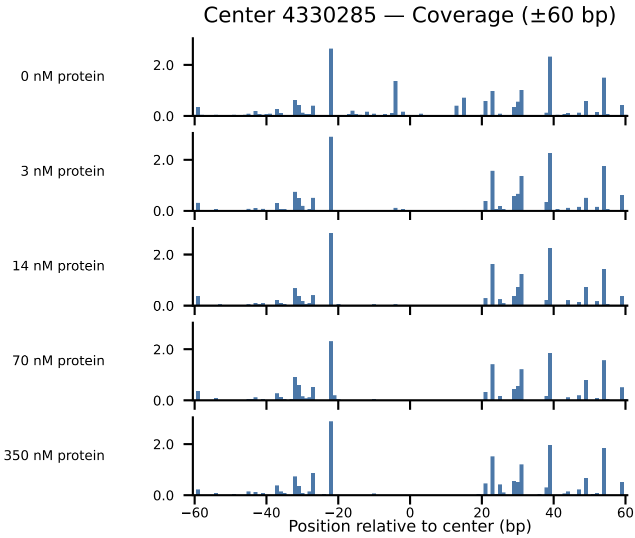

Center 4330285 — Coverage (±60 bp)
0 nM protein
3 nM protein
14 nM protein
70 nM protein
350 nM protein
2.0
0.0
−60
−40
−20
0
20
40
60
Position relative to center (bp)

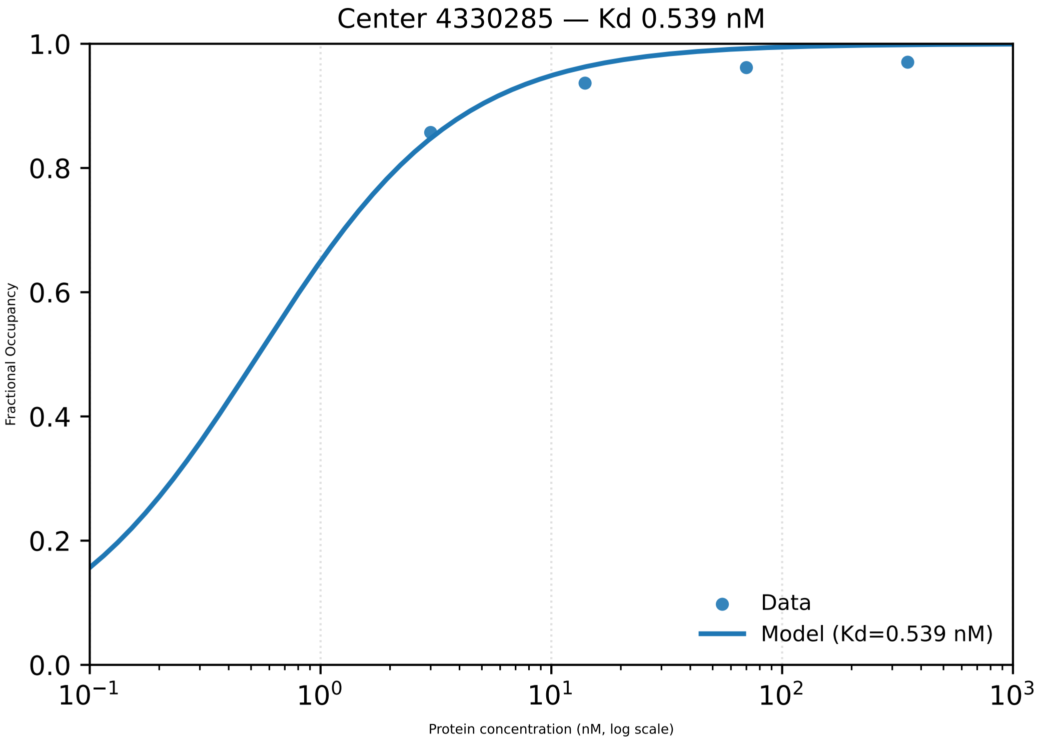

Center 4330285 — Kd 0.539 nM
1.0
0.8
0.6
0.4
0.2
0.0
Fractional Occupancy
$10^{-1}$
$10^{0}$
$10^{1}$
$10^{2}$
$10^{3}$
Protein concentration (nM, log scale)
Data
Model (Kd=0.539 nM)

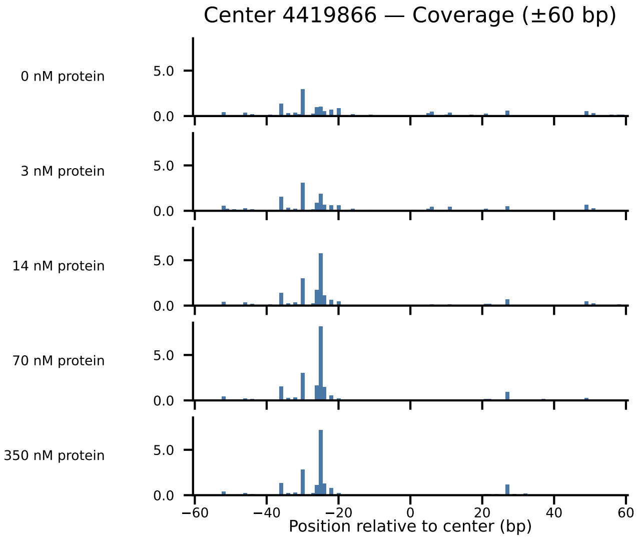

Center 4419866 — Coverage (±60 bp)
0 nM protein
3 nM protein
14 nM protein
70 nM protein
350 nM protein
5.0
0.0
−60
−40
−20
0
20
40
60
Position relative to center (bp)

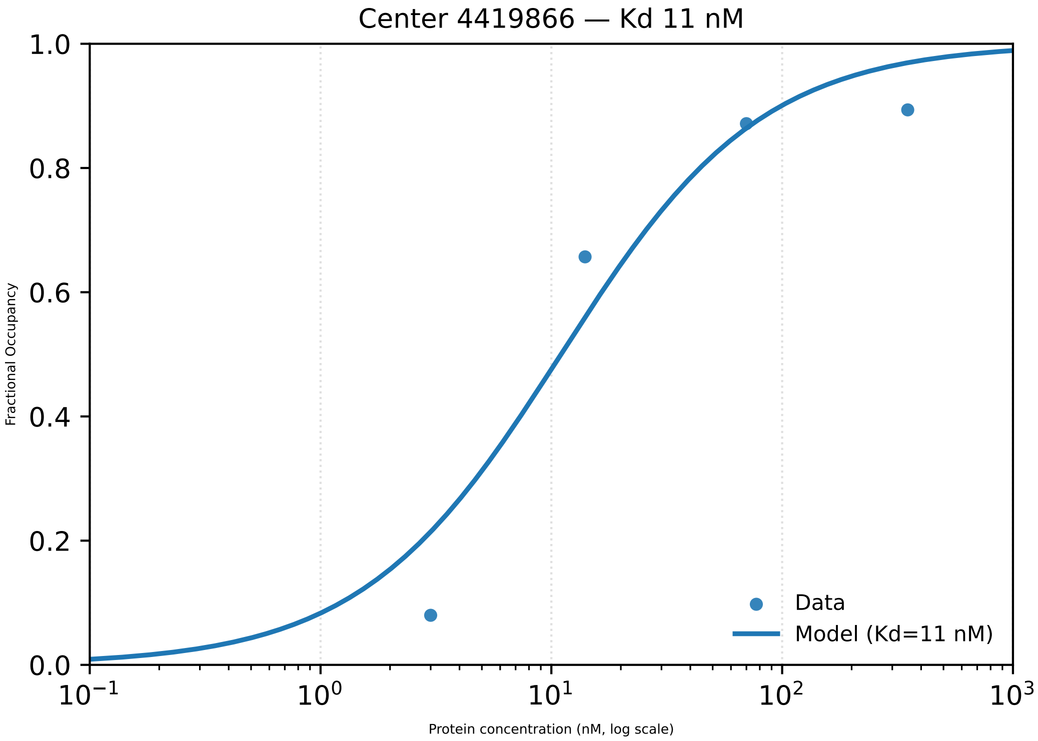

Center 4419866 — Kd 11 nM
1.0
0.8
0.6
0.4
0.2
0.0
Fractional Occupancy
$10^{-1}$
$10^{0}$
$10^{1}$
$10^{2}$
$10^{3}$
Protein concentration (nM, log scale)
Data
Model (Kd=11 nM)

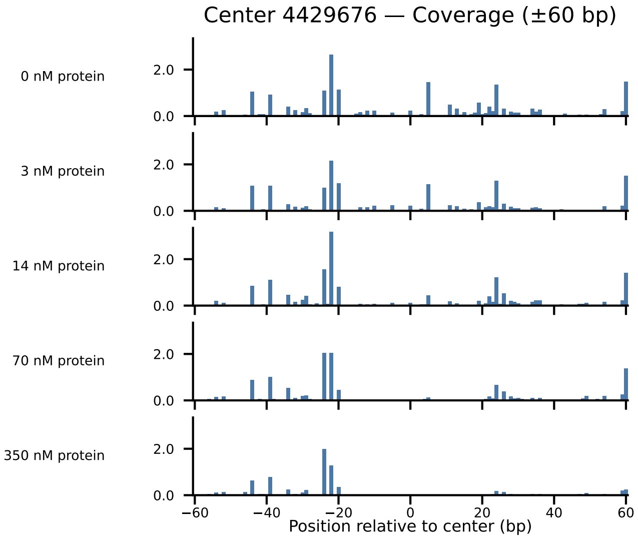

Center 4429676 — Coverage (±60 bp)
0 nM protein
3 nM protein
14 nM protein
70 nM protein
350 nM protein
2.0
0.0
−60
−40
−20
0
20
40
60
Position relative to center (bp)

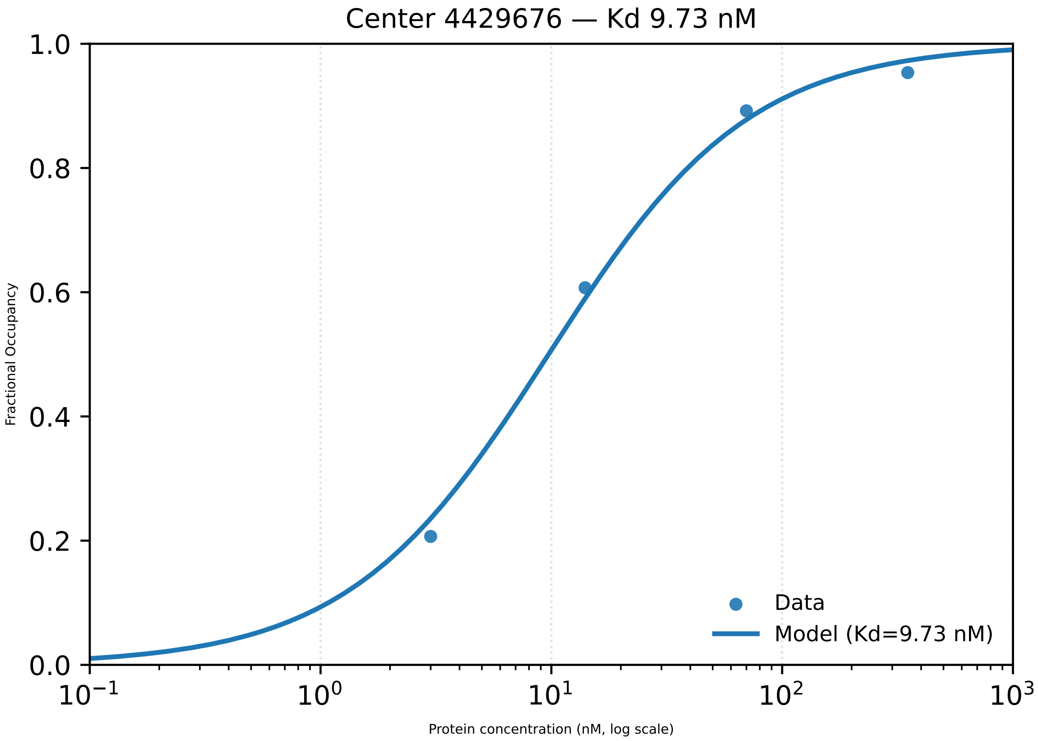

Center 4429676 — Kd 9.73 nM
1.0
0.8
0.6
0.4
0.2
0.0
Fractional Occupancy
$10^{-1}$
$10^{0}$
$10^{1}$
$10^{2}$
$10^{3}$
Protein concentration (nM, log scale)
Data
Model (Kd=9.73 nM)

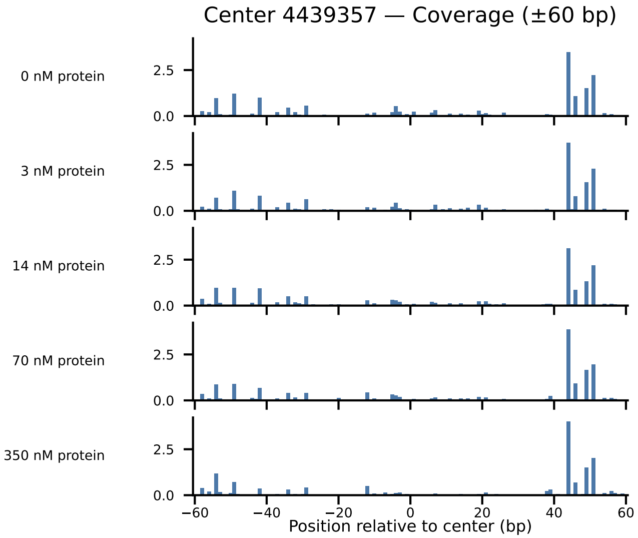

Center 4439357 — Coverage (±60 bp)
0 nM protein
3 nM protein
14 nM protein
70 nM protein
350 nM protein
2.5
0.0
−60
−40
−20
0
20
40
60
Position relative to center (bp)

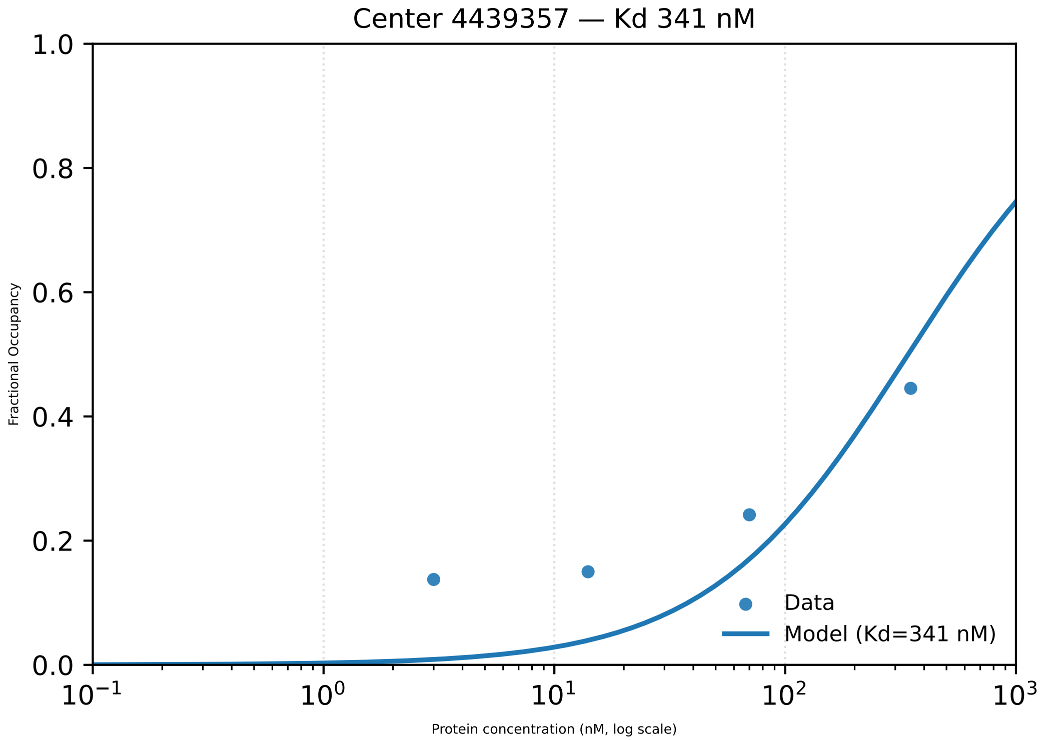

Center 4439357 — Kd 341 nM
1.0
0.8
0.6
0.4
0.2
0.0
Fractional Occupancy
$10^{-1}$
$10^{0}$
$10^{1}$
$10^{2}$
$10^{3}$
Protein concentration (nM, log scale)
Data
Model (Kd=341 nM)

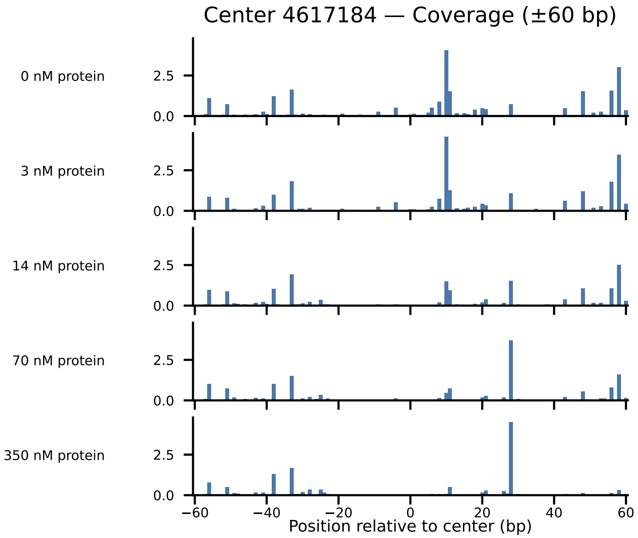

Center 4617184 — Coverage (±60 bp)
0 nM protein
3 nM protein
14 nM protein
70 nM protein
350 nM protein
2.5
0.0
−60
−40
−20
0
20
40
60
Position relative to center (bp)

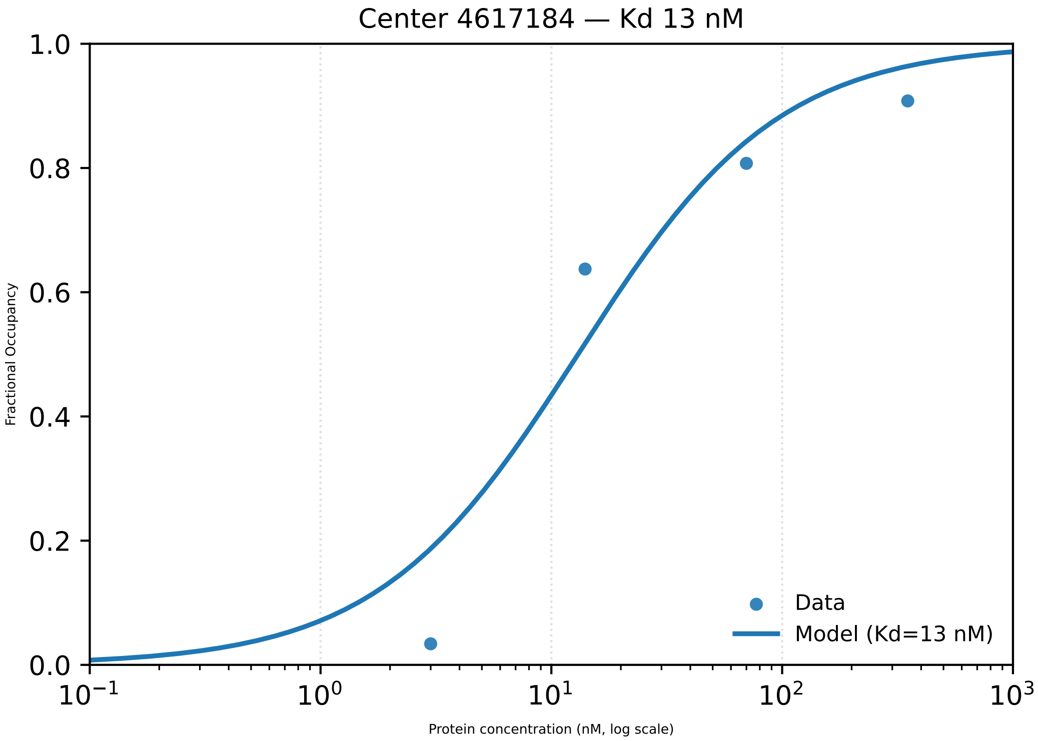

Center 4617184 — Kd 13 nM
Fractional Occupancy
Protein concentration (nM, log scale)
Data
Model (Kd=13 nM)
0.0
0.2
0.4
0.6
0.8
1.0
$10^{-1}$
$10^{0}$
$10^{1}$
$10^{2}$
$10^{3}$

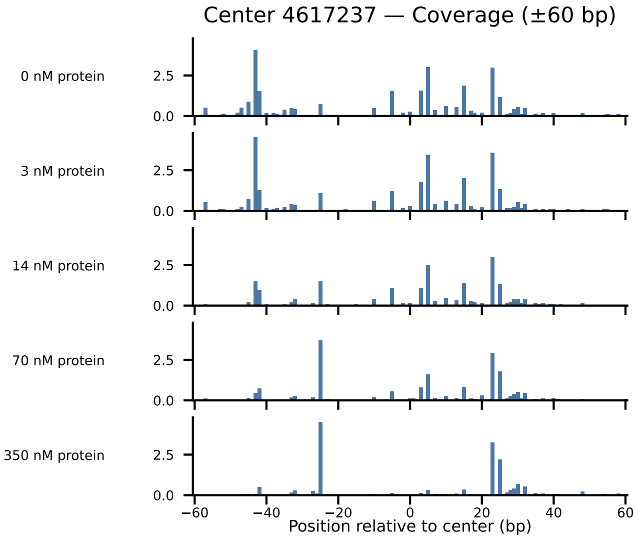
Center 4617237 — Coverage (±60 bp)
0 nM protein
3 nM protein
14 nM protein
70 nM protein
350 nM protein
0.0
2.5
−60
−40
−20
0
20
40
60
Position relative to center (bp)

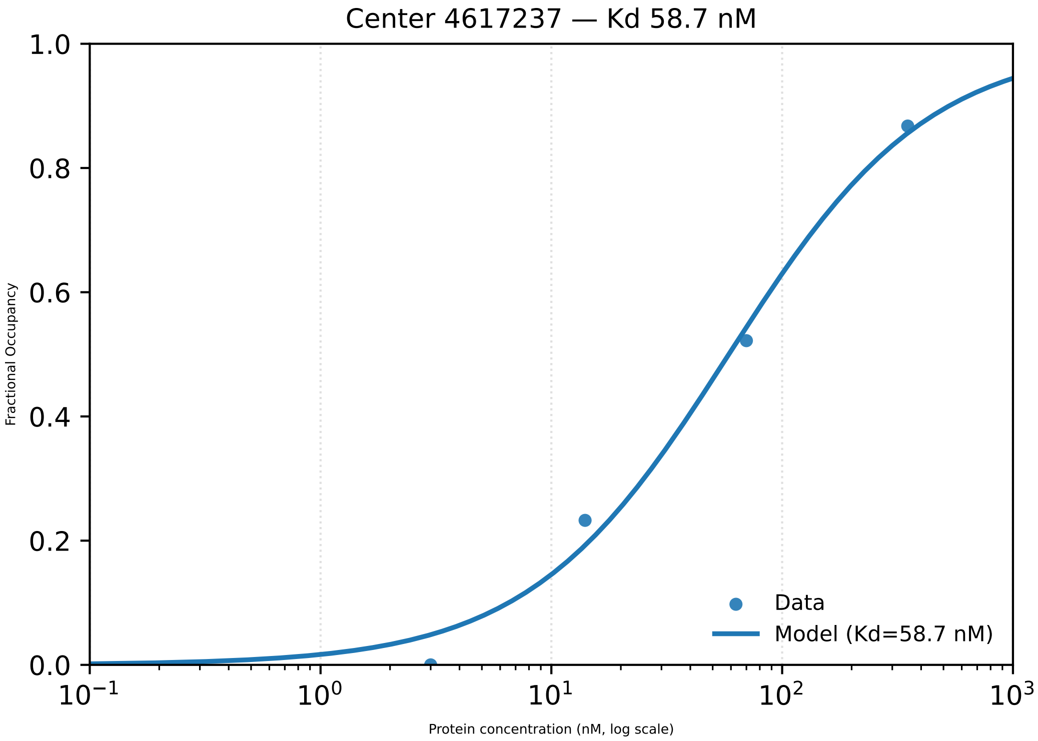

Center 4617237 — Kd 58.7 nM
Fractional Occupancy
1.0
0.8
0.6
0.4
0.2
0.0
$10^{-1}$
$10^{0}$
$10^{1}$
$10^{2}$
$10^{3}$
Protein concentration (nM, log scale)
Data
Model (Kd=58.7 nM)
