## Supplementary material for "Footprint-seq: a simple method to quantitatively map *in vitro* protein-DNA interactions on a genome-wide scale at high spatial resolution": Figure S4

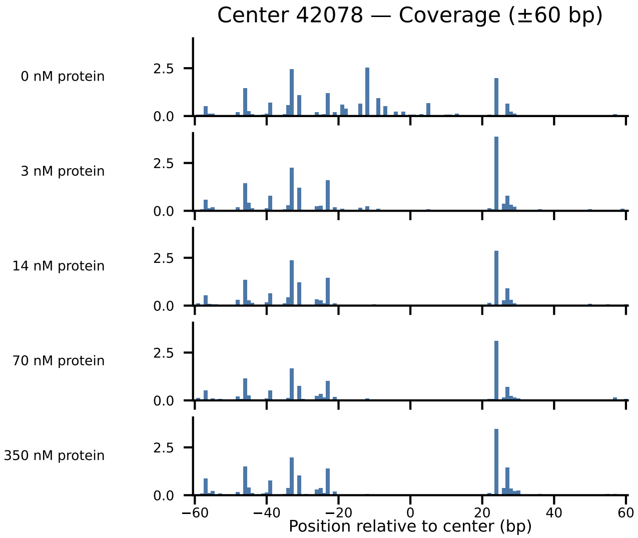

Center 42078 — Coverage (±60 bp)
0 nM protein
3 nM protein
14 nM protein
70 nM protein
350 nM protein
2.5
0.0
−60
−40
−20
0
20
40
60
Position relative to center (bp)

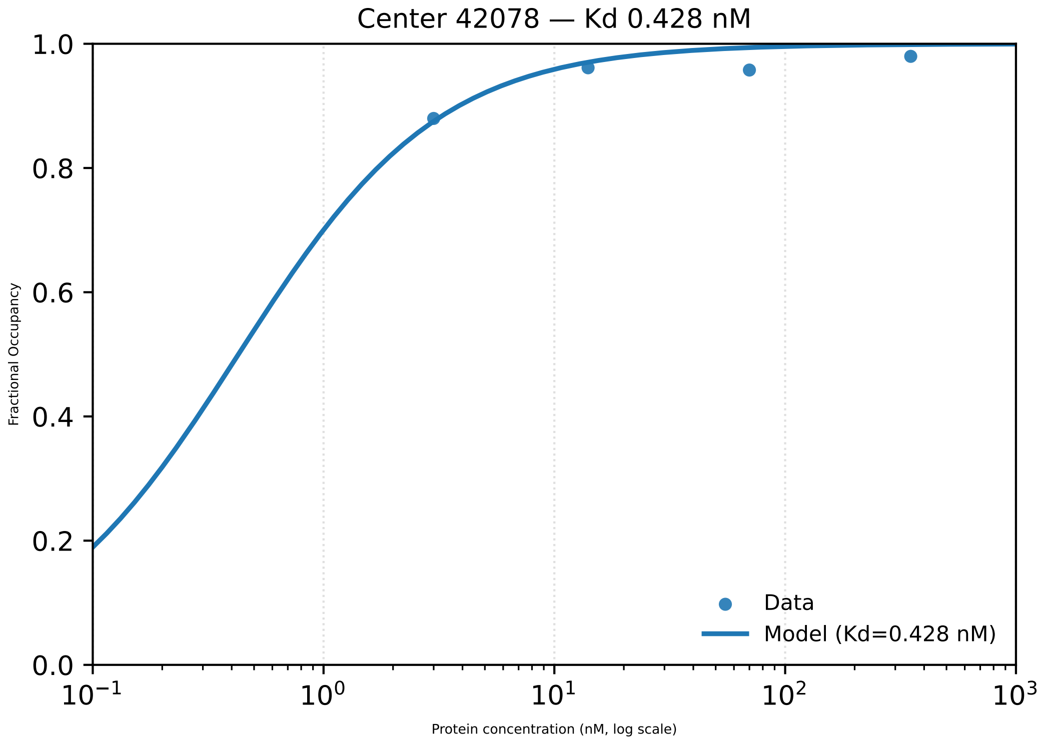

Center 42078 — Kd 0.428 nM
1.0
0.8
0.6
0.4
0.2
0.0
Fractional Occupancy
$10^{-1}$
$10^{0}$
$10^{1}$
$10^{2}$
$10^{3}$
Protein concentration (nM, log scale)
Data
Model (Kd=0.428 nM)

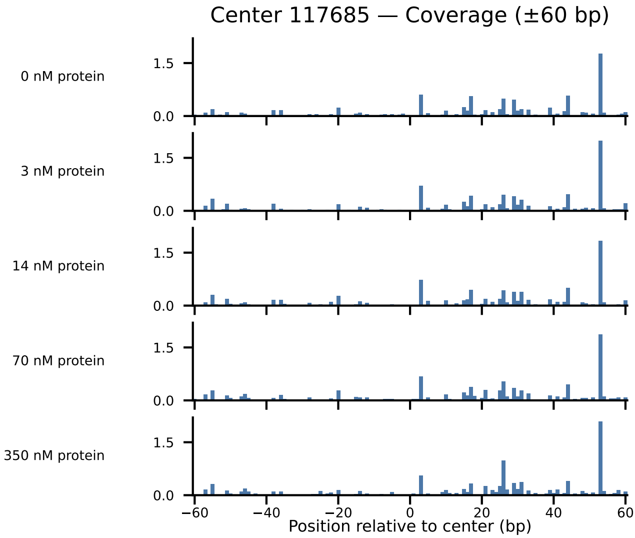

Center 117685 — Coverage (±60 bp)
0 nM protein
3 nM protein
14 nM protein
70 nM protein
350 nM protein
1.5
0.0
−60
−40
−20
0
20
40
60
Position relative to center (bp)

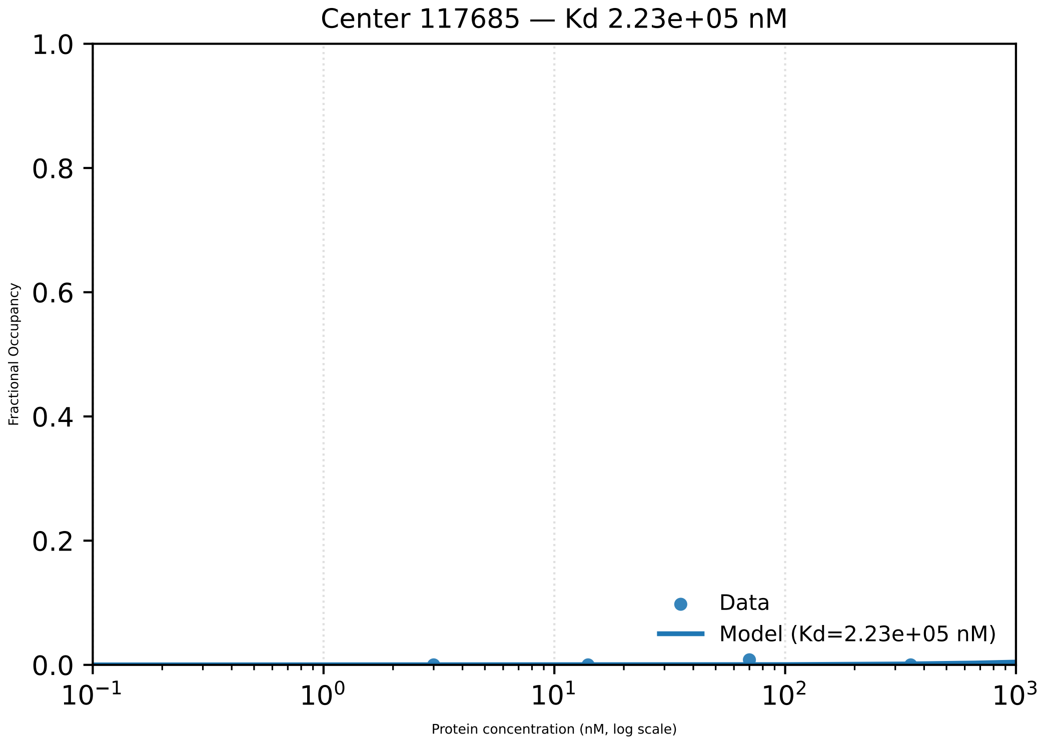

Center 117685 — Kd 2.23e+05 nM
Fractional Occupancy
Protein concentration (nM, log scale)
Data
Model (Kd=2.23e+05 nM)
1.0
0.8
0.6
0.4
0.2
0.0
$10^{-1}$
$10^{0}$
$10^{1}$
$10^{2}$
$10^{3}$

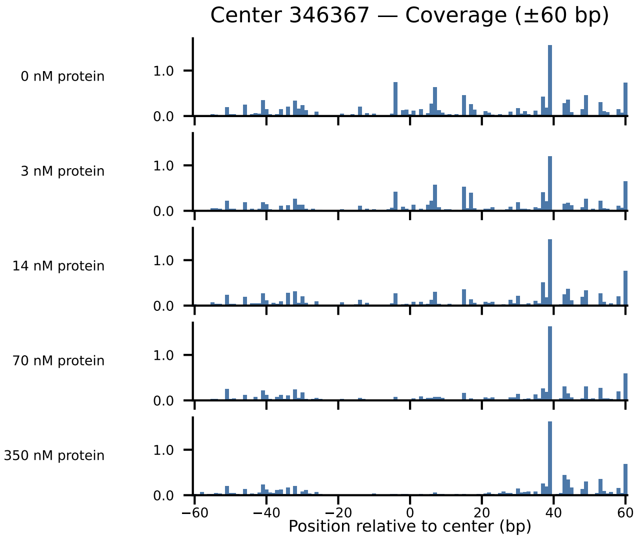

Center 346367 — Coverage (±60 bp)
0 nM protein
3 nM protein
14 nM protein
70 nM protein
350 nM protein
0.0
1.0
−60
−40
−20
0
20
40
60
Position relative to center (bp)

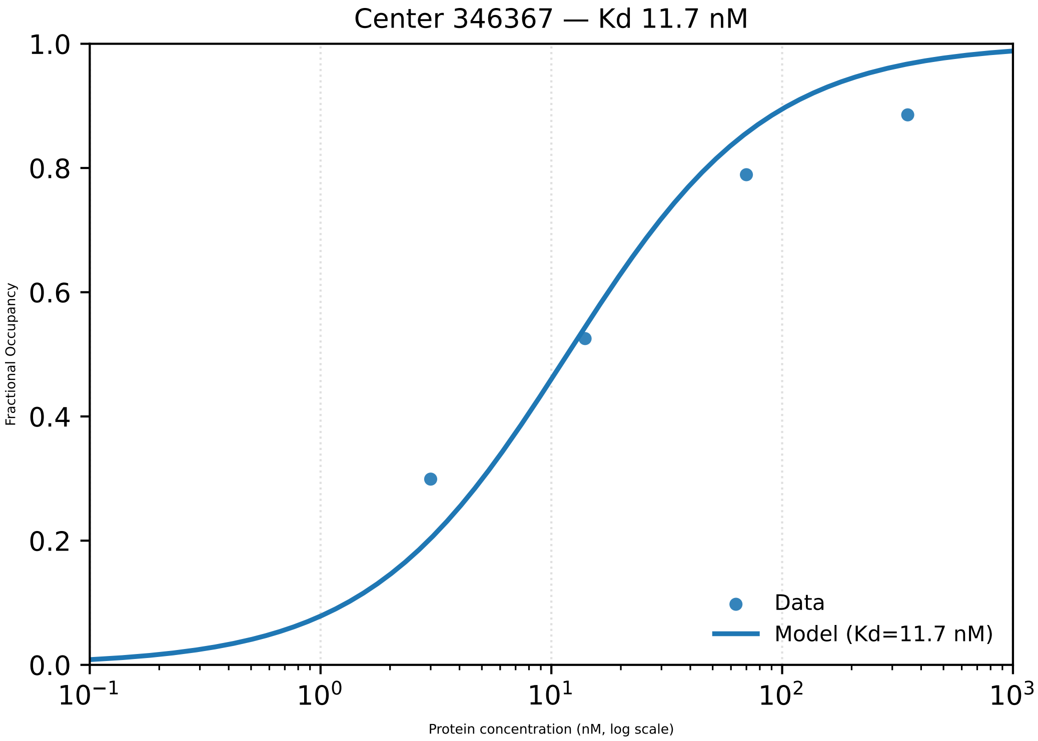

Center 346367 — Kd 11.7 nM
1.0
0.8
0.6
0.4
0.2
0.0
Fractional Occupancy
$10^{-1}$
$10^{0}$
$10^{1}$
$10^{2}$
$10^{3}$
Protein concentration (nM, log scale)
Data
Model (Kd=11.7 nM)

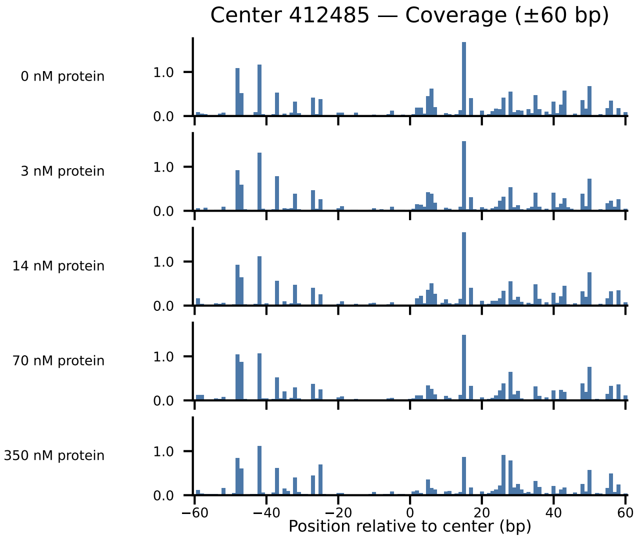

Center 412485 — Coverage (±60 bp)
0 nM protein
3 nM protein
14 nM protein
70 nM protein
350 nM protein
1.0
0.0
−60
−40
−20
0
20
40
60
Position relative to center (bp)

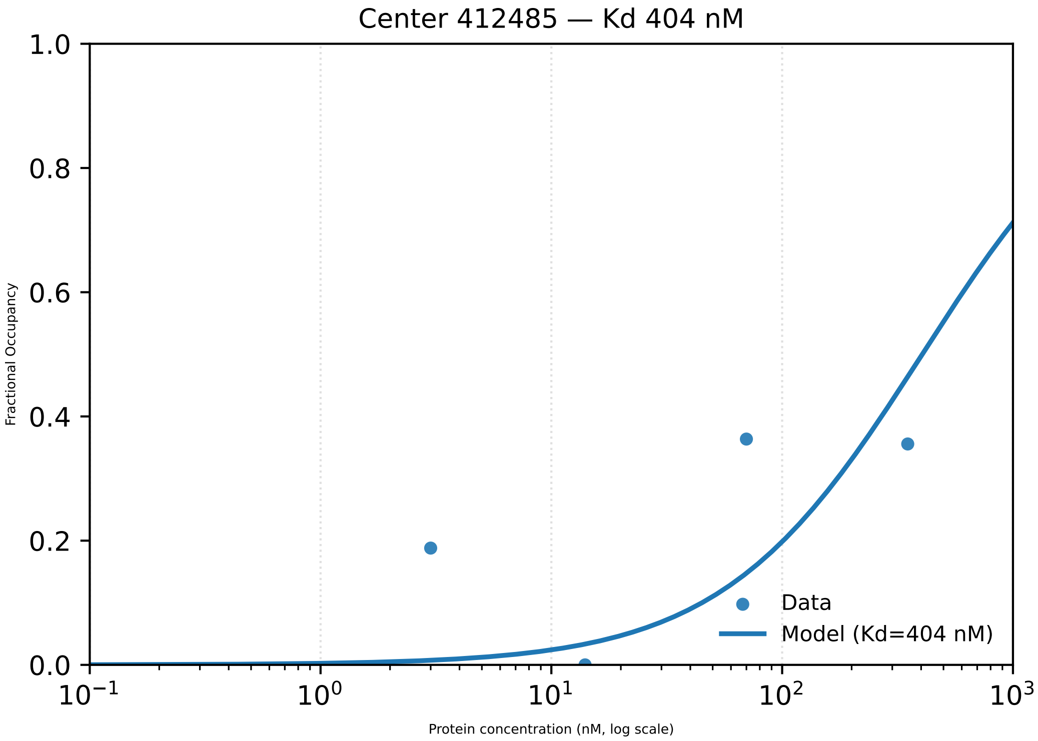

Center 412485 — Kd 404 nM
1.0
0.8
0.6
0.4
0.2
0.0
Fractional Occupancy
$10^{-1}$
$10^{0}$
$10^{1}$
$10^{2}$
$10^{3}$
Protein concentration (nM, log scale)
Data
Model (Kd=404 nM)

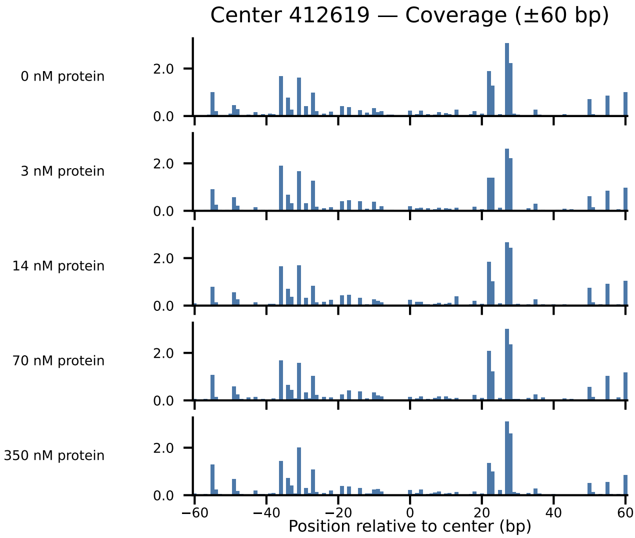
Center 412619 — Coverage (±60 bp)
0 nM protein
3 nM protein
14 nM protein
70 nM protein
350 nM protein
2.0
0.0
−60
−40
−20
0
20
40
60
Position relative to center (bp)

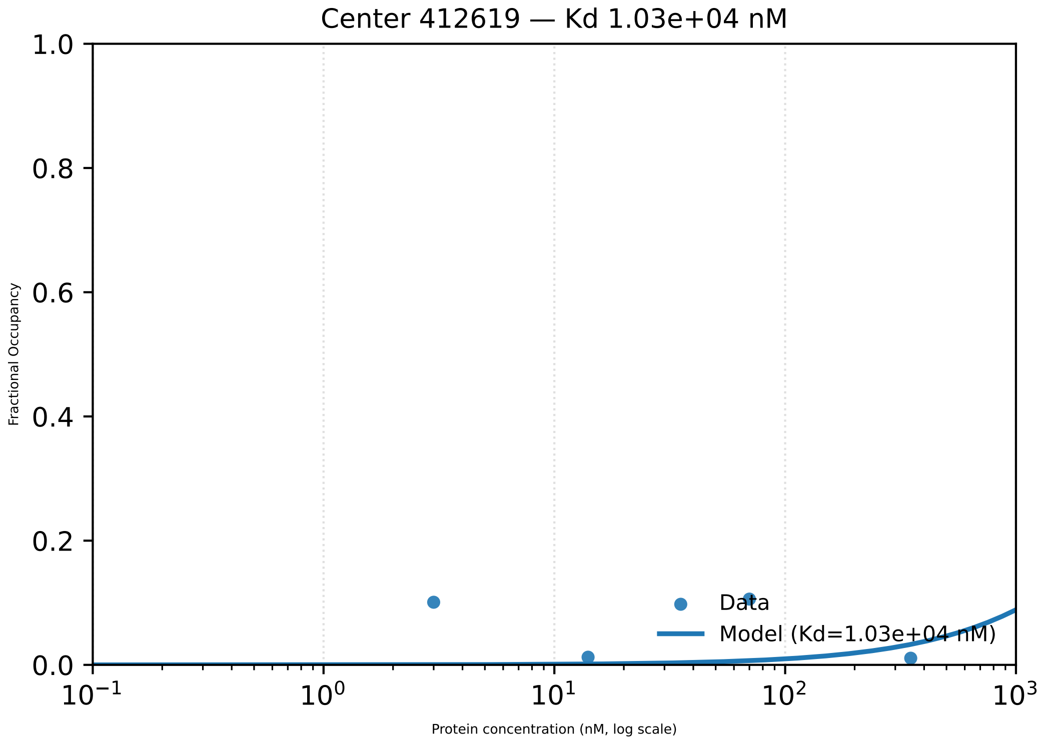

Center 412619 — Kd 1.03e+04 nM
Fractional Occupancy
1.0
0.8
0.6
0.4
0.2
0.0
$10^{-1}$
$10^{0}$
$10^{1}$
$10^{2}$
$10^{3}$
Protein concentration (nM, log scale)
Data
Model (Kd=1.03e+04 nM)

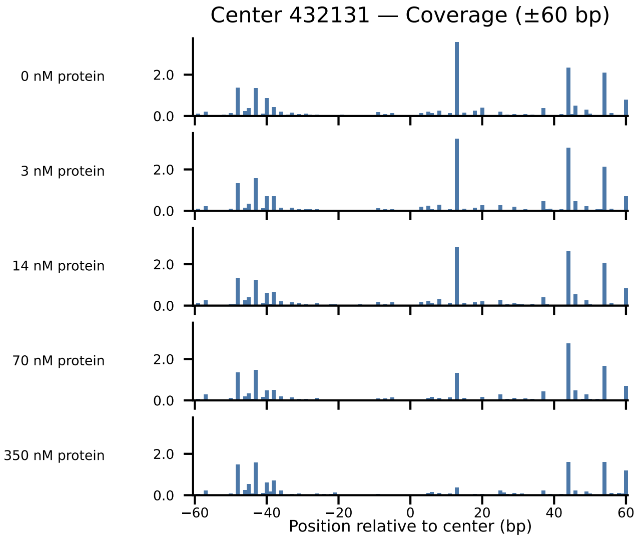

Center 432131 — Coverage (±60 bp)
0 nM protein
3 nM protein
14 nM protein
70 nM protein
350 nM protein
2.0
0.0
−60
−40
−20
0
20
40
60
Position relative to center (bp)

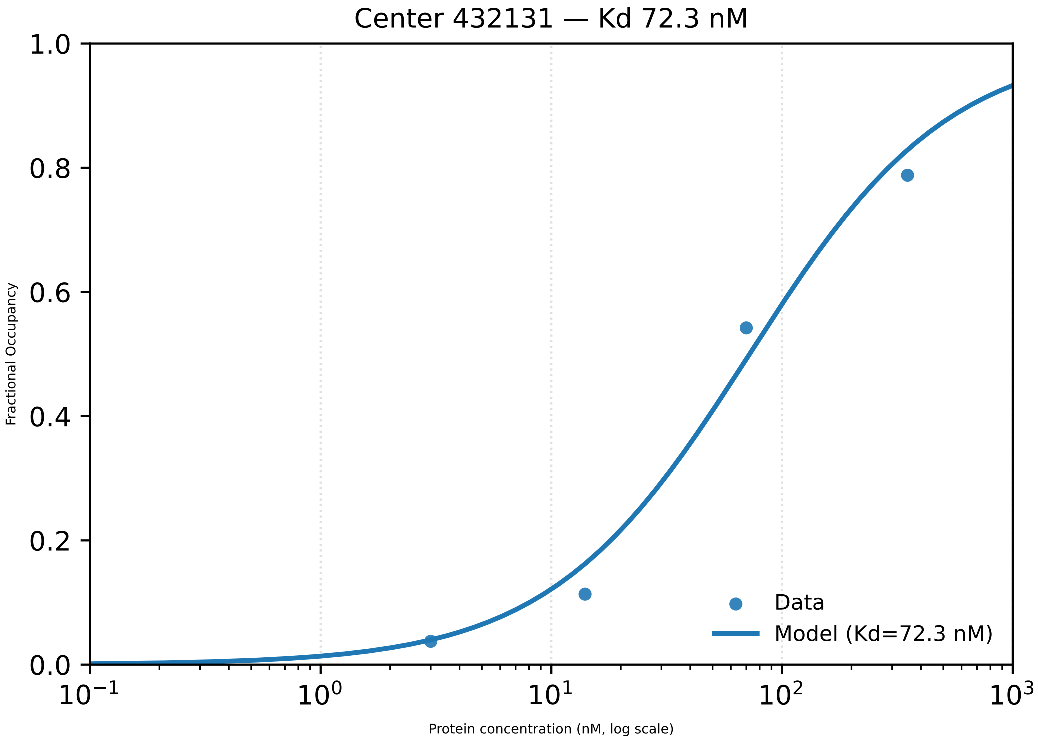

Center 432131 — Kd 72.3 nM
1.0
0.8
0.6
0.4
0.2
0.0
Fractional Occupancy
$10^{-1}$
$10^{0}$
$10^{1}$
$10^{2}$
$10^{3}$
Protein concentration (nM, log scale)
Data
Model (Kd=72.3 nM)

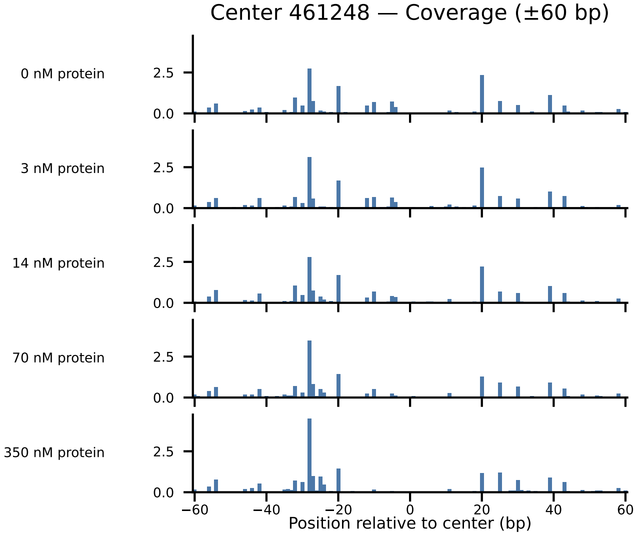

Center 461248 — Coverage (±60 bp)
0 nM protein
3 nM protein
14 nM protein
70 nM protein
350 nM protein
2.5
0.0
−60
−40
−20
0
20
40
60
Position relative to center (bp)

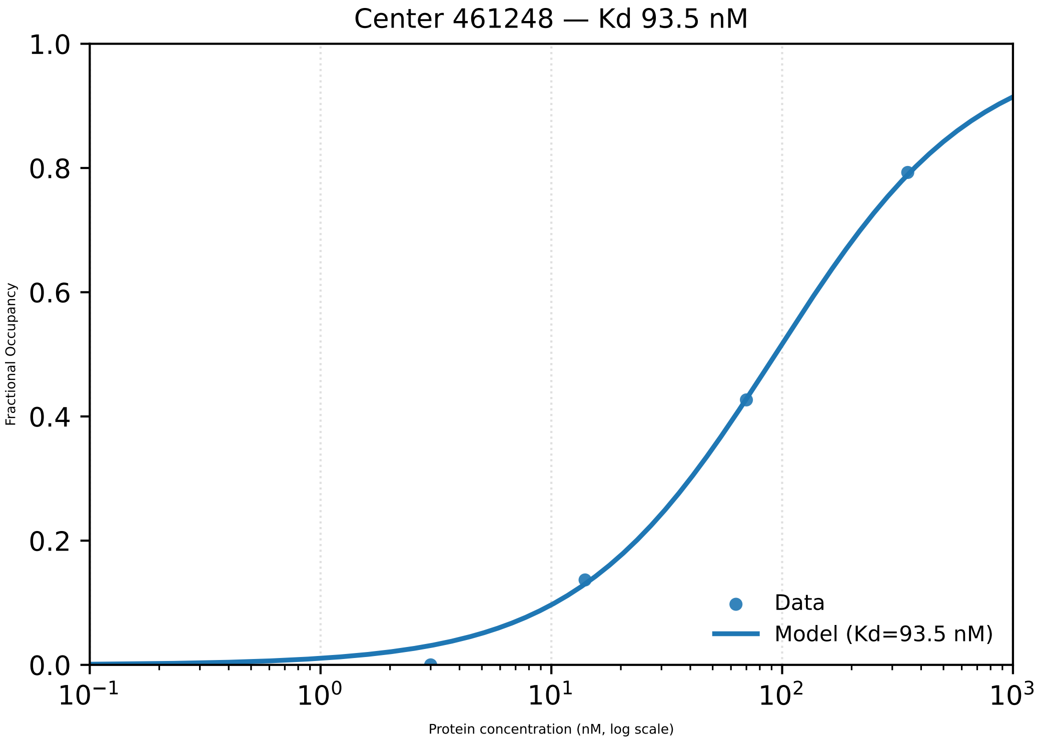

Center 461248 — Kd 93.5 nM
1.0
0.8
0.6
0.4
0.2
0.0
Fractional Occupancy
$10^{-1}$
$10^{0}$
$10^{1}$
$10^{2}$
$10^{3}$
Protein concentration (nM, log scale)
Data
Model (Kd=93.5 nM)

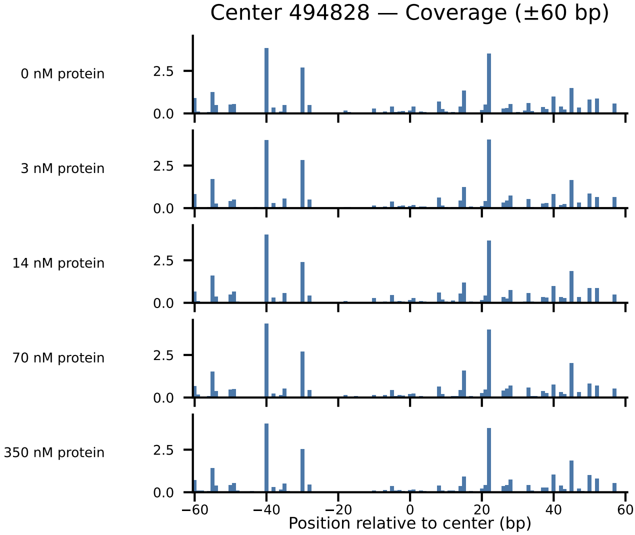

Center 494828 — Coverage (±60 bp)
0 nM protein
3 nM protein
14 nM protein
70 nM protein
350 nM protein
2.5
0.0
−60
−40
−20
0
20
40
60
Position relative to center (bp)

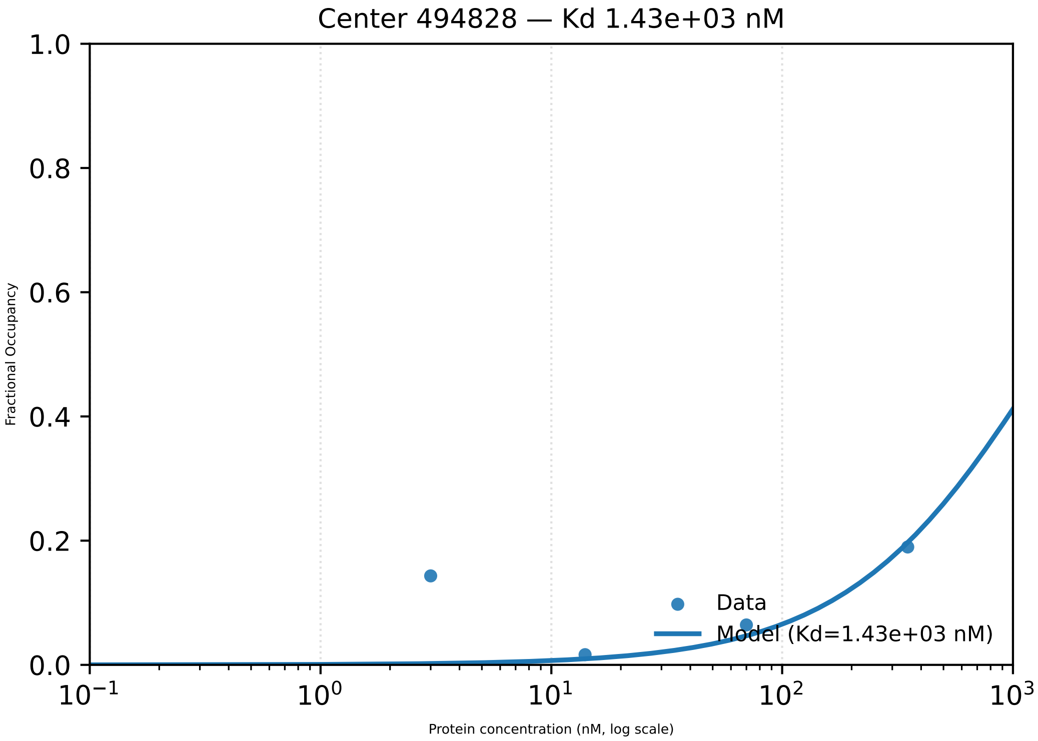

Center 494828 — Kd 1.43e+03 nM
Fractional Occupancy
1.0
0.8
0.6
0.4
0.2
0.0
$10^{-1}$
$10^{0}$
$10^{1}$
$10^{2}$
$10^{3}$
Protein concentration (nM, log scale)
Data
Model (Kd=1.43e+03 nM)

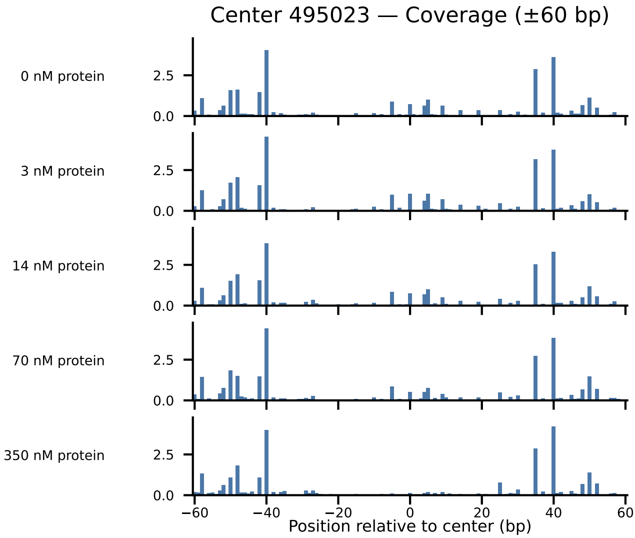

Center 495023 — Coverage (±60 bp)
0 nM protein
3 nM protein
14 nM protein
70 nM protein
350 nM protein
2.5
0.0
−60
−40
−20
0
20
40
60
Position relative to center (bp)

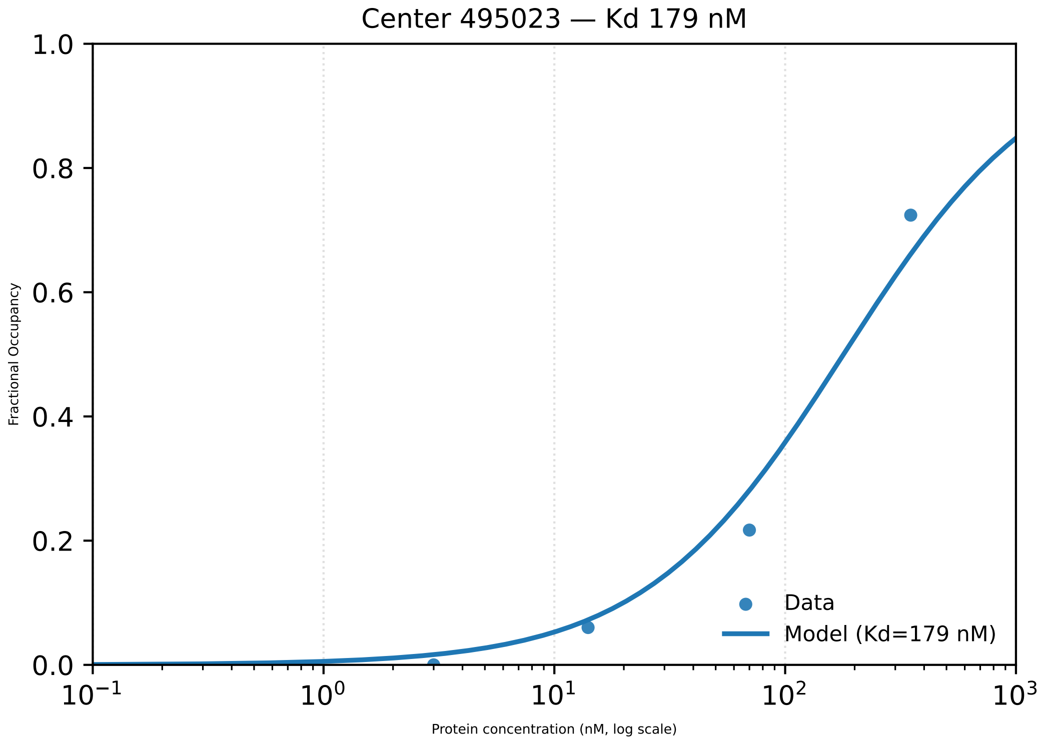
Center 495023 — Kd 179 nM
Fractional Occupancy
1.0
0.8
0.6
0.4
0.2
0.0
$10^{-1}$
$10^{0}$
$10^{1}$
$10^{2}$
$10^{3}$
Protein concentration (nM, log scale)
Data
Model (Kd=179 nM)

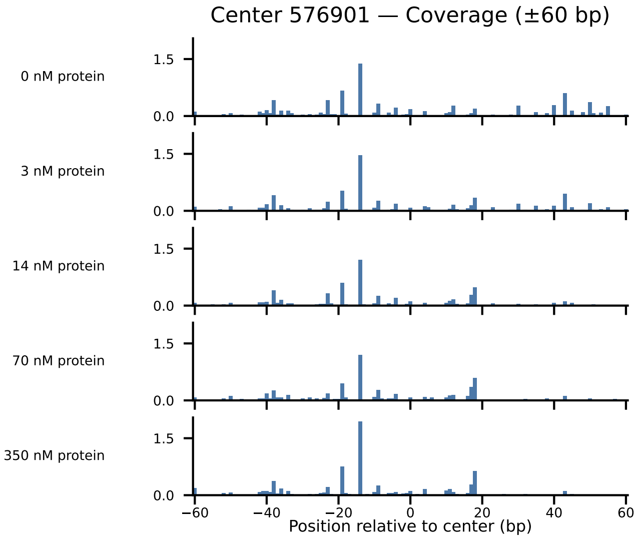
Center 576901 — Coverage (±60 bp)
0 nM protein
3 nM protein
14 nM protein
70 nM protein
350 nM protein
1.5
0.0
−60
−40
−20
0
20
40
60
Position relative to center (bp)

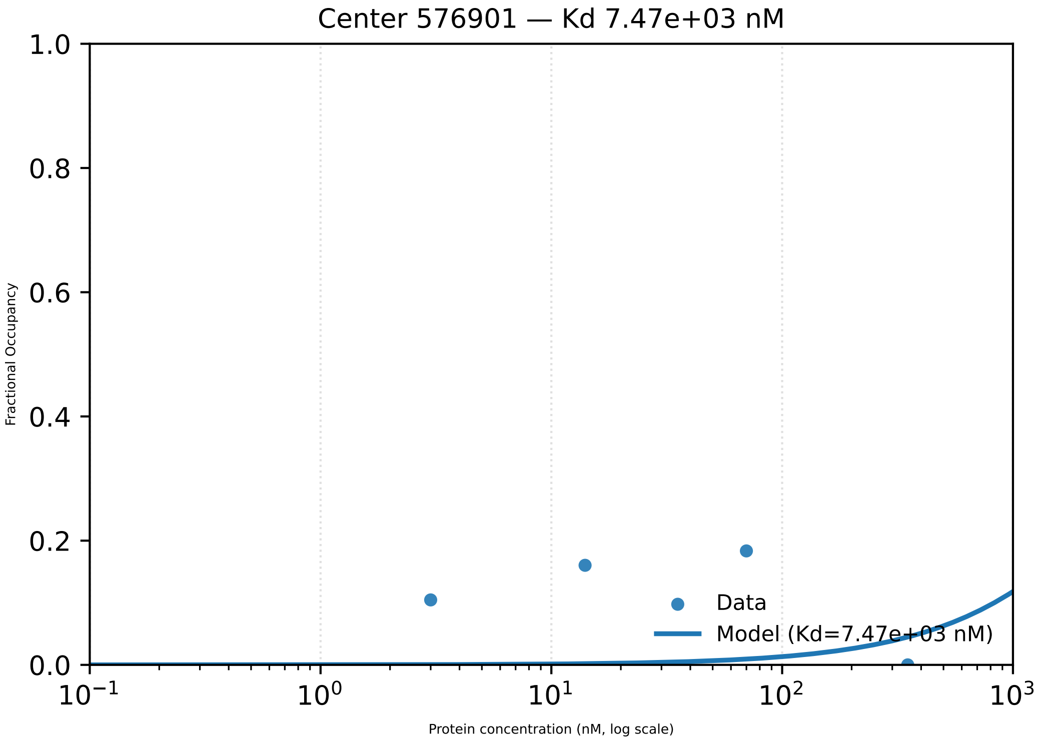
Center 576901 — Kd 7.47e+03 nM
Fractional Occupancy
Protein concentration (nM, log scale)
Data
Model (Kd=7.47e+03 nM)

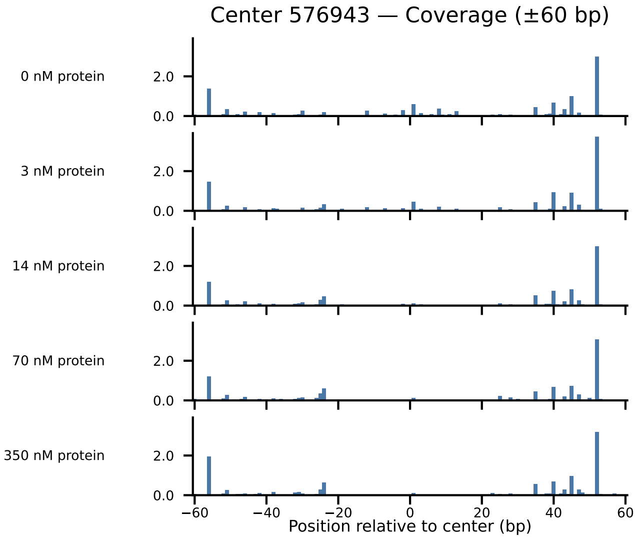

Center 576943 — Coverage (±60 bp)
0 nM protein
3 nM protein
14 nM protein
70 nM protein
350 nM protein
2.0
0.0
−60
−40
−20
0
20
40
60
Position relative to center (bp)

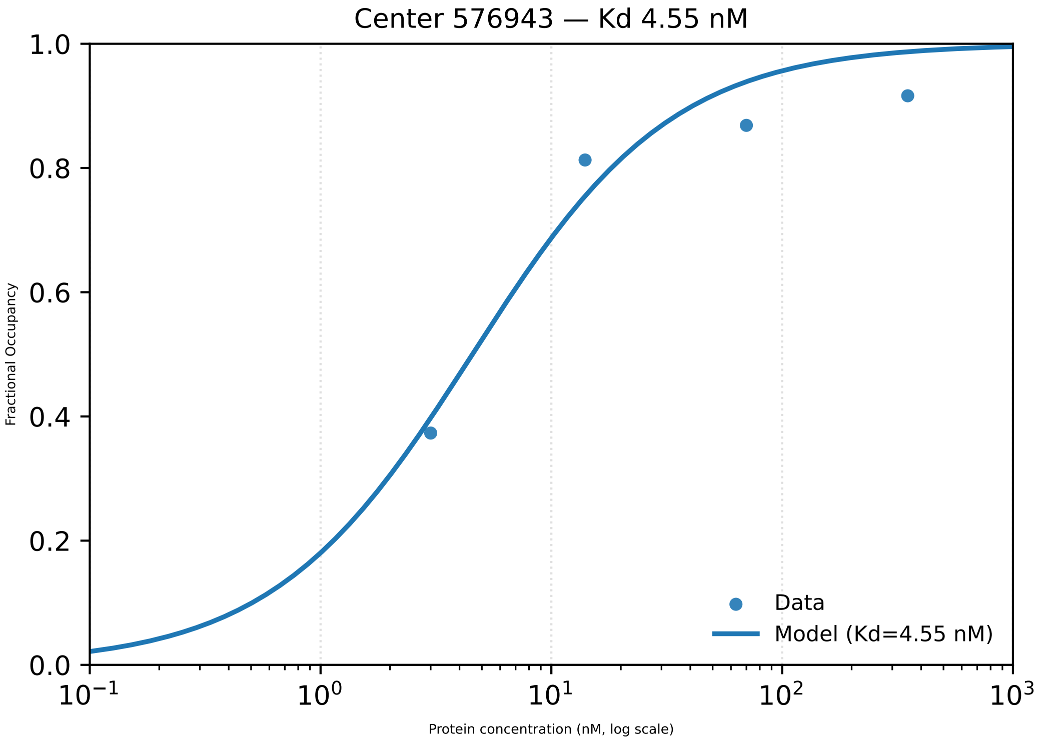

Center 576943 — Kd 4.55 nM
Fractional Occupancy
1.0
0.8
0.6
0.4
0.2
0.0
$10^{-1}$
$10^{0}$
$10^{1}$
$10^{2}$
$10^{3}$
Protein concentration (nM, log scale)
Data
Model (Kd=4.55 nM)

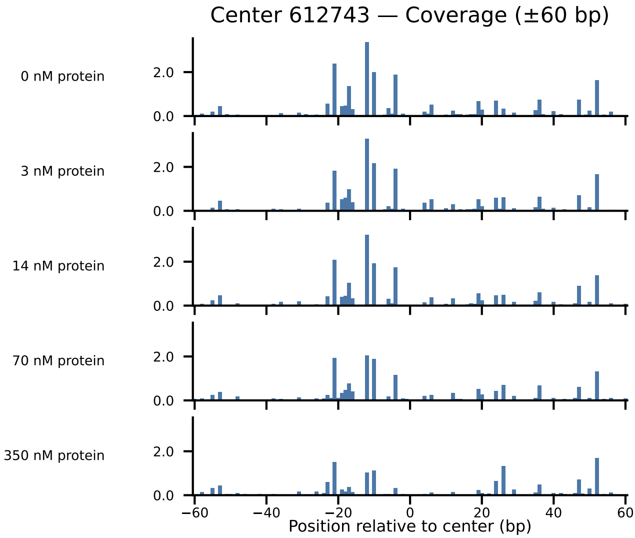
Center 612743 — Coverage (±60 bp)
0 nM protein
3 nM protein
14 nM protein
70 nM protein
350 nM protein
2.0
0.0
−60
−40
−20
0
20
40
60
Position relative to center (bp)

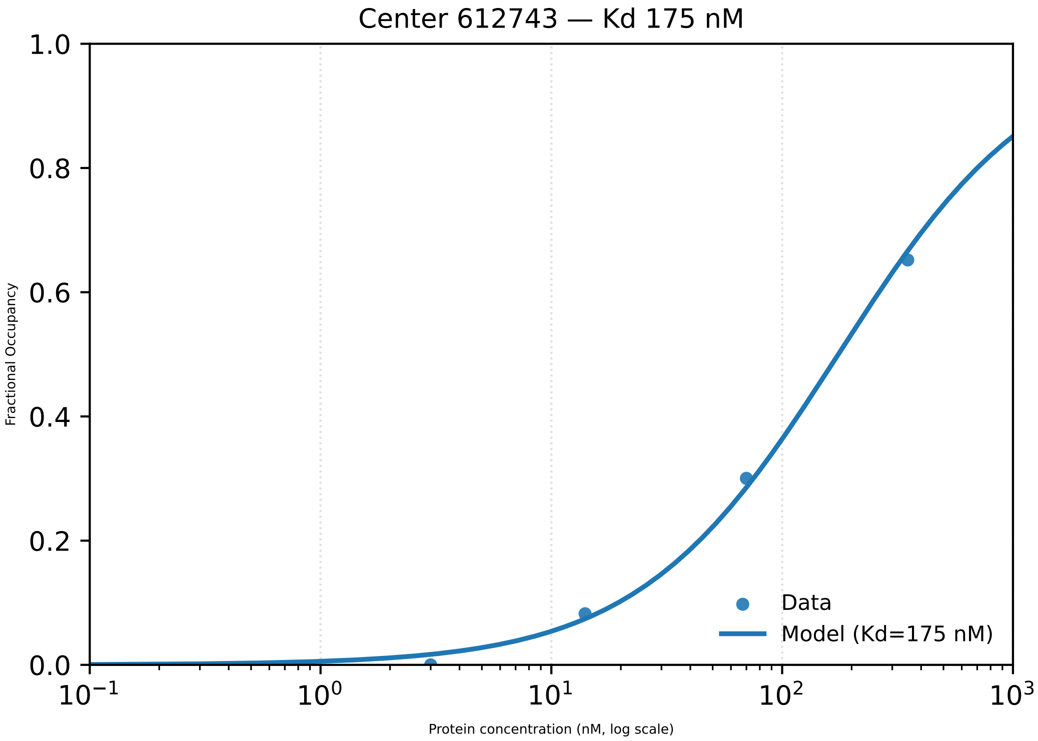

Center 612743 — Kd 175 nM
Fractional Occupancy
1.0
0.8
0.6
0.4
0.2
0.0
$10^{-1}$
$10^{0}$
$10^{1}$
$10^{2}$
$10^{3}$
Protein concentration (nM, log scale)
Data
Model (Kd=175 nM)

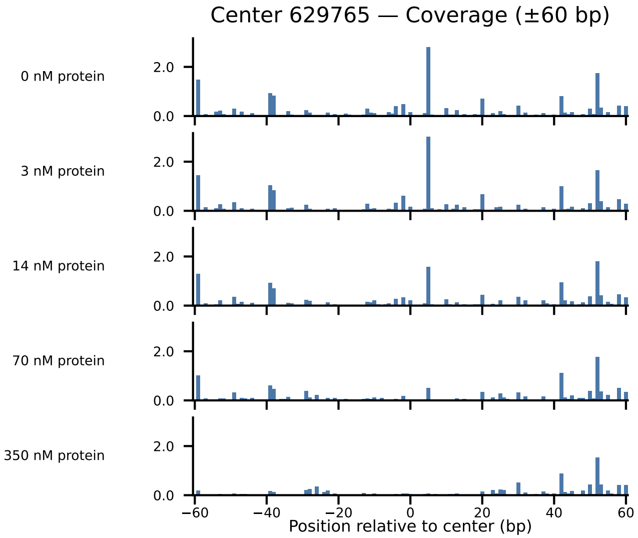

Center 629765 — Coverage (±60 bp)
0 nM protein
3 nM protein
14 nM protein
70 nM protein
350 nM protein
2.0
0.0
−60
−40
−20
0
20
40
60
Position relative to center (bp)

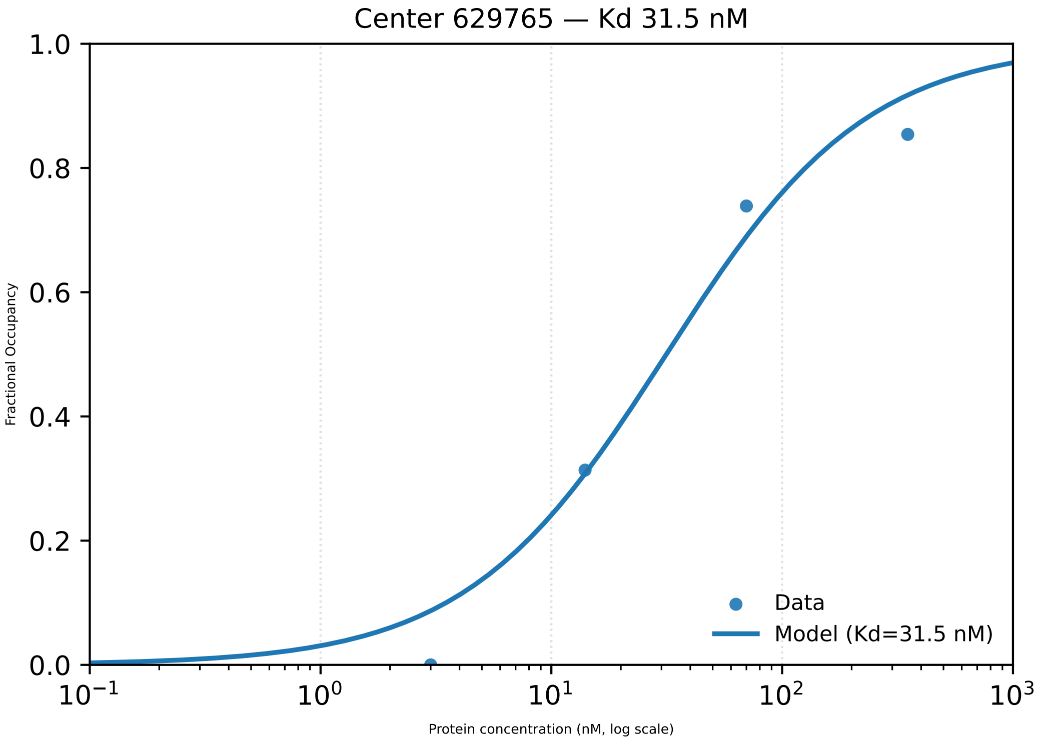

Center 629765 — Kd 31.5 nM
1.0
0.8
0.6
0.4
0.2
0.0
Fractional Occupancy
$10^{-1}$
$10^{0}$
$10^{1}$
$10^{2}$
$10^{3}$
Protein concentration (nM, log scale)
Data
Model (Kd=31.5 nM)

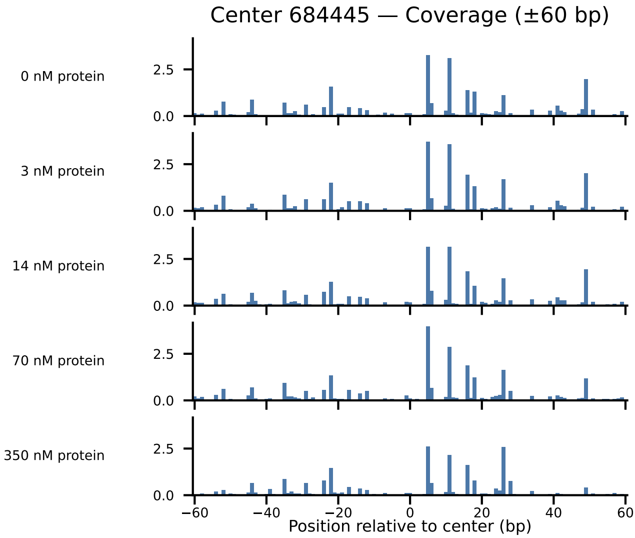

Center 684445 — Coverage (±60 bp)
0 nM protein
3 nM protein
14 nM protein
70 nM protein
350 nM protein
2.5
0.0
−60
−40
−20
0
20
40
60
Position relative to center (bp)

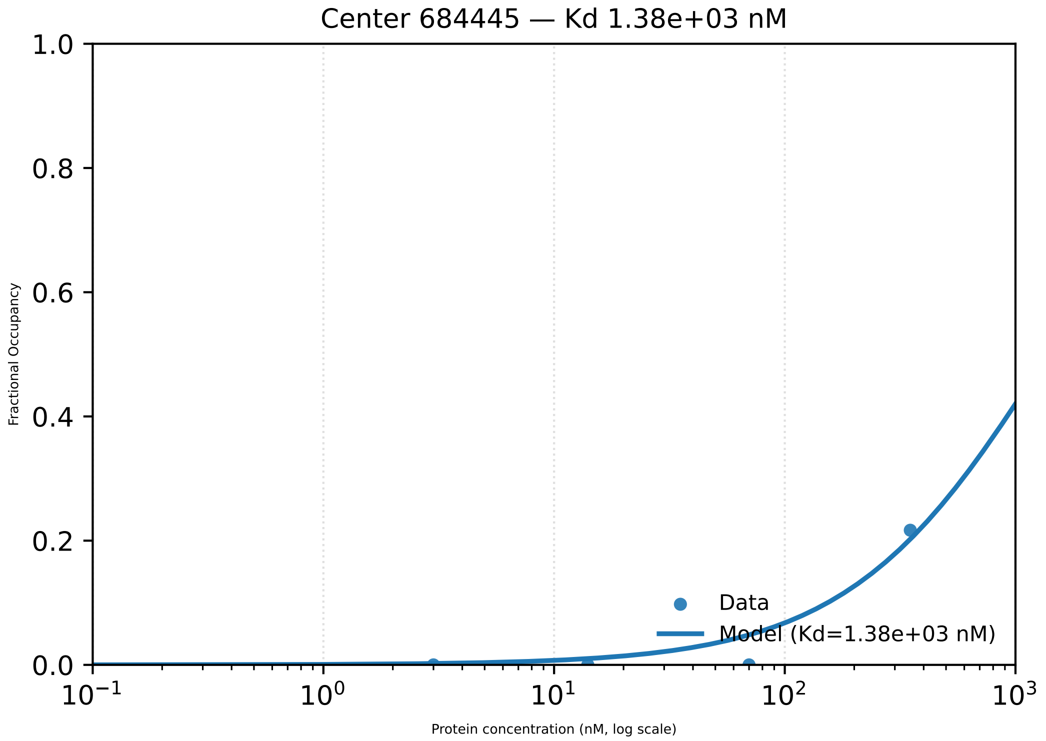

Center 684445 — Kd 1.38e+03 nM
1.0
0.8
0.6
0.4
0.2
0.0
Fractional Occupancy
$10^{-1}$
$10^{0}$
$10^{1}$
$10^{2}$
$10^{3}$
Protein concentration (nM, log scale)
Data
Model (Kd=1.38e+03 nM)

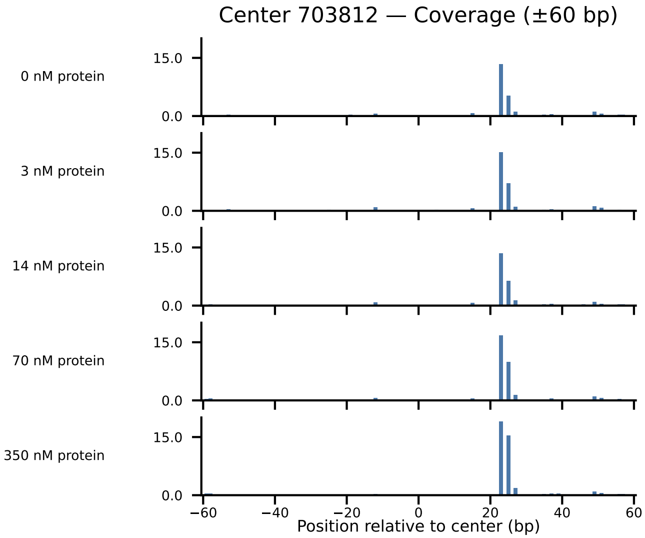

Center 703812 — Coverage (±60 bp)
0 nM protein
3 nM protein
14 nM protein
70 nM protein
350 nM protein
15.0
0.0
−60
−40
−20
0
20
40
60
Position relative to center (bp)

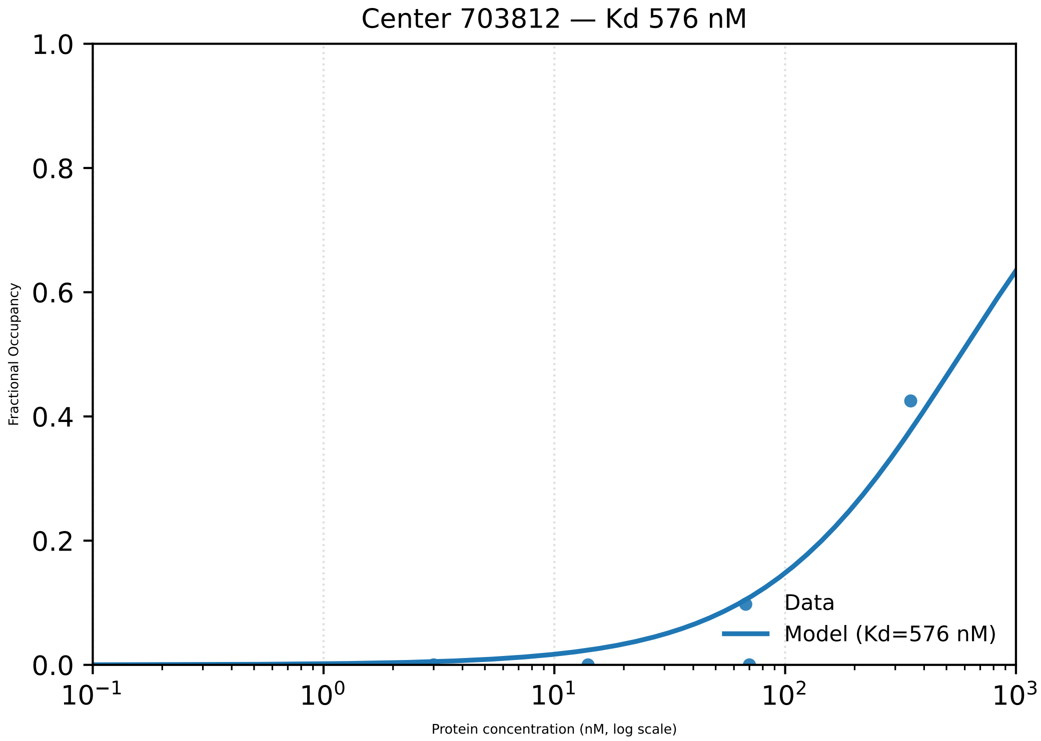

Center 703812 — Kd 576 nM
1.0
0.8
0.6
0.4
0.2
0.0
Fractional Occupancy
$10^{-1}$
$10^{0}$
$10^{1}$
$10^{2}$
$10^{3}$
Protein concentration (nM, log scale)
Data
Model (Kd=576 nM)

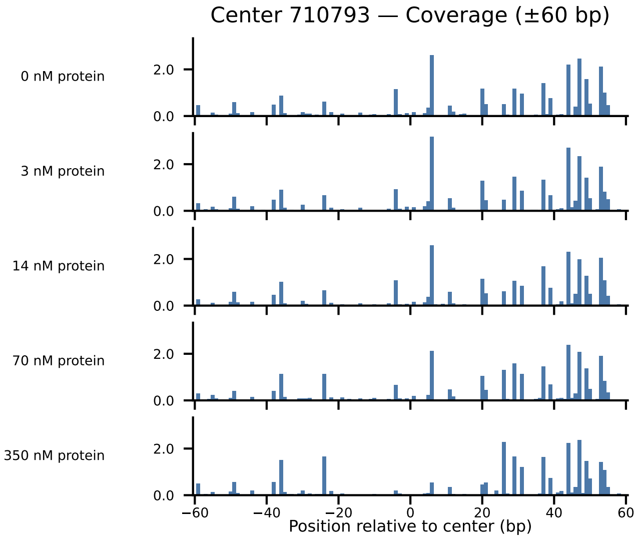
Center 710793 — Coverage (±60 bp)
0 nM protein
3 nM protein
14 nM protein
70 nM protein
350 nM protein
2.0
0.0
−60
−40
−20
0
20
40
60
Position relative to center (bp)

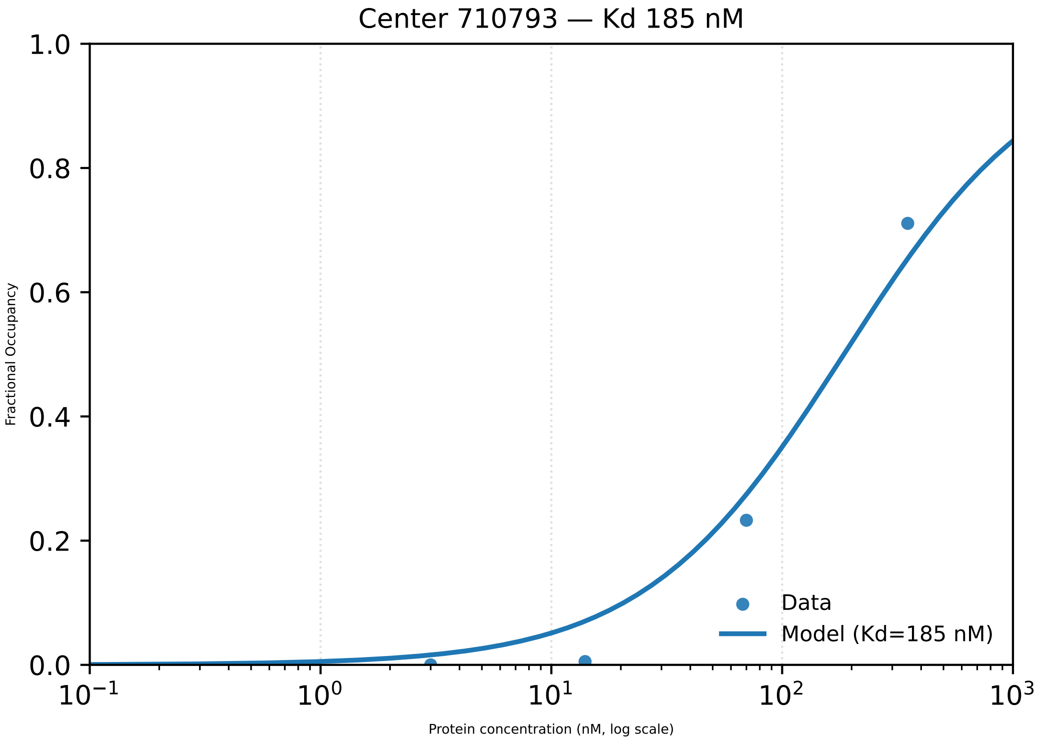

Center 710793 — Kd 185 nM
1.0
0.8
0.6
0.4
0.2
0.0
Fractional Occupancy
$10^{-1}$
$10^{0}$
$10^{1}$
$10^{2}$
$10^{3}$
Protein concentration (nM, log scale)
Data
Model (Kd=185 nM)

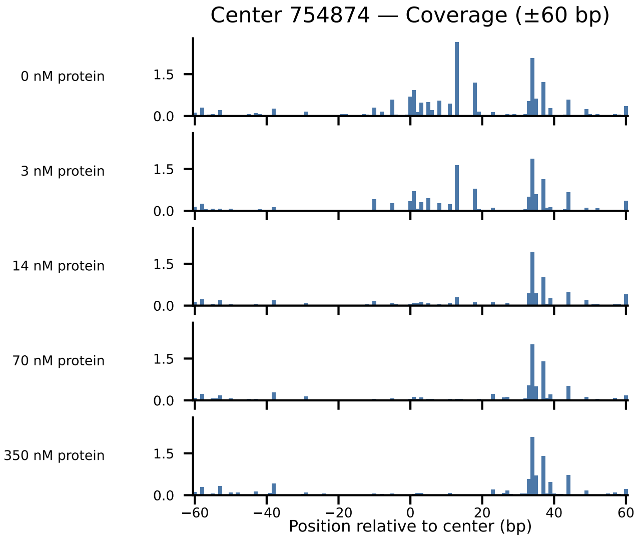
Center 754874 — Coverage (±60 bp)
0 nM protein
3 nM protein
14 nM protein
70 nM protein
350 nM protein
1.5
0.0
−60
−40
−20
0
20
40
60
Position relative to center (bp)

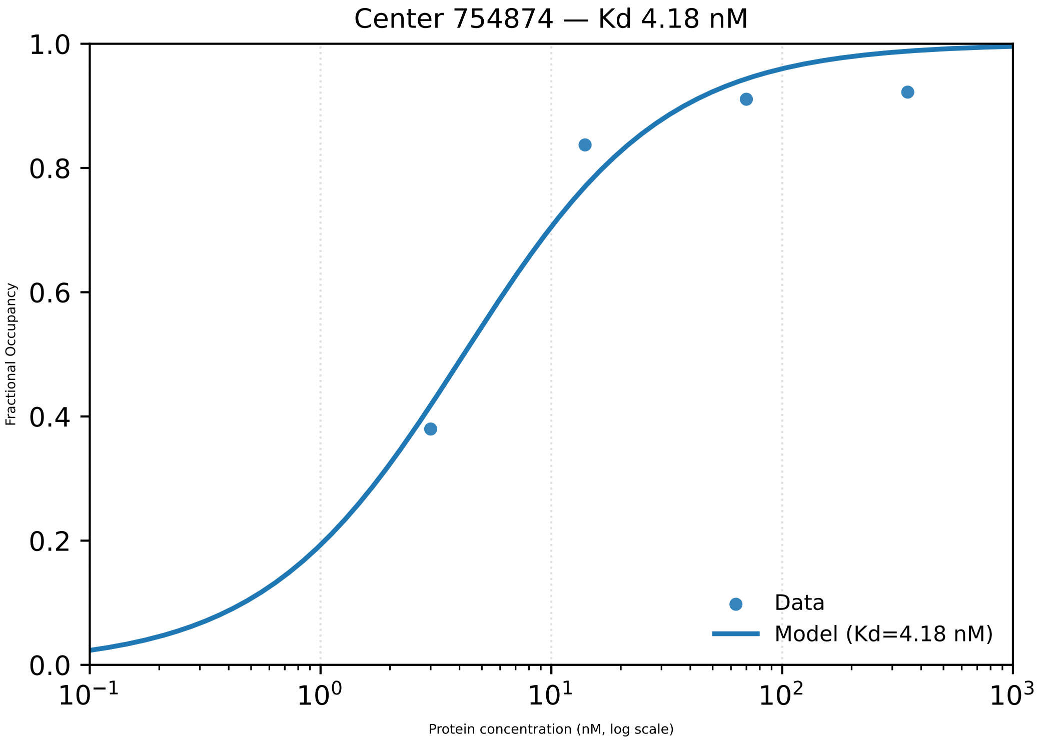

Center 754874 — Kd 4.18 nM
Fractional Occupancy
0.0
0.2
0.4
0.6
0.8
1.0
$10^{-1}$
$10^{0}$
$10^{1}$
$10^{2}$
$10^{3}$
Protein concentration (nM, log scale)
Data
Model (Kd=4.18 nM)

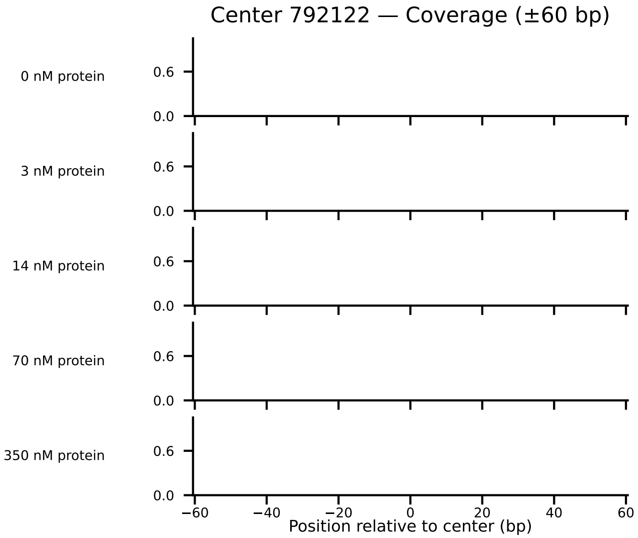

Center 792122 — Coverage (±60 bp)
0 nM protein
3 nM protein
14 nM protein
70 nM protein
350 nM protein
0.6
0.0
−60
−40
−20
0
20
40
60
Position relative to center (bp)

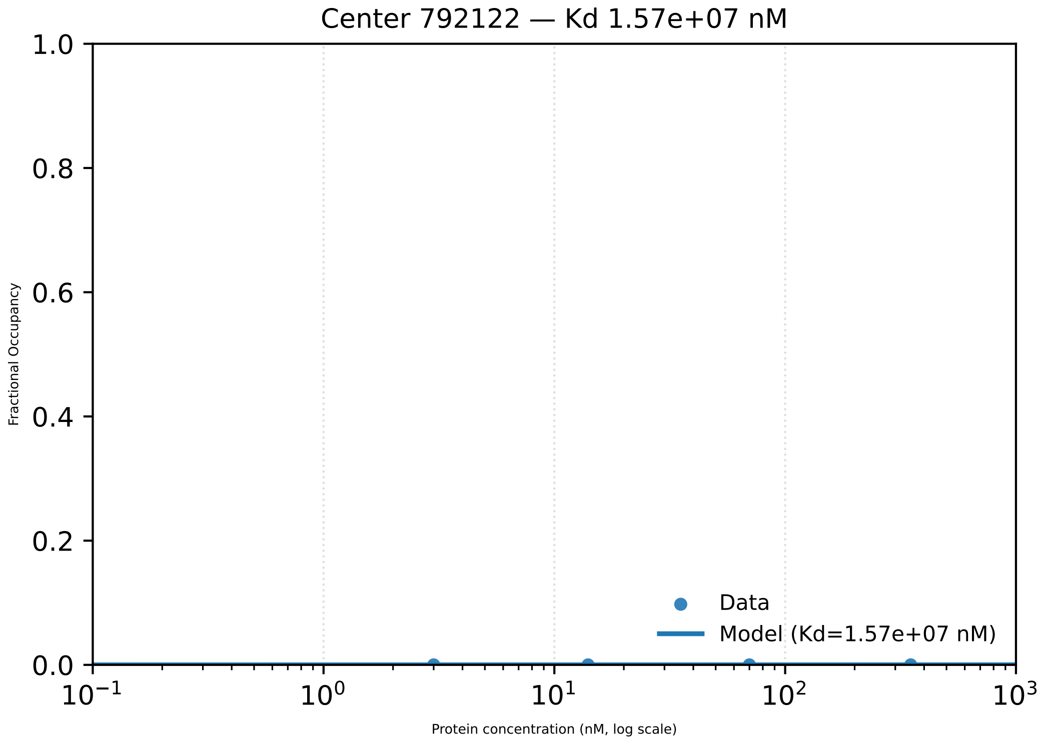
Center 792122 — Kd 1.57e+07 nM
Fractional Occupancy
1.0
0.8
0.6
0.4
0.2
0.0
$10^{-1}$
$10^{0}$
$10^{1}$
$10^{2}$
$10^{3}$
Protein concentration (nM, log scale)
Data
Model (Kd=1.57e+07 nM)

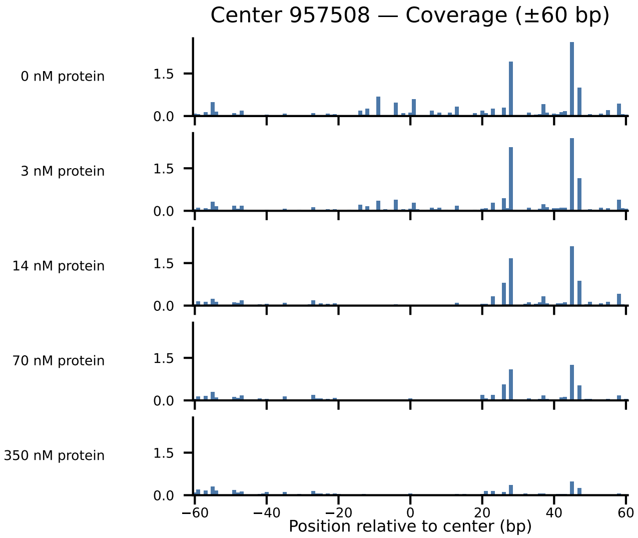

Center 957508 — Coverage (±60 bp)
0 nM protein
3 nM protein
14 nM protein
70 nM protein
350 nM protein
1.5
0.0
−60
−40
−20
0
20
40
60
Position relative to center (bp)

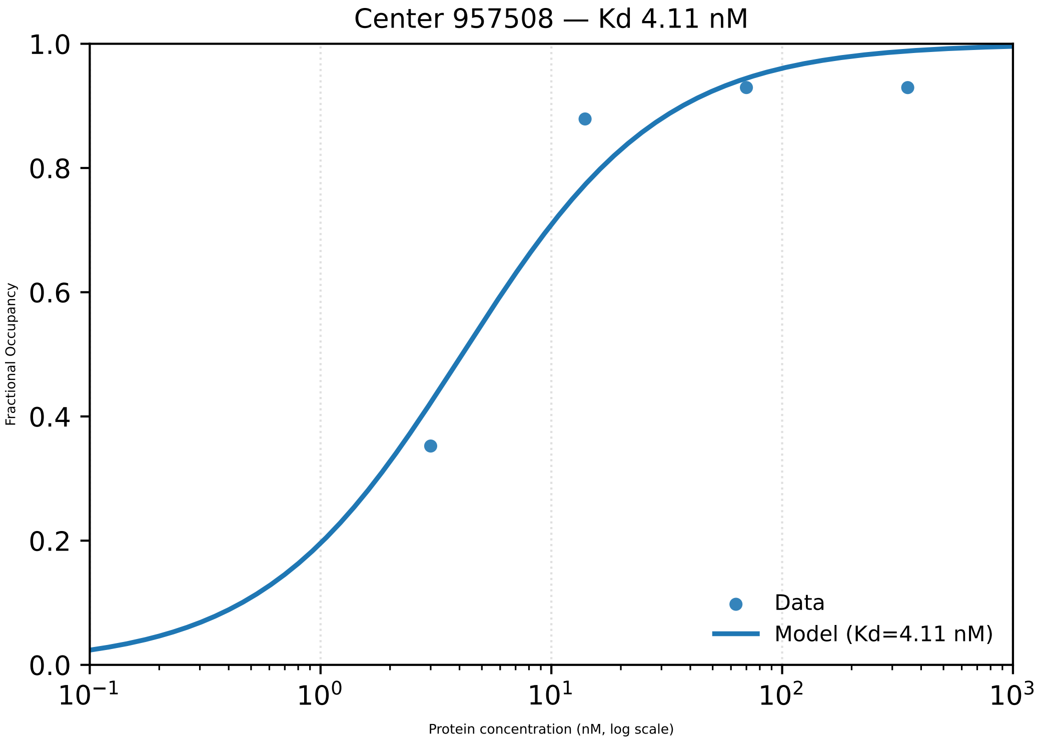

Center 957508 — Kd 4.11 nM
Fractional Occupancy
1.0
0.8
0.6
0.4
0.2
0.0
$10^{-1}$
$10^{0}$
$10^{1}$
$10^{2}$
$10^{3}$
Protein concentration (nM, log scale)
Data
Model (Kd=4.11 nM)

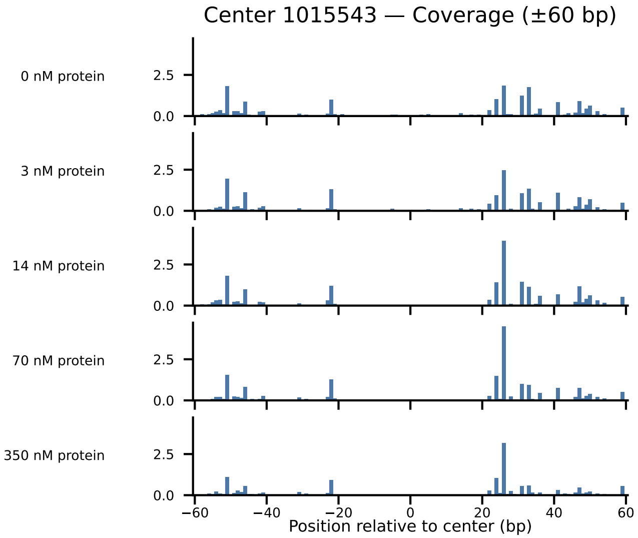

Center 1015543 — Coverage (±60 bp)
0 nM protein
3 nM protein
14 nM protein
70 nM protein
350 nM protein
2.5
0.0
−60
−40
−20
0
20
40
60
Position relative to center (bp)

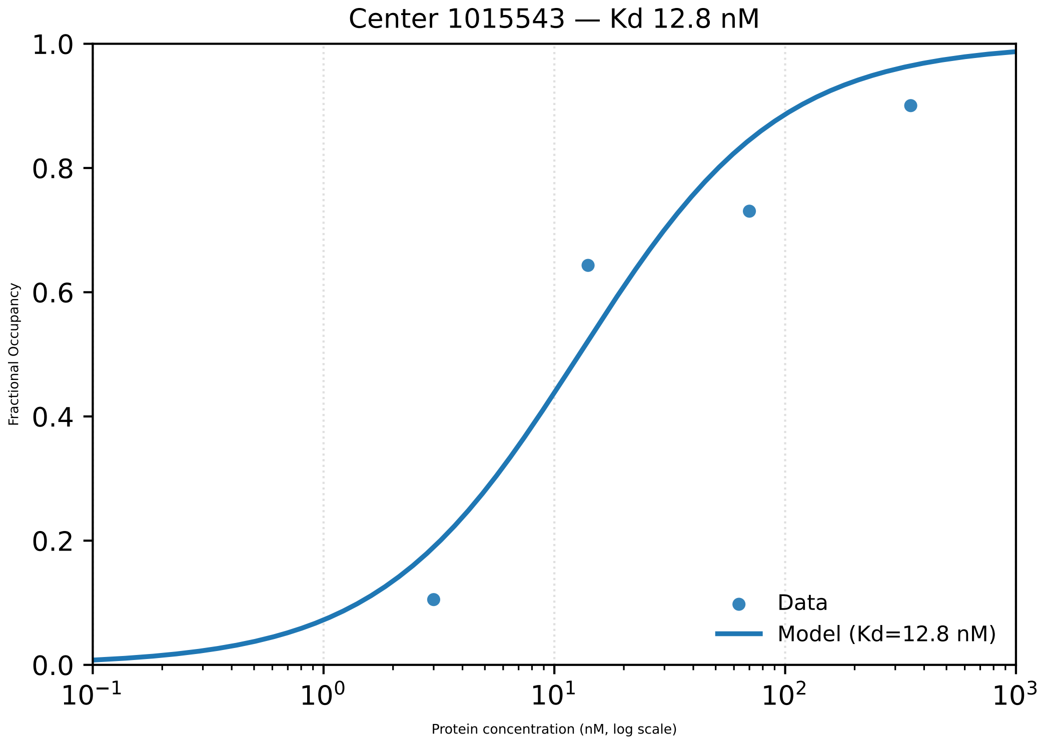

Center 1015543 — Kd 12.8 nM
Fractional Occupancy
Protein concentration (nM, log scale)
1.0
0.8
0.6
0.4
0.2
0.0
$10^{-1}$
$10^{0}$
$10^{1}$
$10^{2}$
$10^{3}$
Data
Model (Kd=12.8 nM)

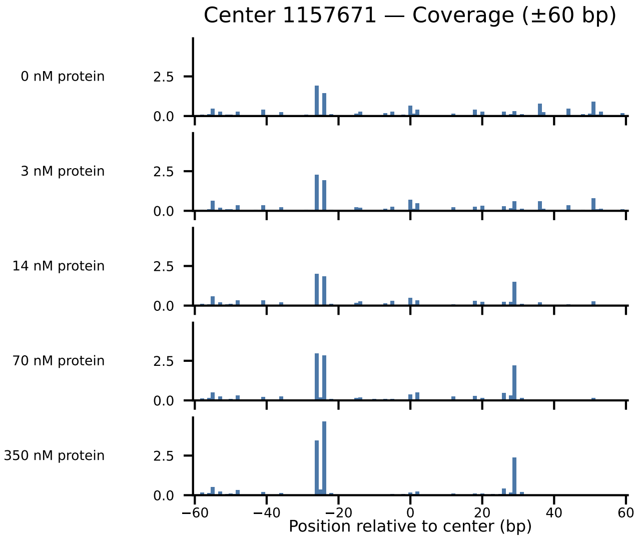

Center 1157671 — Coverage (±60 bp)
0 nM protein
3 nM protein
14 nM protein
70 nM protein
350 nM protein
2.5
0.0
−60
−40
−20
0
20
40
60
Position relative to center (bp)

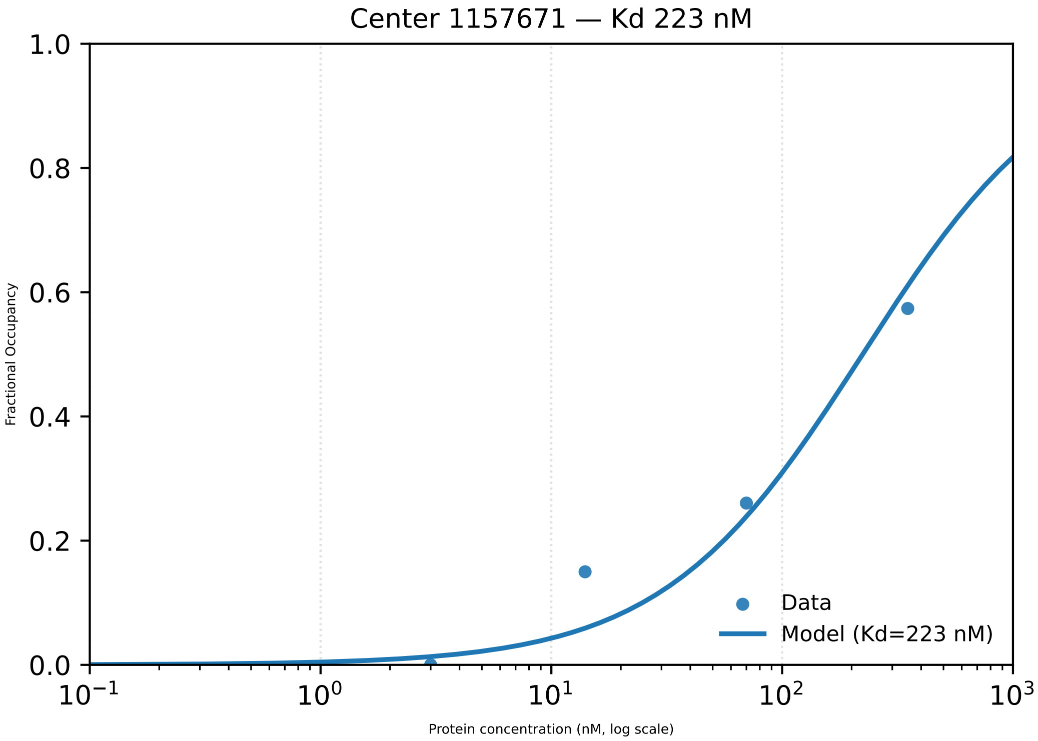

Center 1157671 — Kd 223 nM
1.0
0.8
0.6
0.4
0.2
0.0
Fractional Occupancy
$10^{-1}$
$10^{0}$
$10^{1}$
$10^{2}$
$10^{3}$
Protein concentration (nM, log scale)
Data
Model (Kd=223 nM)

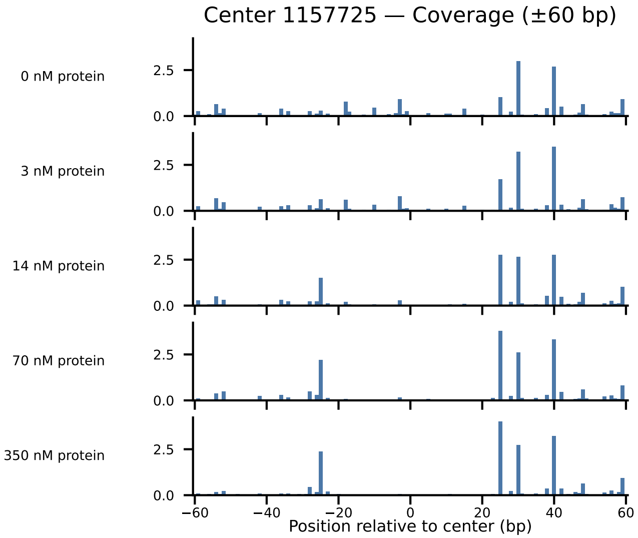

Center 1157725 — Coverage (±60 bp)
0 nM protein
3 nM protein
14 nM protein
70 nM protein
350 nM protein
0.0
2.5
−60
−40
−20
0
20
40
60
Position relative to center (bp)

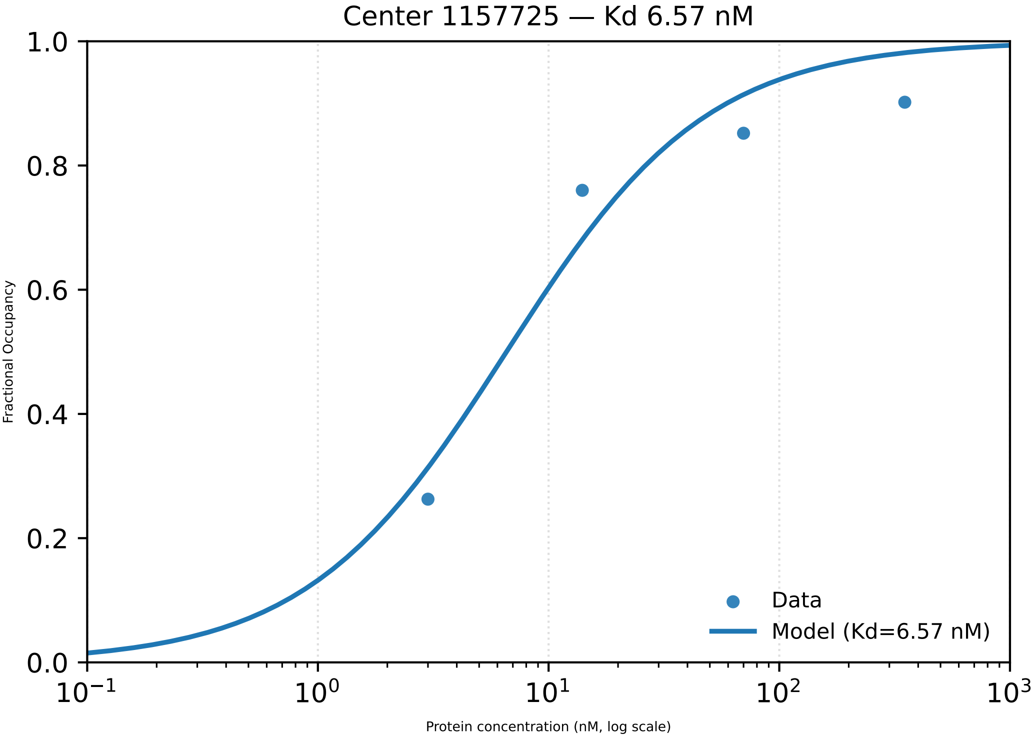

Center 1157725 — Kd 6.57 nM
Fractional Occupancy
1.0
0.8
0.6
0.4
0.2
0.0
$10^{-1}$
$10^{0}$
$10^{1}$
$10^{2}$
$10^{3}$
Protein concentration (nM, log scale)
Data
Model (Kd=6.57 nM)

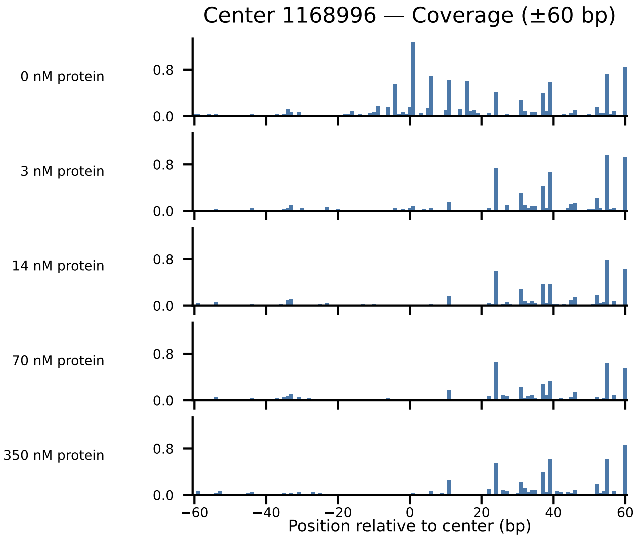
Center 1168996 — Coverage (±60 bp)
0 nM protein
3 nM protein
14 nM protein
70 nM protein
350 nM protein
0.8
0.0
−60
−40
−20
0
20
40
60
Position relative to center (bp)

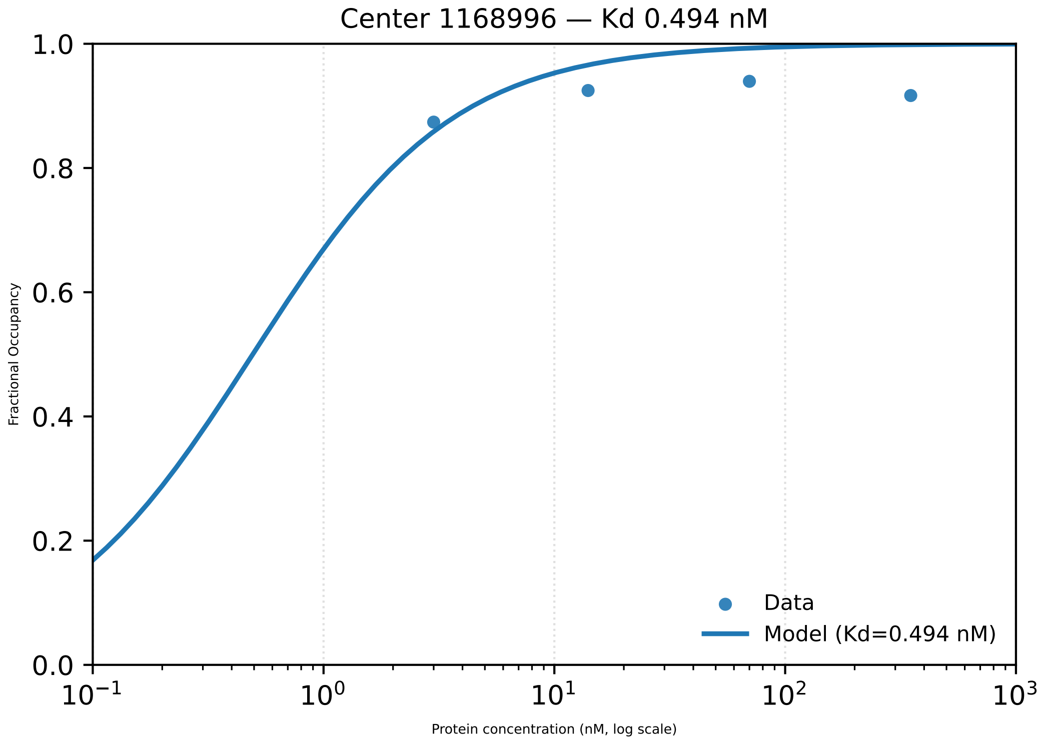

Center 1168996 — Kd 0.494 nM
1.0
0.8
0.6
0.4
0.2
0.0
Fractional Occupancy
$10^{-1}$
$10^{0}$
$10^{1}$
$10^{2}$
$10^{3}$
Protein concentration (nM, log scale)
Data
Model (Kd=0.494 nM)

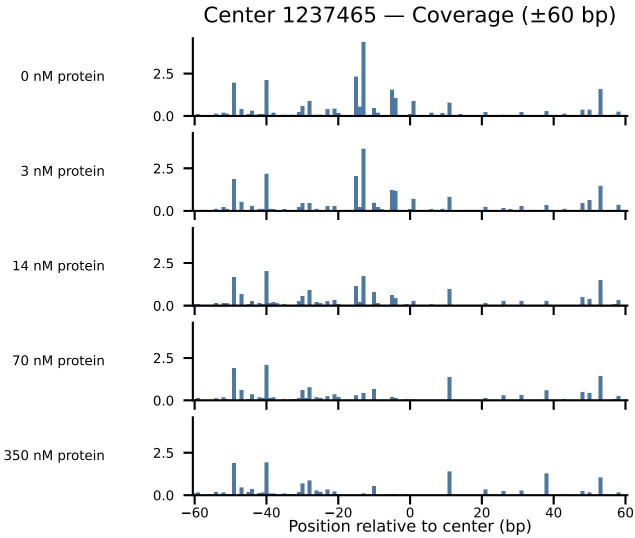

Center 1237465 — Coverage (±60 bp)
0 nM protein
3 nM protein
14 nM protein
70 nM protein
350 nM protein
2.5
0.0
−60
−40
−20
0
20
40
60
Position relative to center (bp)

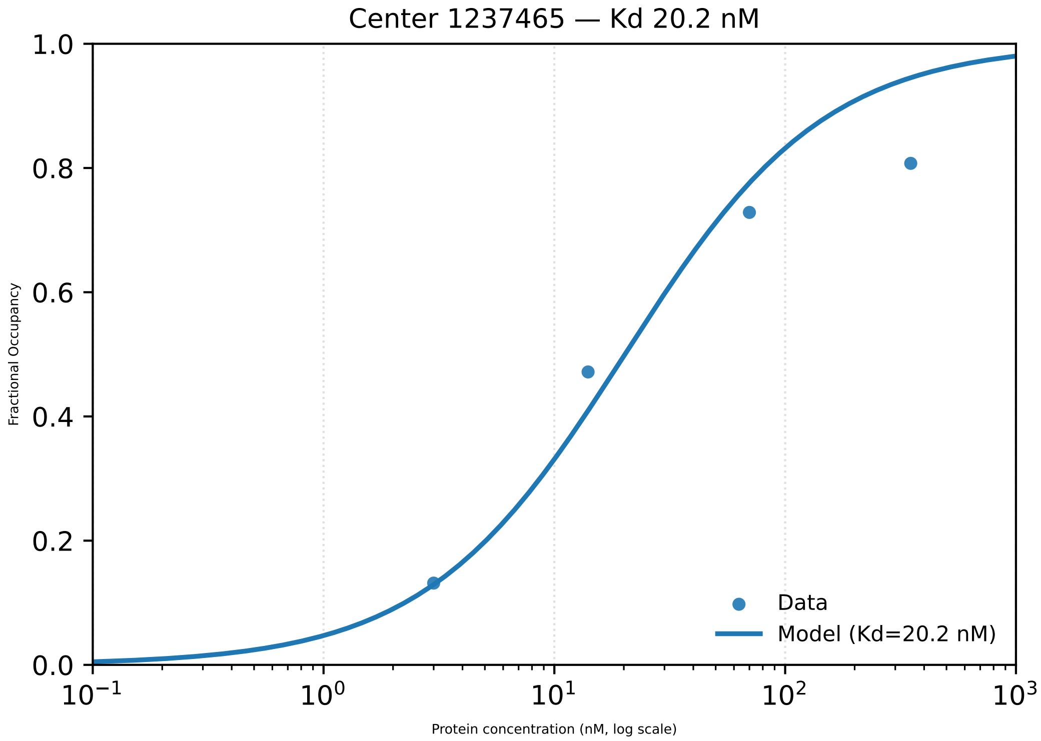

Center 1237465 — Kd 20.2 nM
1.0
0.8
0.6
0.4
0.2
0.0
Fractional Occupancy
$10^{-1}$
$10^{0}$
$10^{1}$
$10^{2}$
$10^{3}$
Protein concentration (nM, log scale)
Data
Model (Kd=20.2 nM)

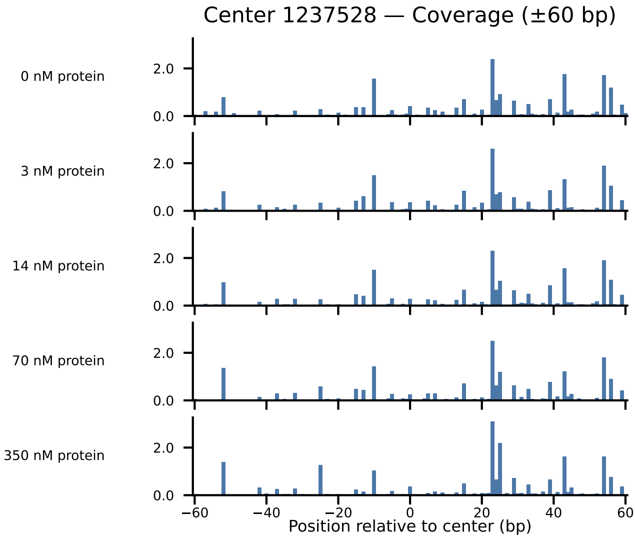

Center 1237528 — Coverage (±60 bp)
0 nM protein
3 nM protein
14 nM protein
70 nM protein
350 nM protein
0.0
2.0
−60
−40
−20
0
20
40
60
Position relative to center (bp)

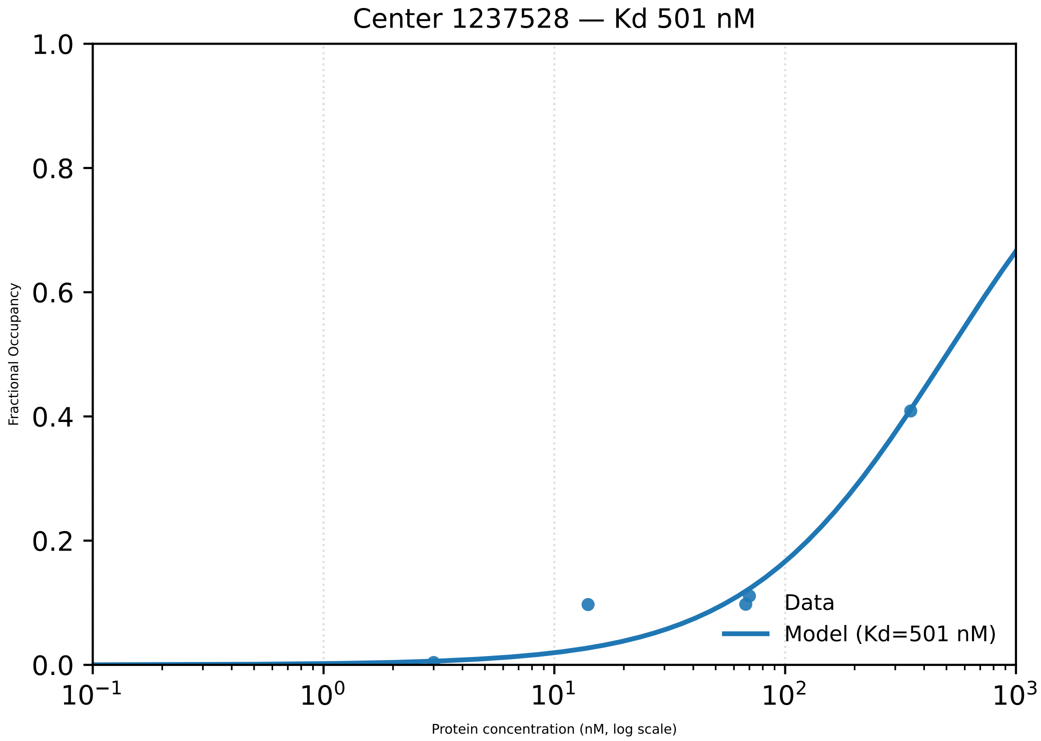

Center 1237528 — Kd 501 nM
1.0
0.8
0.6
0.4
0.2
0.0
Fractional Occupancy
Data
Model (Kd=501 nM)
$10^{-1}$
$10^{0}$
$10^{1}$
$10^{2}$
$10^{3}$
Protein concentration (nM, log scale)

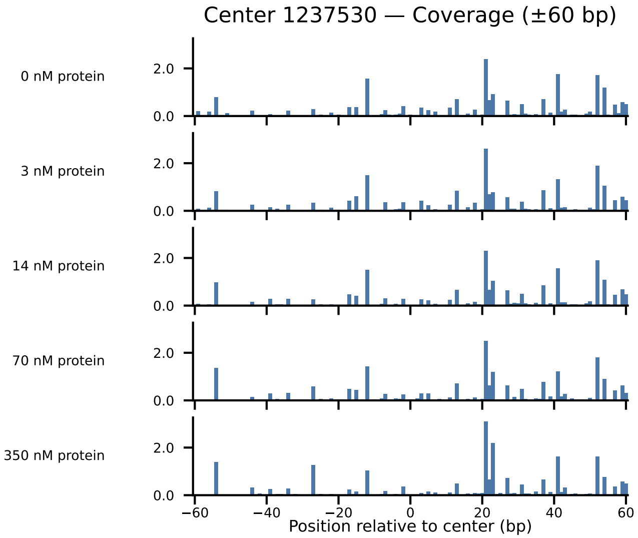
Center 1237530 — Coverage (±60 bp)
0 nM protein
3 nM protein
14 nM protein
70 nM protein
350 nM protein
2.0
0.0
−60
−40
−20
0
20
40
60
Position relative to center (bp)

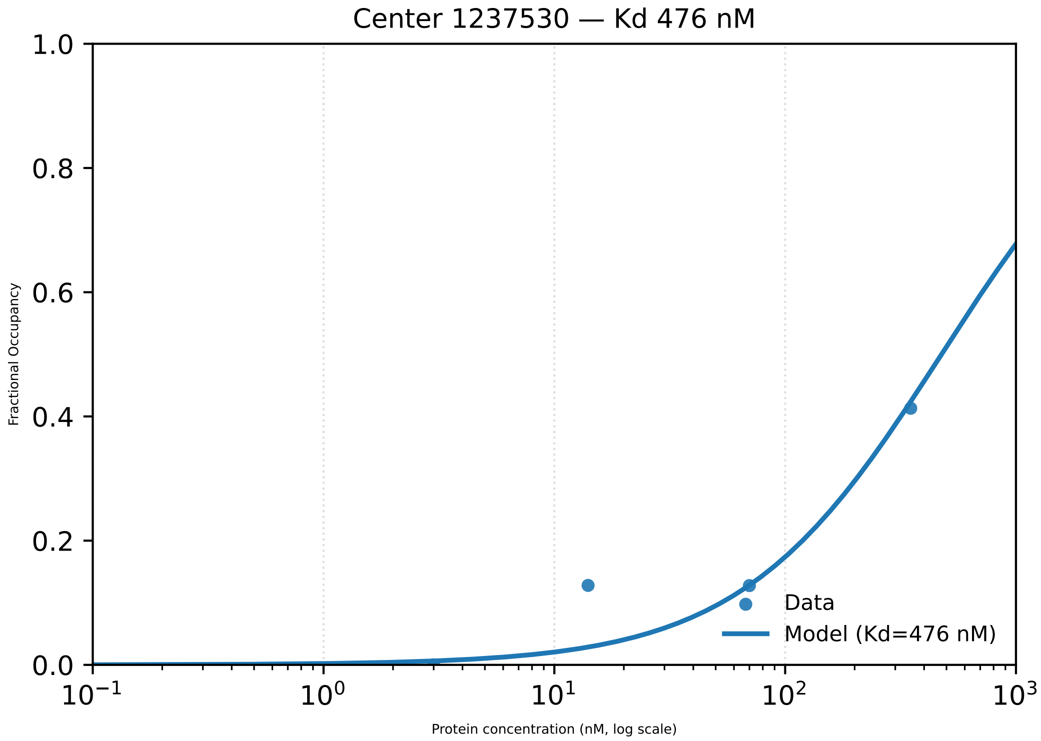

Center 1237530 — Kd 476 nM
1.0
0.8
0.6
0.4
0.2
0.0
Fractional Occupancy
$10^{-1}$
$10^{0}$
$10^{1}$
$10^{2}$
$10^{3}$
Protein concentration (nM, log scale)
Data
Model (Kd=476 nM)

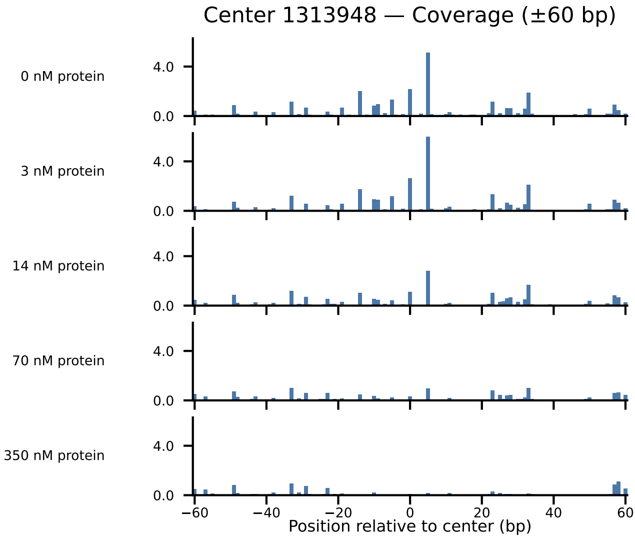

Center 1313948 — Coverage (±60 bp)
0 nM protein
3 nM protein
14 nM protein
70 nM protein
350 nM protein
4.0
0.0
−60
−40
−20
0
20
40
60
Position relative to center (bp)

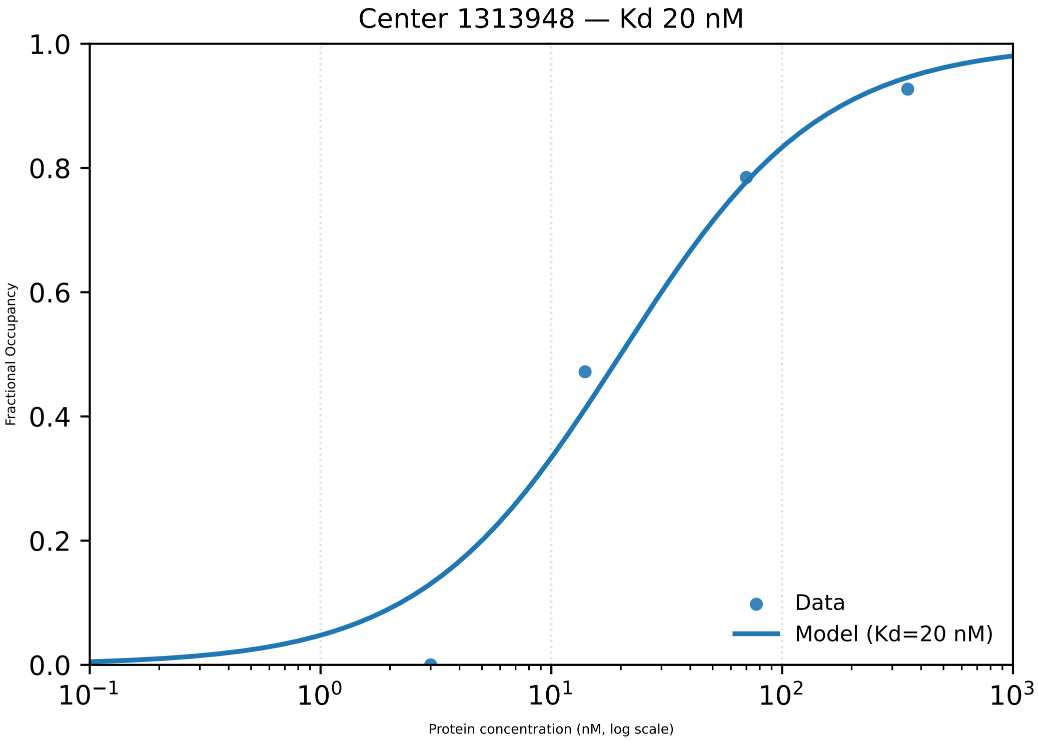

Center 1313948 — Kd 20 nM
1.0
0.8
0.6
0.4
0.2
0.0
Fractional Occupancy
$10^{-1}$
$10^{0}$
$10^{1}$
$10^{2}$
$10^{3}$
Protein concentration (nM, log scale)
Data
Model (Kd=20 nM)

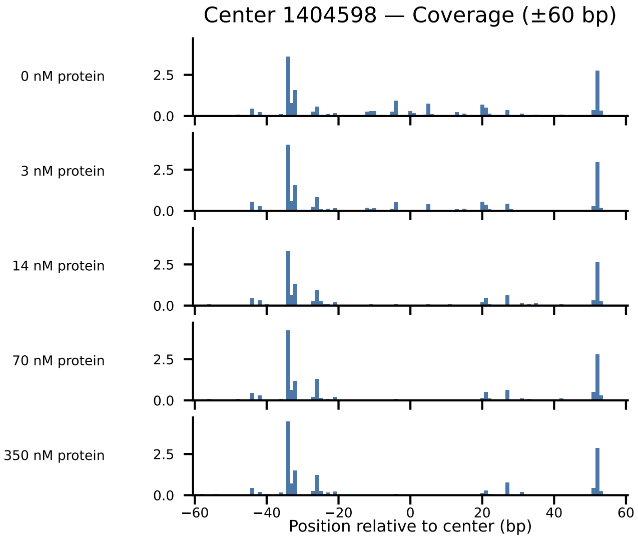

Center 1404598 — Coverage (±60 bp)
0 nM protein
3 nM protein
14 nM protein
70 nM protein
350 nM protein
2.5
0.0
−60
−40
−20
0
20
40
60
Position relative to center (bp)

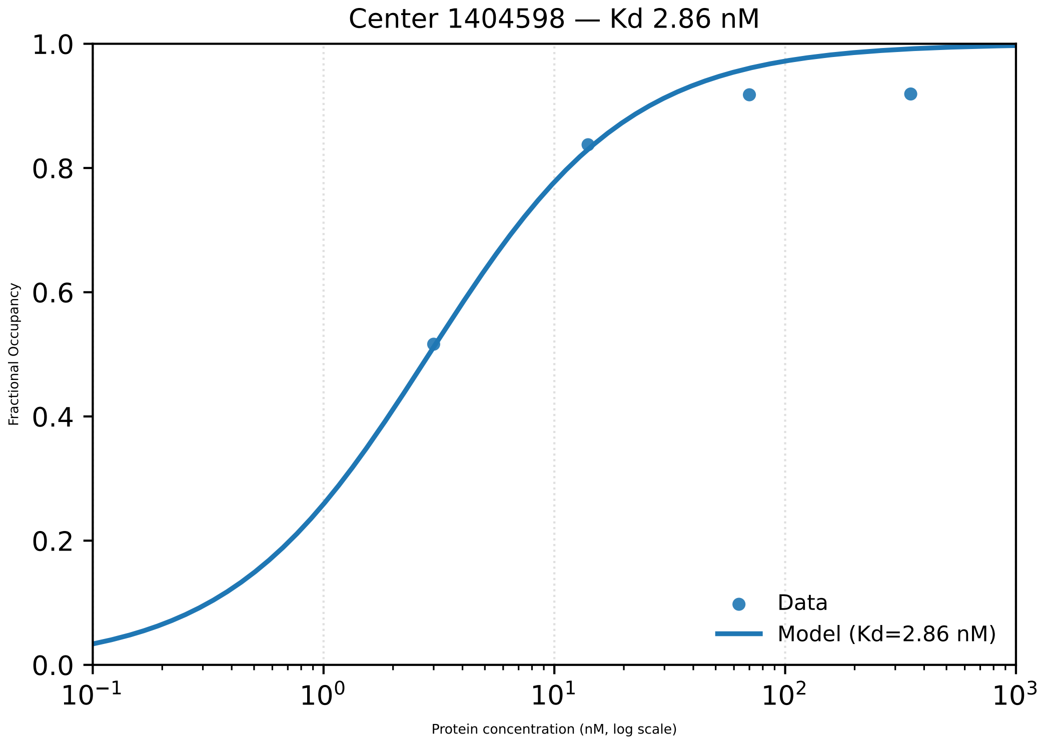

Center 1404598 — Kd 2.86 nM
Fractional Occupancy
1.0
0.8
0.6
0.4
0.2
0.0
$10^{-1}$
$10^{0}$
$10^{1}$
$10^{2}$
$10^{3}$
Protein concentration (nM, log scale)
Data
Model (Kd=2.86 nM)

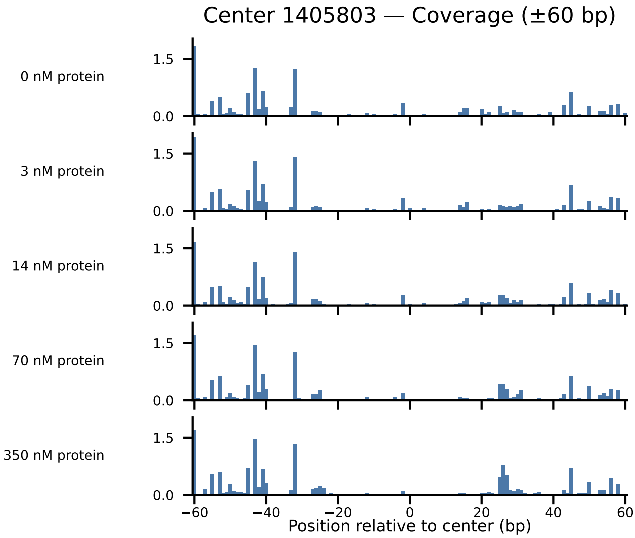
Center 1405803 — Coverage (±60 bp)
0 nM protein
3 nM protein
14 nM protein
70 nM protein
350 nM protein
1.5
0.0
−60
−40
−20
0
20
40
60
Position relative to center (bp)

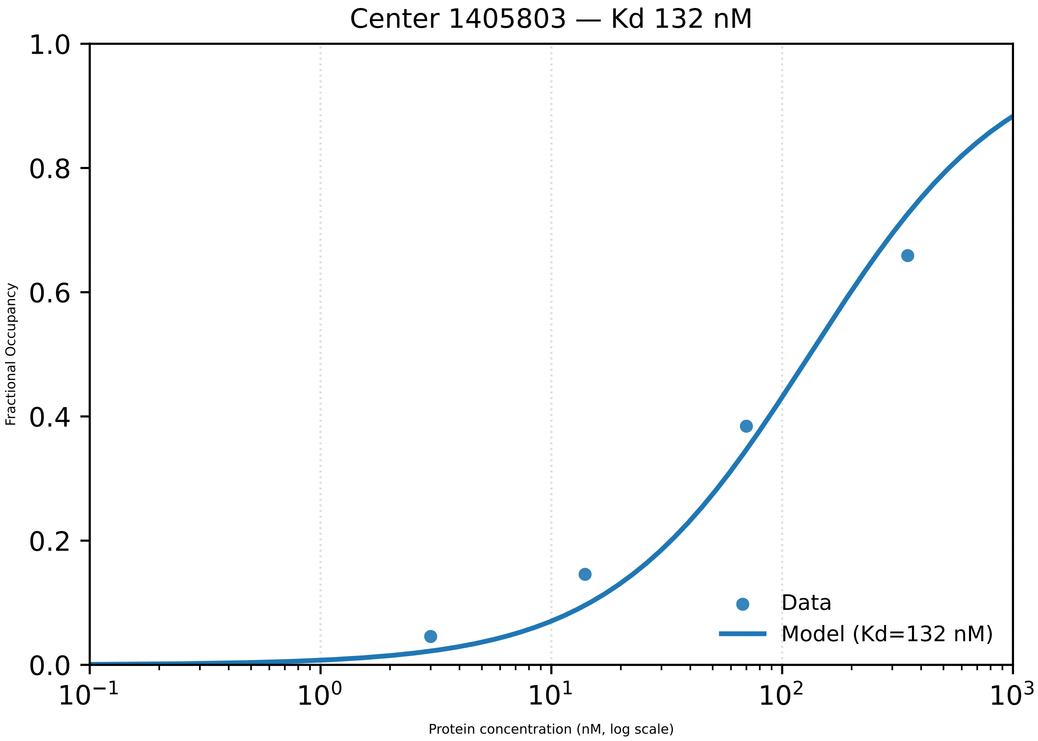

Center 1405803 — Kd 132 nM
1.0
0.8
0.6
0.4
0.2
0.0
Fractional Occupancy
$10^{-1}$
$10^{0}$
$10^{1}$
$10^{2}$
$10^{3}$
Protein concentration (nM, log scale)
Data
Model (Kd=132 nM)

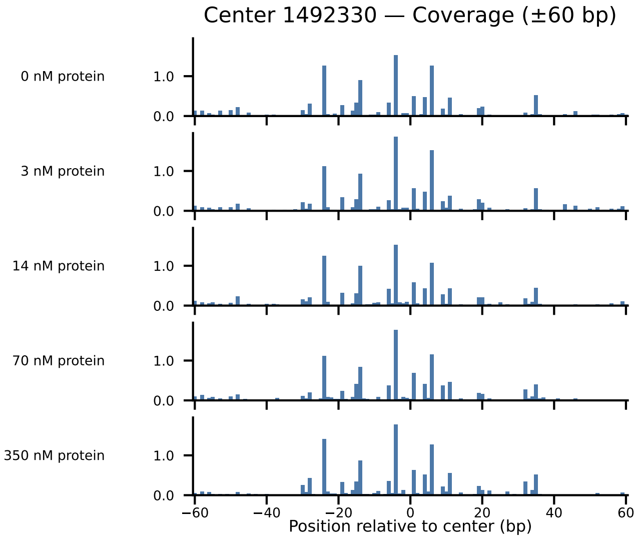

Center 1492330 — Coverage (±60 bp)
0 nM protein
3 nM protein
14 nM protein
70 nM protein
350 nM protein
1.0
0.0
−60
−40
−20
0
20
40
60
Position relative to center (bp)

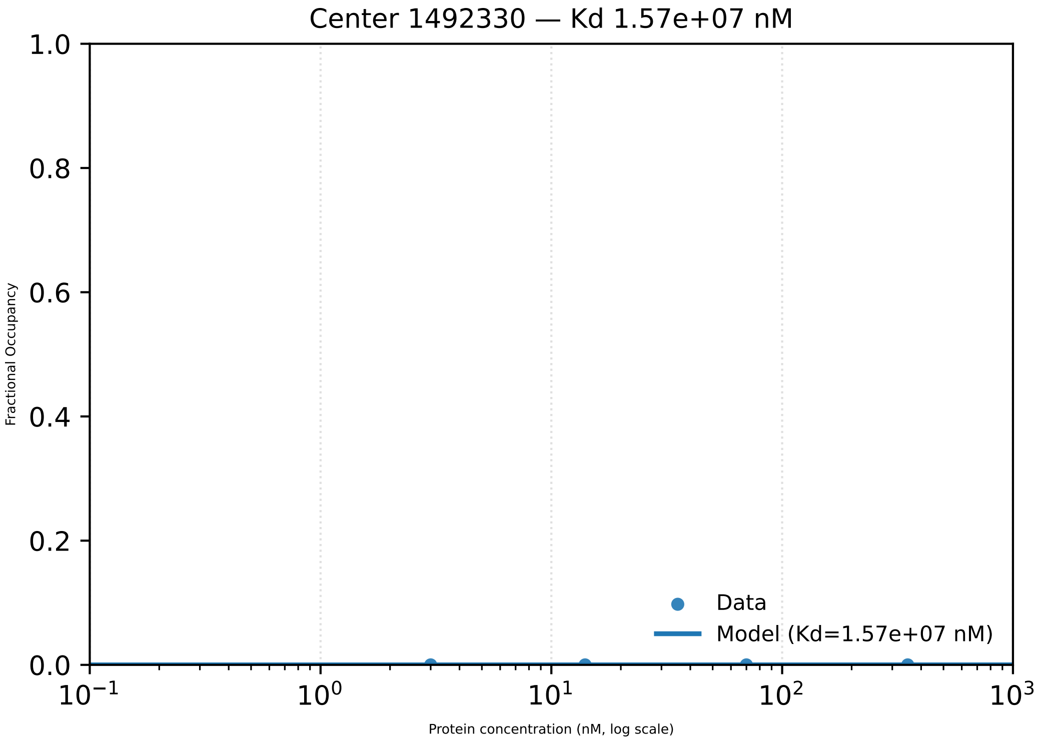
Center 1492330 — Kd 1.57e+07 nM
1.0
0.8
0.6
0.4
0.2
0.0
Fractional Occupancy
Data
Model (Kd=1.57e+07 nM)
$10^{-1}$
$10^{0}$
$10^{1}$
$10^{2}$
$10^{3}$
Protein concentration (nM, log scale)

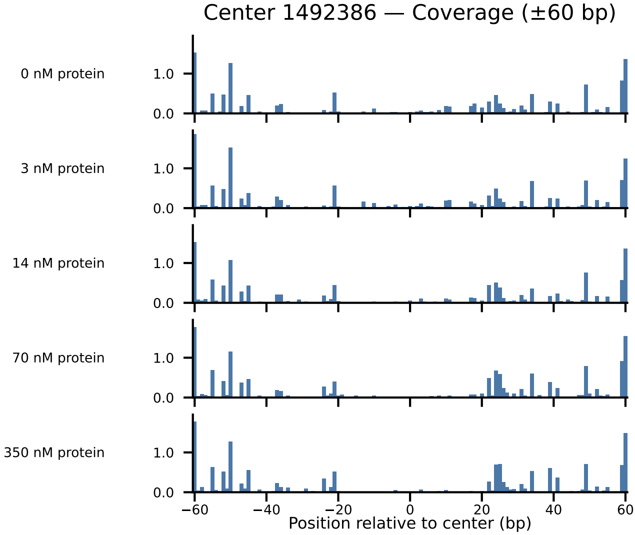
Center 1492386 — Coverage (±60 bp)
0 nM protein
3 nM protein
14 nM protein
70 nM protein
350 nM protein
1.0
0.0
−60
−40
−20
0
20
40
60
Position relative to center (bp)

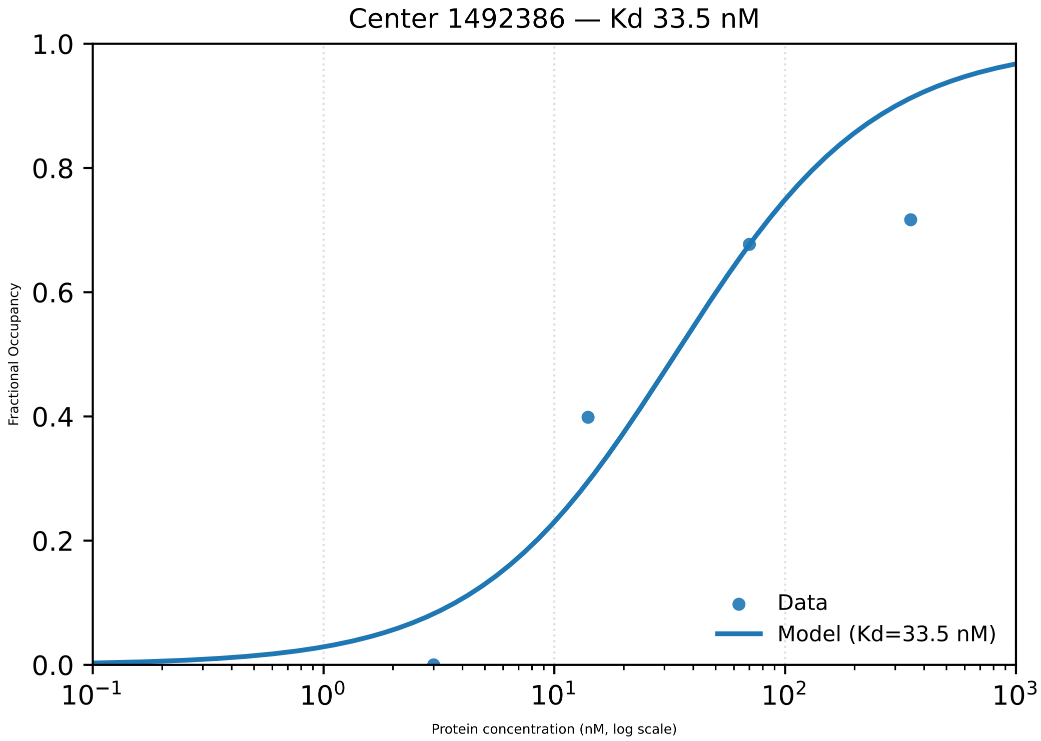

Center 1492386 — Kd 33.5 nM
1.0
0.8
0.6
0.4
0.2
0.0
Fractional Occupancy
$10^{-1}$
$10^{0}$
$10^{1}$
$10^{2}$
$10^{3}$
Protein concentration (nM, log scale)
Data
Model (Kd=33.5 nM)

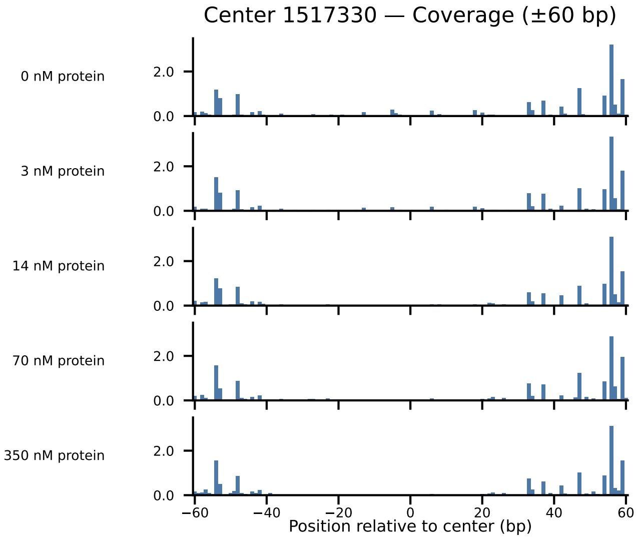

Center 1517330 — Coverage (±60 bp)
0 nM protein
3 nM protein
14 nM protein
70 nM protein
350 nM protein
2.0
0.0
−60
−40
−20
0
20
40
60
Position relative to center (bp)

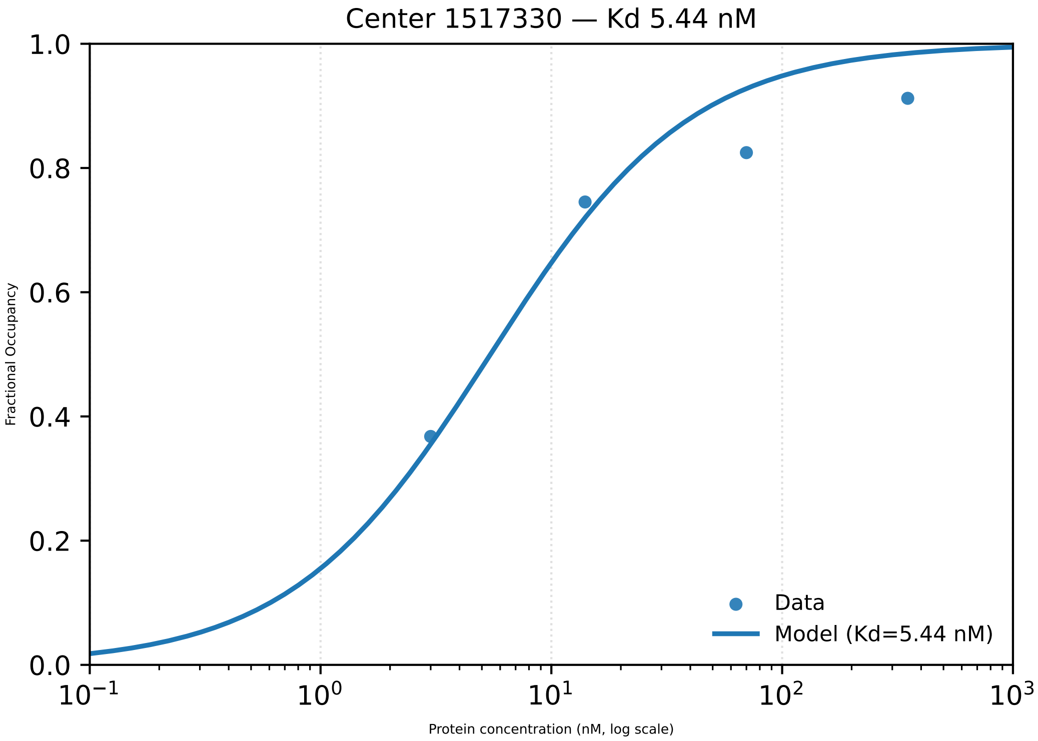
Center 1517330 — Kd 5.44 nM
Fractional Occupancy
1.0
0.8
0.6
0.4
0.2
0.0
$10^{-1}$
$10^{0}$
$10^{1}$
$10^{2}$
$10^{3}$
Protein concentration (nM, log scale)
Data
Model (Kd=5.44 nM)

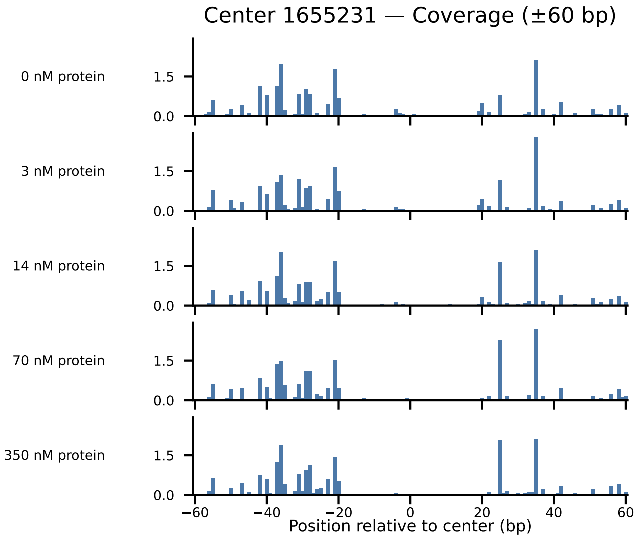

Center 1655231 — Coverage (±60 bp)
0 nM protein
3 nM protein
14 nM protein
70 nM protein
350 nM protein
1.5
0.0
−60
−40
−20
0
20
40
60
Position relative to center (bp)

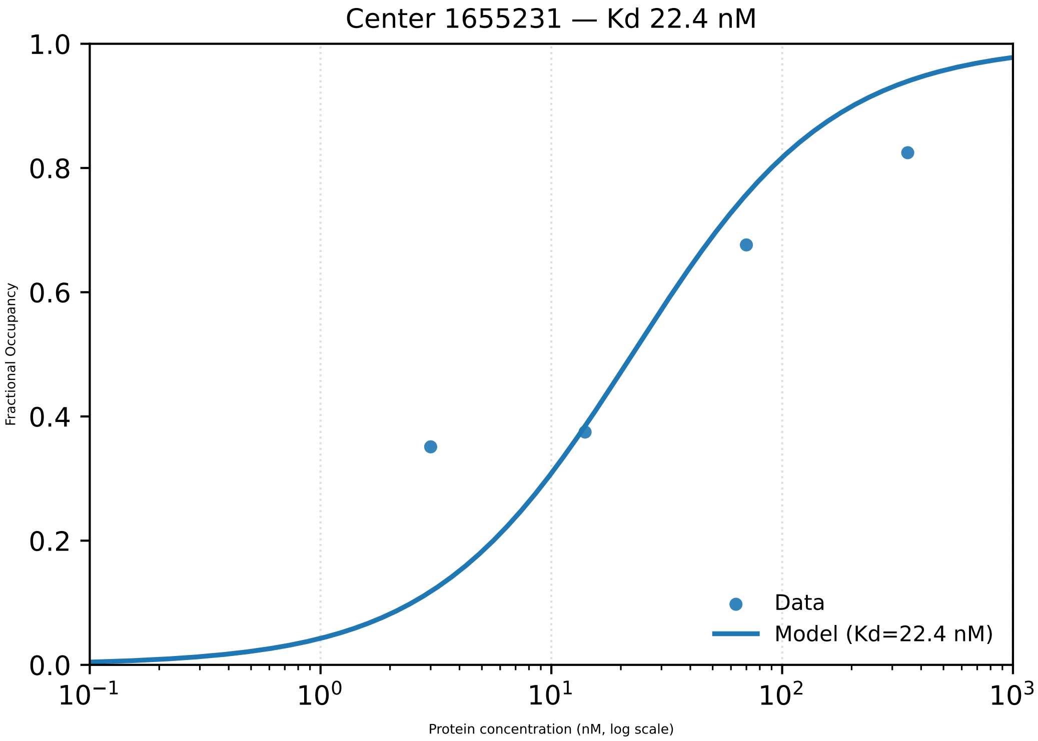

Center 1655231 — Kd 22.4 nM
1.0
0.8
0.6
0.4
0.2
0.0
Fractional Occupancy
$10^{-1}$
$10^{0}$
$10^{1}$
$10^{2}$
$10^{3}$
Protein concentration (nM, log scale)
Data
Model (Kd=22.4 nM)

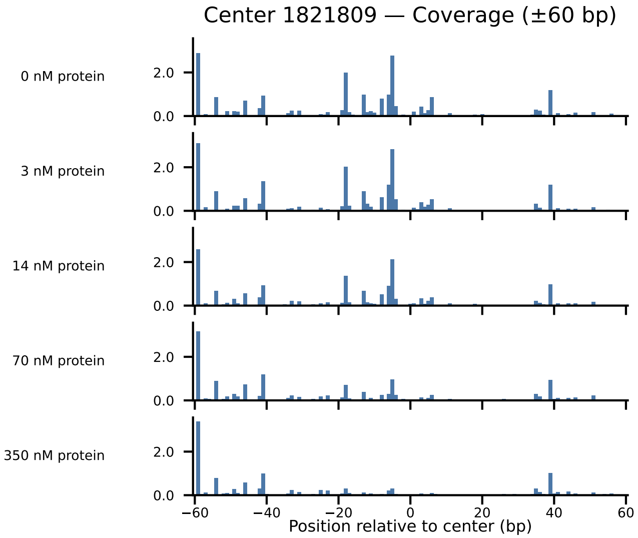

Center 1821809 — Coverage (±60 bp)
0 nM protein
3 nM protein
14 nM protein
70 nM protein
350 nM protein
2.0
0.0
−60
−40
−20
0
20
40
60
Position relative to center (bp)

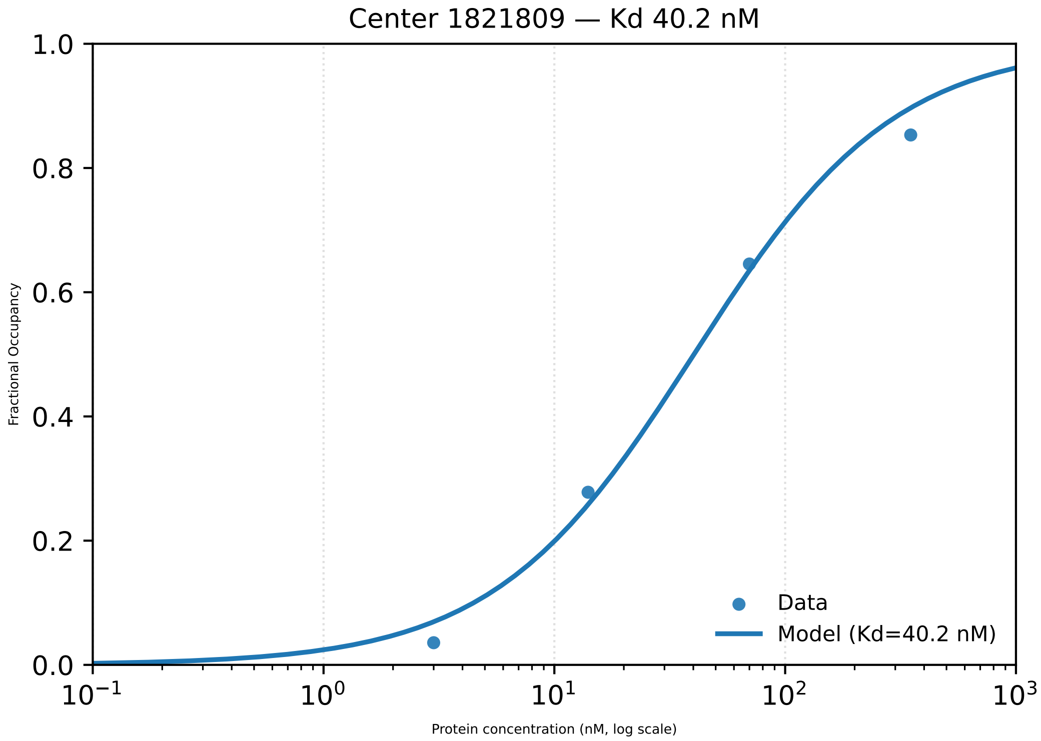

Center 1821809 — Kd 40.2 nM
Fractional Occupancy
Protein concentration (nM, log scale)
Data
Model (Kd=40.2 nM)

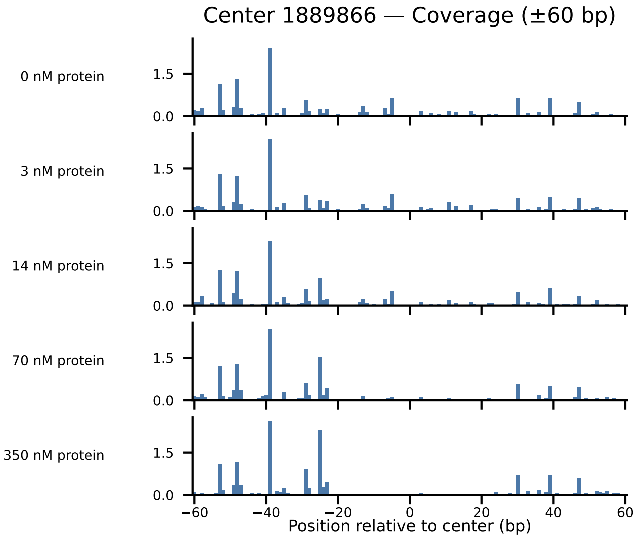
Center 1889866 — Coverage (±60 bp)
0 nM protein
3 nM protein
14 nM protein
70 nM protein
350 nM protein
1.5
0.0
−60
−40
−20
0
20
40
60
Position relative to center (bp)

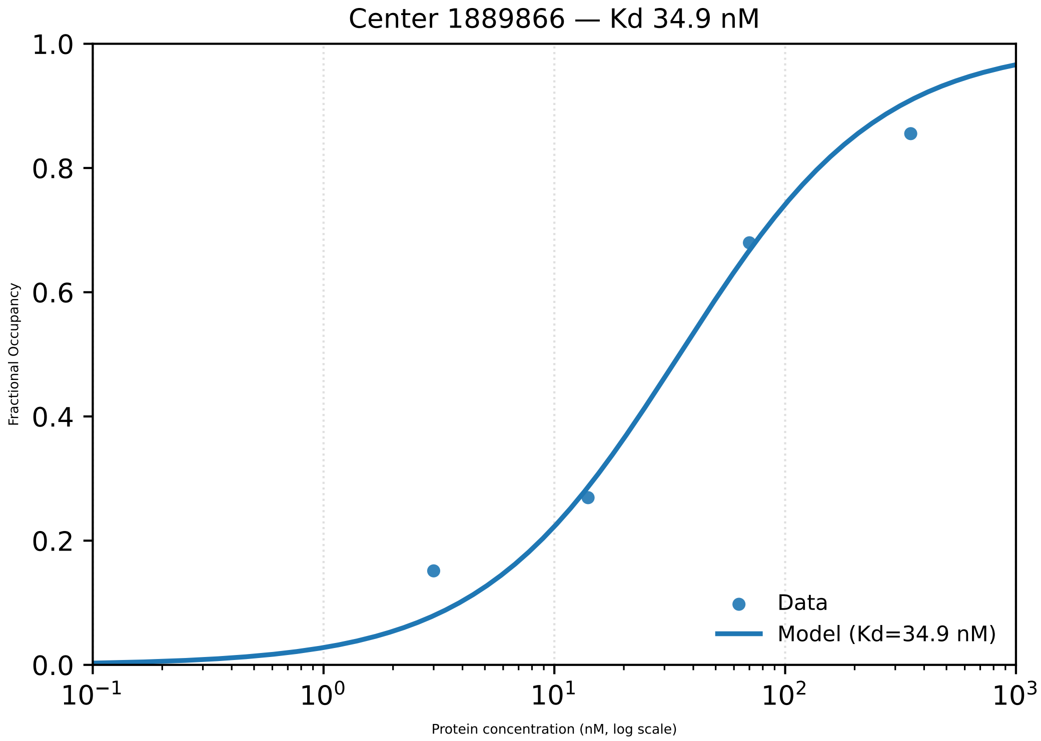

Center 1889866 — Kd 34.9 nM
Fractional Occupancy
1.0
0.8
0.6
0.4
0.2
0.0
$10^{-1}$
$10^{0}$
$10^{1}$
$10^{2}$
$10^{3}$
Protein concentration (nM, log scale)
Data
Model (Kd=34.9 nM)

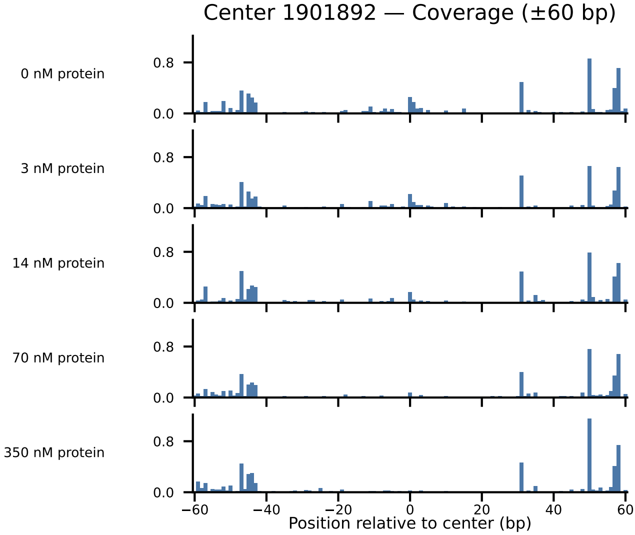
Center 1901892 — Coverage (±60 bp)
0 nM protein
3 nM protein
14 nM protein
70 nM protein
350 nM protein
0.8
0.0
−60
−40
−20
0
20
40
60
Position relative to center (bp)

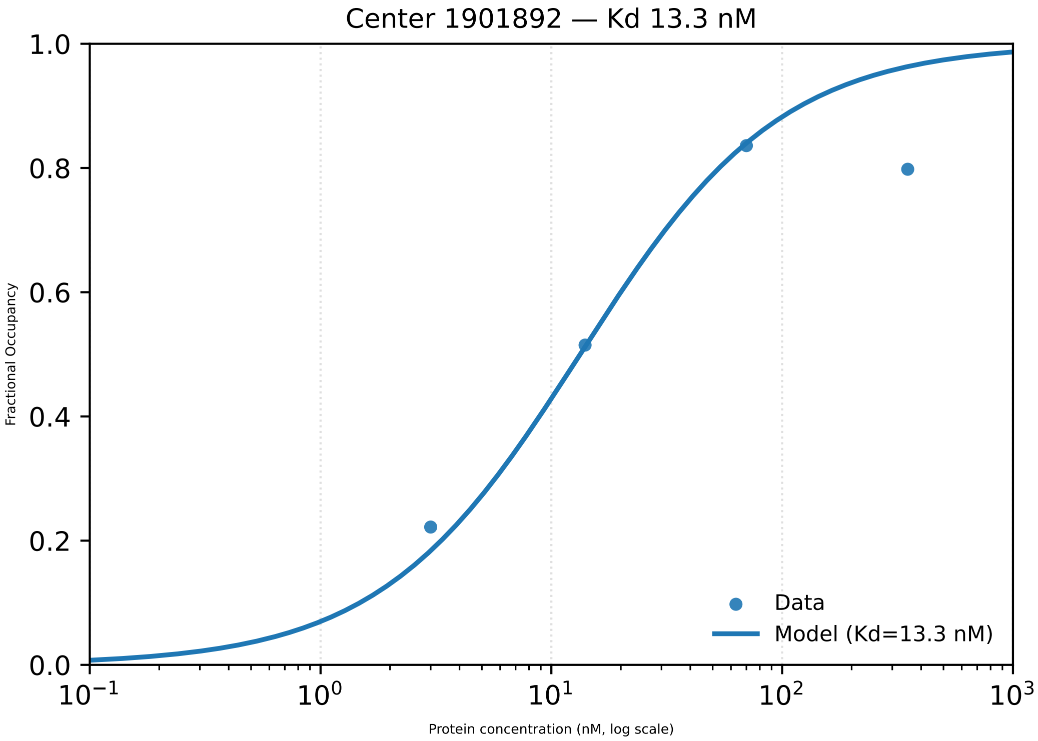

Center 1901892 — Kd 13.3 nM
1.0
0.8
0.6
0.4
0.2
0.0
Fractional Occupancy
$10^{-1}$
$10^{0}$
$10^{1}$
$10^{2}$
$10^{3}$
Protein concentration (nM, log scale)
Data
Model (Kd=13.3 nM)

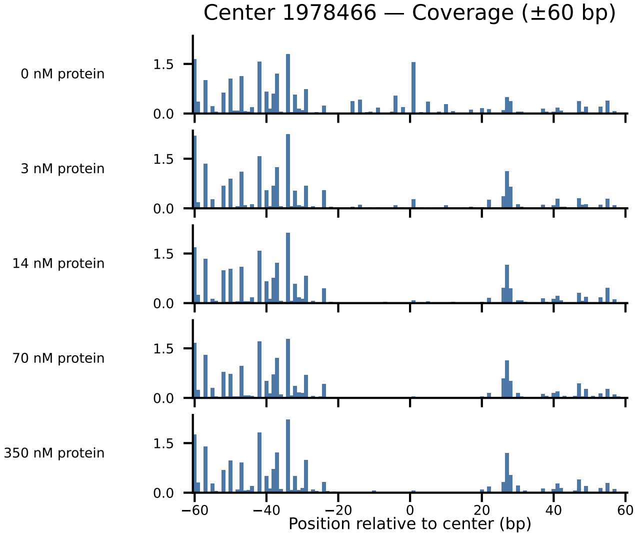
Center 1978466 — Coverage (±60 bp)
0 nM protein
3 nM protein
14 nM protein
70 nM protein
350 nM protein
1.5
0.0
−60
−40
−20
0
20
40
60
Position relative to center (bp)

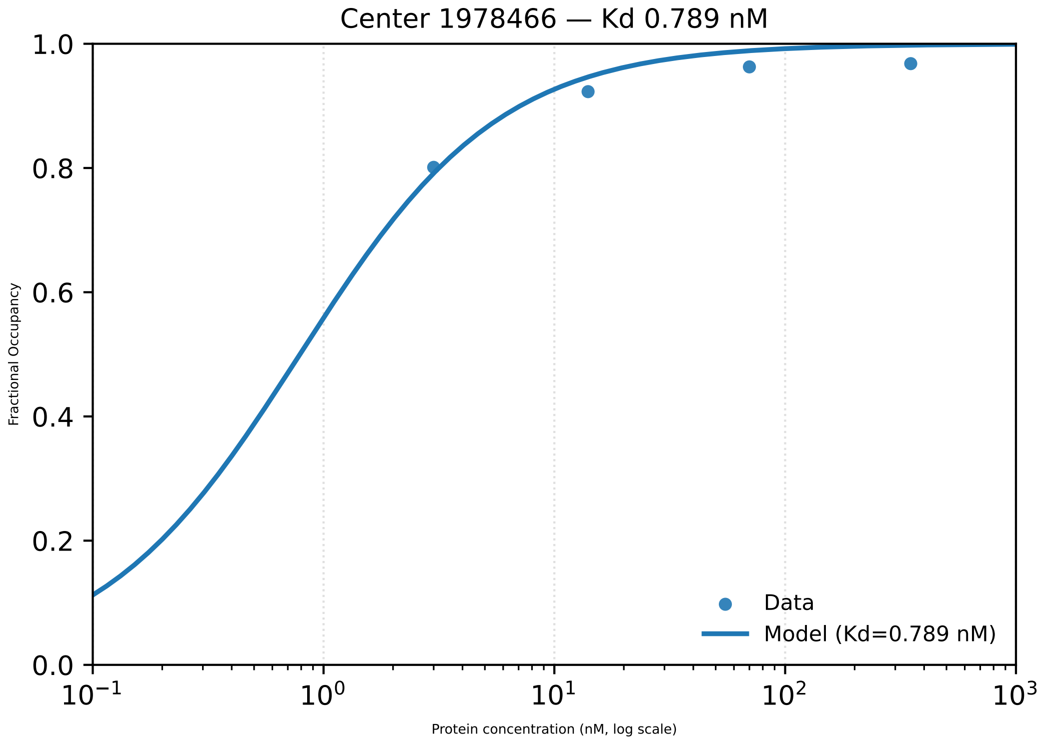

Center 1978466 — Kd 0.789 nM
Fractional Occupancy
0.0
0.2
0.4
0.6
0.8
1.0
$10^{-1}$
$10^{0}$
$10^{1}$
$10^{2}$
$10^{3}$
Protein concentration (nM, log scale)
Data
Model (Kd=0.789 nM)

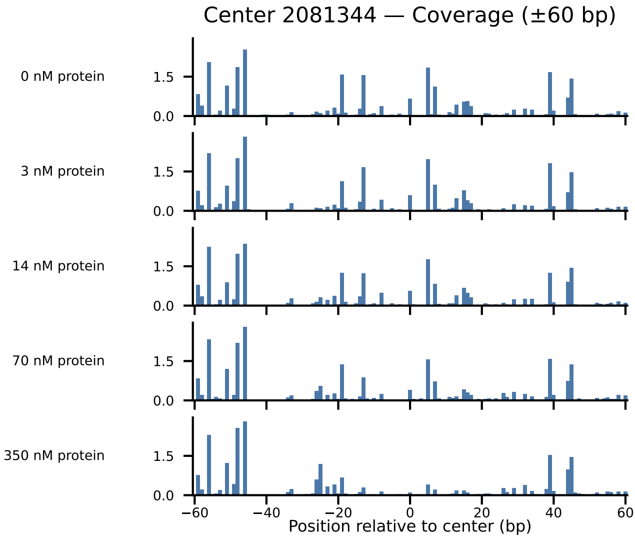

Center 2081344 — Coverage (±60 bp)
0 nM protein
3 nM protein
14 nM protein
70 nM protein
350 nM protein
1.5
0.0
−60
−40
−20
0
20
40
60
Position relative to center (bp)

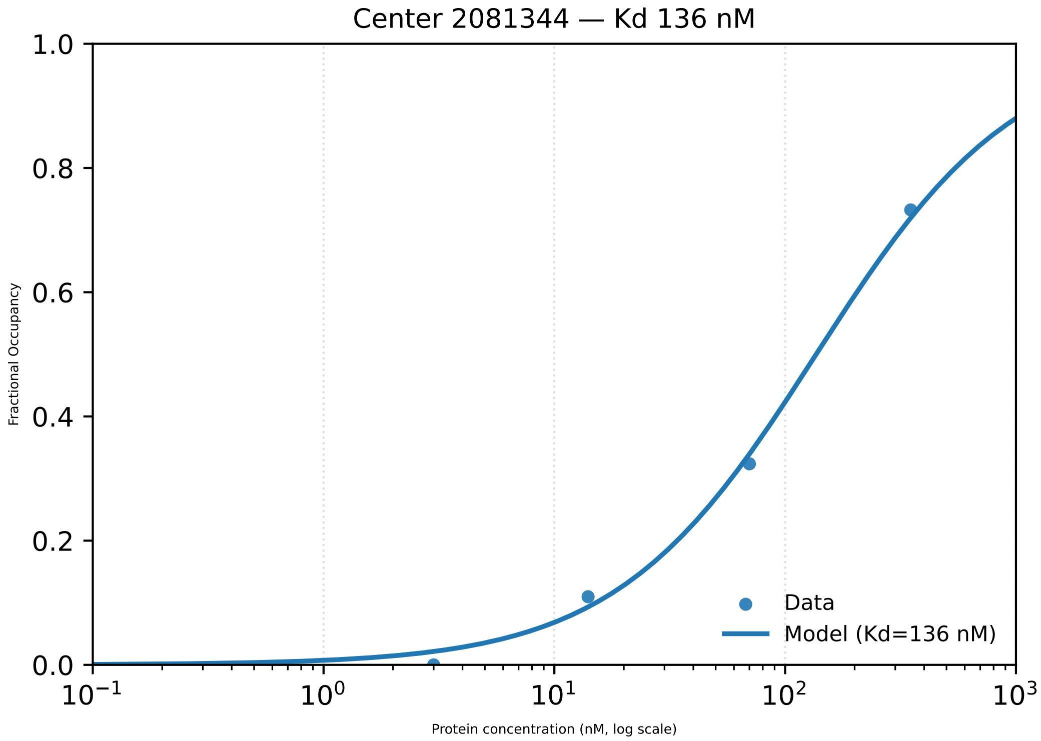

Center 2081344 — Kd 136 nM
Fractional Occupancy
Protein concentration (nM, log scale)
Data
Model (Kd=136 nM)

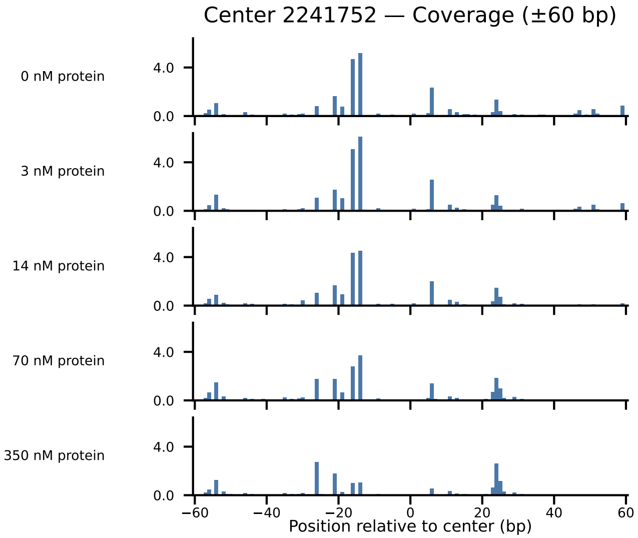

Center 2241752 — Coverage (±60 bp)
0 nM protein
3 nM protein
14 nM protein
70 nM protein
350 nM protein
4.0
0.0
−60
−40
−20
0
20
40
60
Position relative to center (bp)

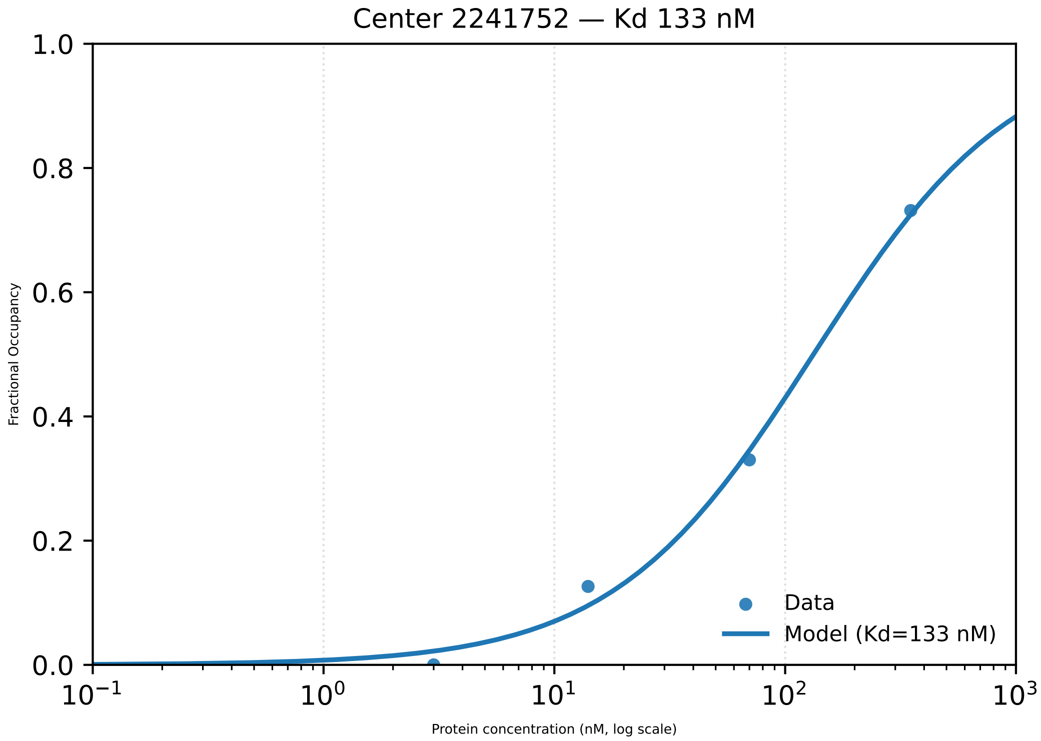

Center 2241752 — Kd 133 nM
Fractional Occupancy
1.0
0.8
0.6
0.4
0.2
0.0
$10^{-1}$
$10^{0}$
$10^{1}$
$10^{2}$
$10^{3}$
Protein concentration (nM, log scale)
Data
Model (Kd=133 nM)

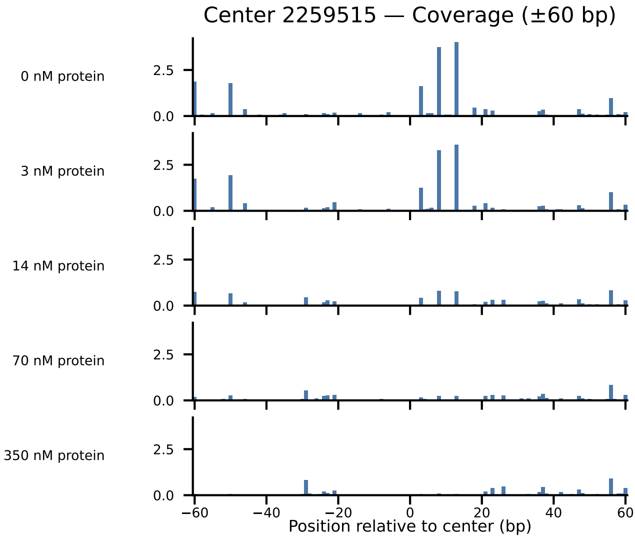

Center 2259515 — Coverage (±60 bp)
0 nM protein
3 nM protein
14 nM protein
70 nM protein
350 nM protein
2.5
0.0
−60
−40
−20
0
20
40
60
Position relative to center (bp)

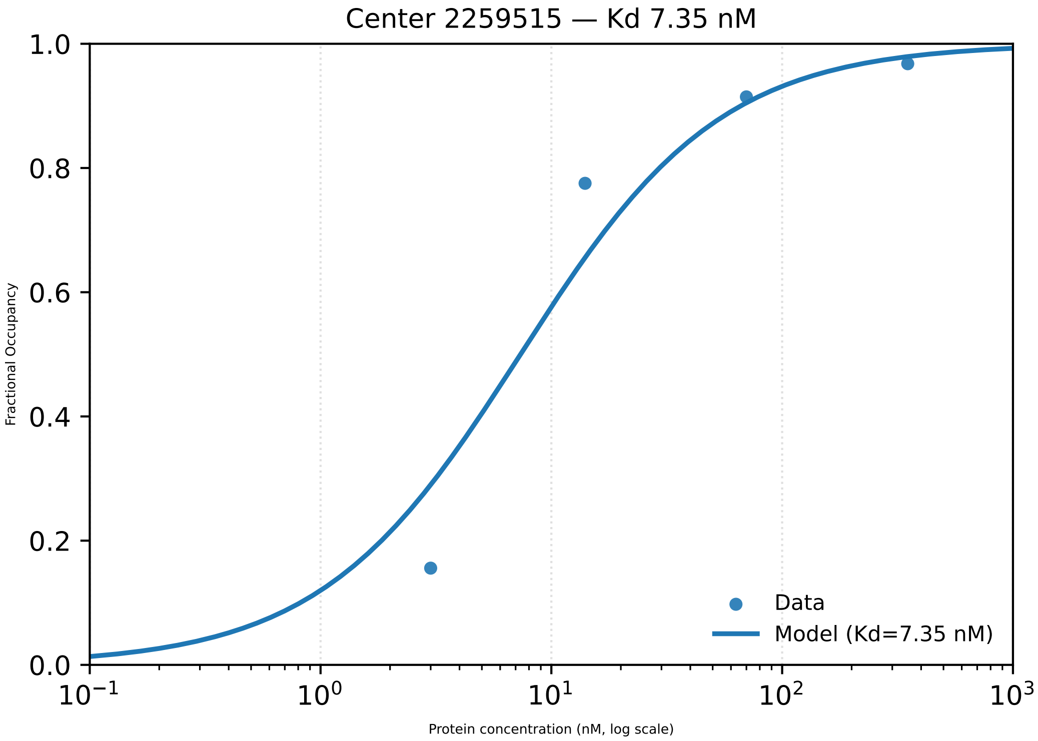
Center 2259515 — Kd 7.35 nM
Fractional Occupancy
1.0
0.8
0.6
0.4
0.2
0.0
$10^{-1}$
$10^{0}$
$10^{1}$
$10^{2}$
$10^{3}$
Protein concentration (nM, log scale)
Data
Model (Kd=7.35 nM)

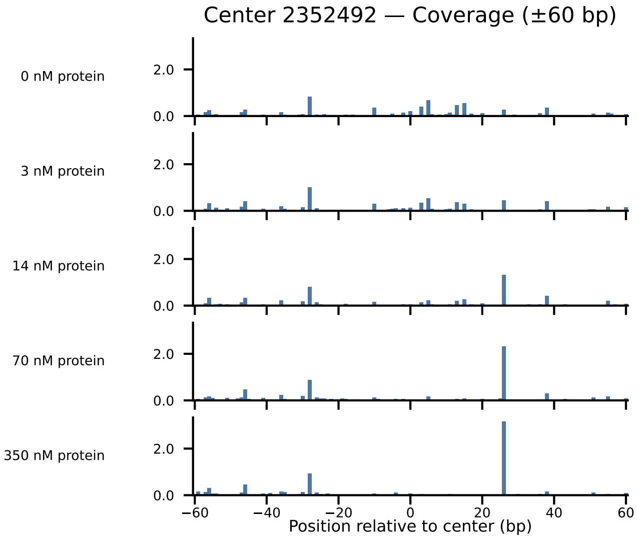

Center 2352492 — Coverage (±60 bp)
0 nM protein
3 nM protein
14 nM protein
70 nM protein
350 nM protein
2.0
0.0
−60
−40
−20
0
20
40
60
Position relative to center (bp)

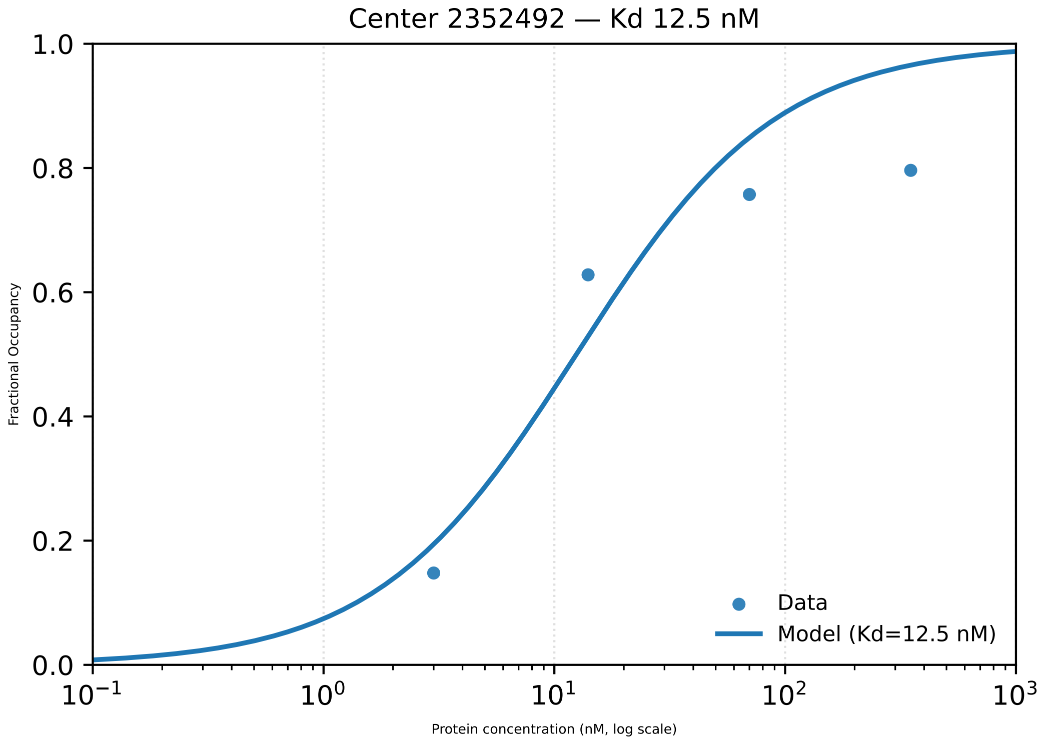

Center 2352492 — Kd 12.5 nM
Fractional Occupancy
1.0
0.8
0.6
0.4
0.2
0.0
$10^{-1}$
$10^{0}$
$10^{1}$
$10^{2}$
$10^{3}$
Protein concentration (nM, log scale)
Data
Model (Kd=12.5 nM)

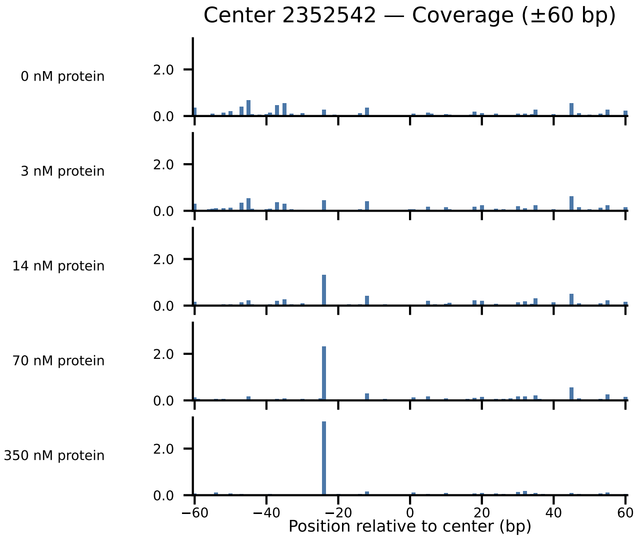

Center 2352542 — Coverage (±60 bp)
0 nM protein
3 nM protein
14 nM protein
70 nM protein
350 nM protein
2.0
0.0
−60
−40
−20
0
20
40
60
Position relative to center (bp)

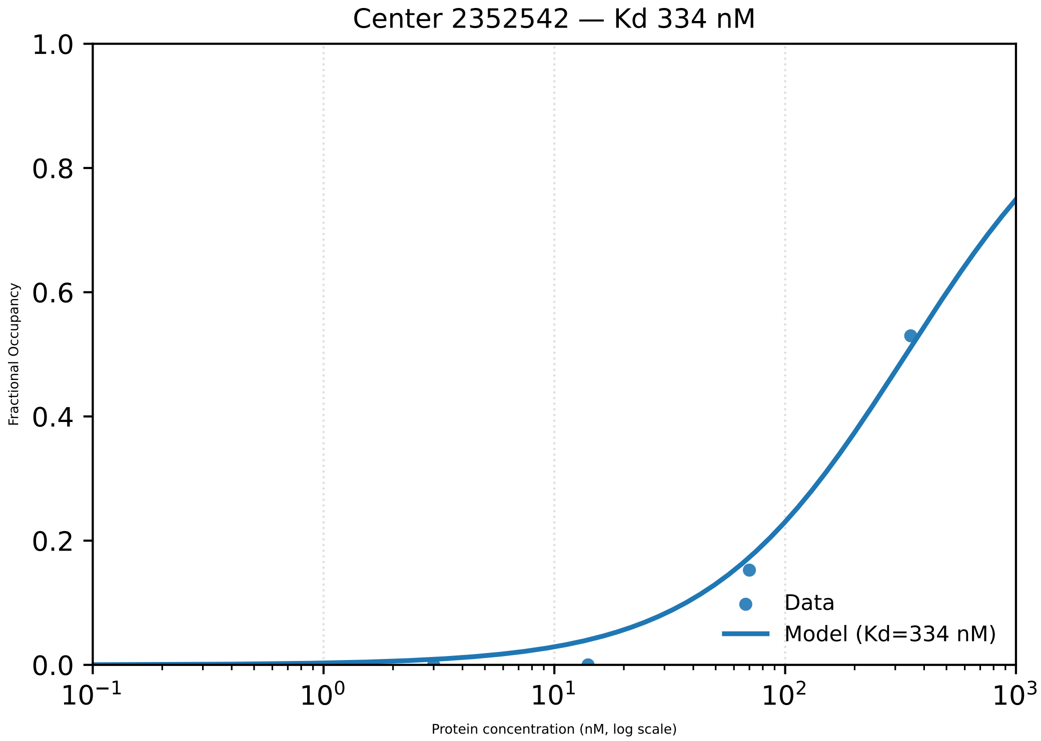

Center 2352542 — Kd 334 nM
1.0
0.8
0.6
0.4
0.2
0.0
Fractional Occupancy
$10^{-1}$
$10^{0}$
$10^{1}$
$10^{2}$
$10^{3}$
Protein concentration (nM, log scale)
Data
Model (Kd=334 nM)

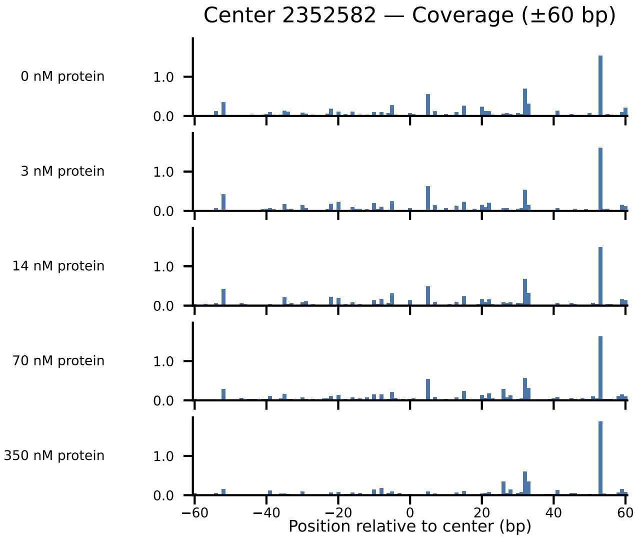

Center 2352582 — Coverage (±60 bp)
0 nM protein
3 nM protein
14 nM protein
70 nM protein
350 nM protein
1.0
0.0
−60
−40
−20
0
20
40
60
Position relative to center (bp)

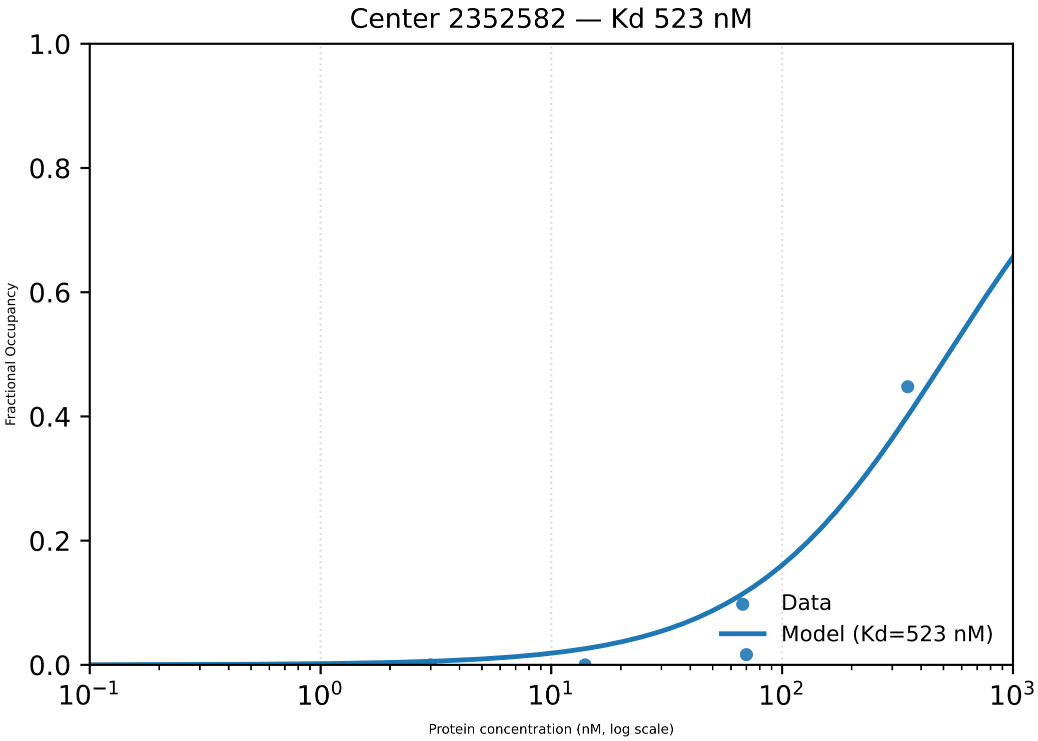
Center 2352582 — Kd 523 nM
1.0
0.8
0.6
0.4
0.2
0.0
Fractional Occupancy
$10^{-1}$
$10^{0}$
$10^{1}$
$10^{2}$
$10^{3}$
Protein concentration (nM, log scale)
Data
Model (Kd=523 nM)

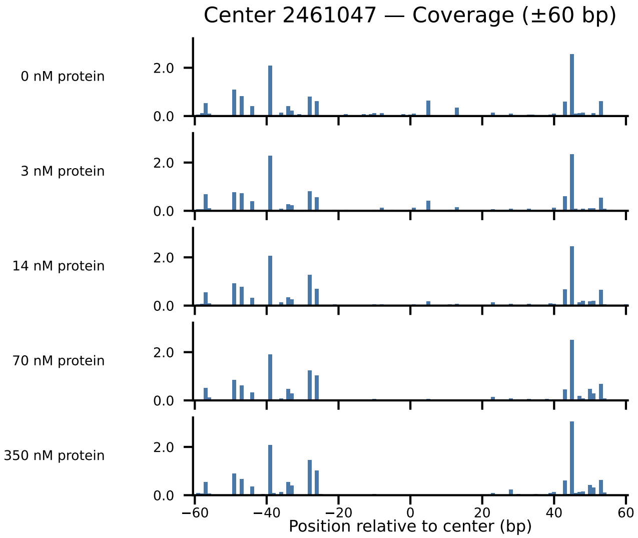
Center 2461047 — Coverage (±60 bp)
0 nM protein
3 nM protein
14 nM protein
70 nM protein
350 nM protein
2.0
0.0
−60
−40
−20
0
20
40
60
Position relative to center (bp)

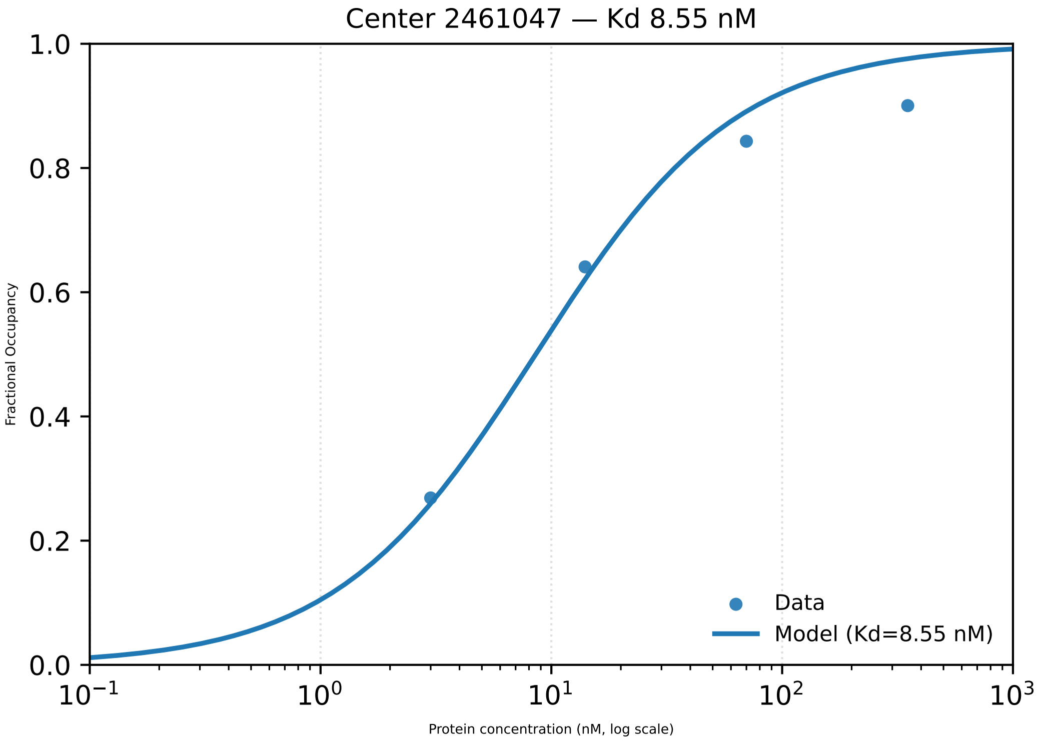

Center 2461047 — Kd 8.55 nM
1.0
0.8
0.6
0.4
0.2
0.0
Fractional Occupancy
$10^{-1}$
$10^{0}$
$10^{1}$
$10^{2}$
$10^{3}$
Protein concentration (nM, log scale)
Data
Model (Kd=8.55 nM)

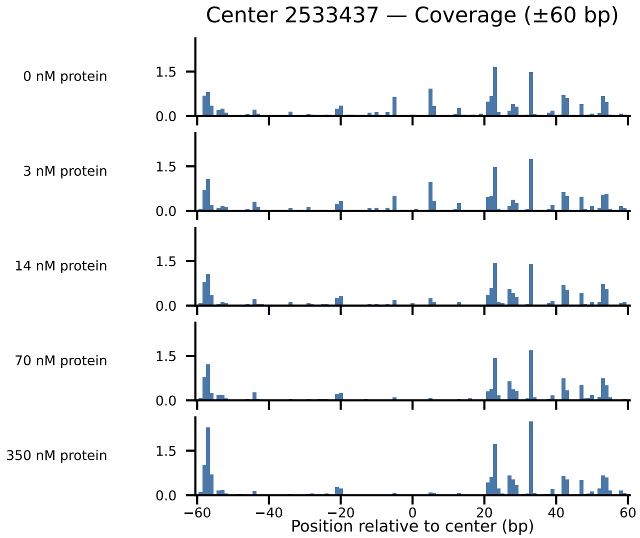
Center 2533437 — Coverage (±60 bp)
0 nM protein
3 nM protein
14 nM protein
70 nM protein
350 nM protein
1.5
0.0
−60
−40
−20
0
20
40
60
Position relative to center (bp)

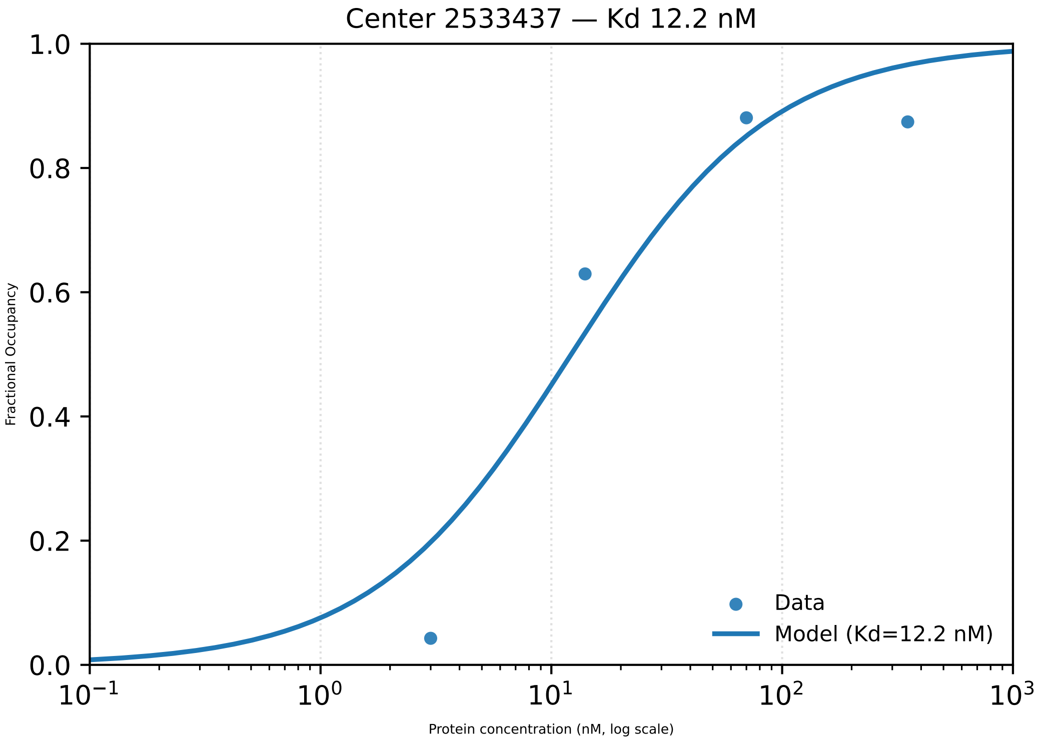

Center 2533437 — Kd 12.2 nM
1.0
0.8
0.6
0.4
0.2
0.0
Fractional Occupancy
$10^{-1}$
$10^{0}$
$10^{1}$
$10^{2}$
$10^{3}$
Protein concentration (nM, log scale)
Data
Model (Kd=12.2 nM)

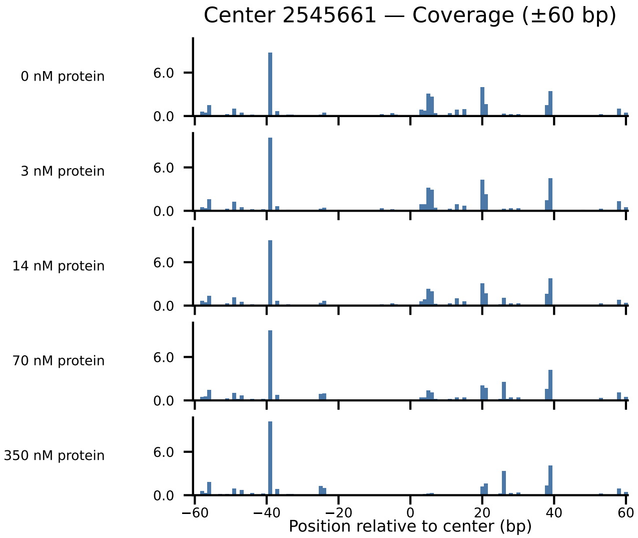
Center 2545661 — Coverage (±60 bp)
0 nM protein
3 nM protein
14 nM protein
70 nM protein
350 nM protein
6.0
0.0
−60
−40
−20
0
20
40
60
Position relative to center (bp)

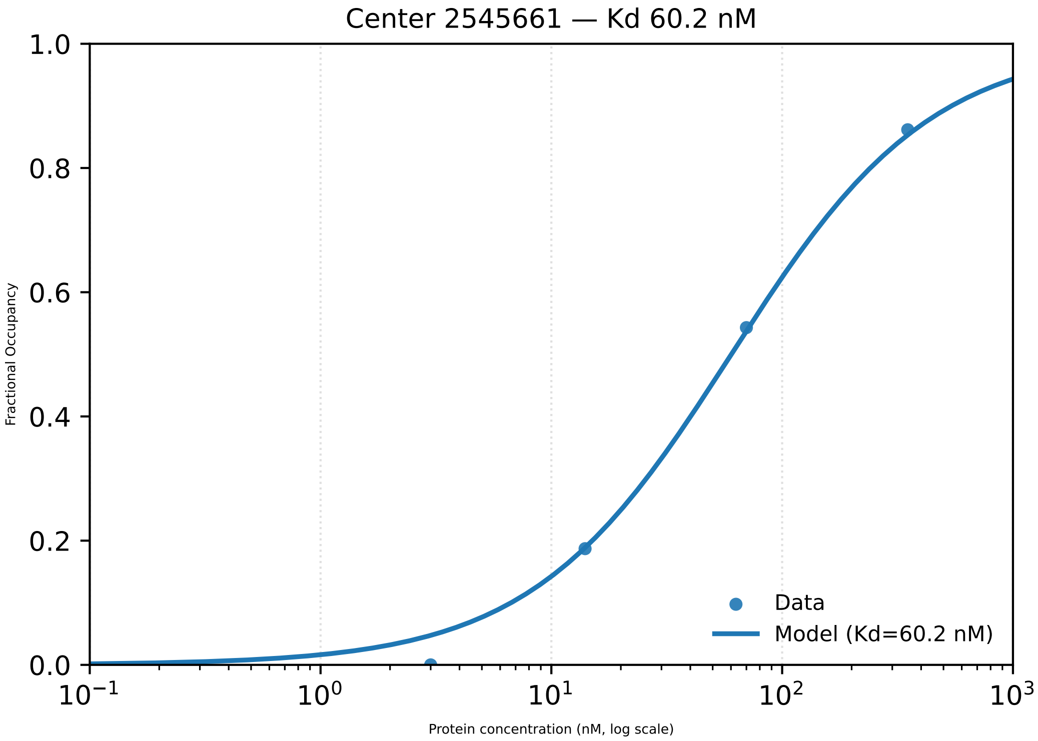

Center 2545661 — Kd 60.2 nM
1.0
0.8
0.6
0.4
0.2
0.0
Fractional Occupancy
$10^{-1}$
$10^{0}$
$10^{1}$
$10^{2}$
$10^{3}$
Protein concentration (nM, log scale)
Data
Model (Kd=60.2 nM)

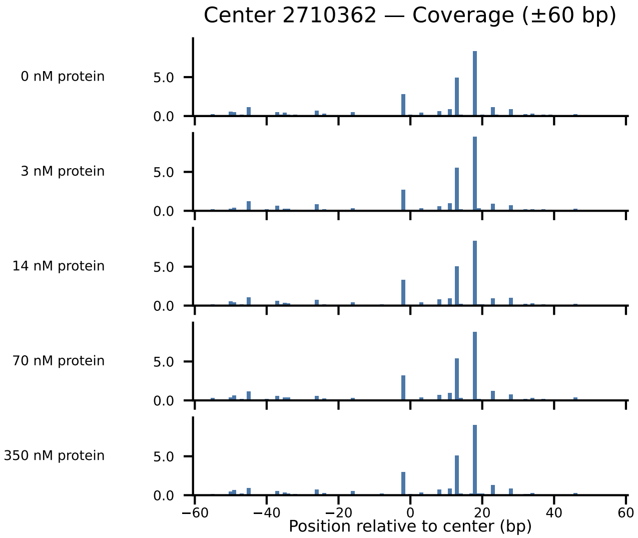

Center 2710362 — Coverage (±60 bp)
0 nM protein
3 nM protein
14 nM protein
70 nM protein
350 nM protein
5.0
0.0
−60
−40
−20
0
20
40
60
Position relative to center (bp)

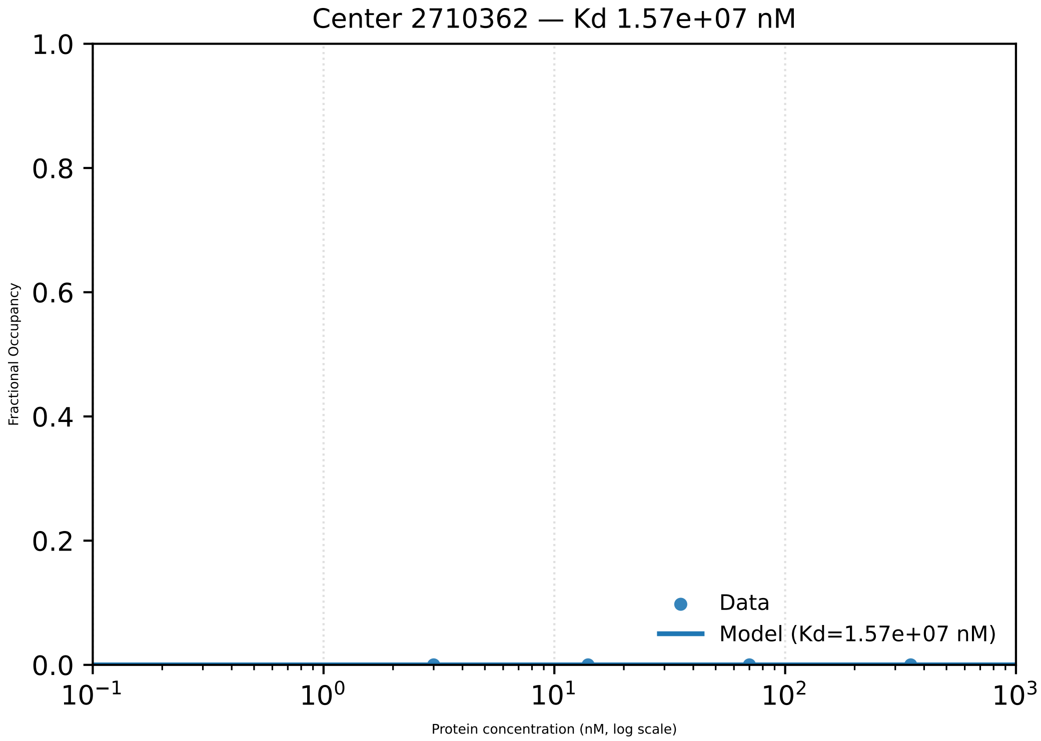

Center 2710362 — Kd 1.57e+07 nM
Fractional Occupancy
1.0
0.8
0.6
0.4
0.2
0.0
Data
Model (Kd=1.57e+07 nM)
$10^{-1}$
$10^{0}$
$10^{1}$
$10^{2}$
$10^{3}$
Protein concentration (nM, log scale)

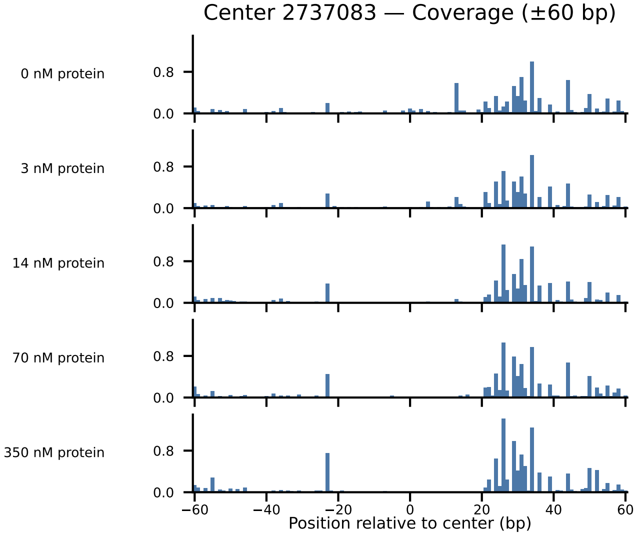

Center 2737083 — Coverage (±60 bp)
0 nM protein
3 nM protein
14 nM protein
70 nM protein
350 nM protein
0.8
0.0
−60
−40
−20
0
20
40
60
Position relative to center (bp)

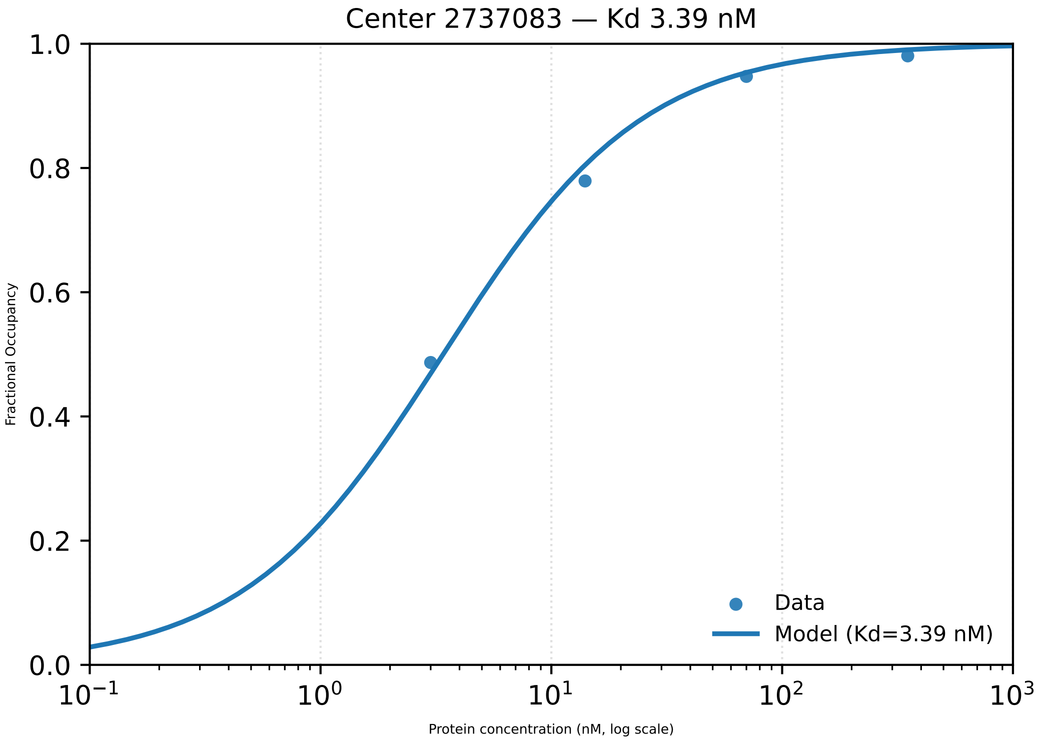

Center 2737083 — Kd 3.39 nM
Fractional Occupancy
1.0
0.8
0.6
0.4
0.2
0.0
$10^{-1}$
$10^{0}$
$10^{1}$
$10^{2}$
$10^{3}$
Protein concentration (nM, log scale)
Data
Model (Kd=3.39 nM)

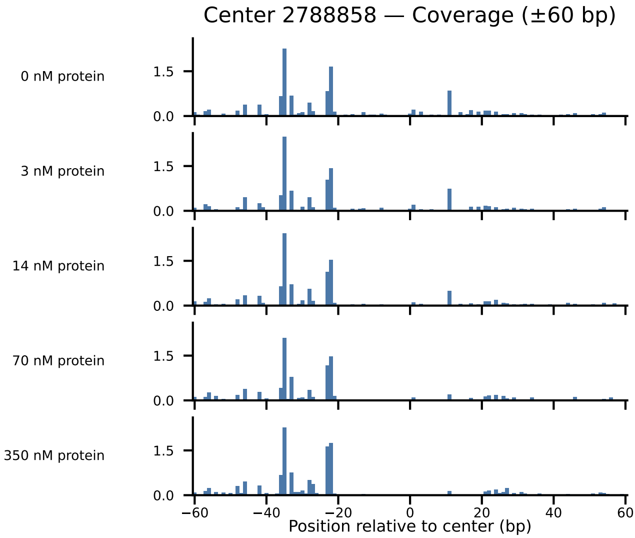

Center 2788858 — Coverage (±60 bp)
0 nM protein
3 nM protein
14 nM protein
70 nM protein
350 nM protein
1.5
0.0
−60
−40
−20
0
20
40
60
Position relative to center (bp)

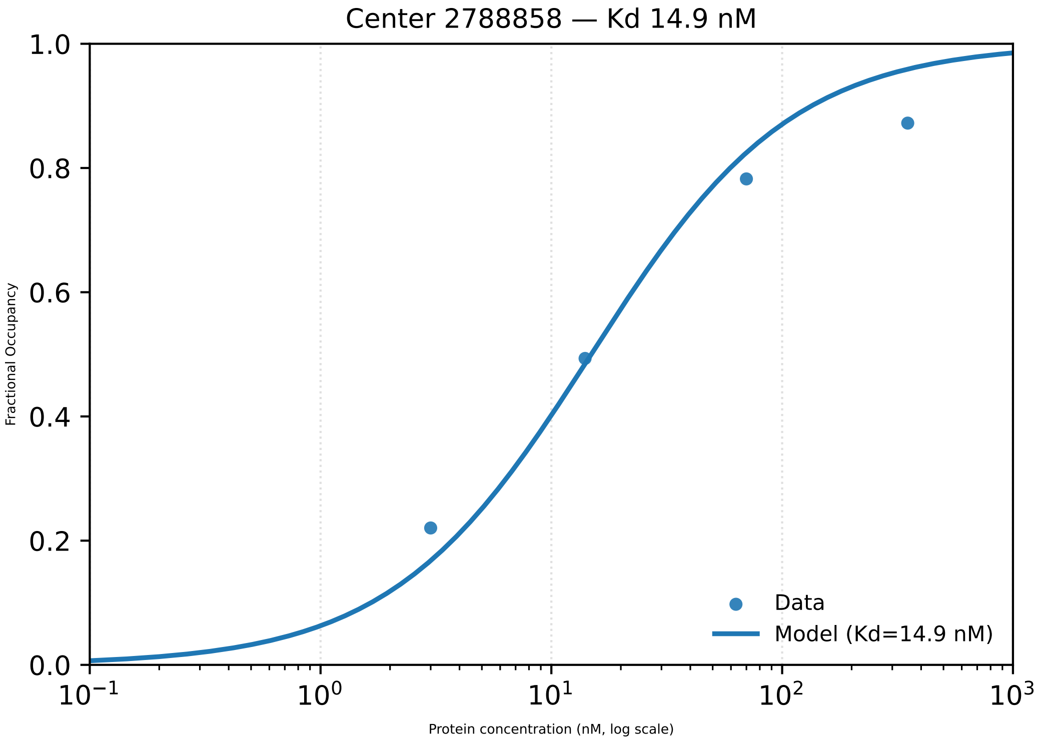

Center 2788858 — Kd 14.9 nM
Fractional Occupancy
Protein concentration (nM, log scale)
1.0
0.8
0.6
0.4
0.2
0.0
$10^{-1}$
$10^{0}$
$10^{1}$
$10^{2}$
$10^{3}$
Data
Model (Kd=14.9 nM)

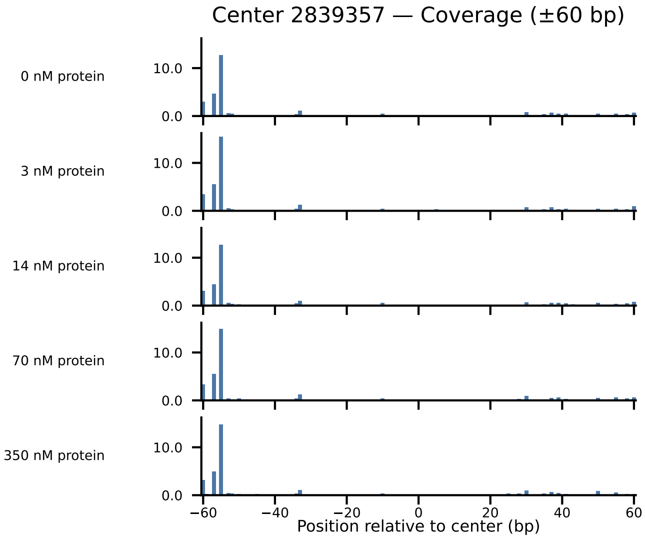

Center 2839357 — Coverage (±60 bp)
0 nM protein
3 nM protein
14 nM protein
70 nM protein
350 nM protein
10.0
0.0
−60
−40
−20
0
20
40
60
Position relative to center (bp)

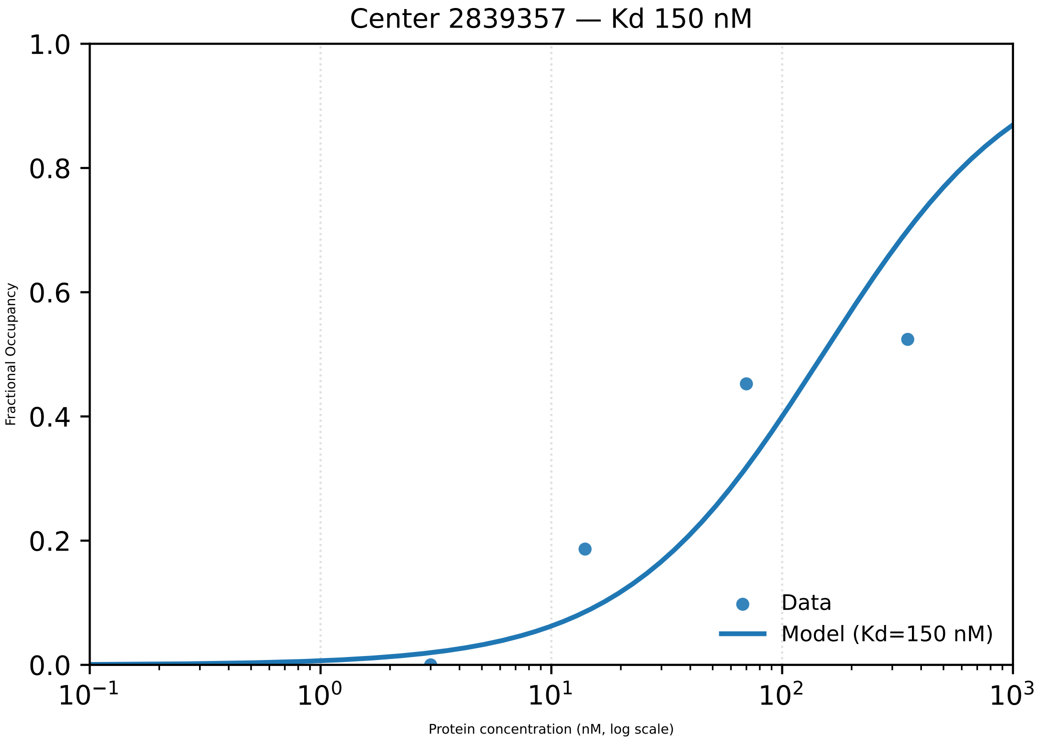

Center 2839357 — Kd 150 nM
1.0
0.8
0.6
0.4
0.2
0.0
Fractional Occupancy
$10^{-1}$
$10^{0}$
$10^{1}$
$10^{2}$
$10^{3}$
Protein concentration (nM, log scale)
Data
Model (Kd=150 nM)

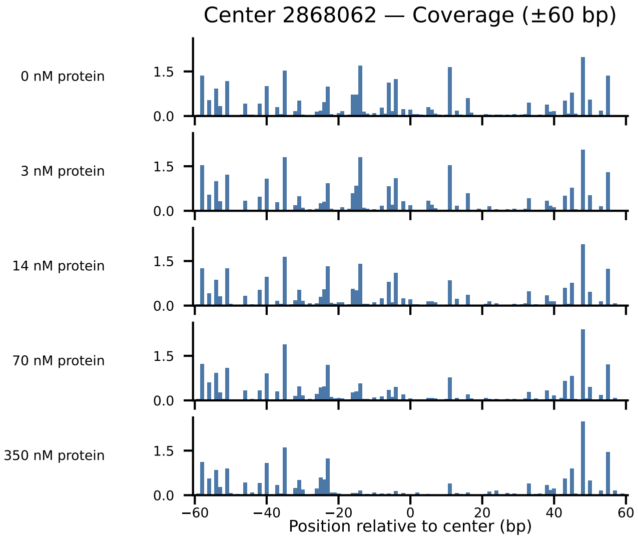

Center 2868062 — Coverage (±60 bp)
0 nM protein
3 nM protein
14 nM protein
70 nM protein
350 nM protein
1.5
0.0
−60
−40
−20
0
20
40
60
Position relative to center (bp)

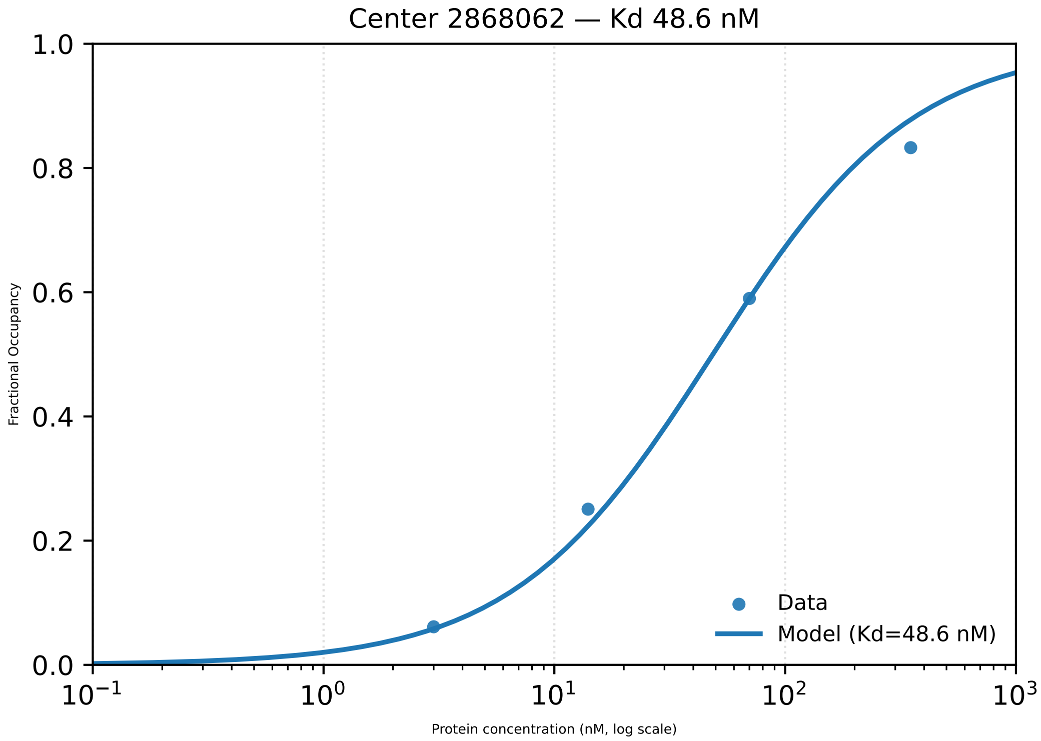

Center 2868062 — Kd 48.6 nM
1.0
0.8
0.6
0.4
0.2
0.0
Fractional Occupancy
$10^{-1}$
$10^{0}$
$10^{1}$
$10^{2}$
$10^{3}$
Protein concentration (nM, log scale)
Data
Model (Kd=48.6 nM)

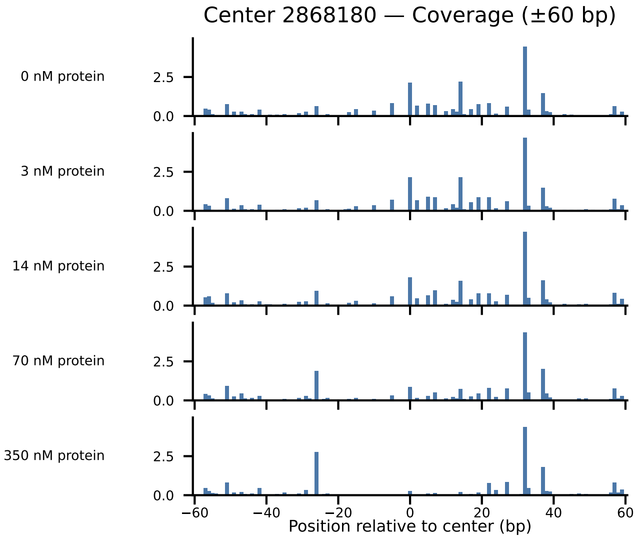
Center 2868180 — Coverage (±60 bp)
0 nM protein
3 nM protein
14 nM protein
70 nM protein
350 nM protein
2.5
0.0
−60
−40
−20
0
20
40
60
Position relative to center (bp)

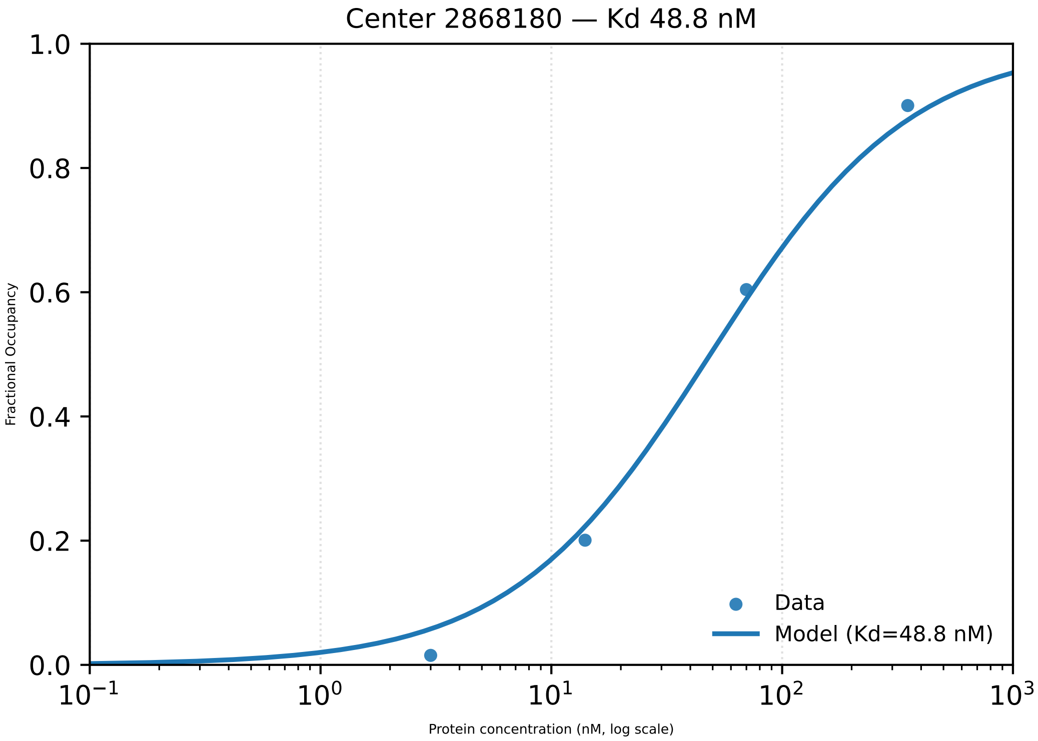

Center 2868180 — Kd 48.8 nM
Fractional Occupancy
1.0
0.8
0.6
0.4
0.2
0.0
$10^{-1}$
$10^{0}$
$10^{1}$
$10^{2}$
$10^{3}$
Protein concentration (nM, log scale)
Data
Model (Kd=48.8 nM)

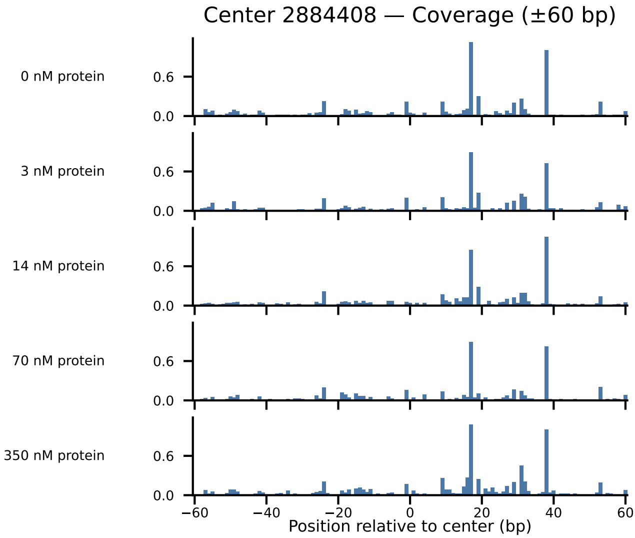

Center 2884408 — Coverage (±60 bp)
0 nM protein
3 nM protein
14 nM protein
70 nM protein
350 nM protein
0.6
0.0
−60
−40
−20
0
20
40
60
Position relative to center (bp)

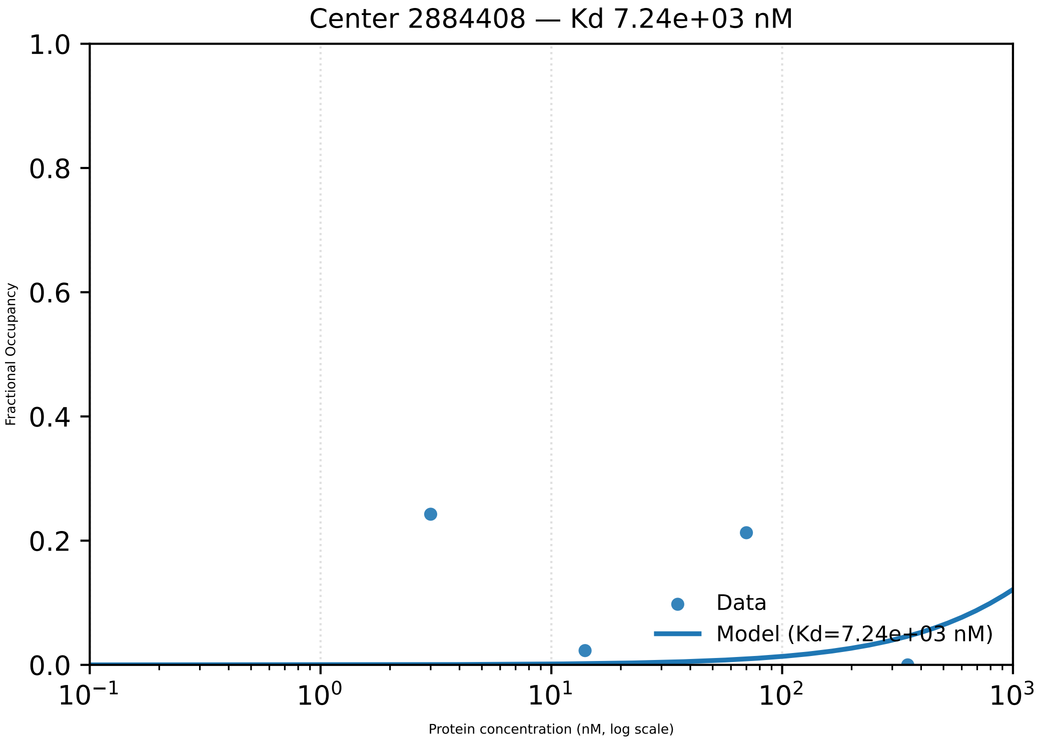

Center 2884408 — Kd 7.24e+03 nM
1.0
0.8
0.6
0.4
0.2
0.0
Fractional Occupancy
Data
Model (Kd=7.24e+03 nM)
$10^{-1}$
$10^{0}$
$10^{1}$
$10^{2}$
$10^{3}$
Protein concentration (nM, log scale)

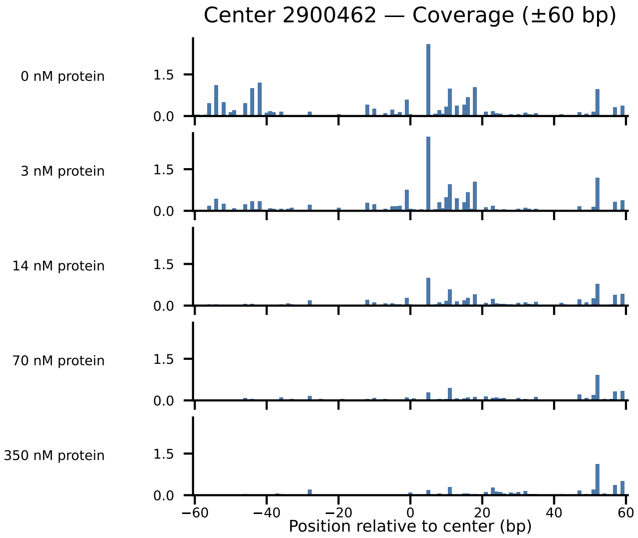

Center 2900462 — Coverage (±60 bp)
0 nM protein
3 nM protein
14 nM protein
70 nM protein
350 nM protein
1.5
0.0
−60
−40
−20
0
20
40
60
Position relative to center (bp)

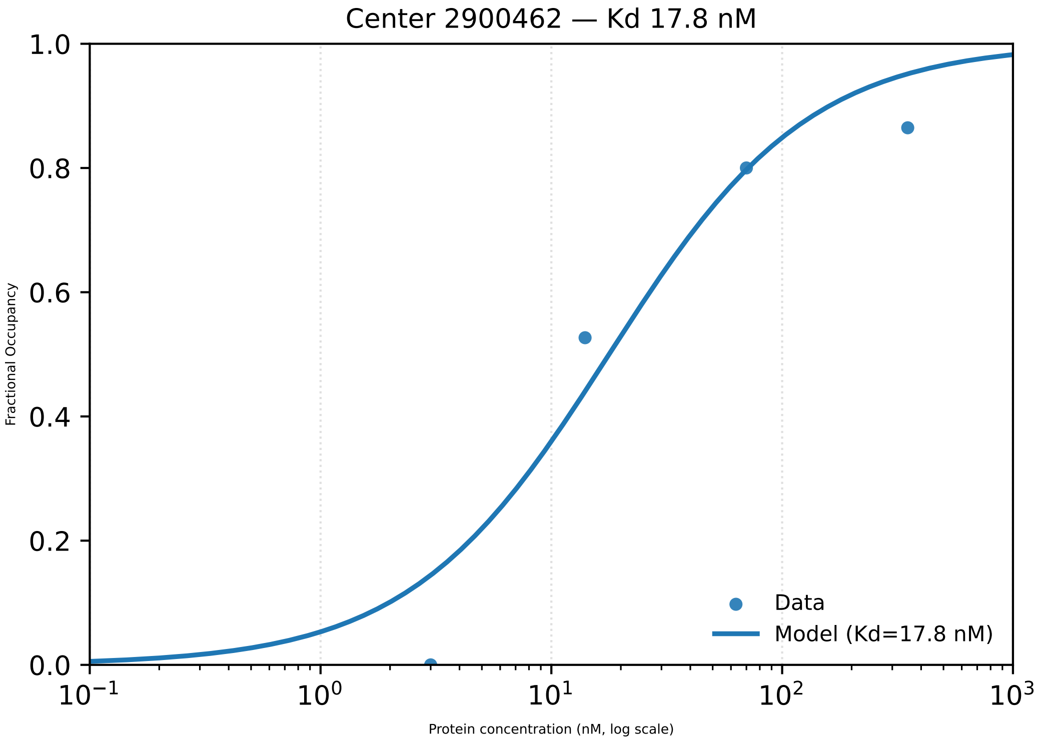

Center 2900462 — Kd 17.8 nM
Fractional Occupancy
Protein concentration (nM, log scale)
Data
Model (Kd=17.8 nM)
0.0
0.2
0.4
0.6
0.8
1.0
$10^{-1}$
$10^{0}$
$10^{1}$
$10^{2}$
$10^{3}$

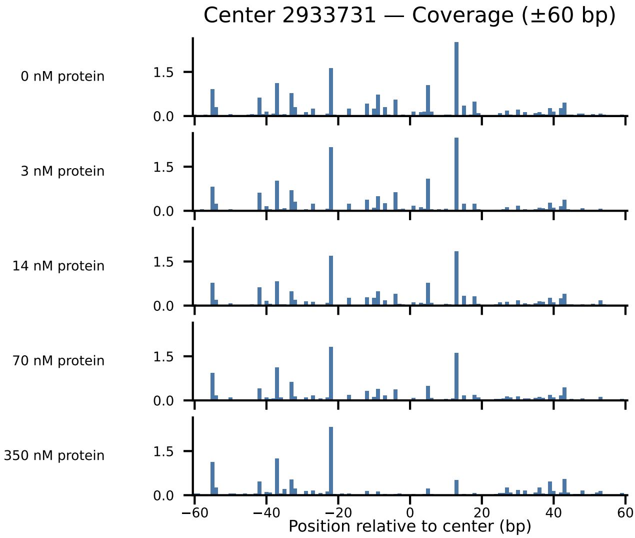
Center 2933731 — Coverage (±60 bp)
0 nM protein
3 nM protein
14 nM protein
70 nM protein
350 nM protein
1.5
0.0
−60
−40
−20
0
20
40
60
Position relative to center (bp)

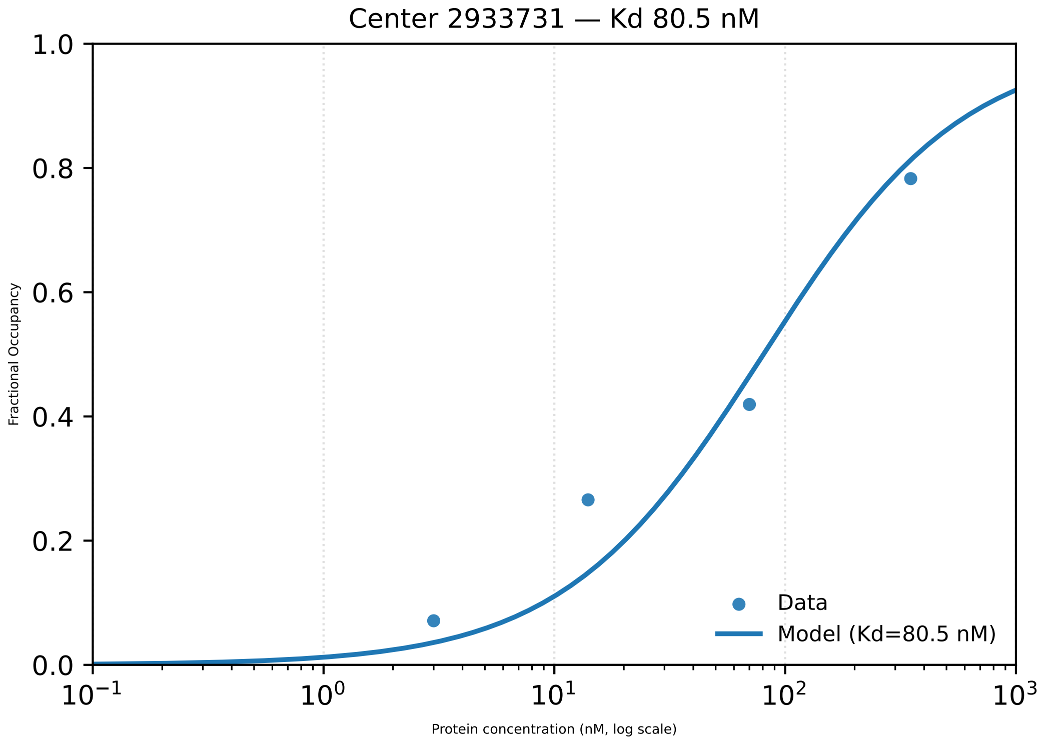

Center 2933731 — Kd 80.5 nM
Fractional Occupancy
1.0
0.8
0.6
0.4
0.2
0.0
$10^{-1}$
$10^{0}$
$10^{1}$
$10^{2}$
$10^{3}$
Protein concentration (nM, log scale)
Data
Model (Kd=80.5 nM)

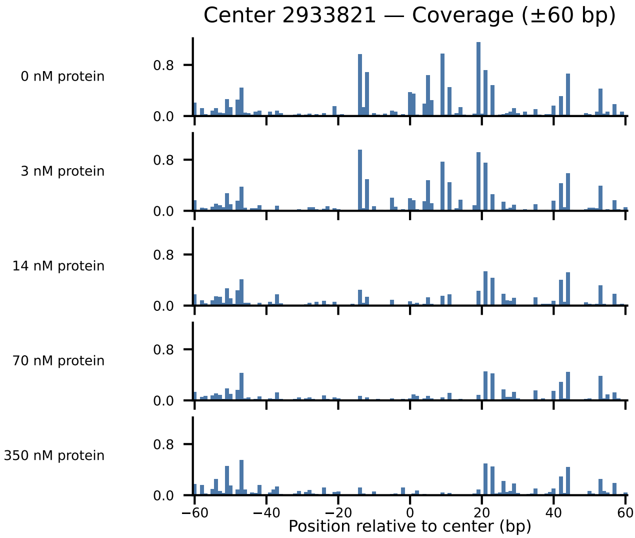
Center 2933821 — Coverage (±60 bp)
0 nM protein
3 nM protein
14 nM protein
70 nM protein
350 nM protein
0.8
0.0
−60
−40
−20
0
20
40
60
Position relative to center (bp)

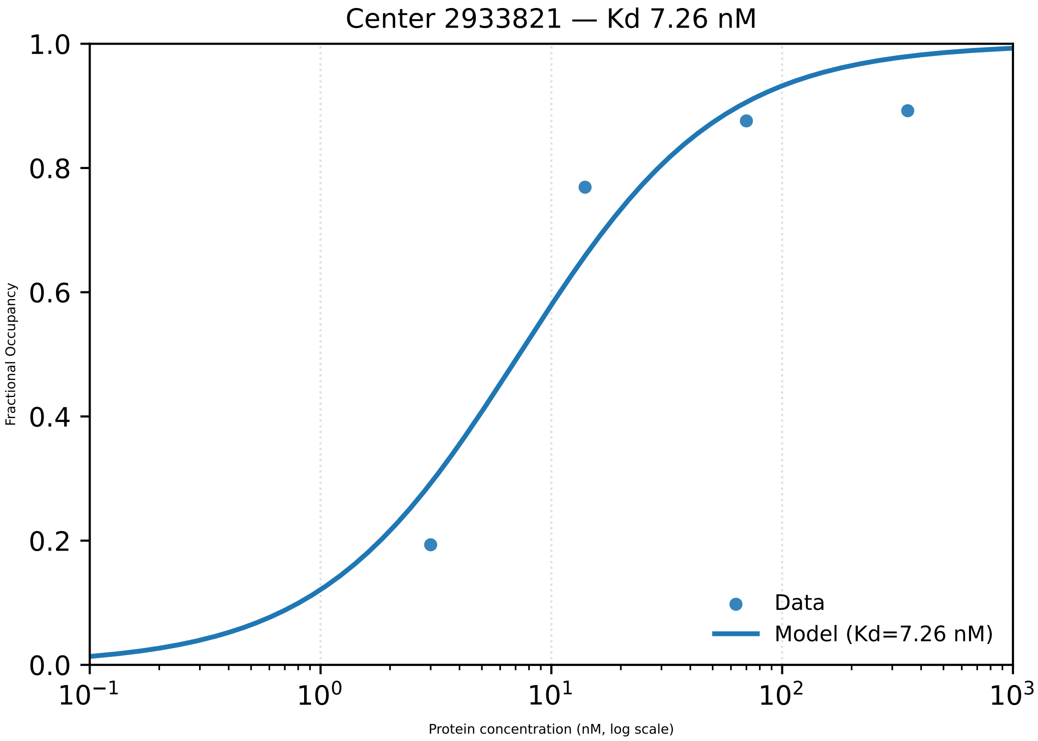

Center 2933821 — Kd 7.26 nM
Fractional Occupancy
1.0
0.8
0.6
0.4
0.2
0.0
$10^{-1}$
$10^{0}$
$10^{1}$
$10^{2}$
$10^{3}$
Protein concentration (nM, log scale)
Data
Model (Kd=7.26 nM)

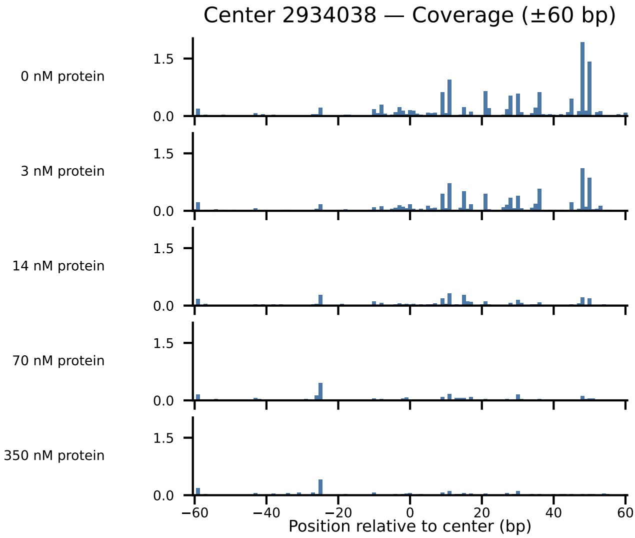
Center 2934038 — Coverage (±60 bp)
0 nM protein
3 nM protein
14 nM protein
70 nM protein
350 nM protein
1.5
0.0
−60
−40
−20
0
20
40
60
Position relative to center (bp)

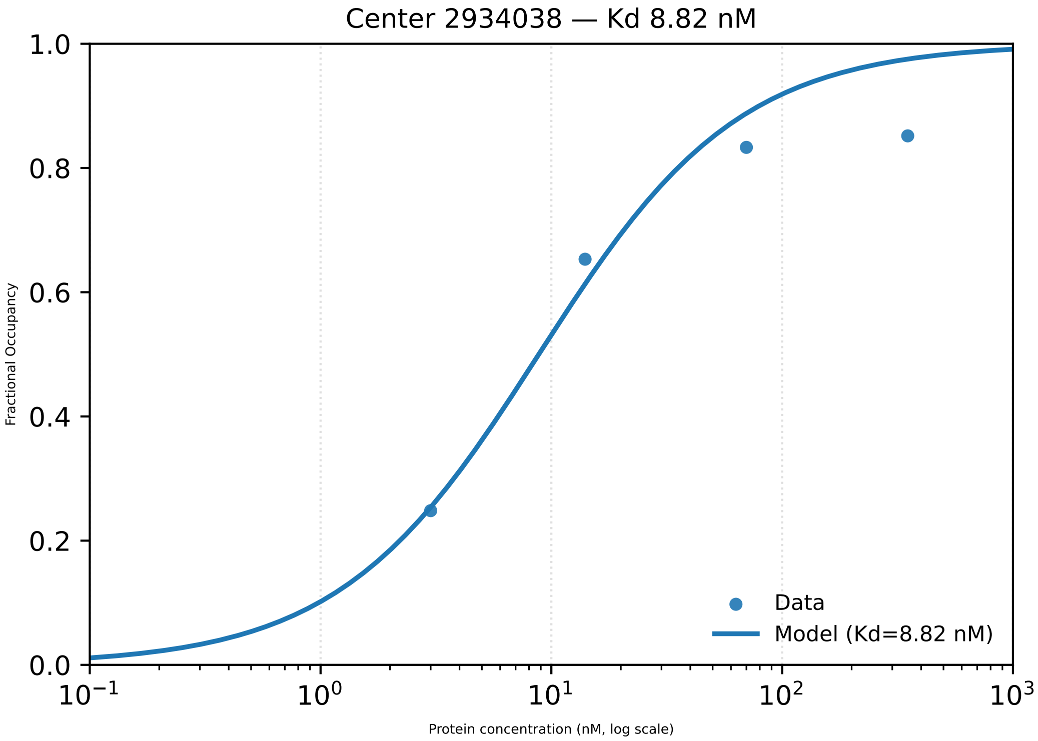

Center 2934038 — Kd 8.82 nM
1.0
0.8
0.6
0.4
0.2
0.0
Fractional Occupancy
$10^{-1}$
$10^{0}$
$10^{1}$
$10^{2}$
$10^{3}$
Protein concentration (nM, log scale)
Data
Model (Kd=8.82 nM)

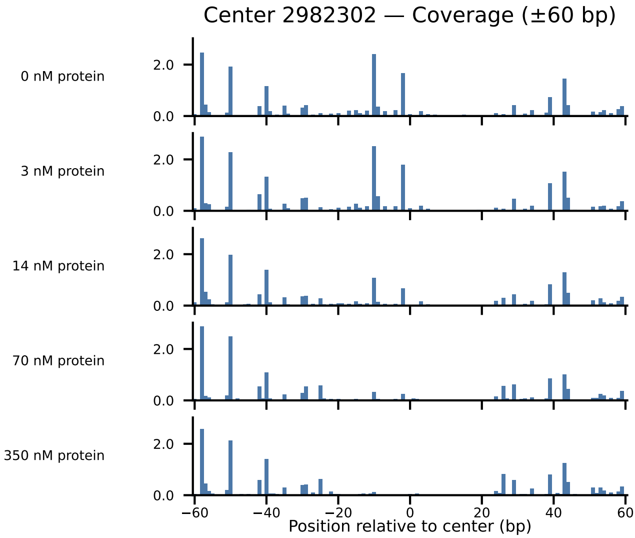
Center 2982302 — Coverage (±60 bp)
0 nM protein
3 nM protein
14 nM protein
70 nM protein
350 nM protein
2.0
0.0
−60
−40
−20
0
20
40
60
Position relative to center (bp)

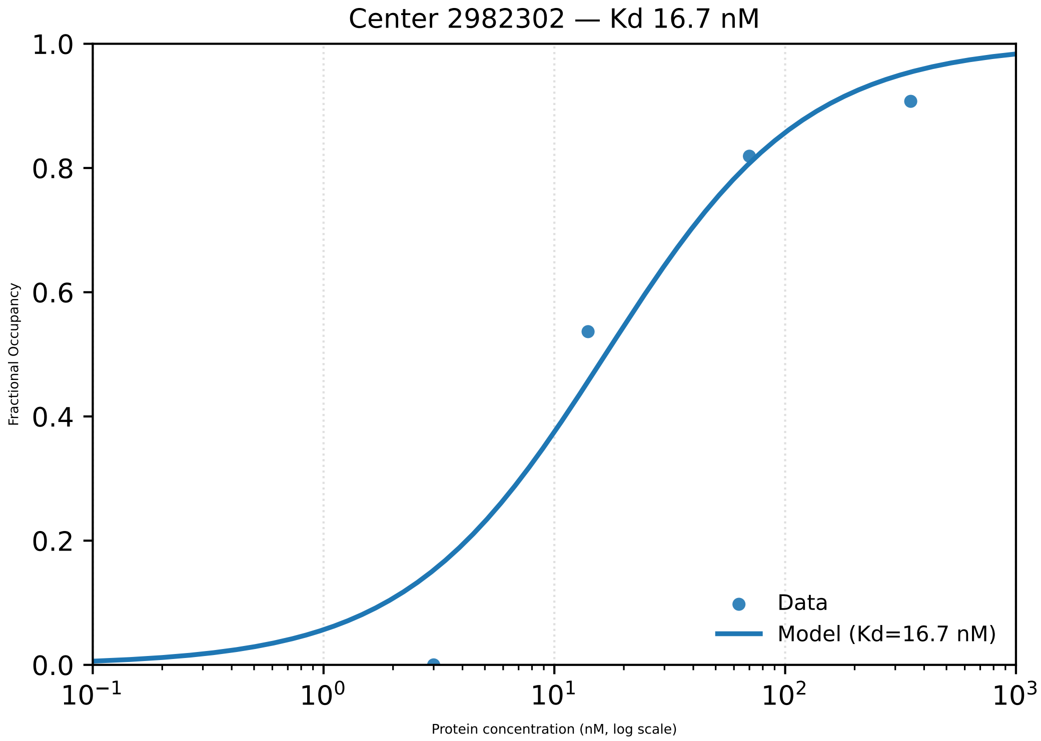

Center 2982302 — Kd 16.7 nM
Fractional Occupancy
1.0
0.8
0.6
0.4
0.2
0.0
$10^{-1}$
$10^{0}$
$10^{1}$
$10^{2}$
$10^{3}$
Protein concentration (nM, log scale)
Data
Model (Kd=16.7 nM)

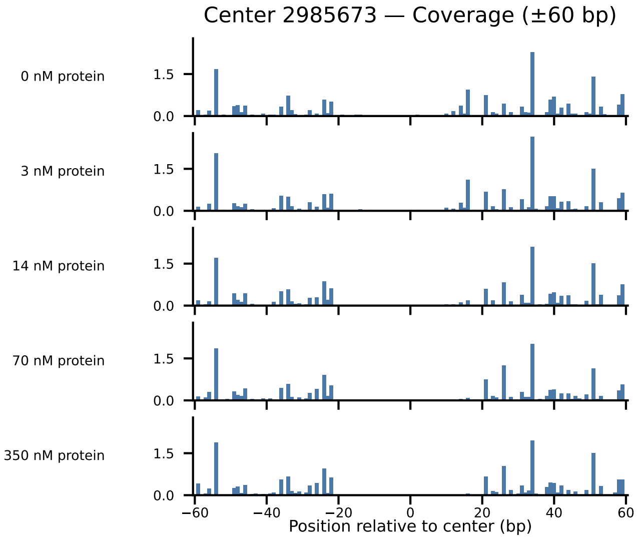
Center 2985673 — Coverage (±60 bp)
0 nM protein
3 nM protein
14 nM protein
70 nM protein
350 nM protein
1.5
0.0
−60
−40
−20
0
20
40
60
Position relative to center (bp)

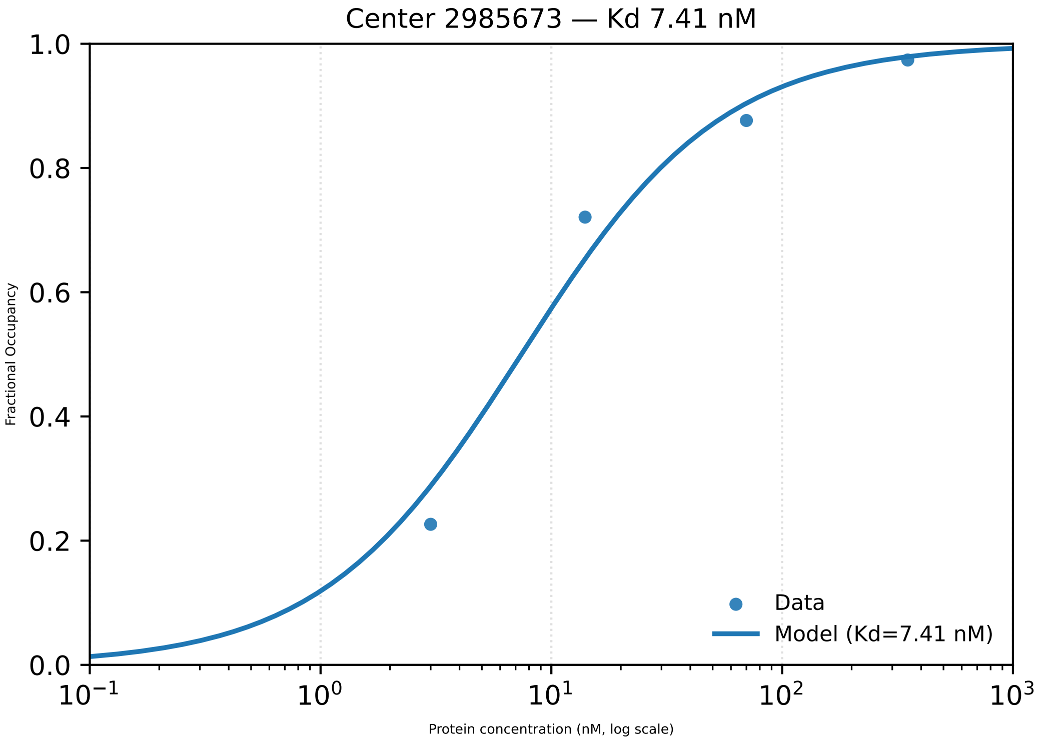

Center 2985673 — Kd 7.41 nM
1.0
0.8
0.6
0.4
0.2
0.0
Fractional Occupancy
$10^{-1}$
$10^{0}$
$10^{1}$
$10^{2}$
$10^{3}$
Protein concentration (nM, log scale)
Data
Model (Kd=7.41 nM)

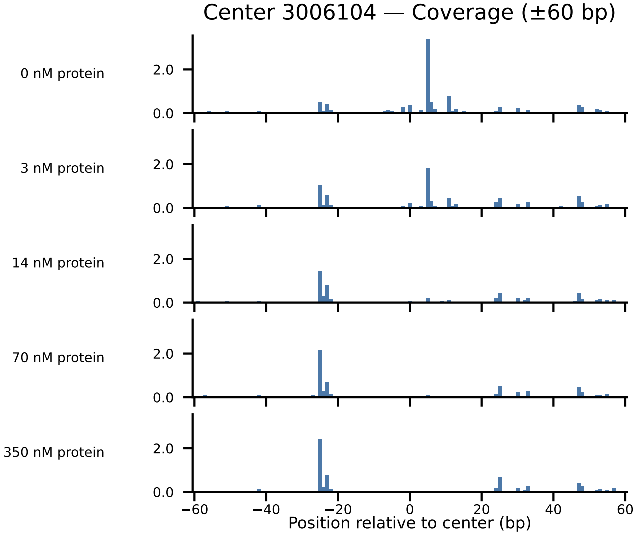

Center 3006104 — Coverage (±60 bp)
0 nM protein
3 nM protein
14 nM protein
70 nM protein
350 nM protein
2.0
0.0
−60
−40
−20
0
20
40
60
Position relative to center (bp)

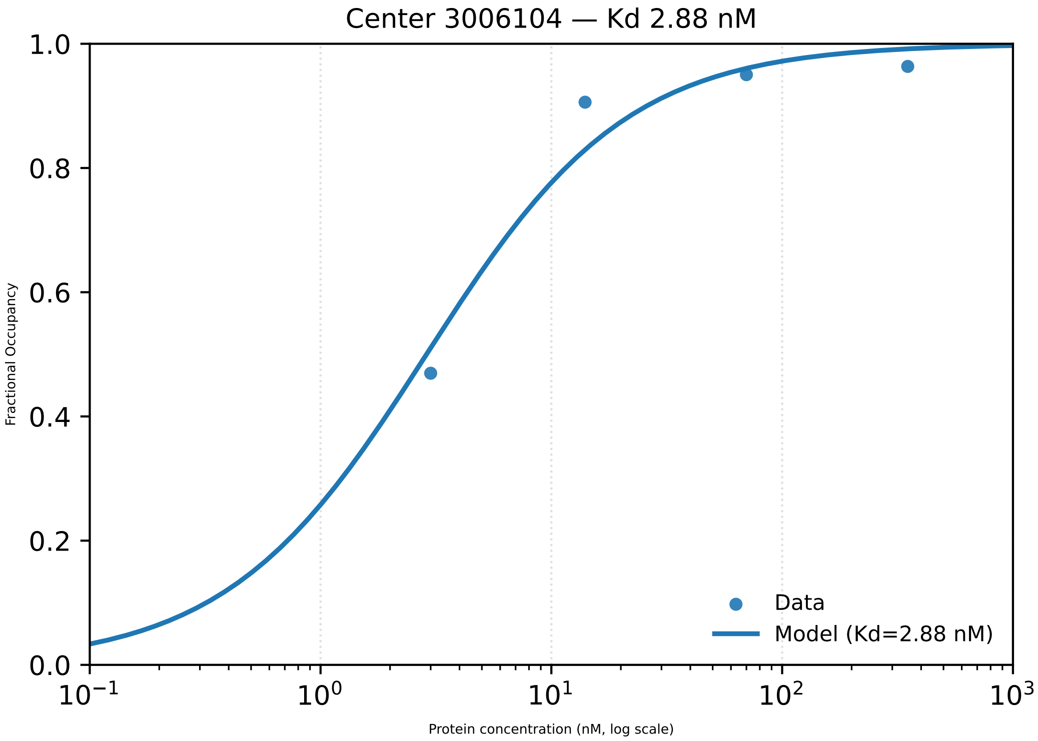

Center 3006104 — Kd 2.88 nM
1.0
0.8
0.6
0.4
0.2
0.0
Fractional Occupancy
$10^{-1}$
$10^{0}$
$10^{1}$
$10^{2}$
$10^{3}$
Protein concentration (nM, log scale)
Data
Model (Kd=2.88 nM)

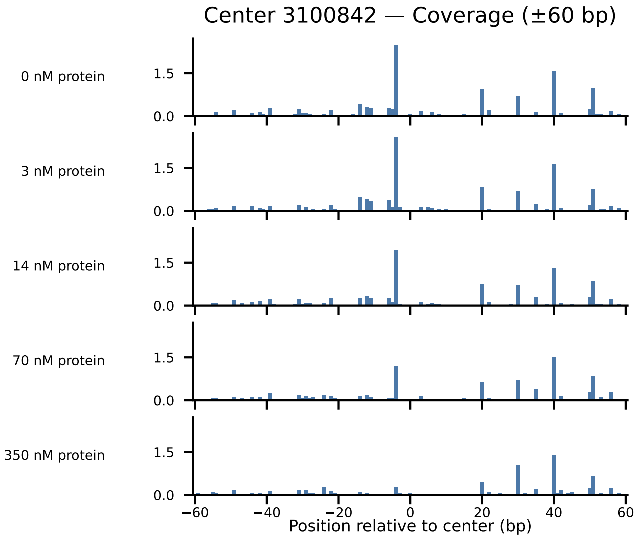

Center 3100842 — Coverage (±60 bp)
0 nM protein
3 nM protein
14 nM protein
70 nM protein
350 nM protein
1.5
0.0
−60
−40
−20
0
20
40
60
Position relative to center (bp)

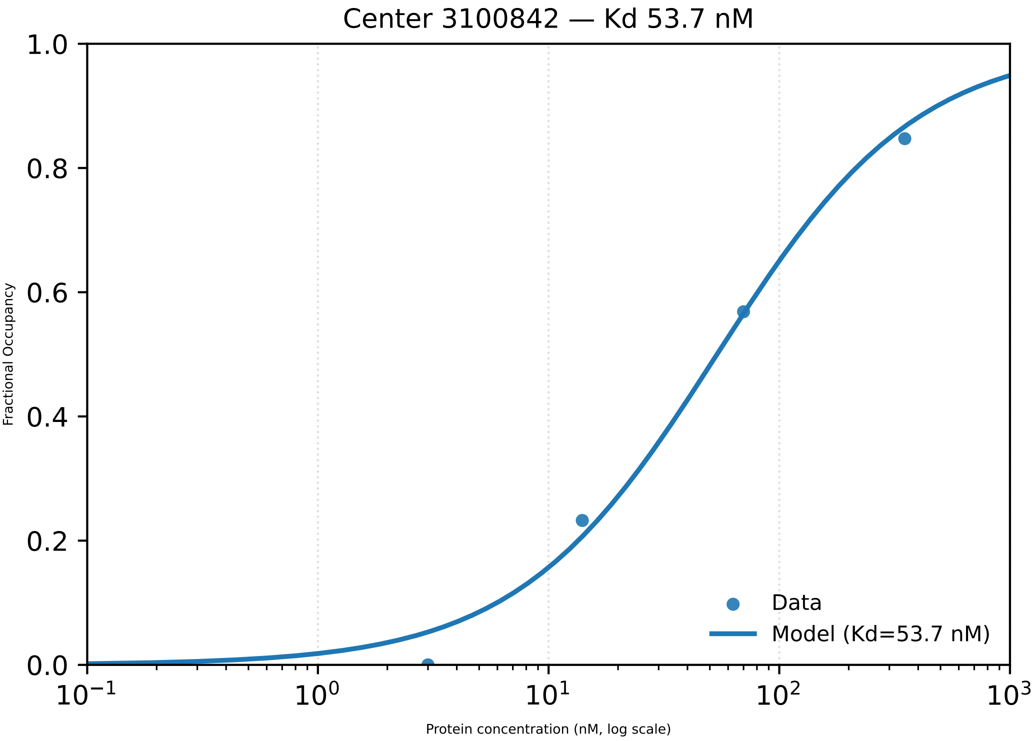

Center 3100842 — Kd 53.7 nM
1.0
0.8
0.6
0.4
0.2
0.0
Fractional Occupancy
Data
Model (Kd=53.7 nM)
$10^{-1}$
$10^{0}$
$10^{1}$
$10^{2}$
$10^{3}$
Protein concentration (nM, log scale)

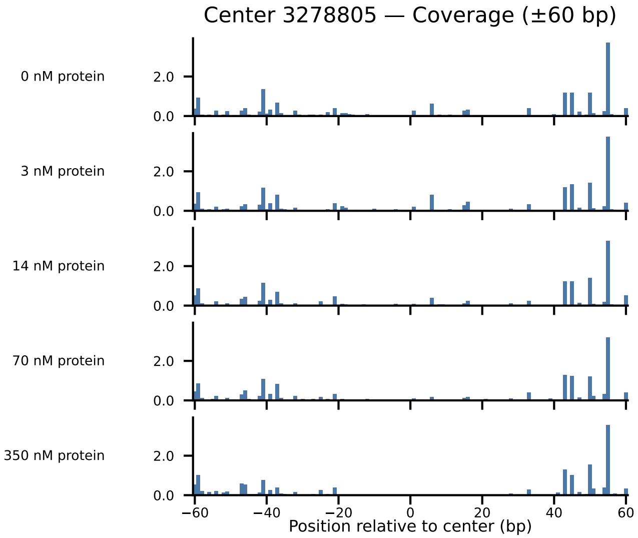

Center 3278805 — Coverage (±60 bp)
0 nM protein
3 nM protein
14 nM protein
70 nM protein
350 nM protein
2.0
0.0
−60
−40
−20
0
20
40
60
Position relative to center (bp)

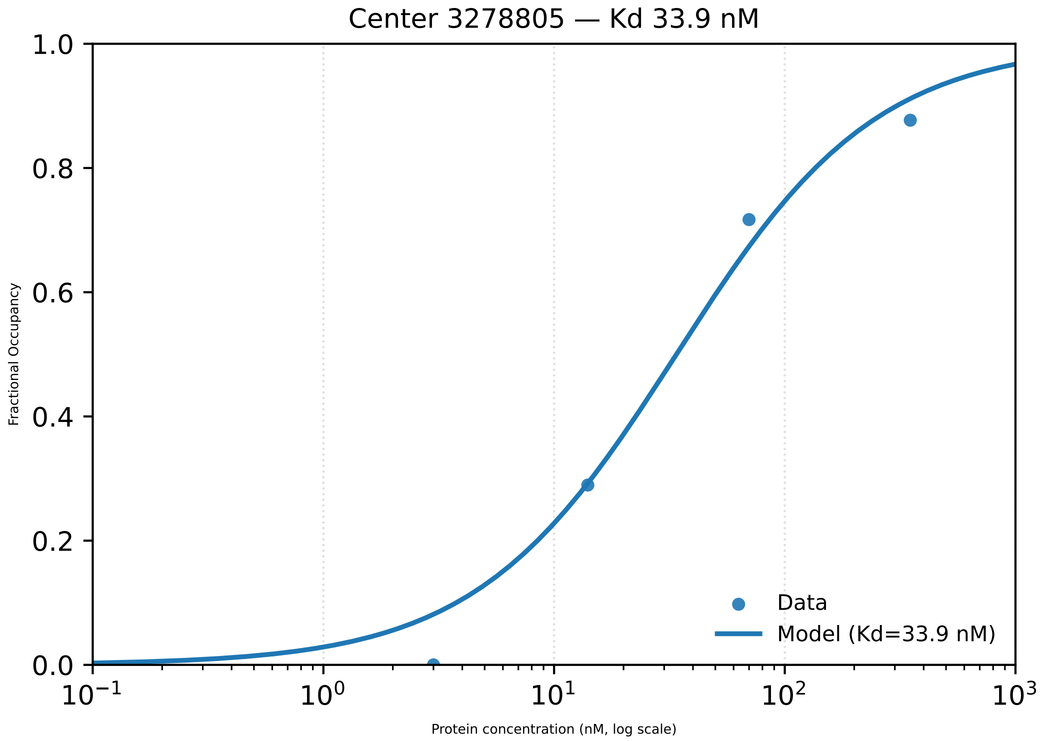

Center 3278805 — Kd 33.9 nM
1.0
0.8
0.6
0.4
0.2
0.0
Fractional Occupancy
$10^{-1}$
$10^{0}$
$10^{1}$
$10^{2}$
$10^{3}$
Protein concentration (nM, log scale)
Data
Model (Kd=33.9 nM)

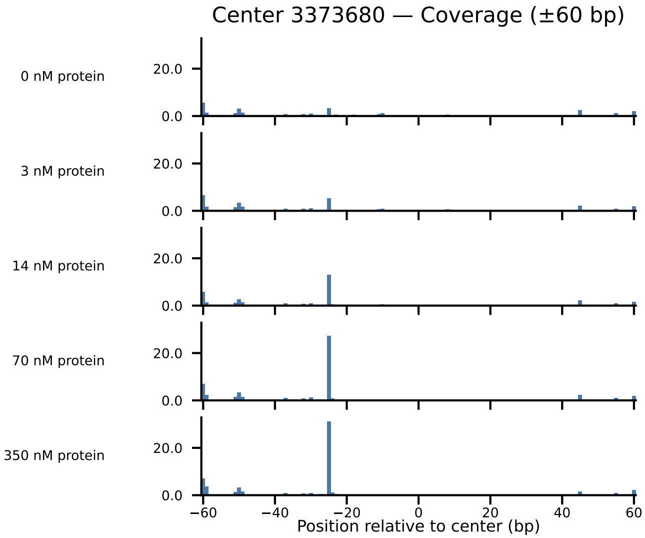

Center 3373680 — Coverage (±60 bp)
0 nM protein
3 nM protein
14 nM protein
70 nM protein
350 nM protein
20.0
0.0
−60
−40
−20
0
20
40
60
Position relative to center (bp)

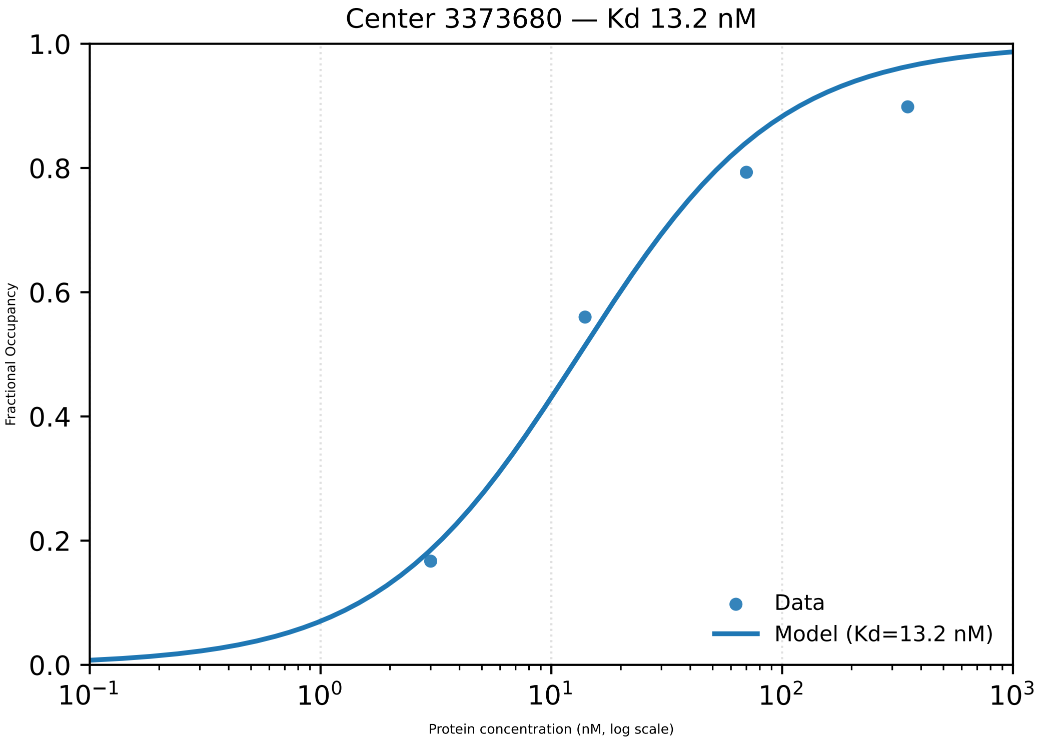

Center 3373680 — Kd 13.2 nM
1.0
0.8
0.6
0.4
0.2
0.0
Fractional Occupancy
Data
Model (Kd=13.2 nM)
$10^{-1}$
$10^{0}$
$10^{1}$
$10^{2}$
$10^{3}$
Protein concentration (nM, log scale)

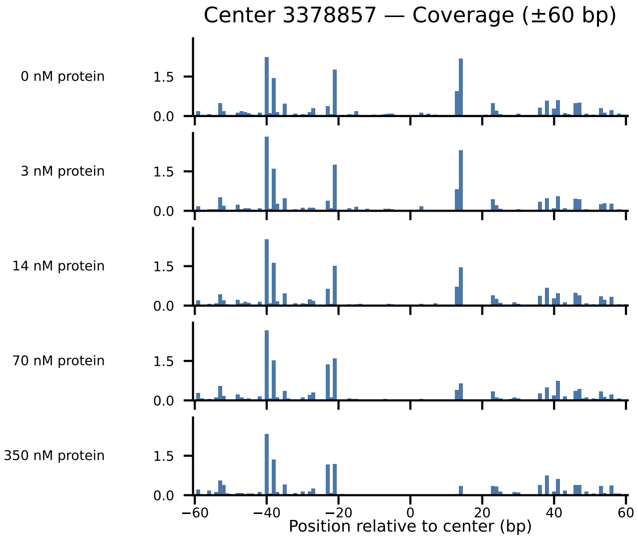

Center 3378857 — Coverage (±60 bp)
0 nM protein
3 nM protein
14 nM protein
70 nM protein
350 nM protein
1.5
0.0
−60
−40
−20
0
20
40
60
Position relative to center (bp)

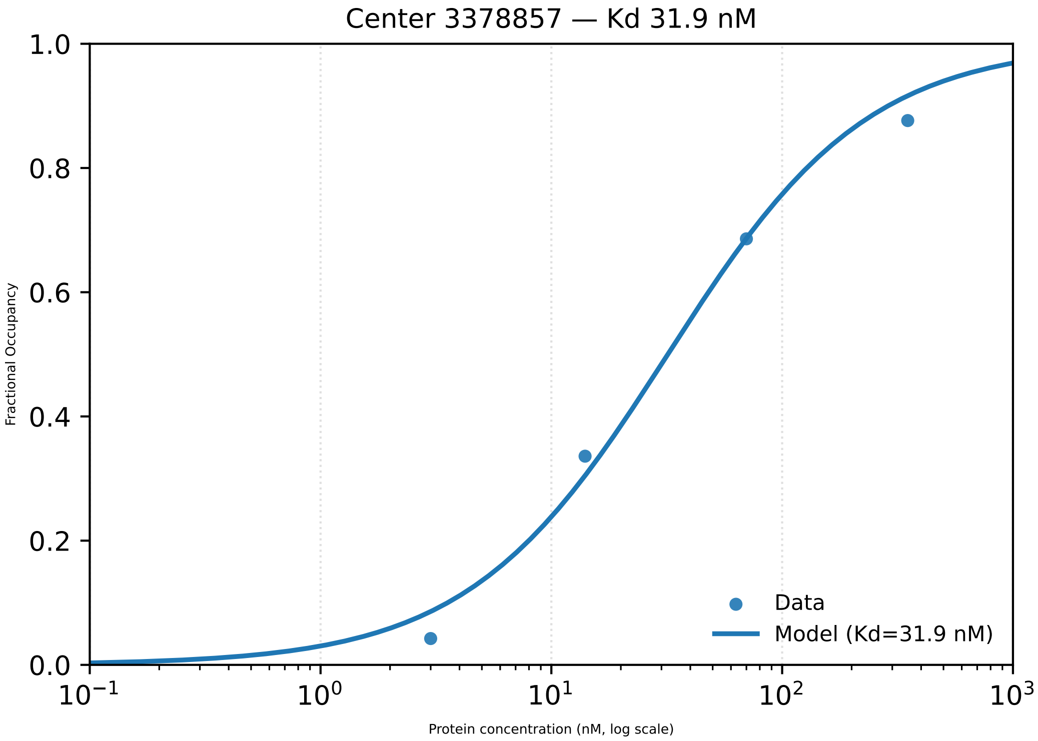

Center 3378857 — Kd 31.9 nM
Fractional Occupancy
Protein concentration (nM, log scale)
Data
Model (Kd=31.9 nM)
0.0
0.2
0.4
0.6
0.8
1.0
$10^{-1}$
$10^{0}$
$10^{1}$
$10^{2}$
$10^{3}$

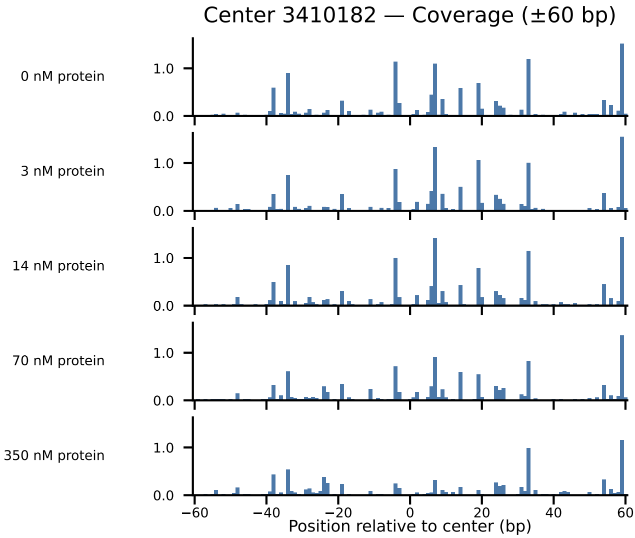
Center 3410182 — Coverage (±60 bp)
0 nM protein
3 nM protein
14 nM protein
70 nM protein
350 nM protein
1.0
0.0
−60
−40
−20
0
20
40
60
Position relative to center (bp)

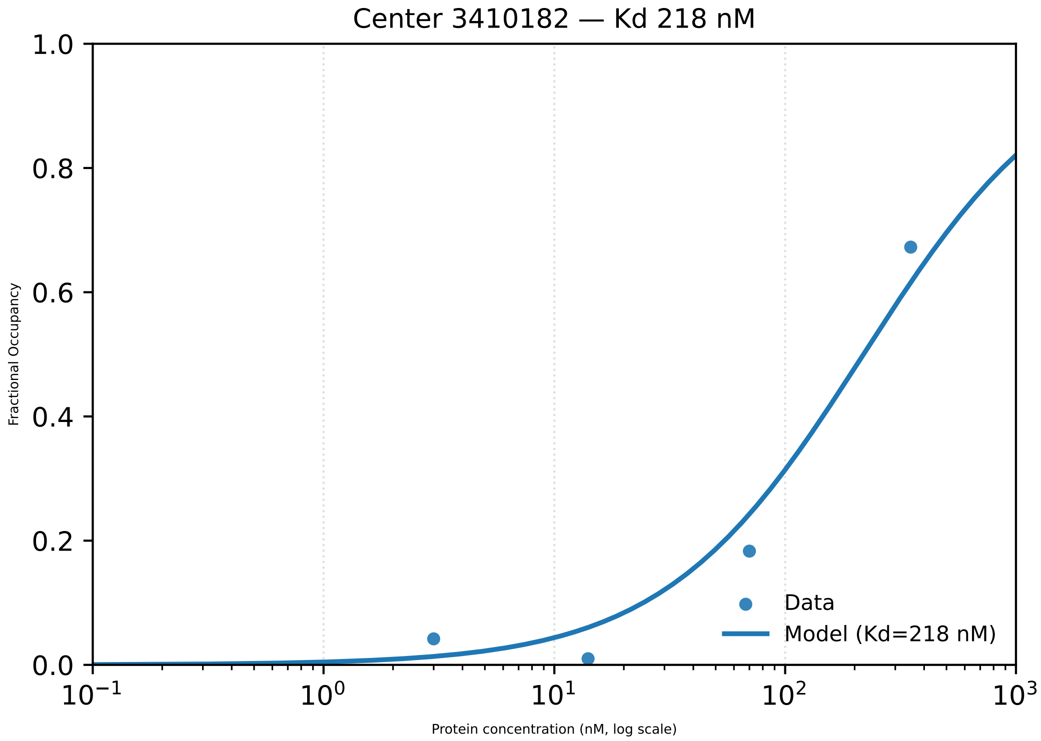
Center 3410182 — Kd 218 nM
Fractional Occupancy
Protein concentration (nM, log scale)
Data
Model (Kd=218 nM)
1.0
0.8
0.6
0.4
0.2
0.0
$10^{-1}$
$10^{0}$
$10^{1}$
$10^{2}$
$10^{3}$

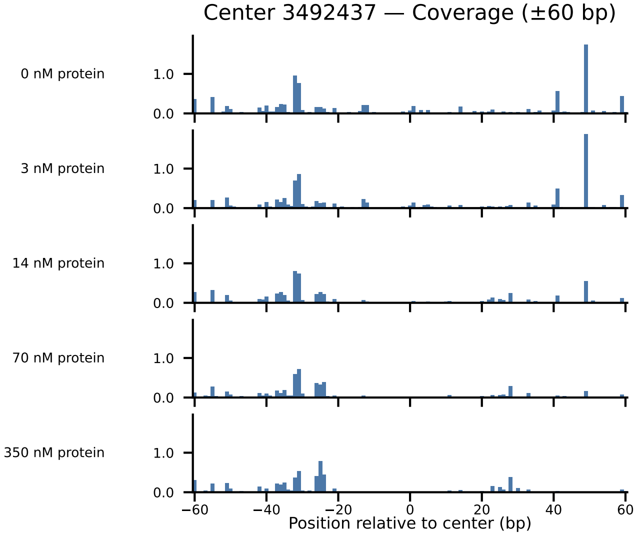
Center 3492437 — Coverage (±60 bp)
0 nM protein
3 nM protein
14 nM protein
70 nM protein
350 nM protein
1.0
0.0
−60
−40
−20
0
20
40
60
Position relative to center (bp)

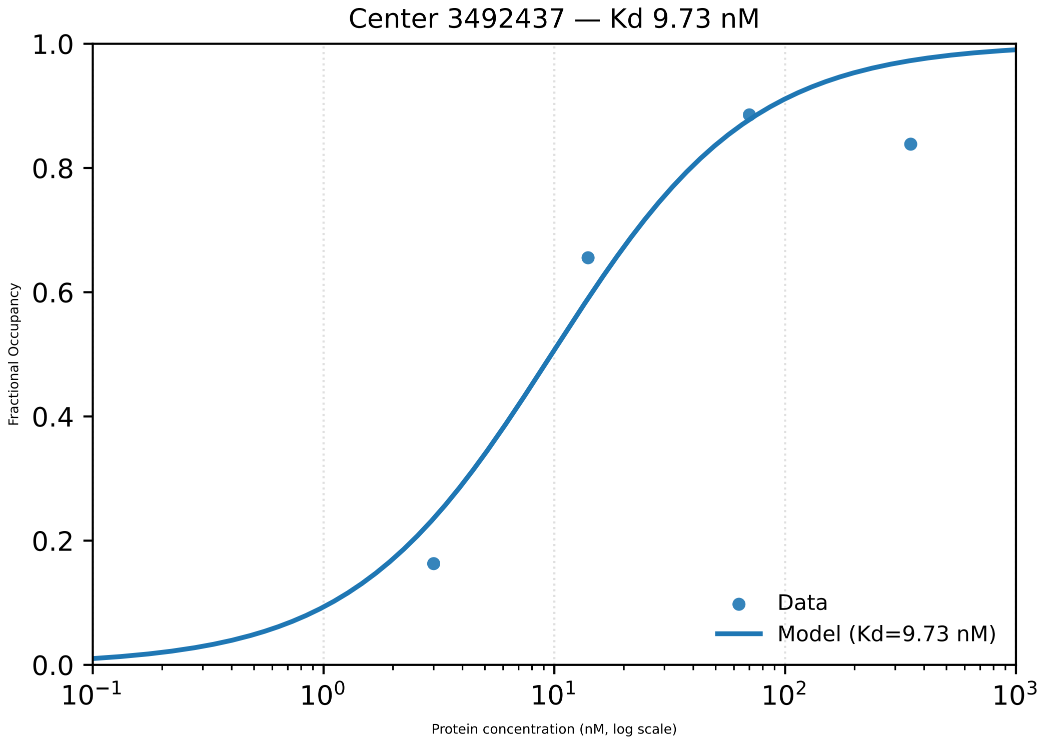

Center 3492437 — Kd 9.73 nM
1.0
0.8
0.6
0.4
0.2
0.0
Fractional Occupancy
$10^{-1}$
$10^{0}$
$10^{1}$
$10^{2}$
$10^{3}$
Protein concentration (nM, log scale)
Data
Model (Kd=9.73 nM)

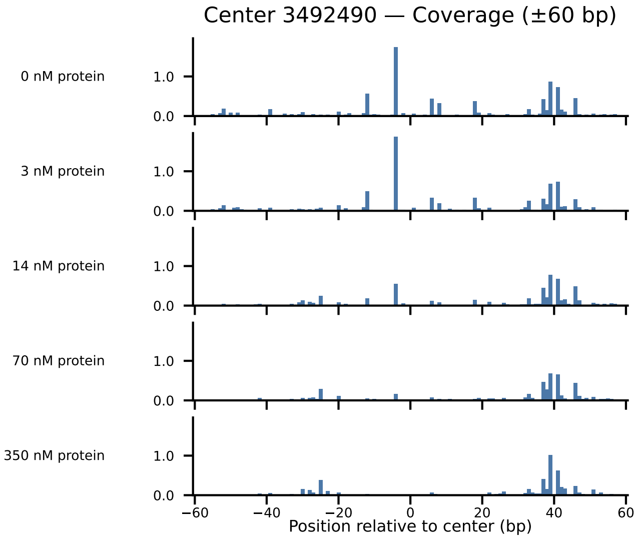

Center 3492490 — Coverage (±60 bp)
0 nM protein
3 nM protein
14 nM protein
70 nM protein
350 nM protein
1.0
0.0
−60
−40
−20
0
20
40
60
Position relative to center (bp)

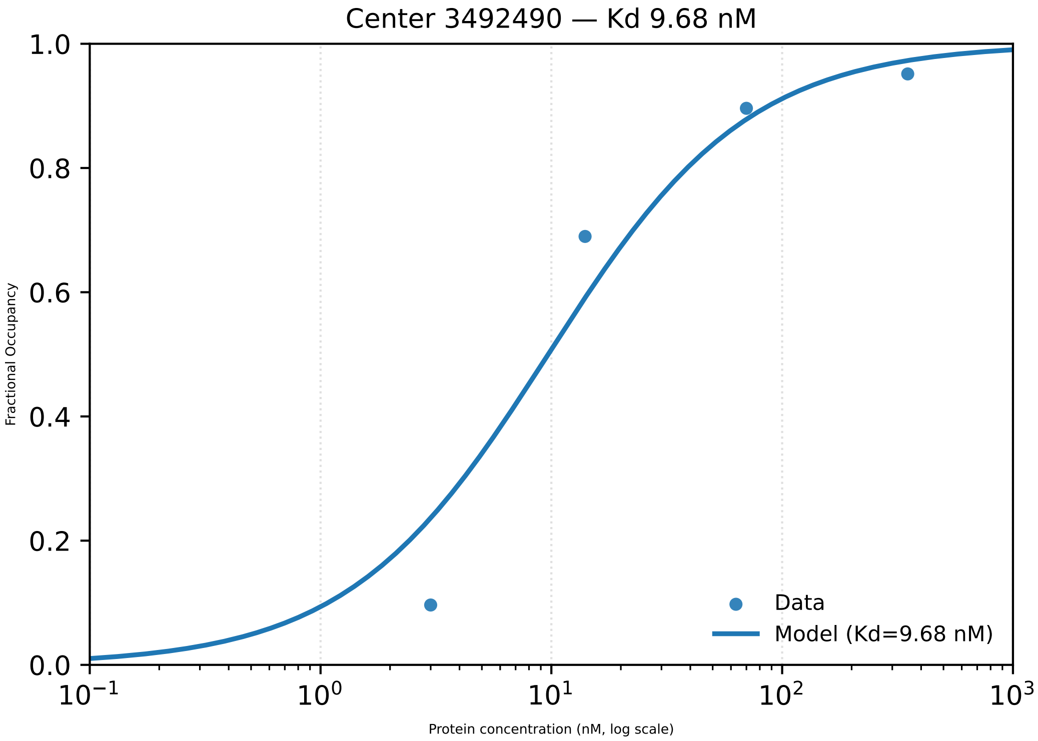

Center 3492490 — Kd 9.68 nM
1.0
0.8
0.6
0.4
0.2
0.0
Fractional Occupancy
$10^{-1}$
$10^{0}$
$10^{1}$
$10^{2}$
$10^{3}$
Protein concentration (nM, log scale)
Data
Model (Kd=9.68 nM)

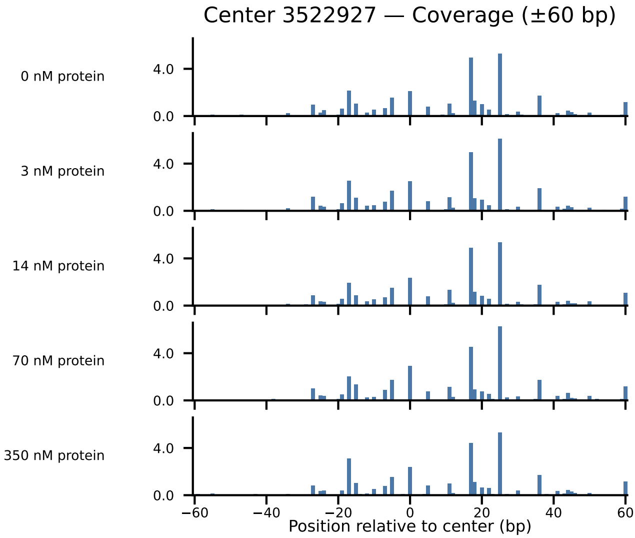
Center 3522927 — Coverage (±60 bp)
0 nM protein
3 nM protein
14 nM protein
70 nM protein
350 nM protein
4.0
0.0
−60
−40
−20
0
20
40
60
Position relative to center (bp)

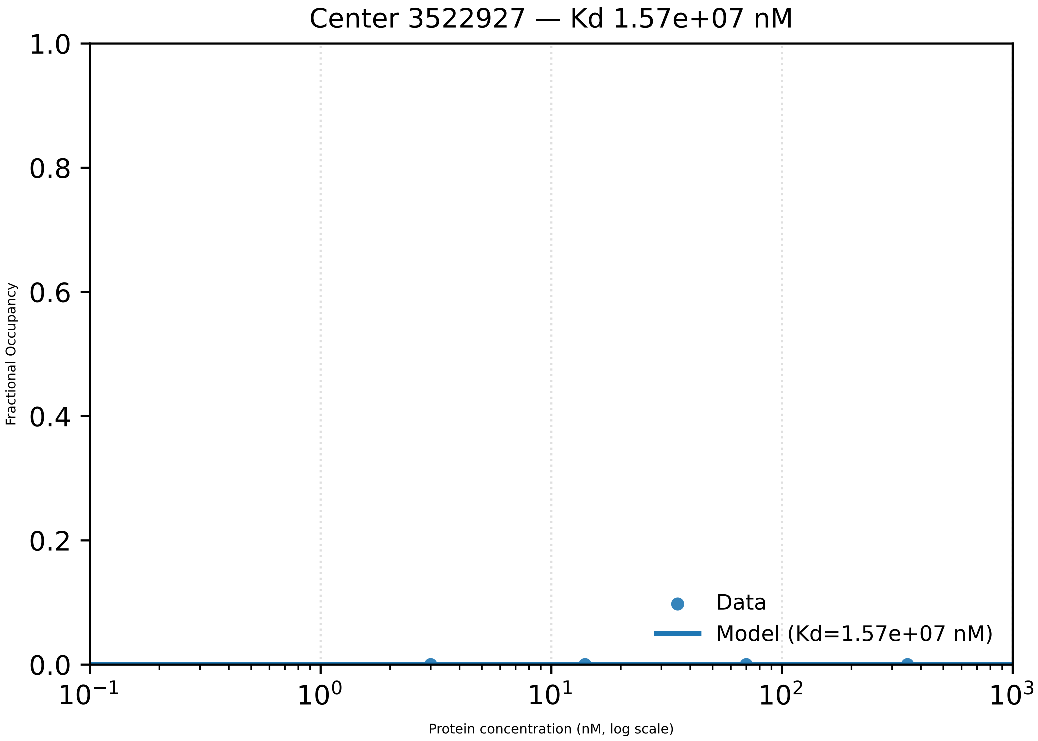

Center 3522927 — Kd 1.57e+07 nM
Fractional Occupancy
1.0
0.8
0.6
0.4
0.2
0.0
$10^{-1}$
$10^{0}$
$10^{1}$
$10^{2}$
$10^{3}$
Protein concentration (nM, log scale)
Data
Model (Kd=1.57e+07 nM)

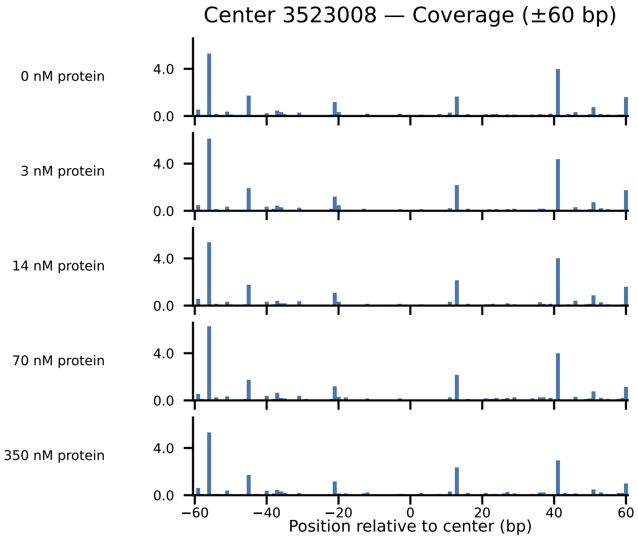

Center 3523008 — Coverage (±60 bp)
0 nM protein
3 nM protein
14 nM protein
70 nM protein
350 nM protein
4.0
0.0
−60
−40
−20
0
20
40
60
Position relative to center (bp)

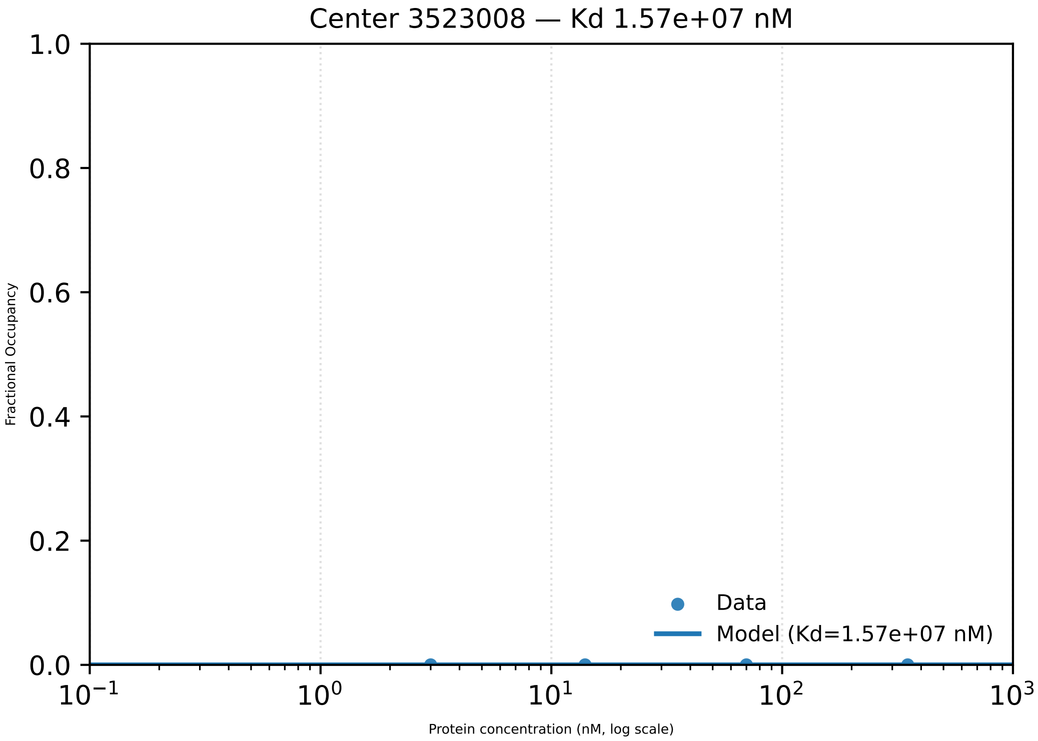

Center 3523008 — Kd 1.57e+07 nM
1.0
0.8
0.6
0.4
0.2
0.0
Fractional Occupancy
Data
Model (Kd=1.57e+07 nM)
$10^{-1}$
$10^{0}$
$10^{1}$
$10^{2}$
$10^{3}$
Protein concentration (nM, log scale)

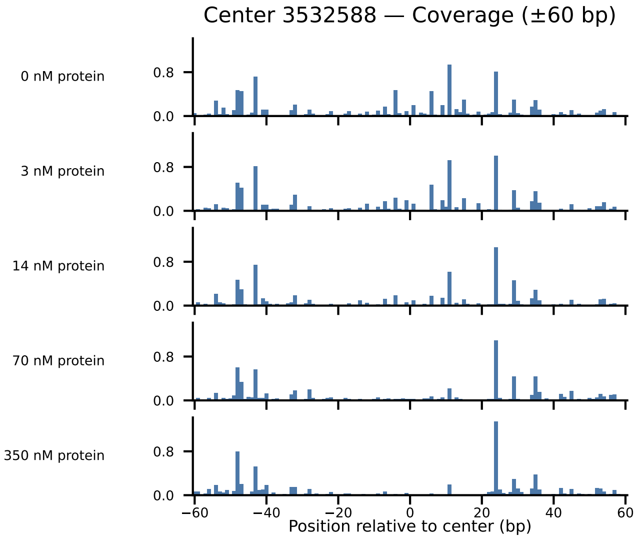

Center 3532588 — Coverage (±60 bp)
0 nM protein
3 nM protein
14 nM protein
70 nM protein
350 nM protein
0.0
0.8
−60
−40
−20
0
20
40
60
Position relative to center (bp)

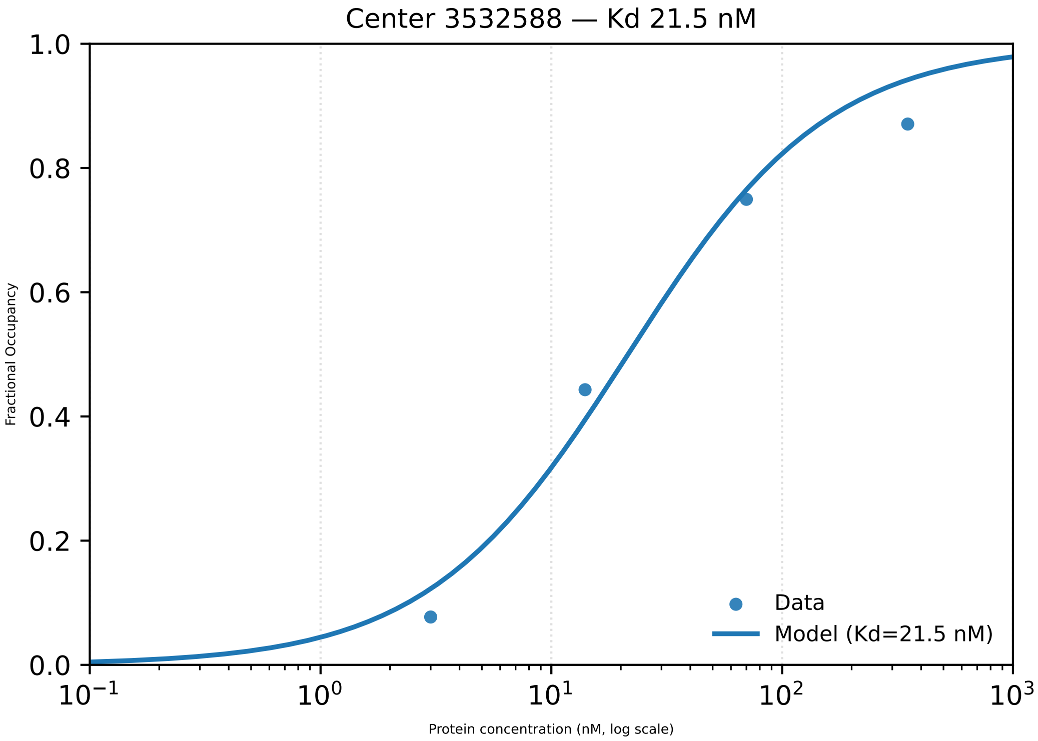

Center 3532588 — Kd 21.5 nM
1.0
0.8
0.6
0.4
0.2
0.0
Fractional Occupancy
$10^{-1}$
$10^{0}$
$10^{1}$
$10^{2}$
$10^{3}$
Protein concentration (nM, log scale)
Data
Model (Kd=21.5 nM)

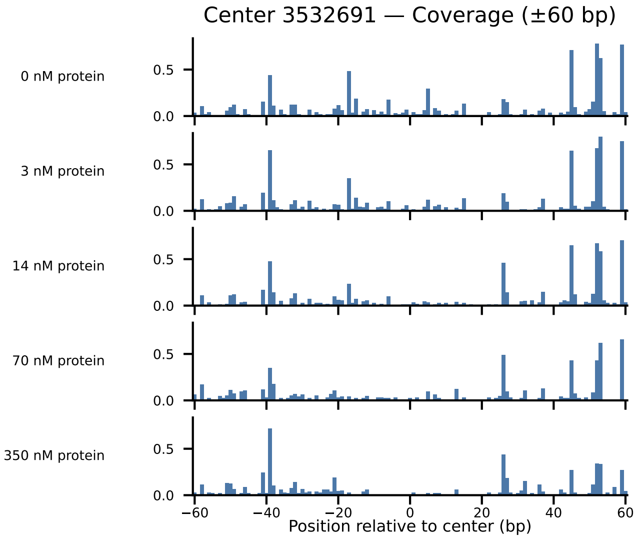

Center 3532691 — Coverage (±60 bp)
0 nM protein
3 nM protein
14 nM protein
70 nM protein
350 nM protein
0.5
0.0
−60
−40
−20
0
20
40
60
Position relative to center (bp)

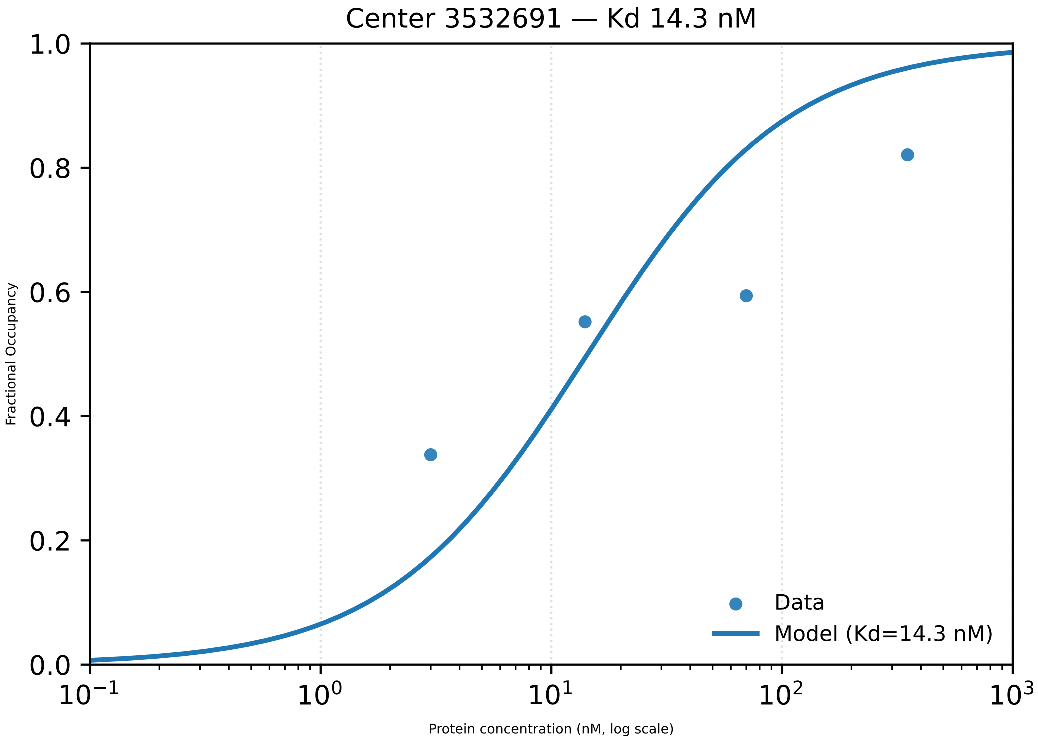

Center 3532691 — Kd 14.3 nM
Fractional Occupancy
1.0
0.8
0.6
0.4
0.2
0.0
$10^{-1}$
$10^{0}$
$10^{1}$
$10^{2}$
$10^{3}$
Protein concentration (nM, log scale)
Data
Model (Kd=14.3 nM)

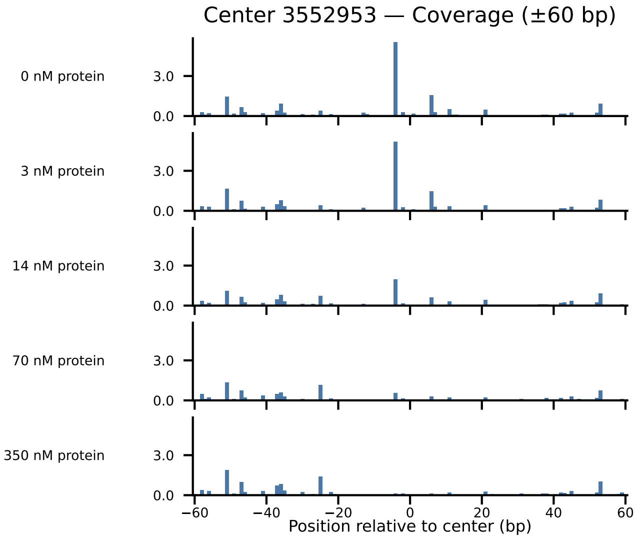

Center 3552953 — Coverage (±60 bp)
0 nM protein
3 nM protein
14 nM protein
70 nM protein
350 nM protein
3.0
0.0
−60
−40
−20
0
20
40
60
Position relative to center (bp)

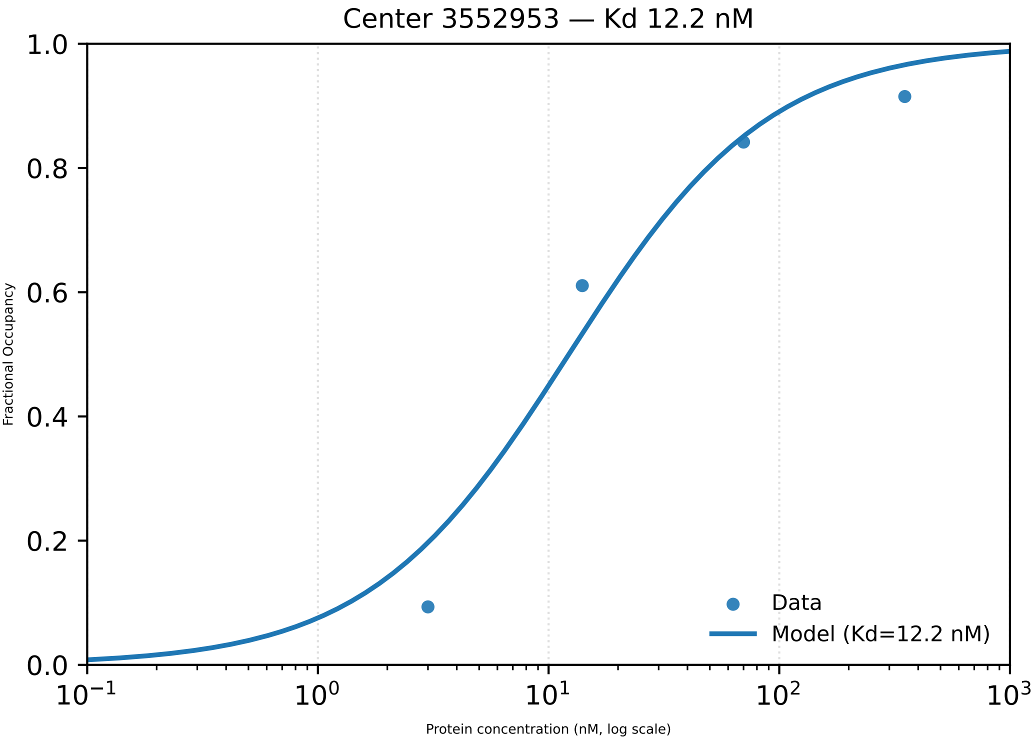

Center 3552953 — Kd 12.2 nM
Fractional Occupancy
1.0
0.8
0.6
0.4
0.2
0.0
$10^{-1}$
$10^{0}$
$10^{1}$
$10^{2}$
$10^{3}$
Protein concentration (nM, log scale)
Data
Model (Kd=12.2 nM)

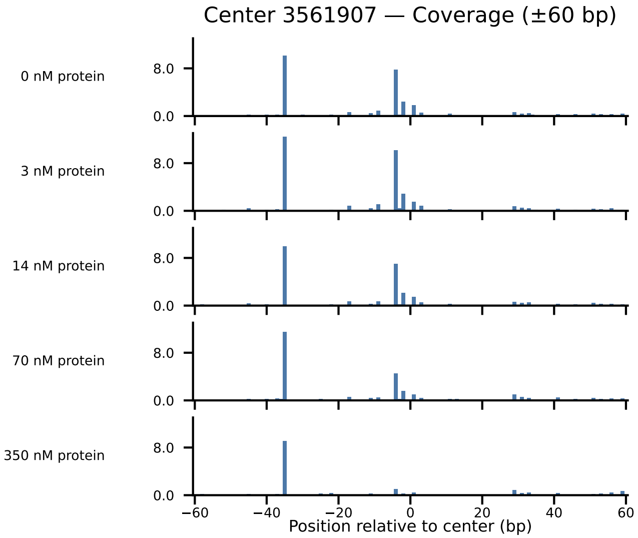

Center 3561907 — Coverage (±60 bp)
0 nM protein
3 nM protein
14 nM protein
70 nM protein
350 nM protein
8.0
0.0
−60
−40
−20
0
20
40
60
Position relative to center (bp)

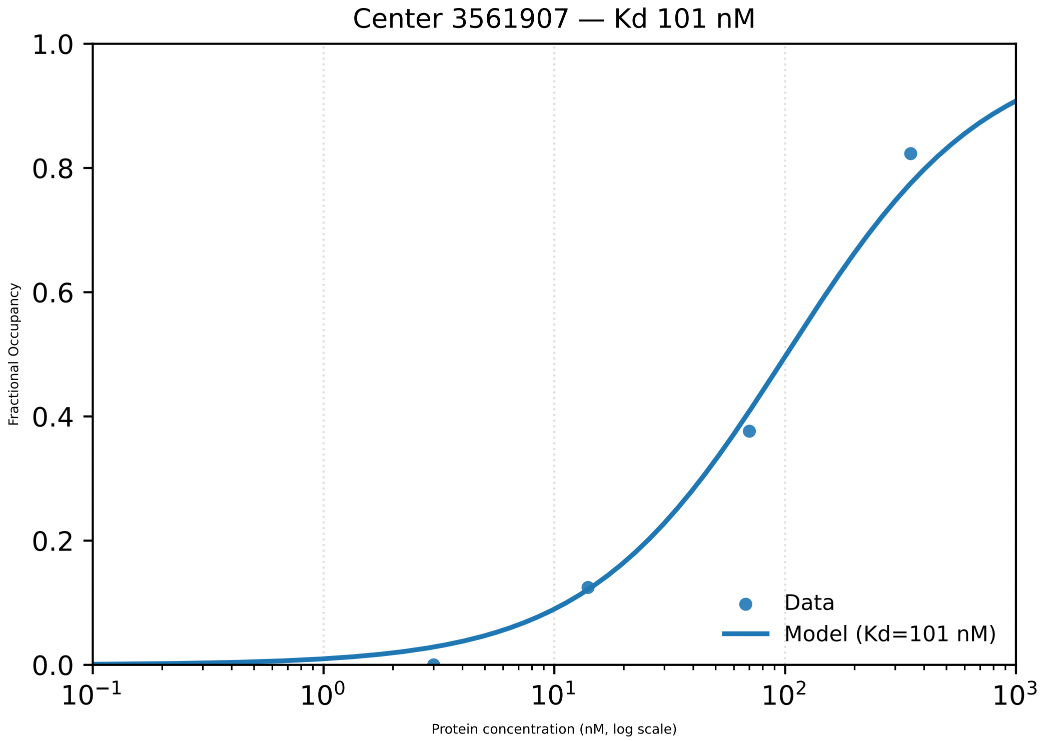

Center 3561907 — Kd 101 nM
1.0
0.8
0.6
0.4
0.2
0.0
Fractional Occupancy
$10^{-1}$
$10^{0}$
$10^{1}$
$10^{2}$
$10^{3}$
Protein concentration (nM, log scale)
Data
Model (Kd=101 nM)

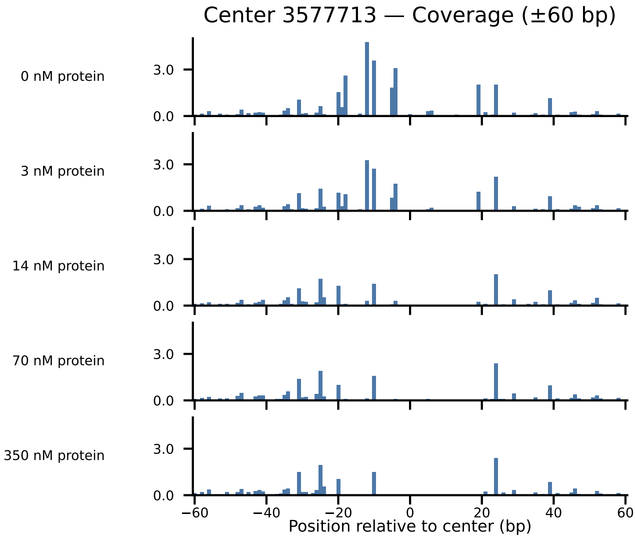

Center 3577713 — Coverage (±60 bp)
0 nM protein
3 nM protein
14 nM protein
70 nM protein
350 nM protein
3.0
0.0
−60
−40
−20
0
20
40
60
Position relative to center (bp)

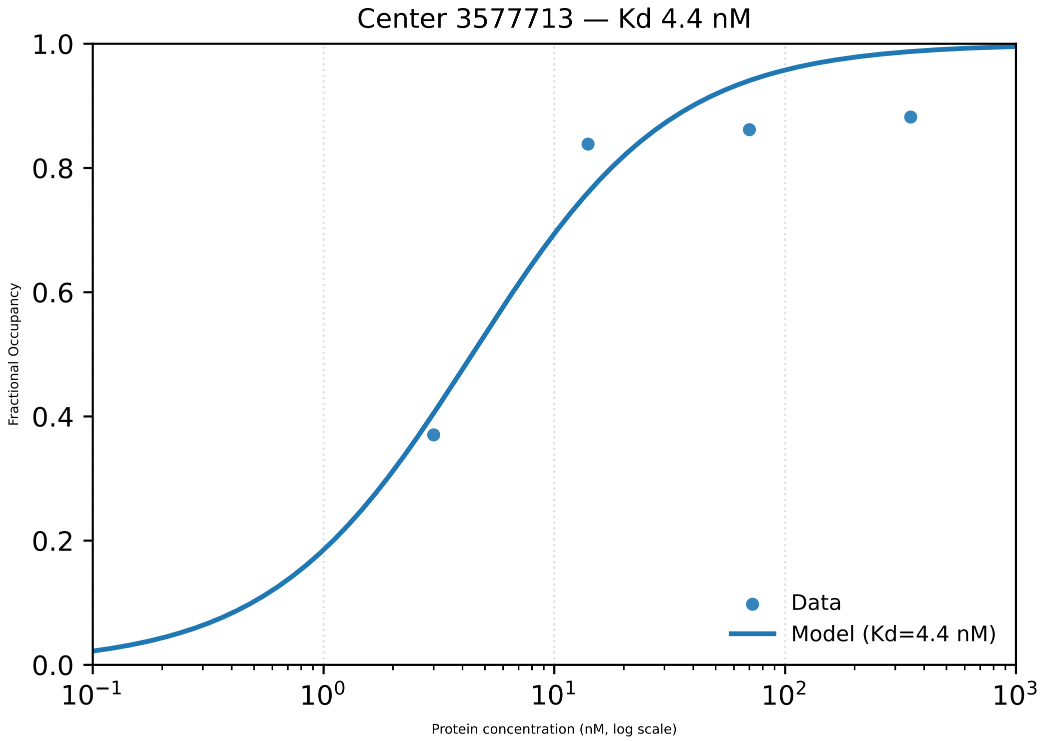

Center 3577713 — Kd 4.4 nM
1.0
0.8
0.6
0.4
0.2
0.0
Fractional Occupancy
$10^{-1}$
$10^{0}$
$10^{1}$
$10^{2}$
$10^{3}$
Protein concentration (nM, log scale)
Data
Model (Kd=4.4 nM)

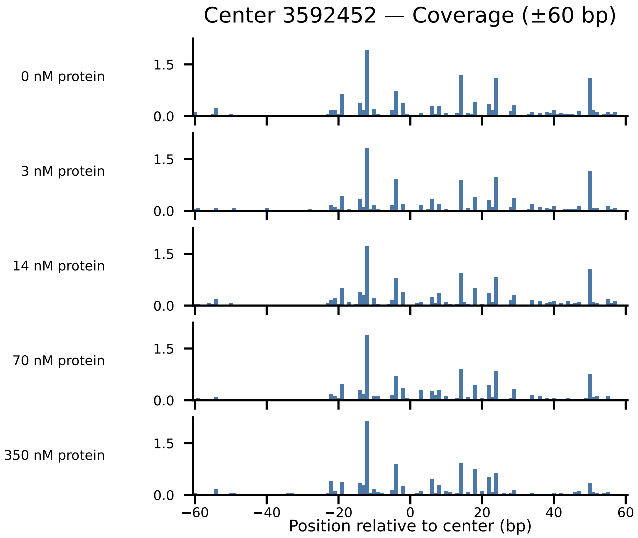
Center 3592452 — Coverage (±60 bp)
0 nM protein
3 nM protein
14 nM protein
70 nM protein
350 nM protein
1.5
0.0
−60
−40
−20
0
20
40
60
Position relative to center (bp)

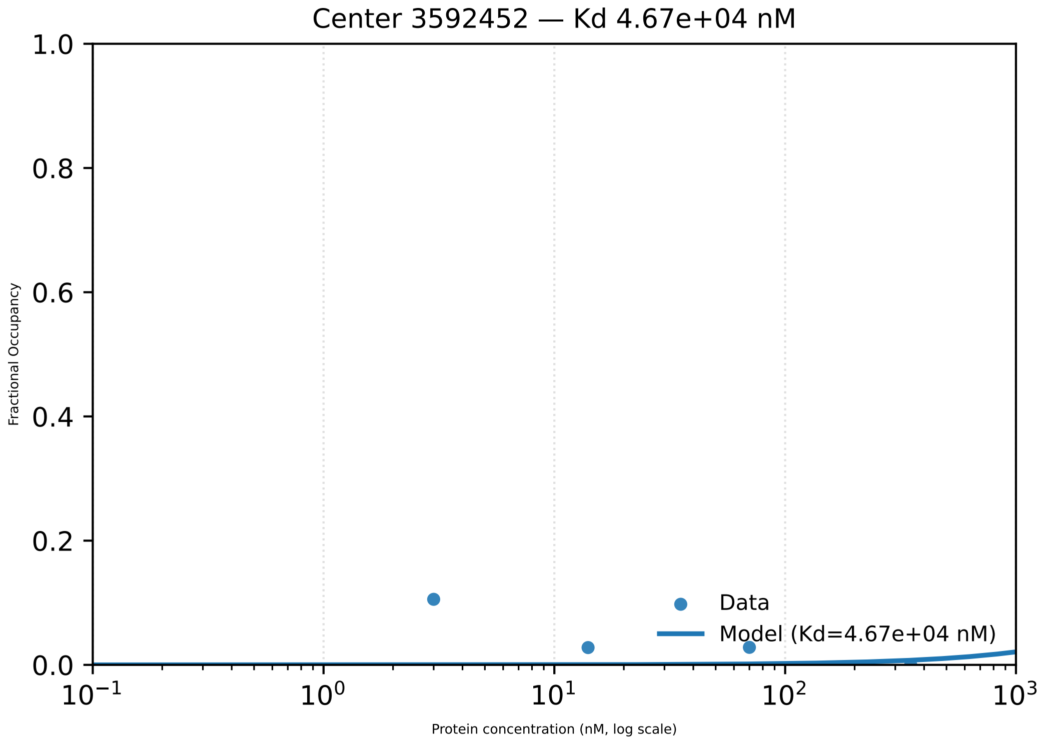

Center 3592452 — Kd 4.67e+04 nM
1.0
0.8
0.6
0.4
0.2
0.0
Fractional Occupancy
$10^{-1}$
$10^{0}$
$10^{1}$
$10^{2}$
$10^{3}$
Protein concentration (nM, log scale)
Data
Model (Kd=4.67e+04 nM)

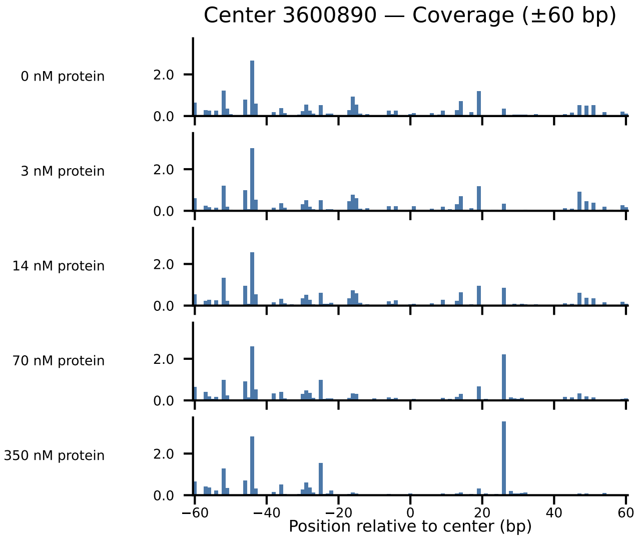

Center 3600890 — Coverage (±60 bp)
0 nM protein
3 nM protein
14 nM protein
70 nM protein
350 nM protein
2.0
0.0
−60
−40
−20
0
20
40
60
Position relative to center (bp)

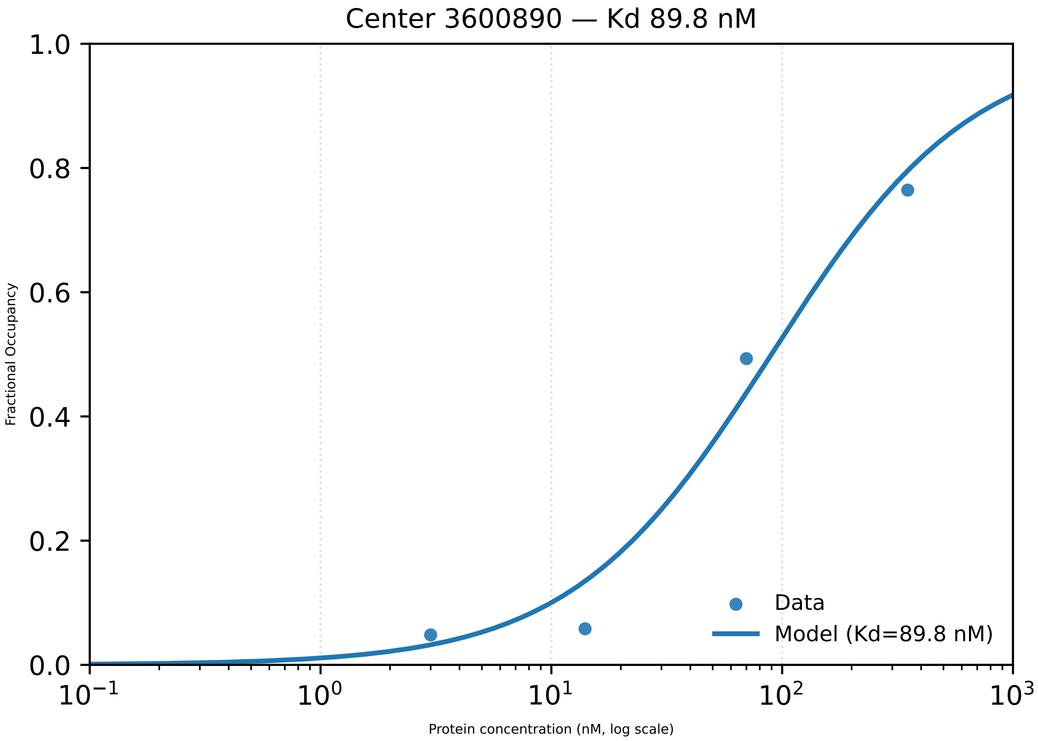

Center 3600890 — Kd 89.8 nM
Fractional Occupancy
0.0
0.2
0.4
0.6
0.8
1.0
$10^{-1}$
$10^{0}$
$10^{1}$
$10^{2}$
$10^{3}$
Protein concentration (nM, log scale)
Data
Model (Kd=89.8 nM)

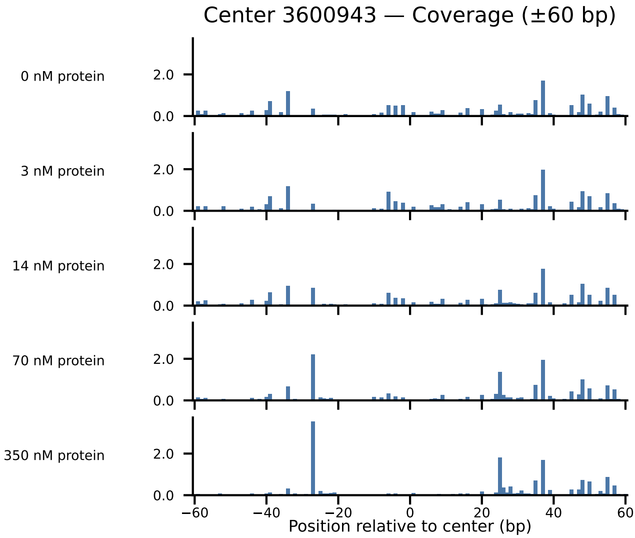

Center 3600943 — Coverage (±60 bp)
0 nM protein
3 nM protein
14 nM protein
70 nM protein
350 nM protein
2.0
0.0
−60
−40
−20
0
20
40
60
Position relative to center (bp)

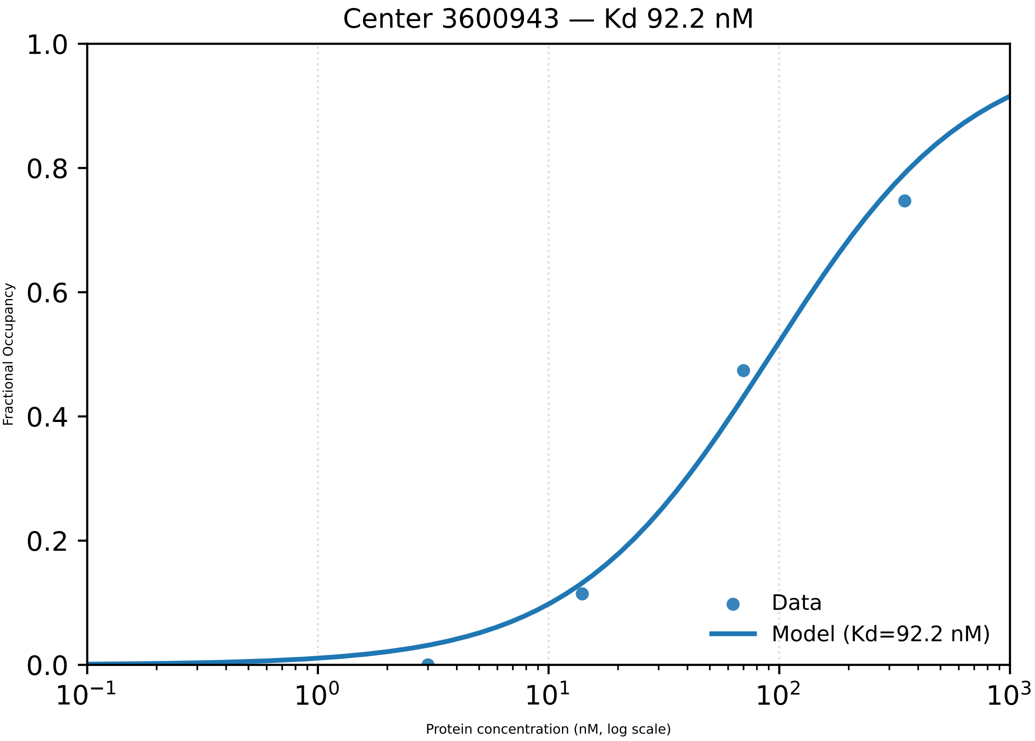

Center 3600943 — Kd 92.2 nM
Fractional Occupancy
1.0
0.8
0.6
0.4
0.2
0.0
$10^{-1}$
$10^{0}$
$10^{1}$
$10^{2}$
$10^{3}$
Protein concentration (nM, log scale)
Data
Model (Kd=92.2 nM)

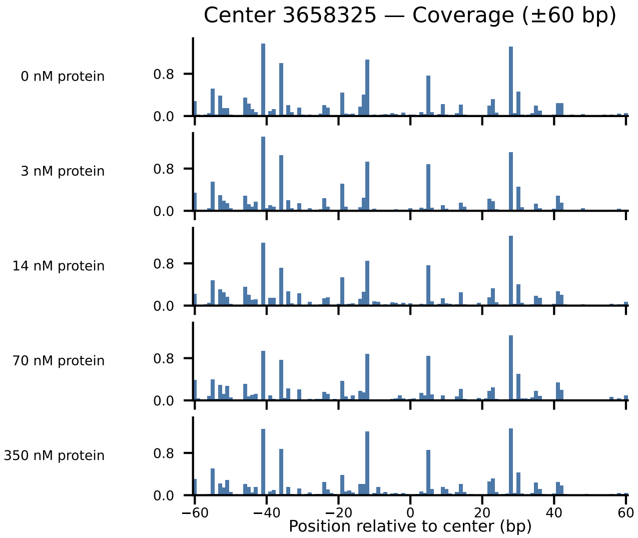

Center 3658325 — Coverage (±60 bp)
0 nM protein
3 nM protein
14 nM protein
70 nM protein
350 nM protein
0.8
0.0
−60
−40
−20
0
20
40
60
Position relative to center (bp)

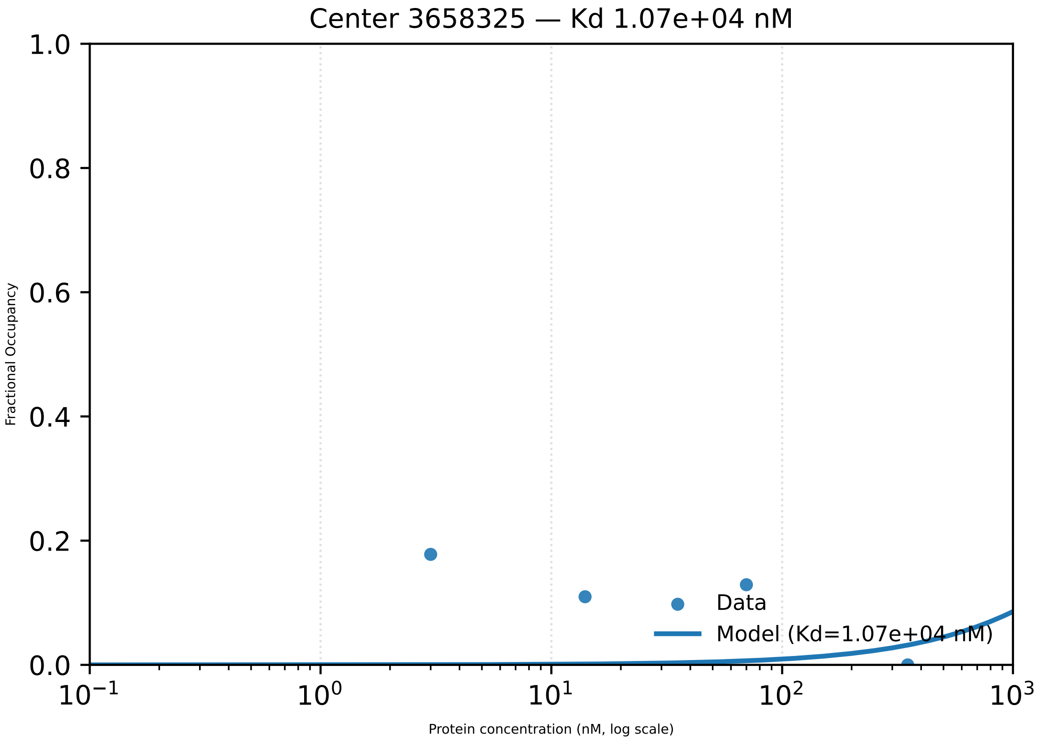

Center 3658325 — Kd 1.07e+04 nM
Fractional Occupancy
1.0
0.8
0.6
0.4
0.2
0.0
10⁻¹
10⁰
10¹
10²
10³
Protein concentration (nM, log scale)
Data
Model (Kd=1.07e+04 nM)

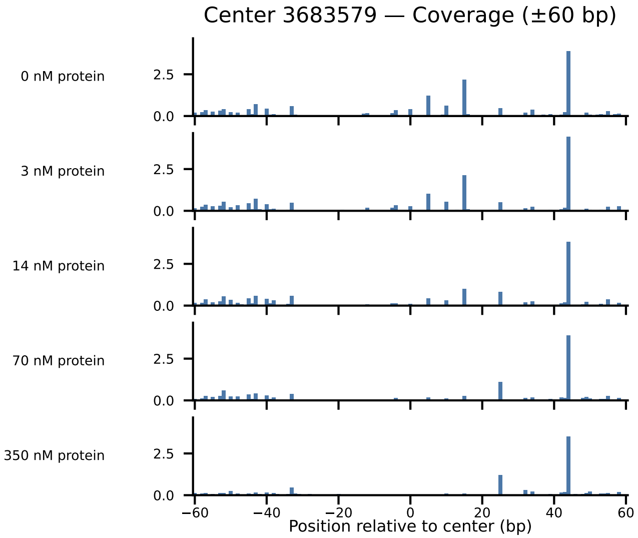

Center 3683579 — Coverage (±60 bp)
0 nM protein
3 nM protein
14 nM protein
70 nM protein
350 nM protein
2.5
0.0
−60
−40
−20
0
20
40
60
Position relative to center (bp)

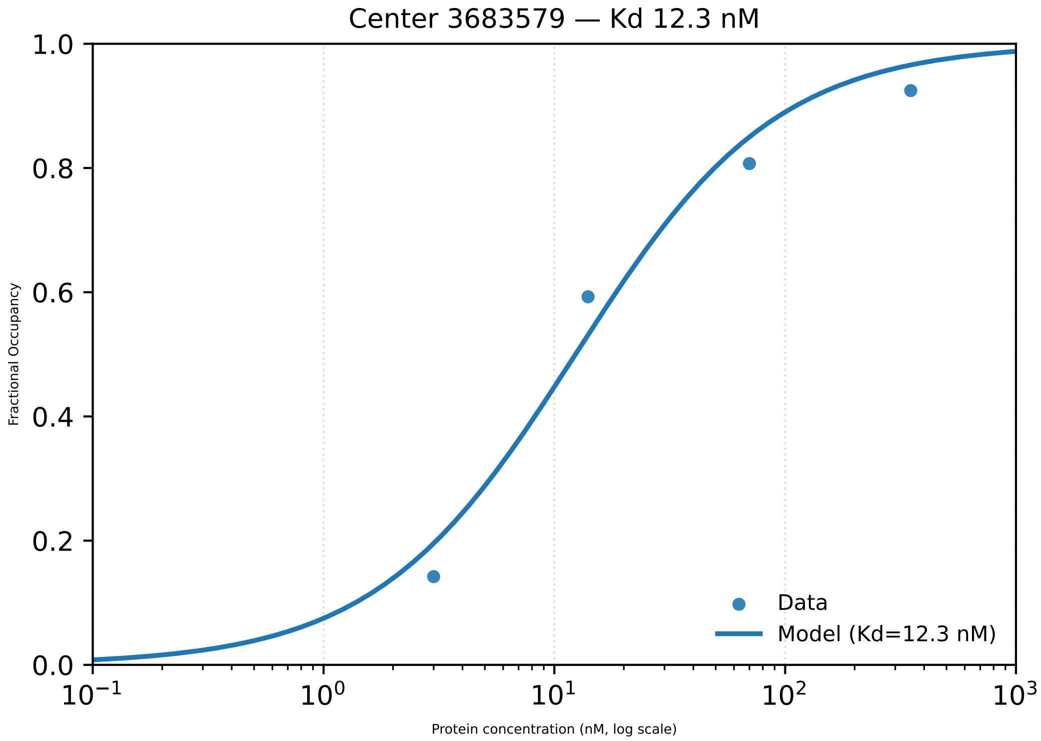

Center 3683579 — Kd 12.3 nM
1.0
0.8
0.6
0.4
0.2
0.0
Fractional Occupancy
$10^{-1}$
$10^{0}$
$10^{1}$
$10^{2}$
$10^{3}$
Protein concentration (nM, log scale)
Data
Model (Kd=12.3 nM)

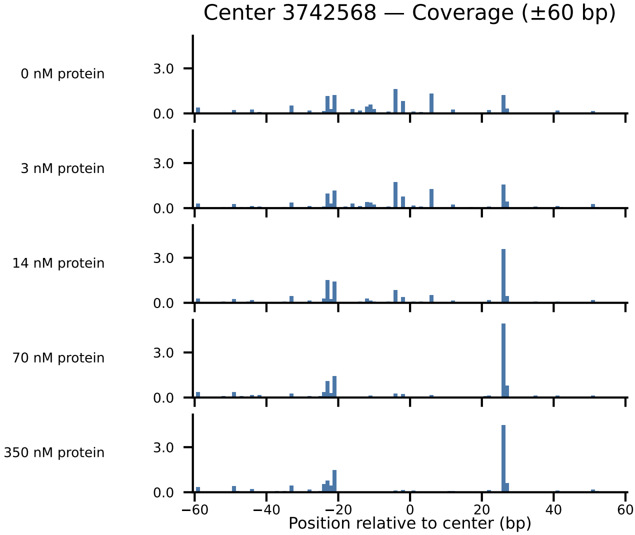
Center 3742568 — Coverage (±60 bp)
0 nM protein
3 nM protein
14 nM protein
70 nM protein
350 nM protein
3.0
0.0
−60
−40
−20
0
20
40
60
Position relative to center (bp)

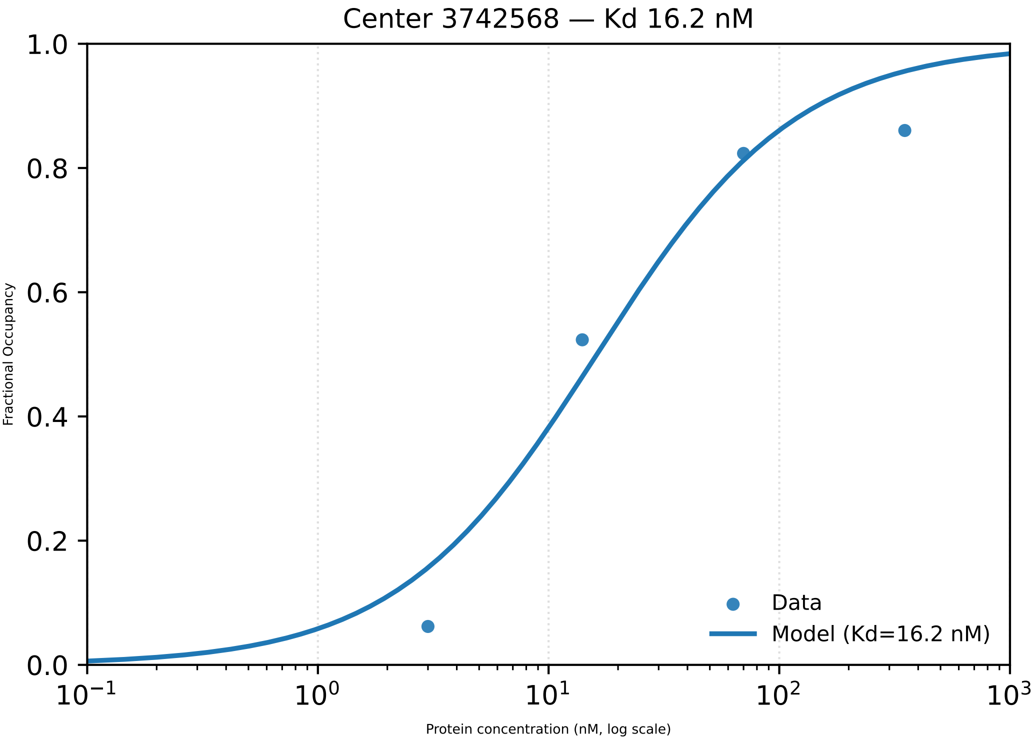

Center 3742568 — Kd 16.2 nM
Fractional Occupancy
1.0
0.8
0.6
0.4
0.2
0.0
$10^{-1}$
$10^{0}$
$10^{1}$
$10^{2}$
$10^{3}$
Protein concentration (nM, log scale)
Data
Model (Kd=16.2 nM)

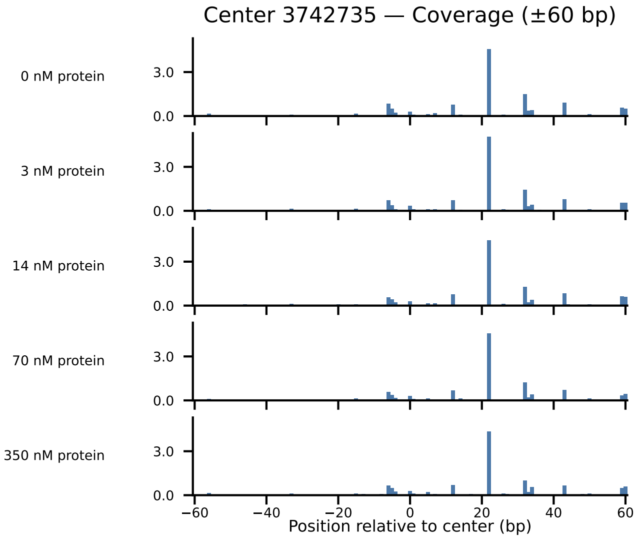

Center 3742735 — Coverage (±60 bp)
0 nM protein
3 nM protein
14 nM protein
70 nM protein
350 nM protein
3.0
0.0
−60
−40
−20
0
20
40
60
Position relative to center (bp)

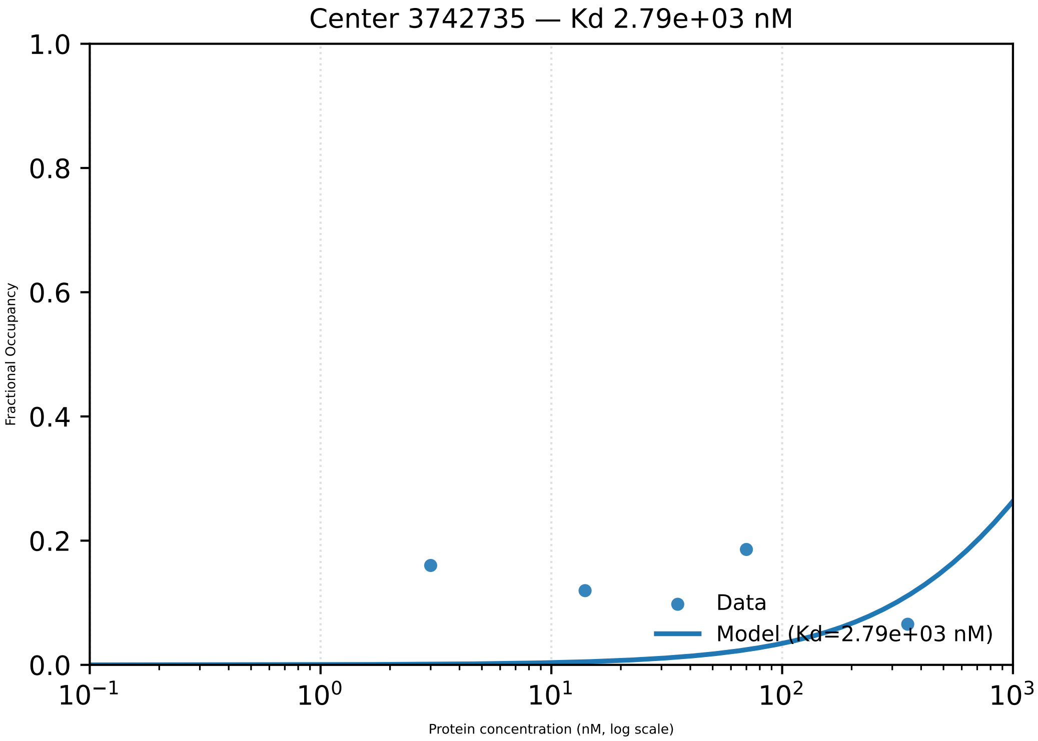

Center 3742735 — Kd 2.79e+03 nM
1.0
0.8
0.6
0.4
0.2
0.0
Fractional Occupancy
$10^{-1}$
$10^{0}$
$10^{1}$
$10^{2}$
$10^{3}$
Protein concentration (nM, log scale)
Data
Model (Kd=2.79e+03 nM)

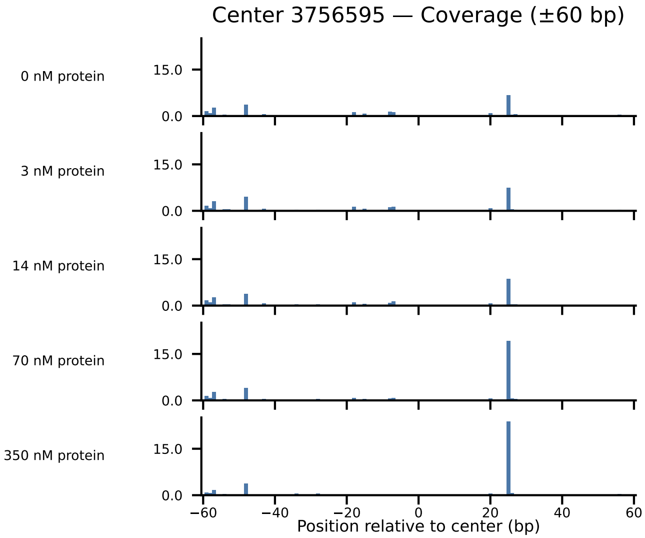
Center 3756595 — Coverage (±60 bp)
0 nM protein
3 nM protein
14 nM protein
70 nM protein
350 nM protein
15.0
0.0
−60
−40
−20
0
20
40
60
Position relative to center (bp)

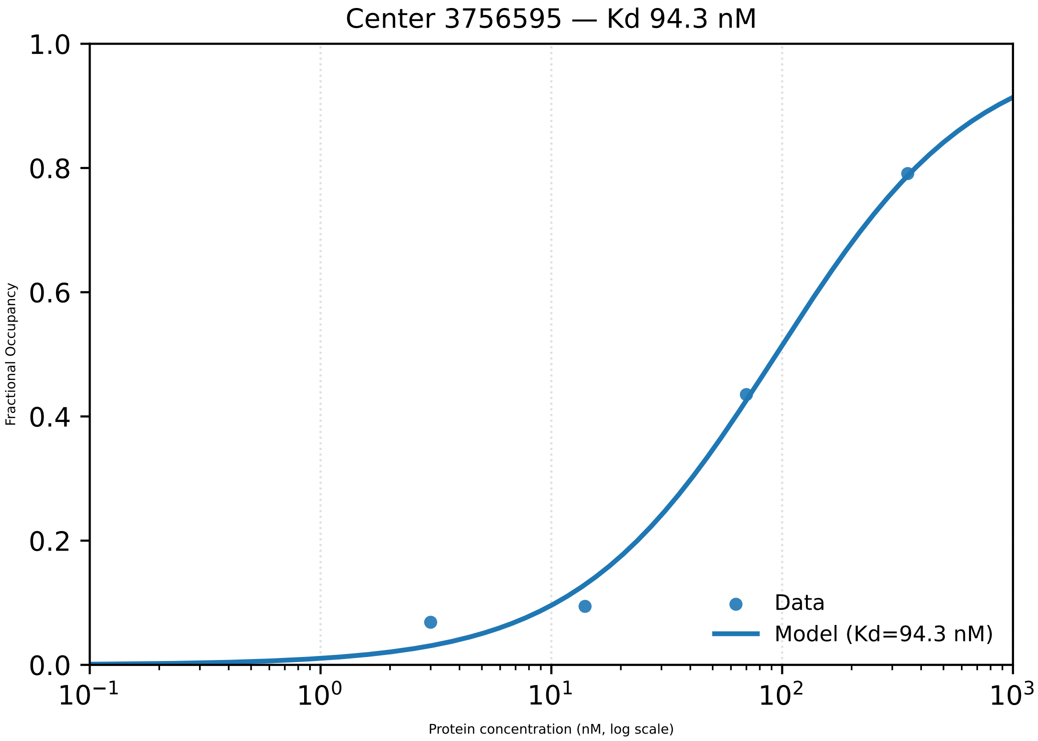

Center 3756595 — Kd 94.3 nM
Fractional Occupancy
1.0
0.8
0.6
0.4
0.2
0.0
$10^{-1}$
$10^{0}$
$10^{1}$
$10^{2}$
$10^{3}$
Protein concentration (nM, log scale)
Data
Model (Kd=94.3 nM)

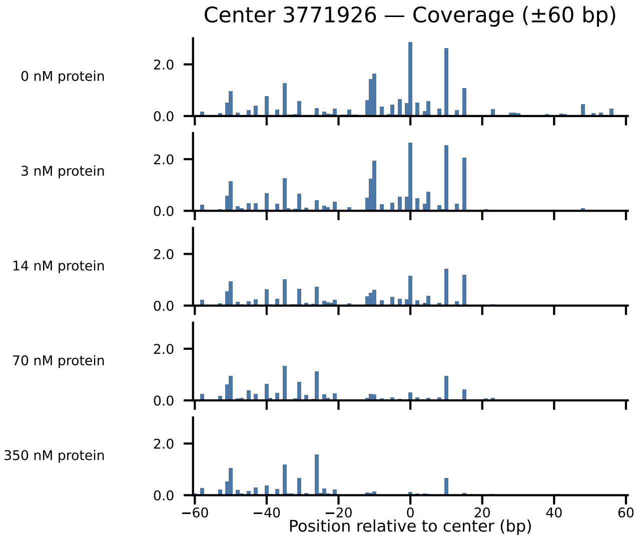
Center 3771926 — Coverage (±60 bp)
0 nM protein
3 nM protein
14 nM protein
70 nM protein
350 nM protein
2.0
0.0
−60
−40
−20
0
20
40
60
Position relative to center (bp)

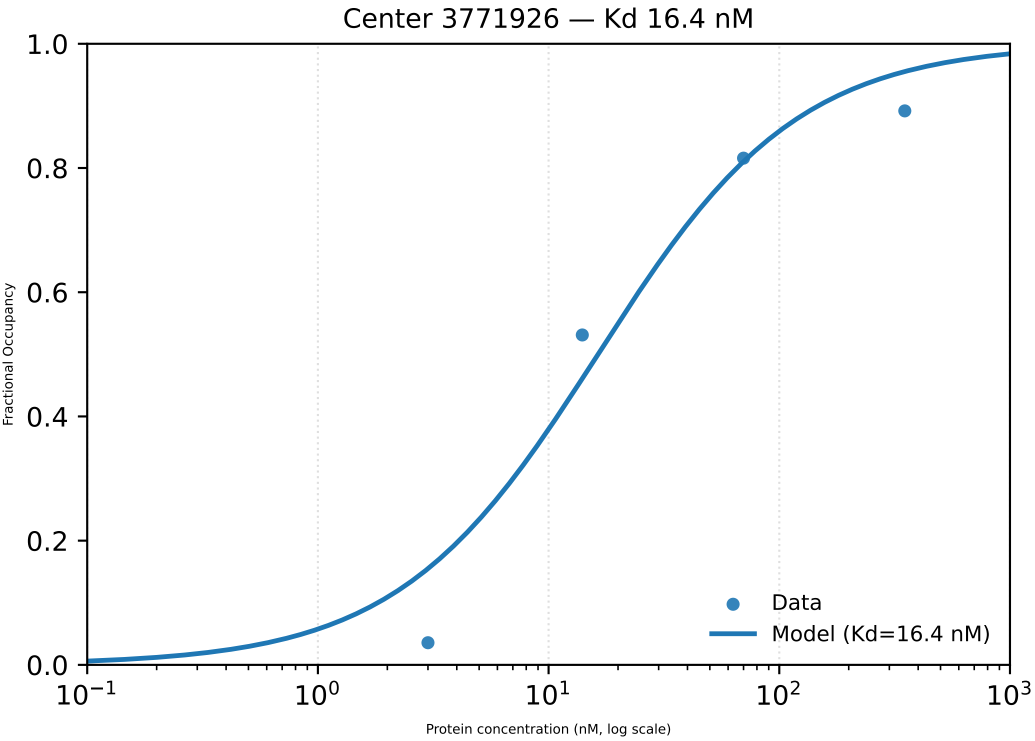

Center 3771926 — Kd 16.4 nM
1.0
0.8
0.6
0.4
0.2
0.0
Fractional Occupancy
$10^{-1}$
$10^{0}$
$10^{1}$
$10^{2}$
$10^{3}$
Protein concentration (nM, log scale)
Data
Model (Kd=16.4 nM)

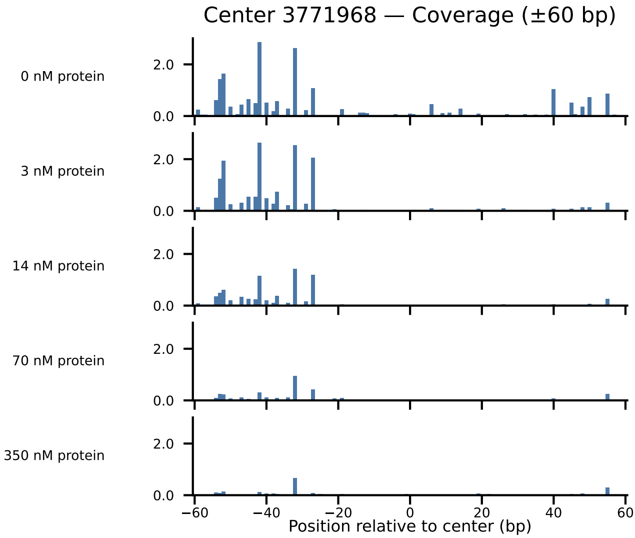

Center 3771968 — Coverage (±60 bp)
0 nM protein
3 nM protein
14 nM protein
70 nM protein
350 nM protein
2.0
0.0
−60
−40
−20
0
20
40
60
Position relative to center (bp)

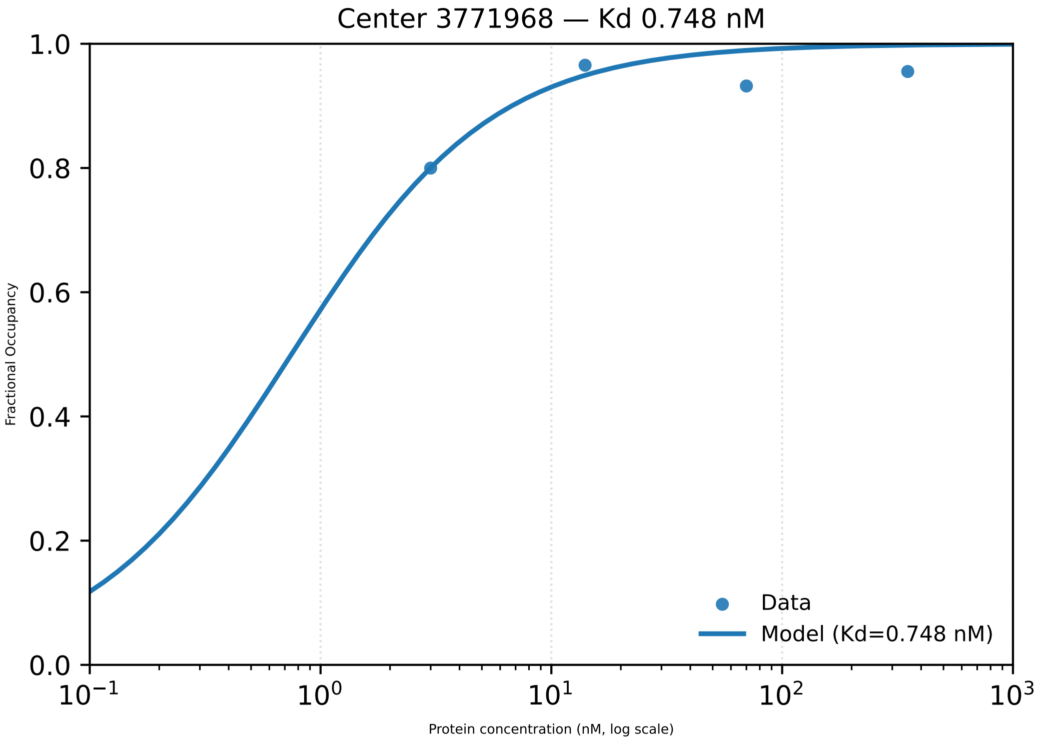

Center 3771968 — Kd 0.748 nM
Fractional Occupancy
1.0
0.8
0.6
0.4
0.2
0.0
$10^{-1}$
$10^{0}$
$10^{1}$
$10^{2}$
$10^{3}$
Protein concentration (nM, log scale)
Data
Model (Kd=0.748 nM)

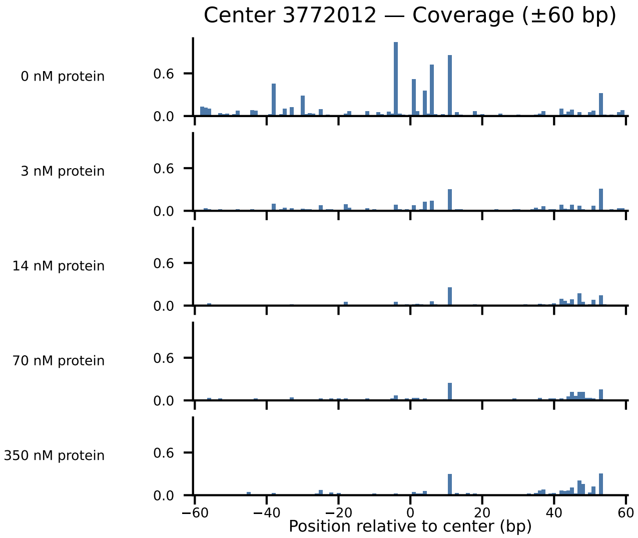

Center 3772012 — Coverage (±60 bp)
0 nM protein
3 nM protein
14 nM protein
70 nM protein
350 nM protein
0.6
0.0
−60
−40
−20
0
20
40
60
Position relative to center (bp)

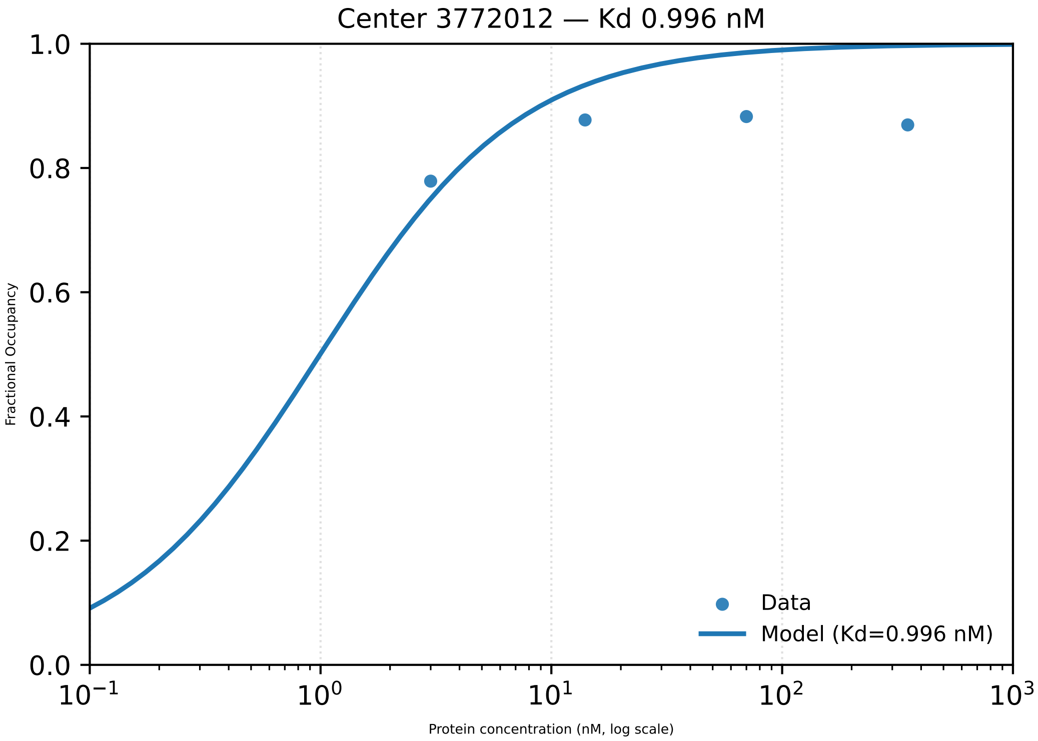

Center 3772012 — Kd 0.996 nM
1.0
0.8
0.6
0.4
0.2
0.0
Fractional Occupancy
$10^{-1}$
$10^{0}$
$10^{1}$
$10^{2}$
$10^{3}$
Protein concentration (nM, log scale)
Data
Model (Kd=0.996 nM)

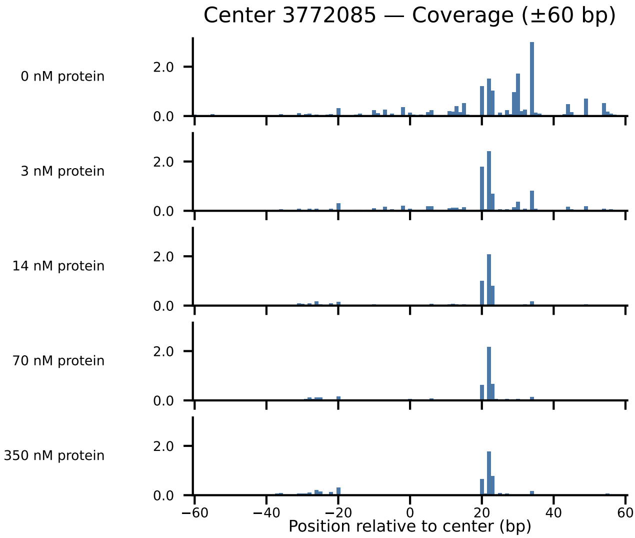

Center 3772085 — Coverage (±60 bp)
0 nM protein
3 nM protein
14 nM protein
70 nM protein
350 nM protein
2.0
0.0
−60
−40
−20
0
20
40
60
Position relative to center (bp)

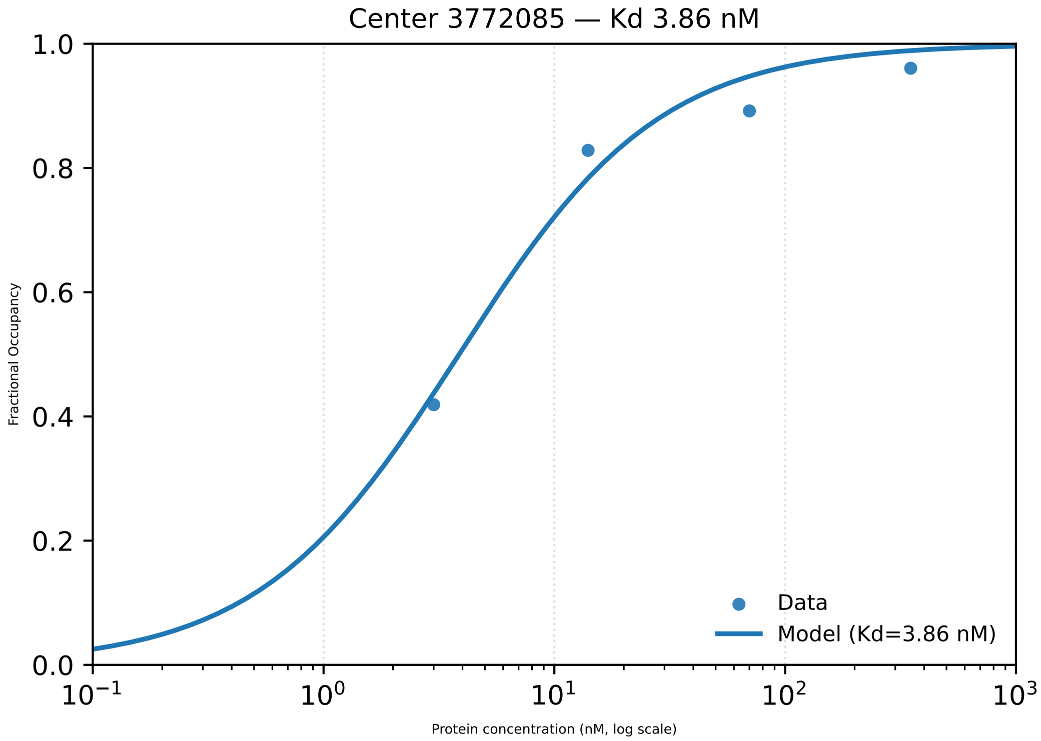

Center 3772085 — Kd 3.86 nM
Fractional Occupancy
1.0
0.8
0.6
0.4
0.2
0.0
$10^{-1}$
$10^{0}$
$10^{1}$
$10^{2}$
$10^{3}$
Protein concentration (nM, log scale)
Data
Model (Kd=3.86 nM)

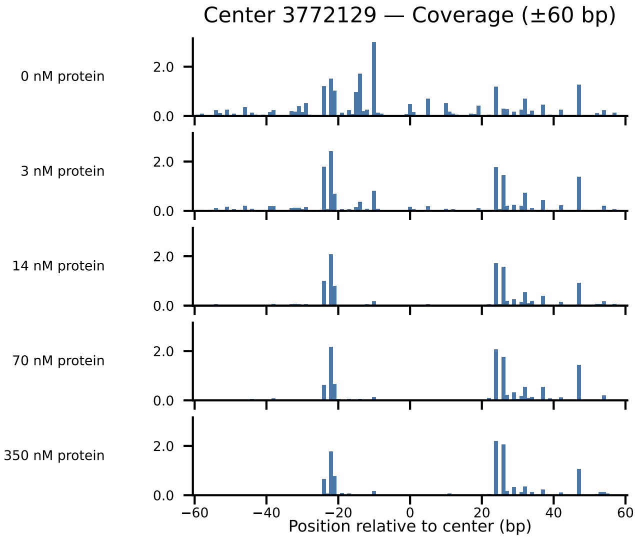
Center 3772129 — Coverage (±60 bp)
0 nM protein
3 nM protein
14 nM protein
70 nM protein
350 nM protein
2.0
0.0
−60
−40
−20
0
20
40
60
Position relative to center (bp)

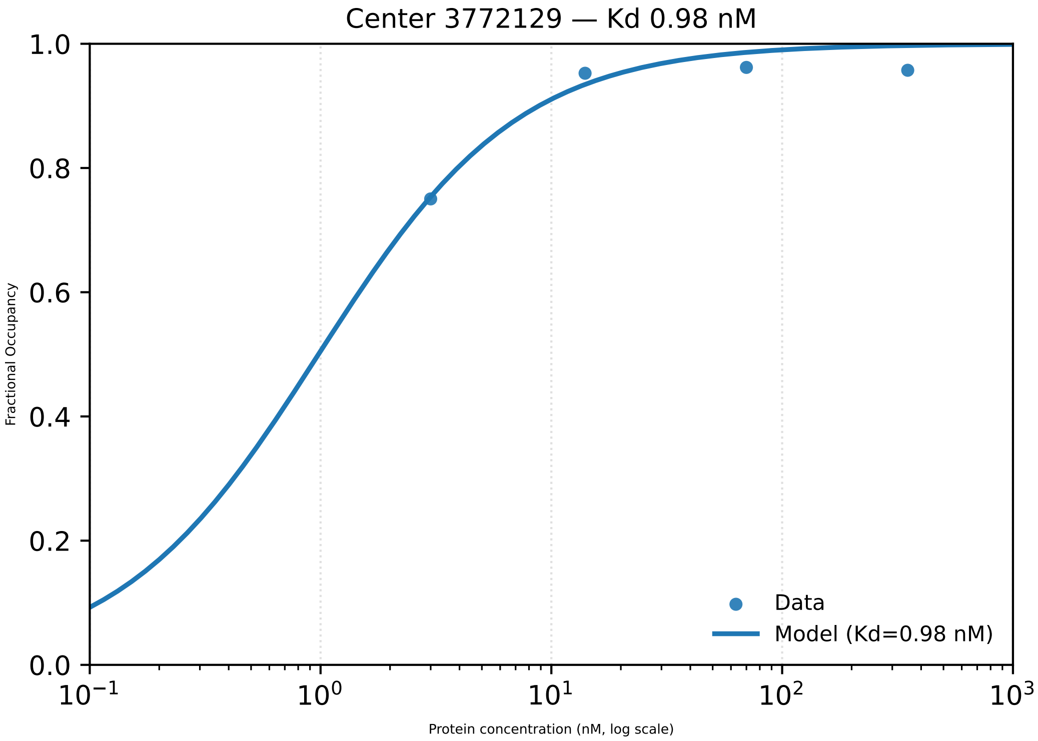

Center 3772129 — Kd 0.98 nM
Fractional Occupancy
1.0
0.8
0.6
0.4
0.2
0.0
$10^{-1}$
$10^{0}$
$10^{1}$
$10^{2}$
$10^{3}$
Protein concentration (nM, log scale)
Data
Model (Kd=0.98 nM)

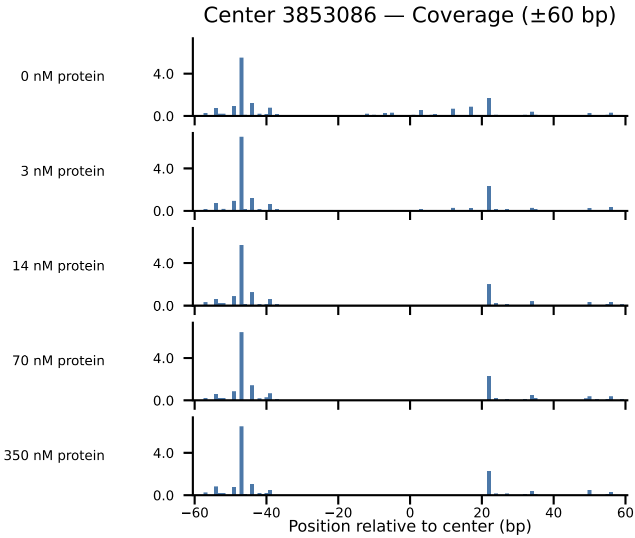
Center 3853086 — Coverage (±60 bp)
0 nM protein
3 nM protein
14 nM protein
70 nM protein
350 nM protein
4.0
0.0
−60
−40
−20
0
20
40
60
Position relative to center (bp)

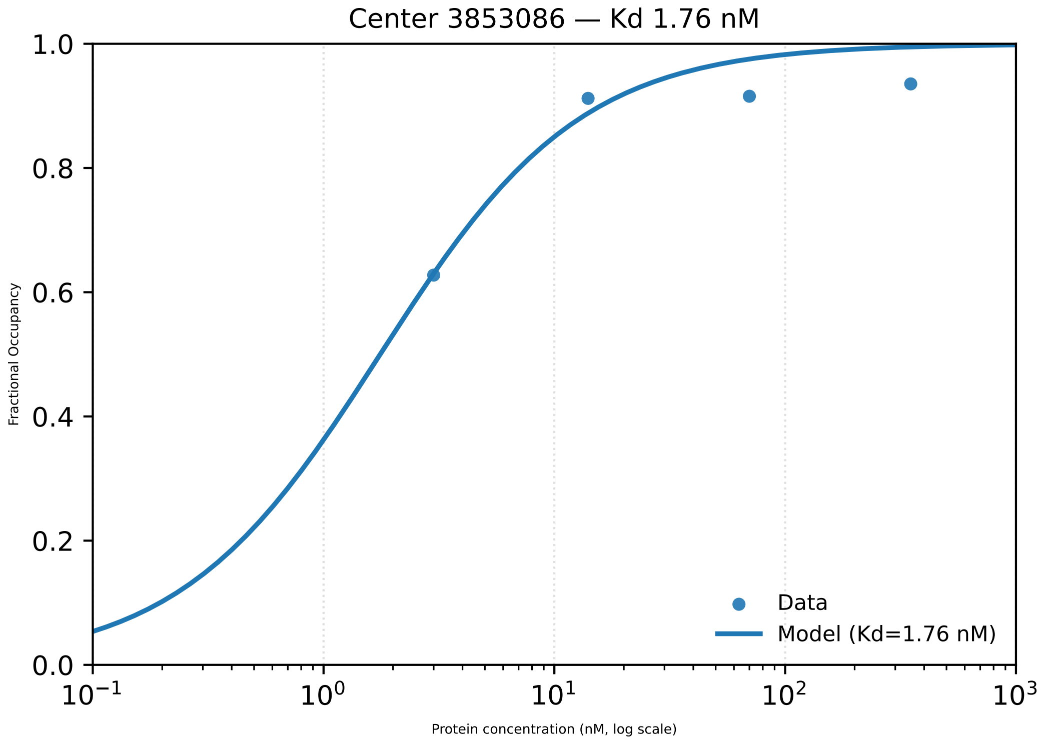

Center 3853086 — Kd 1.76 nM
1.0
0.8
0.6
0.4
0.2
0.0
Fractional Occupancy
$10^{-1}$
$10^{0}$
$10^{1}$
$10^{2}$
$10^{3}$
Protein concentration (nM, log scale)
Data
Model (Kd=1.76 nM)

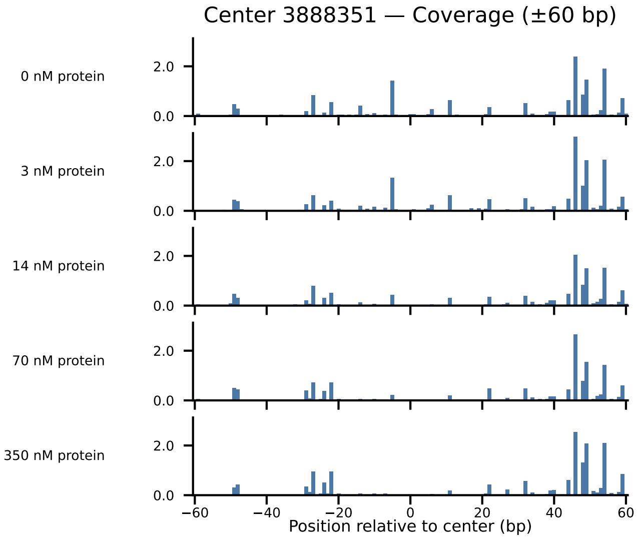

Center 3888351 — Coverage (±60 bp)
0 nM protein
3 nM protein
14 nM protein
70 nM protein
350 nM protein
2.0
0.0
−60
−40
−20
0
20
40
60
Position relative to center (bp)

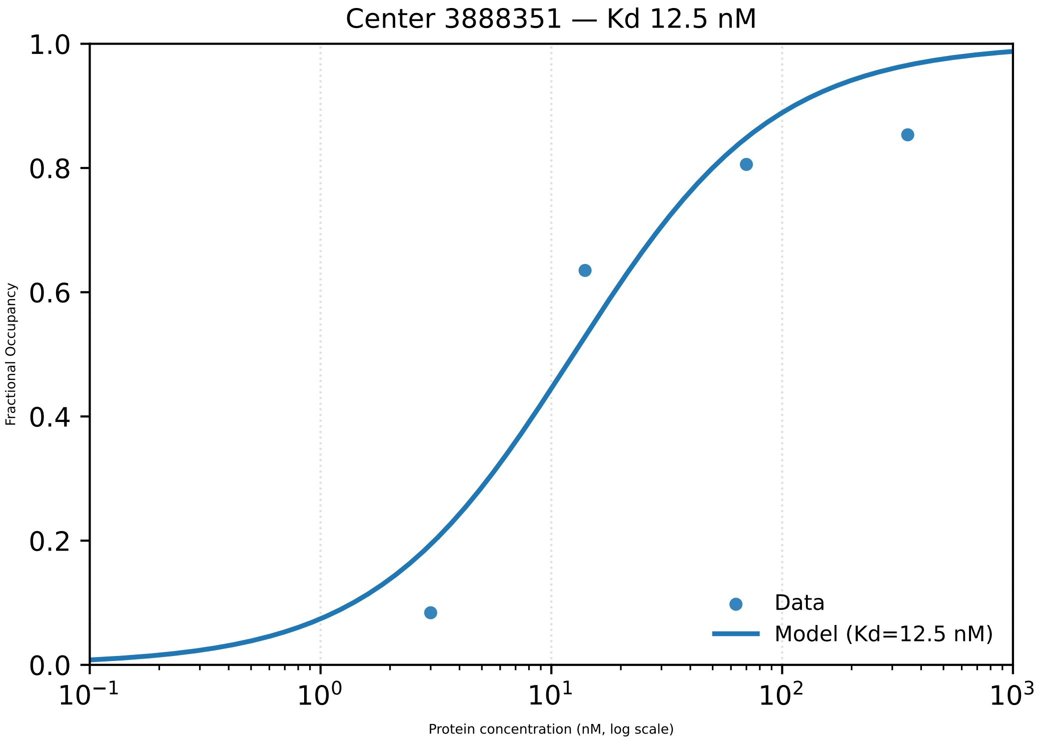
Center 3888351 — Kd 12.5 nM
Fractional Occupancy
1.0
0.8
0.6
0.4
0.2
0.0
$10^{-1}$
$10^{0}$
$10^{1}$
$10^{2}$
$10^{3}$
Protein concentration (nM, log scale)
Data
Model (Kd=12.5 nM)

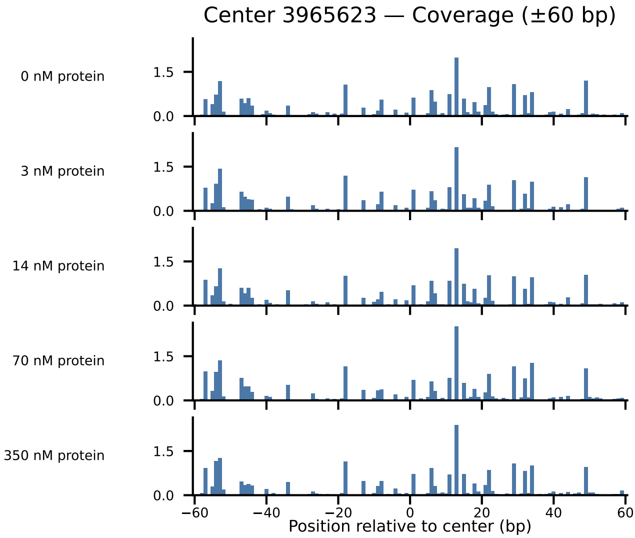

Center 3965623 — Coverage (±60 bp)
0 nM protein
3 nM protein
14 nM protein
70 nM protein
350 nM protein
1.5
0.0
−60
−40
−20
0
20
40
60
Position relative to center (bp)

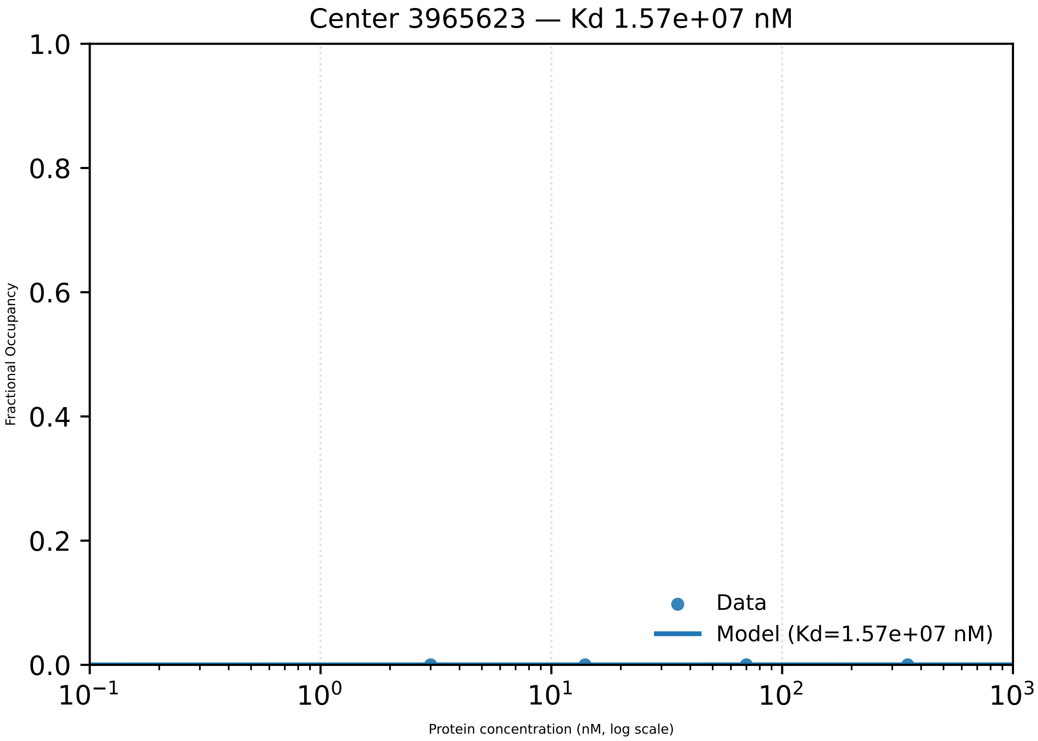

Center 3965623 — Kd 1.57e+07 nM
1.0
0.8
0.6
0.4
0.2
0.0
Fractional Occupancy
Data
Model (Kd=1.57e+07 nM)
$10^{-1}$
$10^{0}$
$10^{1}$
$10^{2}$
$10^{3}$
Protein concentration (nM, log scale)

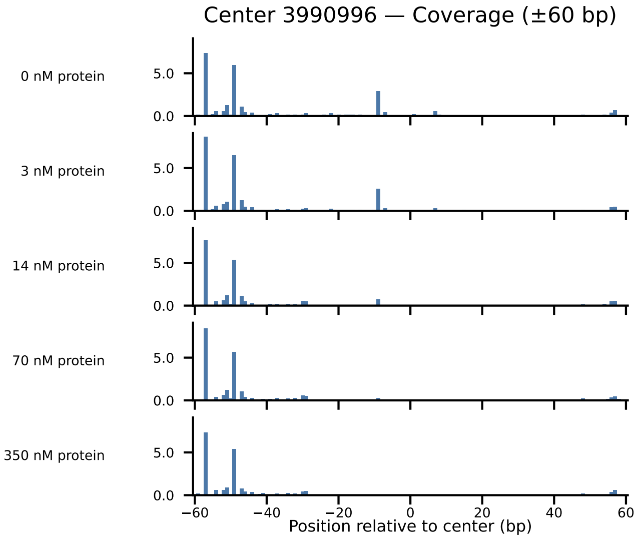
Center 3990996 — Coverage (±60 bp)
0 nM protein
3 nM protein
14 nM protein
70 nM protein
350 nM protein
5.0
0.0
−60
−40
−20
0
20
40
60
Position relative to center (bp)

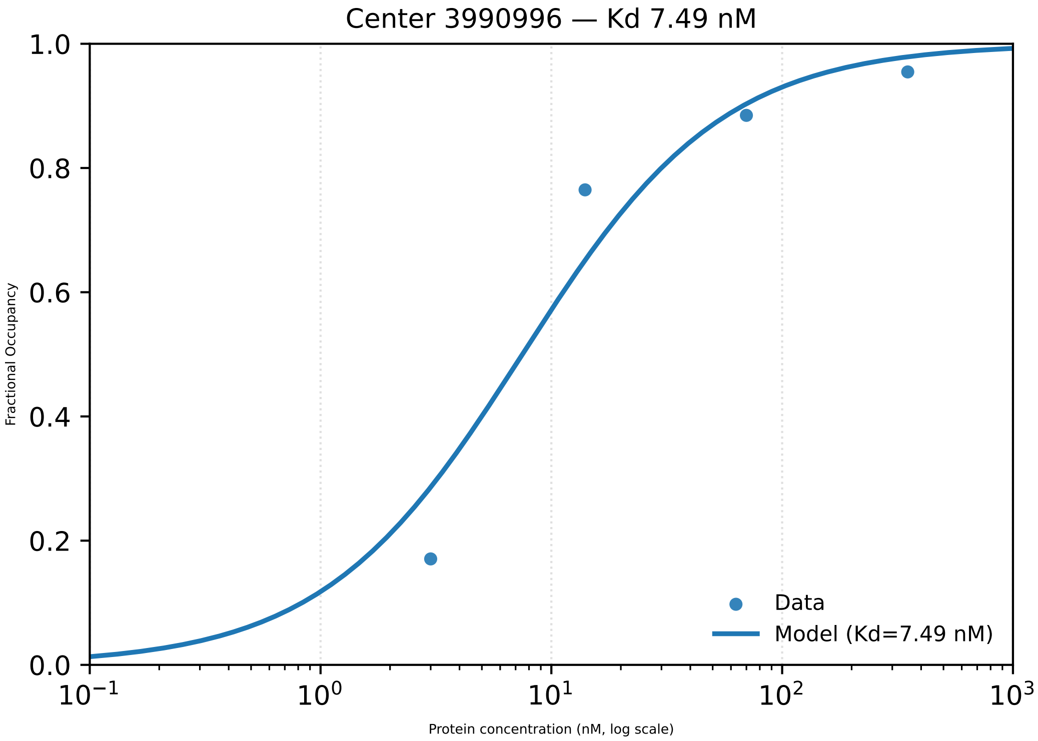

Center 3990996 — Kd 7.49 nM
Fractional Occupancy
1.0
0.8
0.6
0.4
0.2
0.0
$10^{-1}$
$10^{0}$
$10^{1}$
$10^{2}$
$10^{3}$
Protein concentration (nM, log scale)
Data
Model (Kd=7.49 nM)

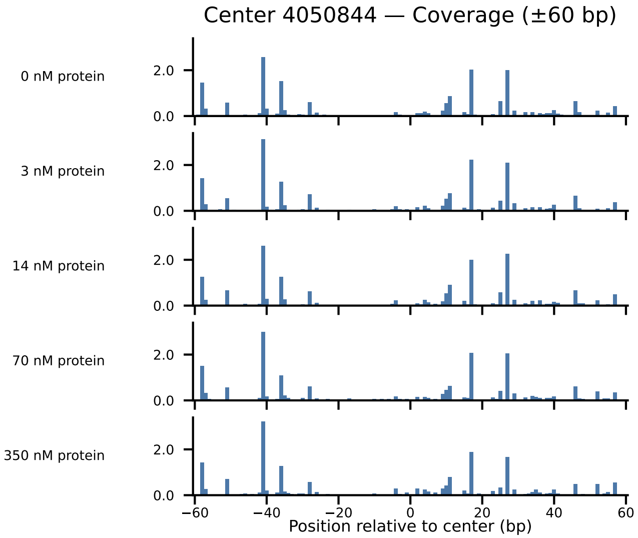

Center 4050844 — Coverage (±60 bp)
0 nM protein
3 nM protein
14 nM protein
70 nM protein
350 nM protein
2.0
0.0
−60
−40
−20
0
20
40
60
Position relative to center (bp)

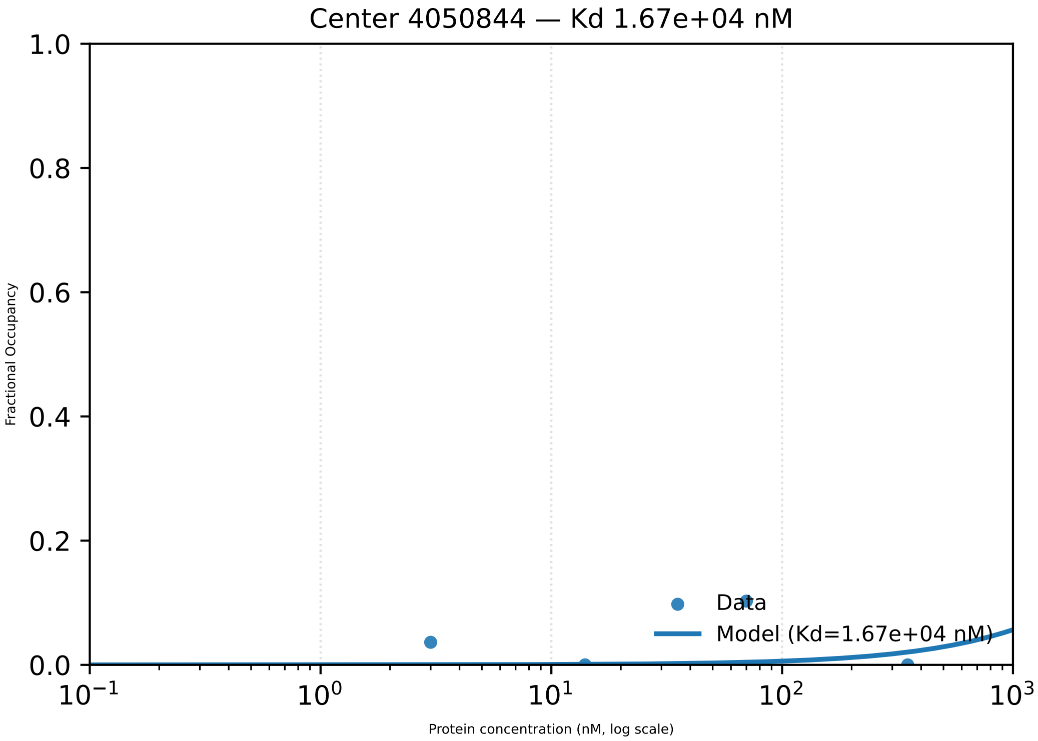

Center 4050844 — Kd 1.67e+04 nM
Fractional Occupancy
1.0
0.8
0.6
0.4
0.2
0.0
$10^{-1}$
$10^{0}$
$10^{1}$
$10^{2}$
$10^{3}$
Protein concentration (nM, log scale)
Data
Model (Kd=1.67e+04 nM)

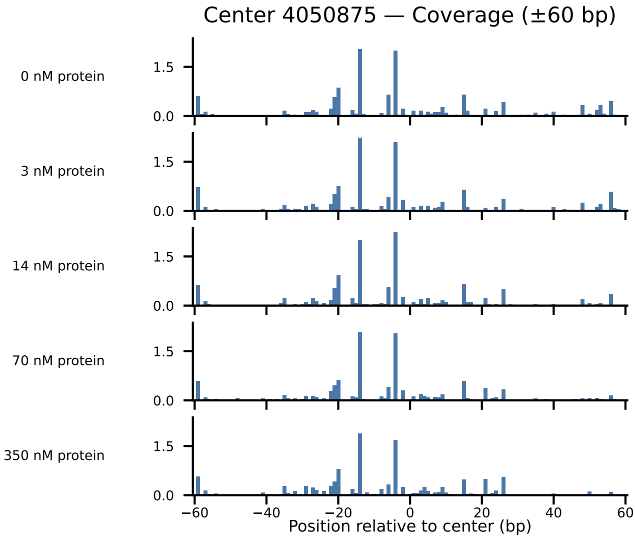
Center 4050875 — Coverage (±60 bp)
0 nM protein
3 nM protein
14 nM protein
70 nM protein
350 nM protein
1.5
0.0
−60
−40
−20
0
20
40
60
Position relative to center (bp)

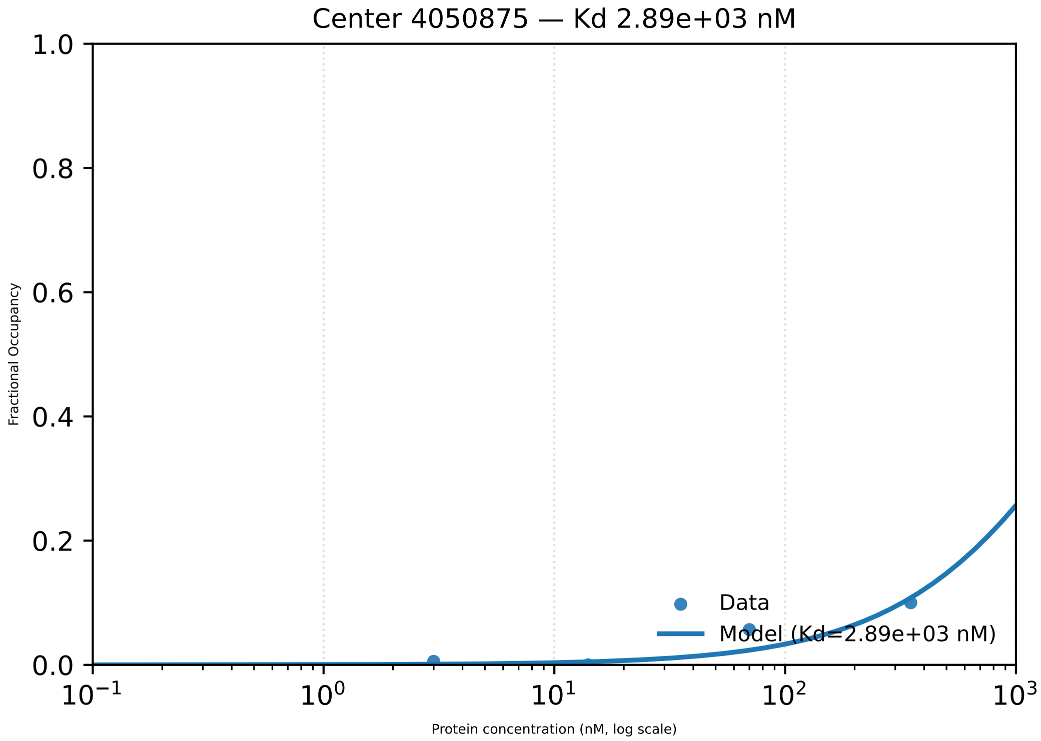

Center 4050875 — Kd 2.89e+03 nM
1.0
0.8
0.6
0.4
0.2
0.0
Fractional Occupancy
$10^{-1}$
$10^{0}$
$10^{1}$
$10^{2}$
$10^{3}$
Protein concentration (nM, log scale)
Data
Model (Kd=2.89e+03 nM)

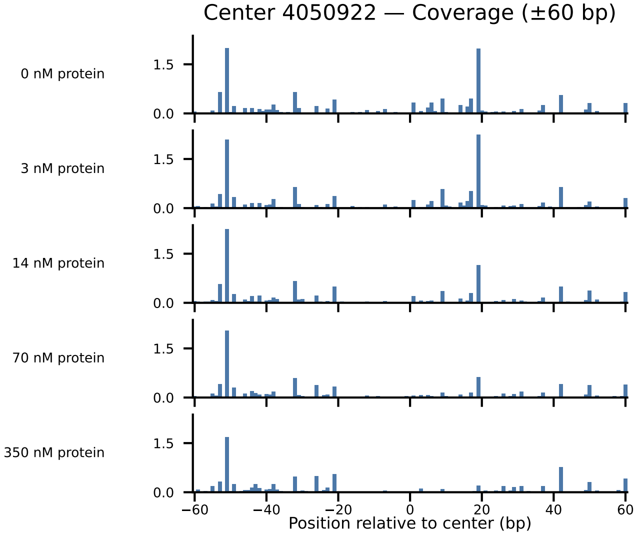
Center 4050922 — Coverage (±60 bp)
0 nM protein
3 nM protein
14 nM protein
70 nM protein
350 nM protein
1.5
0.0
−60
−40
−20
0
20
40
60
Position relative to center (bp)

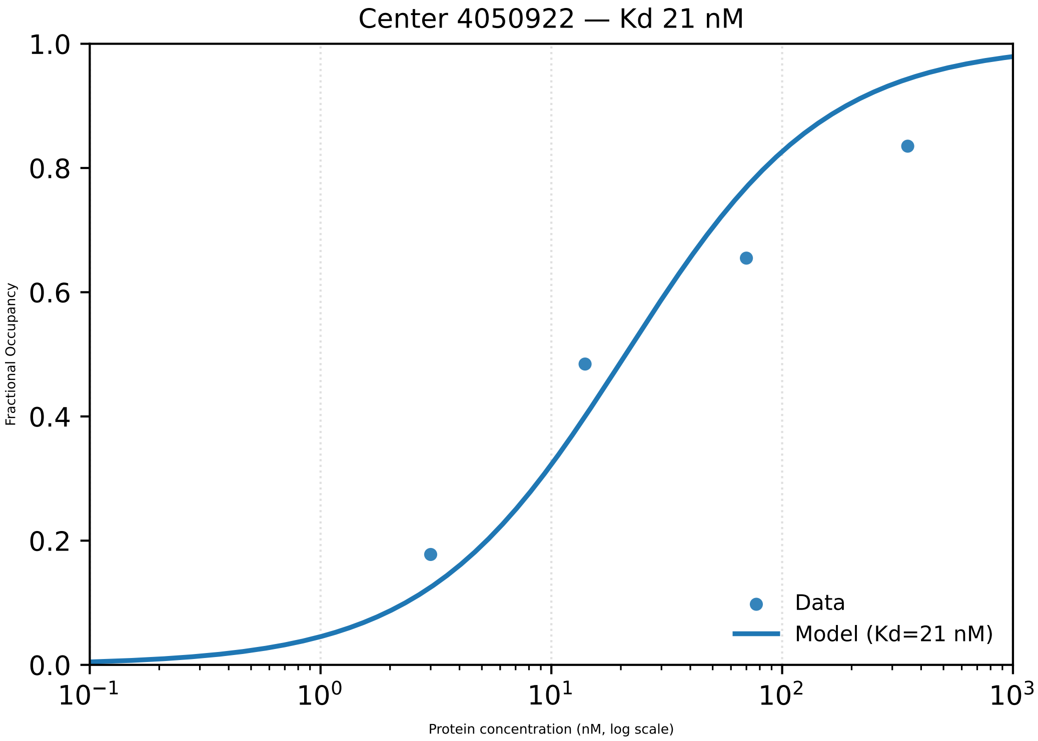

Center 4050922 — Kd 21 nM
Fractional Occupancy
0.0
0.2
0.4
0.6
0.8
1.0
$10^{-1}$
$10^{0}$
$10^{1}$
$10^{2}$
$10^{3}$
Protein concentration (nM, log scale)
Data
Model (Kd=21 nM)

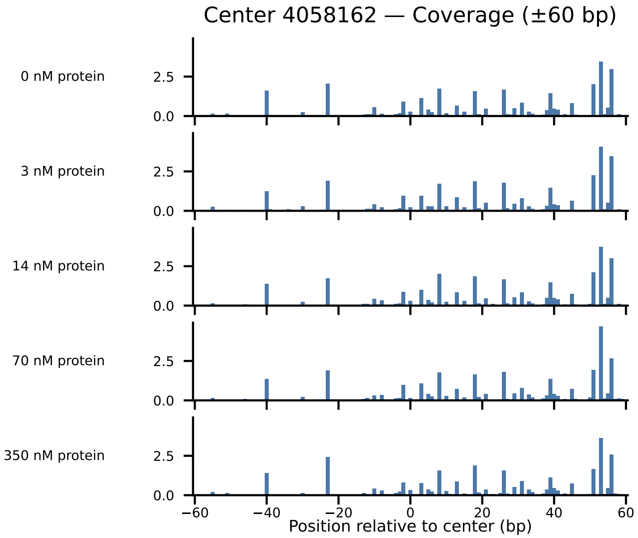
Center 4058162 — Coverage (±60 bp)
0 nM protein
3 nM protein
14 nM protein
70 nM protein
350 nM protein
2.5
0.0
−60
−40
−20
0
20
40
60
Position relative to center (bp)

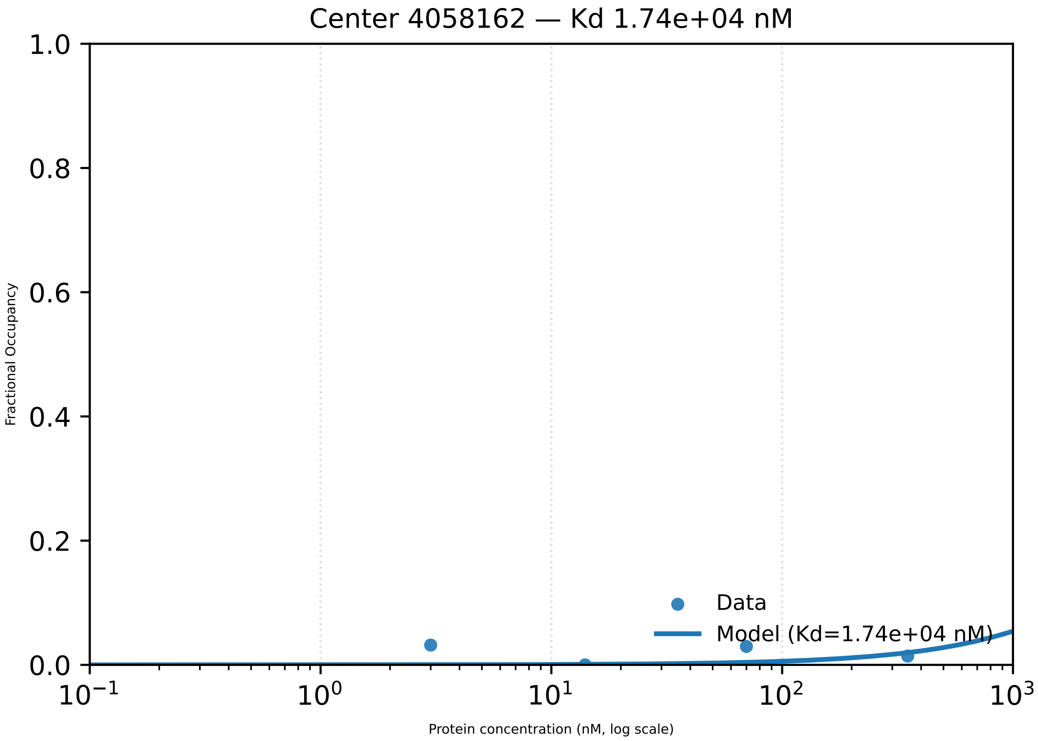
Center 4058162 — Kd 1.74e+04 nM
1.0
0.8
0.6
0.4
0.2
0.0
Fractional Occupancy
$10^{-1}$
$10^{0}$
$10^{1}$
$10^{2}$
$10^{3}$
Protein concentration (nM, log scale)
Data
Model (Kd=1.74e+04 nM)

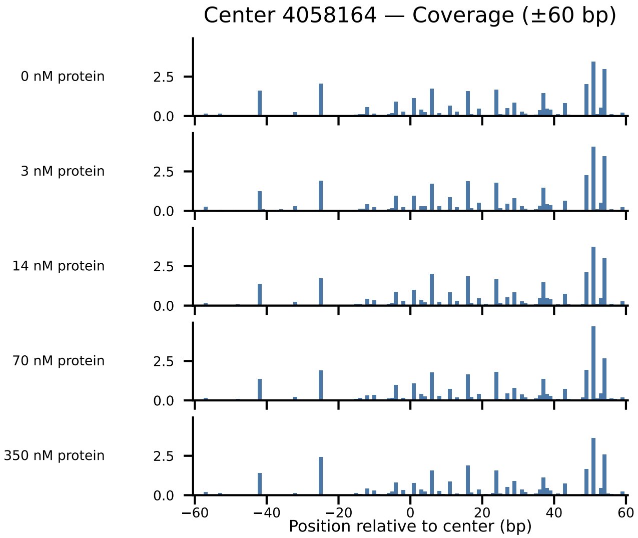

Center 4058164 — Coverage (±60 bp)
0 nM protein
3 nM protein
14 nM protein
70 nM protein
350 nM protein
2.5
0.0
−60
−40
−20
0
20
40
60
Position relative to center (bp)

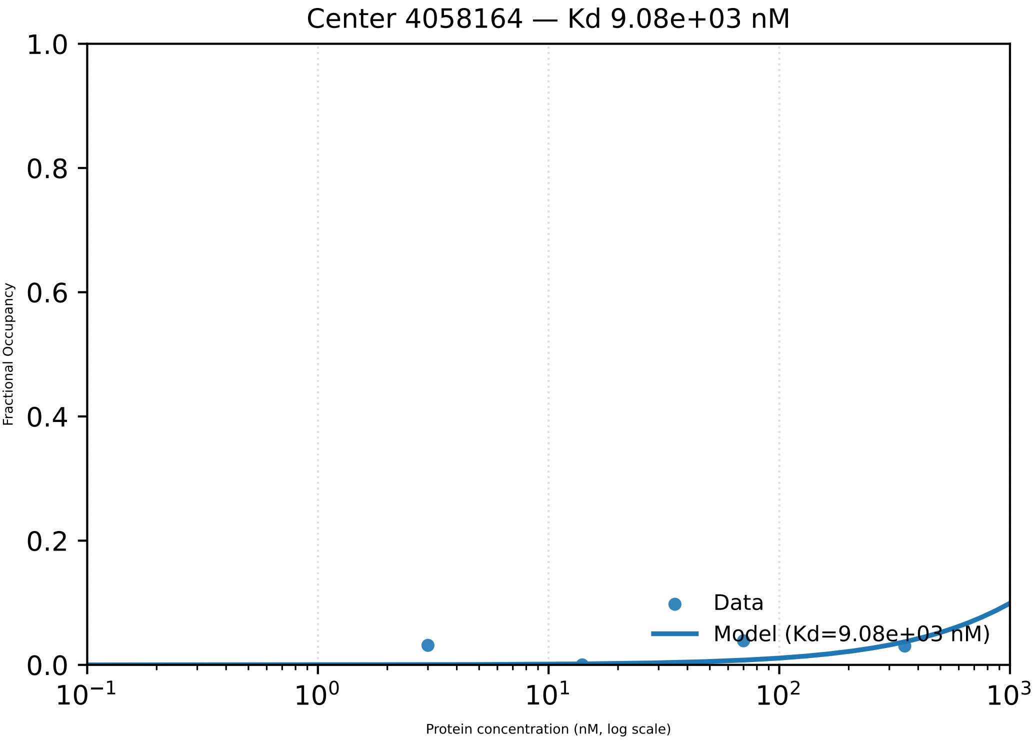

Center 4058164 — Kd 9.08e+03 nM
1.0
0.8
0.6
0.4
0.2
0.0
Fractional Occupancy
Data
Model (Kd=9.08e+03 nM)
$10^{-1}$
$10^{0}$
$10^{1}$
$10^{2}$
$10^{3}$
Protein concentration (nM, log scale)

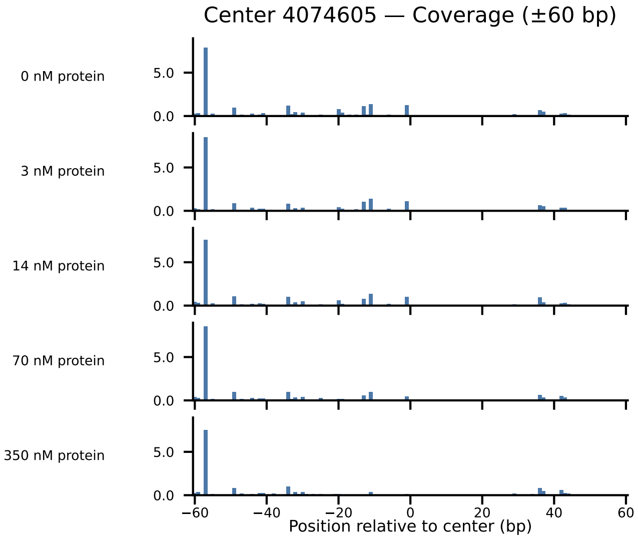
Center 4074605 — Coverage (±60 bp)
0 nM protein
3 nM protein
14 nM protein
70 nM protein
350 nM protein
5.0
0.0
−60
−40
−20
0
20
40
60
Position relative to center (bp)

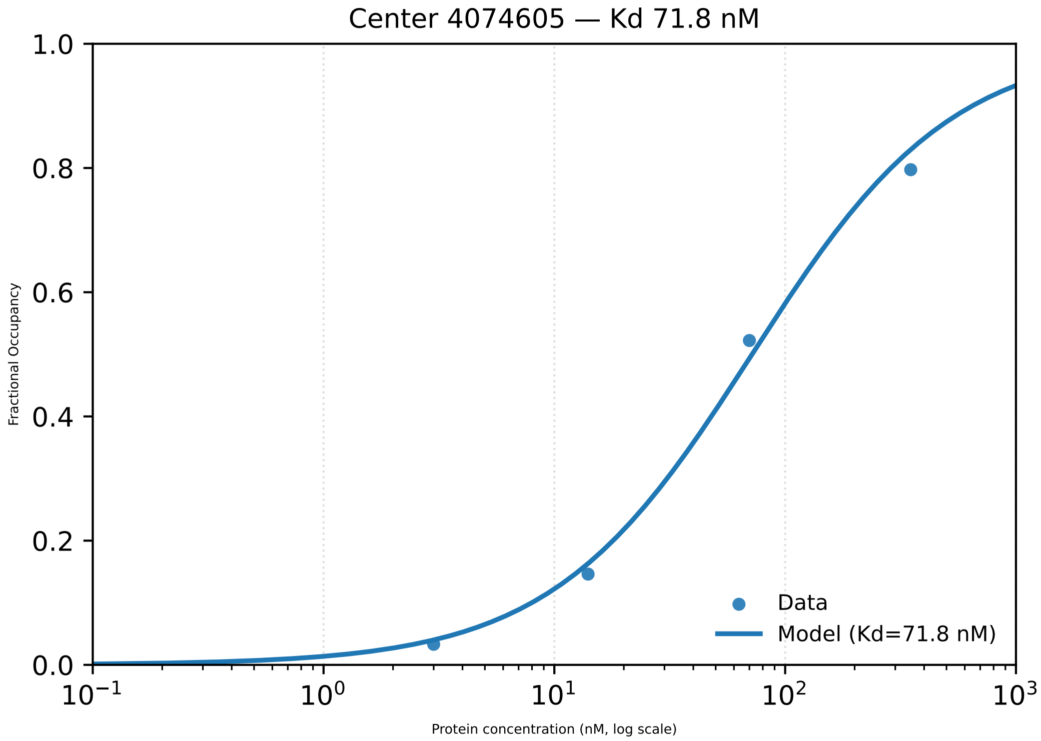

Center 4074605 — Kd 71.8 nM
1.0
0.8
0.6
0.4
0.2
0.0
Fractional Occupancy
$10^{-1}$
$10^{0}$
$10^{1}$
$10^{2}$
$10^{3}$
Protein concentration (nM, log scale)
Data
Model (Kd=71.8 nM)

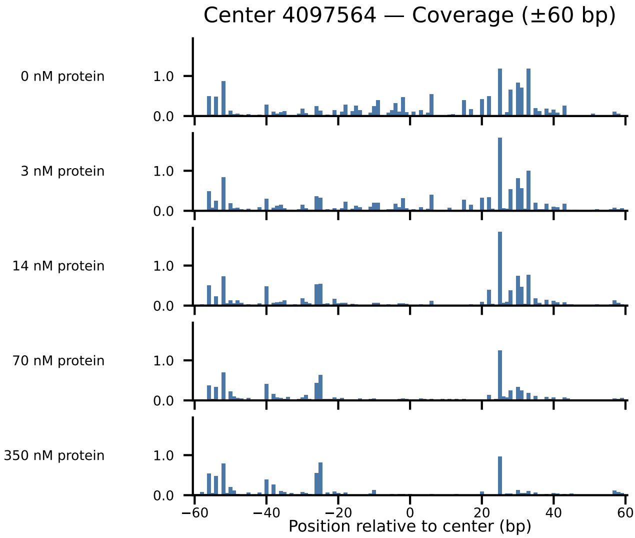
Center 4097564 — Coverage (±60 bp)
0 nM protein
3 nM protein
14 nM protein
70 nM protein
350 nM protein
1.0
0.0
−60
−40
−20
0
20
40
60
Position relative to center (bp)

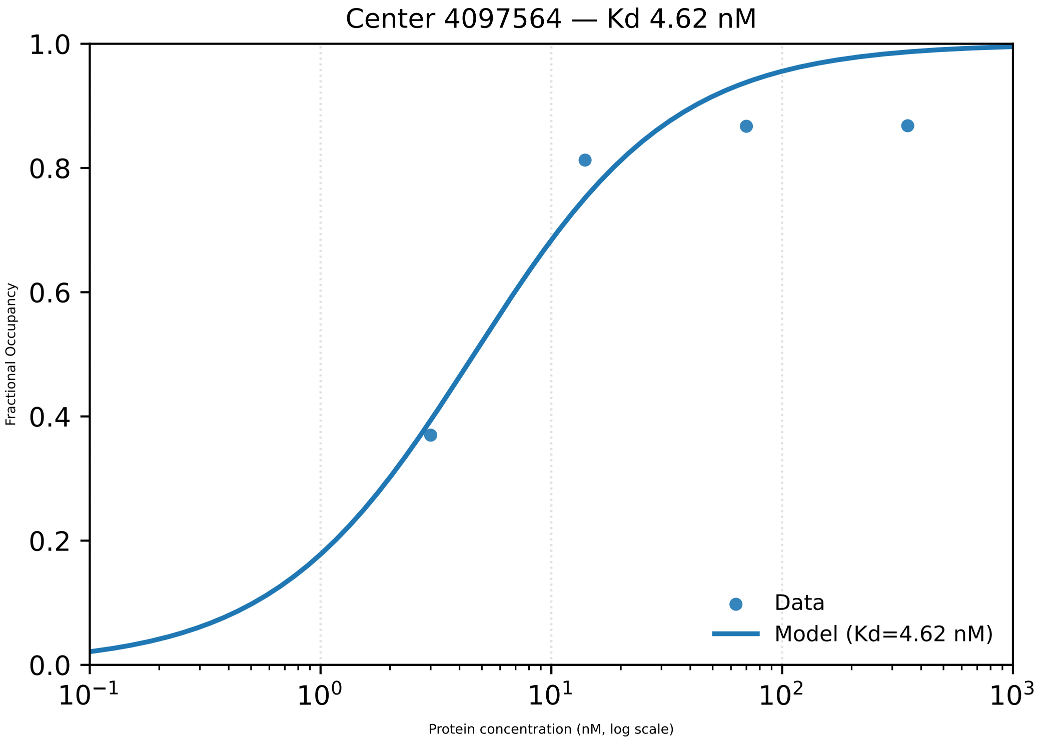
Center 4097564 — Kd 4.62 nM
Fractional Occupancy
1.0
0.8
0.6
0.4
0.2
0.0
$10^{-1}$
$10^{0}$
$10^{1}$
$10^{2}$
$10^{3}$
Protein concentration (nM, log scale)
Data
Model (Kd=4.62 nM)

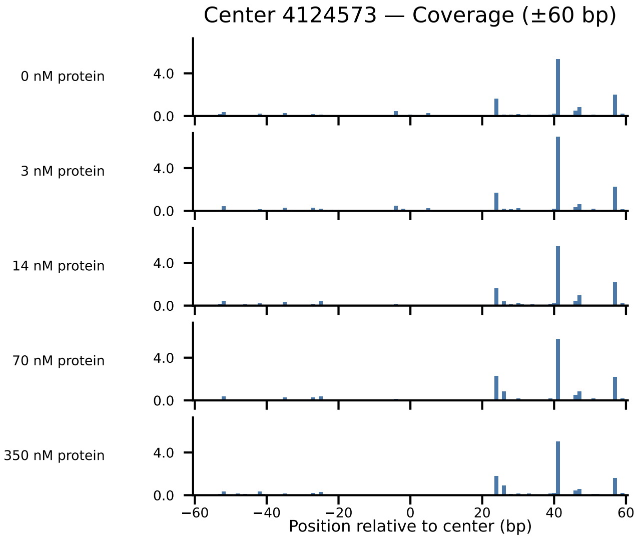

Center 4124573 — Coverage (±60 bp)
0 nM protein
3 nM protein
14 nM protein
70 nM protein
350 nM protein
4.0
0.0
−60
−40
−20
0
20
40
60
Position relative to center (bp)

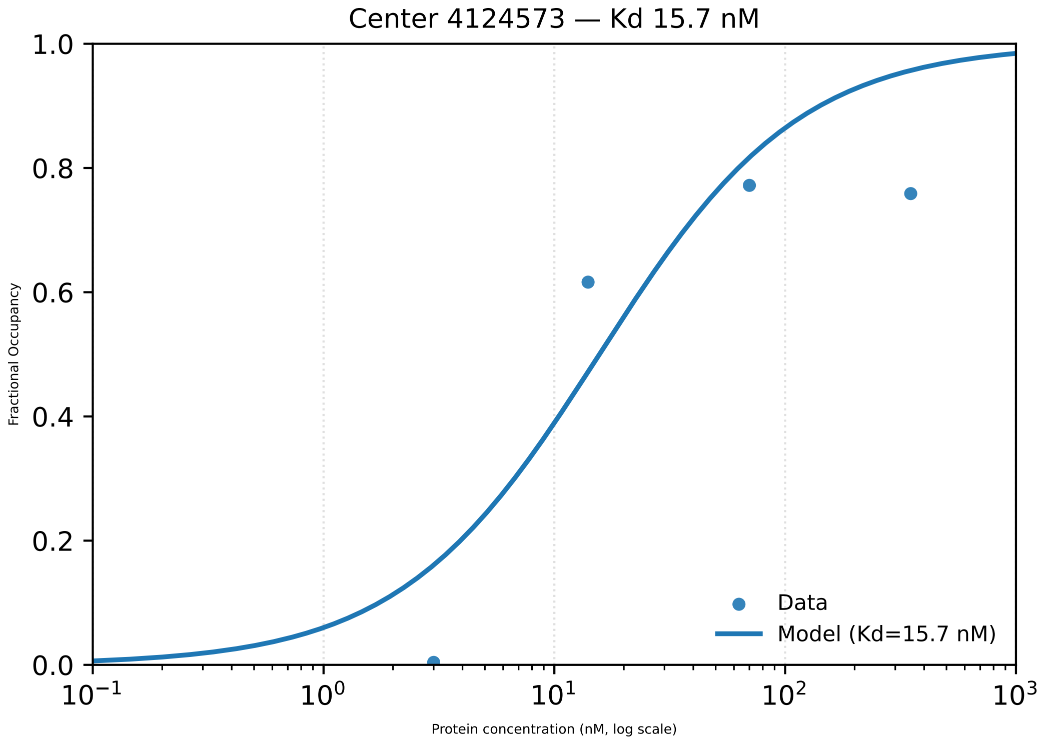

Center 4124573 — Kd 15.7 nM
1.0
0.8
0.6
0.4
0.2
0.0
Fractional Occupancy
$10^{-1}$
$10^{0}$
$10^{1}$
$10^{2}$
$10^{3}$
Protein concentration (nM, log scale)
Data
Model (Kd=15.7 nM)

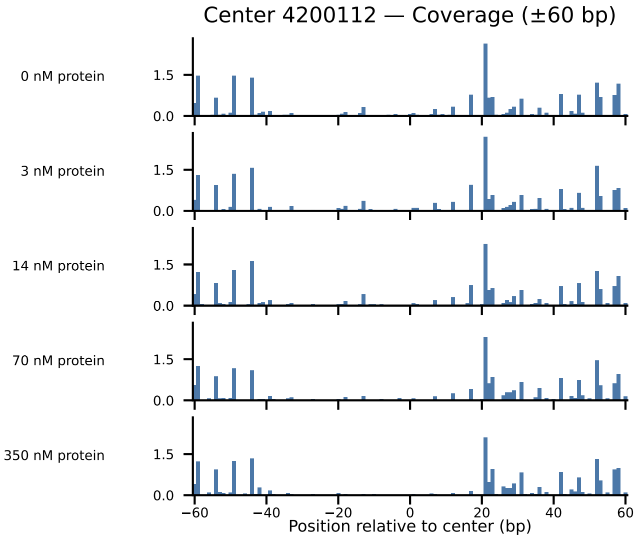
Center 4200112 — Coverage (±60 bp)
0 nM protein
3 nM protein
14 nM protein
70 nM protein
350 nM protein
1.5
0.0
−60
−40
−20
0
20
40
60
Position relative to center (bp)

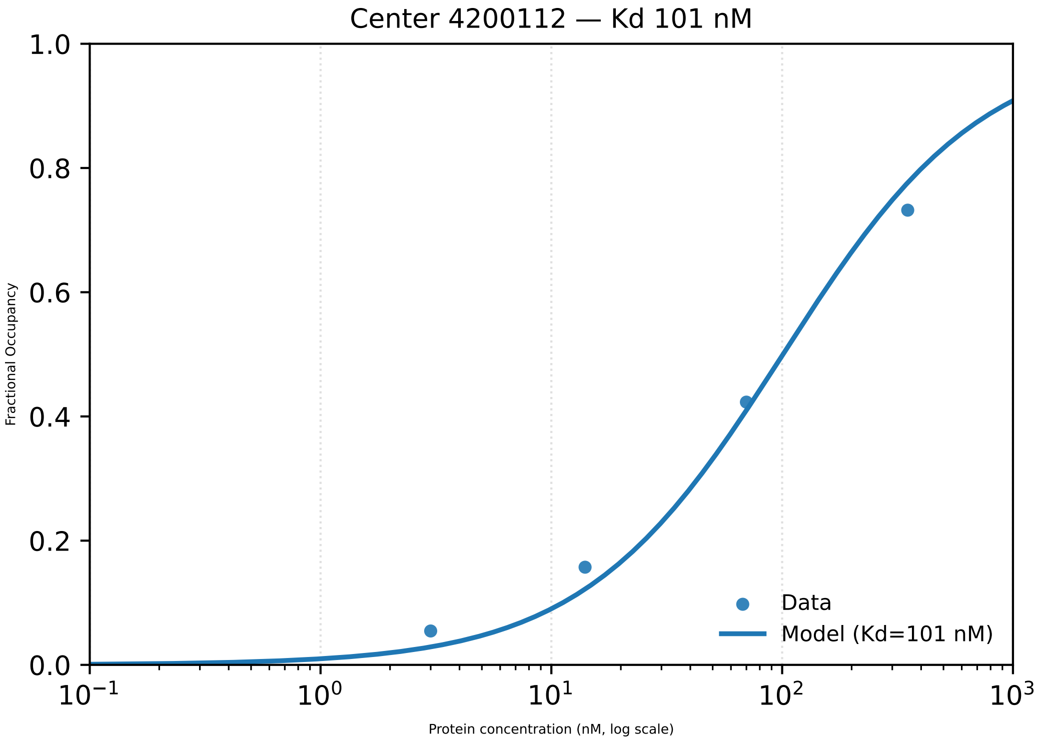

Center 4200112 — Kd 101 nM
Fractional Occupancy
0.0
0.2
0.4
0.6
0.8
1.0
$10^{-1}$
$10^{0}$
$10^{1}$
$10^{2}$
$10^{3}$
Protein concentration (nM, log scale)
Data
Model (Kd=101 nM)

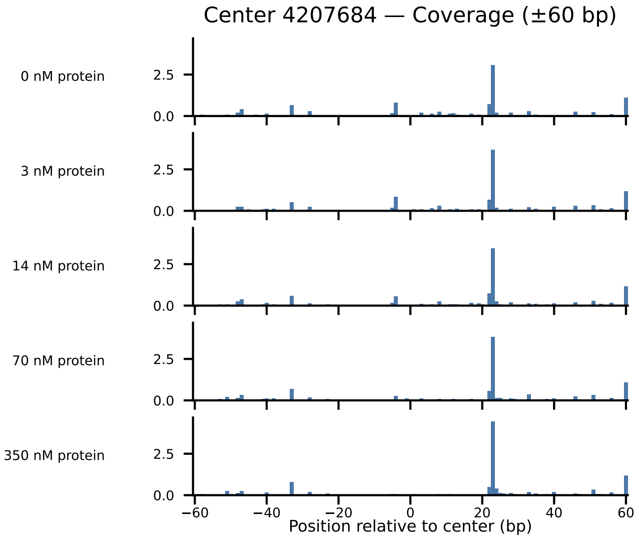

Center 4207684 — Coverage (±60 bp)
0 nM protein
3 nM protein
14 nM protein
70 nM protein
350 nM protein
2.5
0.0
−60
−40
−20
0
20
40
60
Position relative to center (bp)

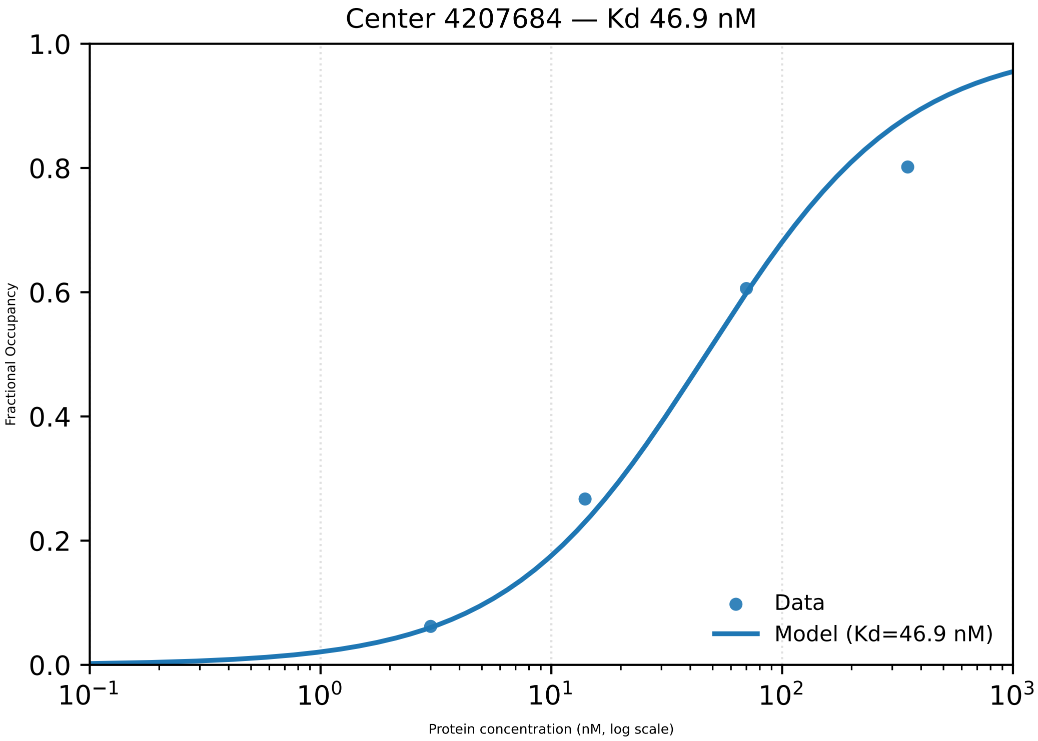

Center 4207684 — Kd 46.9 nM
1.0
0.8
0.6
0.4
0.2
0.0
Fractional Occupancy
$10^{-1}$
$10^{0}$
$10^{1}$
$10^{2}$
$10^{3}$
Protein concentration (nM, log scale)
Data
Model (Kd=46.9 nM)

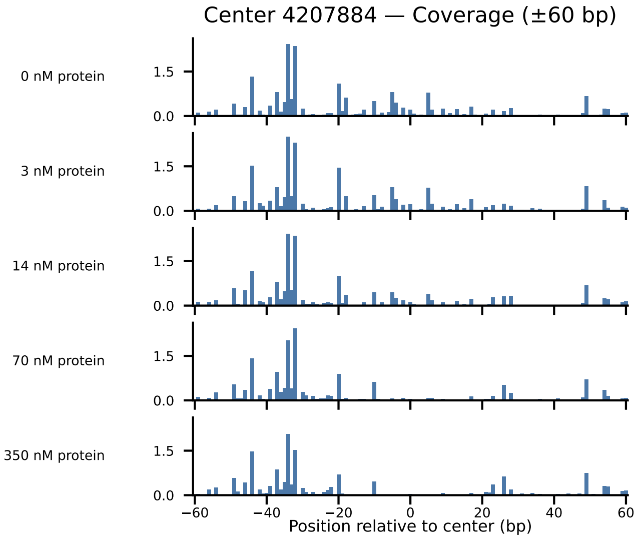

Center 4207884 — Coverage (±60 bp)
0 nM protein
3 nM protein
14 nM protein
70 nM protein
350 nM protein
1.5
0.0
−60
−40
−20
0
20
40
60
Position relative to center (bp)

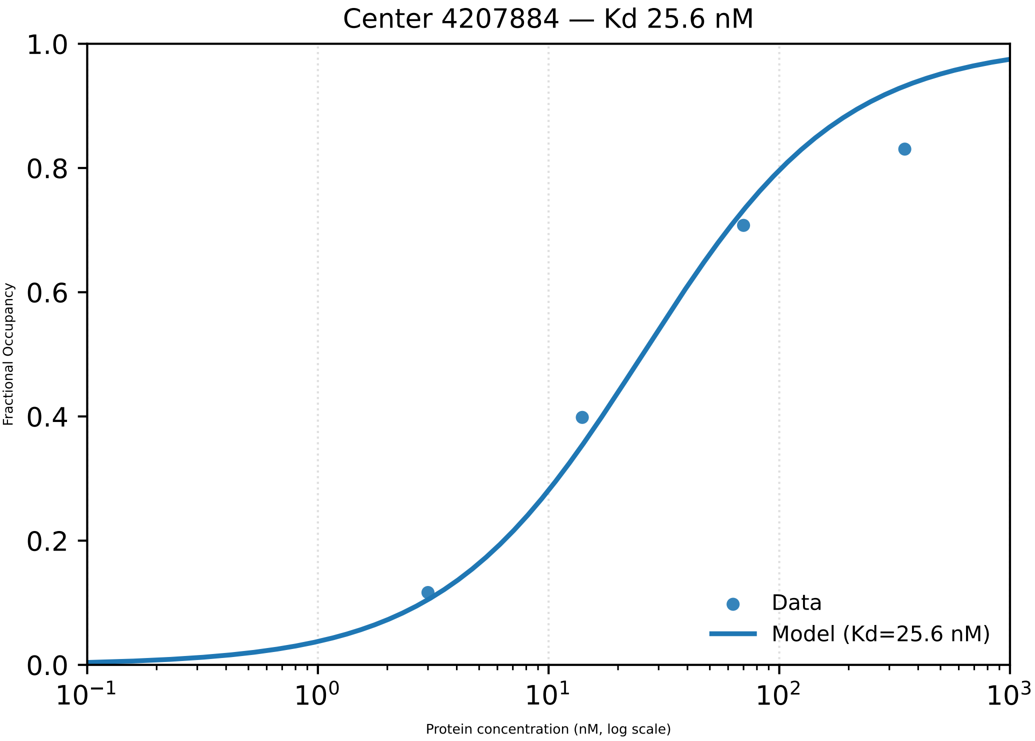

Center 4207884 — Kd 25.6 nM
1.0
0.8
0.6
0.4
0.2
0.0
Fractional Occupancy
$10^{-1}$
$10^{0}$
$10^{1}$
$10^{2}$
$10^{3}$
Protein concentration (nM, log scale)
Data
Model (Kd=25.6 nM)

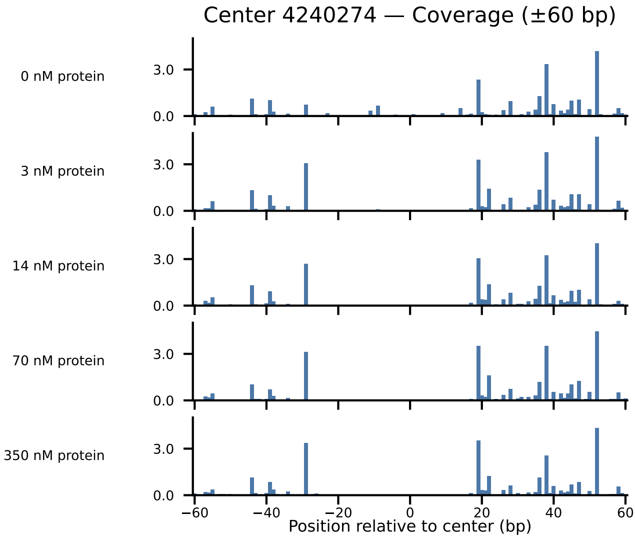

Center 4240274 — Coverage (±60 bp)
0 nM protein
3 nM protein
14 nM protein
70 nM protein
350 nM protein
3.0
0.0
−60
−40
−20
0
20
40
60
Position relative to center (bp)

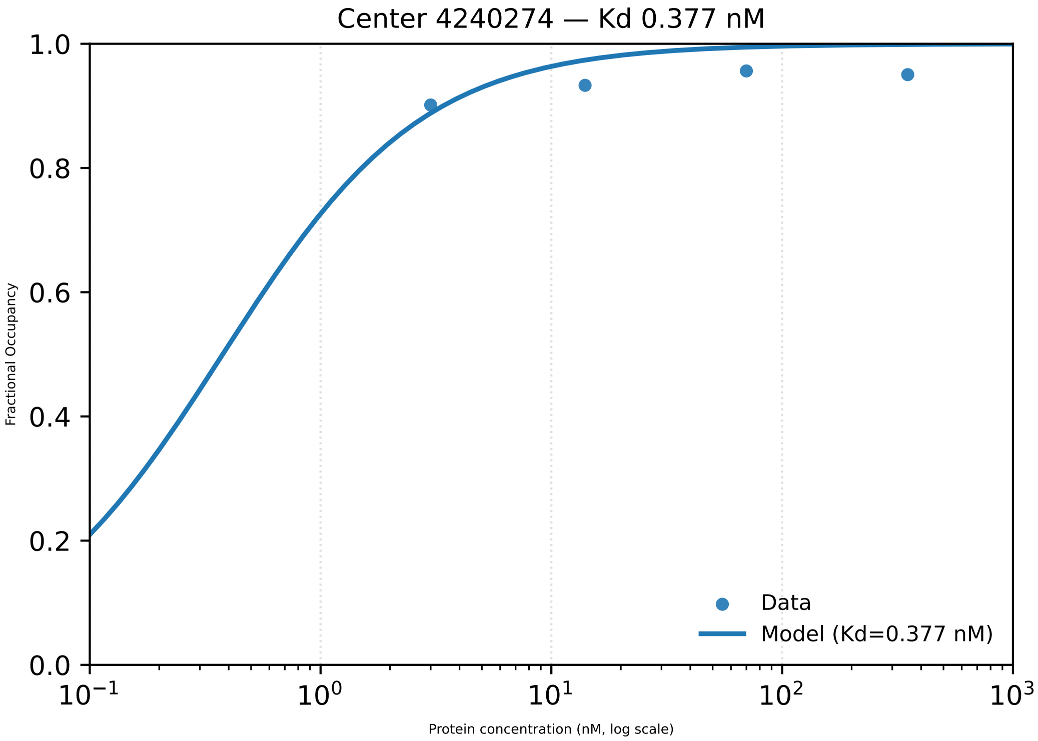

Center 4240274 — Kd 0.377 nM
Fractional Occupancy
1.0
0.8
0.6
0.4
0.2
0.0
$10^{-1}$
$10^{0}$
$10^{1}$
$10^{2}$
$10^{3}$
Protein concentration (nM, log scale)
Data
Model (Kd=0.377 nM)

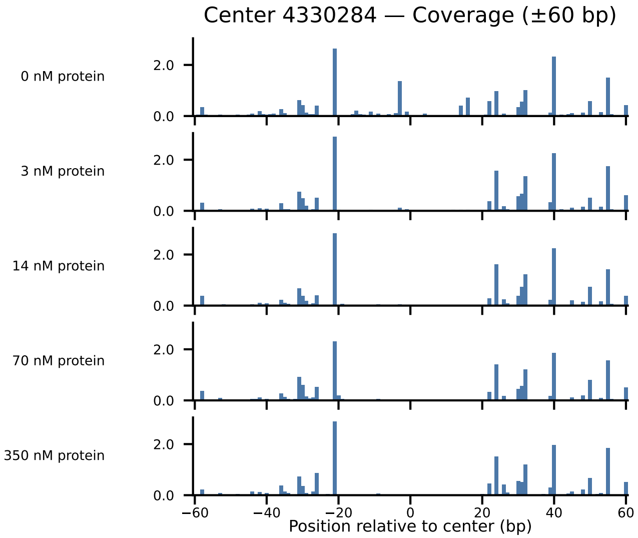

Center 4330284 — Coverage (±60 bp)
0 nM protein
3 nM protein
14 nM protein
70 nM protein
350 nM protein
2.0
0.0
−60
−40
−20
0
20
40
60
Position relative to center (bp)

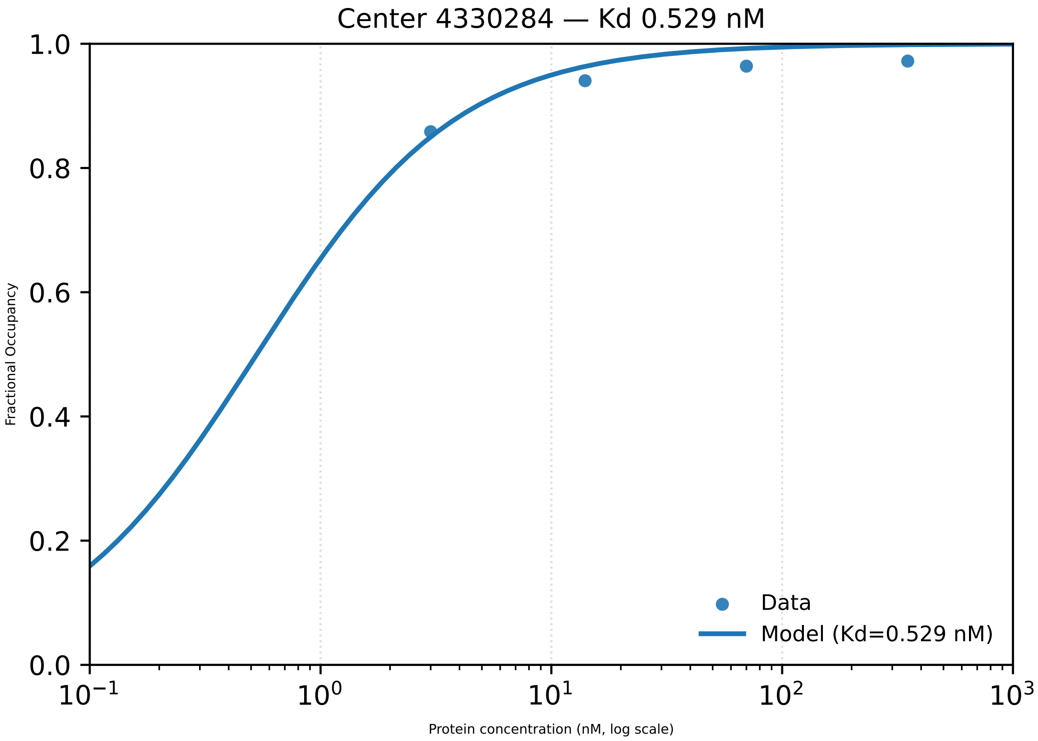

Center 4330284 — Kd 0.529 nM
Fractional Occupancy
1.0
0.8
0.6
0.4
0.2
0.0
$10^{-1}$
$10^{0}$
$10^{1}$
$10^{2}$
$10^{3}$
Protein concentration (nM, log scale)
Data
Model (Kd=0.529 nM)

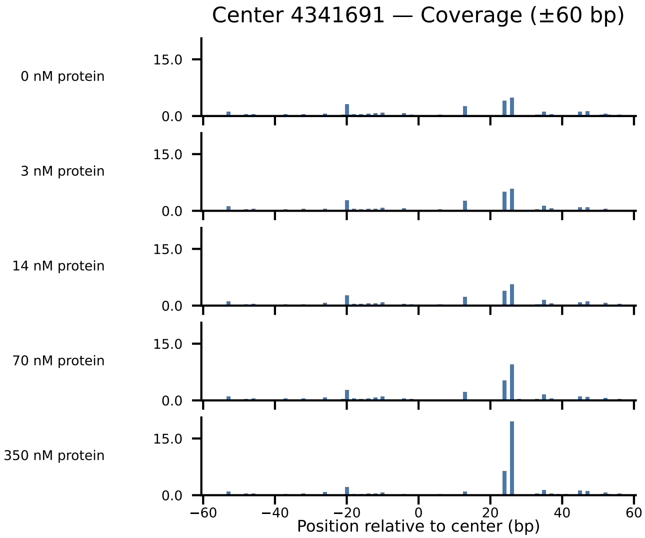

Center 4341691 — Coverage (±60 bp)
0 nM protein
3 nM protein
14 nM protein
70 nM protein
350 nM protein
15.0
0.0
−60
−40
−20
0
20
40
60
Position relative to center (bp)

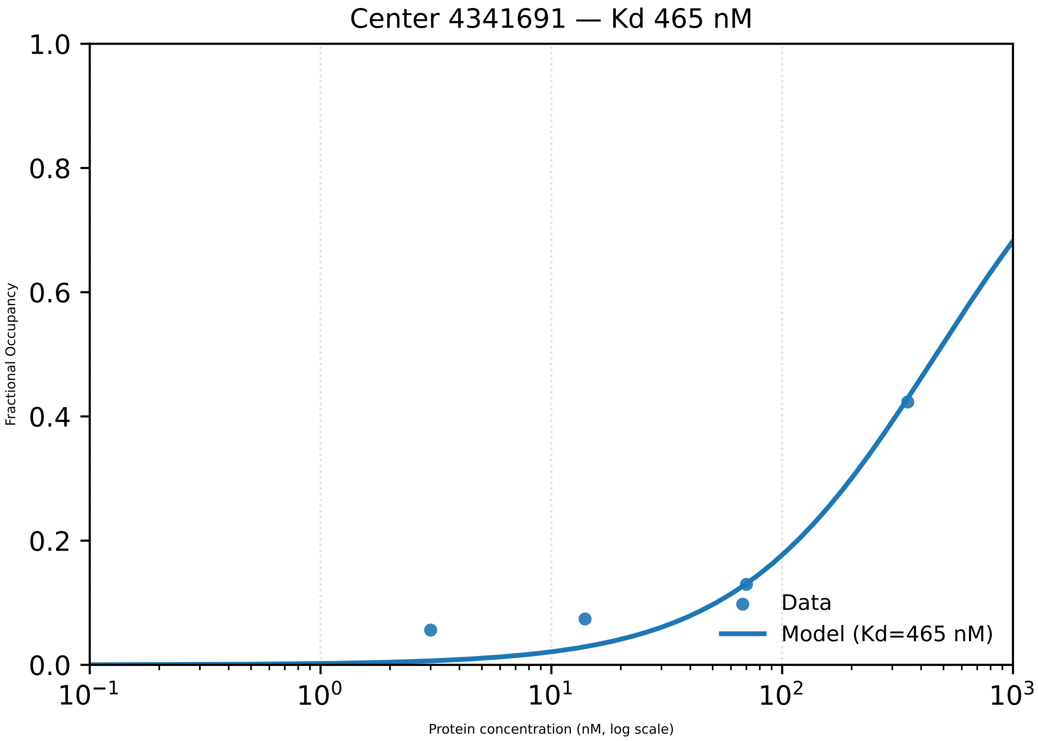

Center 4341691 — Kd 465 nM
1.0
0.8
0.6
0.4
0.2
0.0
Fractional Occupancy
$10^{-1}$
$10^{0}$
$10^{1}$
$10^{2}$
$10^{3}$
Protein concentration (nM, log scale)
Data
Model (Kd=465 nM)

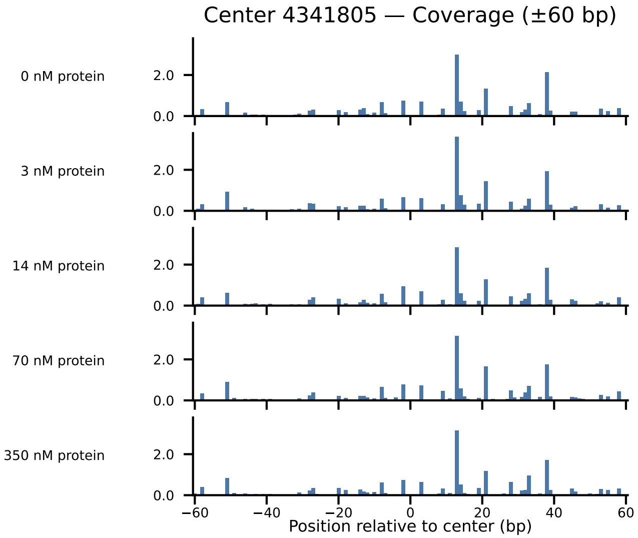

Center 4341805 — Coverage (±60 bp)
0 nM protein
3 nM protein
14 nM protein
70 nM protein
350 nM protein
2.0
0.0
−60
−40
−20
0
20
40
60
Position relative to center (bp)

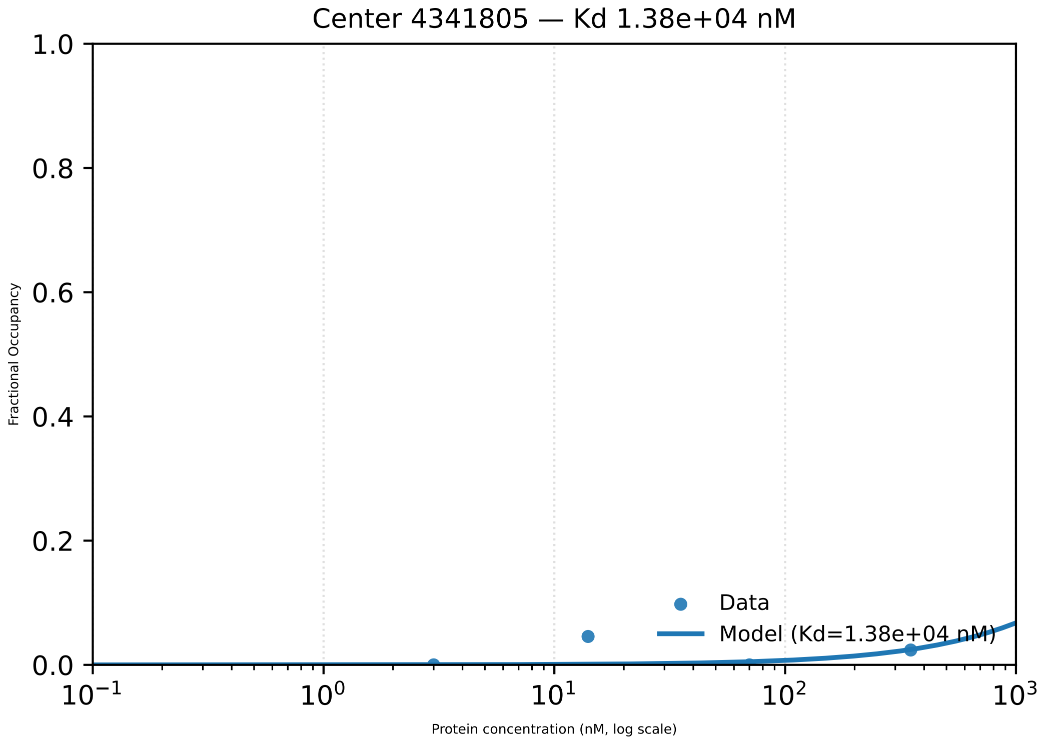

Center 4341805 — Kd 1.38e+04 nM
1.0
0.8
0.6
0.4
0.2
0.0
Fractional Occupancy
$10^{-1}$
$10^{0}$
$10^{1}$
$10^{2}$
$10^{3}$
Protein concentration (nM, log scale)
Data
Model (Kd=1.38e+04 nM)

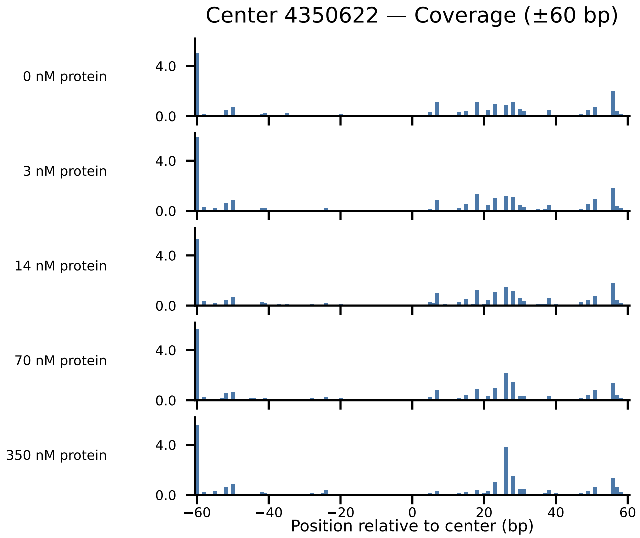
Center 4350622 — Coverage (±60 bp)
0 nM protein
3 nM protein
14 nM protein
70 nM protein
350 nM protein
4.0
0.0
−60
−40
−20
0
20
40
60
Position relative to center (bp)

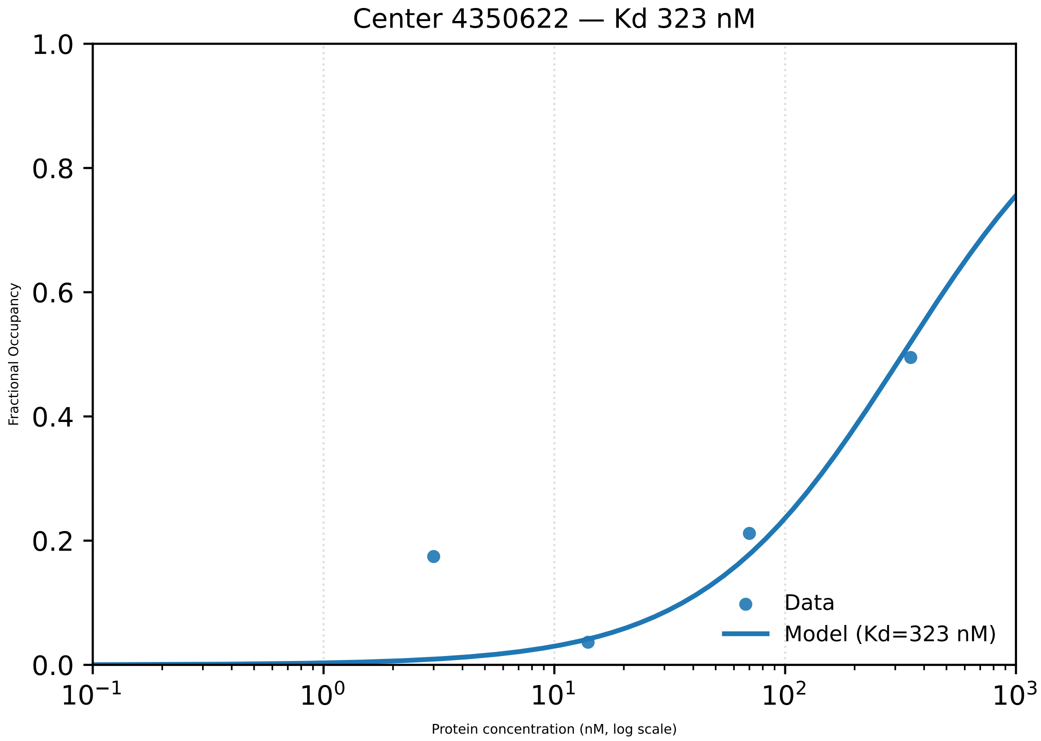

Center 4350622 — Kd 323 nM
1.0
0.8
0.6
0.4
0.2
0.0
Fractional Occupancy
$10^{-1}$
$10^{0}$
$10^{1}$
$10^{2}$
$10^{3}$
Protein concentration (nM, log scale)
Data
Model (Kd=323 nM)

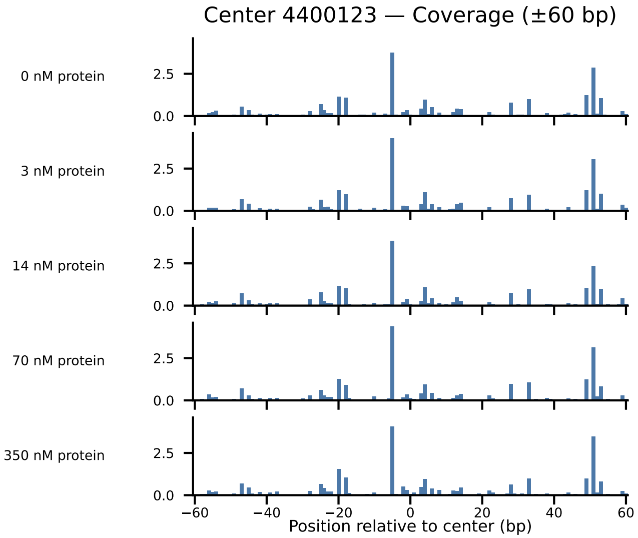

Center 4400123 — Coverage (±60 bp)
0 nM protein
3 nM protein
14 nM protein
70 nM protein
350 nM protein
2.5
0.0
−60
−40
−20
0
20
40
60
Position relative to center (bp)

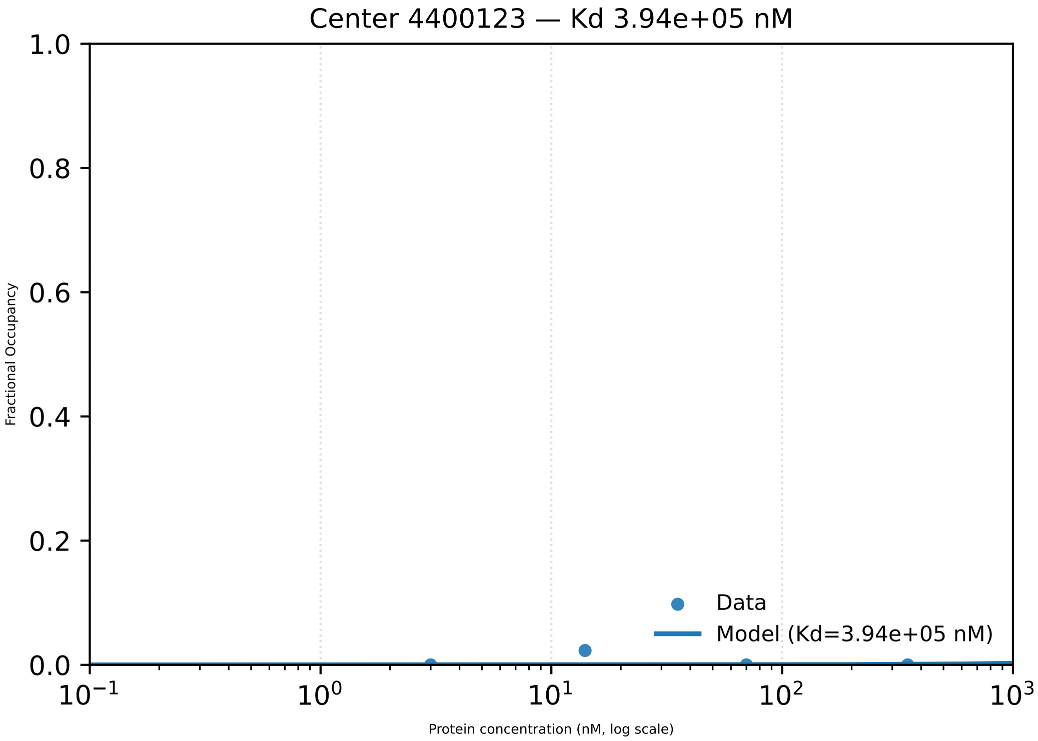

Center 4400123 — Kd 3.94e+05 nM
1.0
0.8
0.6
0.4
0.2
0.0
Fractional Occupancy
Data
Model (Kd=3.94e+05 nM)
$10^{-1}$
$10^{0}$
$10^{1}$
$10^{2}$
$10^{3}$
Protein concentration (nM, log scale)

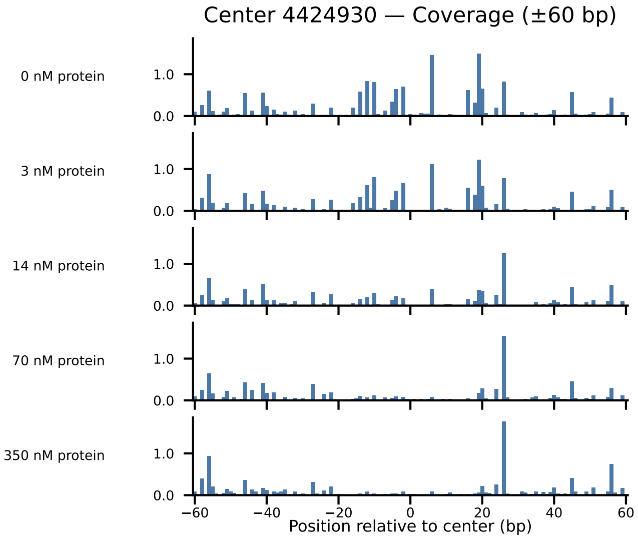

Center 4424930 — Coverage (±60 bp)
0 nM protein
3 nM protein
14 nM protein
70 nM protein
350 nM protein
1.0
0.0
−60
−40
−20
0
20
40
60
Position relative to center (bp)

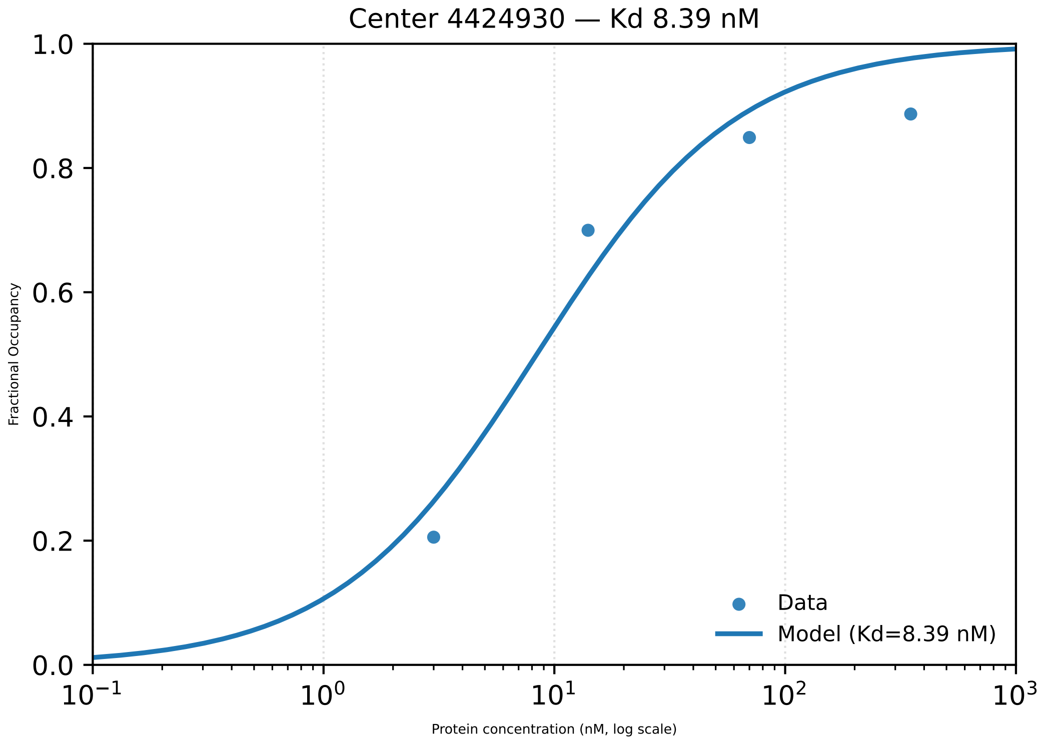

Center 4424930 — Kd 8.39 nM
Fractional Occupancy
1.0
0.8
0.6
0.4
0.2
0.0
$10^{-1}$
$10^{0}$
$10^{1}$
$10^{2}$
$10^{3}$
Protein concentration (nM, log scale)
Data
Model (Kd=8.39 nM)

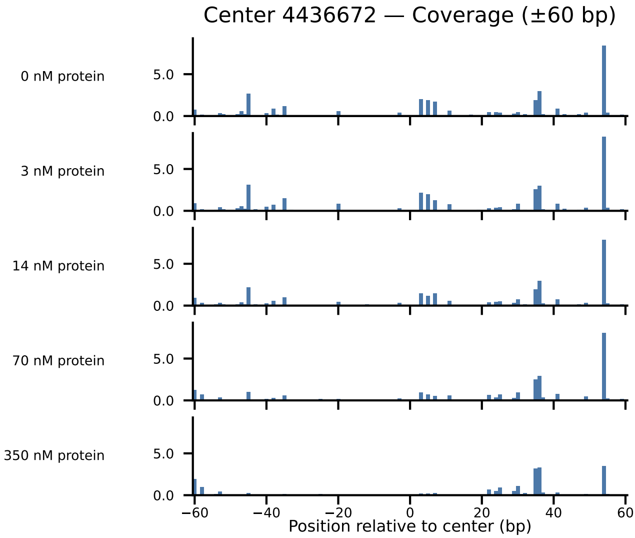

Center 4436672 — Coverage (±60 bp)
0 nM protein
3 nM protein
14 nM protein
70 nM protein
350 nM protein
5.0
0.0
−60
−40
−20
0
20
40
60
Position relative to center (bp)

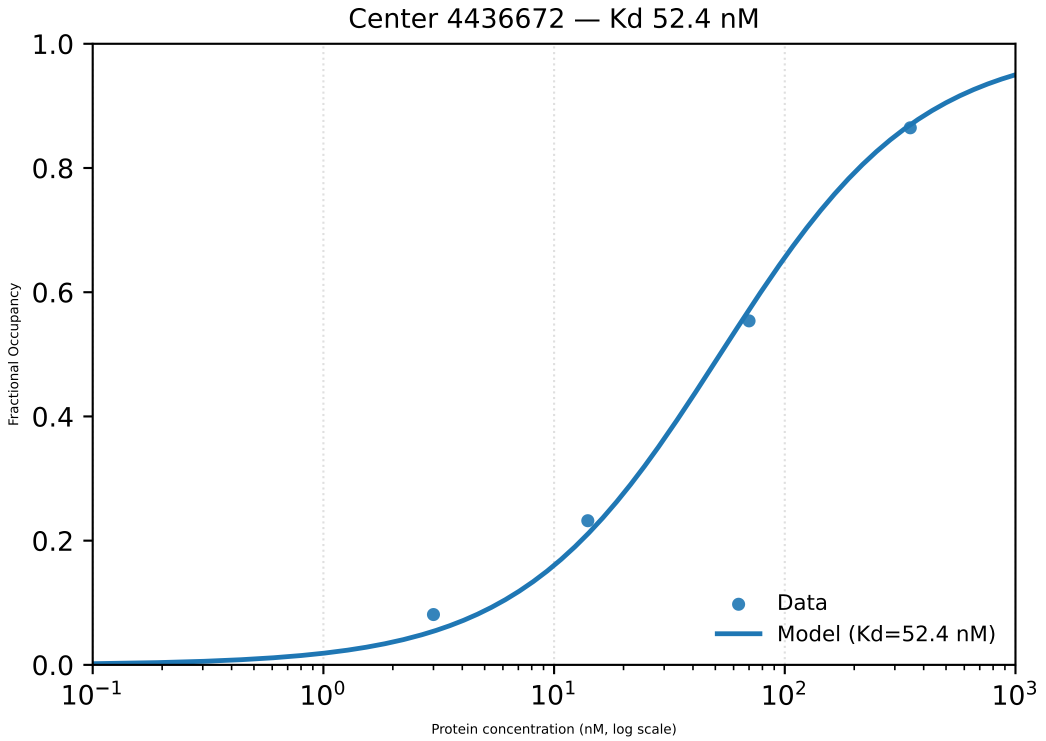

Center 4436672 — Kd 52.4 nM
1.0
0.8
0.6
0.4
0.2
0.0
Fractional Occupancy
$10^{-1}$
$10^{0}$
$10^{1}$
$10^{2}$
$10^{3}$
Protein concentration (nM, log scale)
Data
Model (Kd=52.4 nM)

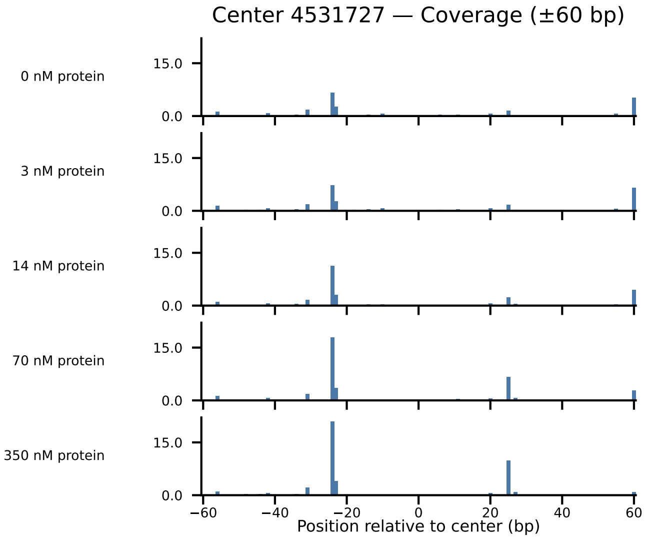

Center 4531727 — Coverage (±60 bp)
0 nM protein
3 nM protein
14 nM protein
70 nM protein
350 nM protein
15.0
0.0
−60
−40
−20
0
20
40
60
Position relative to center (bp)

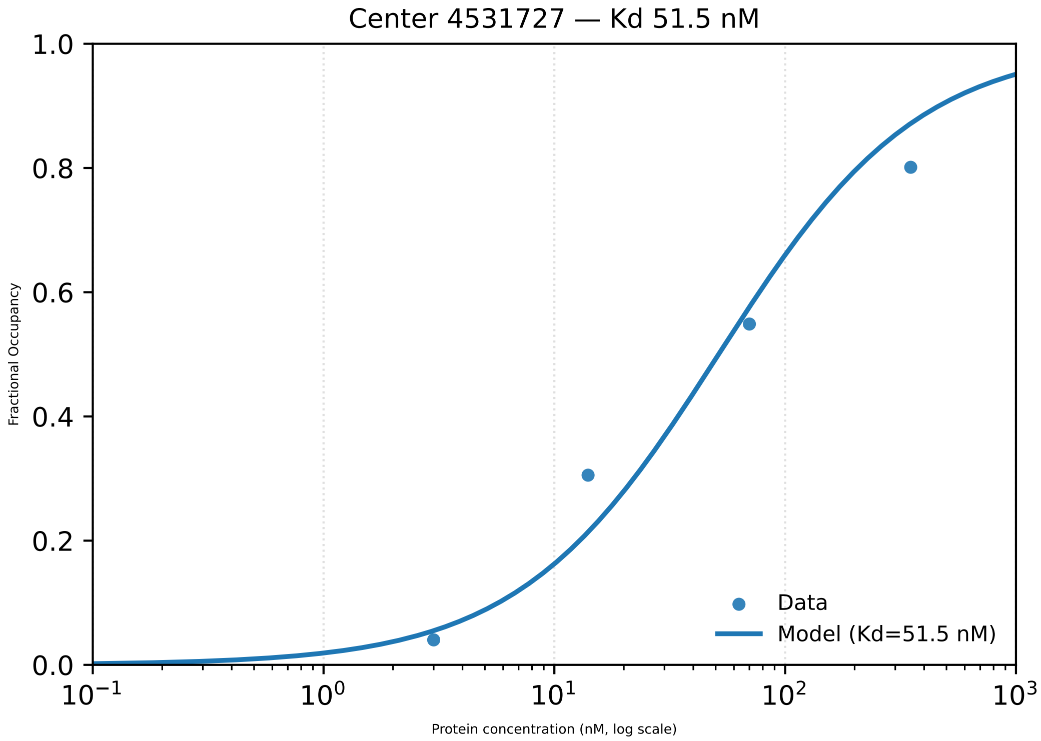

Center 4531727 — Kd 51.5 nM
Fractional Occupancy
1.0
0.8
0.6
0.4
0.2
0.0
$10^{-1}$
$10^{0}$
$10^{1}$
$10^{2}$
$10^{3}$
Protein concentration (nM, log scale)
Data
Model (Kd=51.5 nM)

### Center 4539989 — Coverage (±60 bp)

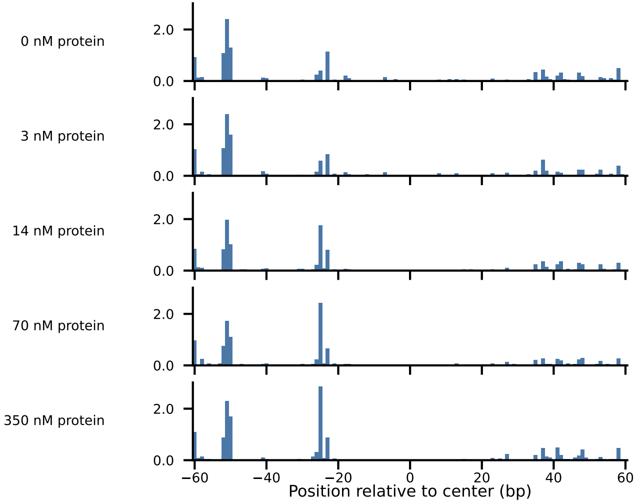

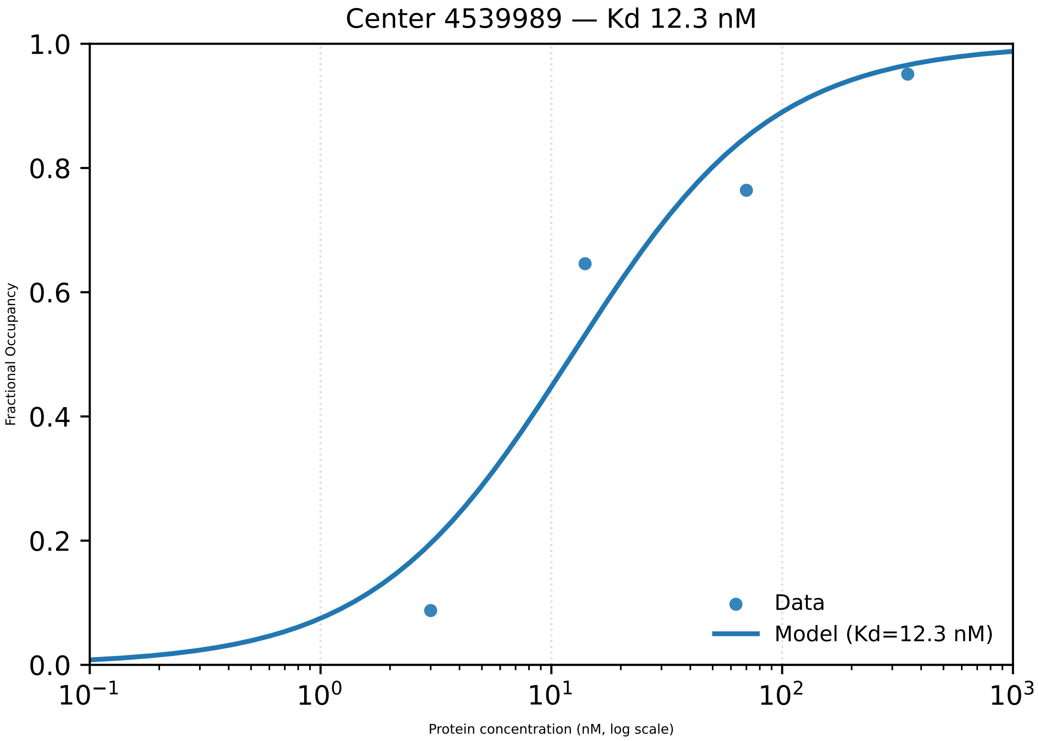

Center 4539989 — Kd 12.3 nM
Fractional Occupancy
1.0
0.8
0.6
0.4
0.2
0.0
$10^{-1}$
$10^{0}$
$10^{1}$
$10^{2}$
$10^{3}$
Protein concentration (nM, log scale)
Data
Model (Kd=12.3 nM)

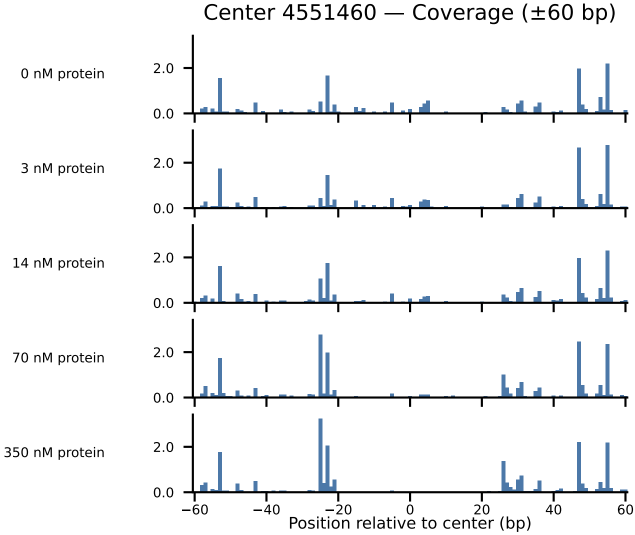

Center 4551460 — Coverage (±60 bp)
0 nM protein
3 nM protein
14 nM protein
70 nM protein
350 nM protein
2.0
0.0
−60
−40
−20
0
20
40
60
Position relative to center (bp)

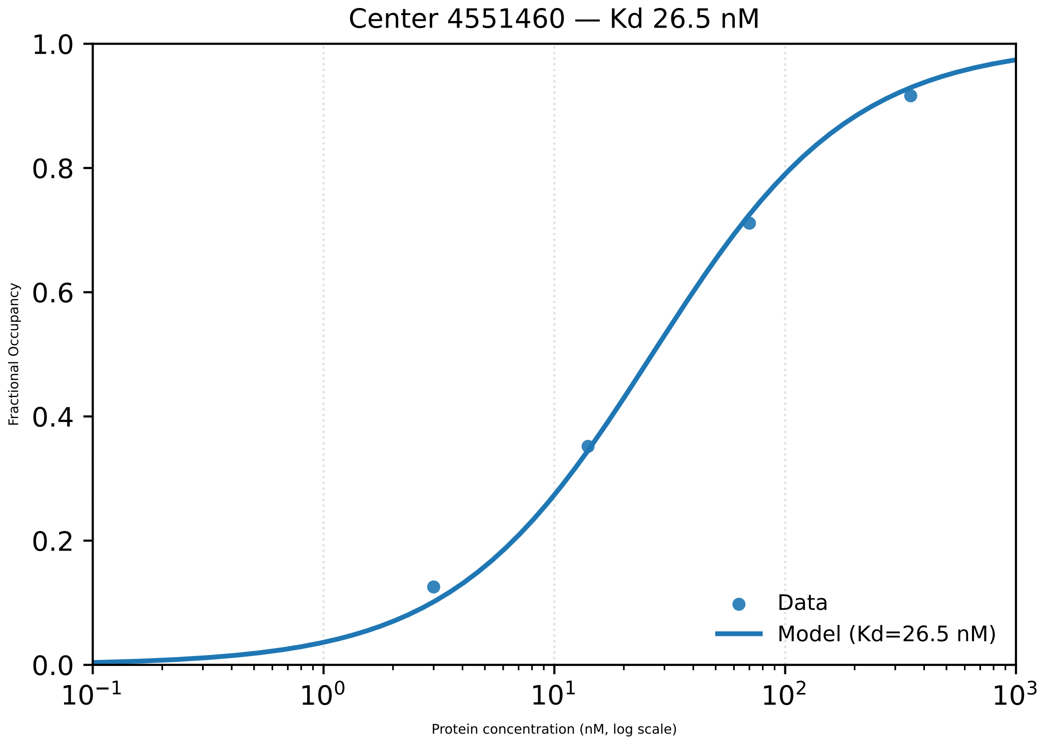

Center 4551460 — Kd 26.5 nM
Fractional Occupancy
1.0
0.8
0.6
0.4
0.2
0.0
$10^{-1}$
$10^{0}$
$10^{1}$
$10^{2}$
$10^{3}$
Protein concentration (nM, log scale)
Data
Model (Kd=26.5 nM)

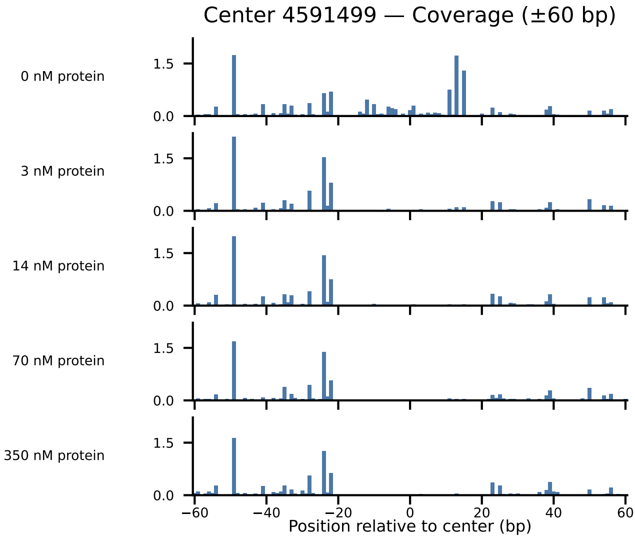
Center 4591499 — Coverage (±60 bp)
0 nM protein
3 nM protein
14 nM protein
70 nM protein
350 nM protein
1.5
0.0
−60
−40
−20
0
20
40
60
Position relative to center (bp)

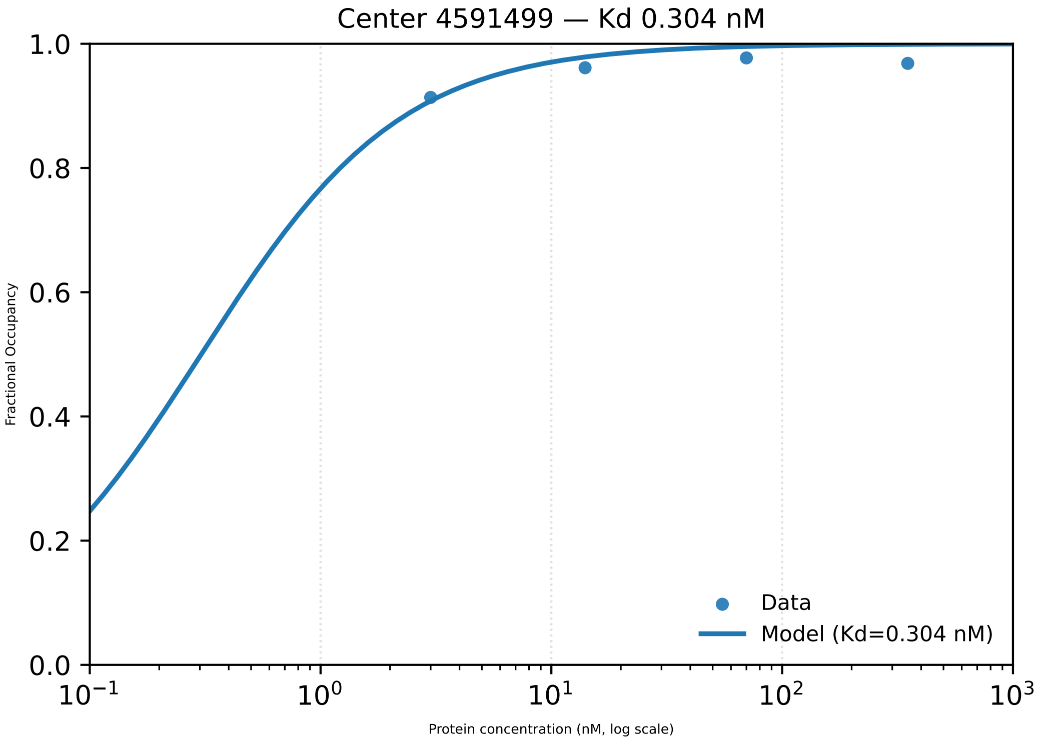
Center 4591499 — Kd 0.304 nM
Fractional Occupancy
Protein concentration (nM, log scale)
Data
Model (Kd=0.304 nM)
0.0
0.2
0.4
0.6
0.8
1.0
$10^{-1}$
$10^{0}$
$10^{1}$
$10^{2}$
$10^{3}$
