## Supplementary material for "Footprint-seq: a simple method to quantitatively map *in vitro* protein-DNA interactions on a genome-wide scale at high spatial resolution": Figure S5

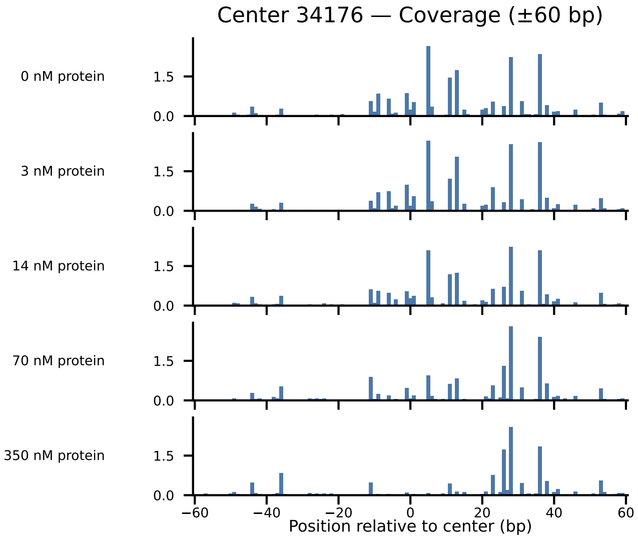
Center 34176 — Coverage (±60 bp)
0 nM protein
3 nM protein
14 nM protein
70 nM protein
350 nM protein
1.5
0.0
−60
−40
−20
0
20
40
60
Position relative to center (bp)

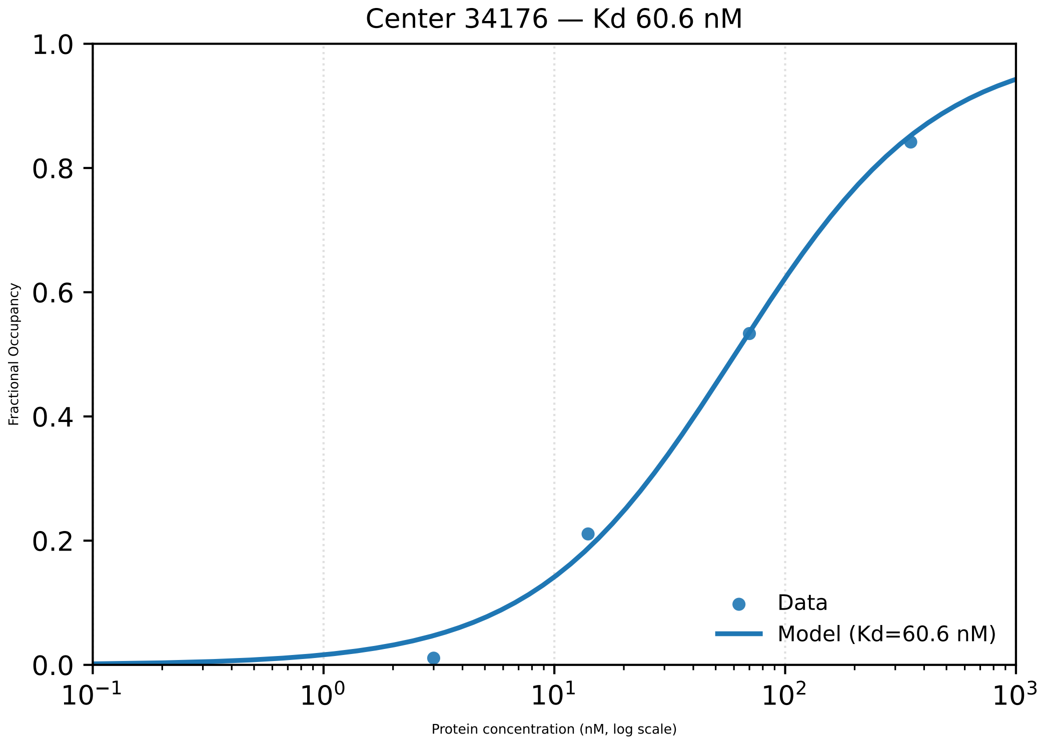

Center 34176 — Kd 60.6 nM
Fractional Occupancy
1.0
0.8
0.6
0.4
0.2
0.0
$10^{-1}$
$10^{0}$
$10^{1}$
$10^{2}$
$10^{3}$
Protein concentration (nM, log scale)
Data
Model (Kd=60.6 nM)

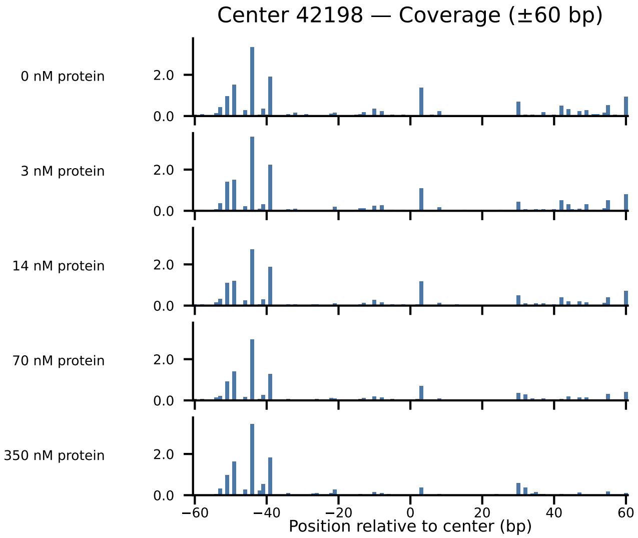

Center 42198 — Coverage (±60 bp)
0 nM protein
3 nM protein
14 nM protein
70 nM protein
350 nM protein
2.0
0.0
−60
−40
−20
0
20
40
60
Position relative to center (bp)

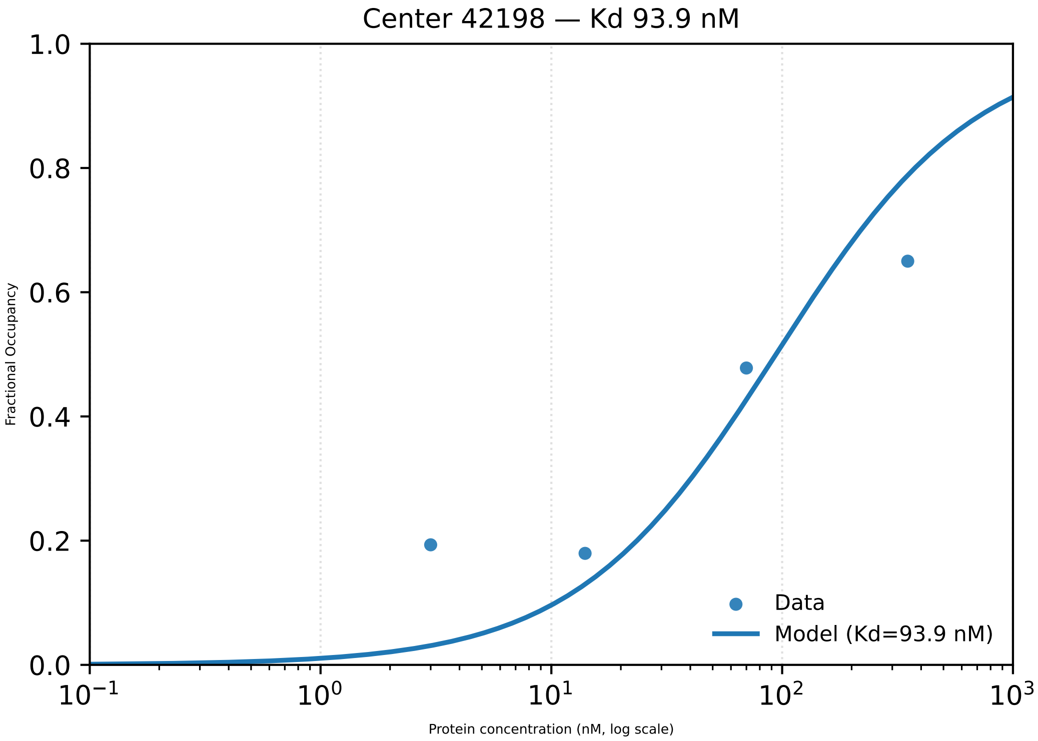

Center 42198 — Kd 93.9 nM
1.0
0.8
0.6
0.4
0.2
0.0
Fractional Occupancy
Data
Model (Kd=93.9 nM)
$10^{-1}$
$10^{0}$
$10^{1}$
$10^{2}$
$10^{3}$
Protein concentration (nM, log scale)

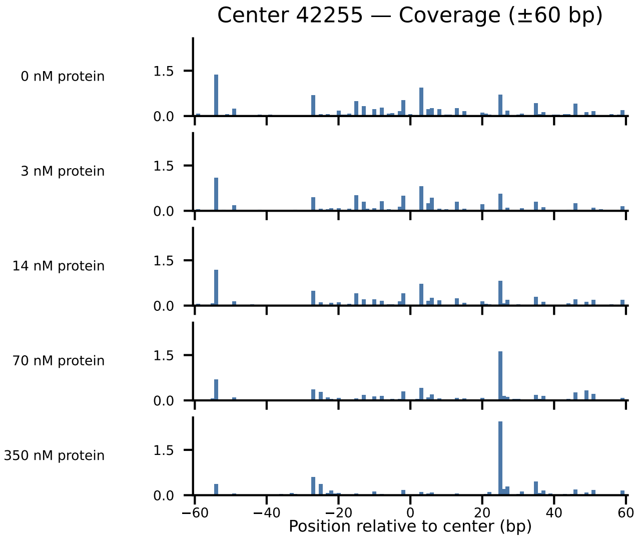
Center 42255 — Coverage (±60 bp)
0 nM protein
3 nM protein
14 nM protein
70 nM protein
350 nM protein
1.5
0.0
−60
−40
−20
0
20
40
60
Position relative to center (bp)

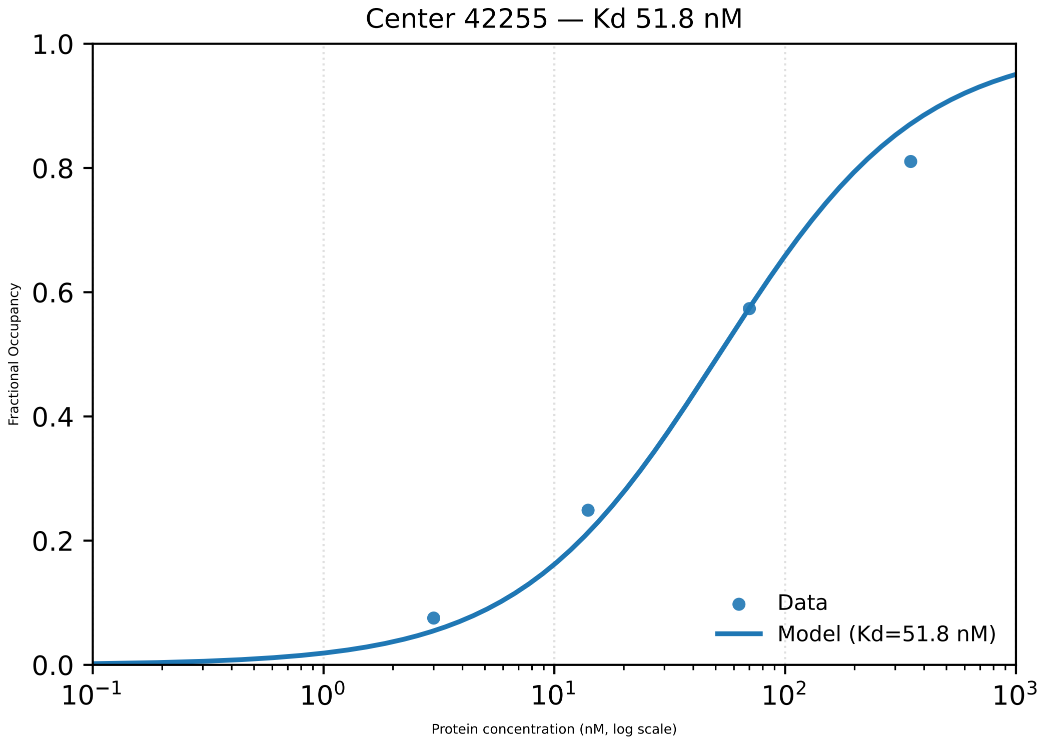

Center 42255 — Kd 51.8 nM
1.0
0.8
0.6
0.4
0.2
0.0
Fractional Occupancy
10−1
100
101
102
103
Protein concentration (nM, log scale)
Data
Model (Kd=51.8 nM)

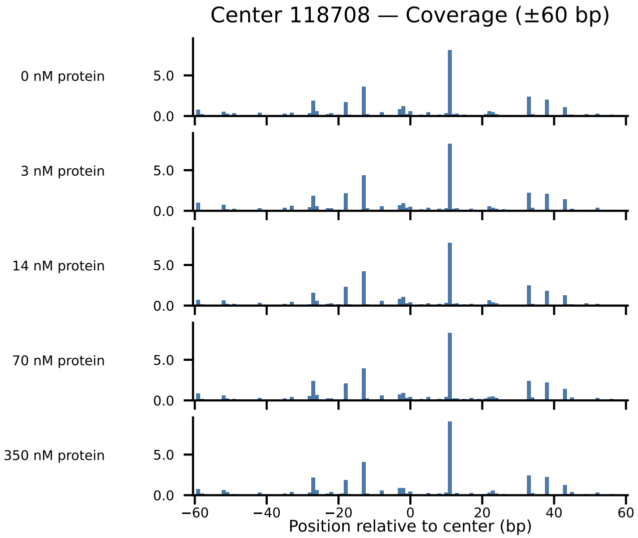
Center 118708 — Coverage (±60 bp)
0 nM protein
3 nM protein
14 nM protein
70 nM protein
350 nM protein
5.0
0.0
−60
−40
−20
0
20
40
60
Position relative to center (bp)

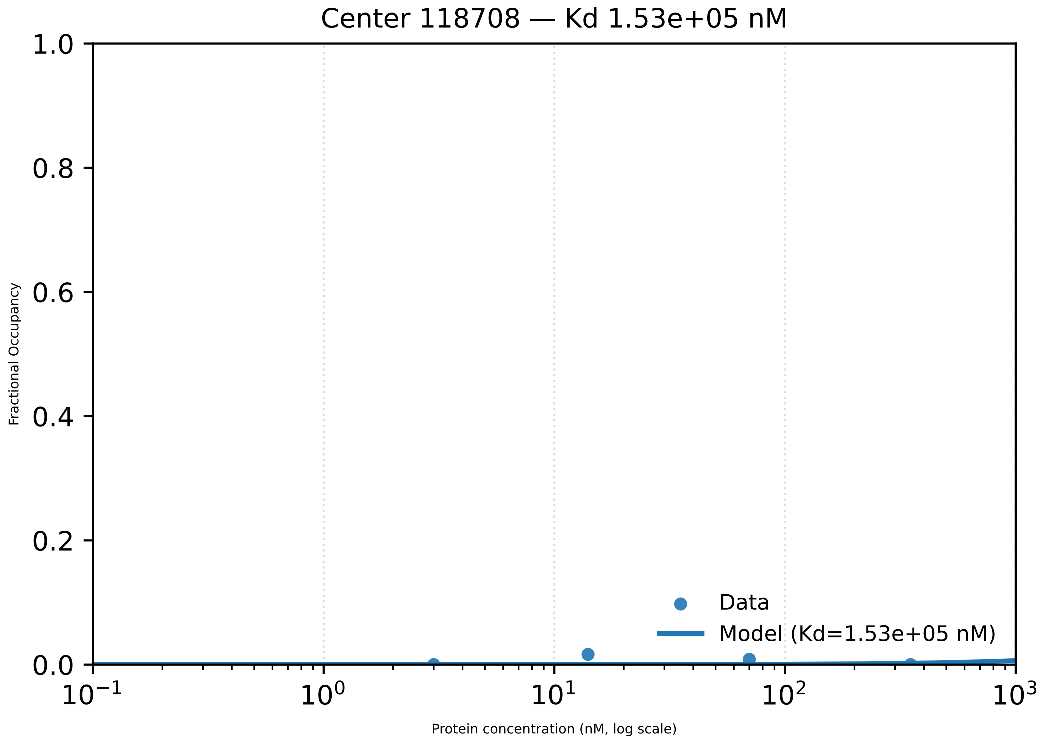
Center 118708 — Kd 1.53e+05 nM
1.0
0.8
0.6
0.4
0.2
0.0
Fractional Occupancy
$10^{-1}$
$10^{0}$
$10^{1}$
$10^{2}$
$10^{3}$
Protein concentration (nM, log scale)
Data
Model (Kd=1.53e+05 nM)

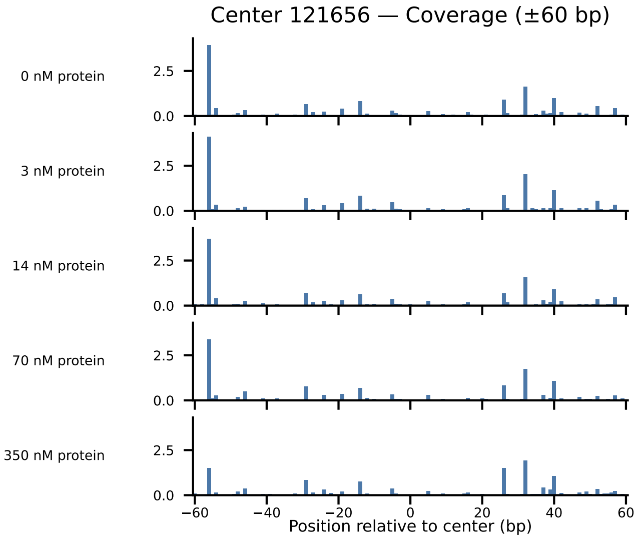

Center 121656 — Coverage (±60 bp)
0 nM protein
3 nM protein
14 nM protein
70 nM protein
350 nM protein
2.5
0.0
−60
−40
−20
0
20
40
60
Position relative to center (bp)

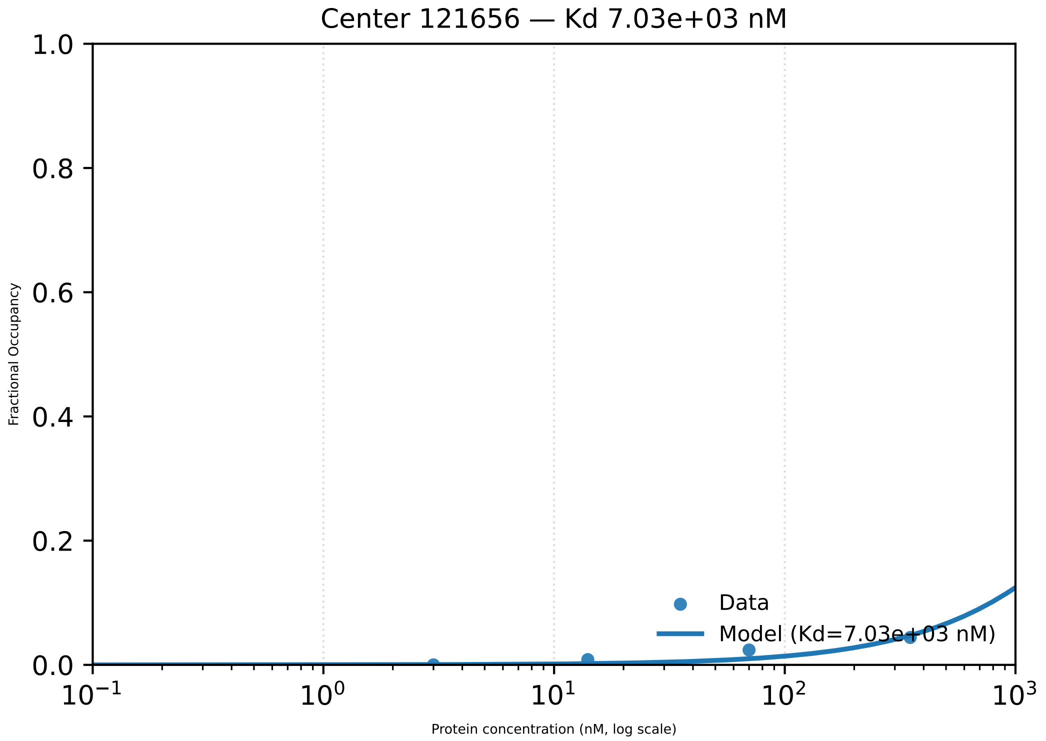
Center 121656 — Kd 7.03e+03 nM
1.0
0.8
0.6
0.4
0.2
0.0
Fractional Occupancy
$10^{-1}$
$10^{0}$
$10^{1}$
$10^{2}$
$10^{3}$
Protein concentration (nM, log scale)
Data
Model (Kd=7.03e+03 nM)

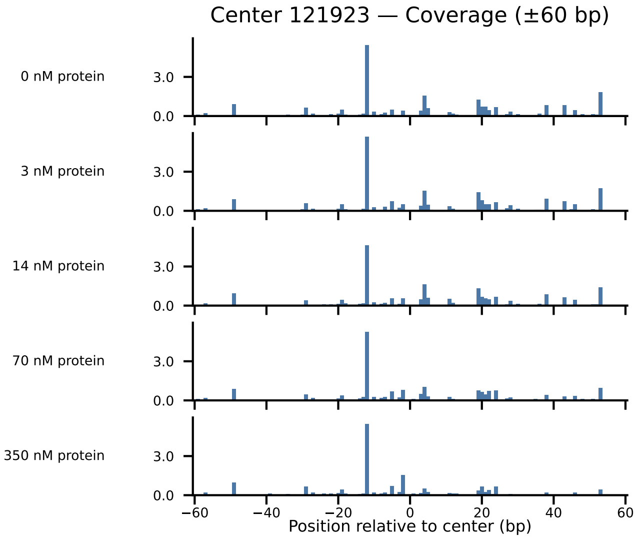
Center 121923 — Coverage (±60 bp)
0 nM protein
3 nM protein
14 nM protein
70 nM protein
350 nM protein
3.0
0.0
−60
−40
−20
0
20
40
60
Position relative to center (bp)

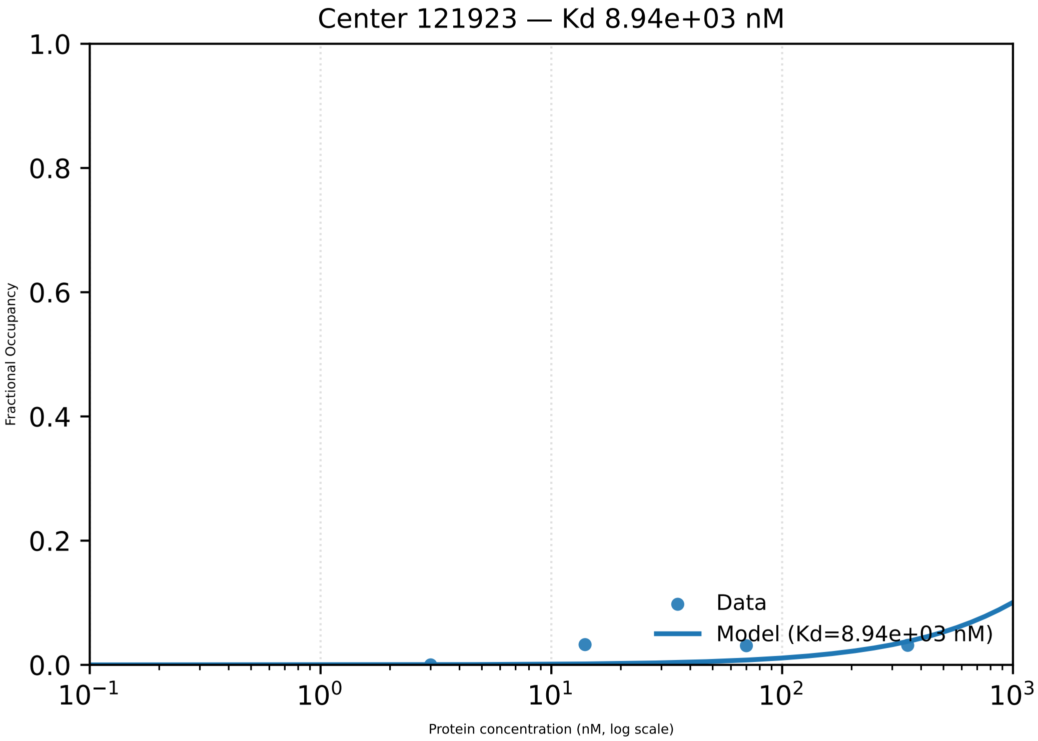

Center 121923 — Kd 8.94e+03 nM
1.0
0.8
0.6
0.4
0.2
0.0
Fractional Occupancy
Data
Model (Kd=8.94e+03 nM)
$10^{-1}$
$10^{0}$
$10^{1}$
$10^{2}$
$10^{3}$
Protein concentration (nM, log scale)

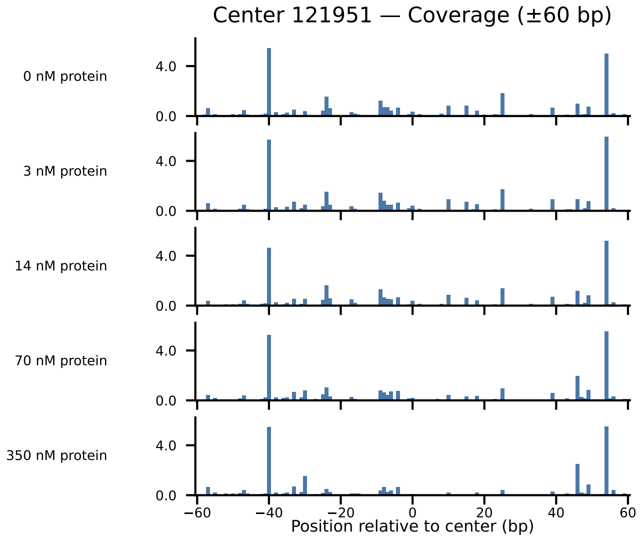

Center 121951 — Coverage (±60 bp)
0 nM protein
3 nM protein
14 nM protein
70 nM protein
350 nM protein
4.0
0.0
−60
−40
−20
0
20
40
60
Position relative to center (bp)

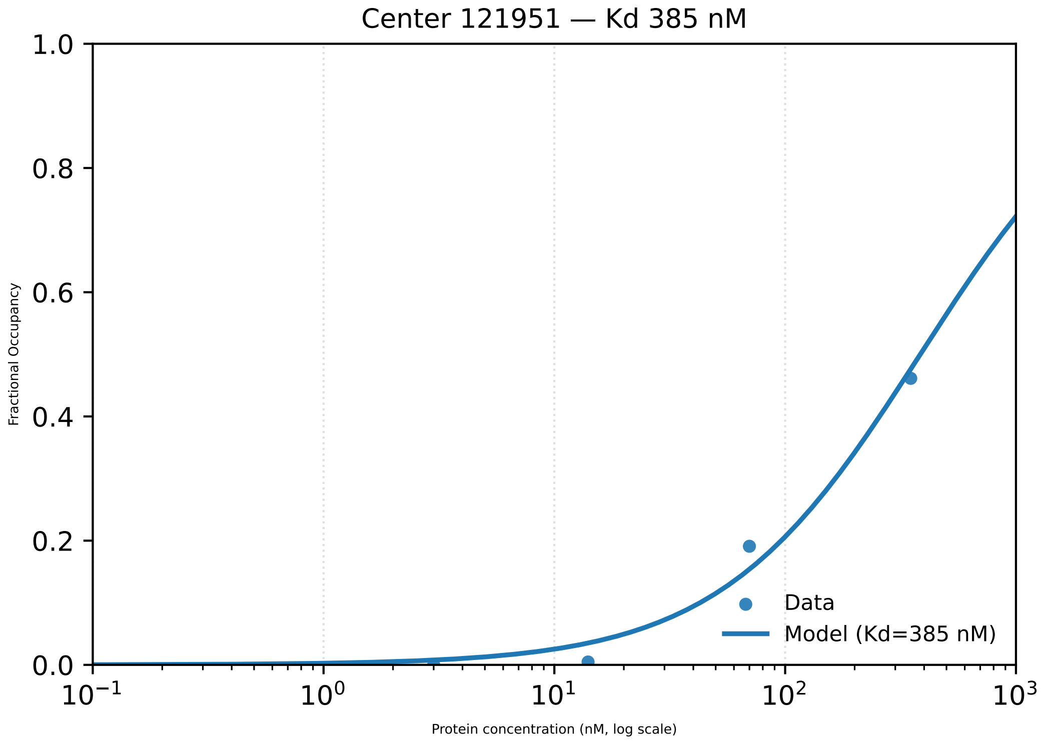

Center 121951 — Kd 385 nM
1.0
0.8
0.6
0.4
0.2
0.0
Fractional Occupancy
$10^{-1}$
$10^{0}$
$10^{1}$
$10^{2}$
$10^{3}$
Protein concentration (nM, log scale)
Data
Model (Kd=385 nM)

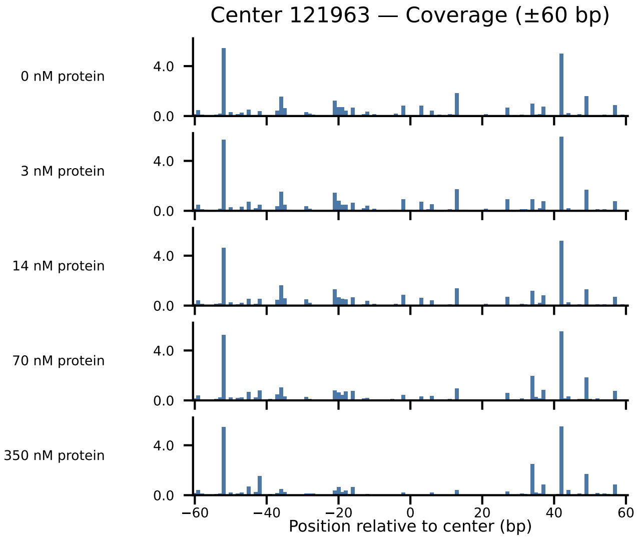
Center 121963 — Coverage (±60 bp)
0 nM protein
3 nM protein
14 nM protein
70 nM protein
350 nM protein
4.0
0.0
−60
−40
−20
0
20
40
60
Position relative to center (bp)

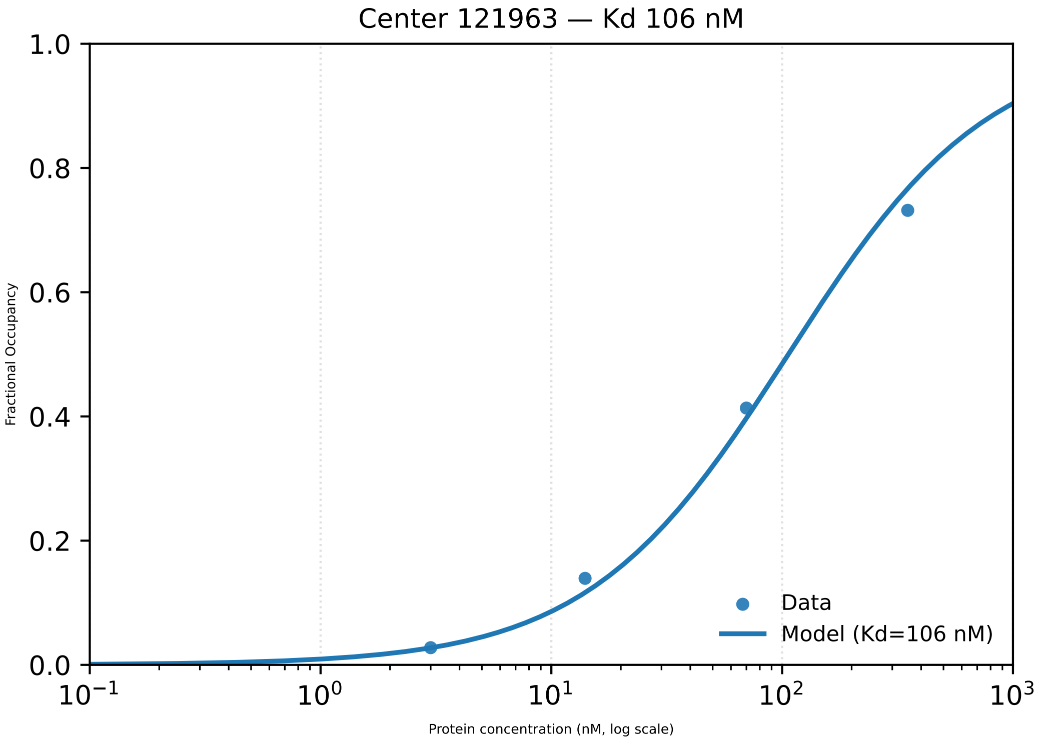

Center 121963 — Kd 106 nM
1.0
0.8
0.6
0.4
0.2
0.0
Fractional Occupancy
$10^{-1}$
$10^{0}$
$10^{1}$
$10^{2}$
$10^{3}$
Protein concentration (nM, log scale)
Data
Model (Kd=106 nM)

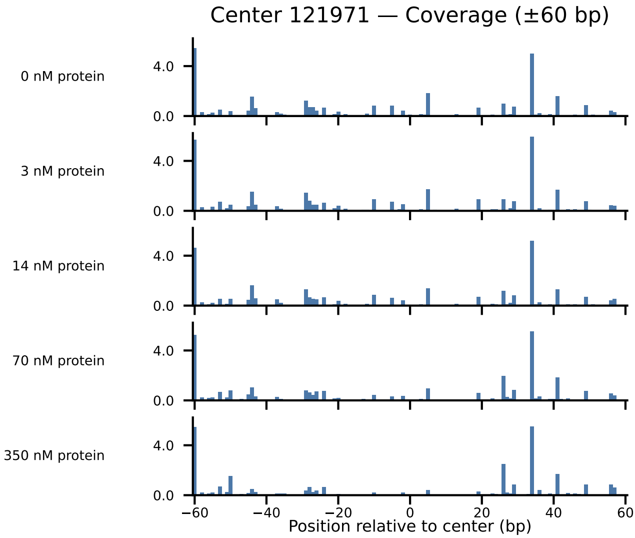
Center 121971 — Coverage (±60 bp)
0 nM protein
3 nM protein
14 nM protein
70 nM protein
350 nM protein
4.0
0.0
−60
−40
−20
0
20
40
60
Position relative to center (bp)

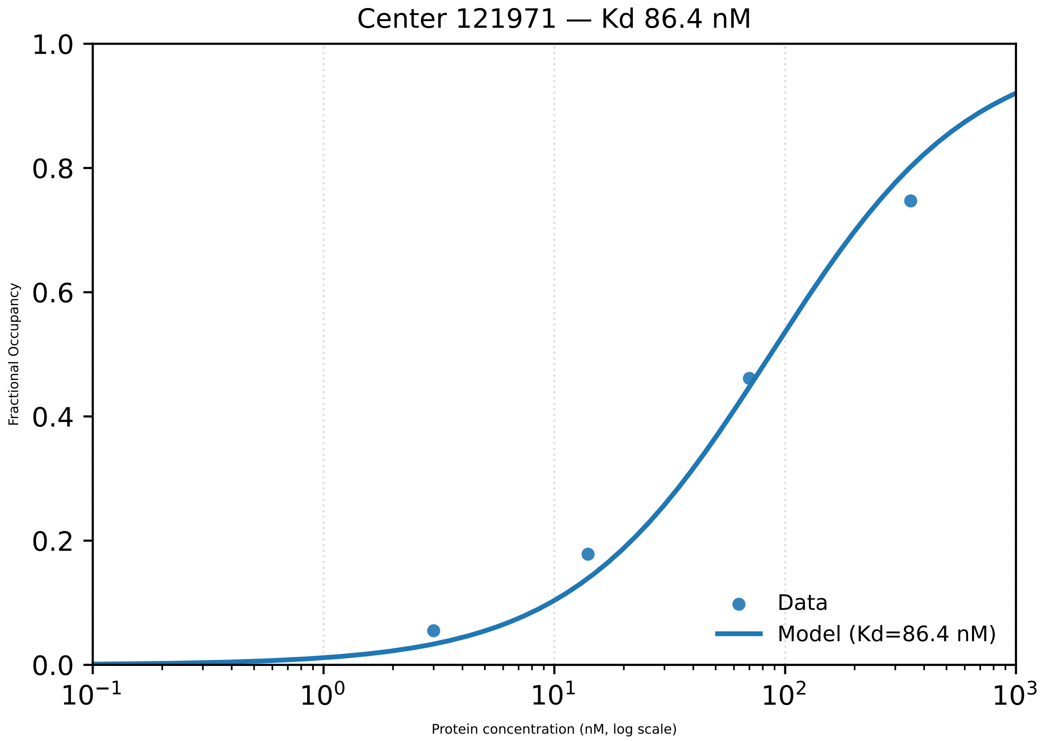

Center 121971 — Kd 86.4 nM
1.0
0.8
0.6
0.4
0.2
0.0
Fractional Occupancy
$10^{-1}$
$10^{0}$
$10^{1}$
$10^{2}$
$10^{3}$
Protein concentration (nM, log scale)
Data
Model (Kd=86.4 nM)

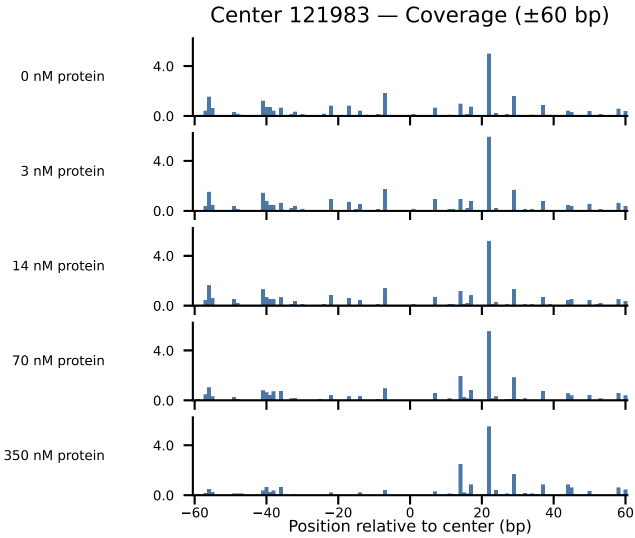
Center 121983 — Coverage (±60 bp)
0 nM protein
3 nM protein
14 nM protein
70 nM protein
350 nM protein
4.0
0.0
−60
−40
−20
0
20
40
60
Position relative to center (bp)

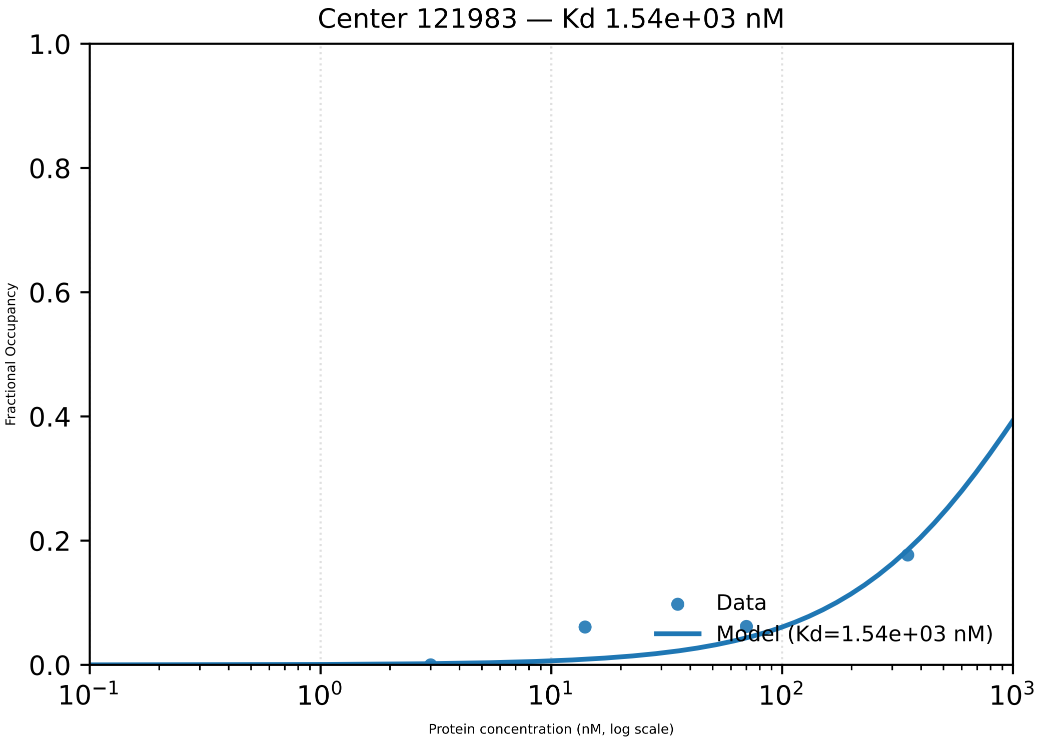
Center 121983 — Kd 1.54e+03 nM
Fractional Occupancy
1.0
0.8
0.6
0.4
0.2
0.0
$10^{-1}$
$10^{0}$
$10^{1}$
$10^{2}$
$10^{3}$
Protein concentration (nM, log scale)
Data
Model (Kd=1.54e+03 nM)

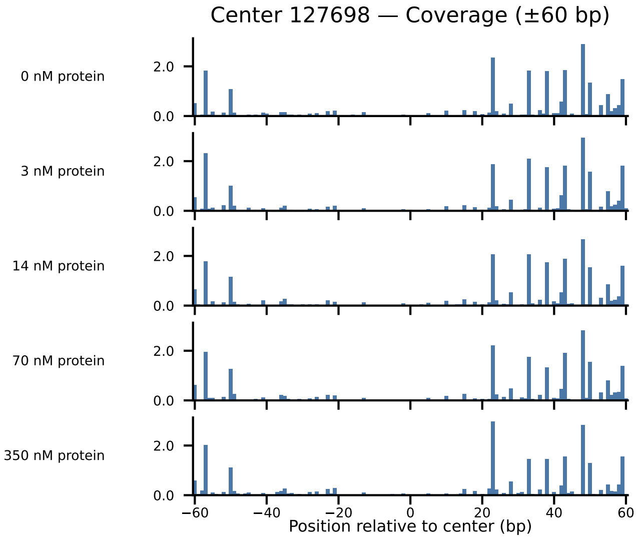
Center 127698 — Coverage (±60 bp)
0 nM protein
3 nM protein
14 nM protein
70 nM protein
350 nM protein
2.0
0.0
−60
−40
−20
0
20
40
60
Position relative to center (bp)

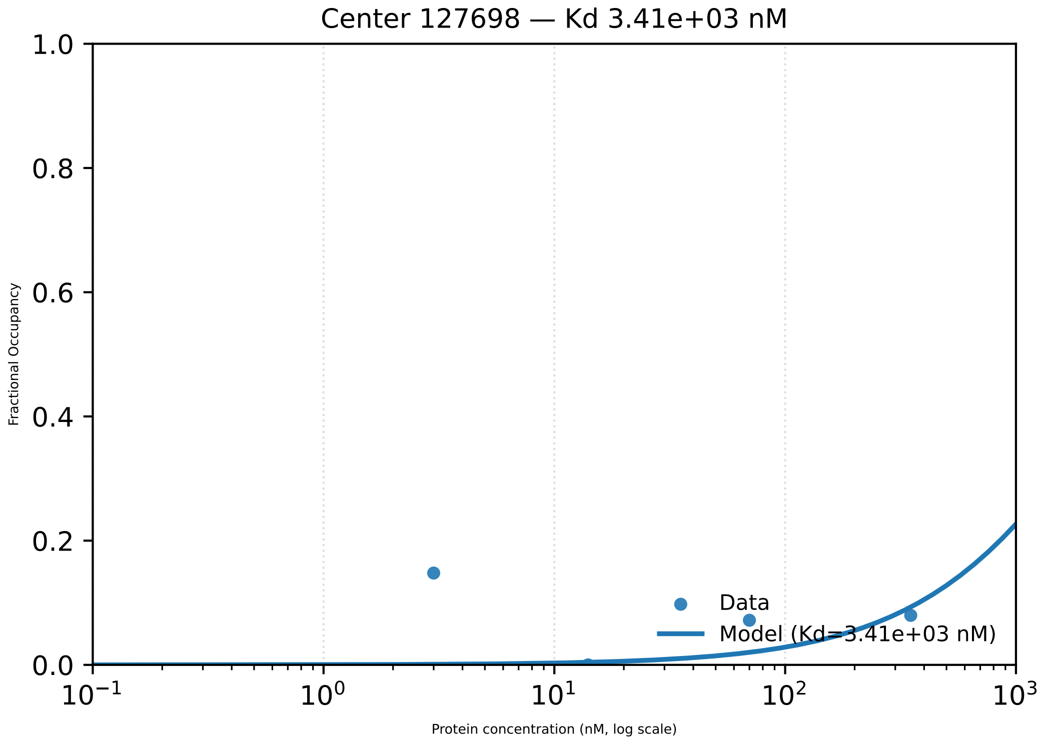

Center 127698 — Kd 3.41e+03 nM
1.0
0.8
0.6
0.4
0.2
0.0
Fractional Occupancy
Data
Model (Kd=3.41e+03 nM)
$10^{-1}$
$10^{0}$
$10^{1}$
$10^{2}$
$10^{3}$
Protein concentration (nM, log scale)

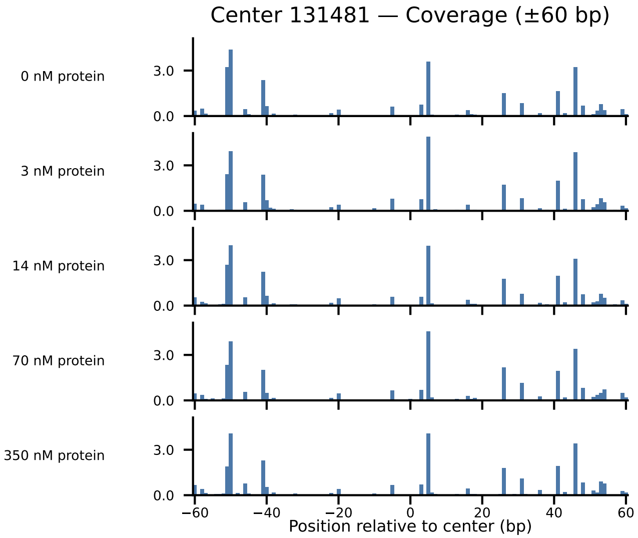
Center 131481 — Coverage (±60 bp)
0 nM protein
3 nM protein
14 nM protein
70 nM protein
350 nM protein
3.0
0.0
−60
−40
−20
0
20
40
60
Position relative to center (bp)

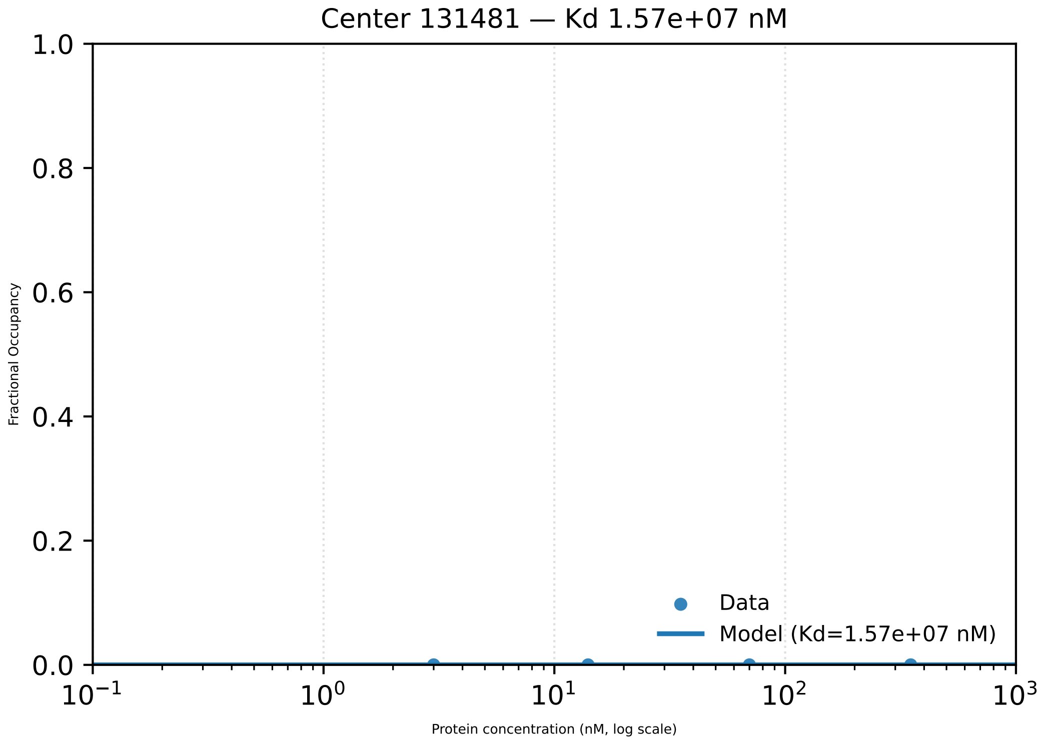

Center 131481 — Kd 1.57e+07 nM
Fractional Occupancy
1.0
0.8
0.6
0.4
0.2
0.0
$10^{-1}$
$10^{0}$
$10^{1}$
$10^{2}$
$10^{3}$
Protein concentration (nM, log scale)
Data
Model (Kd=1.57e+07 nM)

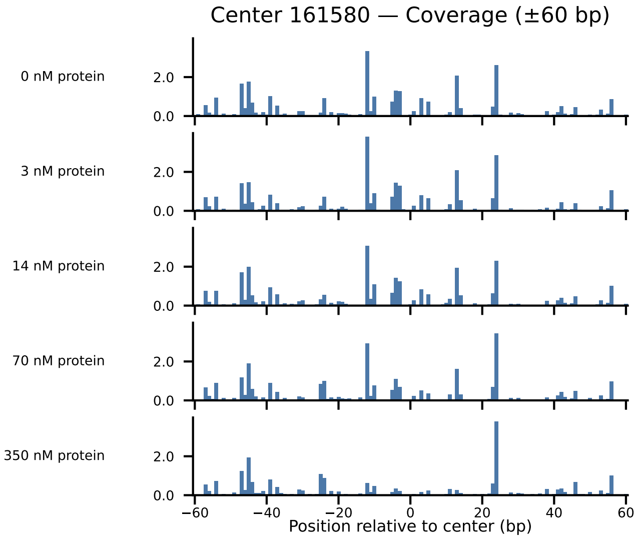

Center 161580 — Coverage (±60 bp)
0 nM protein
3 nM protein
14 nM protein
70 nM protein
350 nM protein
2.0
0.0
−60
−40
−20
0
20
40
60
Position relative to center (bp)

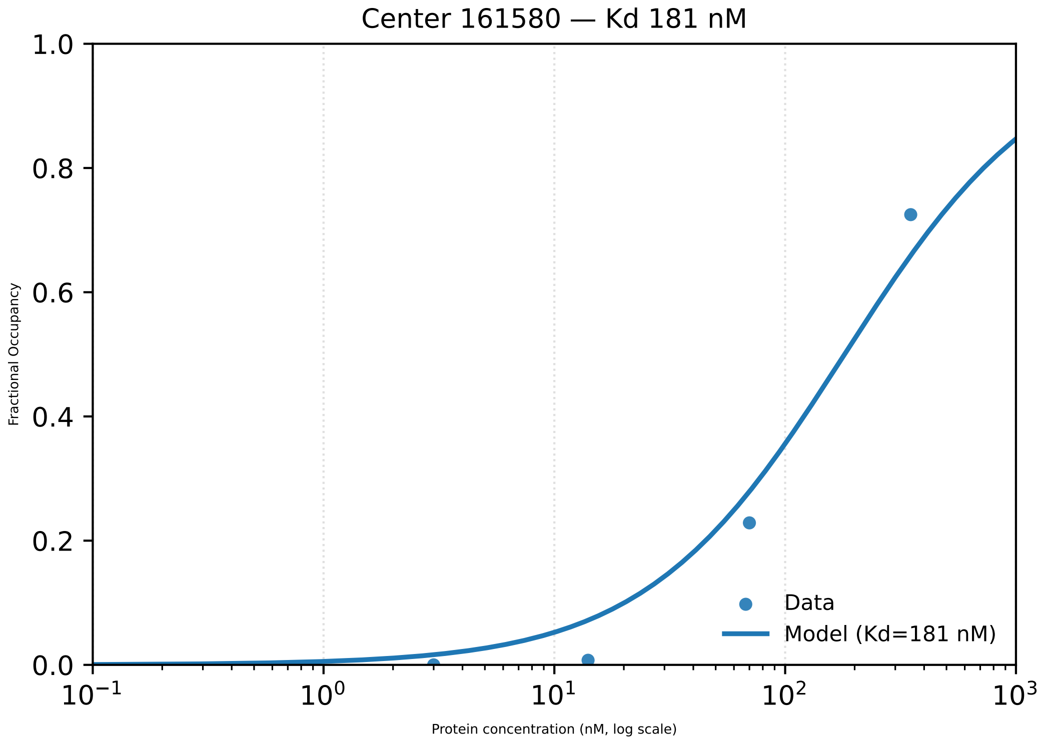
Center 161580 — Kd 181 nM
1.0
0.8
0.6
0.4
0.2
0.0
Fractional Occupancy
$10^{-1}$
$10^{0}$
$10^{1}$
$10^{2}$
$10^{3}$
Protein concentration (nM, log scale)
Data
Model (Kd=181 nM)

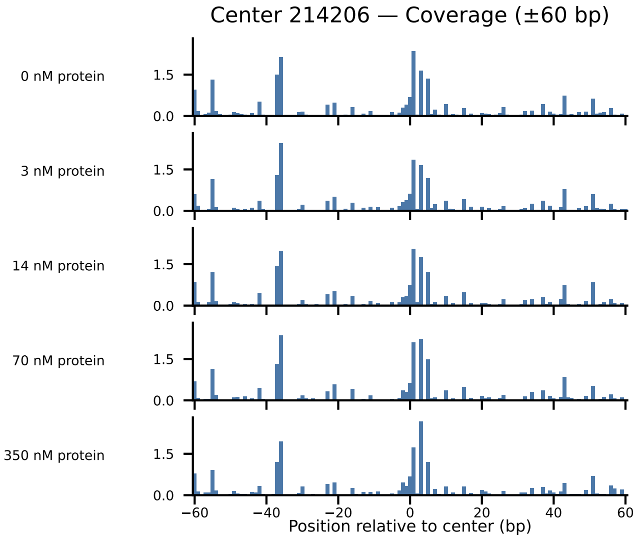

Center 214206 — Coverage (±60 bp)
0 nM protein
3 nM protein
14 nM protein
70 nM protein
350 nM protein
1.5
0.0
−60
−40
−20
0
20
40
60
Position relative to center (bp)

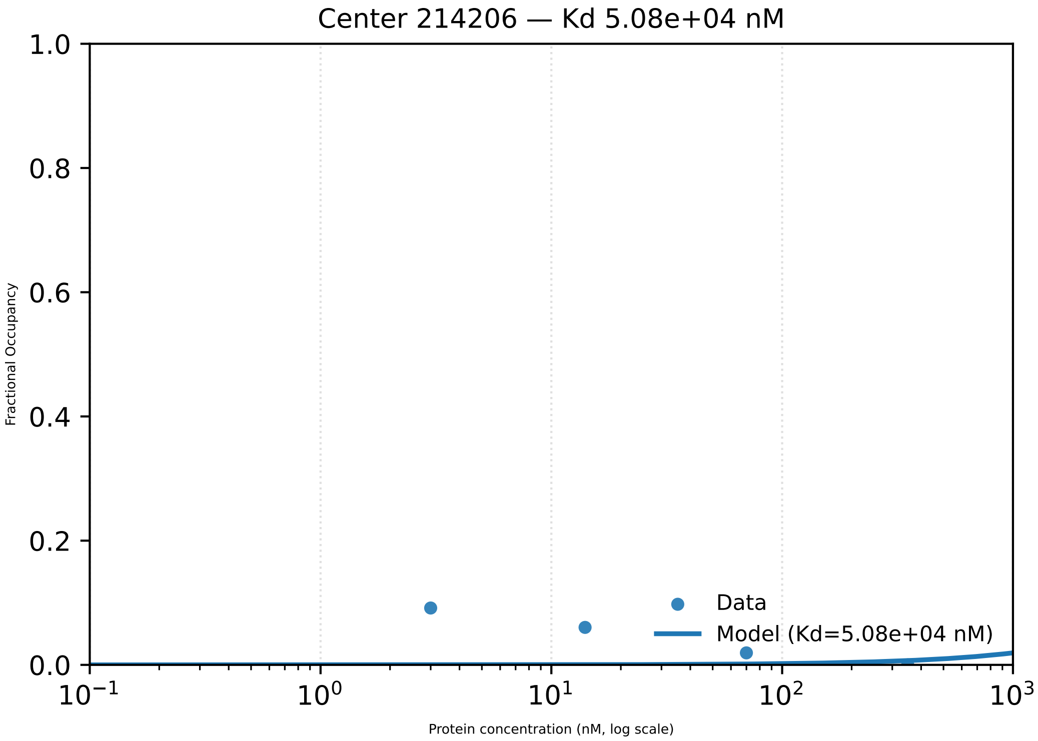
Center 214206 — Kd 5.08e+04 nM
1.0
0.8
0.6
0.4
0.2
0.0
Fractional Occupancy
Data
Model (Kd=5.08e+04 nM)
$10^{-1}$
$10^{0}$
$10^{1}$
$10^{2}$
$10^{3}$
Protein concentration (nM, log scale)

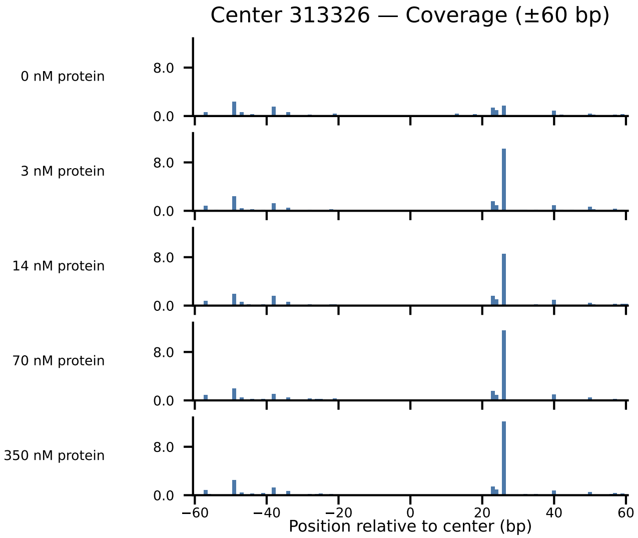

Center 313326 — Coverage (±60 bp)
0 nM protein
3 nM protein
14 nM protein
70 nM protein
350 nM protein
8.0
0.0
−60
−40
−20
0
20
40
60
Position relative to center (bp)

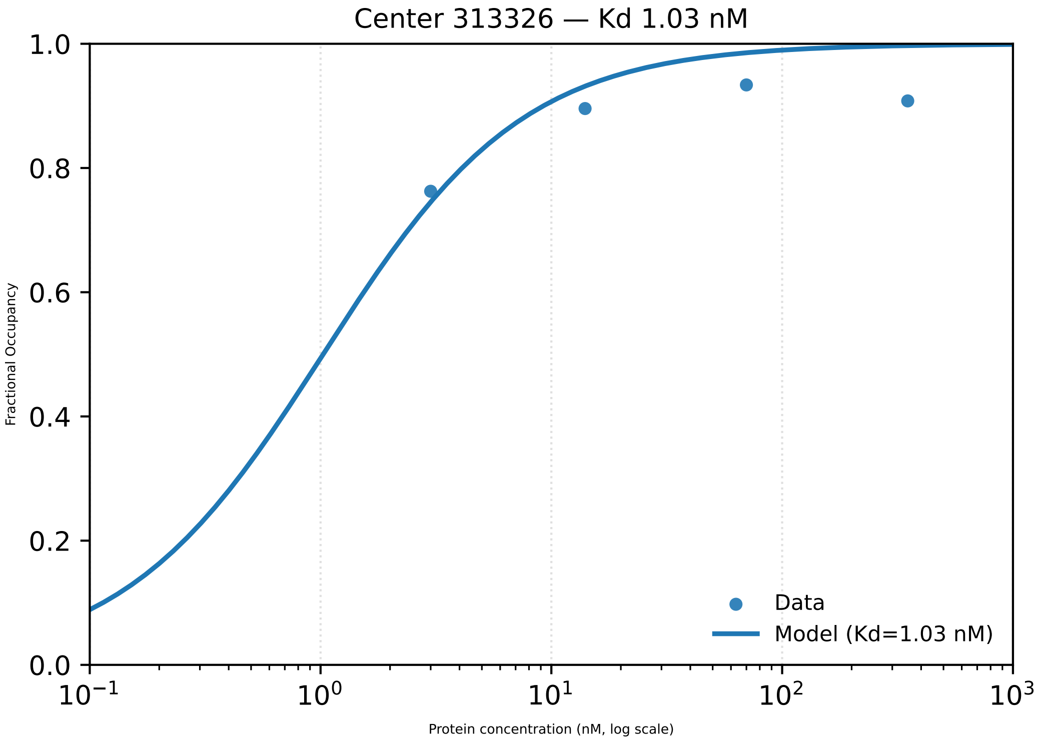
Center 313326 — Kd 1.03 nM
1.0
0.8
0.6
0.4
0.2
0.0
Fractional Occupancy
$10^{-1}$
$10^{0}$
$10^{1}$
$10^{2}$
$10^{3}$
Protein concentration (nM, log scale)
Data
Model (Kd=1.03 nM)

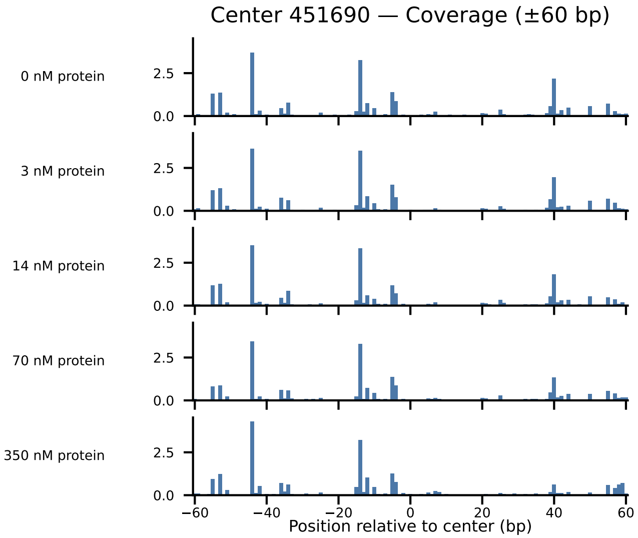

Center 451690 — Coverage (±60 bp)
0 nM protein
3 nM protein
14 nM protein
70 nM protein
350 nM protein
2.5
0.0
−60
−40
−20
0
20
40
60
Position relative to center (bp)

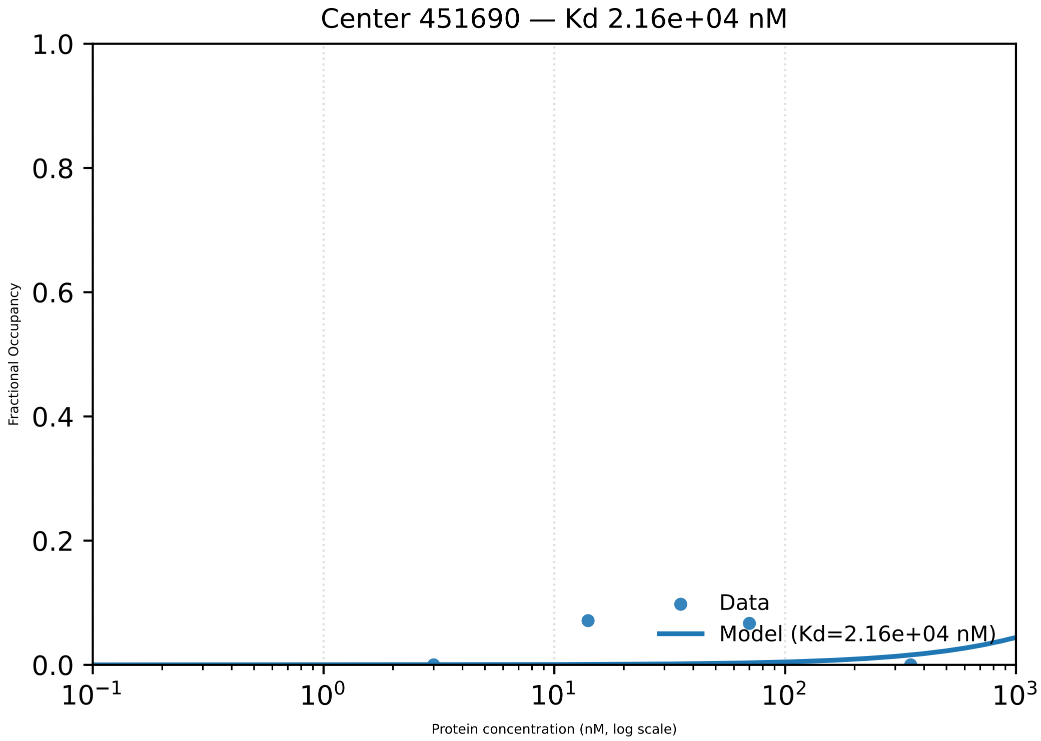

Center 451690 — Kd 2.16e+04 nM
Fractional Occupancy
1.0
0.8
0.6
0.4
0.2
0.0
$10^{-1}$
$10^{0}$
$10^{1}$
$10^{2}$
$10^{3}$
Protein concentration (nM, log scale)
Data
Model (Kd=2.16e+04 nM)

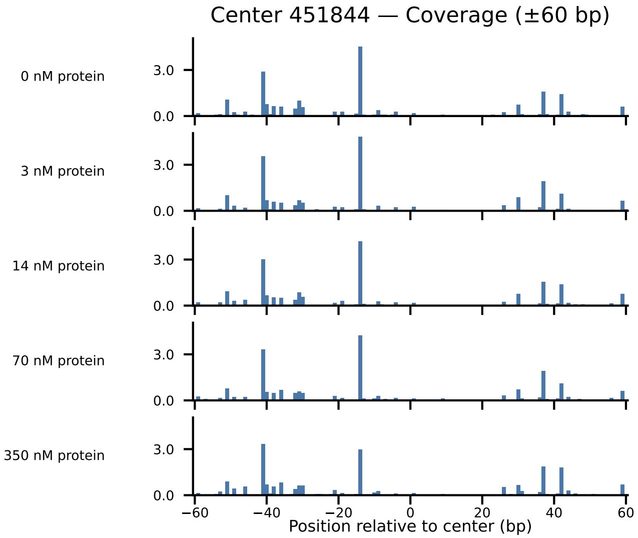

Center 451844 — Coverage (±60 bp)
0 nM protein
3 nM protein
14 nM protein
70 nM protein
350 nM protein
3.0
0.0
−60
−40
−20
0
20
40
60
Position relative to center (bp)

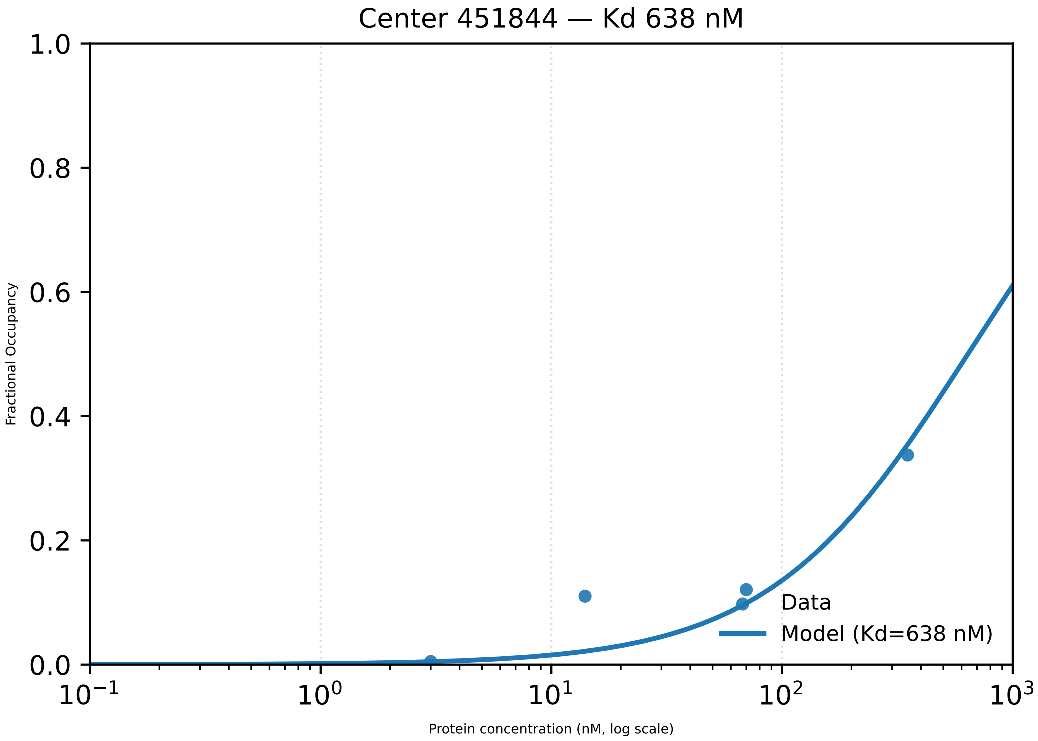

Center 451844 — Kd 638 nM
1.0
0.8
0.6
0.4
0.2
0.0
Fractional Occupancy
$10^{-1}$
$10^{0}$
$10^{1}$
$10^{2}$
$10^{3}$
Protein concentration (nM, log scale)
Data
Model (Kd=638 nM)

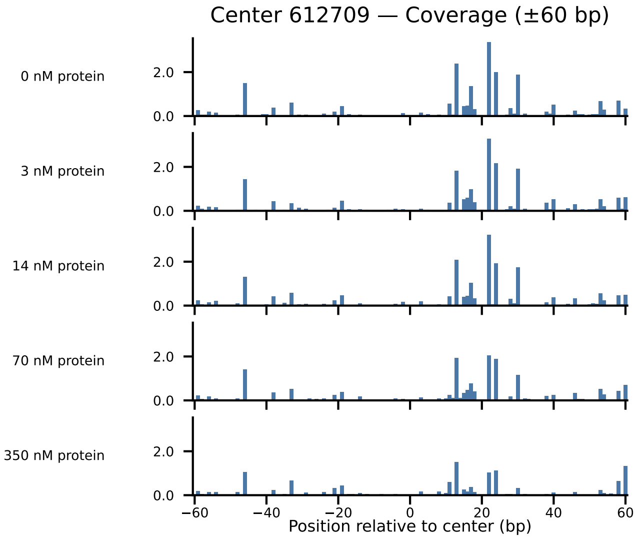
Center 612709 — Coverage (±60 bp)
0 nM protein
3 nM protein
14 nM protein
70 nM protein
350 nM protein
2.0
0.0
−60
−40
−20
0
20
40
60
Position relative to center (bp)

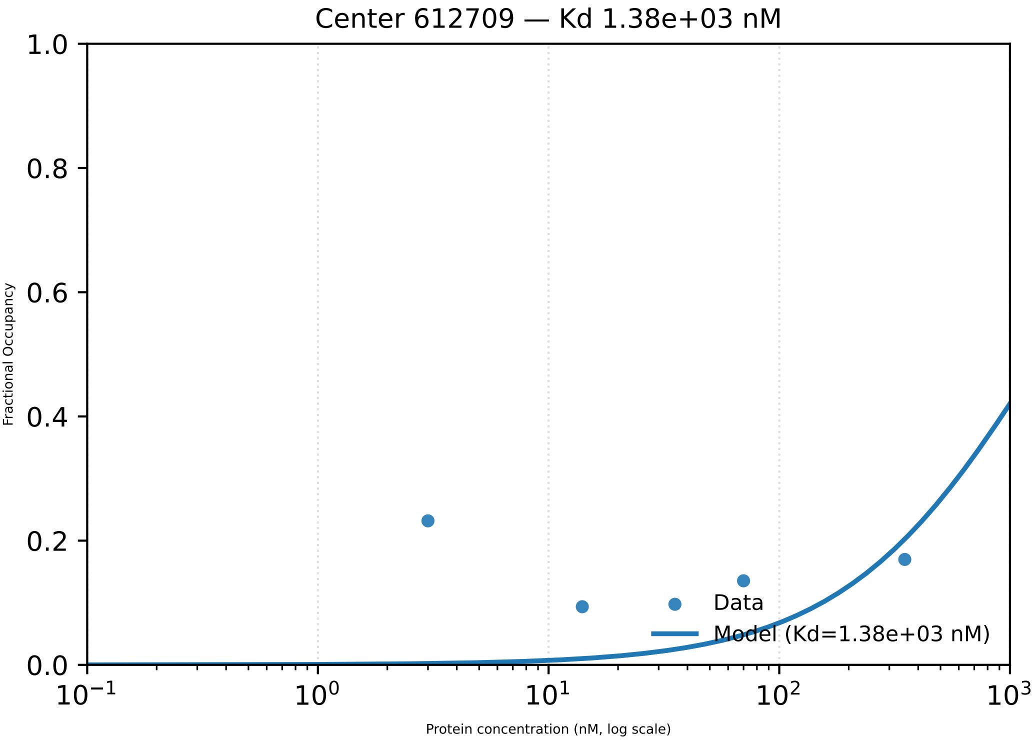

Center 612709 — Kd 1.38e+03 nM
Fractional Occupancy
1.0
0.8
0.6
0.4
0.2
0.0
$10^{-1}$
$10^{0}$
$10^{1}$
$10^{2}$
$10^{3}$
Protein concentration (nM, log scale)
Data
Model (Kd=1.38e+03 nM)

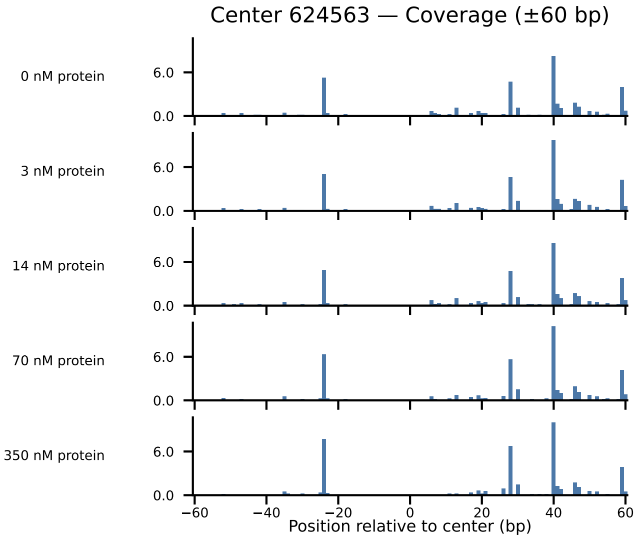
Center 624563 — Coverage (±60 bp)
0 nM protein
3 nM protein
14 nM protein
70 nM protein
350 nM protein
6.0
0.0
−60
−40
−20
0
20
40
60
Position relative to center (bp)

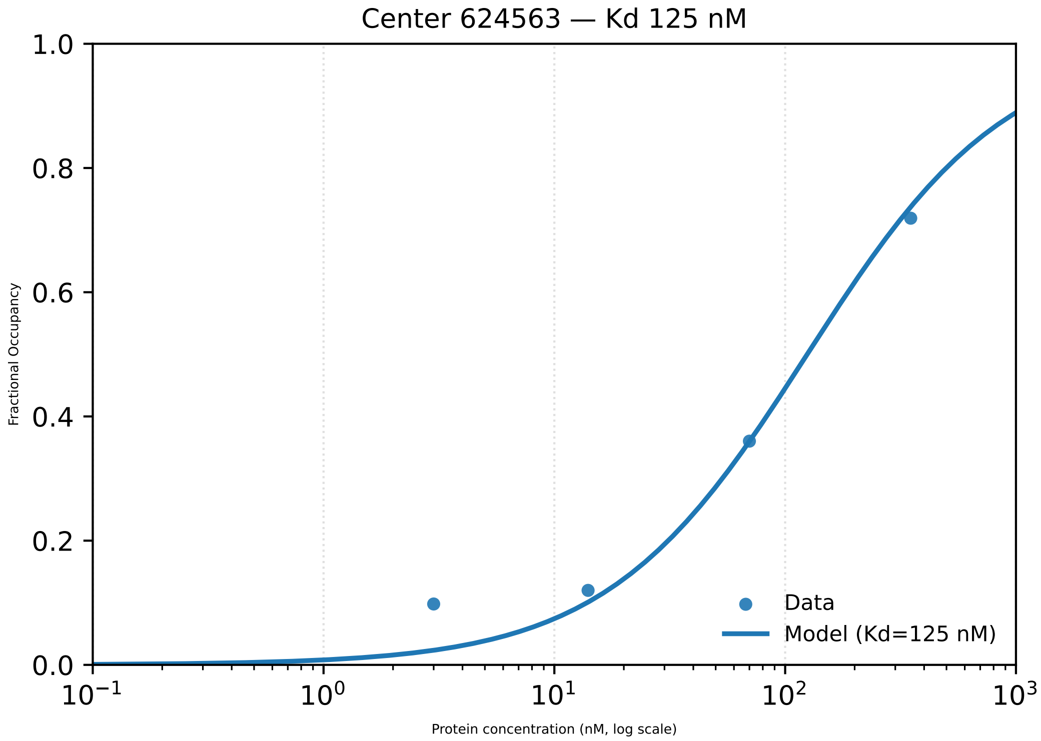

Center 624563 — Kd 125 nM
Fractional Occupancy
1.0
0.8
0.6
0.4
0.2
0.0
$10^{-1}$
$10^{0}$
$10^{1}$
$10^{2}$
$10^{3}$
Protein concentration (nM, log scale)
Data
Model (Kd=125 nM)

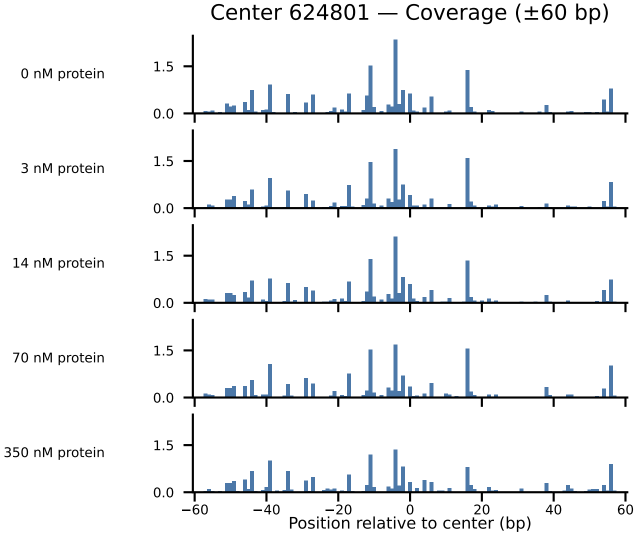

Center 624801 — Coverage (±60 bp)
0 nM protein
3 nM protein
14 nM protein
70 nM protein
350 nM protein
1.5
0.0
−60
−40
−20
0
20
40
60
Position relative to center (bp)

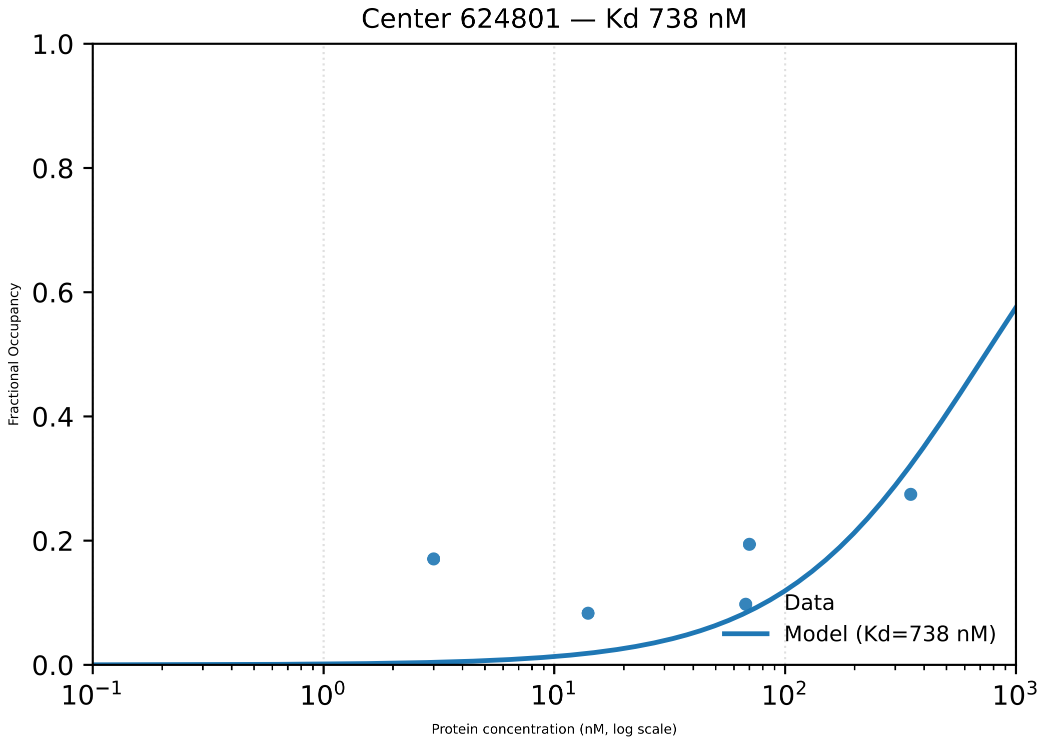

Center 624801 — Kd 738 nM
1.0
0.8
0.6
0.4
0.2
0.0
Fractional Occupancy
$10^{-1}$
$10^{0}$
$10^{1}$
$10^{2}$
$10^{3}$
Protein concentration (nM, log scale)
Data
Model (Kd=738 nM)

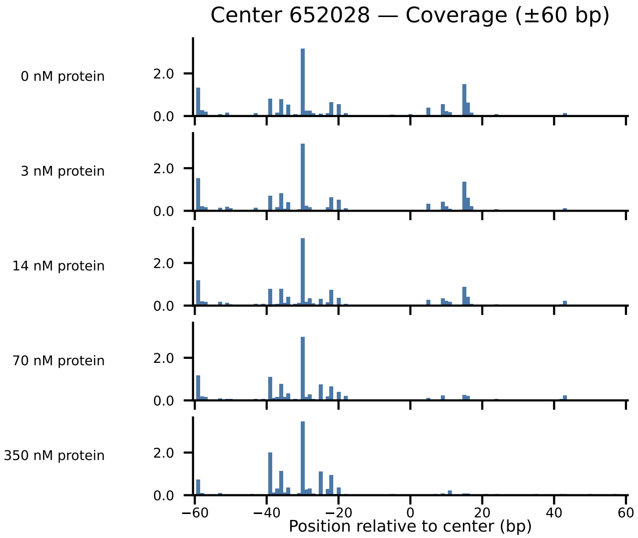

Center 652028 — Coverage (±60 bp)
0 nM protein
3 nM protein
14 nM protein
70 nM protein
350 nM protein
2.0
0.0
−60
−40
−20
0
20
40
60
Position relative to center (bp)

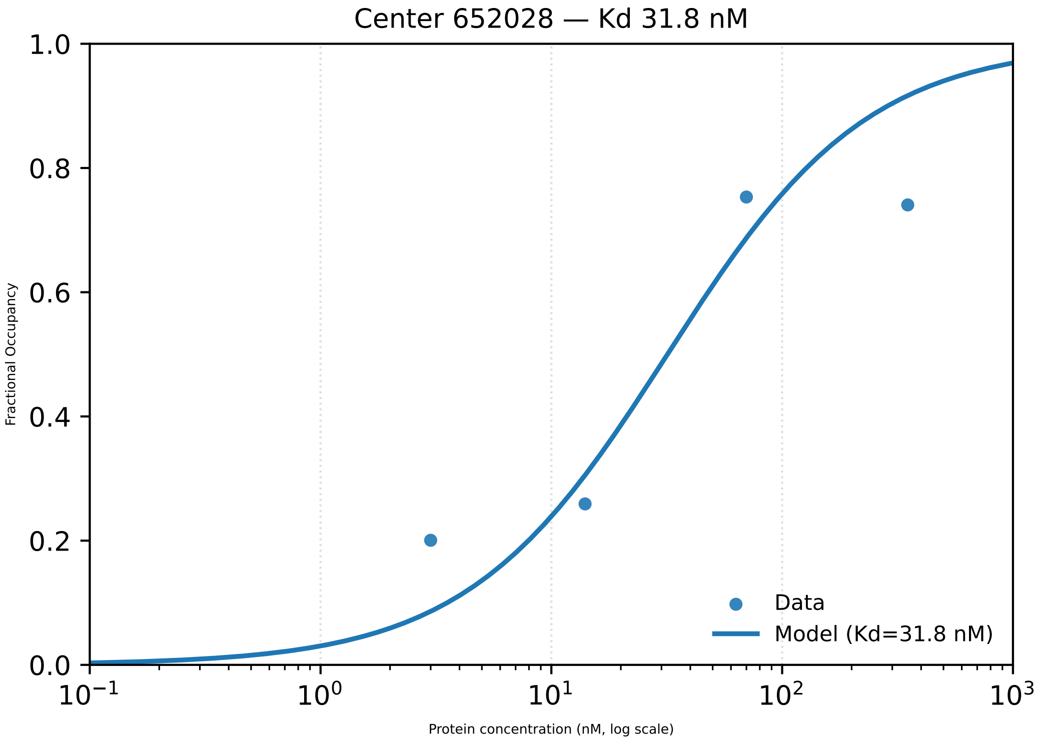
Center 652028 — Kd 31.8 nM
1.0
0.8
0.6
0.4
0.2
0.0
Fractional Occupancy
$10^{-1}$
$10^{0}$
$10^{1}$
$10^{2}$
$10^{3}$
Protein concentration (nM, log scale)
Data
Model (Kd=31.8 nM)

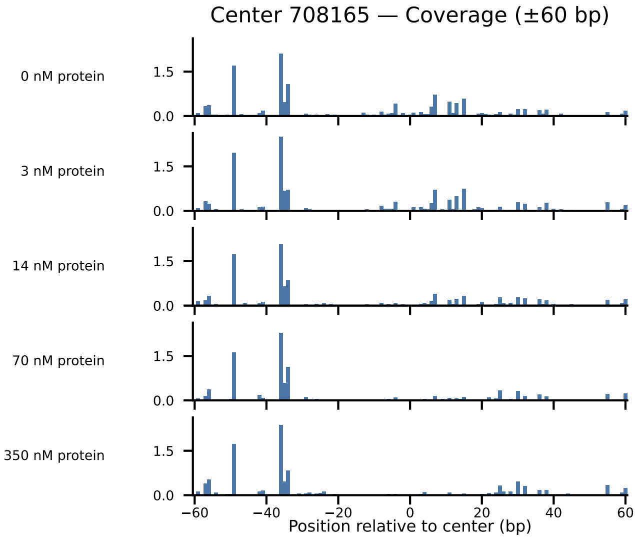
Center 708165 — Coverage (±60 bp)
0 nM protein
3 nM protein
14 nM protein
70 nM protein
350 nM protein
1.5
0.0
−60
−40
−20
0
20
40
60
Position relative to center (bp)

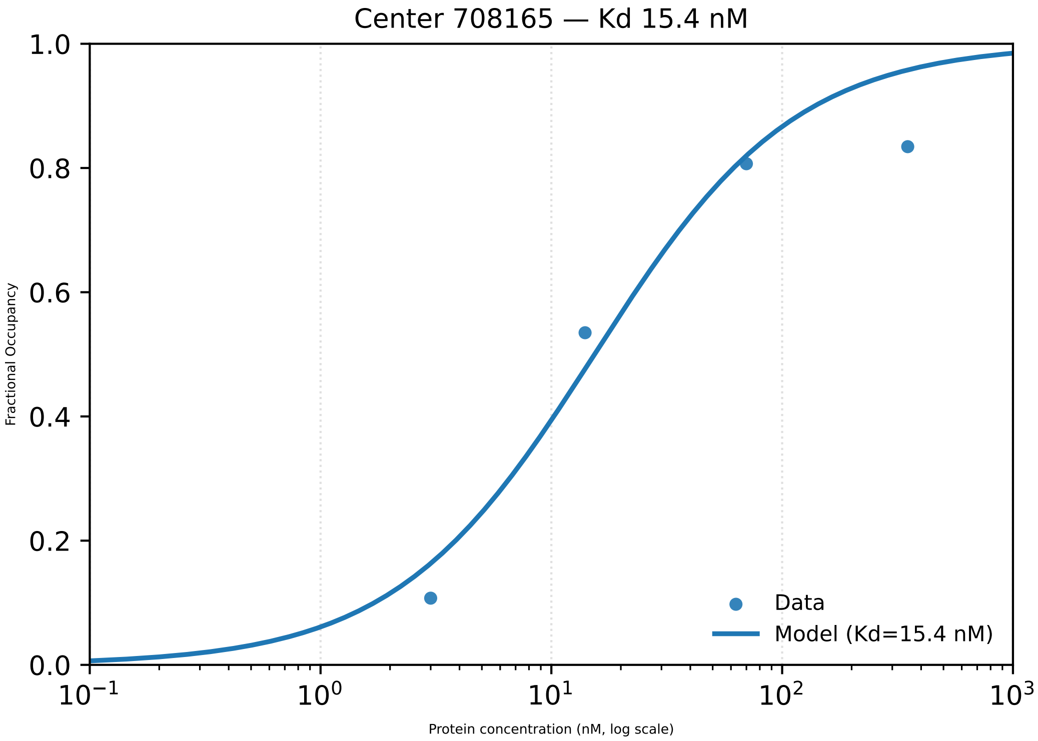

Center 708165 — Kd 15.4 nM
Fractional Occupancy
1.0
0.8
0.6
0.4
0.2
0.0
$10^{-1}$
$10^{0}$
$10^{1}$
$10^{2}$
$10^{3}$
Protein concentration (nM, log scale)
Data
Model (Kd=15.4 nM)

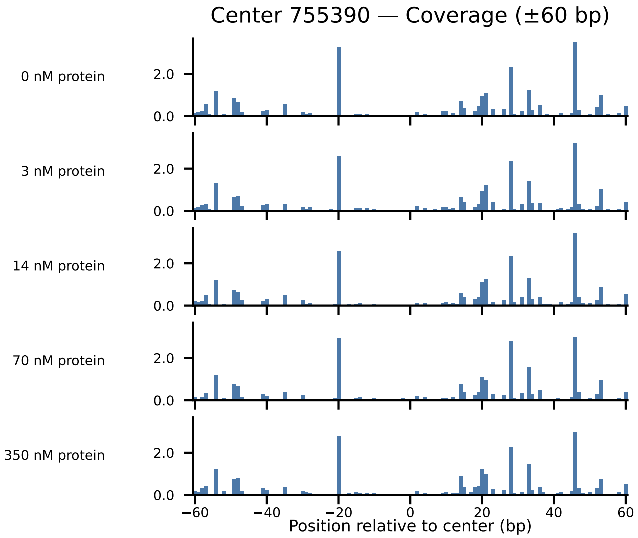
Center 755390 — Coverage (±60 bp)
0 nM protein
3 nM protein
14 nM protein
70 nM protein
350 nM protein
0.0
2.0
−60
−40
−20
0
20
40
60
Position relative to center (bp)

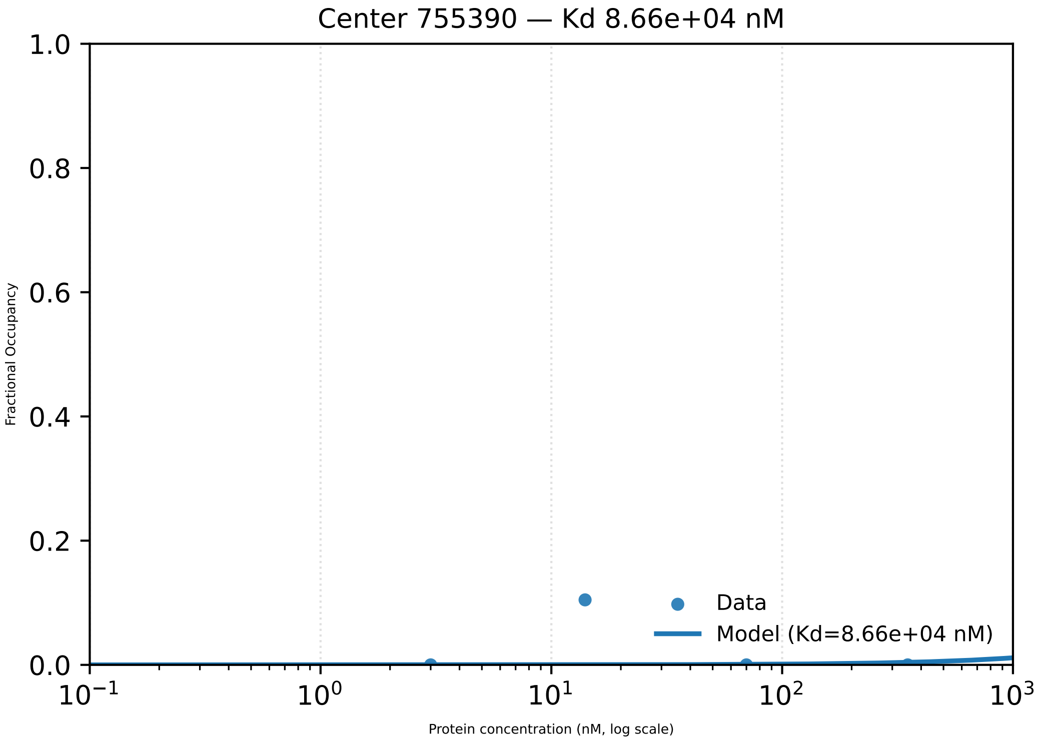

Center 755390 — Kd 8.66e+04 nM
Fractional Occupancy
1.0
0.8
0.6
0.4
0.2
0.0
$10^{-1}$
$10^{0}$
$10^{1}$
$10^{2}$
$10^{3}$
Protein concentration (nM, log scale)
Data
Model (Kd=8.66e+04 nM)

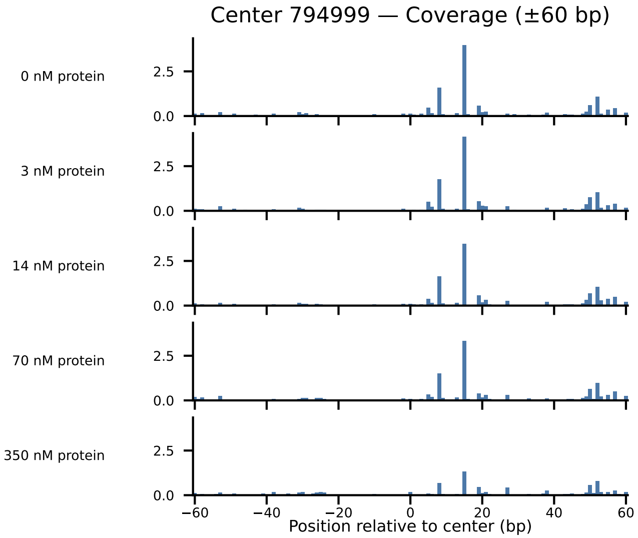
Center 794999 — Coverage (±60 bp)
0 nM protein
3 nM protein
14 nM protein
70 nM protein
350 nM protein
0.0
2.5
−60
−40
−20
0
20
40
60
Position relative to center (bp)

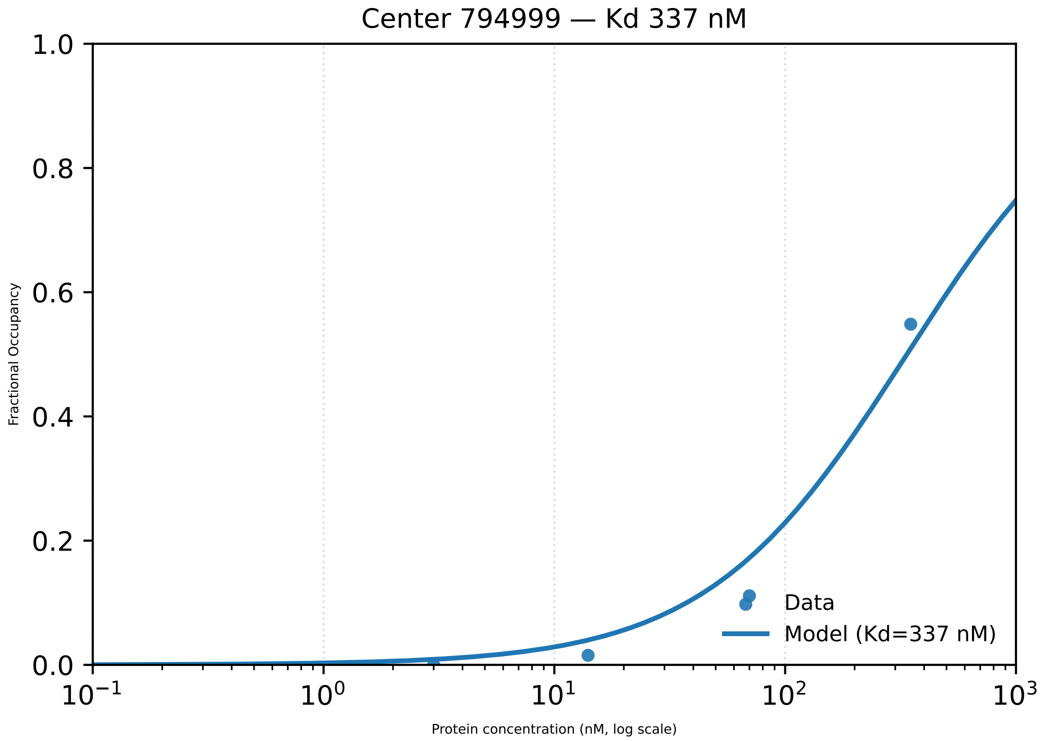

Center 794999 — Kd 337 nM
1.0
0.8
0.6
0.4
0.2
0.0
Fractional Occupancy
$10^{-1}$
$10^{0}$
$10^{1}$
$10^{2}$
$10^{3}$
Protein concentration (nM, log scale)
Data
Model (Kd=337 nM)

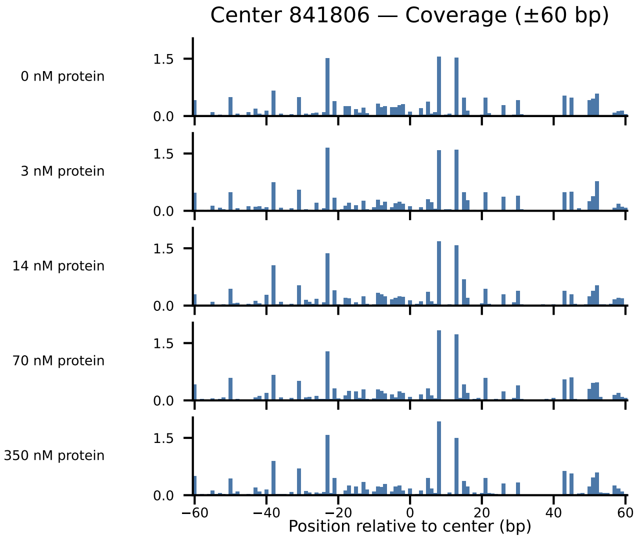
Center 841806 — Coverage (±60 bp)
0 nM protein
3 nM protein
14 nM protein
70 nM protein
350 nM protein
1.5
0.0
−60
−40
−20
0
20
40
60
Position relative to center (bp)

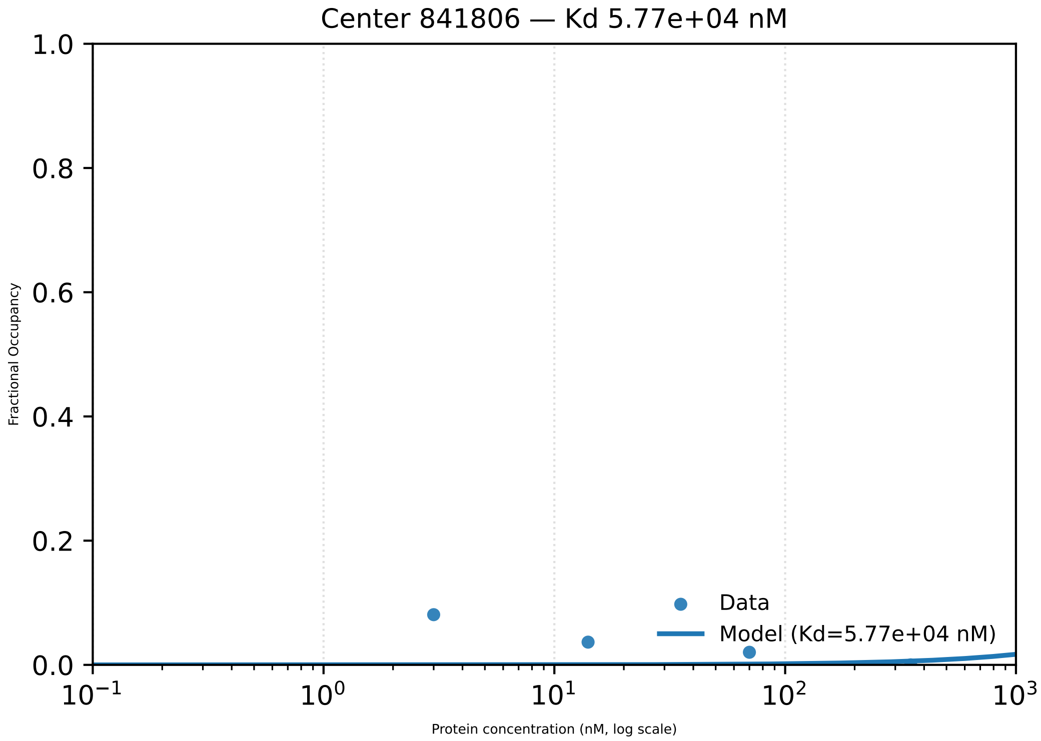
Center 841806 — Kd 5.77e+04 nM
1.0
0.8
0.6
0.4
0.2
0.0
Fractional Occupancy
$10^{-1}$
$10^{0}$
$10^{1}$
$10^{2}$
$10^{3}$
Protein concentration (nM, log scale)
Data
Model (Kd=5.77e+04 nM)

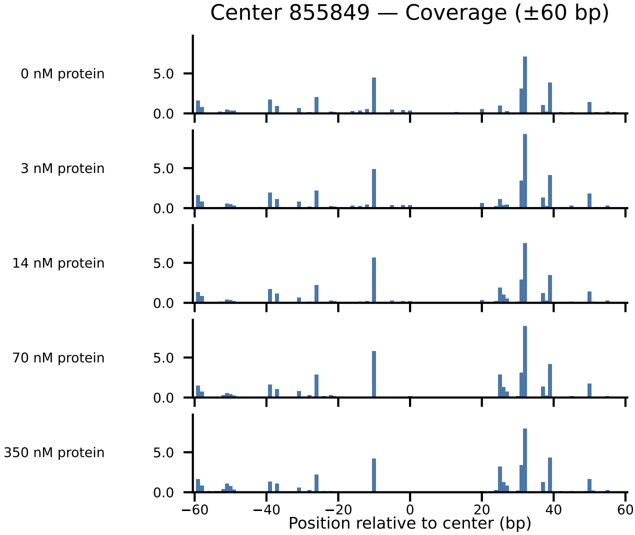

Center 855849 — Coverage (±60 bp)
0 nM protein
3 nM protein
14 nM protein
70 nM protein
350 nM protein
5.0
0.0
−60
−40
−20
0
20
40
60
Position relative to center (bp)

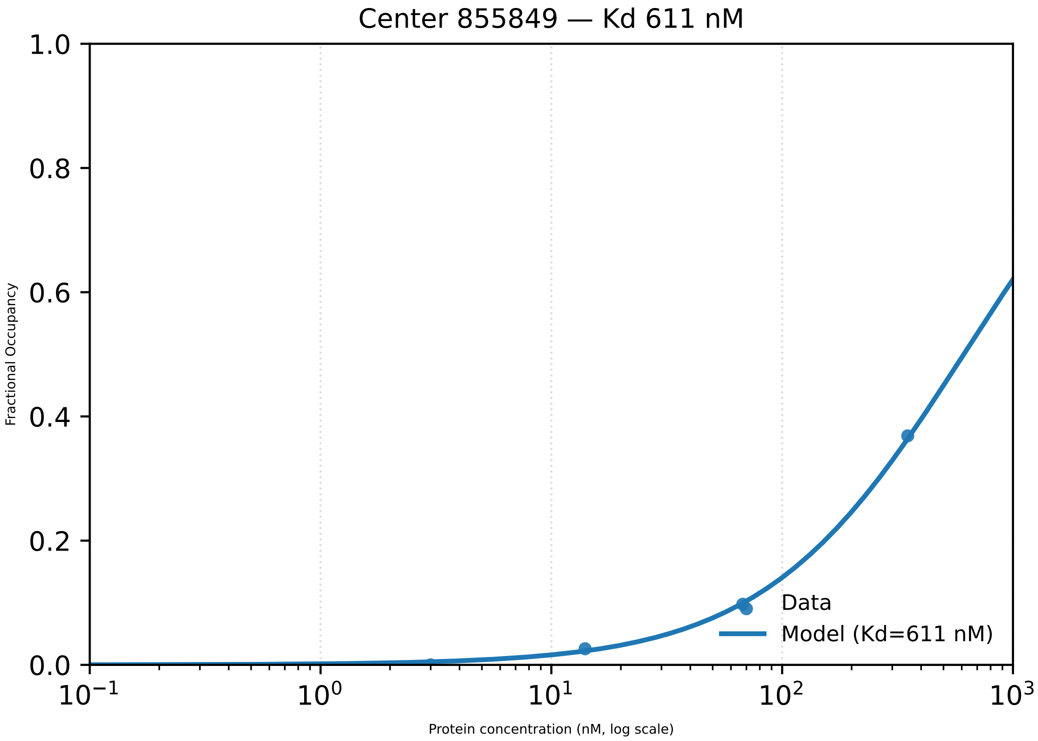
Center 855849 — Kd 611 nM
1.0
0.8
0.6
0.4
0.2
0.0
Fractional Occupancy
$10^{-1}$
$10^{0}$
$10^{1}$
$10^{2}$
$10^{3}$
Protein concentration (nM, log scale)
Data
Model (Kd=611 nM)

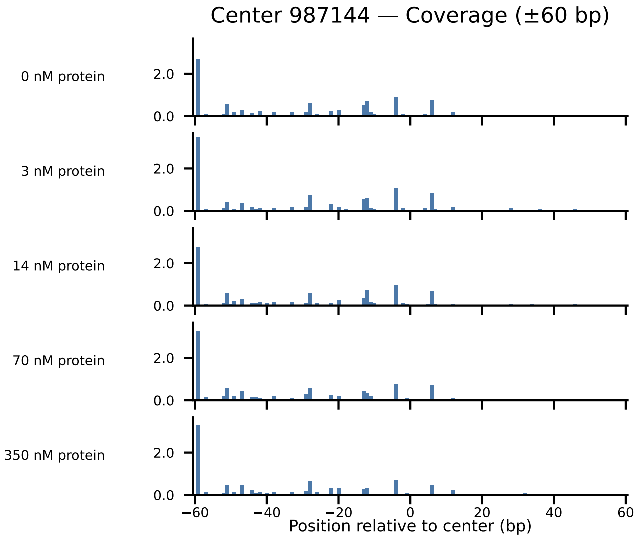

Center 987144 — Coverage (±60 bp)
0 nM protein
3 nM protein
14 nM protein
70 nM protein
350 nM protein
2.0
0.0
−60
−40
−20
0
20
40
60
Position relative to center (bp)

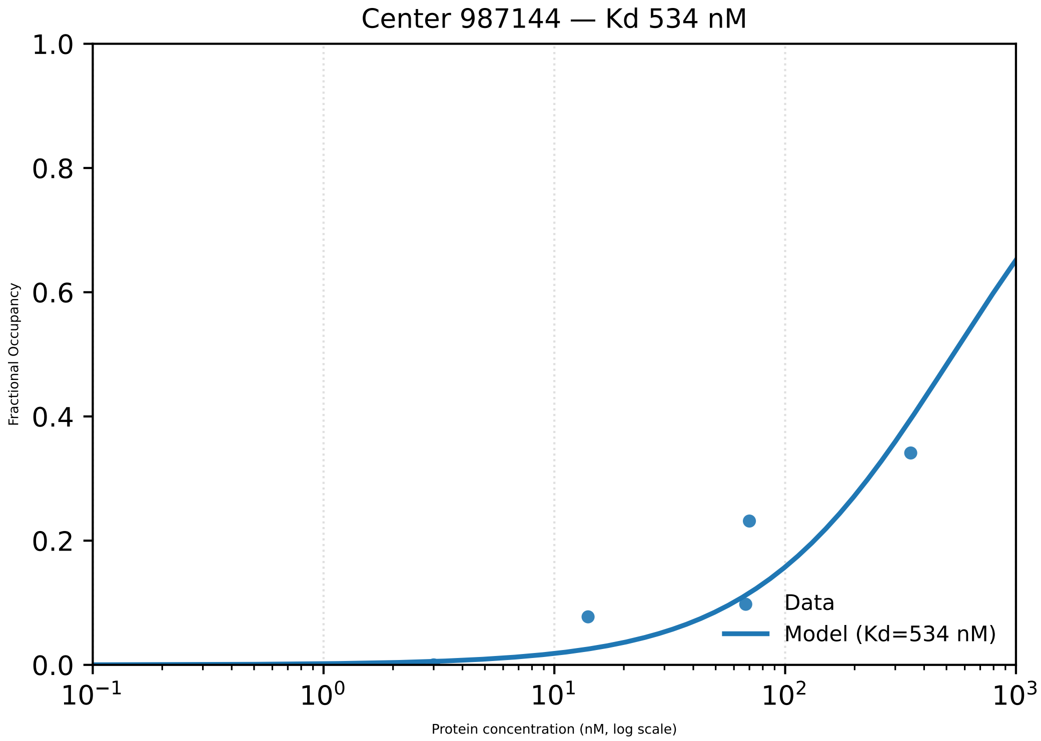

Center 987144 — Kd 534 nM
Fractional Occupancy
1.0
0.8
0.6
0.4
0.2
0.0
$10^{-1}$
$10^{0}$
$10^{1}$
$10^{2}$
$10^{3}$
Protein concentration (nM, log scale)
Data
Model (Kd=534 nM)

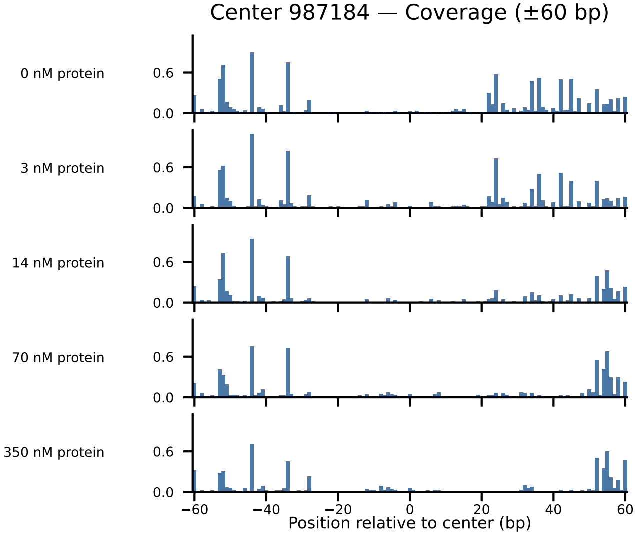
Center 987184 — Coverage (±60 bp)
0 nM protein
3 nM protein
14 nM protein
70 nM protein
350 nM protein
0.6
0.0
−60
−40
−20
0
20
40
60
Position relative to center (bp)

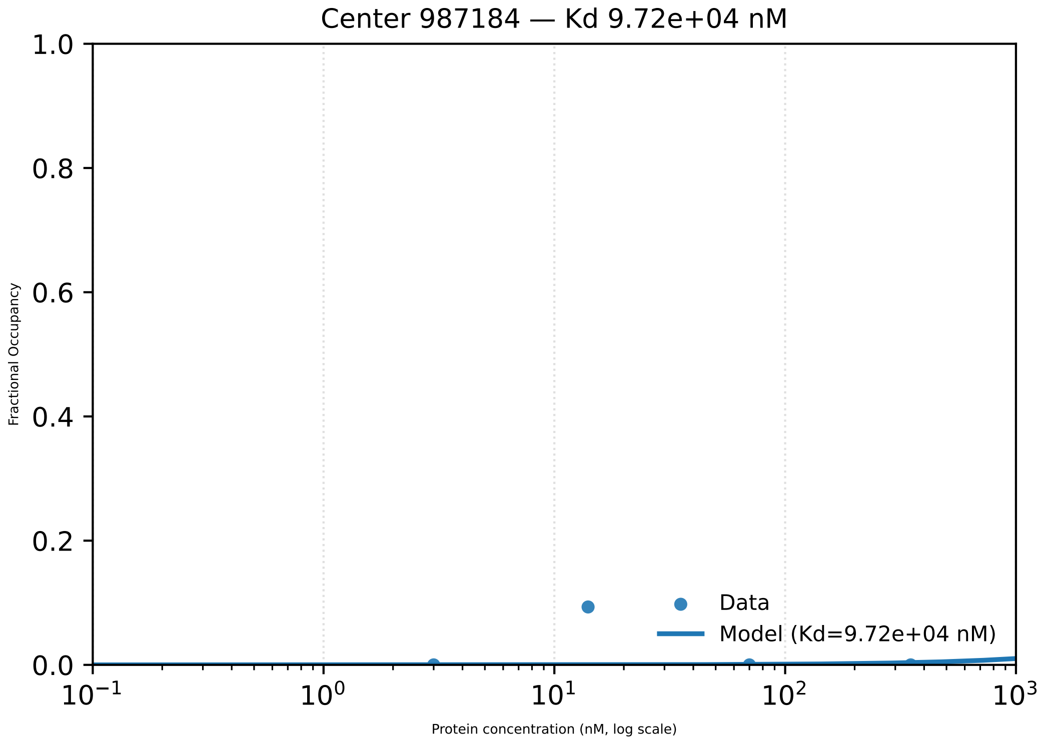

Center 987184 — Kd 9.72e+04 nM
1.0
0.8
0.6
0.4
0.2
0.0
Fractional Occupancy
Data
Model (Kd=9.72e+04 nM)
$10^{-1}$
$10^{0}$
$10^{1}$
$10^{2}$
$10^{3}$
Protein concentration (nM, log scale)

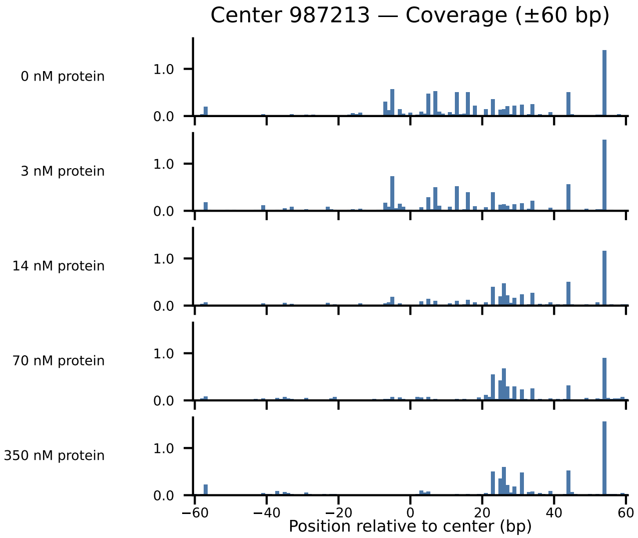

Center 987213 — Coverage (±60 bp)
0 nM protein
3 nM protein
14 nM protein
70 nM protein
350 nM protein
1.0
0.0
−60
−40
−20
0
20
40
60
Position relative to center (bp)

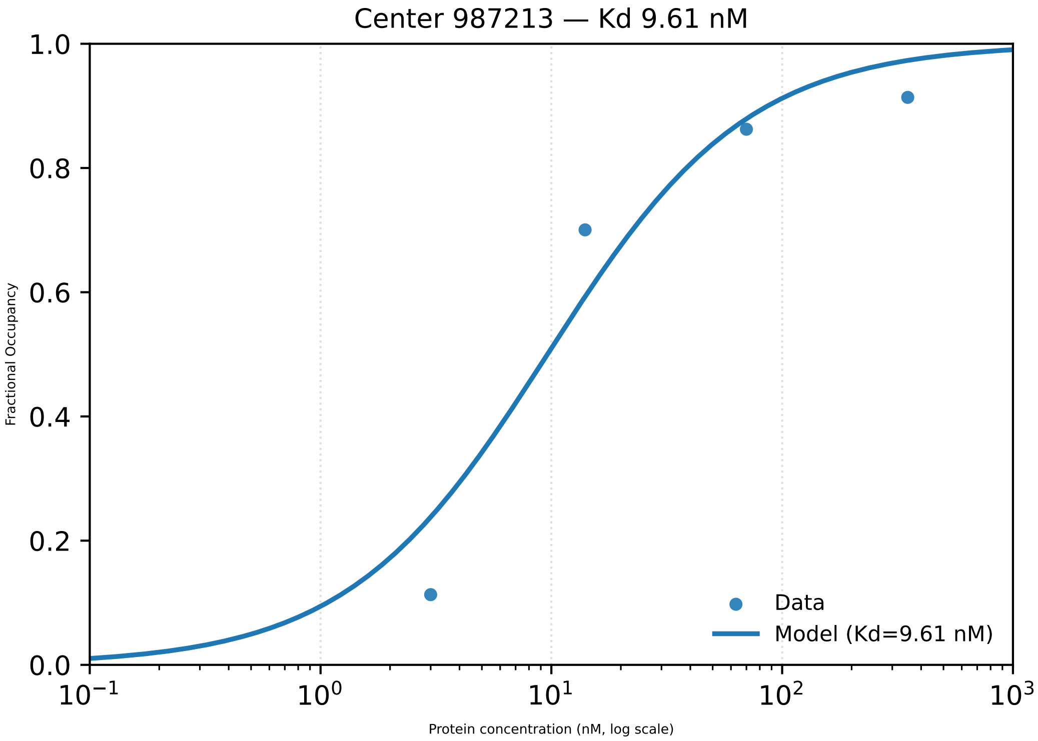
Center 987213 — Kd 9.61 nM
1.0
0.8
0.6
0.4
0.2
0.0
Fractional Occupancy
$10^{-1}$
$10^{0}$
$10^{1}$
$10^{2}$
$10^{3}$
Protein concentration (nM, log scale)
Data
Model (Kd=9.61 nM)

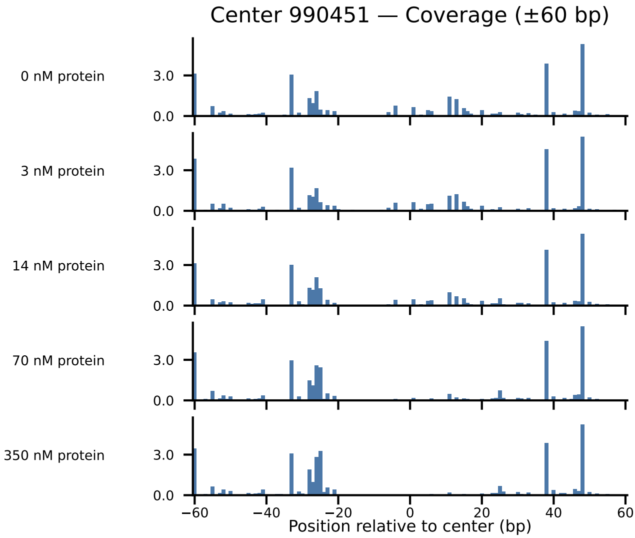

Center 990451 — Coverage (±60 bp)
0 nM protein
3 nM protein
14 nM protein
70 nM protein
350 nM protein
3.0
0.0
−60
−40
−20
0
20
40
60
Position relative to center (bp)

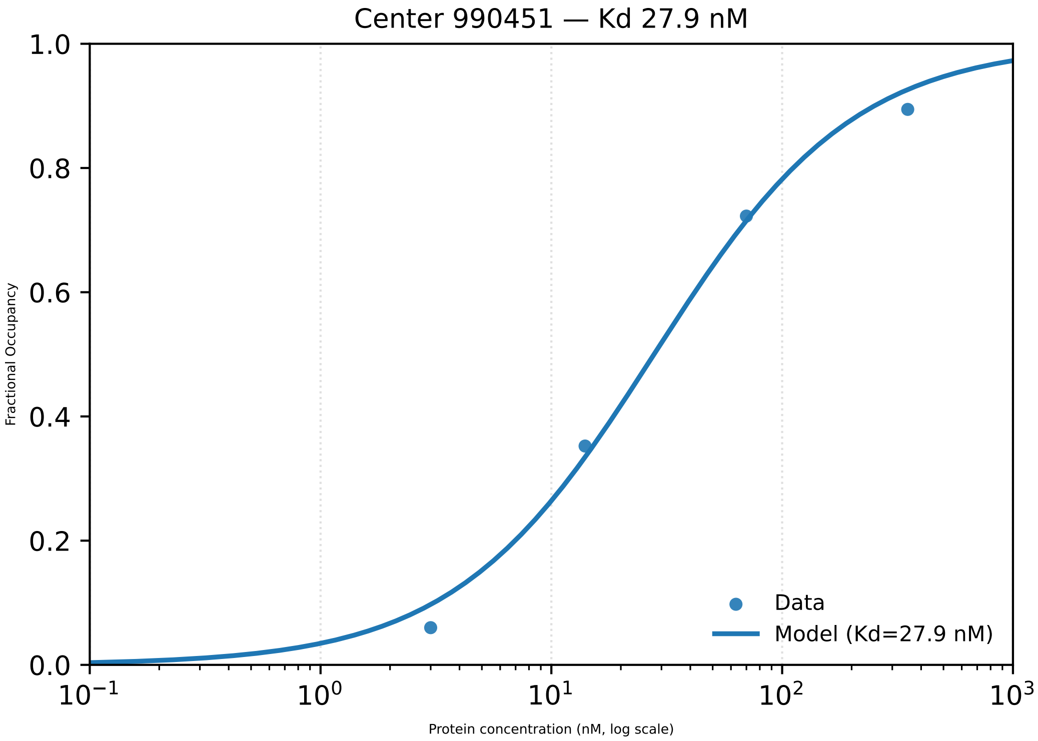
Center 990451 — Kd 27.9 nM
Fractional Occupancy
1.0
0.8
0.6
0.4
0.2
0.0
$10^{-1}$
$10^{0}$
$10^{1}$
$10^{2}$
$10^{3}$
Protein concentration (nM, log scale)
Data
Model (Kd=27.9 nM)

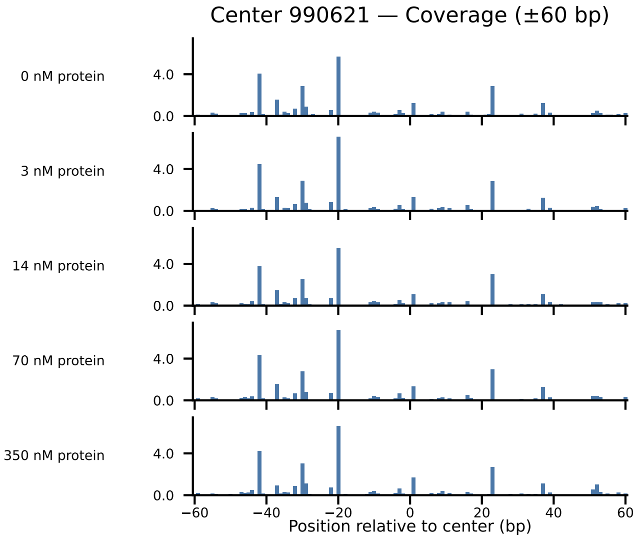

Center 990621 — Coverage (±60 bp)
0 nM protein
3 nM protein
14 nM protein
70 nM protein
350 nM protein
4.0
0.0
−60
−40
−20
0
20
40
60
Position relative to center (bp)

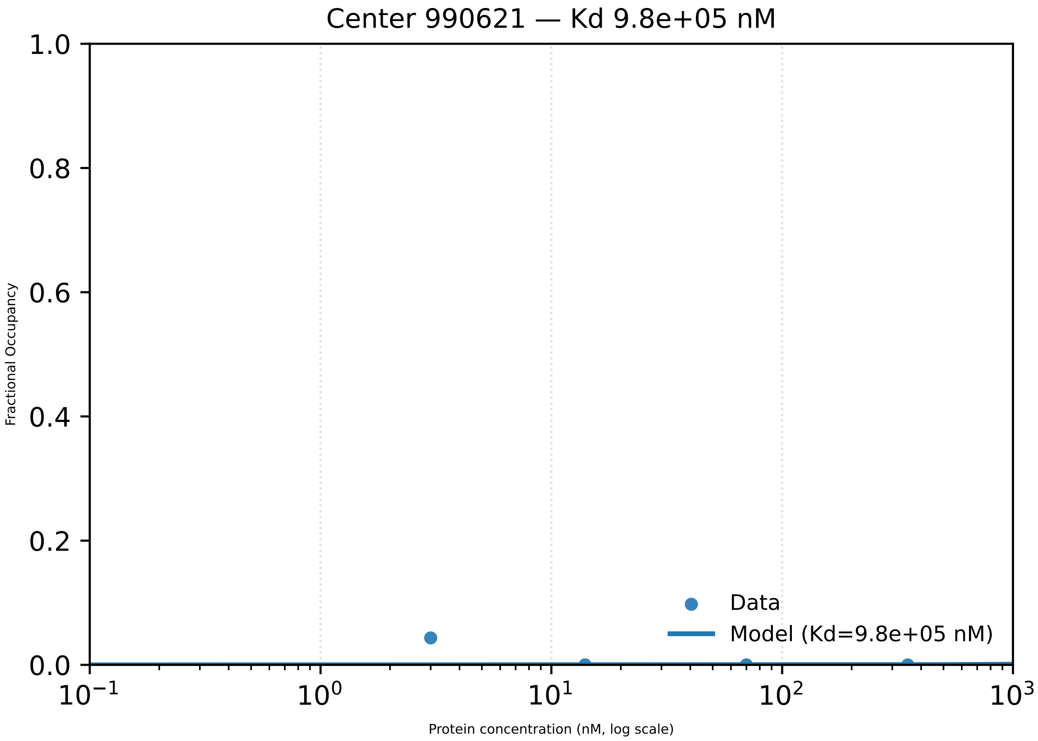

Center 990621 — Kd 9.8e+05 nM
1.0
0.8
0.6
0.4
0.2
0.0
Fractional Occupancy
$10^{-1}$
$10^{0}$
$10^{1}$
$10^{2}$
$10^{3}$
Protein concentration (nM, log scale)
Data
Model (Kd=9.8e+05 nM)

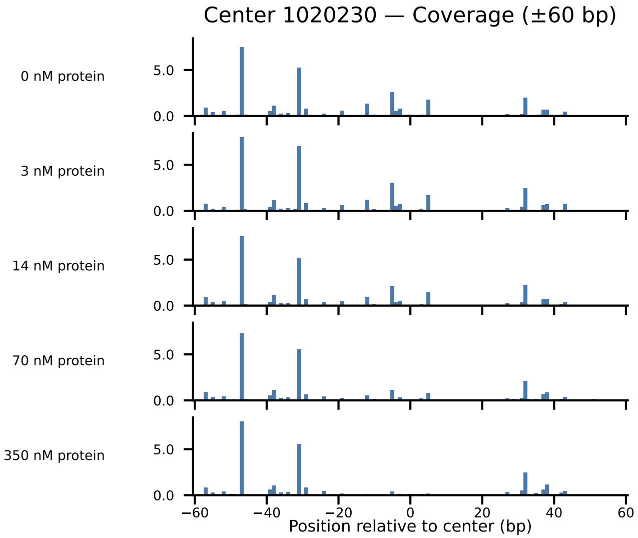
Center 1020230 — Coverage (±60 bp)
0 nM protein
3 nM protein
14 nM protein
70 nM protein
350 nM protein
5.0
0.0
−60
−40
−20
0
20
40
60
Position relative to center (bp)

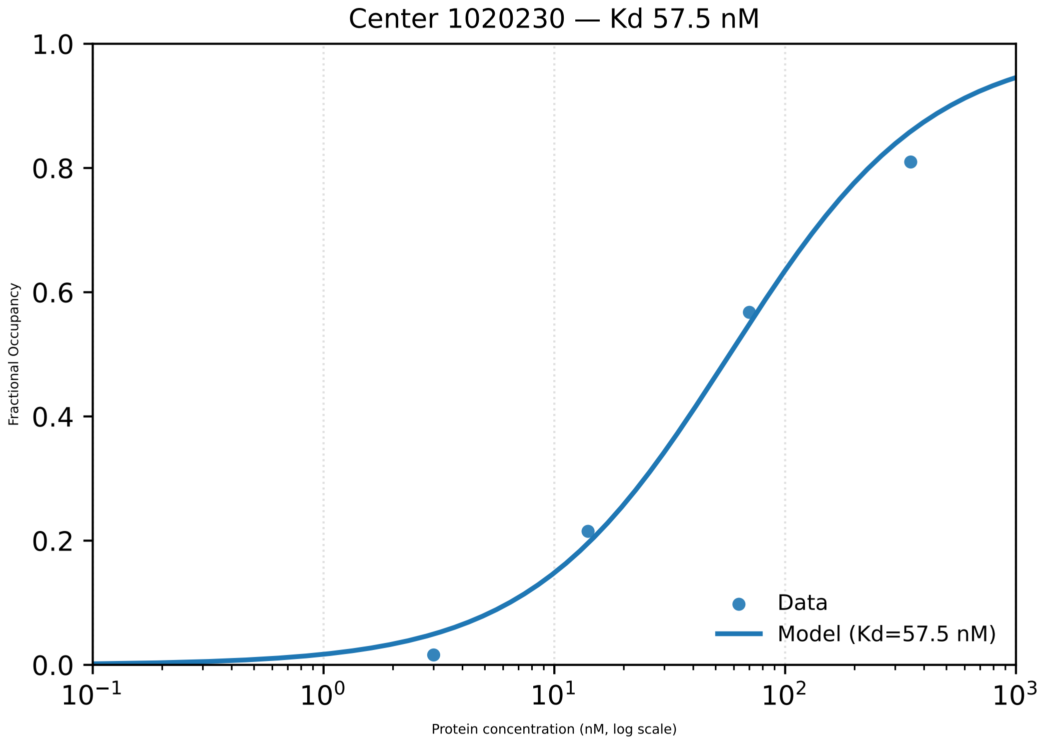

Center 1020230 — Kd 57.5 nM
Fractional Occupancy
1.0
0.8
0.6
0.4
0.2
0.0
10−1
100
101
102
103
Protein concentration (nM, log scale)
Data
Model (Kd=57.5 nM)

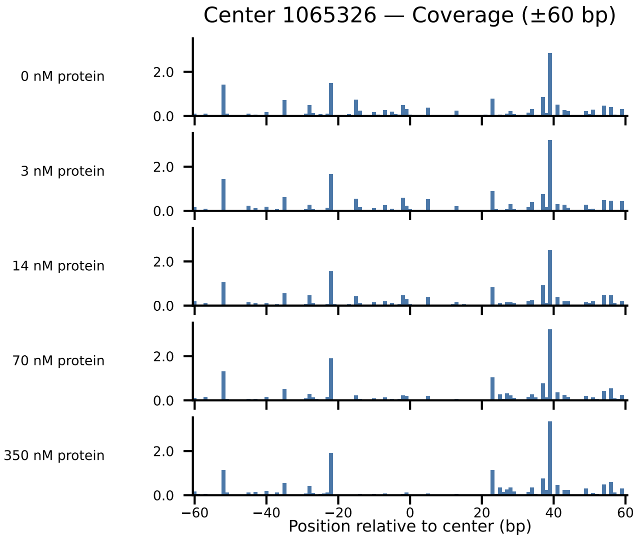

Center 1065326 — Coverage (±60 bp)
0 nM protein
3 nM protein
14 nM protein
70 nM protein
350 nM protein
2.0
0.0
−60
−40
−20
0
20
40
60
Position relative to center (bp)

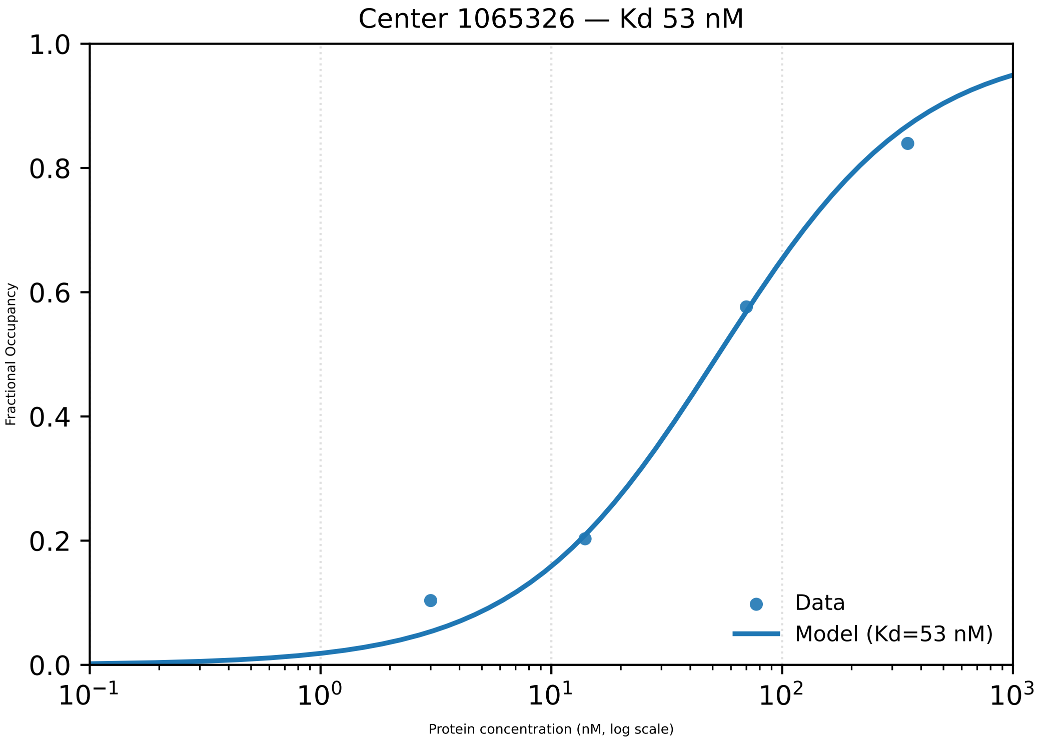

Center 1065326 — Kd 53 nM
1.0
0.8
0.6
0.4
0.2
0.0
Fractional Occupancy
$10^{-1}$
$10^{0}$
$10^{1}$
$10^{2}$
$10^{3}$
Protein concentration (nM, log scale)
Data
Model (Kd=53 nM)

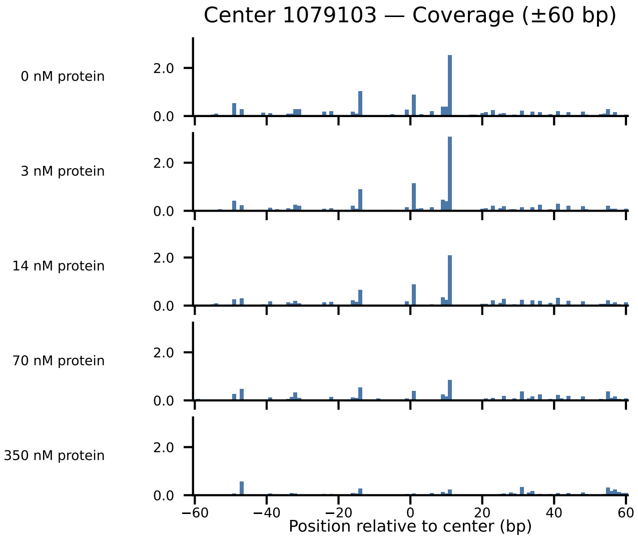
Center 1079103 — Coverage (±60 bp)
0 nM protein
3 nM protein
14 nM protein
70 nM protein
350 nM protein
2.0
0.0
−60
−40
−20
0
20
40
60
Position relative to center (bp)

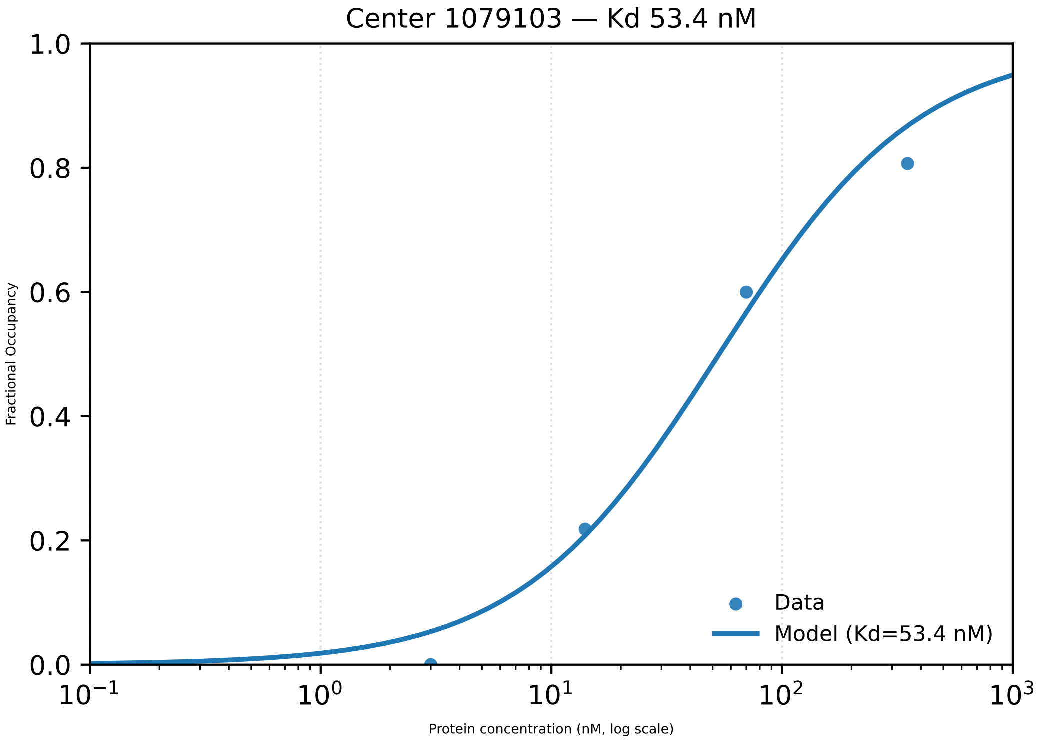

Center 1079103 — Kd 53.4 nM
1.0
0.8
0.6
0.4
0.2
0.0
Fractional Occupancy
$10^{-1}$
$10^{0}$
$10^{1}$
$10^{2}$
$10^{3}$
Protein concentration (nM, log scale)
Data
Model (Kd=53.4 nM)

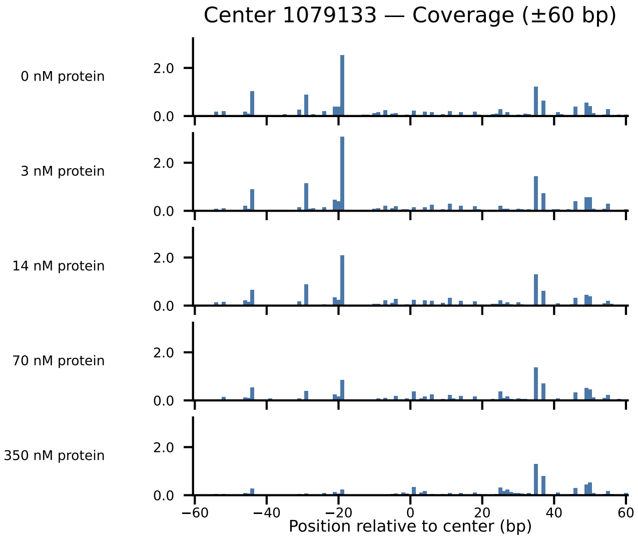

Center 1079133 — Coverage (±60 bp)
0 nM protein
3 nM protein
14 nM protein
70 nM protein
350 nM protein
2.0
0.0
−60
−40
−20
0
20
40
60
Position relative to center (bp)

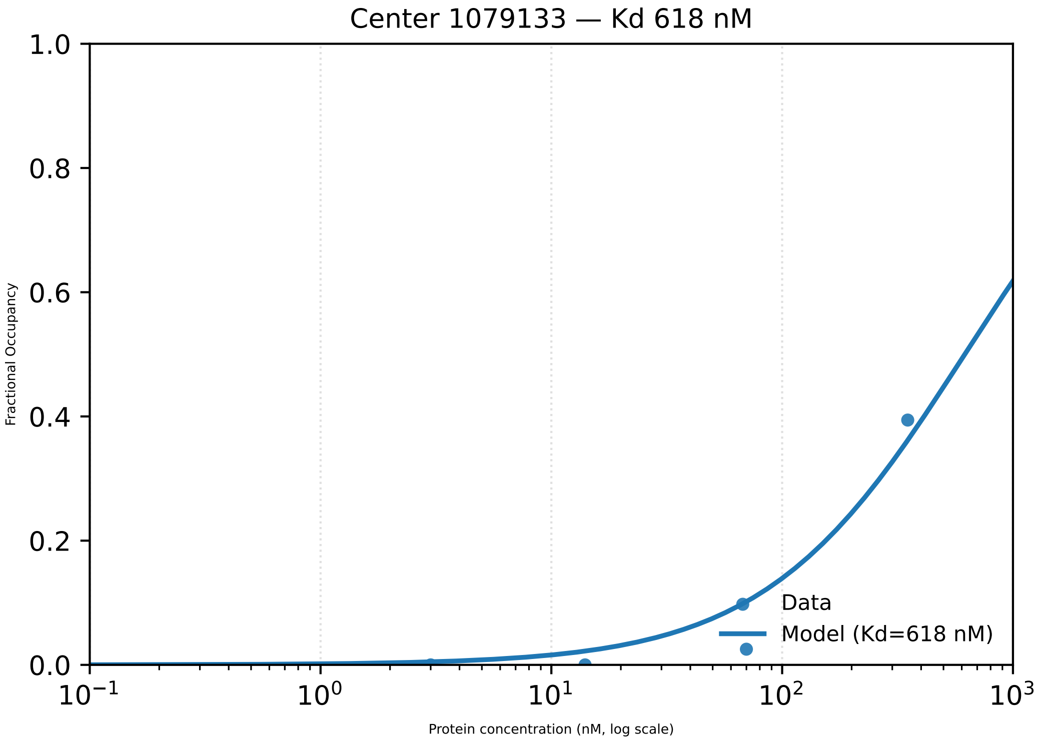

Center 1079133 — Kd 618 nM
Fractional Occupancy
1.0
0.8
0.6
0.4
0.2
0.0
$10^{-1}$
$10^{0}$
$10^{1}$
$10^{2}$
$10^{3}$
Protein concentration (nM, log scale)
Data
Model (Kd=618 nM)

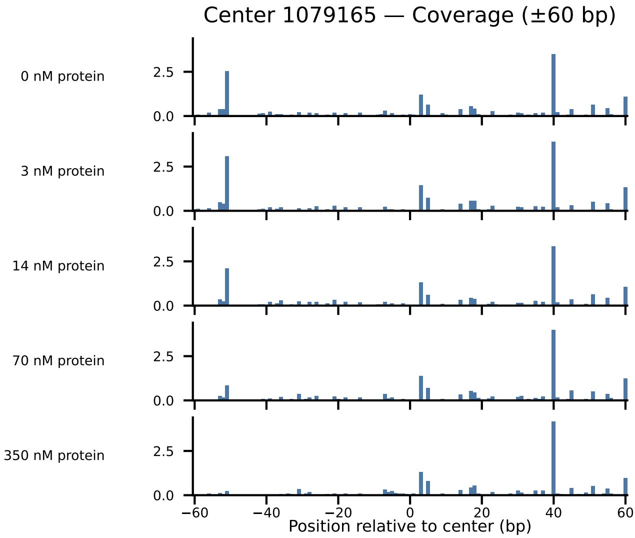

Center 1079165 — Coverage (±60 bp)
0 nM protein
3 nM protein
14 nM protein
70 nM protein
350 nM protein
2.5
0.0
−60
−40
−20
0
20
40
60
Position relative to center (bp)

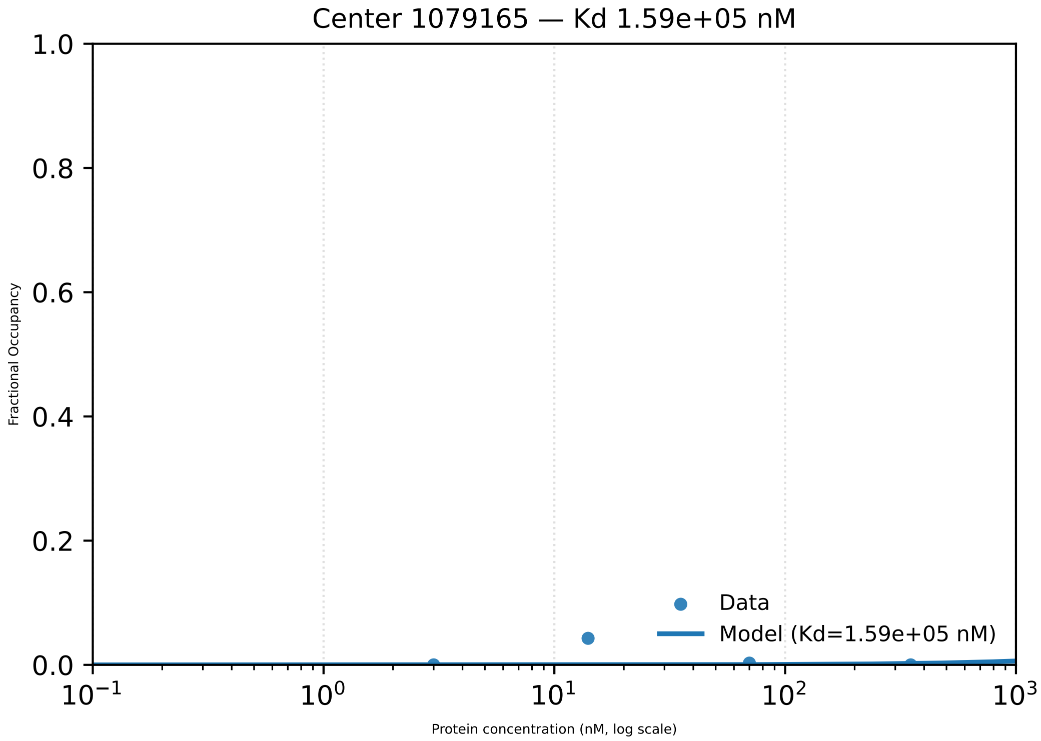

Center 1079165 — Kd 1.59e+05 nM
1.0
0.8
0.6
0.4
0.2
0.0
Fractional Occupancy
Data
Model (Kd=1.59e+05 nM)
$10^{-1}$
$10^{0}$
$10^{1}$
$10^{2}$
$10^{3}$
Protein concentration (nM, log scale)

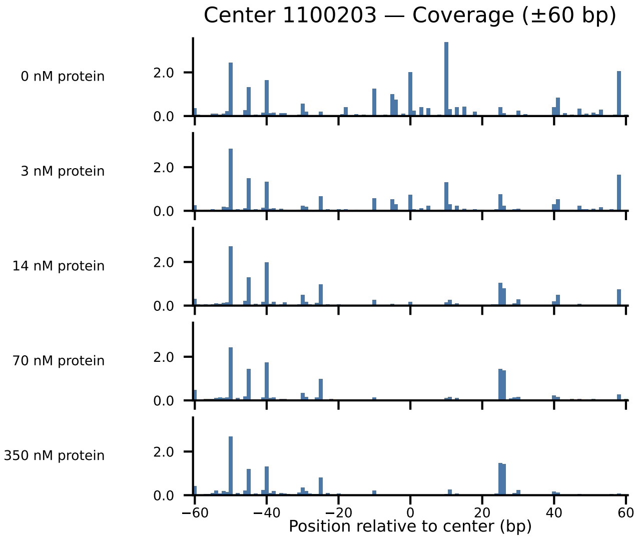
Center 1100203 — Coverage (±60 bp)
0 nM protein
3 nM protein
14 nM protein
70 nM protein
350 nM protein
2.0
0.0
−60
−40
−20
0
20
40
60
Position relative to center (bp)

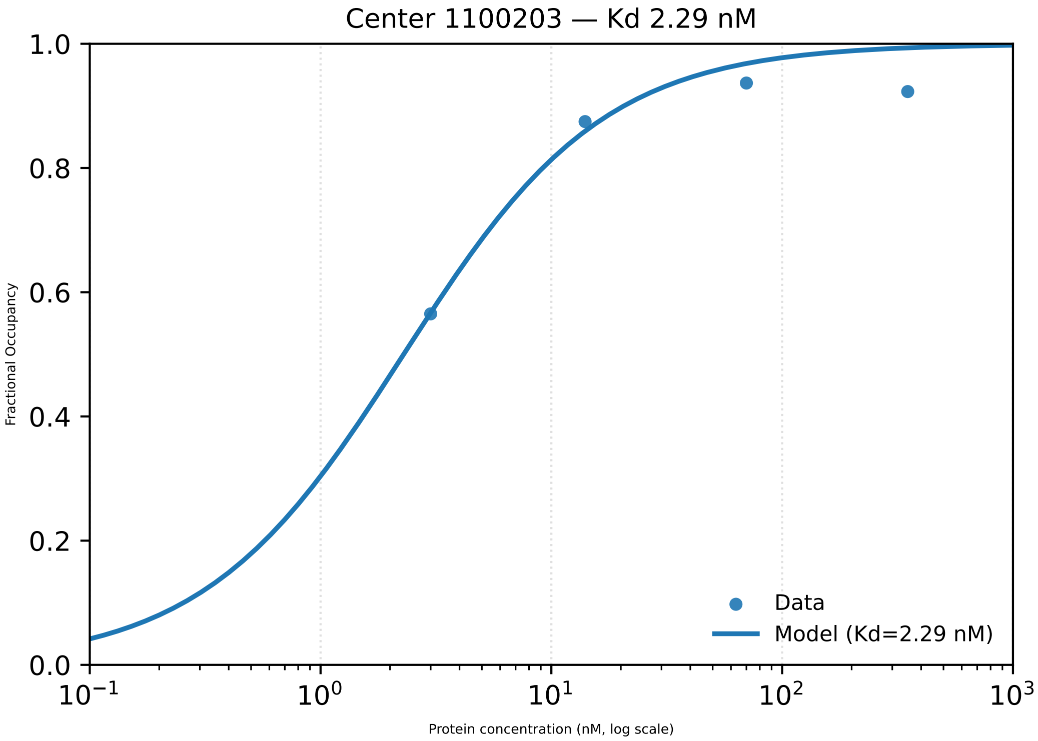
Center 1100203 — Kd 2.29 nM
1.0
0.8
0.6
0.4
0.2
0.0
Fractional Occupancy
$10^{-1}$
$10^{0}$
$10^{1}$
$10^{2}$
$10^{3}$
Protein concentration (nM, log scale)
Data
Model (Kd=2.29 nM)

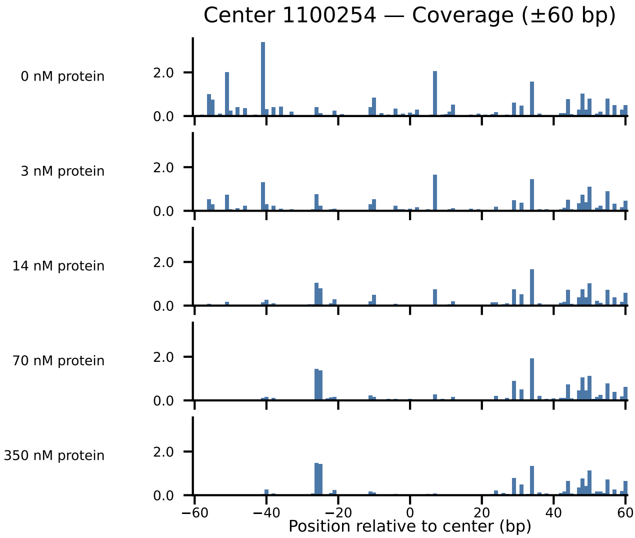
Center 1100254 — Coverage (±60 bp)
0 nM protein
3 nM protein
14 nM protein
70 nM protein
350 nM protein
2.0
0.0
−60
−40
−20
0
20
40
60
Position relative to center (bp)

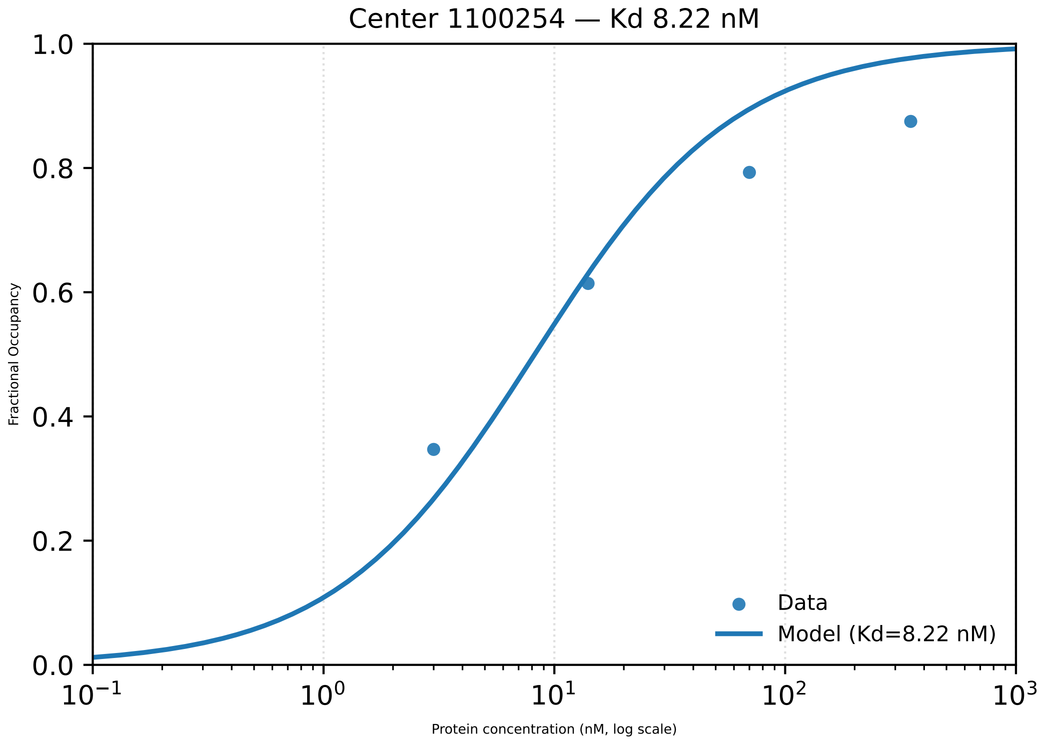

Center 1100254 — Kd 8.22 nM
Fractional Occupancy
1.0
0.8
0.6
0.4
0.2
0.0
$10^{-1}$
$10^{0}$
$10^{1}$
$10^{2}$
$10^{3}$
Protein concentration (nM, log scale)
Data
Model (Kd=8.22 nM)

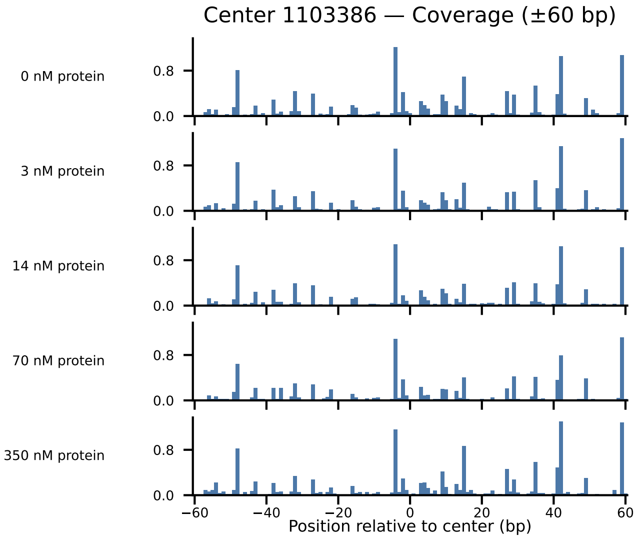

Center 1103386 — Coverage (±60 bp)
0 nM protein
3 nM protein
14 nM protein
70 nM protein
350 nM protein
0.8
0.0
−60
−40
−20
0
20
40
60
Position relative to center (bp)

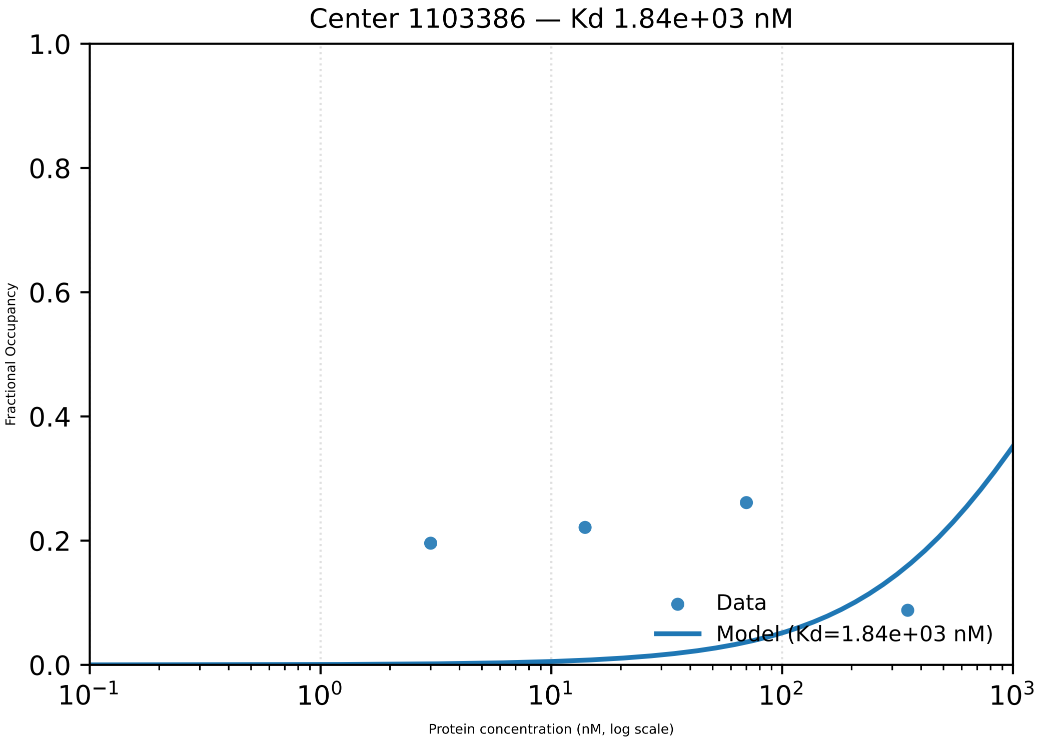

Center 1103386 — Kd 1.84e+03 nM
1.0
0.8
0.6
0.4
0.2
0.0
Fractional Occupancy
$10^{-1}$
$10^{0}$
$10^{1}$
$10^{2}$
$10^{3}$
Protein concentration (nM, log scale)
Data
Model (Kd=1.84e+03 nM)

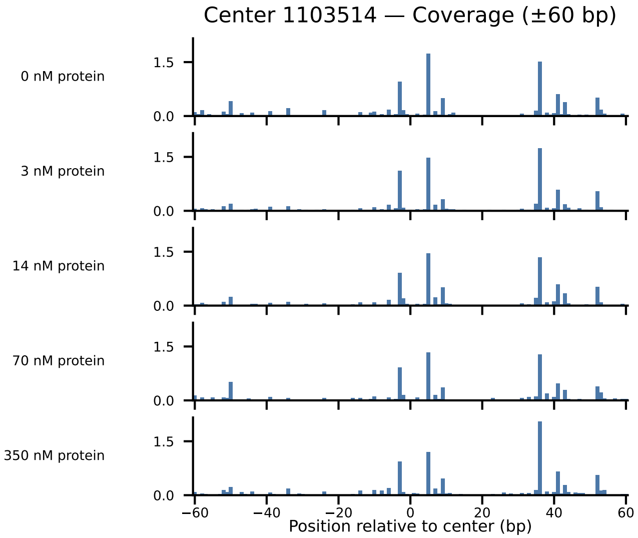
Center 1103514 — Coverage (±60 bp)
0 nM protein
3 nM protein
14 nM protein
70 nM protein
350 nM protein
1.5
0.0
−60
−40
−20
0
20
40
60
Position relative to center (bp)

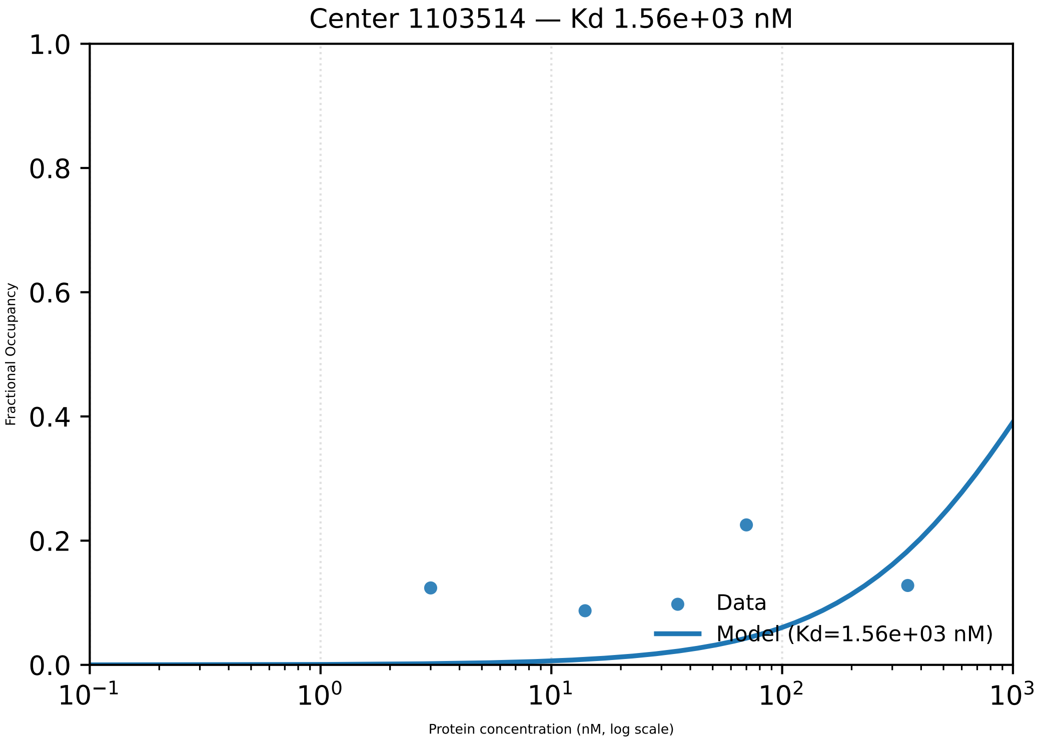
Center 1103514 — Kd 1.56e+03 nM
1.0
0.8
0.6
0.4
0.2
0.0
Fractional Occupancy
$10^{-1}$
$10^{0}$
$10^{1}$
$10^{2}$
$10^{3}$
Protein concentration (nM, log scale)
Data
Model (Kd=1.56e+03 nM)

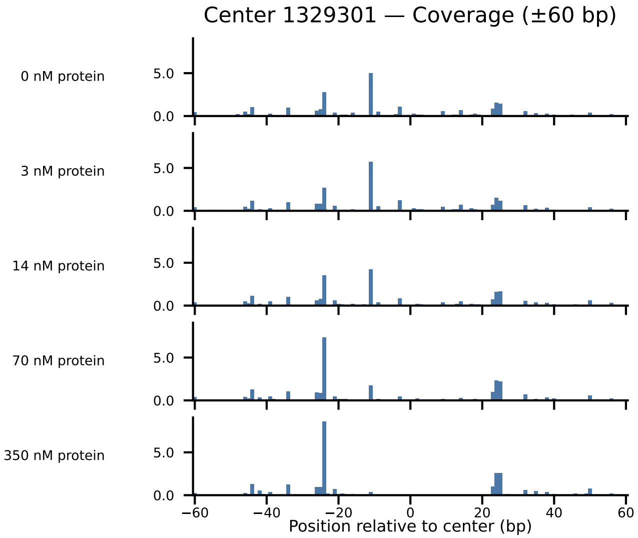

Center 1329301 — Coverage (±60 bp)
0 nM protein
3 nM protein
14 nM protein
70 nM protein
350 nM protein
5.0
0.0
−60
−40
−20
0
20
40
60
Position relative to center (bp)

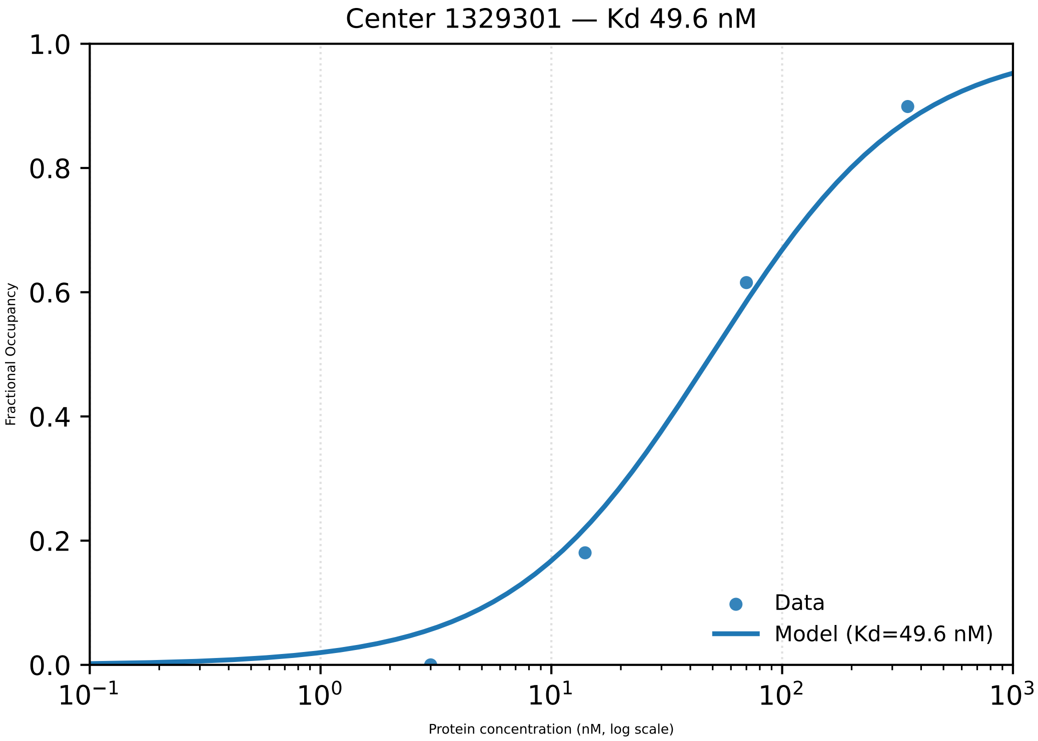
Center 1329301 — Kd 49.6 nM
1.0
0.8
0.6
0.4
0.2
0.0
Fractional Occupancy
$10^{-1}$
$10^{0}$
$10^{1}$
$10^{2}$
$10^{3}$
Protein concentration (nM, log scale)
Data
Model (Kd=49.6 nM)

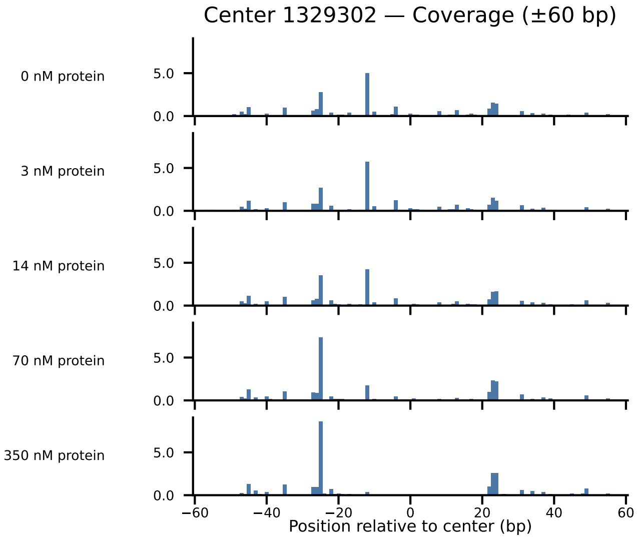

Center 1329302 — Coverage (±60 bp)
0 nM protein
3 nM protein
14 nM protein
70 nM protein
350 nM protein
5.0
0.0
−60
−40
−20
0
20
40
60
Position relative to center (bp)

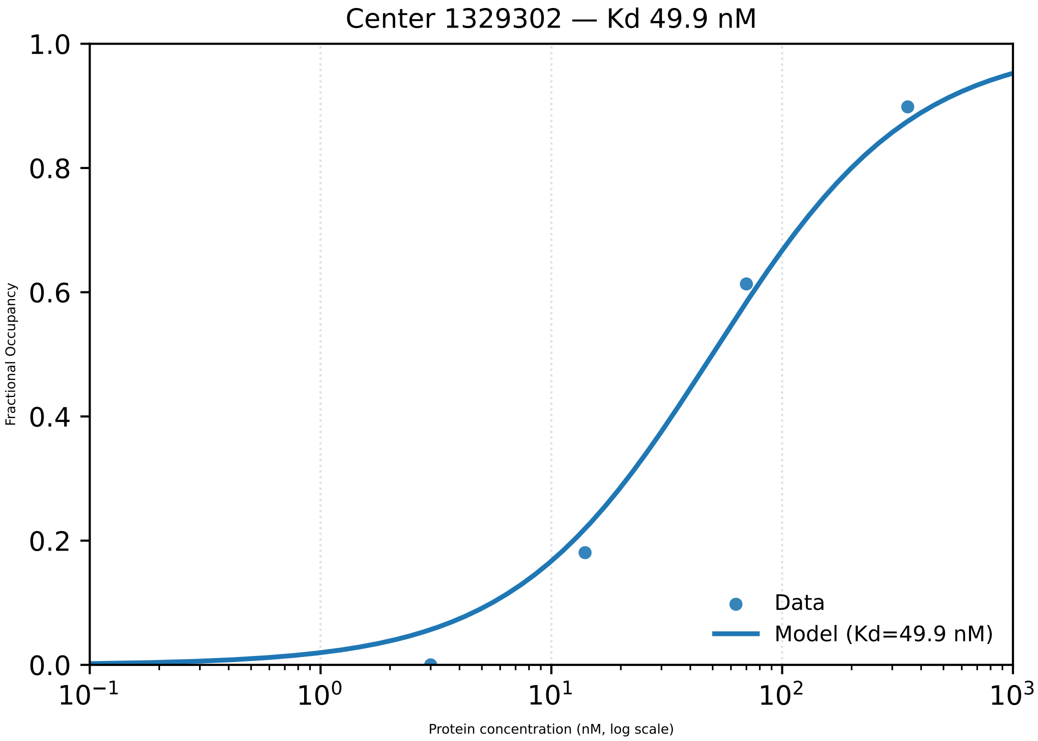

Center 1329302 — Kd 49.9 nM
1.0
0.8
0.6
0.4
0.2
0.0
Fractional Occupancy
$10^{-1}$
$10^{0}$
$10^{1}$
$10^{2}$
$10^{3}$
Protein concentration (nM, log scale)
Data
Model (Kd=49.9 nM)

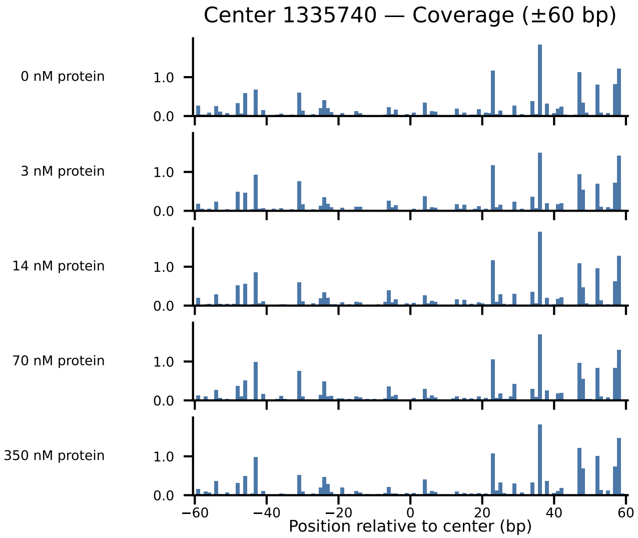

Center 1335740 — Coverage (±60 bp)
0 nM protein
3 nM protein
14 nM protein
70 nM protein
350 nM protein
1.0
0.0
−60
−40
−20
0
20
40
60
Position relative to center (bp)

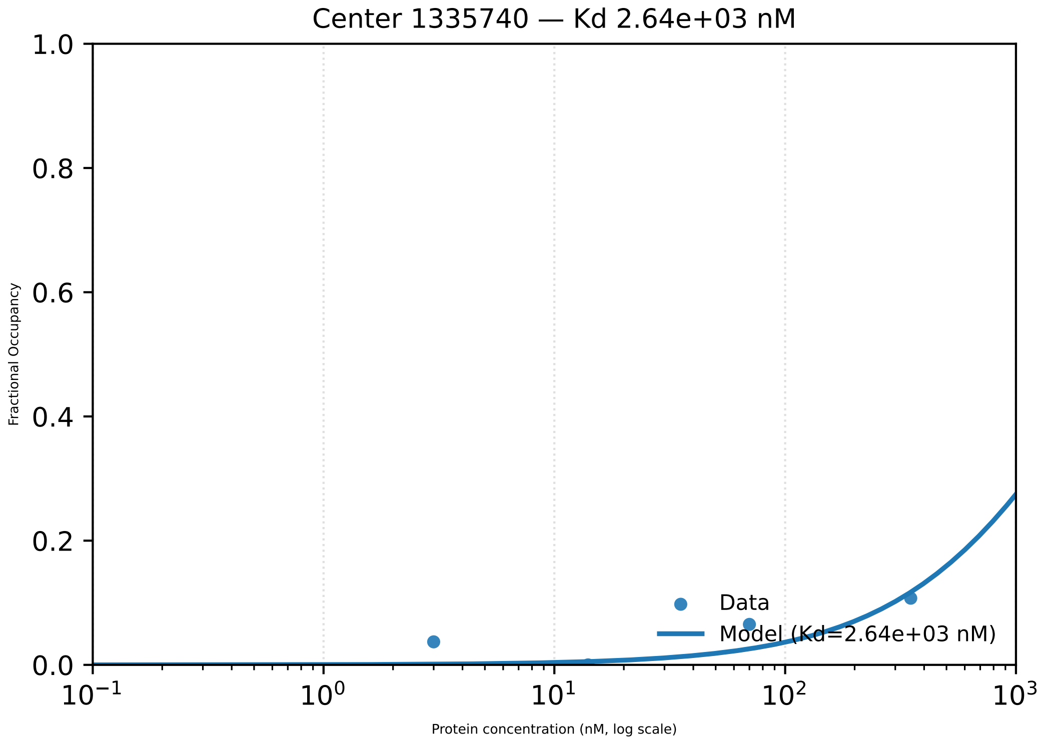

Center 1335740 — Kd 2.64e+03 nM
1.0
0.8
0.6
0.4
0.2
0.0
Fractional Occupancy
$10^{-1}$
$10^{0}$
$10^{1}$
$10^{2}$
$10^{3}$
Protein concentration (nM, log scale)
Data
Model (Kd=2.64e+03 nM)

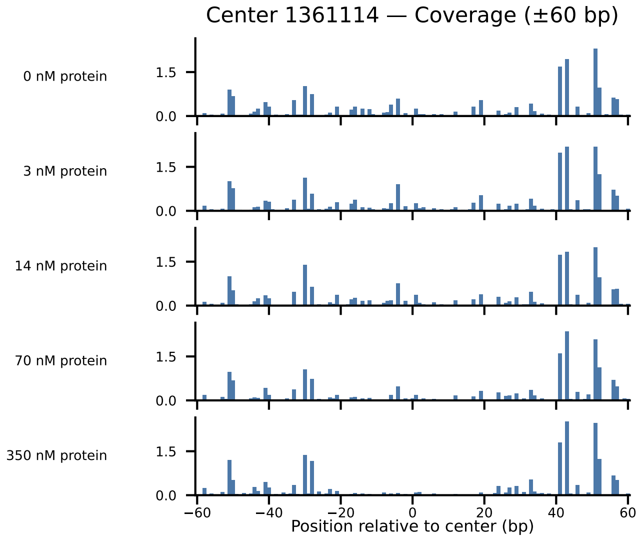

Center 1361114 — Coverage (±60 bp)
0 nM protein
3 nM protein
14 nM protein
70 nM protein
350 nM protein
1.5
0.0
−60
−40
−20
0
20
40
60
Position relative to center (bp)

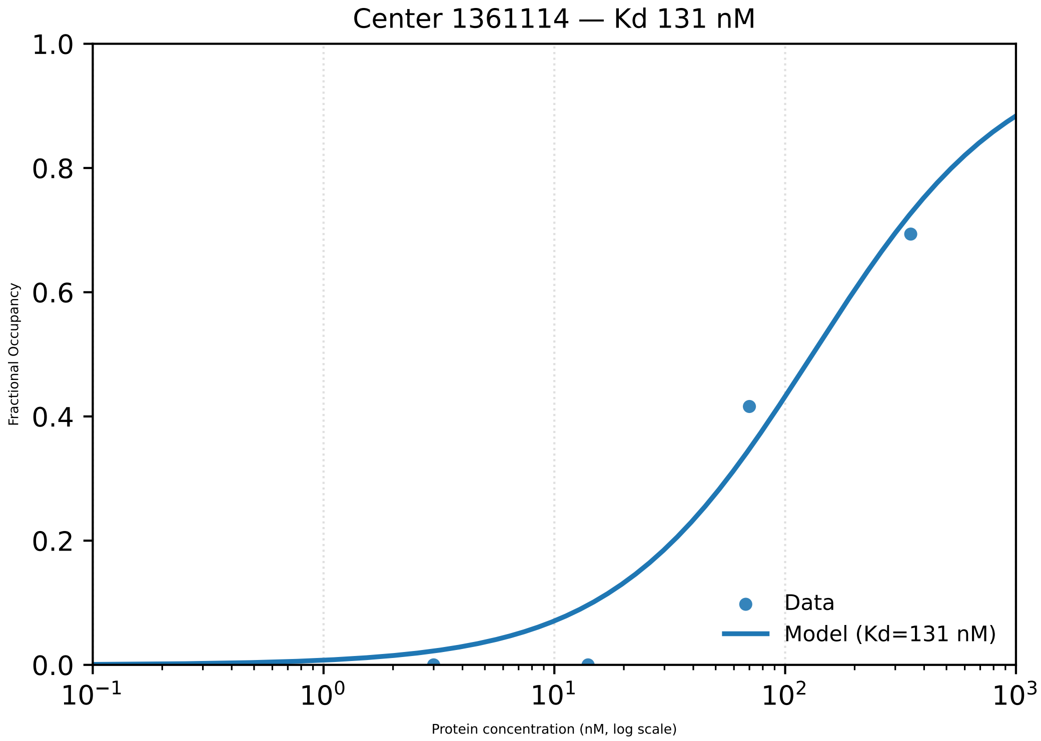
Center 1361114 — Kd 131 nM
Fractional Occupancy
Protein concentration (nM, log scale)
Data
Model (Kd=131 nM)
0.0
0.2
0.4
0.6
0.8
1.0
$10^{-1}$
$10^{0}$
$10^{1}$
$10^{2}$
$10^{3}$

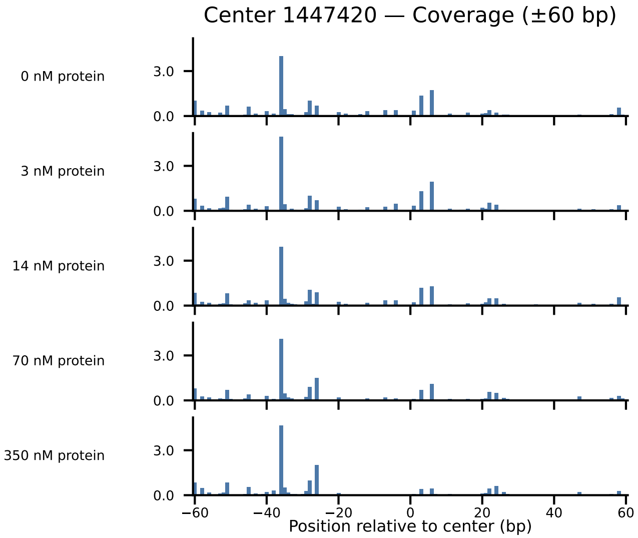

Center 1447420 — Coverage (±60 bp)
0 nM protein
3 nM protein
14 nM protein
70 nM protein
350 nM protein
3.0
0.0
−60
−40
−20
0
20
40
60
Position relative to center (bp)

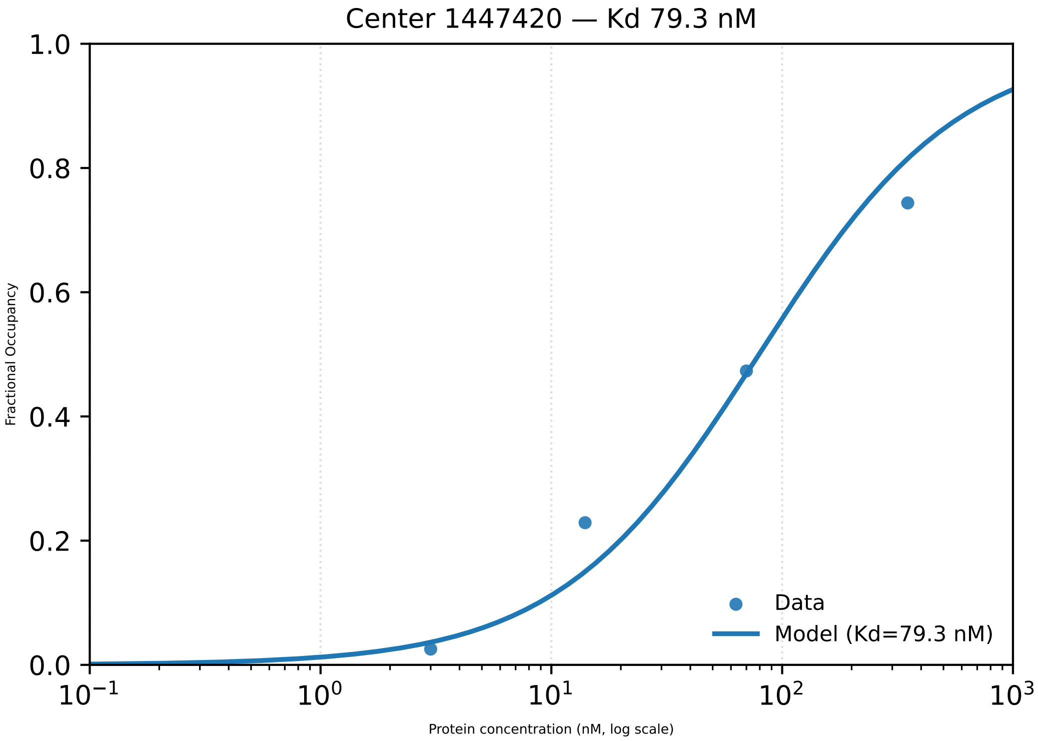
Center 1447420 — Kd 79.3 nM
Fractional Occupancy
Protein concentration (nM, log scale)
Data
Model (Kd=79.3 nM)
1.0
0.8
0.6
0.4
0.2
0.0
$10^{-1}$
$10^{0}$
$10^{1}$
$10^{2}$
$10^{3}$

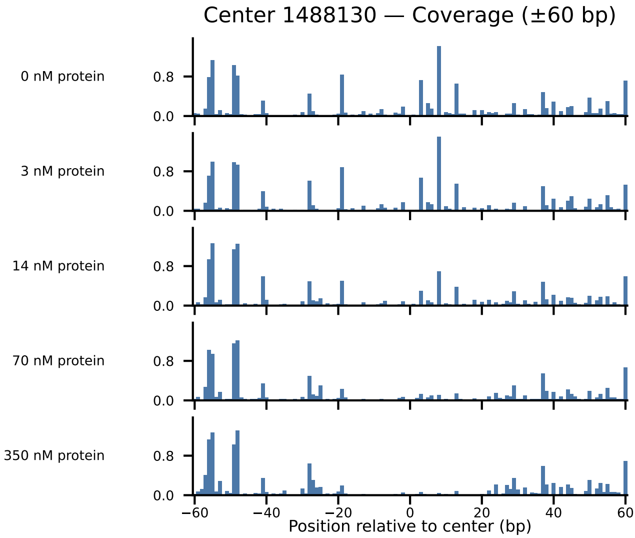

Center 1488130 — Coverage (±60 bp)
0 nM protein
3 nM protein
14 nM protein
70 nM protein
350 nM protein
0.8
0.0
−60
−40
−20
0
20
40
60
Position relative to center (bp)

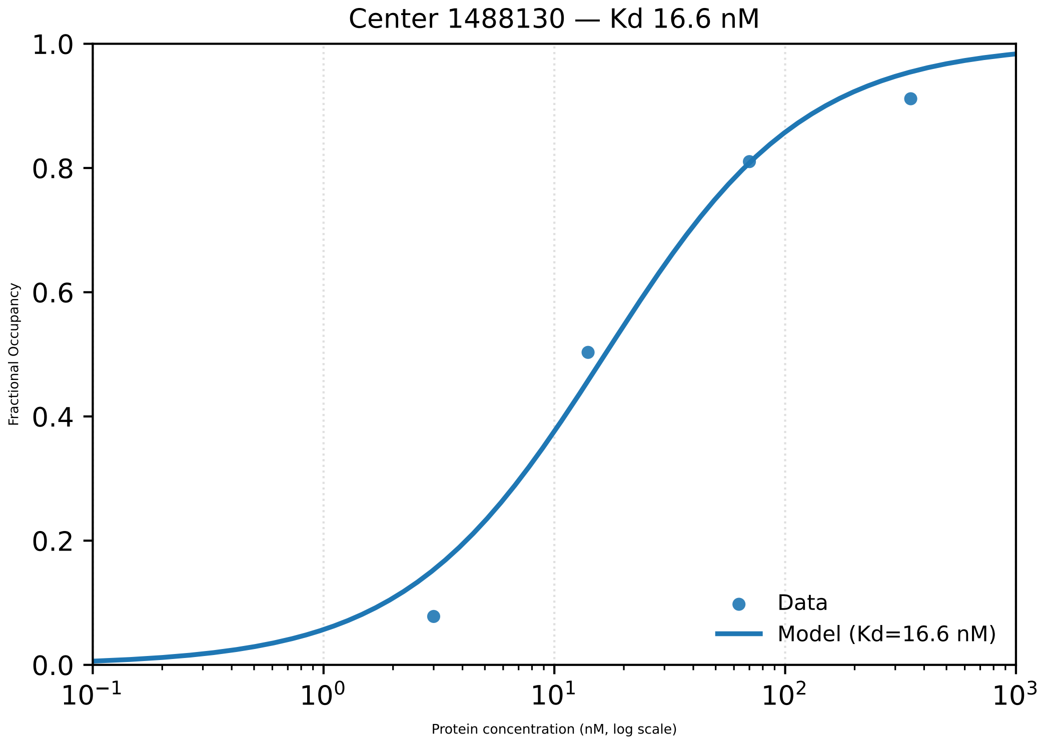

Center 1488130 — Kd 16.6 nM
1.0
0.8
0.6
0.4
0.2
0.0
Fractional Occupancy
Data
Model (Kd=16.6 nM)
$10^{-1}$
$10^{0}$
$10^{1}$
$10^{2}$
$10^{3}$
Protein concentration (nM, log scale)

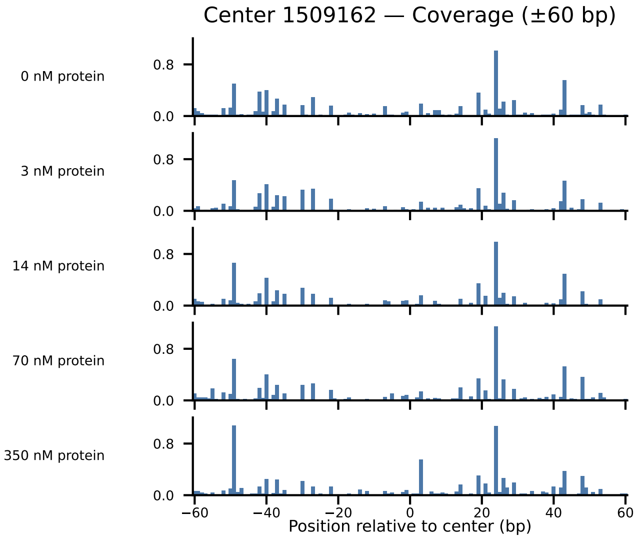

Center 1509162 — Coverage (±60 bp)
0 nM protein
3 nM protein
14 nM protein
70 nM protein
350 nM protein
0.8
0.0
−60
−40
−20
0
20
40
60
Position relative to center (bp)

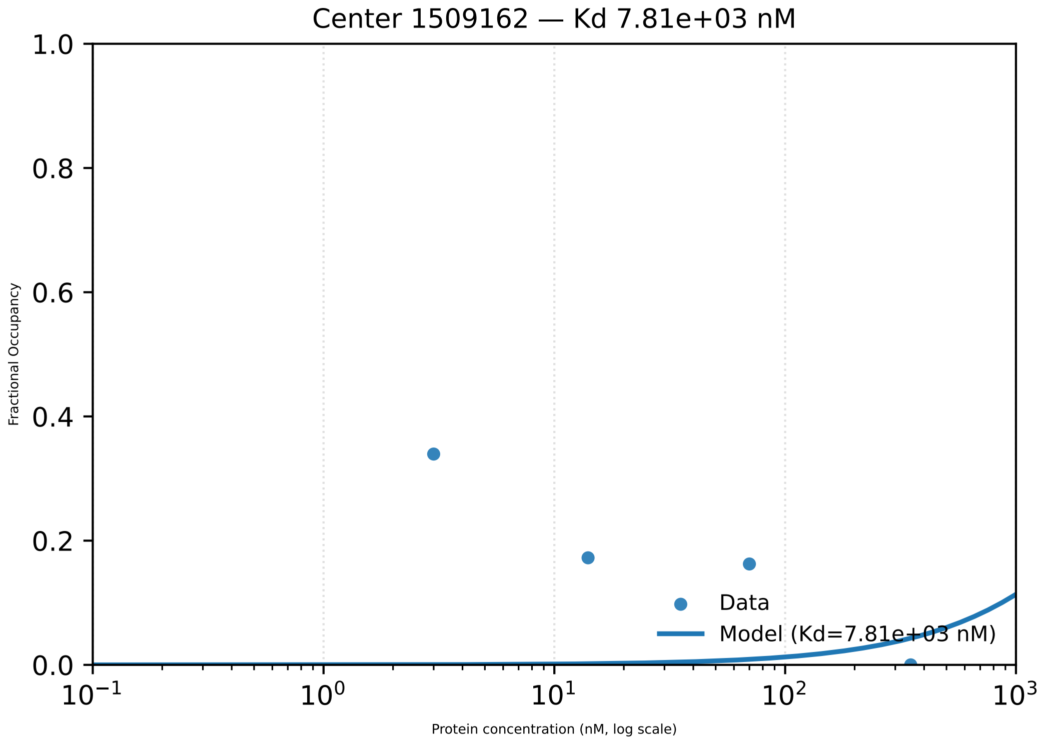

Center 1509162 — Kd 7.81e+03 nM
1.0
0.8
0.6
0.4
0.2
0.0
Fractional Occupancy
$10^{-1}$
$10^{0}$
$10^{1}$
$10^{2}$
$10^{3}$
Protein concentration (nM, log scale)
Data
Model (Kd=7.81e+03 nM)

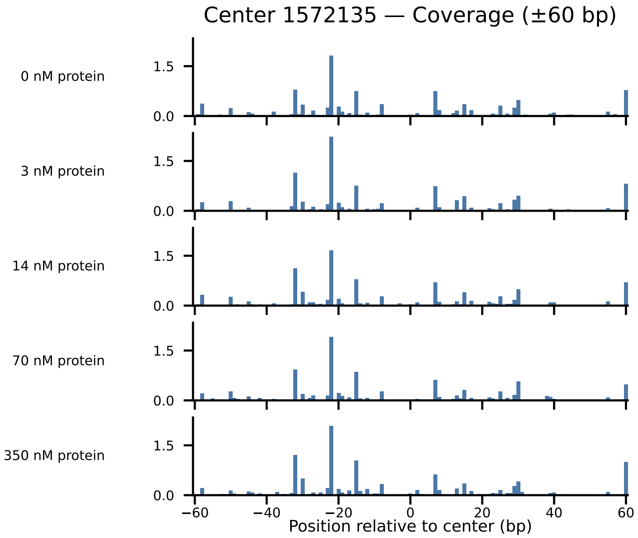

Center 1572135 — Coverage (±60 bp)
0 nM protein
3 nM protein
14 nM protein
70 nM protein
350 nM protein
1.5
0.0
−60
−40
−20
0
20
40
60
Position relative to center (bp)

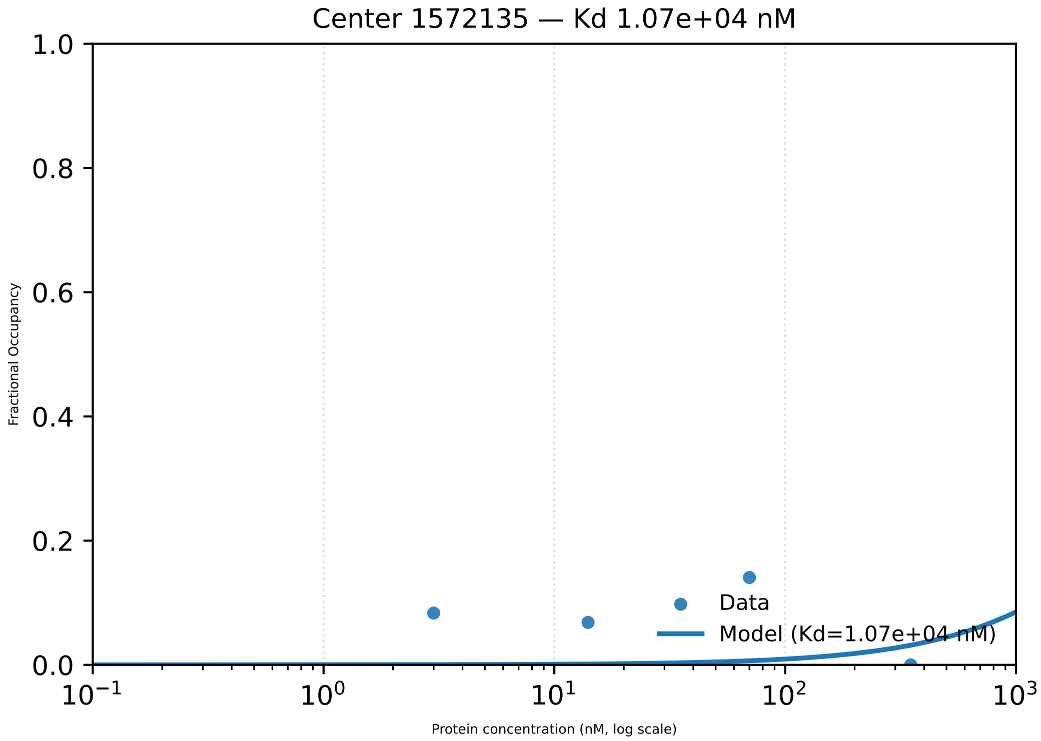

Center 1572135 — Kd 1.07e+04 nM
Fractional Occupancy
1.0
0.8
0.6
0.4
0.2
0.0
$10^{-1}$
$10^{0}$
$10^{1}$
$10^{2}$
$10^{3}$
Protein concentration (nM, log scale)
Data
Model (Kd=1.07e+04 nM)

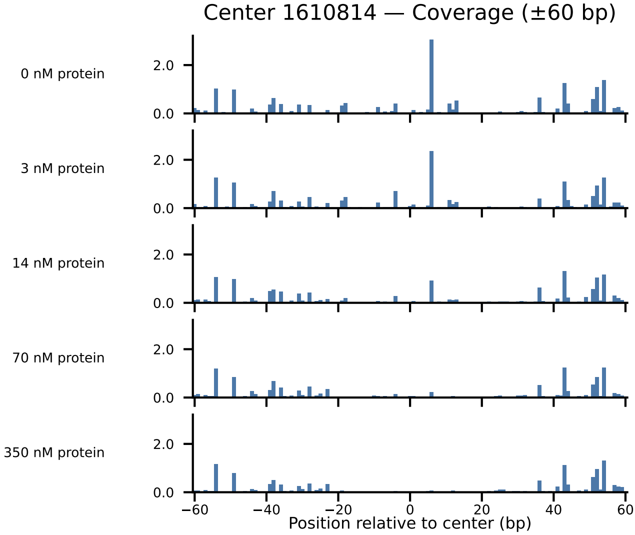
Center 1610814 — Coverage (±60 bp)
0 nM protein
3 nM protein
14 nM protein
70 nM protein
350 nM protein
2.0
0.0
−60
−40
−20
0
20
40
60
Position relative to center (bp)

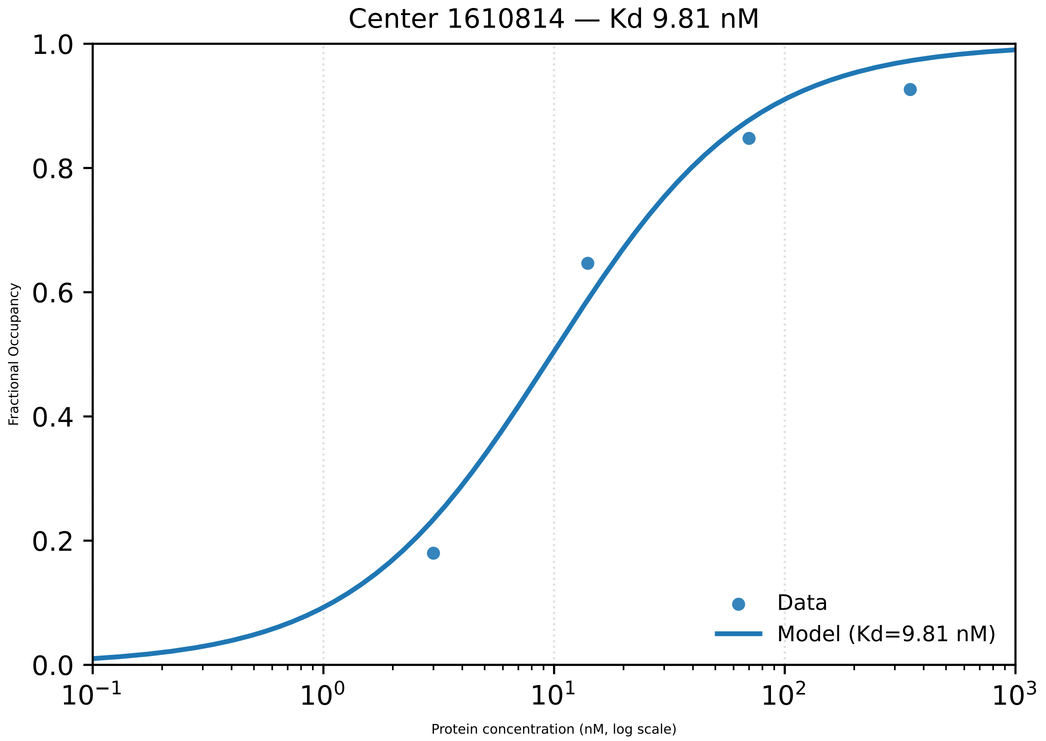

Center 1610814 — Kd 9.81 nM
Fractional Occupancy
0.0
0.2
0.4
0.6
0.8
1.0
$10^{-1}$
$10^{0}$
$10^{1}$
$10^{2}$
$10^{3}$
Protein concentration (nM, log scale)
Data
Model (Kd=9.81 nM)

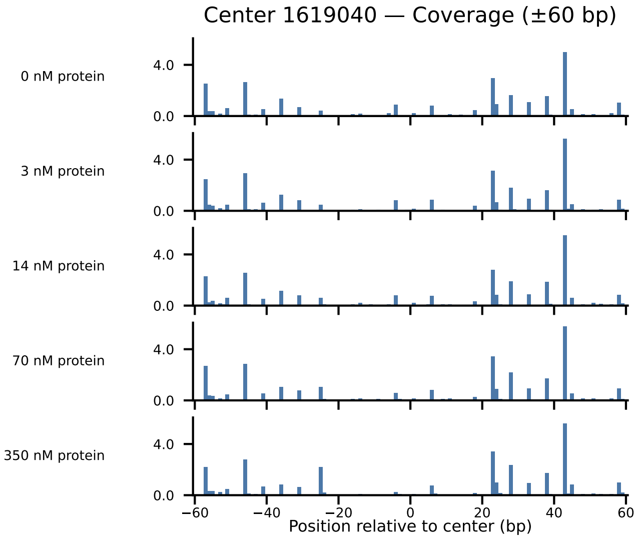

Center 1619040 — Coverage (±60 bp)
0 nM protein
3 nM protein
14 nM protein
70 nM protein
350 nM protein
4.0
0.0
−60
−40
−20
0
20
40
60
Position relative to center (bp)

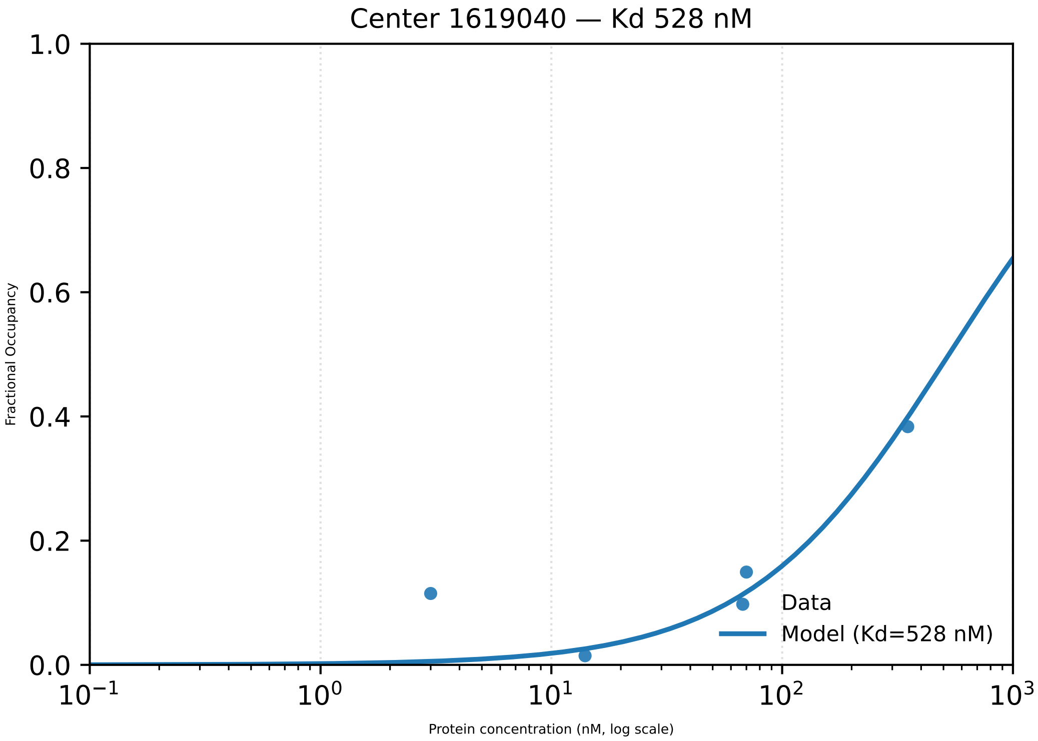

Center 1619040 — Kd 528 nM
1.0
0.8
0.6
0.4
0.2
0.0
Fractional Occupancy
$10^{-1}$
$10^{0}$
$10^{1}$
$10^{2}$
$10^{3}$
Protein concentration (nM, log scale)
Data
Model (Kd=528 nM)

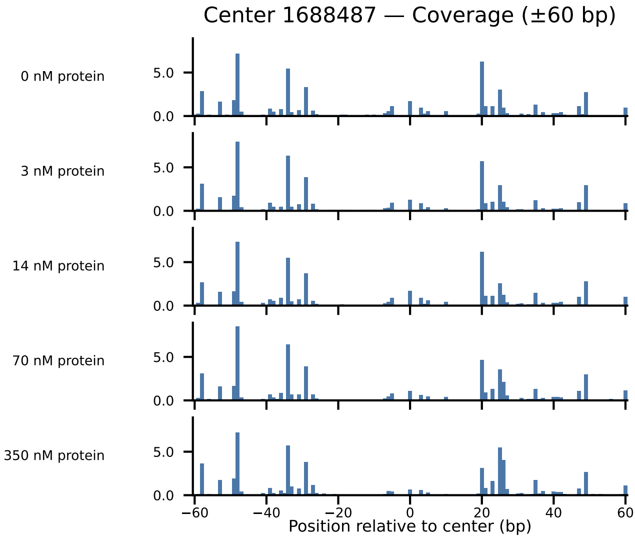

Center 1688487 — Coverage (±60 bp)
0 nM protein
3 nM protein
14 nM protein
70 nM protein
350 nM protein
5.0
0.0
−60
−40
−20
0
20
40
60
Position relative to center (bp)

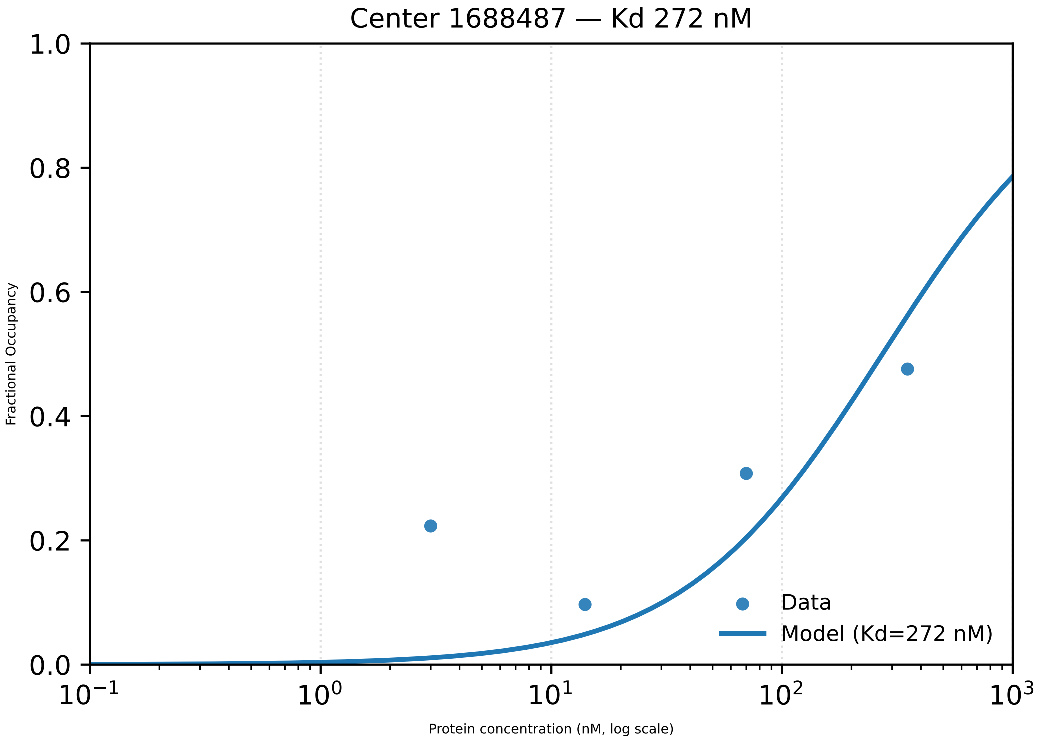

Center 1688487 — Kd 272 nM
1.0
0.8
0.6
0.4
0.2
0.0
Fractional Occupancy
$10^{-1}$
$10^{0}$
$10^{1}$
$10^{2}$
$10^{3}$
Protein concentration (nM, log scale)
Data
Model (Kd=272 nM)

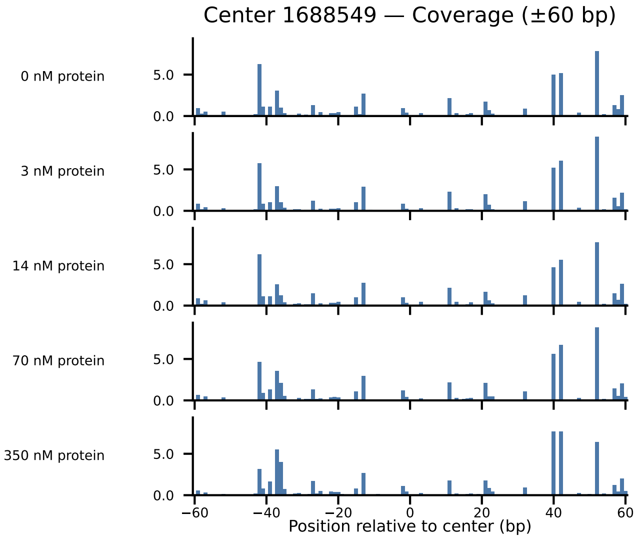
Center 1688549 — Coverage (±60 bp)
0 nM protein
3 nM protein
14 nM protein
70 nM protein
350 nM protein
5.0
0.0
−60
−40
−20
0
20
40
60
Position relative to center (bp)

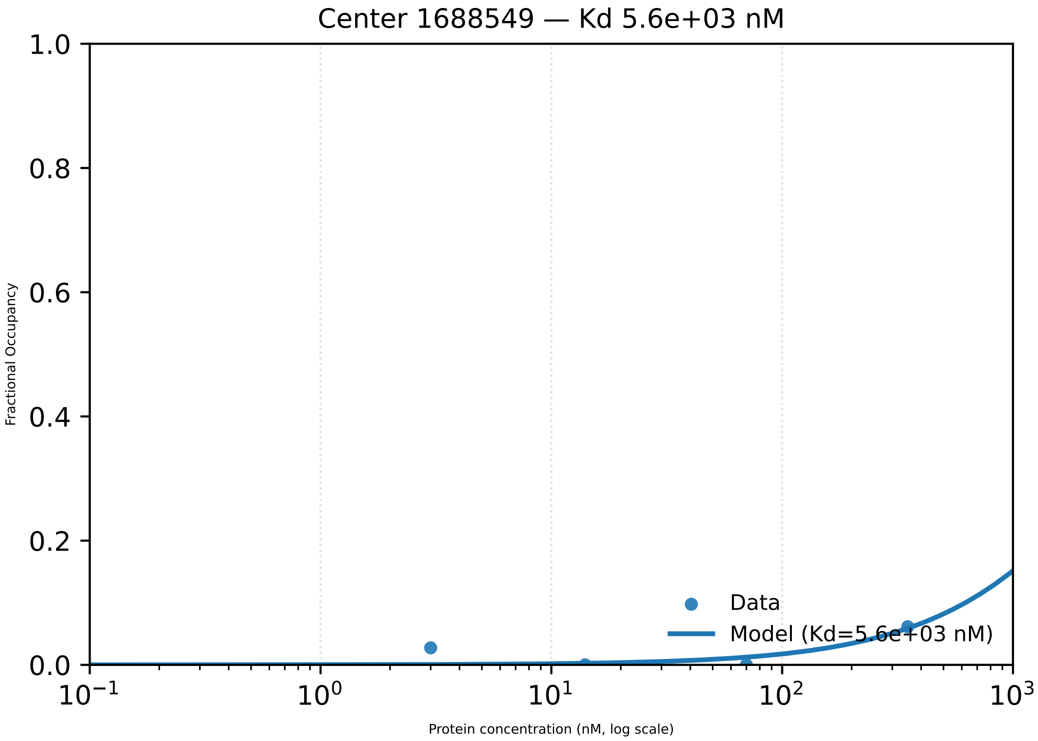
Center 1688549 — Kd 5.6e+03 nM
Fractional Occupancy
1.0
0.8
0.6
0.4
0.2
0.0
$10^{-1}$
$10^{0}$
$10^{1}$
$10^{2}$
$10^{3}$
Protein concentration (nM, log scale)
Data
Model (Kd=5.6e+03 nM)

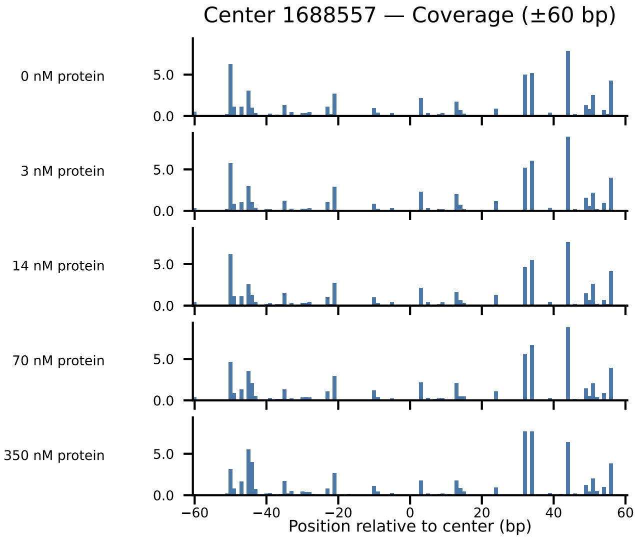

Center 1688557 — Coverage (±60 bp)
0 nM protein
3 nM protein
14 nM protein
70 nM protein
350 nM protein
5.0
0.0
−60
−40
−20
0
20
40
60
Position relative to center (bp)

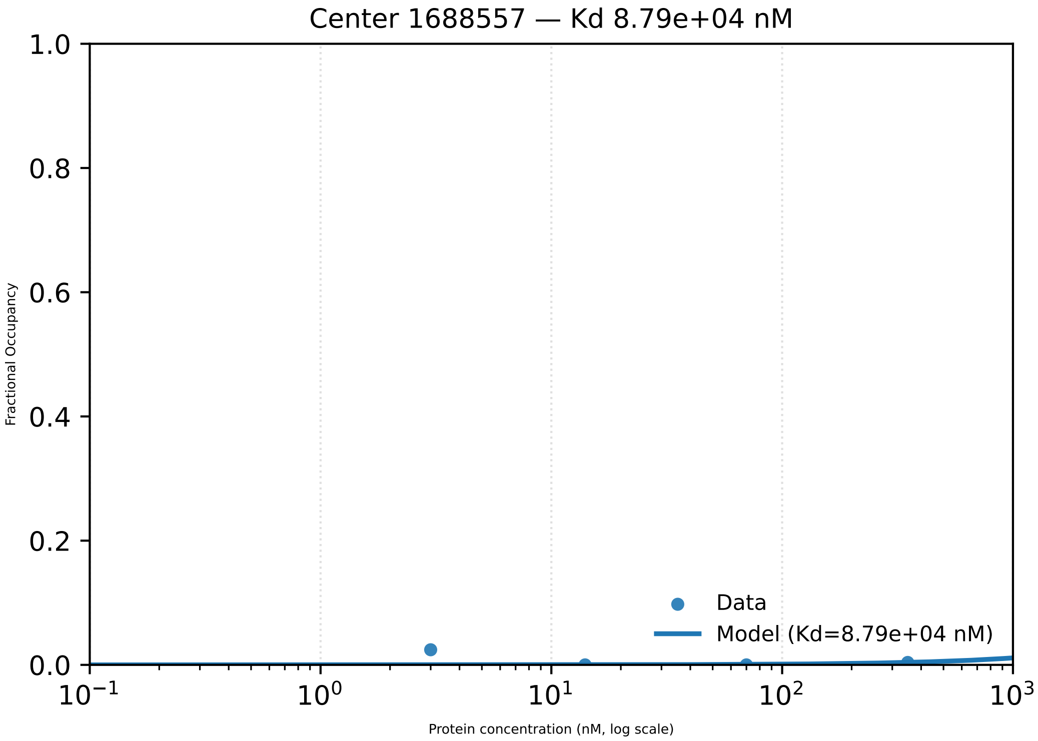
Center 1688557 — Kd 8.79e+04 nM
1.0
0.8
0.6
0.4
0.2
0.0
Fractional Occupancy
Data
Model (Kd=8.79e+04 nM)
$10^{-1}$
$10^{0}$
$10^{1}$
$10^{2}$
$10^{3}$
Protein concentration (nM, log scale)

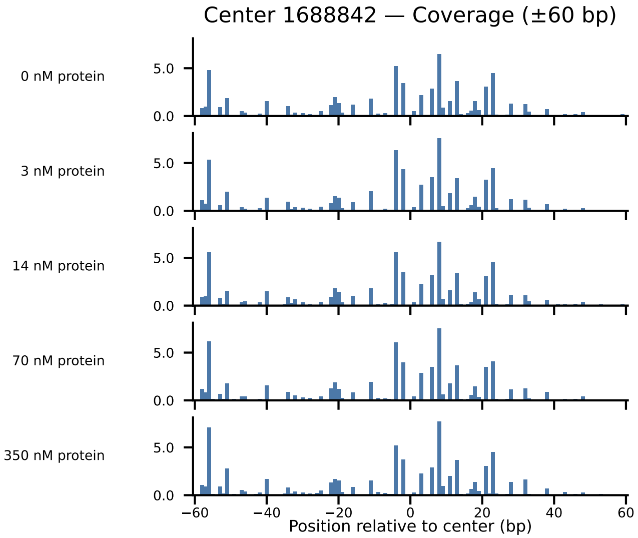

Center 1688842 — Coverage (±60 bp)
0 nM protein
3 nM protein
14 nM protein
70 nM protein
350 nM protein
5.0
0.0
−60
−40
−20
0
20
40
60
Position relative to center (bp)

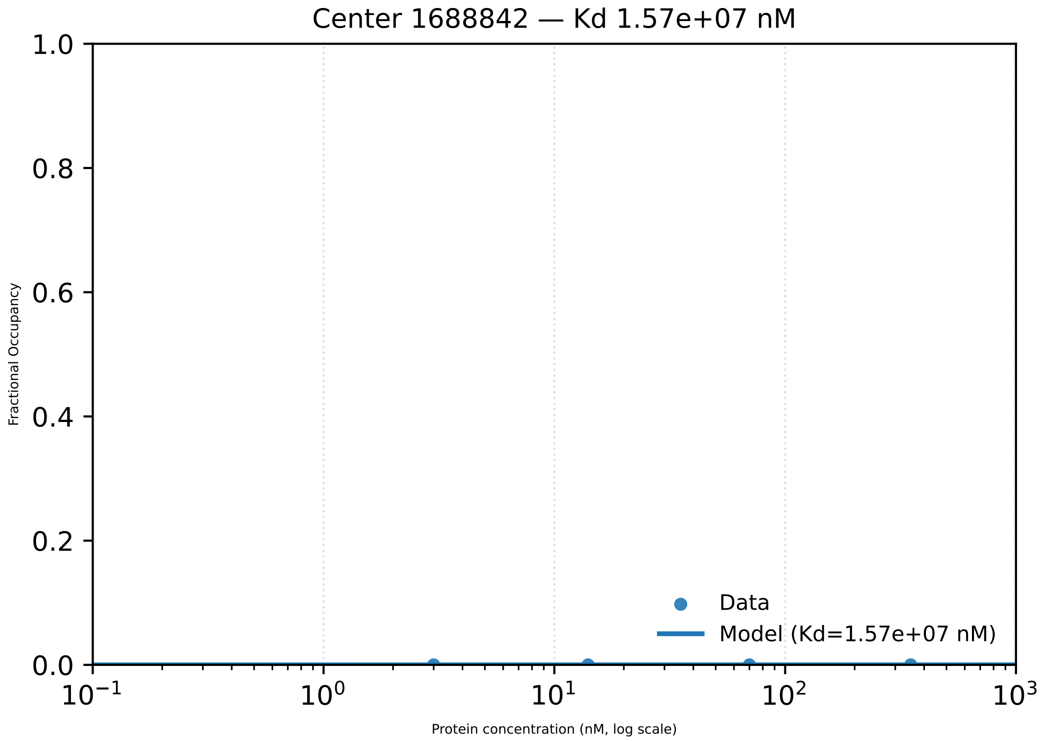
Center 1688842 — Kd 1.57e+07 nM
Fractional Occupancy
1.0
0.8
0.6
0.4
0.2
0.0
$10^{-1}$
$10^{0}$
$10^{1}$
$10^{2}$
$10^{3}$
Protein concentration (nM, log scale)
Data
Model (Kd=1.57e+07 nM)

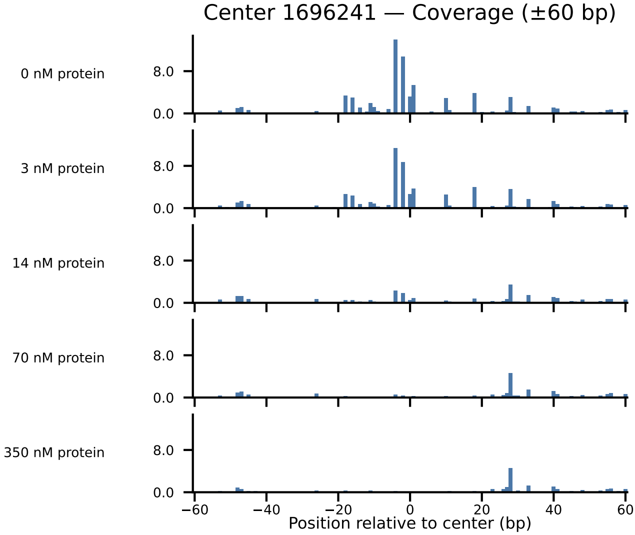

Center 1696241 — Coverage (±60 bp)
0 nM protein
3 nM protein
14 nM protein
70 nM protein
350 nM protein
8.0
0.0
−60
−40
−20
0
20
40
60
Position relative to center (bp)

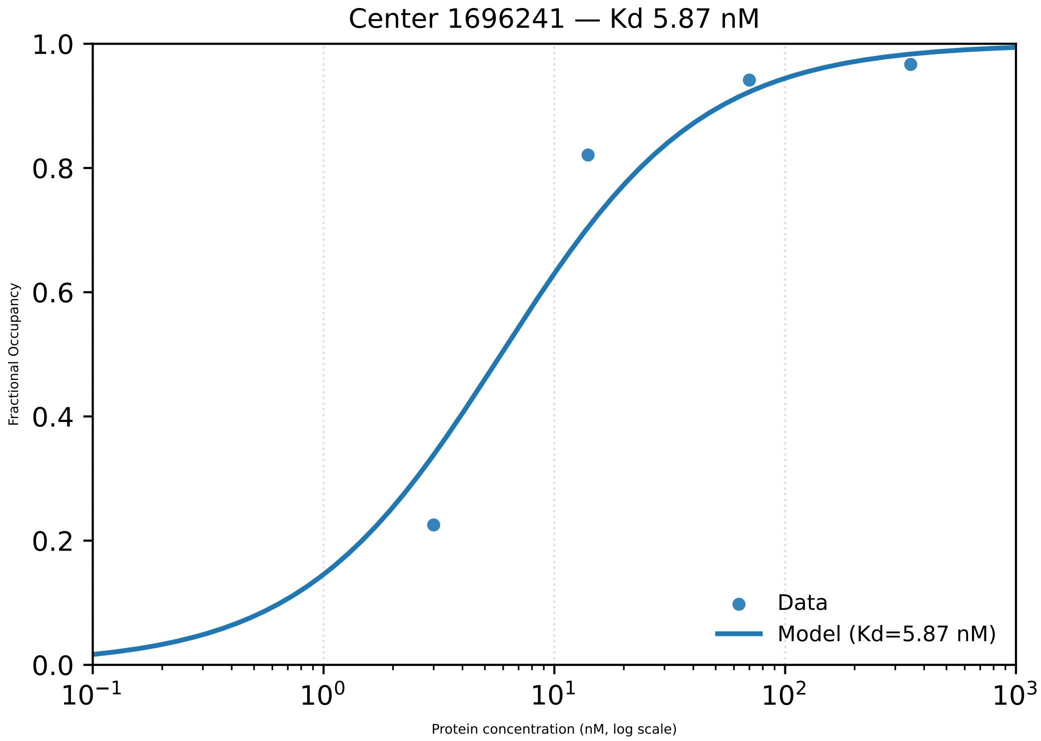

Center 1696241 — Kd 5.87 nM
Fractional Occupancy
1.0
0.8
0.6
0.4
0.2
0.0
$10^{-1}$
$10^{0}$
$10^{1}$
$10^{2}$
$10^{3}$
Protein concentration (nM, log scale)
Data
Model (Kd=5.87 nM)

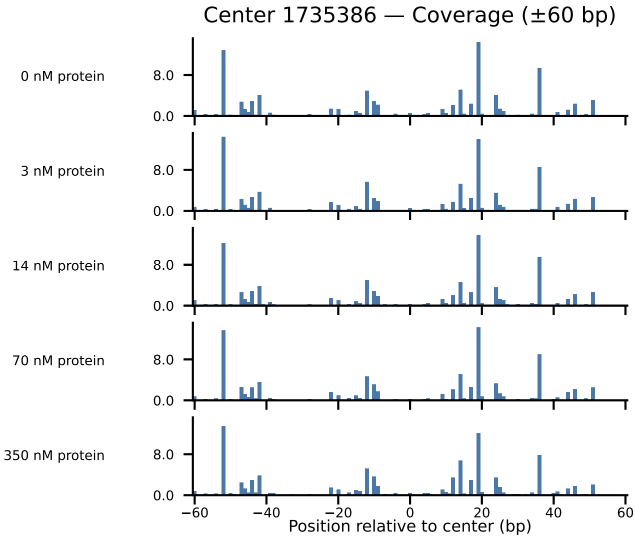
Center 1735386 — Coverage (±60 bp)
0 nM protein
3 nM protein
14 nM protein
70 nM protein
350 nM protein
8.0
0.0
−60
−40
−20
0
20
40
60
Position relative to center (bp)

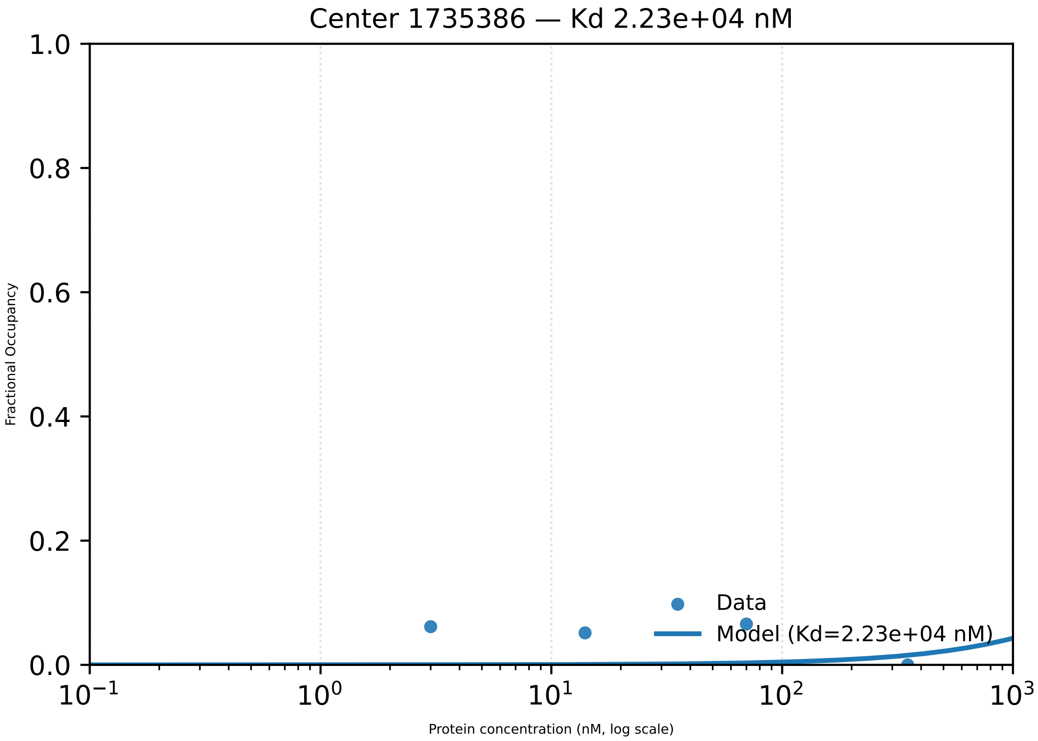

Center 1735386 — Kd 2.23e+04 nM
1.0
0.8
0.6
0.4
0.2
0.0
Fractional Occupancy
Data
Model (Kd=2.23e+04 nM)
$10^{-1}$
$10^{0}$
$10^{1}$
$10^{2}$
$10^{3}$
Protein concentration (nM, log scale)

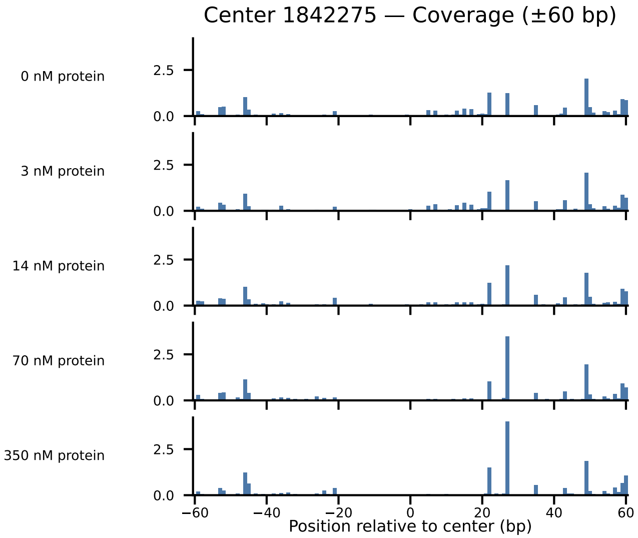
Center 1842275 — Coverage (±60 bp)
0 nM protein
3 nM protein
14 nM protein
70 nM protein
350 nM protein
2.5
0.0
−60
−40
−20
0
20
40
60
Position relative to center (bp)

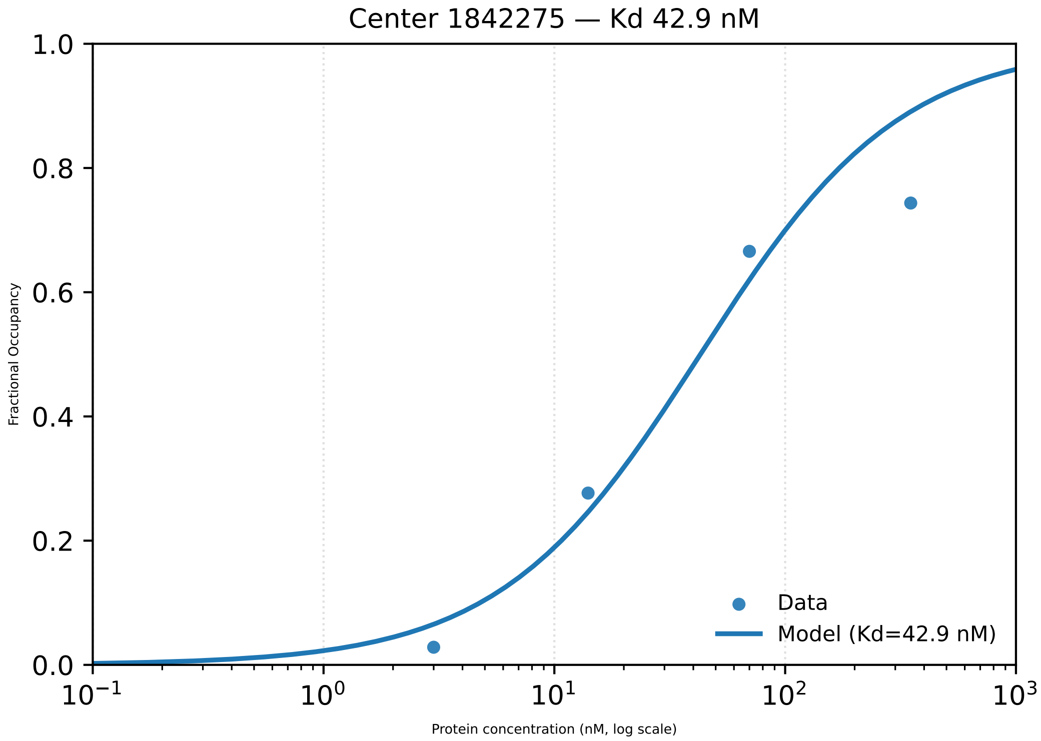

Center 1842275 — Kd 42.9 nM
1.0
0.8
0.6
0.4
0.2
0.0
Fractional Occupancy
$10^{-1}$
$10^{0}$
$10^{1}$
$10^{2}$
$10^{3}$
Protein concentration (nM, log scale)
Data
Model (Kd=42.9 nM)

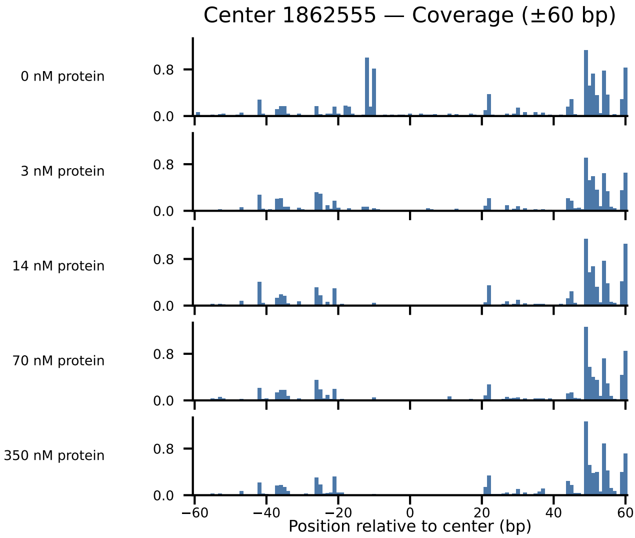

Center 1862555 — Coverage (±60 bp)
0 nM protein
3 nM protein
14 nM protein
70 nM protein
350 nM protein
0.8
0.0
−60
−40
−20
0
20
40
60
Position relative to center (bp)

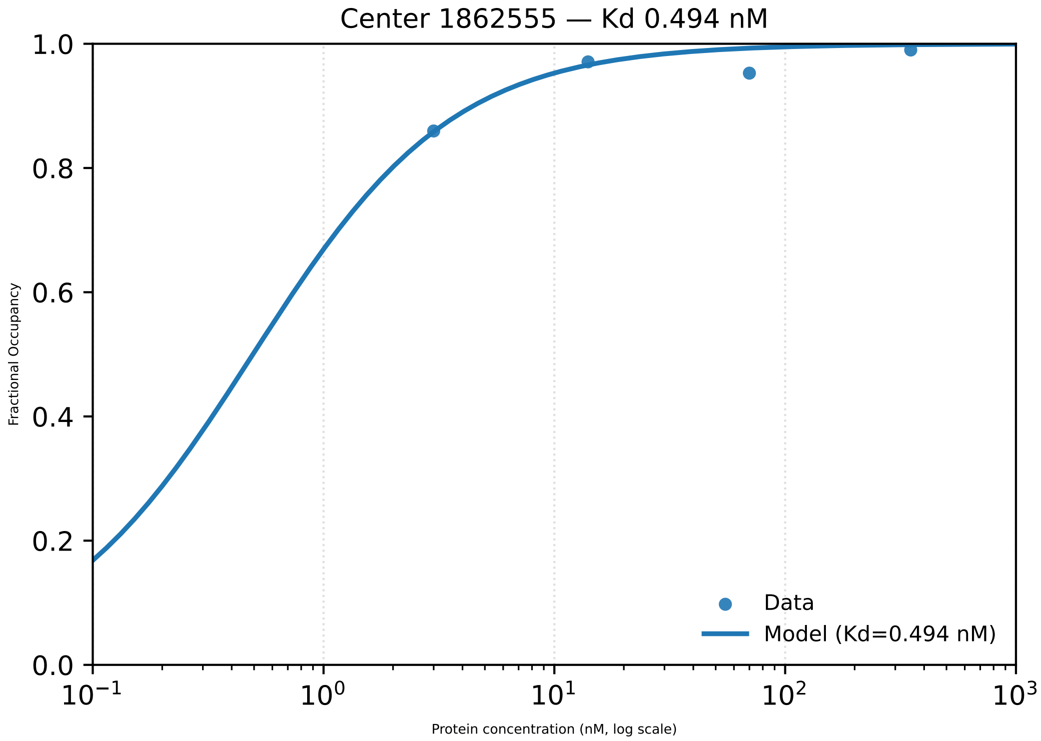
Center 1862555 — Kd 0.494 nM
1.0
0.8
0.6
0.4
0.2
0.0
Fractional Occupancy
$10^{-1}$
$10^{0}$
$10^{1}$
$10^{2}$
$10^{3}$
Protein concentration (nM, log scale)
Data
Model (Kd=0.494 nM)

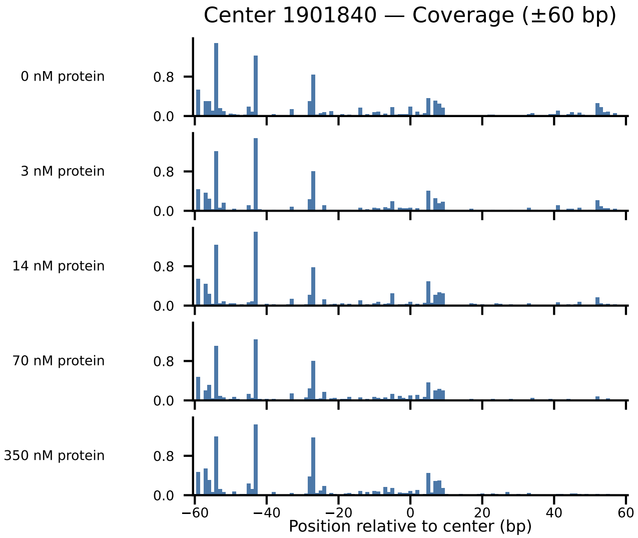

Center 1901840 — Coverage (±60 bp)
0 nM protein
3 nM protein
14 nM protein
70 nM protein
350 nM protein
0.8
0.0
−60
−40
−20
0
20
40
60
Position relative to center (bp)

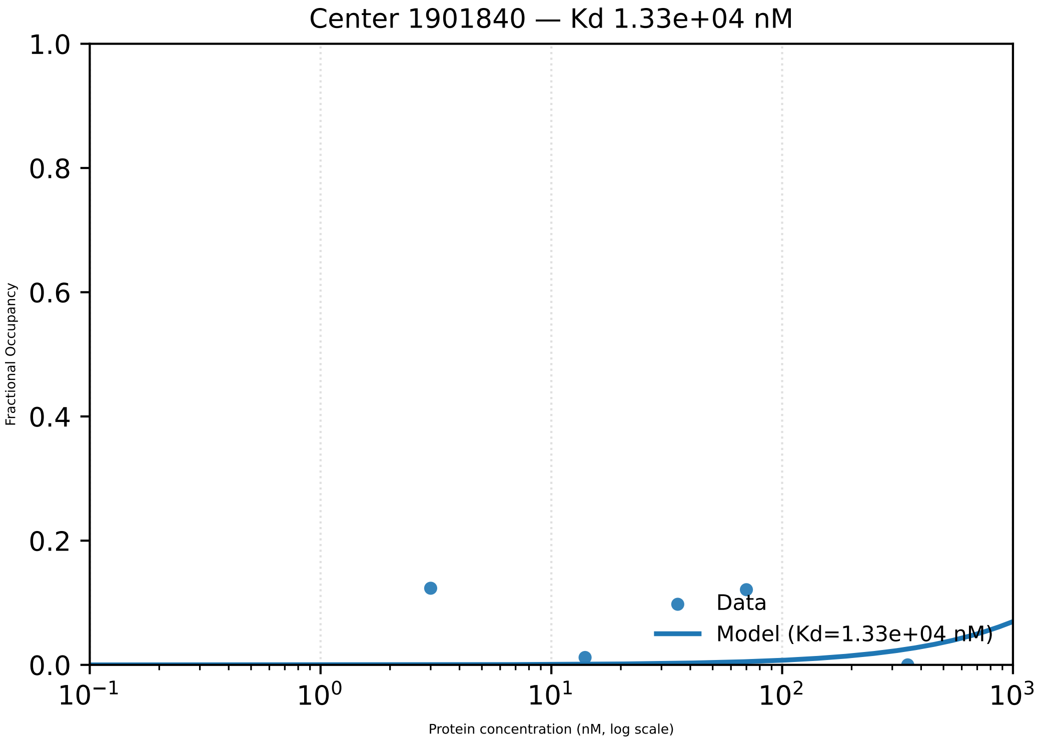

Center 1901840 — Kd 1.33e+04 nM
1.0
0.8
0.6
0.4
0.2
0.0
Fractional Occupancy
$10^{-1}$
$10^{0}$
$10^{1}$
$10^{2}$
$10^{3}$
Protein concentration (nM, log scale)
Data
Model (Kd=1.33e+04 nM)

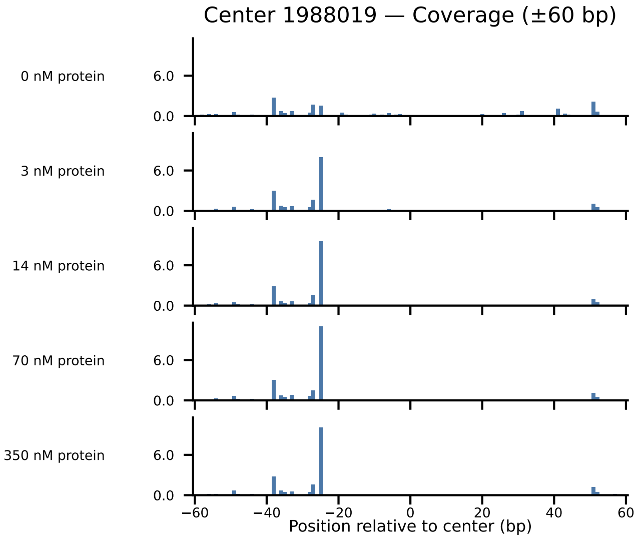
Center 1988019 — Coverage (±60 bp)
0 nM protein
3 nM protein
14 nM protein
70 nM protein
350 nM protein
6.0
0.0
−60
−40
−20
0
20
40
60
Position relative to center (bp)

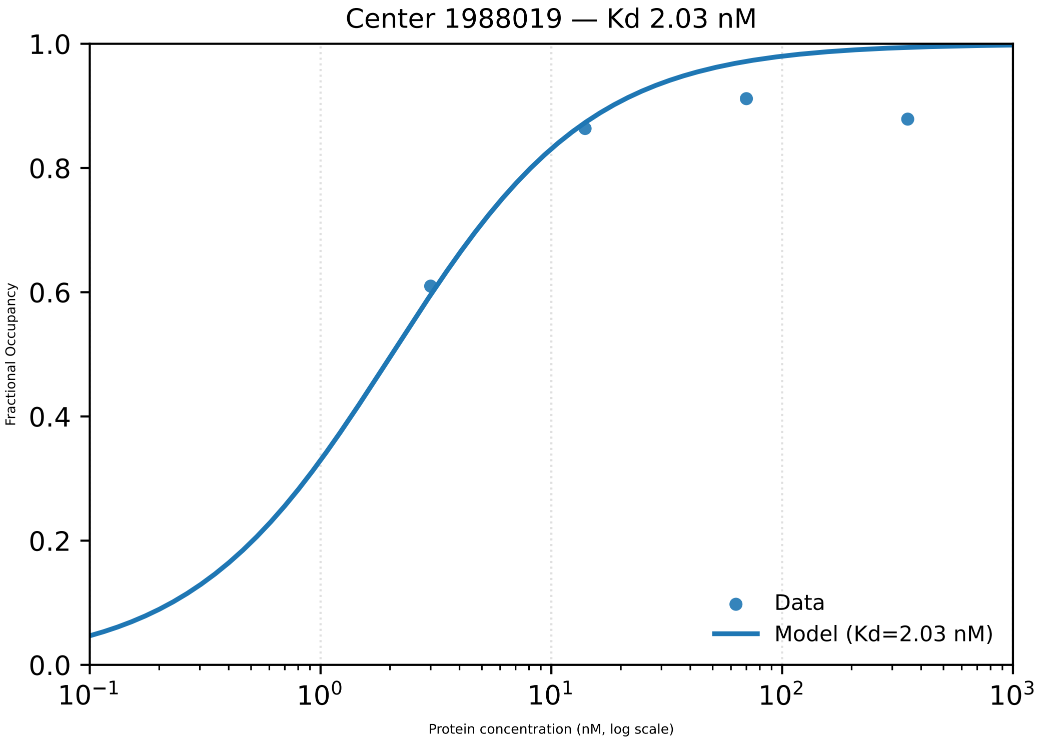

Center 1988019 — Kd 2.03 nM
1.0
0.8
0.6
0.4
0.2
0.0
Fractional Occupancy
$10^{-1}$
$10^{0}$
$10^{1}$
$10^{2}$
$10^{3}$
Protein concentration (nM, log scale)
Data
Model (Kd=2.03 nM)

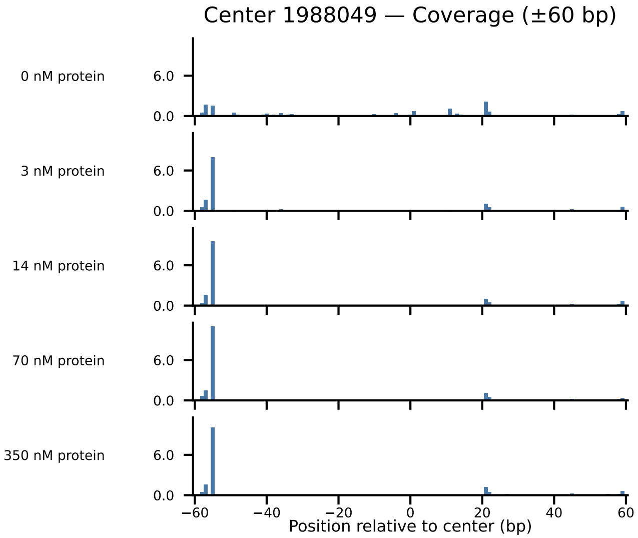

Center 1988049 — Coverage (±60 bp)
0 nM protein
3 nM protein
14 nM protein
70 nM protein
350 nM protein
6.0
0.0
−60
−40
−20
0
20
40
60
Position relative to center (bp)

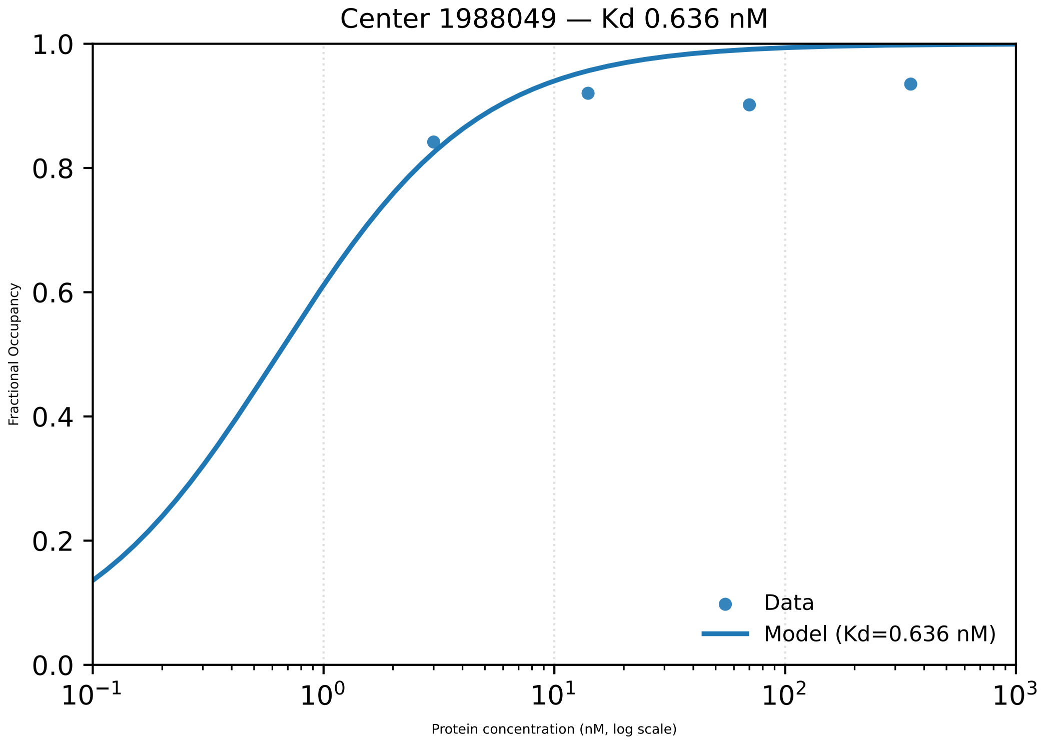

Center 1988049 — Kd 0.636 nM
1.0
0.8
0.6
0.4
0.2
0.0
Fractional Occupancy
$10^{-1}$
$10^{0}$
$10^{1}$
$10^{2}$
$10^{3}$
Protein concentration (nM, log scale)
Data
Model (Kd=0.636 nM)

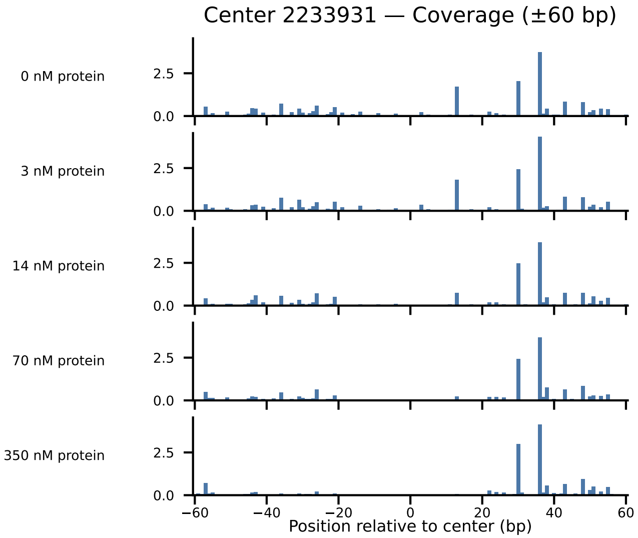

Center 2233931 — Coverage (±60 bp)
0 nM protein
3 nM protein
14 nM protein
70 nM protein
350 nM protein
2.5
0.0
−60
−40
−20
0
20
40
60
Position relative to center (bp)

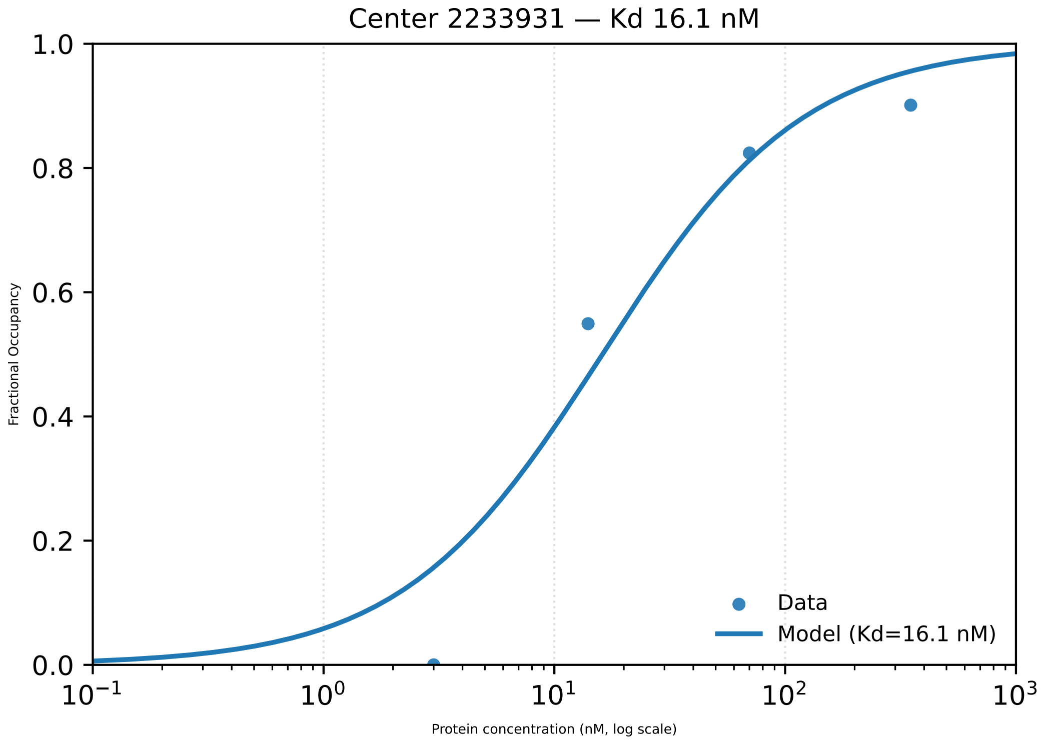

Center 2233931 — Kd 16.1 nM
Fractional Occupancy
1.0
0.8
0.6
0.4
0.2
0.0
$10^{-1}$
$10^{0}$
$10^{1}$
$10^{2}$
$10^{3}$
Protein concentration (nM, log scale)
Data
Model (Kd=16.1 nM)

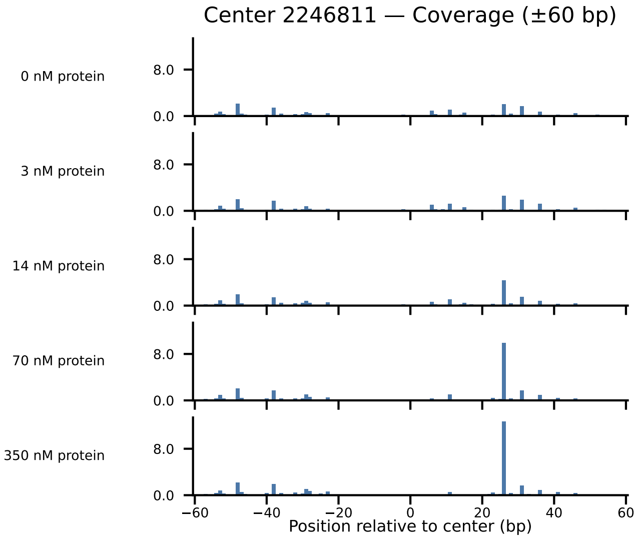
Center 2246811 — Coverage (±60 bp)
0 nM protein
3 nM protein
14 nM protein
70 nM protein
350 nM protein
8.0
0.0
−60
−40
−20
0
20
40
60
Position relative to center (bp)

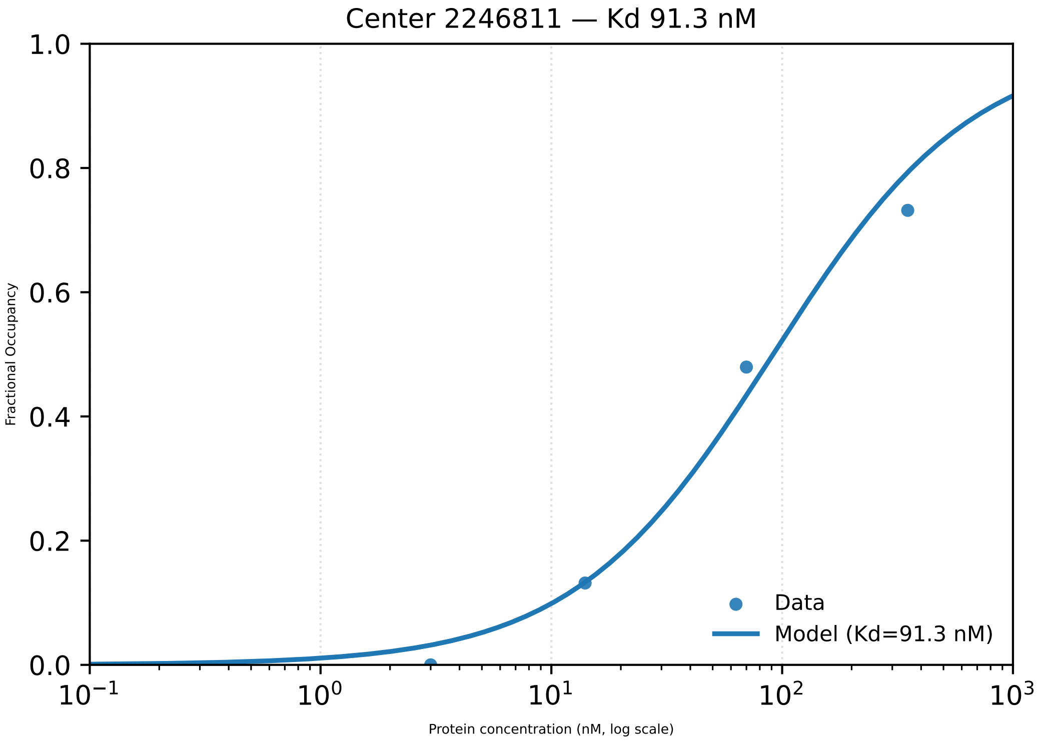

Center 2246811 — Kd 91.3 nM
Fractional Occupancy
1.0
0.8
0.6
0.4
0.2
0.0
$10^{-1}$
$10^{0}$
$10^{1}$
$10^{2}$
$10^{3}$
Protein concentration (nM, log scale)
Data
Model (Kd=91.3 nM)

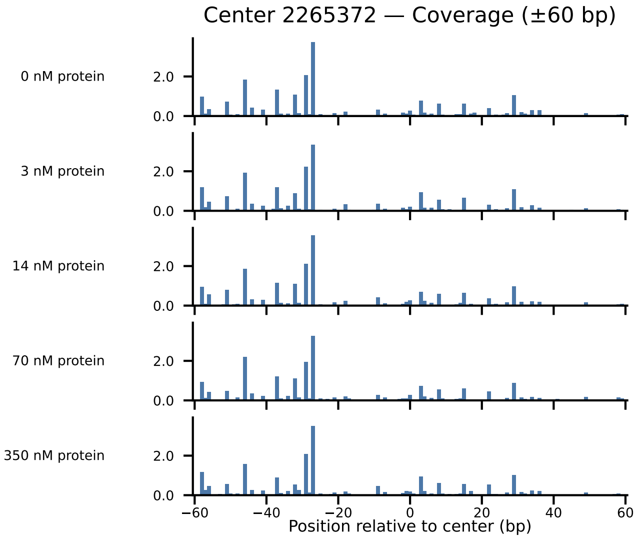
Center 2265372 — Coverage (±60 bp)
0 nM protein
3 nM protein
14 nM protein
70 nM protein
350 nM protein
2.0
0.0
−60
−40
−20
0
20
40
60
Position relative to center (bp)

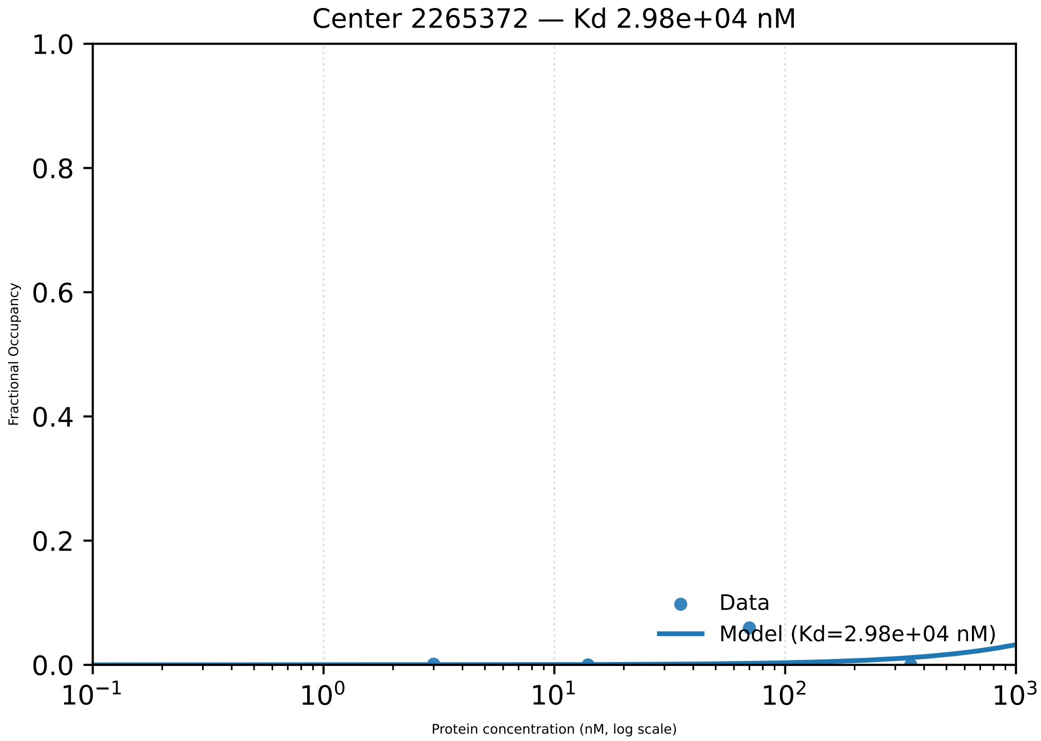
Center 2265372 — Kd 2.98e+04 nM
1.0
0.8
0.6
0.4
0.2
0.0
Fractional Occupancy
Data
Model (Kd=2.98e+04 nM)
$10^{-1}$
$10^{0}$
$10^{1}$
$10^{2}$
$10^{3}$
Protein concentration (nM, log scale)

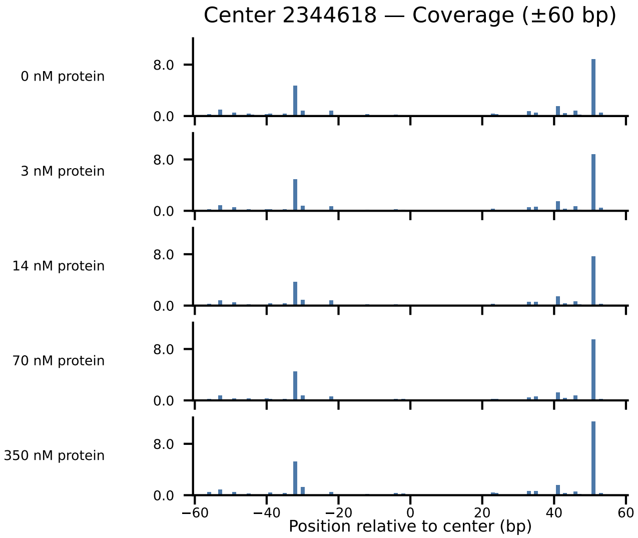

Center 2344618 — Coverage (±60 bp)
0 nM protein
3 nM protein
14 nM protein
70 nM protein
350 nM protein
8.0
0.0
−60
−40
−20
0
20
40
60
Position relative to center (bp)

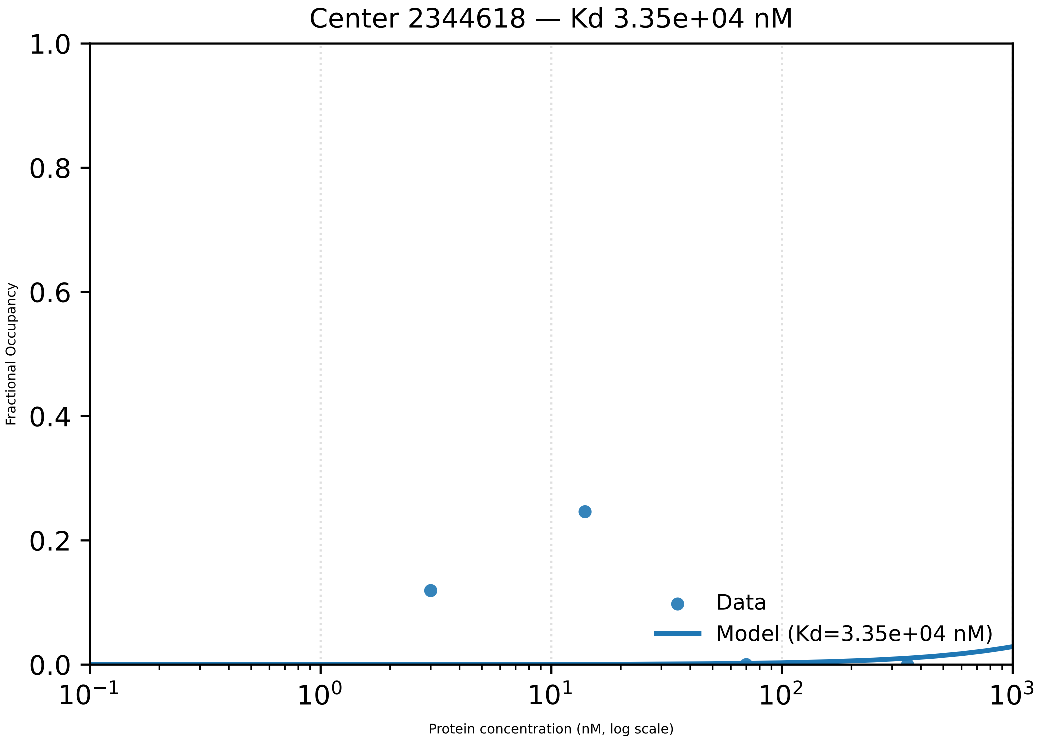

Center 2344618 — Kd 3.35e+04 nM
1.0
0.8
0.6
0.4
0.2
0.0
Fractional Occupancy
Data
Model (Kd=3.35e+04 nM)
$10^{-1}$
$10^{0}$
$10^{1}$
$10^{2}$
$10^{3}$
Protein concentration (nM, log scale)

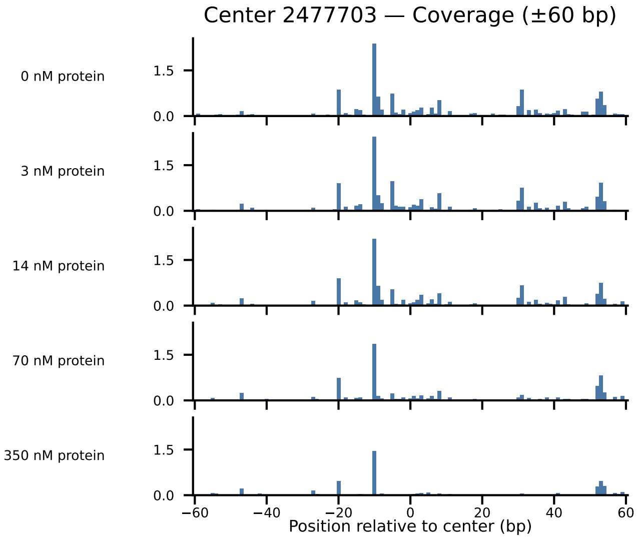

Center 2477703 — Coverage (±60 bp)
0 nM protein
3 nM protein
14 nM protein
70 nM protein
350 nM protein
1.5
0.0
−60
−40
−20
0
20
40
60
Position relative to center (bp)

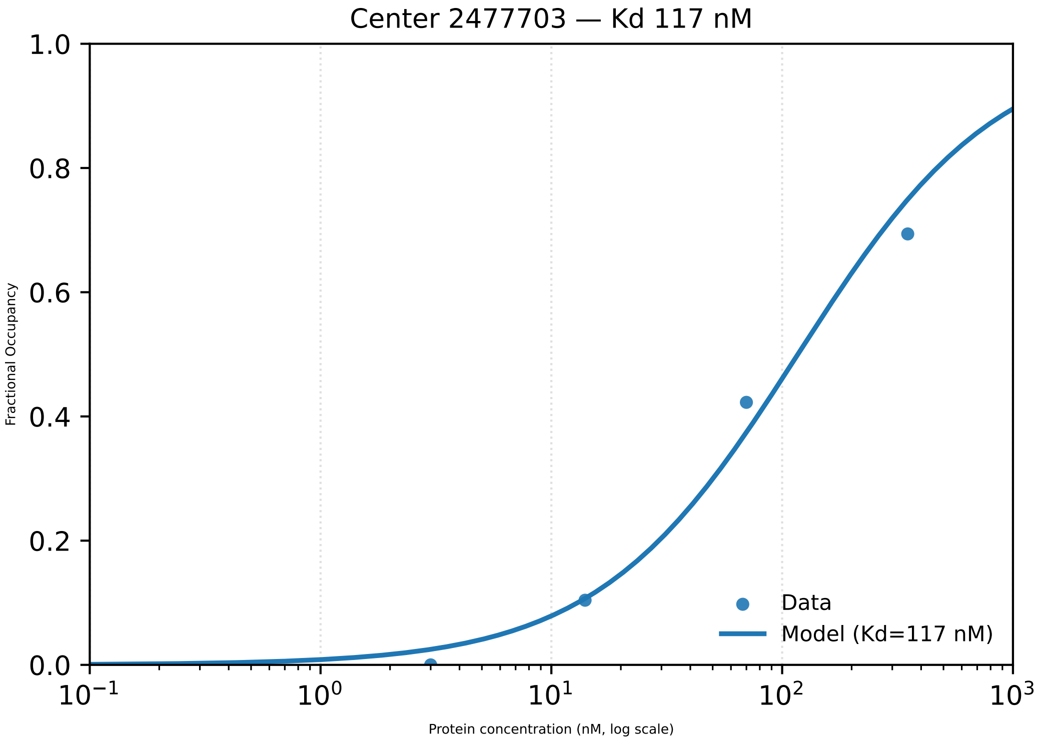

Center 2477703 — Kd 117 nM
1.0
0.8
0.6
0.4
0.2
0.0
Fractional Occupancy
$10^{-1}$
$10^{0}$
$10^{1}$
$10^{2}$
$10^{3}$
Protein concentration (nM, log scale)
Data
Model (Kd=117 nM)

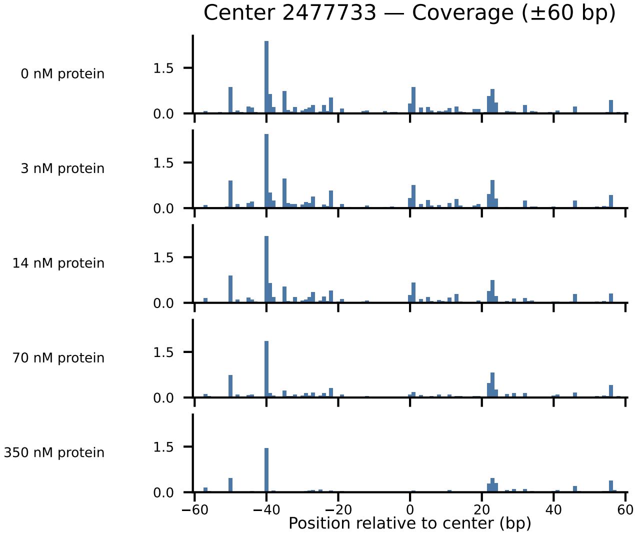

Center 2477733 — Coverage (±60 bp)
0 nM protein
3 nM protein
14 nM protein
70 nM protein
350 nM protein
1.5
0.0
−60
−40
−20
0
20
40
60
Position relative to center (bp)

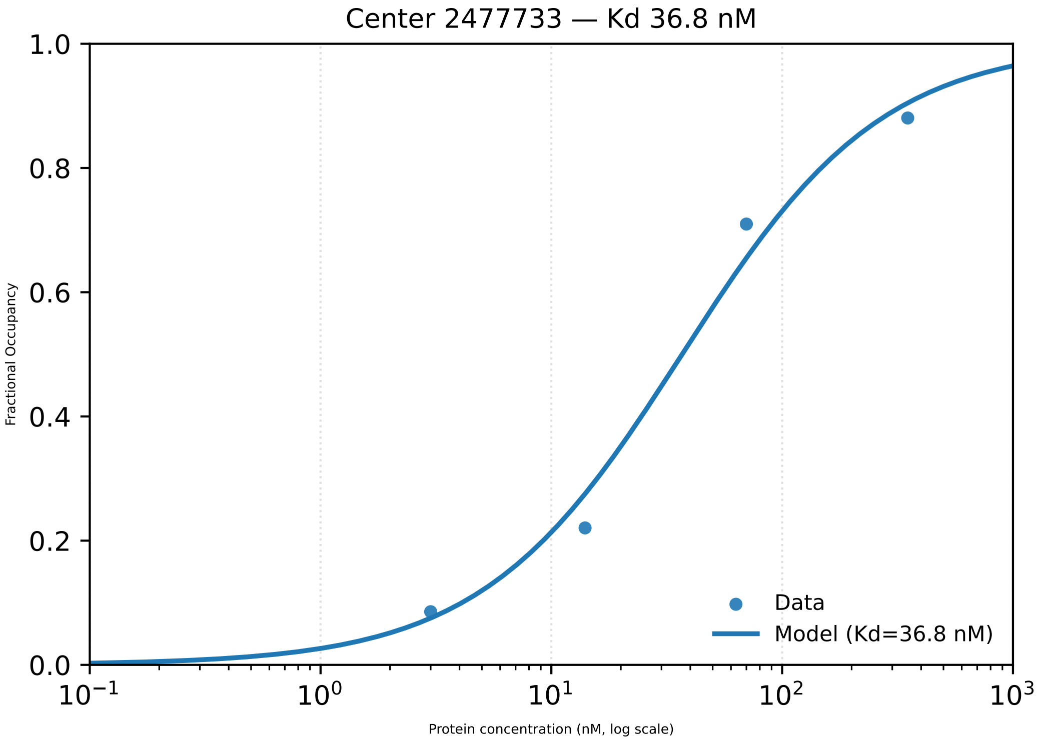
Center 2477733 — Kd 36.8 nM
Fractional Occupancy
1.0
0.8
0.6
0.4
0.2
0.0
$10^{-1}$
$10^{0}$
$10^{1}$
$10^{2}$
$10^{3}$
Protein concentration (nM, log scale)
Data
Model (Kd=36.8 nM)

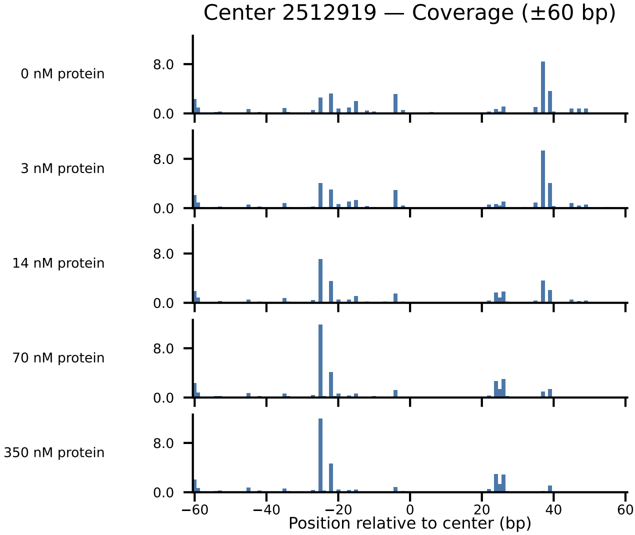
Center 2512919 — Coverage (±60 bp)
0 nM protein
3 nM protein
14 nM protein
70 nM protein
350 nM protein
8.0
0.0
−60
−40
−20
0
20
40
60
Position relative to center (bp)

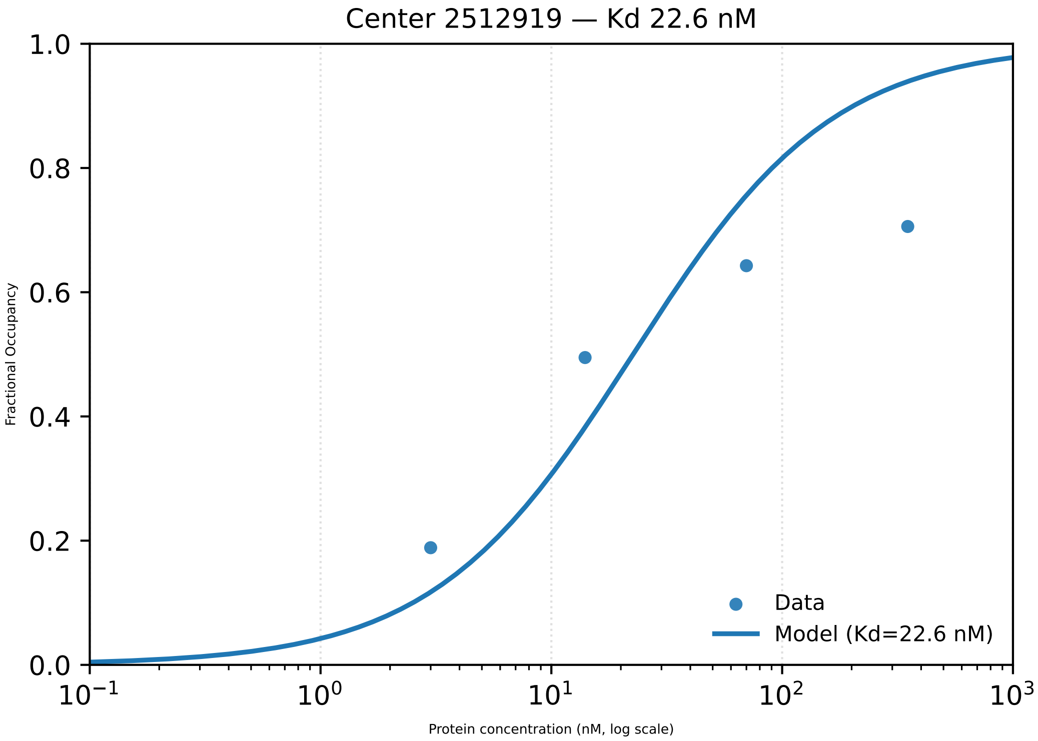

Center 2512919 — Kd 22.6 nM
Fractional Occupancy
1.0
0.8
0.6
0.4
0.2
0.0
$10^{-1}$
$10^{0}$
$10^{1}$
$10^{2}$
$10^{3}$
Protein concentration (nM, log scale)
Data
Model (Kd=22.6 nM)

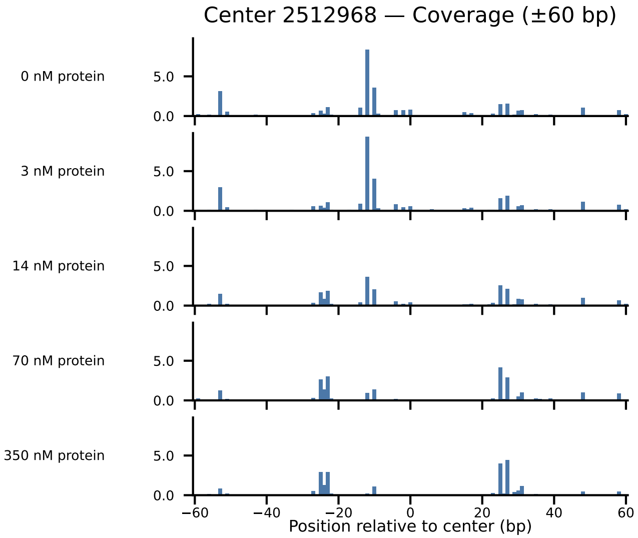
Center 2512968 — Coverage (±60 bp)
0 nM protein
3 nM protein
14 nM protein
70 nM protein
350 nM protein
5.0
0.0
−60
−40
−20
0
20
40
60
Position relative to center (bp)

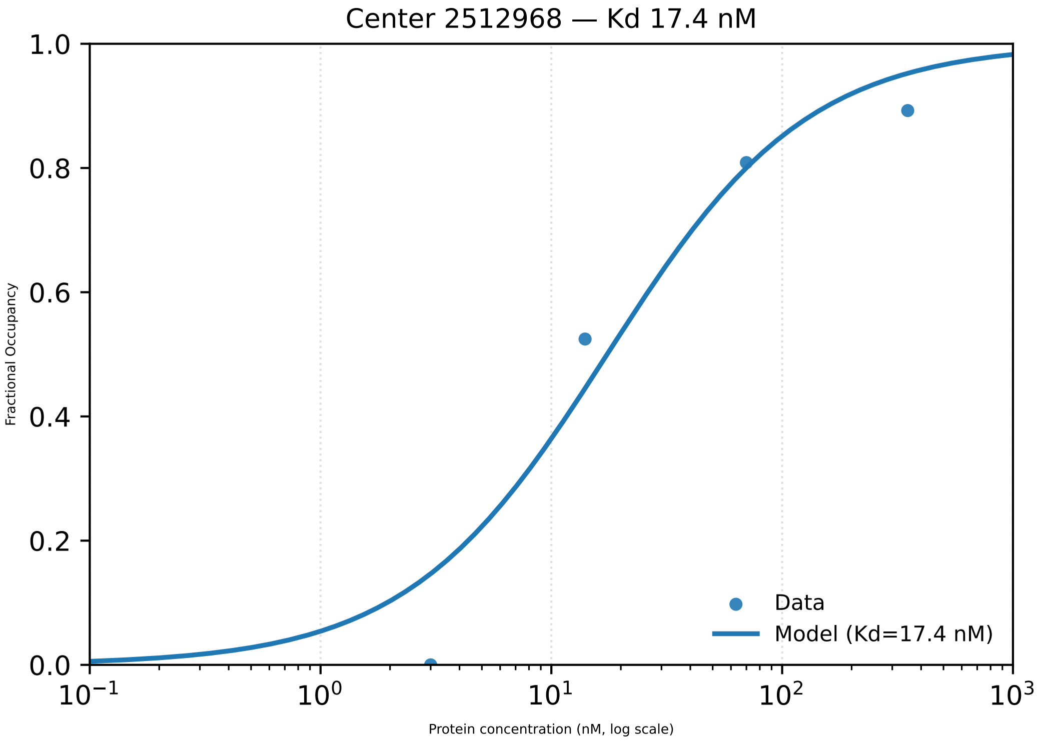

Center 2512968 — Kd 17.4 nM
Fractional Occupancy
1.0
0.8
0.6
0.4
0.2
0.0
$10^{-1}$
$10^{0}$
$10^{1}$
$10^{2}$
$10^{3}$
Protein concentration (nM, log scale)
Data
Model (Kd=17.4 nM)

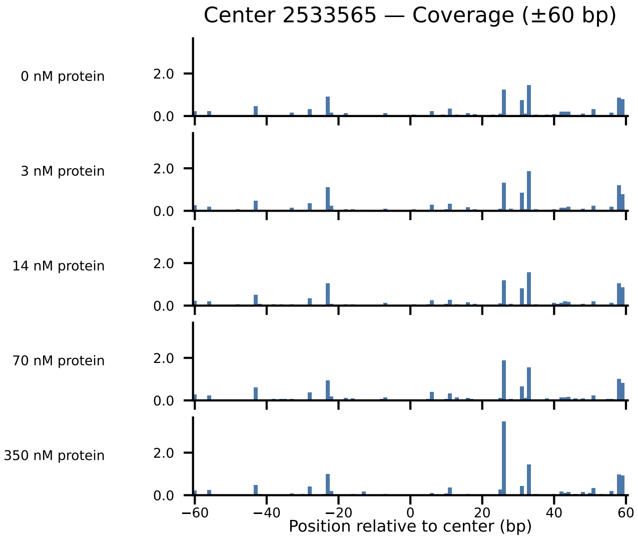

Center 2533565 — Coverage (±60 bp)
0 nM protein
3 nM protein
14 nM protein
70 nM protein
350 nM protein
2.0
0.0
−60
−40
−20
0
20
40
60
Position relative to center (bp)

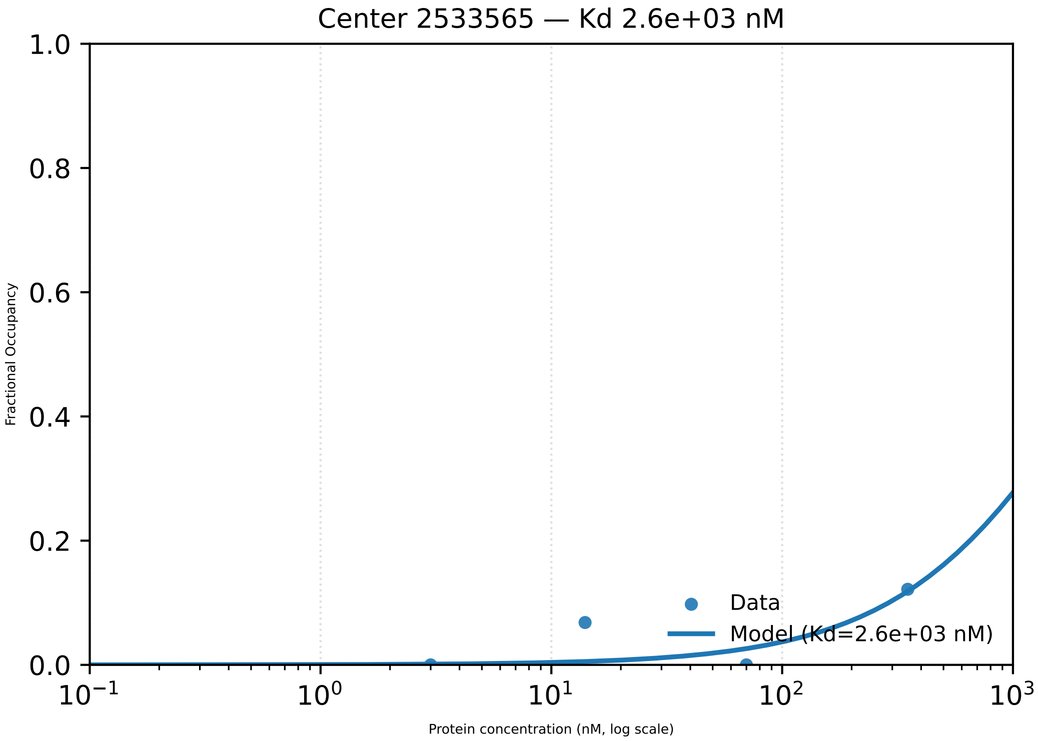

Center 2533565 — Kd 2.6e+03 nM
1.0
0.8
0.6
0.4
0.2
0.0
Fractional Occupancy
$10^{-1}$
$10^{0}$
$10^{1}$
$10^{2}$
$10^{3}$
Protein concentration (nM, log scale)
Data
Model (Kd=2.6e+03 nM)

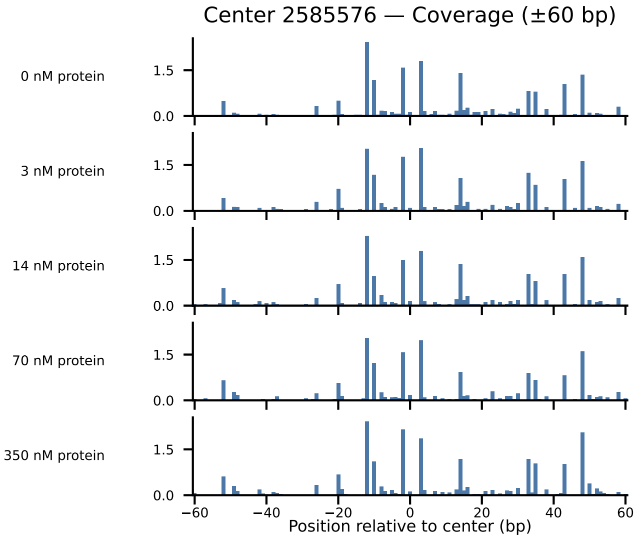

Center 2585576 — Coverage (±60 bp)
0 nM protein
3 nM protein
14 nM protein
70 nM protein
350 nM protein
1.5
0.0
−60
−40
−20
0
20
40
60
Position relative to center (bp)

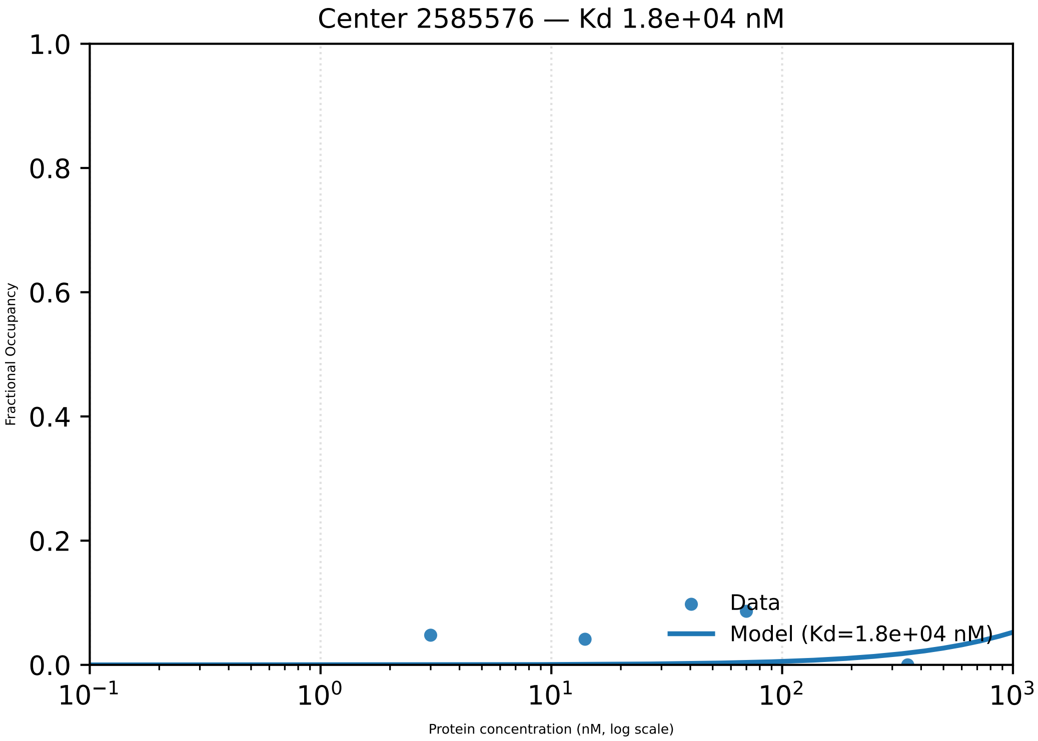
Center 2585576 — Kd 1.8e+04 nM
Fractional Occupancy
Protein concentration (nM, log scale)
1.0
0.8
0.6
0.4
0.2
0.0
$10^{-1}$
$10^{0}$
$10^{1}$
$10^{2}$
$10^{3}$
Data
Model (Kd=1.8e+04 nM)

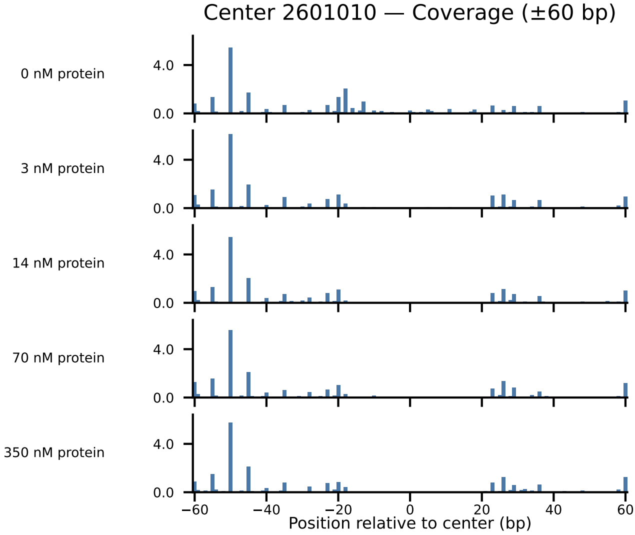

Center 2601010 — Coverage (±60 bp)
0 nM protein
3 nM protein
14 nM protein
70 nM protein
350 nM protein
4.0
0.0
−60
−40
−20
0
20
40
60
Position relative to center (bp)

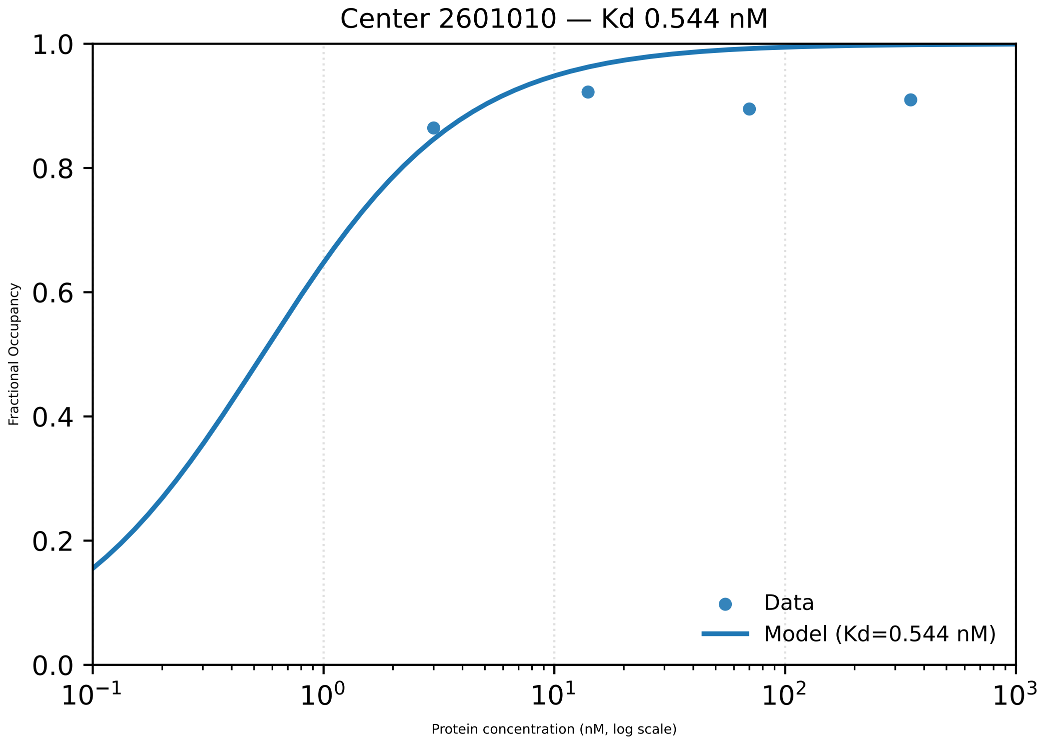
Center 2601010 — Kd 0.544 nM
1.0
0.8
0.6
0.4
0.2
0.0
Fractional Occupancy
10⁻¹
10⁰
10¹
10²
10³
Protein concentration (nM, log scale)
Data
Model (Kd=0.544 nM)

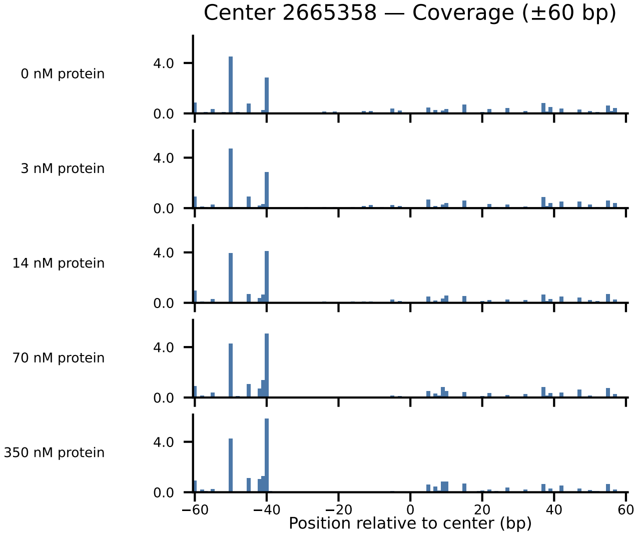
Center 2665358 — Coverage (±60 bp)
0 nM protein
3 nM protein
14 nM protein
70 nM protein
350 nM protein
4.0
0.0
−60
−40
−20
0
20
40
60
Position relative to center (bp)

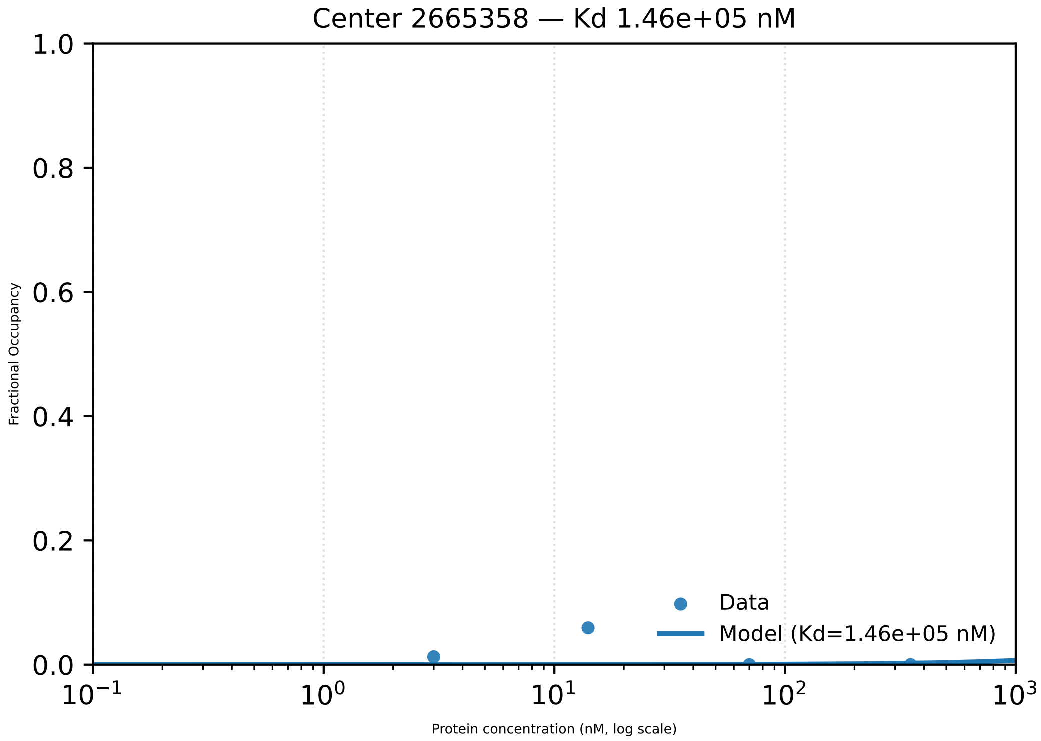
Center 2665358 — Kd 1.46e+05 nM
1.0
0.8
0.6
0.4
0.2
0.0
Fractional Occupancy
$10^{-1}$
$10^{0}$
$10^{1}$
$10^{2}$
$10^{3}$
Protein concentration (nM, log scale)
Data
Model (Kd=1.46e+05 nM)

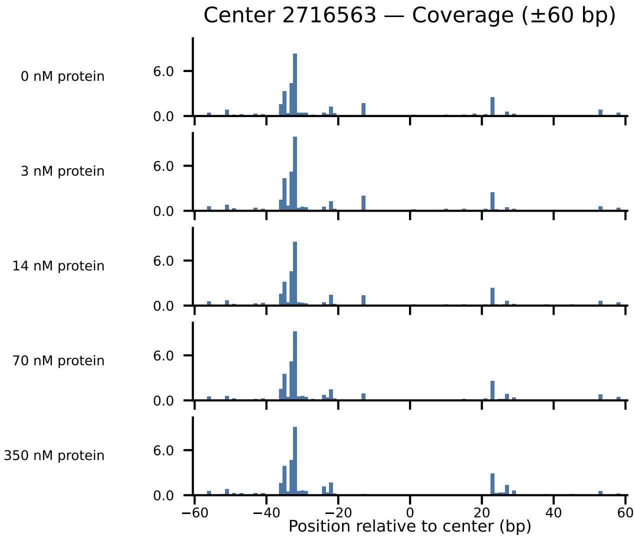

Center 2716563 — Coverage (±60 bp)
0 nM protein
3 nM protein
14 nM protein
70 nM protein
350 nM protein
6.0
0.0
−60
−40
−20
0
20
40
60
Position relative to center (bp)

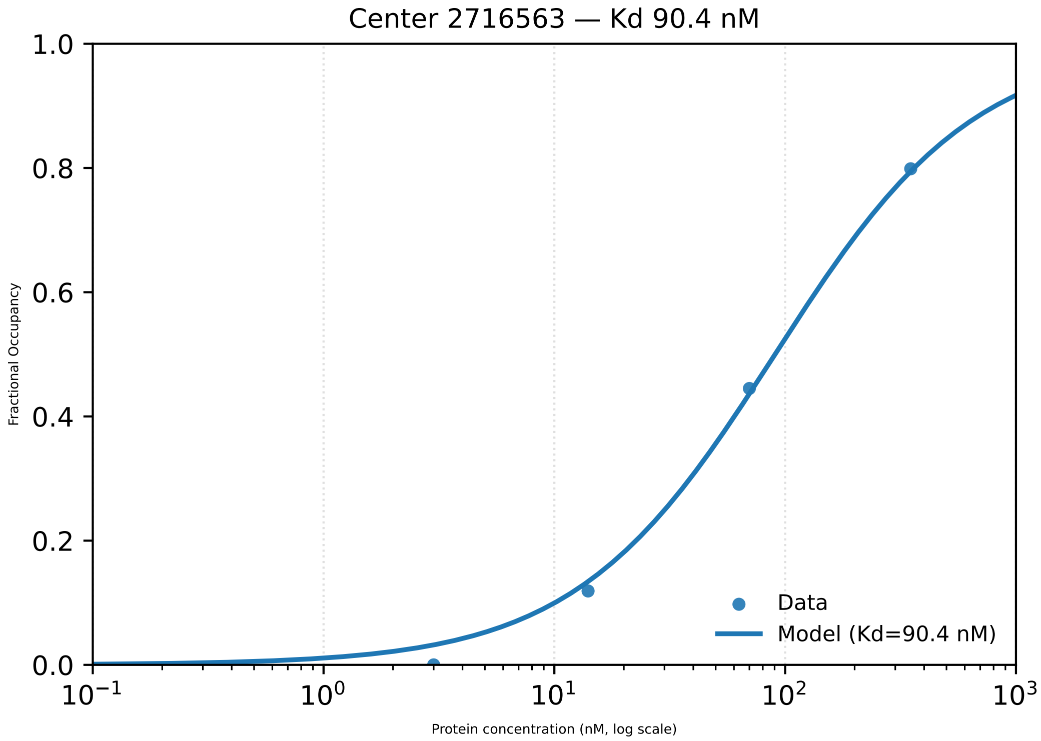
Center 2716563 — Kd 90.4 nM
1.0
0.8
0.6
0.4
0.2
0.0
Fractional Occupancy
$10^{-1}$
$10^{0}$
$10^{1}$
$10^{2}$
$10^{3}$
Protein concentration (nM, log scale)
Data
Model (Kd=90.4 nM)

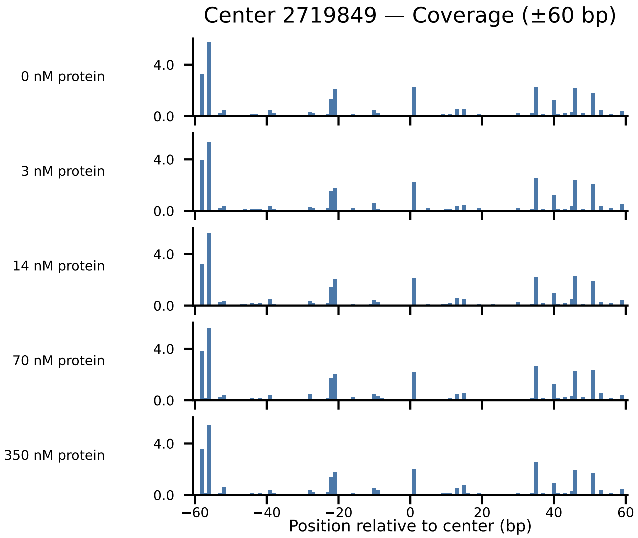

Center 2719849 — Coverage (±60 bp)
0 nM protein
3 nM protein
14 nM protein
70 nM protein
350 nM protein
4.0
0.0
−60
−40
−20
0
20
40
60
Position relative to center (bp)

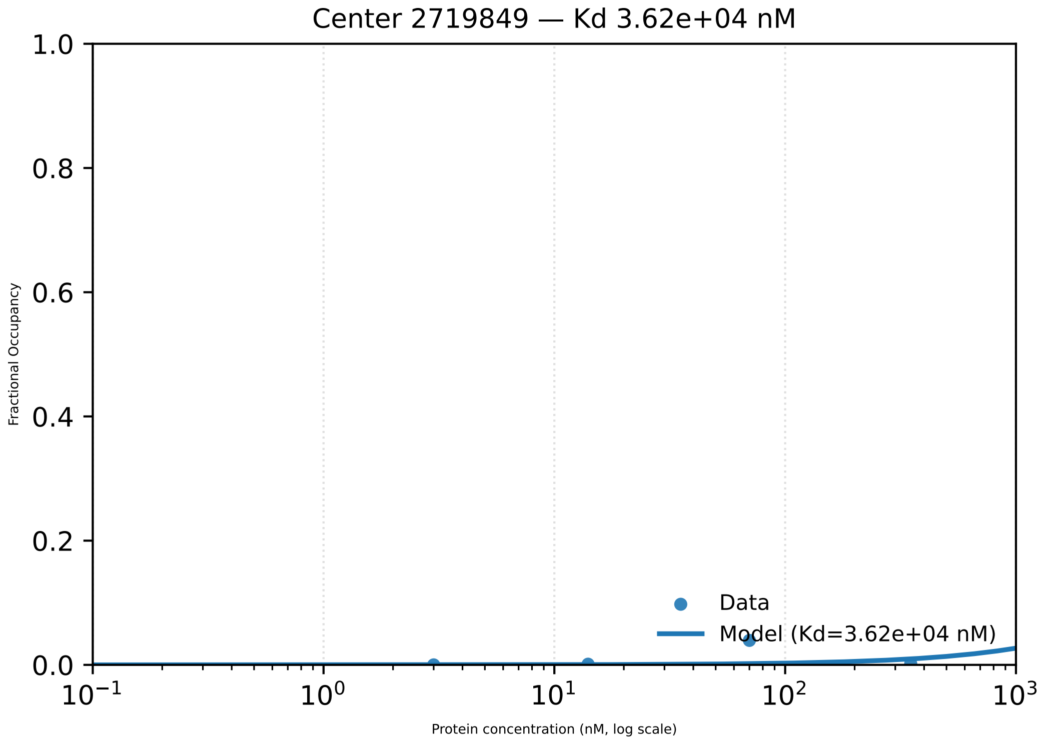

Center 2719849 — Kd 3.62e+04 nM
Fractional Occupancy
1.0
0.8
0.6
0.4
0.2
0.0
$10^{-1}$
$10^{0}$
$10^{1}$
$10^{2}$
$10^{3}$
Protein concentration (nM, log scale)
Data
Model (Kd=3.62e+04 nM)

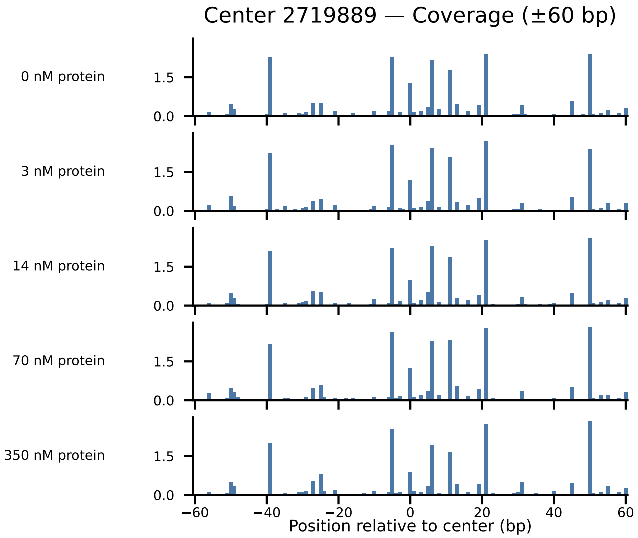
Center 2719889 — Coverage (±60 bp)
0 nM protein
3 nM protein
14 nM protein
70 nM protein
350 nM protein
0.0
1.5
−60
−40
−20
0
20
40
60
Position relative to center (bp)

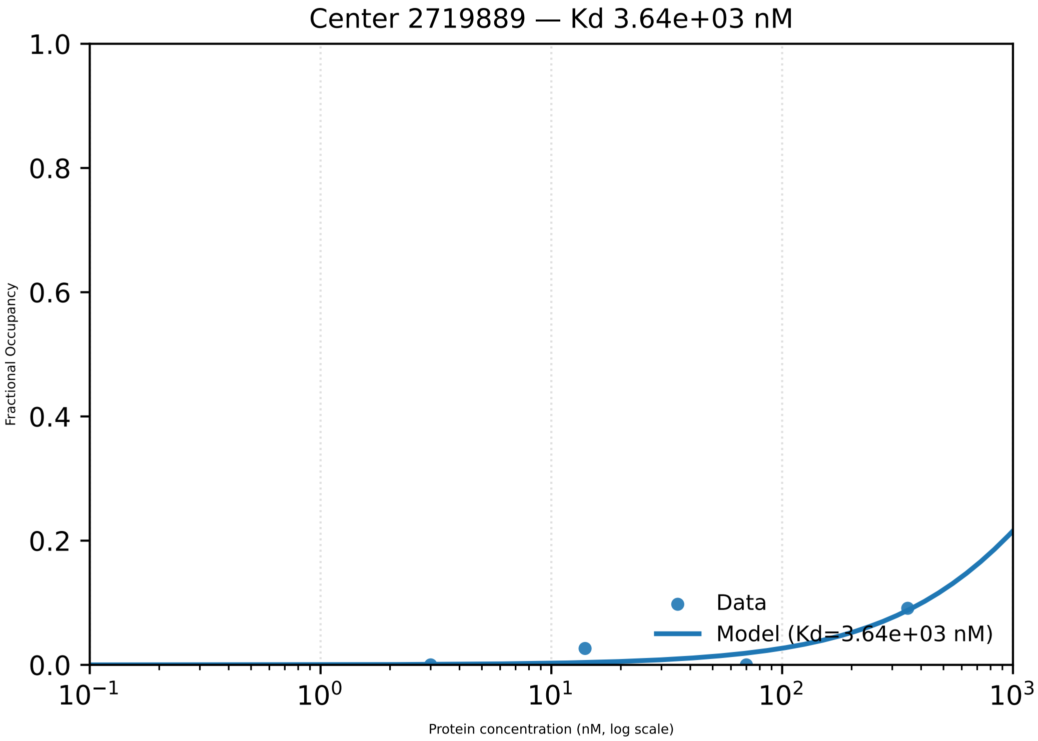

Center 2719889 — Kd 3.64e+03 nM
1.0
0.8
0.6
0.4
0.2
0.0
Fractional Occupancy
Data
Model (Kd=3.64e+03 nM)
$10^{-1}$
$10^{0}$
$10^{1}$
$10^{2}$
$10^{3}$
Protein concentration (nM, log scale)

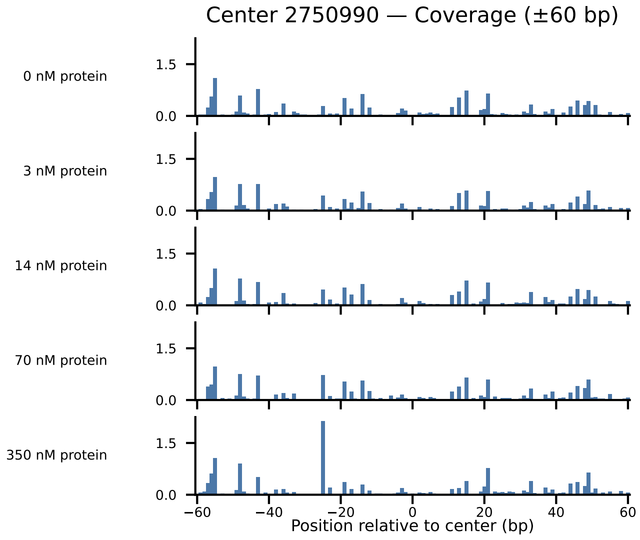
Center 2750990 — Coverage (±60 bp)
0 nM protein
3 nM protein
14 nM protein
70 nM protein
350 nM protein
1.5
0.0
−60
−40
−20
0
20
40
60
Position relative to center (bp)

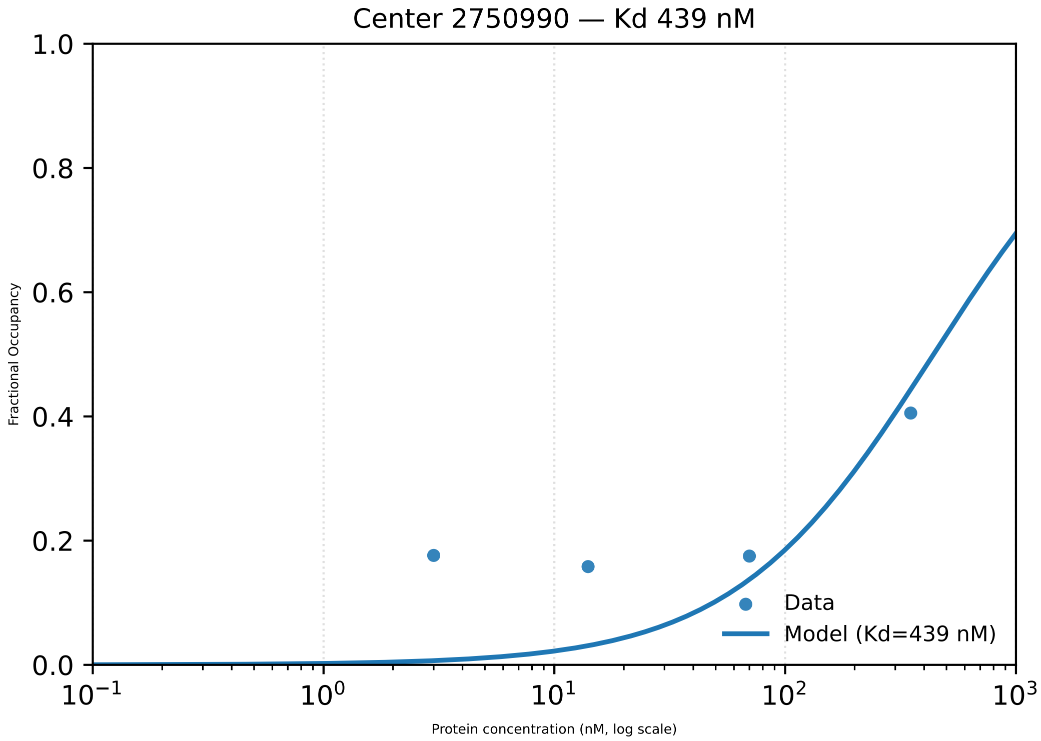

Center 2750990 — Kd 439 nM
1.0
0.8
0.6
0.4
0.2
0.0
Fractional Occupancy
$10^{-1}$
$10^{0}$
$10^{1}$
$10^{2}$
$10^{3}$
Protein concentration (nM, log scale)
Data
Model (Kd=439 nM)

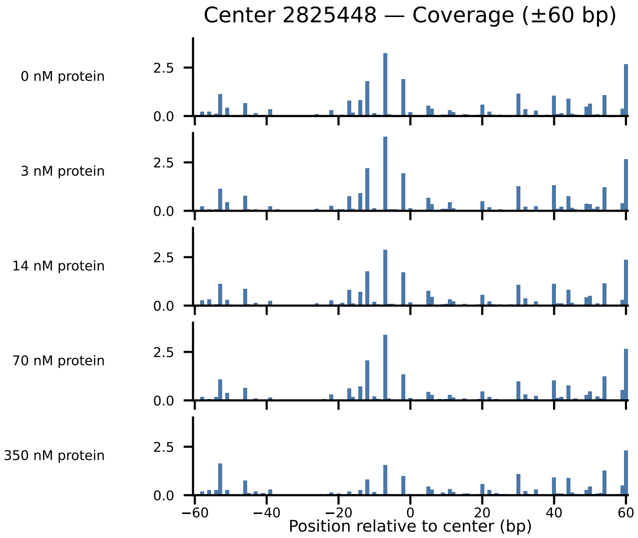
Center 2825448 — Coverage (±60 bp)
0 nM protein
3 nM protein
14 nM protein
70 nM protein
350 nM protein
2.5
0.0
−60
−40
−20
0
20
40
60
Position relative to center (bp)

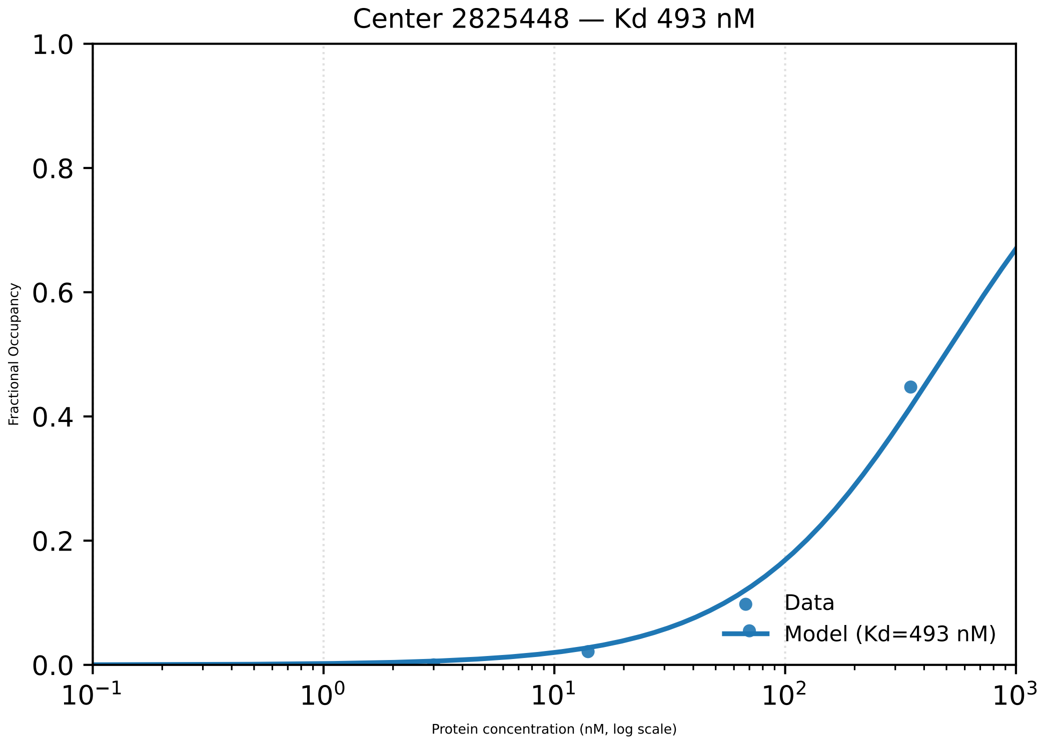

Center 2825448 — Kd 493 nM
Fractional Occupancy
Protein concentration (nM, log scale)
1.0
0.8
0.6
0.4
0.2
0.0
$10^{-1}$
$10^{0}$
$10^{1}$
$10^{2}$
$10^{3}$
Data
Model (Kd=493 nM)

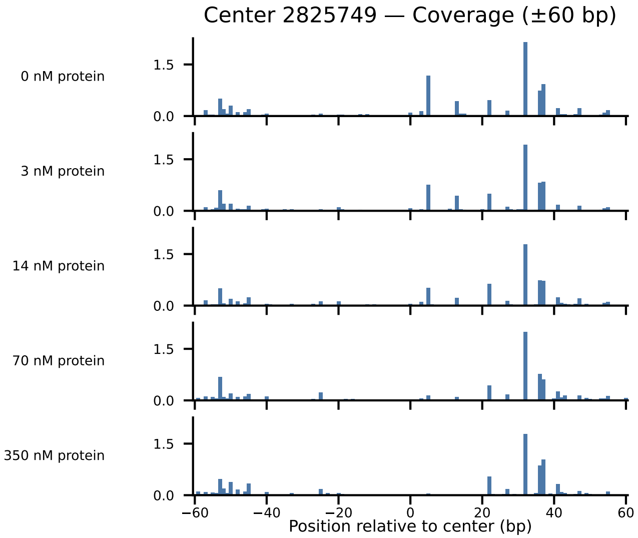

Center 2825749 — Coverage (±60 bp)
0 nM protein
3 nM protein
14 nM protein
70 nM protein
350 nM protein
1.5
0.0
−60
−40
−20
0
20
40
60
Position relative to center (bp)

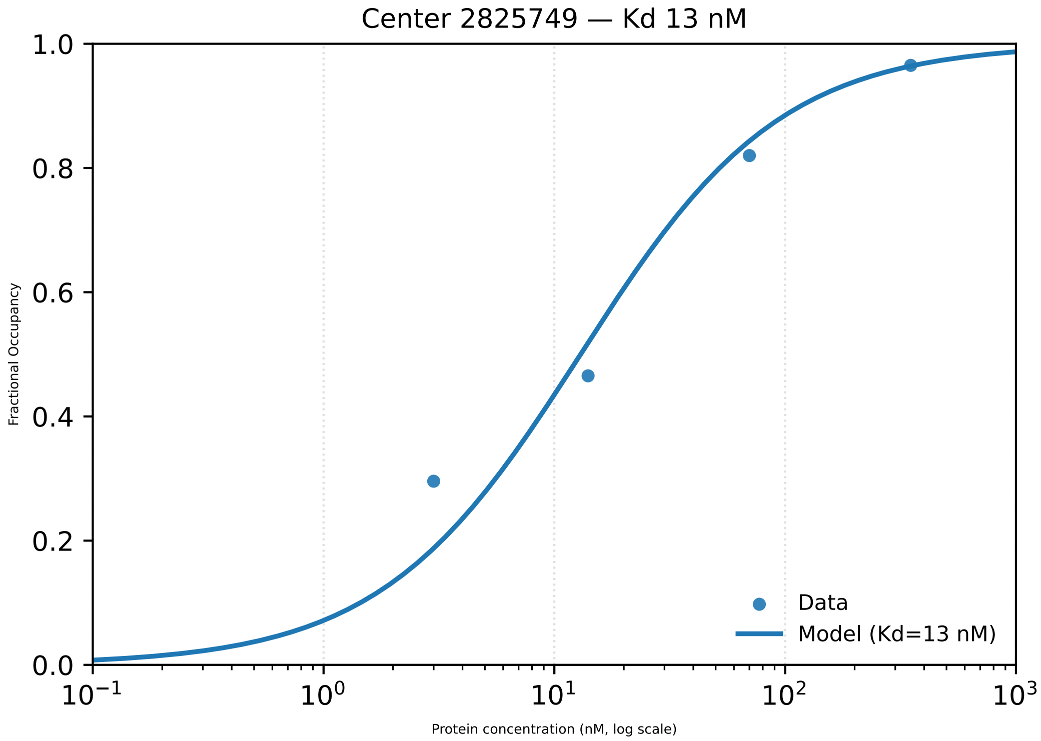

Center 2825749 — Kd 13 nM
Fractional Occupancy
1.0
0.8
0.6
0.4
0.2
0.0
$10^{-1}$
$10^{0}$
$10^{1}$
$10^{2}$
$10^{3}$
Protein concentration (nM, log scale)
Data
Model (Kd=13 nM)

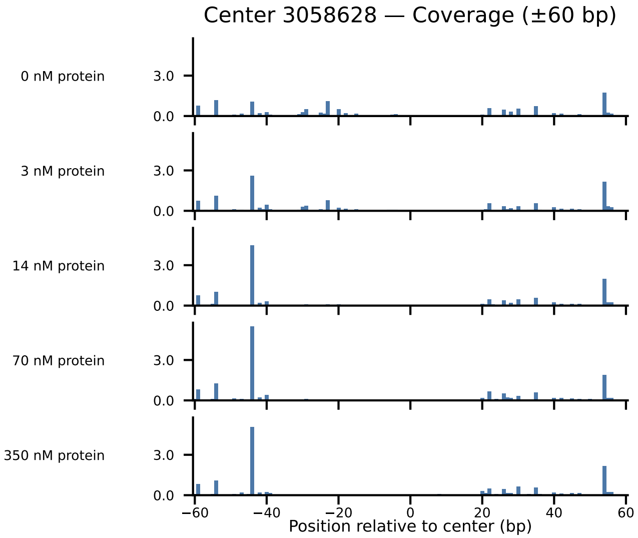

Center 3058628 — Coverage (±60 bp)
0 nM protein
3 nM protein
14 nM protein
70 nM protein
350 nM protein
3.0
0.0
−60
−40
−20
0
20
40
60
Position relative to center (bp)

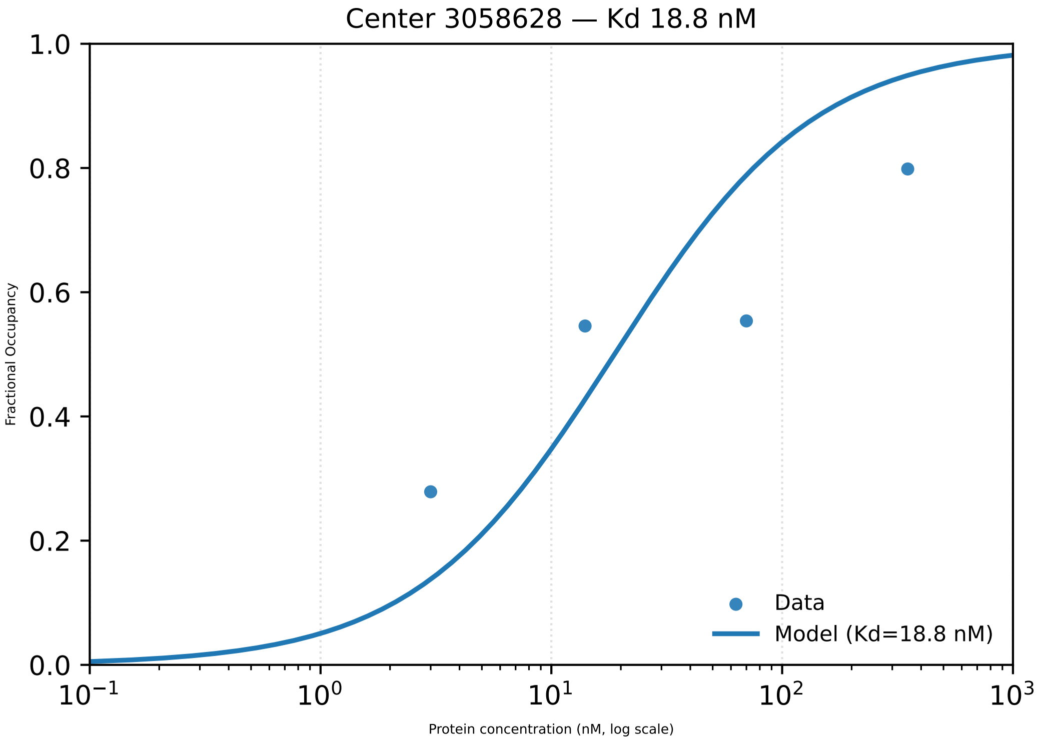

Center 3058628 — Kd 18.8 nM
1.0
0.8
0.6
0.4
0.2
0.0
Fractional Occupancy
$10^{-1}$
$10^{0}$
$10^{1}$
$10^{2}$
$10^{3}$
Protein concentration (nM, log scale)
Data
Model (Kd=18.8 nM)

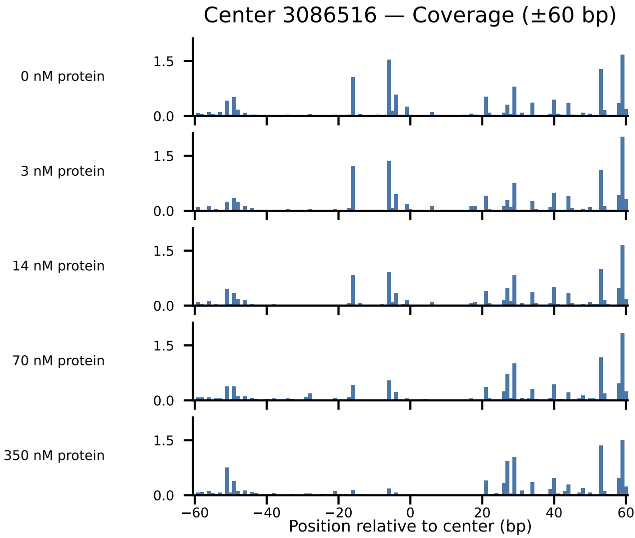

Center 3086516 — Coverage (±60 bp)
0 nM protein
3 nM protein
14 nM protein
70 nM protein
350 nM protein
1.5
0.0
−60
−40
−20
0
20
40
60
Position relative to center (bp)

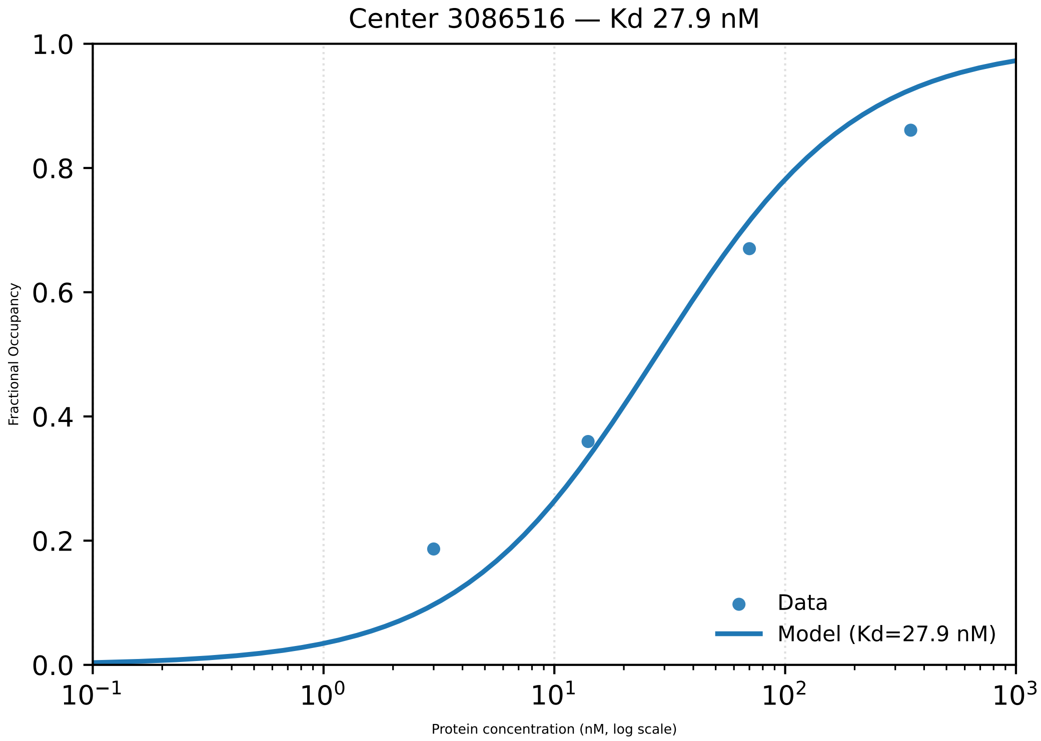

Center 3086516 — Kd 27.9 nM
1.0
0.8
0.6
0.4
0.2
0.0
Fractional Occupancy
$10^{-1}$
$10^{0}$
$10^{1}$
$10^{2}$
$10^{3}$
Protein concentration (nM, log scale)
Data
Model (Kd=27.9 nM)

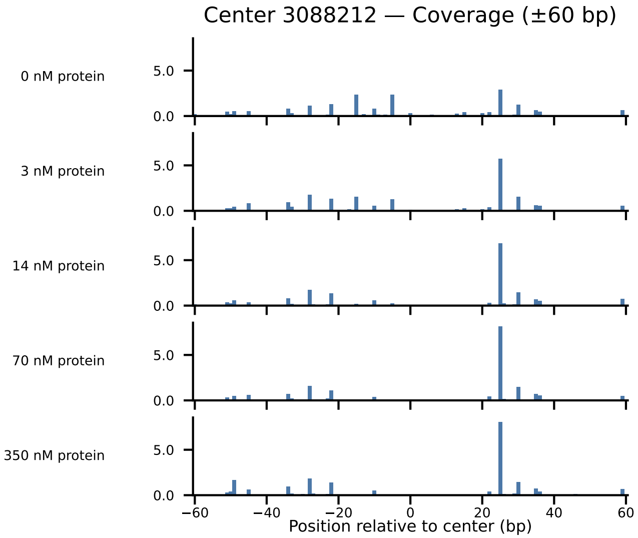
Center 3088212 — Coverage (±60 bp)
0 nM protein
3 nM protein
14 nM protein
70 nM protein
350 nM protein
5.0
0.0
−60
−40
−20
0
20
40
60
Position relative to center (bp)

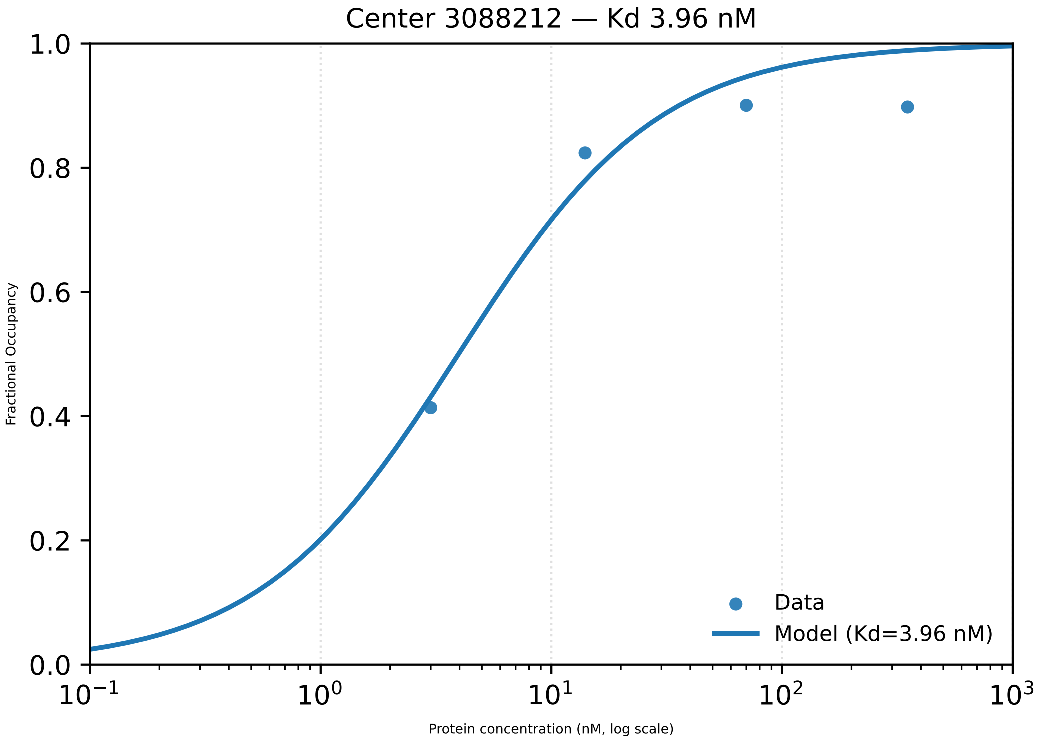

Center 3088212 — Kd 3.96 nM
Fractional Occupancy
1.0
0.8
0.6
0.4
0.2
0.0
$10^{-1}$
$10^{0}$
$10^{1}$
$10^{2}$
$10^{3}$
Protein concentration (nM, log scale)
Data
Model (Kd=3.96 nM)

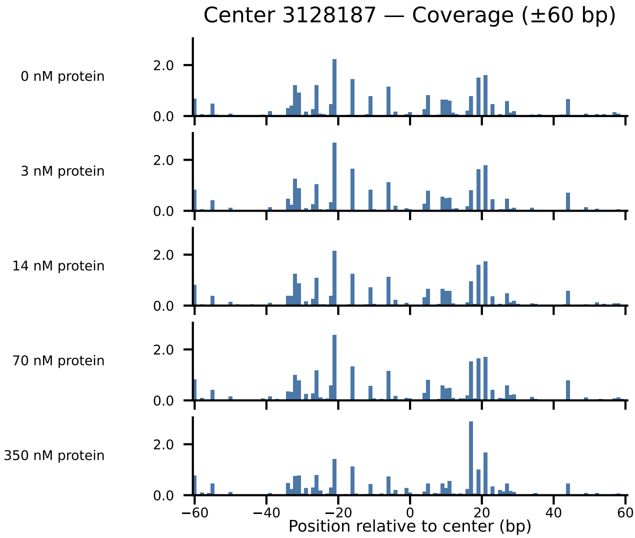

Center 3128187 — Coverage (±60 bp)
0 nM protein
3 nM protein
14 nM protein
70 nM protein
350 nM protein
2.0
0.0
−60
−40
−20
0
20
40
60
Position relative to center (bp)

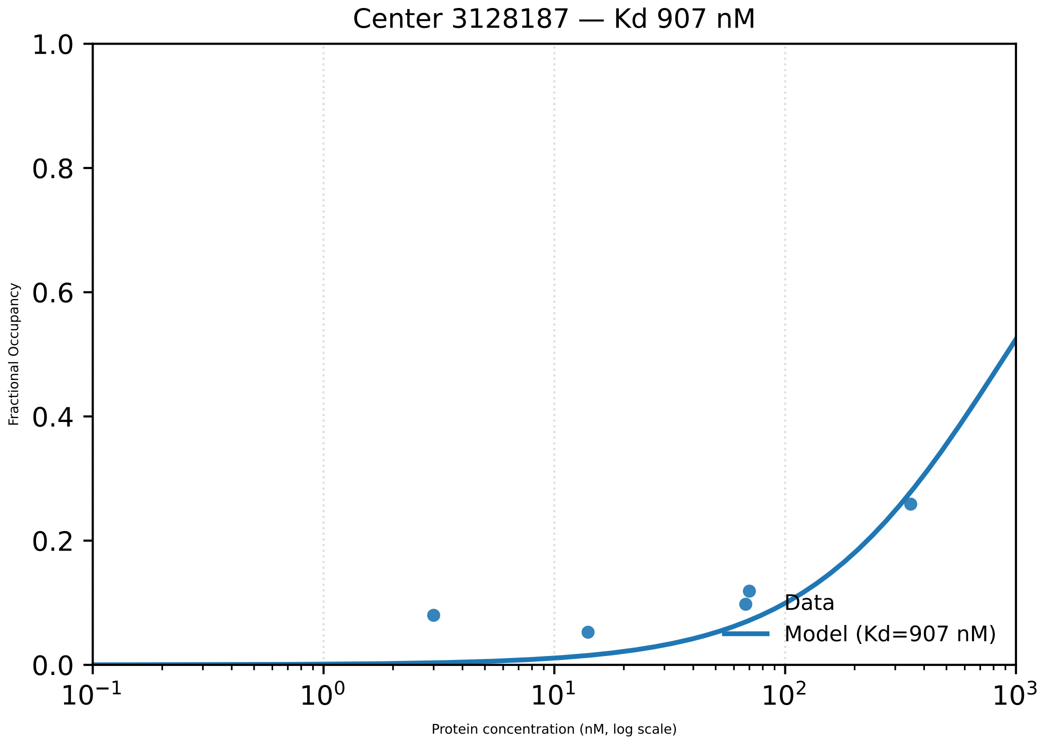

Center 3128187 — Kd 907 nM
1.0
0.8
0.6
0.4
0.2
0.0
Fractional Occupancy
$10^{-1}$
$10^{0}$
$10^{1}$
$10^{2}$
$10^{3}$
Protein concentration (nM, log scale)
Data
Model (Kd=907 nM)

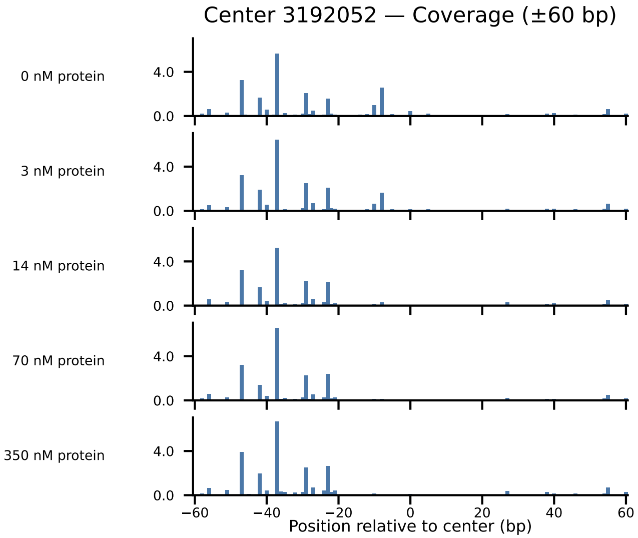
Center 3192052 — Coverage (±60 bp)
0 nM protein
3 nM protein
14 nM protein
70 nM protein
350 nM protein
4.0
0.0
−60
−40
−20
0
20
40
60
Position relative to center (bp)

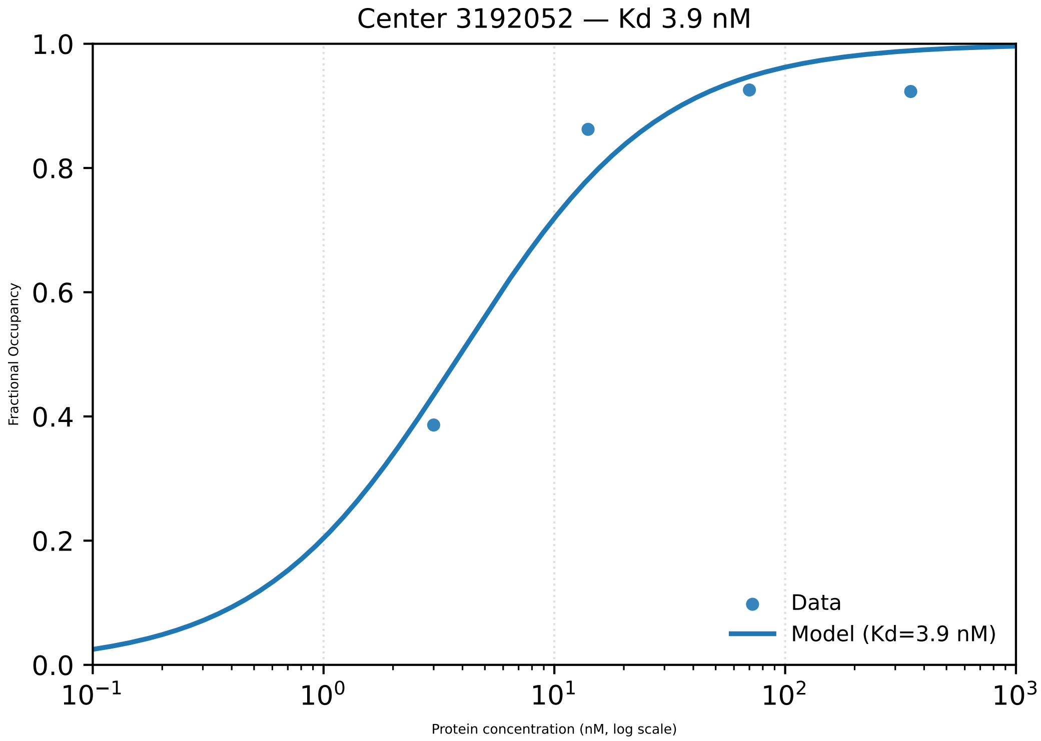

Center 3192052 — Kd 3.9 nM
1.0
0.8
0.6
0.4
0.2
0.0
Fractional Occupancy
$10^{-1}$
$10^{0}$
$10^{1}$
$10^{2}$
$10^{3}$
Protein concentration (nM, log scale)
Data
Model (Kd=3.9 nM)

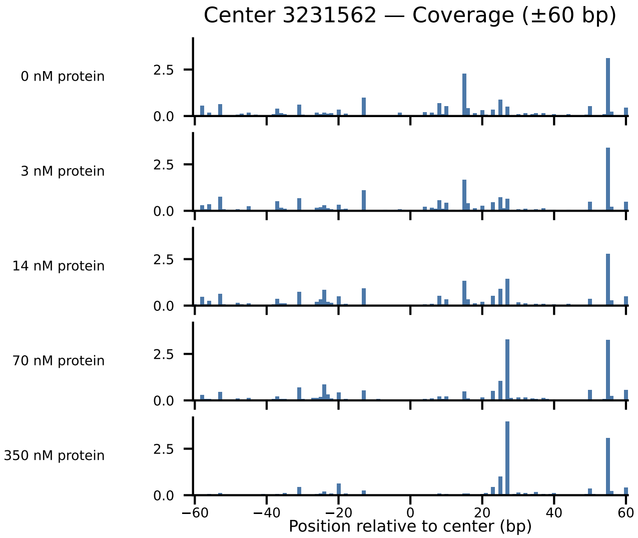

Center 3231562 — Coverage (±60 bp)
0 nM protein
3 nM protein
14 nM protein
70 nM protein
350 nM protein
2.5
0.0
−60
−40
−20
0
20
40
60
Position relative to center (bp)

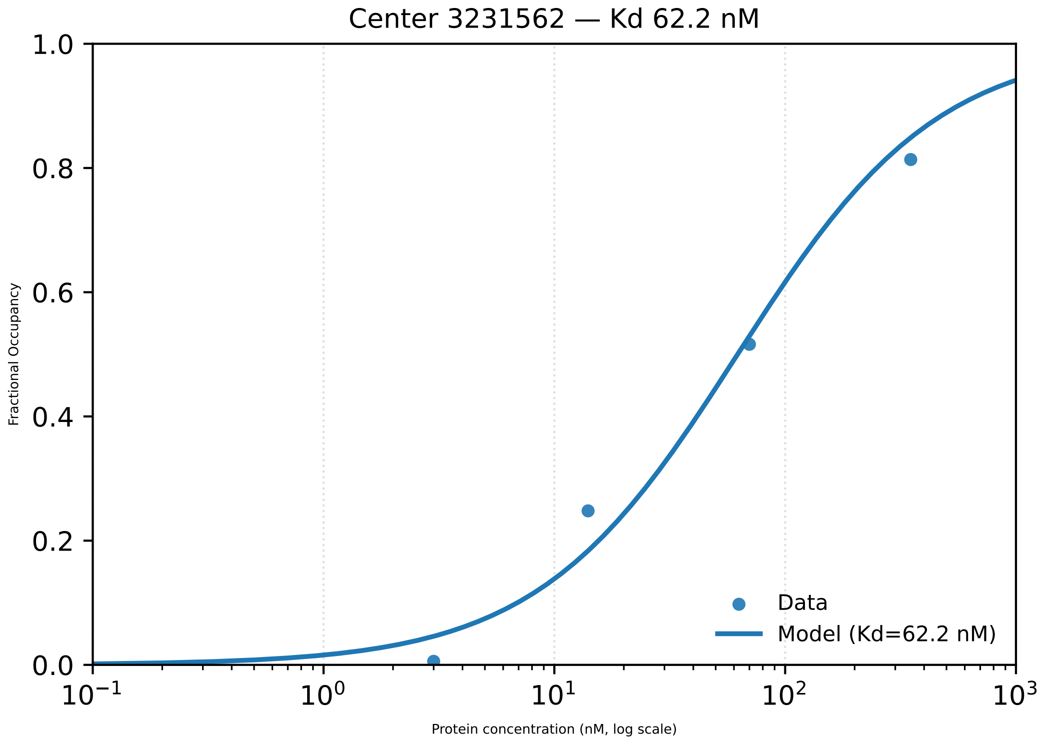

Center 3231562 — Kd 62.2 nM
Fractional Occupancy
1.0
0.8
0.6
0.4
0.2
0.0
$10^{-1}$
$10^{0}$
$10^{1}$
$10^{2}$
$10^{3}$
Protein concentration (nM, log scale)
Data
Model (Kd=62.2 nM)

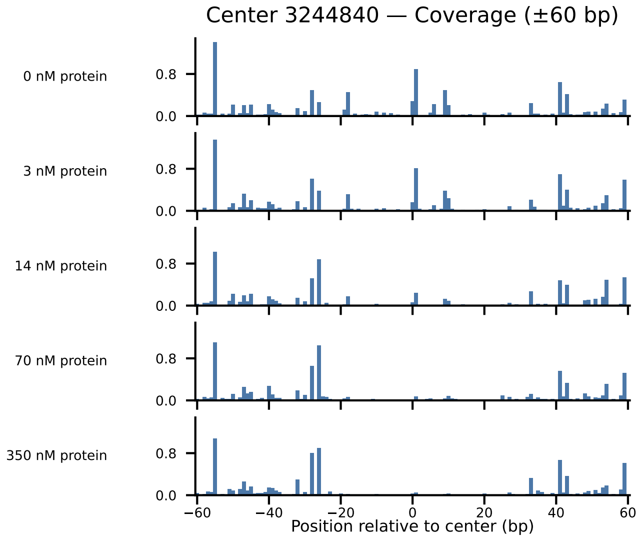

Center 3244840 — Coverage (±60 bp)
0 nM protein
3 nM protein
14 nM protein
70 nM protein
350 nM protein
0.8
0.0
−60
−40
−20
0
20
40
60
Position relative to center (bp)

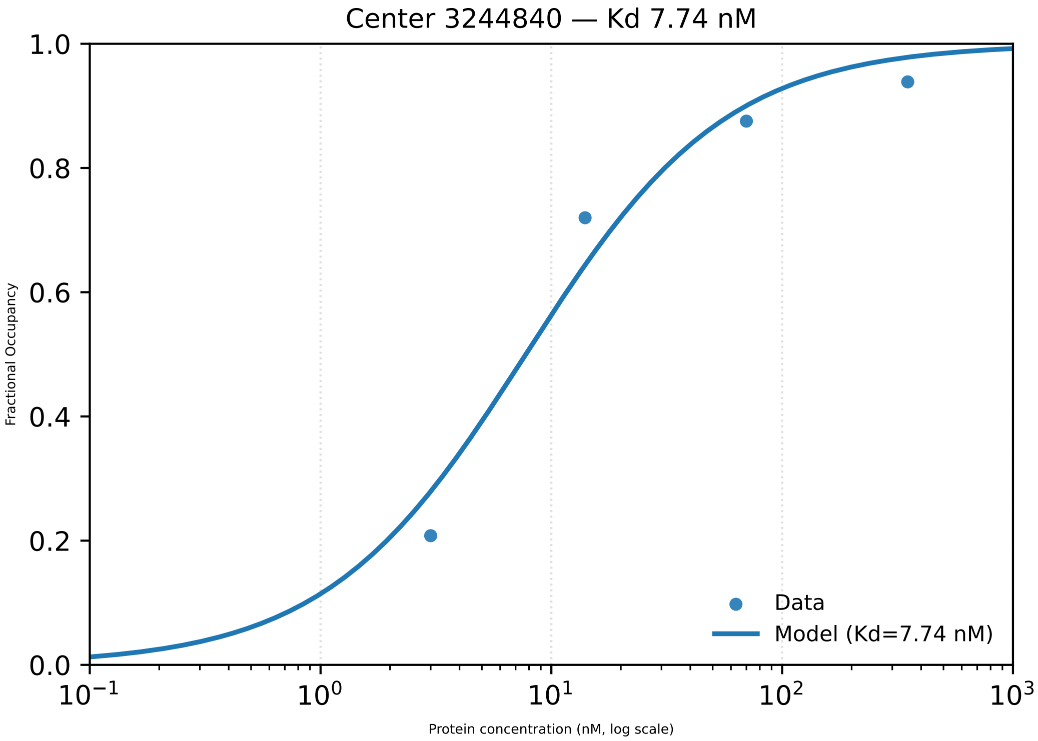

Center 3244840 — Kd 7.74 nM
Fractional Occupancy
1.0
0.8
0.6
0.4
0.2
0.0
$10^{-1}$
$10^{0}$
$10^{1}$
$10^{2}$
$10^{3}$
Protein concentration (nM, log scale)
Data
Model (Kd=7.74 nM)

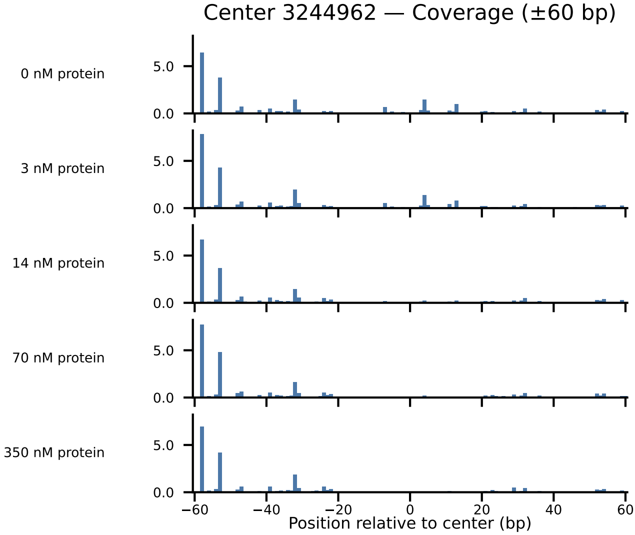

Center 3244962 — Coverage (±60 bp)
0 nM protein
3 nM protein
14 nM protein
70 nM protein
350 nM protein
5.0
0.0
−60
−40
−20
0
20
40
60
Position relative to center (bp)

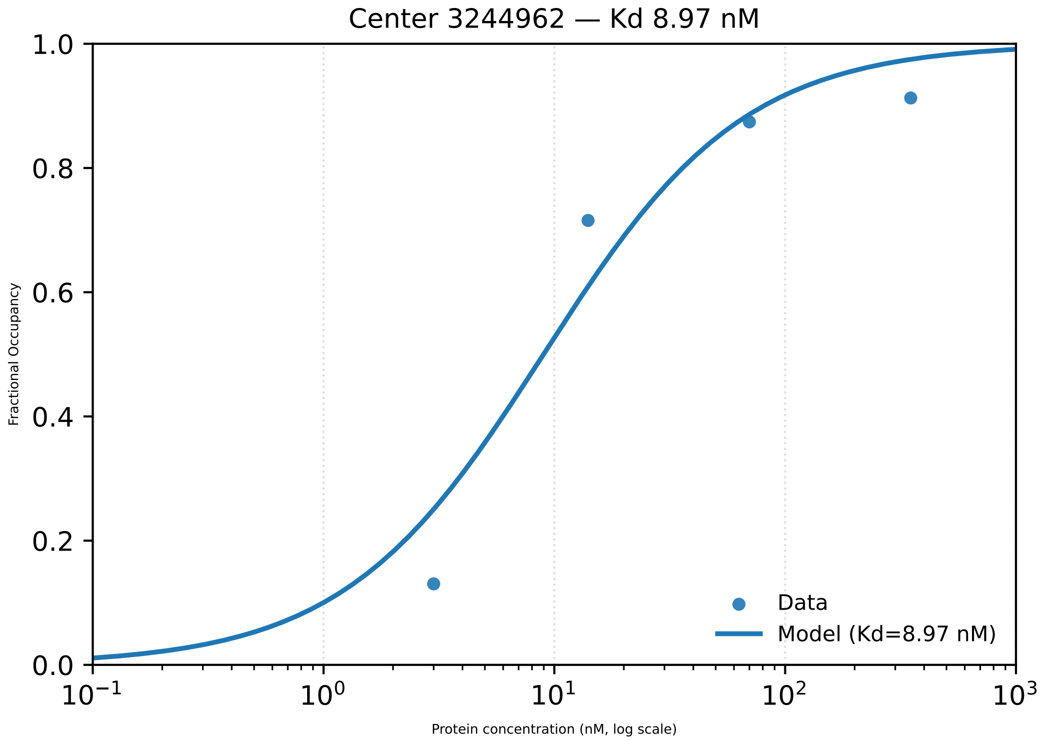

Center 3244962 — Kd 8.97 nM
1.0
0.8
0.6
0.4
0.2
0.0
Fractional Occupancy
$10^{-1}$
$10^{0}$
$10^{1}$
$10^{2}$
$10^{3}$
Protein concentration (nM, log scale)
Data
Model (Kd=8.97 nM)

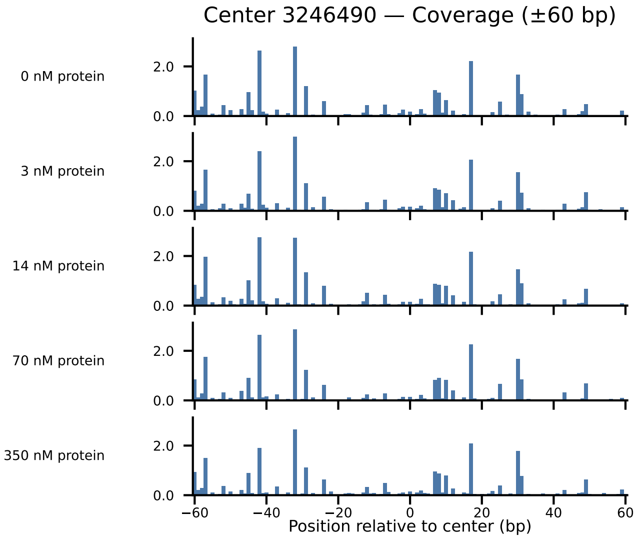

Center 3246490 — Coverage (±60 bp)
0 nM protein
3 nM protein
14 nM protein
70 nM protein
350 nM protein
2.0
0.0
−60
−40
−20
0
20
40
60
Position relative to center (bp)

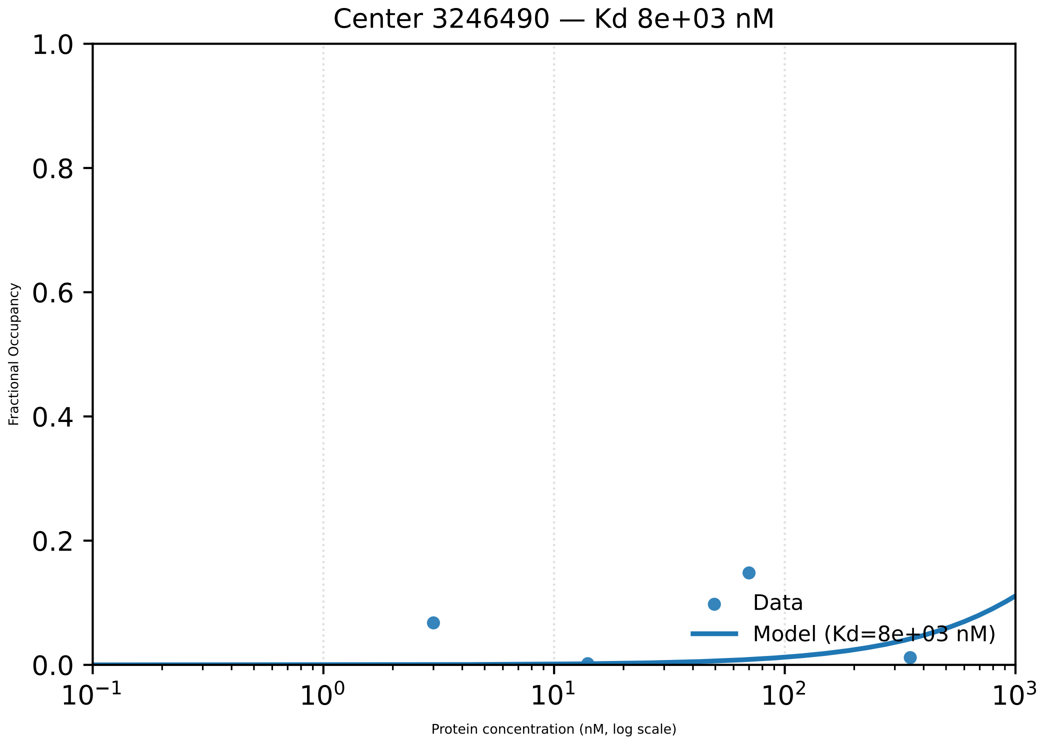

Center 3246490 — Kd 8e+03 nM
1.0
0.8
0.6
0.4
0.2
0.0
Fractional Occupancy
Data
Model (Kd=8e+03 nM)
$10^{-1}$
$10^{0}$
$10^{1}$
$10^{2}$
$10^{3}$
Protein concentration (nM, log scale)

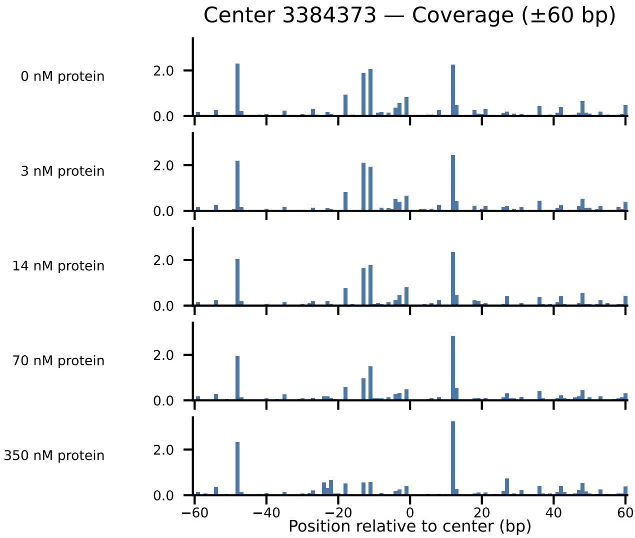
Center 3384373 — Coverage (±60 bp)
0 nM protein
3 nM protein
14 nM protein
70 nM protein
350 nM protein
2.0
0.0
−60
−40
−20
0
20
40
60
Position relative to center (bp)

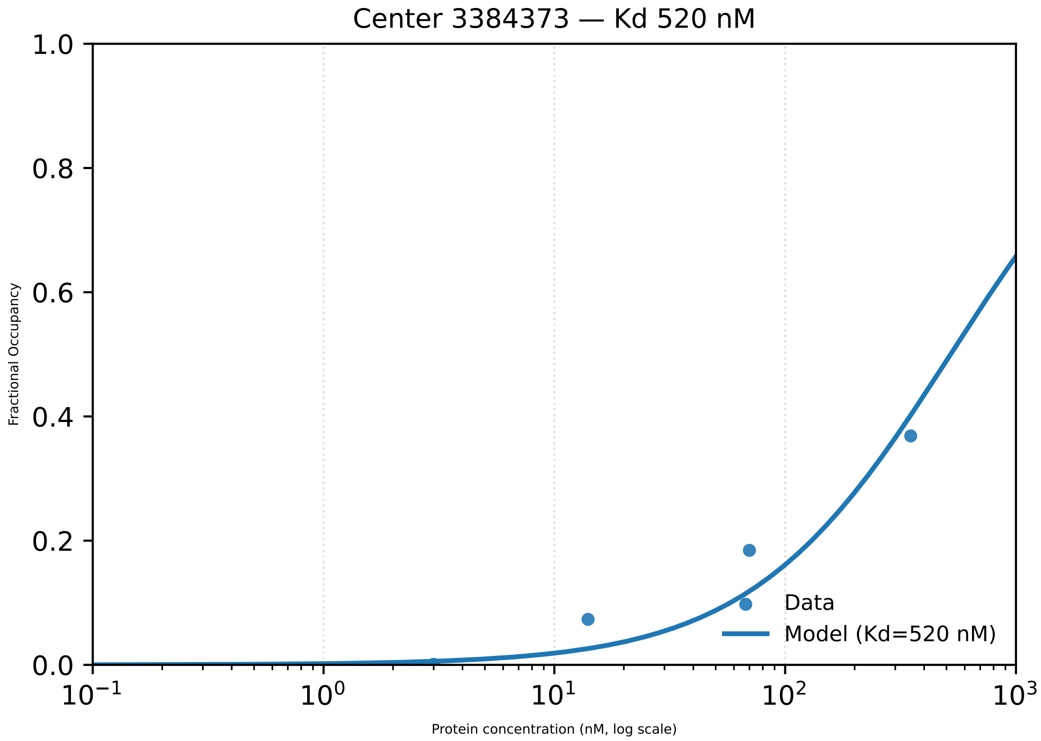

Center 3384373 — Kd 520 nM
Fractional Occupancy
1.0
0.8
0.6
0.4
0.2
0.0
$10^{-1}$
$10^{0}$
$10^{1}$
$10^{2}$
$10^{3}$
Protein concentration (nM, log scale)
Data
Model (Kd=520 nM)

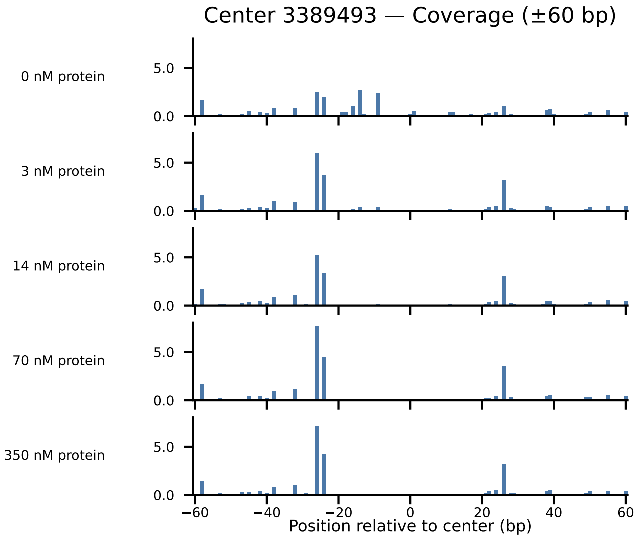
Center 3389493 — Coverage (±60 bp)
0 nM protein
3 nM protein
14 nM protein
70 nM protein
350 nM protein
5.0
0.0
−60
−40
−20
0
20
40
60
Position relative to center (bp)

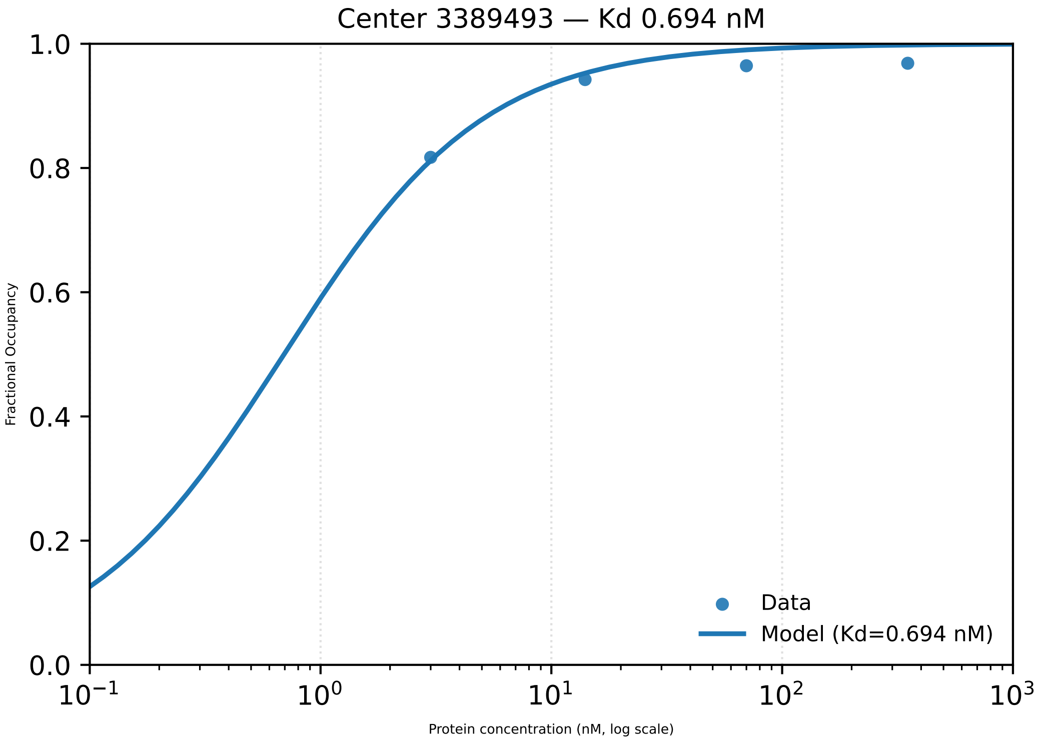

Center 3389493 — Kd 0.694 nM
1.0
0.8
0.6
0.4
0.2
0.0
Fractional Occupancy
$10^{-1}$
$10^{0}$
$10^{1}$
$10^{2}$
$10^{3}$
Protein concentration (nM, log scale)
Data
Model (Kd=0.694 nM)

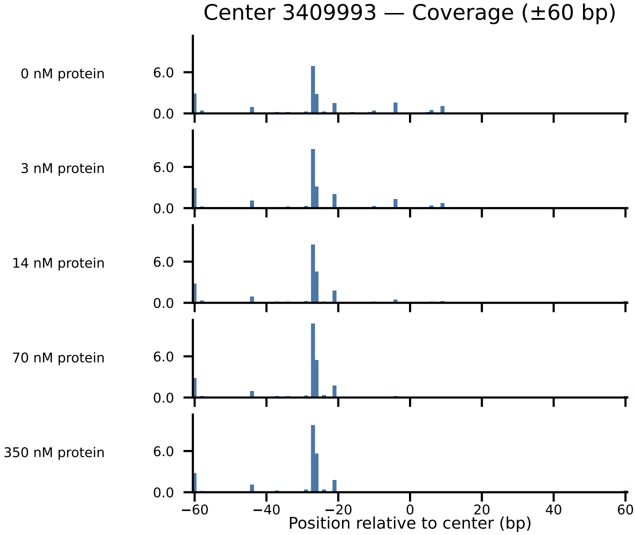

Center 3409993 — Coverage (±60 bp)
0 nM protein
3 nM protein
14 nM protein
70 nM protein
350 nM protein
6.0
0.0
−60
−40
−20
0
20
40
60
Position relative to center (bp)

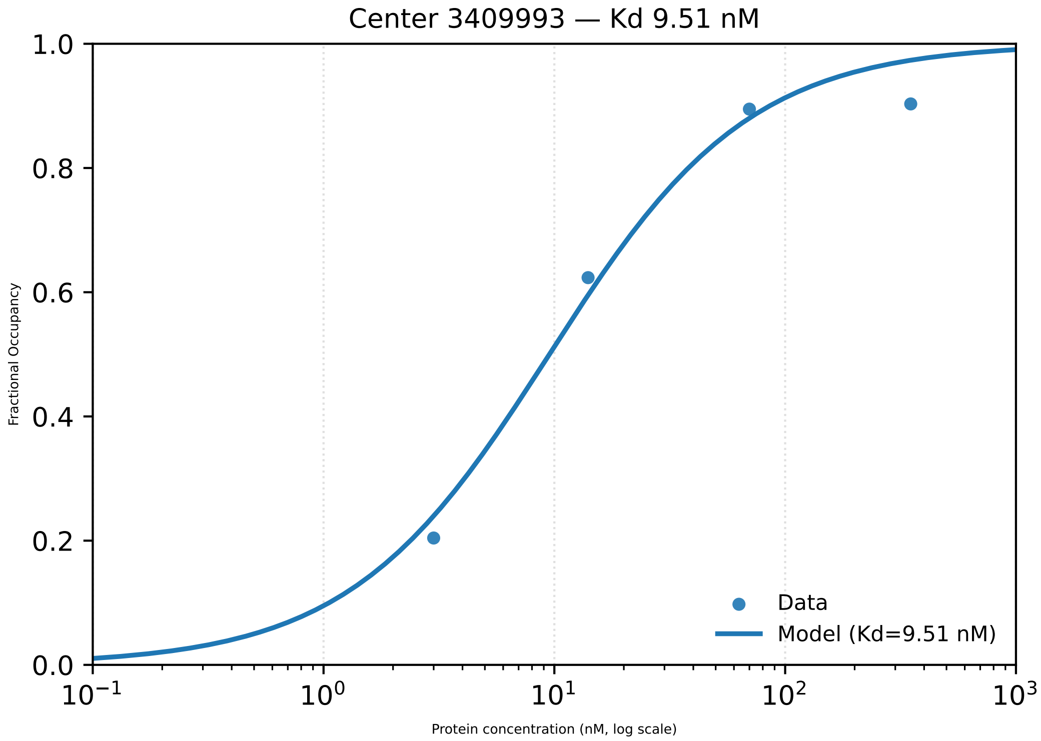

Center 3409993 — Kd 9.51 nM
Fractional Occupancy
1.0
0.8
0.6
0.4
0.2
0.0
$10^{-1}$
$10^{0}$
$10^{1}$
$10^{2}$
$10^{3}$
Protein concentration (nM, log scale)
Data
Model (Kd=9.51 nM)

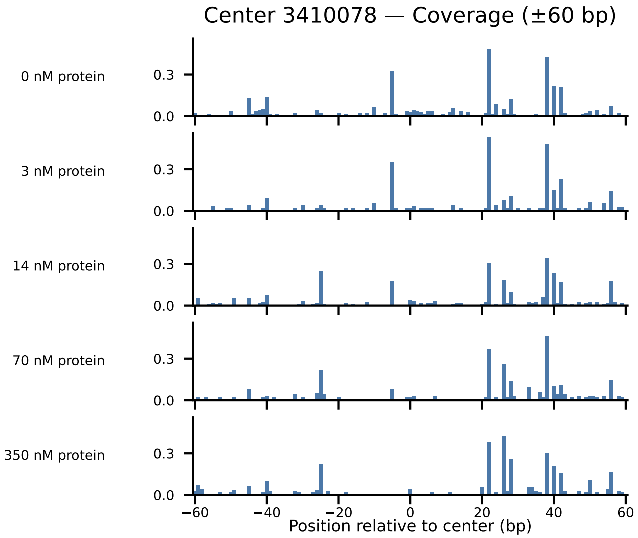

Center 3410078 — Coverage (±60 bp)
0 nM protein
3 nM protein
14 nM protein
70 nM protein
350 nM protein
0.3
0.0
−60
−40
−20
0
20
40
60
Position relative to center (bp)

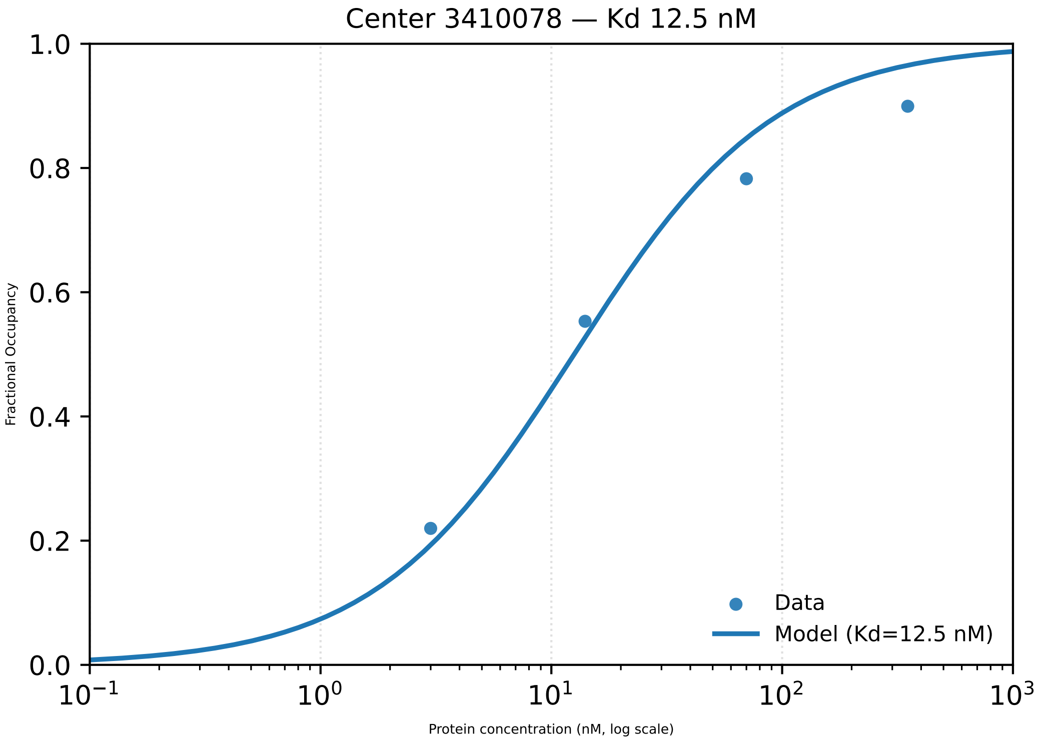

Center 3410078 — Kd 12.5 nM
Fractional Occupancy
1.0
0.8
0.6
0.4
0.2
0.0
$10^{-1}$
$10^{0}$
$10^{1}$
$10^{2}$
$10^{3}$
Protein concentration (nM, log scale)
Data
Model (Kd=12.5 nM)

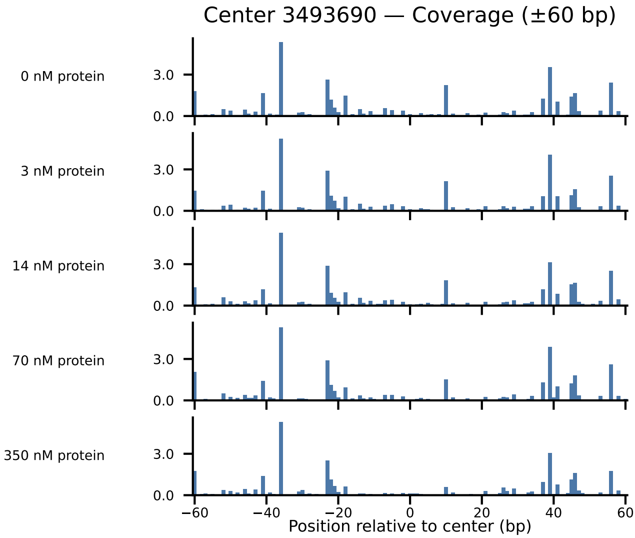

Center 3493690 — Coverage (±60 bp)
0 nM protein
3 nM protein
14 nM protein
70 nM protein
350 nM protein
3.0
0.0
−60
−40
−20
0
20
40
60
Position relative to center (bp)

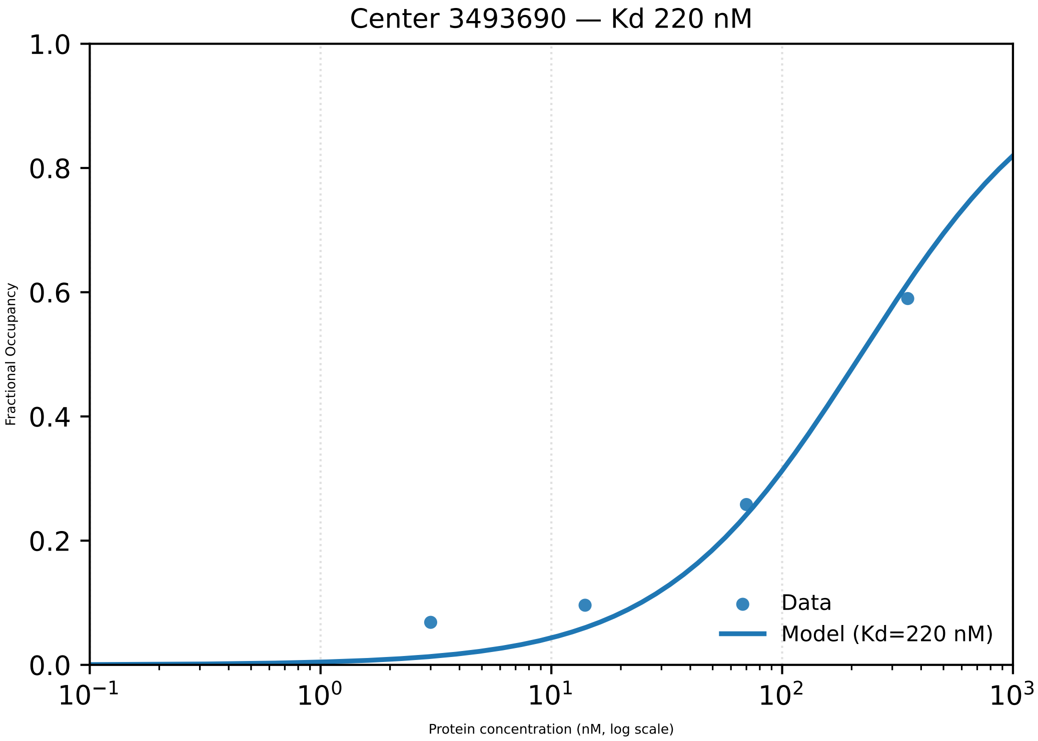
Center 3493690 — Kd 220 nM
Fractional Occupancy
1.0
0.8
0.6
0.4
0.2
0.0
$10^{-1}$
$10^{0}$
$10^{1}$
$10^{2}$
$10^{3}$
Protein concentration (nM, log scale)
Data
Model (Kd=220 nM)

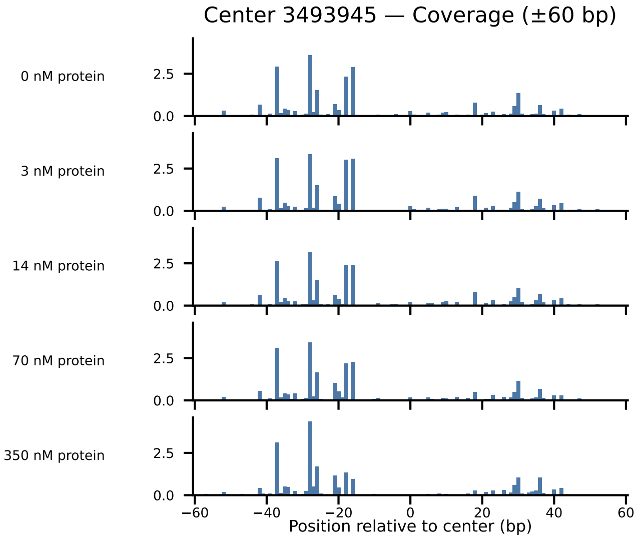
Center 3493945 — Coverage (±60 bp)
0 nM protein
3 nM protein
14 nM protein
70 nM protein
350 nM protein
2.5
0.0
−60
−40
−20
0
20
40
60
Position relative to center (bp)

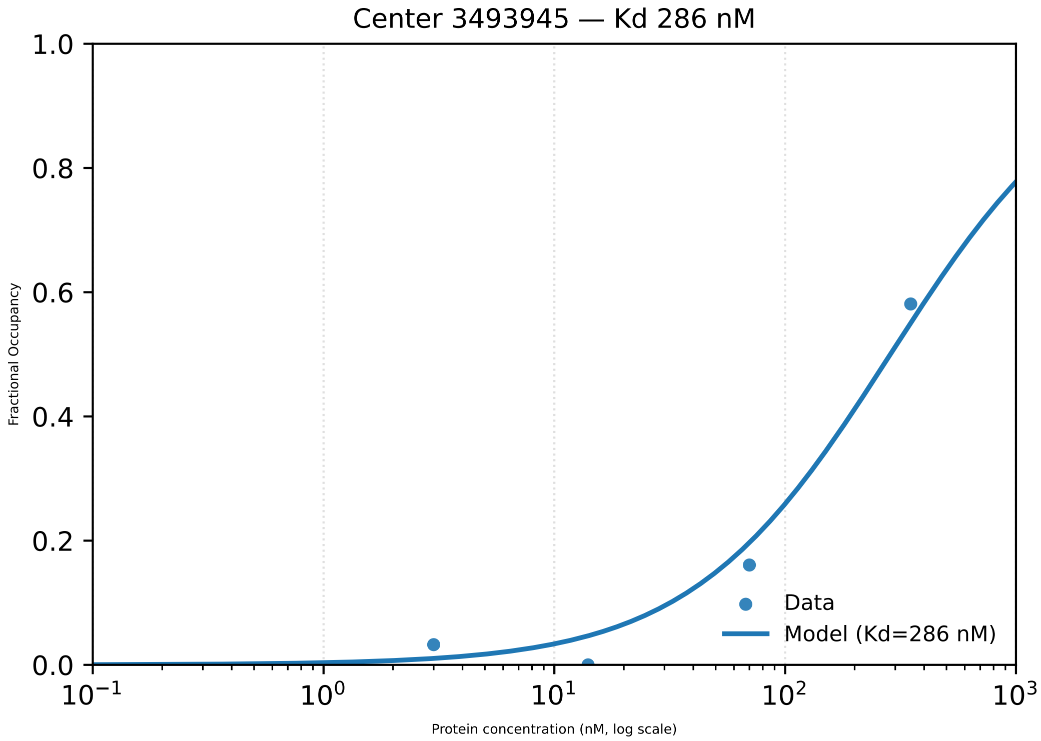
Center 3493945 — Kd 286 nM
Fractional Occupancy
1.0
0.8
0.6
0.4
0.2
0.0
$10^{-1}$
$10^{0}$
$10^{1}$
$10^{2}$
$10^{3}$
Protein concentration (nM, log scale)
Data
Model (Kd=286 nM)

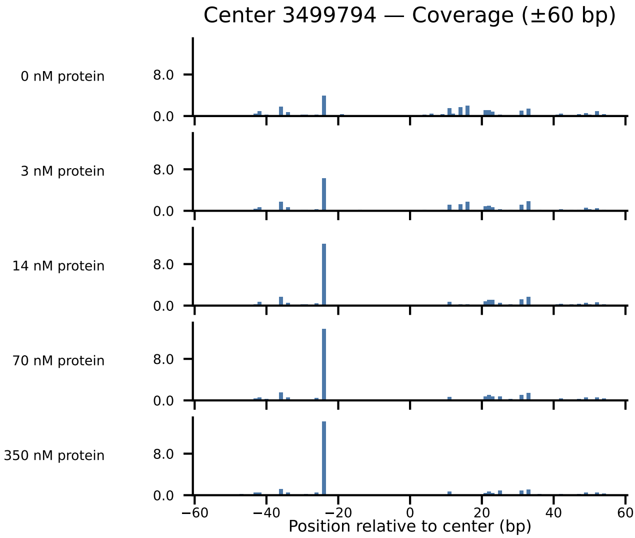
Center 3499794 — Coverage (±60 bp)
0 nM protein
3 nM protein
14 nM protein
70 nM protein
350 nM protein
8.0
0.0
−60
−40
−20
0
20
40
60
Position relative to center (bp)

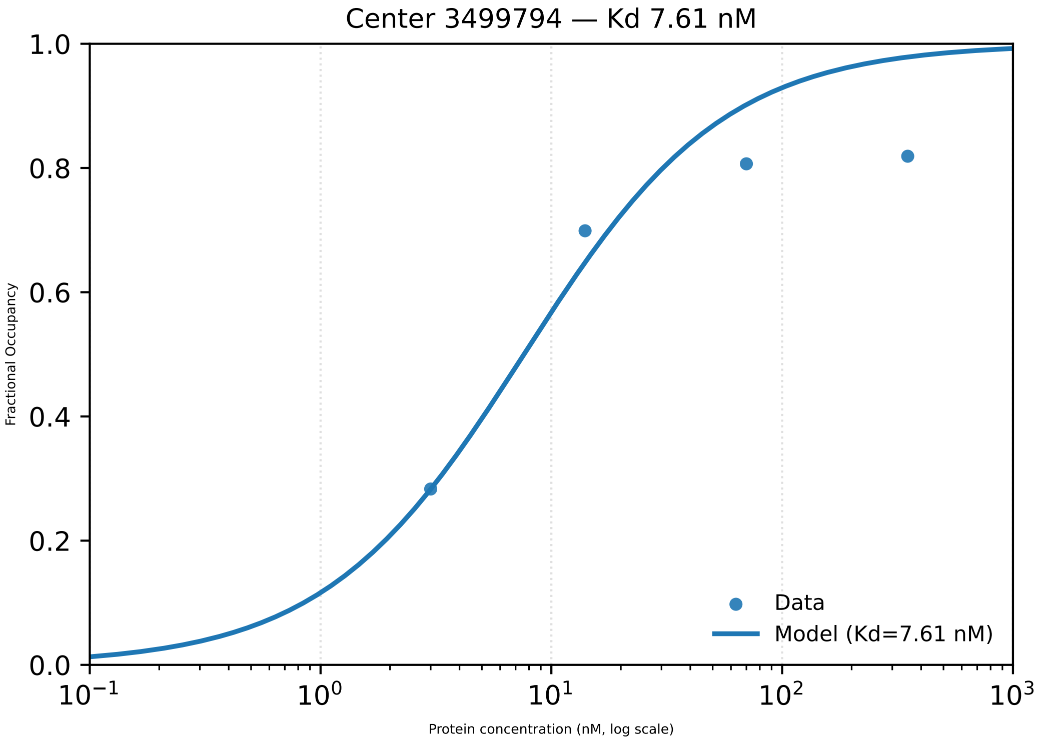
Center 3499794 — Kd 7.61 nM
1.0
0.8
0.6
0.4
0.2
0.0
Fractional Occupancy
$10^{-1}$
$10^{0}$
$10^{1}$
$10^{2}$
$10^{3}$
Protein concentration (nM, log scale)
Data
Model (Kd=7.61 nM)

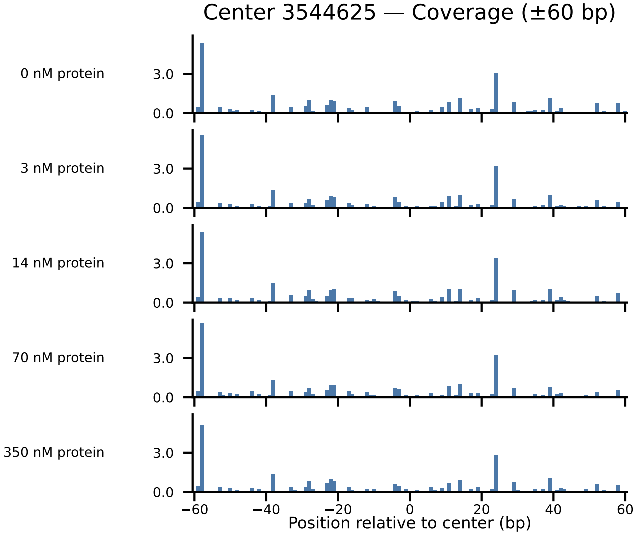
Center 3544625 — Coverage (±60 bp)
0 nM protein
3 nM protein
14 nM protein
70 nM protein
350 nM protein
3.0
0.0
−60
−40
−20
0
20
40
60
Position relative to center (bp)

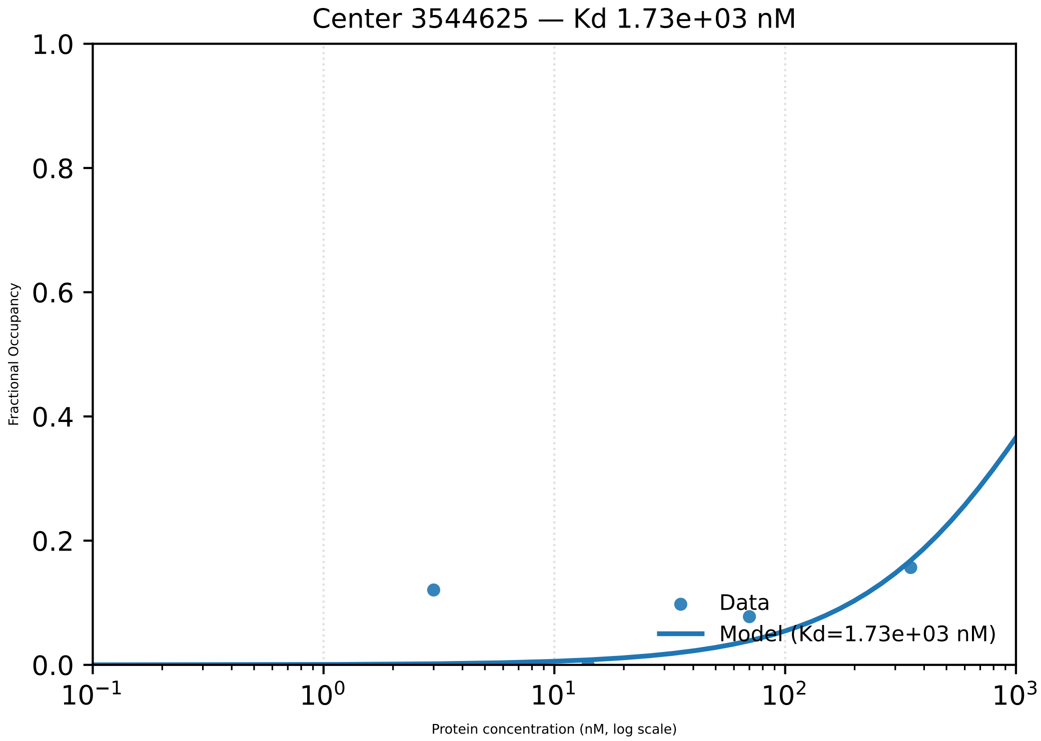
Center 3544625 — Kd 1.73e+03 nM
1.0
0.8
0.6
0.4
0.2
0.0
Fractional Occupancy
Data
Model (Kd=1.73e+03 nM)
$10^{-1}$
$10^{0}$
$10^{1}$
$10^{2}$
$10^{3}$
Protein concentration (nM, log scale)

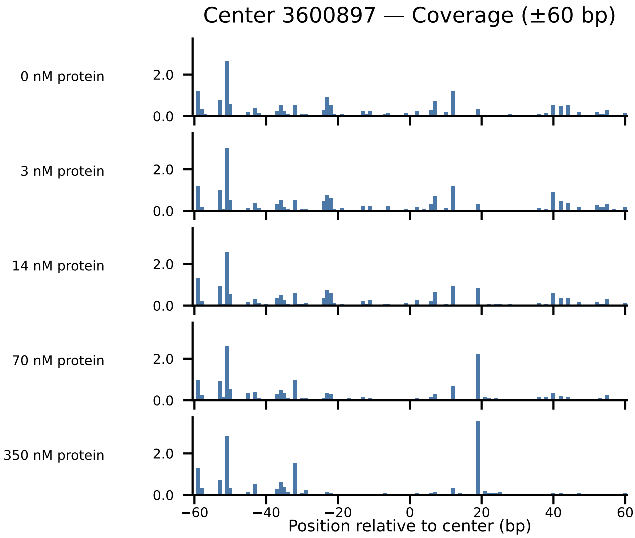
Center 3600897 — Coverage (±60 bp)
0 nM protein
3 nM protein
14 nM protein
70 nM protein
350 nM protein
2.0
0.0
−60
−40
−20
0
20
40
60
Position relative to center (bp)

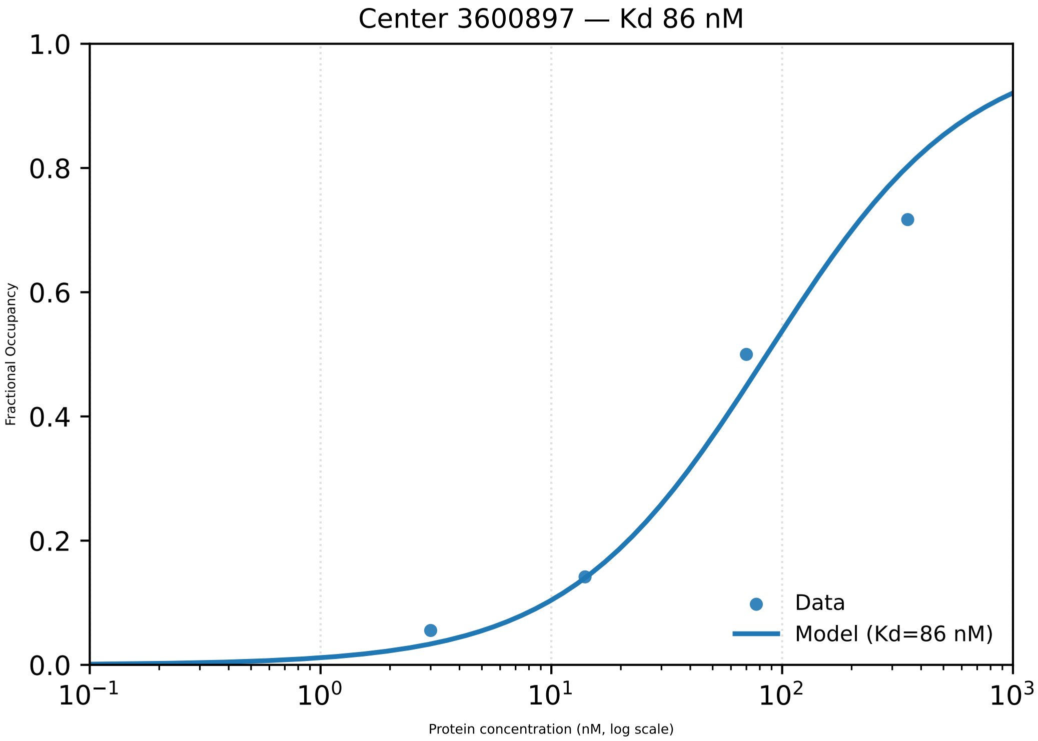

Center 3600897 — Kd 86 nM
1.0
0.8
0.6
0.4
0.2
0.0
Fractional Occupancy
$10^{-1}$
$10^{0}$
$10^{1}$
$10^{2}$
$10^{3}$
Protein concentration (nM, log scale)
Data
Model (Kd=86 nM)

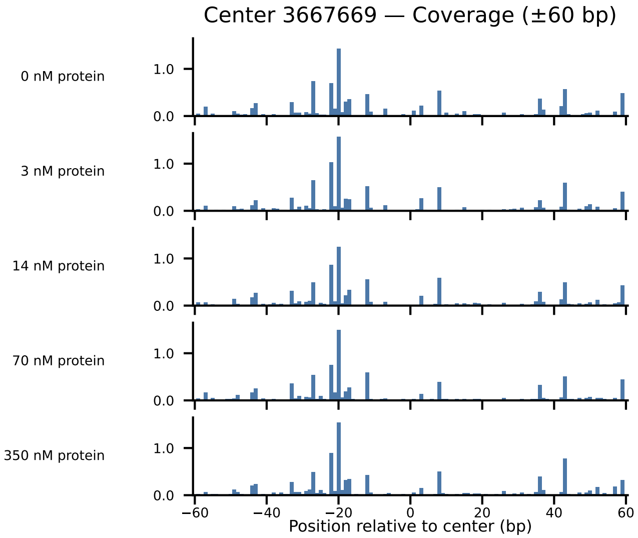
Center 3667669 — Coverage (±60 bp)
0 nM protein
3 nM protein
14 nM protein
70 nM protein
350 nM protein
1.0
0.0
−60
−40
−20
0
20
40
60
Position relative to center (bp)

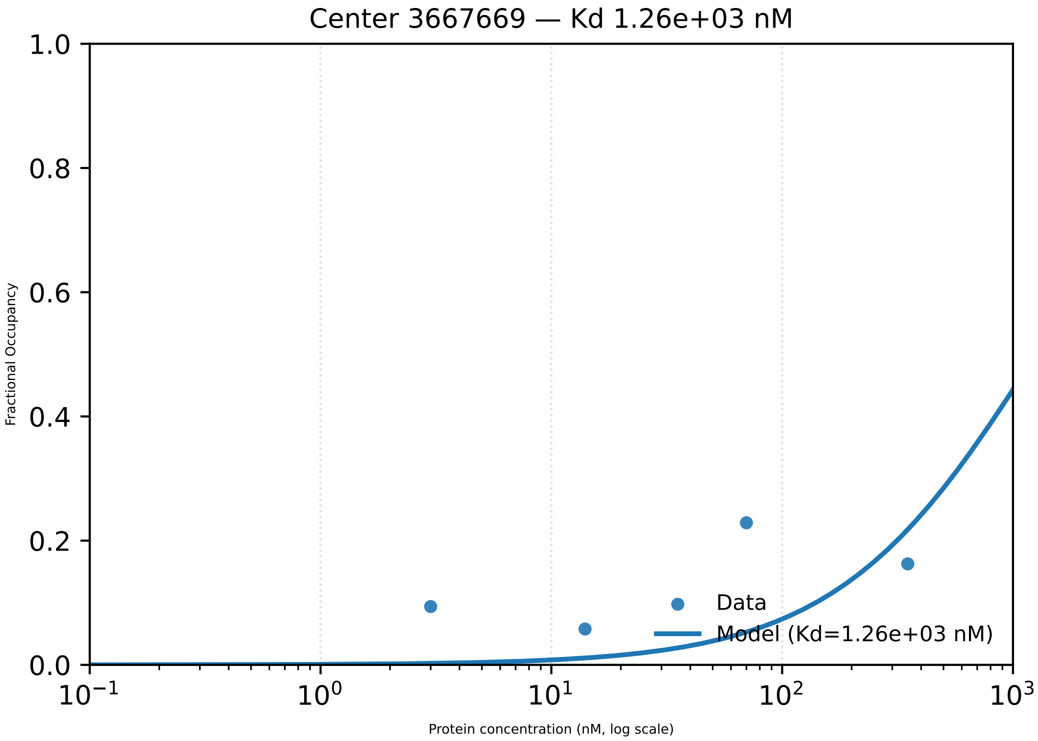

Center 3667669 — Kd 1.26e+03 nM
Fractional Occupancy
1.0
0.8
0.6
0.4
0.2
0.0
$10^{-1}$
$10^{0}$
$10^{1}$
$10^{2}$
$10^{3}$
Protein concentration (nM, log scale)
Data
Model (Kd=1.26e+03 nM)

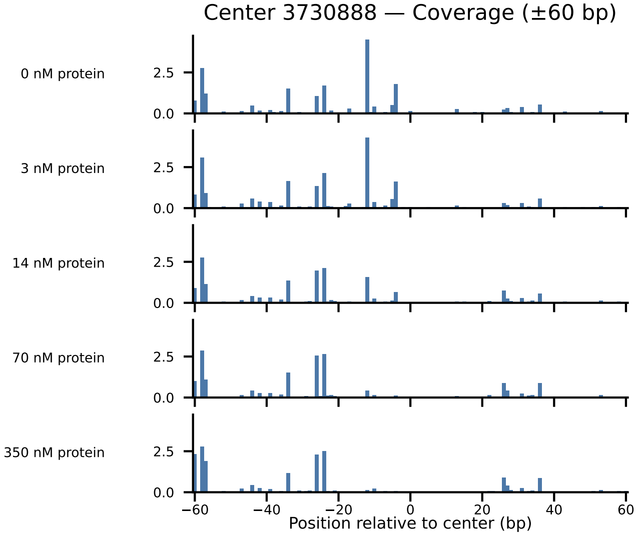
Center 3730888 — Coverage (±60 bp)
0 nM protein
3 nM protein
14 nM protein
70 nM protein
350 nM protein
2.5
0.0
−60
−40
−20
0
20
40
60
Position relative to center (bp)

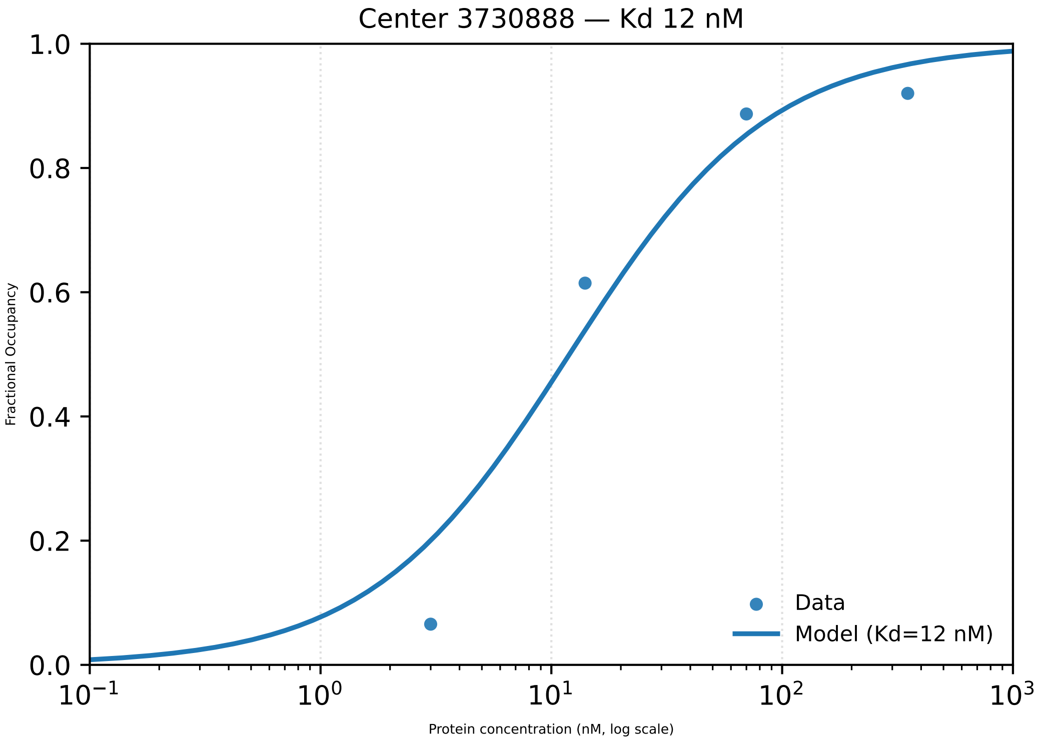

Center 3730888 — Kd 12 nM
Fractional Occupancy
1.0
0.8
0.6
0.4
0.2
0.0
$10^{-1}$
$10^{0}$
$10^{1}$
$10^{2}$
$10^{3}$
Protein concentration (nM, log scale)
Data
Model (Kd=12 nM)

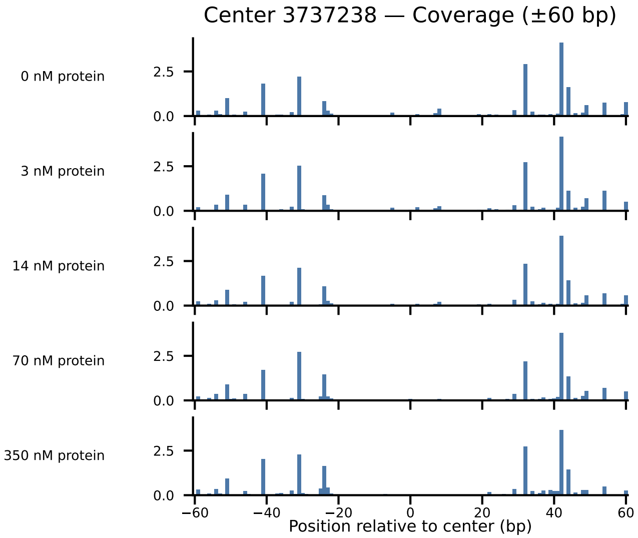

Center 3737238 — Coverage (±60 bp)
0 nM protein
3 nM protein
14 nM protein
70 nM protein
350 nM protein
2.5
0.0
−60
−40
−20
0
20
40
60
Position relative to center (bp)

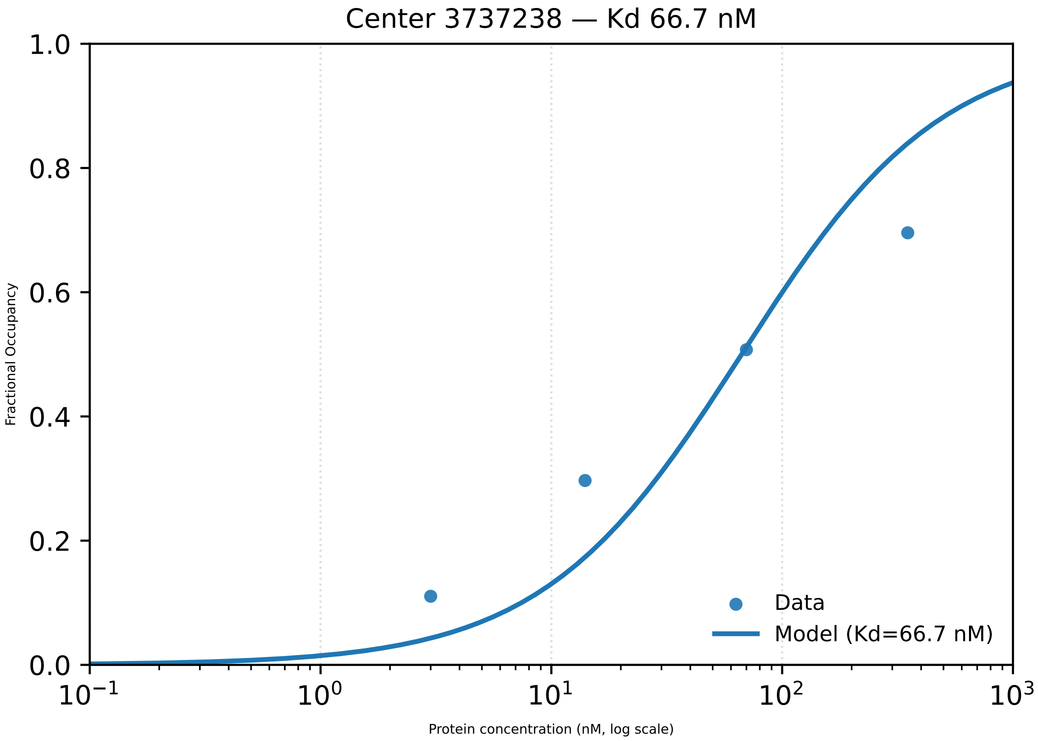

Center 3737238 — Kd 66.7 nM
Fractional Occupancy
Protein concentration (nM, log scale)
Data
Model (Kd=66.7 nM)
1.0
0.8
0.6
0.4
0.2
0.0
$10^{-1}$
$10^{0}$
$10^{1}$
$10^{2}$
$10^{3}$

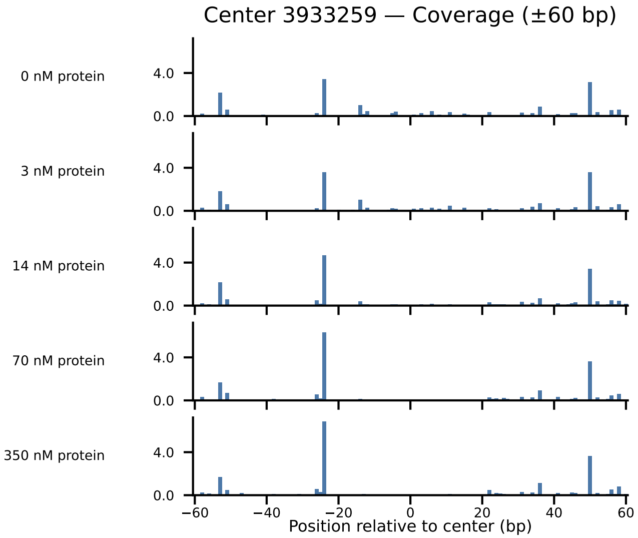

Center 3933259 — Coverage (±60 bp)
0 nM protein
3 nM protein
14 nM protein
70 nM protein
350 nM protein
4.0
0.0
−60
−40
−20
0
20
40
60
Position relative to center (bp)

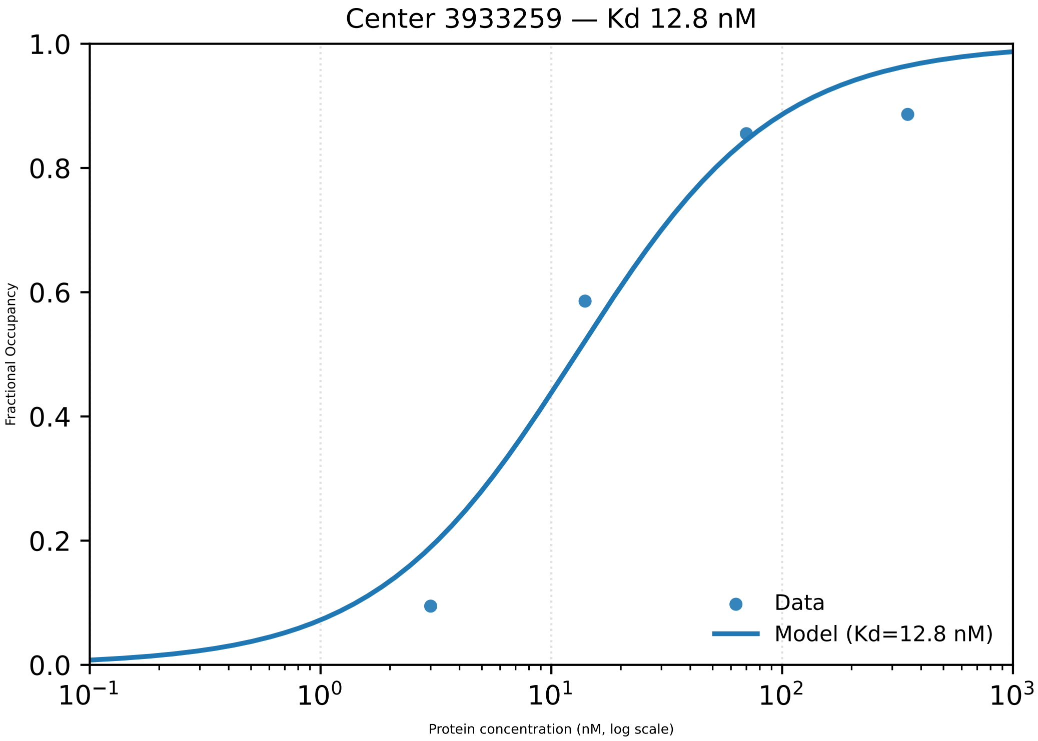
Center 3933259 — Kd 12.8 nM
1.0
0.8
0.6
0.4
0.2
0.0
Fractional Occupancy
$10^{-1}$
$10^{0}$
$10^{1}$
$10^{2}$
$10^{3}$
Protein concentration (nM, log scale)
Data
Model (Kd=12.8 nM)

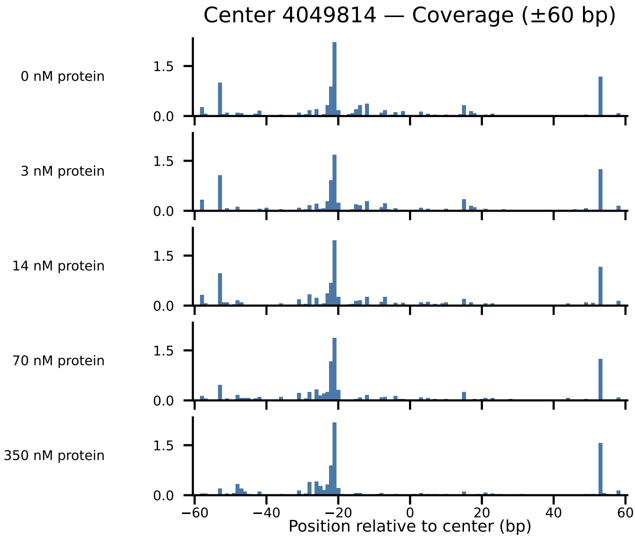

Center 4049814 — Coverage (±60 bp)
0 nM protein
3 nM protein
14 nM protein
70 nM protein
350 nM protein
1.5
0.0
−60
−40
−20
0
20
40
60
Position relative to center (bp)

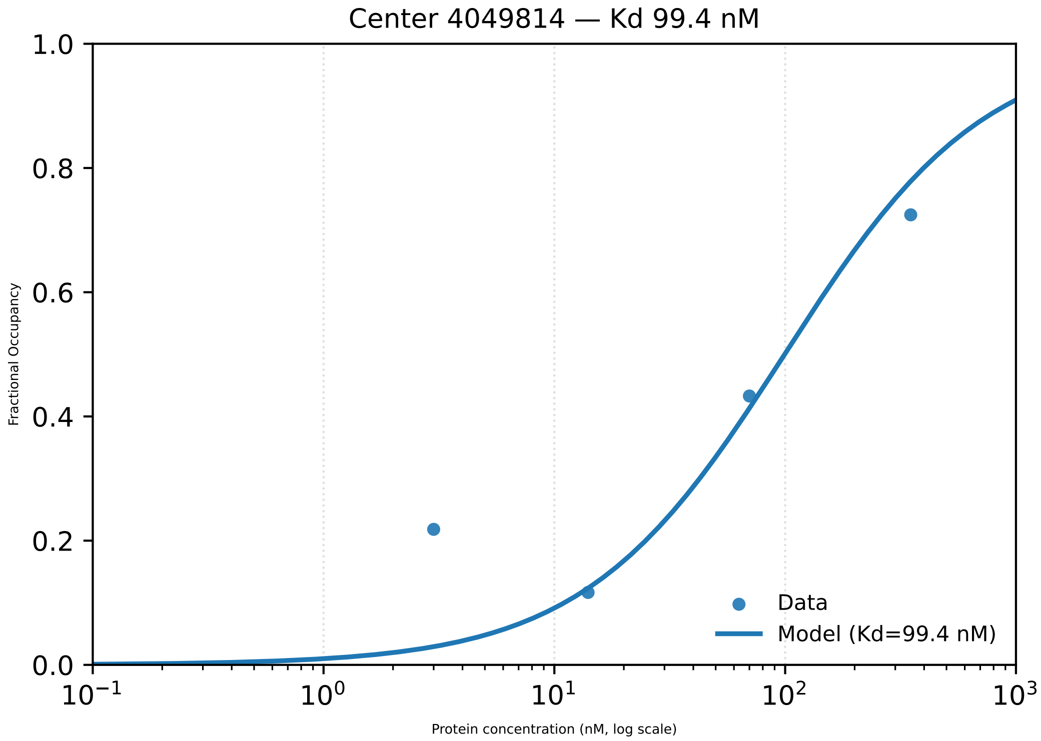
Center 4049814 — Kd 99.4 nM
1.0
0.8
0.6
0.4
0.2
0.0
Fractional Occupancy
$10^{-1}$
$10^{0}$
$10^{1}$
$10^{2}$
$10^{3}$
Protein concentration (nM, log scale)
Data
Model (Kd=99.4 nM)

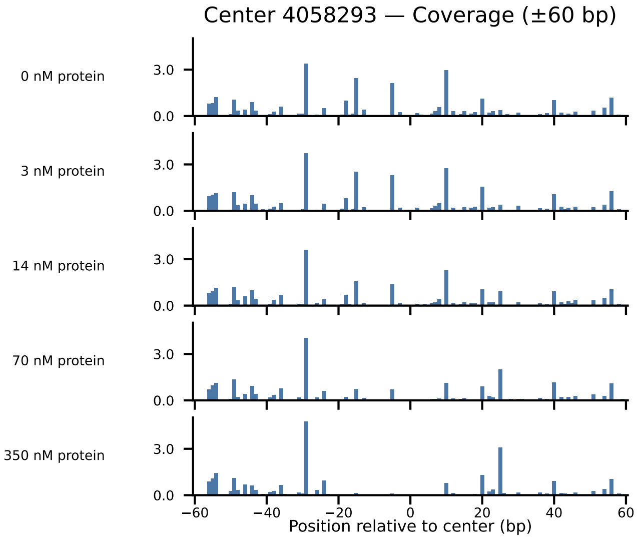
Center 4058293 — Coverage (±60 bp)
0 nM protein
3 nM protein
14 nM protein
70 nM protein
350 nM protein
3.0
0.0
−60
−40
−20
0
20
40
60
Position relative to center (bp)

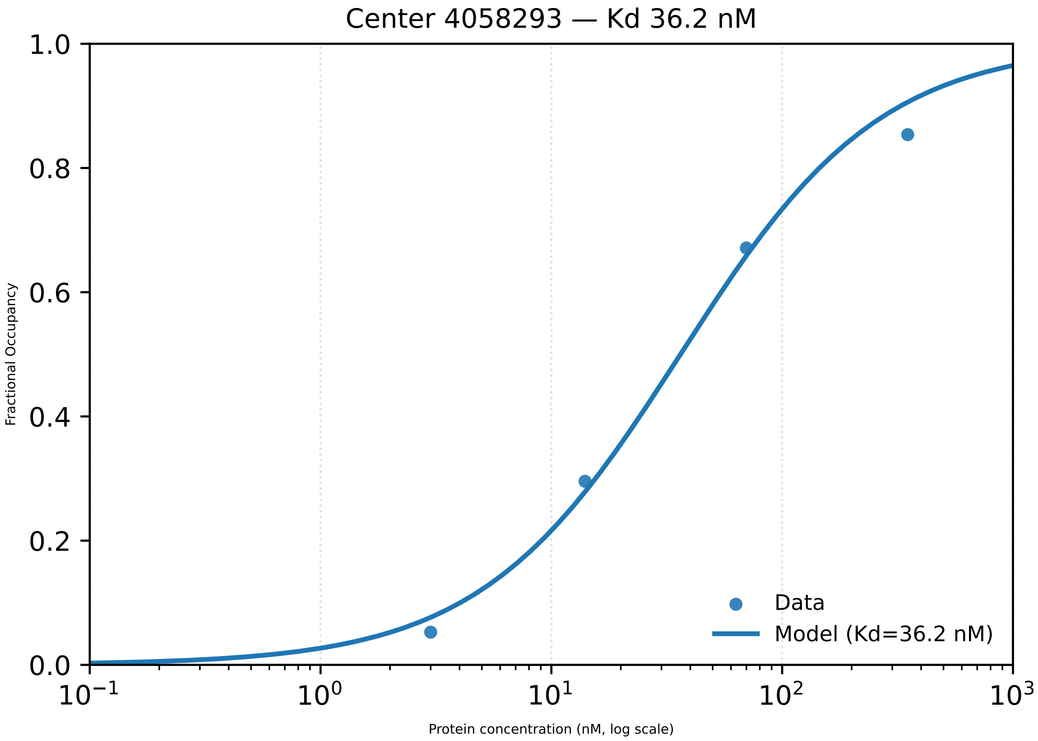

Center 4058293 — Kd 36.2 nM
Fractional Occupancy
1.0
0.8
0.6
0.4
0.2
0.0
$10^{-1}$
$10^{0}$
$10^{1}$
$10^{2}$
$10^{3}$
Protein concentration (nM, log scale)
Data
Model (Kd=36.2 nM)

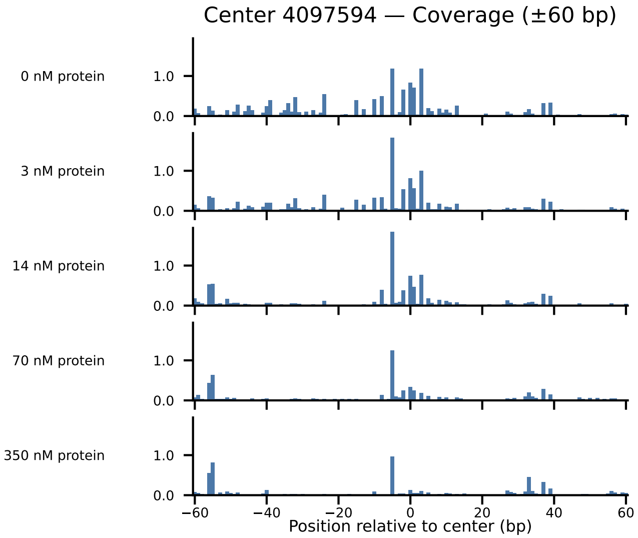

Center 4097594 — Coverage (±60 bp)
0 nM protein
3 nM protein
14 nM protein
70 nM protein
350 nM protein
1.0
0.0
−60
−40
−20
0
20
40
60
Position relative to center (bp)

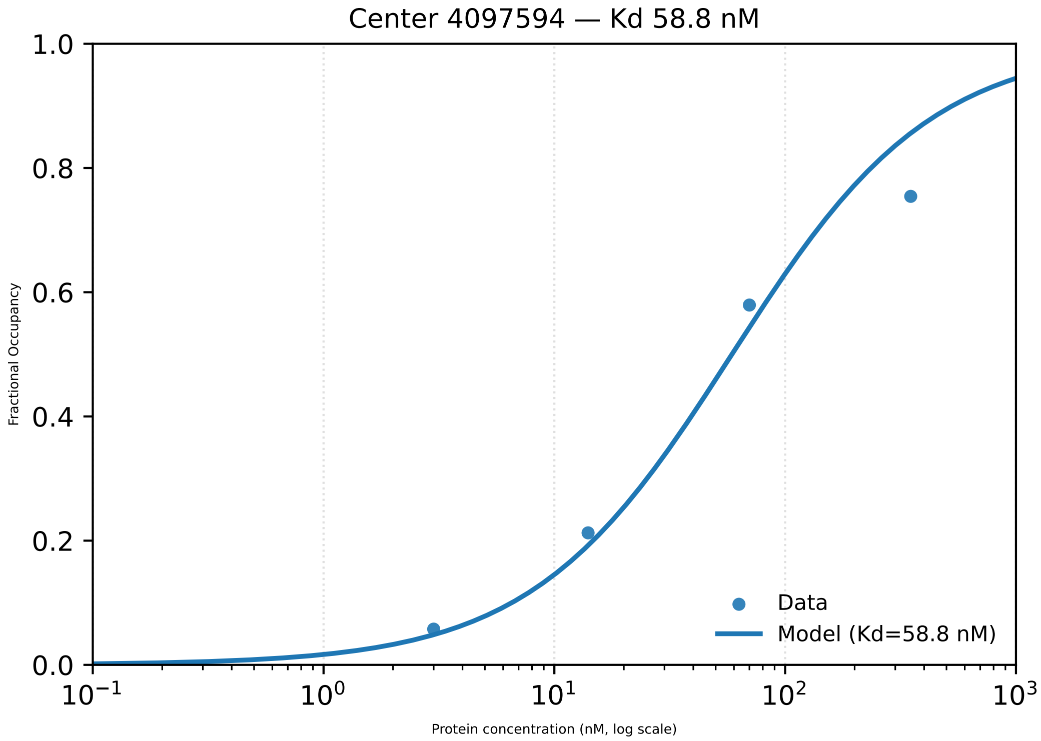

Center 4097594 — Kd 58.8 nM
1.0
0.8
0.6
0.4
0.2
0.0
Fractional Occupancy
$10^{-1}$
$10^{0}$
$10^{1}$
$10^{2}$
$10^{3}$
Protein concentration (nM, log scale)
Data
Model (Kd=58.8 nM)

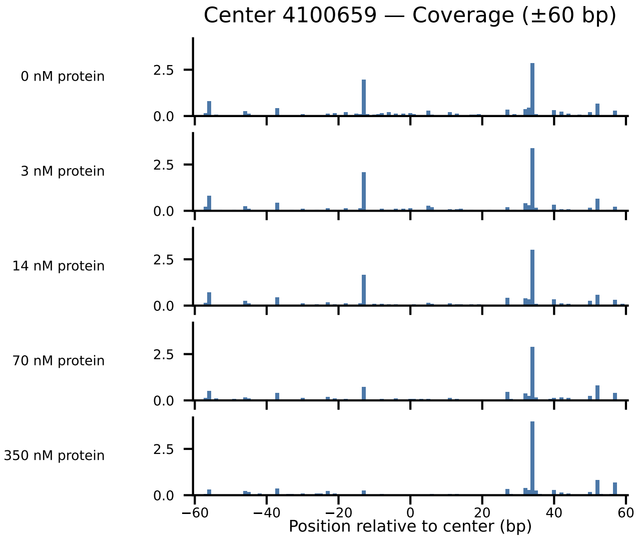

Center 4100659 — Coverage (±60 bp)
0 nM protein
3 nM protein
14 nM protein
70 nM protein
350 nM protein
2.5
0.0
−60
−40
−20
0
20
40
60
Position relative to center (bp)

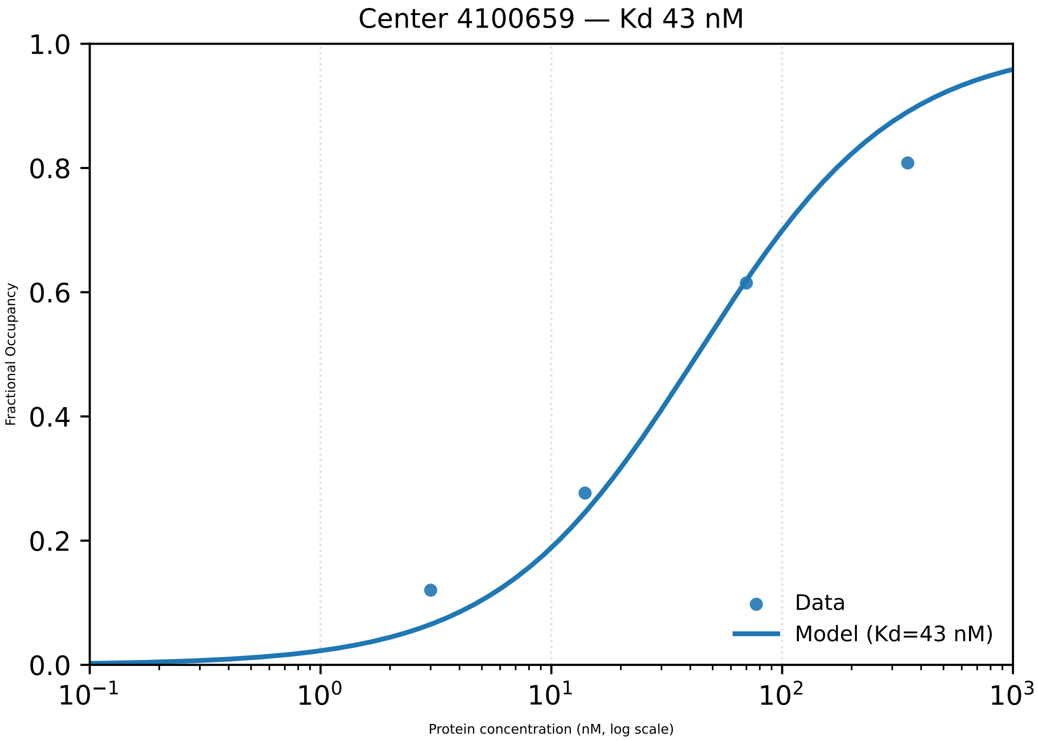

Center 4100659 — Kd 43 nM
Fractional Occupancy
Protein concentration (nM, log scale)
Data
Model (Kd=43 nM)
1.0
0.8
0.6
0.4
0.2
0.0
$10^{-1}$
$10^{0}$
$10^{1}$
$10^{2}$
$10^{3}$

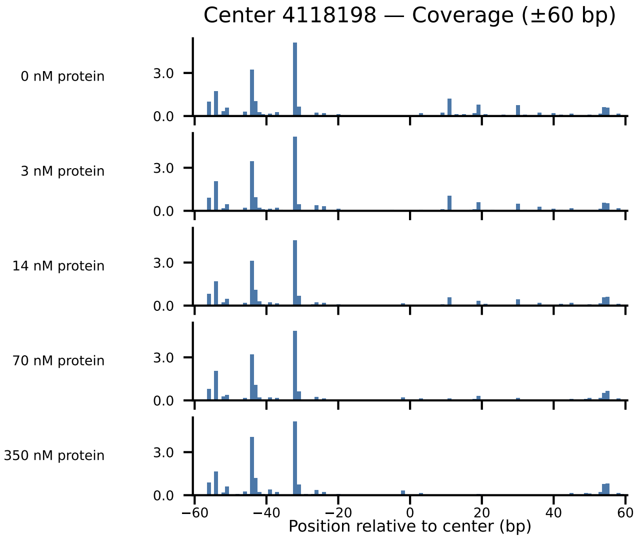

Center 4118198 — Coverage (±60 bp)
0 nM protein
3 nM protein
14 nM protein
70 nM protein
350 nM protein
3.0
0.0
−60
−40
−20
0
20
40
60
Position relative to center (bp)

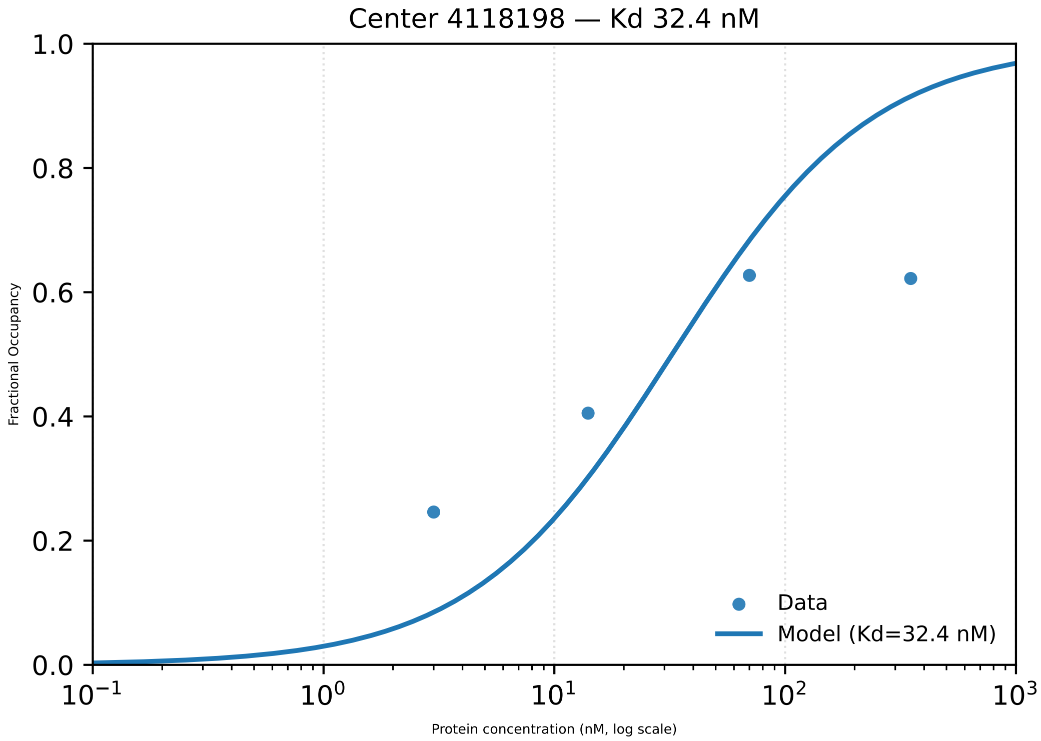
Center 4118198 — Kd 32.4 nM
Fractional Occupancy
1.0
0.8
0.6
0.4
0.2
0.0
$10^{-1}$
$10^{0}$
$10^{1}$
$10^{2}$
$10^{3}$
Protein concentration (nM, log scale)
Data
Model (Kd=32.4 nM)

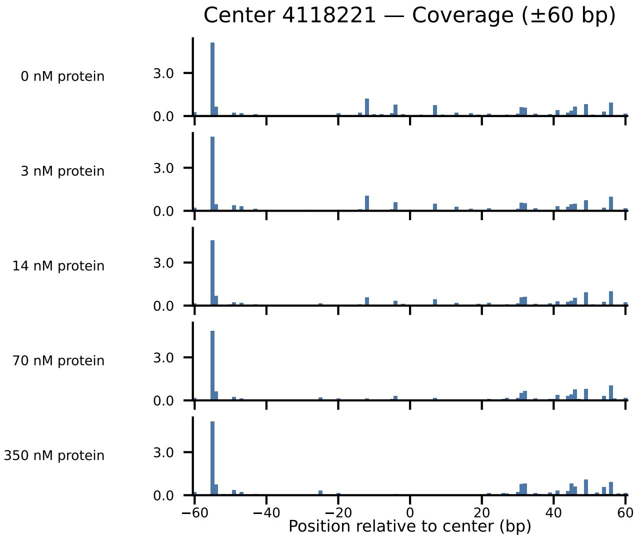

Center 4118221 — Coverage (±60 bp)
0 nM protein
3 nM protein
14 nM protein
70 nM protein
350 nM protein
3.0
0.0
−60
−40
−20
0
20
40
60
Position relative to center (bp)

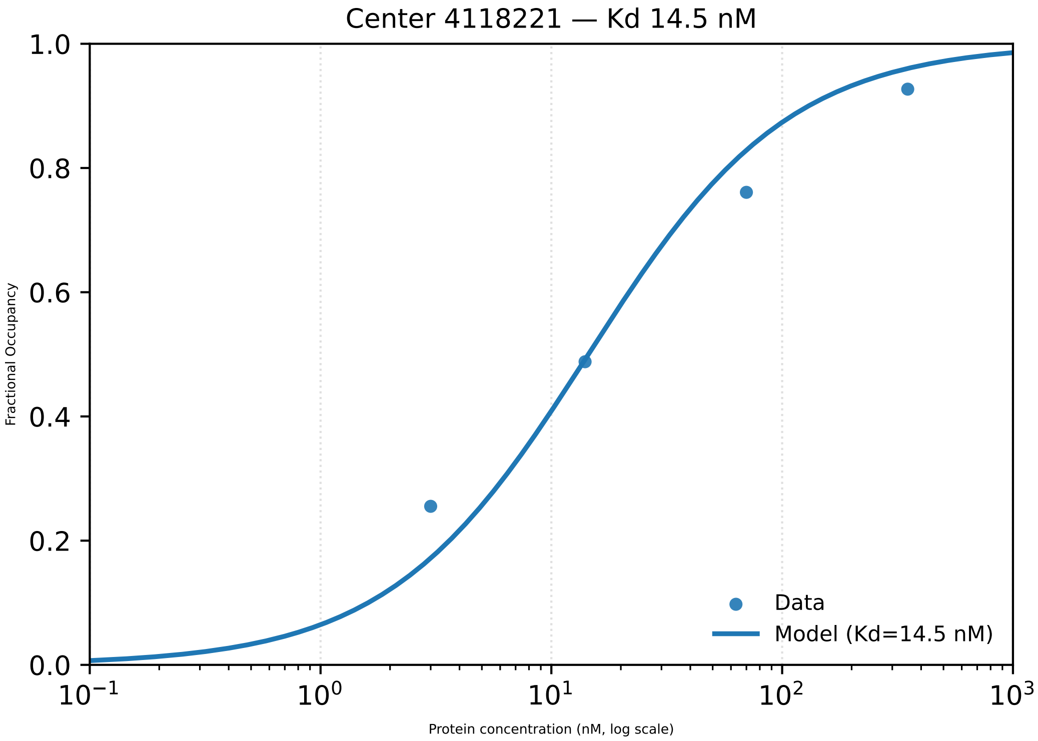

Center 4118221 — Kd 14.5 nM
1.0
0.8
0.6
0.4
0.2
0.0
Fractional Occupancy
$10^{-1}$
$10^{0}$
$10^{1}$
$10^{2}$
$10^{3}$
Protein concentration (nM, log scale)
Data
Model (Kd=14.5 nM)

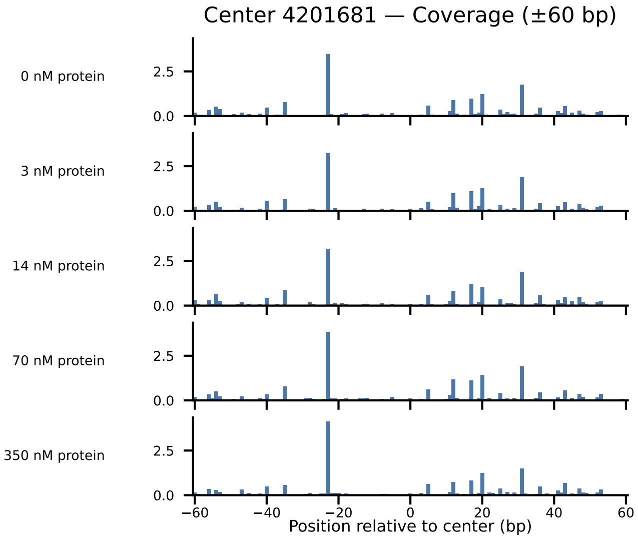
Center 4201681 — Coverage (±60 bp)
0 nM protein
3 nM protein
14 nM protein
70 nM protein
350 nM protein
2.5
0.0
−60
−40
−20
0
20
40
60
Position relative to center (bp)

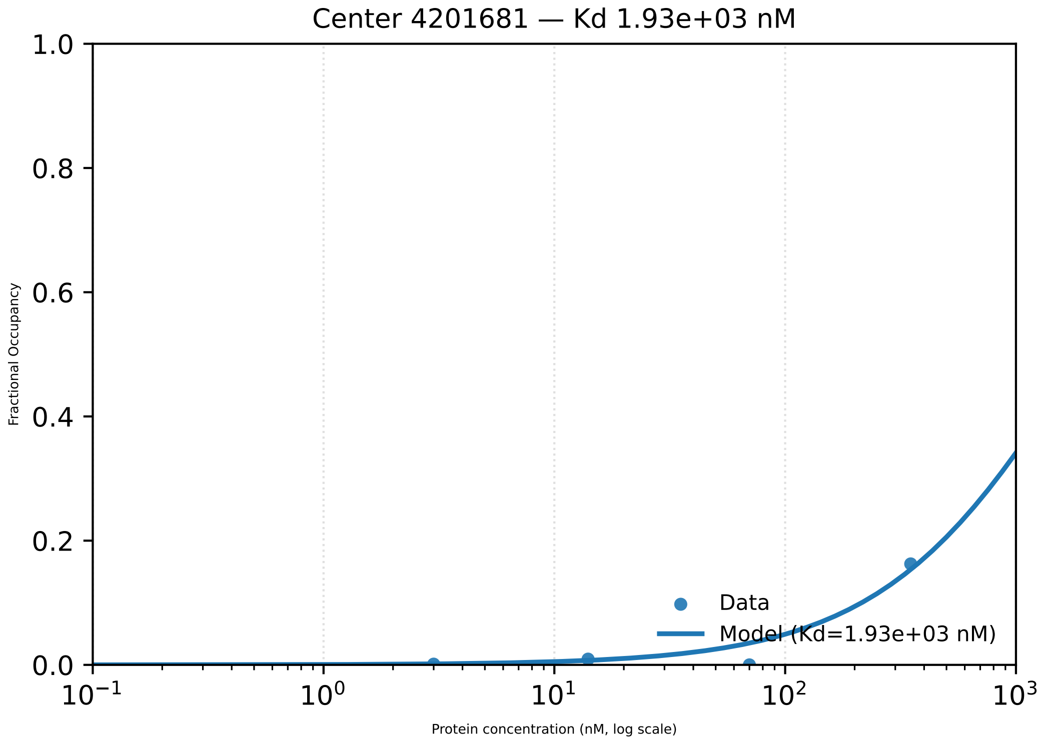

Center 4201681 — Kd 1.93e+03 nM
1.0
0.8
0.6
0.4
0.2
0.0
Fractional Occupancy
Data
Model (Kd=1.93e+03 nM)
$10^{-1}$
$10^{0}$
$10^{1}$
$10^{2}$
$10^{3}$
Protein concentration (nM, log scale)

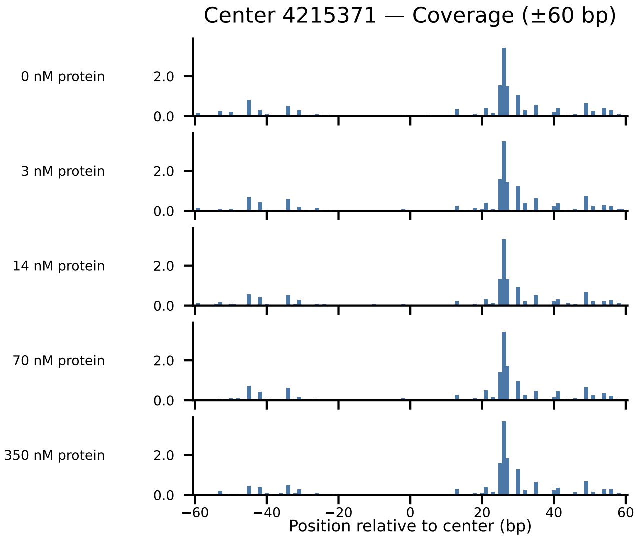

Center 4215371 — Coverage (±60 bp)
0 nM protein
3 nM protein
14 nM protein
70 nM protein
350 nM protein
2.0
0.0
−60
−40
−20
0
20
40
60
Position relative to center (bp)

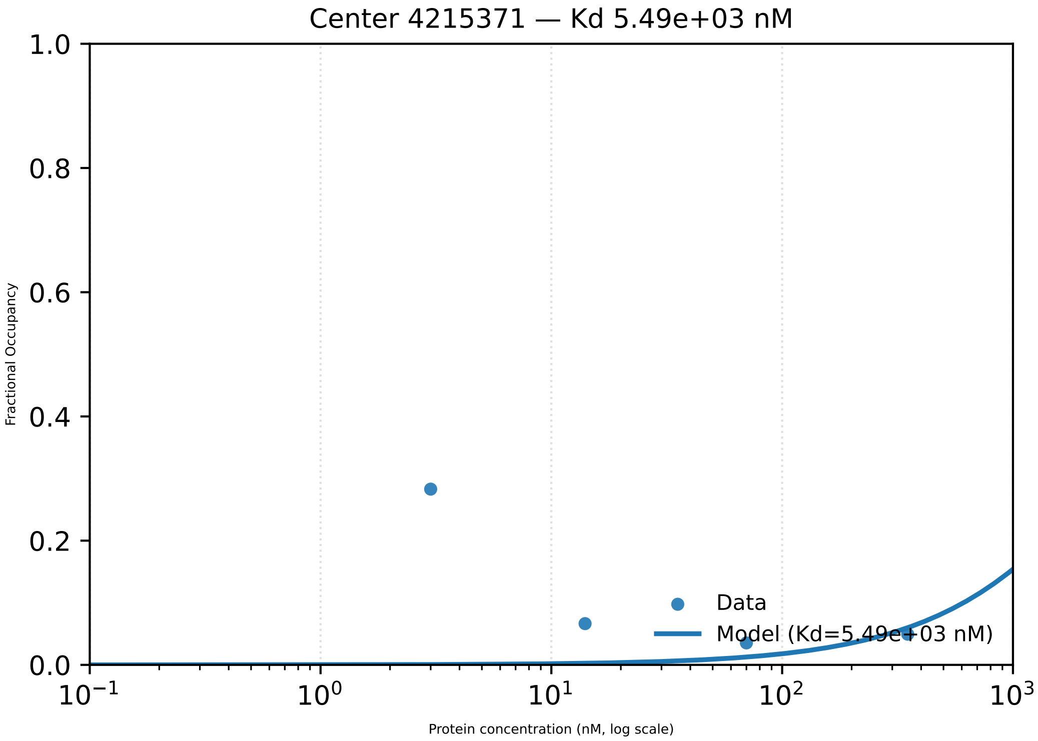

Center 4215371 — Kd 5.49e+03 nM
1.0
0.8
0.6
0.4
0.2
0.0
Fractional Occupancy
Data
Model (Kd=5.49e+03 nM)
$10^{-1}$
$10^{0}$
$10^{1}$
$10^{2}$
$10^{3}$
Protein concentration (nM, log scale)

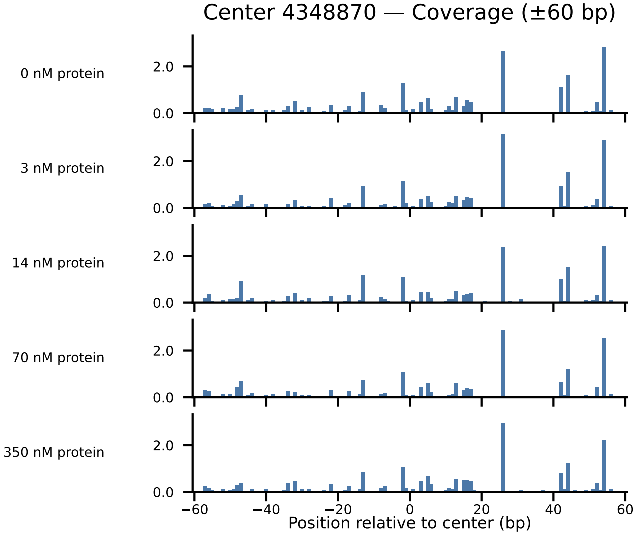
Center 4348870 — Coverage (±60 bp)
0 nM protein
3 nM protein
14 nM protein
70 nM protein
350 nM protein
2.0
0.0
−60
−40
−20
0
20
40
60
Position relative to center (bp)

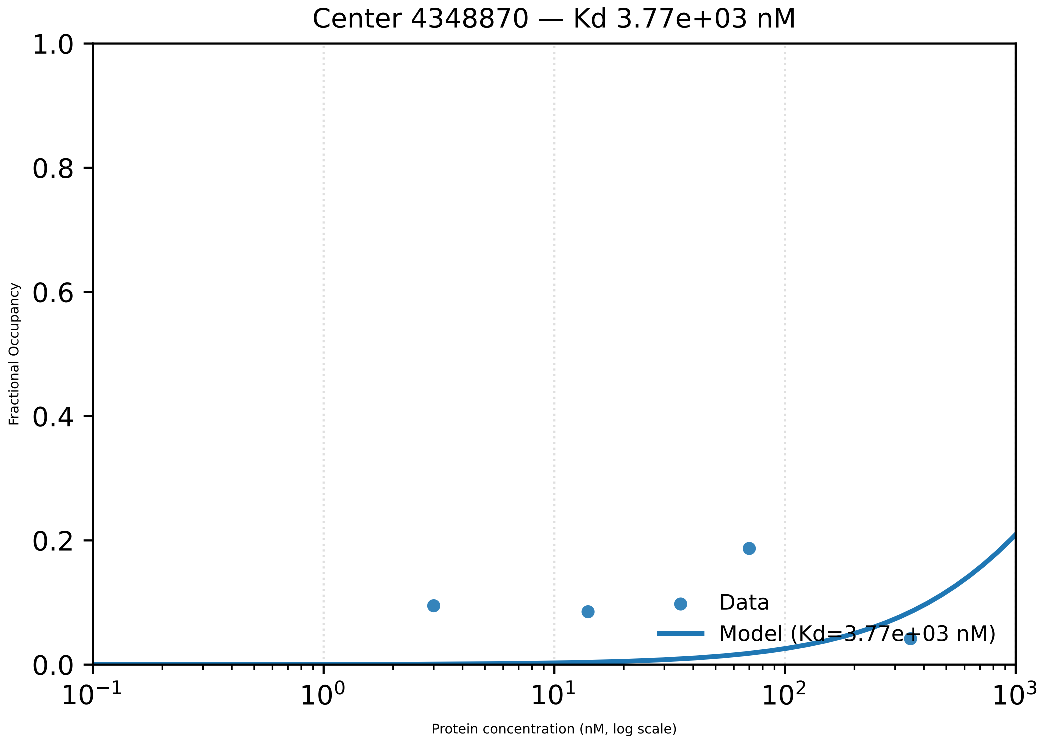

Center 4348870 — Kd 3.77e+03 nM
1.0
0.8
0.6
0.4
0.2
0.0
Fractional Occupancy
$10^{-1}$
$10^{0}$
$10^{1}$
$10^{2}$
$10^{3}$
Protein concentration (nM, log scale)
Data
Model (Kd=3.77e+03 nM)

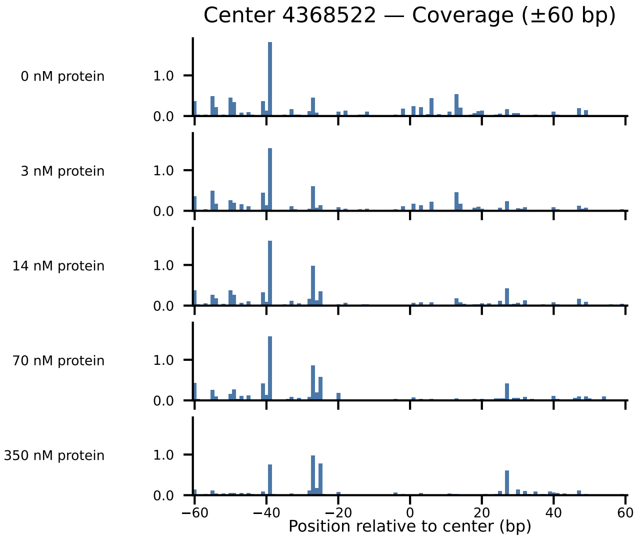

Center 4368522 — Coverage (±60 bp)
0 nM protein
3 nM protein
14 nM protein
70 nM protein
350 nM protein
1.0
0.0
−60
−40
−20
0
20
40
60
Position relative to center (bp)

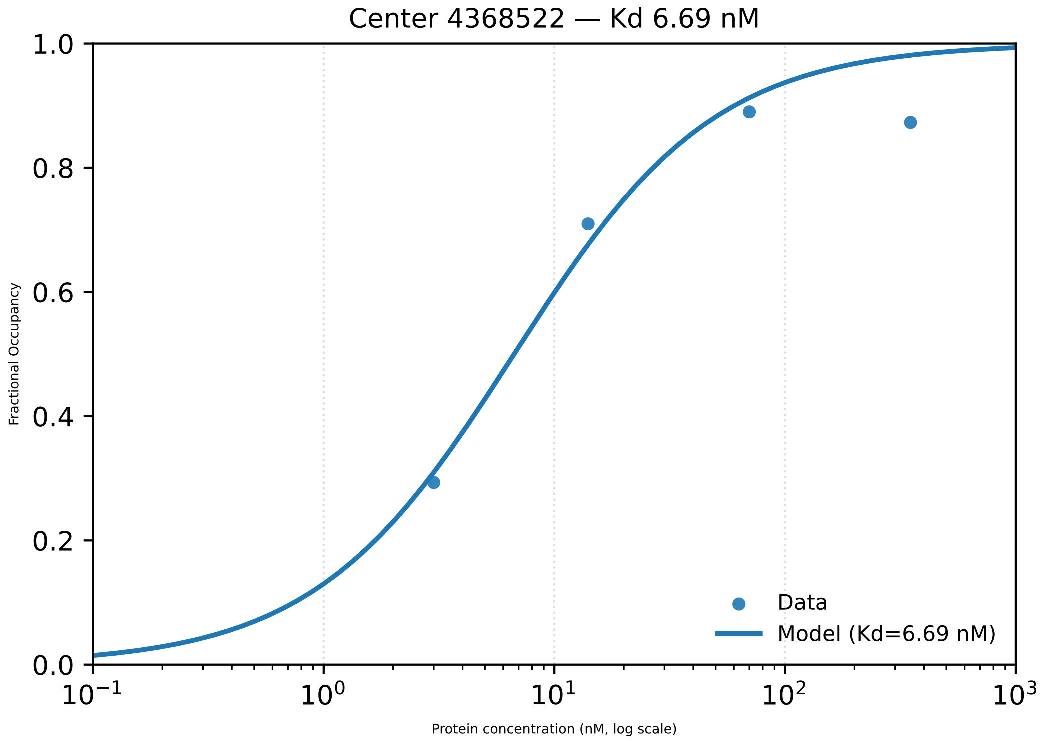
Center 4368522 — Kd 6.69 nM
1.0
0.8
0.6
0.4
0.2
0.0
Fractional Occupancy
$10^{-1}$
$10^{0}$
$10^{1}$
$10^{2}$
$10^{3}$
Protein concentration (nM, log scale)
Data
Model (Kd=6.69 nM)

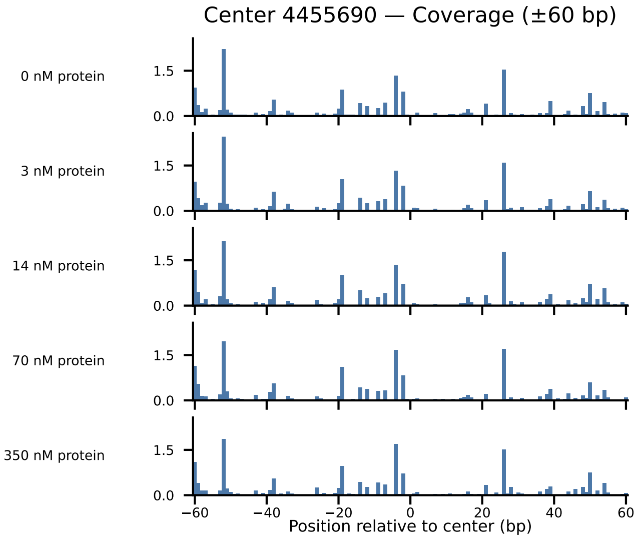

Center 4455690 — Coverage (±60 bp)
0 nM protein
3 nM protein
14 nM protein
70 nM protein
350 nM protein
1.5
0.0
−60
−40
−20
0
20
40
60
Position relative to center (bp)

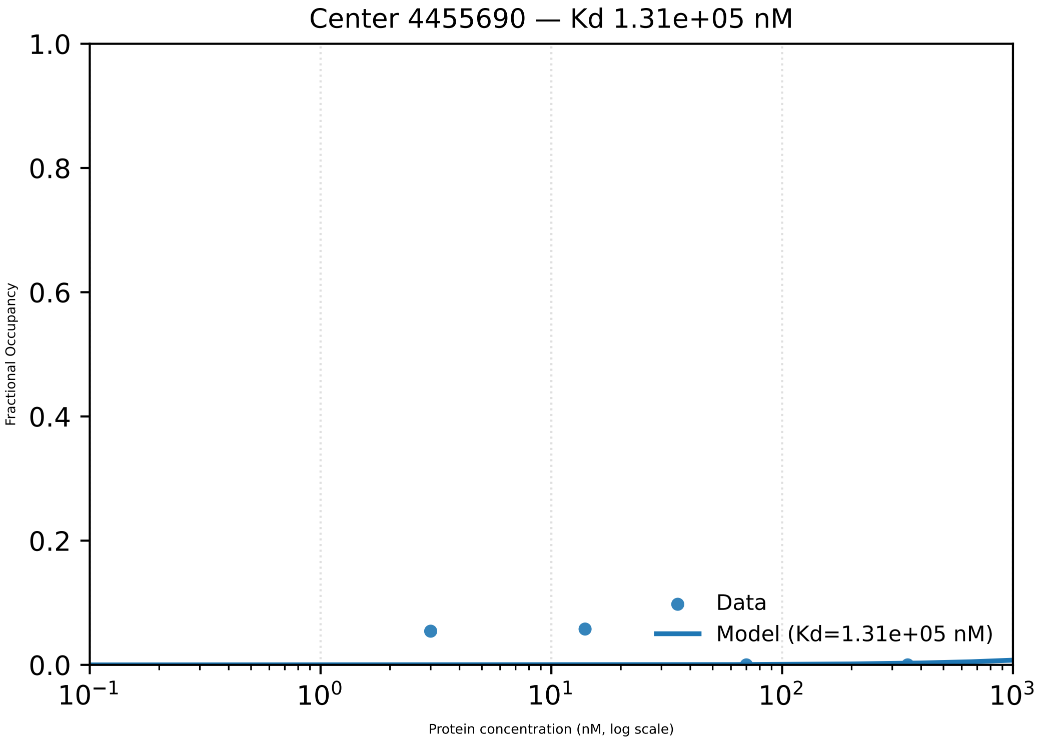

Center 4455690 — Kd 1.31e+05 nM
Fractional Occupancy
1.0
0.8
0.6
0.4
0.2
0.0
Data
Model (Kd=1.31e+05 nM)
$10^{-1}$
$10^{0}$
$10^{1}$
$10^{2}$
$10^{3}$
Protein concentration (nM, log scale)

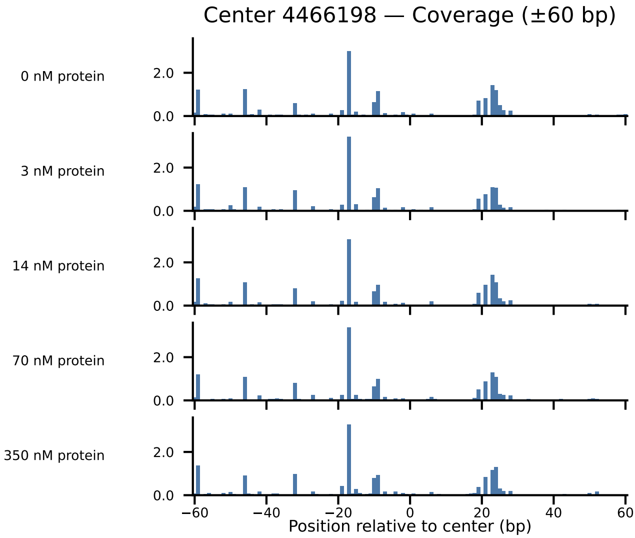

Center 4466198 — Coverage (±60 bp)
0 nM protein
3 nM protein
14 nM protein
70 nM protein
350 nM protein
2.0
0.0
−60
−40
−20
0
20
40
60
Position relative to center (bp)

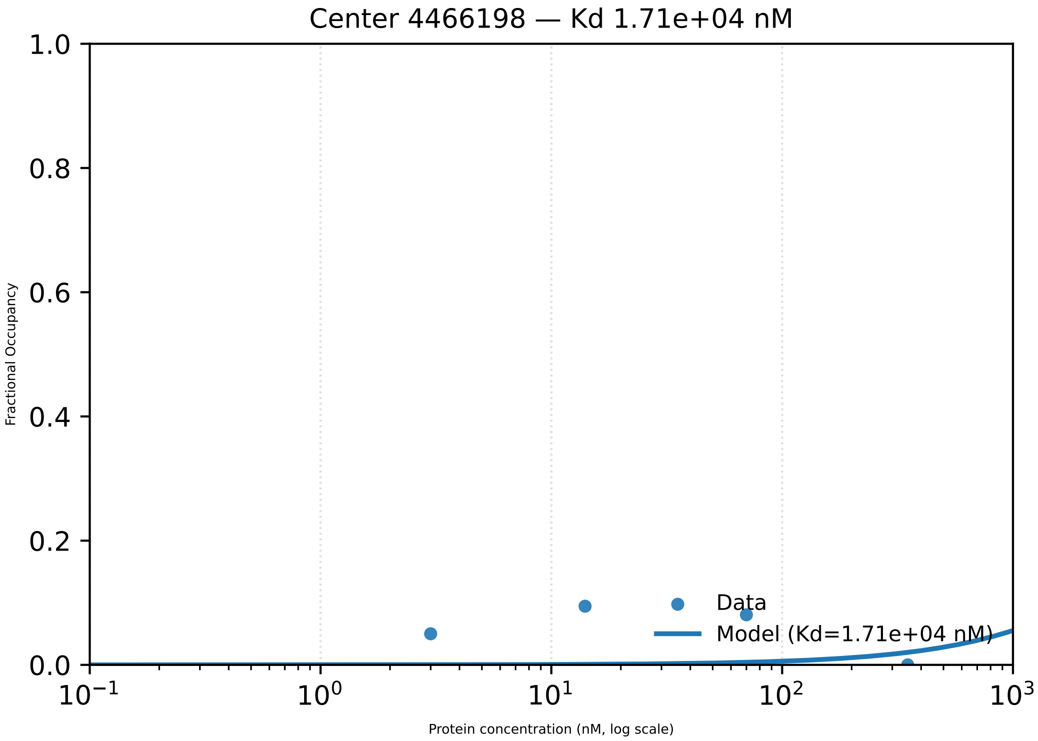
Center 4466198 — Kd 1.71e+04 nM
1.0
0.8
0.6
0.4
0.2
0.0
Fractional Occupancy
$10^{-1}$
$10^{0}$
$10^{1}$
$10^{2}$
$10^{3}$
Protein concentration (nM, log scale)
Data
Model (Kd=1.71e+04 nM)

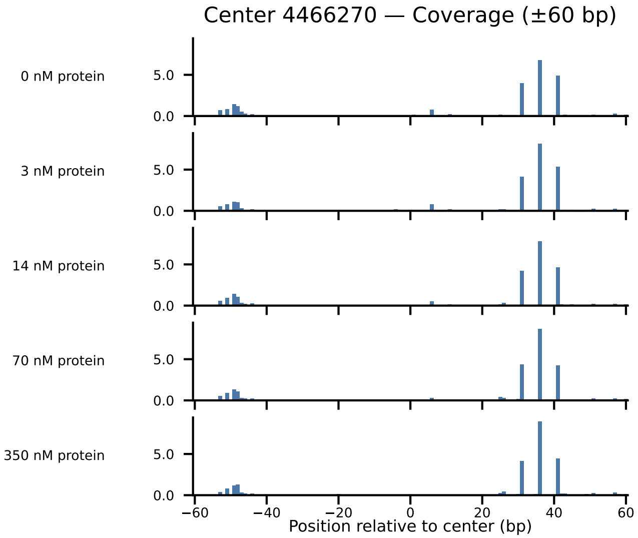

Center 4466270 — Coverage (±60 bp)
0 nM protein
3 nM protein
14 nM protein
70 nM protein
350 nM protein
5.0
0.0
−60
−40
−20
0
20
40
60
Position relative to center (bp)

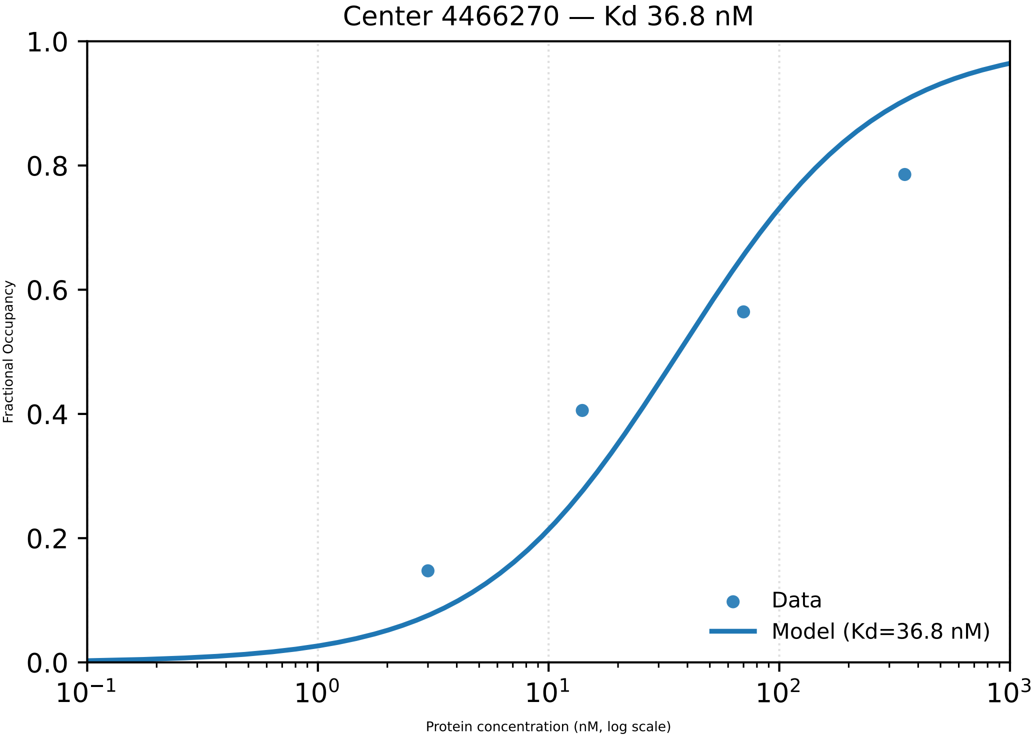
Center 4466270 — Kd 36.8 nM
Fractional Occupancy
1.0
0.8
0.6
0.4
0.2
0.0
$10^{-1}$
$10^{0}$
$10^{1}$
$10^{2}$
$10^{3}$
Protein concentration (nM, log scale)
Data
Model (Kd=36.8 nM)

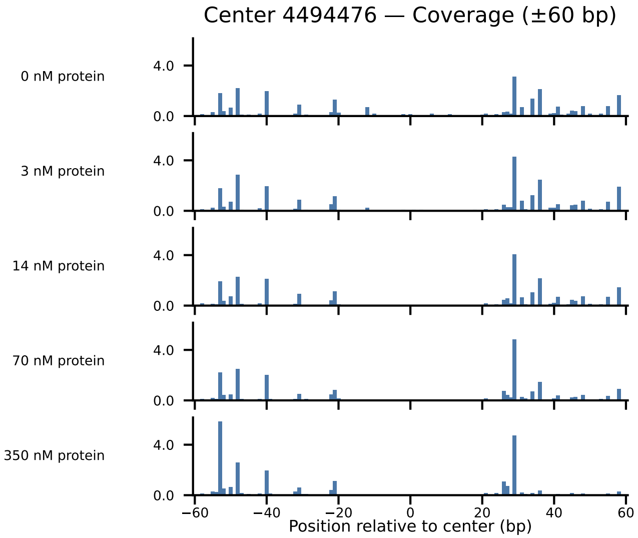

Center 4494476 — Coverage (±60 bp)
0 nM protein
3 nM protein
14 nM protein
70 nM protein
350 nM protein
4.0
0.0
−60
−40
−20
0
20
40
60
Position relative to center (bp)

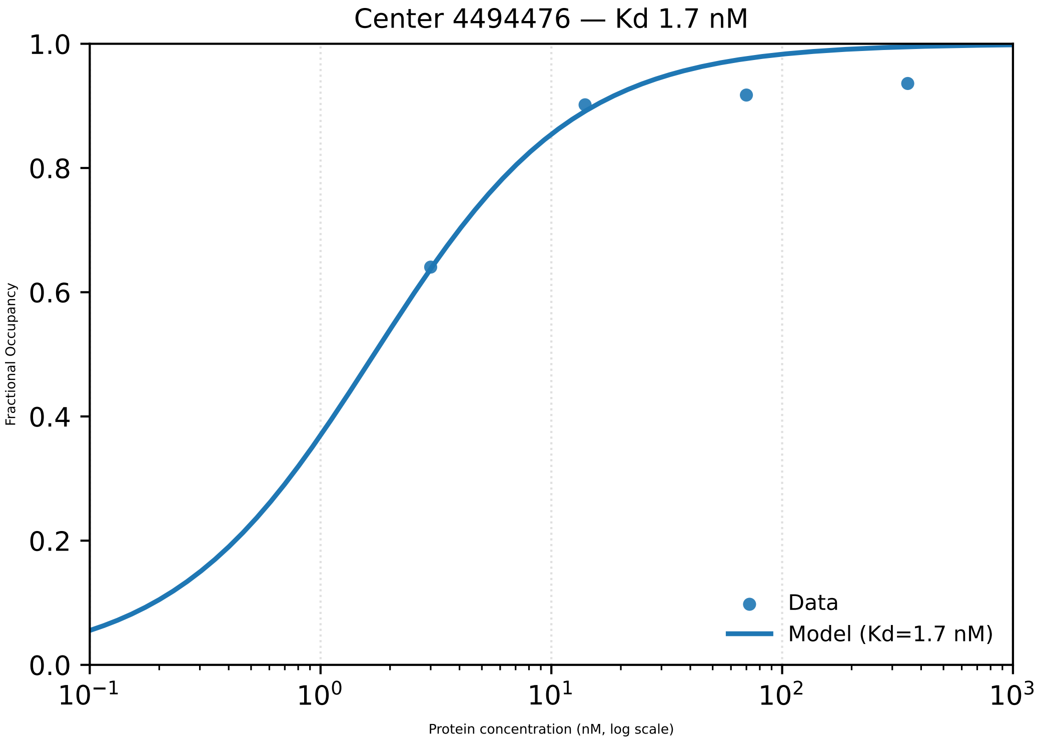

Center 4494476 — Kd 1.7 nM
1.0
0.8
0.6
0.4
0.2
0.0
Fractional Occupancy
$10^{-1}$
$10^{0}$
$10^{1}$
$10^{2}$
$10^{3}$
Protein concentration (nM, log scale)
Data
Model (Kd=1.7 nM)

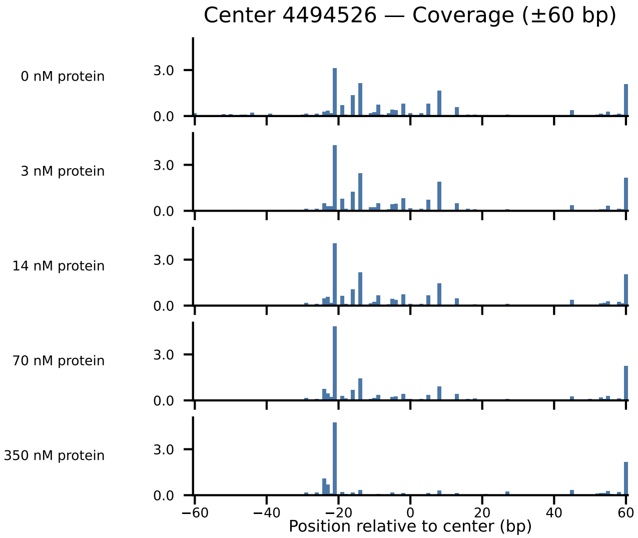

Center 4494526 — Coverage (±60 bp)
0 nM protein
3 nM protein
14 nM protein
70 nM protein
350 nM protein
3.0
0.0
−60
−40
−20
0
20
40
60
Position relative to center (bp)

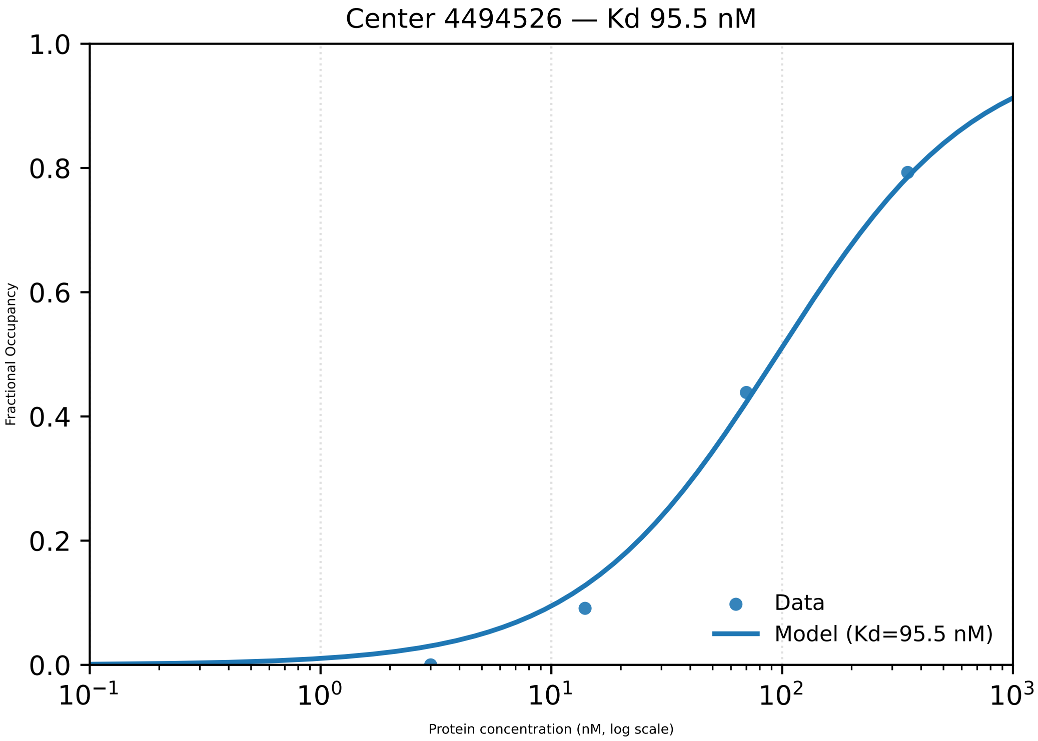

Center 4494526 — Kd 95.5 nM
Fractional Occupancy
1.0
0.8
0.6
0.4
0.2
0.0
$10^{-1}$
$10^{0}$
$10^{1}$
$10^{2}$
$10^{3}$
Protein concentration (nM, log scale)
Data
Model (Kd=95.5 nM)

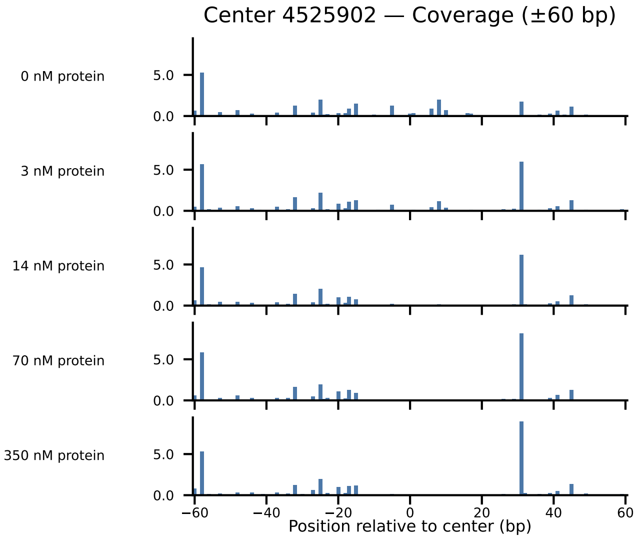

Center 4525902 — Coverage (±60 bp)
0 nM protein
3 nM protein
14 nM protein
70 nM protein
350 nM protein
5.0
0.0
−60
−40
−20
0
20
40
60
Position relative to center (bp)

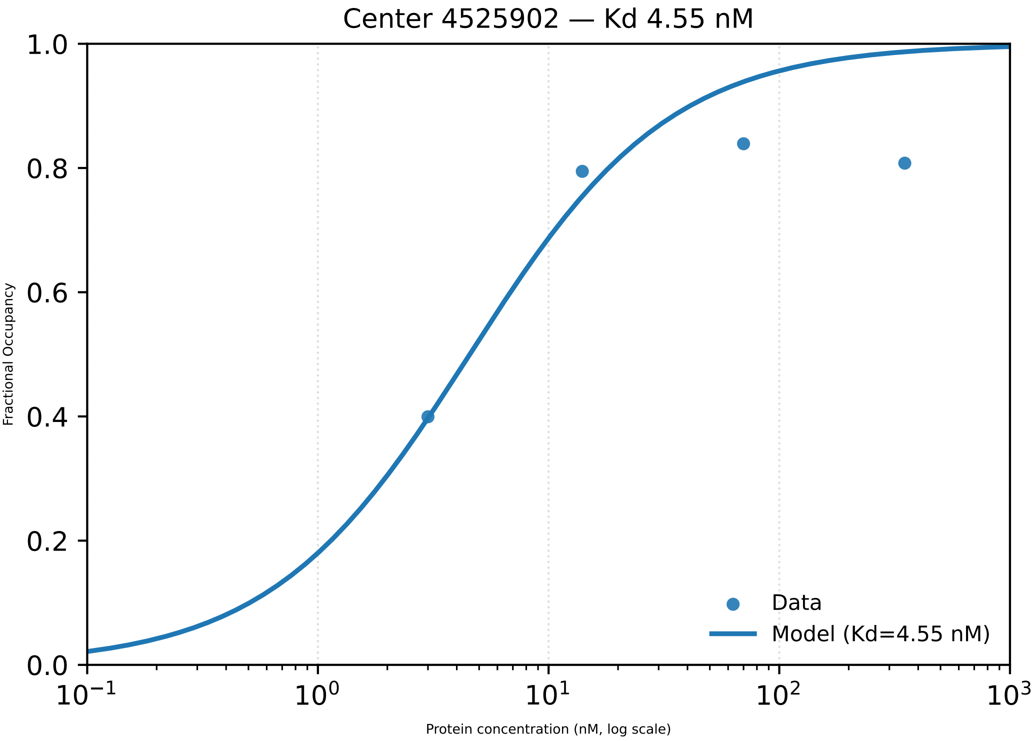

Center 4525902 — Kd 4.55 nM
Fractional Occupancy
Protein concentration (nM, log scale)
0.0
0.2
0.4
0.6
0.8
1.0
$10^{-1}$
$10^{0}$
$10^{1}$
$10^{2}$
$10^{3}$
Data
Model (Kd=4.55 nM)

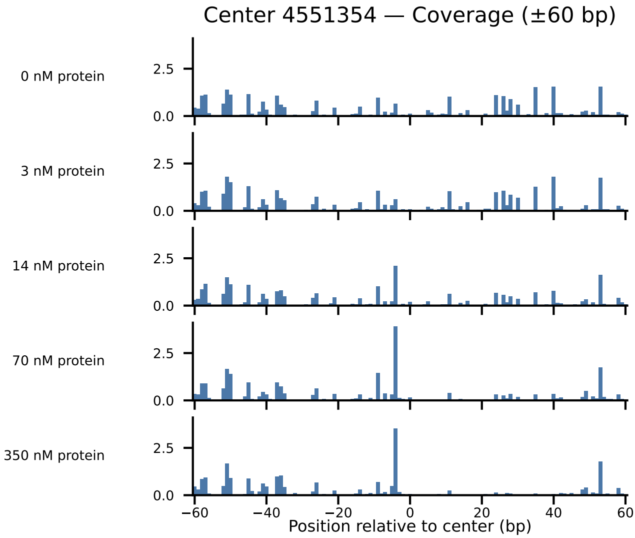

Center 4551354 — Coverage (±60 bp)
0 nM protein
3 nM protein
14 nM protein
70 nM protein
350 nM protein
2.5
0.0
−60
−40
−20
0
20
40
60
Position relative to center (bp)

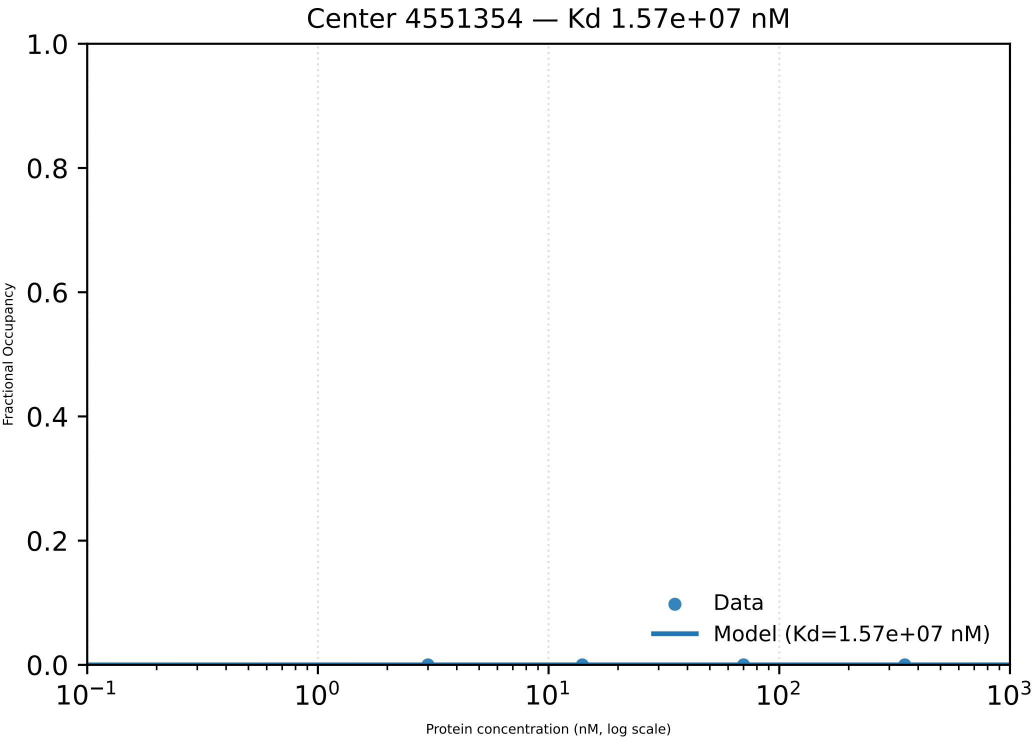

Center 4551354 — Kd 1.57e+07 nM
1.0
0.8
0.6
0.4
0.2
0.0
Fractional Occupancy
Data
Model (Kd=1.57e+07 nM)
$10^{-1}$
$10^{0}$
$10^{1}$
$10^{2}$
$10^{3}$
Protein concentration (nM, log scale)

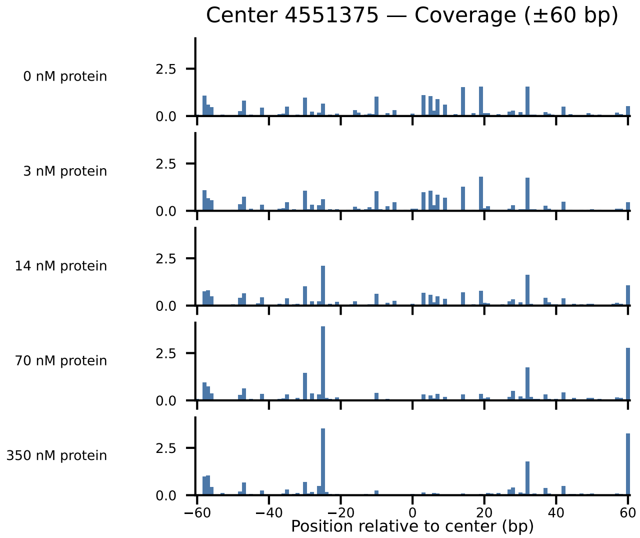
Center 4551375 — Coverage (±60 bp)
0 nM protein
3 nM protein
14 nM protein
70 nM protein
350 nM protein
2.5
0.0
−60
−40
−20
0
20
40
60
Position relative to center (bp)

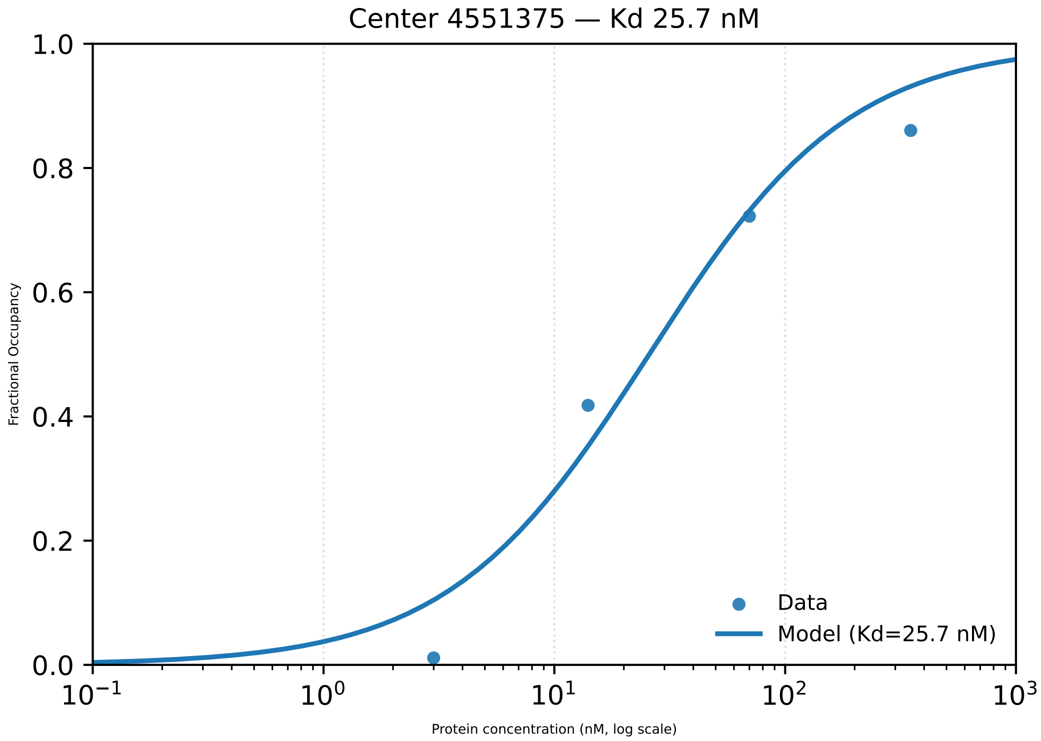

Center 4551375 — Kd 25.7 nM
1.0
0.8
0.6
0.4
0.2
0.0
Fractional Occupancy
Data
Model (Kd=25.7 nM)
$10^{-1}$
$10^{0}$
$10^{1}$
$10^{2}$
$10^{3}$
Protein concentration (nM, log scale)

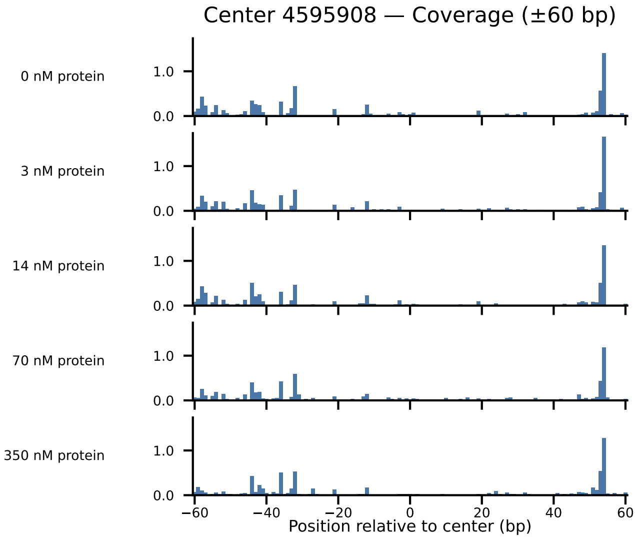

Center 4595908 — Coverage (±60 bp)
0 nM protein
3 nM protein
14 nM protein
70 nM protein
350 nM protein
1.0
0.0
−60
−40
−20
0
20
40
60
Position relative to center (bp)

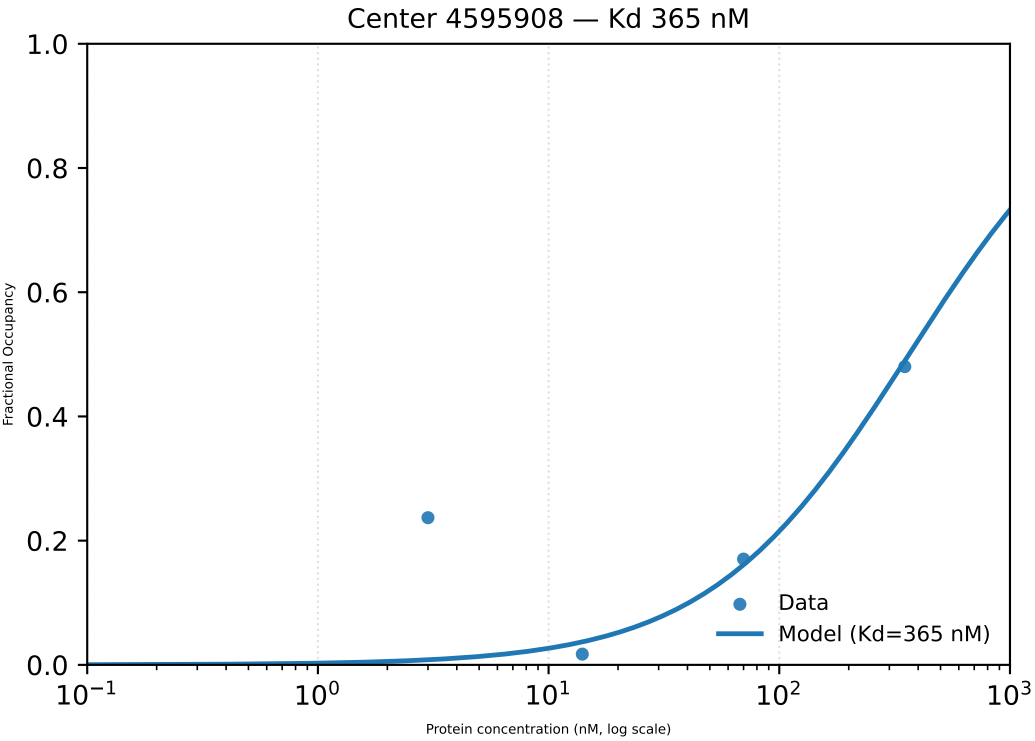

Center 4595908 — Kd 365 nM
1.0
0.8
0.6
0.4
0.2
0.0
Fractional Occupancy
10−1
10^0
10^1
10^2
10^3
Protein concentration (nM, log scale)
Data
Model (Kd=365 nM)

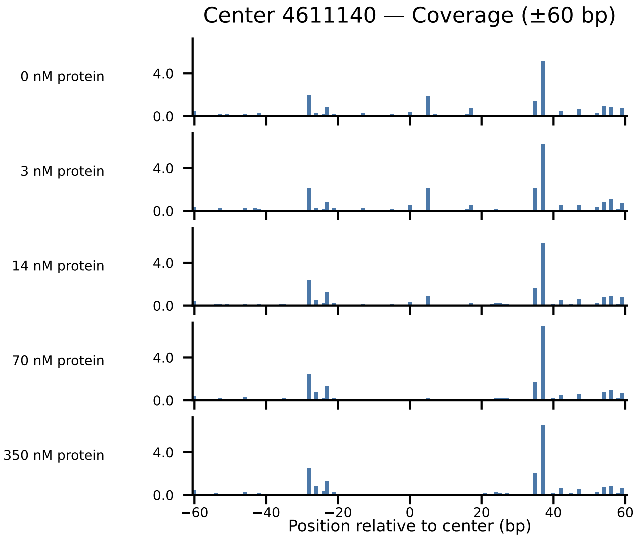

Center 4611140 — Coverage (±60 bp)
0 nM protein
3 nM protein
14 nM protein
70 nM protein
350 nM protein
4.0
0.0
−60
−40
−20
0
20
40
60
Position relative to center (bp)

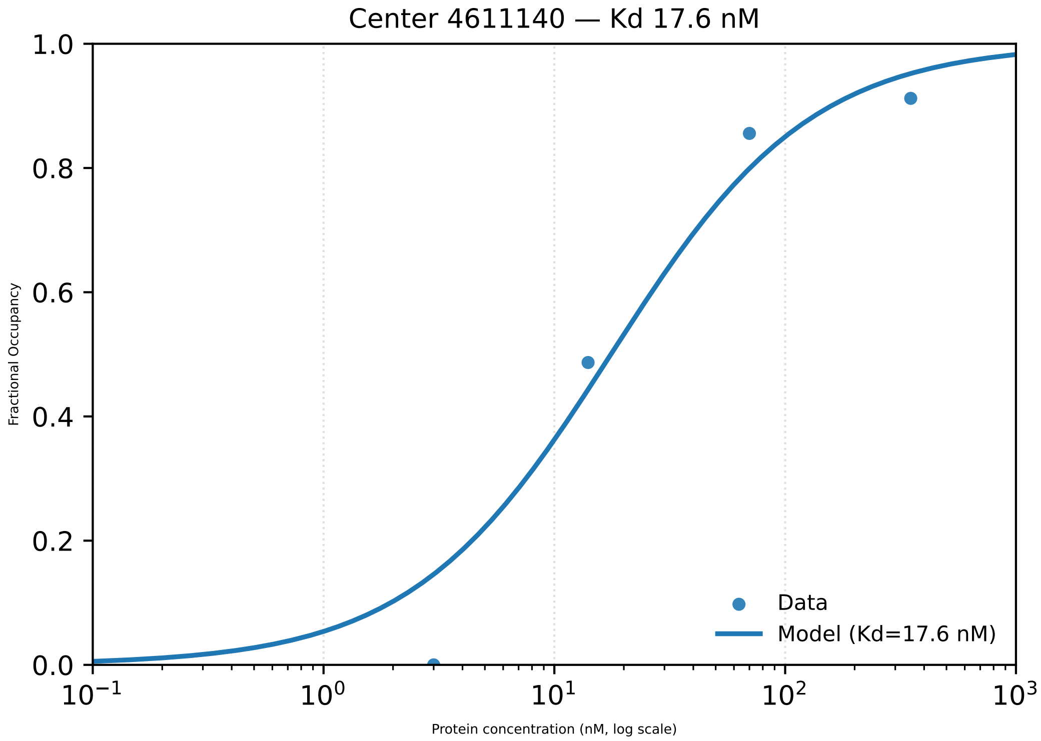

Center 4611140 — Kd 17.6 nM
1.0
0.8
0.6
0.4
0.2
0.0
Fractional Occupancy
$10^{-1}$
$10^{0}$
$10^{1}$
$10^{2}$
$10^{3}$
Protein concentration (nM, log scale)
Data
Model (Kd=17.6 nM)

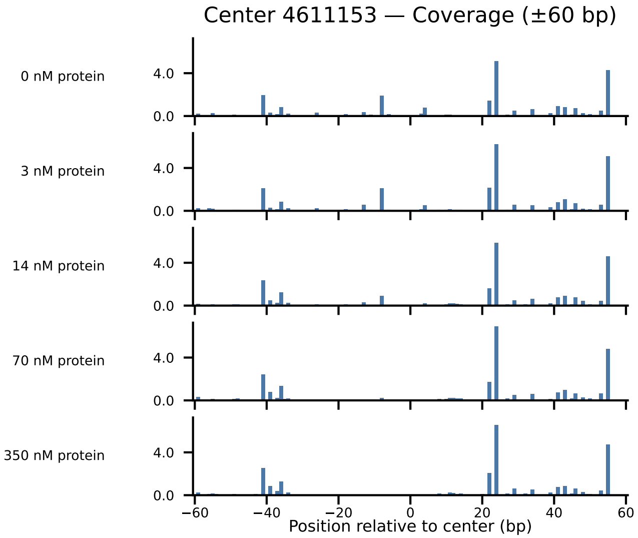

Center 4611153 — Coverage (±60 bp)
0 nM protein
3 nM protein
14 nM protein
70 nM protein
350 nM protein
4.0
0.0
−60
−40
−20
0
20
40
60
Position relative to center (bp)

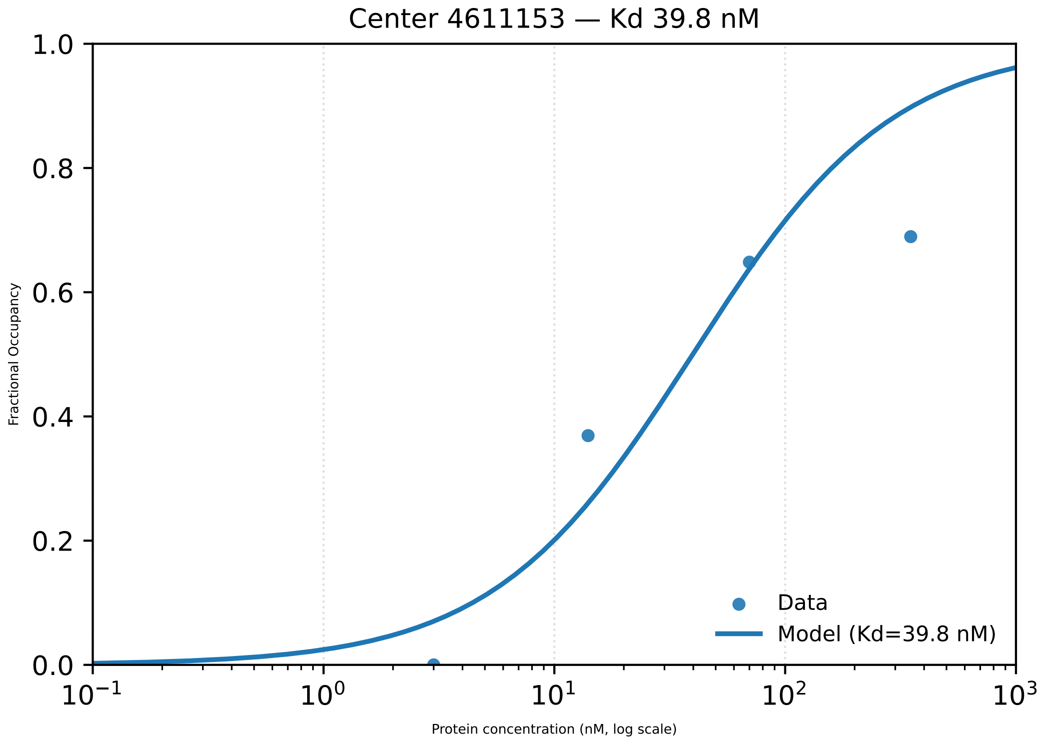

Center 4611153 — Kd 39.8 nM
1.0
0.8
0.6
0.4
0.2
0.0
Fractional Occupancy
$10^{-1}$
$10^{0}$
$10^{1}$
$10^{2}$
$10^{3}$
Protein concentration (nM, log scale)
Data
Model (Kd=39.8 nM)

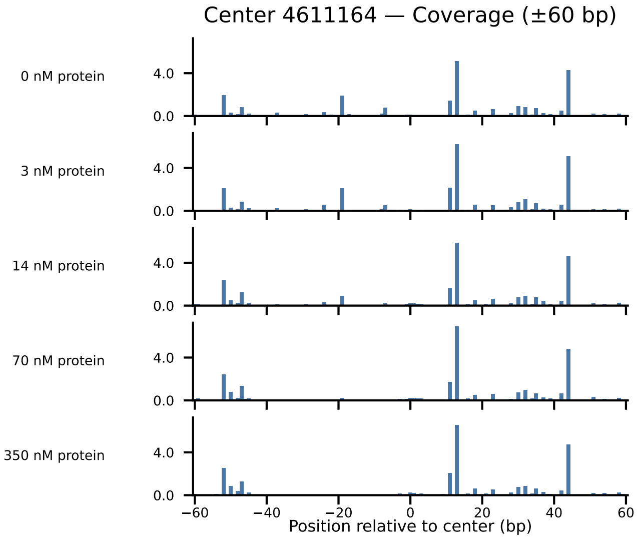

Center 4611164 — Coverage (±60 bp)
0 nM protein
3 nM protein
14 nM protein
70 nM protein
350 nM protein
4.0
0.0
−60
−40
−20
0
20
40
60
Position relative to center (bp)

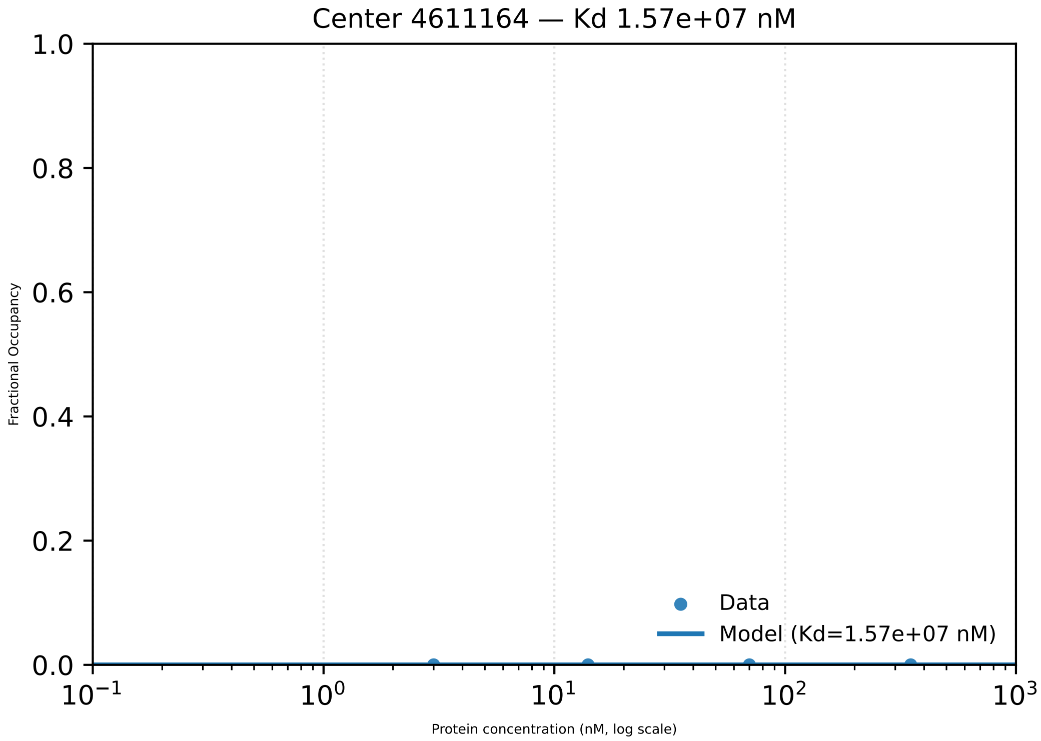

Center 4611164 — Kd 1.57e+07 nM
1.0
0.8
0.6
0.4
0.2
0.0
Fractional Occupancy
Data
Model (Kd=1.57e+07 nM)
$10^{-1}$
$10^{0}$
$10^{1}$
$10^{2}$
$10^{3}$
Protein concentration (nM, log scale)
