## Supplementary material for "Footprint-seq: a simple method to quantitatively map *in vitro* protein-DNA interactions on a genome-wide scale at high spatial resolution": Figure S8

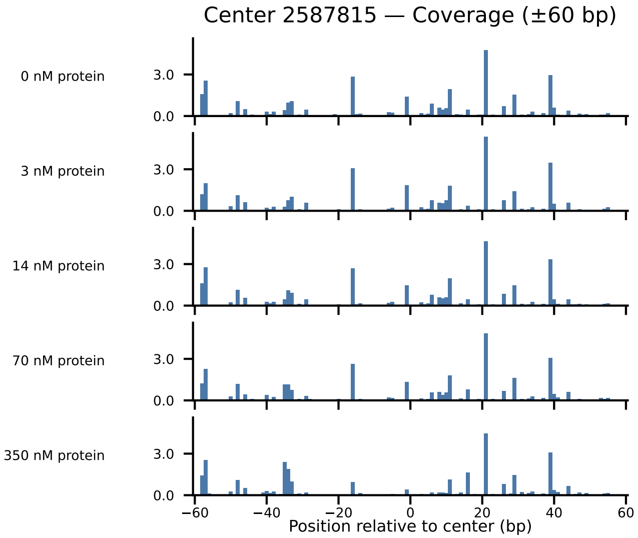

Center 2587815 — Coverage (±60 bp)
0 nM protein
3 nM protein
14 nM protein
70 nM protein
350 nM protein
3.0
0.0
−60
−40
−20
0
20
40
60
Position relative to center (bp)

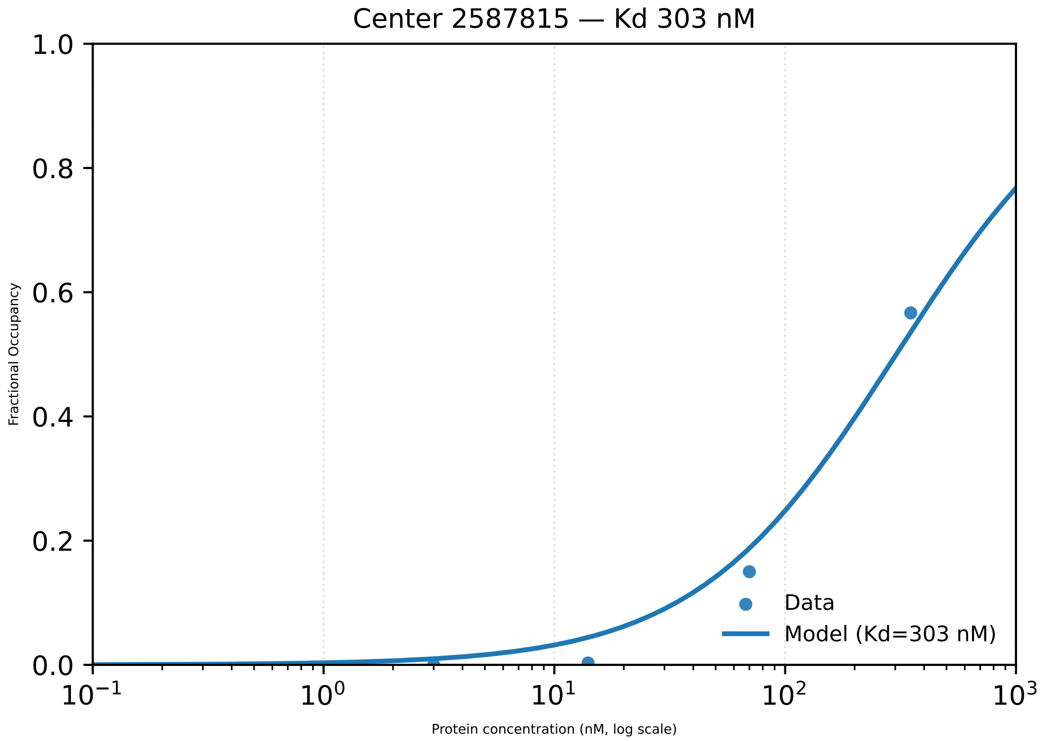

Center 2587815 — Kd 303 nM
1.0
0.8
0.6
0.4
0.2
0.0
Fractional Occupancy
$10^{-1}$
$10^{0}$
$10^{1}$
$10^{2}$
$10^{3}$
Protein concentration (nM, log scale)
Data
Model (Kd=303 nM)

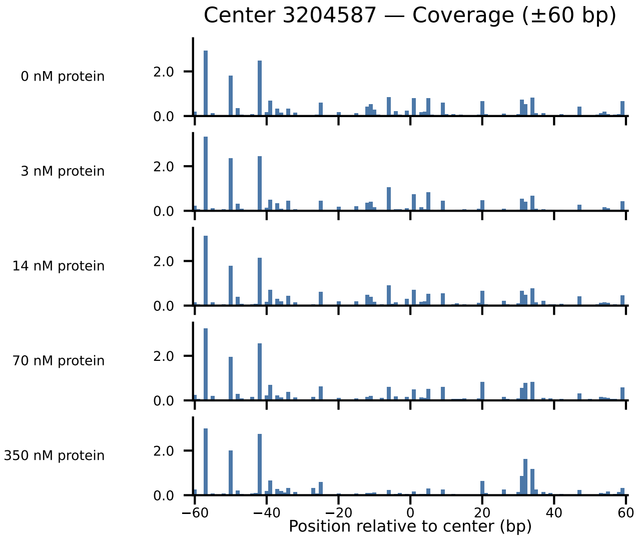

Center 3204587 — Coverage (±60 bp)
0 nM protein
3 nM protein
14 nM protein
70 nM protein
350 nM protein
2.0
0.0
−60
−40
−20
0
20
40
60
Position relative to center (bp)

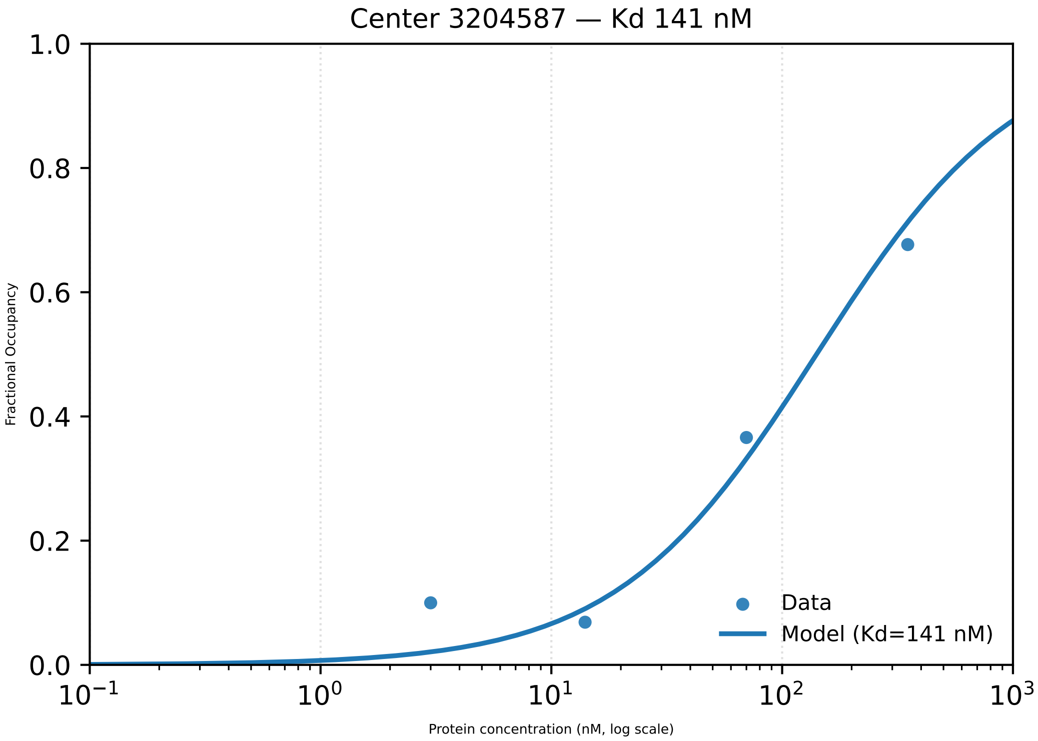

Center 3204587 — Kd 141 nM
Fractional Occupancy
1.0
0.8
0.6
0.4
0.2
0.0
$10^{-1}$
$10^{0}$
$10^{1}$
$10^{2}$
$10^{3}$
Protein concentration (nM, log scale)
Data
Model (Kd=141 nM)

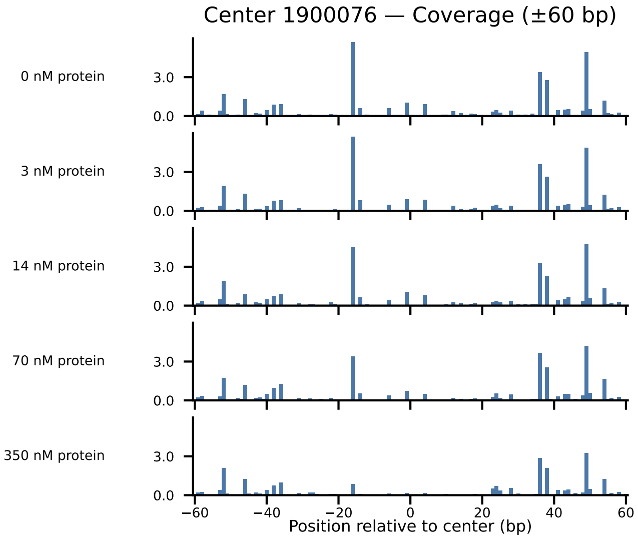
Center 1900076 — Coverage (±60 bp)
0 nM protein
3 nM protein
14 nM protein
70 nM protein
350 nM protein
3.0
0.0
−60
−40
−20
0
20
40
60
Position relative to center (bp)

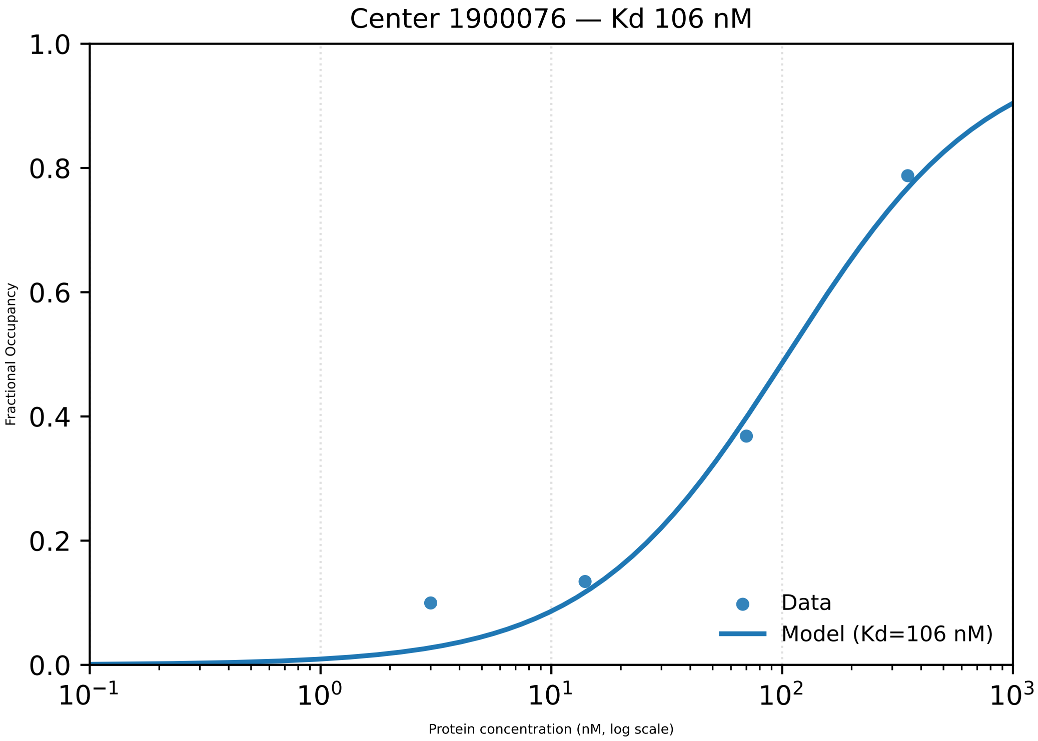

Center 1900076 — Kd 106 nM
Fractional Occupancy
1.0
0.8
0.6
0.4
0.2
0.0
$10^{-1}$
$10^{0}$
$10^{1}$
$10^{2}$
$10^{3}$
Protein concentration (nM, log scale)
Data
Model (Kd=106 nM)

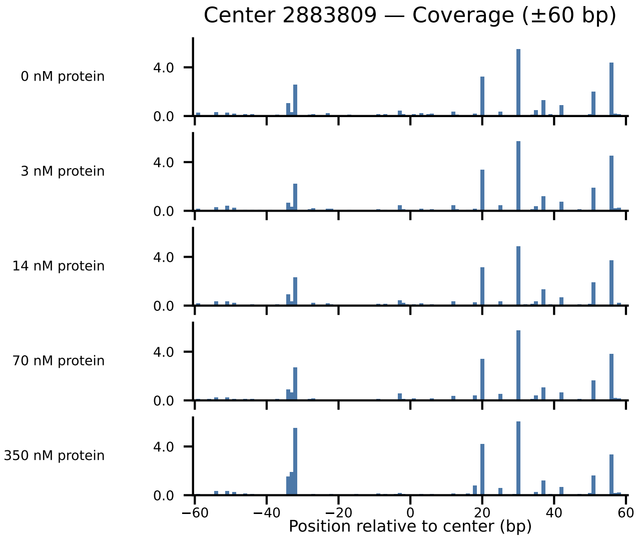
Center 2883809 — Coverage (±60 bp)
0 nM protein
3 nM protein
14 nM protein
70 nM protein
350 nM protein
4.0
0.0
−60
−40
−20
0
20
40
60
Position relative to center (bp)

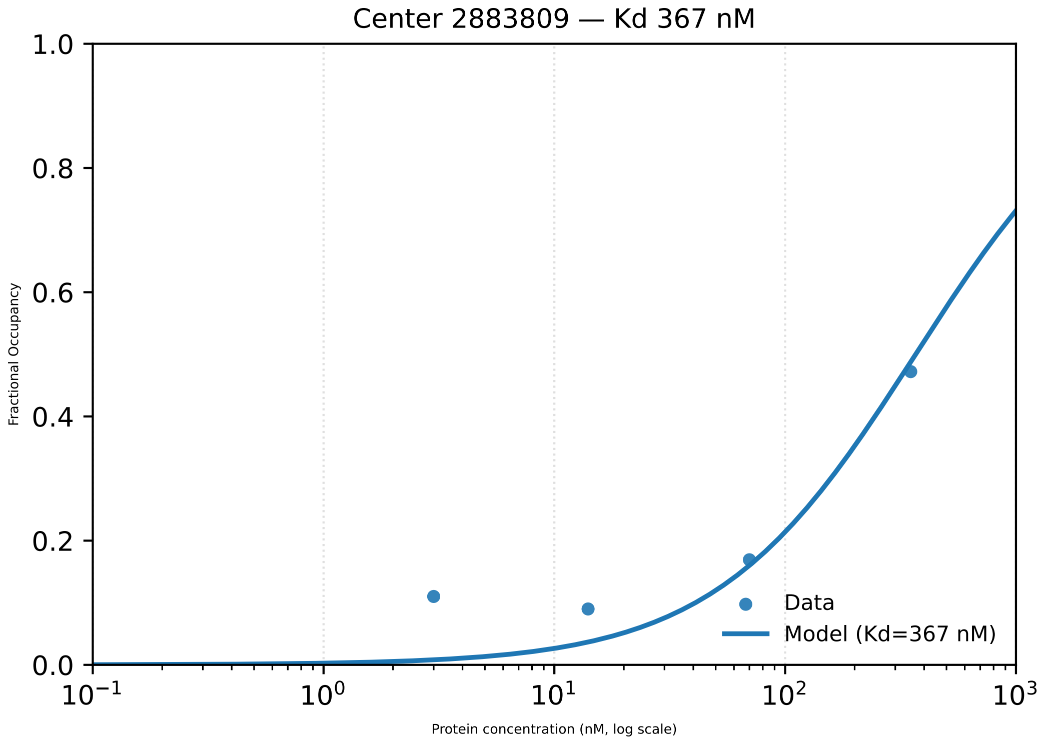
Center 2883809 — Kd 367 nM
Fractional Occupancy
1.0
0.8
0.6
0.4
0.2
0.0
$10^{-1}$
$10^{0}$
$10^{1}$
$10^{2}$
$10^{3}$
Protein concentration (nM, log scale)
Data
Model (Kd=367 nM)

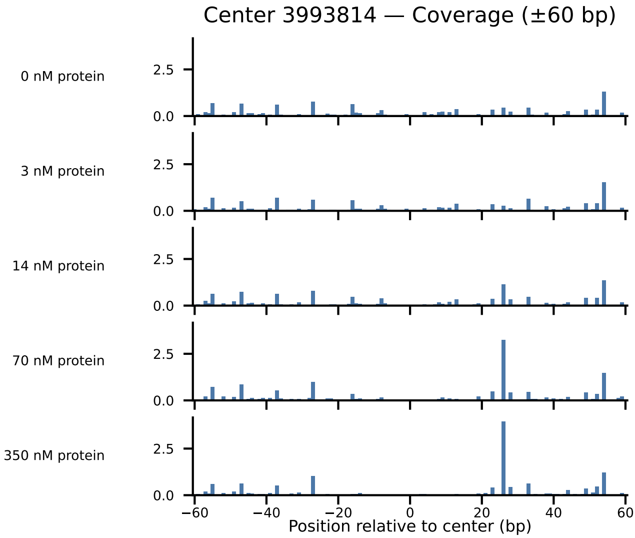

Center 3993814 — Coverage (±60 bp)
0 nM protein
3 nM protein
14 nM protein
70 nM protein
350 nM protein
2.5
0.0
−60
−40
−20
0
20
40
60
Position relative to center (bp)

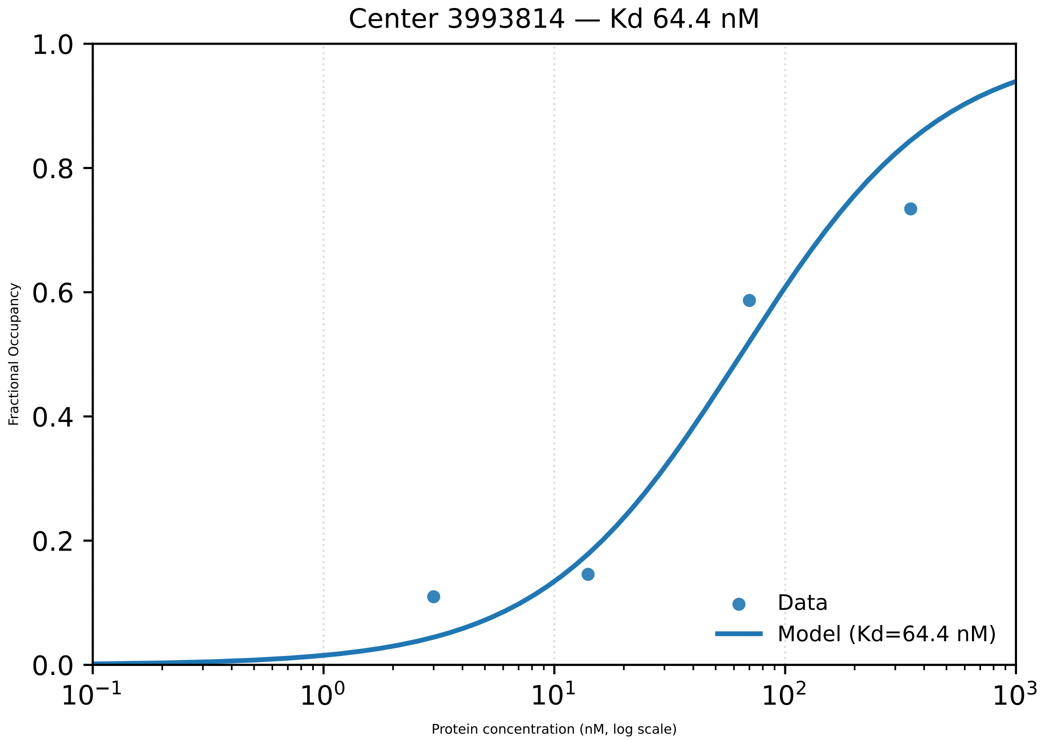

Center 3993814 — Kd 64.4 nM
Fractional Occupancy
1.0
0.8
0.6
0.4
0.2
0.0
$10^{-1}$
$10^{0}$
$10^{1}$
$10^{2}$
$10^{3}$
Protein concentration (nM, log scale)
Data
Model (Kd=64.4 nM)

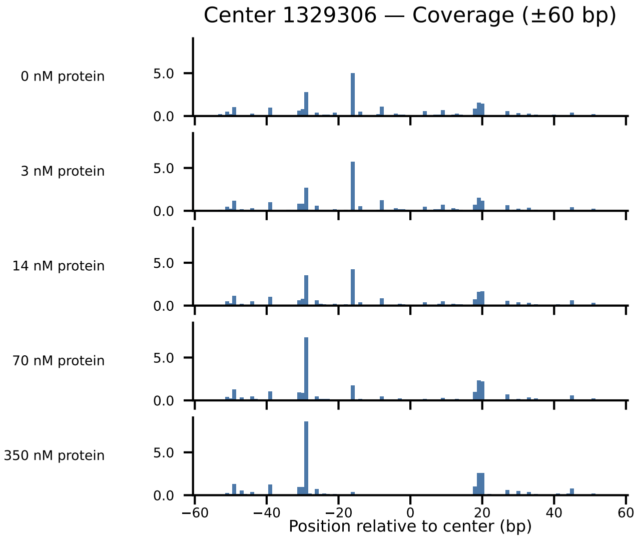

Center 1329306 — Coverage (±60 bp)
0 nM protein
3 nM protein
14 nM protein
70 nM protein
350 nM protein
5.0
0.0
−60
−40
−20
0
20
40
60
Position relative to center (bp)

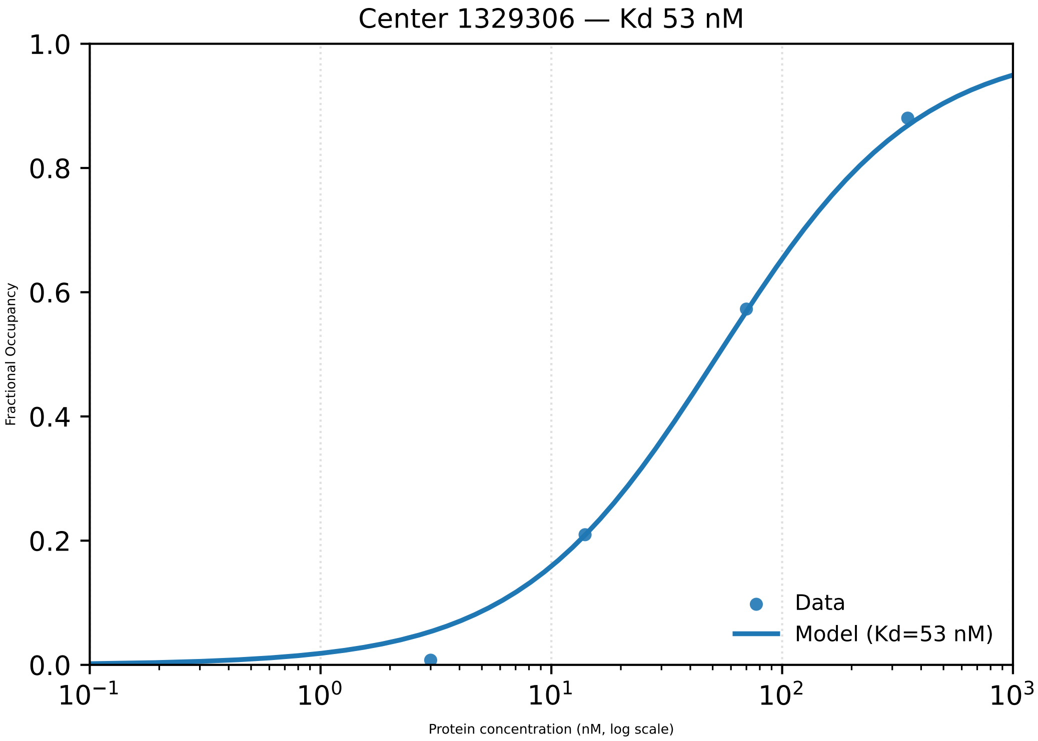
Center 1329306 — Kd 53 nM
Fractional Occupancy
1.0
0.8
0.6
0.4
0.2
0.0
$10^{-1}$
$10^{0}$
$10^{1}$
$10^{2}$
$10^{3}$
Protein concentration (nM, log scale)
Data
Model (Kd=53 nM)

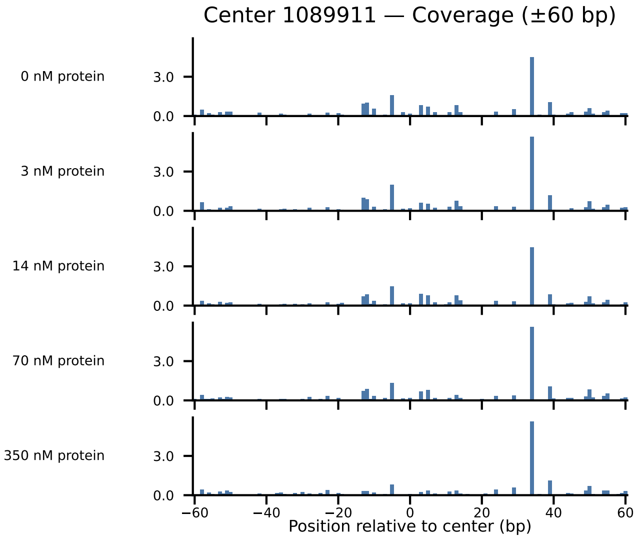

Center 1089911 — Coverage (±60 bp)
0 nM protein
3 nM protein
14 nM protein
70 nM protein
350 nM protein
3.0
0.0
−60
−40
−20
0
20
40
60
Position relative to center (bp)

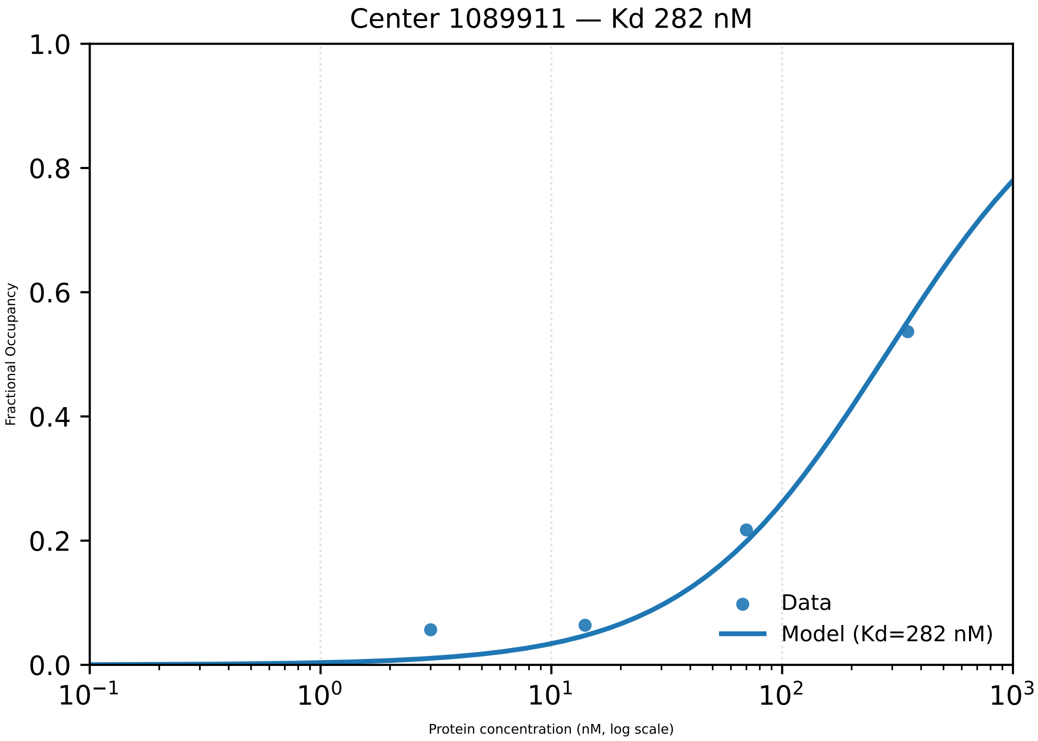

Center 1089911 — Kd 282 nM
Fractional Occupancy
1.0
0.8
0.6
0.4
0.2
0.0
$10^{-1}$
$10^{0}$
$10^{1}$
$10^{2}$
$10^{3}$
Protein concentration (nM, log scale)
Data
Model (Kd=282 nM)

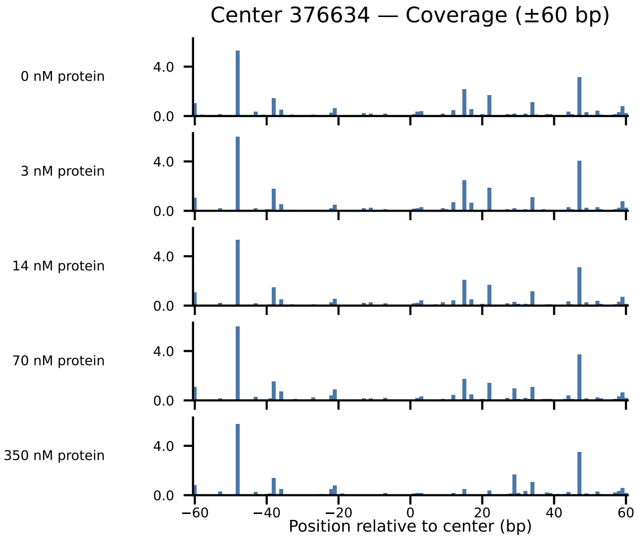
Center 376634 — Coverage (±60 bp)
0 nM protein
3 nM protein
14 nM protein
70 nM protein
350 nM protein
4.0
0.0
−60
−40
−20
0
20
40
60
Position relative to center (bp)

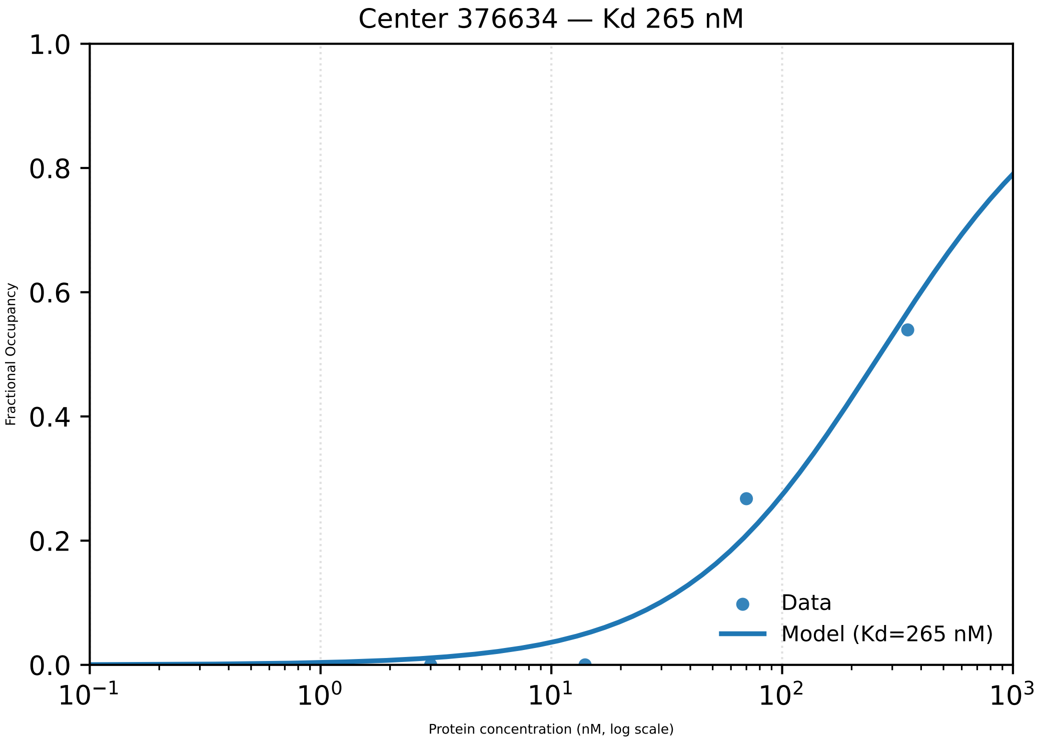

Center 376634 — Kd 265 nM
1.0
0.8
0.6
0.4
0.2
0.0
Fractional Occupancy
$10^{-1}$
$10^{0}$
$10^{1}$
$10^{2}$
$10^{3}$
Protein concentration (nM, log scale)
Data
Model (Kd=265 nM)

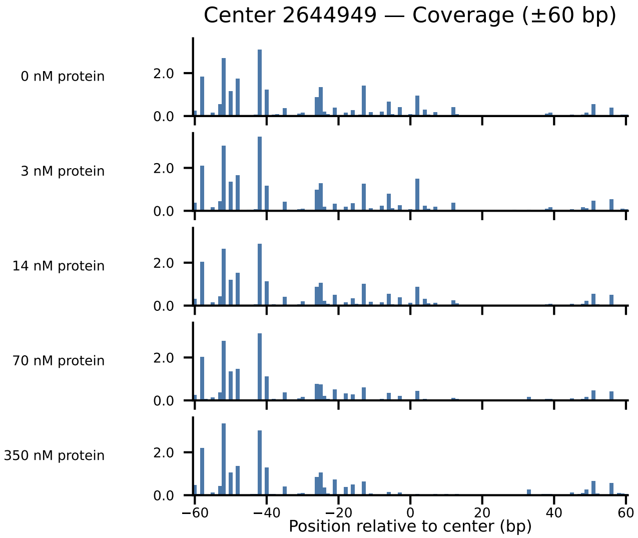

Center 2644949 — Coverage (±60 bp)
0 nM protein
3 nM protein
14 nM protein
70 nM protein
350 nM protein
2.0
0.0
−60
−40
−20
0
20
40
60
Position relative to center (bp)

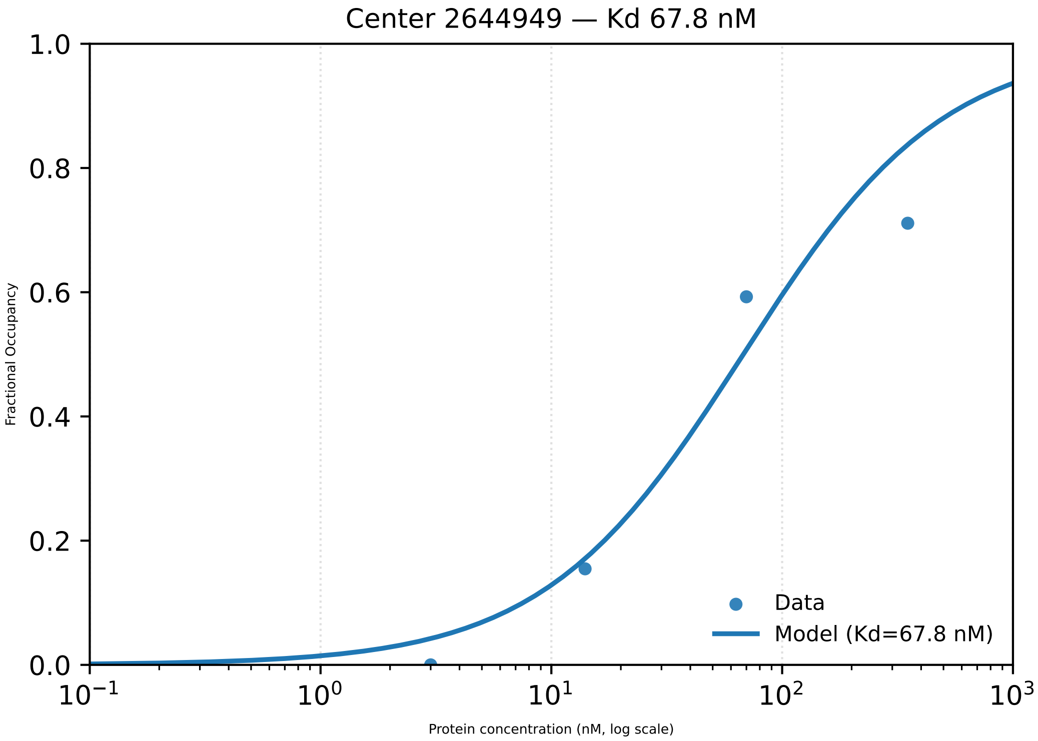
Center 2644949 — Kd 67.8 nM
Fractional Occupancy
1.0
0.8
0.6
0.4
0.2
0.0
$10^{-1}$
$10^{0}$
$10^{1}$
$10^{2}$
$10^{3}$
Protein concentration (nM, log scale)
Data
Model (Kd=67.8 nM)

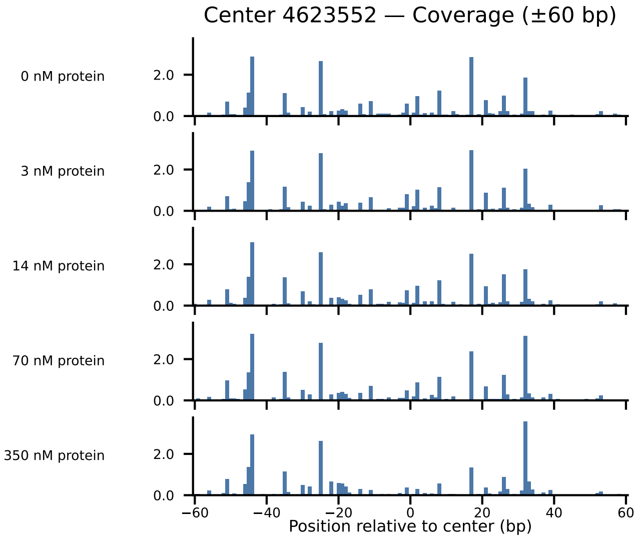

Center 4623552 — Coverage (±60 bp)
0 nM protein
3 nM protein
14 nM protein
70 nM protein
350 nM protein
2.0
0.0
−60
−40
−20
0
20
40
60
Position relative to center (bp)

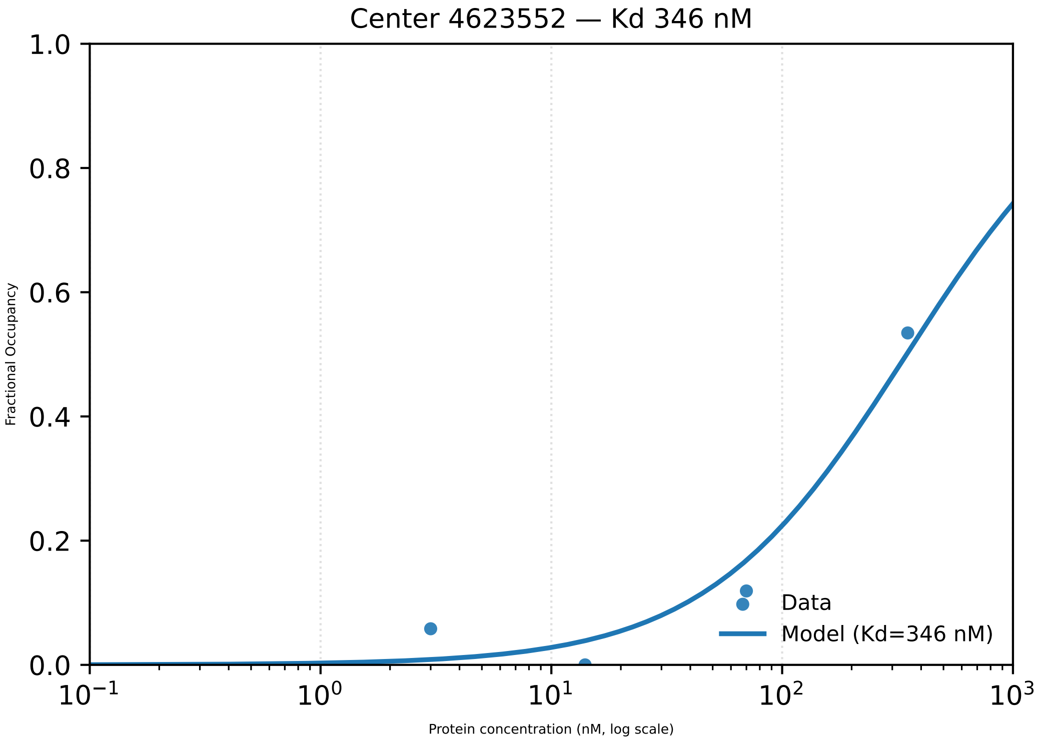
Center 4623552 — Kd 346 nM
Fractional Occupancy
Protein concentration (nM, log scale)
1.0
0.8
0.6
0.4
0.2
0.0
$10^{-1}$
$10^{0}$
$10^{1}$
$10^{2}$
$10^{3}$
Data
Model (Kd=346 nM)

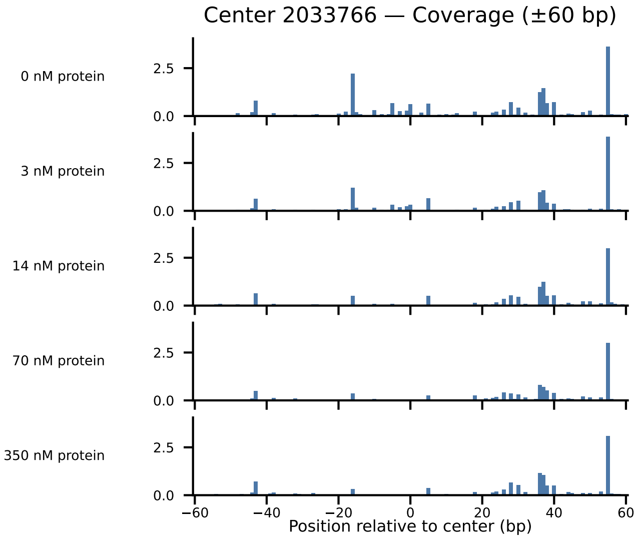

Center 2033766 — Coverage (±60 bp)
0 nM protein
3 nM protein
14 nM protein
70 nM protein
350 nM protein
2.5
0.0
−60
−40
−20
0
20
40
60
Position relative to center (bp)

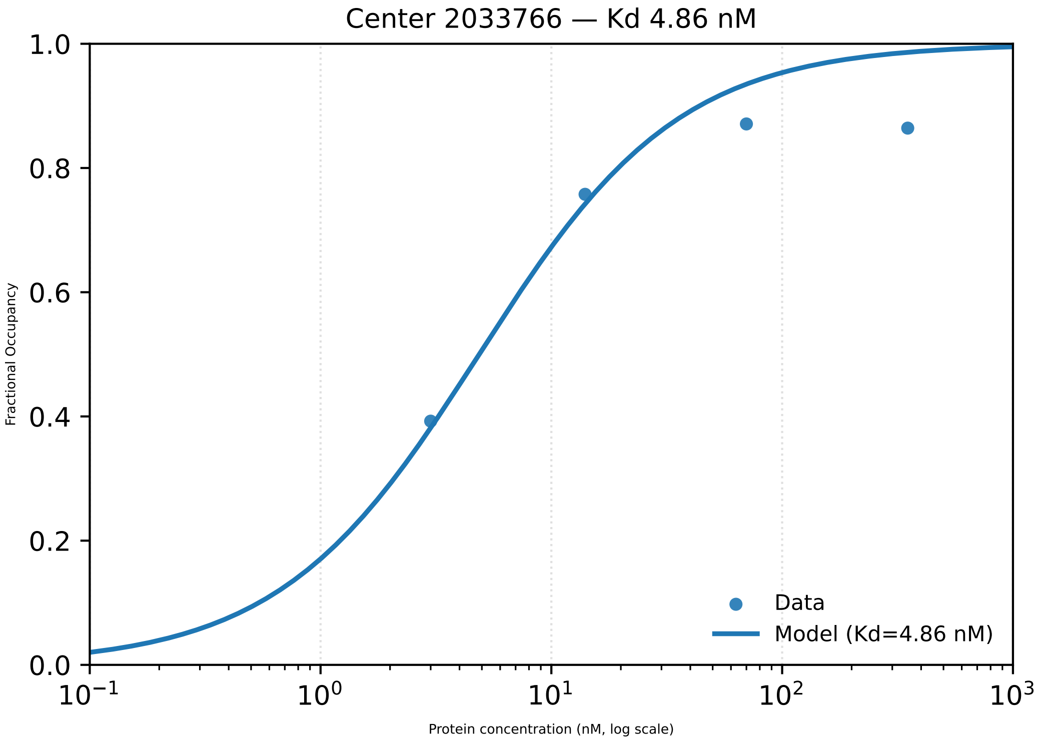

Center 2033766 — Kd 4.86 nM
Fractional Occupancy
1.0
0.8
0.6
0.4
0.2
0.0
$10^{-1}$
$10^{0}$
$10^{1}$
$10^{2}$
$10^{3}$
Protein concentration (nM, log scale)
Data
Model (Kd=4.86 nM)

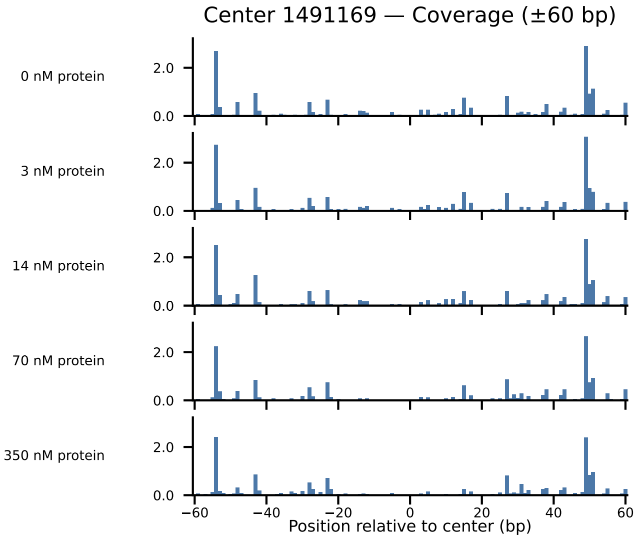
Center 1491169 — Coverage (±60 bp)
0 nM protein
3 nM protein
14 nM protein
70 nM protein
350 nM protein
2.0
0.0
−60
−40
−20
0
20
40
60
Position relative to center (bp)

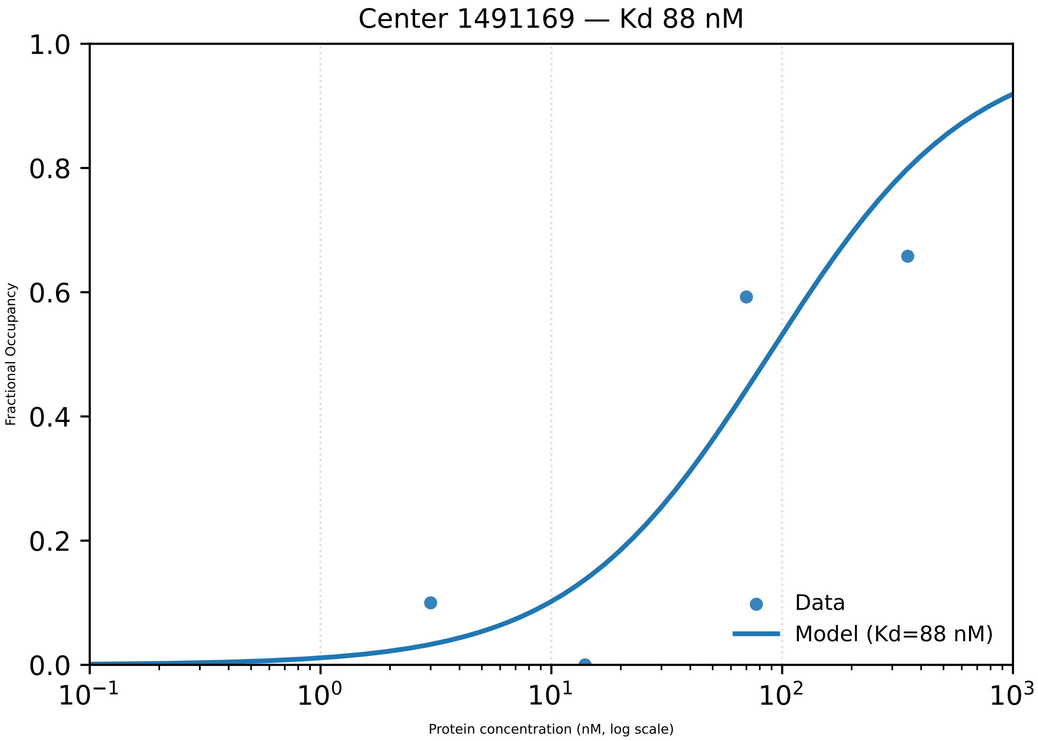
Center 1491169 — Kd 88 nM
Fractional Occupancy
Protein concentration (nM, log scale)
Data
Model (Kd=88 nM)
0.0
0.2
0.4
0.6
0.8
1.0
$10^{-1}$
$10^{0}$
$10^{1}$
$10^{2}$
$10^{3}$

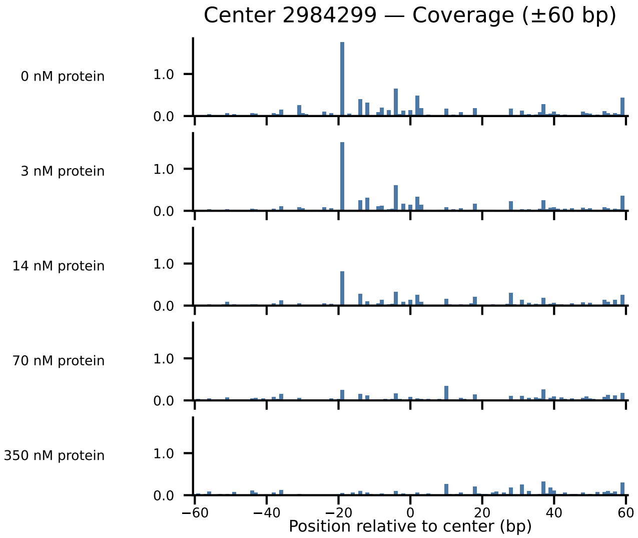

Center 2984299 — Coverage (±60 bp)
0 nM protein
3 nM protein
14 nM protein
70 nM protein
350 nM protein
1.0
0.0
−60
−40
−20
0
20
40
60
Position relative to center (bp)

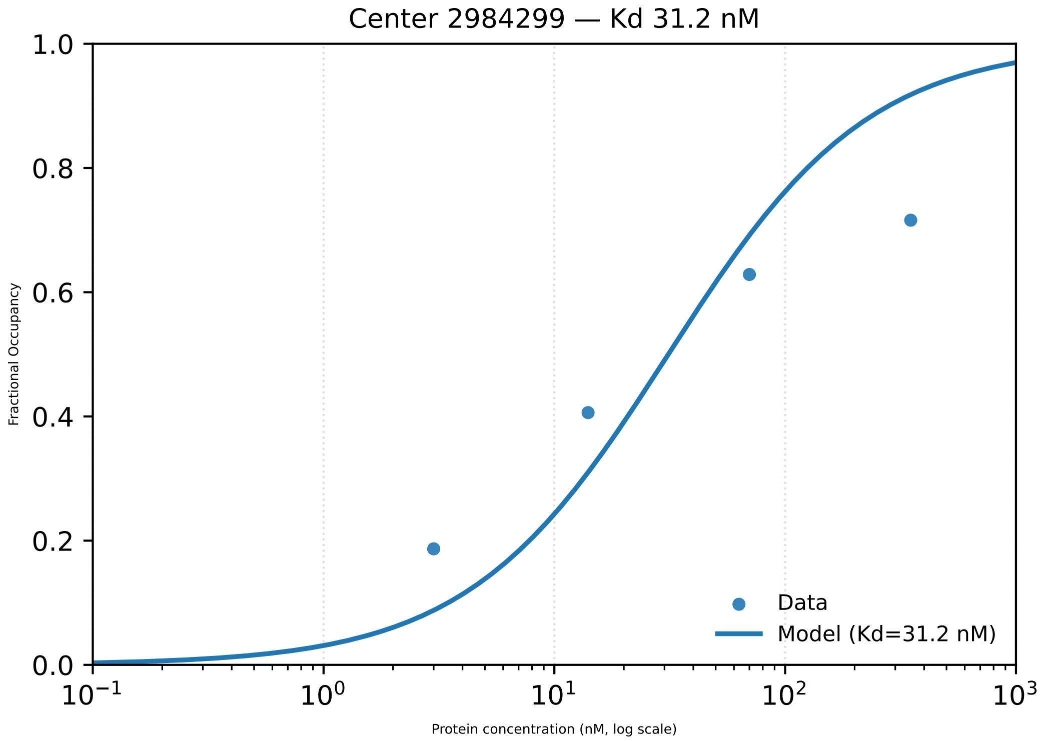
Center 2984299 — Kd 31.2 nM
1.0
0.8
0.6
0.4
0.2
0.0
Fractional Occupancy
$10^{-1}$
$10^{0}$
$10^{1}$
$10^{2}$
$10^{3}$
Protein concentration (nM, log scale)
Data
Model (Kd=31.2 nM)

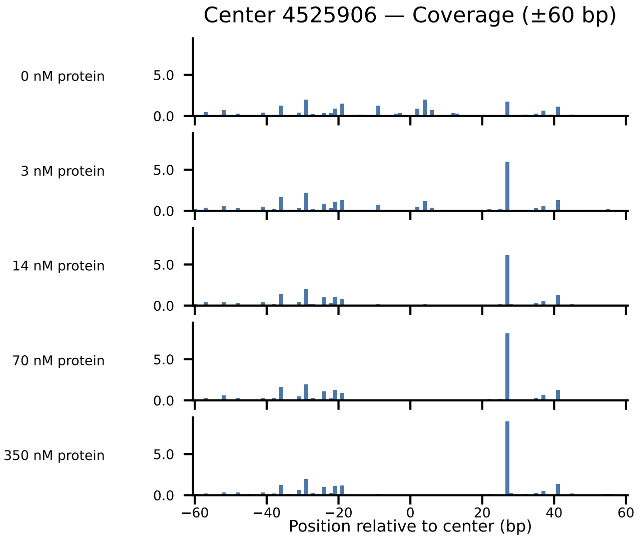
Center 4525906 — Coverage (±60 bp)
0 nM protein
3 nM protein
14 nM protein
70 nM protein
350 nM protein
5.0
0.0
−60
−40
−20
0
20
40
60
Position relative to center (bp)

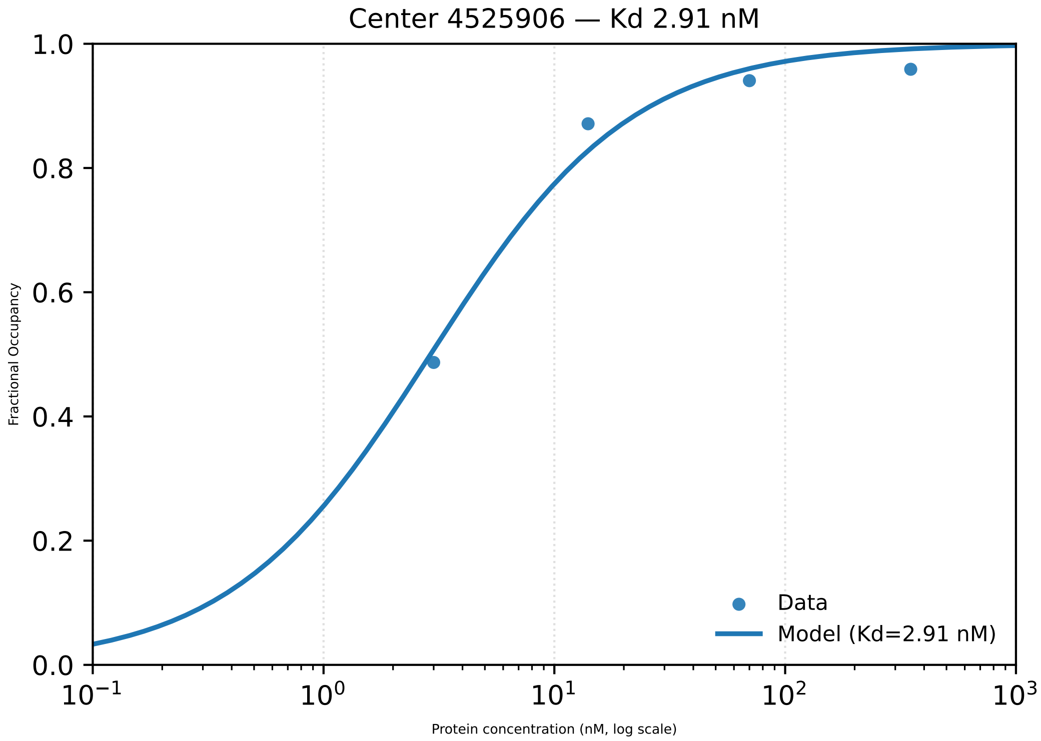
Center 4525906 — Kd 2.91 nM
Fractional Occupancy
1.0
0.8
0.6
0.4
0.2
0.0
$10^{-1}$
$10^{0}$
$10^{1}$
$10^{2}$
$10^{3}$
Protein concentration (nM, log scale)
Data
Model (Kd=2.91 nM)

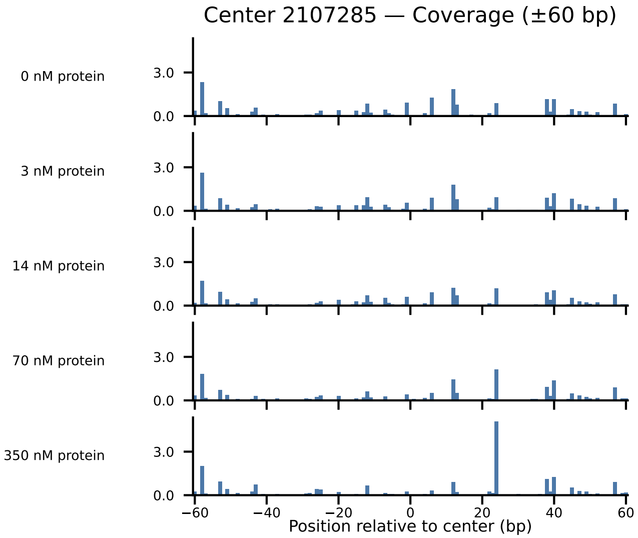
Center 2107285 — Coverage (±60 bp)
0 nM protein
3 nM protein
14 nM protein
70 nM protein
350 nM protein
3.0
0.0
−60
−40
−20
0
20
40
60
Position relative to center (bp)

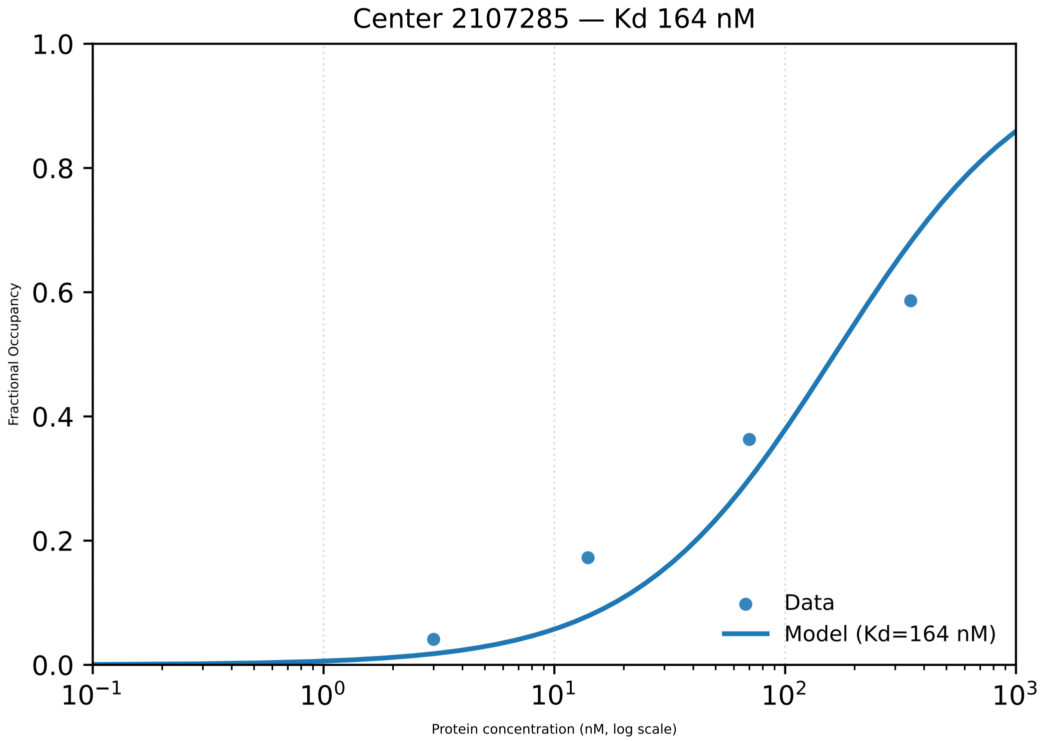

Center 2107285 — Kd 164 nM
1.0
0.8
0.6
0.4
0.2
0.0
Fractional Occupancy
$10^{-1}$
$10^{0}$
$10^{1}$
$10^{2}$
$10^{3}$
Protein concentration (nM, log scale)
Data
Model (Kd=164 nM)

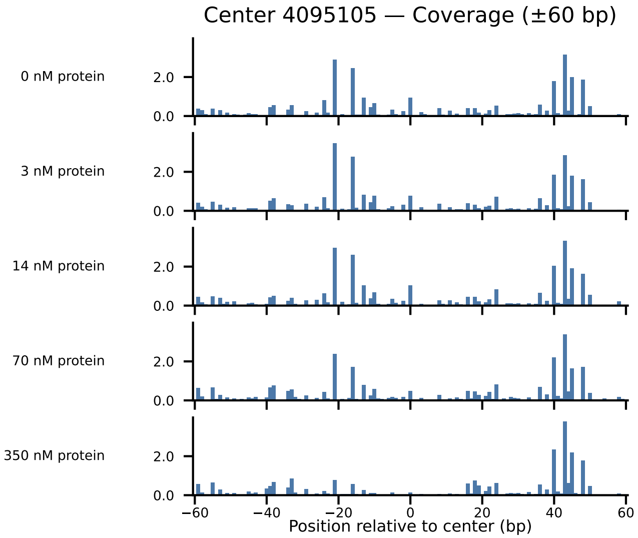
Center 4095105 — Coverage (±60 bp)
0 nM protein
3 nM protein
14 nM protein
70 nM protein
350 nM protein
2.0
0.0
−60
−40
−20
0
20
40
60
Position relative to center (bp)

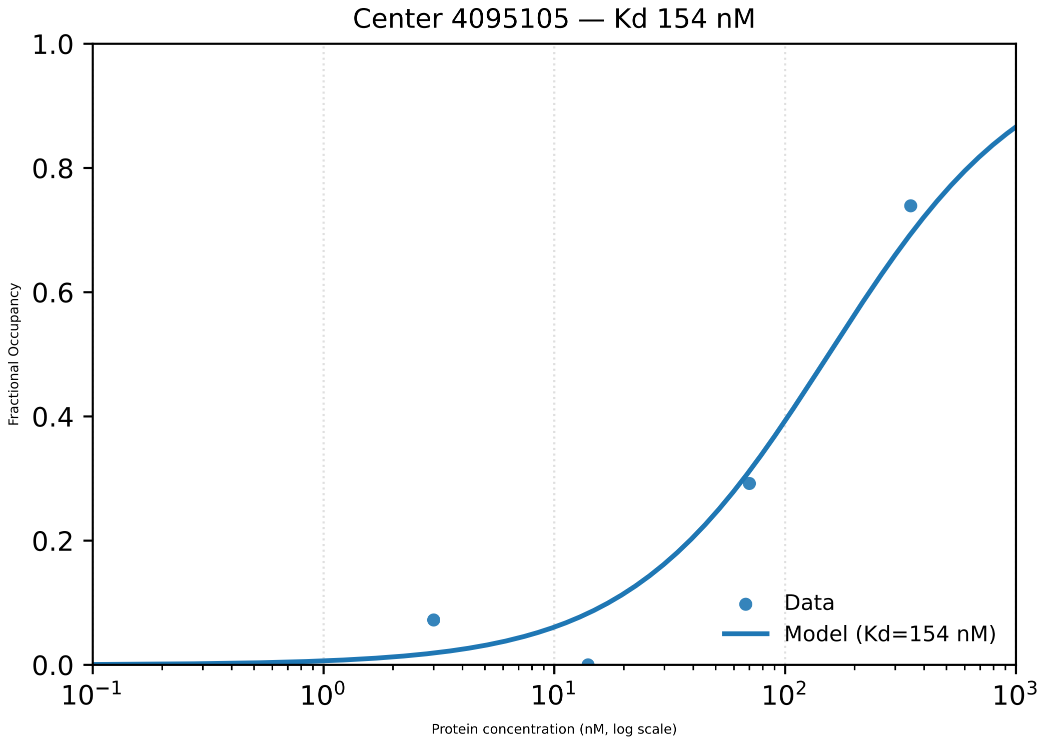
Center 4095105 — Kd 154 nM
1.0
0.8
0.6
0.4
0.2
0.0
Fractional Occupancy
$10^{-1}$
$10^{0}$
$10^{1}$
$10^{2}$
$10^{3}$
Protein concentration (nM, log scale)
Data
Model (Kd=154 nM)

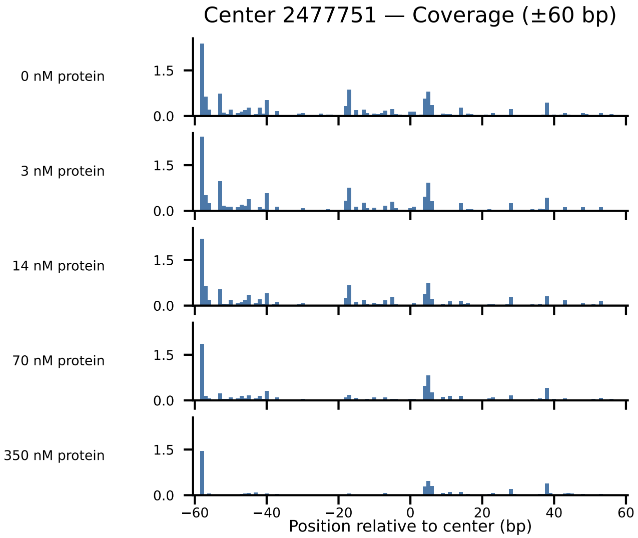
Center 2477751 — Coverage (±60 bp)
0 nM protein
3 nM protein
14 nM protein
70 nM protein
350 nM protein
1.5
0.0
−60
−40
−20
0
20
40
60
Position relative to center (bp)

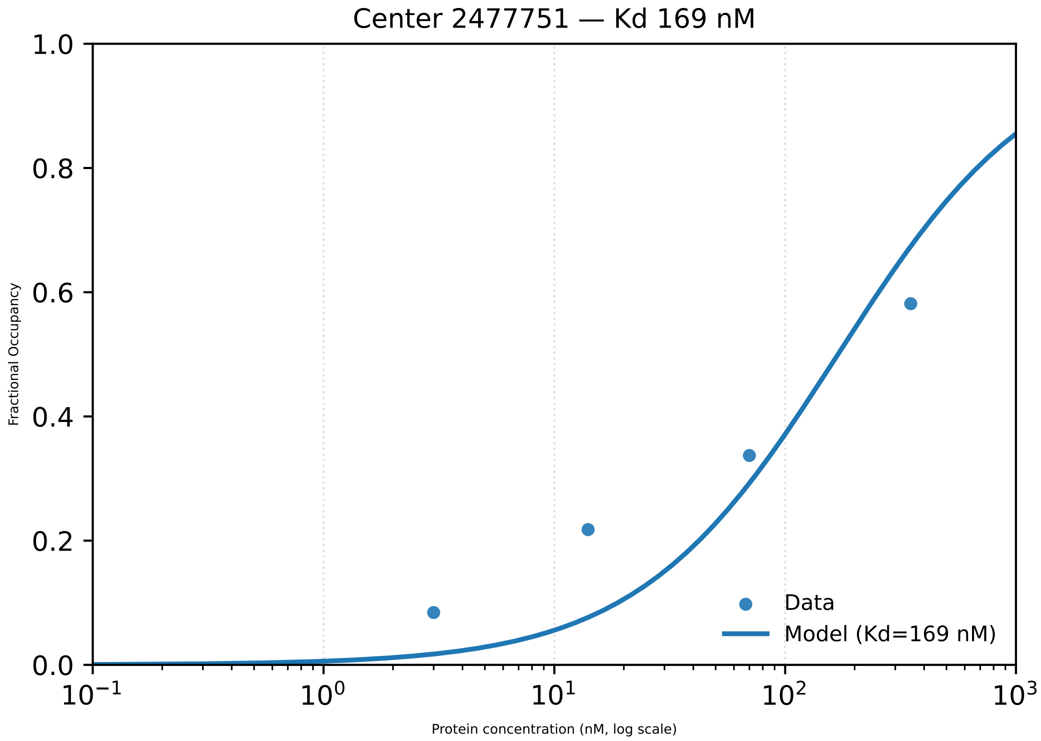

Center 2477751 — Kd 169 nM
1.0
0.8
0.6
0.4
0.2
0.0
Fractional Occupancy
Data
Model (Kd=169 nM)
$10^{-1}$
$10^{0}$
$10^{1}$
$10^{2}$
$10^{3}$
Protein concentration (nM, log scale)

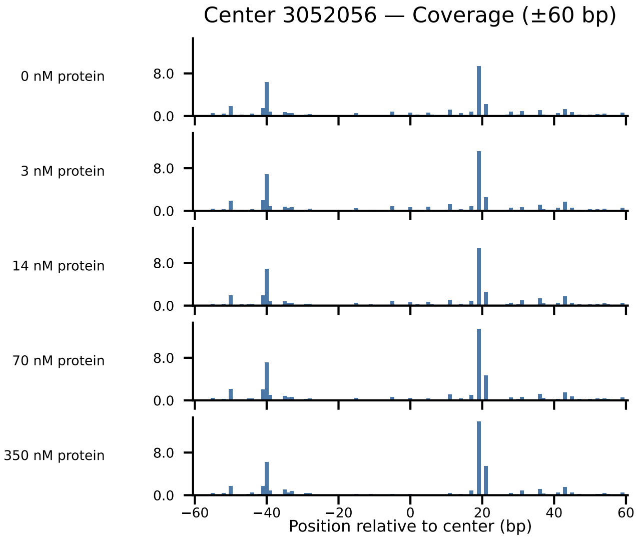
Center 3052056 — Coverage (±60 bp)
0 nM protein
3 nM protein
14 nM protein
70 nM protein
350 nM protein
8.0
0.0
−60
−40
−20
0
20
40
60
Position relative to center (bp)

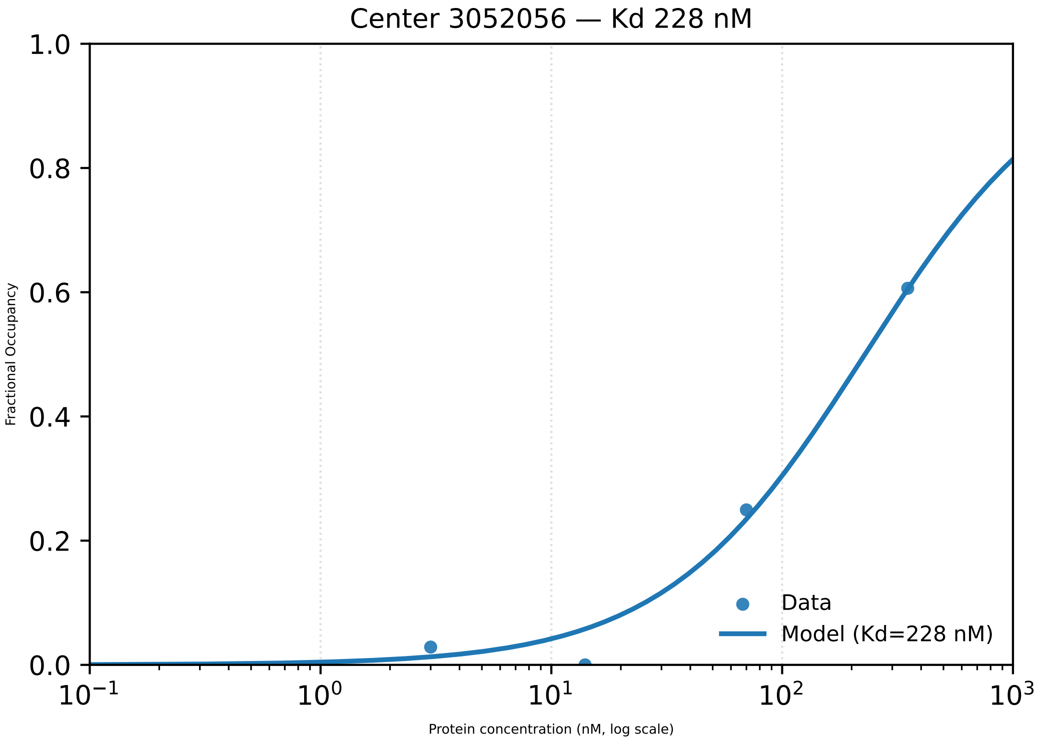

Center 3052056 — Kd 228 nM
1.0
0.8
0.6
0.4
0.2
0.0
Fractional Occupancy
$10^{-1}$
$10^{0}$
$10^{1}$
$10^{2}$
$10^{3}$
Protein concentration (nM, log scale)
Data
Model (Kd=228 nM)

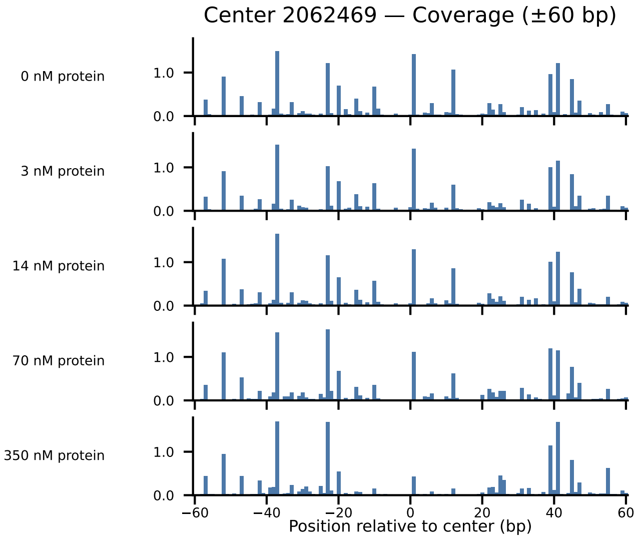
Center 2062469 — Coverage (±60 bp)
0 nM protein
3 nM protein
14 nM protein
70 nM protein
350 nM protein
1.0
0.0
−60
−40
−20
0
20
40
60
Position relative to center (bp)

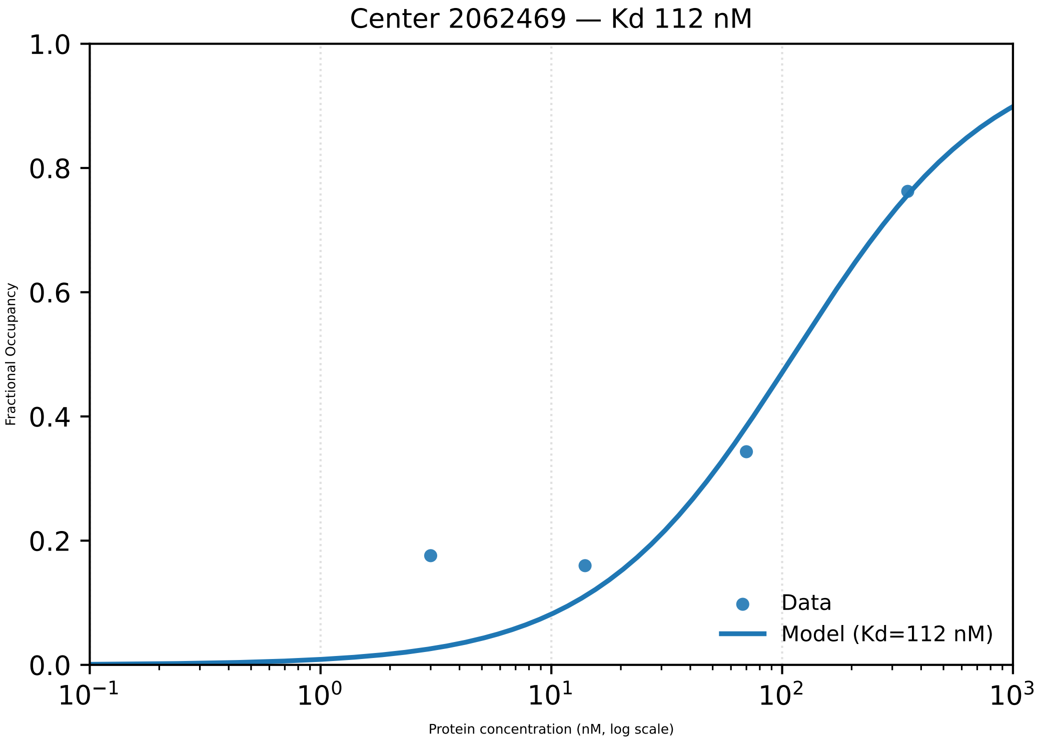
Center 2062469 — Kd 112 nM
1.0
0.8
0.6
0.4
0.2
0.0
Fractional Occupancy
$10^{-1}$
$10^{0}$
$10^{1}$
$10^{2}$
$10^{3}$
Protein concentration (nM, log scale)
Data
Model (Kd=112 nM)

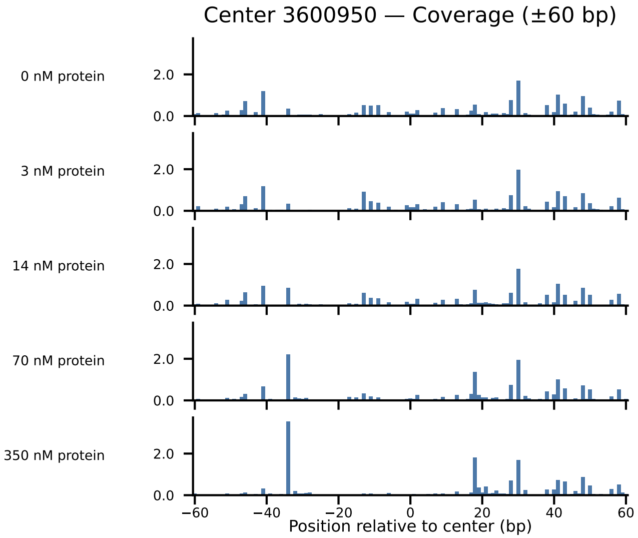
Center 3600950 — Coverage (±60 bp)
0 nM protein
3 nM protein
14 nM protein
70 nM protein
350 nM protein
2.0
0.0
−60
−40
−20
0
20
40
60
Position relative to center (bp)

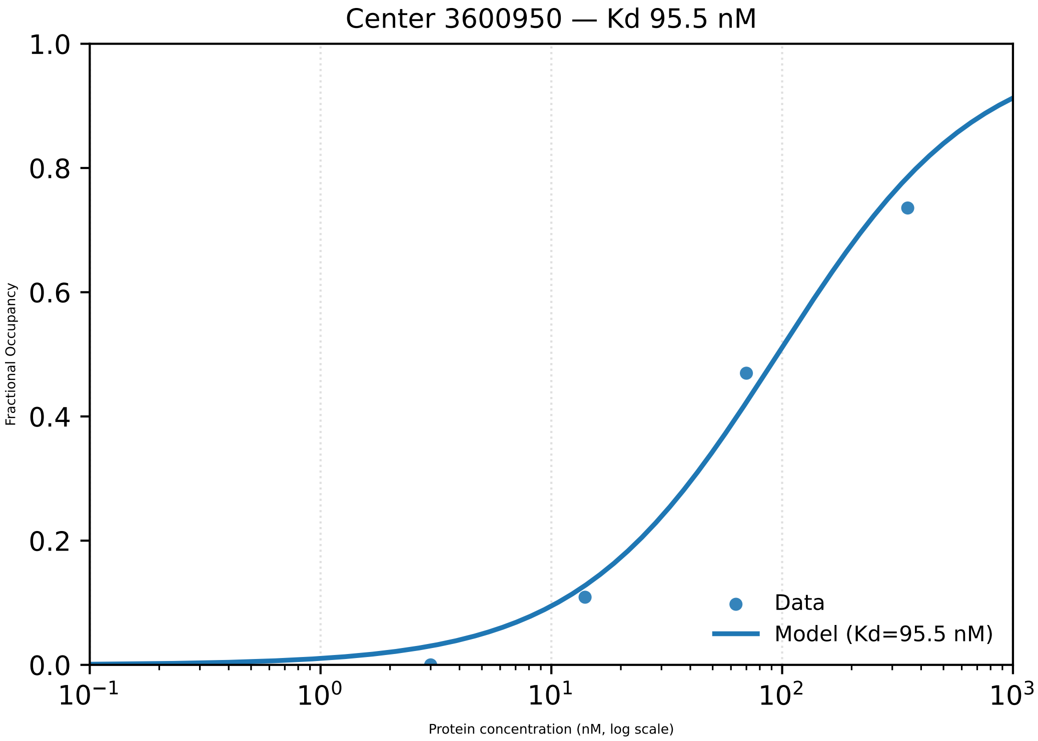

Center 3600950 — Kd 95.5 nM
Fractional Occupancy
1.0
0.8
0.6
0.4
0.2
0.0
$10^{-1}$
$10^{0}$
$10^{1}$
$10^{2}$
$10^{3}$
Protein concentration (nM, log scale)
Data
Model (Kd=95.5 nM)

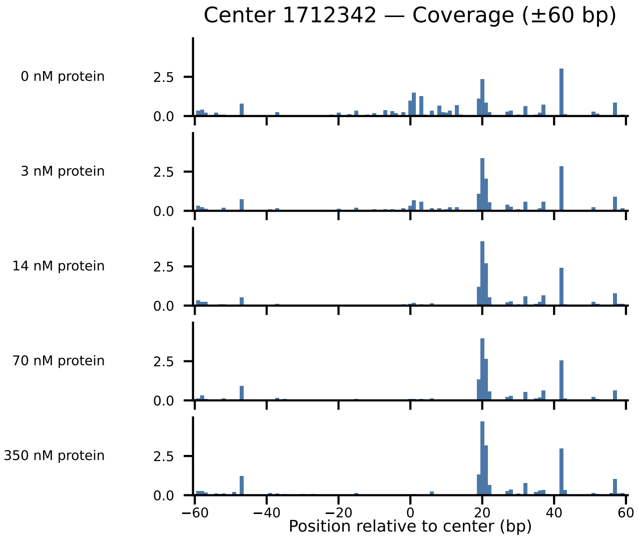
Center 1712342 — Coverage (±60 bp)
0 nM protein
3 nM protein
14 nM protein
70 nM protein
350 nM protein
2.5
0.0
−60
−40
−20
0
20
40
60
Position relative to center (bp)

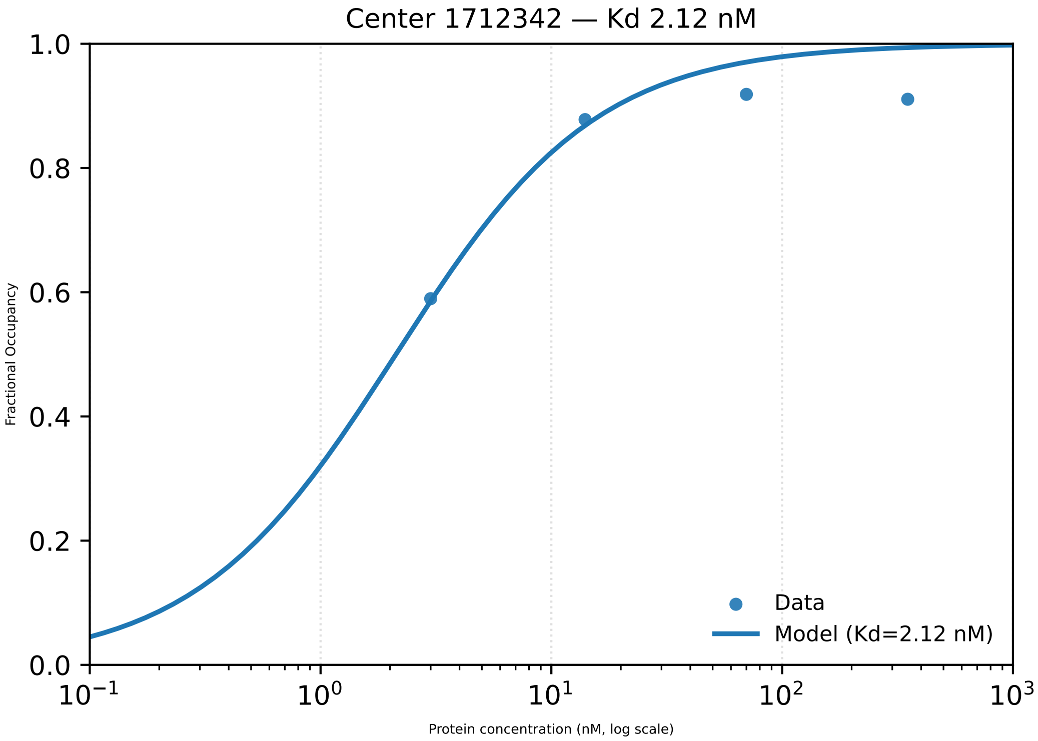
Center 1712342 — Kd 2.12 nM
Fractional Occupancy
Protein concentration (nM, log scale)
1.0
0.8
0.6
0.4
0.2
0.0
$10^{-1}$
$10^{0}$
$10^{1}$
$10^{2}$
$10^{3}$
Data
Model (Kd=2.12 nM)

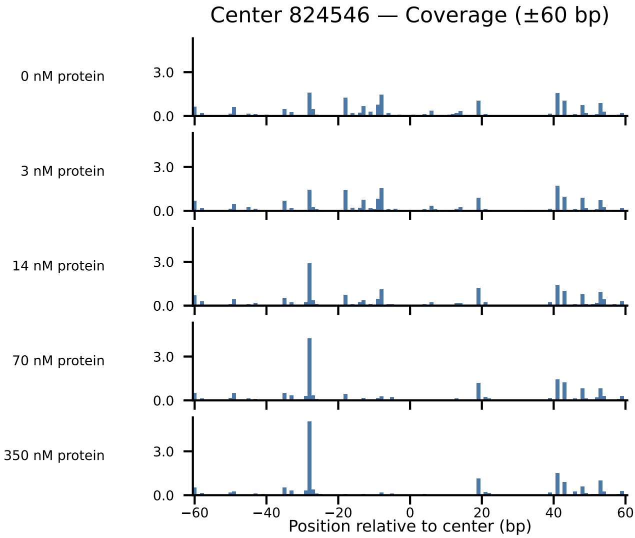

Center 824546 — Coverage (±60 bp)
0 nM protein
3 nM protein
14 nM protein
70 nM protein
350 nM protein
3.0
0.0
−60
−40
−20
0
20
40
60
Position relative to center (bp)

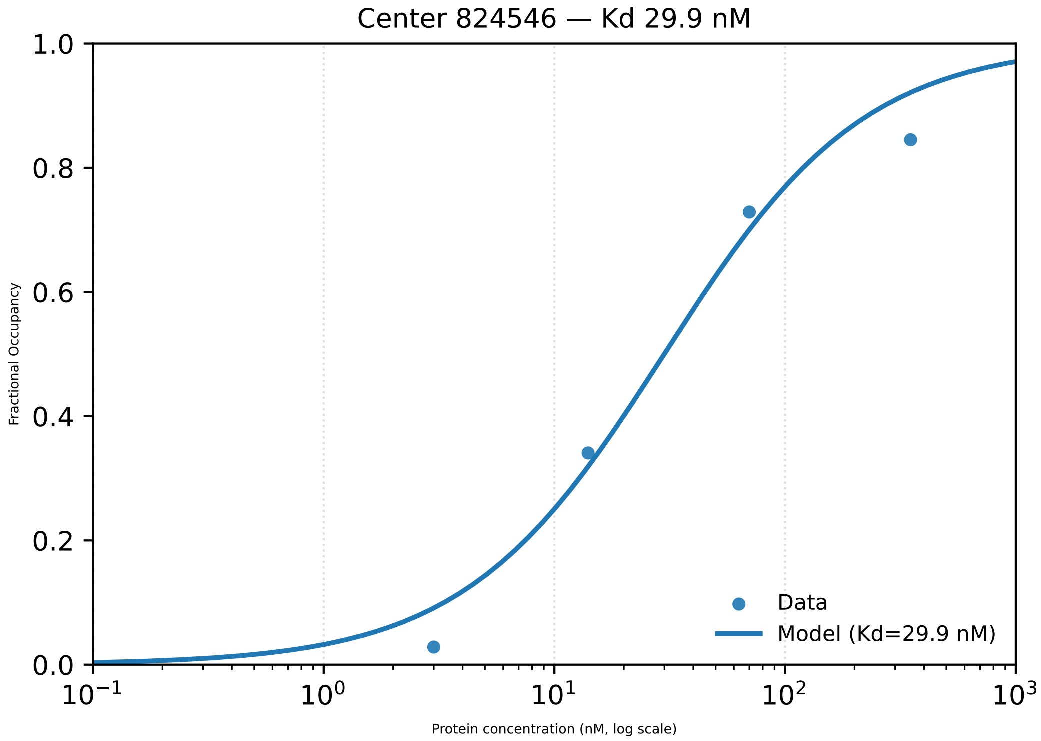

Center 824546 — Kd 29.9 nM
Fractional Occupancy
1.0
0.8
0.6
0.4
0.2
0.0
$10^{-1}$
$10^{0}$
$10^{1}$
$10^{2}$
$10^{3}$
Protein concentration (nM, log scale)
Data
Model (Kd=29.9 nM)

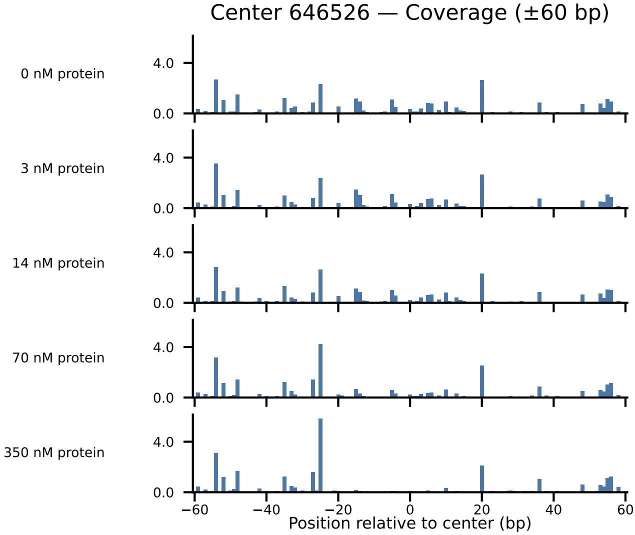

Center 646526 — Coverage (±60 bp)
0 nM protein
3 nM protein
14 nM protein
70 nM protein
350 nM protein
4.0
0.0
−60
−40
−20
0
20
40
60
Position relative to center (bp)

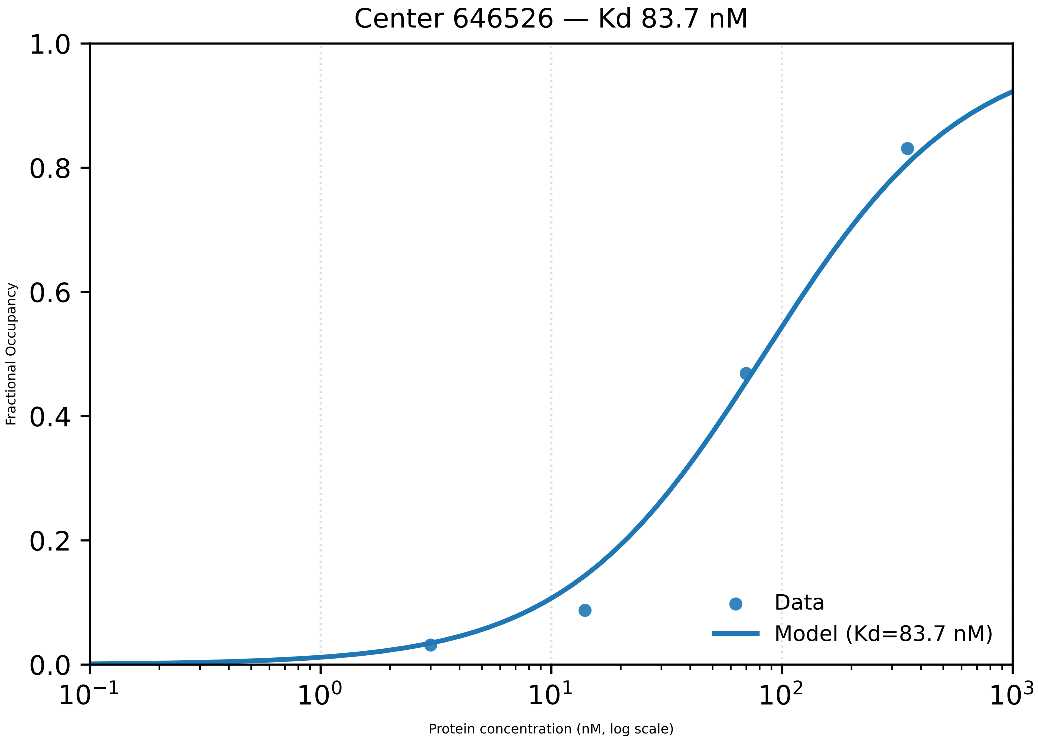

Center 646526 — Kd 83.7 nM
1.0
0.8
0.6
0.4
0.2
0.0
Fractional Occupancy
$10^{-1}$
$10^{0}$
$10^{1}$
$10^{2}$
$10^{3}$
Protein concentration (nM, log scale)
Data
Model (Kd=83.7 nM)

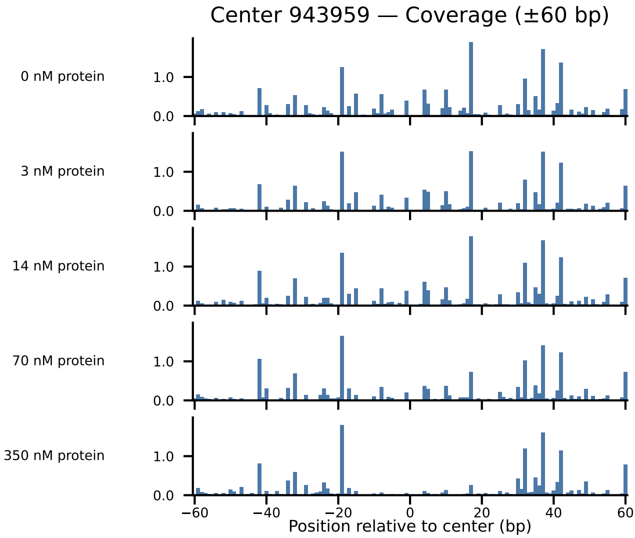
Center 943959 — Coverage (±60 bp)
0 nM protein
3 nM protein
14 nM protein
70 nM protein
350 nM protein
1.0
0.0
−60
−40
−20
0
20
40
60
Position relative to center (bp)

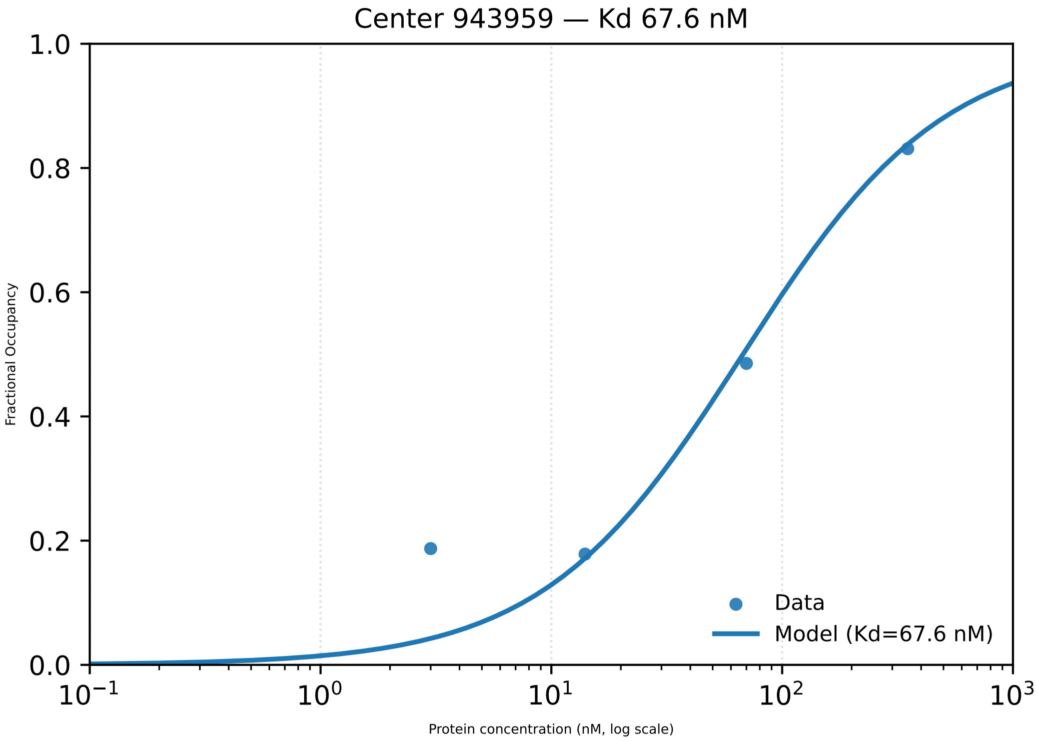
Center 943959 — Kd 67.6 nM
1.0
0.8
0.6
0.4
0.2
0.0
Fractional Occupancy
$10^{-1}$
$10^{0}$
$10^{1}$
$10^{2}$
$10^{3}$
Protein concentration (nM, log scale)
Data
Model (Kd=67.6 nM)

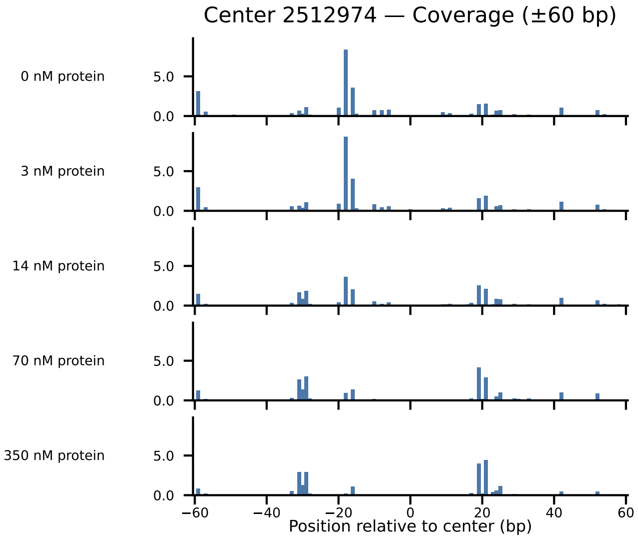

Center 2512974 — Coverage (±60 bp)
0 nM protein
3 nM protein
14 nM protein
70 nM protein
350 nM protein
5.0
0.0
−60
−40
−20
0
20
40
60
Position relative to center (bp)

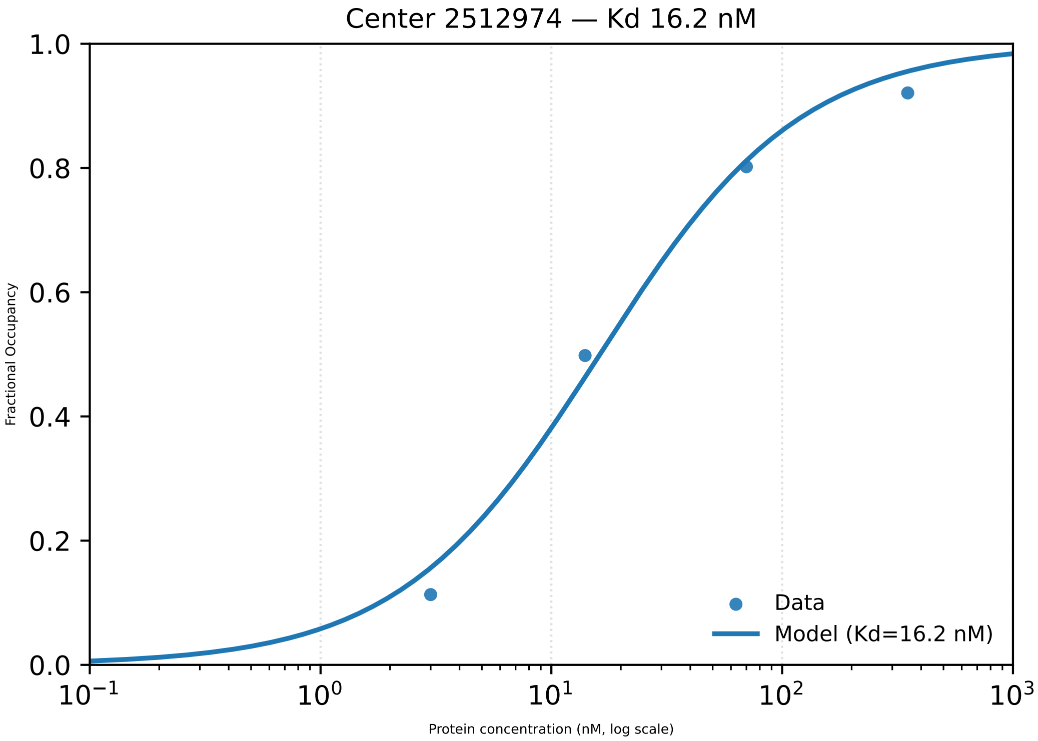

Center 2512974 — Kd 16.2 nM
1.0
0.8
0.6
0.4
0.2
0.0
Fractional Occupancy
$10^{-1}$
$10^{0}$
$10^{1}$
$10^{2}$
$10^{3}$
Protein concentration (nM, log scale)
Data
Model (Kd=16.2 nM)

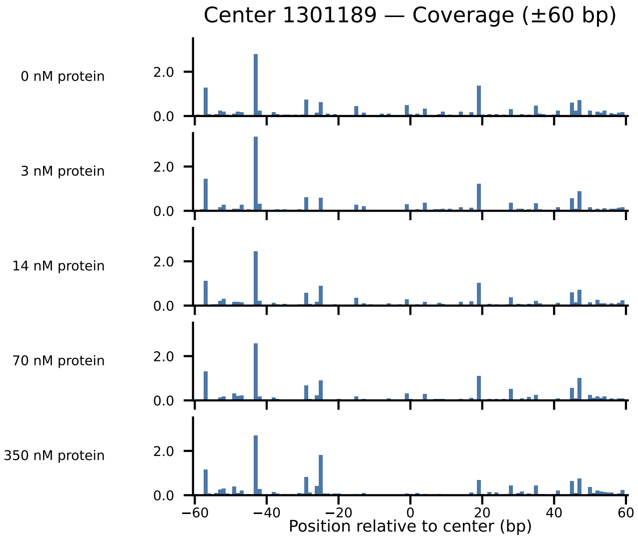

Center 1301189 — Coverage (±60 bp)
0 nM protein
3 nM protein
14 nM protein
70 nM protein
350 nM protein
2.0
0.0
−60
−40
−20
0
20
40
60
Position relative to center (bp)

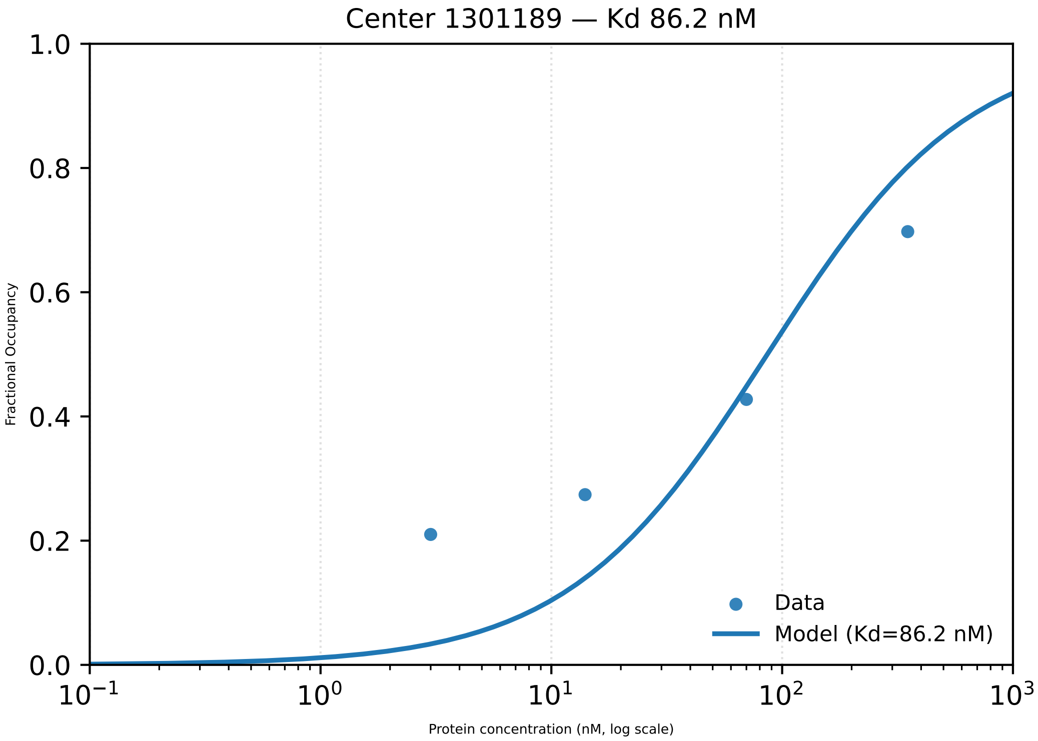

Center 1301189 — Kd 86.2 nM
1.0
0.8
0.6
0.4
0.2
0.0
Fractional Occupancy
$10^{-1}$
$10^{0}$
$10^{1}$
$10^{2}$
$10^{3}$
Protein concentration (nM, log scale)
Data
Model (Kd=86.2 nM)

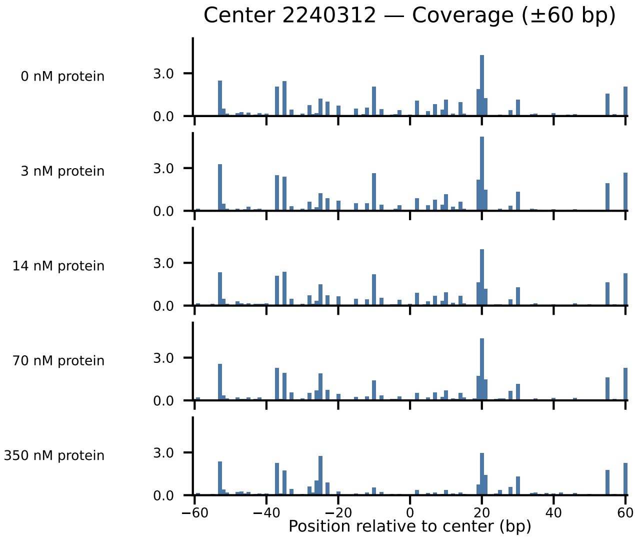

Center 2240312 — Coverage (±60 bp)
0 nM protein
3 nM protein
14 nM protein
70 nM protein
350 nM protein
3.0
0.0
−60
−40
−20
0
20
40
60
Position relative to center (bp)

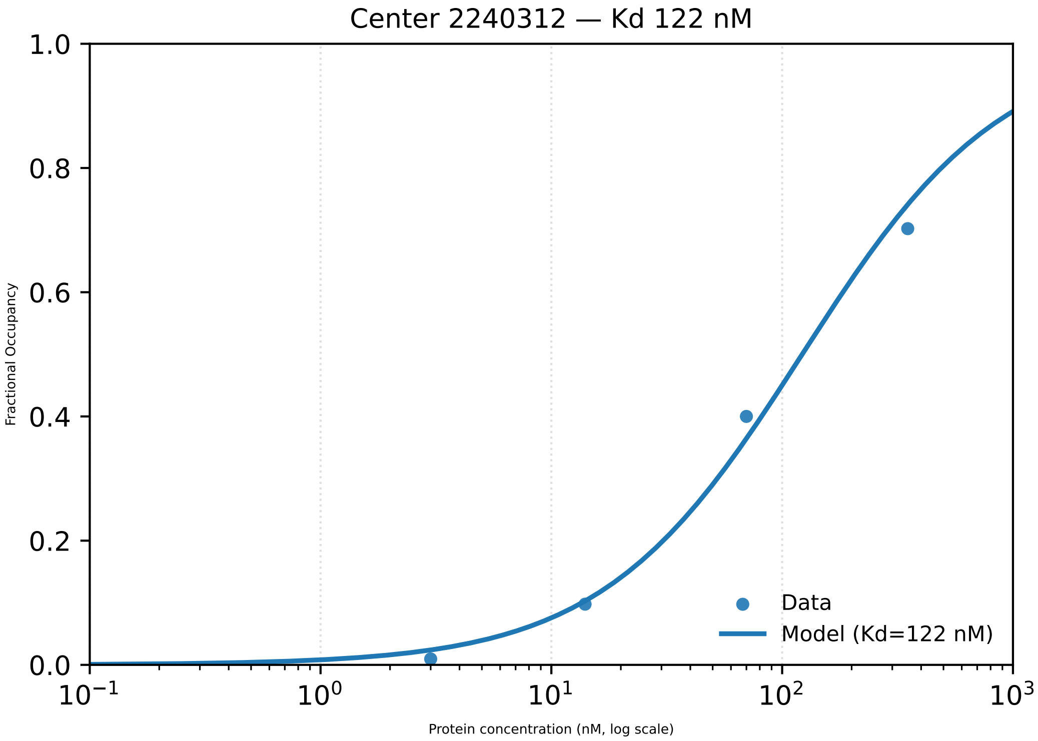

Center 2240312 — Kd 122 nM
Fractional Occupancy
1.0
0.8
0.6
0.4
0.2
0.0
$10^{-1}$
$10^{0}$
$10^{1}$
$10^{2}$
$10^{3}$
Protein concentration (nM, log scale)
Data
Model (Kd=122 nM)

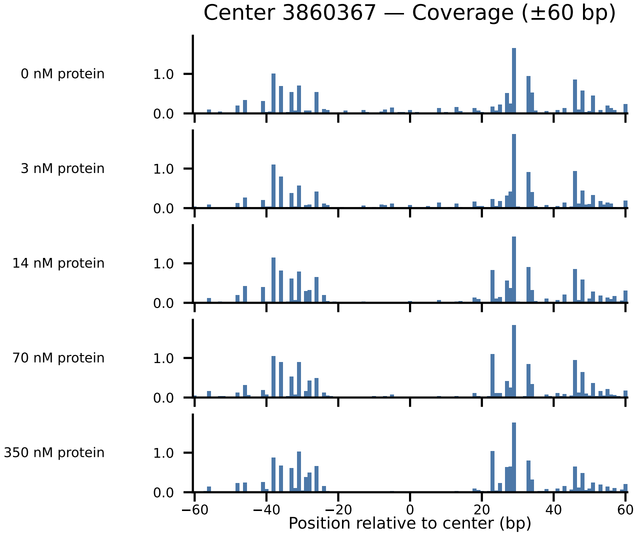
Center 3860367 — Coverage (±60 bp)
0 nM protein
3 nM protein
14 nM protein
70 nM protein
350 nM protein
1.0
0.0
−60
−40
−20
0
20
40
60
Position relative to center (bp)

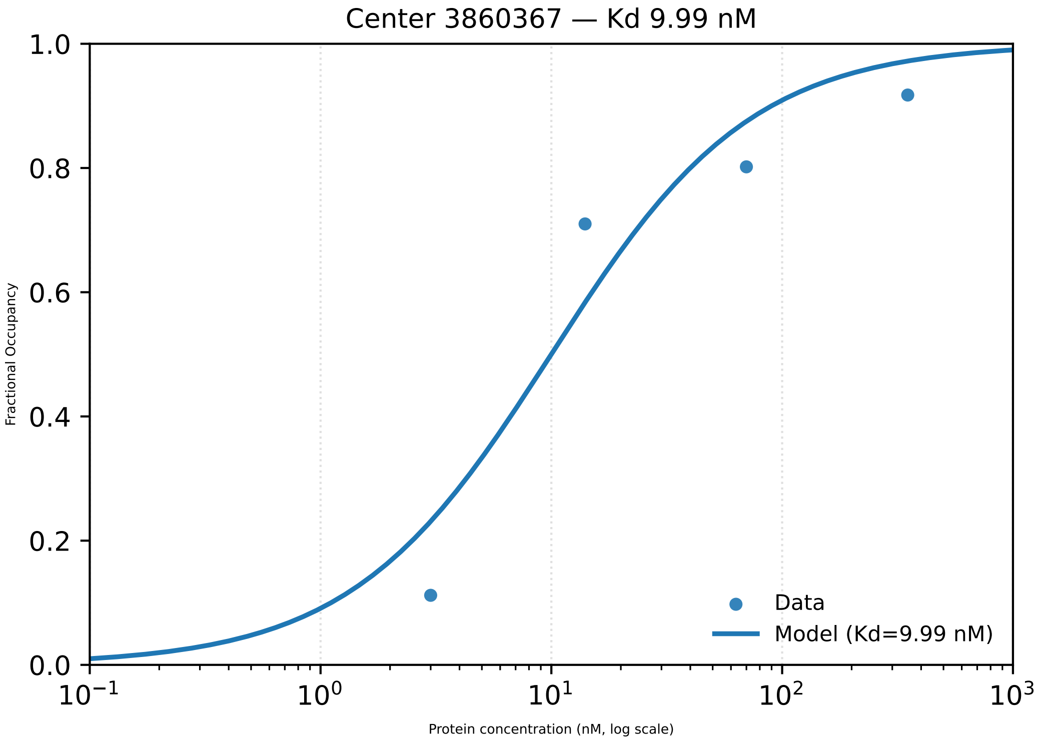
Center 3860367 — Kd 9.99 nM
1.0
0.8
0.6
0.4
0.2
0.0
Fractional Occupancy
$10^{-1}$
$10^{0}$
$10^{1}$
$10^{2}$
$10^{3}$
Protein concentration (nM, log scale)
Data
Model (Kd=9.99 nM)

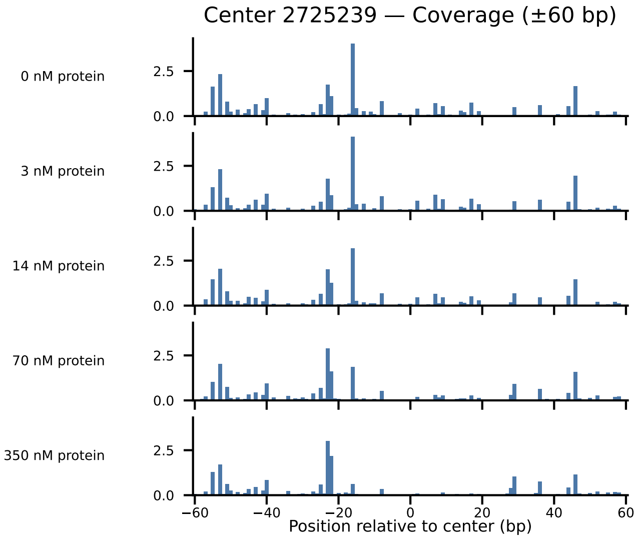
Center 2725239 — Coverage (±60 bp)
0 nM protein
3 nM protein
14 nM protein
70 nM protein
350 nM protein
2.5
0.0
−60
−40
−20
0
20
40
60
Position relative to center (bp)

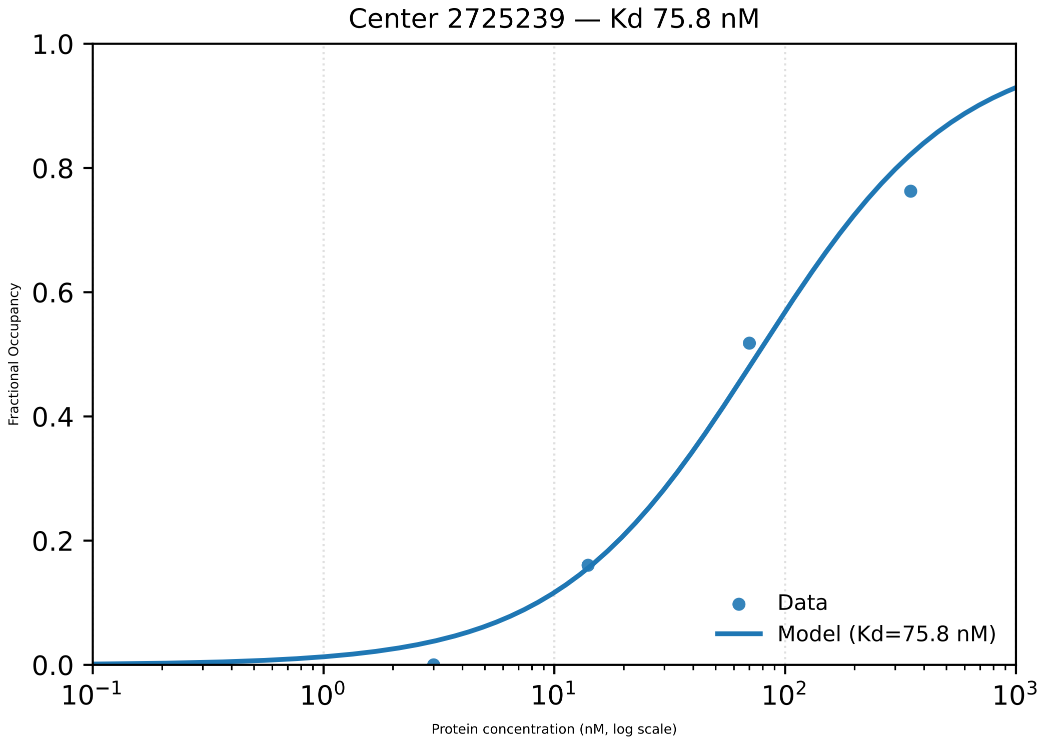

Center 2725239 — Kd 75.8 nM
1.0
0.8
0.6
0.4
0.2
0.0
Fractional Occupancy
$10^{-1}$
$10^{0}$
$10^{1}$
$10^{2}$
$10^{3}$
Protein concentration (nM, log scale)
Data
Model (Kd=75.8 nM)

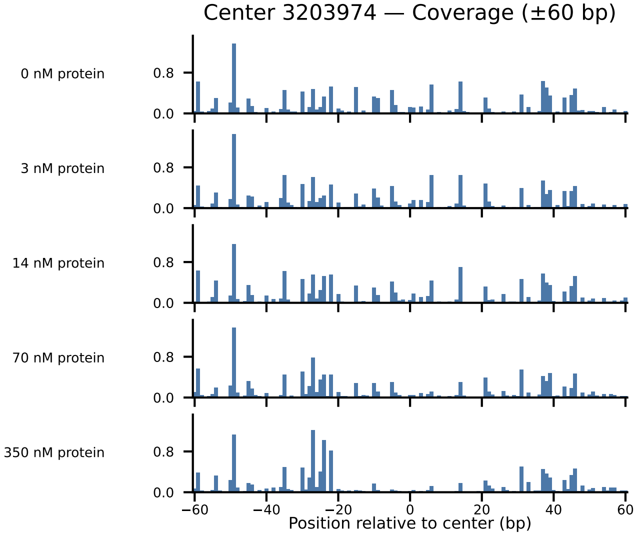
Center 3203974 — Coverage (±60 bp)
0 nM protein
3 nM protein
14 nM protein
70 nM protein
350 nM protein
0.8
0.0
−60
−40
−20
0
20
40
60
Position relative to center (bp)

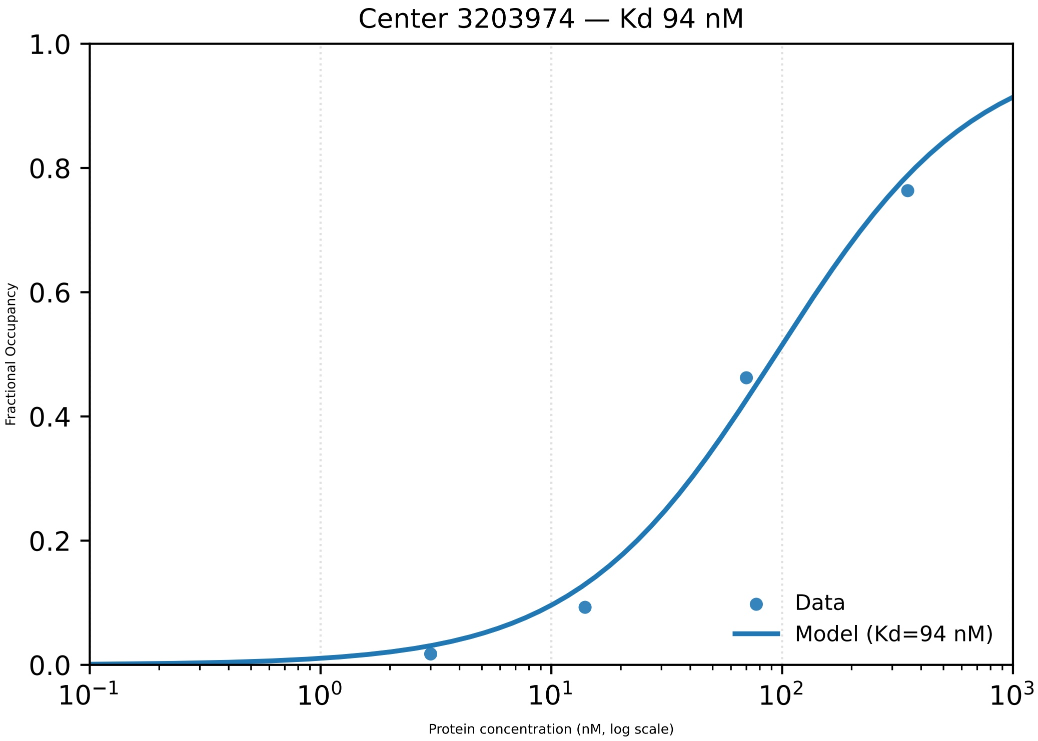

Center 3203974 — Kd 94 nM
1.0
0.8
0.6
0.4
0.2
0.0
Fractional Occupancy
$10^{-1}$
$10^{0}$
$10^{1}$
$10^{2}$
$10^{3}$
Protein concentration (nM, log scale)
Data
Model (Kd=94 nM)

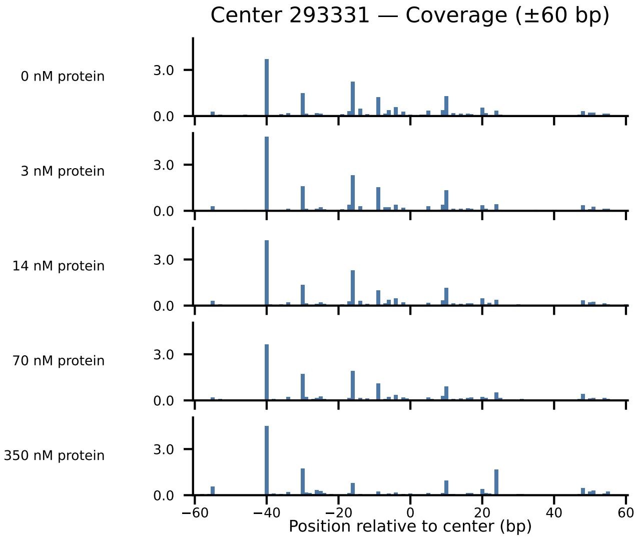

Center 293331 — Coverage (±60 bp)
0 nM protein
3 nM protein
14 nM protein
70 nM protein
350 nM protein
3.0
0.0
−60
−40
−20
0
20
40
60
Position relative to center (bp)

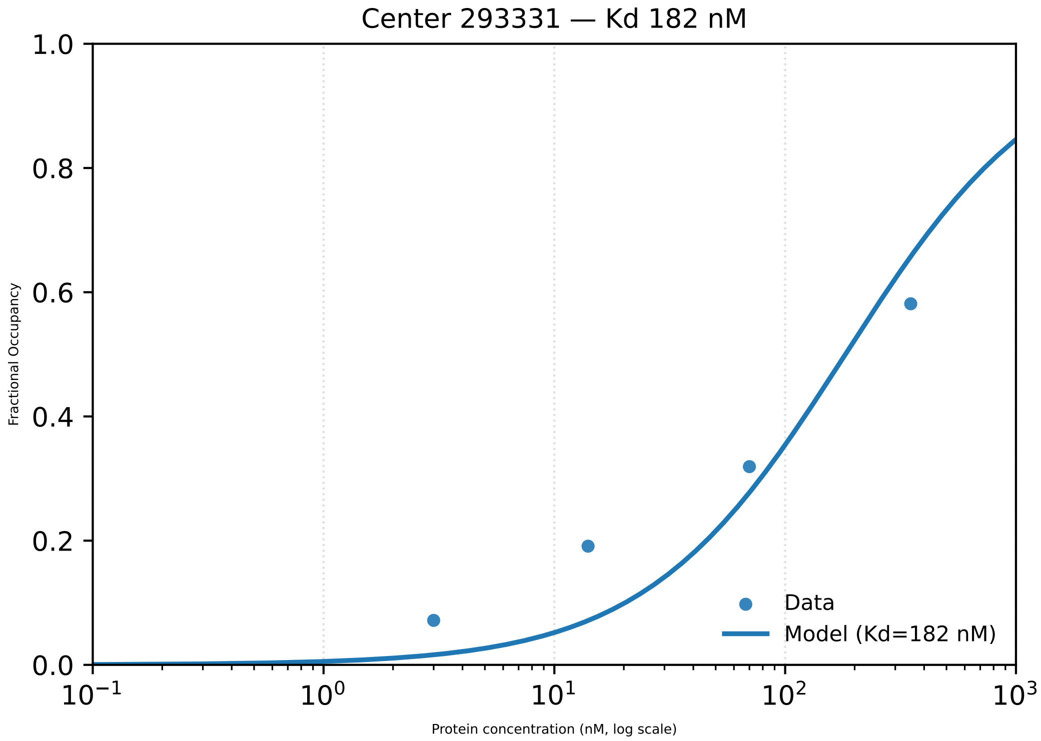

Center 293331 — Kd 182 nM
1.0
0.8
0.6
0.4
0.2
0.0
Fractional Occupancy
Data
Model (Kd=182 nM)
$10^{-1}$
$10^{0}$
$10^{1}$
$10^{2}$
$10^{3}$
Protein concentration (nM, log scale)

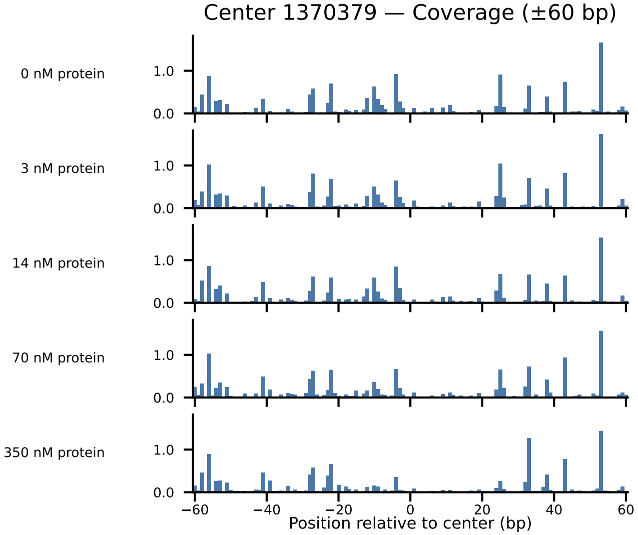
Center 1370379 — Coverage (±60 bp)
0 nM protein
3 nM protein
14 nM protein
70 nM protein
350 nM protein
1.0
0.0
−60
−40
−20
0
20
40
60
Position relative to center (bp)

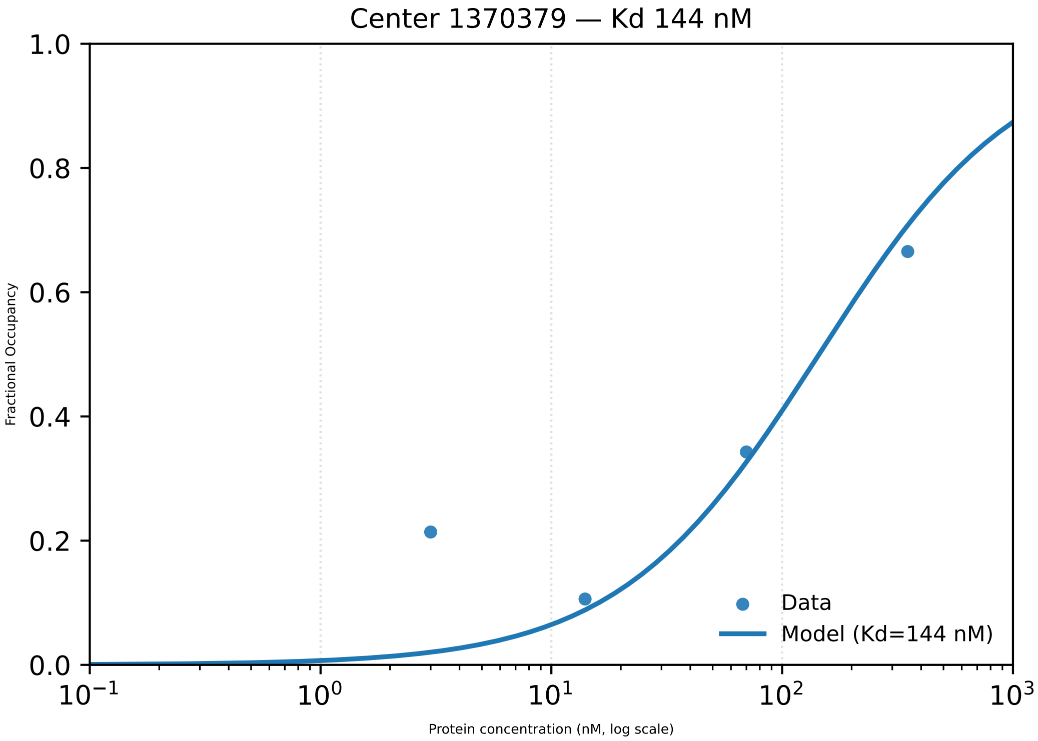

Center 1370379 — Kd 144 nM
Fractional Occupancy
Protein concentration (nM, log scale)
1.0
0.8
0.6
0.4
0.2
0.0
$10^{-1}$
$10^{0}$
$10^{1}$
$10^{2}$
$10^{3}$
Data
Model (Kd=144 nM)

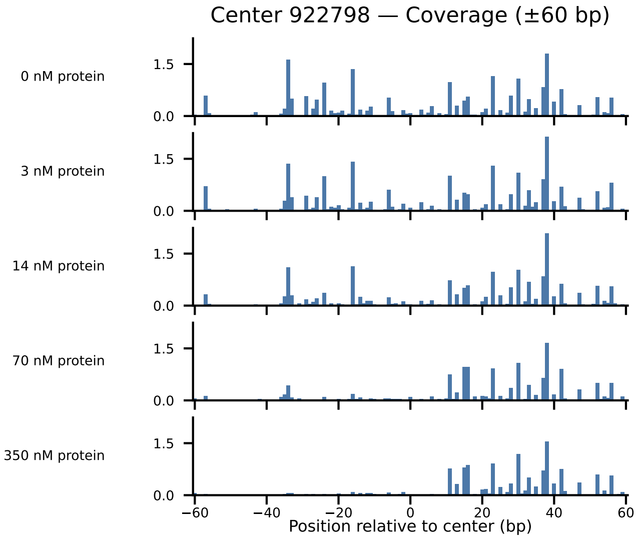
Center 922798 — Coverage (±60 bp)
0 nM protein
3 nM protein
14 nM protein
70 nM protein
350 nM protein
1.5
0.0
−60
−40
−20
0
20
40
60
Position relative to center (bp)

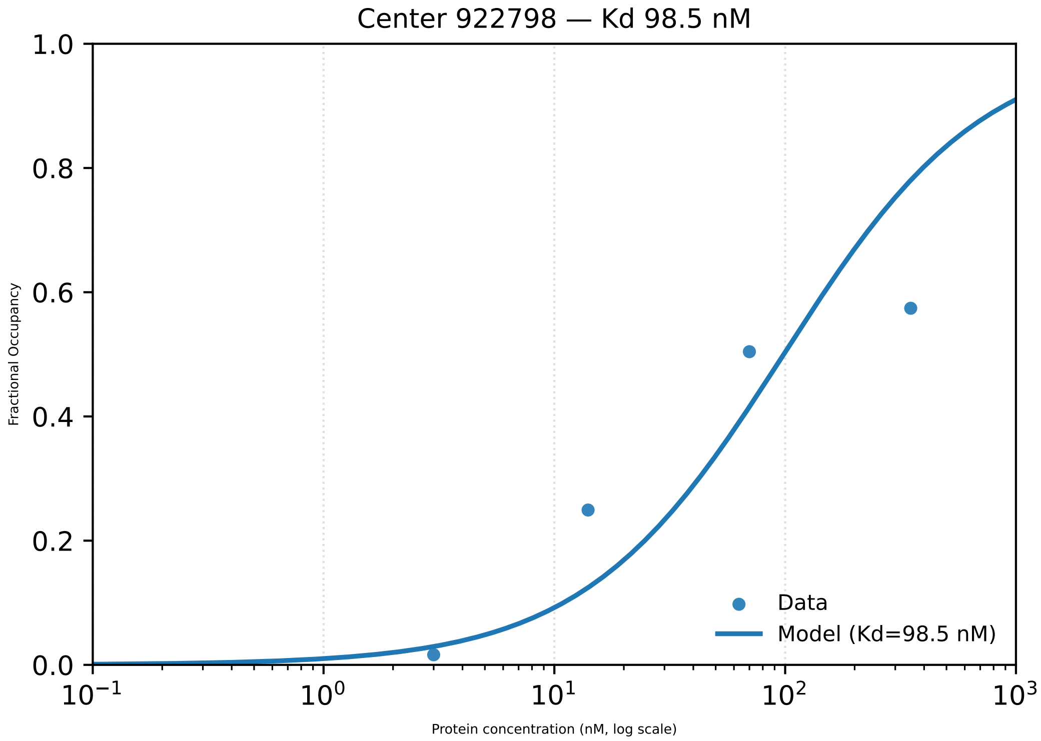

Center 922798 — Kd 98.5 nM
Fractional Occupancy
Protein concentration (nM, log scale)
Data
Model (Kd=98.5 nM)

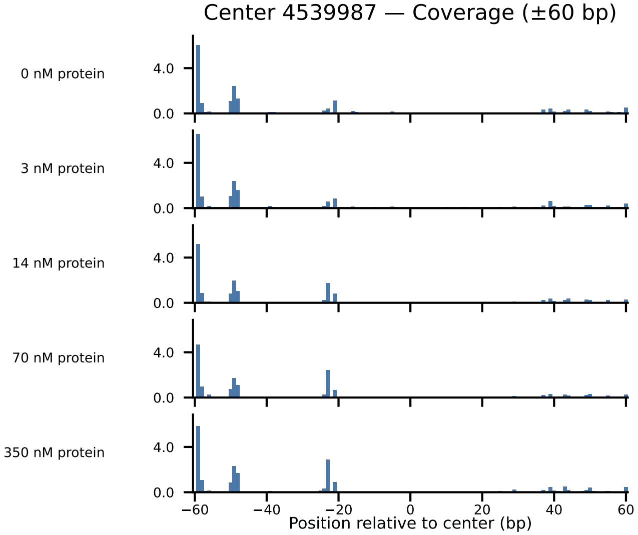
Center 4539987 — Coverage (±60 bp)
0 nM protein
3 nM protein
14 nM protein
70 nM protein
350 nM protein
4.0
0.0
−60
−40
−20
0
20
40
60
Position relative to center (bp)

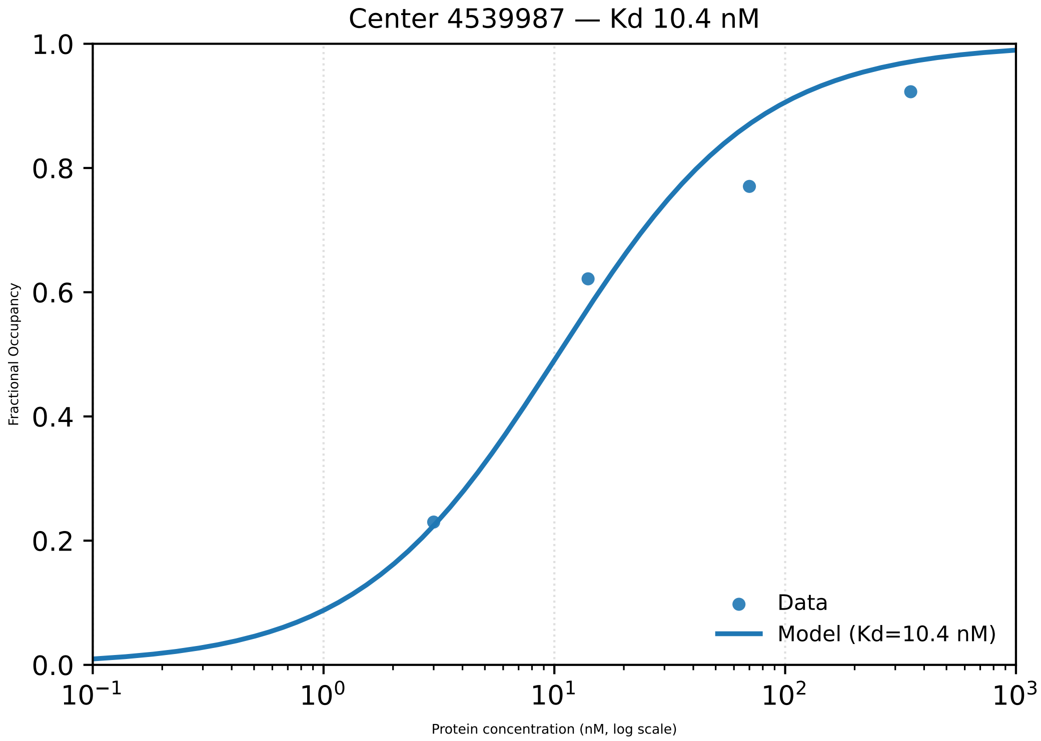

Center 4539987 — Kd 10.4 nM
1.0
0.8
0.6
0.4
0.2
0.0
Fractional Occupancy
$10^{-1}$
$10^{0}$
$10^{1}$
$10^{2}$
$10^{3}$
Protein concentration (nM, log scale)
Data
Model (Kd=10.4 nM)

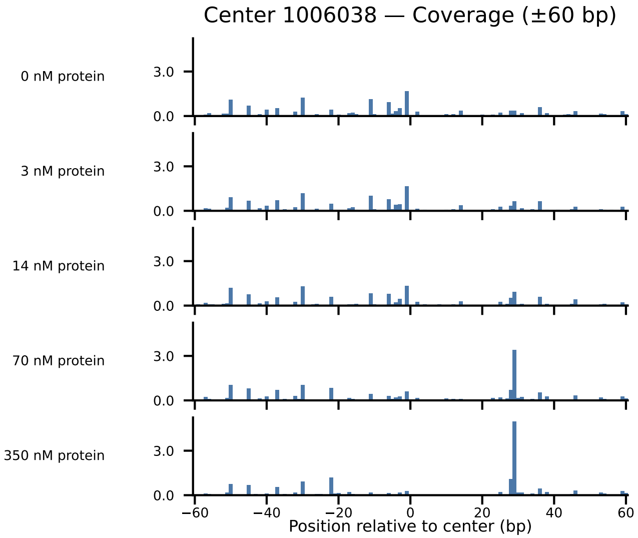
Center 1006038 — Coverage (±60 bp)
0 nM protein
3 nM protein
14 nM protein
70 nM protein
350 nM protein
3.0
0.0
−60
−40
−20
0
20
40
60
Position relative to center (bp)

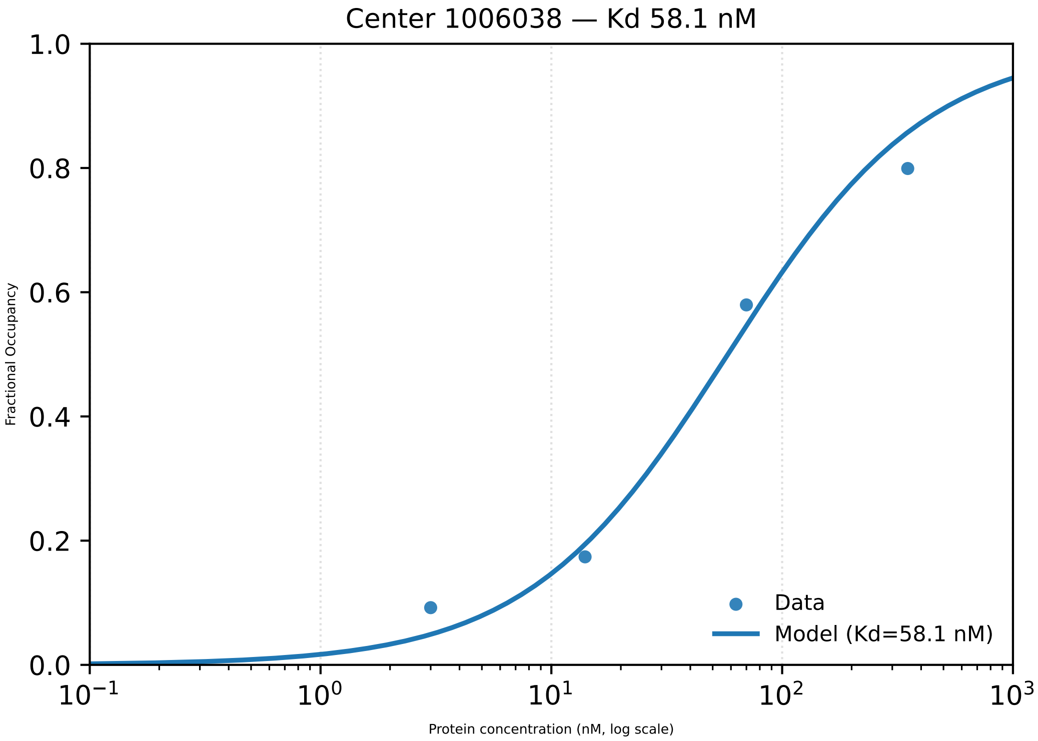

Center 1006038 — Kd 58.1 nM
1.0
0.8
0.6
0.4
0.2
0.0
Fractional Occupancy
Data
Model (Kd=58.1 nM)
$10^{-1}$
$10^{0}$
$10^{1}$
$10^{2}$
$10^{3}$
Protein concentration (nM, log scale)

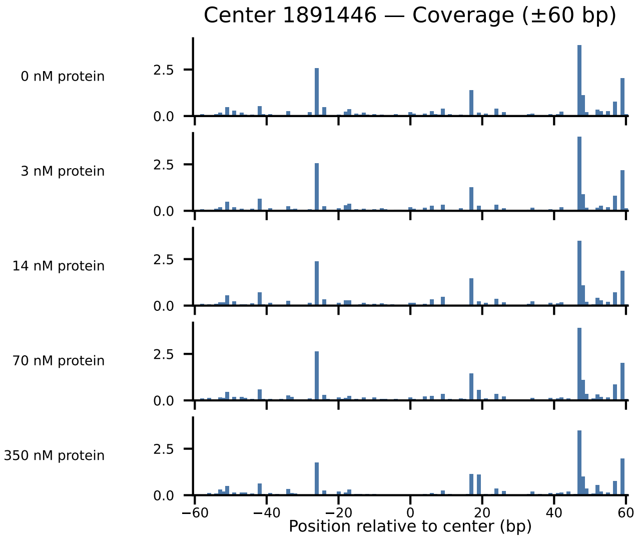

Center 1891446 — Coverage (±60 bp)
0 nM protein
3 nM protein
14 nM protein
70 nM protein
350 nM protein
2.5
0.0
−60
−40
−20
0
20
40
60
Position relative to center (bp)

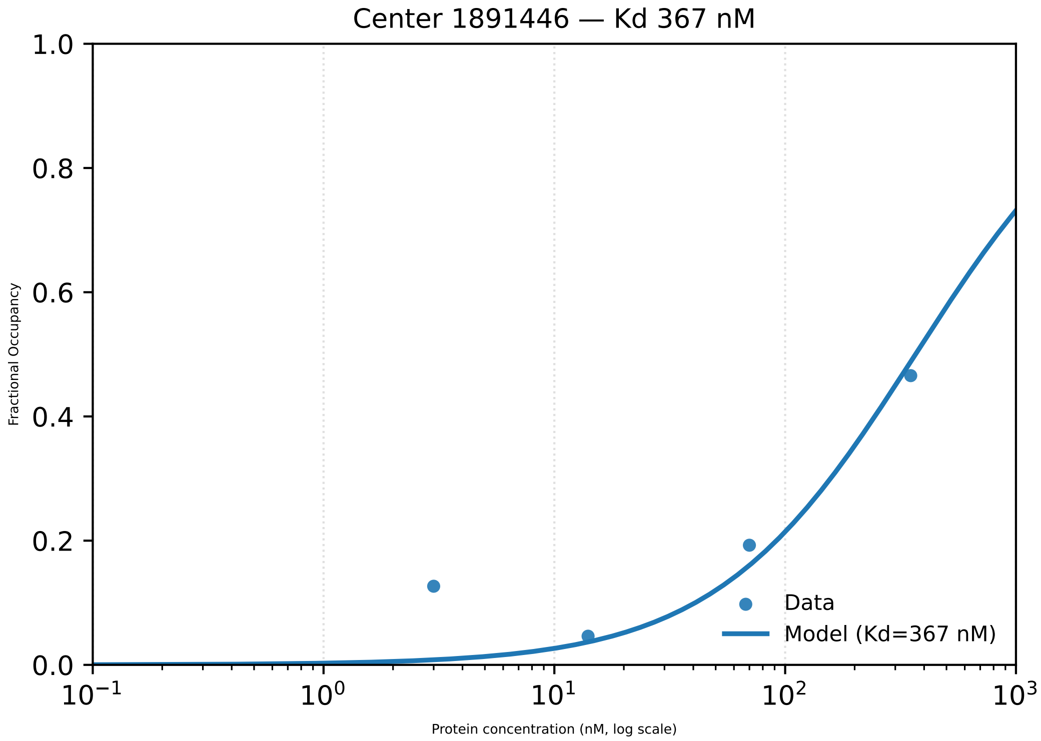

Center 1891446 — Kd 367 nM
1.0
0.8
0.6
0.4
0.2
0.0
Fractional Occupancy
$10^{-1}$
$10^{0}$
$10^{1}$
$10^{2}$
$10^{3}$
Protein concentration (nM, log scale)
Data
Model (Kd=367 nM)

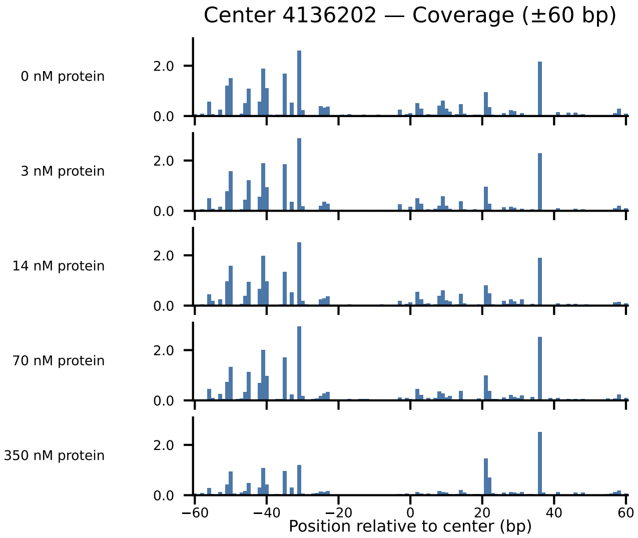

Center 4136202 — Coverage (±60 bp)
0 nM protein
3 nM protein
14 nM protein
70 nM protein
350 nM protein
2.0
0.0
−60
−40
−20
0
20
40
60
Position relative to center (bp)

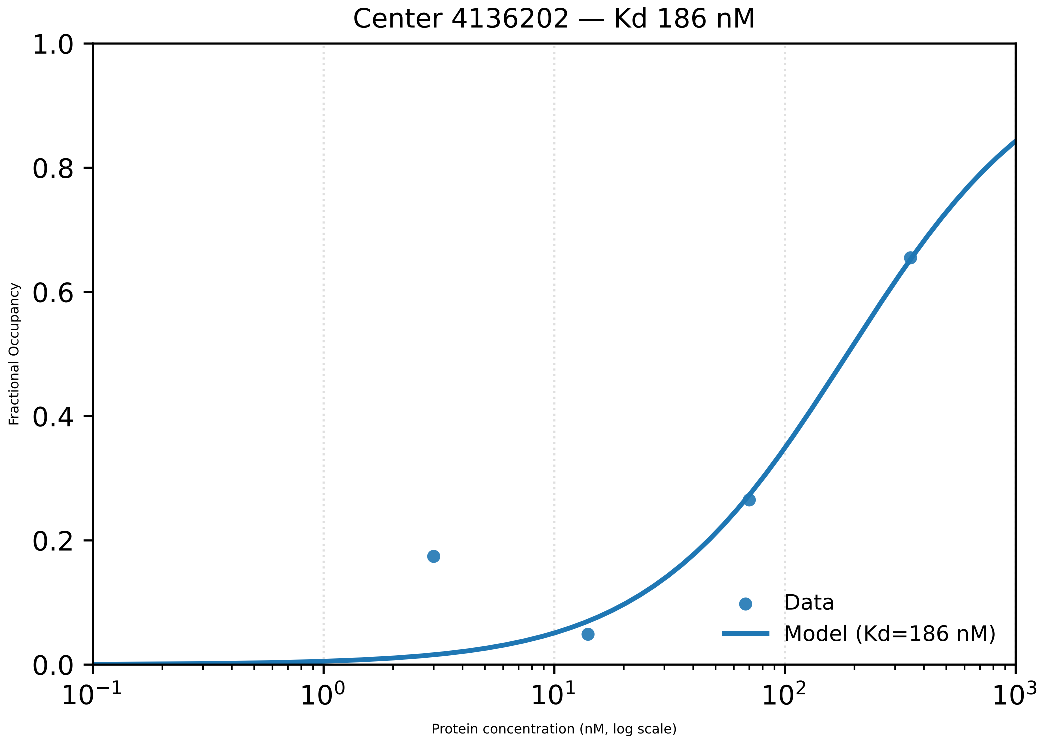
Center 4136202 — Kd 186 nM
Fractional Occupancy
Protein concentration (nM, log scale)
Data
Model (Kd=186 nM)
0.0
0.2
0.4
0.6
0.8
1.0
10−1
100
101
102
103

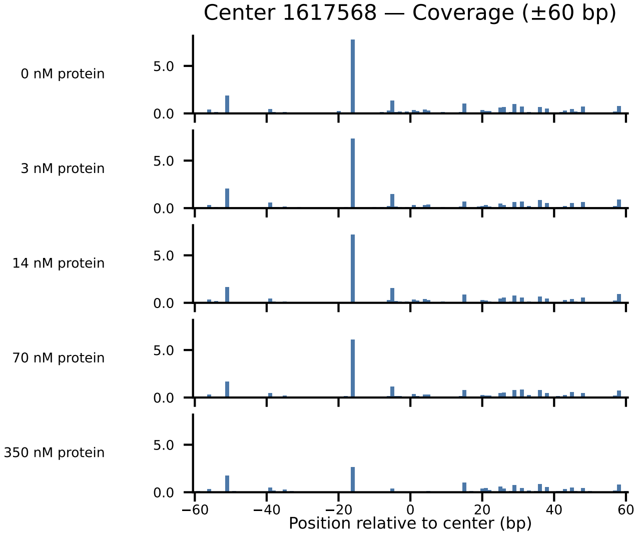

Center 1617568 — Coverage (±60 bp)
0 nM protein
3 nM protein
14 nM protein
70 nM protein
350 nM protein
5.0
0.0
−60
−40
−20
0
20
40
60
Position relative to center (bp)

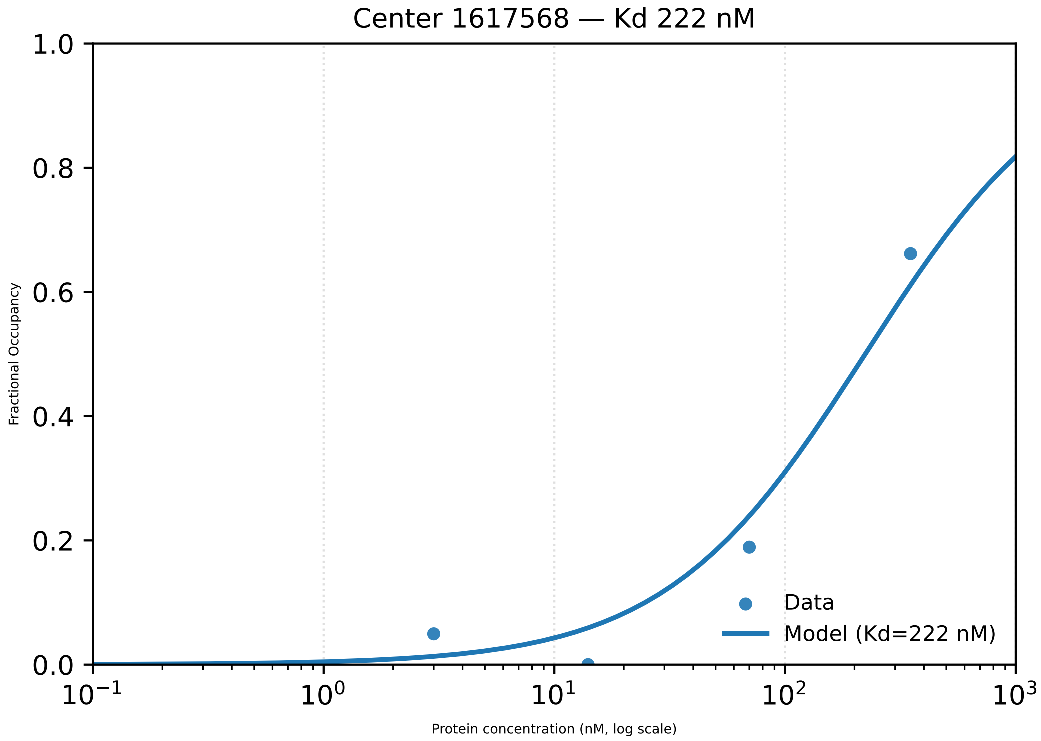

Center 1617568 — Kd 222 nM
Fractional Occupancy
1.0
0.8
0.6
0.4
0.2
0.0
$10^{-1}$
$10^{0}$
$10^{1}$
$10^{2}$
$10^{3}$
Protein concentration (nM, log scale)
Data
Model (Kd=222 nM)

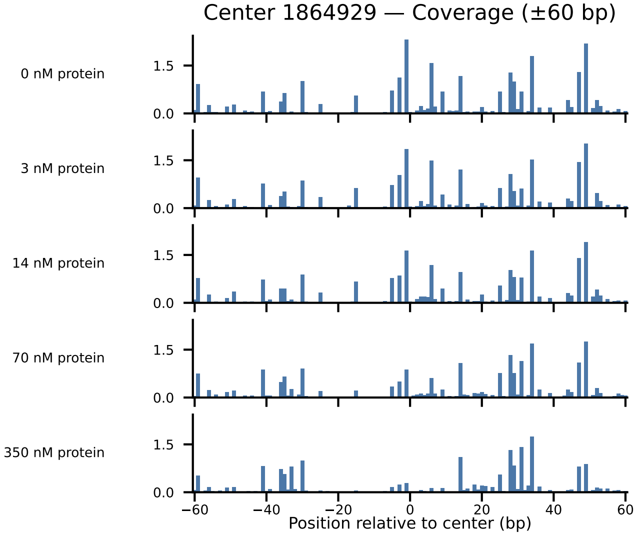

Center 1864929 — Coverage (±60 bp)
0 nM protein
3 nM protein
14 nM protein
70 nM protein
350 nM protein
1.5
0.0
−60
−40
−20
0
20
40
60
Position relative to center (bp)

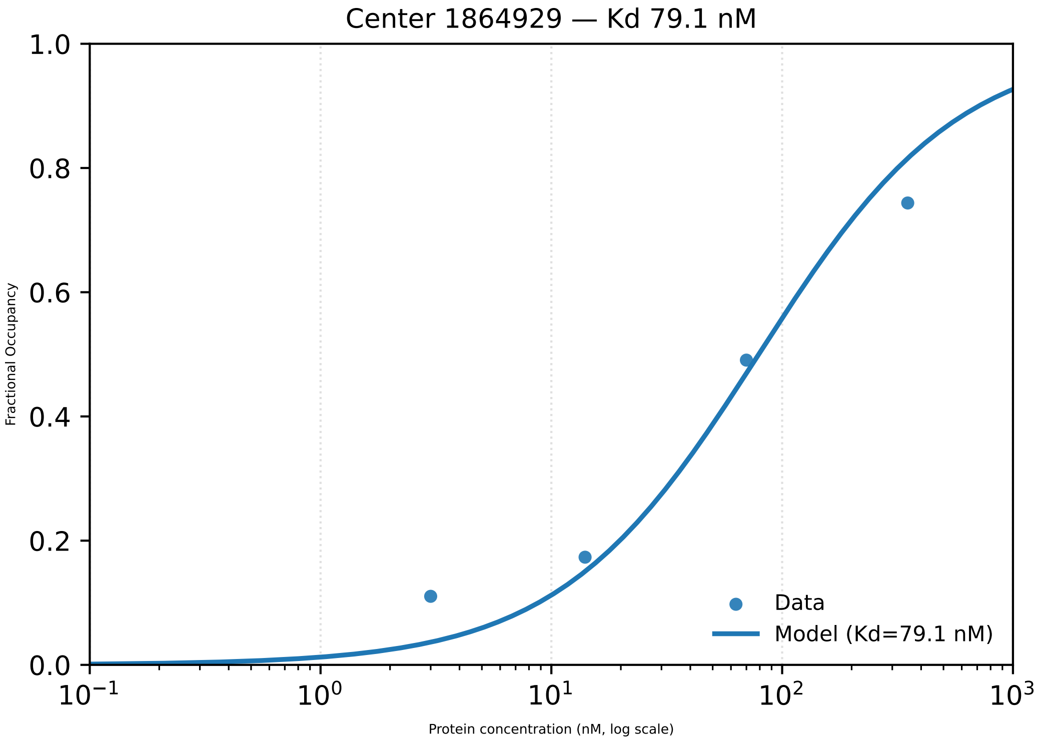
Center 1864929 — Kd 79.1 nM
1.0
0.8
0.6
0.4
0.2
0.0
Fractional Occupancy
$10^{-1}$
$10^{0}$
$10^{1}$
$10^{2}$
$10^{3}$
Protein concentration (nM, log scale)
Data
Model (Kd=79.1 nM)

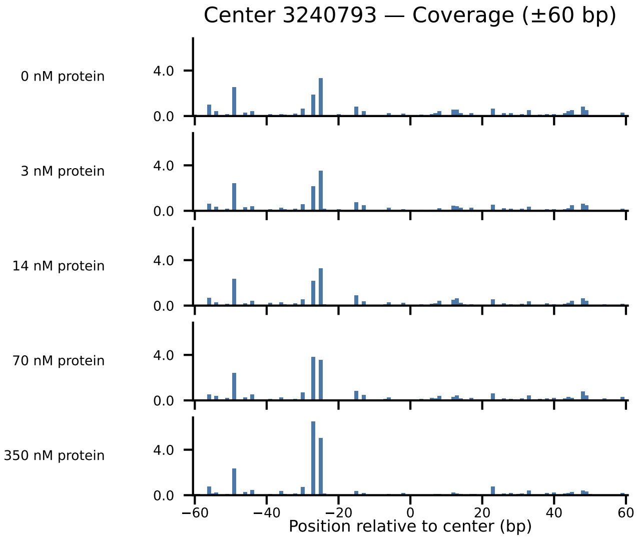
Center 3240793 — Coverage (±60 bp)
0 nM protein
3 nM protein
14 nM protein
70 nM protein
350 nM protein
4.0
0.0
−60
−40
−20
0
20
40
60
Position relative to center (bp)

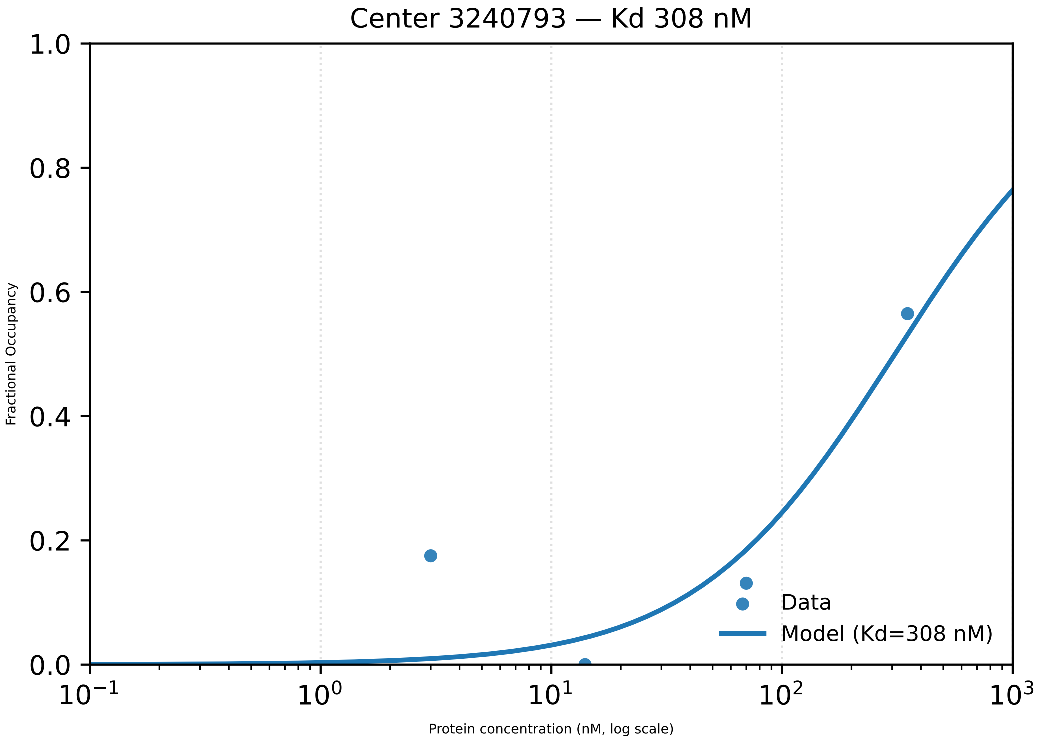

Center 3240793 — Kd 308 nM
1.0
0.8
0.6
0.4
0.2
0.0
Fractional Occupancy
10⁻¹
10⁰
10¹
10²
10³
Protein concentration (nM, log scale)
Data
Model (Kd=308 nM)

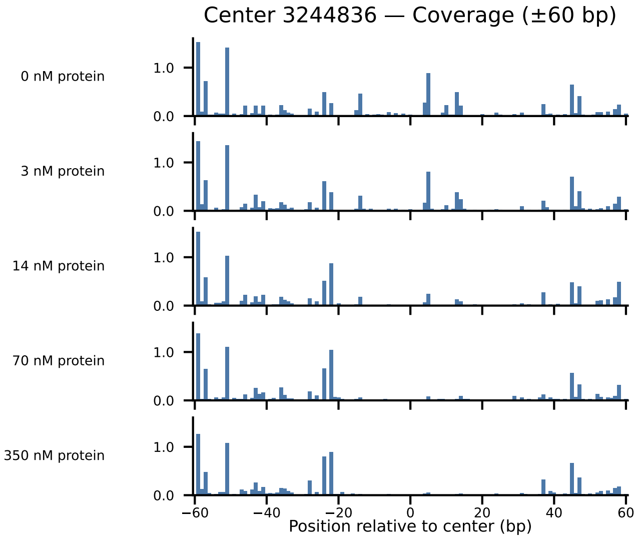

Center 3244836 — Coverage (±60 bp)
0 nM protein
3 nM protein
14 nM protein
70 nM protein
350 nM protein
0.0
1.0
−60
−40
−20
0
20
40
60
Position relative to center (bp)

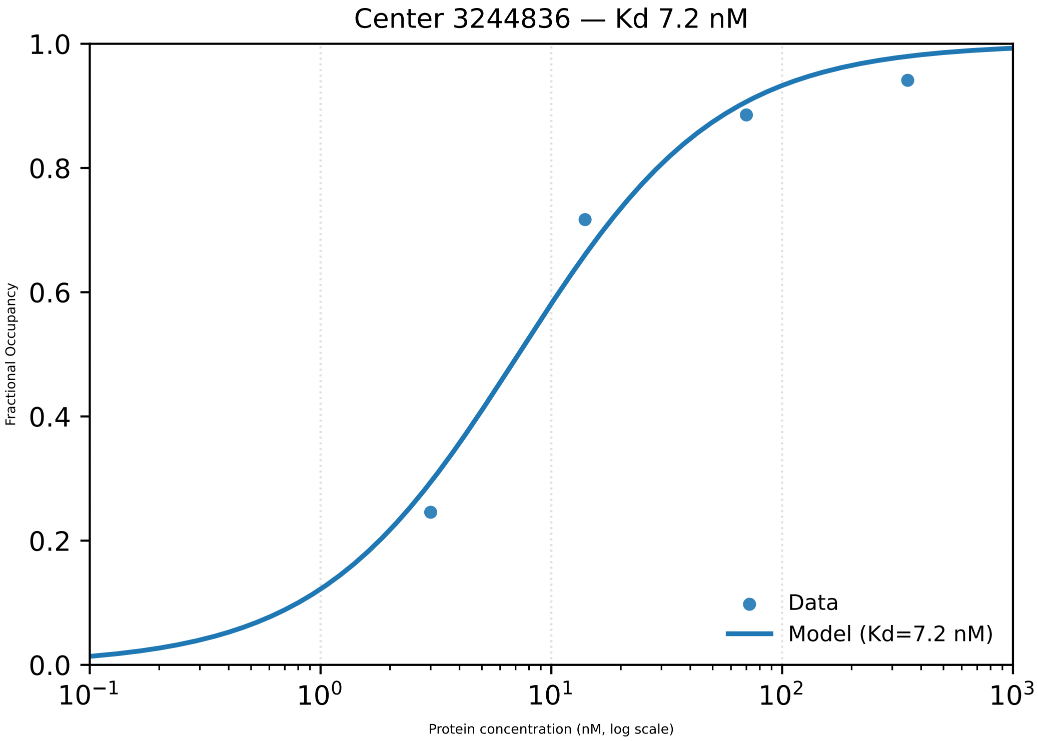
Center 3244836 — Kd 7.2 nM
1.0
0.8
0.6
0.4
0.2
0.0
Fractional Occupancy
$10^{-1}$
$10^{0}$
$10^{1}$
$10^{2}$
$10^{3}$
Protein concentration (nM, log scale)
Data
Model (Kd=7.2 nM)

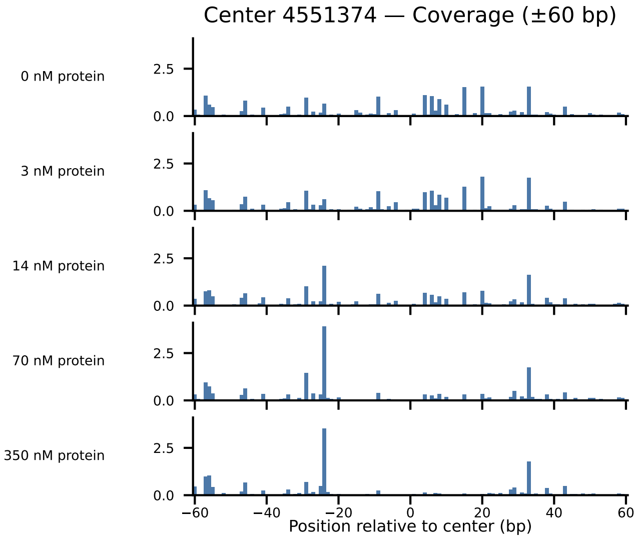
Center 4551374 — Coverage (±60 bp)
0 nM protein
3 nM protein
14 nM protein
70 nM protein
350 nM protein
2.5
0.0
−60
−40
−20
0
20
40
60
Position relative to center (bp)

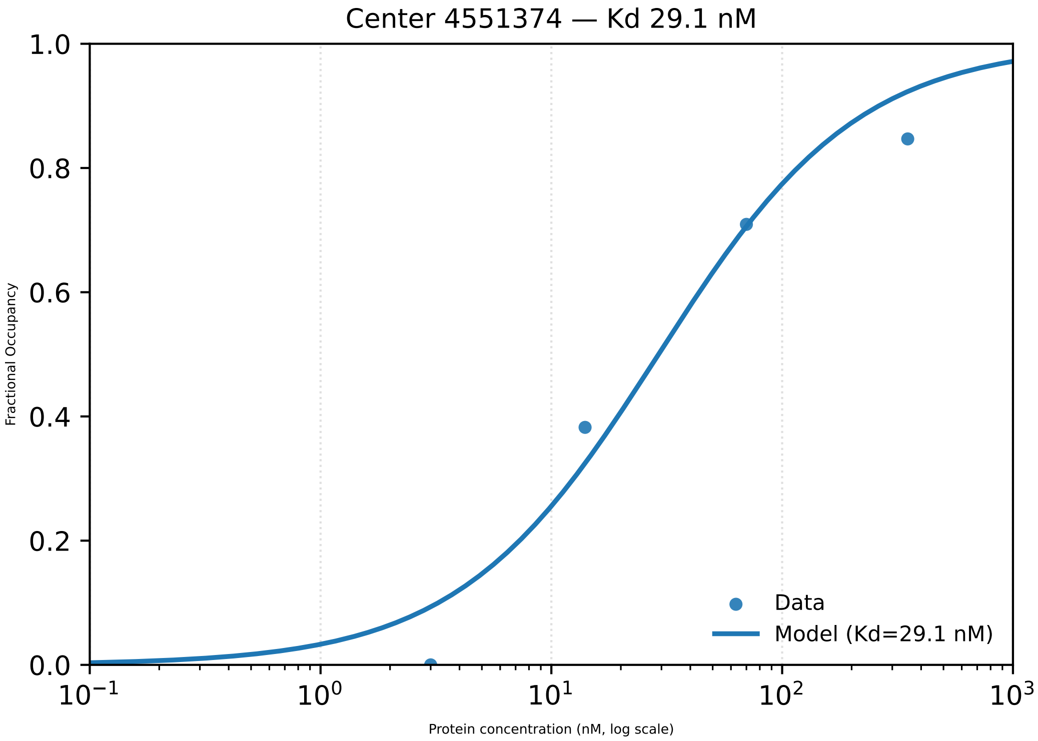

Center 4551374 — Kd 29.1 nM
1.0
0.8
0.6
0.4
0.2
0.0
Fractional Occupancy
$10^{-1}$
$10^{0}$
$10^{1}$
$10^{2}$
$10^{3}$
Protein concentration (nM, log scale)
Data
Model (Kd=29.1 nM)

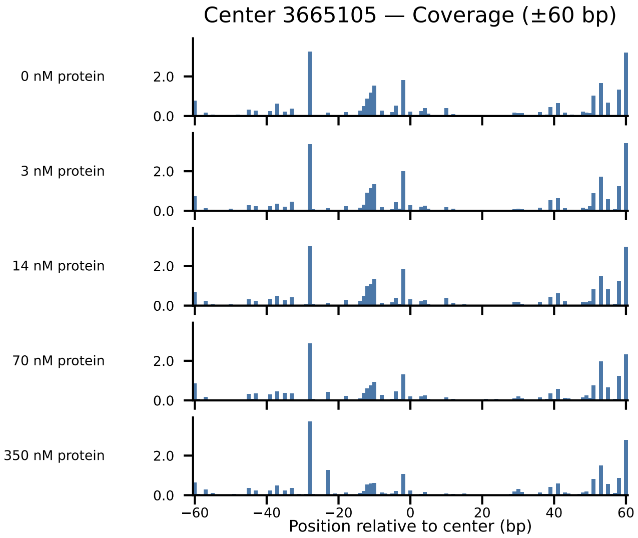

Center 3665105 — Coverage (±60 bp)
0 nM protein
3 nM protein
14 nM protein
70 nM protein
350 nM protein
2.0
0.0
−60
−40
−20
0
20
40
60
Position relative to center (bp)

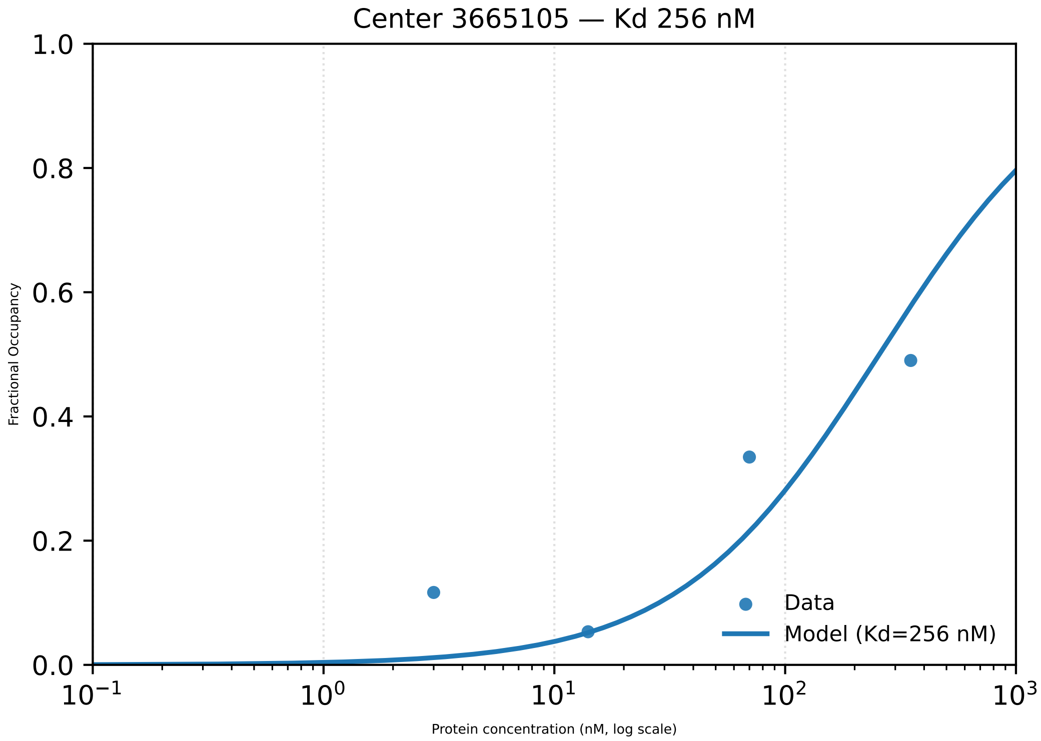

Center 3665105 — Kd 256 nM
Fractional Occupancy
1.0
0.8
0.6
0.4
0.2
0.0
$10^{-1}$
$10^{0}$
$10^{1}$
$10^{2}$
$10^{3}$
Protein concentration (nM, log scale)
Data
Model (Kd=256 nM)

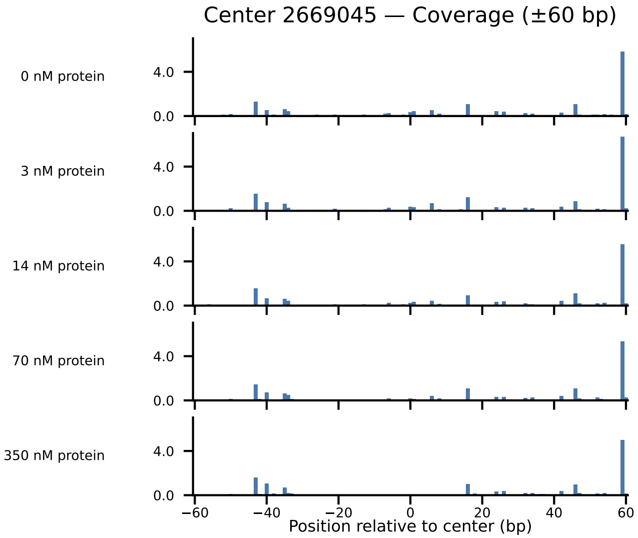

Center 2669045 — Coverage (±60 bp)
0 nM protein
3 nM protein
14 nM protein
70 nM protein
350 nM protein
4.0
0.0
−60
−40
−20
0
20
40
60
Position relative to center (bp)

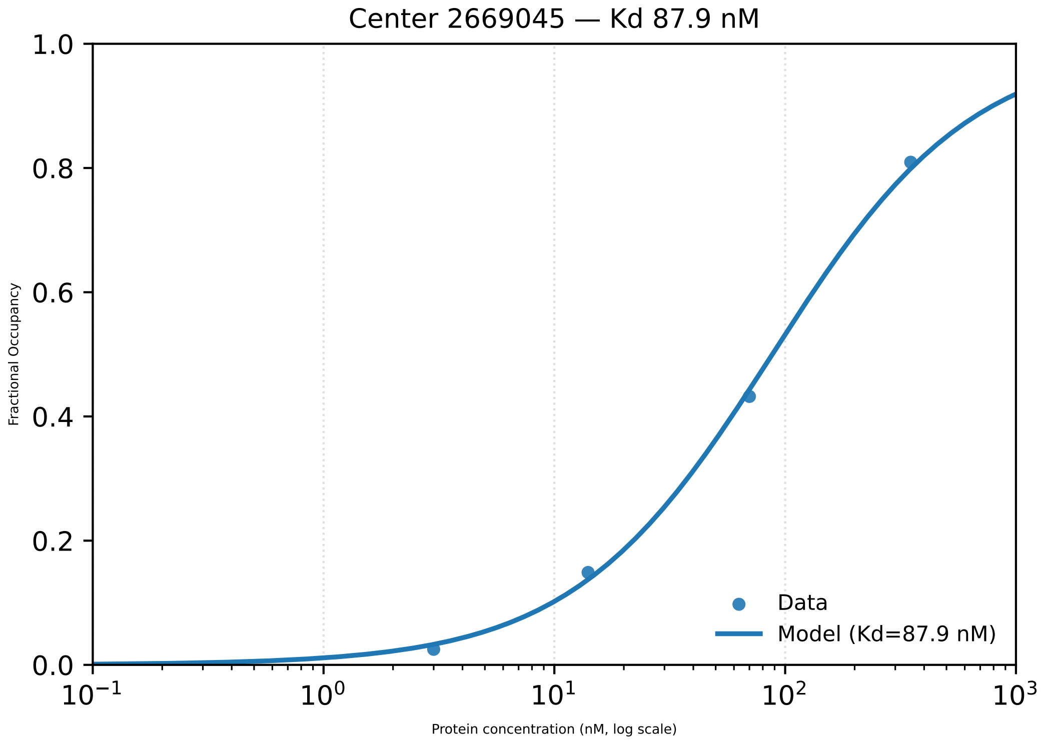

Center 2669045 — Kd 87.9 nM
Fractional Occupancy
1.0
0.8
0.6
0.4
0.2
0.0
$10^{-1}$
$10^{0}$
$10^{1}$
$10^{2}$
$10^{3}$
Protein concentration (nM, log scale)
Data
Model (Kd=87.9 nM)

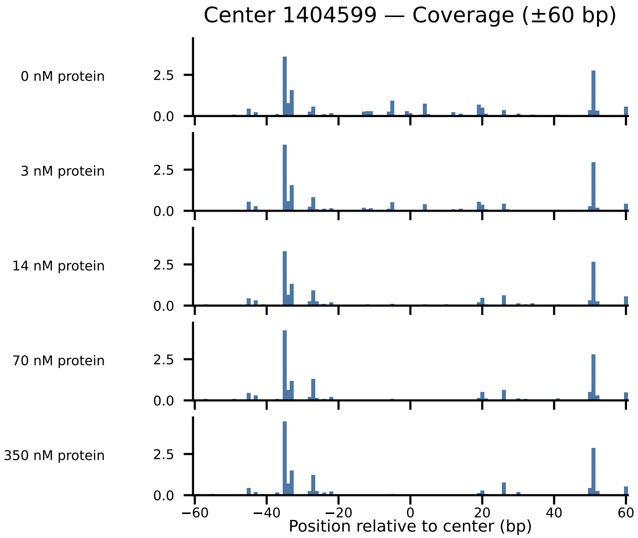

Center 1404599 — Coverage (±60 bp)
0 nM protein
3 nM protein
14 nM protein
70 nM protein
350 nM protein
2.5
0.0
−60
−40
−20
0
20
40
60
Position relative to center (bp)

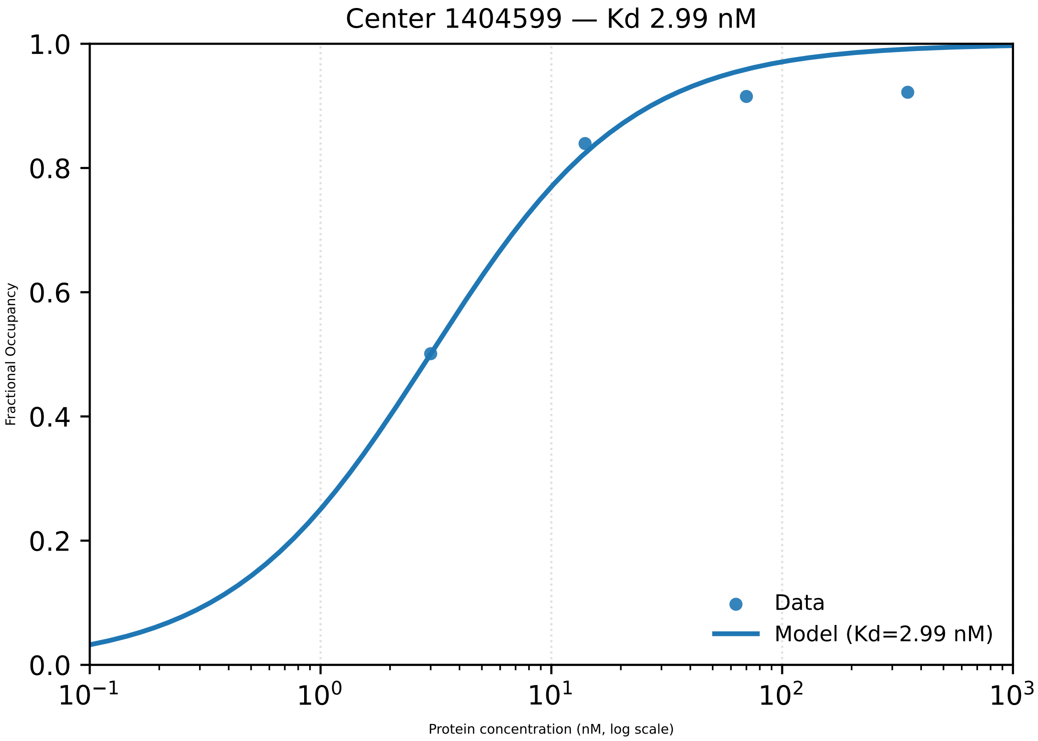

Center 1404599 — Kd 2.99 nM
Fractional Occupancy
Protein concentration (nM, log scale)
Data
Model (Kd=2.99 nM)
1.0
0.8
0.6
0.4
0.2
0.0
$10^{-1}$
$10^{0}$
$10^{1}$
$10^{2}$
$10^{3}$

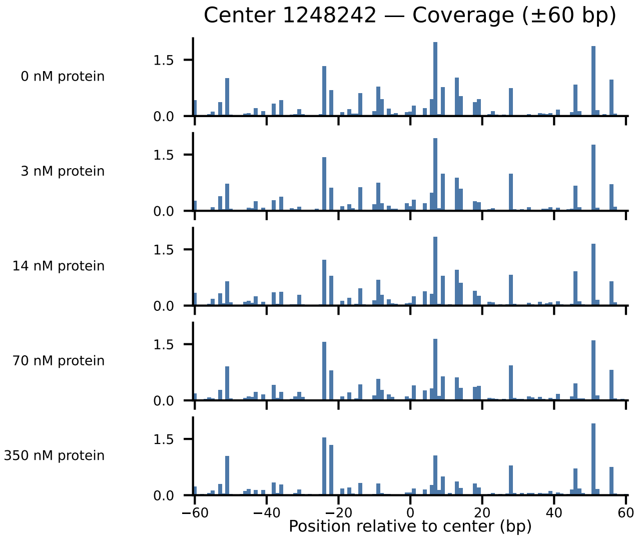
Center 1248242 — Coverage (±60 bp)
0 nM protein
3 nM protein
14 nM protein
70 nM protein
350 nM protein
1.5
0.0
−60
−40
−20
0
20
40
60
Position relative to center (bp)

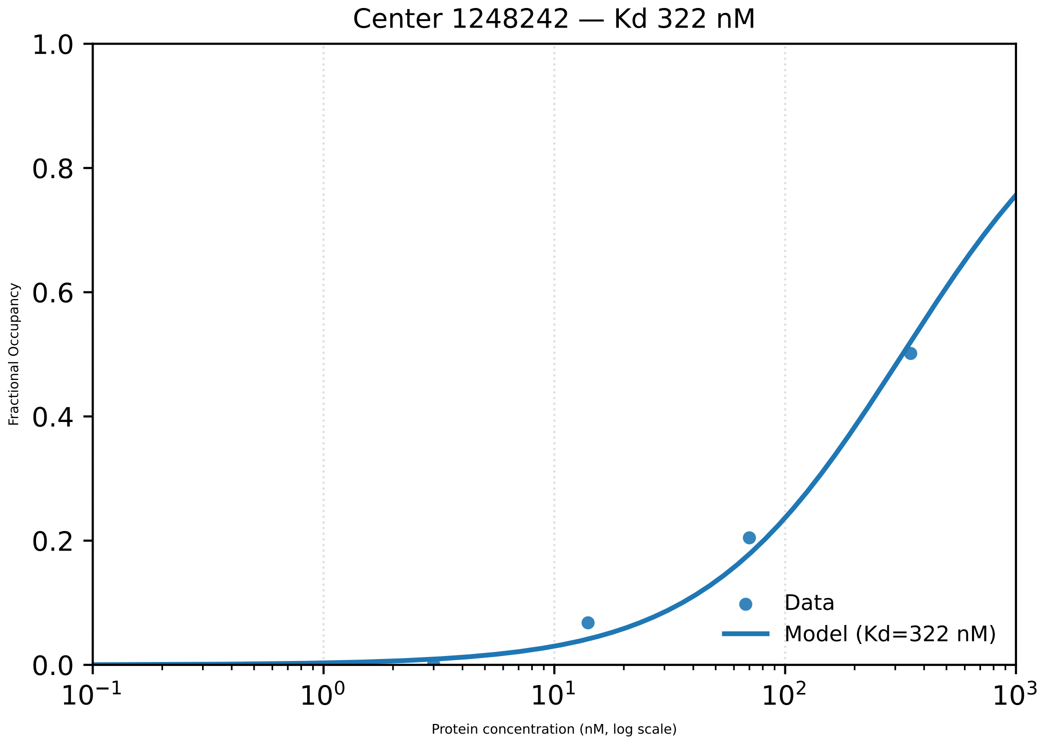
Center 1248242 — Kd 322 nM
1.0
0.8
0.6
0.4
0.2
0.0
Fractional Occupancy
$10^{-1}$
$10^{0}$
$10^{1}$
$10^{2}$
$10^{3}$
Protein concentration (nM, log scale)
Data
Model (Kd=322 nM)

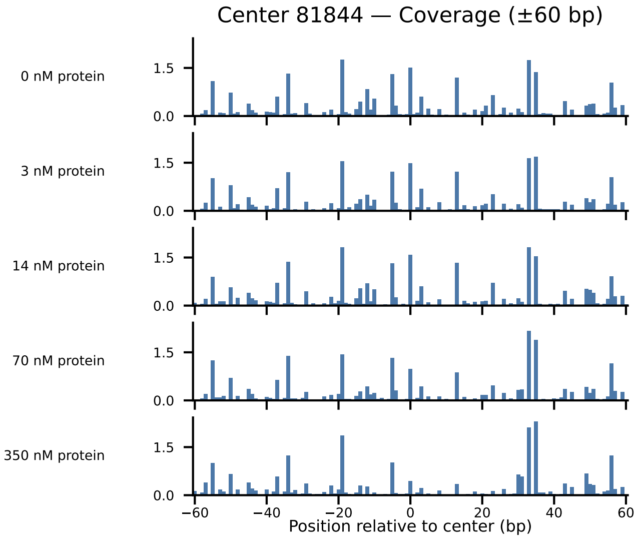

Center 81844 — Coverage (±60 bp)
0 nM protein
3 nM protein
14 nM protein
70 nM protein
350 nM protein
1.5
0.0
−60
−40
−20
0
20
40
60
Position relative to center (bp)

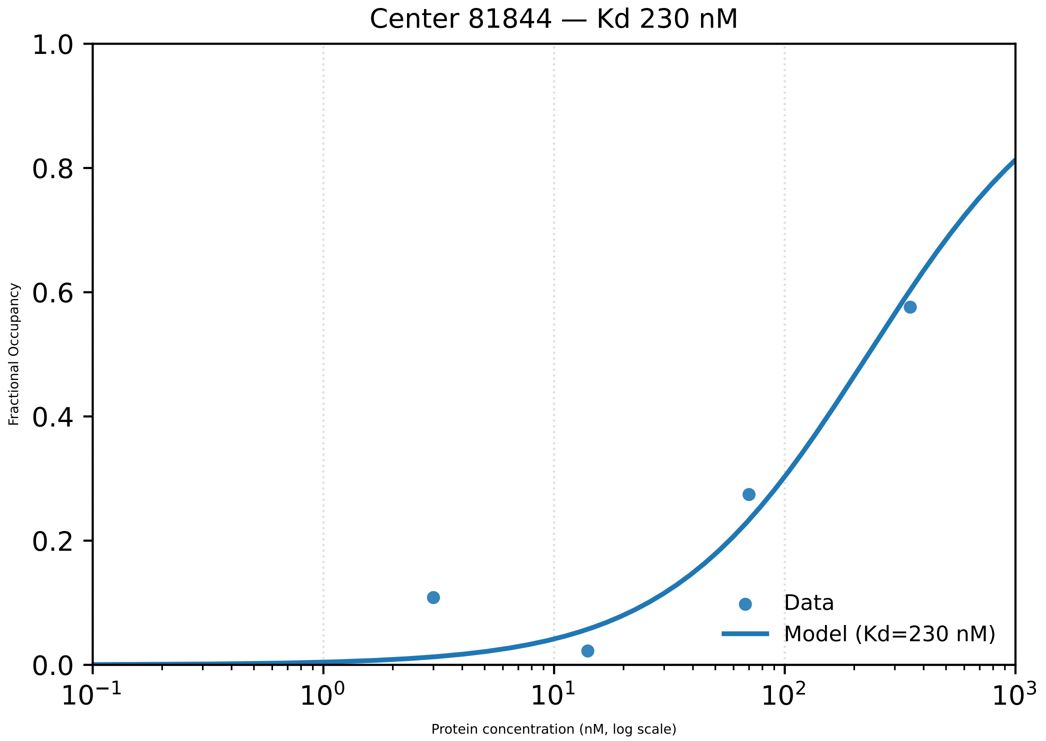

Center 81844 — Kd 230 nM
1.0
0.8
0.6
0.4
0.2
0.0
Fractional Occupancy
$10^{-1}$
$10^{0}$
$10^{1}$
$10^{2}$
$10^{3}$
Protein concentration (nM, log scale)
Data
Model (Kd=230 nM)

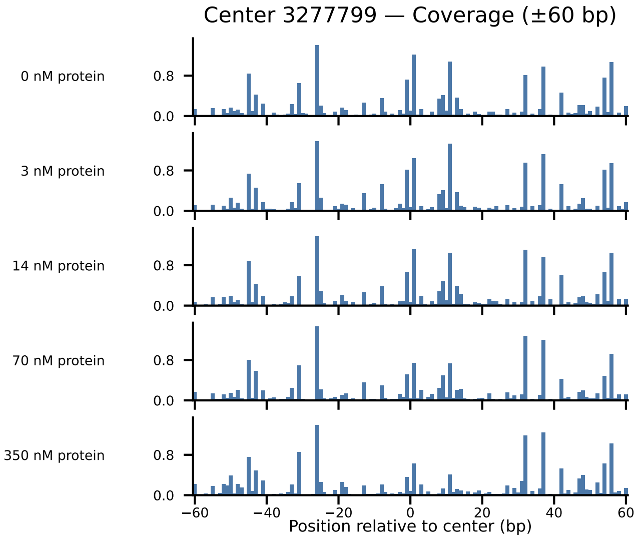
Center 3277799 — Coverage (±60 bp)
0 nM protein
3 nM protein
14 nM protein
70 nM protein
350 nM protein
0.8
0.0
−60
−40
−20
0
20
40
60
Position relative to center (bp)

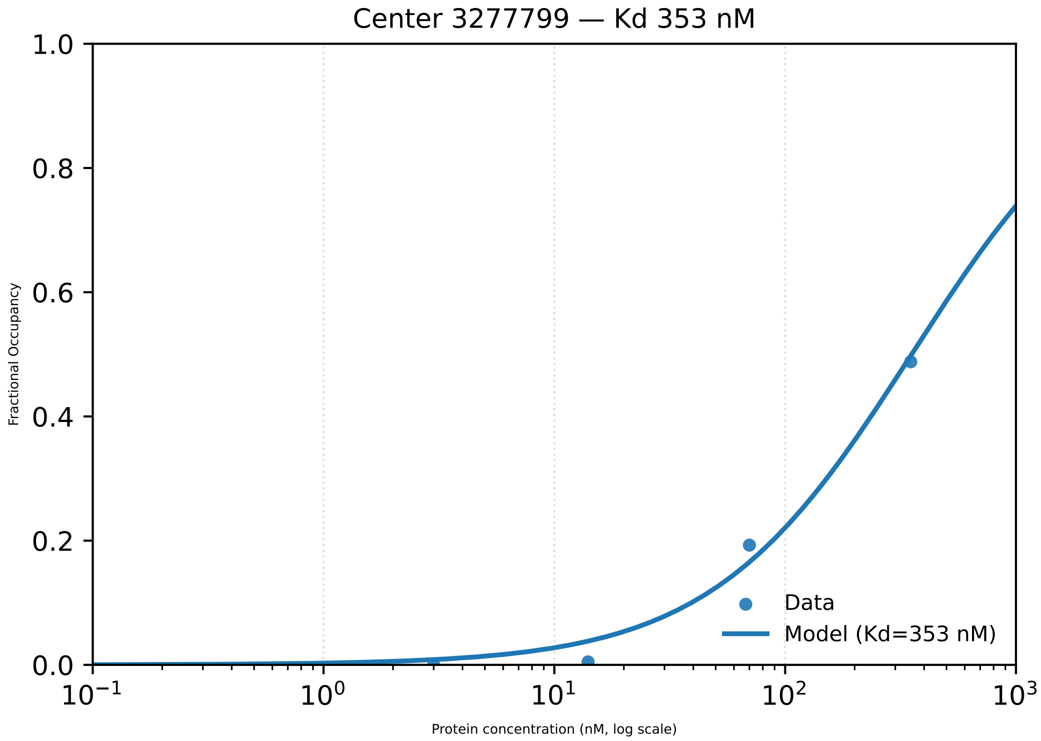

Center 3277799 — Kd 353 nM
1.0
0.8
0.6
0.4
0.2
0.0
Fractional Occupancy
$10^{-1}$
$10^{0}$
$10^{1}$
$10^{2}$
$10^{3}$
Protein concentration (nM, log scale)
Data
Model (Kd=353 nM)

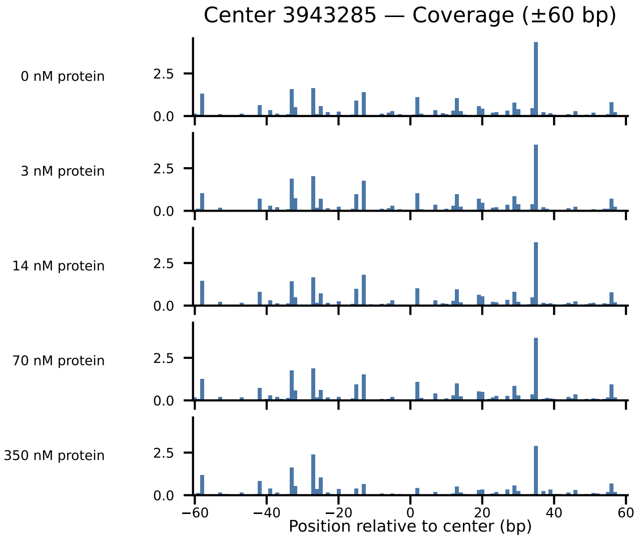
Center 3943285 — Coverage (±60 bp)
0 nM protein
3 nM protein
14 nM protein
70 nM protein
350 nM protein
2.5
0.0
−60
−40
−20
0
20
40
60
Position relative to center (bp)

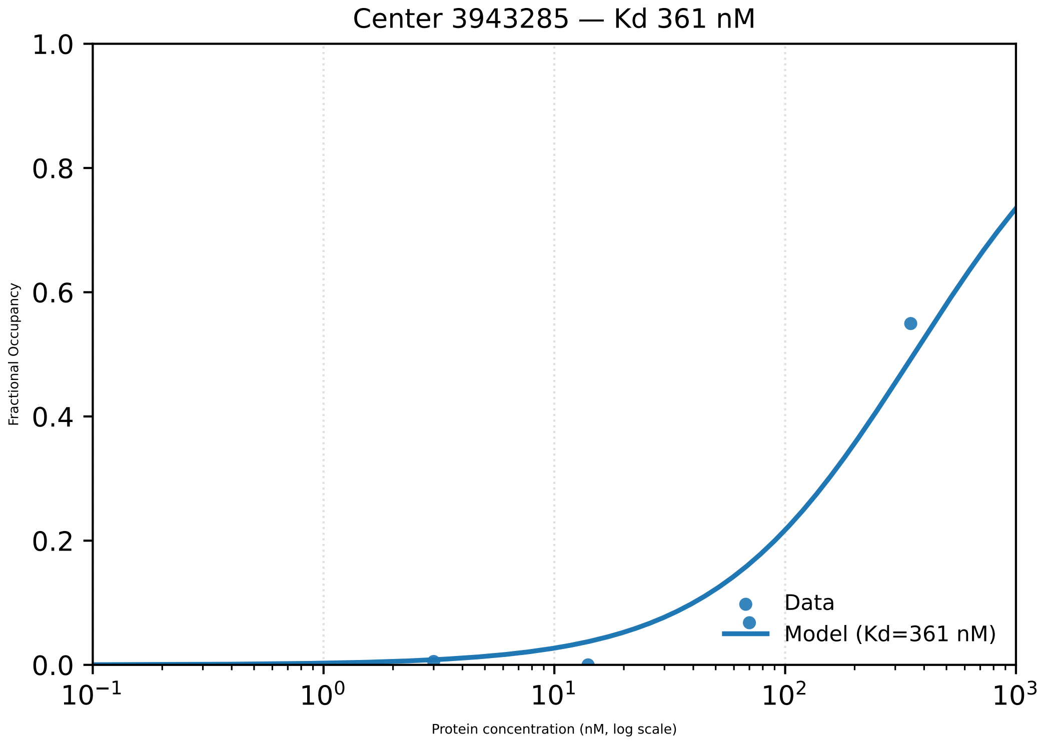

Center 3943285 — Kd 361 nM
1.0
0.8
0.6
0.4
0.2
0.0
Fractional Occupancy
$10^{-1}$
$10^{0}$
$10^{1}$
$10^{2}$
$10^{3}$
Protein concentration (nM, log scale)
Data
Model (Kd=361 nM)

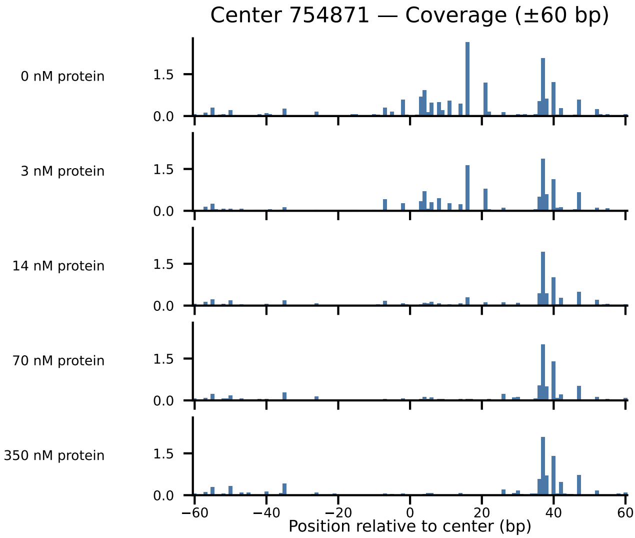

Center 754871 — Coverage (±60 bp)
0 nM protein
3 nM protein
14 nM protein
70 nM protein
350 nM protein
1.5
0.0
−60
−40
−20
0
20
40
60
Position relative to center (bp)

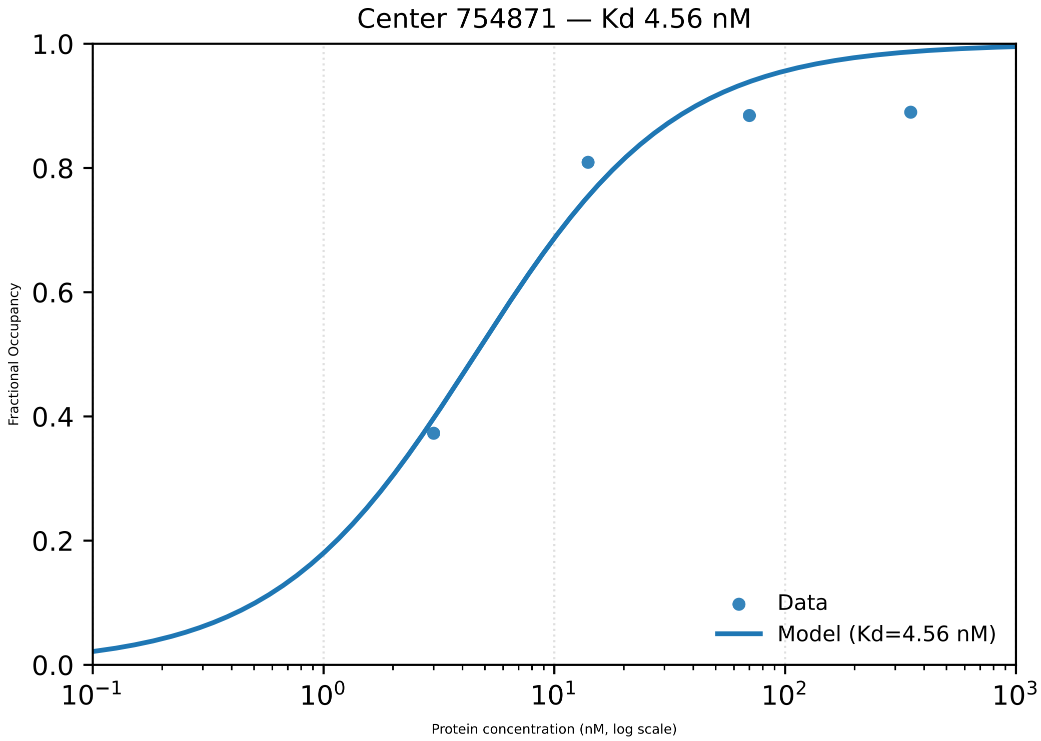

Center 754871 — Kd 4.56 nM
1.0
0.8
0.6
0.4
0.2
0.0
Fractional Occupancy
$10^{-1}$
$10^{0}$
$10^{1}$
$10^{2}$
$10^{3}$
Protein concentration (nM, log scale)
Data
Model (Kd=4.56 nM)

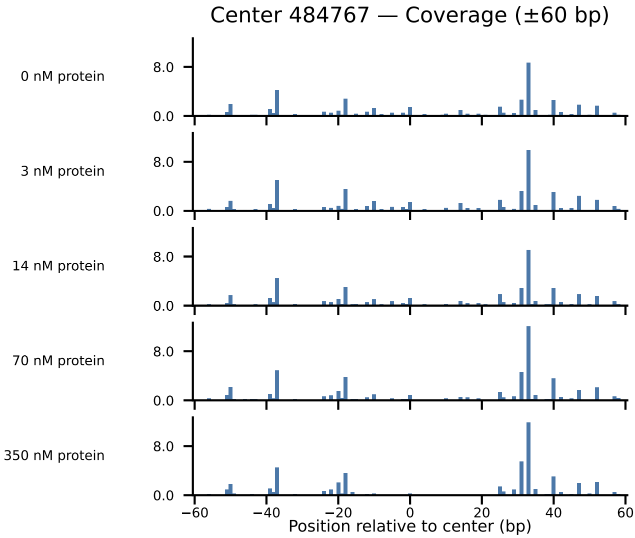
Center 484767 — Coverage (±60 bp)
0 nM protein
3 nM protein
14 nM protein
70 nM protein
350 nM protein
8.0
0.0
−60
−40
−20
0
20
40
60
Position relative to center (bp)

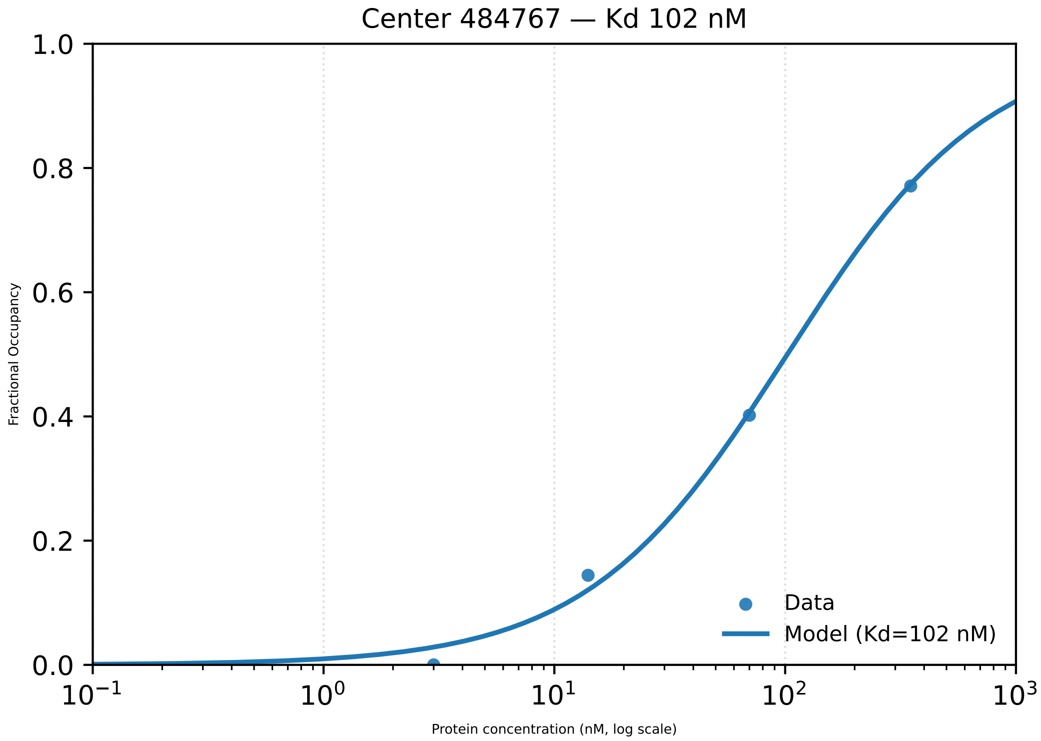

Center 484767 — Kd 102 nM
Fractional Occupancy
1.0
0.8
0.6
0.4
0.2
0.0
$10^{-1}$
$10^{0}$
$10^{1}$
$10^{2}$
$10^{3}$
Protein concentration (nM, log scale)
Data
Model (Kd=102 nM)

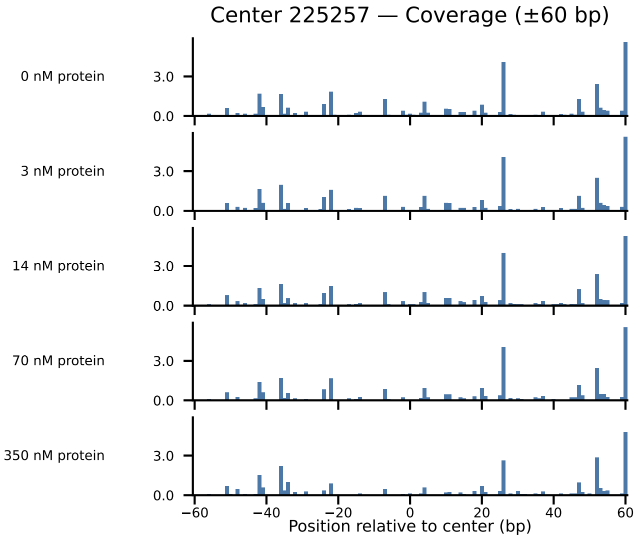

Center 225257 — Coverage (±60 bp)
0 nM protein
3 nM protein
14 nM protein
70 nM protein
350 nM protein
3.0
0.0
−60
−40
−20
0
20
40
60
Position relative to center (bp)

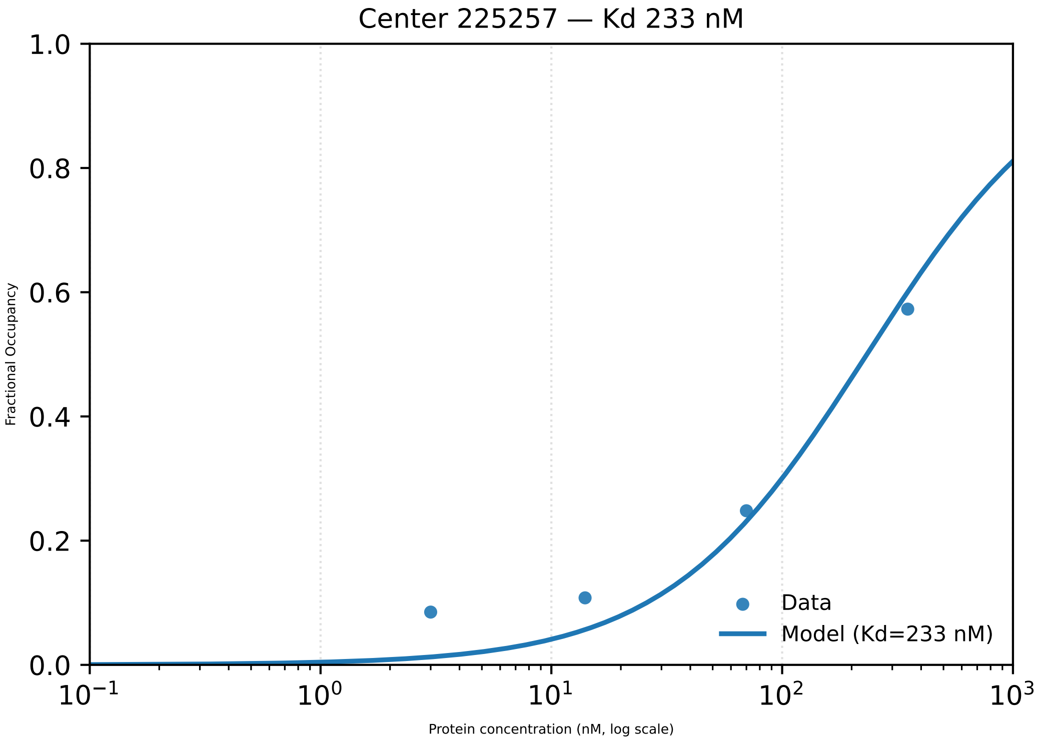

Center 225257 — Kd 233 nM
1.0
0.8
0.6
0.4
0.2
0.0
Fractional Occupancy
$10^{-1}$
$10^{0}$
$10^{1}$
$10^{2}$
$10^{3}$
Protein concentration (nM, log scale)
Data
Model (Kd=233 nM)

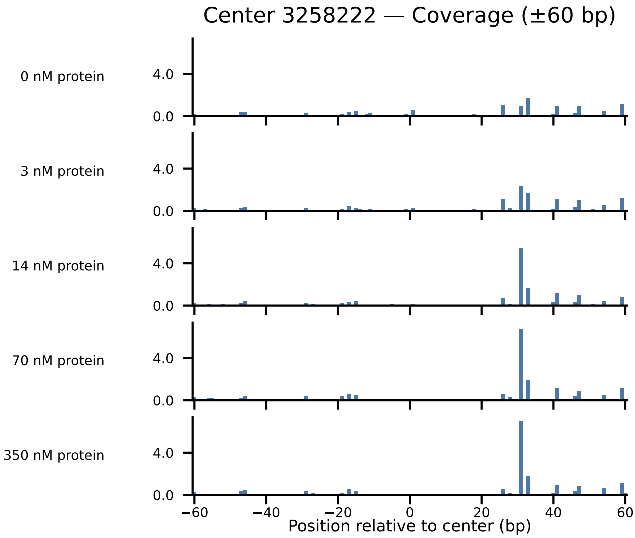
Center 3258222 — Coverage (±60 bp)
0 nM protein
3 nM protein
14 nM protein
70 nM protein
350 nM protein
4.0
0.0
−60
−40
−20
0
20
40
60
Position relative to center (bp)

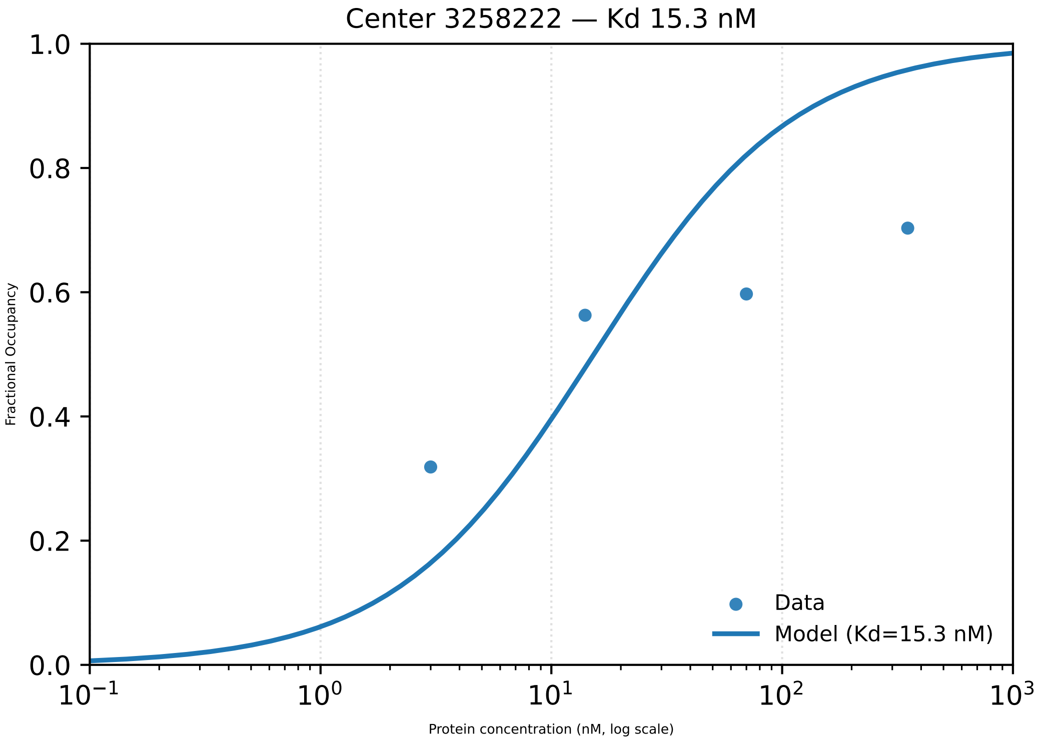

Center 3258222 — Kd 15.3 nM
Fractional Occupancy
Protein concentration (nM, log scale)
1.0
0.8
0.6
0.4
0.2
0.0
$10^{-1}$
$10^{0}$
$10^{1}$
$10^{2}$
$10^{3}$
Data
Model (Kd=15.3 nM)

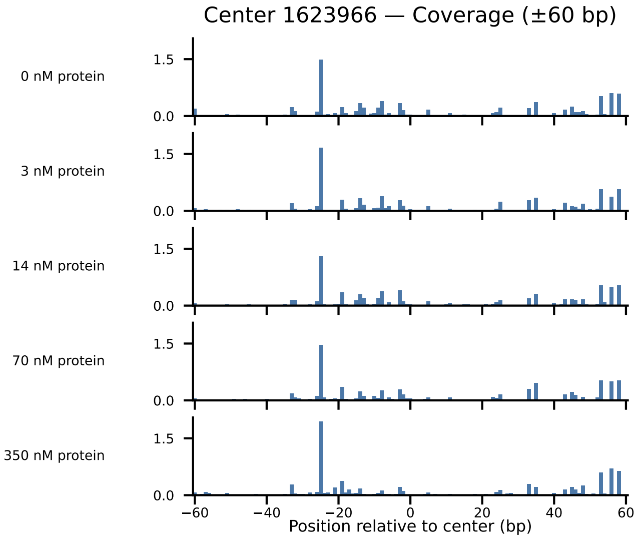
Center 1623966 — Coverage (±60 bp)
0 nM protein
3 nM protein
14 nM protein
70 nM protein
350 nM protein
1.5
0.0
−60
−40
−20
0
20
40
60
Position relative to center (bp)

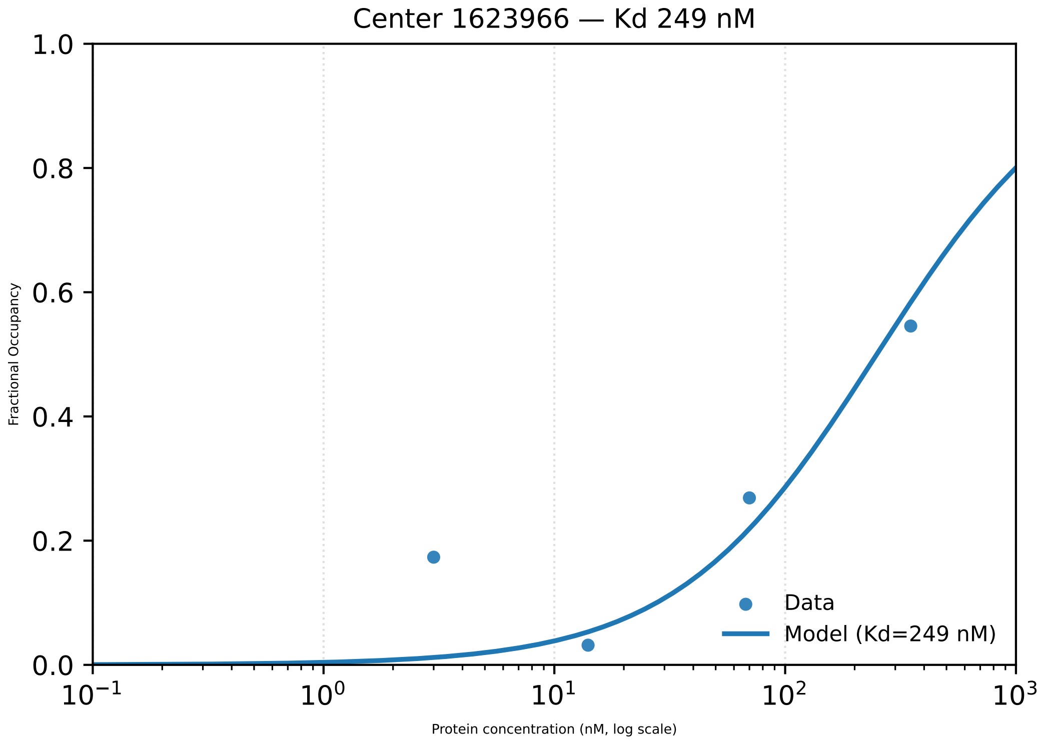

Center 1623966 — Kd 249 nM
Fractional Occupancy
Protein concentration (nM, log scale)
1.0
0.8
0.6
0.4
0.2
0.0
$10^{-1}$
$10^{0}$
$10^{1}$
$10^{2}$
$10^{3}$
Data
Model (Kd=249 nM)

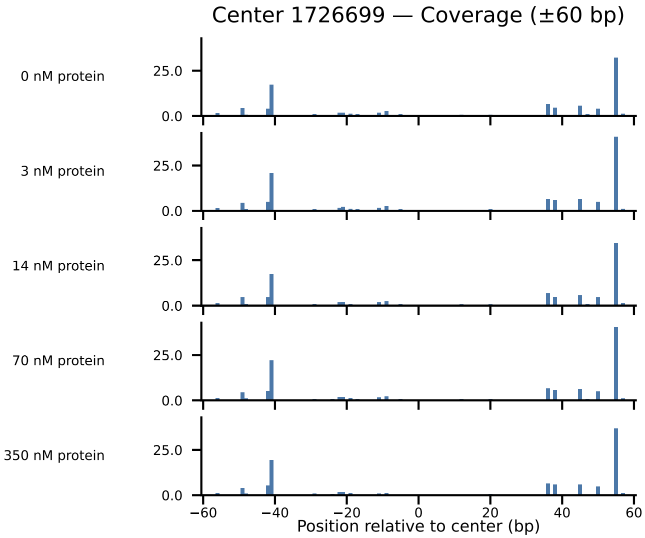

Center 1726699 — Coverage (±60 bp)
0 nM protein
3 nM protein
14 nM protein
70 nM protein
350 nM protein
25.0
0.0
−60
−40
−20
0
20
40
60
Position relative to center (bp)

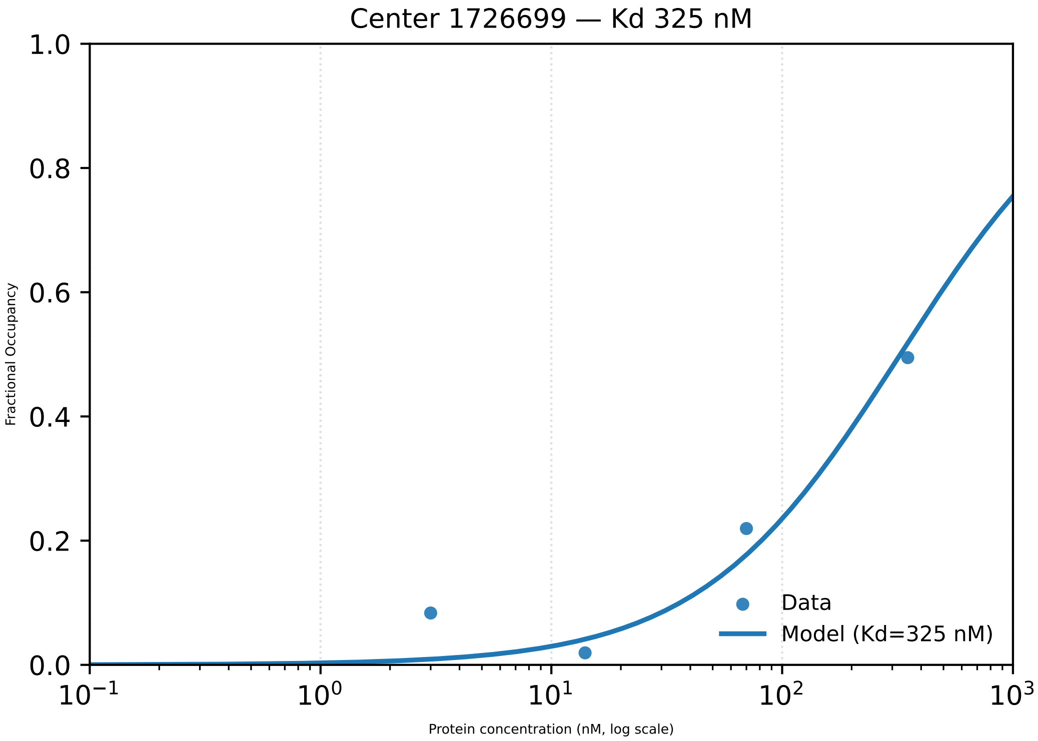

Center 1726699 — Kd 325 nM
Fractional Occupancy
1.0
0.8
0.6
0.4
0.2
0.0
$10^{-1}$
$10^{0}$
$10^{1}$
$10^{2}$
$10^{3}$
Protein concentration (nM, log scale)
Data
Model (Kd=325 nM)

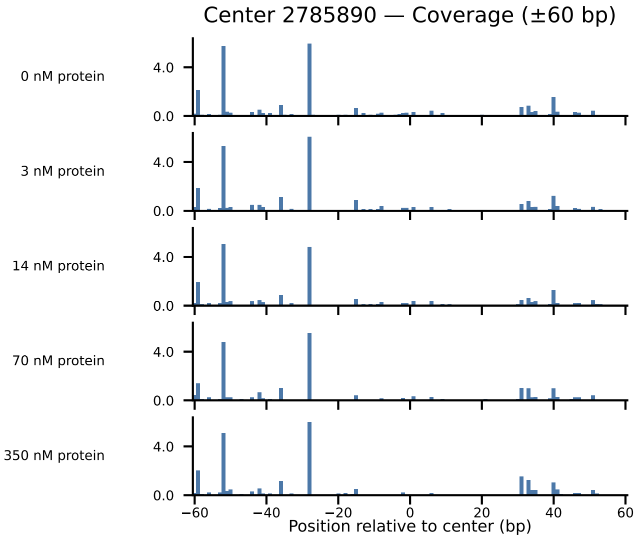

Center 2785890 — Coverage (±60 bp)
0 nM protein
3 nM protein
14 nM protein
70 nM protein
350 nM protein
4.0
0.0
−60
−40
−20
0
20
40
60
Position relative to center (bp)

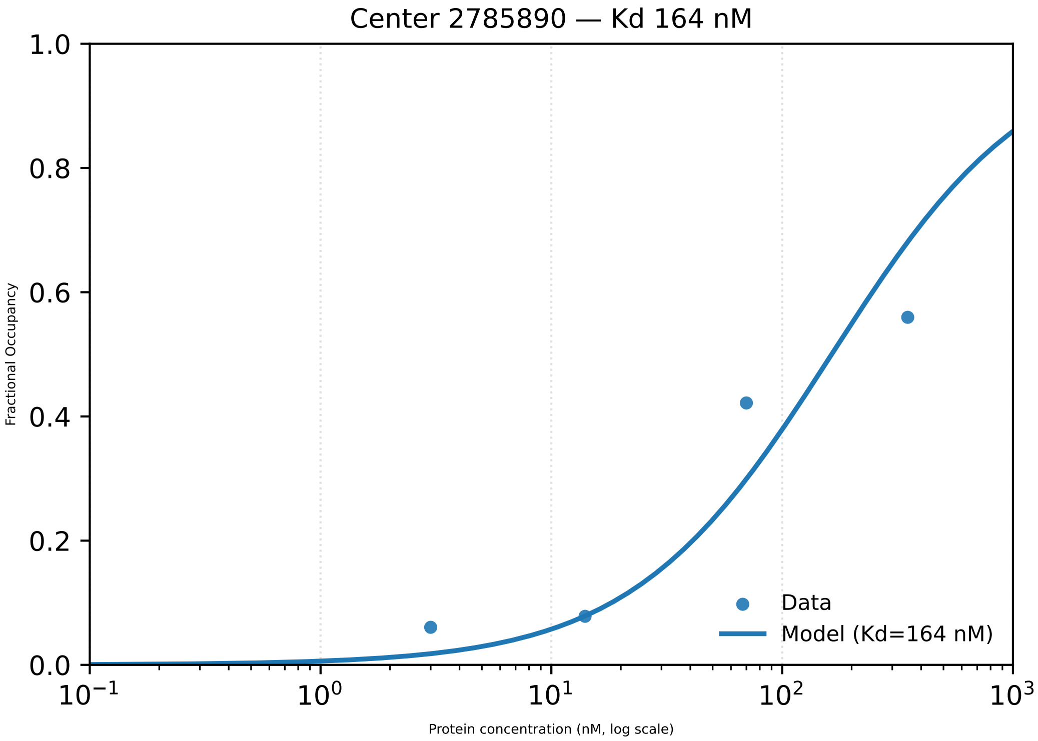

Center 2785890 — Kd 164 nM
Fractional Occupancy
Protein concentration (nM, log scale)
Data
Model (Kd=164 nM)
0.0
0.2
0.4
0.6
0.8
1.0
$10^{-1}$
$10^{0}$
$10^{1}$
$10^{2}$
$10^{3}$

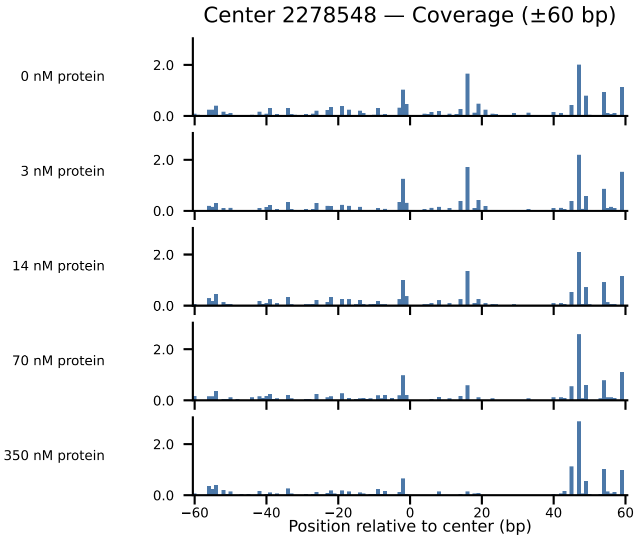
Center 2278548 — Coverage (±60 bp)
0 nM protein
3 nM protein
14 nM protein
70 nM protein
350 nM protein
2.0
0.0
−60
−40
−20
0
20
40
60
Position relative to center (bp)

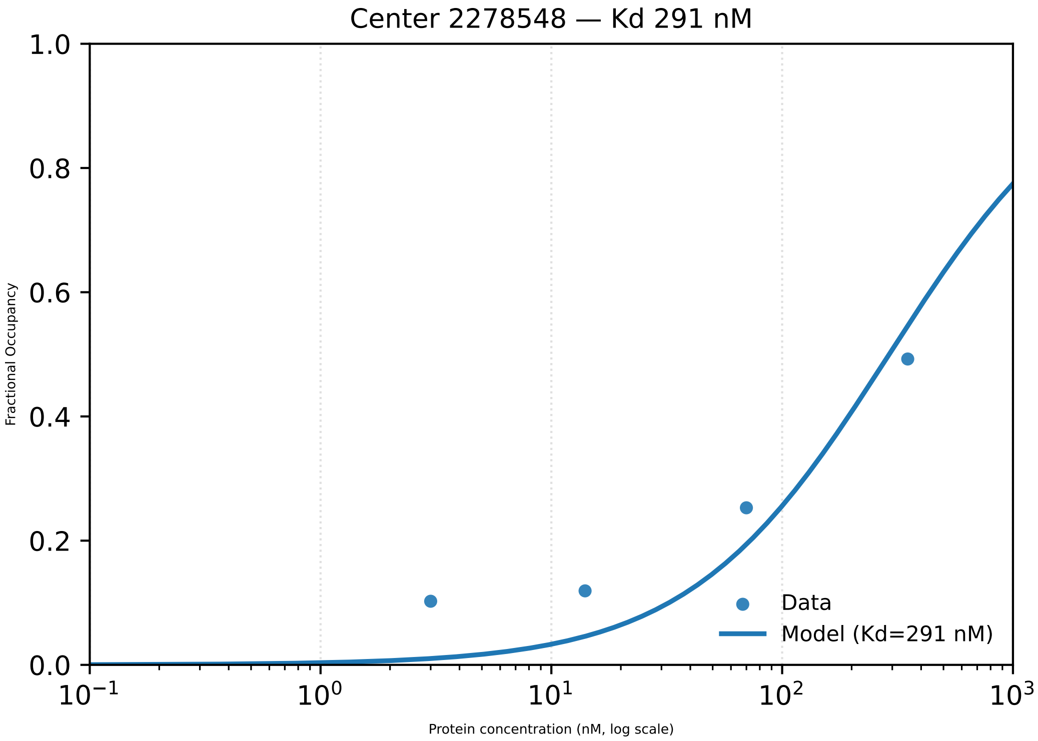
Center 2278548 — Kd 291 nM
Fractional Occupancy
Protein concentration (nM, log scale)
Data
Model (Kd=291 nM)
1.0
0.8
0.6
0.4
0.2
0.0
$10^{-1}$
$10^{0}$
$10^{1}$
$10^{2}$
$10^{3}$

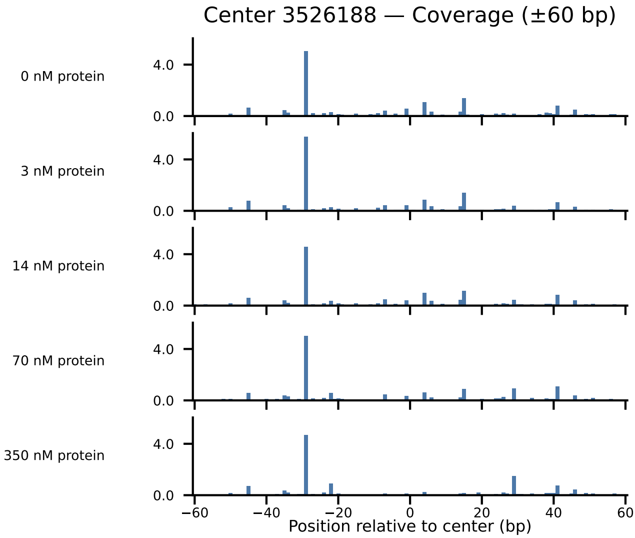

Center 3526188 — Coverage (±60 bp)
0 nM protein
3 nM protein
14 nM protein
70 nM protein
350 nM protein
4.0
0.0
−60
−40
−20
0
20
40
60
Position relative to center (bp)

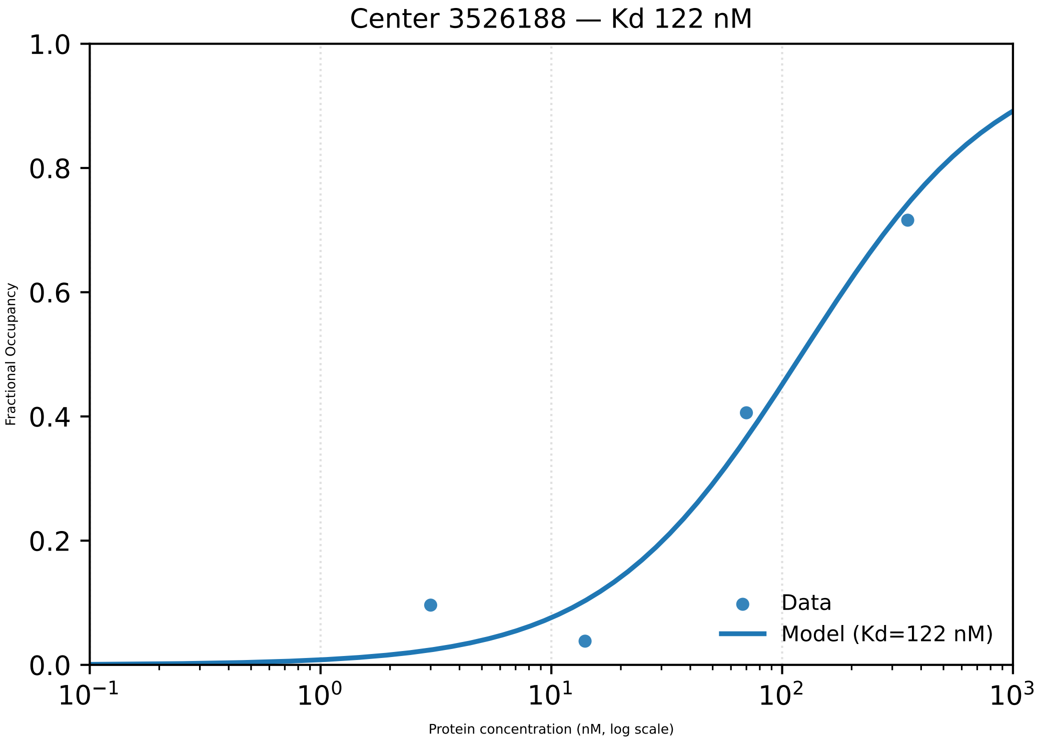

Center 3526188 — Kd 122 nM
Fractional Occupancy
1.0
0.8
0.6
0.4
0.2
0.0
$10^{-1}$
$10^{0}$
$10^{1}$
$10^{2}$
$10^{3}$
Protein concentration (nM, log scale)
Data
Model (Kd=122 nM)

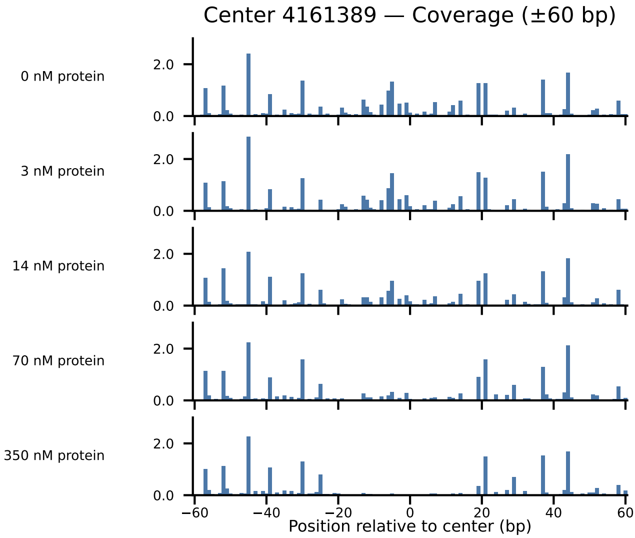

Center 4161389 — Coverage (±60 bp)
0 nM protein
3 nM protein
14 nM protein
70 nM protein
350 nM protein
2.0
0.0
−60
−40
−20
0
20
40
60
Position relative to center (bp)

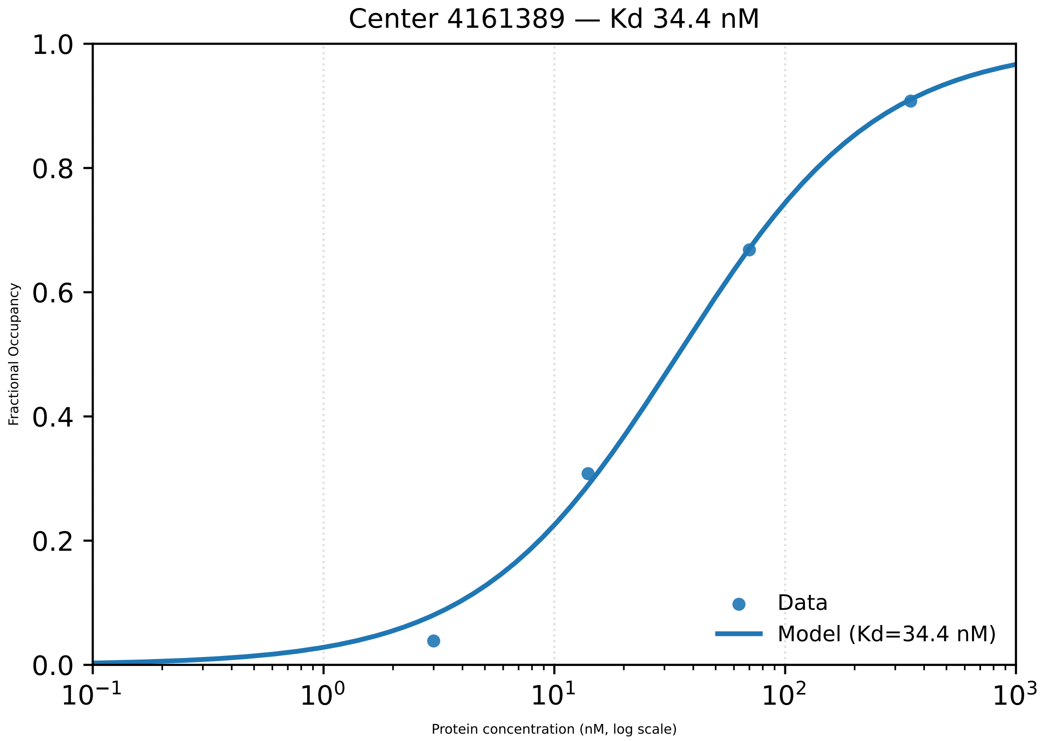

Center 4161389 — Kd 34.4 nM
Fractional Occupancy
Protein concentration (nM, log scale)
1.0
0.8
0.6
0.4
0.2
0.0
$10^{-1}$
$10^{0}$
$10^{1}$
$10^{2}$
$10^{3}$
Data
Model (Kd=34.4 nM)

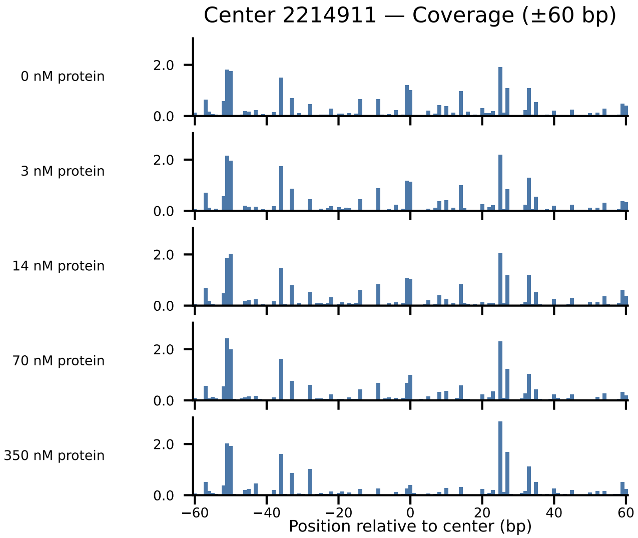

Center 2214911 — Coverage (±60 bp)
0 nM protein
3 nM protein
14 nM protein
70 nM protein
350 nM protein
2.0
0.0
−60
−40
−20
0
20
40
60
Position relative to center (bp)

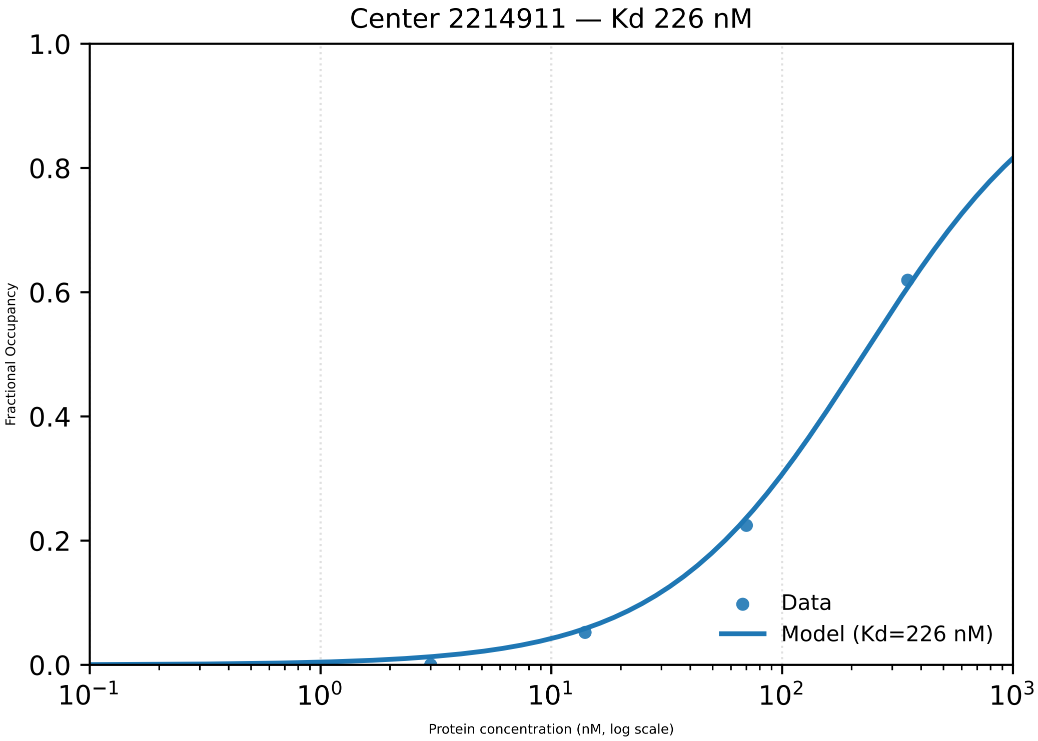

Center 2214911 — Kd 226 nM
1.0
0.8
0.6
0.4
0.2
0.0
Fractional Occupancy
$10^{-1}$
$10^{0}$
$10^{1}$
$10^{2}$
$10^{3}$
Protein concentration (nM, log scale)
Data
Model (Kd=226 nM)

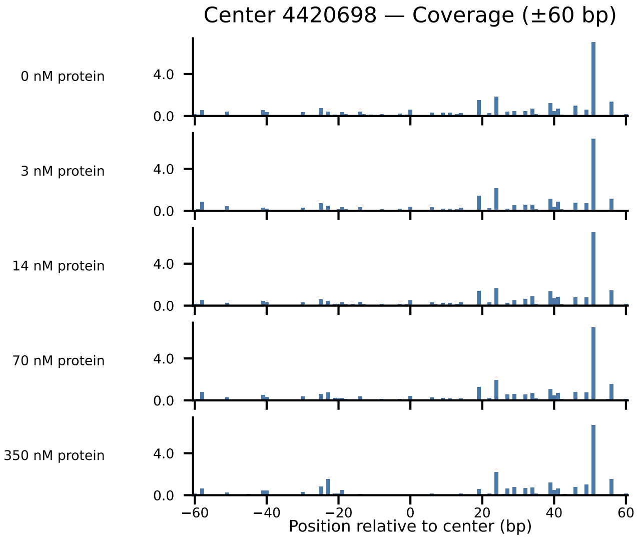
Center 4420698 — Coverage (±60 bp)
0 nM protein
3 nM protein
14 nM protein
70 nM protein
350 nM protein
4.0
0.0
−60
−40
−20
0
20
40
60
Position relative to center (bp)

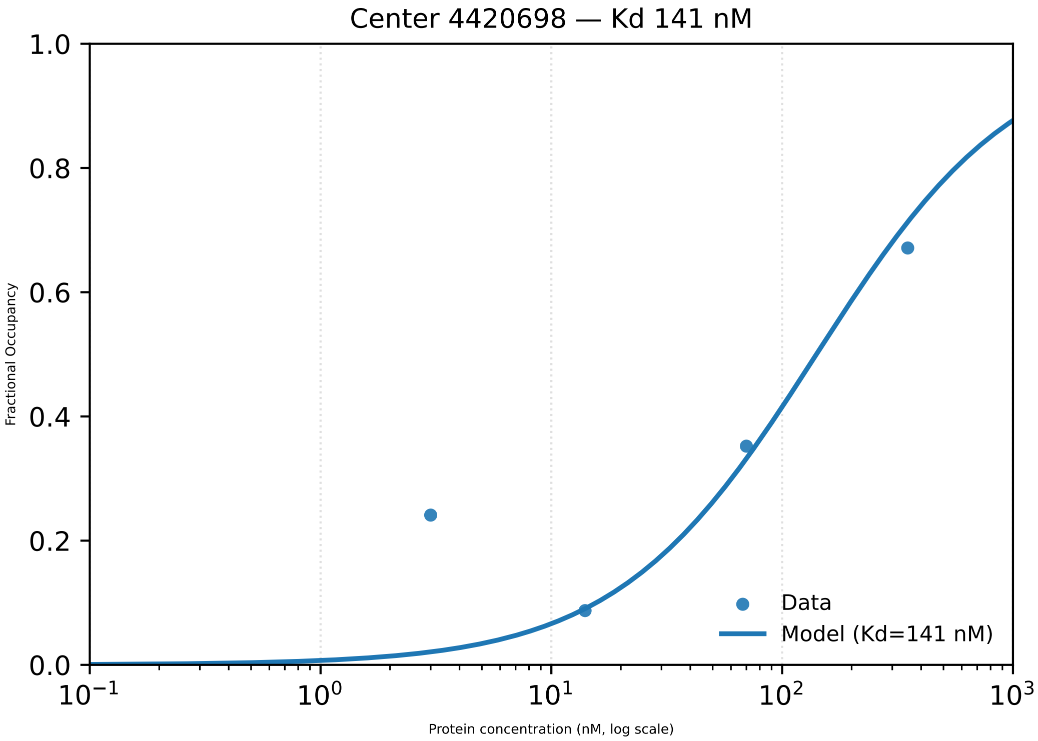

Center 4420698 — Kd 141 nM
1.0
0.8
0.6
0.4
0.2
0.0
Fractional Occupancy
$10^{-1}$
$10^{0}$
$10^{1}$
$10^{2}$
$10^{3}$
Protein concentration (nM, log scale)
Data
Model (Kd=141 nM)

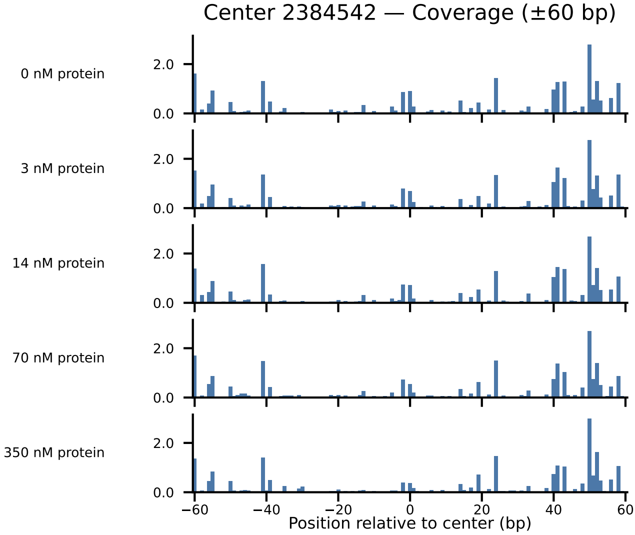
Center 2384542 — Coverage (±60 bp)
0 nM protein
3 nM protein
14 nM protein
70 nM protein
350 nM protein
2.0
0.0
−60
−40
−20
0
20
40
60
Position relative to center (bp)

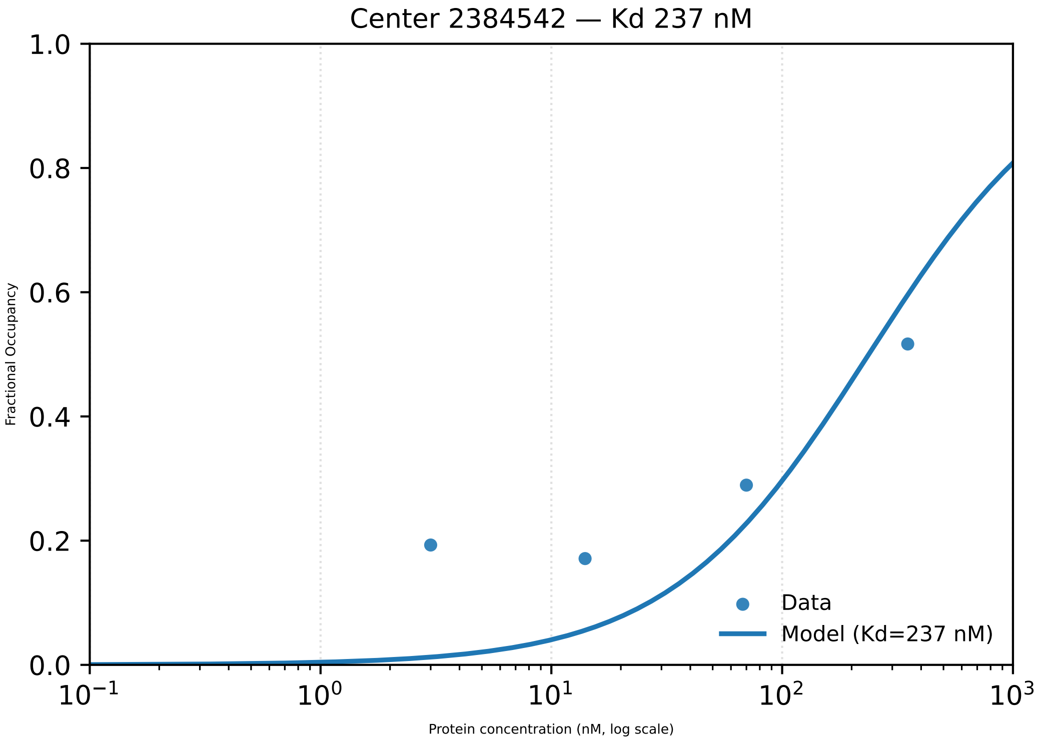

Center 2384542 — Kd 237 nM
Fractional Occupancy
1.0
0.8
0.6
0.4
0.2
0.0
$10^{-1}$
$10^{0}$
$10^{1}$
$10^{2}$
$10^{3}$
Protein concentration (nM, log scale)
Data
Model (Kd=237 nM)

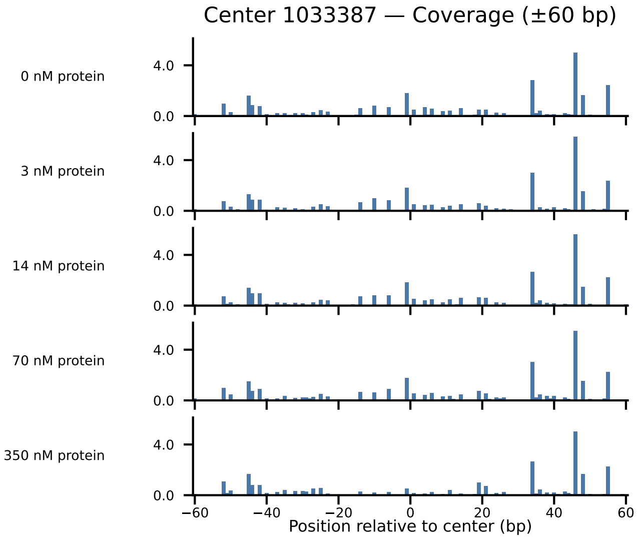

Center 1033387 — Coverage (±60 bp)
0 nM protein
3 nM protein
14 nM protein
70 nM protein
350 nM protein
4.0
0.0
−60
−40
−20
0
20
40
60
Position relative to center (bp)

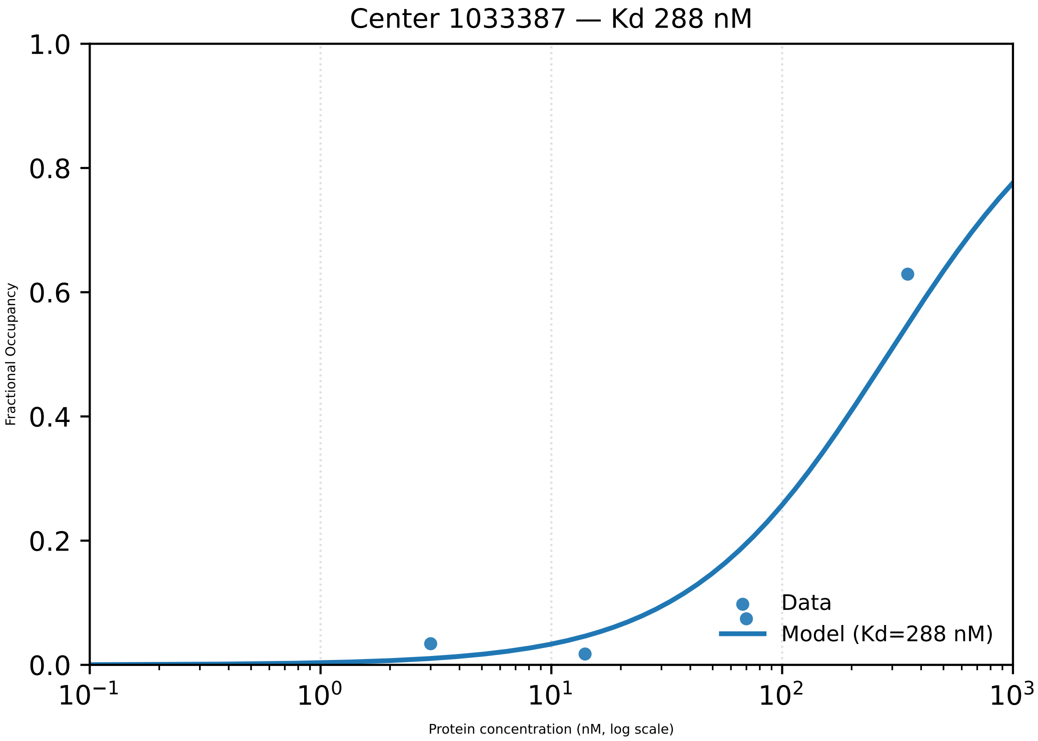

Center 1033387 — Kd 288 nM
1.0
0.8
0.6
0.4
0.2
0.0
Fractional Occupancy
$10^{-1}$
$10^{0}$
$10^{1}$
$10^{2}$
$10^{3}$
Protein concentration (nM, log scale)
Data
Model (Kd=288 nM)

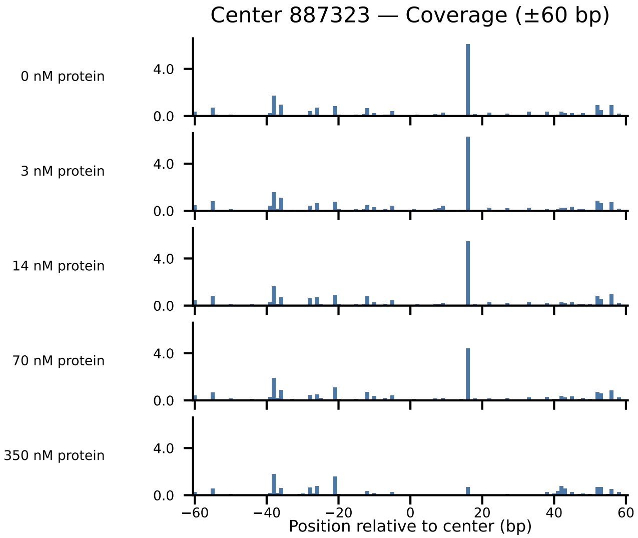
Center 887323 — Coverage (±60 bp)
0 nM protein
3 nM protein
14 nM protein
70 nM protein
350 nM protein
4.0
0.0
−60
−40
−20
0
20
40
60
Position relative to center (bp)

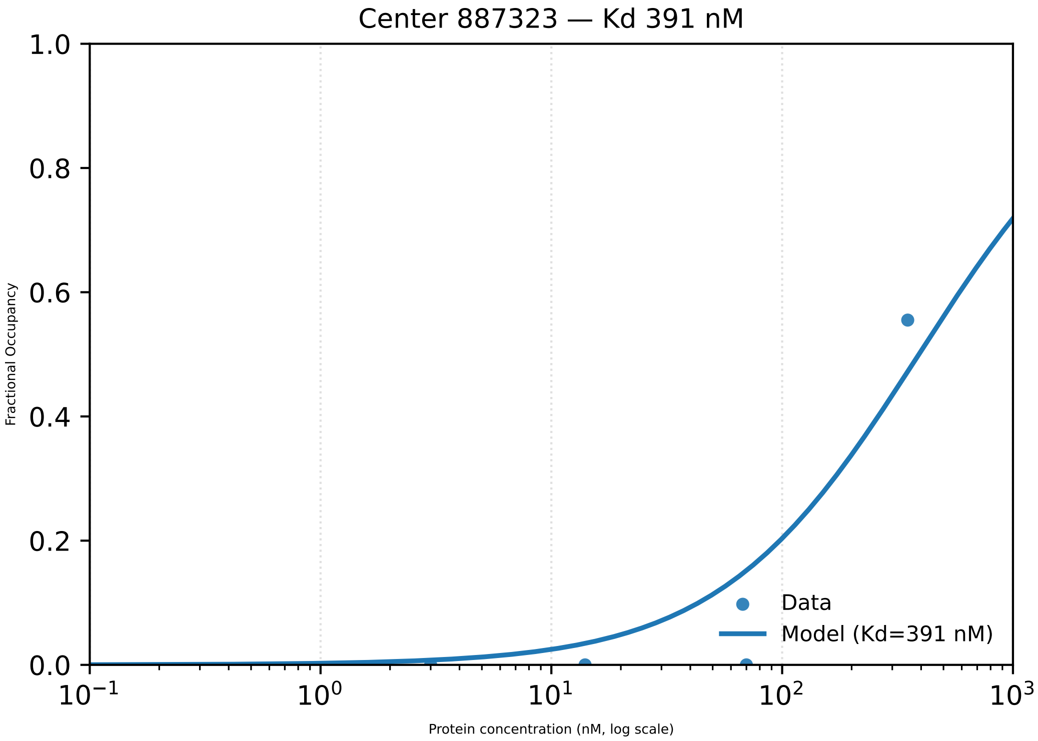
Center 887323 — Kd 391 nM
1.0
0.8
0.6
0.4
0.2
0.0
Fractional Occupancy
$10^{-1}$
$10^{0}$
$10^{1}$
$10^{2}$
$10^{3}$
Protein concentration (nM, log scale)
Data
Model (Kd=391 nM)

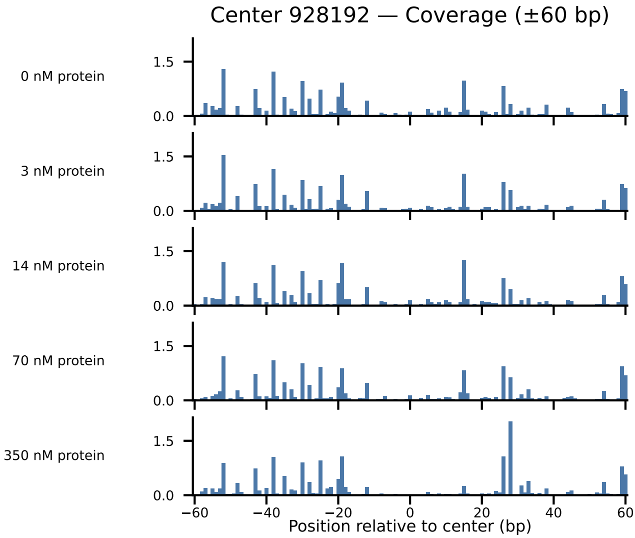
Center 928192 — Coverage (±60 bp)
0 nM protein
3 nM protein
14 nM protein
70 nM protein
350 nM protein
1.5
0.0
−60
−40
−20
0
20
40
60
Position relative to center (bp)

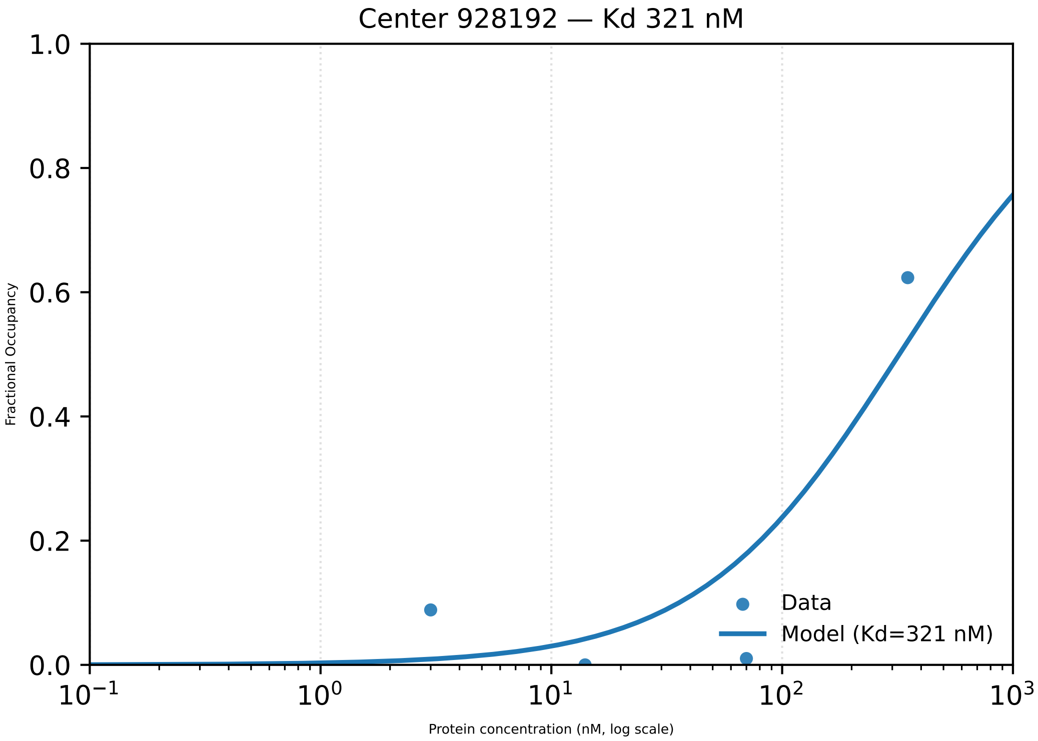

Center 928192 — Kd 321 nM
Fractional Occupancy
Protein concentration (nM, log scale)
Data
Model (Kd=321 nM)
1.0
0.8
0.6
0.4
0.2
0.0
$10^{-1}$
$10^{0}$
$10^{1}$
$10^{2}$
$10^{3}$

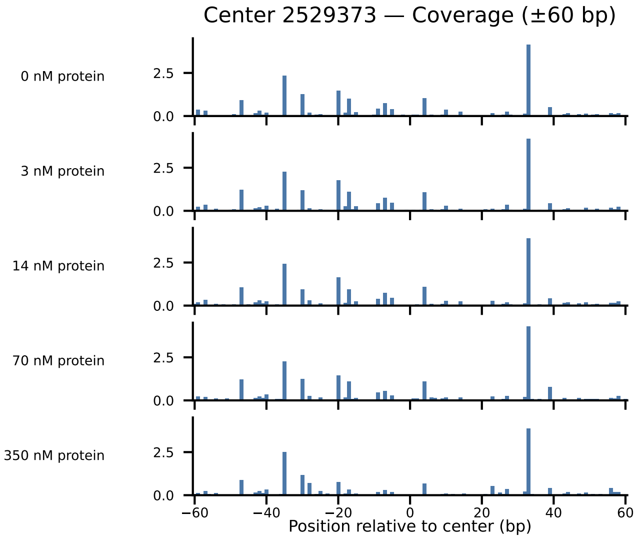

Center 2529373 — Coverage (±60 bp)
0 nM protein
3 nM protein
14 nM protein
70 nM protein
350 nM protein
2.5
0.0
−60
−40
−20
0
20
40
60
Position relative to center (bp)

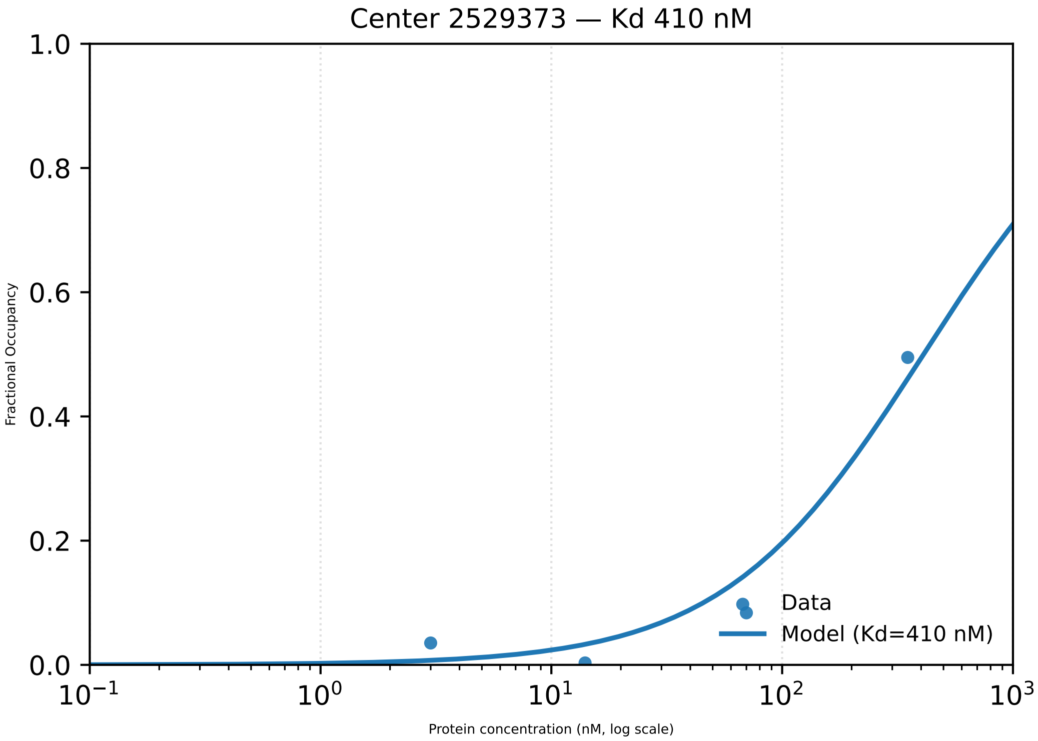

Center 2529373 — Kd 410 nM
1.0
0.8
0.6
0.4
0.2
0.0
Fractional Occupancy
$10^{-1}$
$10^{0}$
$10^{1}$
$10^{2}$
$10^{3}$
Protein concentration (nM, log scale)
Data
Model (Kd=410 nM)

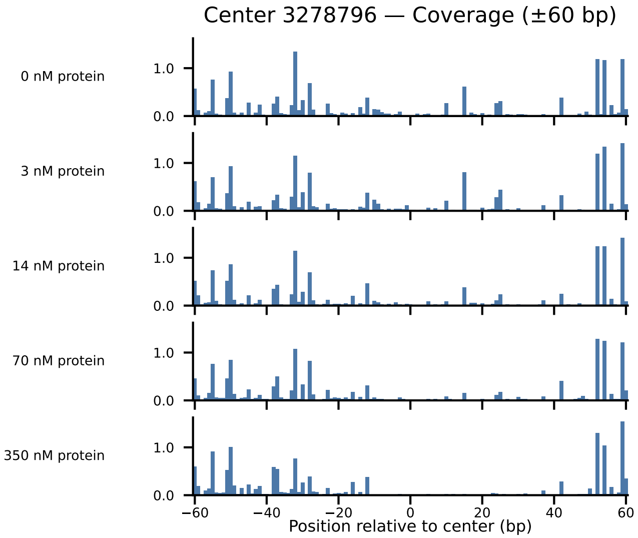

Center 3278796 — Coverage (±60 bp)
0 nM protein
3 nM protein
14 nM protein
70 nM protein
350 nM protein
1.0
0.0
−60
−40
−20
0
20
40
60
Position relative to center (bp)

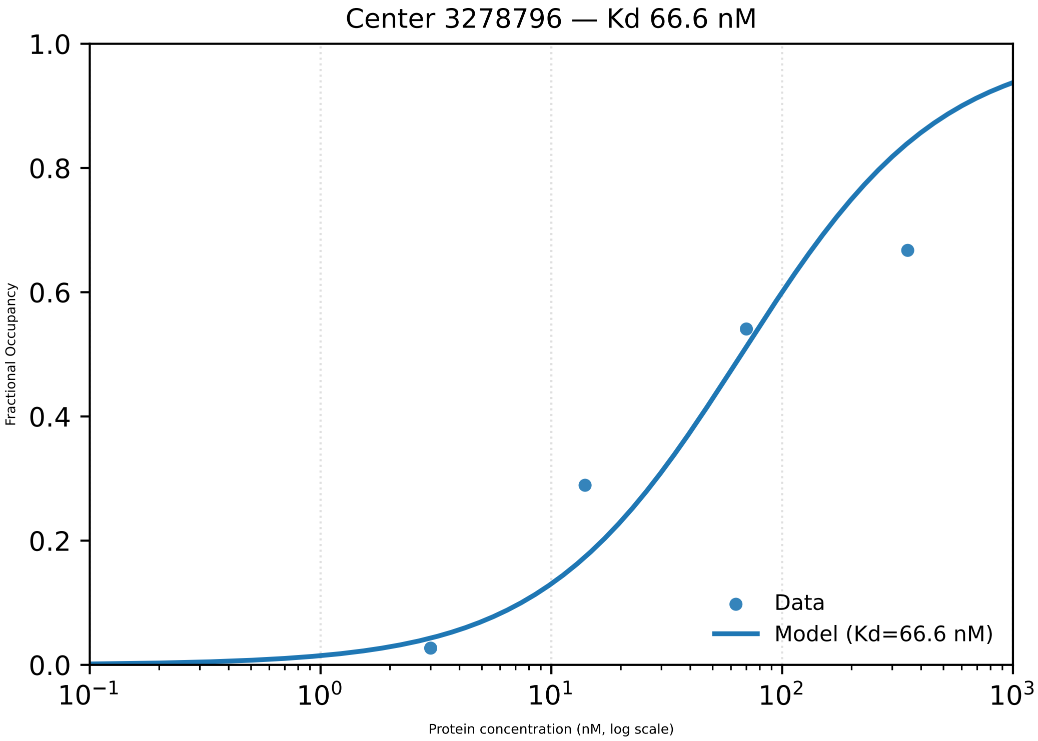

Center 3278796 — Kd 66.6 nM
Fractional Occupancy
1.0
0.8
0.6
0.4
0.2
0.0
$10^{-1}$
$10^{0}$
$10^{1}$
$10^{2}$
$10^{3}$
Protein concentration (nM, log scale)
Data
Model (Kd=66.6 nM)

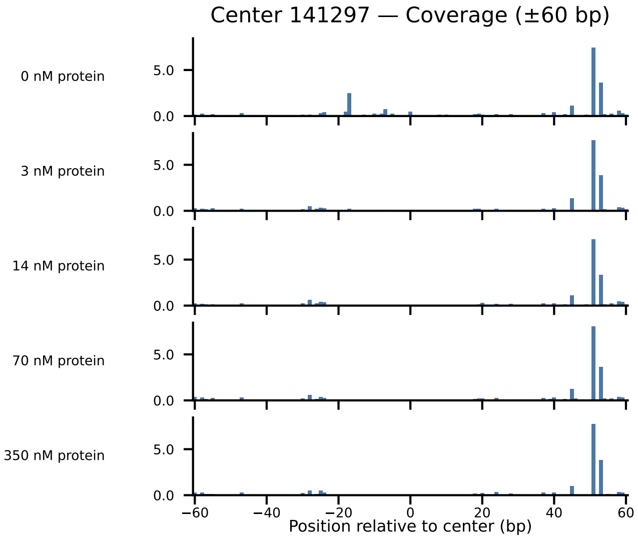

Center 141297 — Coverage (±60 bp)
0 nM protein
3 nM protein
14 nM protein
70 nM protein
350 nM protein
5.0
0.0
−60
−40
−20
0
20
40
60
Position relative to center (bp)

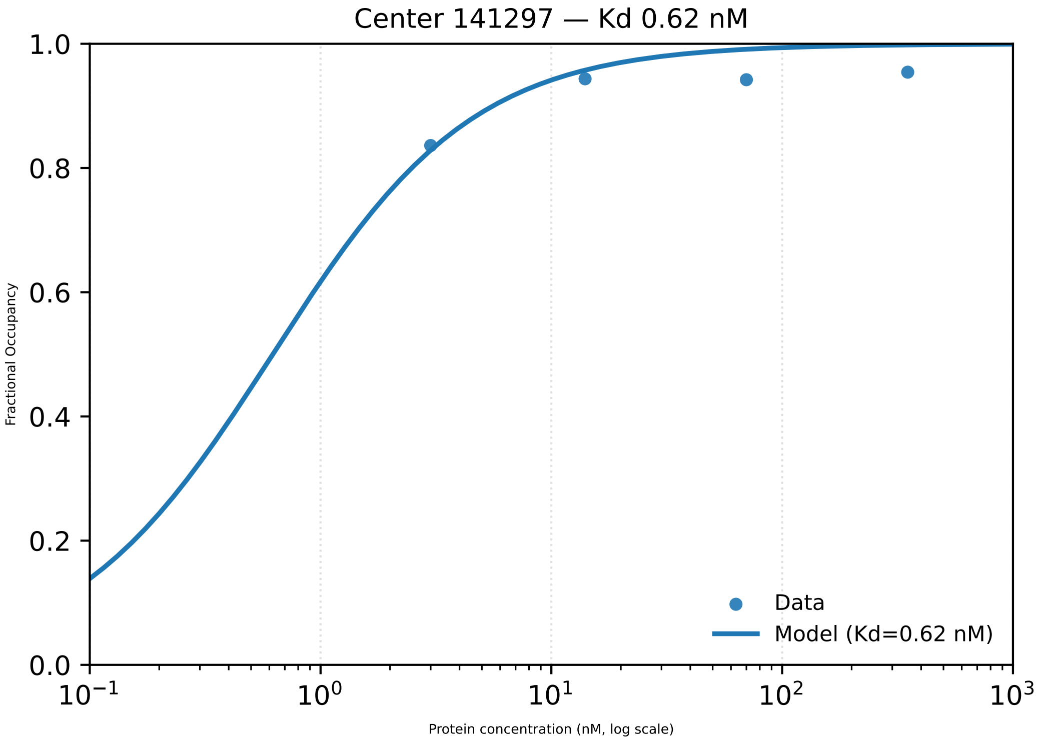

Center 141297 — Kd 0.62 nM
Fractional Occupancy
0.0
0.2
0.4
0.6
0.8
1.0
$10^{-1}$
$10^{0}$
$10^{1}$
$10^{2}$
$10^{3}$
Protein concentration (nM, log scale)
Data
Model (Kd=0.62 nM)

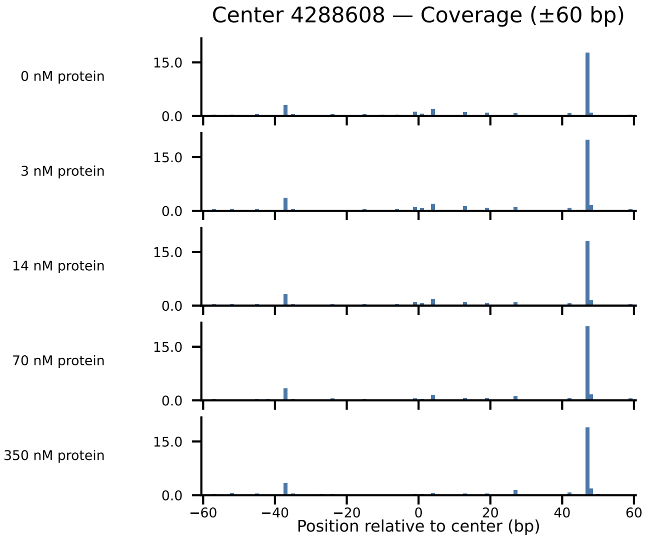

Center 4288608 — Coverage (±60 bp)
0 nM protein
3 nM protein
14 nM protein
70 nM protein
350 nM protein
15.0
0.0
−60
−40
−20
0
20
40
60
Position relative to center (bp)

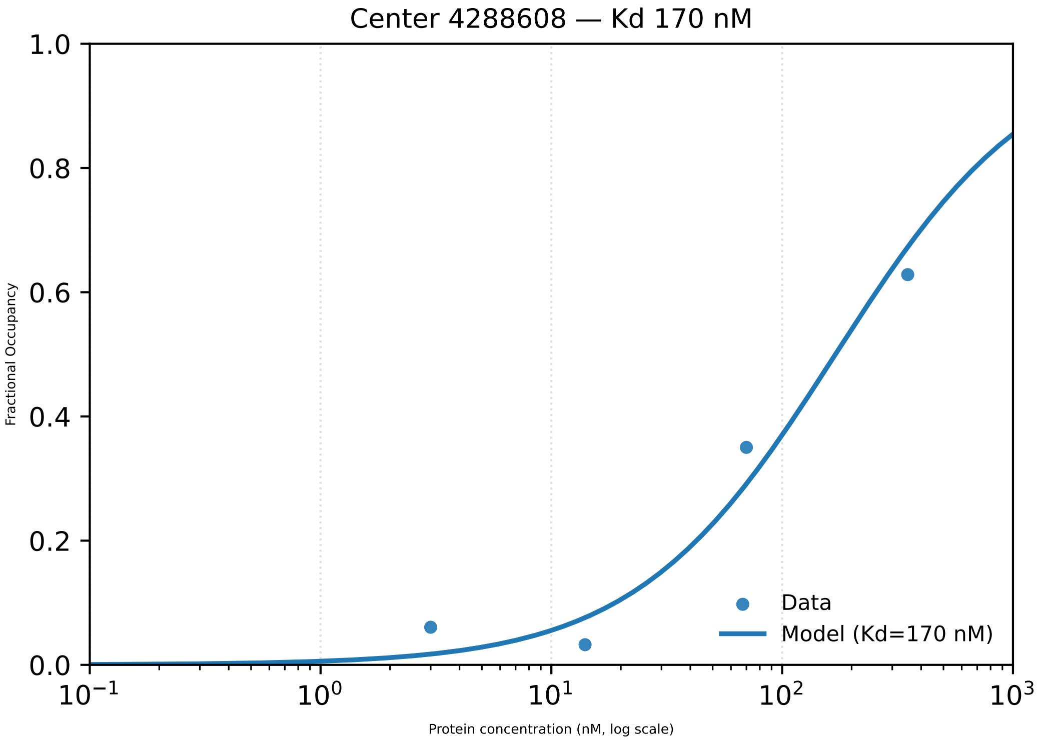

Center 4288608 — Kd 170 nM
Fractional Occupancy
1.0
0.8
0.6
0.4
0.2
0.0
$10^{-1}$
$10^{0}$
$10^{1}$
$10^{2}$
$10^{3}$
Protein concentration (nM, log scale)
Data
Model (Kd=170 nM)

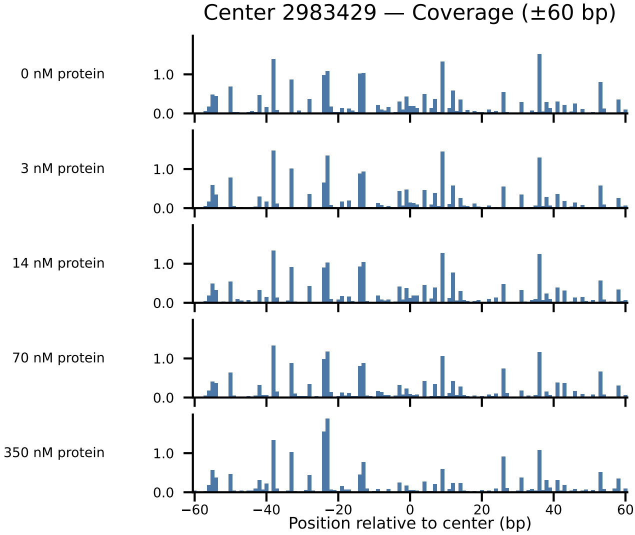

Center 2983429 — Coverage (±60 bp)
0 nM protein
3 nM protein
14 nM protein
70 nM protein
350 nM protein
1.0
0.0
−60
−40
−20
0
20
40
60
Position relative to center (bp)

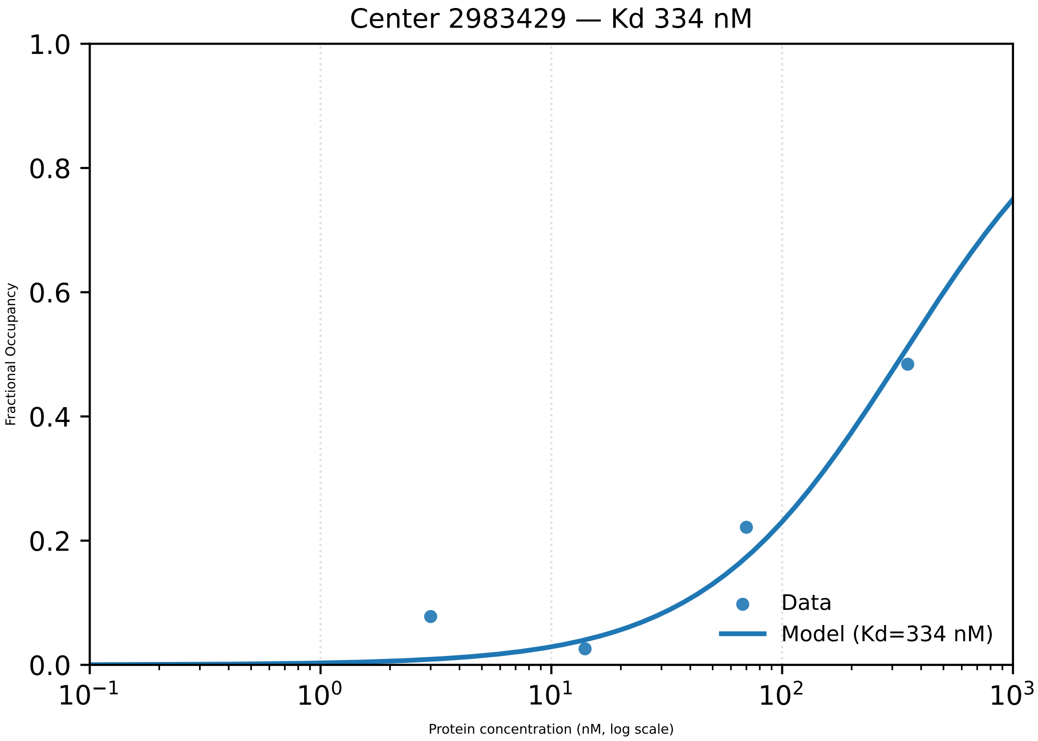

Center 2983429 — Kd 334 nM
Fractional Occupancy
1.0
0.8
0.6
0.4
0.2
0.0
$10^{-1}$
$10^{0}$
$10^{1}$
$10^{2}$
$10^{3}$
Protein concentration (nM, log scale)
Data
Model (Kd=334 nM)

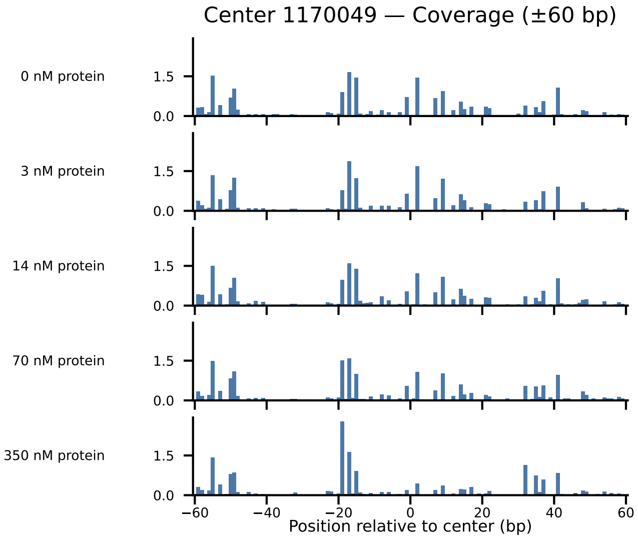

Center 1170049 — Coverage (±60 bp)
0 nM protein
3 nM protein
14 nM protein
70 nM protein
350 nM protein
1.5
0.0
−60
−40
−20
0
20
40
60
Position relative to center (bp)

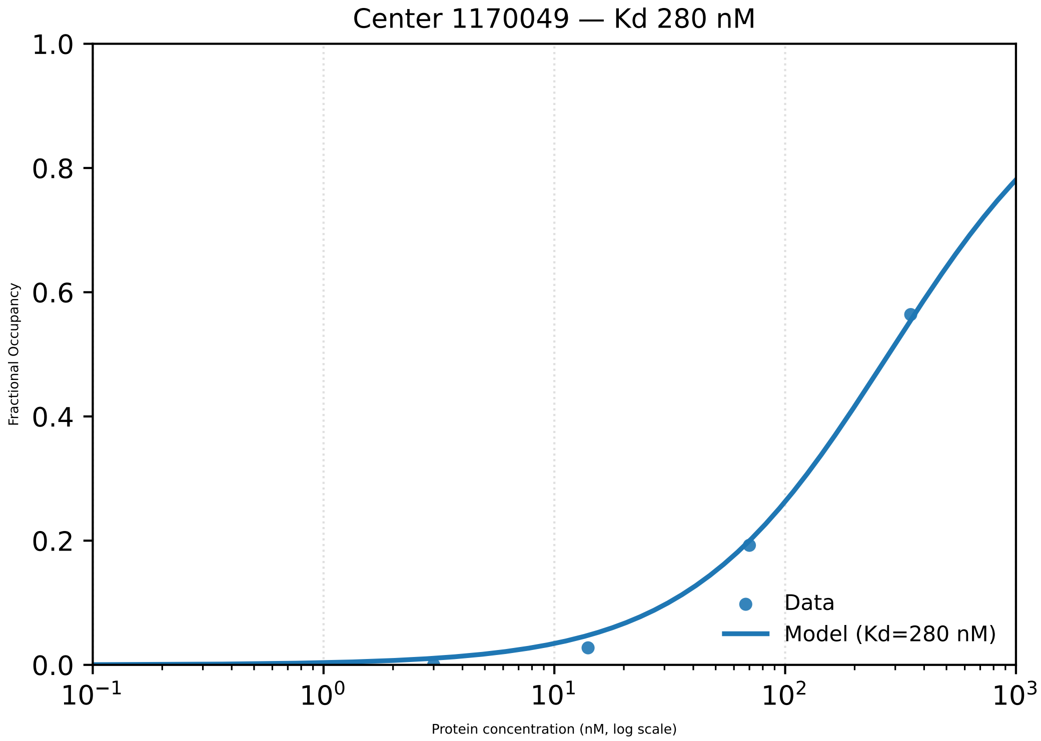

Center 1170049 — Kd 280 nM
Fractional Occupancy
1.0
0.8
0.6
0.4
0.2
0.0
$10^{-1}$
$10^{0}$
$10^{1}$
$10^{2}$
$10^{3}$
Protein concentration (nM, log scale)
Data
Model (Kd=280 nM)

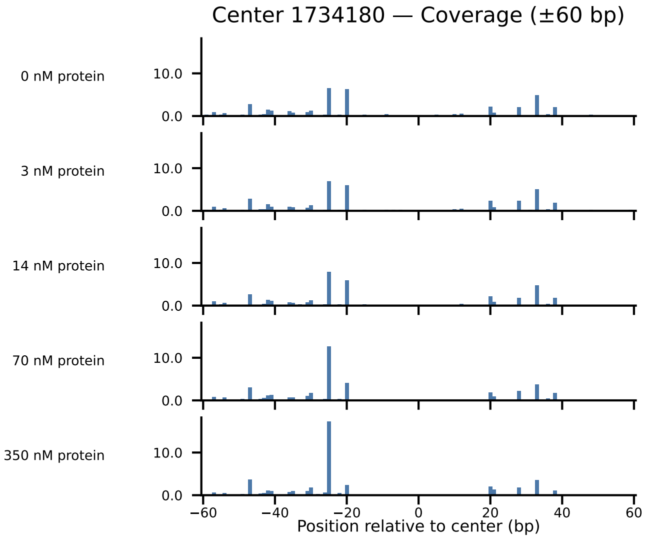
Center 1734180 — Coverage (±60 bp)
0 nM protein
3 nM protein
14 nM protein
70 nM protein
350 nM protein
10.0
0.0
−60
−40
−20
0
20
40
60
Position relative to center (bp)

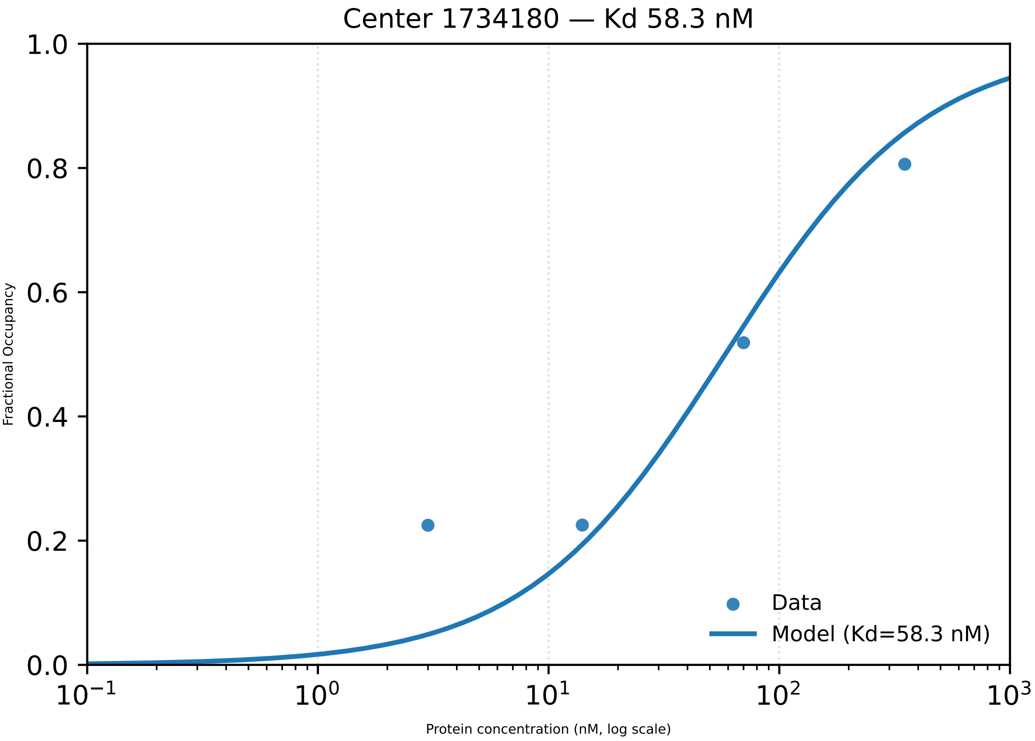

Center 1734180 — Kd 58.3 nM
Fractional Occupancy
1.0
0.8
0.6
0.4
0.2
0.0
$10^{-1}$
$10^{0}$
$10^{1}$
$10^{2}$
$10^{3}$
Protein concentration (nM, log scale)
Data
Model (Kd=58.3 nM)

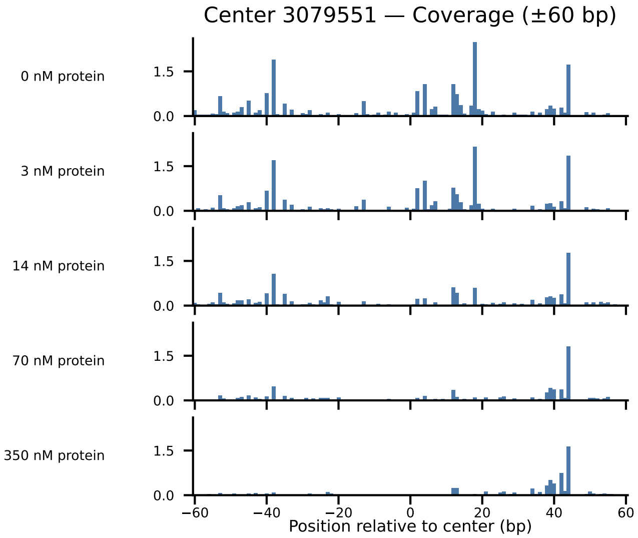

Center 3079551 — Coverage (±60 bp)
0 nM protein
3 nM protein
14 nM protein
70 nM protein
350 nM protein
1.5
0.0
−60
−40
−20
0
20
40
60
Position relative to center (bp)

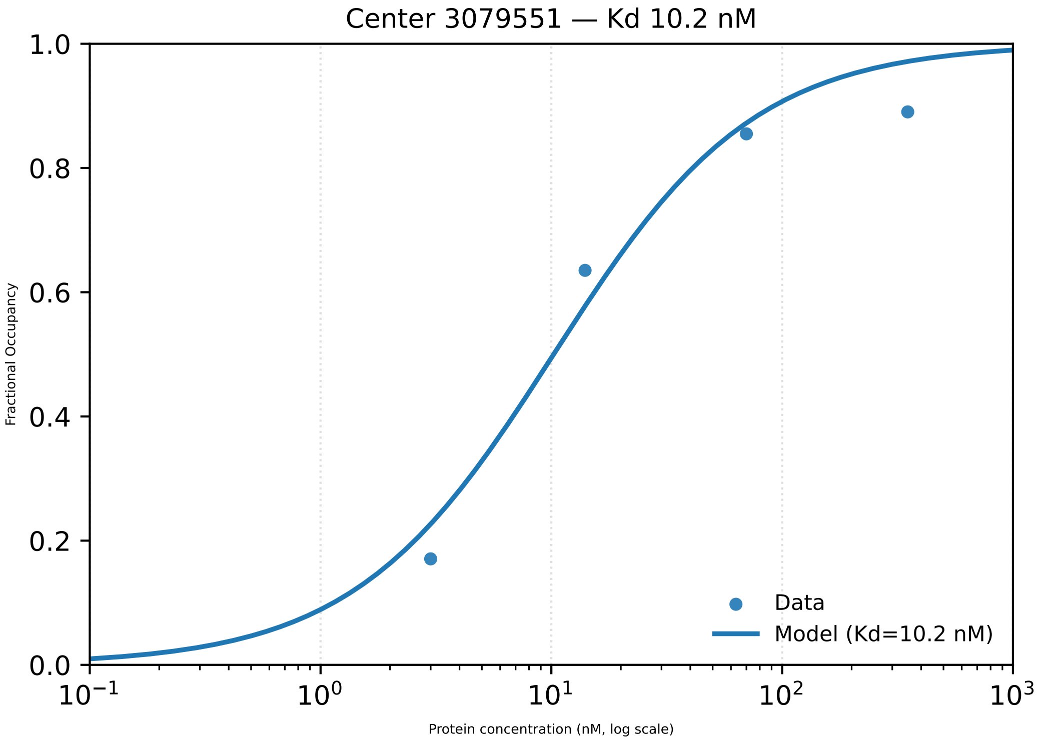

Center 3079551 — Kd 10.2 nM
1.0
0.8
0.6
0.4
0.2
0.0
Fractional Occupancy
$10^{-1}$
$10^{0}$
$10^{1}$
$10^{2}$
$10^{3}$
Protein concentration (nM, log scale)
Data
Model (Kd=10.2 nM)

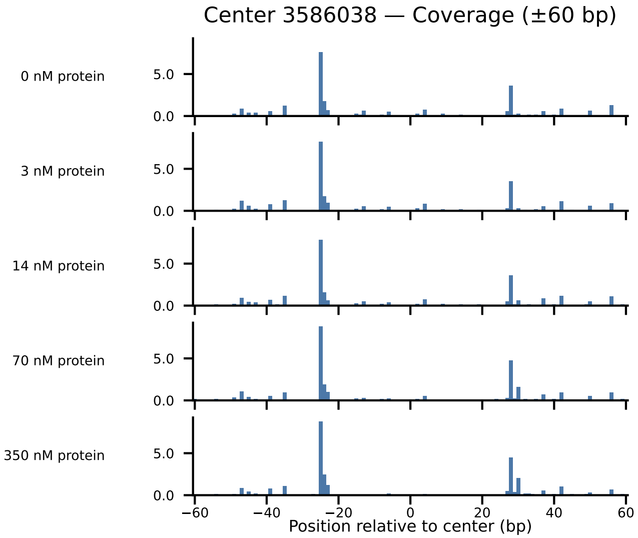
Center 3586038 — Coverage (±60 bp)
0 nM protein
3 nM protein
14 nM protein
70 nM protein
350 nM protein
5.0
0.0
−60
−40
−20
0
20
40
60
Position relative to center (bp)

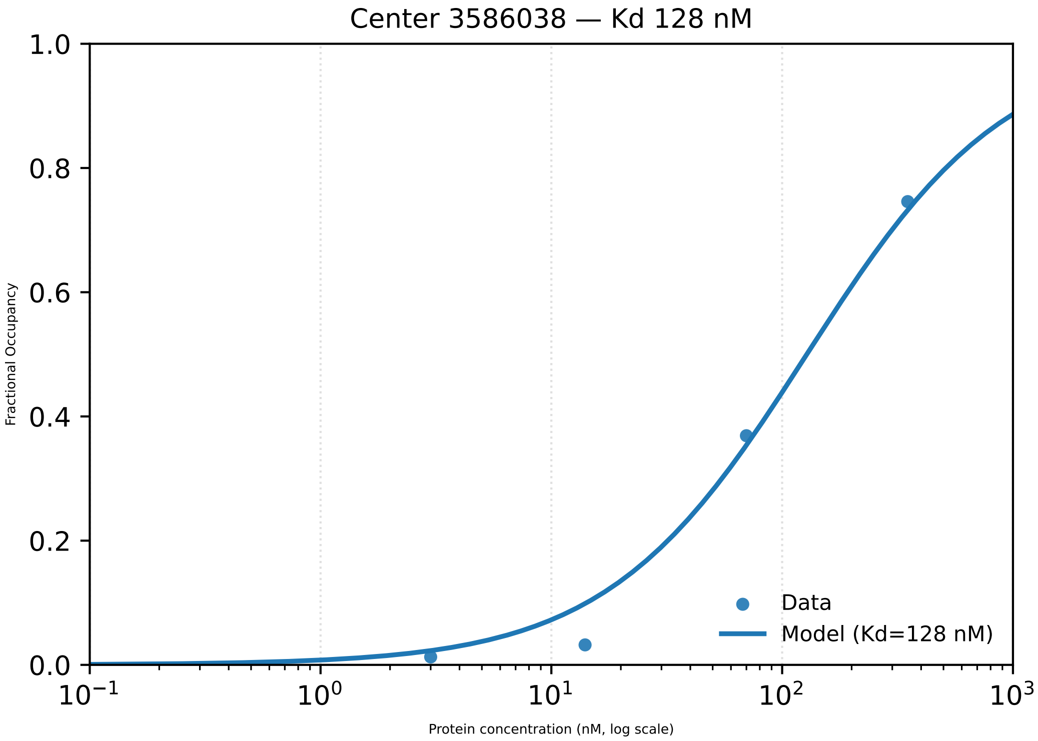
Center 3586038 — Kd 128 nM
1.0
0.8
0.6
0.4
0.2
0.0
Fractional Occupancy
10$^{-1}$
10$^{0}$
10$^{1}$
10$^{2}$
10$^{3}$
Protein concentration (nM, log scale)
Data
Model (Kd=128 nM)

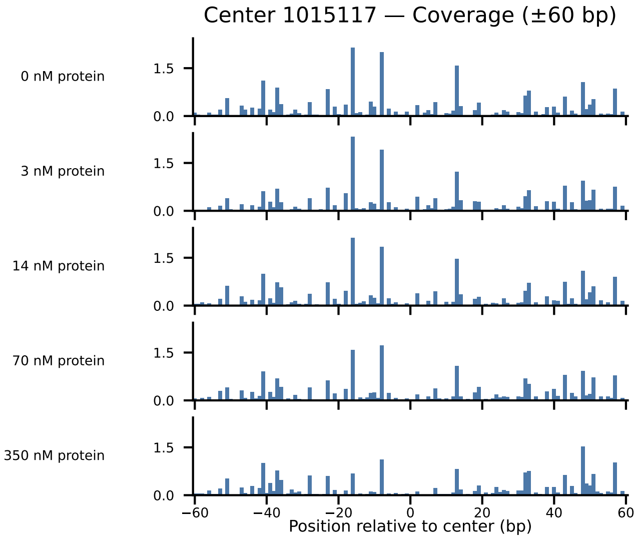
Center 1015117 — Coverage (±60 bp)
0 nM protein
3 nM protein
14 nM protein
70 nM protein
350 nM protein
1.5
0.0
−60
−40
−20
0
20
40
60
Position relative to center (bp)

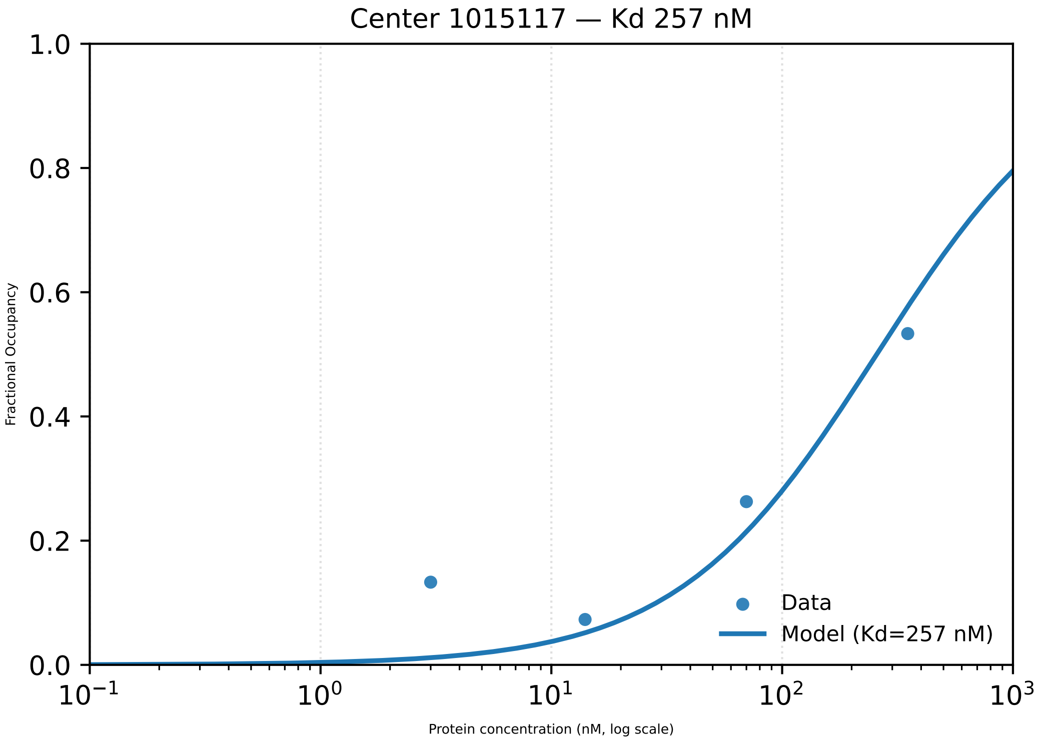

Center 1015117 — Kd 257 nM
Fractional Occupancy
1.0
0.8
0.6
0.4
0.2
0.0
$10^{-1}$
$10^{0}$
$10^{1}$
$10^{2}$
$10^{3}$
Protein concentration (nM, log scale)
Data
Model (Kd=257 nM)

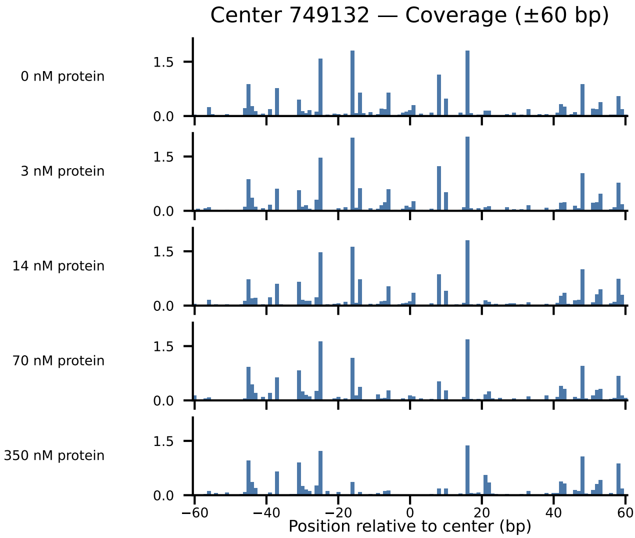
Center 749132 — Coverage (±60 bp)
0 nM protein
3 nM protein
14 nM protein
70 nM protein
350 nM protein
1.5
0.0
−60
−40
−20
0
20
40
60
Position relative to center (bp)

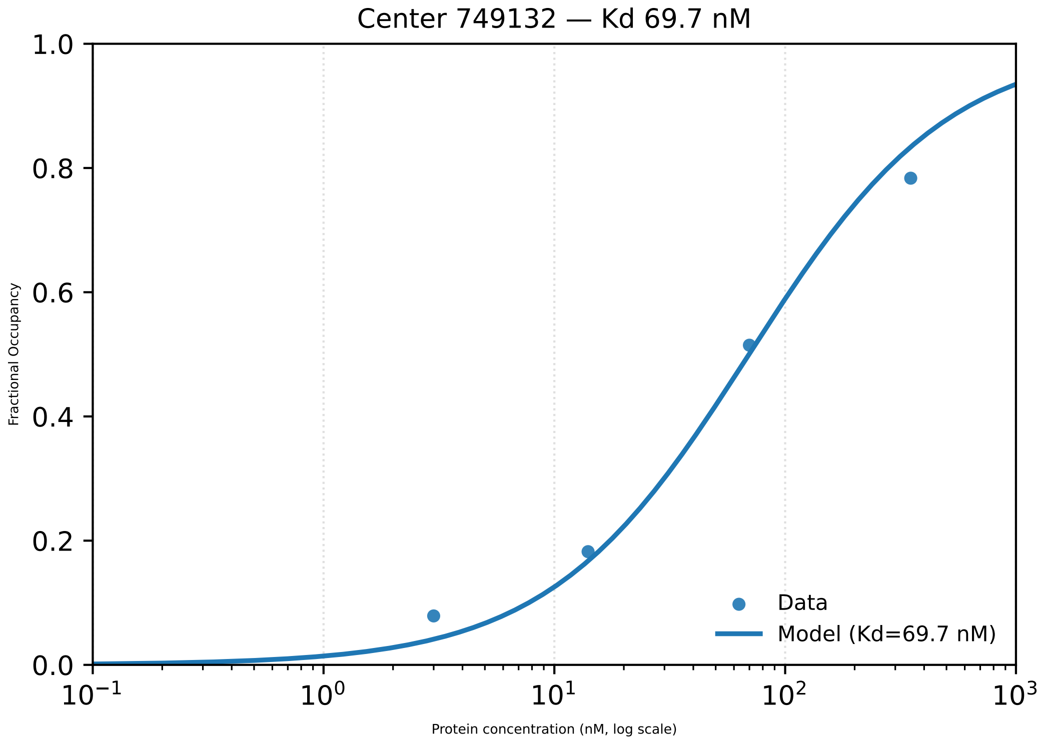

Center 749132 — Kd 69.7 nM
1.0
0.8
0.6
0.4
0.2
0.0
Fractional Occupancy
$10^{-1}$
$10^{0}$
$10^{1}$
$10^{2}$
$10^{3}$
Protein concentration (nM, log scale)
Data
Model (Kd=69.7 nM)

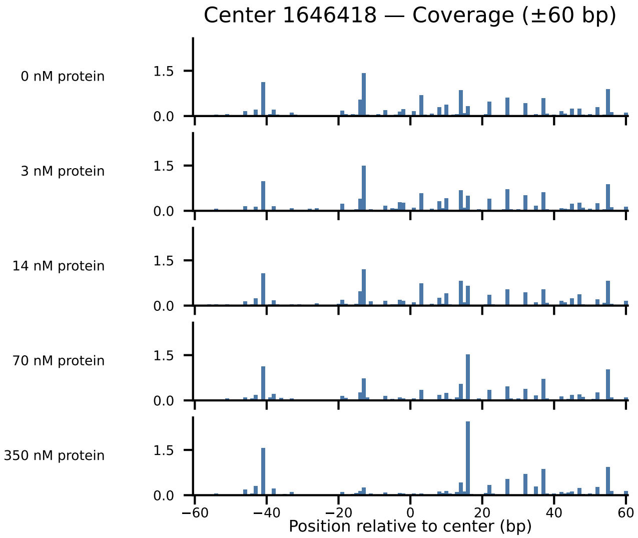

Center 1646418 — Coverage (±60 bp)
0 nM protein
3 nM protein
14 nM protein
70 nM protein
350 nM protein
1.5
0.0
−60
−40
−20
0
20
40
60
Position relative to center (bp)

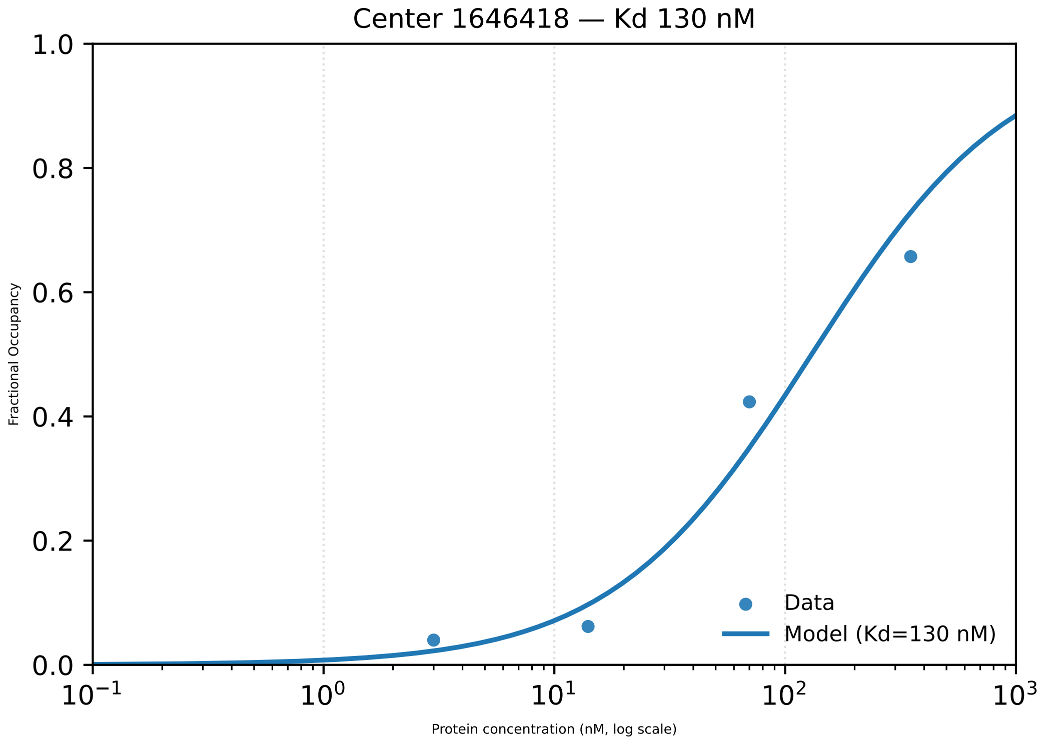

Center 1646418 — Kd 130 nM
1.0
0.8
0.6
0.4
0.2
0.0
Fractional Occupancy
$10^{-1}$
$10^{0}$
$10^{1}$
$10^{2}$
$10^{3}$
Protein concentration (nM, log scale)
Data
Model (Kd=130 nM)

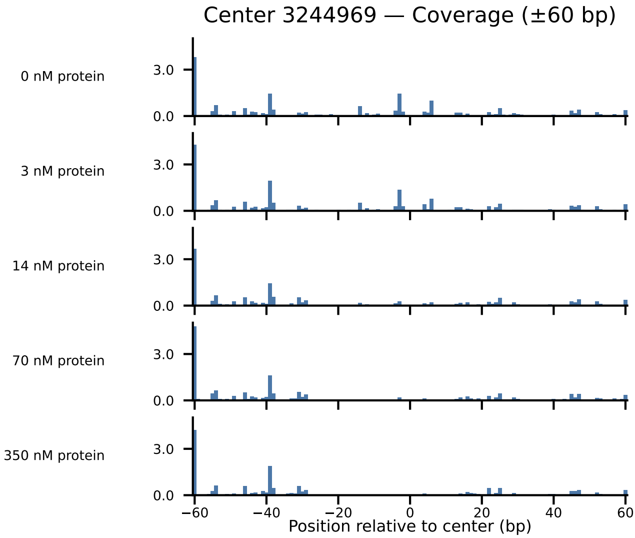
Center 3244969 — Coverage (±60 bp)
0 nM protein
3 nM protein
14 nM protein
70 nM protein
350 nM protein
3.0
0.0
−60
−40
−20
0
20
40
60
Position relative to center (bp)

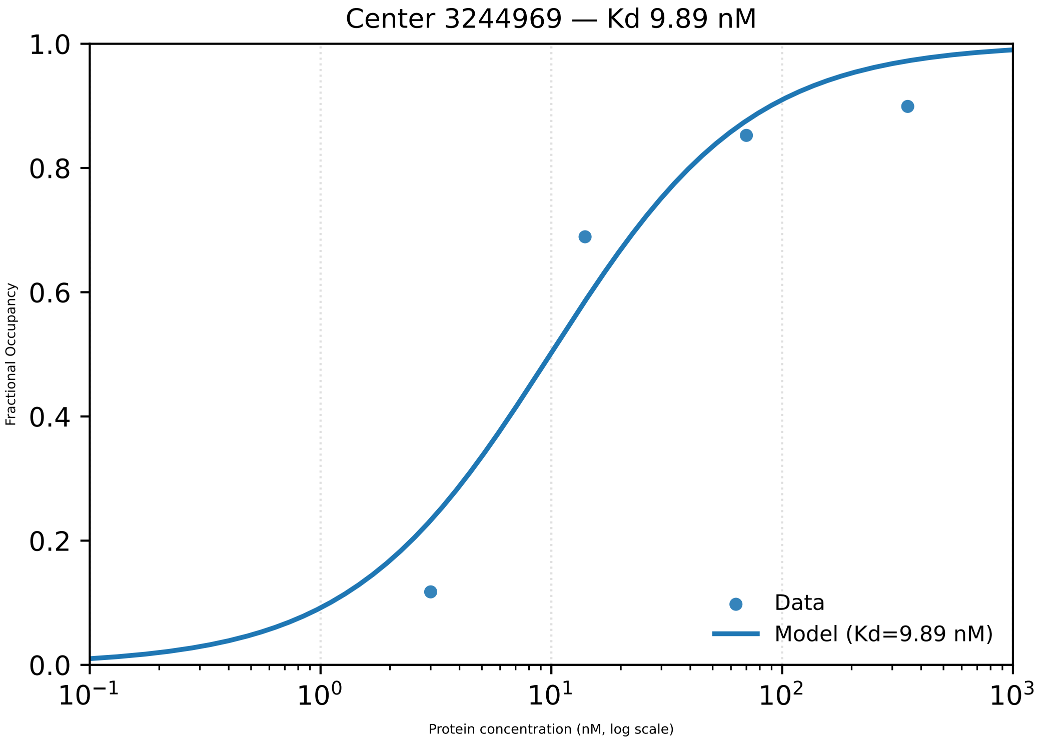
Center 3244969 — Kd 9.89 nM
Fractional Occupancy
0.0
0.2
0.4
0.6
0.8
1.0
$10^{-1}$
$10^{0}$
$10^{1}$
$10^{2}$
$10^{3}$
Protein concentration (nM, log scale)
Data
Model (Kd=9.89 nM)

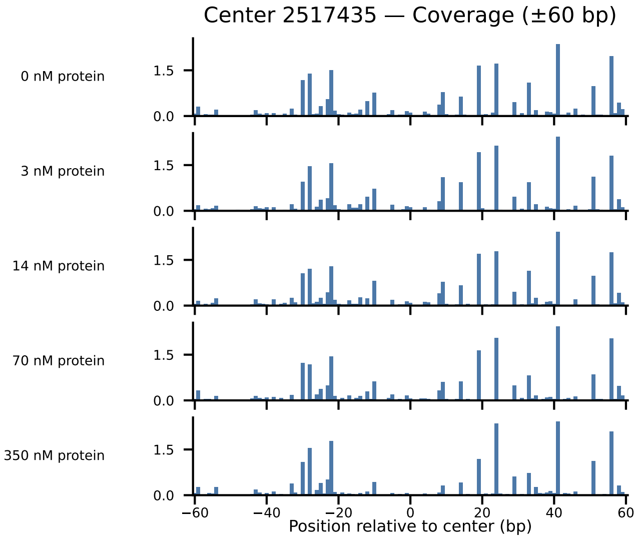
Center 2517435 — Coverage (±60 bp)
0 nM protein
3 nM protein
14 nM protein
70 nM protein
350 nM protein
1.5
0.0
−60
−40
−20
0
20
40
60
Position relative to center (bp)

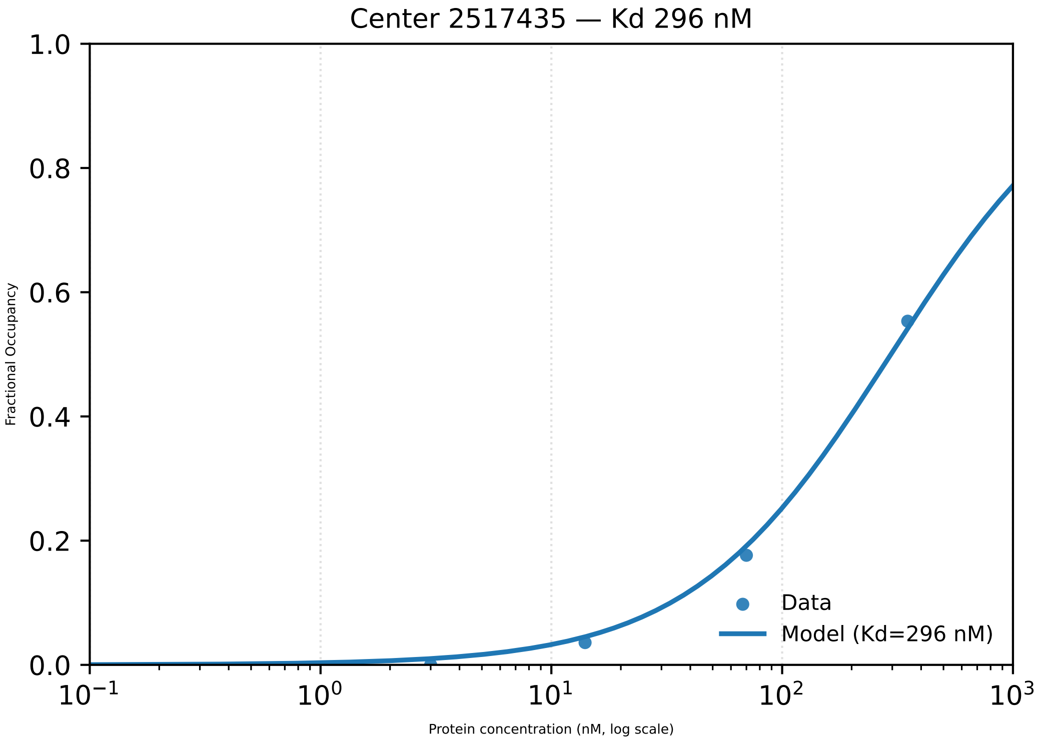

Center 2517435 — Kd 296 nM
1.0
0.8
0.6
0.4
0.2
0.0
Fractional Occupancy
$10^{-1}$
$10^{0}$
$10^{1}$
$10^{2}$
$10^{3}$
Protein concentration (nM, log scale)
Data
Model (Kd=296 nM)

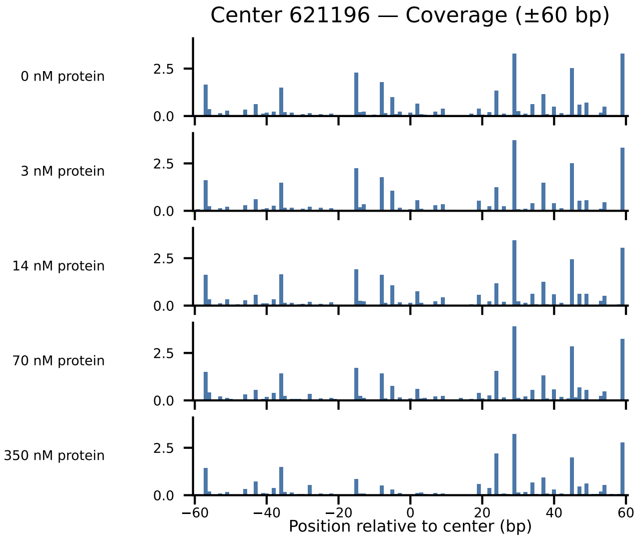

Center 621196 — Coverage (±60 bp)
0 nM protein
3 nM protein
14 nM protein
70 nM protein
350 nM protein
2.5
0.0
−60
−40
−20
0
20
40
60
Position relative to center (bp)

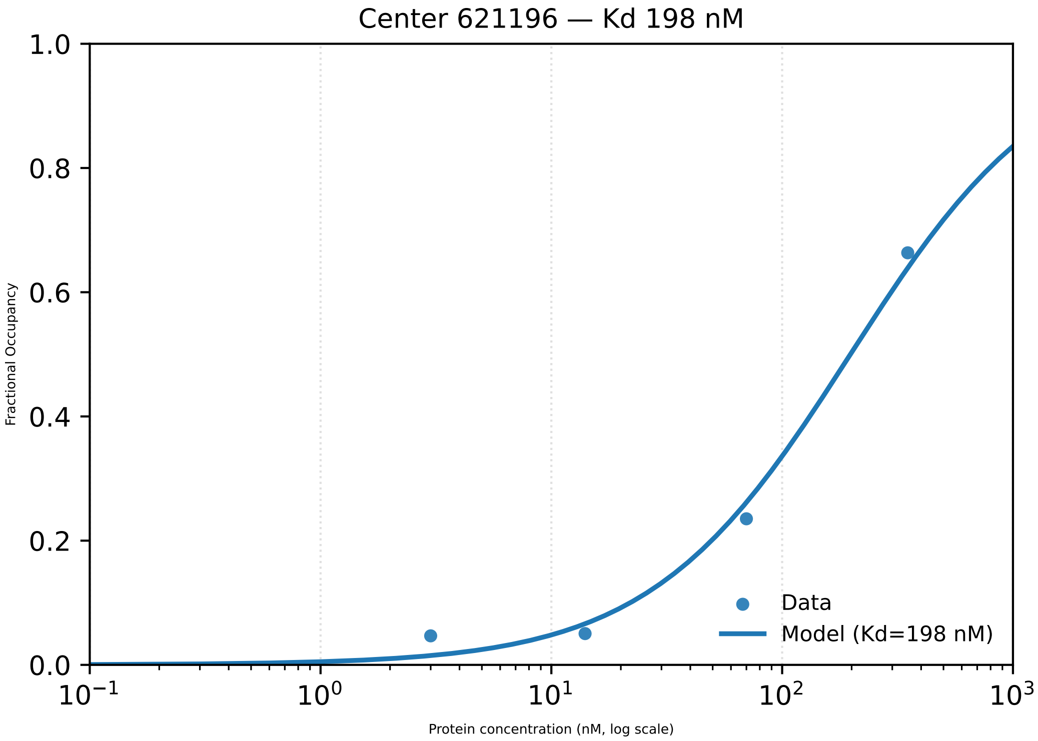

Center 621196 — Kd 198 nM
Fractional Occupancy
1.0
0.8
0.6
0.4
0.2
0.0
$10^{-1}$
$10^{0}$
$10^{1}$
$10^{2}$
$10^{3}$
Protein concentration (nM, log scale)
Data
Model (Kd=198 nM)

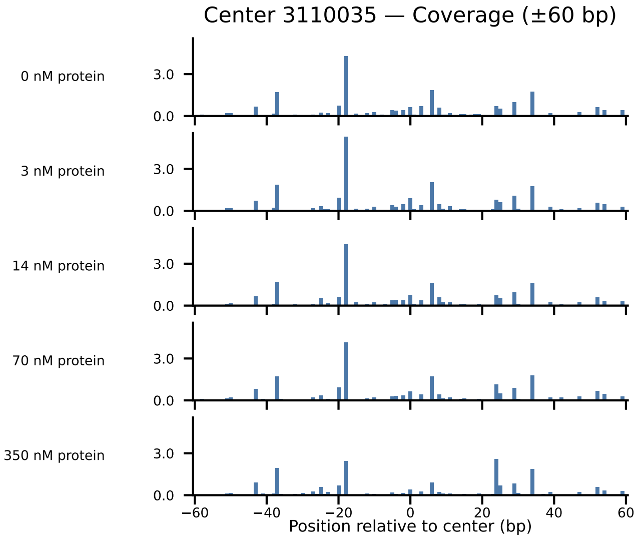

Center 3110035 — Coverage (±60 bp)
0 nM protein
3 nM protein
14 nM protein
70 nM protein
350 nM protein
3.0
0.0
−60
−40
−20
0
20
40
60
Position relative to center (bp)

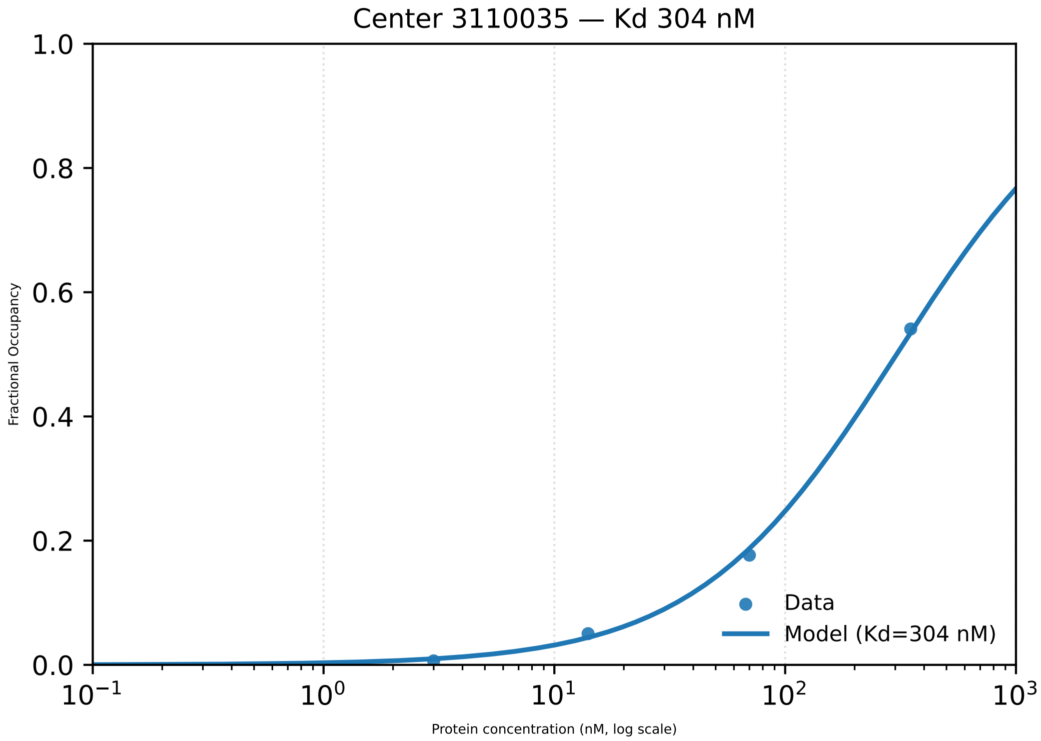

Center 3110035 — Kd 304 nM
Fractional Occupancy
1.0
0.8
0.6
0.4
0.2
0.0
$10^{-1}$
$10^{0}$
$10^{1}$
$10^{2}$
$10^{3}$
Protein concentration (nM, log scale)
Data
Model (Kd=304 nM)

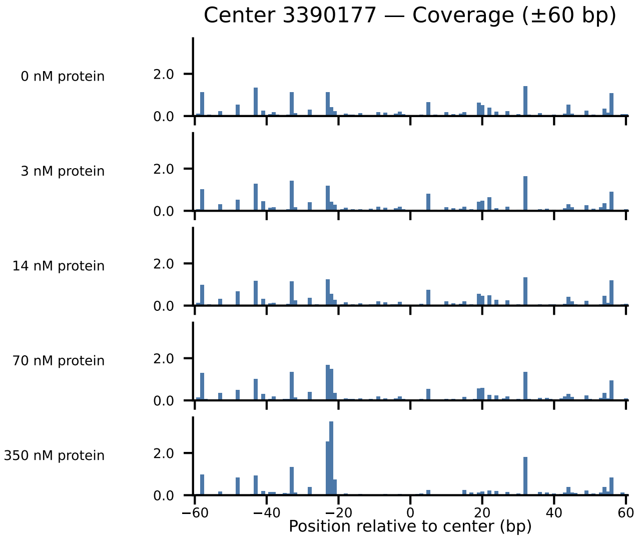
Center 3390177 — Coverage (±60 bp)
0 nM protein
3 nM protein
14 nM protein
70 nM protein
350 nM protein
2.0
0.0
−60
−40
−20
0
20
40
60
Position relative to center (bp)

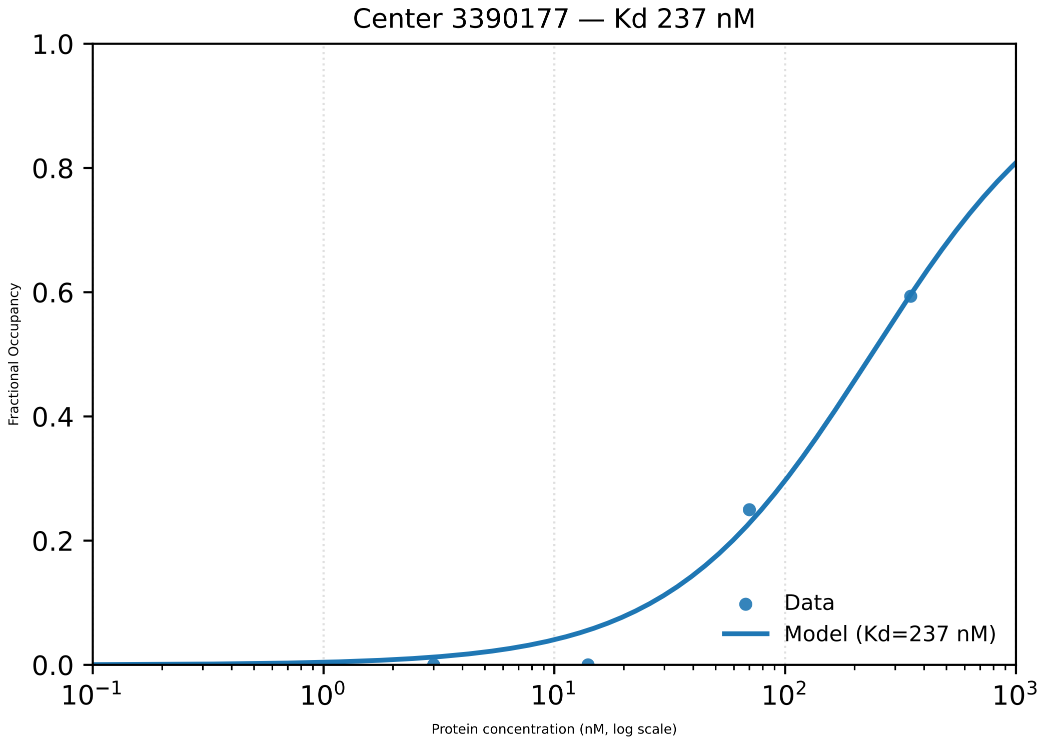

Center 3390177 — Kd 237 nM
Fractional Occupancy
1.0
0.8
0.6
0.4
0.2
0.0
$10^{-1}$
$10^{0}$
$10^{1}$
$10^{2}$
$10^{3}$
Protein concentration (nM, log scale)
Data
Model (Kd=237 nM)

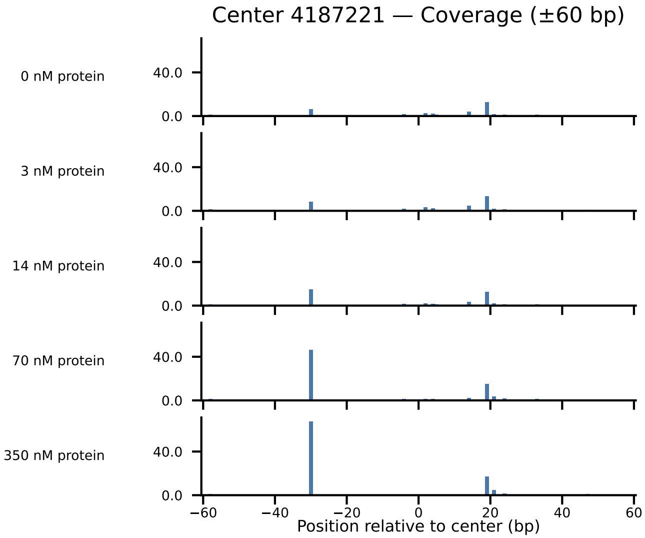
Center 4187221 — Coverage (±60 bp)
0 nM protein
3 nM protein
14 nM protein
70 nM protein
350 nM protein
40.0
0.0
−60
−40
−20
0
20
40
60
Position relative to center (bp)

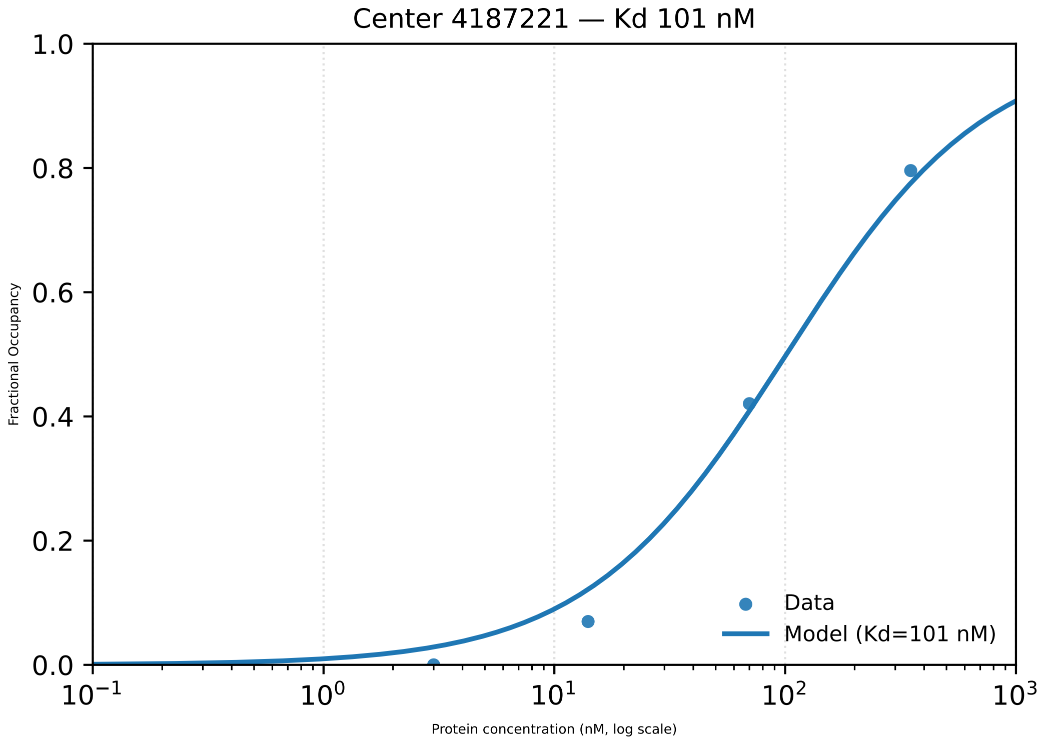

Center 4187221 — Kd 101 nM
1.0
0.8
0.6
0.4
0.2
0.0
Fractional Occupancy
Data
Model (Kd=101 nM)
$10^{-1}$
$10^{0}$
$10^{1}$
$10^{2}$
$10^{3}$
Protein concentration (nM, log scale)

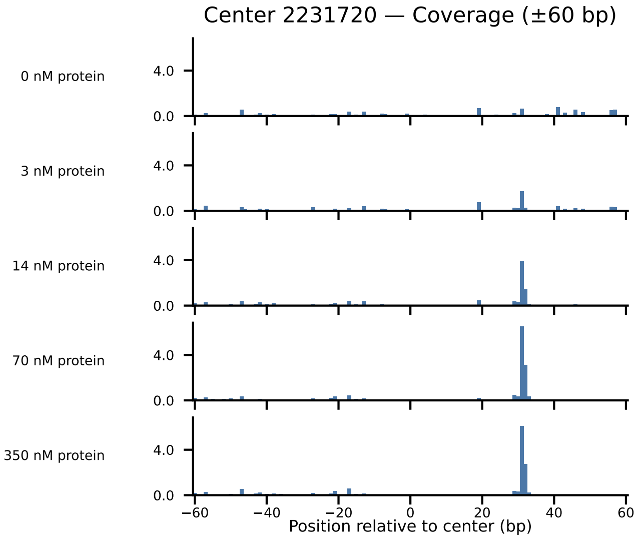

Center 2231720 — Coverage (±60 bp)
0 nM protein
3 nM protein
14 nM protein
70 nM protein
350 nM protein
4.0
0.0
−60
−40
−20
0
20
40
60
Position relative to center (bp)

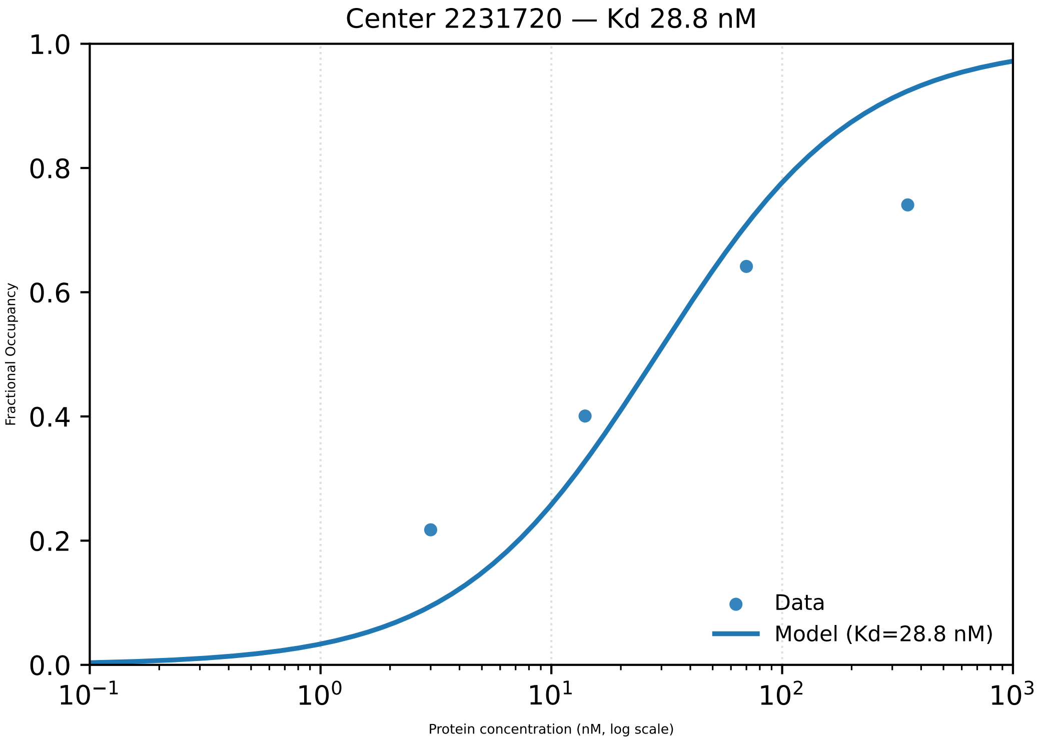

Center 2231720 — Kd 28.8 nM
1.0
0.8
0.6
0.4
0.2
0.0
Fractional Occupancy
$10^{-1}$
$10^{0}$
$10^{1}$
$10^{2}$
$10^{3}$
Protein concentration (nM, log scale)
Data
Model (Kd=28.8 nM)

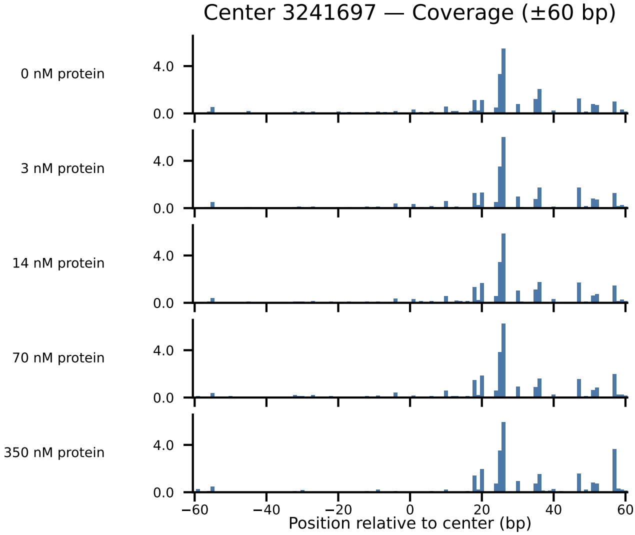
Center 3241697 — Coverage (±60 bp)
0 nM protein
3 nM protein
14 nM protein
70 nM protein
350 nM protein
4.0
0.0
−60
−40
−20
0
20
40
60
Position relative to center (bp)

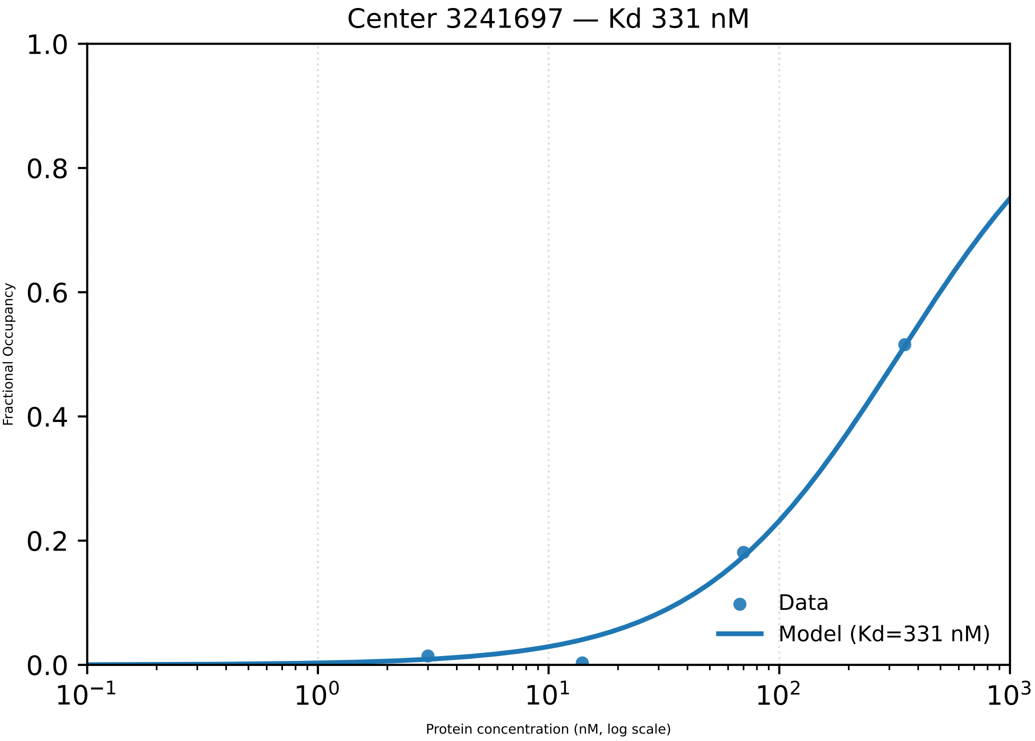

Center 3241697 — Kd 331 nM
Fractional Occupancy
1.0
0.8
0.6
0.4
0.2
0.0
$10^{-1}$
$10^{0}$
$10^{1}$
$10^{2}$
$10^{3}$
Protein concentration (nM, log scale)
Data
Model (Kd=331 nM)

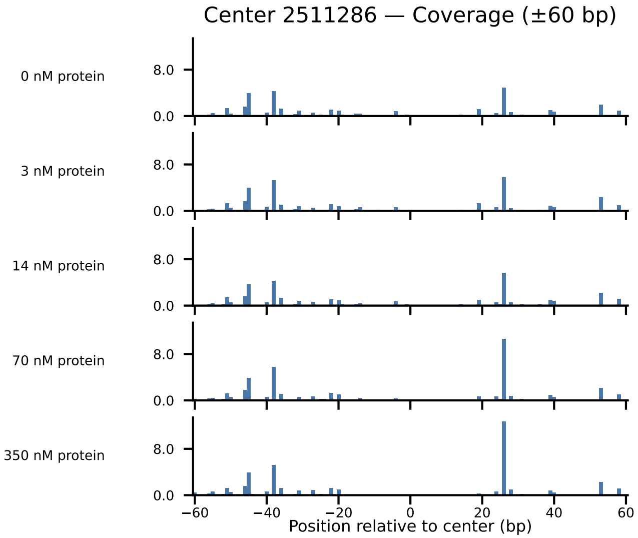
Center 2511286 — Coverage (±60 bp)
0 nM protein
3 nM protein
14 nM protein
70 nM protein
350 nM protein
8.0
0.0
−60
−40
−20
0
20
40
60
Position relative to center (bp)

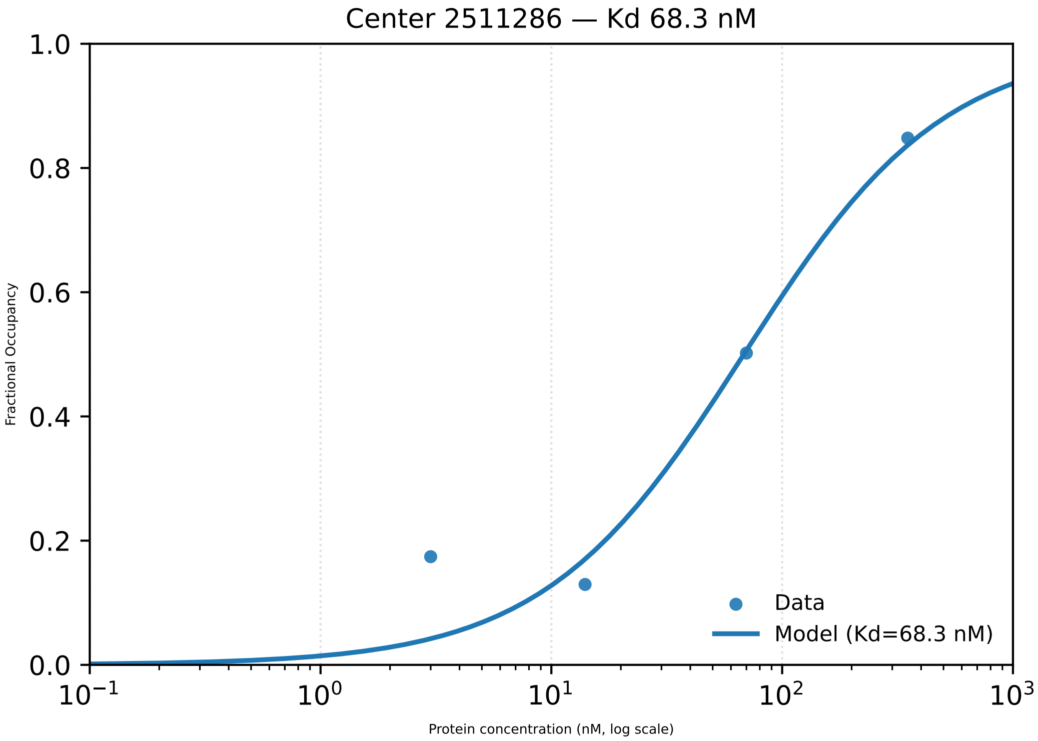
Center 2511286 — Kd 68.3 nM
1.0
0.8
0.6
0.4
0.2
0.0
Fractional Occupancy
$10^{-1}$
$10^{0}$
$10^{1}$
$10^{2}$
$10^{3}$
Protein concentration (nM, log scale)
Data
Model (Kd=68.3 nM)

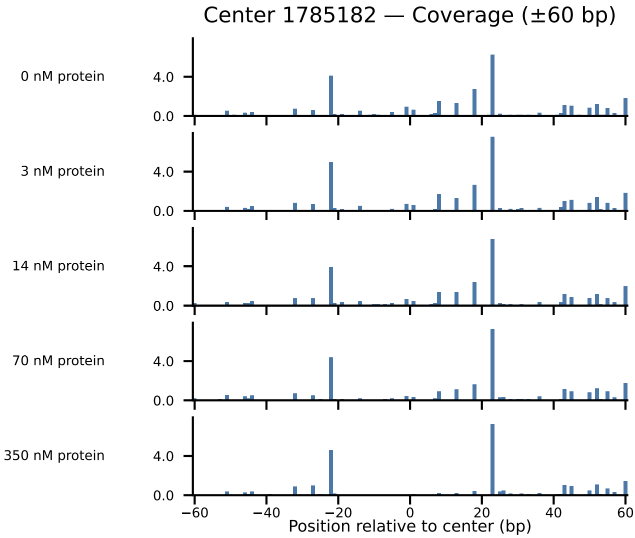

Center 1785182 — Coverage (±60 bp)
0 nM protein
3 nM protein
14 nM protein
70 nM protein
350 nM protein
4.0
0.0
−60
−40
−20
0
20
40
60
Position relative to center (bp)

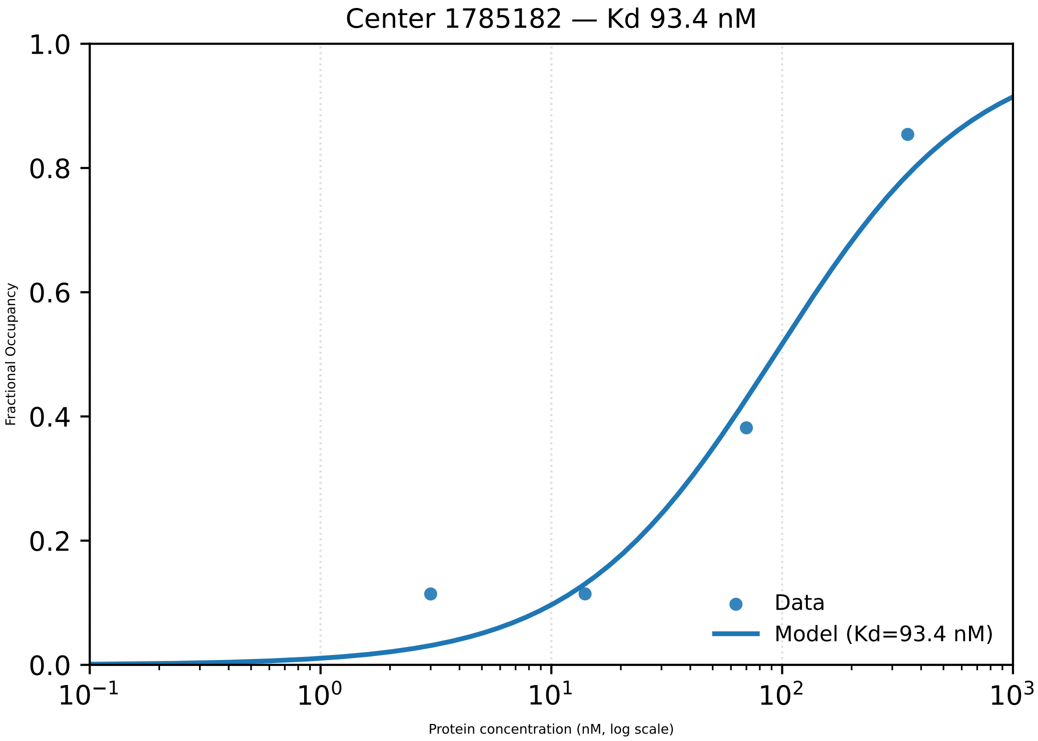

Center 1785182 — Kd 93.4 nM
Fractional Occupancy
1.0
0.8
0.6
0.4
0.2
0.0
$10^{-1}$
$10^{0}$
$10^{1}$
$10^{2}$
$10^{3}$
Protein concentration (nM, log scale)
Data
Model (Kd=93.4 nM)

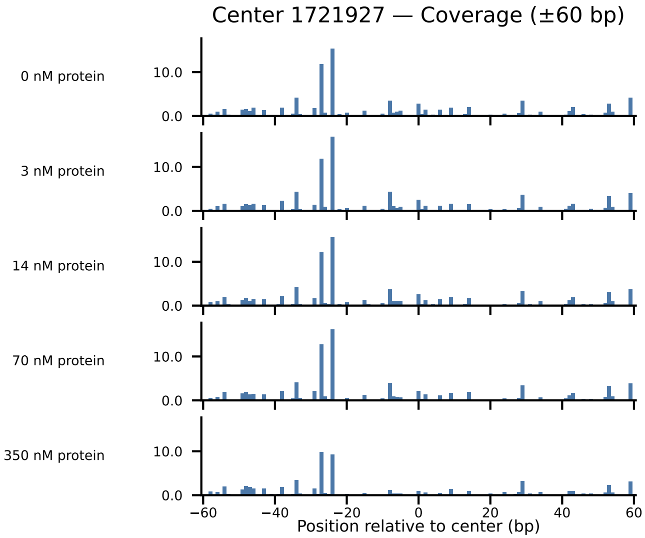

Center 1721927 — Coverage (±60 bp)
0 nM protein
3 nM protein
14 nM protein
70 nM protein
350 nM protein
10.0
0.0
−60
−40
−20
0
20
40
60
Position relative to center (bp)

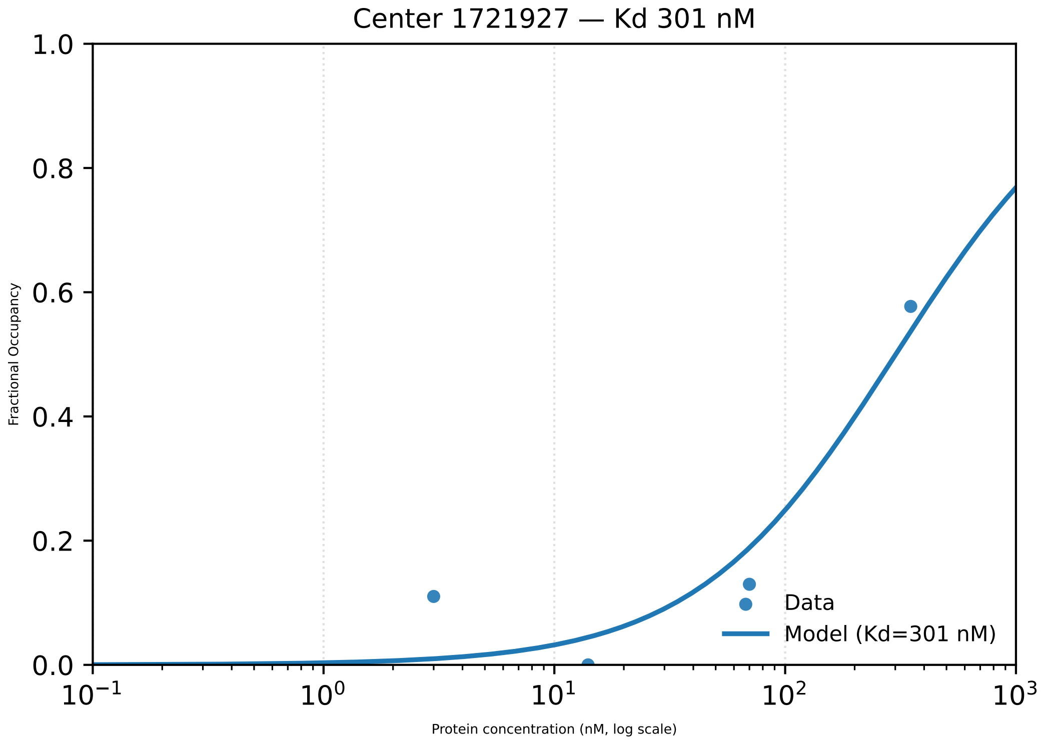

Center 1721927 — Kd 301 nM
Fractional Occupancy
1.0
0.8
0.6
0.4
0.2
0.0
$10^{-1}$
$10^{0}$
$10^{1}$
$10^{2}$
$10^{3}$
Protein concentration (nM, log scale)
Data
Model (Kd=301 nM)

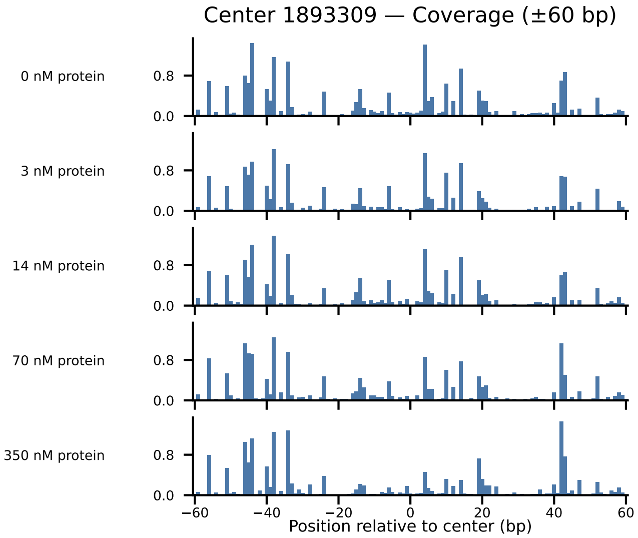
Center 1893309 — Coverage (±60 bp)
0 nM protein
3 nM protein
14 nM protein
70 nM protein
350 nM protein
0.8
0.0
−60
−40
−20
0
20
40
60
Position relative to center (bp)

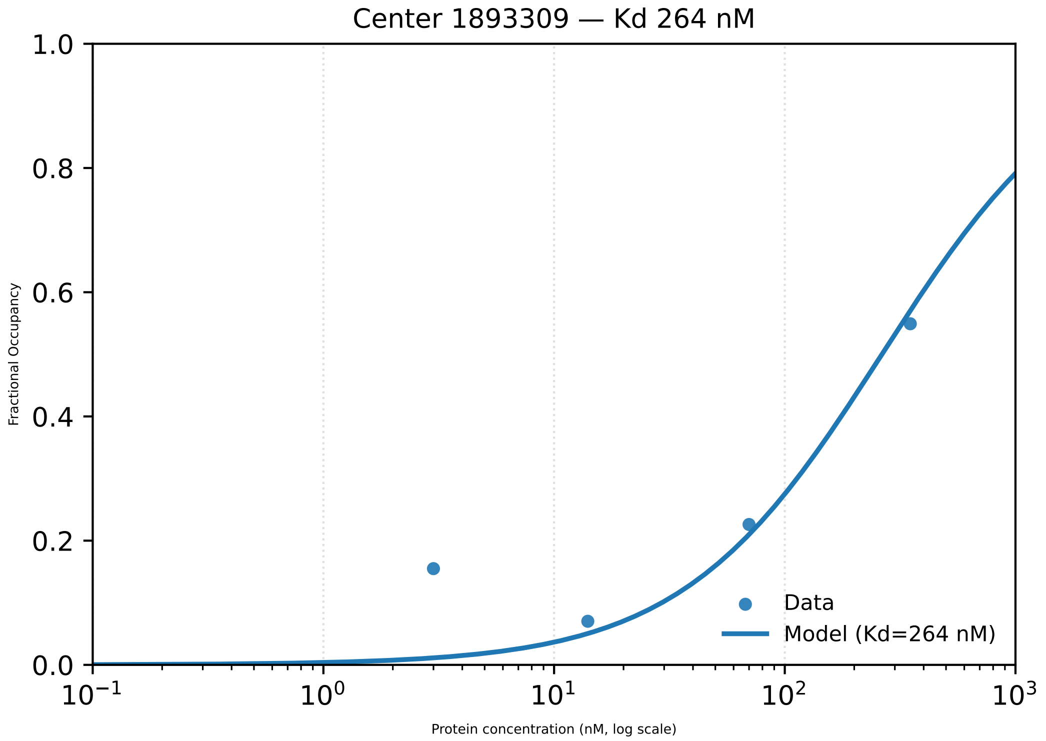
Center 1893309 — Kd 264 nM
Fractional Occupancy
Protein concentration (nM, log scale)
Data
Model (Kd=264 nM)
1.0
0.8
0.6
0.4
0.2
0.0
$10^{-1}$
$10^{0}$
$10^{1}$
$10^{2}$
$10^{3}$

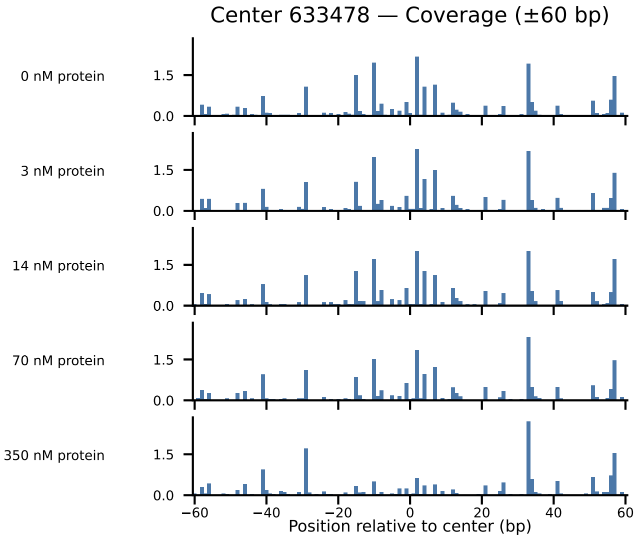
Center 633478 — Coverage (±60 bp)
0 nM protein
3 nM protein
14 nM protein
70 nM protein
350 nM protein
1.5
0.0
−60
−40
−20
0
20
40
60
Position relative to center (bp)

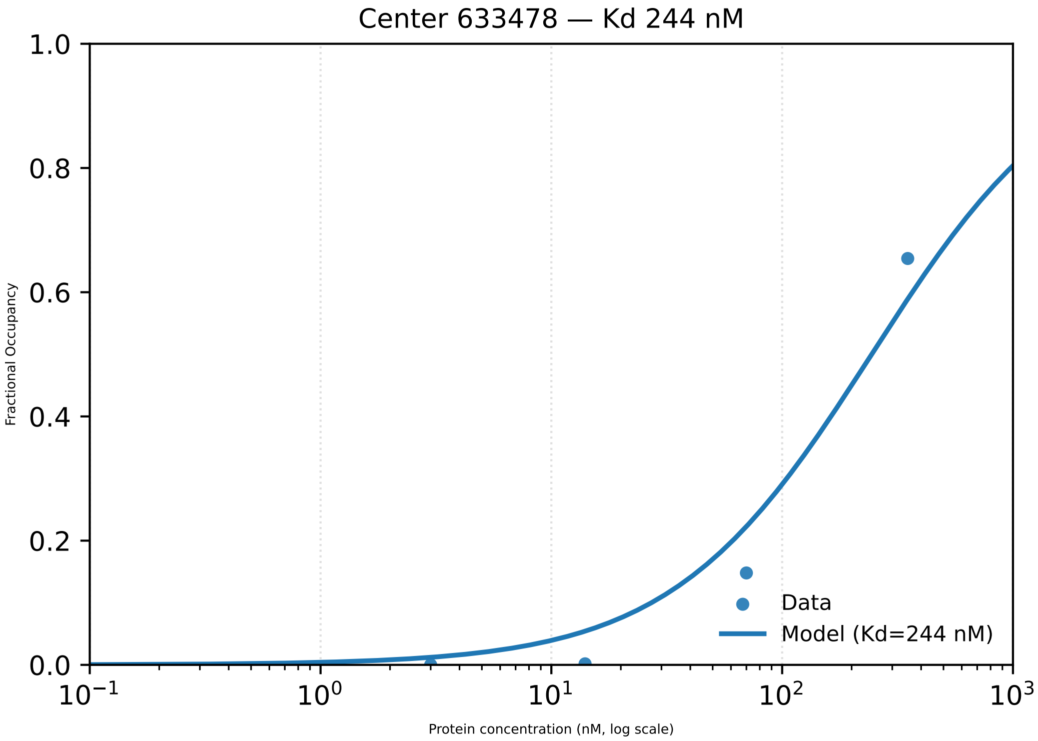

Center 633478 — Kd 244 nM
Fractional Occupancy
1.0
0.8
0.6
0.4
0.2
0.0
$10^{-1}$
$10^{0}$
$10^{1}$
$10^{2}$
$10^{3}$
Protein concentration (nM, log scale)
Data
Model (Kd=244 nM)

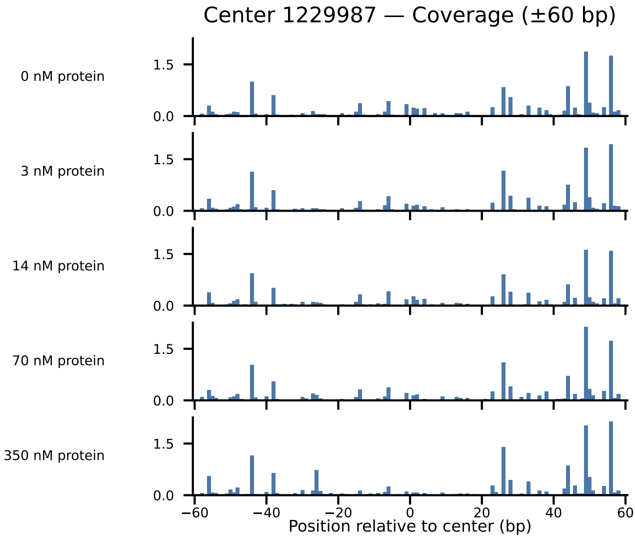
Center 1229987 — Coverage (±60 bp)
0 nM protein
3 nM protein
14 nM protein
70 nM protein
350 nM protein
1.5
0.0
−60
−40
−20
0
20
40
60
Position relative to center (bp)

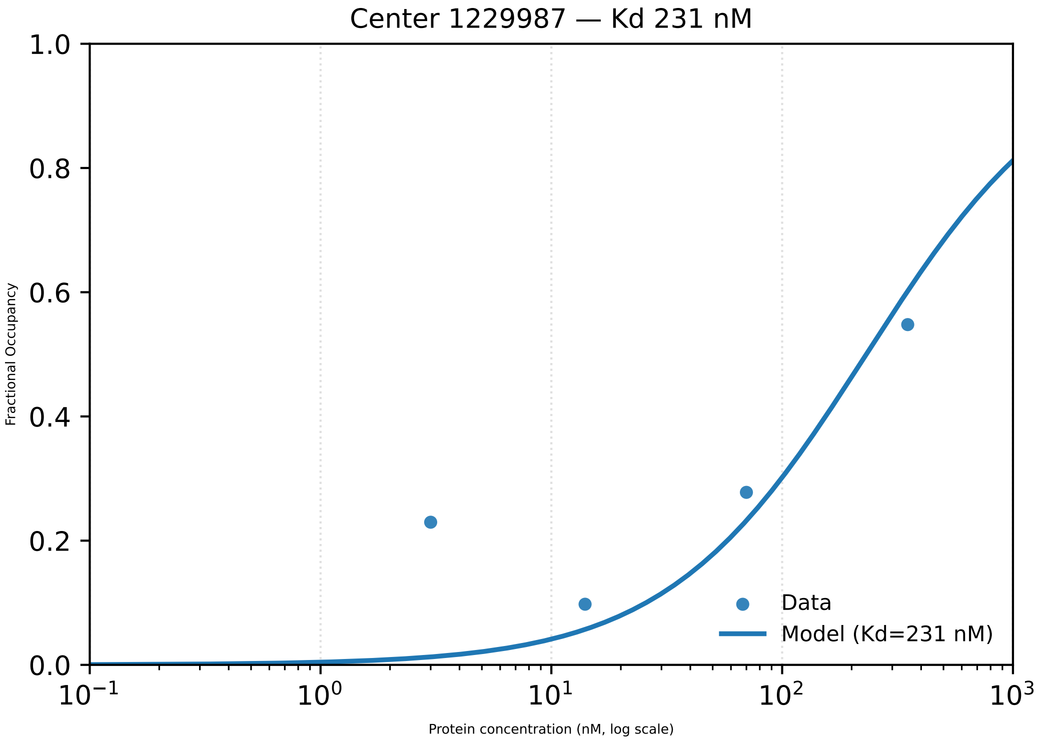

Center 1229987 — Kd 231 nM
1.0
0.8
0.6
0.4
0.2
0.0
Fractional Occupancy
$10^{-1}$
$10^{0}$
$10^{1}$
$10^{2}$
$10^{3}$
Protein concentration (nM, log scale)
Data
Model (Kd=231 nM)

Center 4081632 — Coverage (±60 bp)

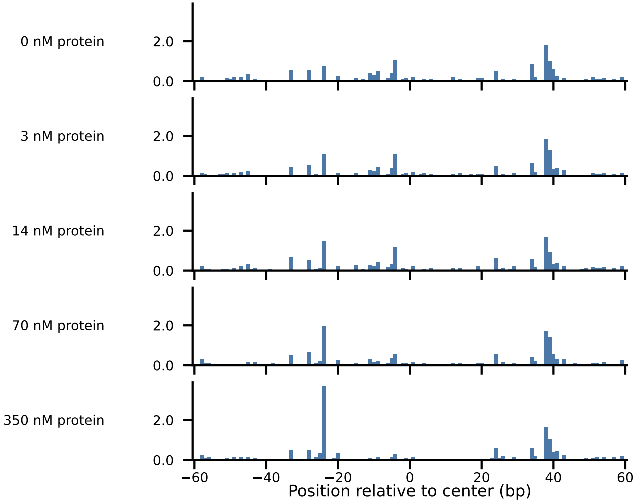

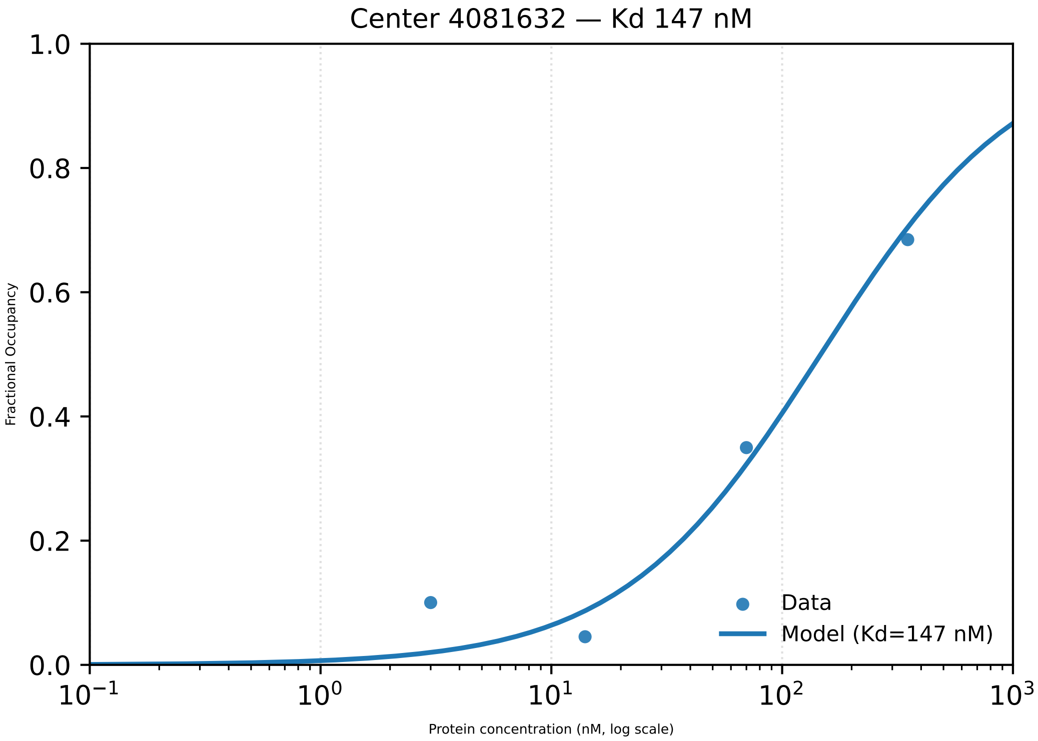

Center 4081632 — Kd 147 nM
Fractional Occupancy
1.0
0.8
0.6
0.4
0.2
0.0
$10^{-1}$
$10^{0}$
$10^{1}$
$10^{2}$
$10^{3}$
Protein concentration (nM, log scale)
Data
Model (Kd=147 nM)

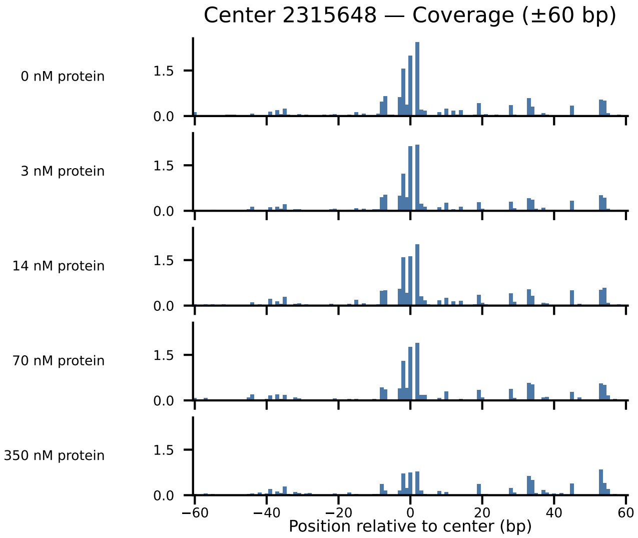

Center 2315648 — Coverage (±60 bp)
0 nM protein
3 nM protein
14 nM protein
70 nM protein
350 nM protein
1.5
0.0
−60
−40
−20
0
20
40
60
Position relative to center (bp)

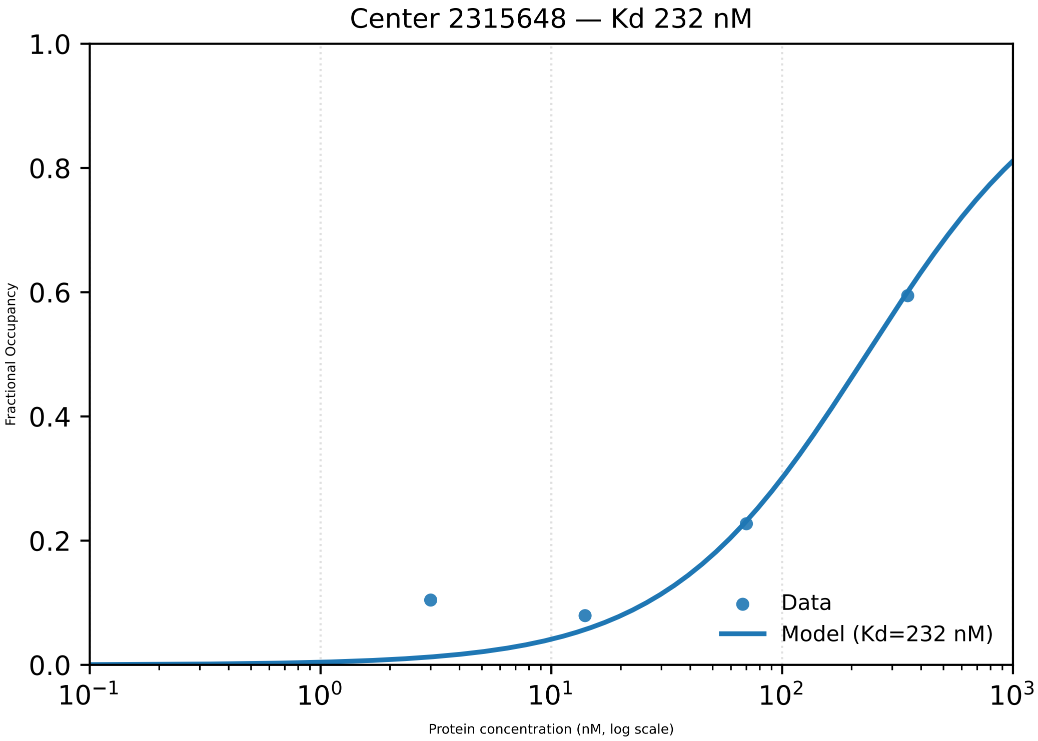
Center 2315648 — Kd 232 nM
1.0
0.8
0.6
0.4
0.2
0.0
Fractional Occupancy
$10^{-1}$
$10^{0}$
$10^{1}$
$10^{2}$
$10^{3}$
Protein concentration (nM, log scale)
Data
Model (Kd=232 nM)

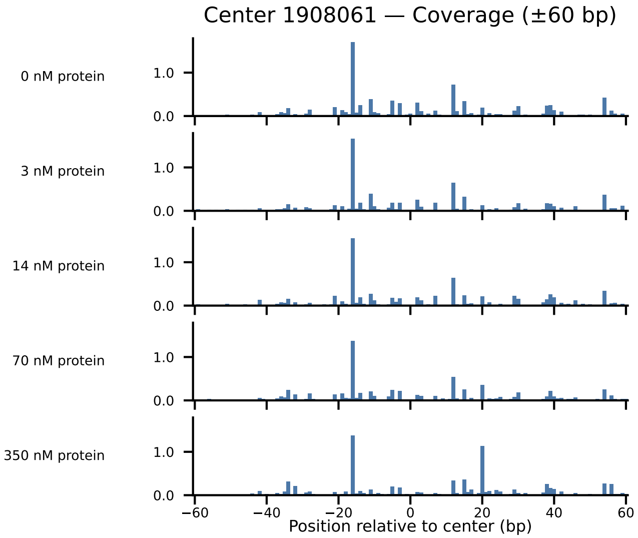
Center 1908061 — Coverage (±60 bp)
0 nM protein
3 nM protein
14 nM protein
70 nM protein
350 nM protein
1.0
0.0
−60
−40
−20
0
20
40
60
Position relative to center (bp)

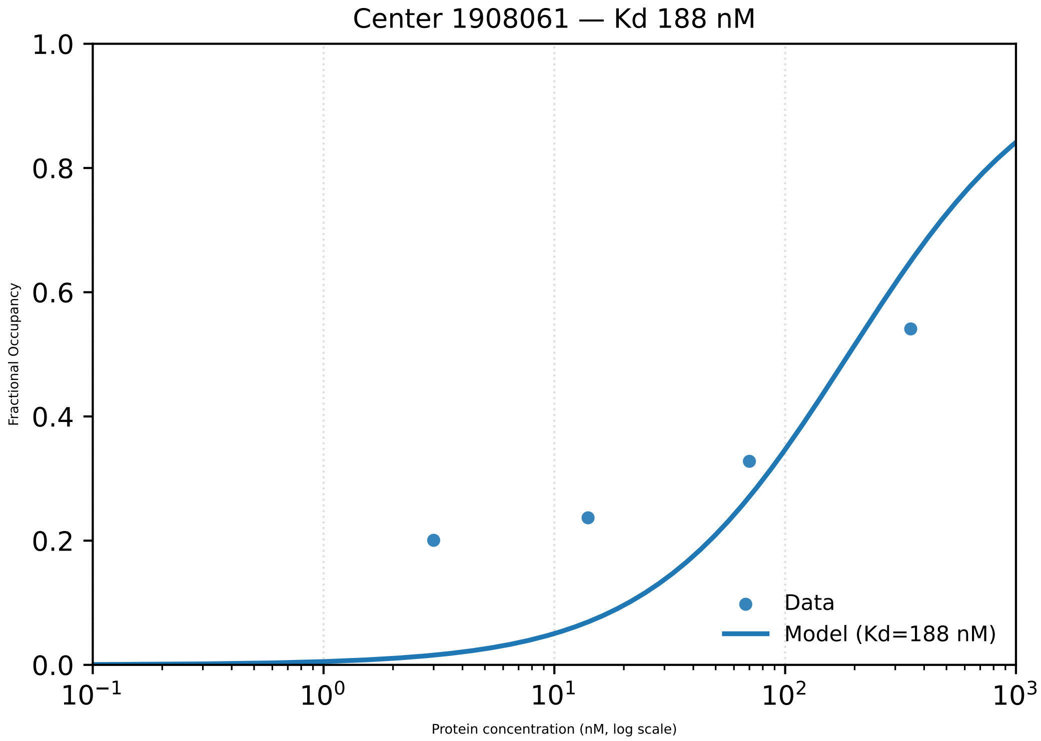

Center 1908061 — Kd 188 nM
1.0
0.8
0.6
0.4
0.2
0.0
Fractional Occupancy
$10^{-1}$
$10^{0}$
$10^{1}$
$10^{2}$
$10^{3}$
Protein concentration (nM, log scale)
Data
Model (Kd=188 nM)

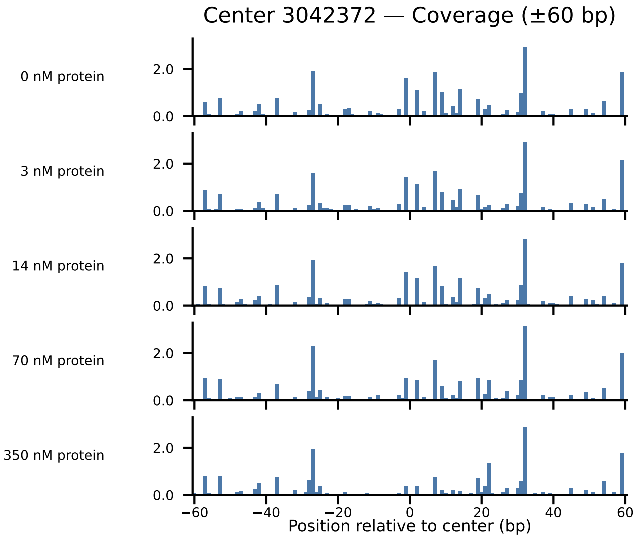
Center 3042372 — Coverage (±60 bp)
0 nM protein
3 nM protein
14 nM protein
70 nM protein
350 nM protein
2.0
0.0
−60
−40
−20
0
20
40
60
Position relative to center (bp)

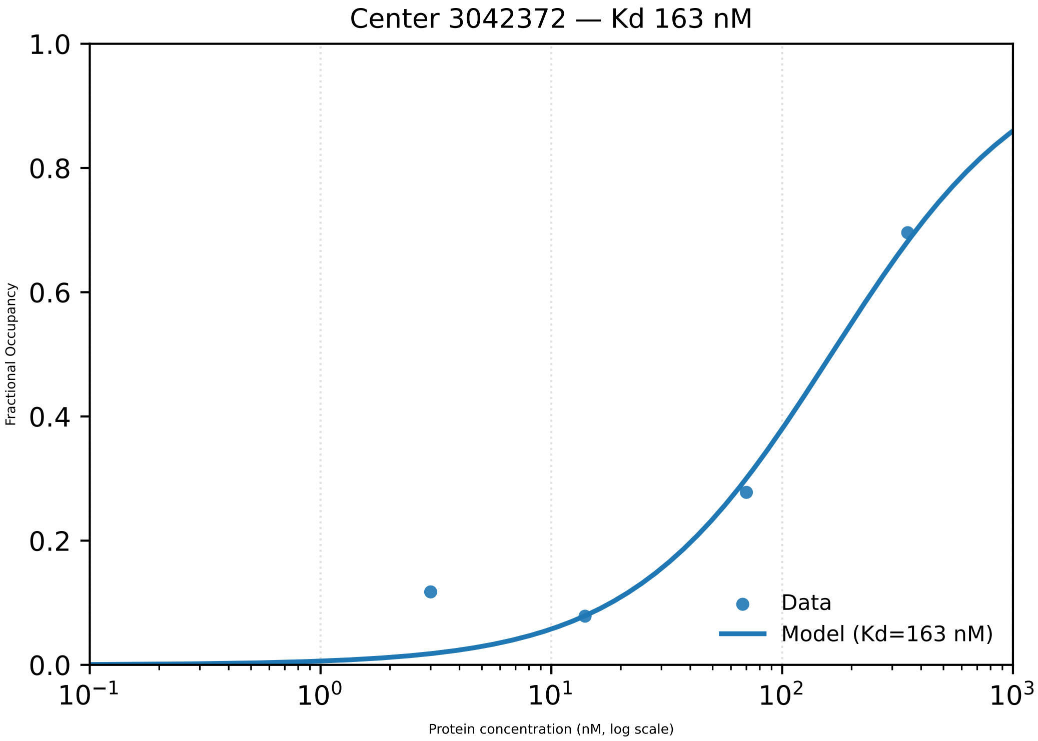

Center 3042372 — Kd 163 nM
1.0
0.8
0.6
0.4
0.2
0.0
Fractional Occupancy
$10^{-1}$
$10^{0}$
$10^{1}$
$10^{2}$
$10^{3}$
Protein concentration (nM, log scale)
Data
Model (Kd=163 nM)

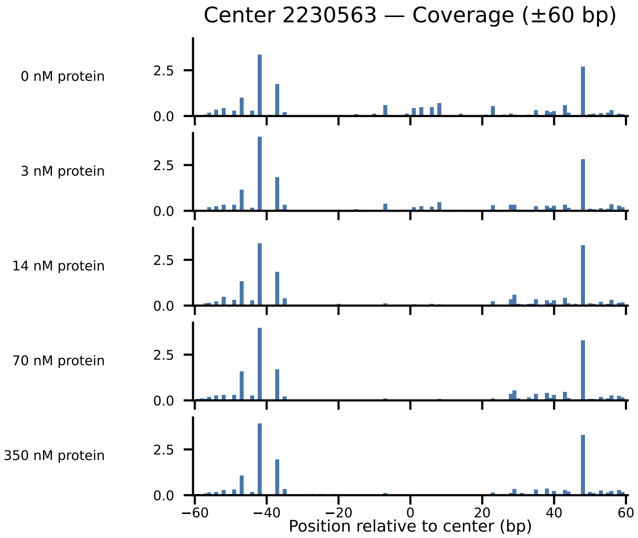
Center 2230563 — Coverage (±60 bp)
0 nM protein
3 nM protein
14 nM protein
70 nM protein
350 nM protein
2.5
0.0
−60
−40
−20
0
20
40
60
Position relative to center (bp)

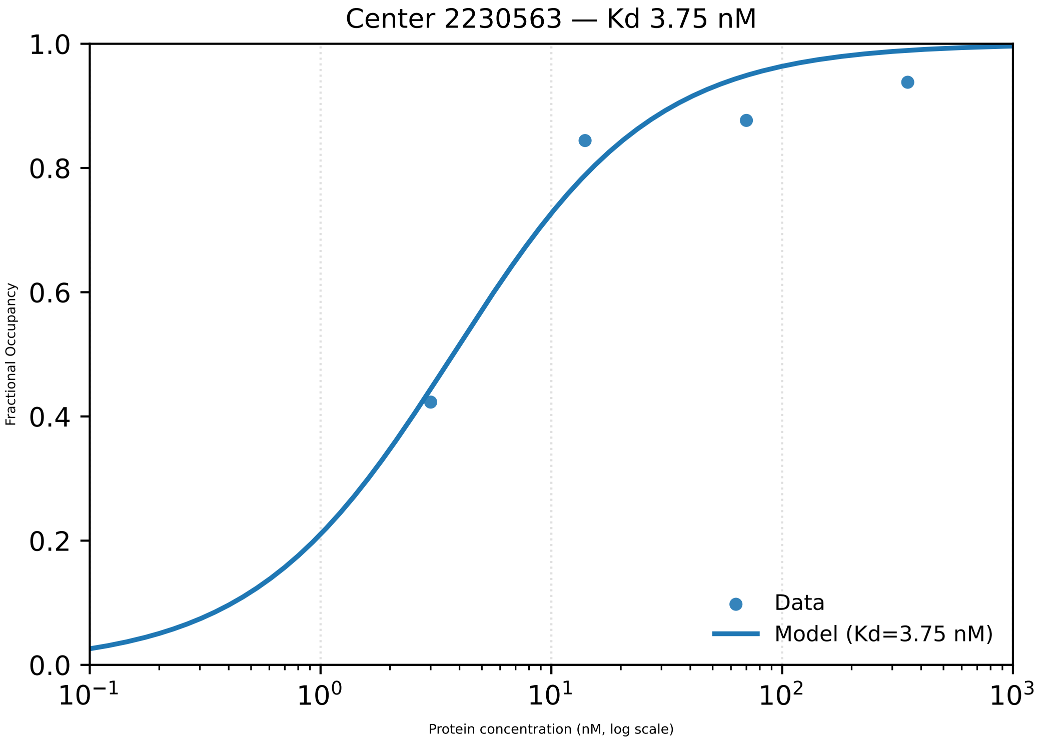

Center 2230563 — Kd 3.75 nM
Fractional Occupancy
1.0
0.8
0.6
0.4
0.2
0.0
$10^{-1}$
$10^{0}$
$10^{1}$
$10^{2}$
$10^{3}$
Protein concentration (nM, log scale)
Data
Model (Kd=3.75 nM)

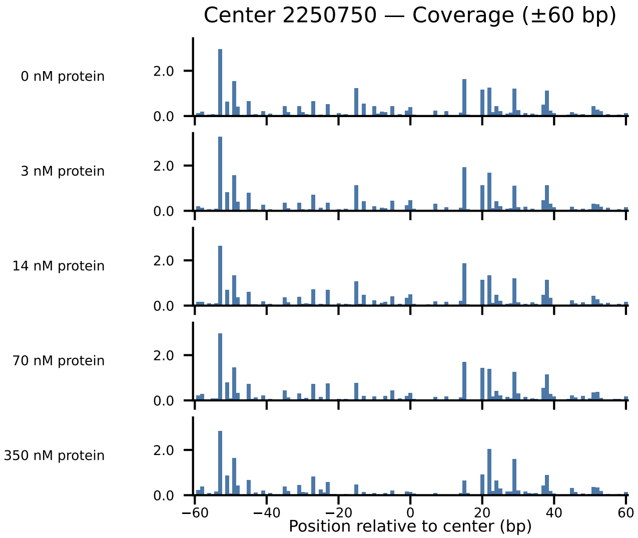
Center 2250750 — Coverage (±60 bp)
0 nM protein
3 nM protein
14 nM protein
70 nM protein
350 nM protein
2.0
0.0
−60
−40
−20
0
20
40
60
Position relative to center (bp)

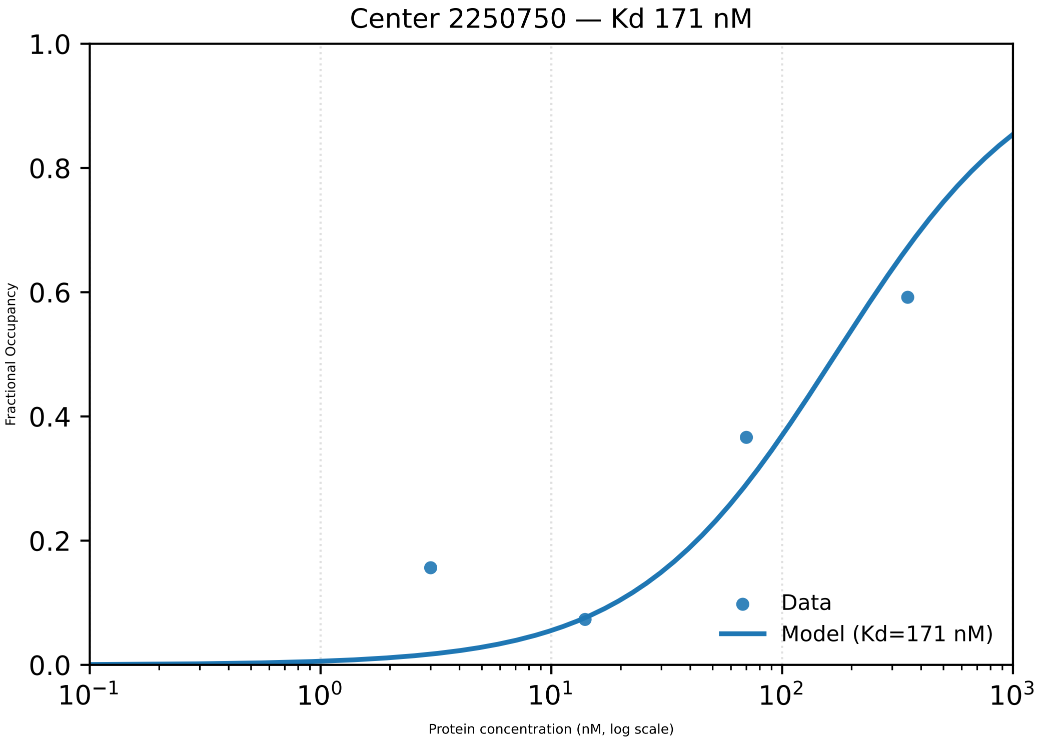
Center 2250750 — Kd 171 nM
1.0
0.8
0.6
0.4
0.2
0.0
Fractional Occupancy
$10^{-1}$
$10^{0}$
$10^{1}$
$10^{2}$
$10^{3}$
Protein concentration (nM, log scale)
Data
Model (Kd=171 nM)

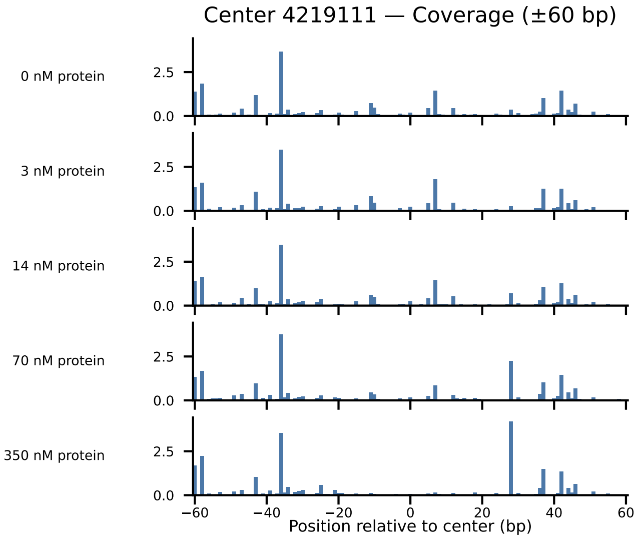

Center 4219111 — Coverage (±60 bp)
0 nM protein
3 nM protein
14 nM protein
70 nM protein
350 nM protein
2.5
0.0
−60
−40
−20
0
20
40
60
Position relative to center (bp)

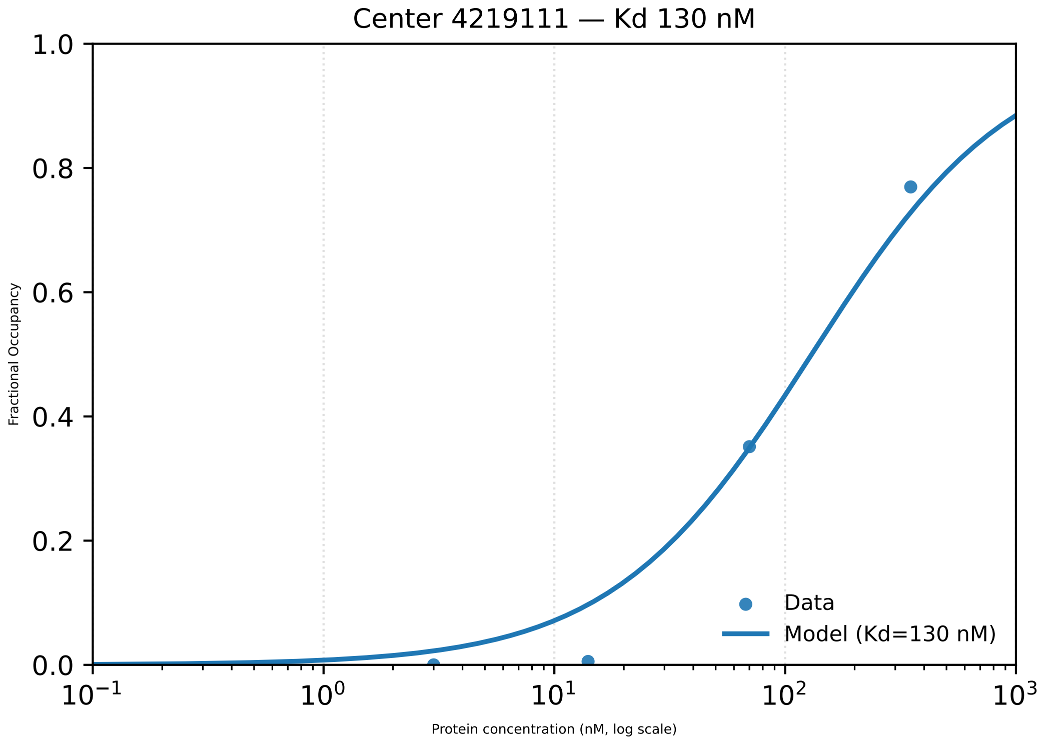

Center 4219111 — Kd 130 nM
Fractional Occupancy
1.0
0.8
0.6
0.4
0.2
0.0
$10^{-1}$
$10^{0}$
$10^{1}$
$10^{2}$
$10^{3}$
Protein concentration (nM, log scale)
Data
Model (Kd=130 nM)

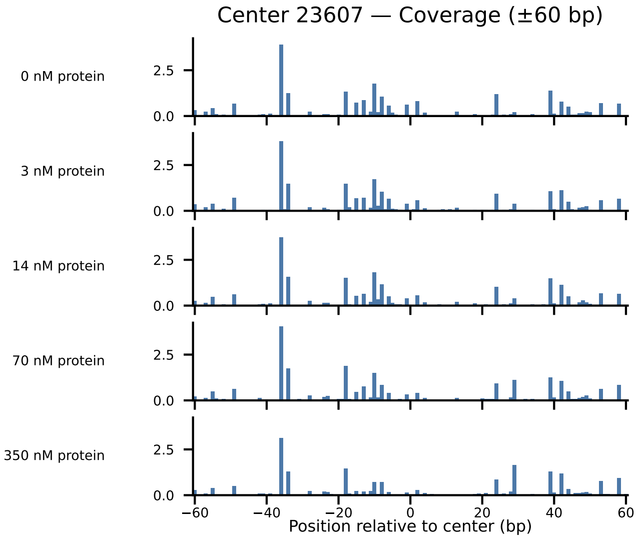

Center 23607 — Coverage (±60 bp)
0 nM protein
3 nM protein
14 nM protein
70 nM protein
350 nM protein
0.0
2.5
−60
−40
−20
0
20
40
60
Position relative to center (bp)

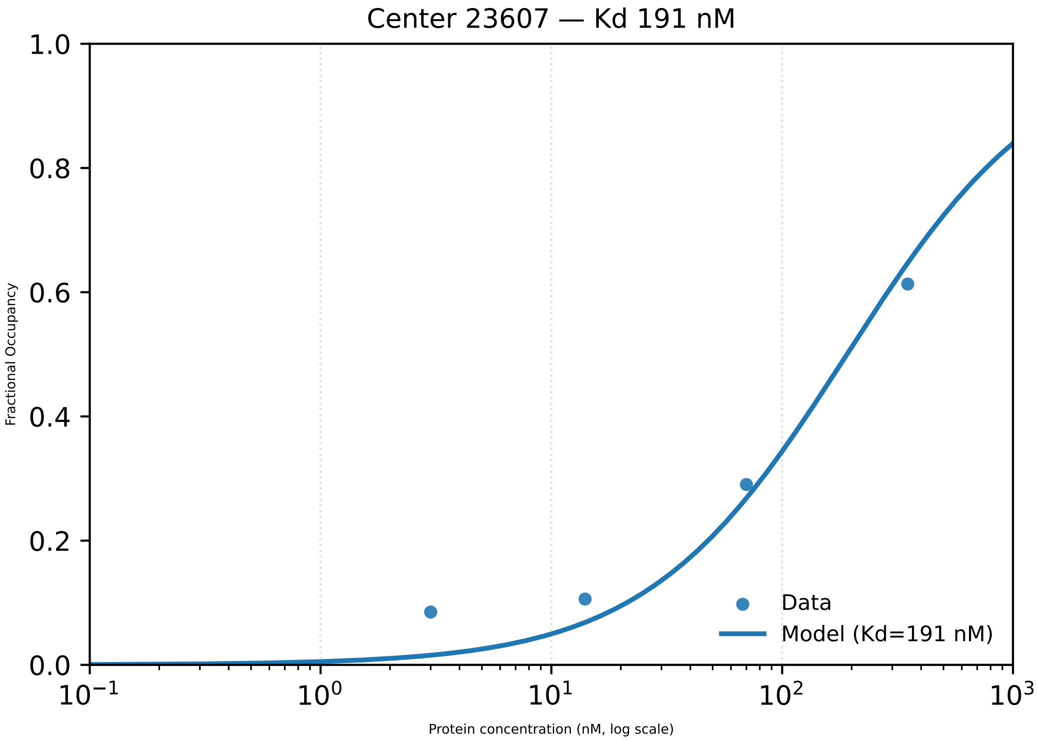
Center 23607 — Kd 191 nM
Fractional Occupancy
Protein concentration (nM, log scale)
1.0
0.8
0.6
0.4
0.2
0.0
$10^{-1}$
$10^{0}$
$10^{1}$
$10^{2}$
$10^{3}$
Data
Model (Kd=191 nM)

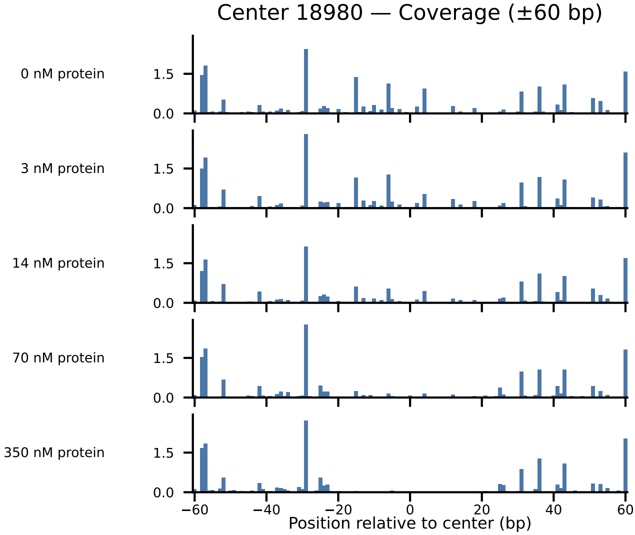
Center 18980 — Coverage (±60 bp)
0 nM protein
3 nM protein
14 nM protein
70 nM protein
350 nM protein
1.5
0.0
−60
−40
−20
0
20
40
60
Position relative to center (bp)

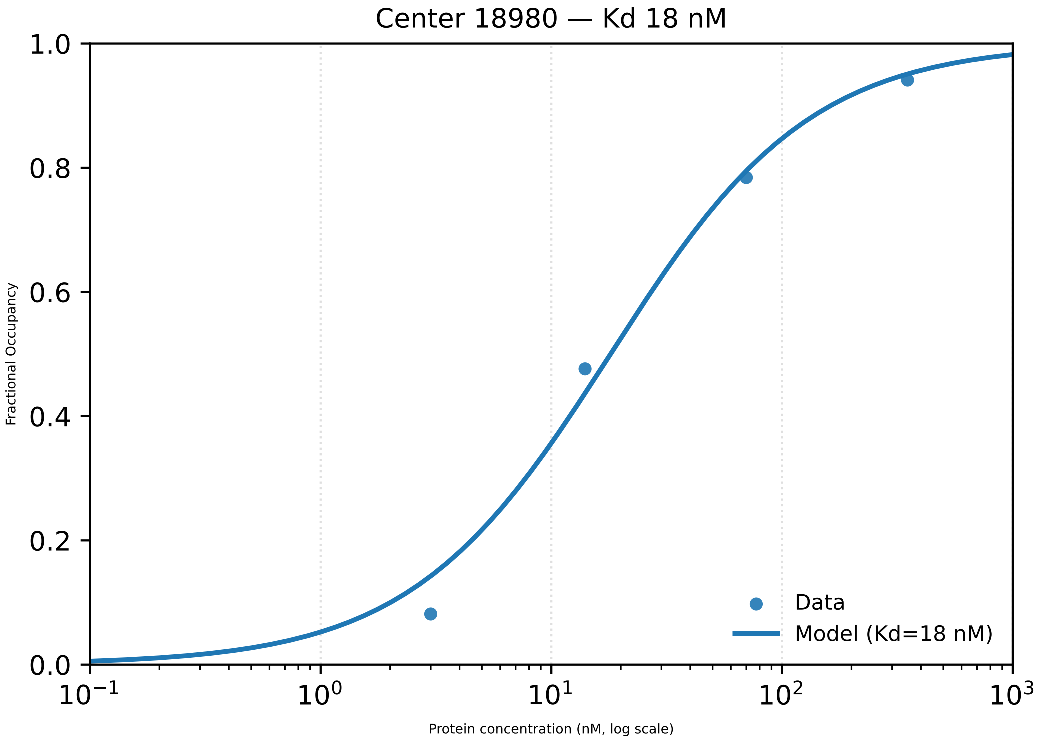

Center 18980 — Kd 18 nM
1.0
0.8
0.6
0.4
0.2
0.0
Fractional Occupancy
$10^{-1}$
$10^{0}$
$10^{1}$
$10^{2}$
$10^{3}$
Protein concentration (nM, log scale)
Data
Model (Kd=18 nM)
