## Supplementary material for "Footprint-seq: a simple method to quantitatively map *in vitro* protein-DNA interactions on a genome-wide scale at high spatial resolution": Figure S10

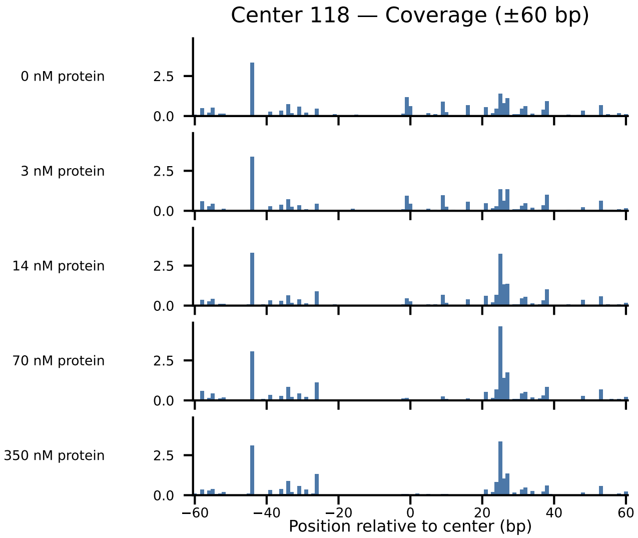

Center 118 — Coverage (±60 bp)
0 nM protein
3 nM protein
14 nM protein
70 nM protein
350 nM protein
2.5
0.0
−60
−40
−20
0
20
40
60
Position relative to center (bp)

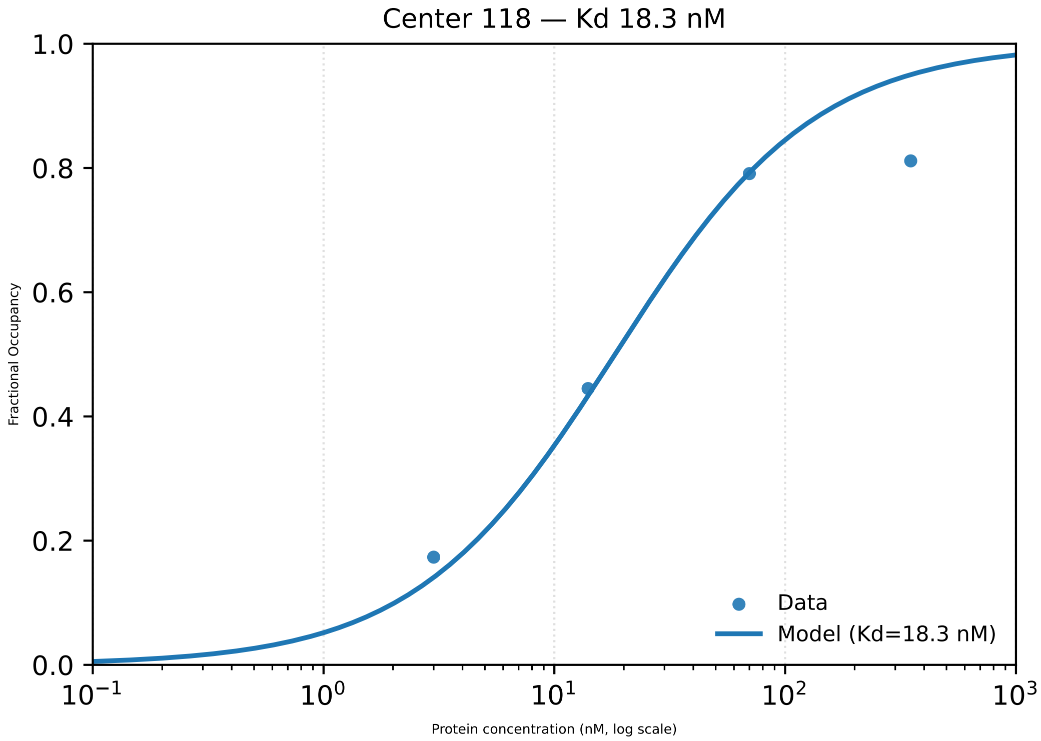

Center 118 — Kd 18.3 nM
1.0
0.8
0.6
0.4
0.2
0.0
Fractional Occupancy
10^−1
10^0
10^1
10^2
10^3
Protein concentration (nM, log scale)
Data
Model (Kd=18.3 nM)

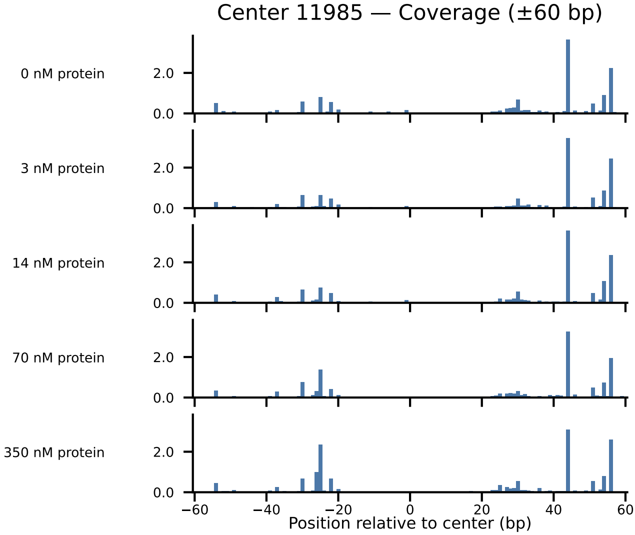
Center 11985 — Coverage (±60 bp)
0 nM protein
3 nM protein
14 nM protein
70 nM protein
350 nM protein
2.0
0.0
−60
−40
−20
0
20
40
60
Position relative to center (bp)

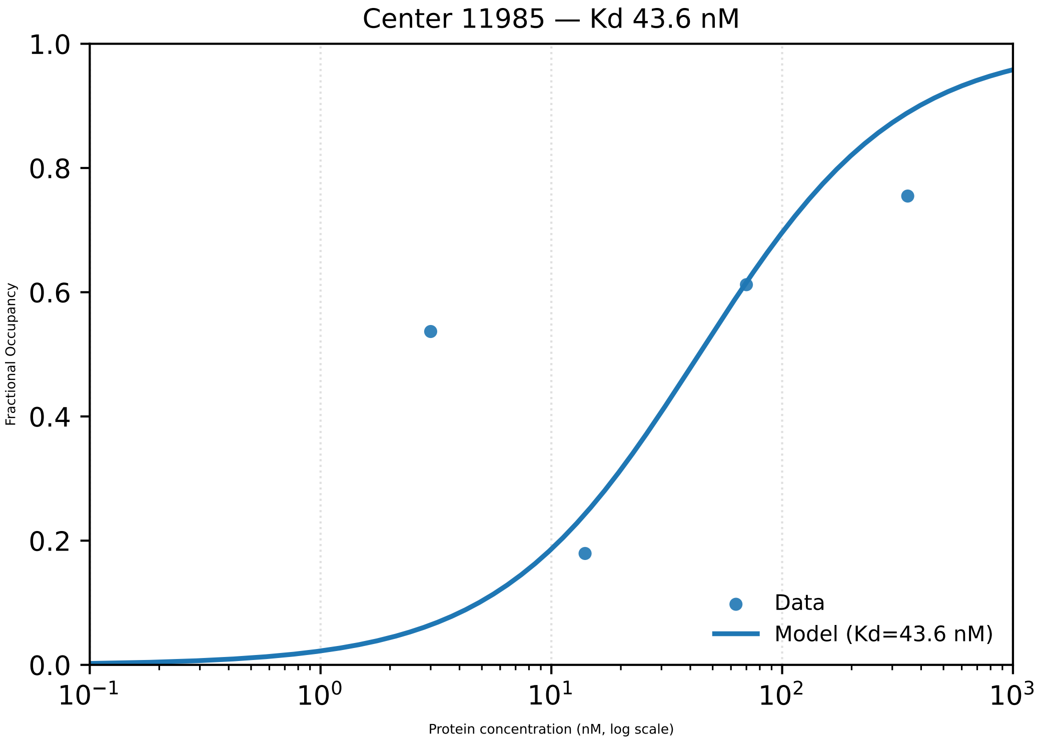
Center 11985 — Kd 43.6 nM
Fractional Occupancy
1.0
0.8
0.6
0.4
0.2
0.0
$10^{-1}$
$10^{0}$
$10^{1}$
$10^{2}$
$10^{3}$
Protein concentration (nM, log scale)
Data
Model (Kd=43.6 nM)

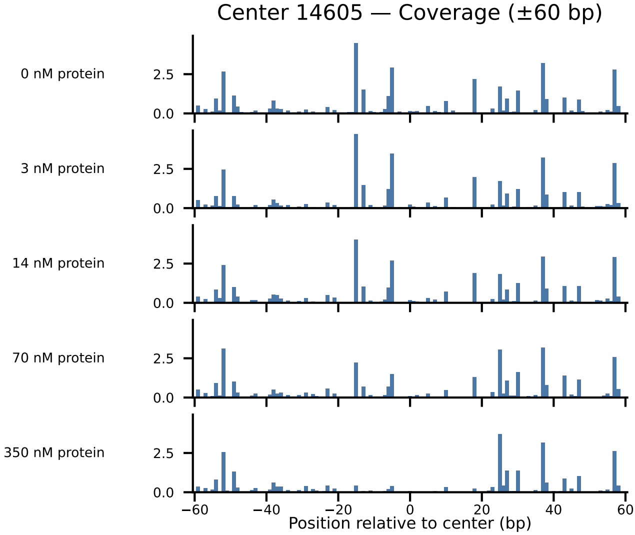

Center 14605 — Coverage (±60 bp)
0 nM protein
3 nM protein
14 nM protein
70 nM protein
350 nM protein
2.5
0.0
−60
−40
−20
0
20
40
60
Position relative to center (bp)

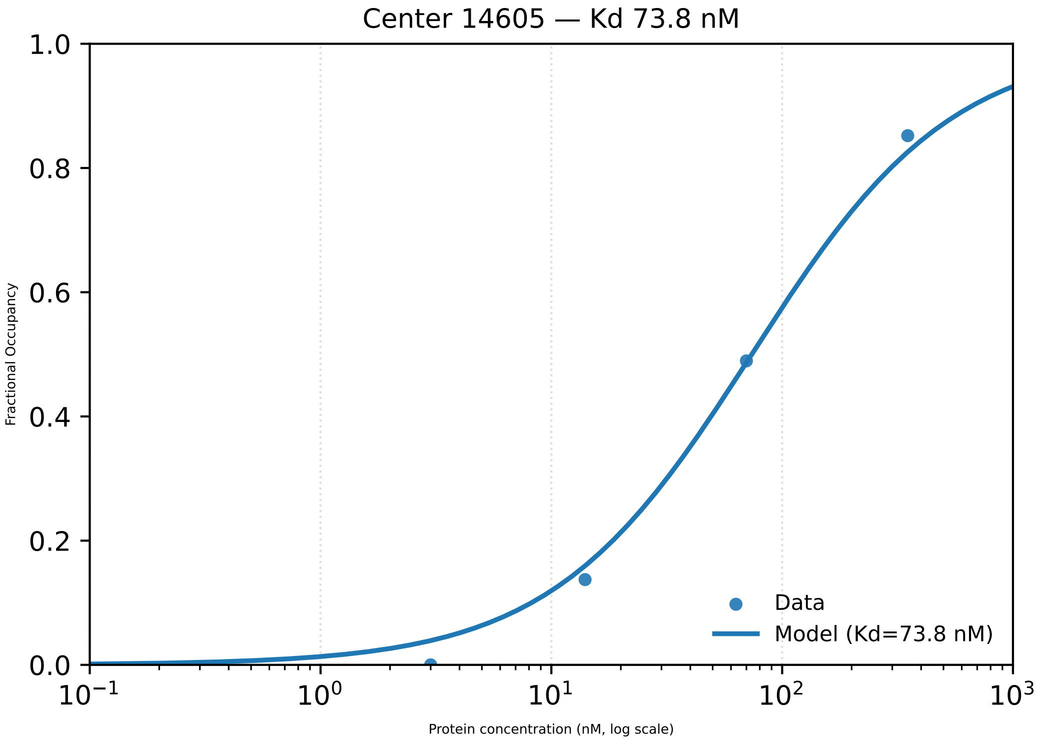

Center 14605 — Kd 73.8 nM
Fractional Occupancy
1.0
0.8
0.6
0.4
0.2
0.0
$10^{-1}$
$10^{0}$
$10^{1}$
$10^{2}$
$10^{3}$
Protein concentration (nM, log scale)
Data
Model (Kd=73.8 nM)

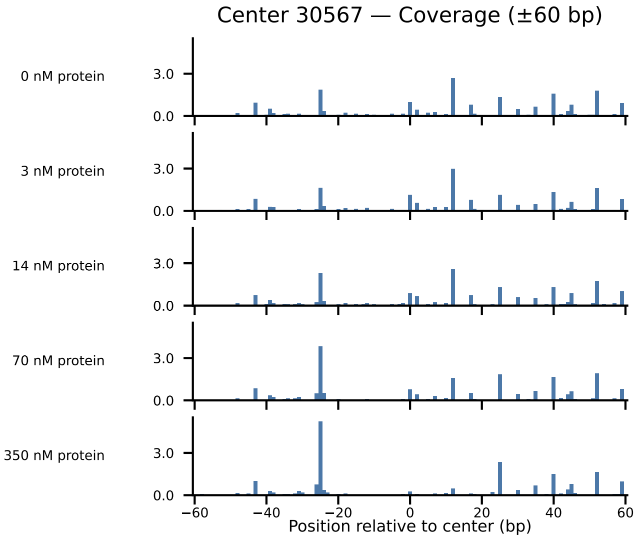

Center 30567 — Coverage (±60 bp)
0 nM protein
3 nM protein
14 nM protein
70 nM protein
350 nM protein
3.0
0.0
−60
−40
−20
0
20
40
60
Position relative to center (bp)

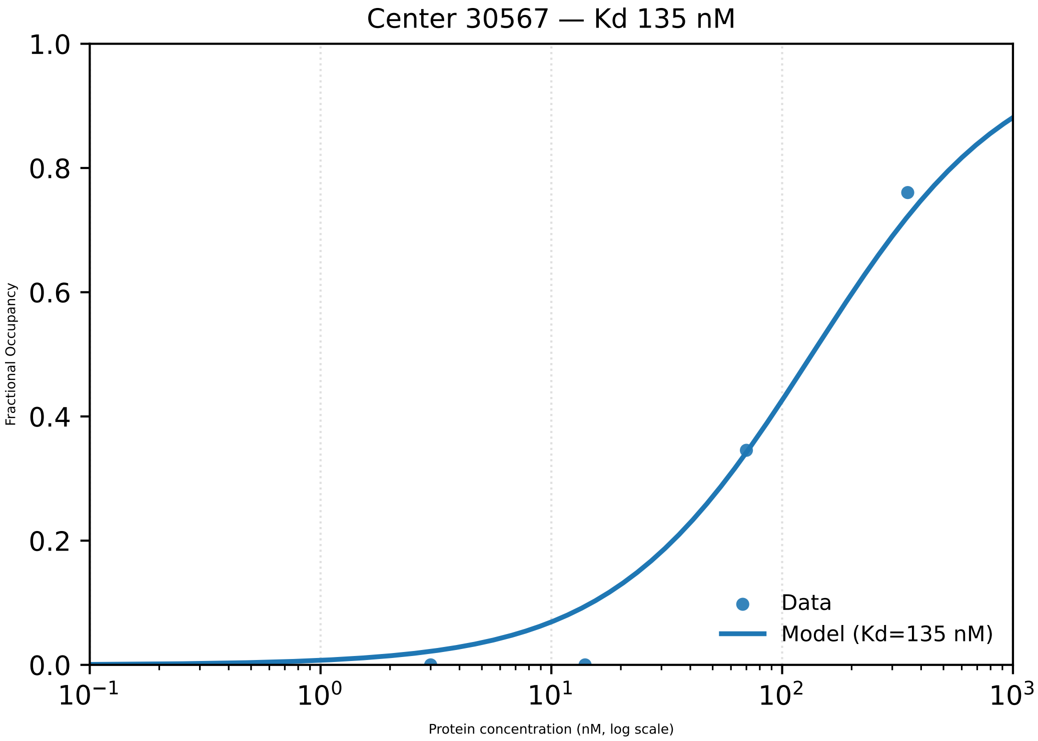
Center 30567 — Kd 135 nM
1.0
0.8
0.6
0.4
0.2
0.0
Fractional Occupancy
$10^{-1}$
$10^{0}$
$10^{1}$
$10^{2}$
$10^{3}$
Protein concentration (nM, log scale)
Data
Model (Kd=135 nM)

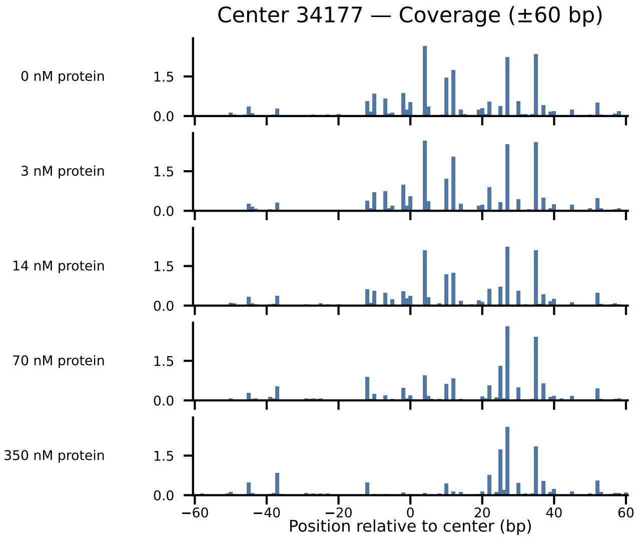
Center 34177 — Coverage (±60 bp)
0 nM protein
3 nM protein
14 nM protein
70 nM protein
350 nM protein
0.0
1.5
−60
−40
−20
0
20
40
60
Position relative to center (bp)

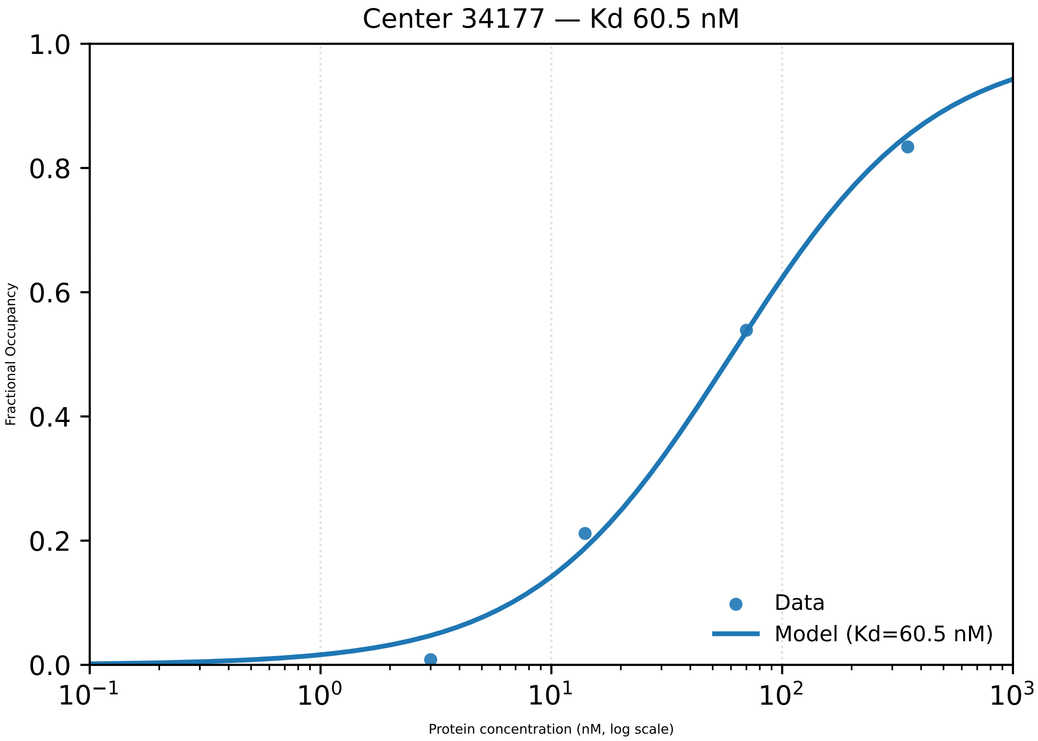
Center 34177 — Kd 60.5 nM
Fractional Occupancy
Protein concentration (nM, log scale)
Data
Model (Kd=60.5 nM)
0.0
0.2
0.4
0.6
0.8
1.0
$10^{-1}$
$10^{0}$
$10^{1}$
$10^{2}$
$10^{3}$

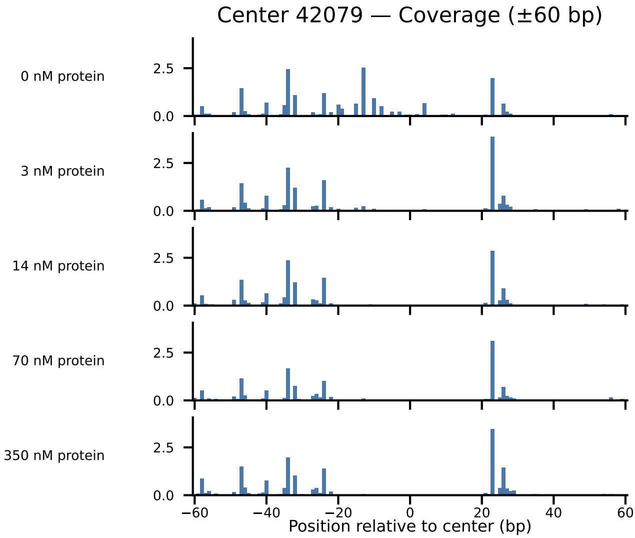

Center 42079 — Coverage (±60 bp)
0 nM protein
3 nM protein
14 nM protein
70 nM protein
350 nM protein
2.5
0.0
−60
−40
−20
0
20
40
60
Position relative to center (bp)

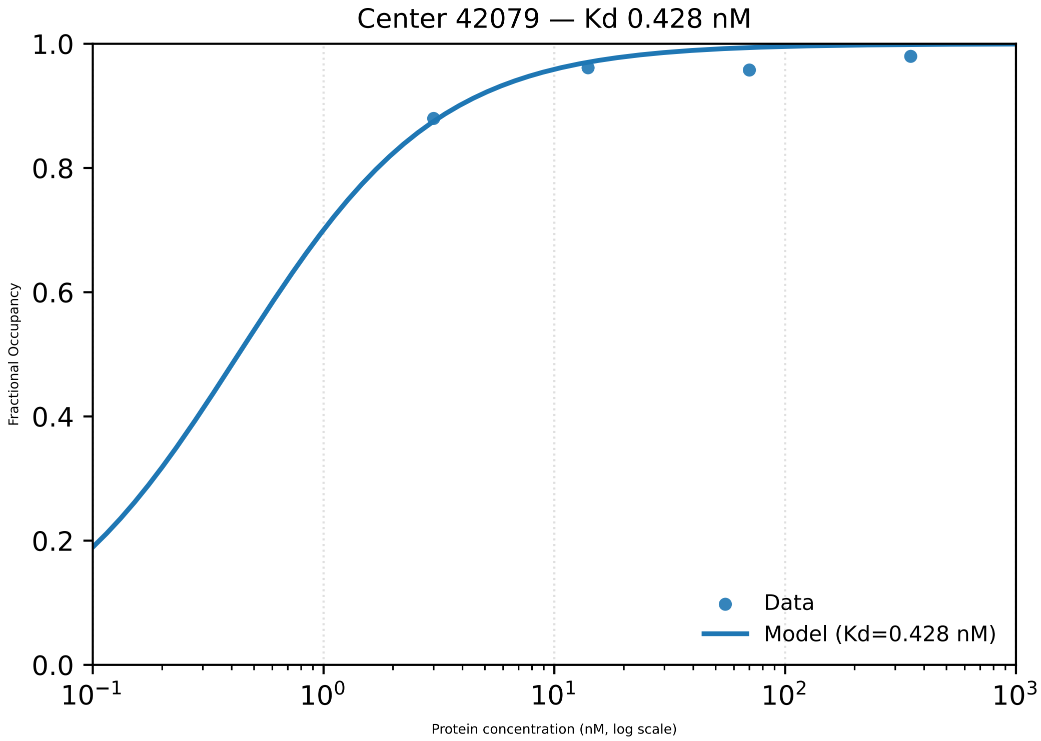

Center 42079 — Kd 0.428 nM
Fractional Occupancy
Protein concentration (nM, log scale)
1.0
0.8
0.6
0.4
0.2
0.0
$10^{-1}$
$10^{0}$
$10^{1}$
$10^{2}$
$10^{3}$
Data
Model (Kd=0.428 nM)

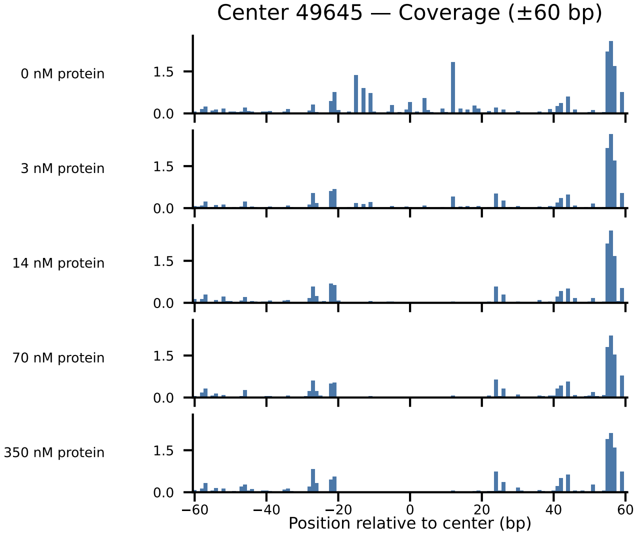
Center 49645 — Coverage (±60 bp)
0 nM protein
3 nM protein
14 nM protein
70 nM protein
350 nM protein
1.5
0.0
−60
−40
−20
0
20
40
60
Position relative to center (bp)

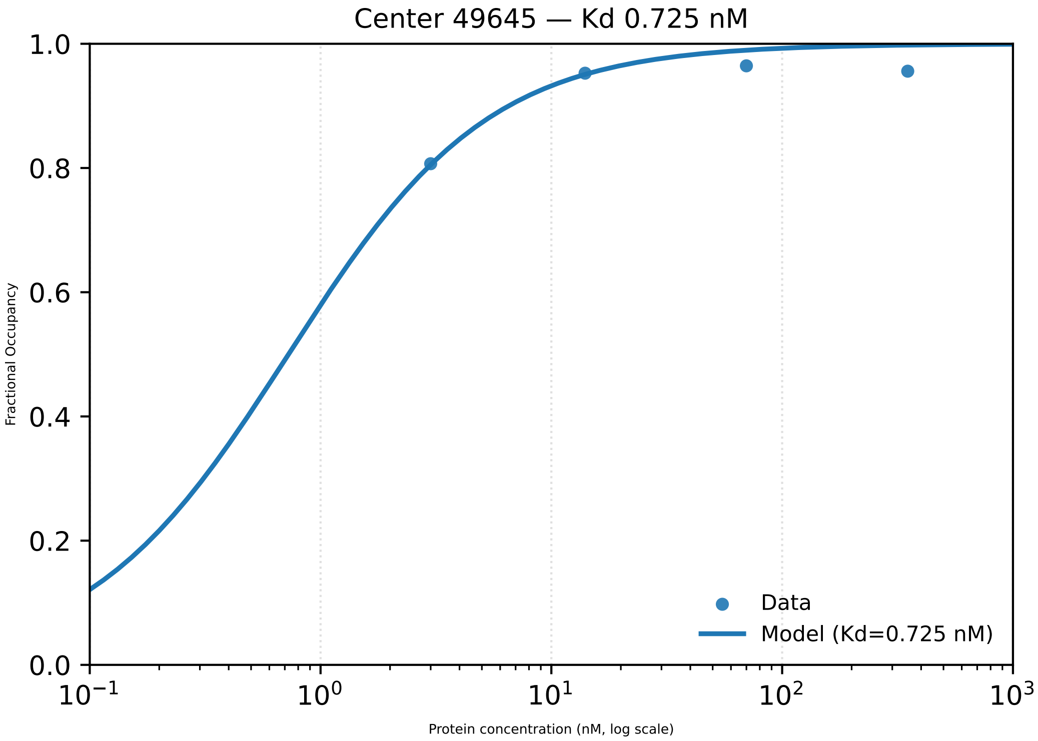

Center 49645 — Kd 0.725 nM
1.0
0.8
0.6
0.4
0.2
0.0
Fractional Occupancy
$10^{-1}$
$10^{0}$
$10^{1}$
$10^{2}$
$10^{3}$
Protein concentration (nM, log scale)
Data
Model (Kd=0.725 nM)

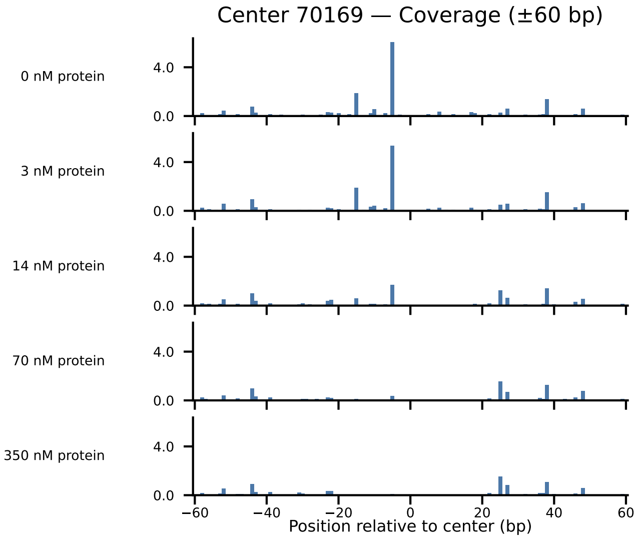

Center 70169 — Coverage (±60 bp)
0 nM protein
3 nM protein
14 nM protein
70 nM protein
350 nM protein
4.0
0.0
−60
−40
−20
0
20
40
60
Position relative to center (bp)

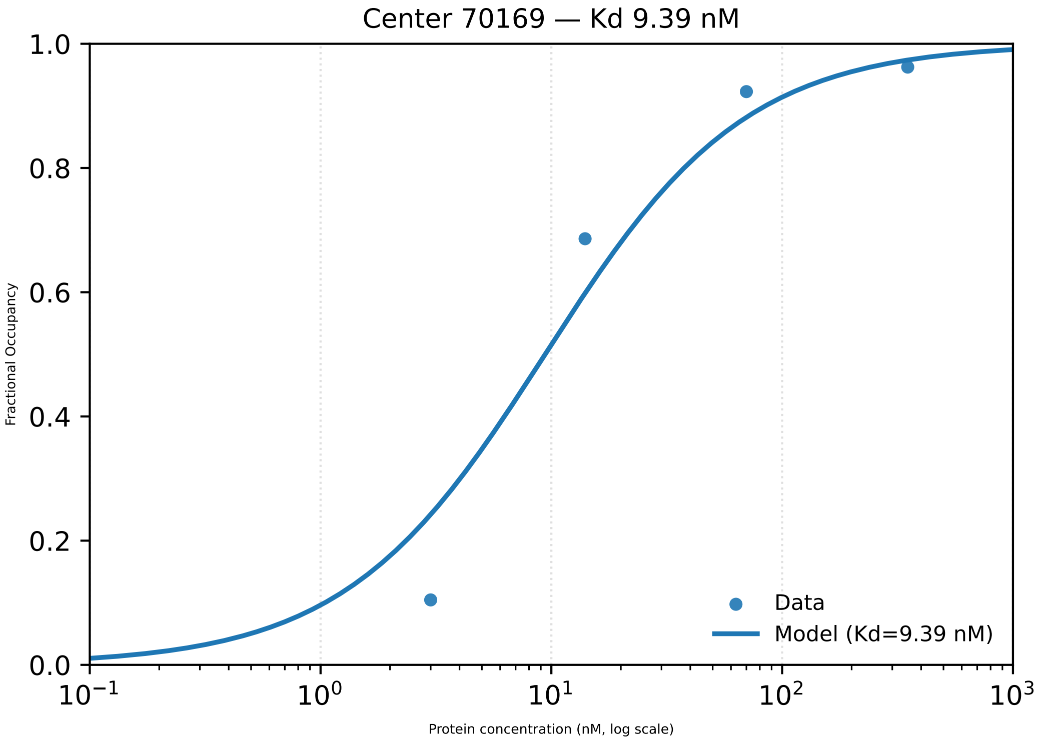

Center 70169 — Kd 9.39 nM
Fractional Occupancy
1.0
0.8
0.6
0.4
0.2
0.0
$10^{-1}$
$10^{0}$
$10^{1}$
$10^{2}$
$10^{3}$
Protein concentration (nM, log scale)
Data
Model (Kd=9.39 nM)

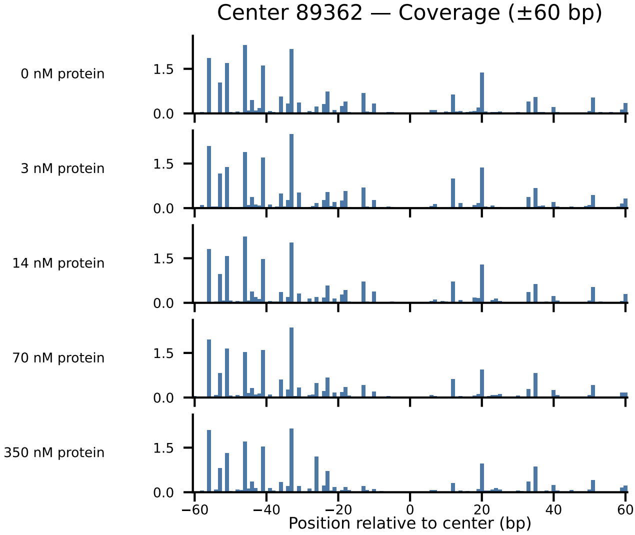
Center 89362 — Coverage (±60 bp)
0 nM protein
3 nM protein
14 nM protein
70 nM protein
350 nM protein
1.5
0.0
−60
−40
−20
0
20
40
60
Position relative to center (bp)

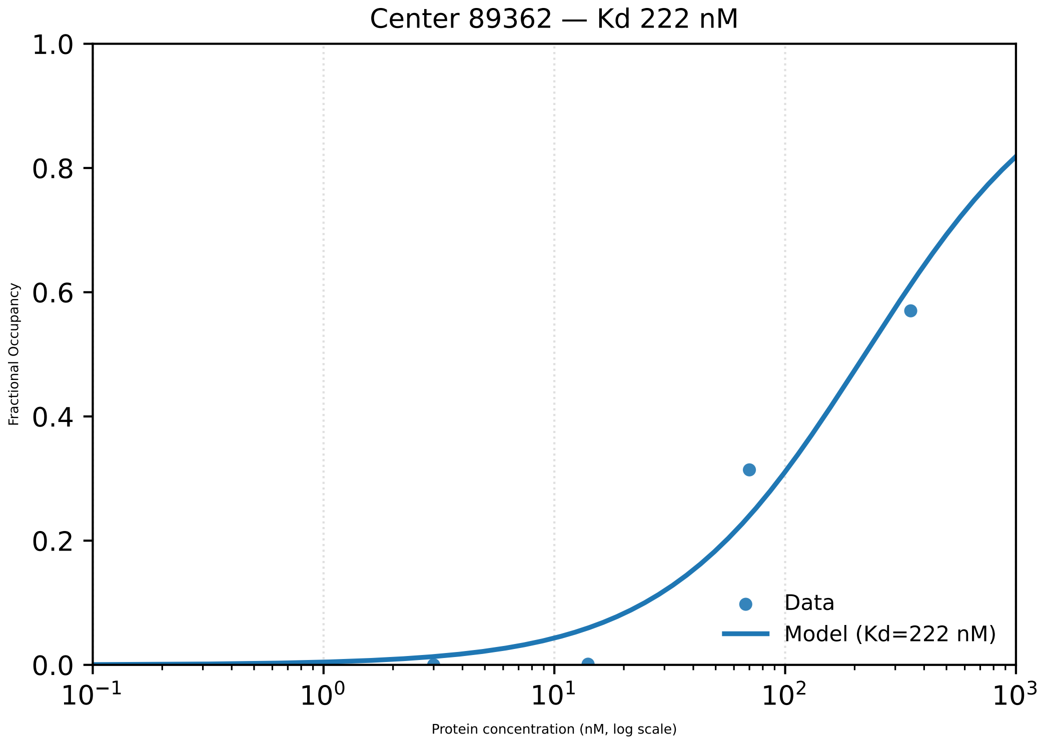

Center 89362 — Kd 222 nM
Fractional Occupancy
Protein concentration (nM, log scale)
1.0
0.8
0.6
0.4
0.2
0.0
$10^{-1}$
$10^{0}$
$10^{1}$
$10^{2}$
$10^{3}$
Data
Model (Kd=222 nM)

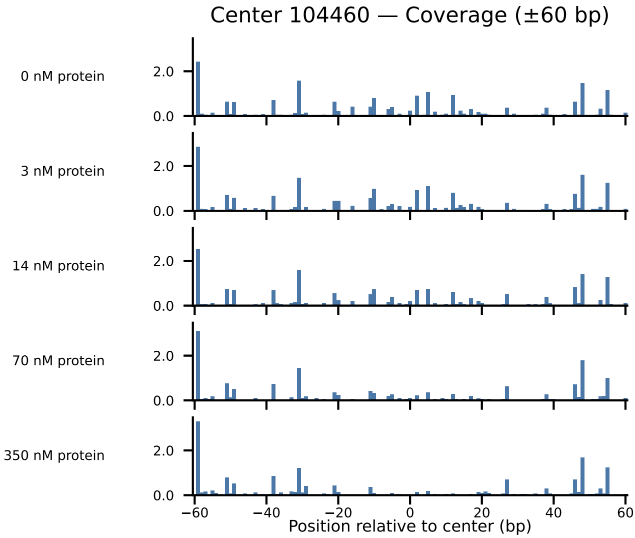

Center 104460 — Coverage (±60 bp)
0 nM protein
3 nM protein
14 nM protein
70 nM protein
350 nM protein
2.0
0.0
−60
−40
−20
0
20
40
60
Position relative to center (bp)

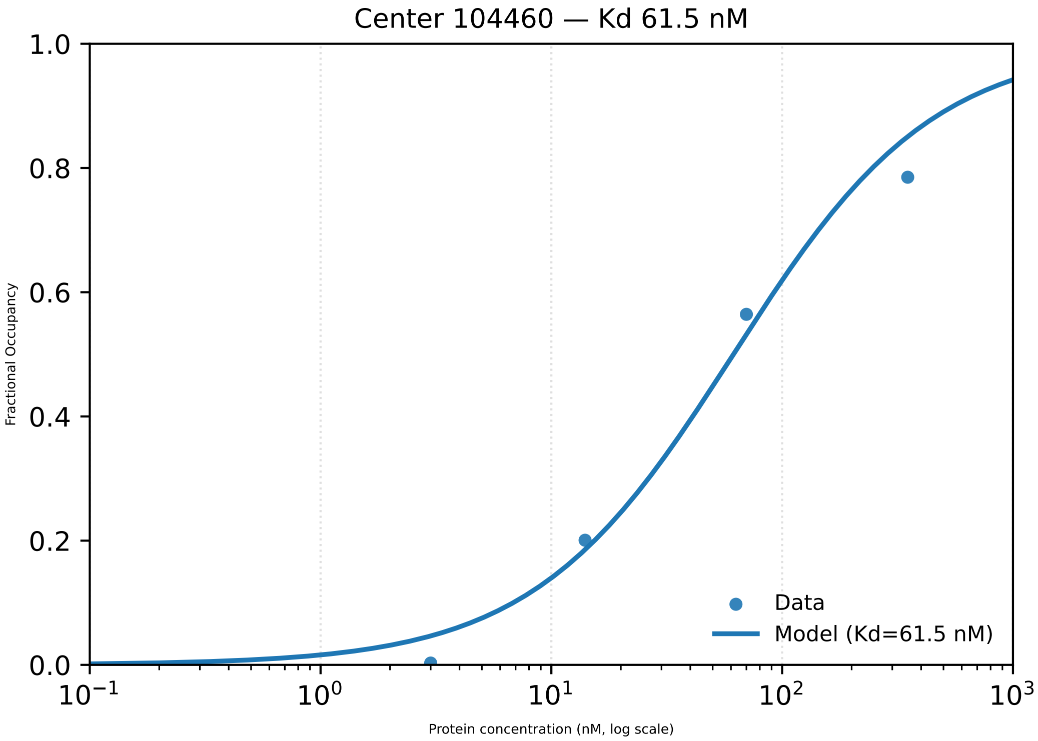

Center 104460 — Kd 61.5 nM
Fractional Occupancy
Protein concentration (nM, log scale)
1.0
0.8
0.6
0.4
0.2
0.0
$10^{-1}$
$10^{0}$
$10^{1}$
$10^{2}$
$10^{3}$
Data
Model (Kd=61.5 nM)

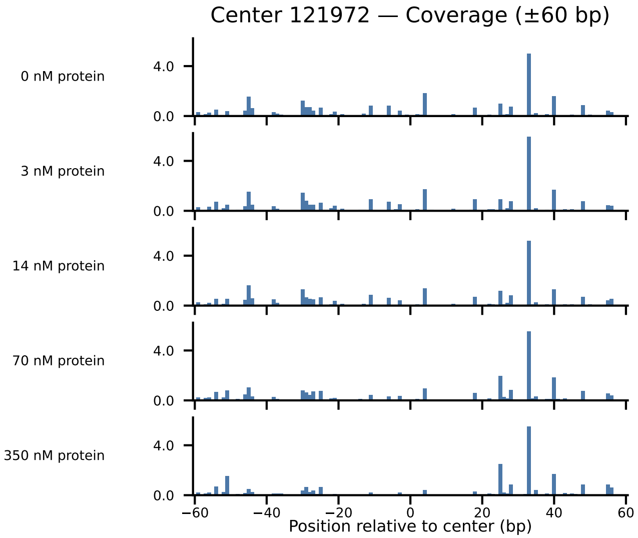

Center 121972 — Coverage (±60 bp)
0 nM protein
3 nM protein
14 nM protein
70 nM protein
350 nM protein
4.0
0.0
−60
−40
−20
0
20
40
60
Position relative to center (bp)

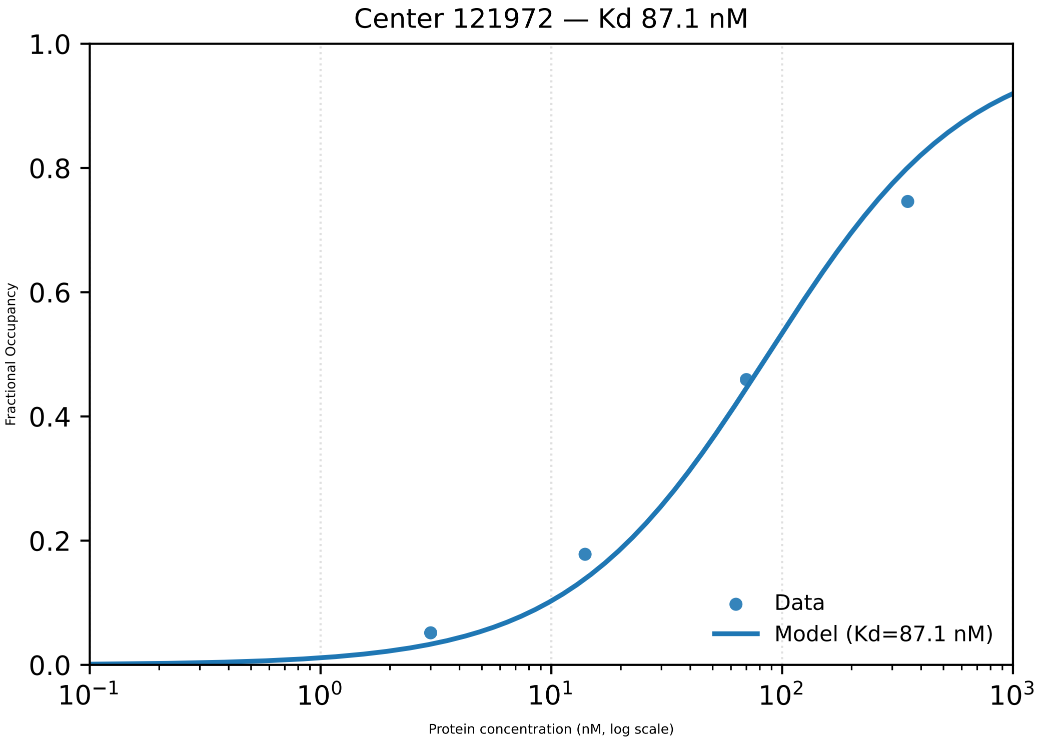

Center 121972 — Kd 87.1 nM
1.0
0.8
0.6
0.4
0.2
0.0
Fractional Occupancy
$10^{-1}$
$10^{0}$
$10^{1}$
$10^{2}$
$10^{3}$
Protein concentration (nM, log scale)
Data
Model (Kd=87.1 nM)

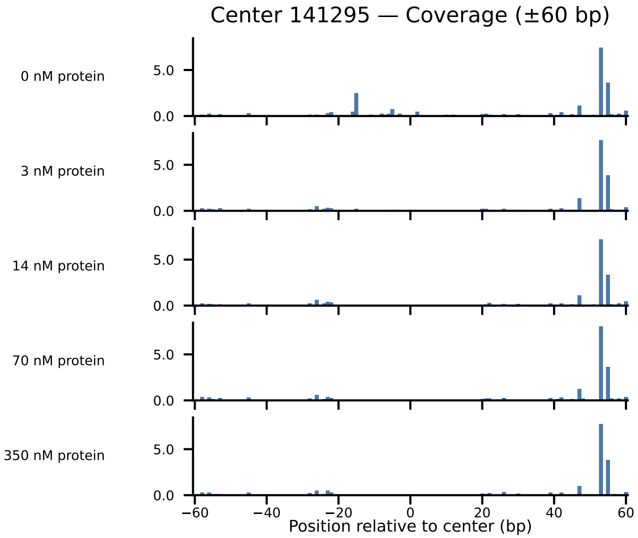
Center 141295 — Coverage (±60 bp)
0 nM protein
3 nM protein
14 nM protein
70 nM protein
350 nM protein
5.0
0.0
−60
−40
−20
0
20
40
60
Position relative to center (bp)

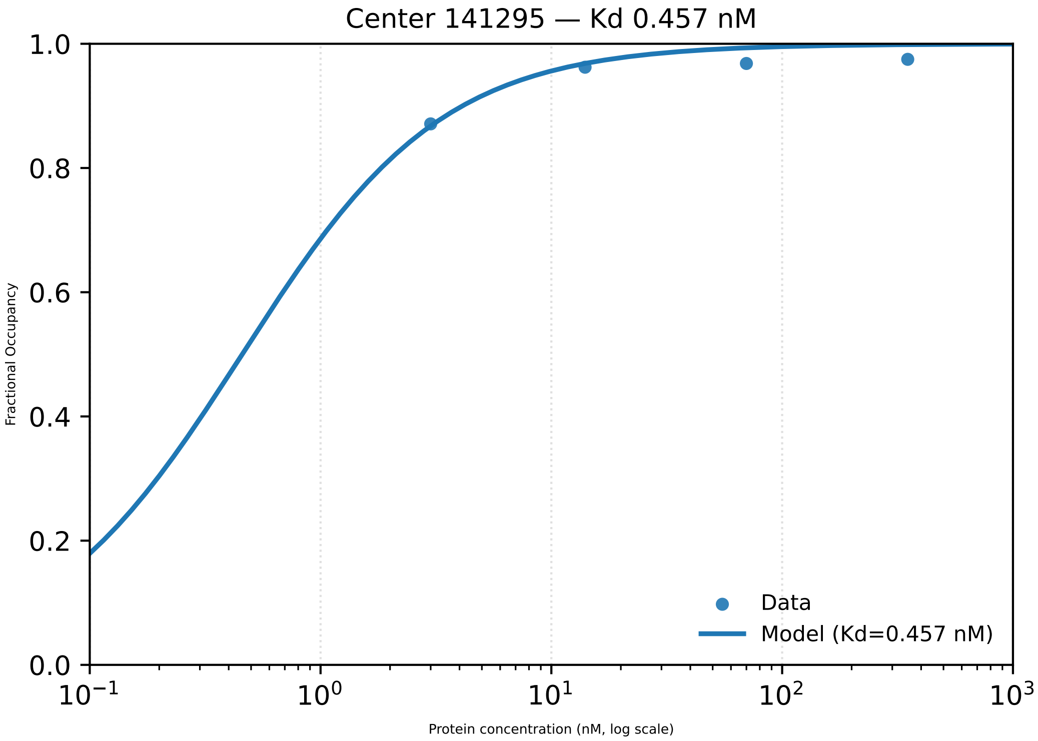

Center 141295 — Kd 0.457 nM
Fractional Occupancy
0.0
0.2
0.4
0.6
0.8
1.0
$10^{-1}$
$10^{0}$
$10^{1}$
$10^{2}$
$10^{3}$
Protein concentration (nM, log scale)
Data
Model (Kd=0.457 nM)

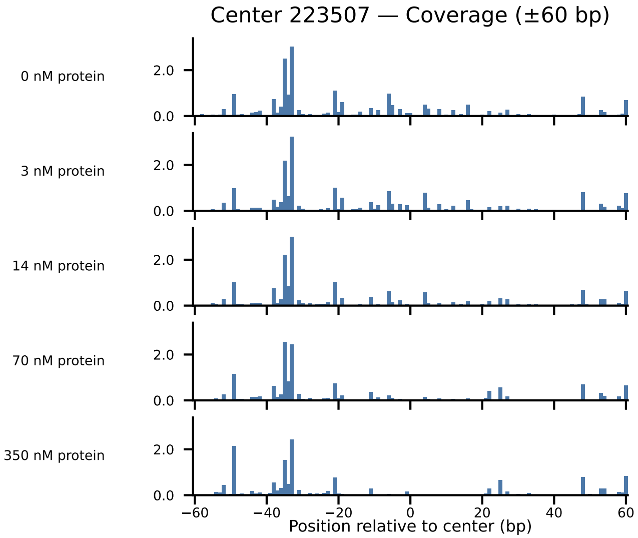
Center 223507 — Coverage (±60 bp)
0 nM protein
3 nM protein
14 nM protein
70 nM protein
350 nM protein
2.0
0.0
−60
−40
−20
0
20
40
60
Position relative to center (bp)

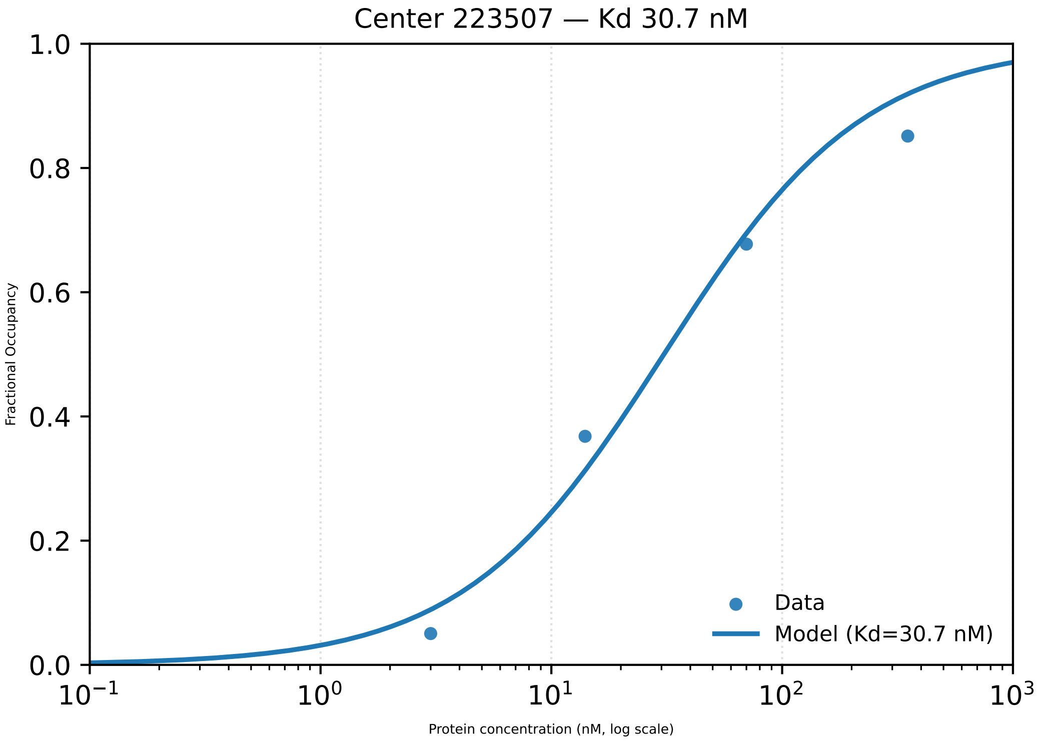

Center 223507 — Kd 30.7 nM
1.0
0.8
0.6
0.4
0.2
0.0
Fractional Occupancy
$10^{-1}$
$10^{0}$
$10^{1}$
$10^{2}$
$10^{3}$
Protein concentration (nM, log scale)
Data
Model (Kd=30.7 nM)

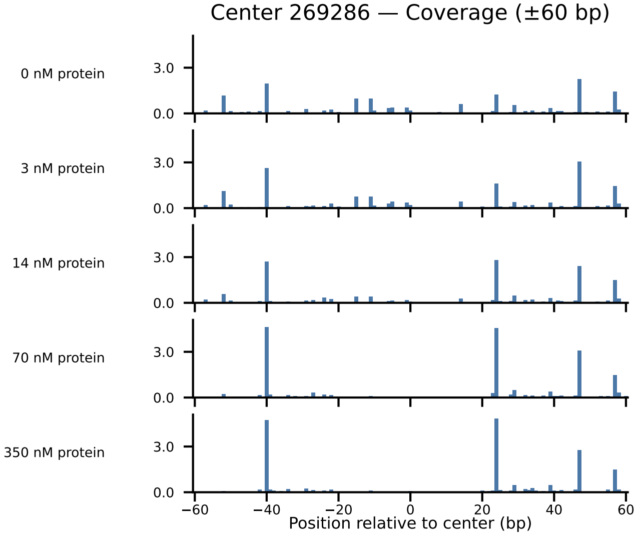

Center 269286 — Coverage (±60 bp)
0 nM protein
3 nM protein
14 nM protein
70 nM protein
350 nM protein
3.0
0.0
−60
−40
−20
0
20
40
60
Position relative to center (bp)

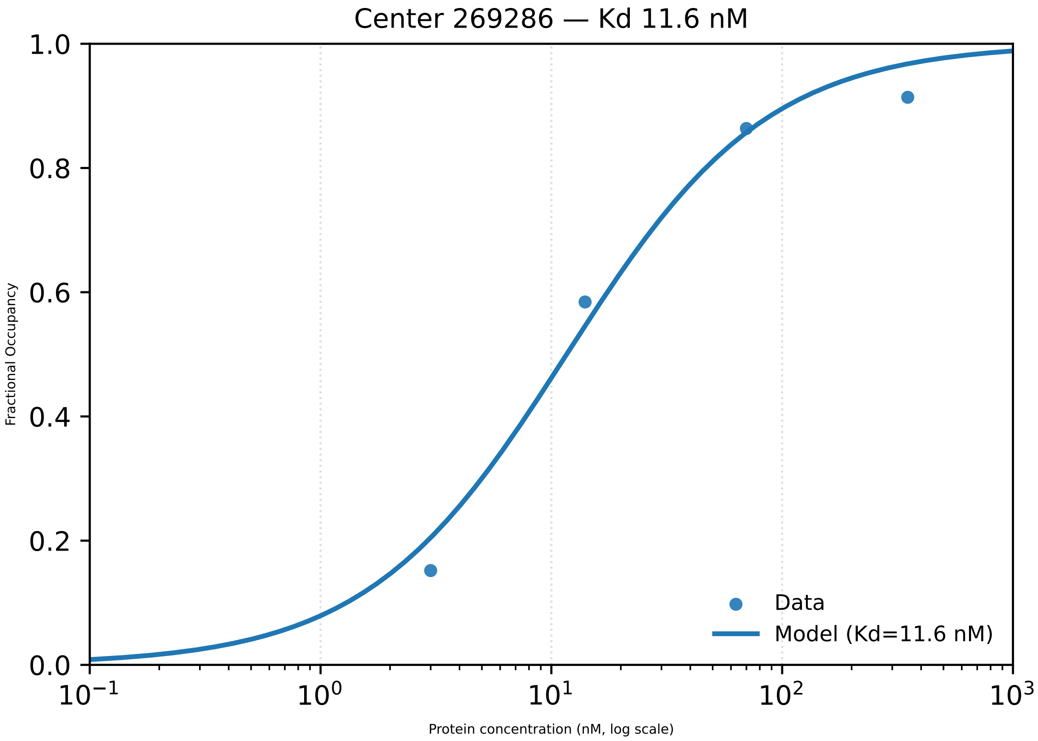

Center 269286 — Kd 11.6 nM
Fractional Occupancy
1.0
0.8
0.6
0.4
0.2
0.0
$10^{-1}$
$10^{0}$
$10^{1}$
$10^{2}$
$10^{3}$
Protein concentration (nM, log scale)
Data
Model (Kd=11.6 nM)

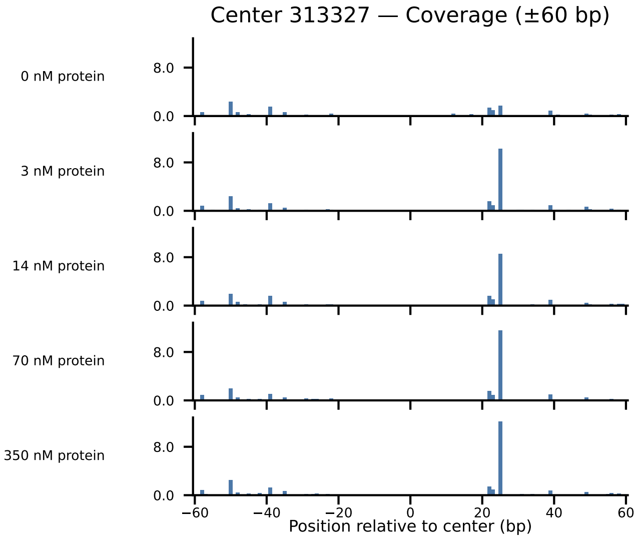

Center 313327 — Coverage (±60 bp)
0 nM protein
3 nM protein
14 nM protein
70 nM protein
350 nM protein
8.0
0.0
−60
−40
−20
0
20
40
60
Position relative to center (bp)

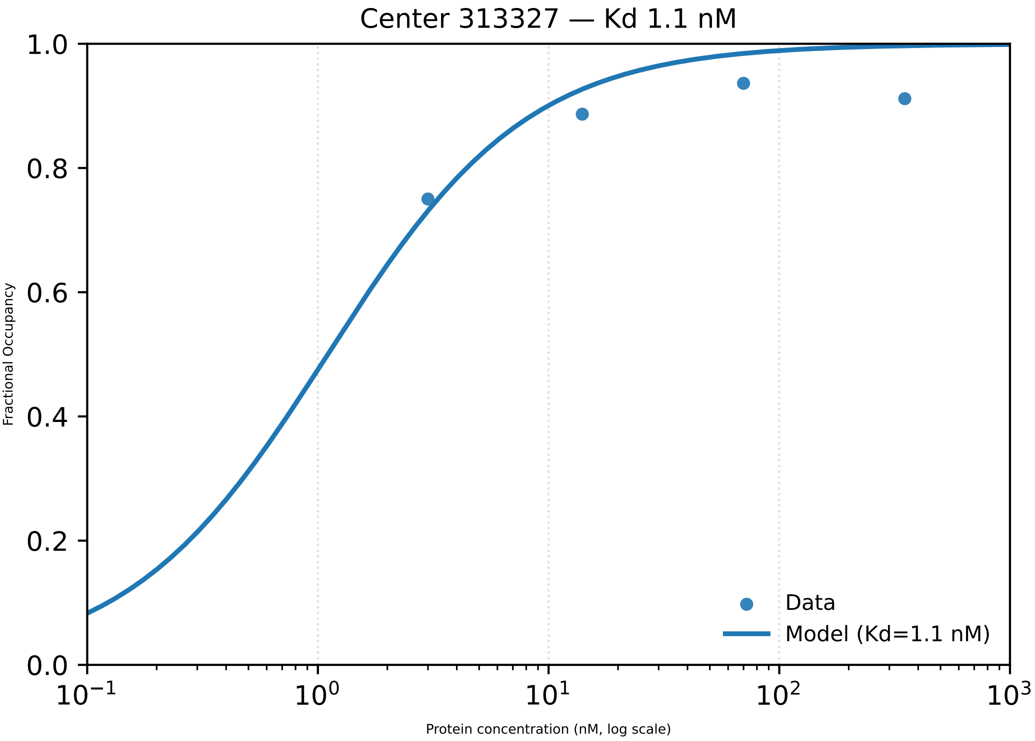

Center 313327 — Kd 1.1 nM
Fractional Occupancy
Protein concentration (nM, log scale)
1.0
0.8
0.6
0.4
0.2
0.0
$10^{-1}$
$10^{0}$
$10^{1}$
$10^{2}$
$10^{3}$
Data
Model (Kd=1.1 nM)

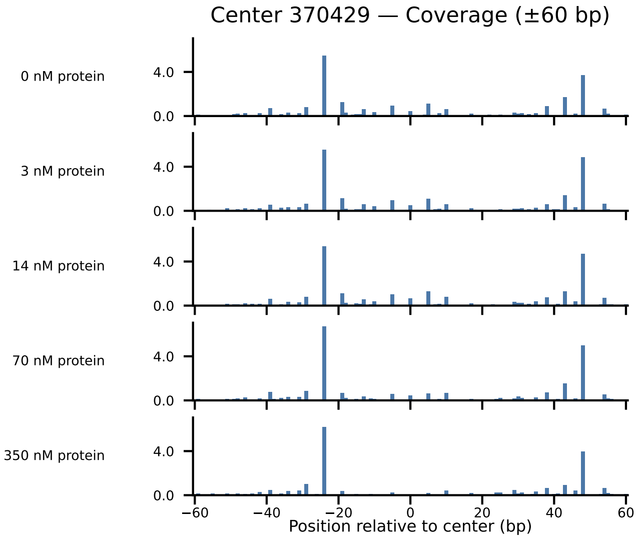

Center 370429 — Coverage (±60 bp)
0 nM protein
3 nM protein
14 nM protein
70 nM protein
350 nM protein
4.0
0.0
−60
−40
−20
0
20
40
60
Position relative to center (bp)

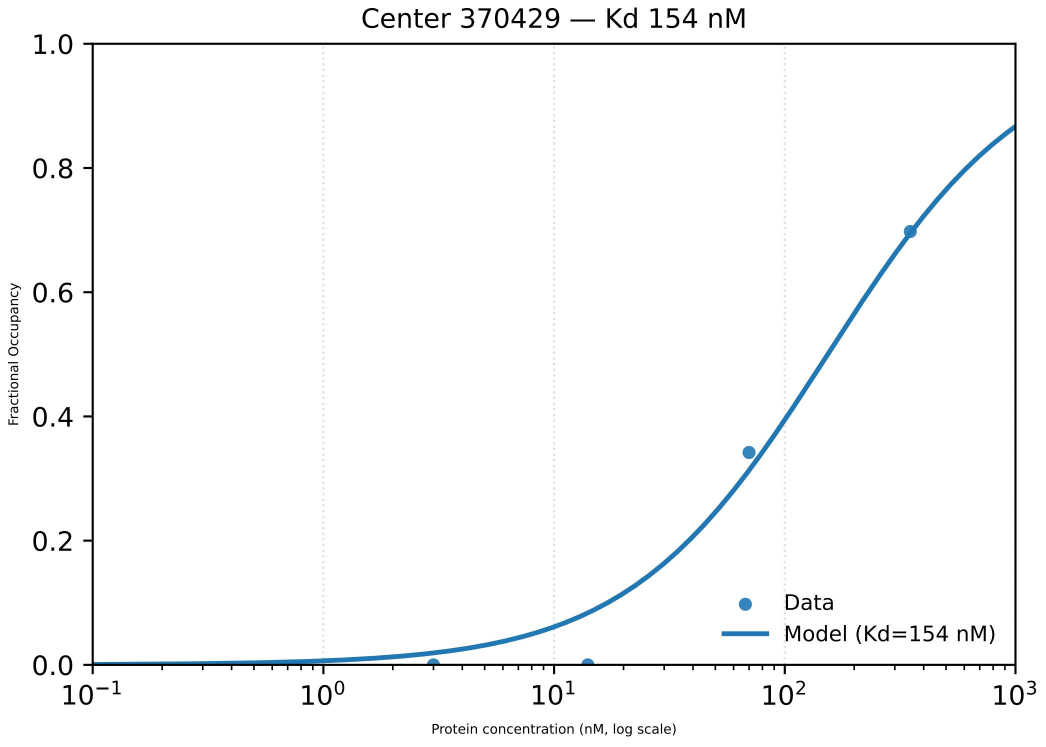

Center 370429 — Kd 154 nM
Fractional Occupancy
1.0
0.8
0.6
0.4
0.2
0.0
$10^{-1}$
$10^{0}$
$10^{1}$
$10^{2}$
$10^{3}$
Protein concentration (nM, log scale)
Data
Model (Kd=154 nM)

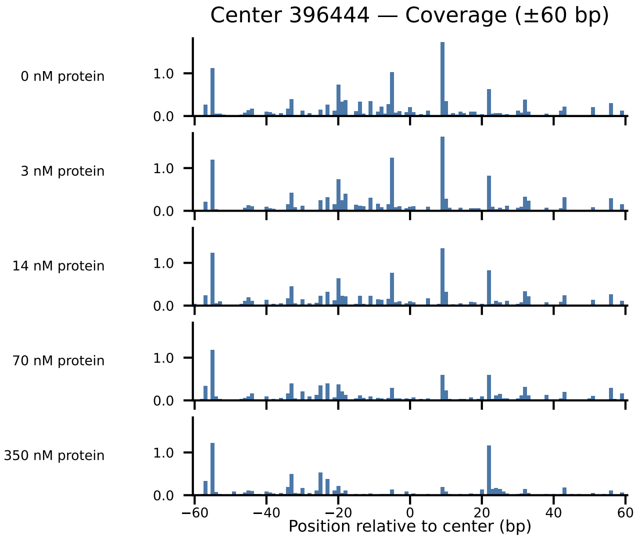

Center 396444 — Coverage (±60 bp)
0 nM protein
3 nM protein
14 nM protein
70 nM protein
350 nM protein
1.0
0.0
−60
−40
−20
0
20
40
60
Position relative to center (bp)

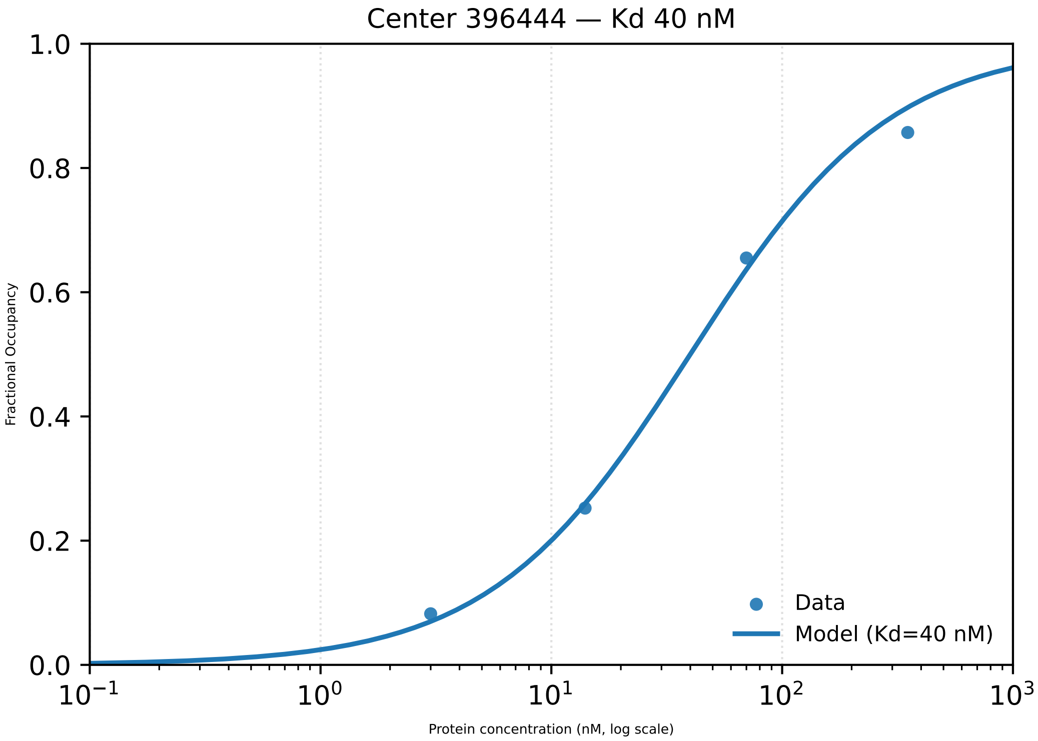

Center 396444 — Kd 40 nM
Fractional Occupancy
Protein concentration (nM, log scale)
0.0
0.2
0.4
0.6
0.8
1.0
$10^{-1}$
$10^{0}$
$10^{1}$
$10^{2}$
$10^{3}$
Data
Model (Kd=40 nM)

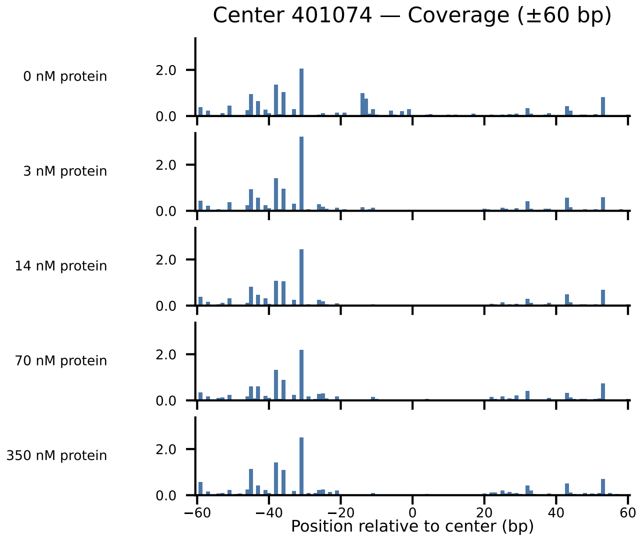

Center 401074 — Coverage (±60 bp)
0 nM protein
3 nM protein
14 nM protein
70 nM protein
350 nM protein
0.0
2.0
−60
−40
−20
0
20
40
60
Position relative to center (bp)

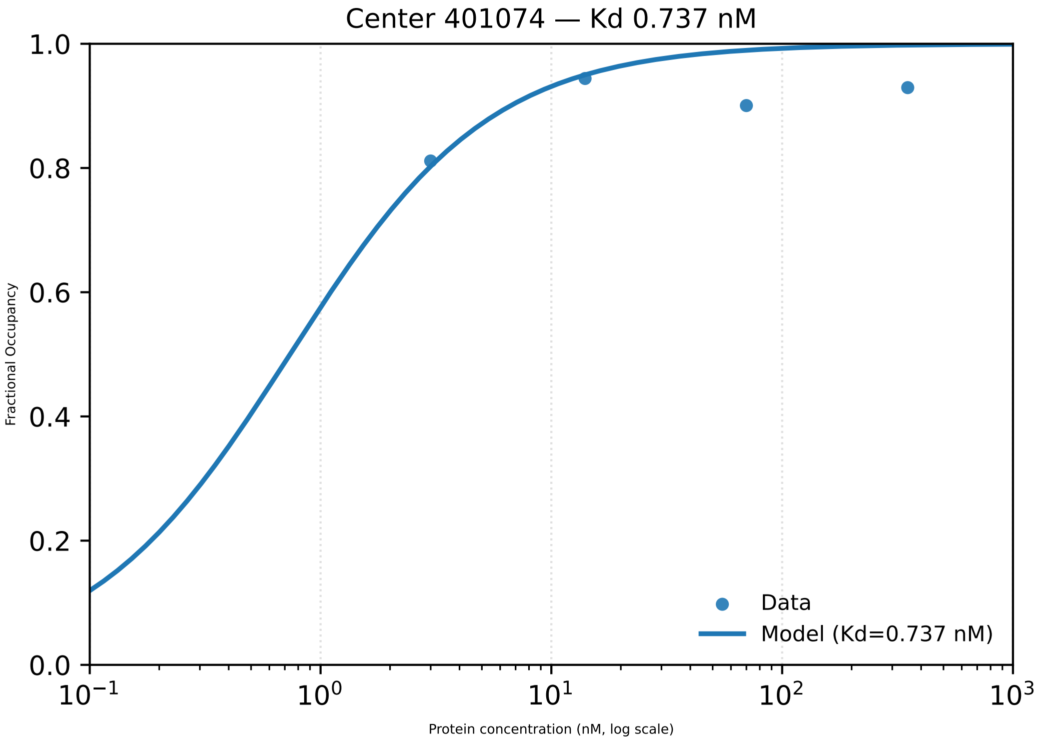
Center 401074 — Kd 0.737 nM
1.0
0.8
0.6
0.4
0.2
0.0
Fractional Occupancy
$10^{-1}$
$10^{0}$
$10^{1}$
$10^{2}$
$10^{3}$
Protein concentration (nM, log scale)
Data
Model (Kd=0.737 nM)

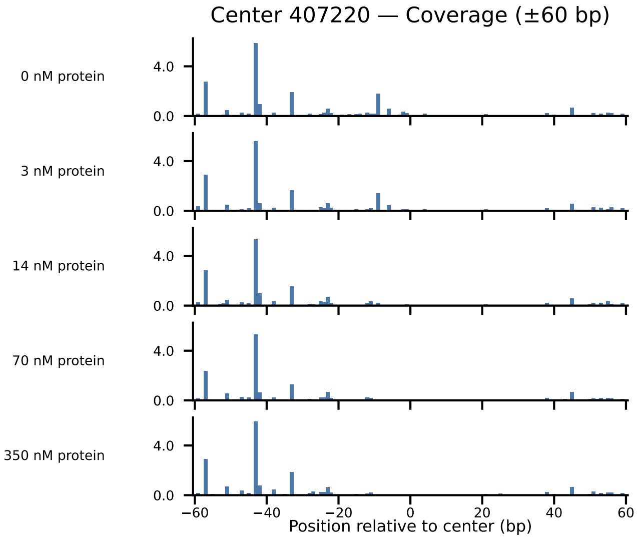

Center 407220 — Coverage (±60 bp)
0 nM protein
3 nM protein
14 nM protein
70 nM protein
350 nM protein
4.0
0.0
−60
−40
−20
0
20
40
60
Position relative to center (bp)

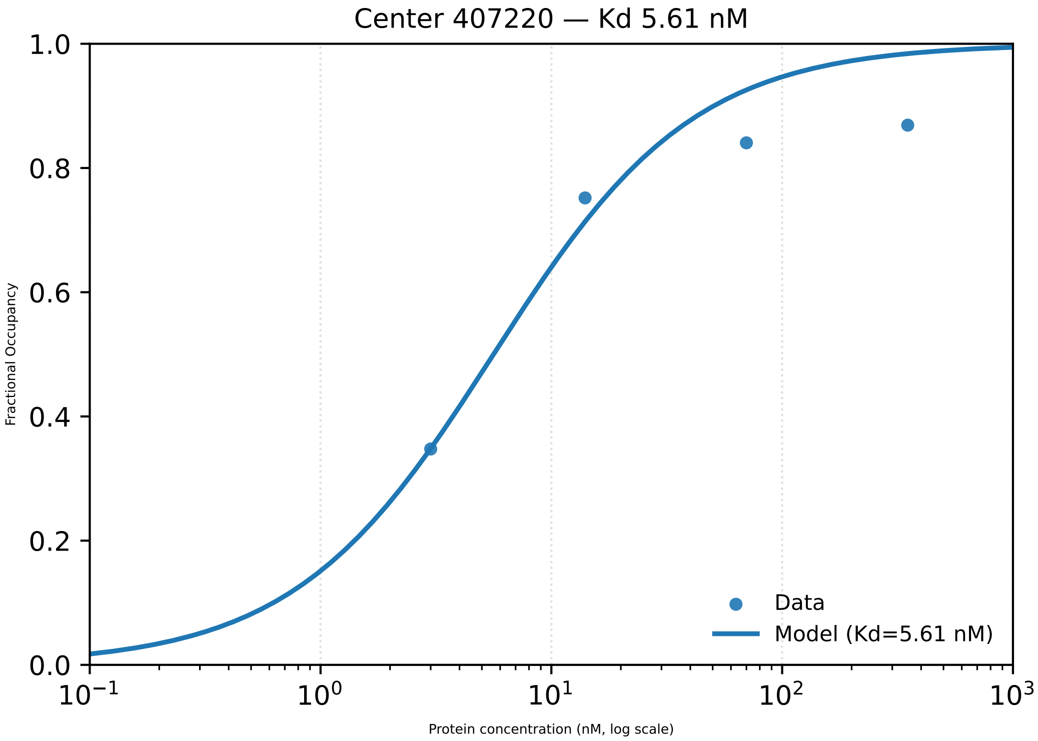

Center 407220 — Kd 5.61 nM
Fractional Occupancy
1.0
0.8
0.6
0.4
0.2
0.0
10−1
100
101
102
103
Protein concentration (nM, log scale)
Data
Model (Kd=5.61 nM)

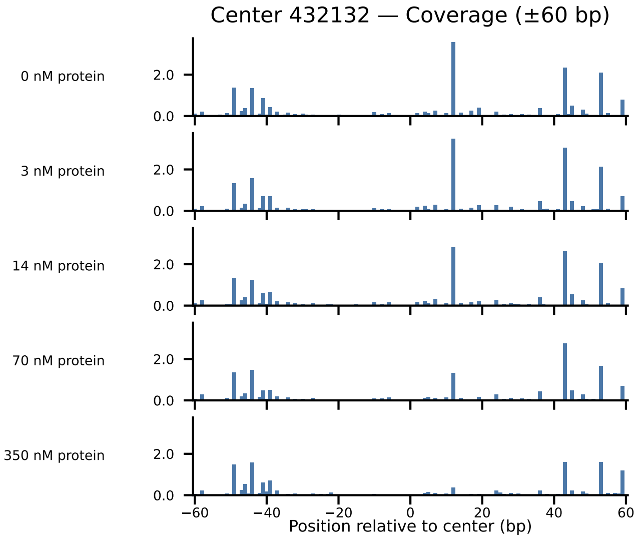

Center 432132 — Coverage (±60 bp)
0 nM protein
3 nM protein
14 nM protein
70 nM protein
350 nM protein
2.0
0.0
−60
−40
−20
0
20
40
60
Position relative to center (bp)

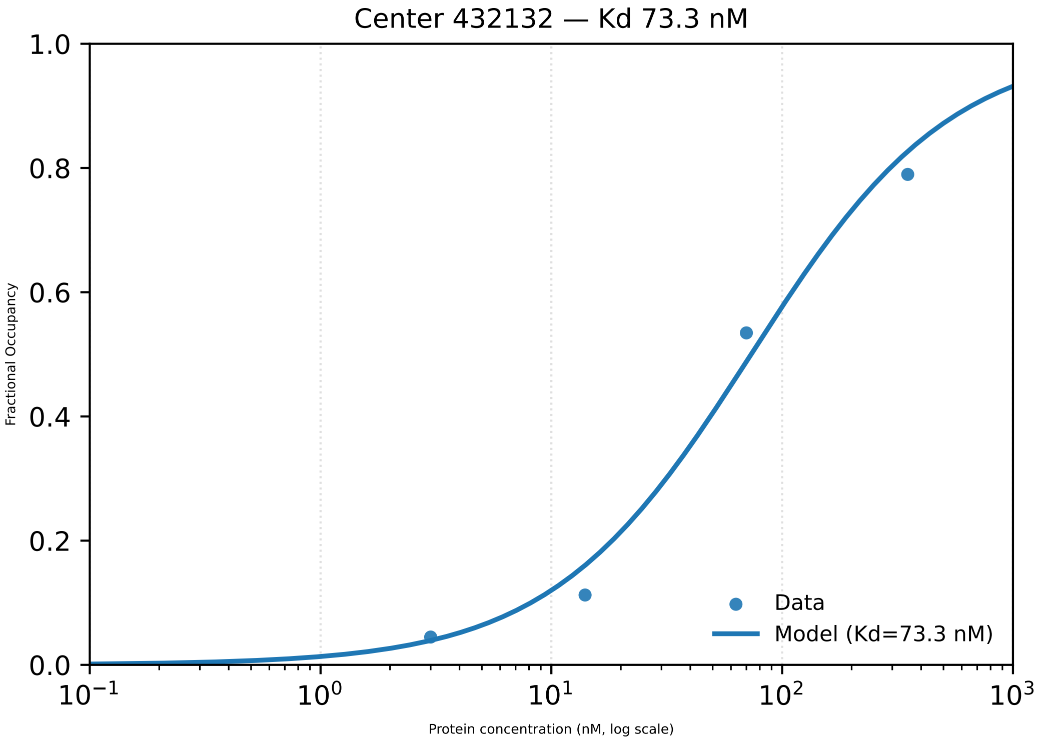

Center 432132 — Kd 73.3 nM
1.0
0.8
0.6
0.4
0.2
0.0
Fractional Occupancy
$10^{-1}$
$10^{0}$
$10^{1}$
$10^{2}$
$10^{3}$
Protein concentration (nM, log scale)
Data
Model (Kd=73.3 nM)

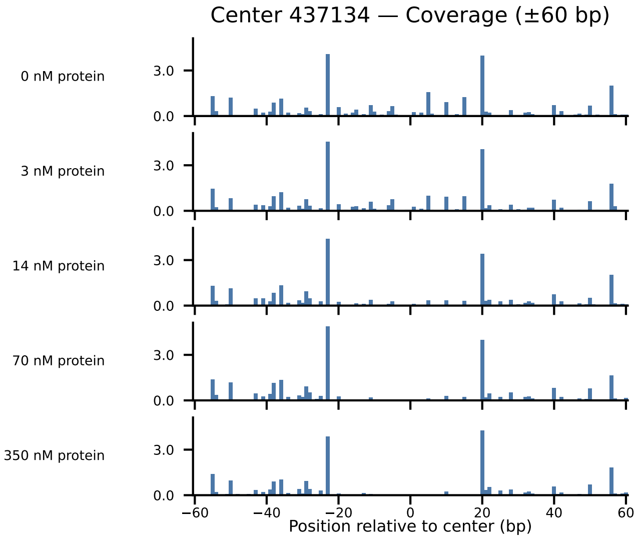

Center 437134 — Coverage (±60 bp)
0 nM protein
3 nM protein
14 nM protein
70 nM protein
350 nM protein
3.0
0.0
−60
−40
−20
0
20
40
60
Position relative to center (bp)

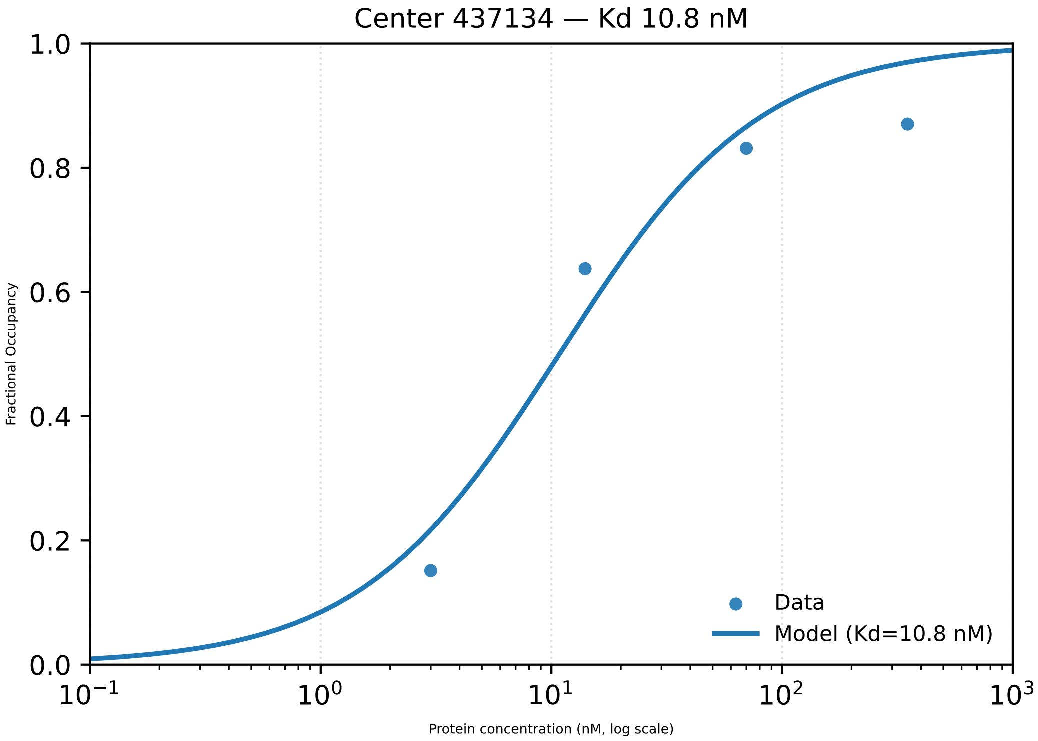

Center 437134 — Kd 10.8 nM
1.0
0.8
0.6
0.4
0.2
0.0
Fractional Occupancy
$10^{-1}$
$10^{0}$
$10^{1}$
$10^{2}$
$10^{3}$
Protein concentration (nM, log scale)
Data
Model (Kd=10.8 nM)

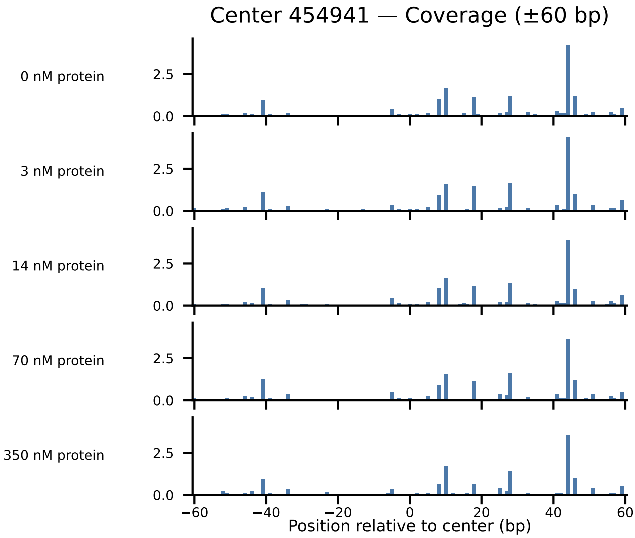
Center 454941 — Coverage (±60 bp)
0 nM protein
3 nM protein
14 nM protein
70 nM protein
350 nM protein
2.5
0.0
−60
−40
−20
0
20
40
60
Position relative to center (bp)

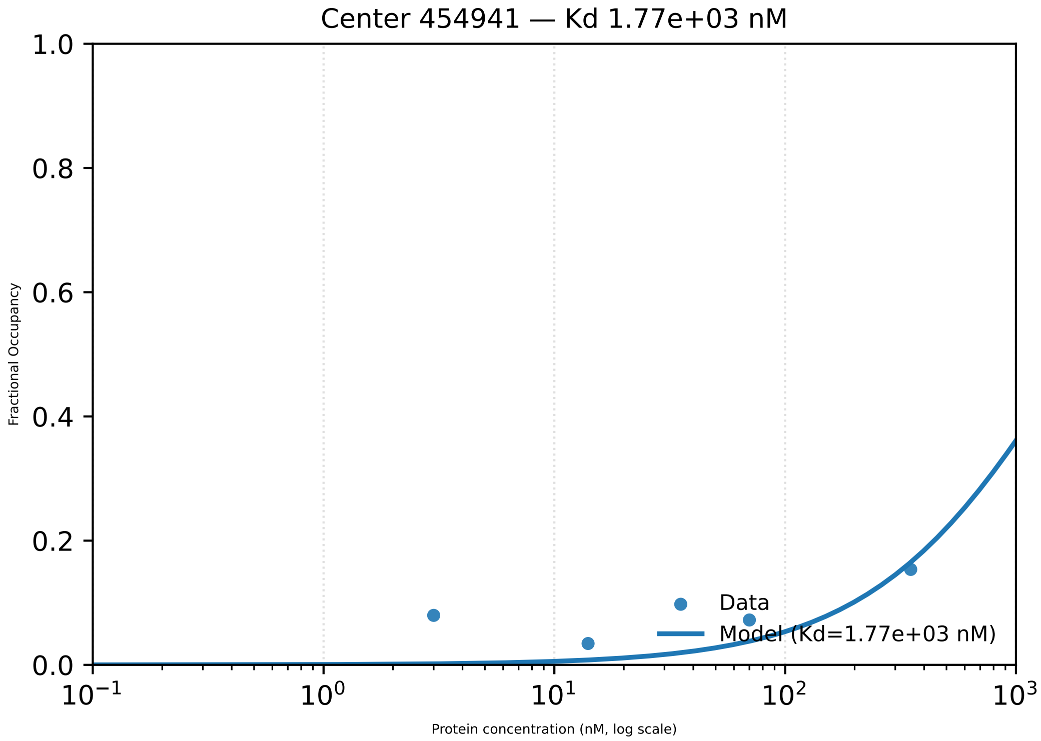

Center 454941 — Kd 1.77e+03 nM
Fractional Occupancy
1.0
0.8
0.6
0.4
0.2
0.0
$10^{-1}$
$10^{0}$
$10^{1}$
$10^{2}$
$10^{3}$
Protein concentration (nM, log scale)
Data
Model (Kd=1.77e+03 nM)

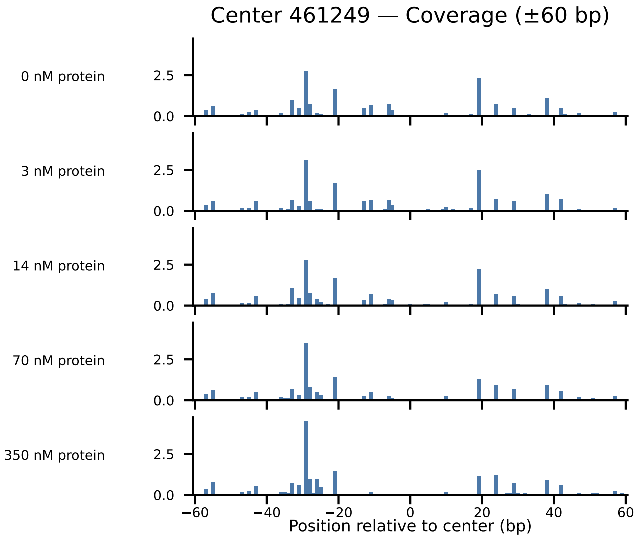

Center 461249 — Coverage (±60 bp)
0 nM protein
3 nM protein
14 nM protein
70 nM protein
350 nM protein
2.5
0.0
−60
−40
−20
0
20
40
60
Position relative to center (bp)

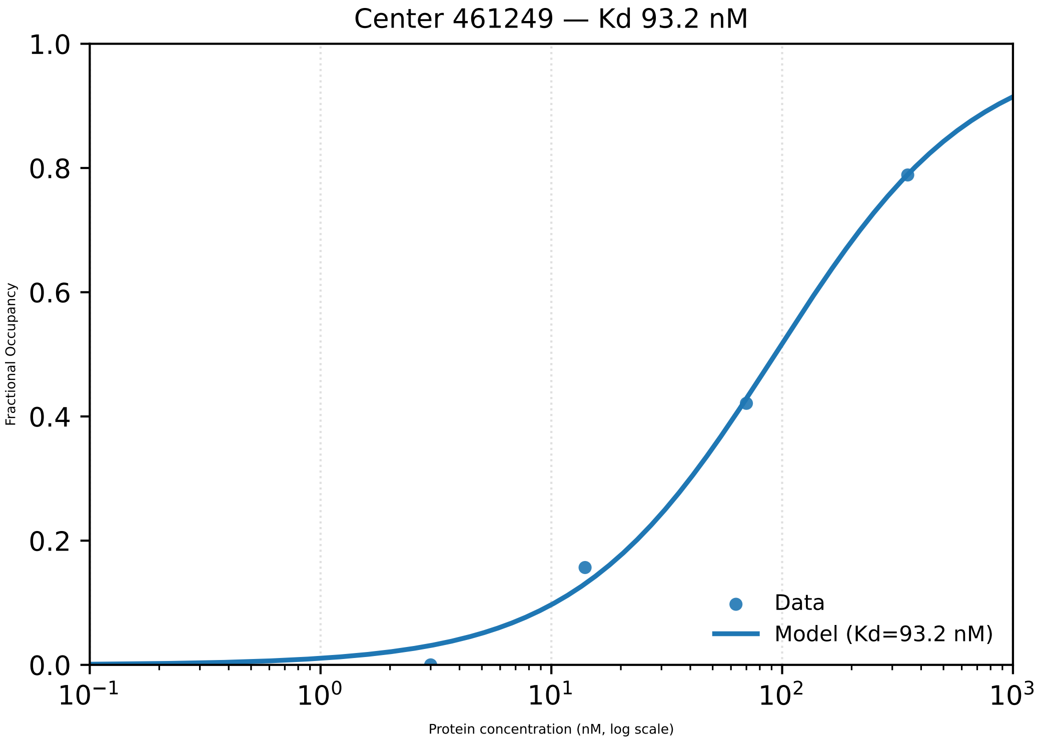

Center 461249 — Kd 93.2 nM
Fractional Occupancy
1.0
0.8
0.6
0.4
0.2
0.0
$10^{-1}$
$10^{0}$
$10^{1}$
$10^{2}$
$10^{3}$
Protein concentration (nM, log scale)
Data
Model (Kd=93.2 nM)

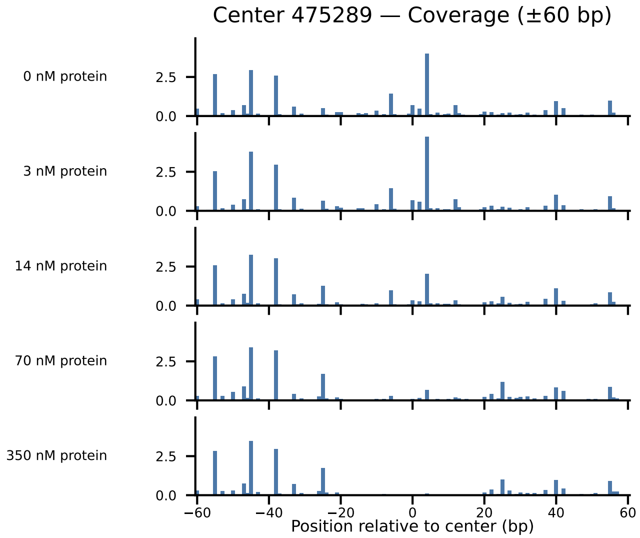

Center 475289 — Coverage (±60 bp)
0 nM protein
3 nM protein
14 nM protein
70 nM protein
350 nM protein
2.5
0.0
−60
−40
−20
0
20
40
60
Position relative to center (bp)

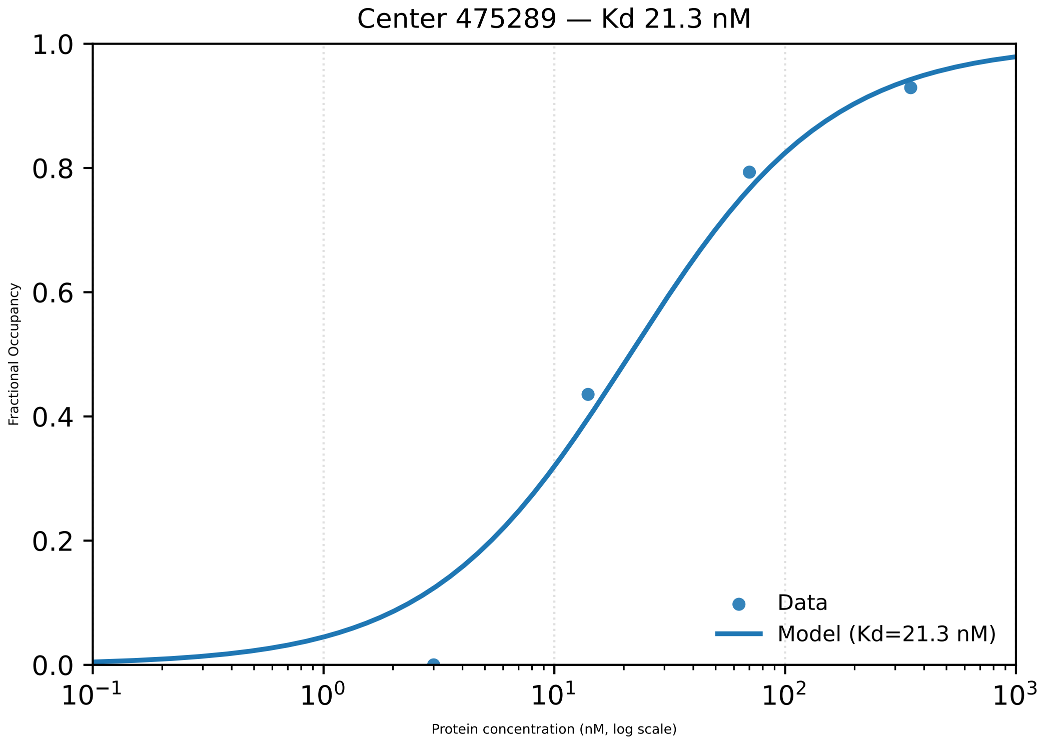

Center 475289 — Kd 21.3 nM
Fractional Occupancy
1.0
0.8
0.6
0.4
0.2
0.0
$10^{-1}$
$10^{0}$
$10^{1}$
$10^{2}$
$10^{3}$
Protein concentration (nM, log scale)
Data
Model (Kd=21.3 nM)

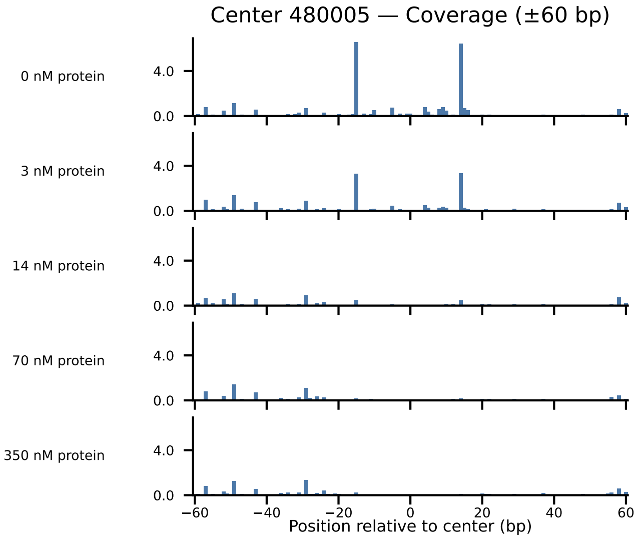

Center 480005 — Coverage (±60 bp)
0 nM protein
3 nM protein
14 nM protein
70 nM protein
350 nM protein
4.0
0.0
−60
−40
−20
0
20
40
60
Position relative to center (bp)

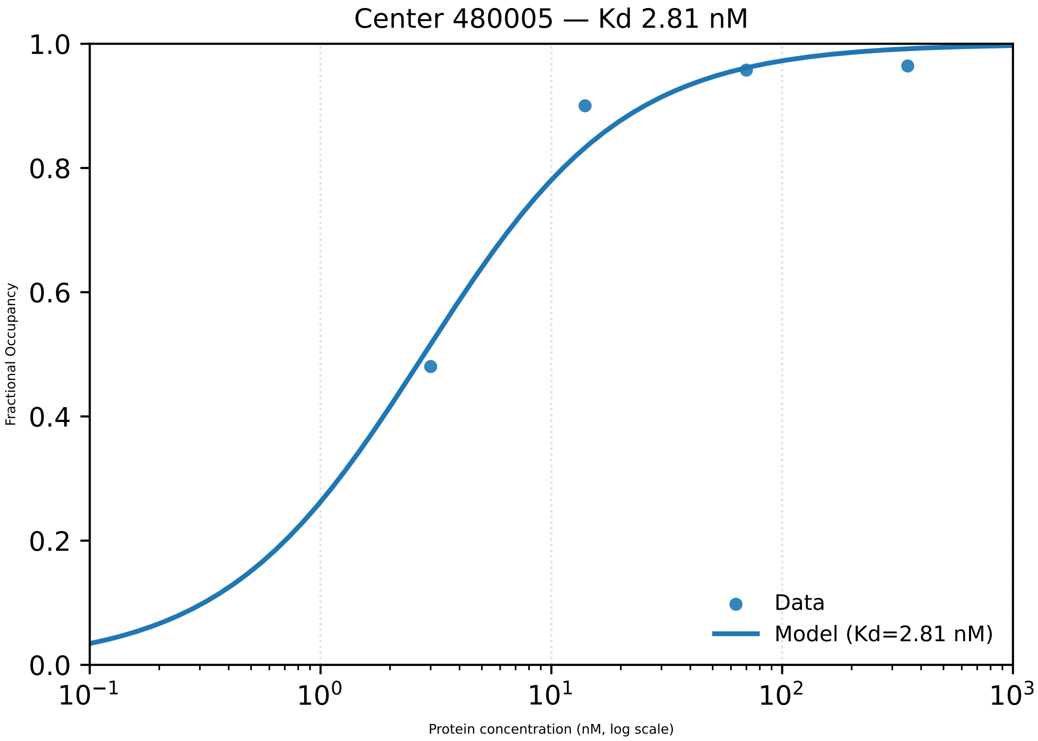

Center 480005 — Kd 2.81 nM
Fractional Occupancy
Protein concentration (nM, log scale)
Data
Model (Kd=2.81 nM)

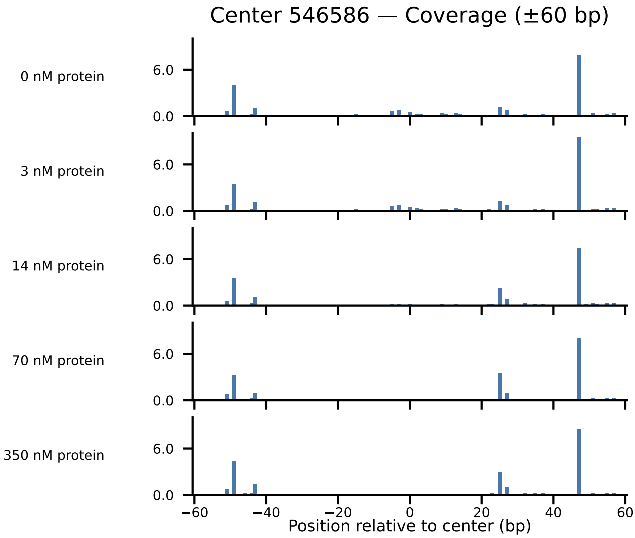

Center 546586 — Coverage (±60 bp)
0 nM protein
3 nM protein
14 nM protein
70 nM protein
350 nM protein
6.0
0.0
−60
−40
−20
0
20
40
60
Position relative to center (bp)

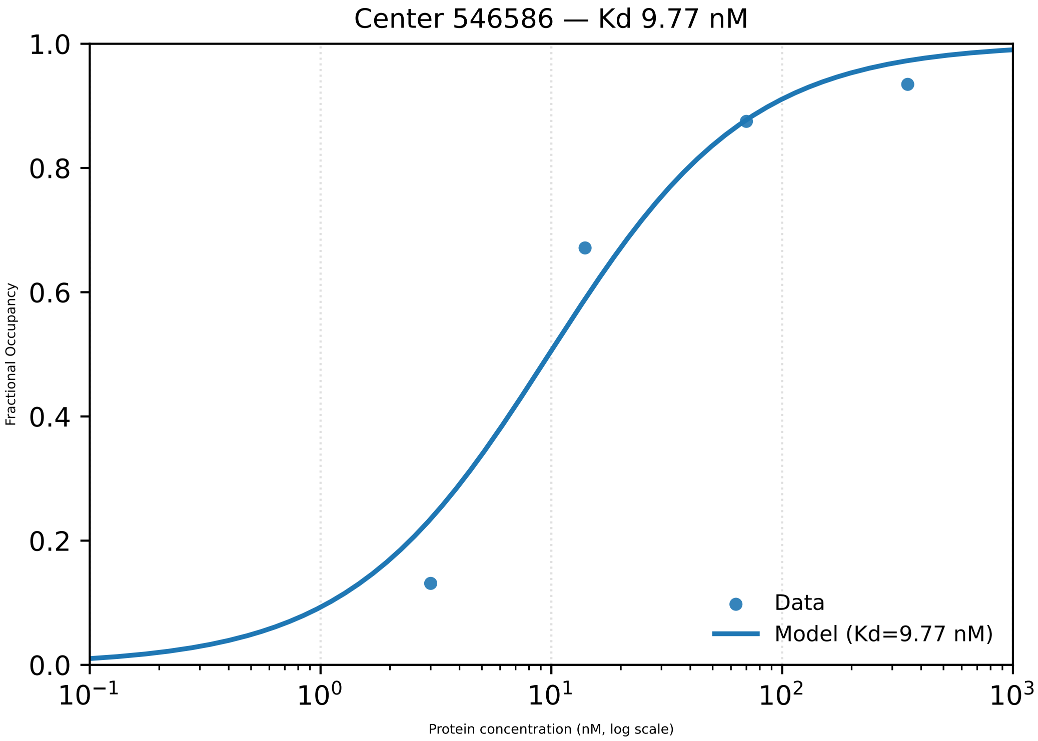

Center 546586 — Kd 9.77 nM
1.0
0.8
0.6
0.4
0.2
0.0
Fractional Occupancy
$10^{-1}$
$10^{0}$
$10^{1}$
$10^{2}$
$10^{3}$
Protein concentration (nM, log scale)
Data
Model (Kd=9.77 nM)

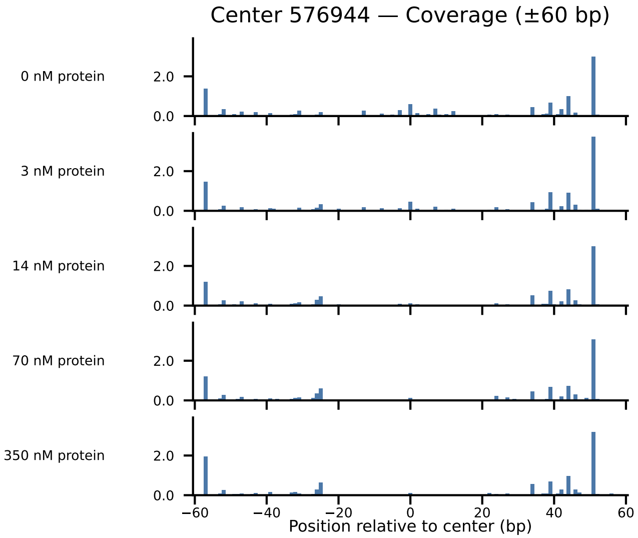
Center 576944 — Coverage (±60 bp)
0 nM protein
3 nM protein
14 nM protein
70 nM protein
350 nM protein
2.0
0.0
−60
−40
−20
0
20
40
60
Position relative to center (bp)

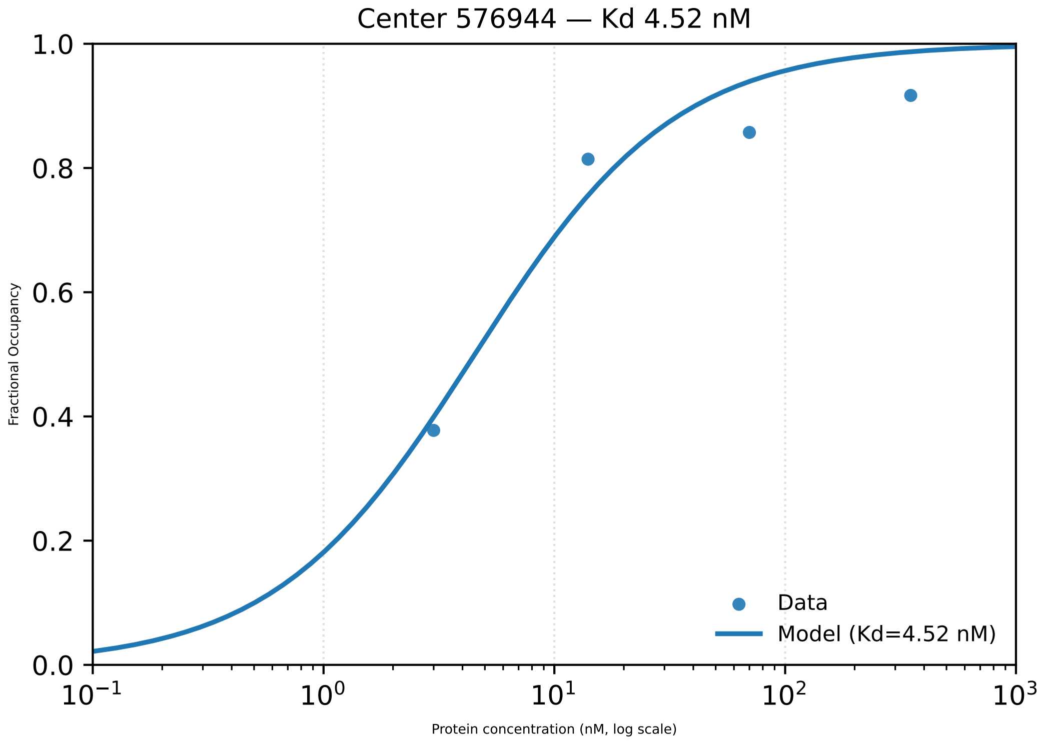

Center 576944 — Kd 4.52 nM
Fractional Occupancy
1.0
0.8
0.6
0.4
0.2
0.0
$10^{-1}$
$10^{0}$
$10^{1}$
$10^{2}$
$10^{3}$
Protein concentration (nM, log scale)
Data
Model (Kd=4.52 nM)

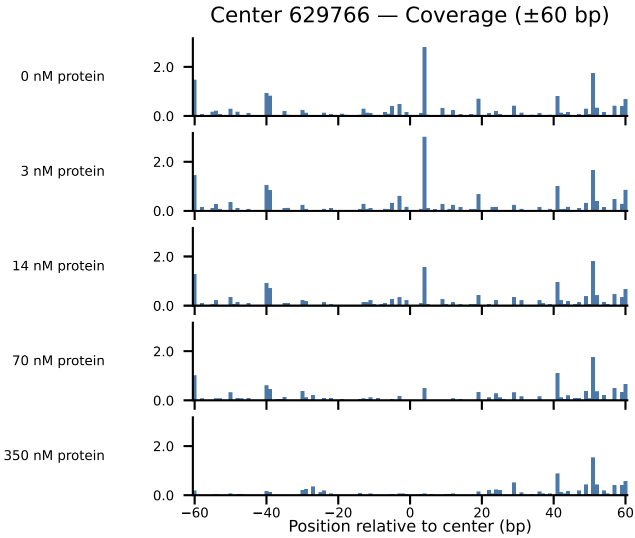

Center 629766 — Coverage (±60 bp)
0 nM protein
3 nM protein
14 nM protein
70 nM protein
350 nM protein
2.0
0.0
−60
−40
−20
0
20
40
60
Position relative to center (bp)

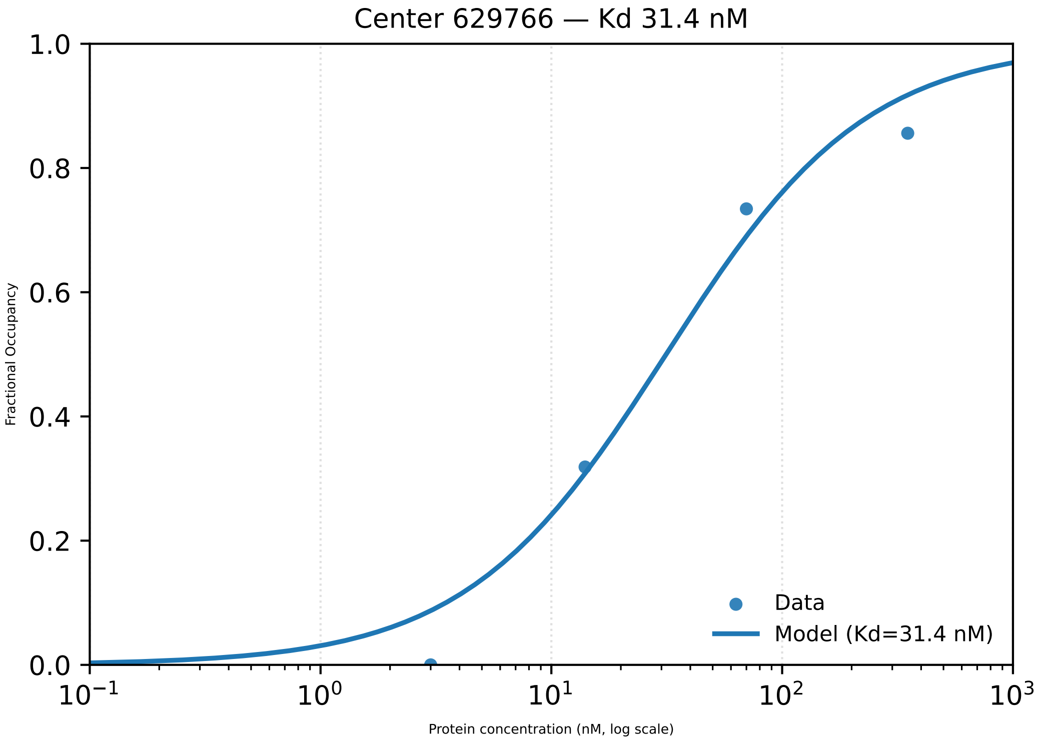
Center 629766 — Kd 31.4 nM
Fractional Occupancy
Protein concentration (nM, log scale)
Data
Model (Kd=31.4 nM)
1.0
0.8
0.6
0.4
0.2
0.0
$10^{-1}$
$10^{0}$
$10^{1}$
$10^{2}$
$10^{3}$

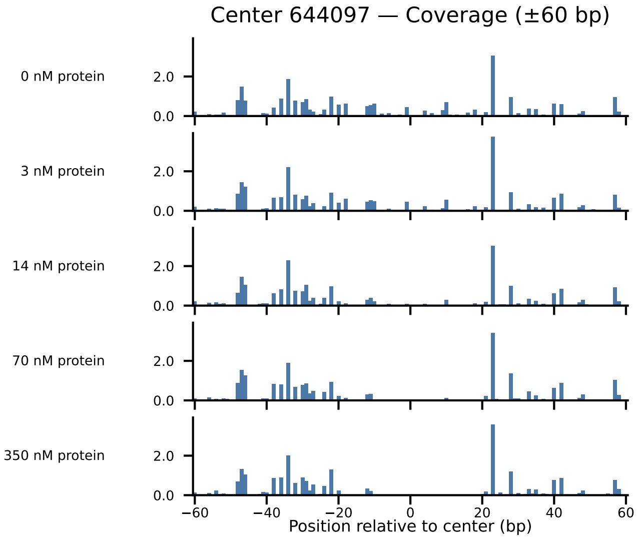
Center 644097 — Coverage (±60 bp)
0 nM protein
3 nM protein
14 nM protein
70 nM protein
350 nM protein
2.0
0.0
−60
−40
−20
0
20
40
60
Position relative to center (bp)

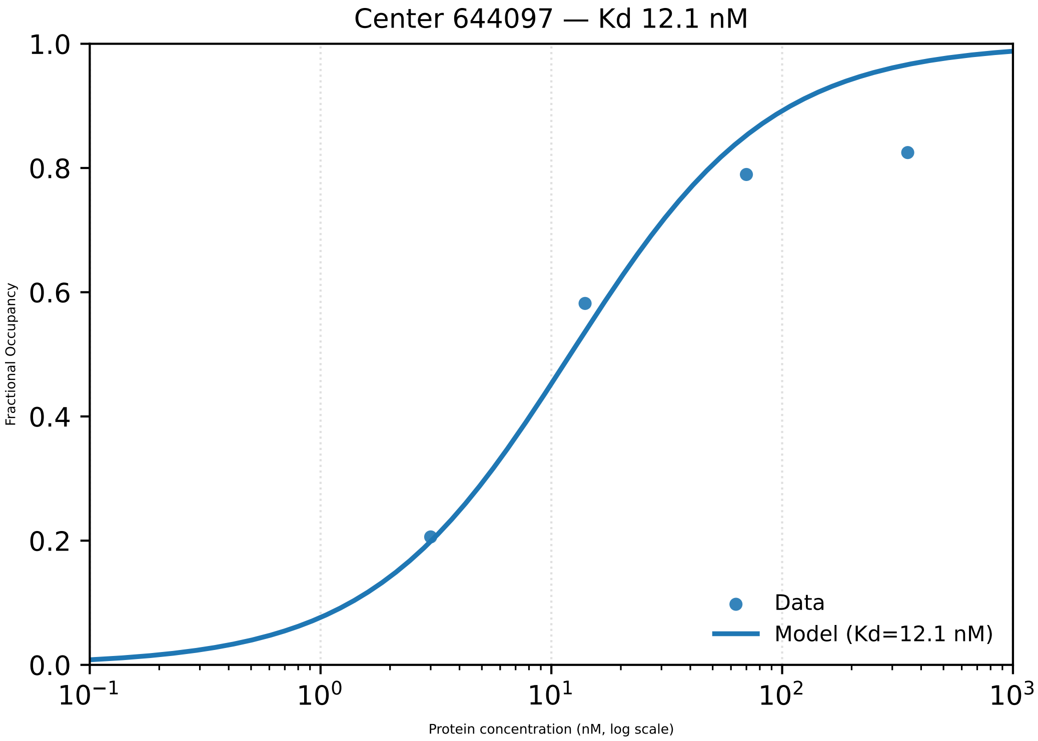

Center 644097 — Kd 12.1 nM
Fractional Occupancy
Protein concentration (nM, log scale)
1.0
0.8
0.6
0.4
0.2
0.0
$10^{-1}$
$10^{0}$
$10^{1}$
$10^{2}$
$10^{3}$
Data
Model (Kd=12.1 nM)

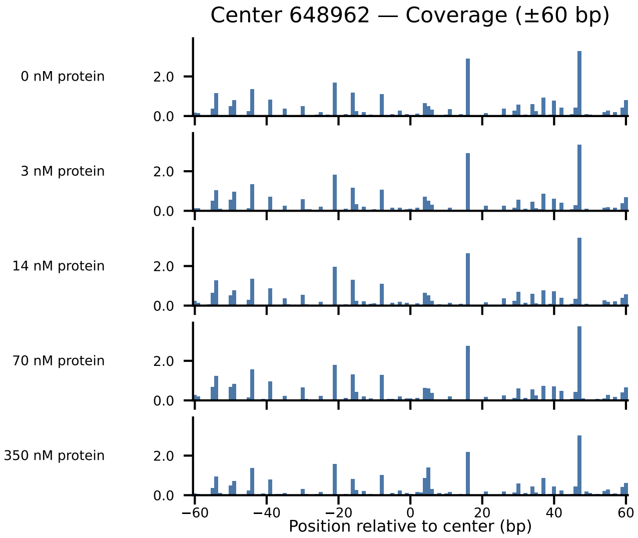

Center 648962 — Coverage (±60 bp)
0 nM protein
3 nM protein
14 nM protein
70 nM protein
350 nM protein
2.0
0.0
−60
−40
−20
0
20
40
60
Position relative to center (bp)

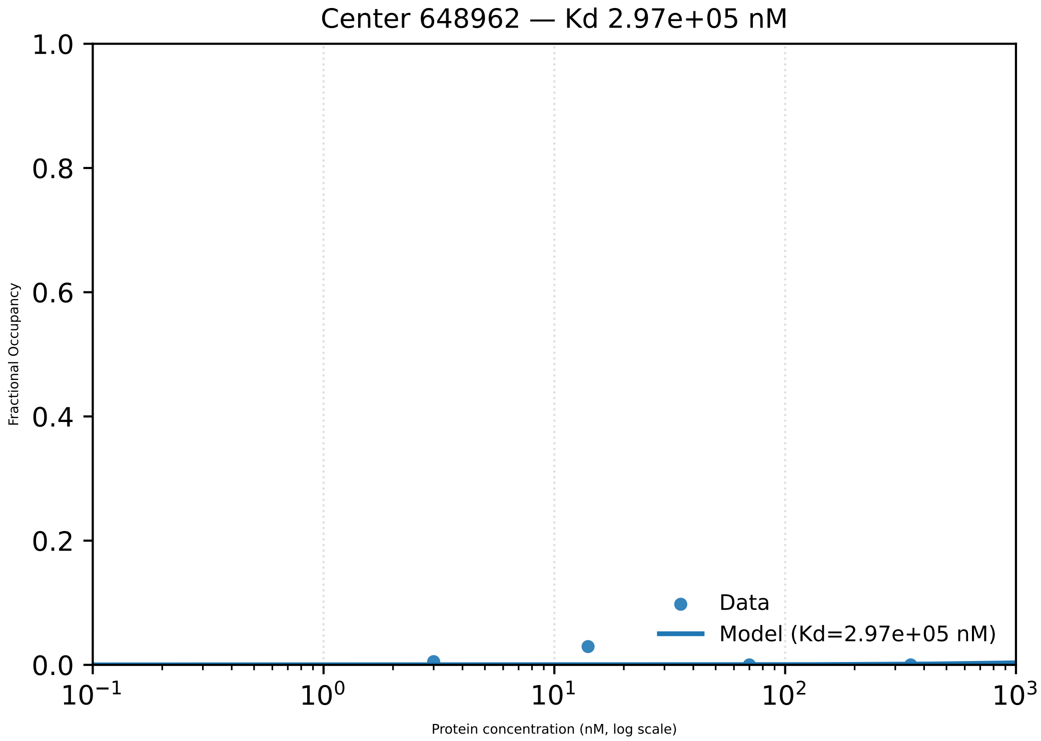
Center 648962 — Kd 2.97e+05 nM
1.0
0.8
0.6
0.4
0.2
0.0
Fractional Occupancy
$10^{-1}$
$10^{0}$
$10^{1}$
$10^{2}$
$10^{3}$
Protein concentration (nM, log scale)
Data
Model (Kd=2.97e+05 nM)

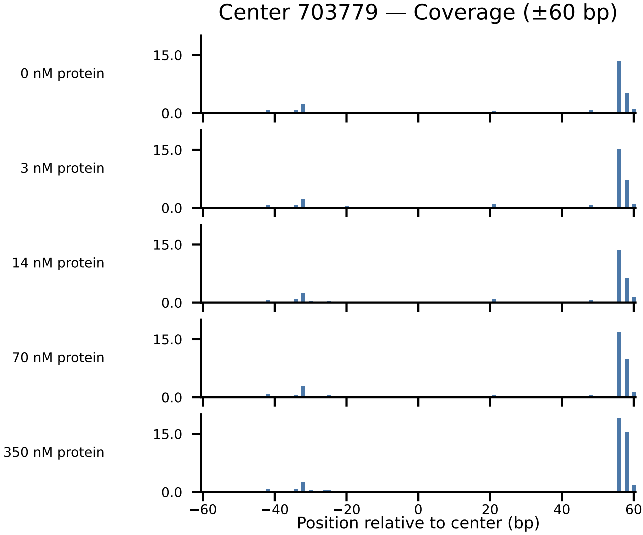
Center 703779 — Coverage (±60 bp)
0 nM protein
3 nM protein
14 nM protein
70 nM protein
350 nM protein
15.0
0.0
−60
−40
−20
0
20
40
60
Position relative to center (bp)

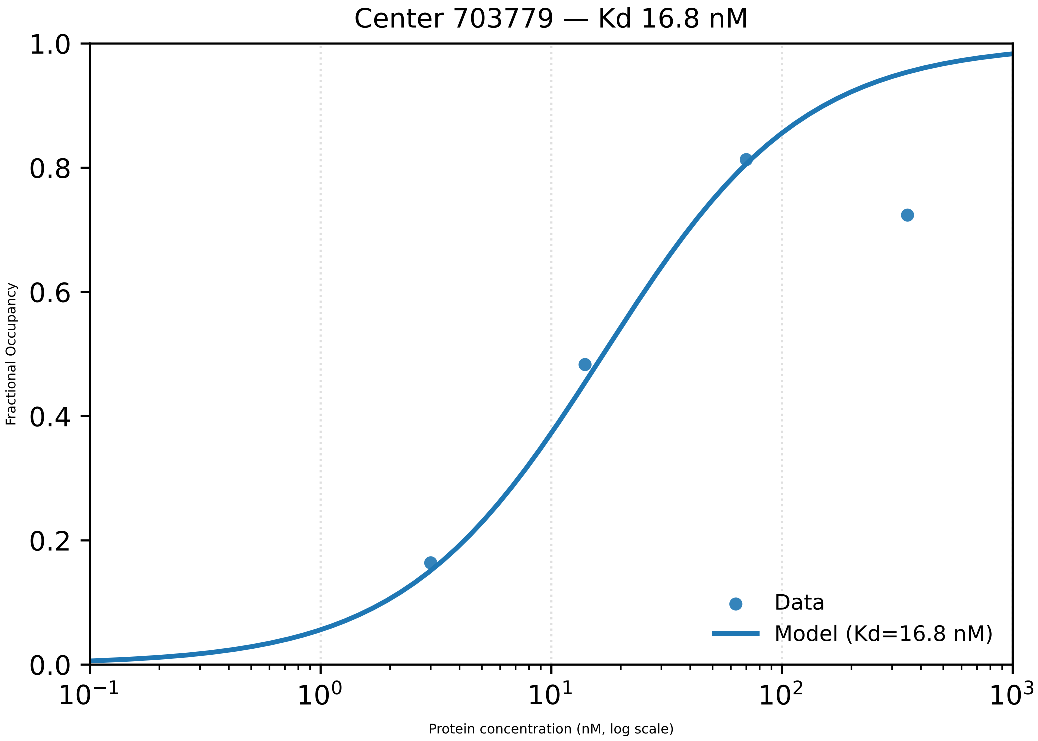

Center 703779 — Kd 16.8 nM
1.0
0.8
0.6
0.4
0.2
0.0
Fractional Occupancy
10−1
100
101
102
103
Protein concentration (nM, log scale)
Data
Model (Kd=16.8 nM)

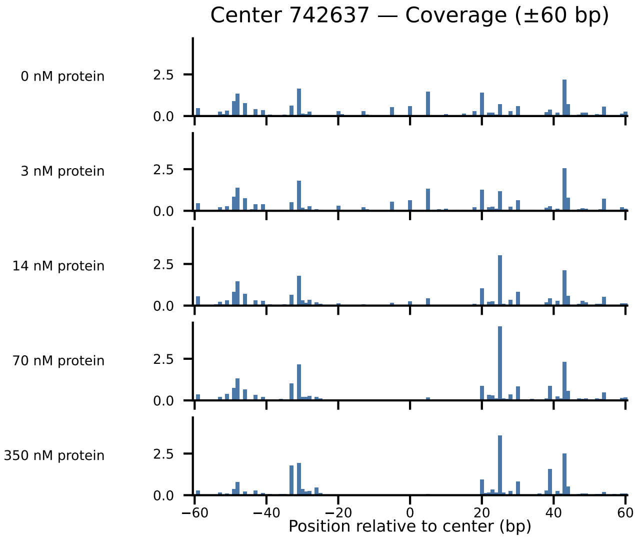

Center 742637 — Coverage (±60 bp)
0 nM protein
3 nM protein
14 nM protein
70 nM protein
350 nM protein
2.5
0.0
−60
−40
−20
0
20
40
60
Position relative to center (bp)

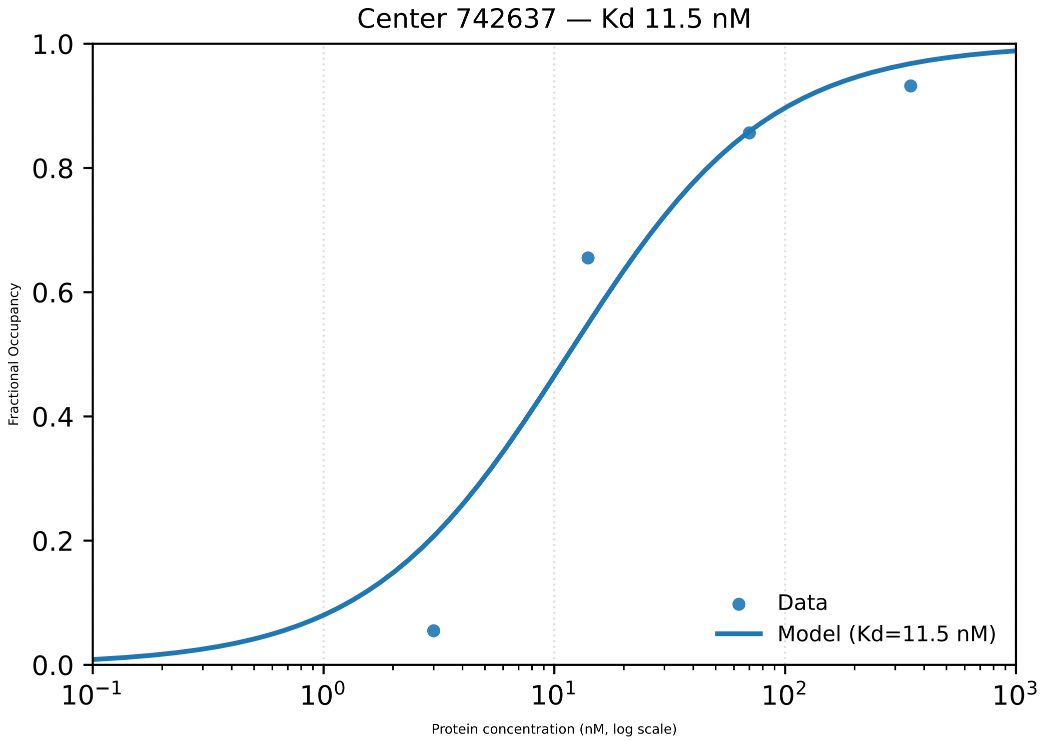

Center 742637 — Kd 11.5 nM
Fractional Occupancy
1.0
0.8
0.6
0.4
0.2
0.0
$10^{-1}$
$10^{0}$
$10^{1}$
$10^{2}$
$10^{3}$
Protein concentration (nM, log scale)
Data
Model (Kd=11.5 nM)

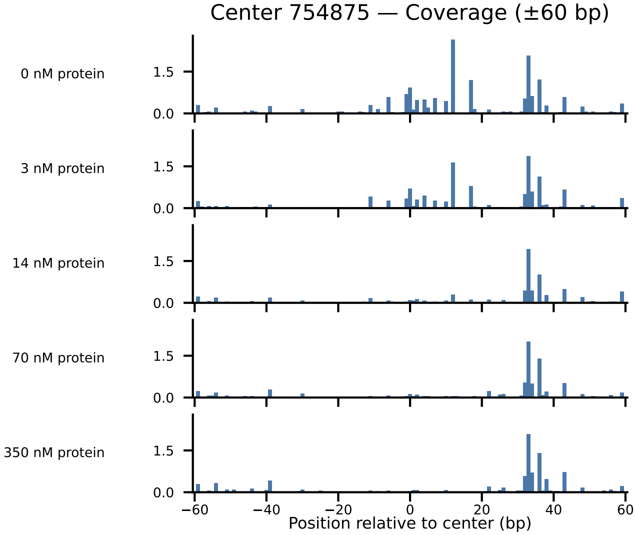

Center 754875 — Coverage (±60 bp)
0 nM protein
3 nM protein
14 nM protein
70 nM protein
350 nM protein
1.5
0.0
−60
−40
−20
0
20
40
60
Position relative to center (bp)

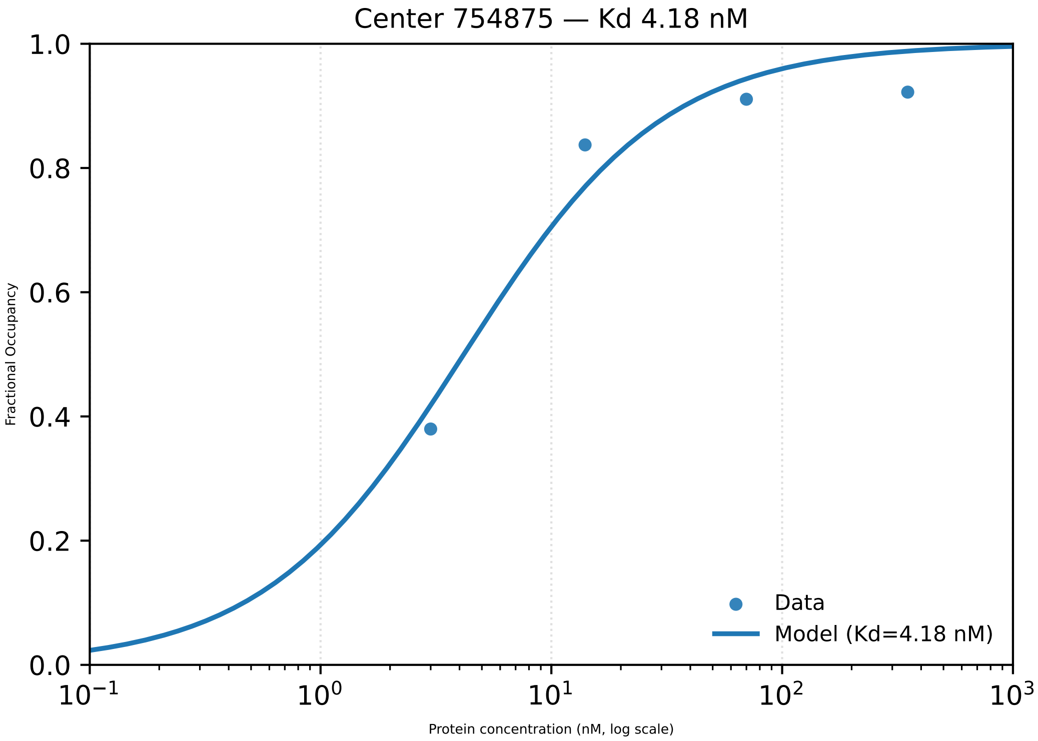

Center 754875 — Kd 4.18 nM
1.0
0.8
0.6
0.4
0.2
0.0
Fractional Occupancy
$10^{-1}$
$10^{0}$
$10^{1}$
$10^{2}$
$10^{3}$
Protein concentration (nM, log scale)
Data
Model (Kd=4.18 nM)

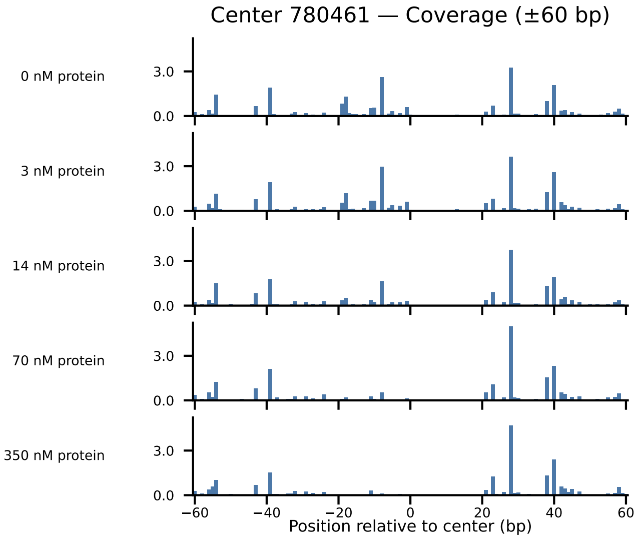

Center 780461 — Coverage (±60 bp)
0 nM protein
3 nM protein
14 nM protein
70 nM protein
350 nM protein
3.0
0.0
−60
−40
−20
0
20
40
60
Position relative to center (bp)

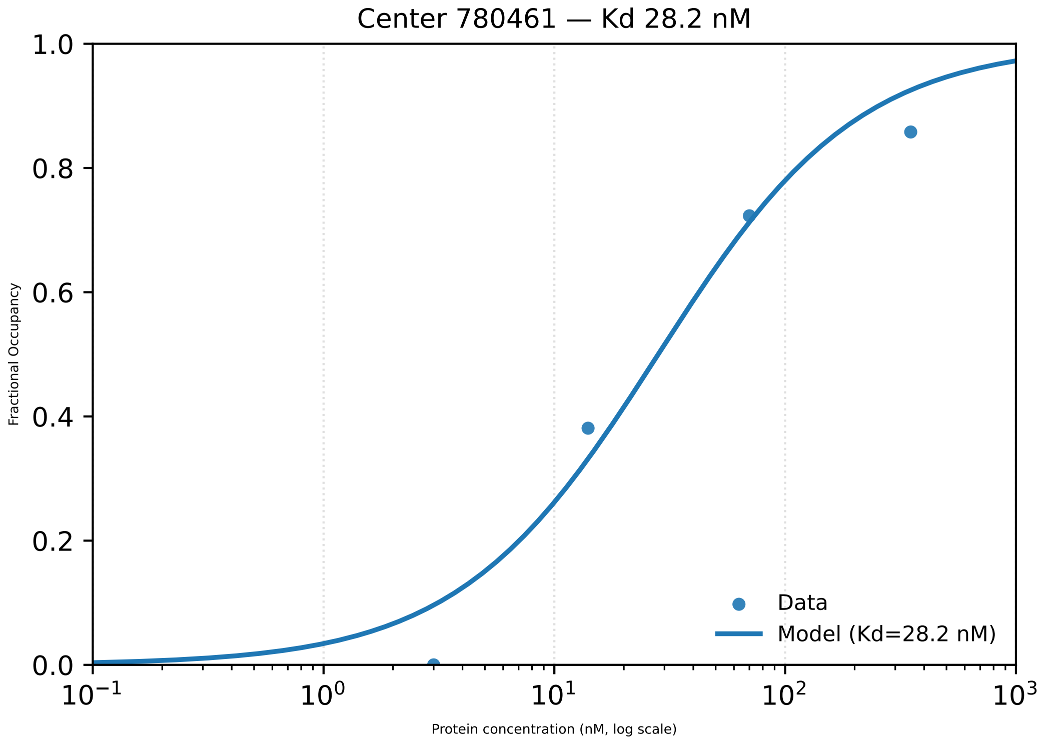

Center 780461 — Kd 28.2 nM
Fractional Occupancy
Protein concentration (nM, log scale)
1.0
0.8
0.6
0.4
0.2
0.0
$10^{-1}$
$10^{0}$
$10^{1}$
$10^{2}$
$10^{3}$
Data
Model (Kd=28.2 nM)

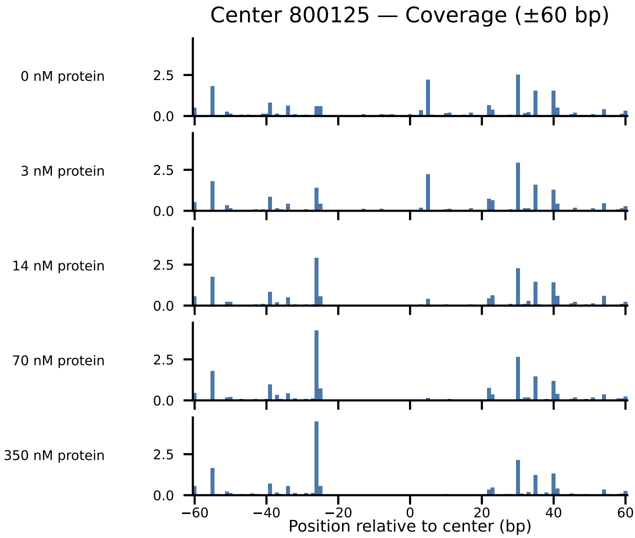

Center 800125 — Coverage (±60 bp)
0 nM protein
3 nM protein
14 nM protein
70 nM protein
350 nM protein
2.5
0.0
−60
−40
−20
0
20
40
60
Position relative to center (bp)

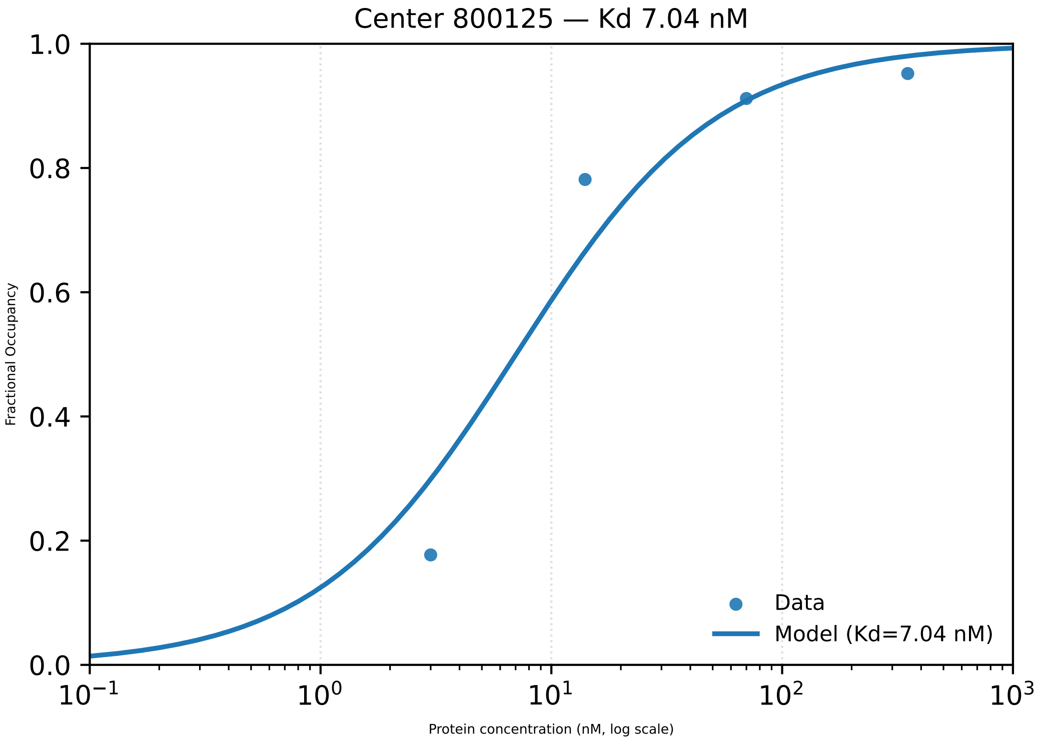

Center 800125 — Kd 7.04 nM
1.0
0.8
0.6
0.4
0.2
0.0
Fractional Occupancy
$10^{-1}$
$10^{0}$
$10^{1}$
$10^{2}$
$10^{3}$
Protein concentration (nM, log scale)
Data
Model (Kd=7.04 nM)

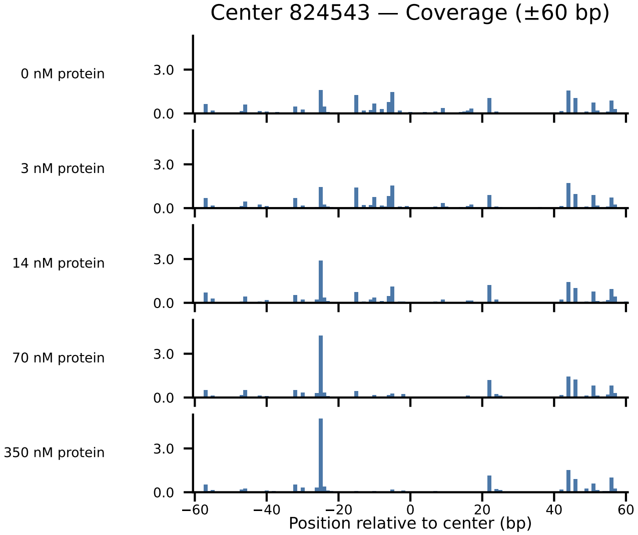
Center 824543 — Coverage (±60 bp)
0 nM protein
3 nM protein
14 nM protein
70 nM protein
350 nM protein
3.0
0.0
−60
−40
−20
0
20
40
60
Position relative to center (bp)

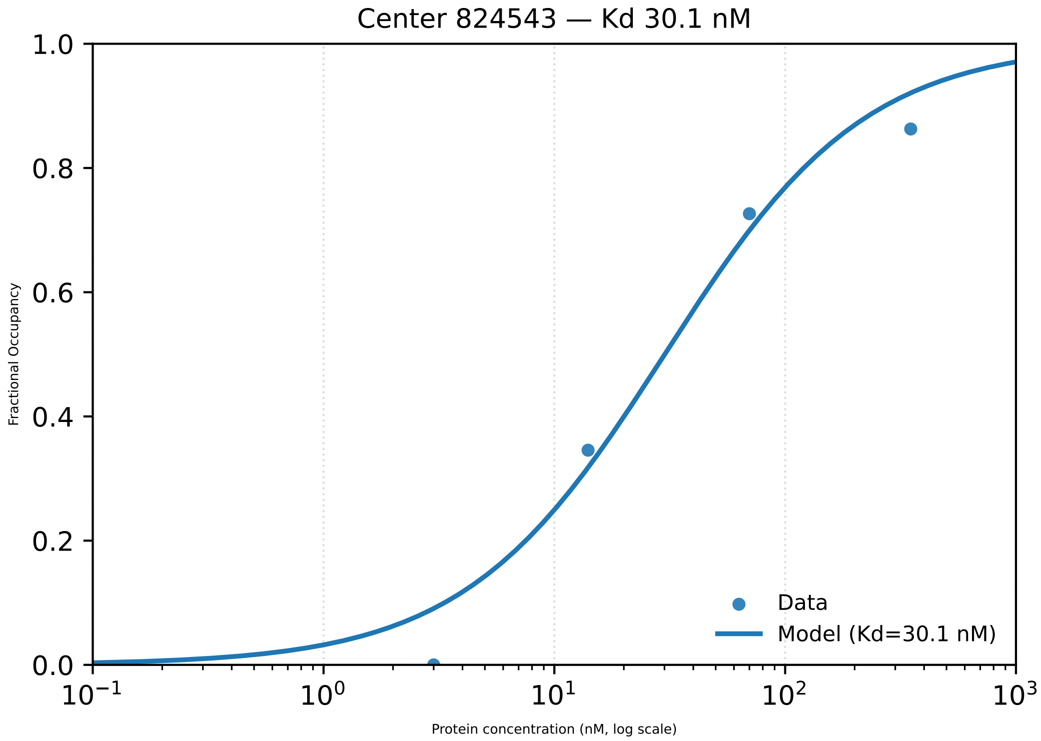

Center 824543 — Kd 30.1 nM
1.0
0.8
0.6
0.4
0.2
0.0
Fractional Occupancy
Data
Model (Kd=30.1 nM)
$10^{-1}$
$10^{0}$
$10^{1}$
$10^{2}$
$10^{3}$
Protein concentration (nM, log scale)

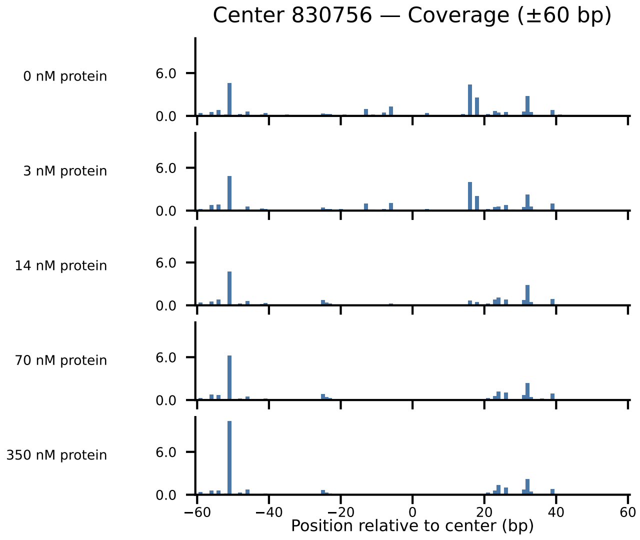

Center 830756 — Coverage (±60 bp)
0 nM protein
3 nM protein
14 nM protein
70 nM protein
350 nM protein
6.0
0.0
−60
−40
−20
0
20
40
60
Position relative to center (bp)

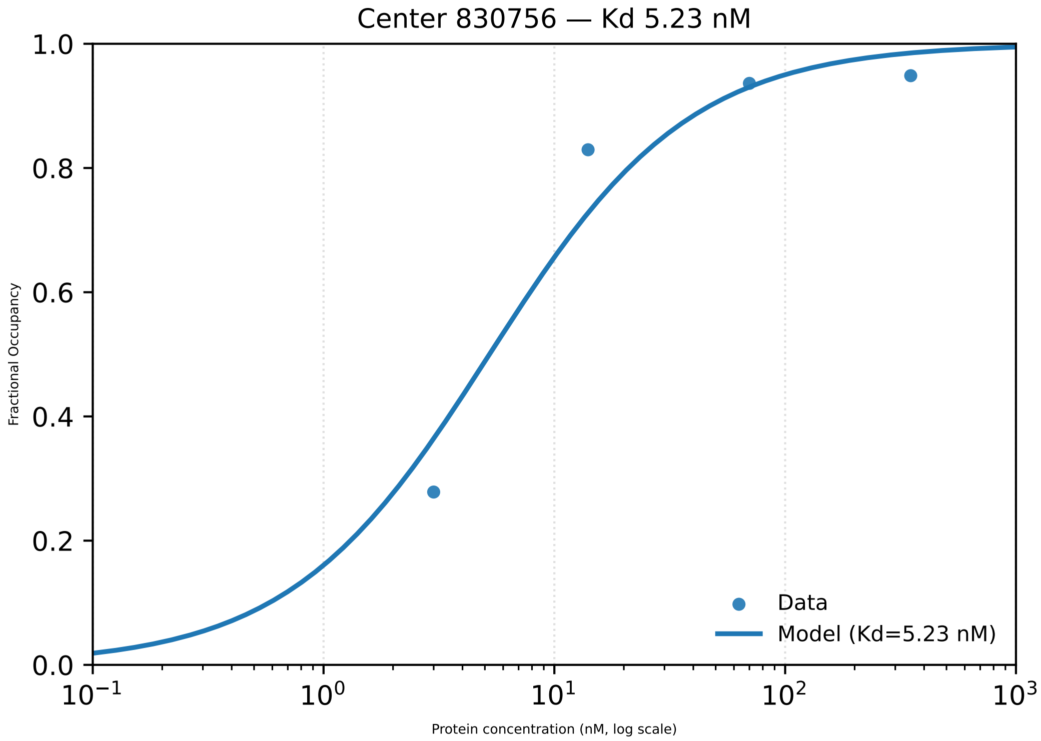

Center 830756 — Kd 5.23 nM
1.0
0.8
0.6
0.4
0.2
0.0
Fractional Occupancy
$10^{-1}$
$10^{0}$
$10^{1}$
$10^{2}$
$10^{3}$
Protein concentration (nM, log scale)
Data
Model (Kd=5.23 nM)

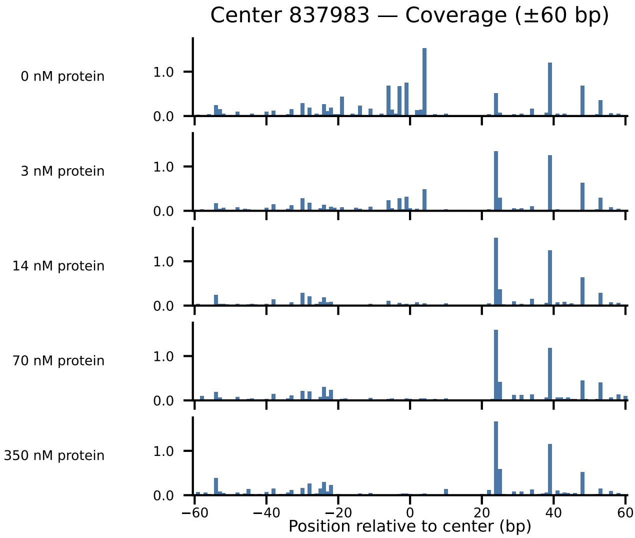

Center 837983 — Coverage (±60 bp)
0 nM protein
3 nM protein
14 nM protein
70 nM protein
350 nM protein
1.0
0.0
−60
−40
−20
0
20
40
60
Position relative to center (bp)

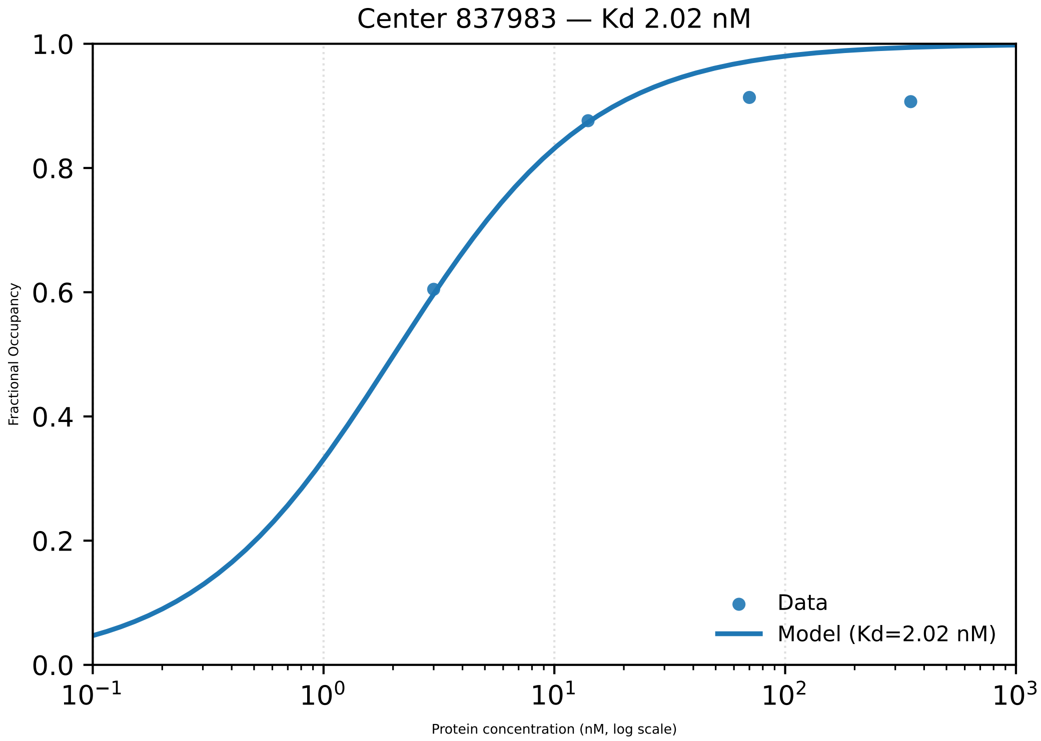

Center 837983 — Kd 2.02 nM
Fractional Occupancy
Protein concentration (nM, log scale)
Data
Model (Kd=2.02 nM)
1.0
0.8
0.6
0.4
0.2
0.0
$10^{-1}$
$10^{0}$
$10^{1}$
$10^{2}$
$10^{3}$

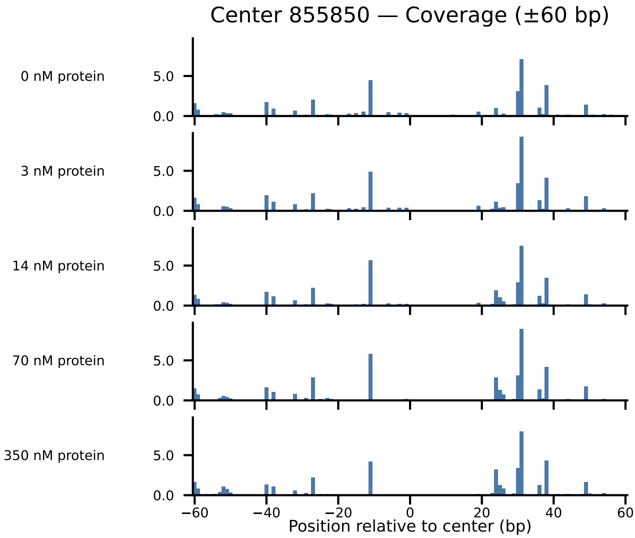

Center 855850 — Coverage (±60 bp)
0 nM protein
3 nM protein
14 nM protein
70 nM protein
350 nM protein
5.0
0.0
−60
−40
−20
0
20
40
60
Position relative to center (bp)

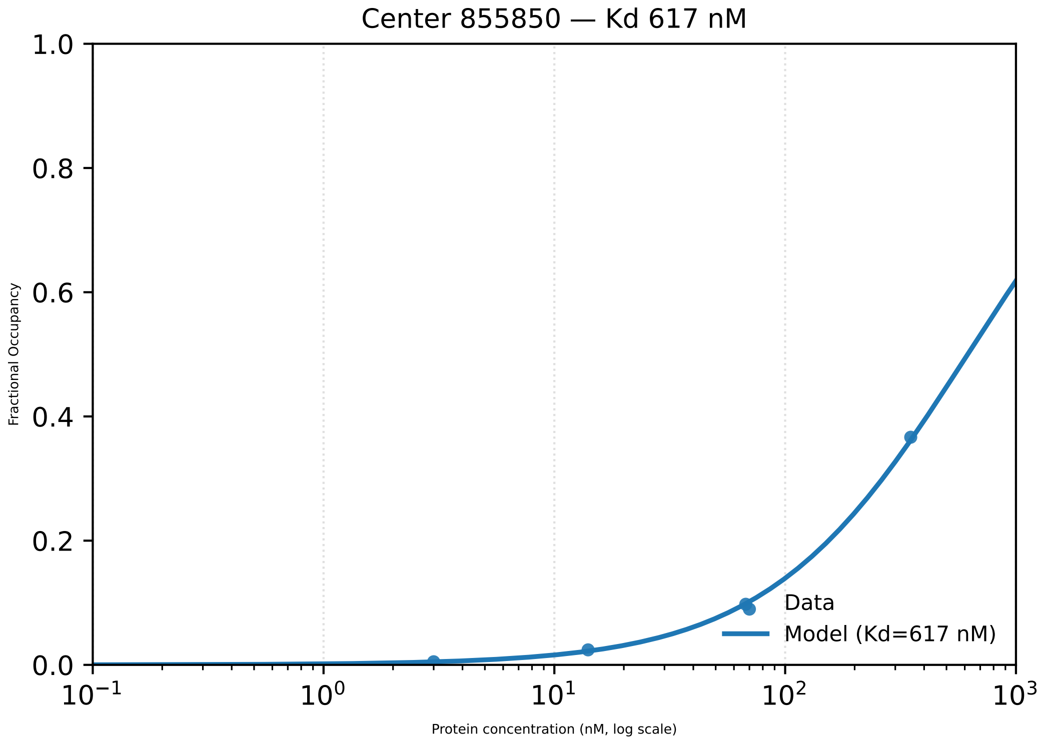

Center 855850 — Kd 617 nM
1.0
0.8
0.6
0.4
0.2
0.0
Fractional Occupancy
$10^{-1}$
$10^{0}$
$10^{1}$
$10^{2}$
$10^{3}$
Protein concentration (nM, log scale)
Data
Model (Kd=617 nM)

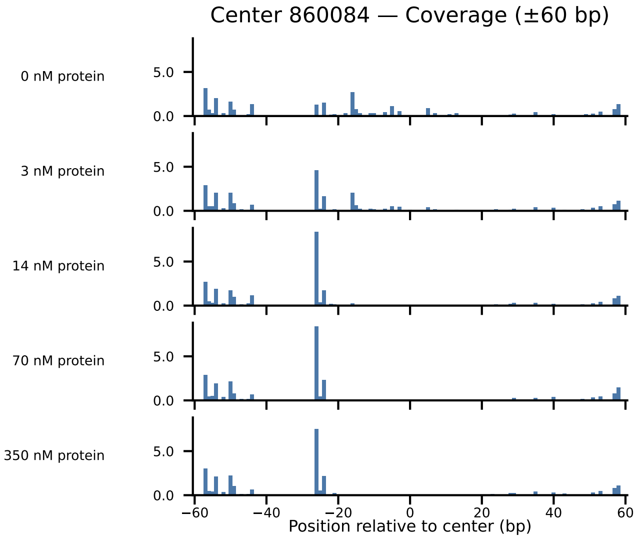

Center 860084 — Coverage (±60 bp)
0 nM protein
3 nM protein
14 nM protein
70 nM protein
350 nM protein
5.0
0.0
−60
−40
−20
0
20
40
60
Position relative to center (bp)

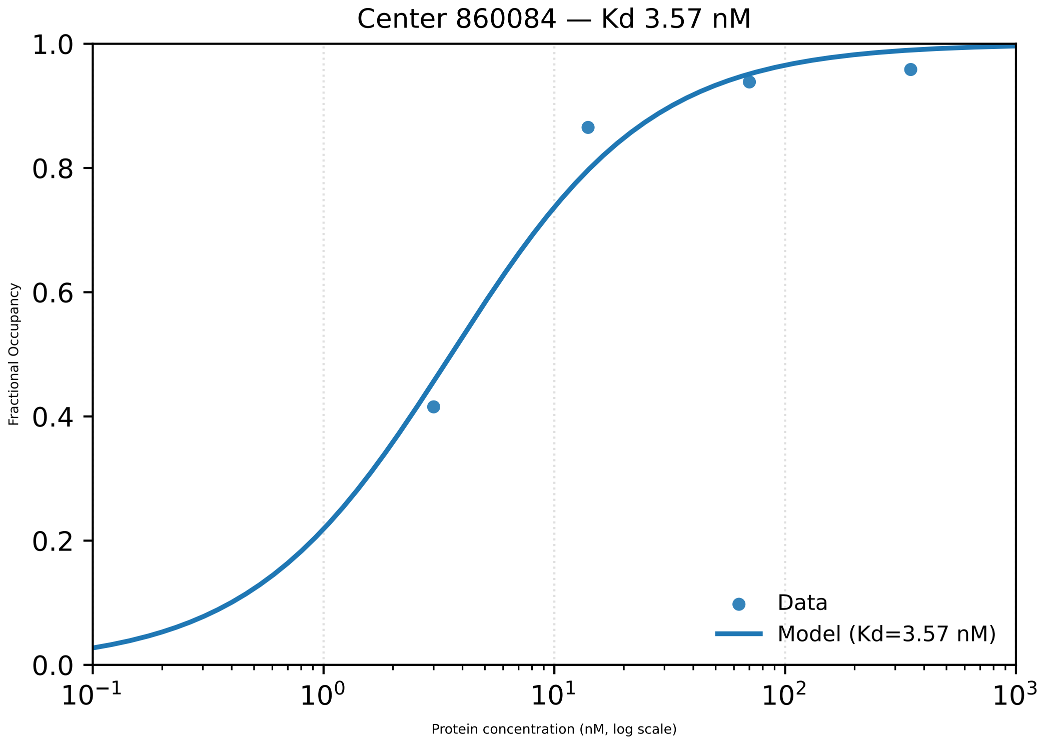

Center 860084 — Kd 3.57 nM
1.0
0.8
0.6
0.4
0.2
0.0
Fractional Occupancy
$10^{-1}$
$10^{0}$
$10^{1}$
$10^{2}$
$10^{3}$
Protein concentration (nM, log scale)
Data
Model (Kd=3.57 nM)

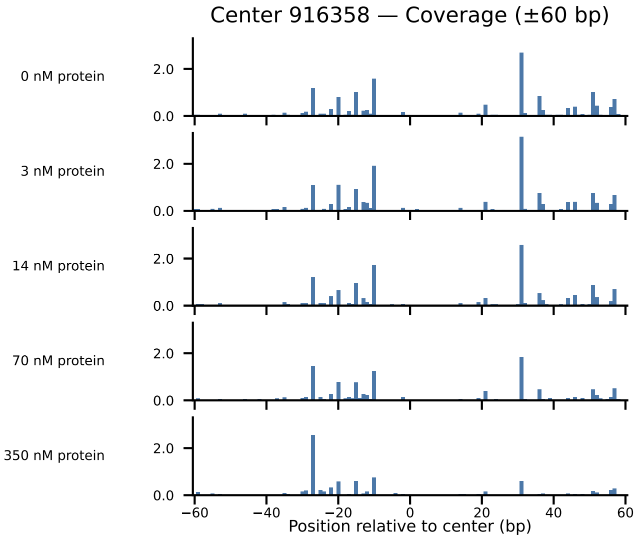
Center 916358 — Coverage (±60 bp)
0 nM protein
3 nM protein
14 nM protein
70 nM protein
350 nM protein
2.0
0.0
−60
−40
−20
0
20
40
60
Position relative to center (bp)

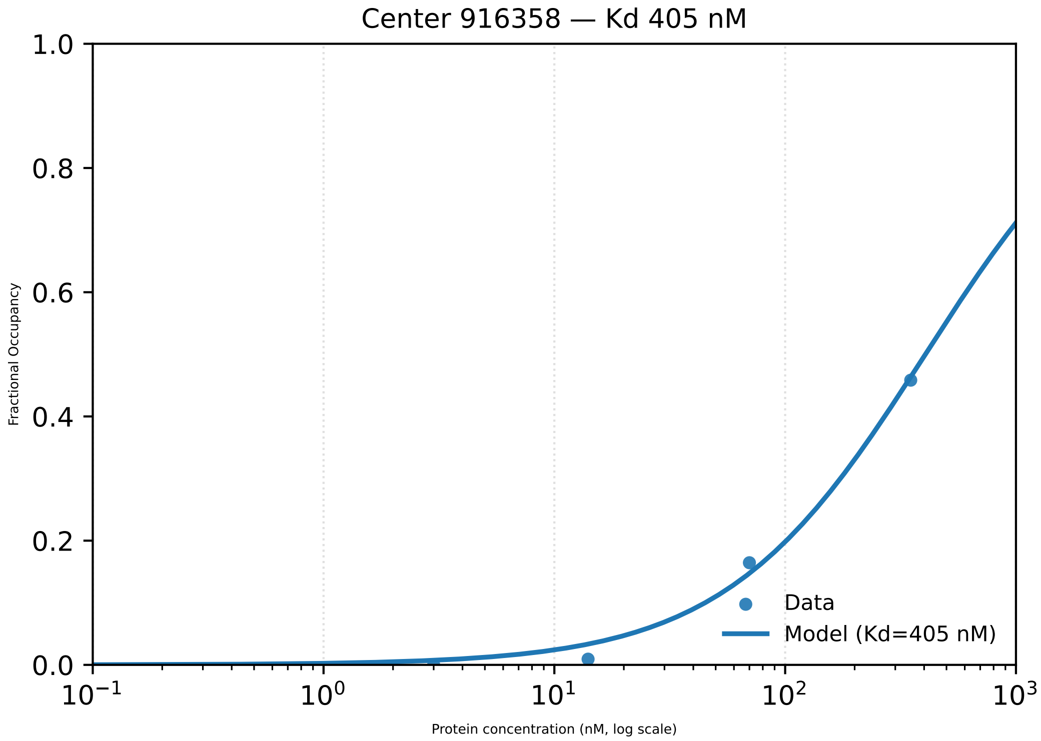
Center 916358 — Kd 405 nM
1.0
0.8
0.6
0.4
0.2
0.0
Fractional Occupancy
$10^{-1}$
$10^{0}$
$10^{1}$
$10^{2}$
$10^{3}$
Protein concentration (nM, log scale)
Data
Model (Kd=405 nM)

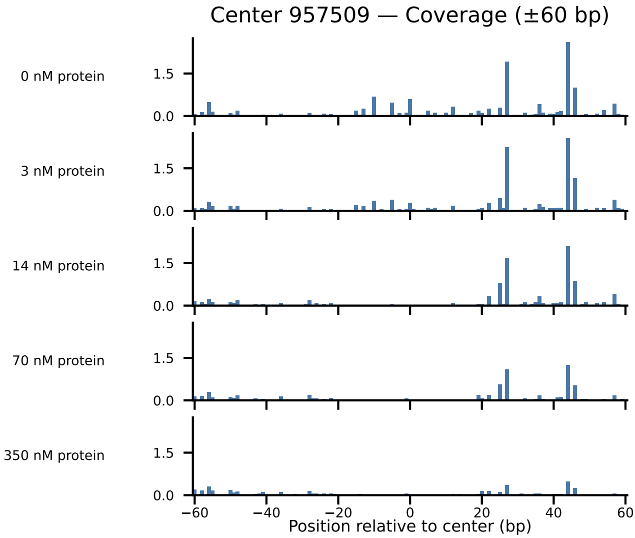

Center 957509 — Coverage (±60 bp)
0 nM protein
3 nM protein
14 nM protein
70 nM protein
350 nM protein
1.5
0.0
−60
−40
−20
0
20
40
60
Position relative to center (bp)

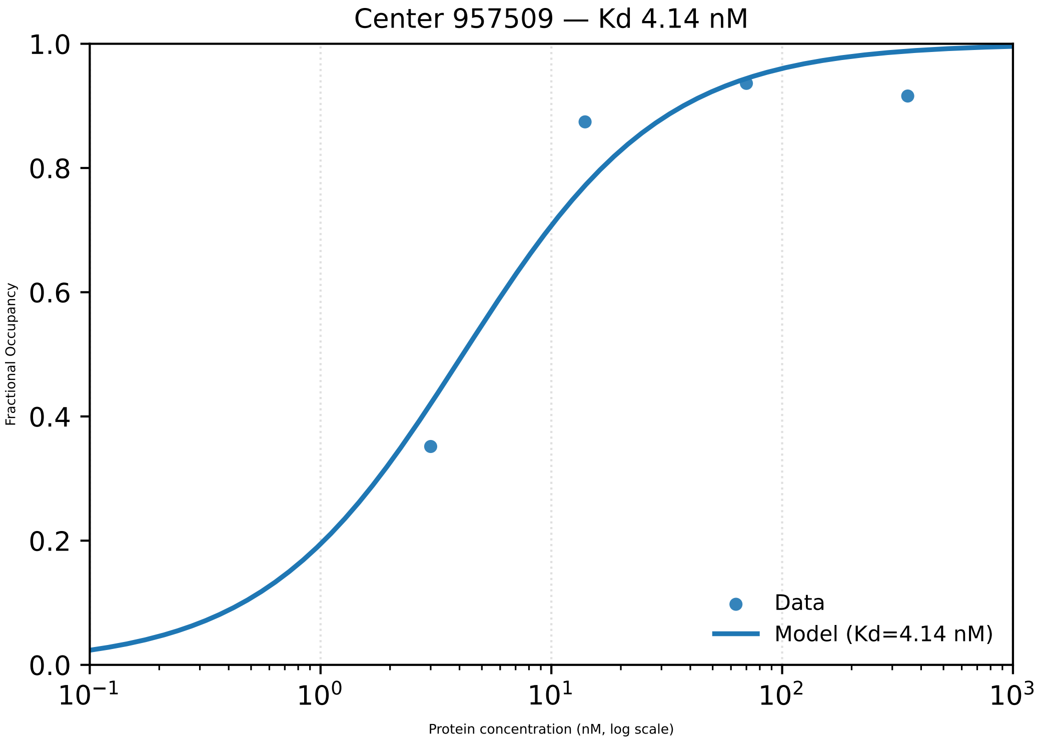

Center 957509 — Kd 4.14 nM
1.0
0.8
0.6
0.4
0.2
0.0
Fractional Occupancy
$10^{-1}$
$10^{0}$
$10^{1}$
$10^{2}$
$10^{3}$
Protein concentration (nM, log scale)
Data
Model (Kd=4.14 nM)

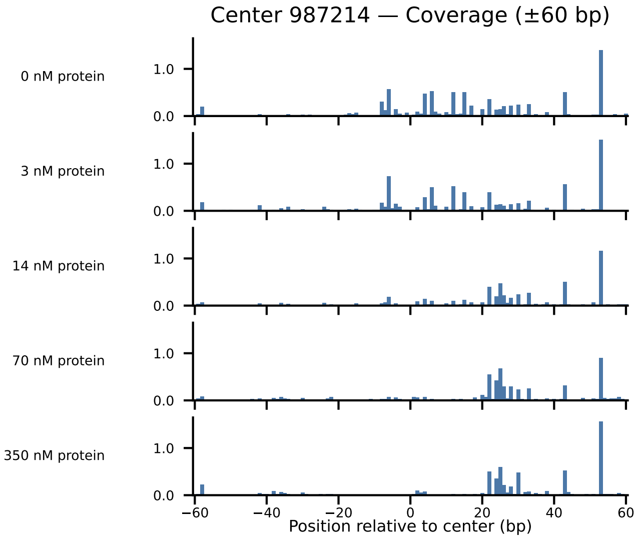
Center 987214 — Coverage (±60 bp)
0 nM protein
3 nM protein
14 nM protein
70 nM protein
350 nM protein
1.0
0.0
−60
−40
−20
0
20
40
60
Position relative to center (bp)

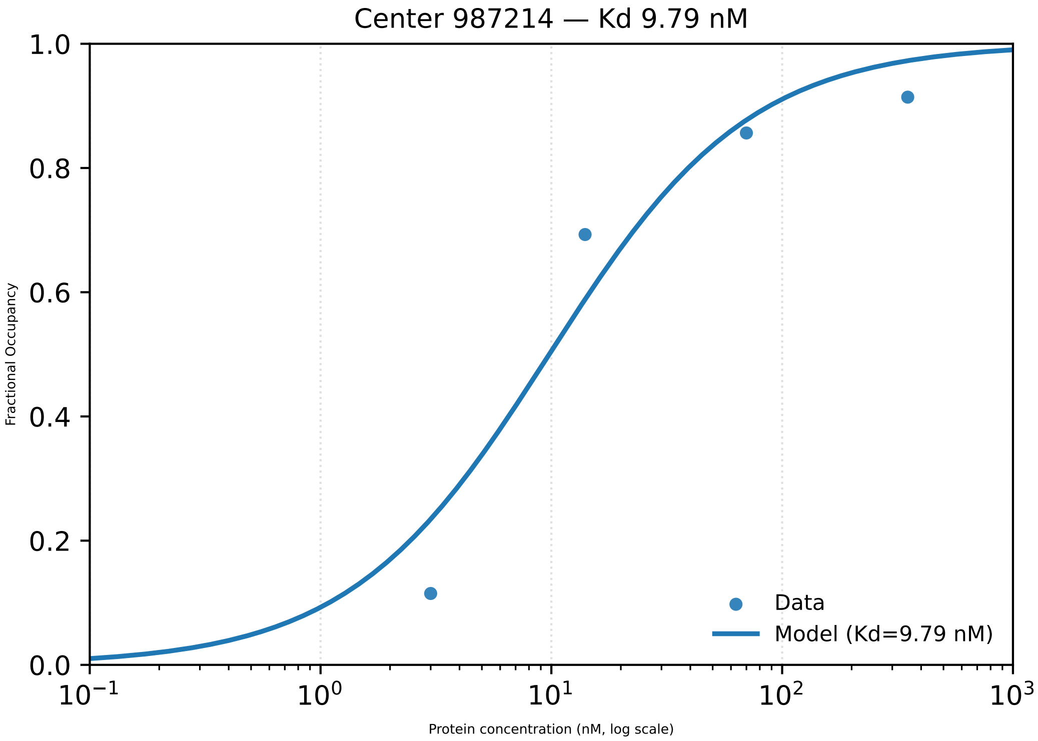

Center 987214 — Kd 9.79 nM
Fractional Occupancy
0.0
0.2
0.4
0.6
0.8
1.0
$10^{-1}$
$10^{0}$
$10^{1}$
$10^{2}$
$10^{3}$
Protein concentration (nM, log scale)
Data
Model (Kd=9.79 nM)

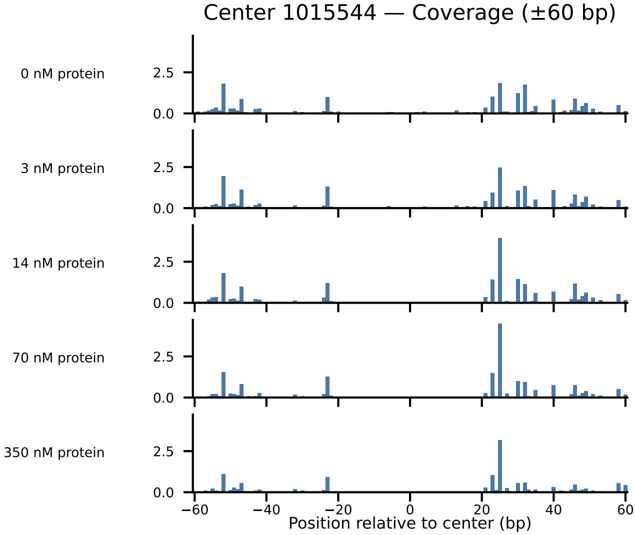

Center 1015544 — Coverage (±60 bp)
0 nM protein
3 nM protein
14 nM protein
70 nM protein
350 nM protein
2.5
0.0
−60
−40
−20
0
20
40
60
Position relative to center (bp)

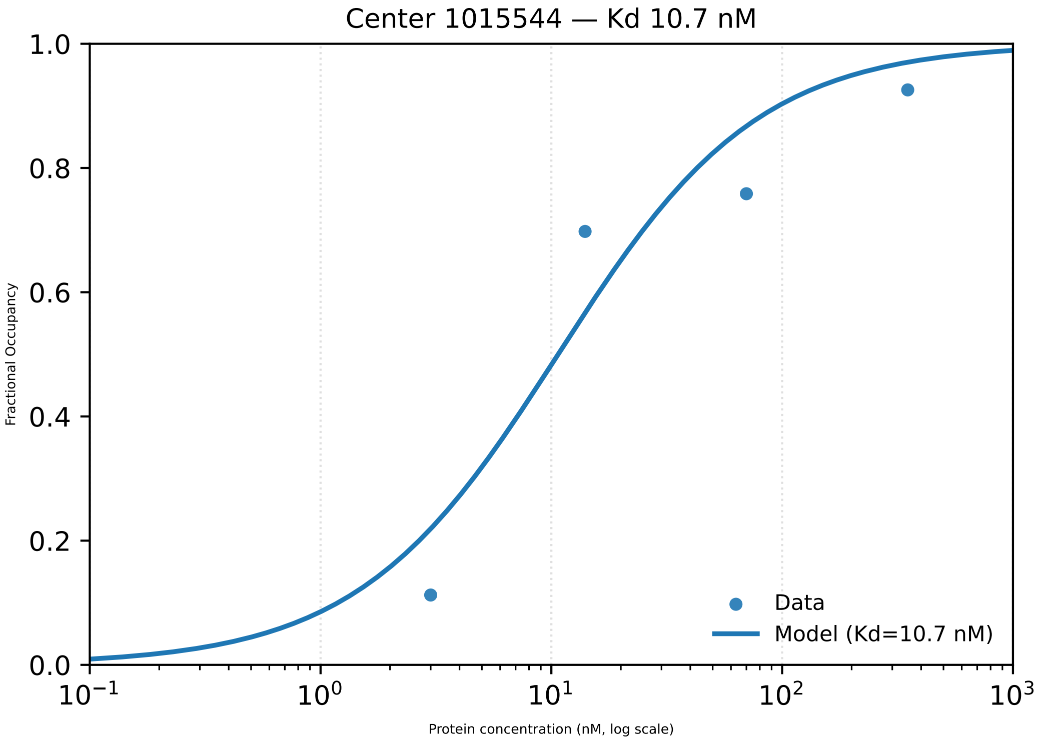

Center 1015544 — Kd 10.7 nM
1.0
0.8
0.6
0.4
0.2
0.0
Fractional Occupancy
$10^{-1}$
$10^{0}$
$10^{1}$
$10^{2}$
$10^{3}$
Protein concentration (nM, log scale)
Data
Model (Kd=10.7 nM)

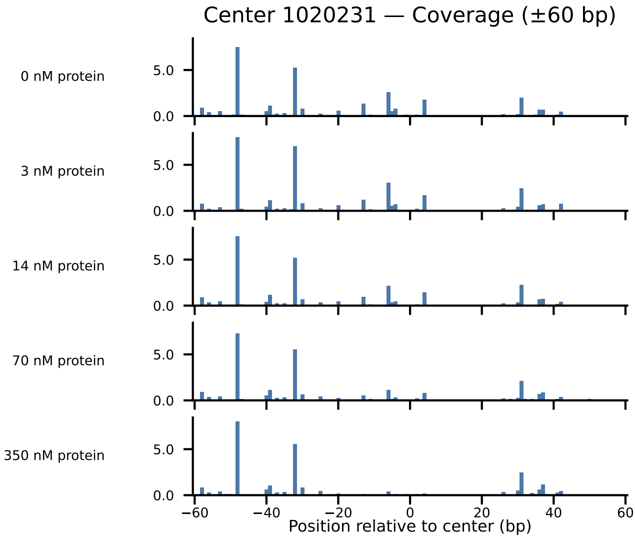

Center 1020231 — Coverage (±60 bp)
0 nM protein
3 nM protein
14 nM protein
70 nM protein
350 nM protein
5.0
0.0
−60
−40
−20
0
20
40
60
Position relative to center (bp)

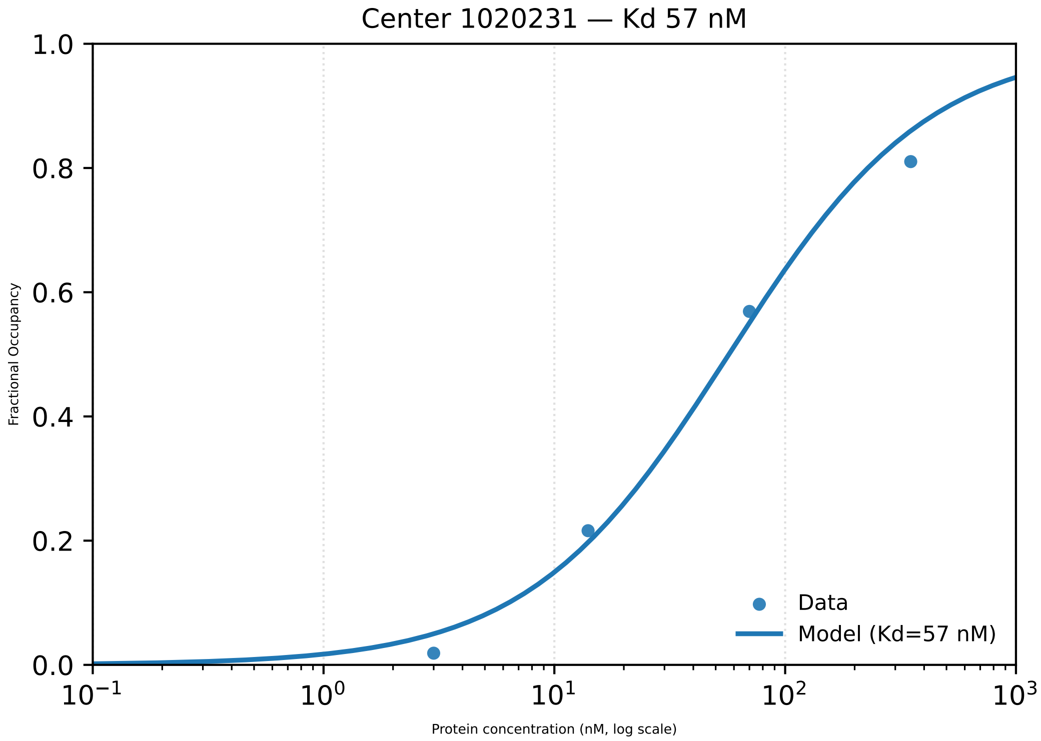

Center 1020231 — Kd 57 nM
Fractional Occupancy
0.0
0.2
0.4
0.6
0.8
1.0
$10^{-1}$
$10^{0}$
$10^{1}$
$10^{2}$
$10^{3}$
Protein concentration (nM, log scale)
Data
Model (Kd=57 nM)

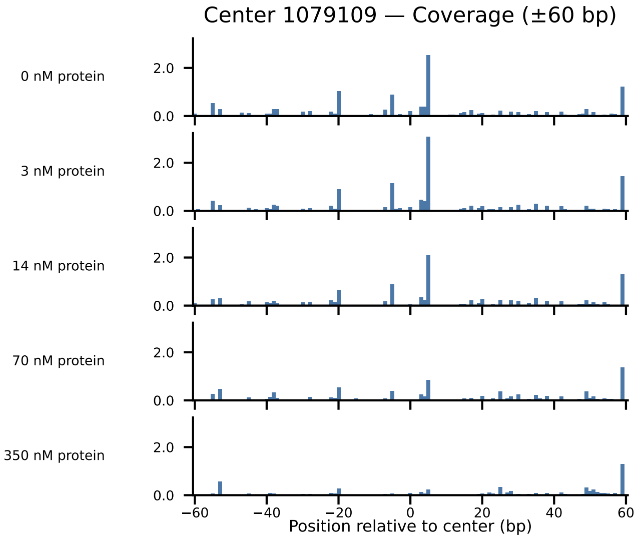
Center 1079109 — Coverage (±60 bp)
0 nM protein
3 nM protein
14 nM protein
70 nM protein
350 nM protein
2.0
0.0
−60
−40
−20
0
20
40
60
Position relative to center (bp)

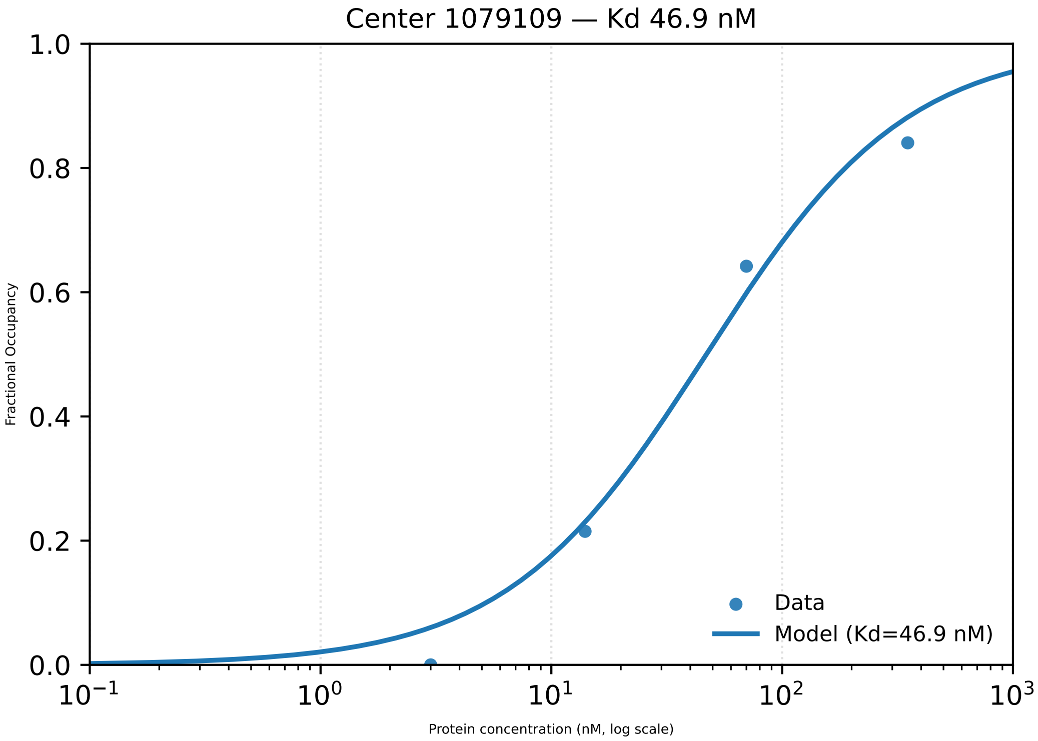
Center 1079109 — Kd 46.9 nM
1.0
0.8
0.6
0.4
0.2
0.0
Fractional Occupancy
$10^{-1}$
$10^{0}$
$10^{1}$
$10^{2}$
$10^{3}$
Protein concentration (nM, log scale)
Data
Model (Kd=46.9 nM)

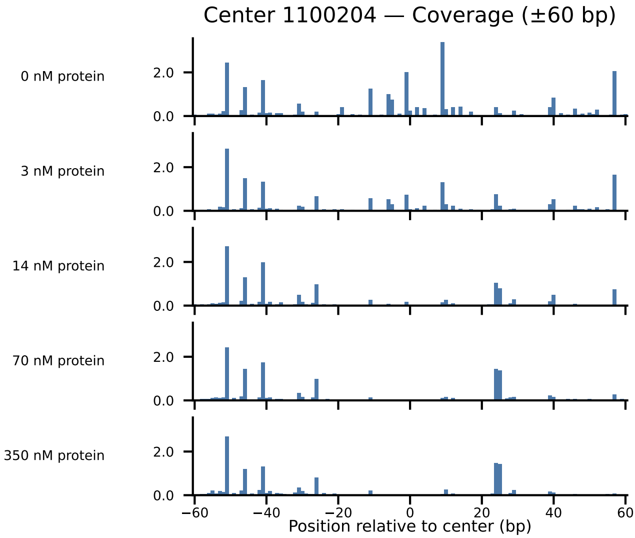
Center 1100204 — Coverage (±60 bp)
0 nM protein
3 nM protein
14 nM protein
70 nM protein
350 nM protein
2.0
0.0
−60
−40
−20
0
20
40
60
Position relative to center (bp)

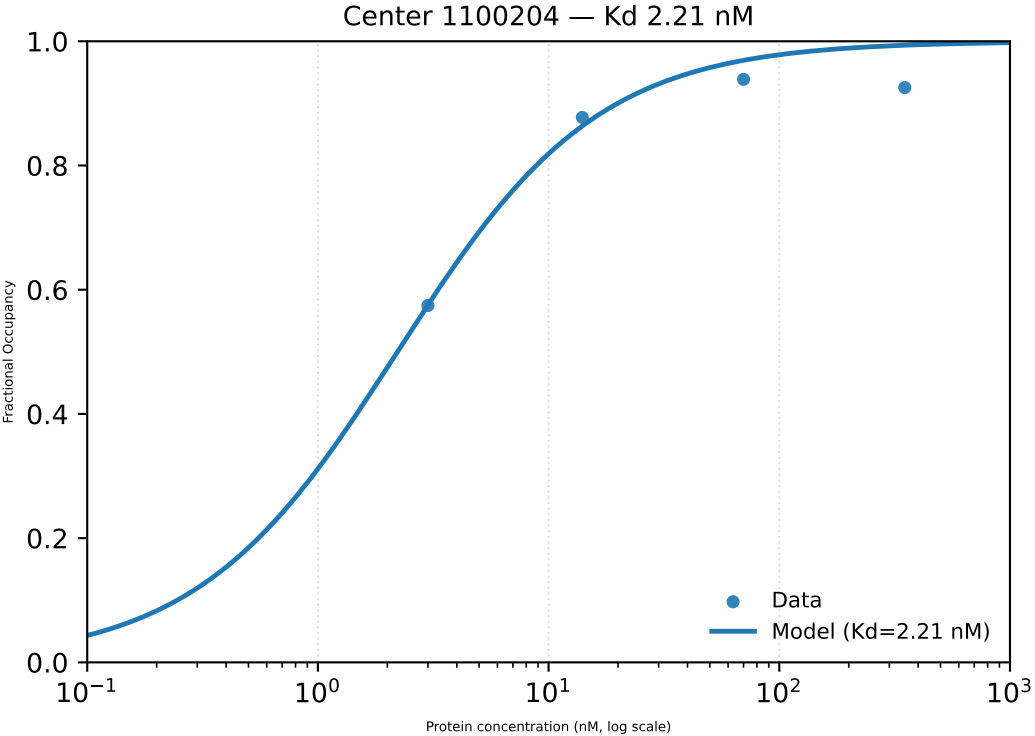

Center 1100204 — Kd 2.21 nM
1.0
0.8
0.6
0.4
0.2
0.0
Fractional Occupancy
Data
Model (Kd=2.21 nM)
$10^{-1}$
$10^{0}$
$10^{1}$
$10^{2}$
$10^{3}$
Protein concentration (nM, log scale)

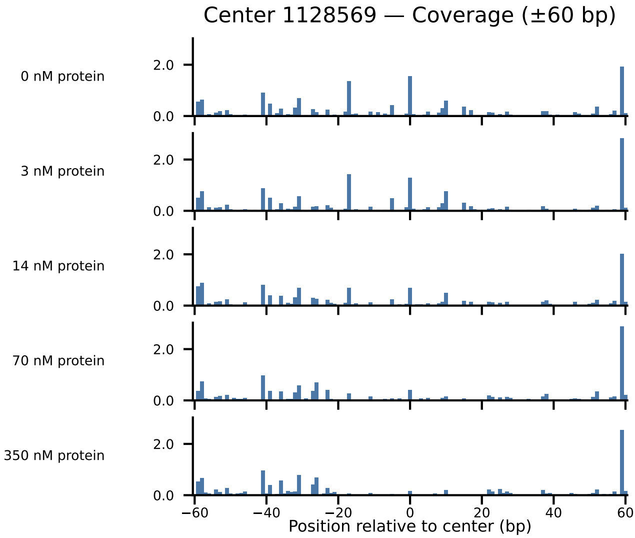
Center 1128569 — Coverage (±60 bp)
0 nM protein
3 nM protein
14 nM protein
70 nM protein
350 nM protein
0.0
2.0
−60
−40
−20
0
20
40
60
Position relative to center (bp)

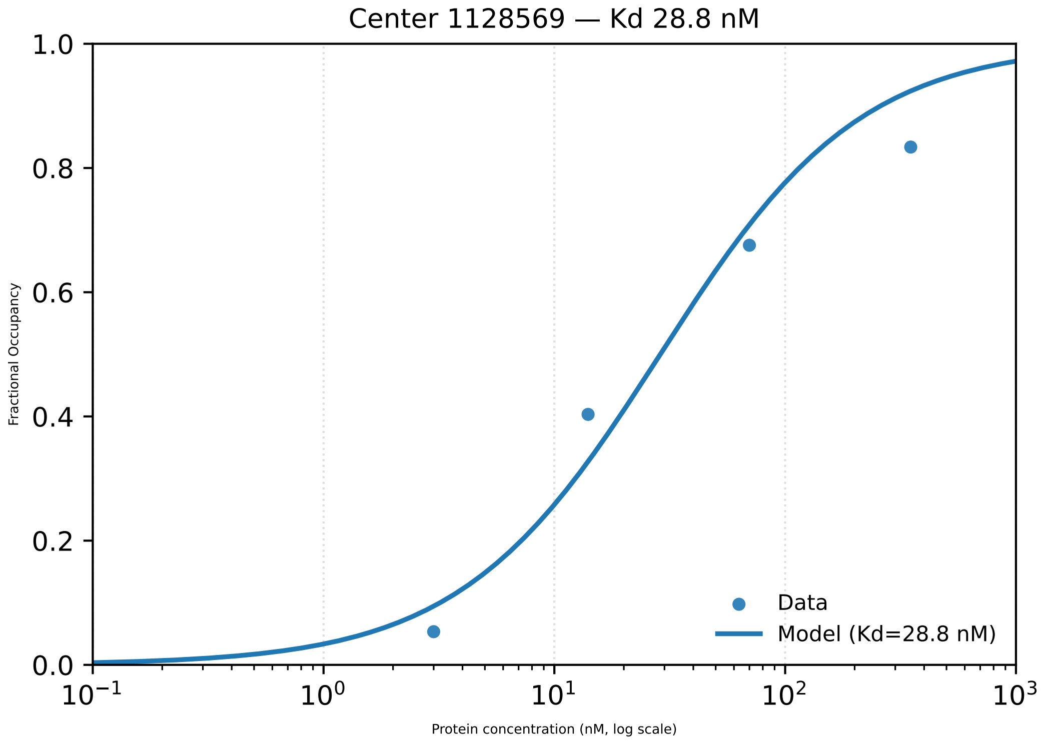
Center 1128569 — Kd 28.8 nM
Fractional Occupancy
1.0
0.8
0.6
0.4
0.2
0.0
$10^{-1}$
$10^{0}$
$10^{1}$
$10^{2}$
$10^{3}$
Protein concentration (nM, log scale)
Data
Model (Kd=28.8 nM)

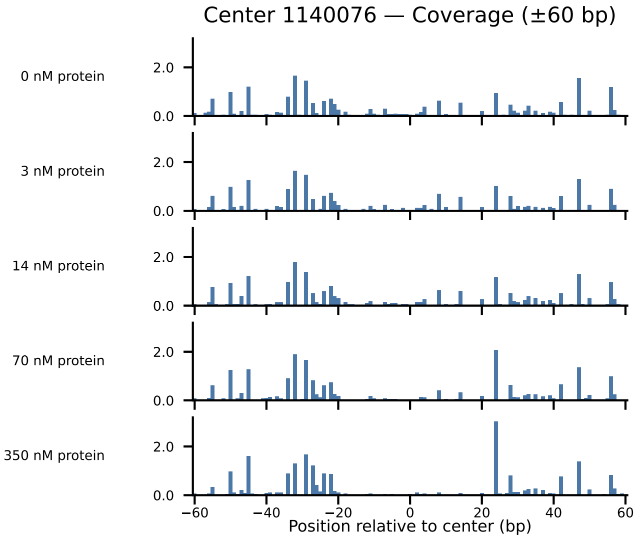
Center 1140076 — Coverage (±60 bp)
0 nM protein
3 nM protein
14 nM protein
70 nM protein
350 nM protein
2.0
0.0
−60
−40
−20
0
20
40
60
Position relative to center (bp)

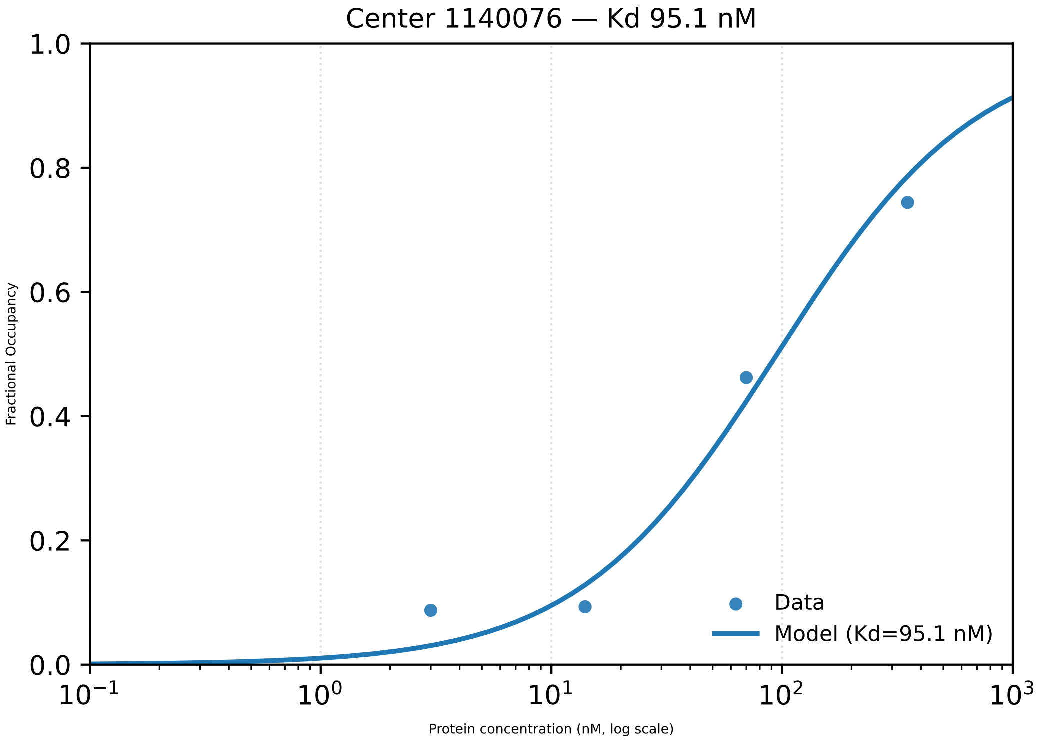
Center 1140076 — Kd 95.1 nM
Fractional Occupancy
1.0
0.8
0.6
0.4
0.2
0.0
$10^{-1}$
$10^{0}$
$10^{1}$
$10^{2}$
$10^{3}$
Protein concentration (nM, log scale)
Data
Model (Kd=95.1 nM)

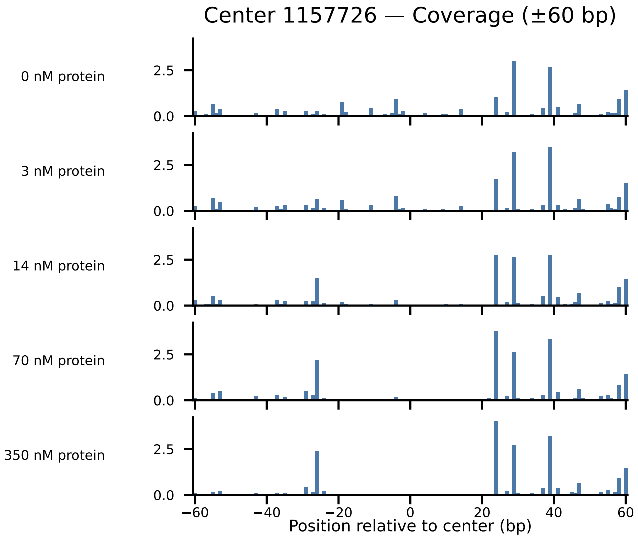
Center 1157726 — Coverage (±60 bp)
0 nM protein
3 nM protein
14 nM protein
70 nM protein
350 nM protein
0.0
2.5
−60
−40
−20
0
20
40
60
Position relative to center (bp)

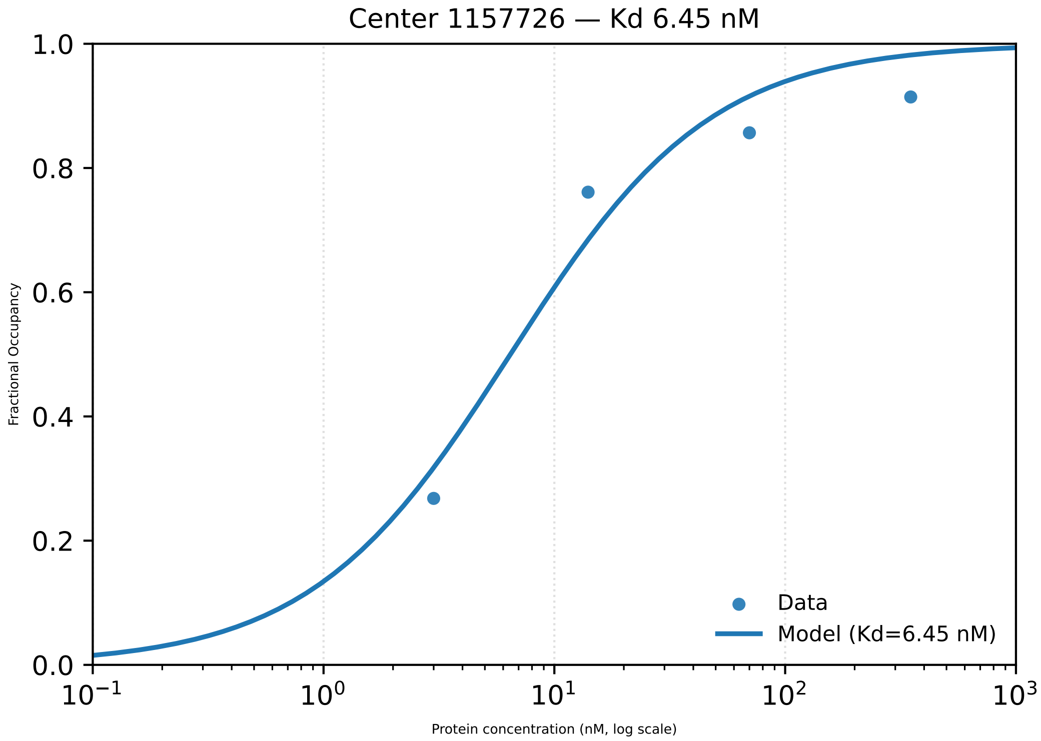

Center 1157726 — Kd 6.45 nM
1.0
0.8
0.6
0.4
0.2
0.0
Fractional Occupancy
$10^{-1}$
$10^{0}$
$10^{1}$
$10^{2}$
$10^{3}$
Protein concentration (nM, log scale)
Data
Model (Kd=6.45 nM)

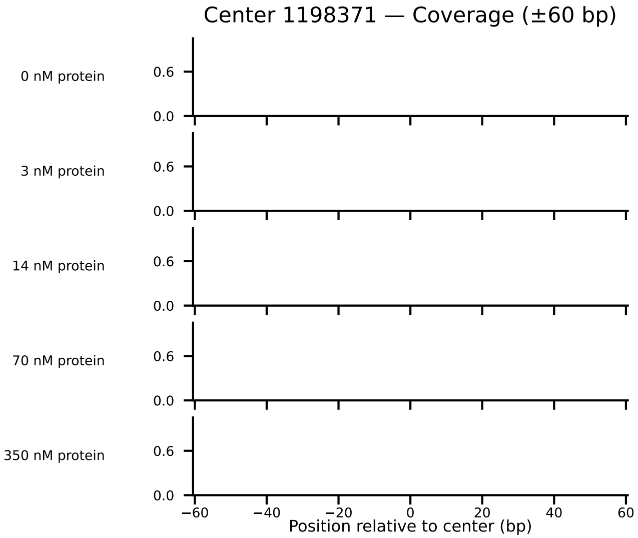

Center 1198371 — Coverage (±60 bp)
0 nM protein
3 nM protein
14 nM protein
70 nM protein
350 nM protein
0.6
0.0
−60
−40
−20
0
20
40
60
Position relative to center (bp)

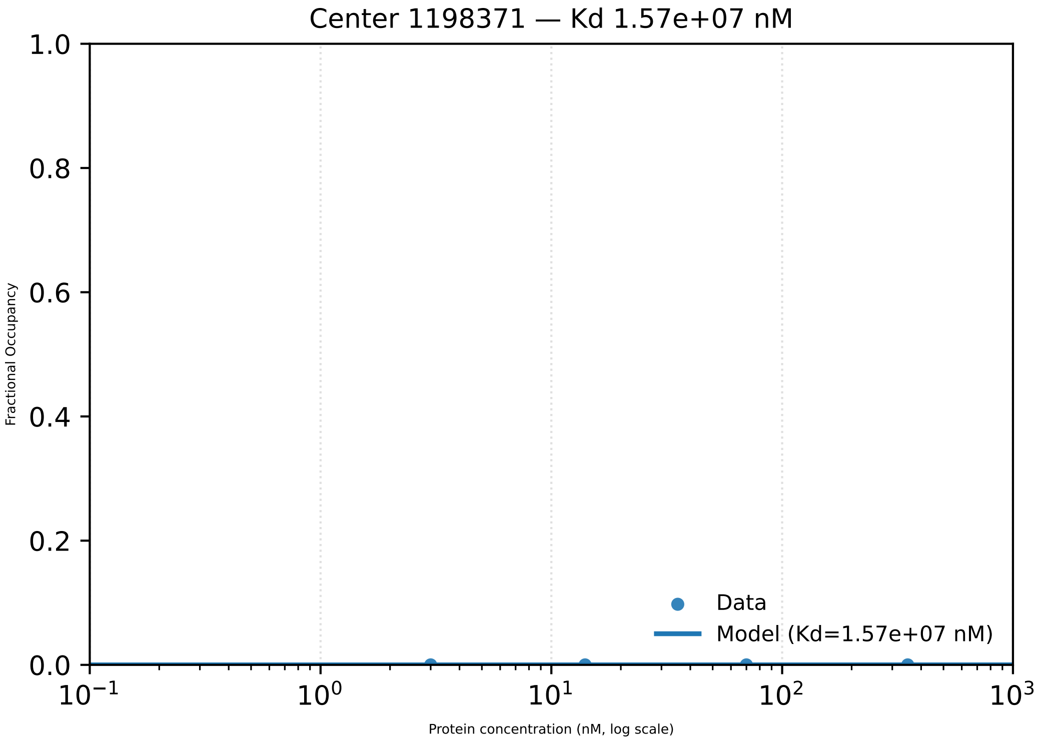

Center 1198371 — Kd 1.57e+07 nM
1.0
0.8
0.6
0.4
0.2
0.0
Fractional Occupancy
$10^{-1}$
$10^{0}$
$10^{1}$
$10^{2}$
$10^{3}$
Protein concentration (nM, log scale)
Data
Model (Kd=1.57e+07 nM)

Center 1219418 — Coverage (±60 bp)

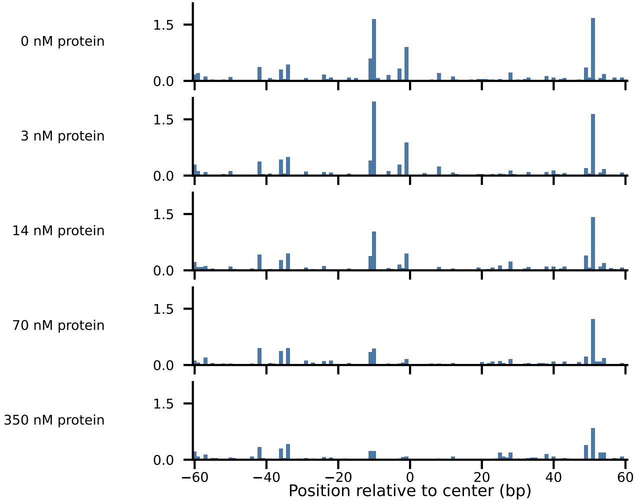

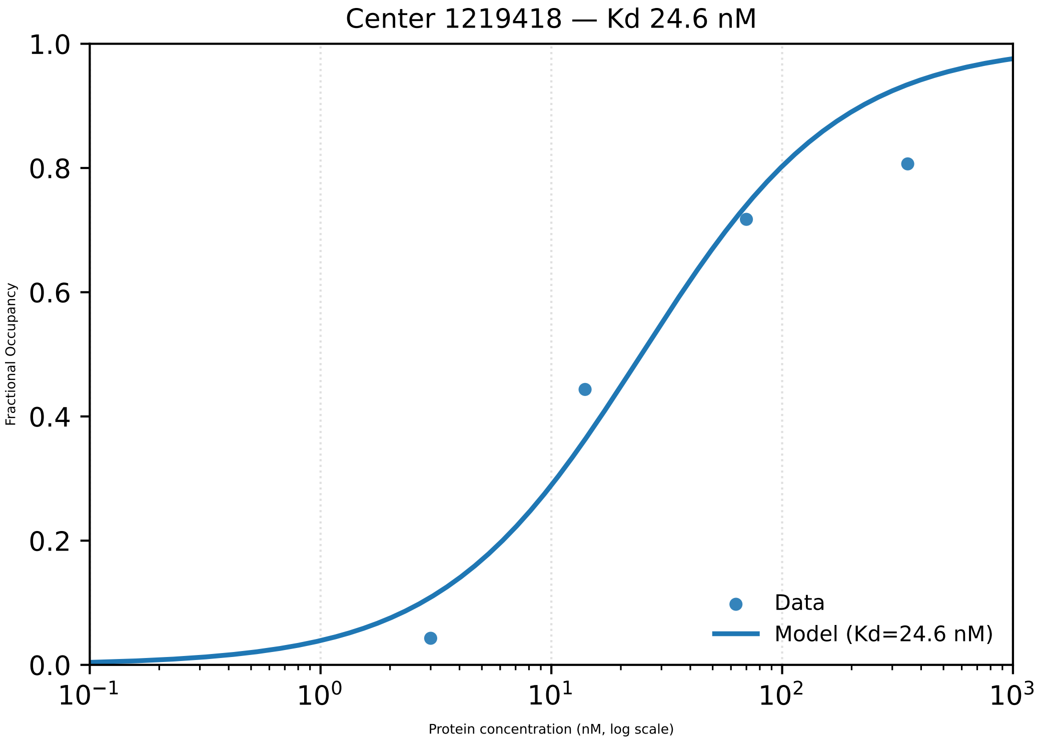

Center 1219418 — Kd 24.6 nM
Fractional Occupancy
1.0
0.8
0.6
0.4
0.2
0.0
$10^{-1}$
$10^{0}$
$10^{1}$
$10^{2}$
$10^{3}$
Protein concentration (nM, log scale)
Data
Model (Kd=24.6 nM)

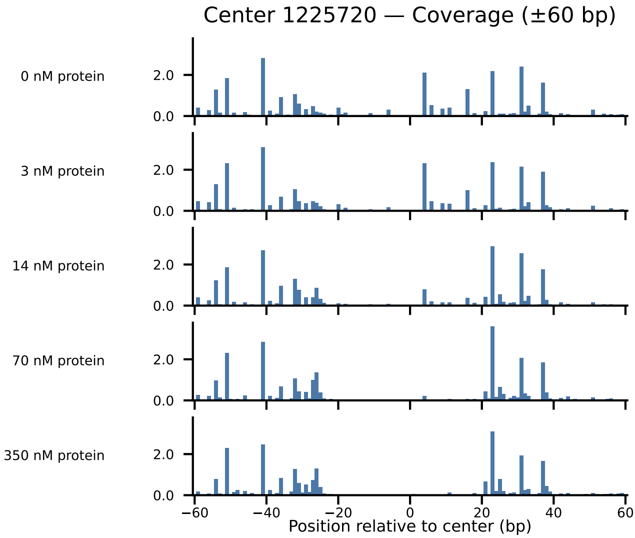

Center 1225720 — Coverage (±60 bp)
0 nM protein
3 nM protein
14 nM protein
70 nM protein
350 nM protein
2.0
0.0
−60
−40
−20
0
20
40
60
Position relative to center (bp)

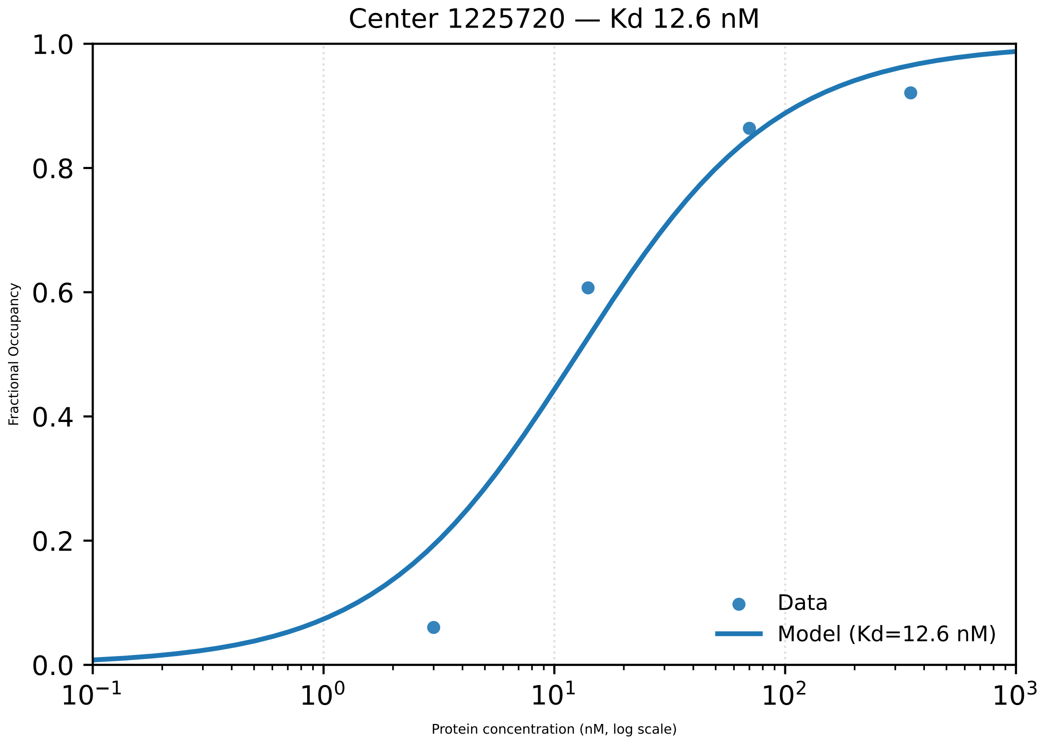

Center 1225720 — Kd 12.6 nM
1.0
0.8
0.6
0.4
0.2
0.0
Fractional Occupancy
$10^{-1}$
$10^{0}$
$10^{1}$
$10^{2}$
$10^{3}$
Protein concentration (nM, log scale)
Data
Model (Kd=12.6 nM)

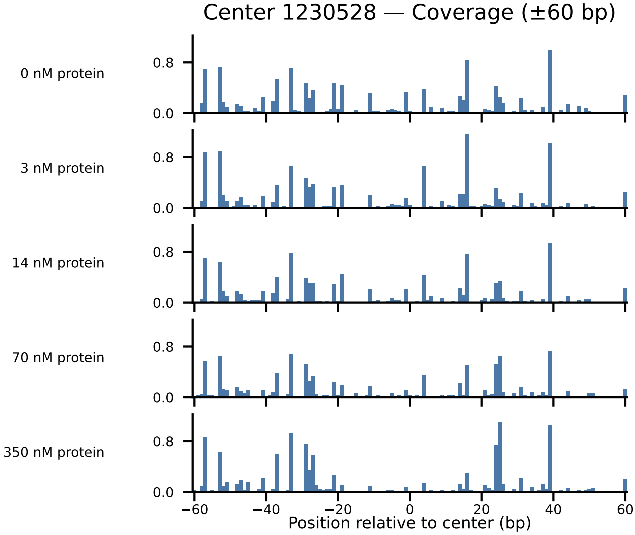

Center 1230528 — Coverage (±60 bp)
0 nM protein
3 nM protein
14 nM protein
70 nM protein
350 nM protein
0.8
0.0
−60
−40
−20
0
20
40
60
Position relative to center (bp)

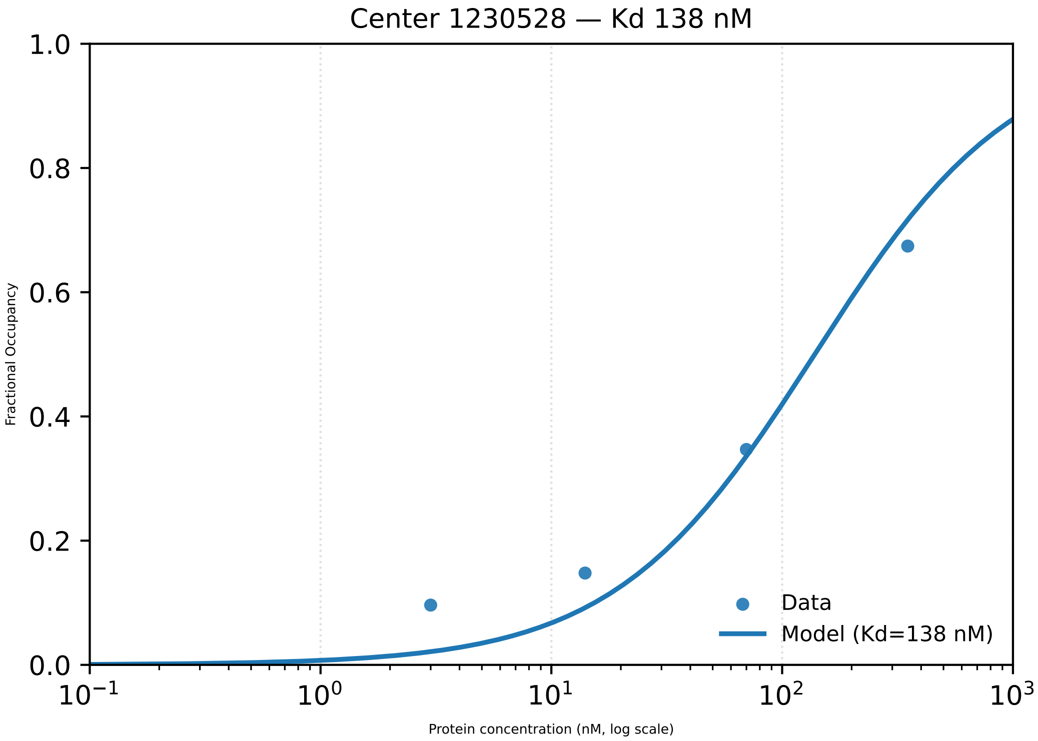

Center 1230528 — Kd 138 nM
Fractional Occupancy
1.0
0.8
0.6
0.4
0.2
0.0
$10^{-1}$
$10^{0}$
$10^{1}$
$10^{2}$
$10^{3}$
Protein concentration (nM, log scale)
Data
Model (Kd=138 nM)

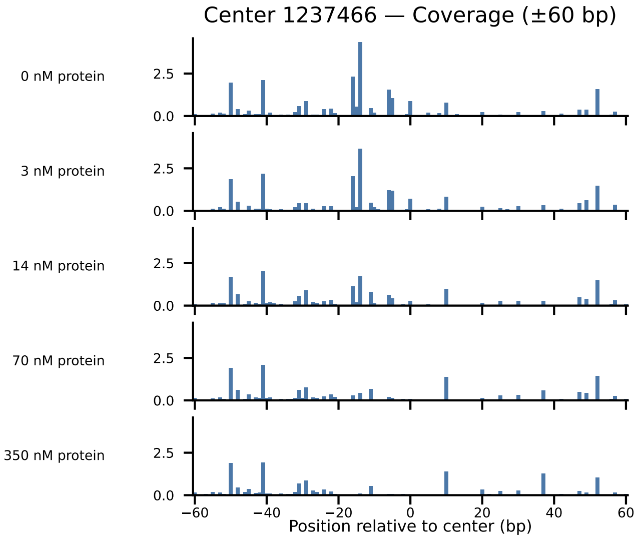

Center 1237466 — Coverage (±60 bp)
0 nM protein
3 nM protein
14 nM protein
70 nM protein
350 nM protein
2.5
0.0
−60
−40
−20
0
20
40
60
Position relative to center (bp)

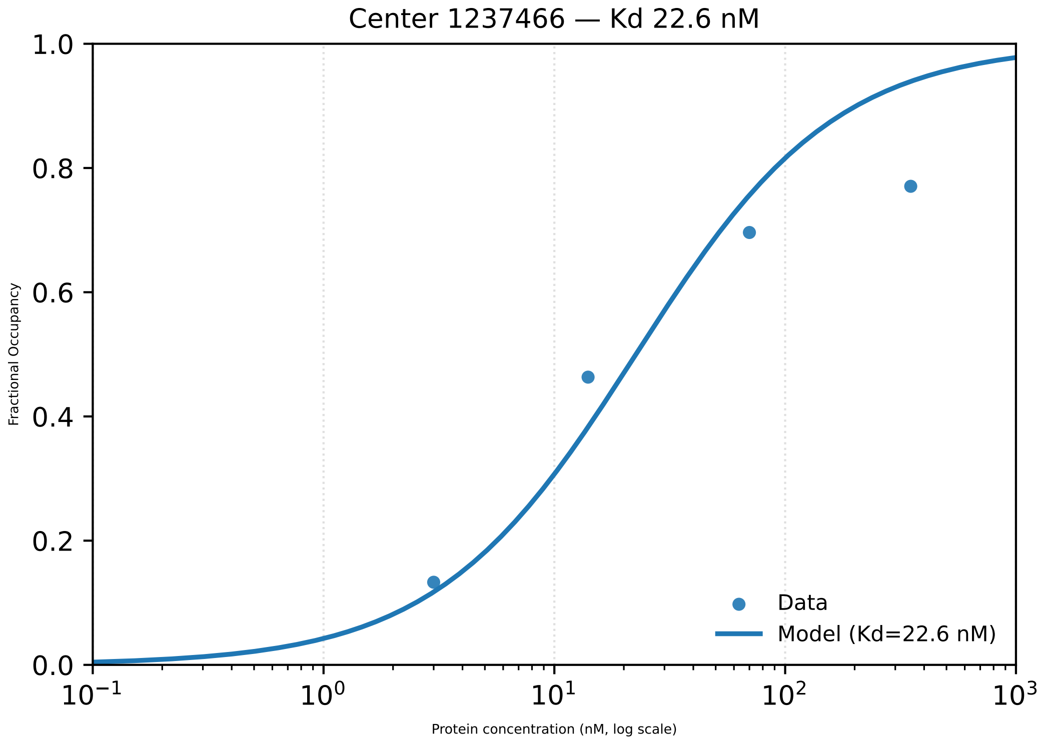
Center 1237466 — Kd 22.6 nM
Fractional Occupancy
Protein concentration (nM, log scale)
Data
Model (Kd=22.6 nM)
0.0
0.2
0.4
0.6
0.8
1.0
$10^{-1}$
$10^{0}$
$10^{1}$
$10^{2}$
$10^{3}$

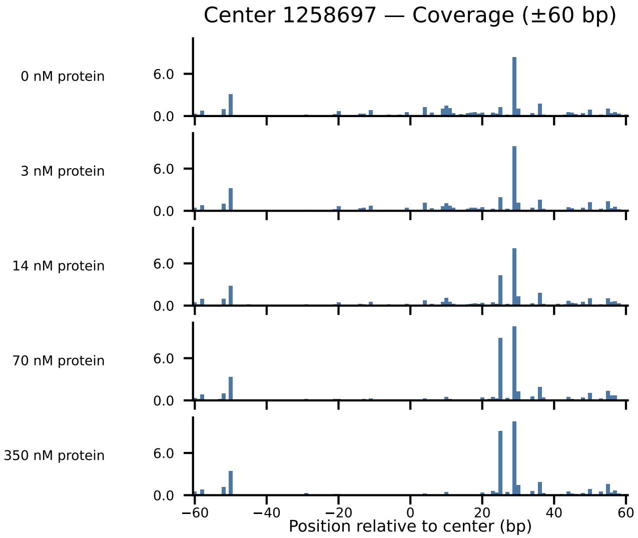

Center 1258697 — Coverage (±60 bp)
0 nM protein
3 nM protein
14 nM protein
70 nM protein
350 nM protein
6.0
0.0
−60
−40
−20
0
20
40
60
Position relative to center (bp)

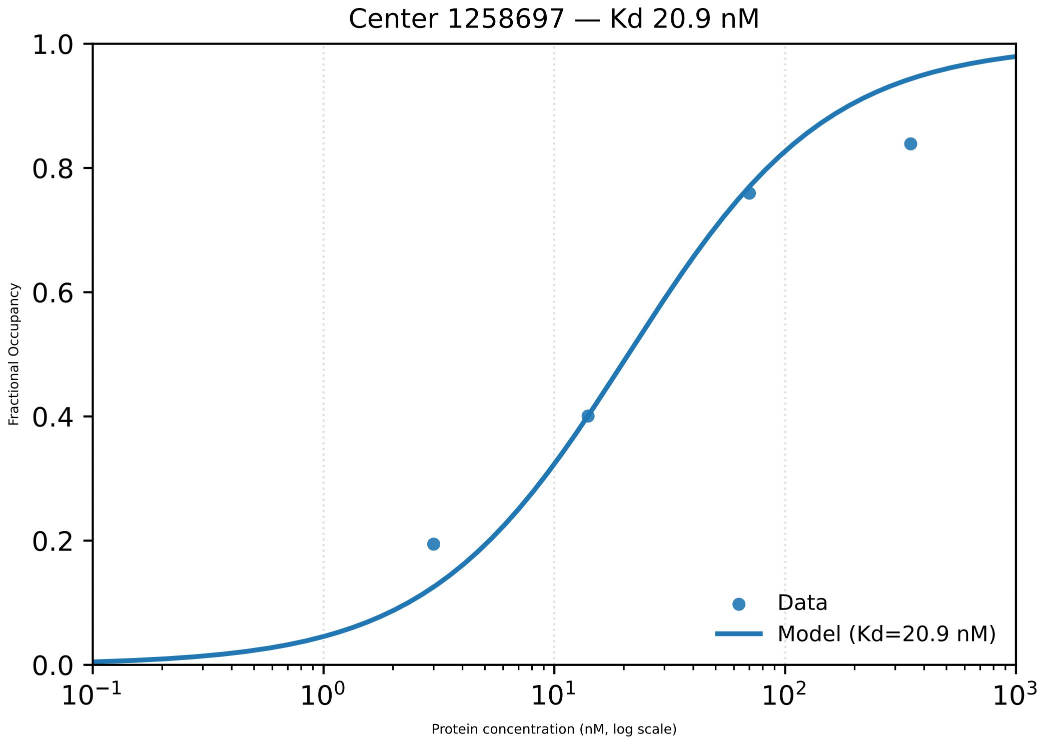

Center 1258697 — Kd 20.9 nM
1.0
0.8
0.6
0.4
0.2
0.0
Fractional Occupancy
$10^{-1}$
$10^{0}$
$10^{1}$
$10^{2}$
$10^{3}$
Protein concentration (nM, log scale)
Data
Model (Kd=20.9 nM)

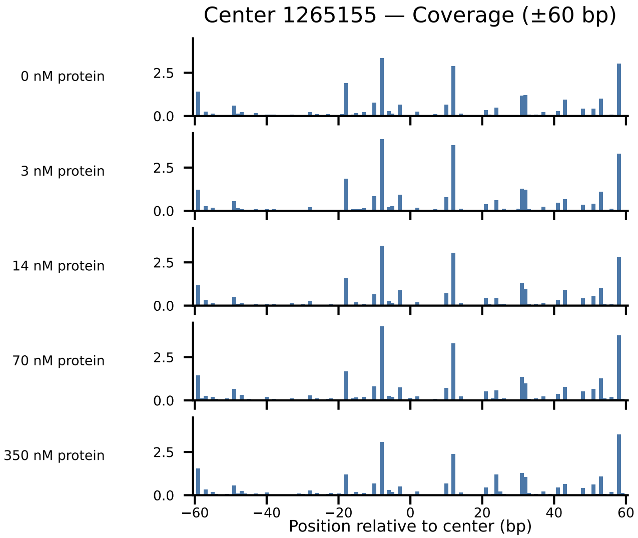

Center 1265155 — Coverage (±60 bp)
0 nM protein
3 nM protein
14 nM protein
70 nM protein
350 nM protein
2.5
0.0
−60
−40
−20
0
20
40
60
Position relative to center (bp)

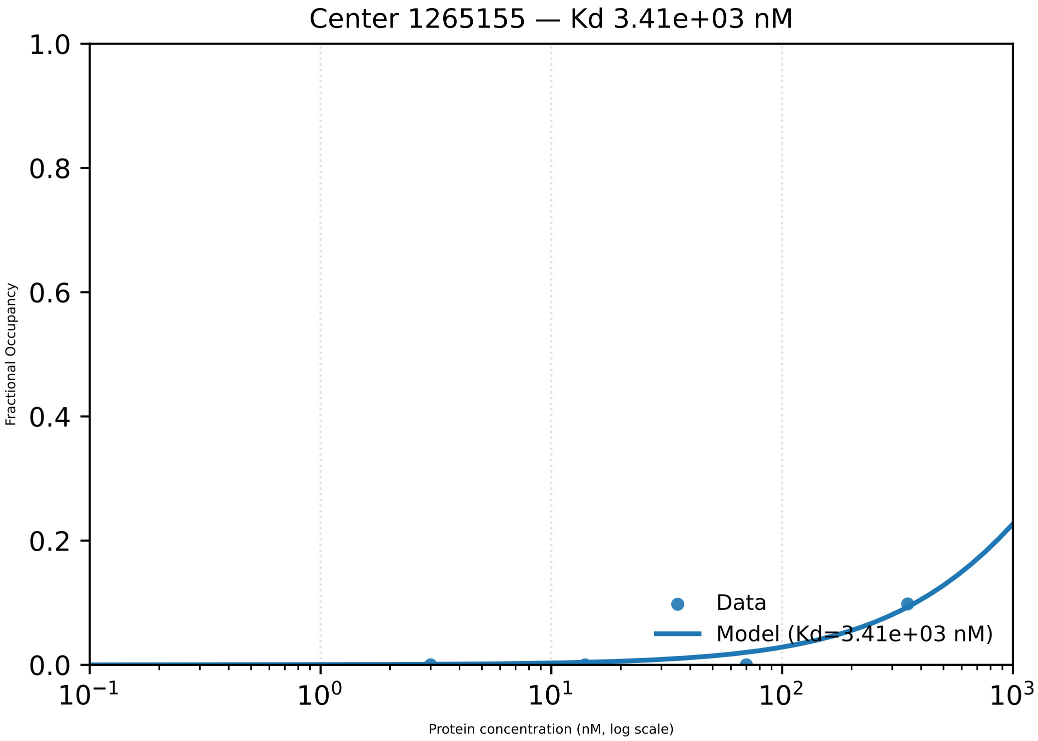
Center 1265155 — Kd 3.41e+03 nM
Fractional Occupancy
1.0
0.8
0.6
0.4
0.2
0.0
$10^{-1}$
$10^{0}$
$10^{1}$
$10^{2}$
$10^{3}$
Protein concentration (nM, log scale)
Data
Model (Kd=3.41e+03 nM)

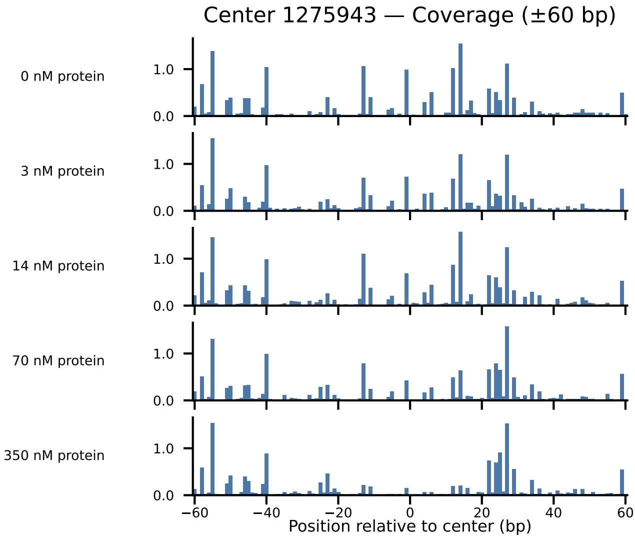
Center 1275943 — Coverage (±60 bp)
0 nM protein
3 nM protein
14 nM protein
70 nM protein
350 nM protein
1.0
0.0
−60
−40
−20
0
20
40
60
Position relative to center (bp)

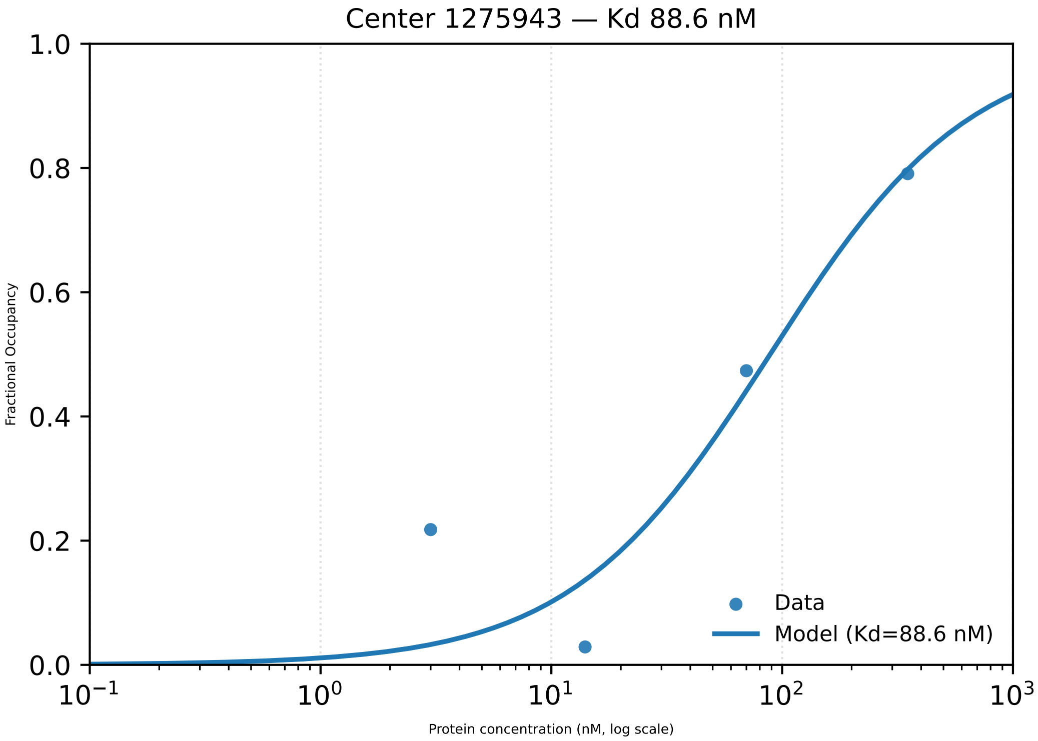

Center 1275943 — Kd 88.6 nM
Fractional Occupancy
1.0
0.8
0.6
0.4
0.2
0.0
$10^{-1}$
$10^{0}$
$10^{1}$
$10^{2}$
$10^{3}$
Protein concentration (nM, log scale)
Data
Model (Kd=88.6 nM)

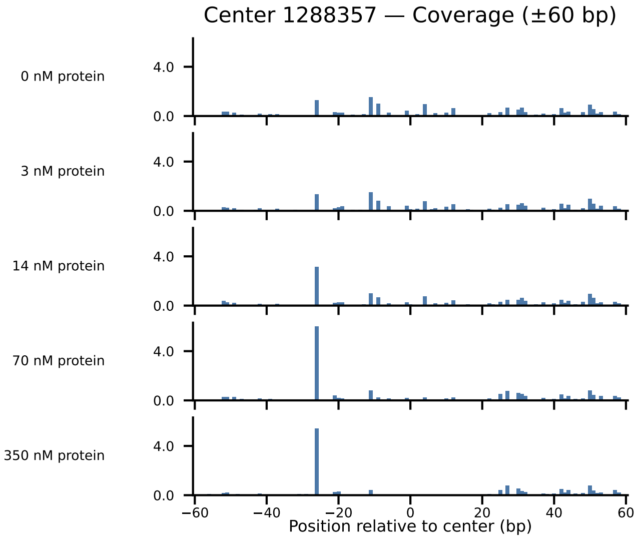

Center 1288357 — Coverage (±60 bp)
0 nM protein
3 nM protein
14 nM protein
70 nM protein
350 nM protein
4.0
0.0
−60
−40
−20
0
20
40
60
Position relative to center (bp)

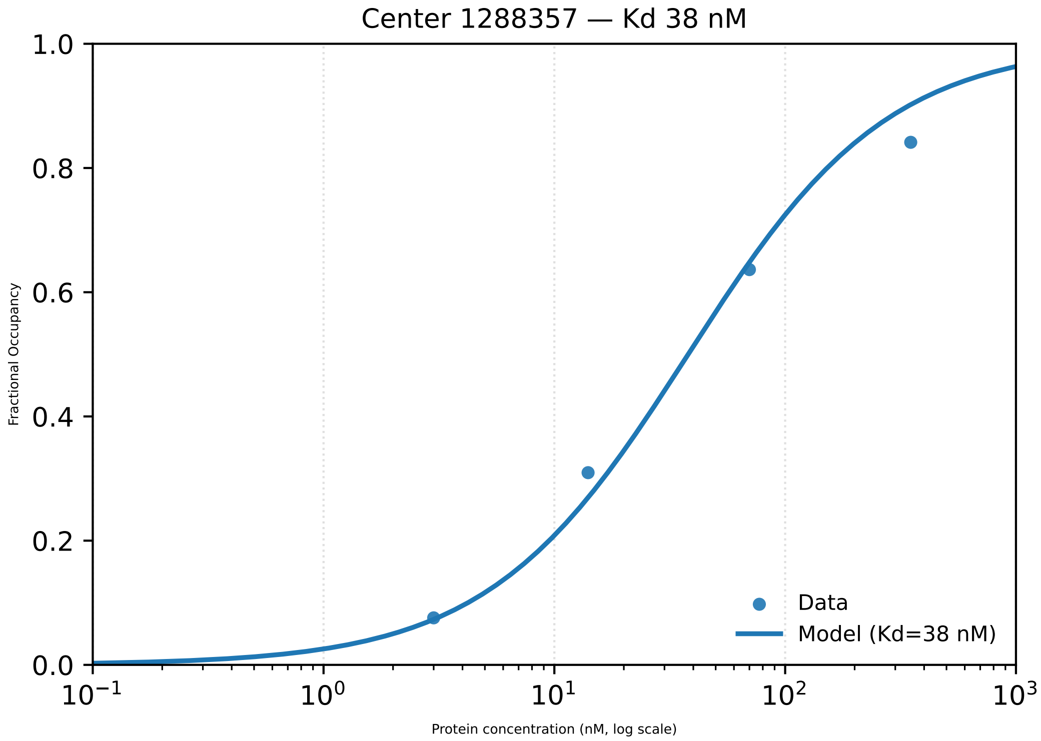
Center 1288357 — Kd 38 nM
1.0
0.8
0.6
0.4
0.2
0.0
Fractional Occupancy
$10^{-1}$
$10^{0}$
$10^{1}$
$10^{2}$
$10^{3}$
Protein concentration (nM, log scale)
Data
Model (Kd=38 nM)

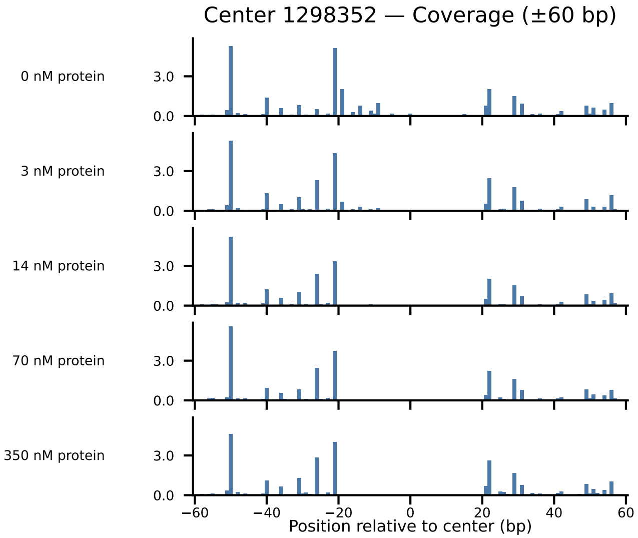

Center 1298352 — Coverage (±60 bp)
0 nM protein
3 nM protein
14 nM protein
70 nM protein
350 nM protein
3.0
0.0
−60
−40
−20
0
20
40
60
Position relative to center (bp)

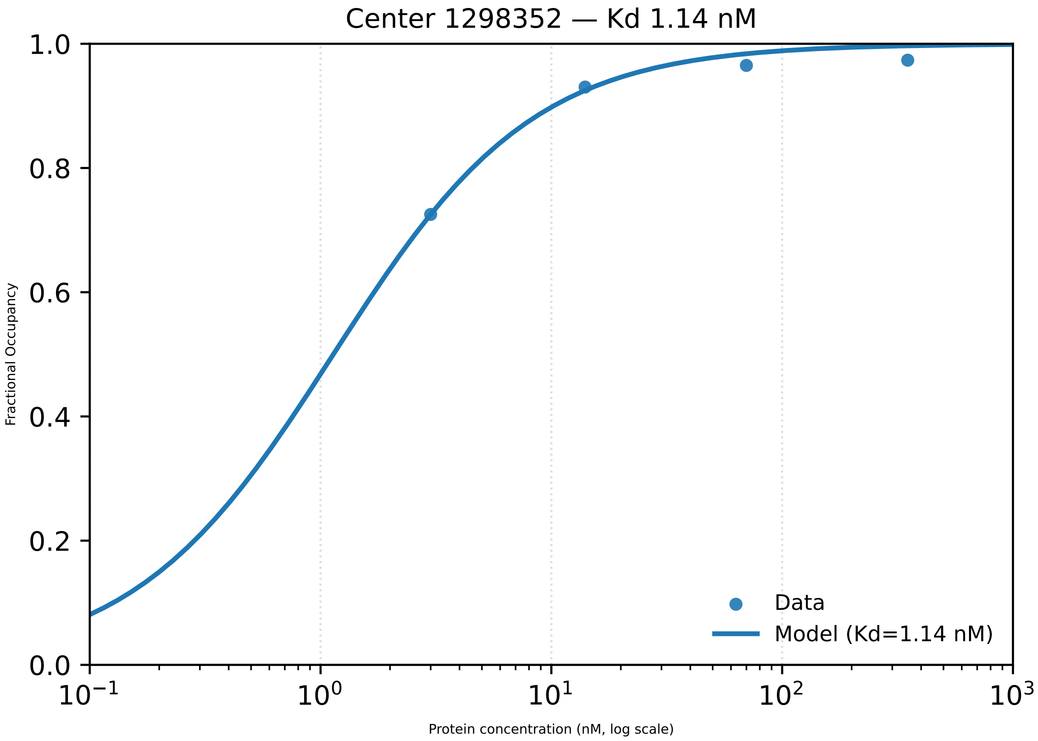

Center 1298352 — Kd 1.14 nM
Fractional Occupancy
Protein concentration (nM, log scale)
Data
Model (Kd=1.14 nM)
0.0
0.2
0.4
0.6
0.8
1.0
$10^{-1}$
$10^{0}$
$10^{1}$
$10^{2}$
$10^{3}$

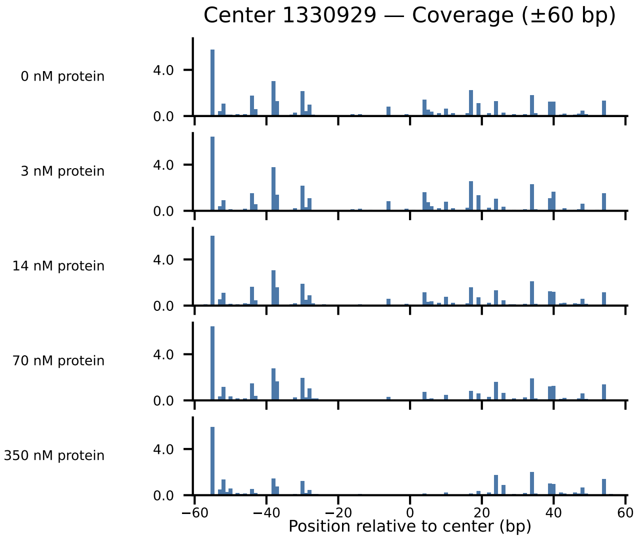

Center 1330929 — Coverage (±60 bp)
0 nM protein
3 nM protein
14 nM protein
70 nM protein
350 nM protein
4.0
0.0
−60
−40
−20
0
20
40
60
Position relative to center (bp)

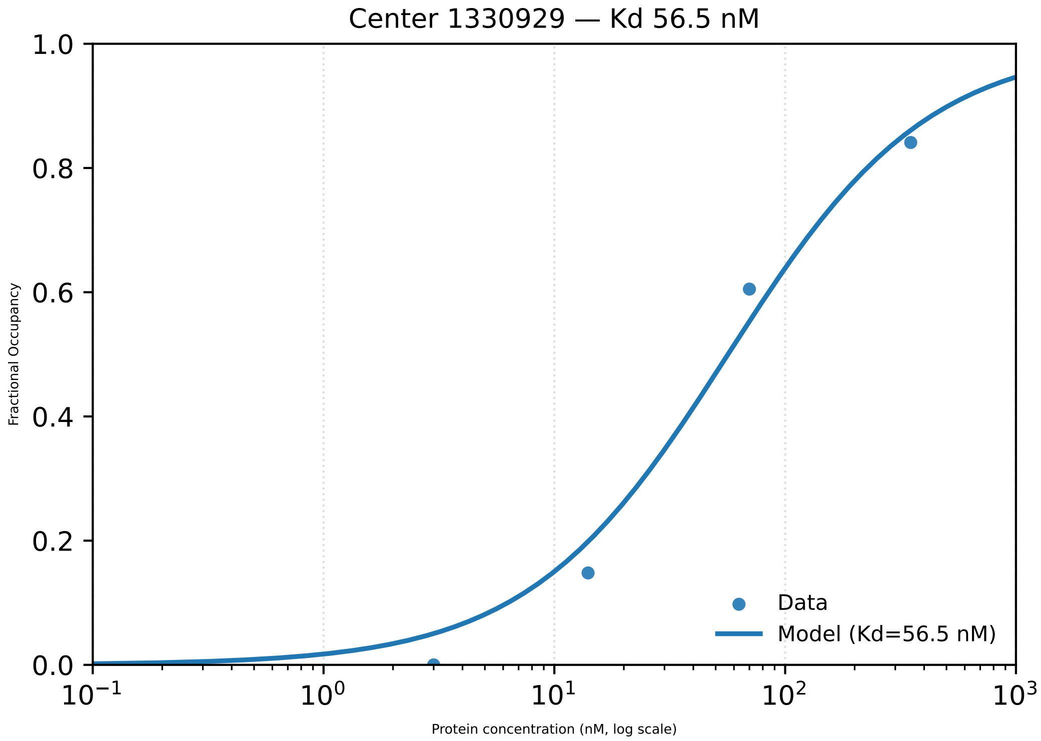

Center 1330929 — Kd 56.5 nM
1.0
0.8
0.6
0.4
0.2
0.0
Fractional Occupancy
$10^{-1}$
$10^{0}$
$10^{1}$
$10^{2}$
$10^{3}$
Protein concentration (nM, log scale)
Data
Model (Kd=56.5 nM)

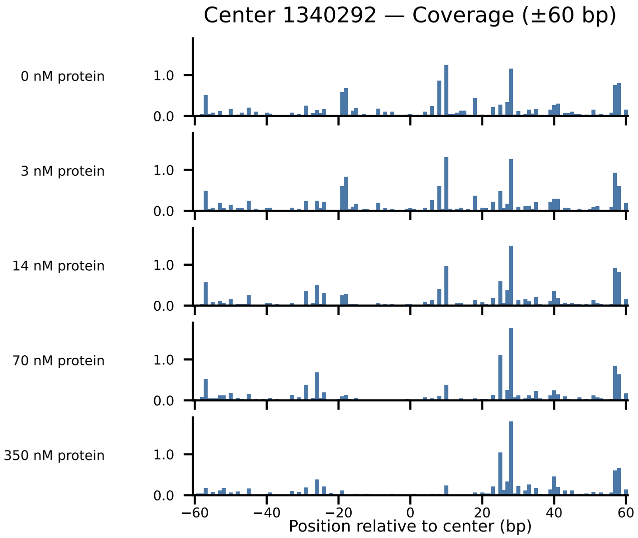

Center 1340292 — Coverage (±60 bp)
0 nM protein
3 nM protein
14 nM protein
70 nM protein
350 nM protein
1.0
0.0
−60
−40
−20
0
20
40
60
Position relative to center (bp)

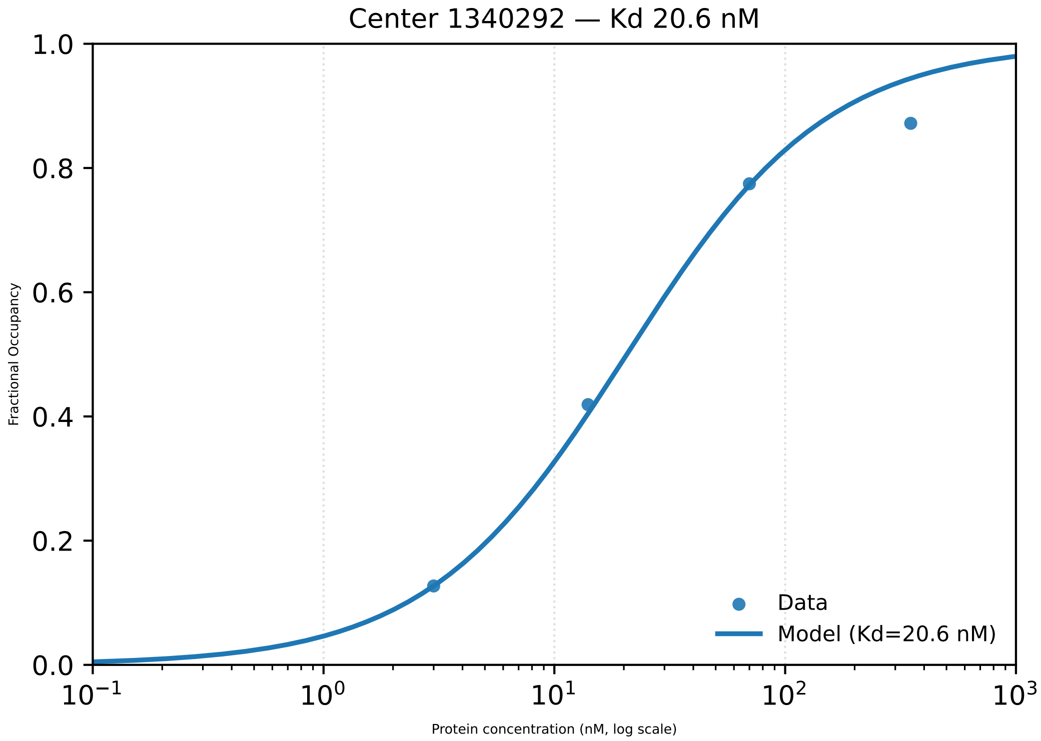

Center 1340292 — Kd 20.6 nM
1.0
0.8
0.6
0.4
0.2
0.0
Fractional Occupancy
$10^{-1}$
$10^{0}$
$10^{1}$
$10^{2}$
$10^{3}$
Protein concentration (nM, log scale)
Data
Model (Kd=20.6 nM)

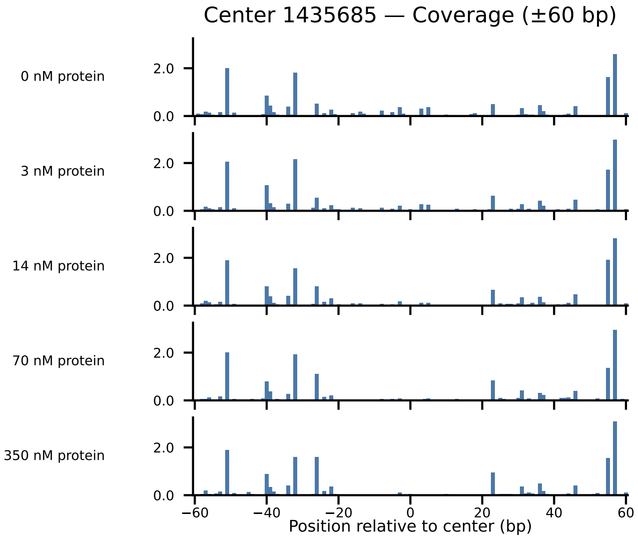

Center 1435685 — Coverage (±60 bp)
0 nM protein
3 nM protein
14 nM protein
70 nM protein
350 nM protein
2.0
0.0
−60
−40
−20
0
20
40
60
Position relative to center (bp)

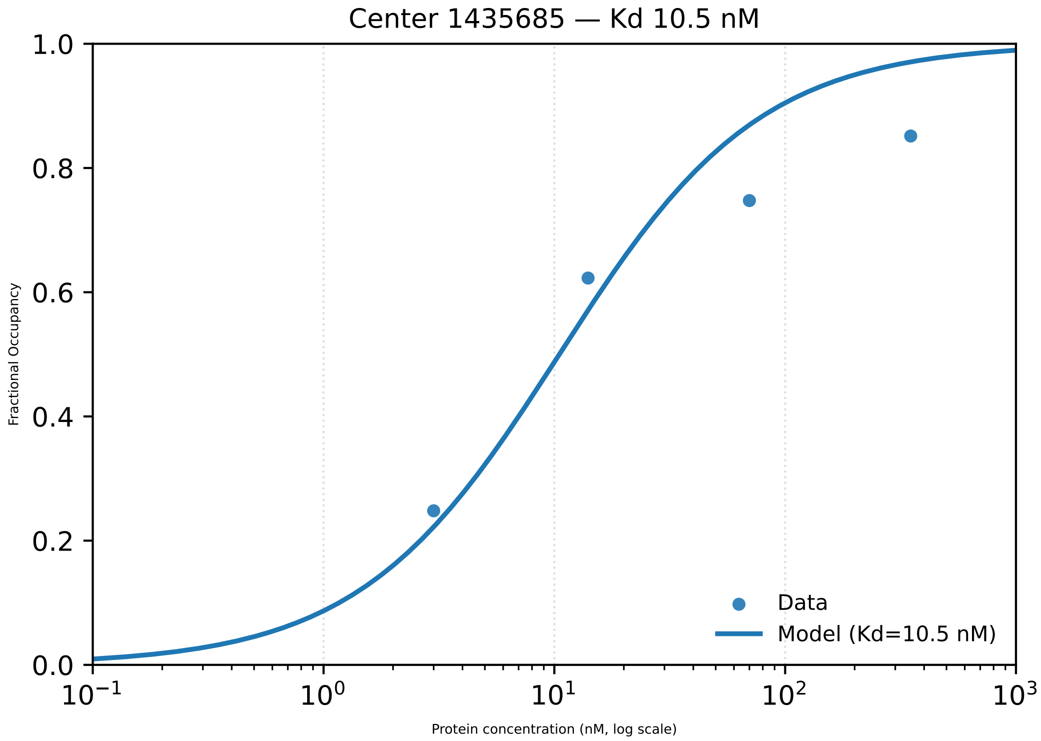
Center 1435685 — Kd 10.5 nM
Fractional Occupancy
1.0
0.8
0.6
0.4
0.2
0.0
$10^{-1}$
$10^{0}$
$10^{1}$
$10^{2}$
$10^{3}$
Protein concentration (nM, log scale)
Data
Model (Kd=10.5 nM)

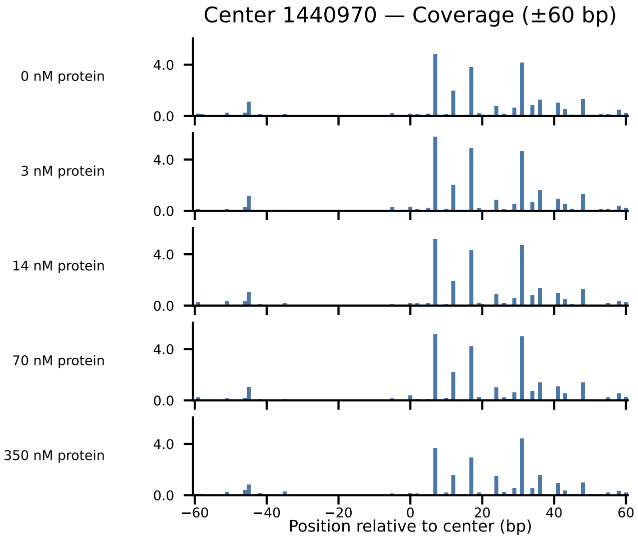

Center 1440970 — Coverage (±60 bp)
0 nM protein
3 nM protein
14 nM protein
70 nM protein
350 nM protein
4.0
0.0
−60
−40
−20
0
20
40
60
Position relative to center (bp)

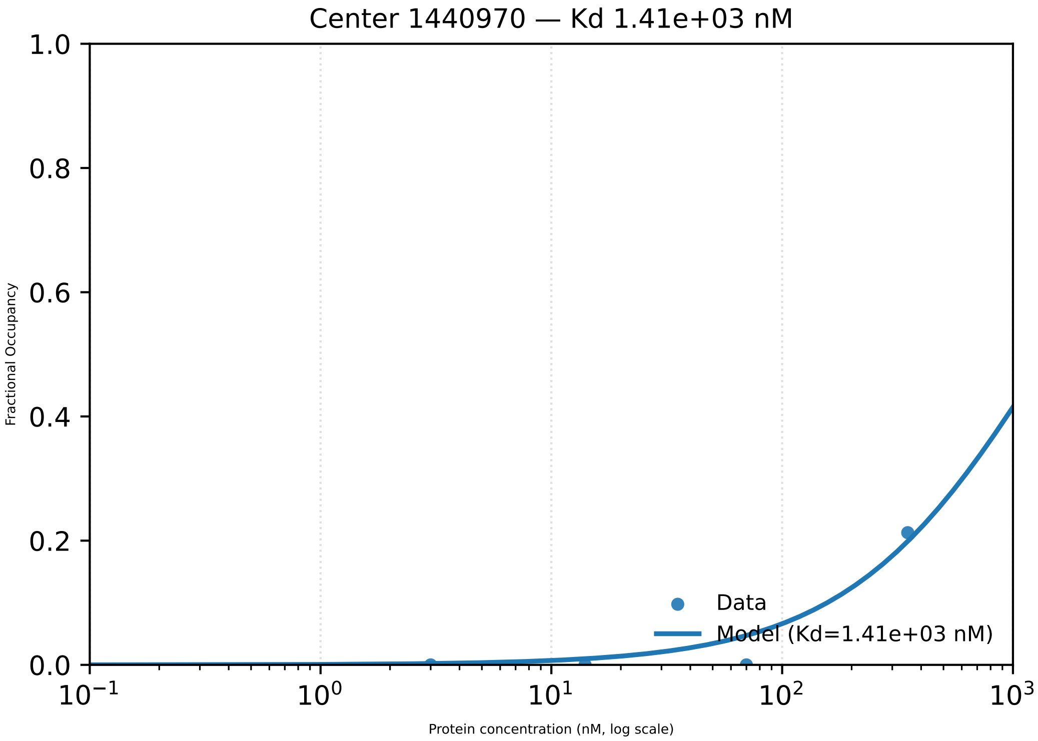

Center 1440970 — Kd 1.41e+03 nM
Fractional Occupancy
Protein concentration (nM, log scale)
Data
Model (Kd=1.41e+03 nM)
1.0
0.8
0.6
0.4
0.2
0.0
$10^{-1}$
$10^{0}$
$10^{1}$
$10^{2}$
$10^{3}$

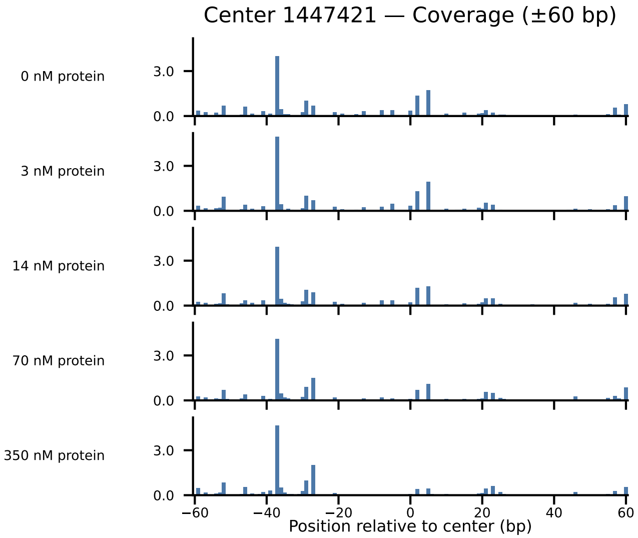

Center 1447421 — Coverage (±60 bp)
0 nM protein
3 nM protein
14 nM protein
70 nM protein
350 nM protein
3.0
0.0
−60
−40
−20
0
20
40
60
Position relative to center (bp)

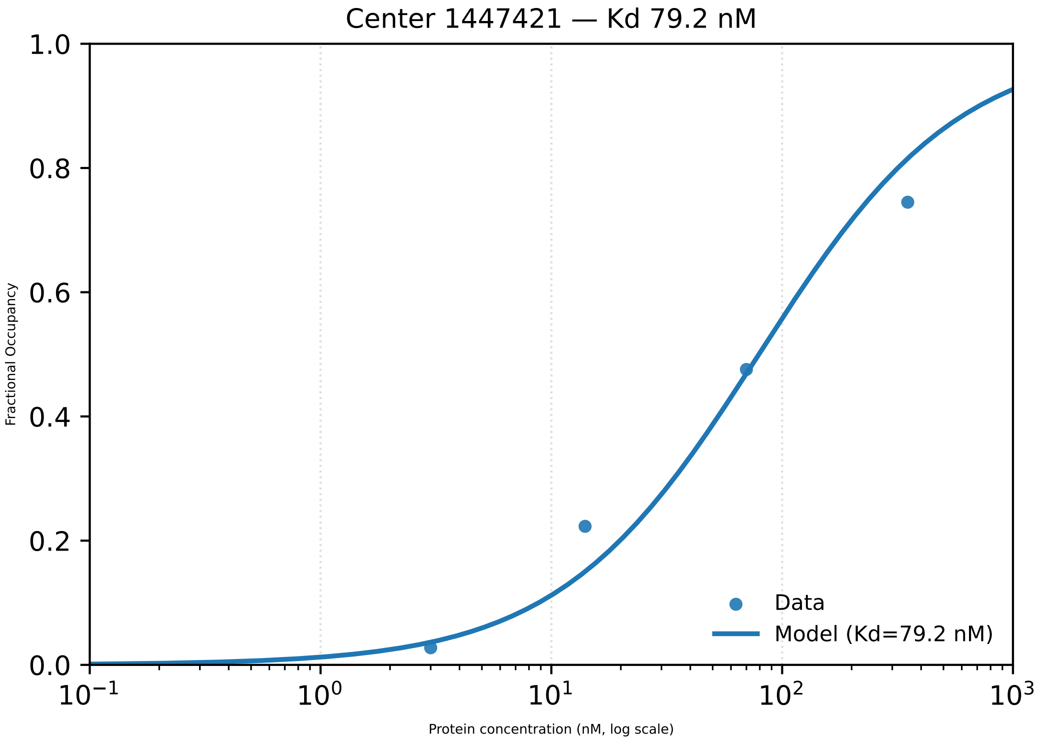

Center 1447421 — Kd 79.2 nM
Fractional Occupancy
Protein concentration (nM, log scale)
1.0
0.8
0.6
0.4
0.2
0.0
$10^{-1}$
$10^{0}$
$10^{1}$
$10^{2}$
$10^{3}$
Data
Model (Kd=79.2 nM)

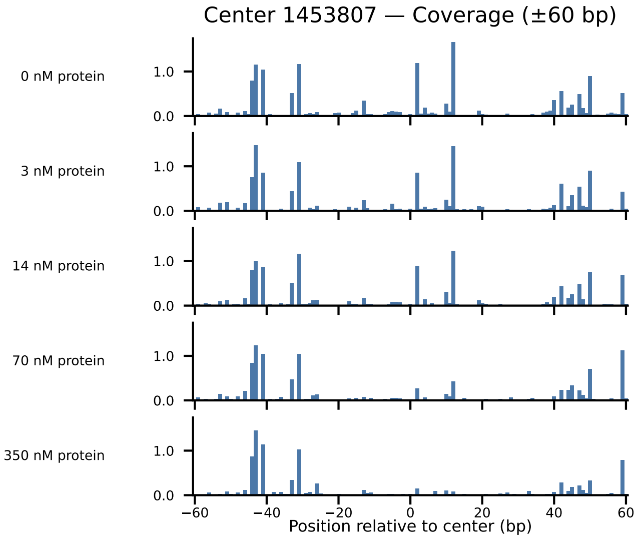

Center 1453807 — Coverage (±60 bp)
0 nM protein
3 nM protein
14 nM protein
70 nM protein
350 nM protein
1.0
0.0
−60
−40
−20
0
20
40
60
Position relative to center (bp)

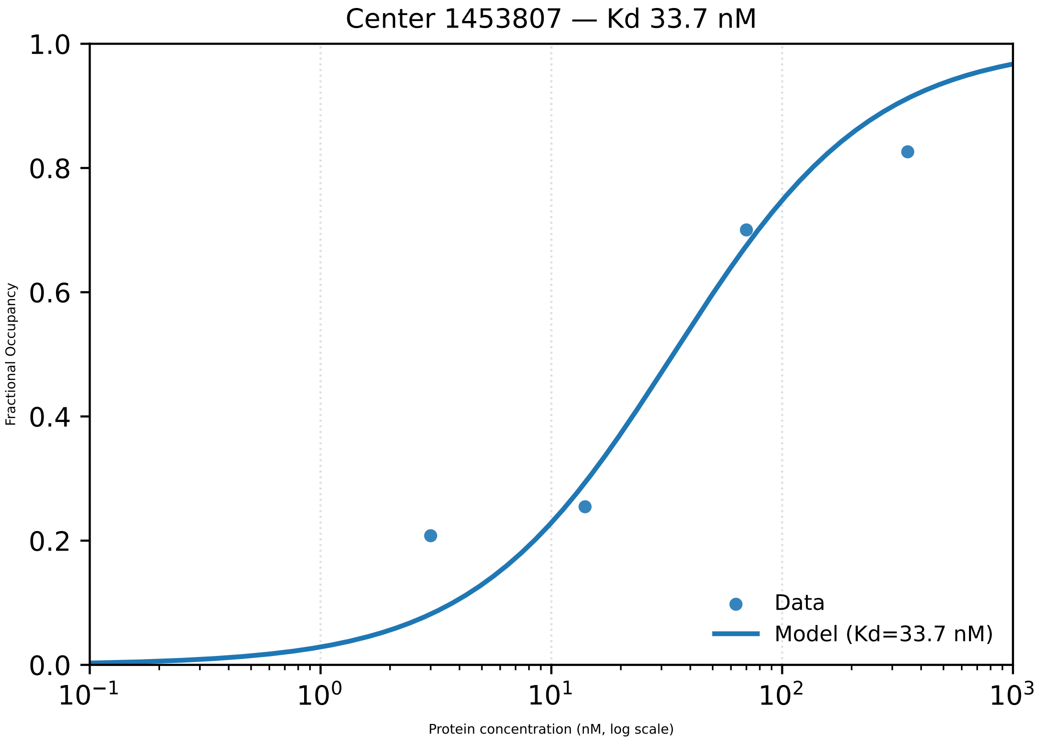

Center 1453807 — Kd 33.7 nM
Fractional Occupancy
1.0
0.8
0.6
0.4
0.2
0.0
$10^{-1}$
$10^{0}$
$10^{1}$
$10^{2}$
$10^{3}$
Protein concentration (nM, log scale)
Data
Model (Kd=33.7 nM)

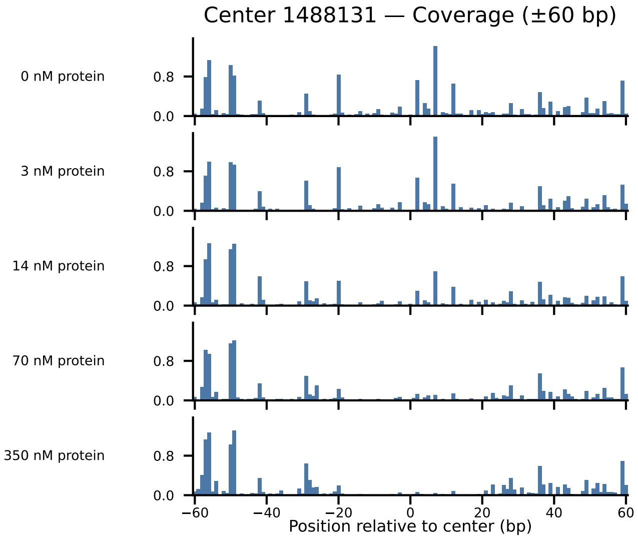
Center 1488131 — Coverage (±60 bp)
0 nM protein
3 nM protein
14 nM protein
70 nM protein
350 nM protein
0.8
0.0
−60
−40
−20
0
20
40
60
Position relative to center (bp)

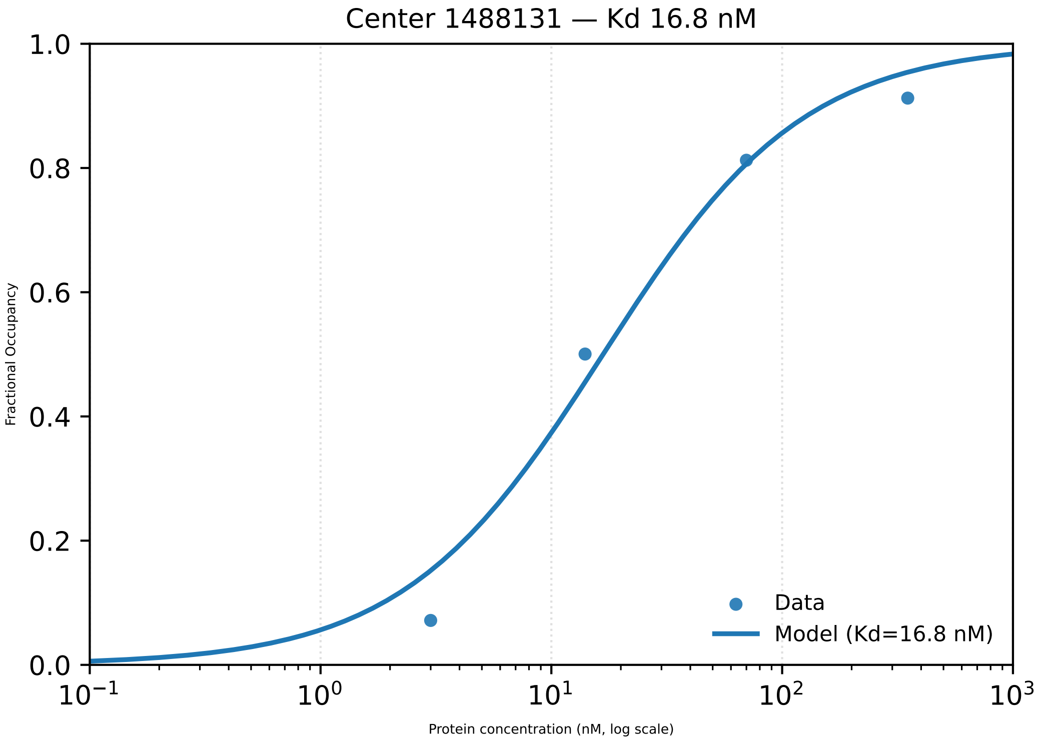

Center 1488131 — Kd 16.8 nM
Fractional Occupancy
Protein concentration (nM, log scale)
0.0
0.2
0.4
0.6
0.8
1.0
$10^{-1}$
$10^{0}$
$10^{1}$
$10^{2}$
$10^{3}$
Data
Model (Kd=16.8 nM)

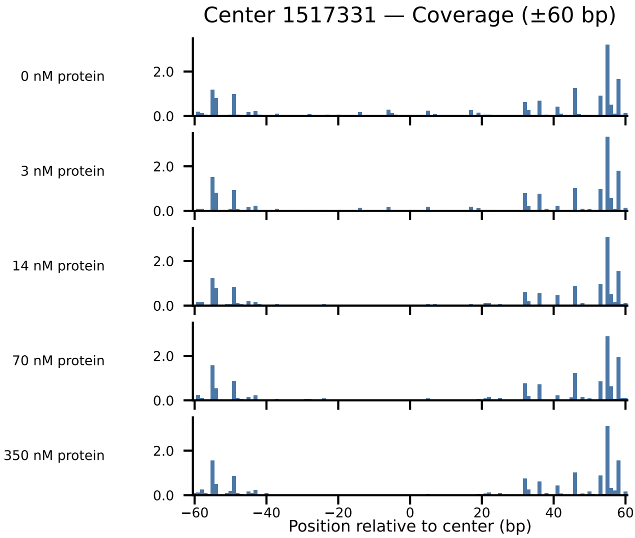

Center 1517331 — Coverage (±60 bp)
0 nM protein
3 nM protein
14 nM protein
70 nM protein
350 nM protein
2.0
0.0
−60
−40
−20
0
20
40
60
Position relative to center (bp)

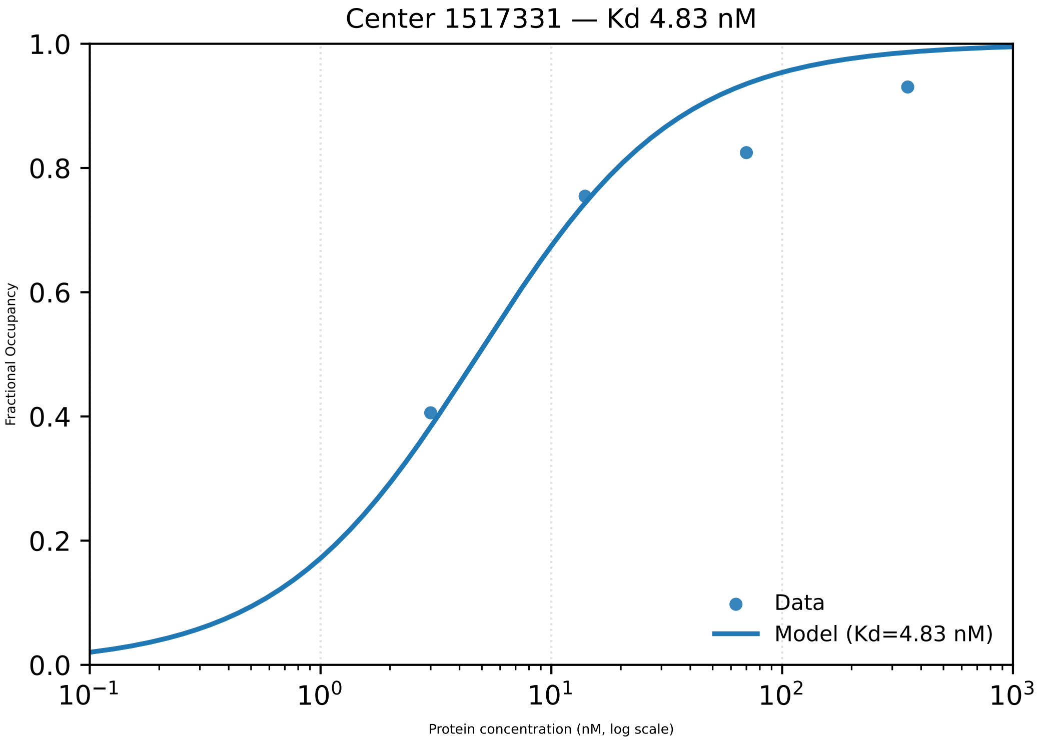

Center 1517331 — Kd 4.83 nM
1.0
0.8
0.6
0.4
0.2
0.0
Fractional Occupancy
$10^{-1}$
$10^{0}$
$10^{1}$
$10^{2}$
$10^{3}$
Protein concentration (nM, log scale)
Data
Model (Kd=4.83 nM)

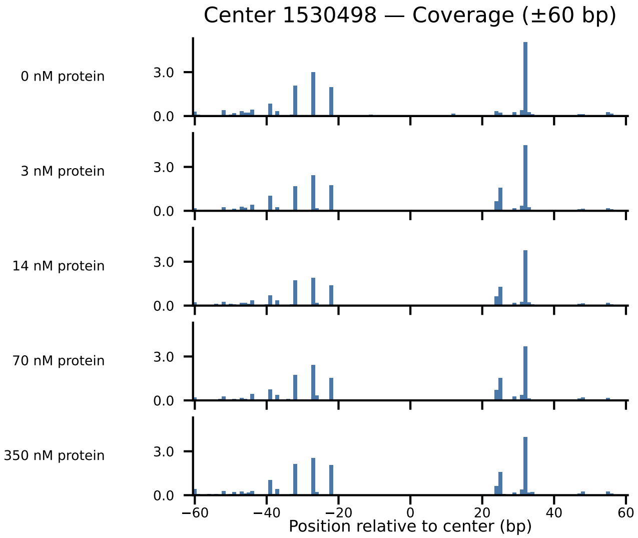

Center 1530498 — Coverage (±60 bp)
0 nM protein
3 nM protein
14 nM protein
70 nM protein
350 nM protein
3.0
0.0
−60
−40
−20
0
20
40
60
Position relative to center (bp)

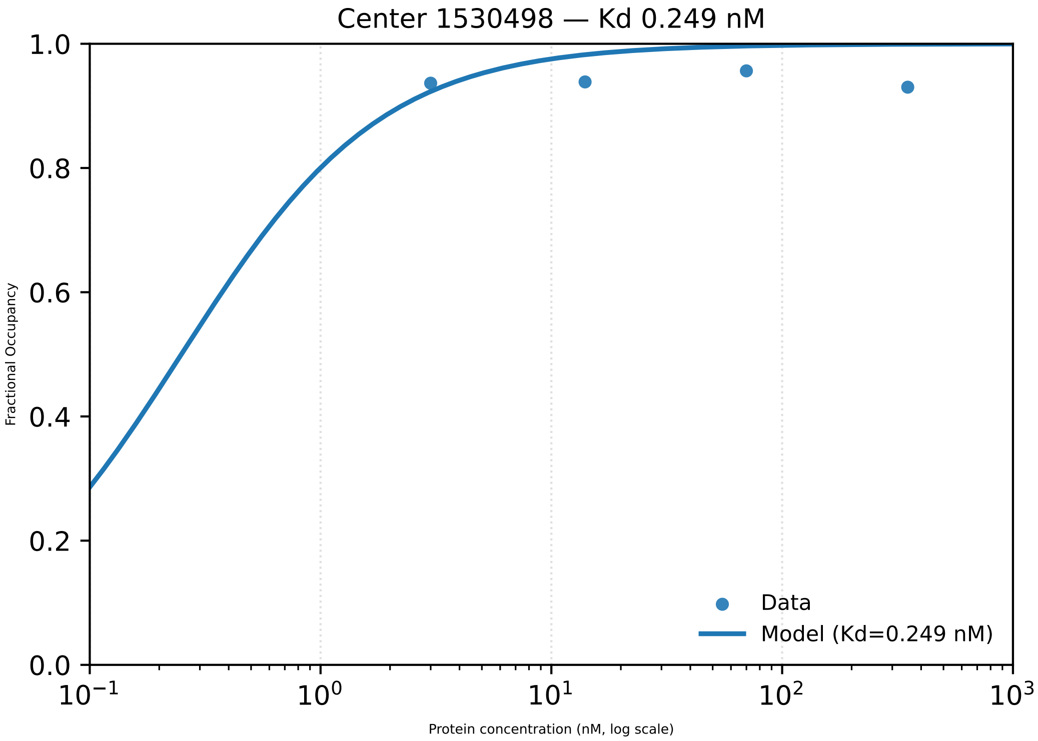

Center 1530498 — Kd 0.249 nM
1.0
0.8
0.6
0.4
0.2
0.0
Fractional Occupancy
$10^{-1}$
$10^{0}$
$10^{1}$
$10^{2}$
$10^{3}$
Protein concentration (nM, log scale)
Data
Model (Kd=0.249 nM)

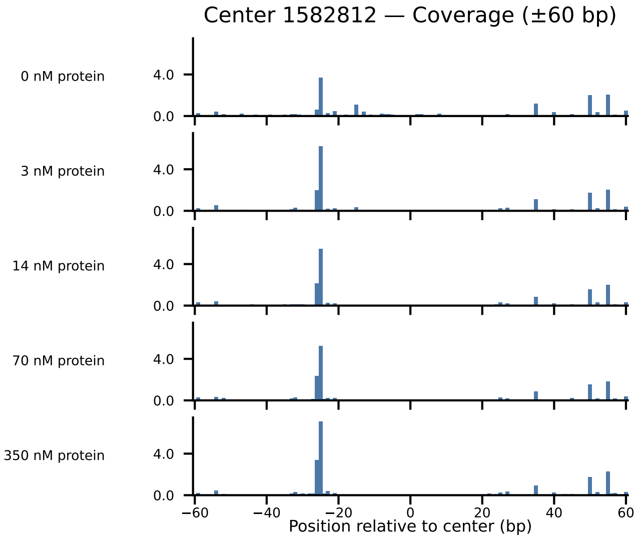
Center 1582812 — Coverage (±60 bp)
0 nM protein
3 nM protein
14 nM protein
70 nM protein
350 nM protein
4.0
0.0
−60
−40
−20
0
20
40
60
Position relative to center (bp)

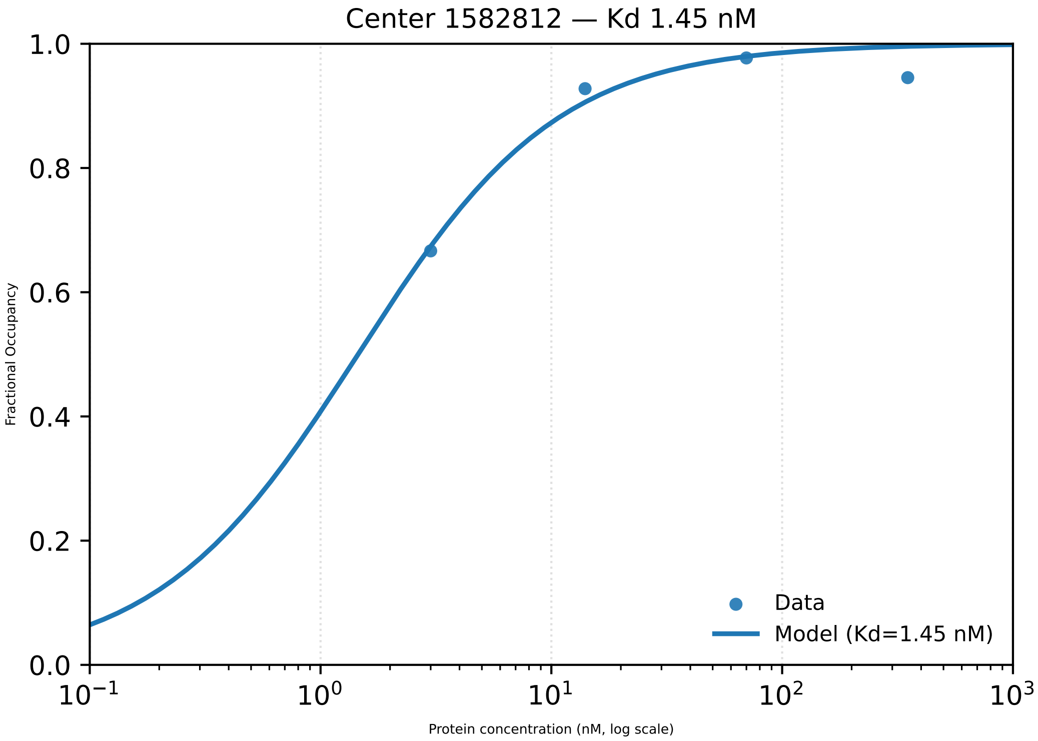
Center 1582812 — Kd 1.45 nM
Fractional Occupancy
Protein concentration (nM, log scale)
Data
Model (Kd=1.45 nM)
0.0
0.2
0.4
0.6
0.8
1.0
$10^{-1}$
$10^{0}$
$10^{1}$
$10^{2}$
$10^{3}$

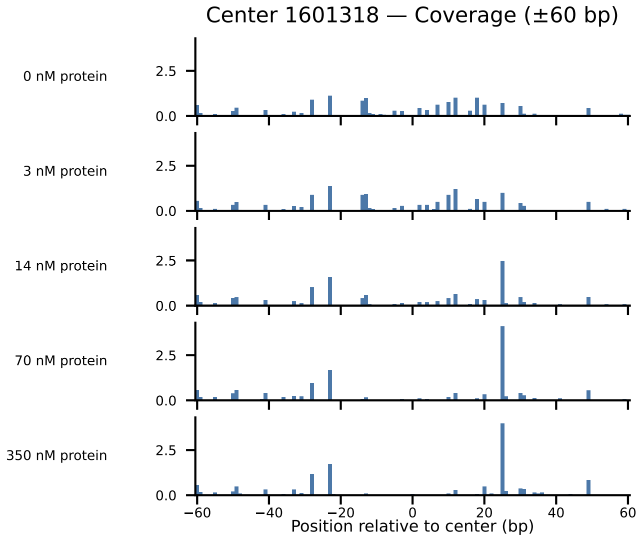

Center 1601318 — Coverage (±60 bp)
0 nM protein
3 nM protein
14 nM protein
70 nM protein
350 nM protein
2.5
0.0
−60
−40
−20
0
20
40
60
Position relative to center (bp)

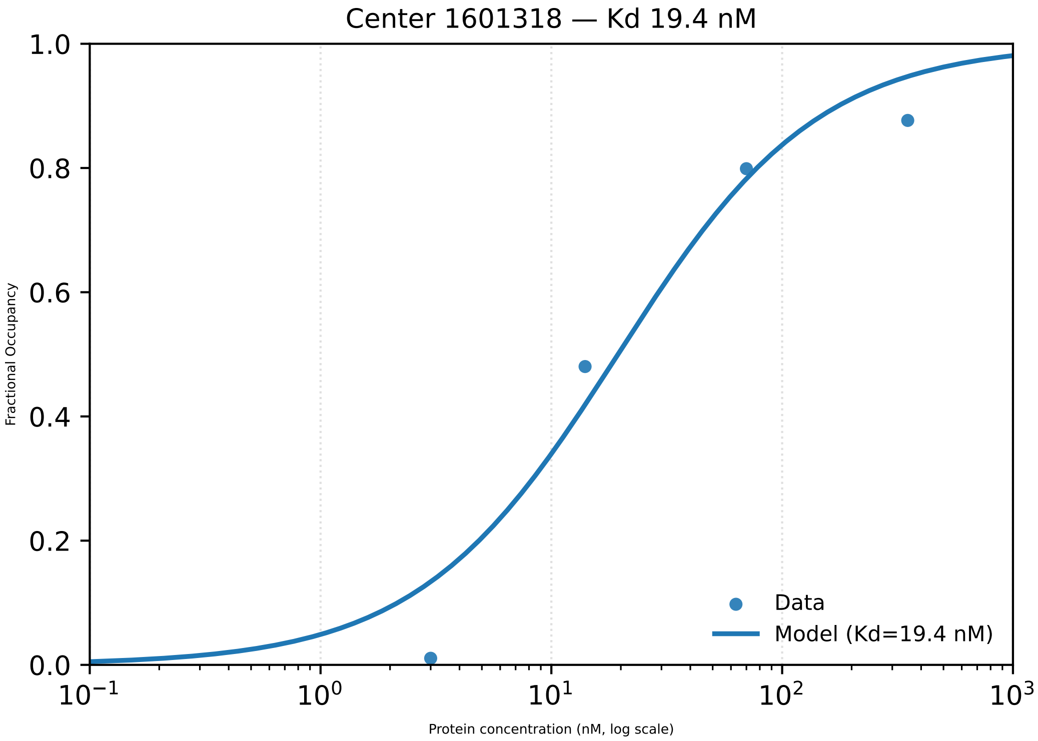

Center 1601318 — Kd 19.4 nM
1.0
0.8
0.6
0.4
0.2
0.0
Fractional Occupancy
Data
Model (Kd=19.4 nM)
$10^{-1}$
$10^{0}$
$10^{1}$
$10^{2}$
$10^{3}$
Protein concentration (nM, log scale)

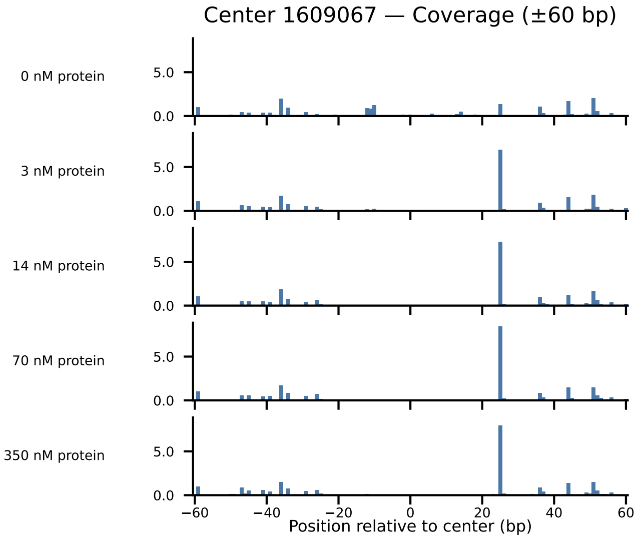

Center 1609067 — Coverage (±60 bp)
0 nM protein
3 nM protein
14 nM protein
70 nM protein
350 nM protein
5.0
0.0
−60
−40
−20
0
20
40
60
Position relative to center (bp)

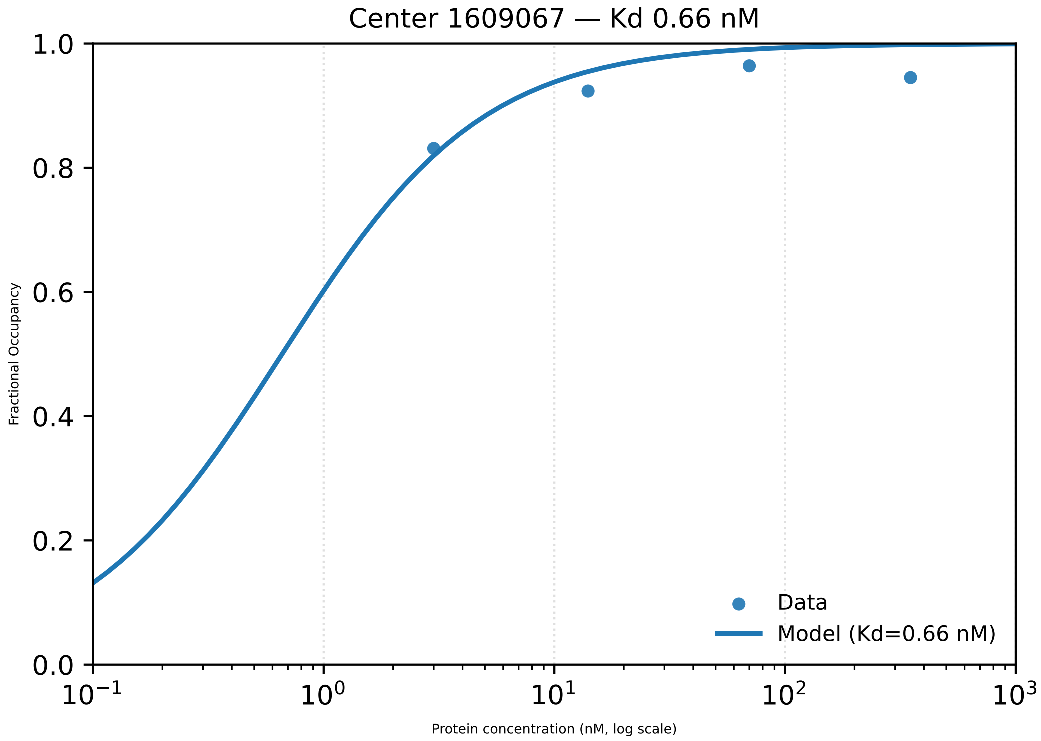
Center 1609067 — Kd 0.66 nM
1.0
0.8
0.6
0.4
0.2
0.0
Fractional Occupancy
$10^{-1}$
$10^{0}$
$10^{1}$
$10^{2}$
$10^{3}$
Protein concentration (nM, log scale)
Data
Model (Kd=0.66 nM)

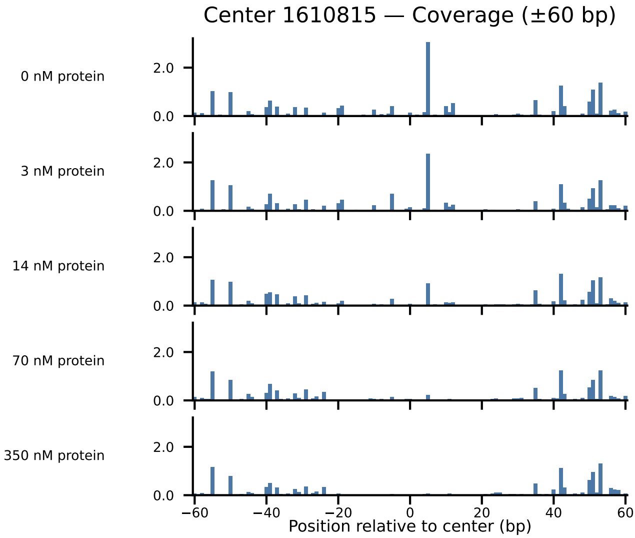

Center 1610815 — Coverage (±60 bp)
0 nM protein
3 nM protein
14 nM protein
70 nM protein
350 nM protein
2.0
0.0
−60
−40
−20
0
20
40
60
Position relative to center (bp)

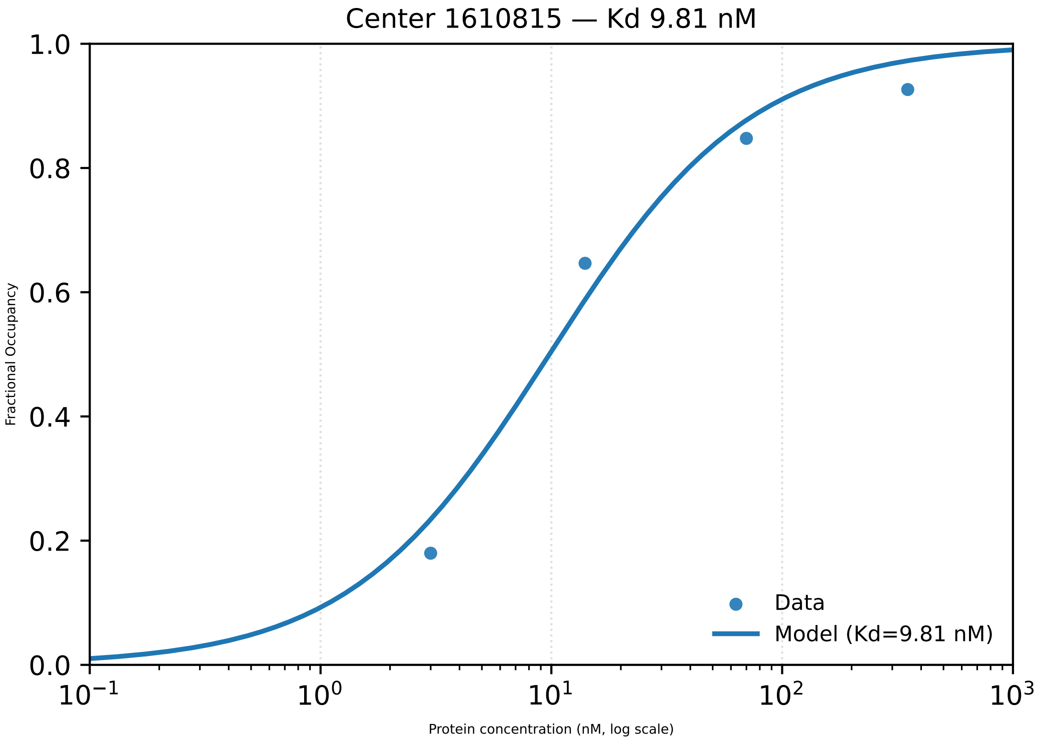

Center 1610815 — Kd 9.81 nM
1.0
0.8
0.6
0.4
0.2
0.0
Fractional Occupancy
$10^{-1}$
$10^{0}$
$10^{1}$
$10^{2}$
$10^{3}$
Protein concentration (nM, log scale)
Data
Model (Kd=9.81 nM)

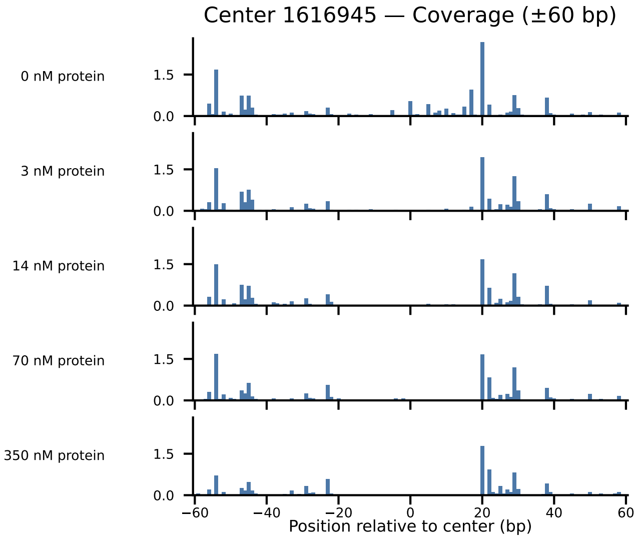

Center 1616945 — Coverage (±60 bp)
0 nM protein
3 nM protein
14 nM protein
70 nM protein
350 nM protein
1.5
0.0
−60
−40
−20
0
20
40
60
Position relative to center (bp)

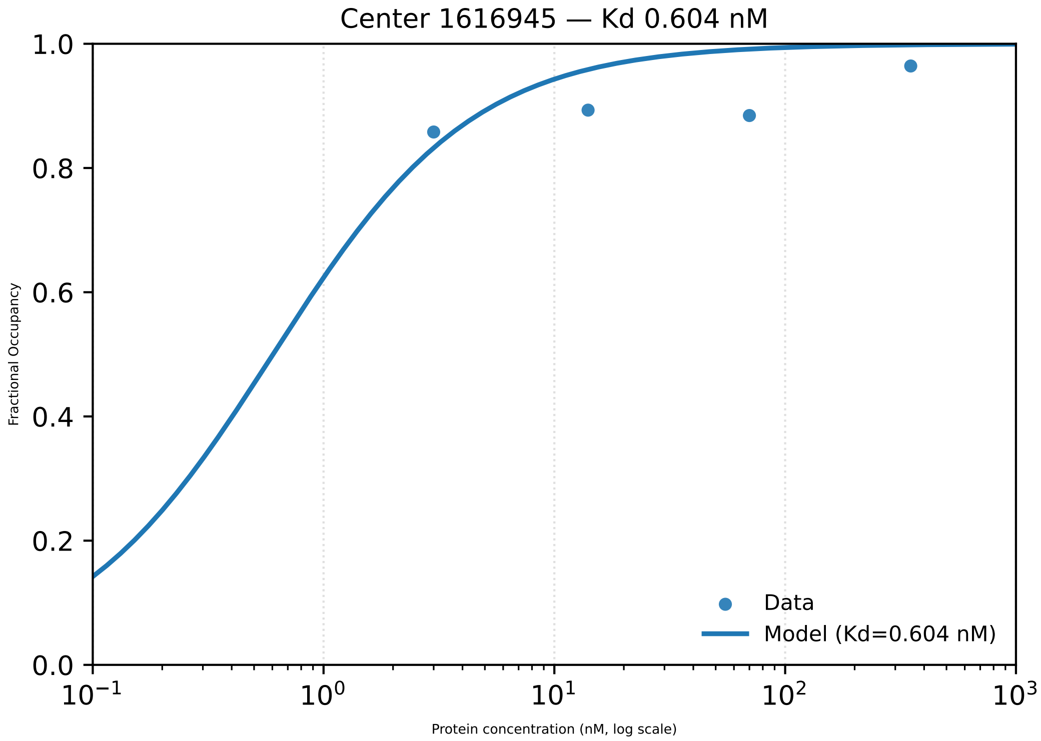

Center 1616945 — Kd 0.604 nM
1.0
0.8
0.6
0.4
0.2
0.0
Fractional Occupancy
$10^{-1}$
$10^{0}$
$10^{1}$
$10^{2}$
$10^{3}$
Protein concentration (nM, log scale)
Data
Model (Kd=0.604 nM)

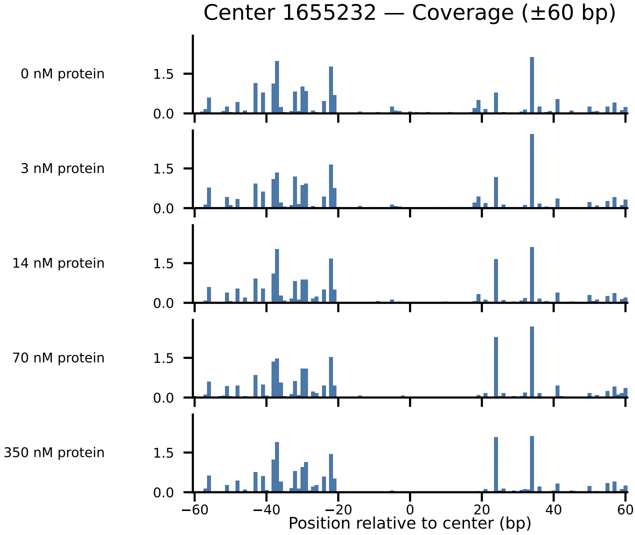
Center 1655232 — Coverage (±60 bp)
0 nM protein
3 nM protein
14 nM protein
70 nM protein
350 nM protein
1.5
0.0
−60
−40
−20
0
20
40
60
Position relative to center (bp)

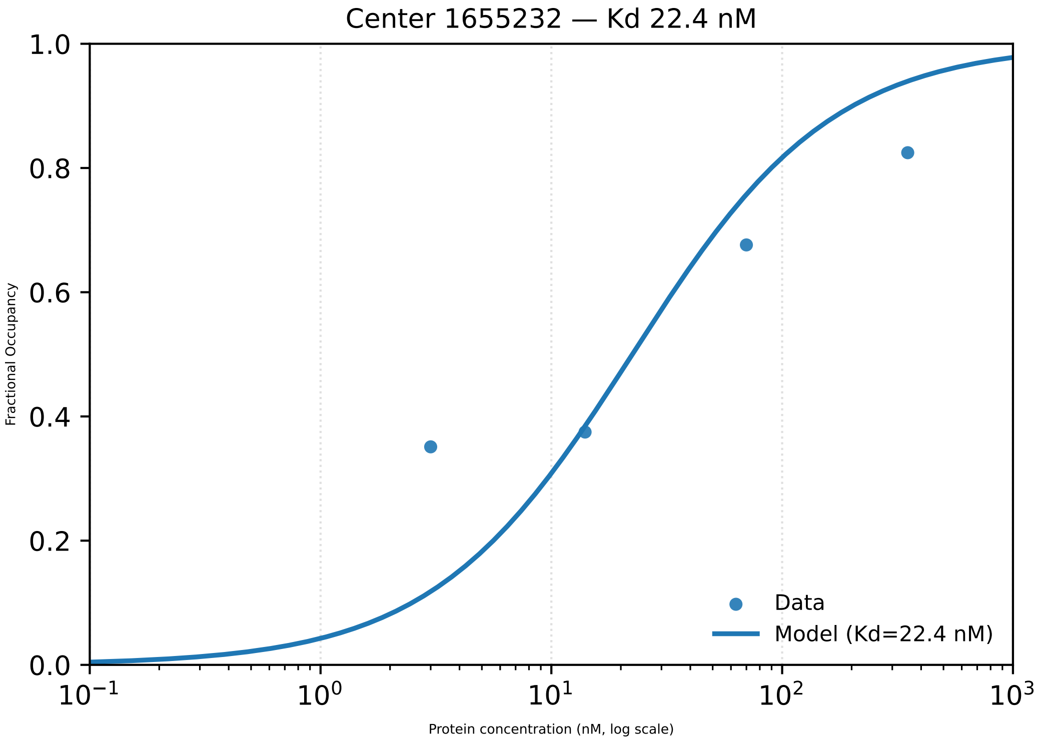

Center 1655232 — Kd 22.4 nM
1.0
0.8
0.6
0.4
0.2
0.0
Fractional Occupancy
$10^{-1}$
$10^{0}$
$10^{1}$
$10^{2}$
$10^{3}$
Protein concentration (nM, log scale)
Data
Model (Kd=22.4 nM)

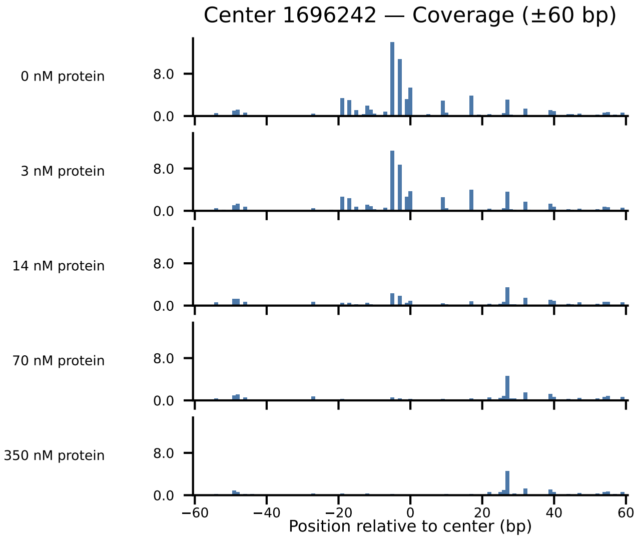

Center 1696242 — Coverage (±60 bp)
0 nM protein
3 nM protein
14 nM protein
70 nM protein
350 nM protein
8.0
0.0
−60
−40
−20
0
20
40
60
Position relative to center (bp)

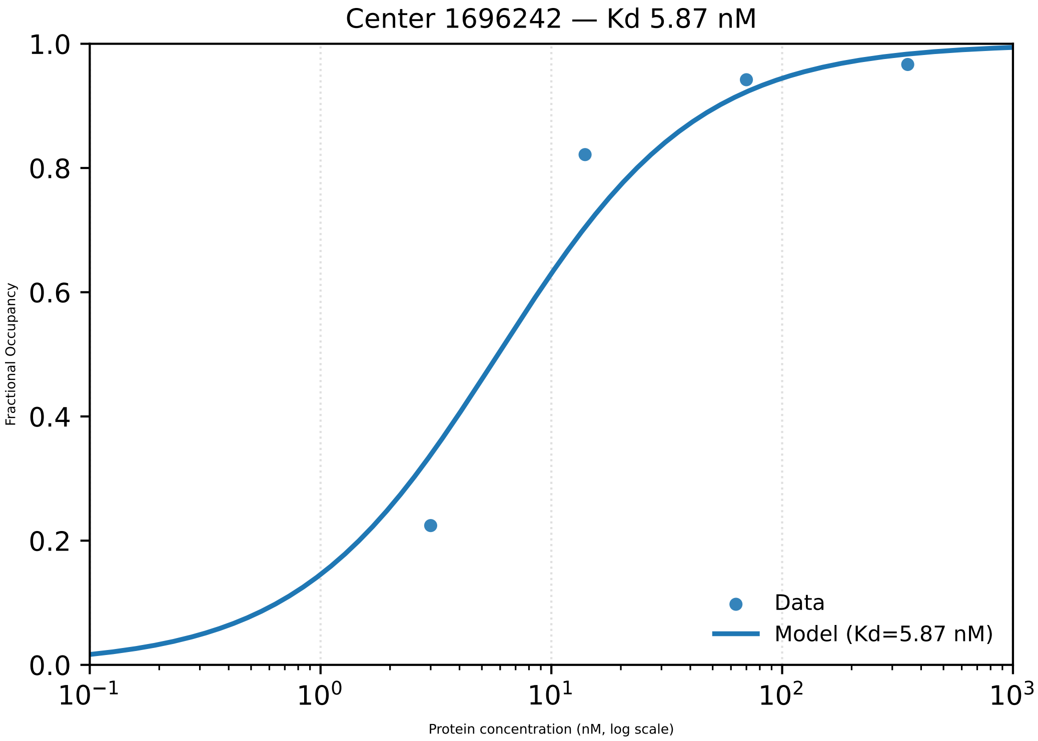
Center 1696242 — Kd 5.87 nM
1.0
0.8
0.6
0.4
0.2
0.0
Fractional Occupancy
$10^{-1}$
$10^{0}$
$10^{1}$
$10^{2}$
$10^{3}$
Protein concentration (nM, log scale)
Data
Model (Kd=5.87 nM)

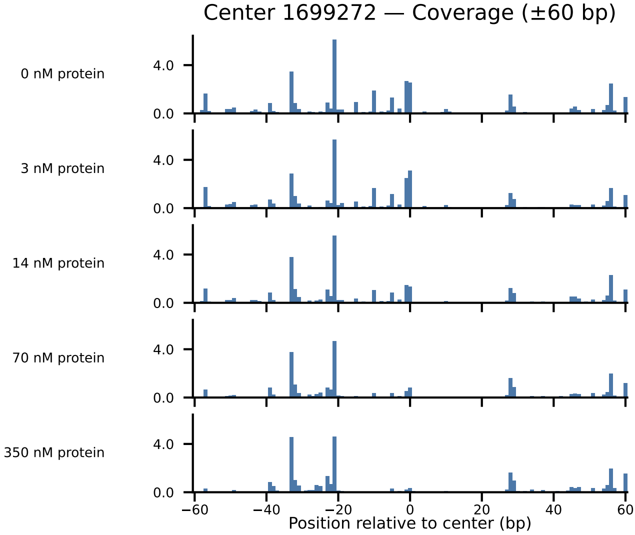
Center 1699272 — Coverage (±60 bp)
0 nM protein
3 nM protein
14 nM protein
70 nM protein
350 nM protein
4.0
0.0
−60
−40
−20
0
20
40
60
Position relative to center (bp)

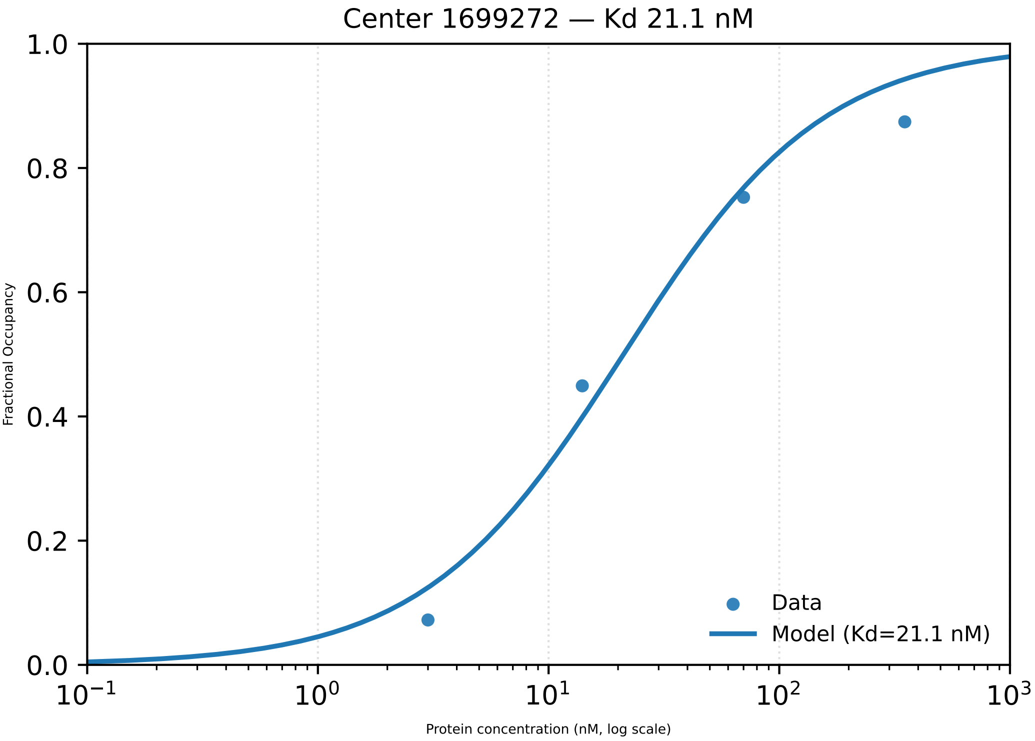

Center 1699272 — Kd 21.1 nM
1.0
0.8
0.6
0.4
0.2
0.0
Fractional Occupancy
Data
Model (Kd=21.1 nM)
$10^{-1}$
$10^{0}$
$10^{1}$
$10^{2}$
$10^{3}$
Protein concentration (nM, log scale)

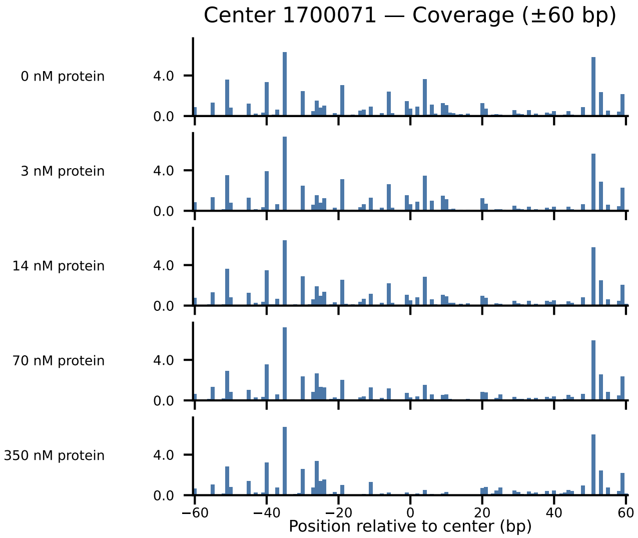
Center 1700071 — Coverage (±60 bp)
0 nM protein
3 nM protein
14 nM protein
70 nM protein
350 nM protein
4.0
0.0
−60
−40
−20
0
20
40
60
Position relative to center (bp)

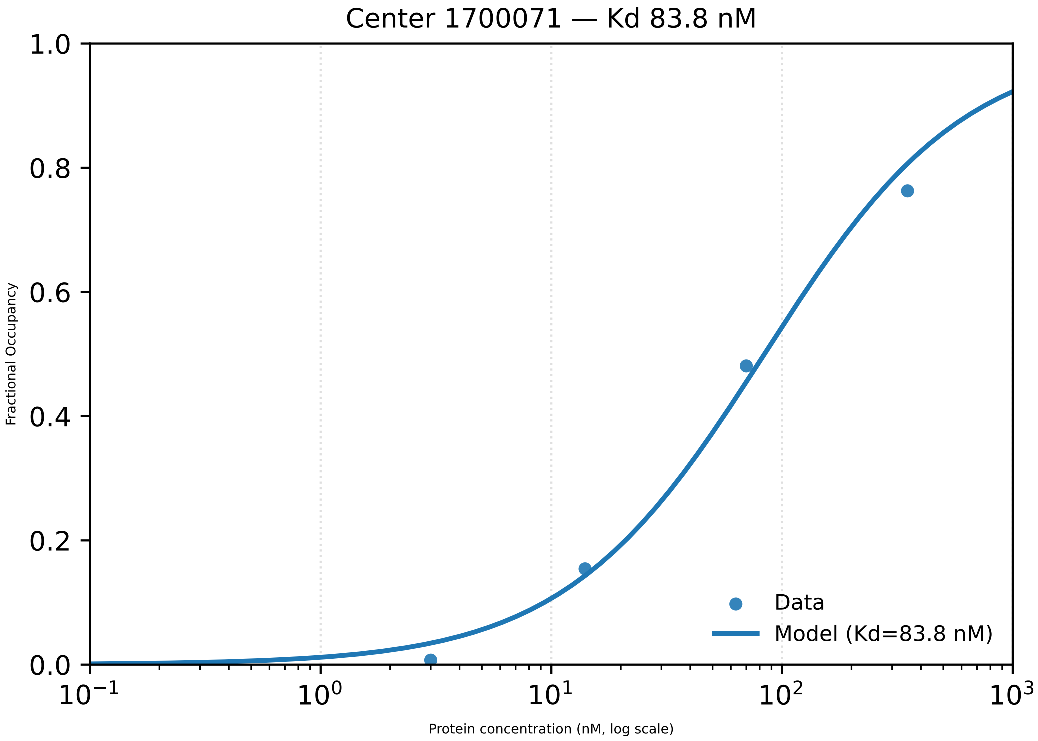

Center 1700071 — Kd 83.8 nM
1.0
0.8
0.6
0.4
0.2
0.0
Fractional Occupancy
$10^{-1}$
$10^{0}$
$10^{1}$
$10^{2}$
$10^{3}$
Protein concentration (nM, log scale)
Data
Model (Kd=83.8 nM)

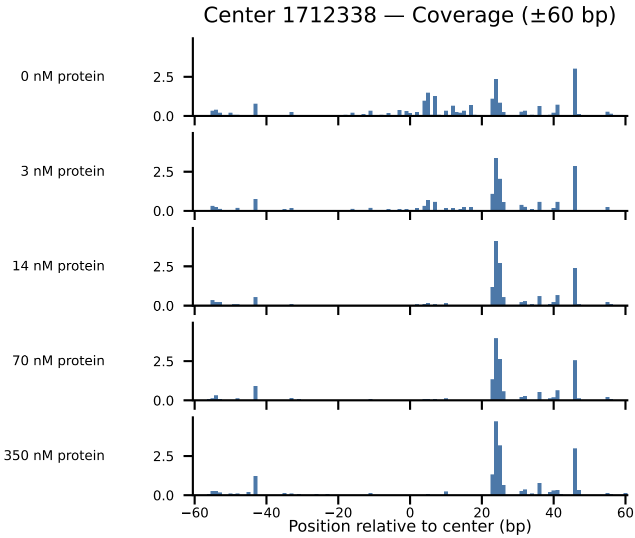

Center 1712338 — Coverage (±60 bp)
0 nM protein
3 nM protein
14 nM protein
70 nM protein
350 nM protein
0.0
2.5
−60
−40
−20
0
20
40
60
Position relative to center (bp)

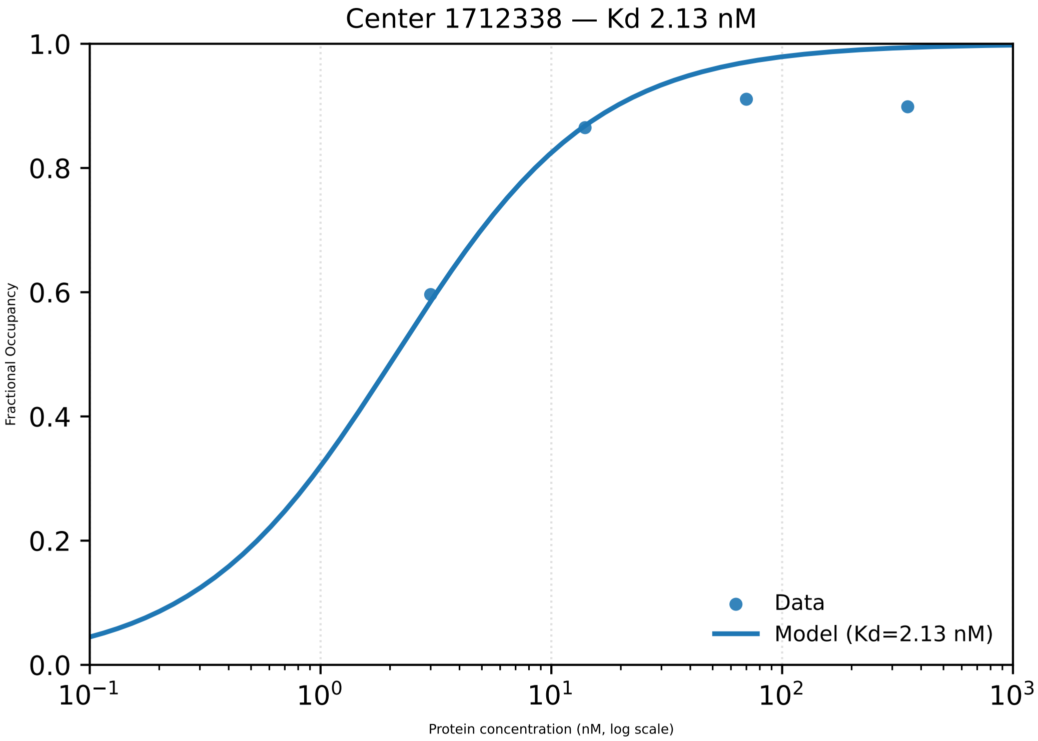

Center 1712338 — Kd 2.13 nM
1.0
0.8
0.6
0.4
0.2
0.0
Fractional Occupancy
Data
Model (Kd=2.13 nM)
$10^{-1}$
$10^{0}$
$10^{1}$
$10^{2}$
$10^{3}$
Protein concentration (nM, log scale)

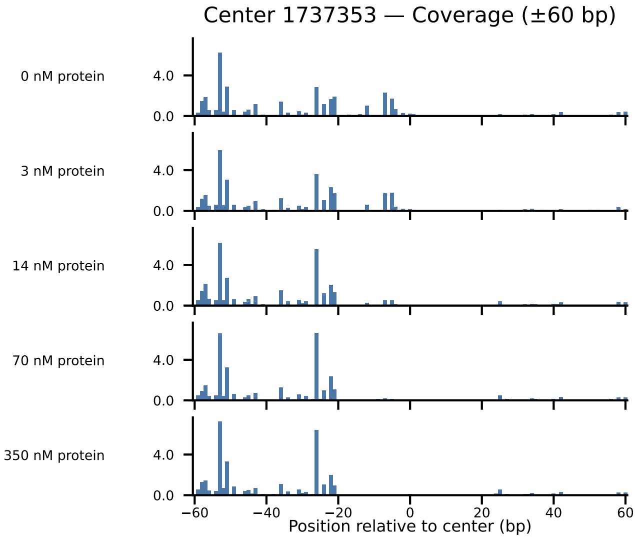

Center 1737353 — Coverage (±60 bp)
0 nM protein
3 nM protein
14 nM protein
70 nM protein
350 nM protein
4.0
0.0
−60
−40
−20
0
20
40
60
Position relative to center (bp)

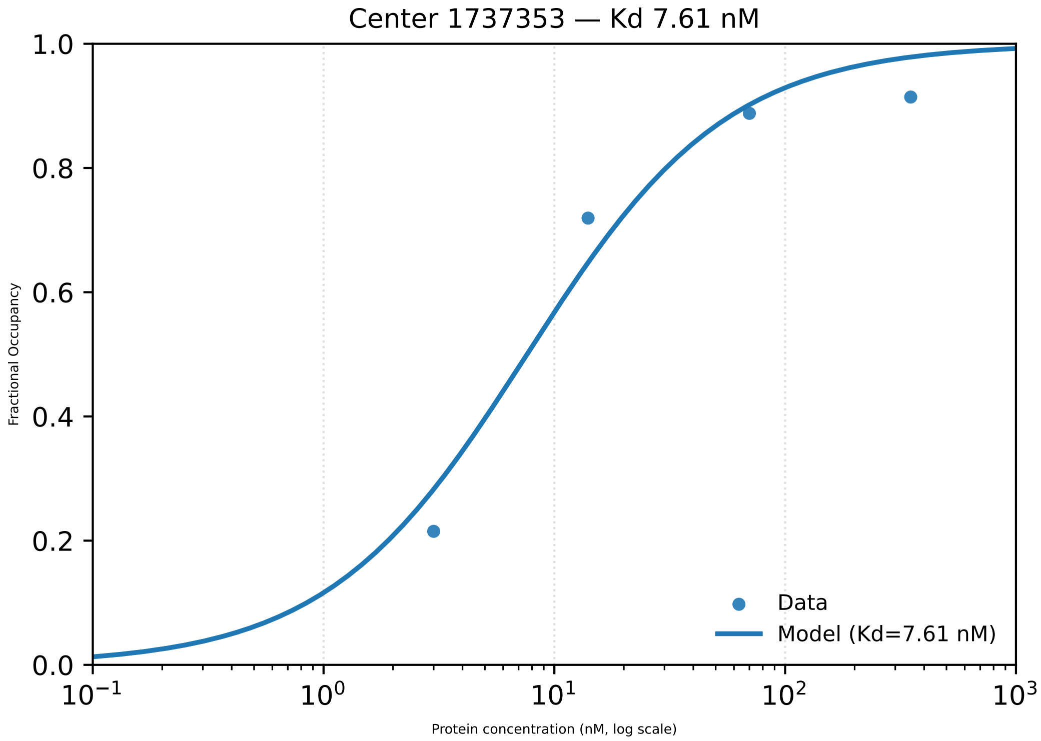

Center 1737353 — Kd 7.61 nM
1.0
0.8
0.6
0.4
0.2
0.0
Fractional Occupancy
$10^{-1}$
$10^{0}$
$10^{1}$
$10^{2}$
$10^{3}$
Protein concentration (nM, log scale)
Data
Model (Kd=7.61 nM)

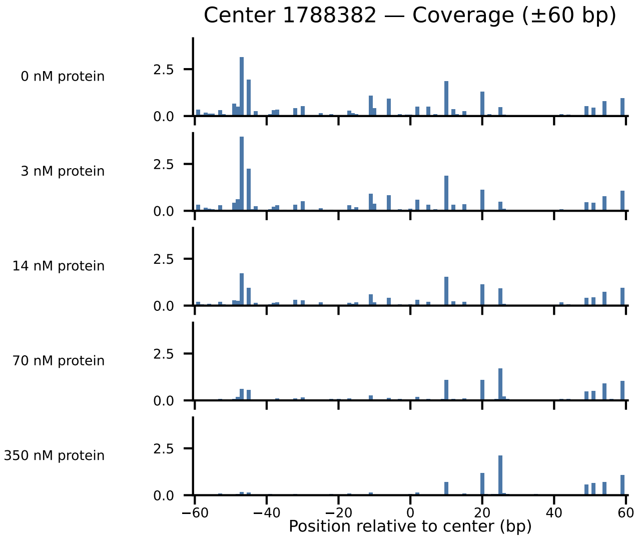

Center 1788382 — Coverage (±60 bp)
0 nM protein
3 nM protein
14 nM protein
70 nM protein
350 nM protein
2.5
0.0
−60
−40
−20
0
20
40
60
Position relative to center (bp)

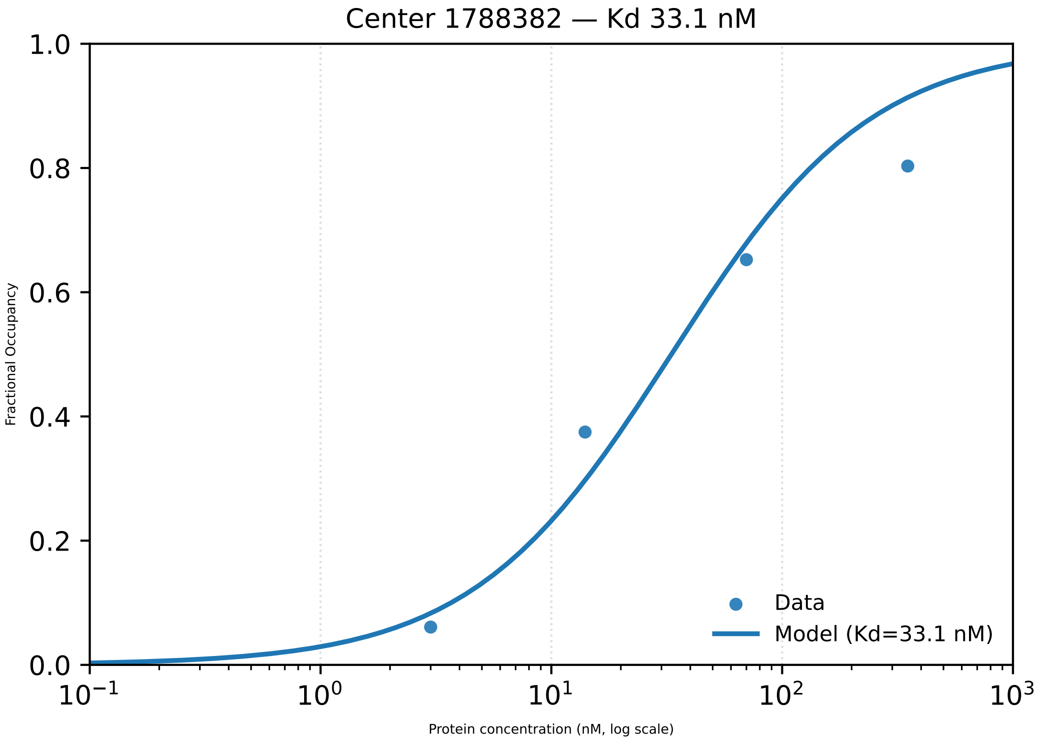

Center 1788382 — Kd 33.1 nM
Fractional Occupancy
1.0
0.8
0.6
0.4
0.2
0.0
$10^{-1}$
$10^{0}$
$10^{1}$
$10^{2}$
$10^{3}$
Protein concentration (nM, log scale)
Data
Model (Kd=33.1 nM)

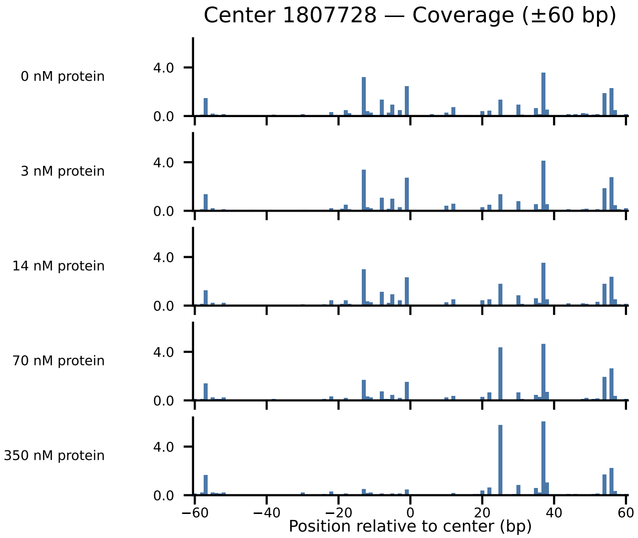

Center 1807728 — Coverage (±60 bp)
0 nM protein
3 nM protein
14 nM protein
70 nM protein
350 nM protein
4.0
0.0
−60
−40
−20
0
20
40
60
Position relative to center (bp)

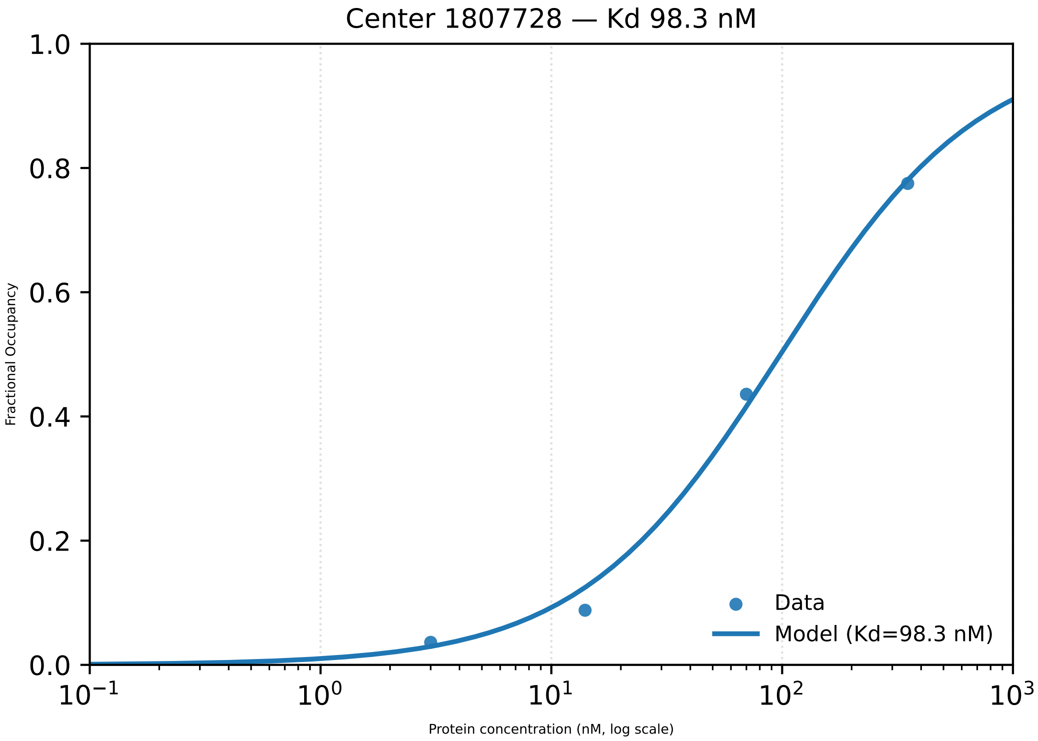

Center 1807728 — Kd 98.3 nM
1.0
0.8
0.6
0.4
0.2
0.0
Fractional Occupancy
Data
Model (Kd=98.3 nM)
$10^{-1}$
$10^{0}$
$10^{1}$
$10^{2}$
$10^{3}$
Protein concentration (nM, log scale)

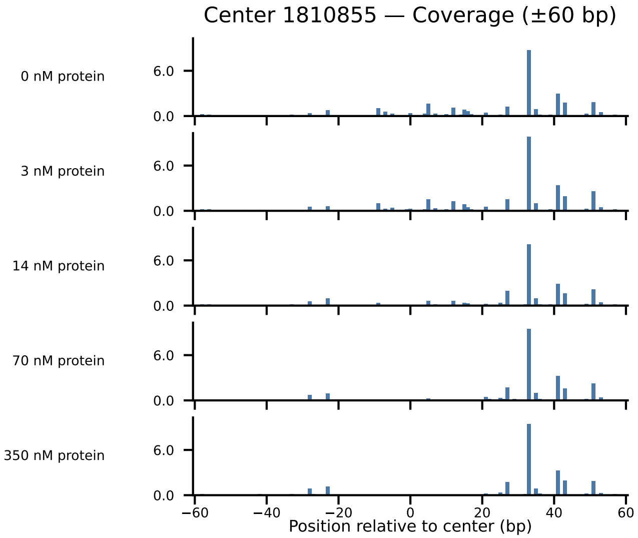

Center 1810855 — Coverage (±60 bp)
0 nM protein
3 nM protein
14 nM protein
70 nM protein
350 nM protein
6.0
0.0
−60
−40
−20
0
20
40
60
Position relative to center (bp)

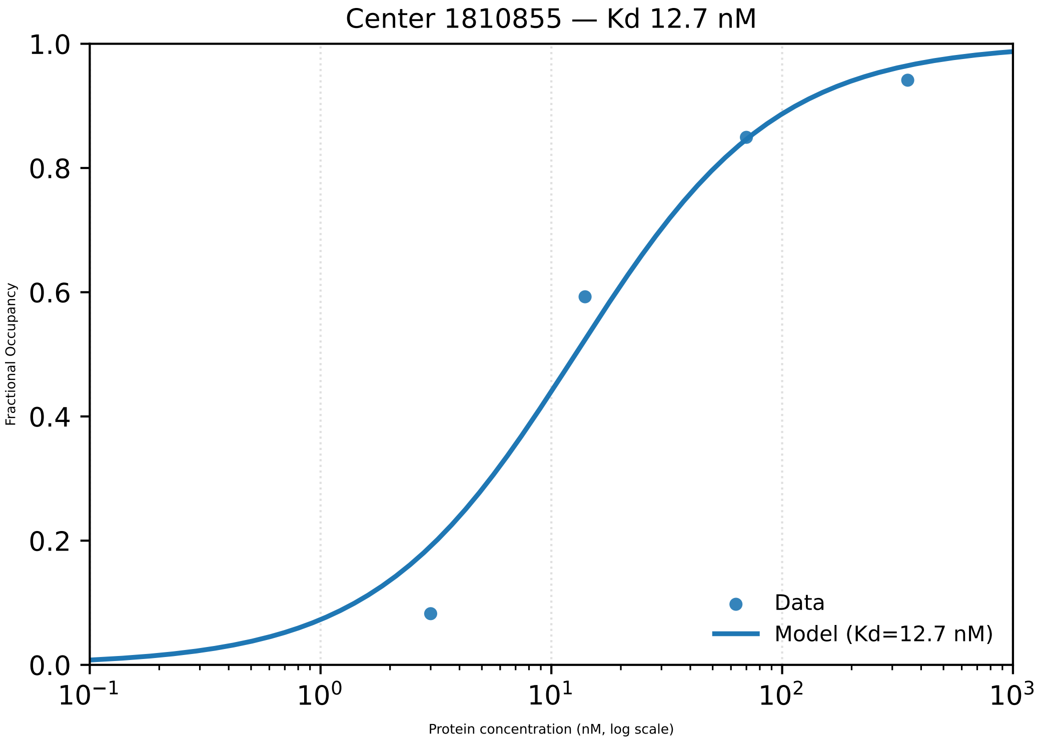

Center 1810855 — Kd 12.7 nM
1.0
0.8
0.6
0.4
0.2
0.0
Fractional Occupancy
$10^{-1}$
$10^{0}$
$10^{1}$
$10^{2}$
$10^{3}$
Protein concentration (nM, log scale)
Data
Model (Kd=12.7 nM)

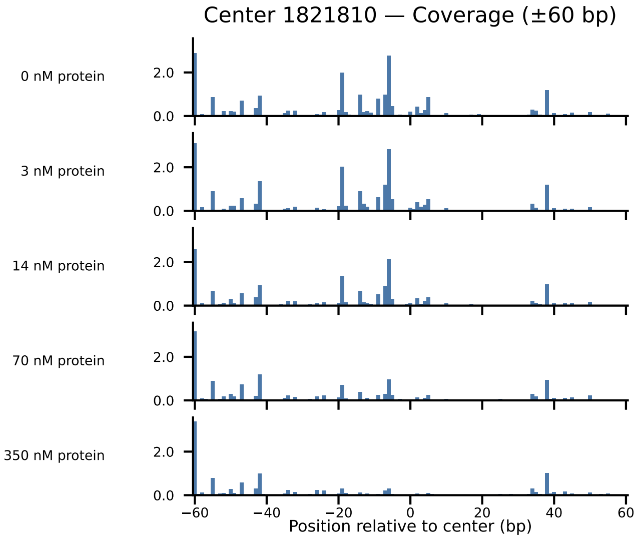

Center 1821810 — Coverage (±60 bp)
0 nM protein
3 nM protein
14 nM protein
70 nM protein
350 nM protein
2.0
0.0
−60
−40
−20
0
20
40
60
Position relative to center (bp)

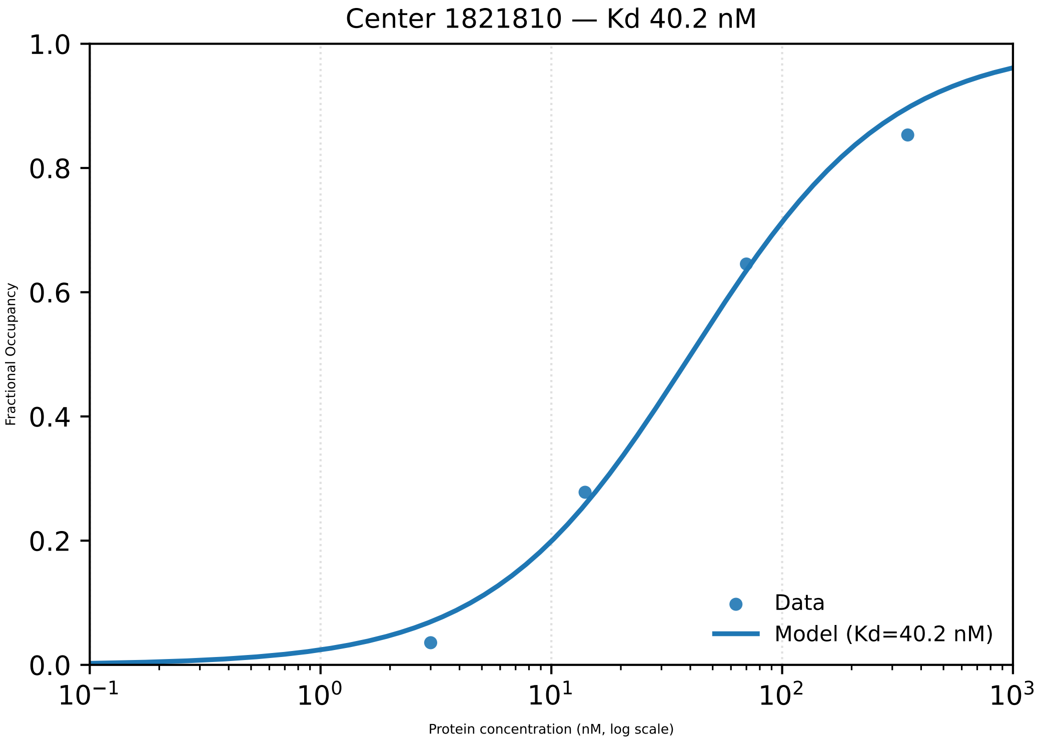

Center 1821810 — Kd 40.2 nM
1.0
0.8
0.6
0.4
0.2
0.0
Fractional Occupancy
Data
Model (Kd=40.2 nM)
$10^{-1}$
$10^{0}$
$10^{1}$
$10^{2}$
$10^{3}$
Protein concentration (nM, log scale)

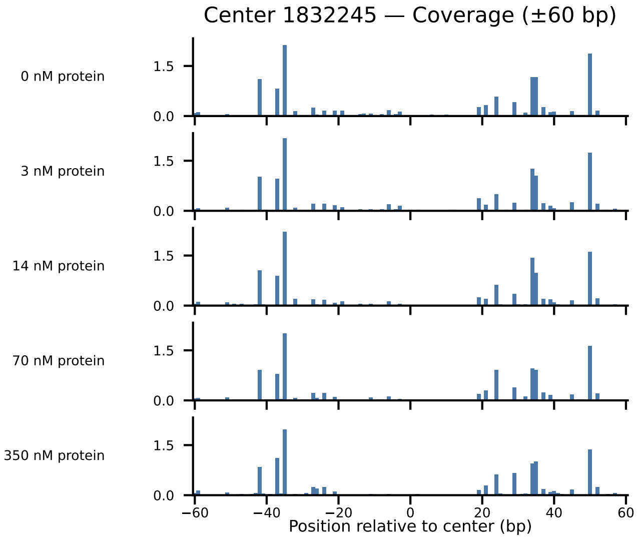

Center 1832245 — Coverage (±60 bp)
0 nM protein
3 nM protein
14 nM protein
70 nM protein
350 nM protein
1.5
0.0
−60
−40
−20
0
20
40
60
Position relative to center (bp)

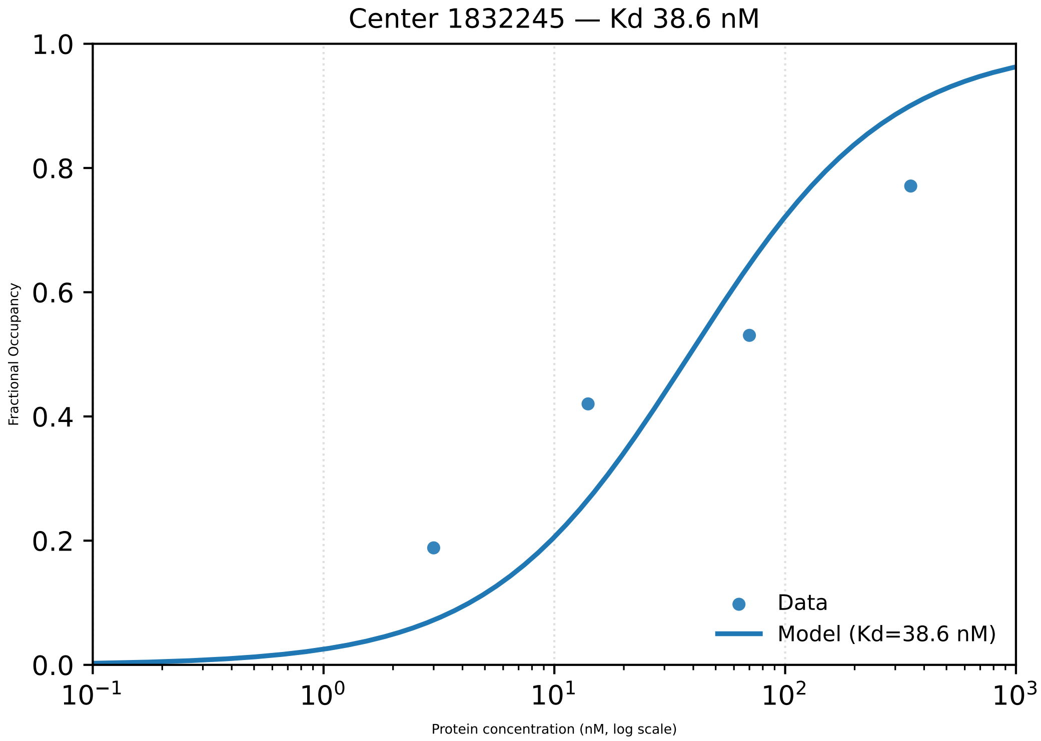

Center 1832245 — Kd 38.6 nM
1.0
0.8
0.6
0.4
0.2
0.0
Fractional Occupancy
$10^{-1}$
$10^{0}$
$10^{1}$
$10^{2}$
$10^{3}$
Protein concentration (nM, log scale)
Data
Model (Kd=38.6 nM)

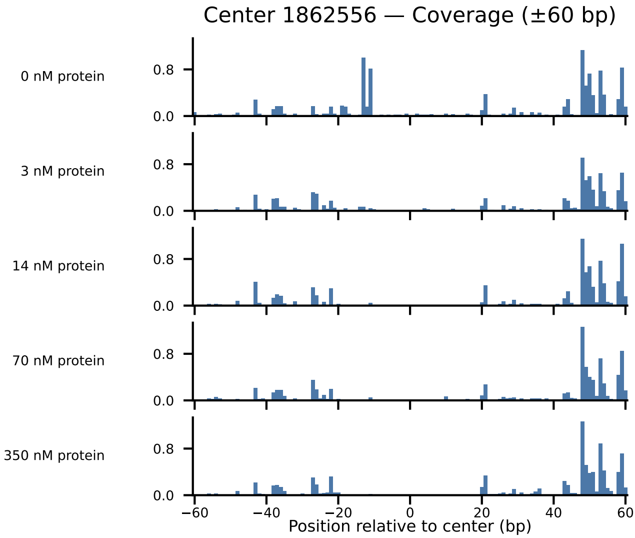

Center 1862556 — Coverage (±60 bp)
0 nM protein
3 nM protein
14 nM protein
70 nM protein
350 nM protein
0.8
0.0
−60
−40
−20
0
20
40
60
Position relative to center (bp)

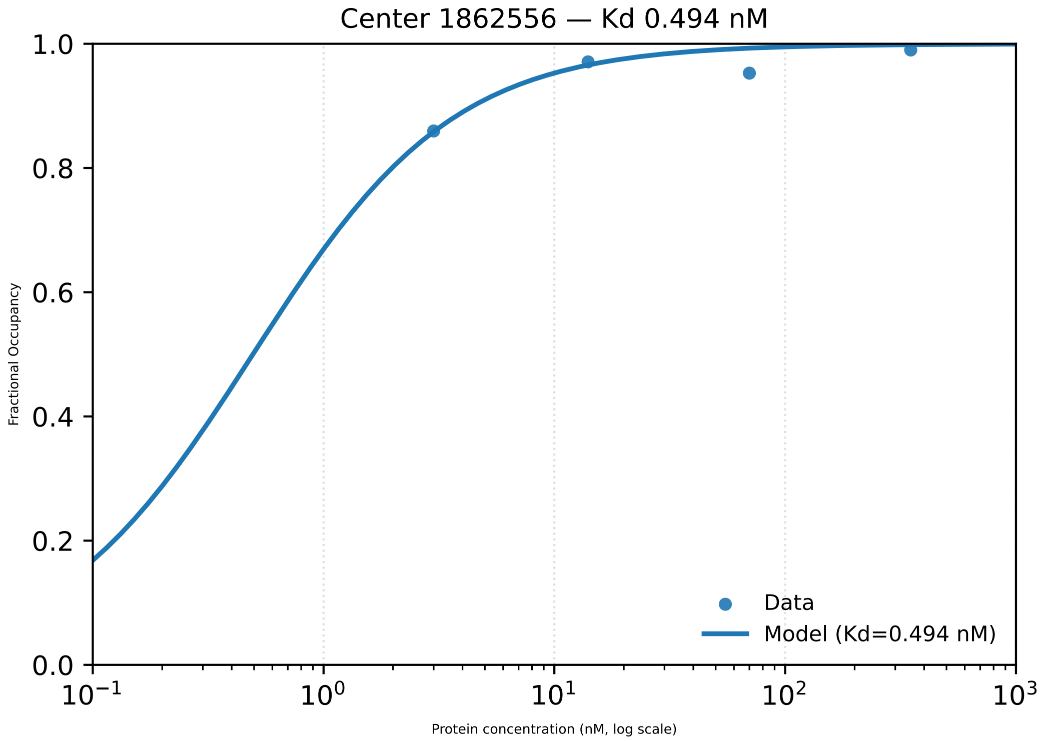

Center 1862556 — Kd 0.494 nM
Fractional Occupancy
1.0
0.8
0.6
0.4
0.2
0.0
$10^{-1}$
$10^{0}$
$10^{1}$
$10^{2}$
$10^{3}$
Protein concentration (nM, log scale)
Data
Model (Kd=0.494 nM)

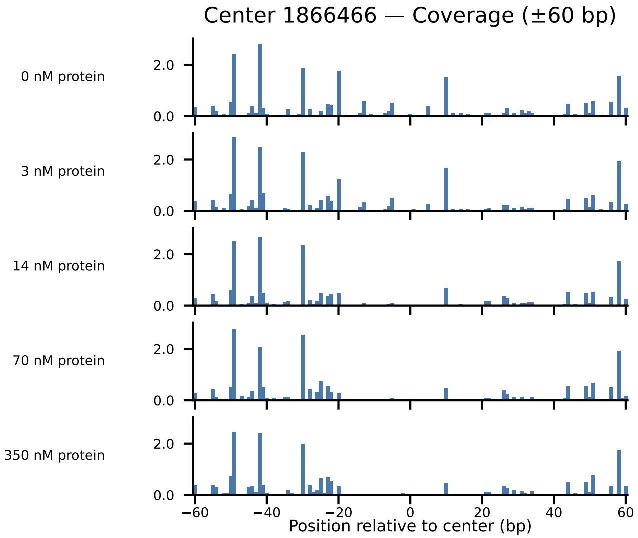

Center 1866466 — Coverage (±60 bp)
0 nM protein
3 nM protein
14 nM protein
70 nM protein
350 nM protein
2.0
0.0
−60
−40
−20
0
20
40
60
Position relative to center (bp)

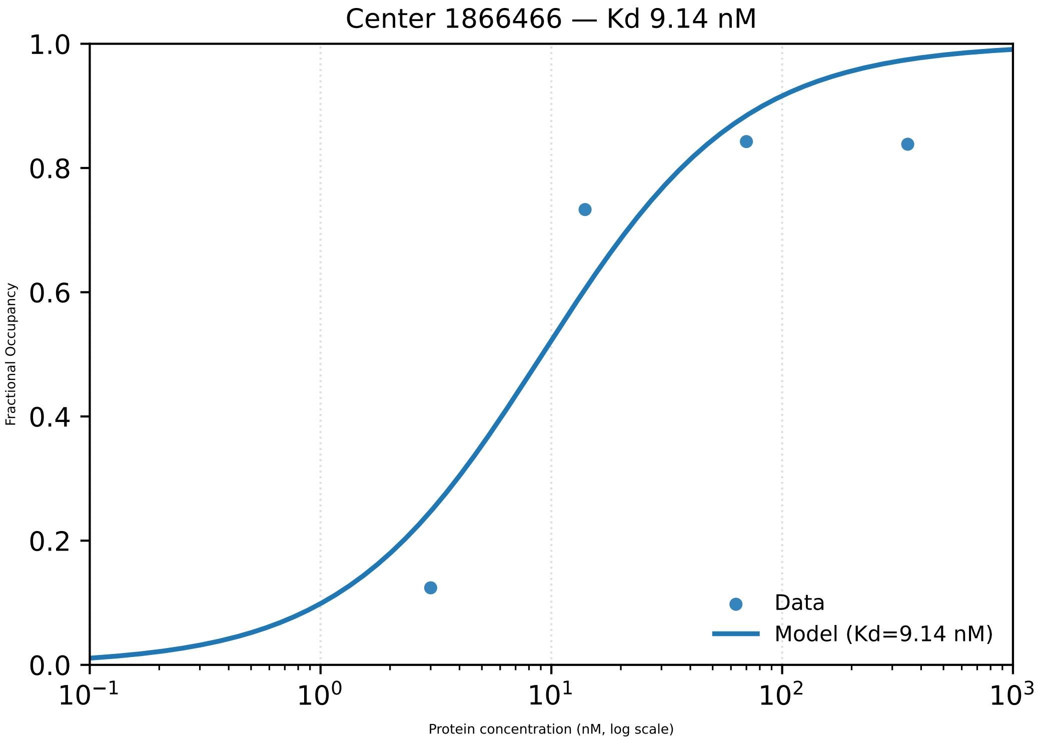
Center 1866466 — Kd 9.14 nM
1.0
0.8
0.6
0.4
0.2
0.0
Fractional Occupancy
10−1
100
101
102
103
Protein concentration (nM, log scale)
Data
Model (Kd=9.14 nM)

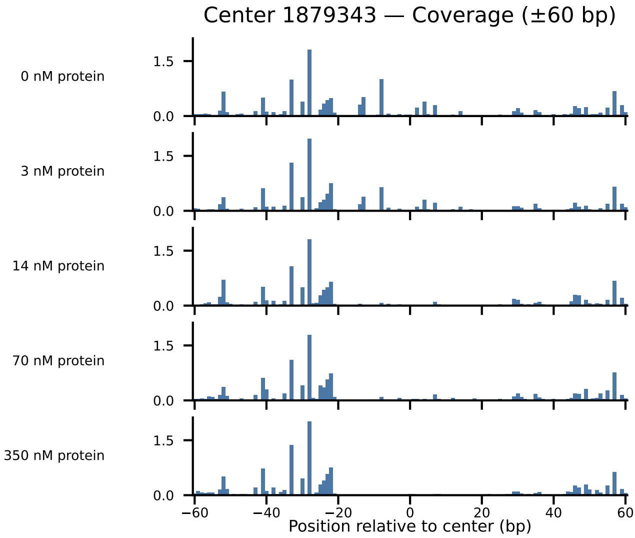

Center 1879343 — Coverage (±60 bp)
0 nM protein
3 nM protein
14 nM protein
70 nM protein
350 nM protein
1.5
0.0
−60
−40
−20
0
20
40
60
Position relative to center (bp)

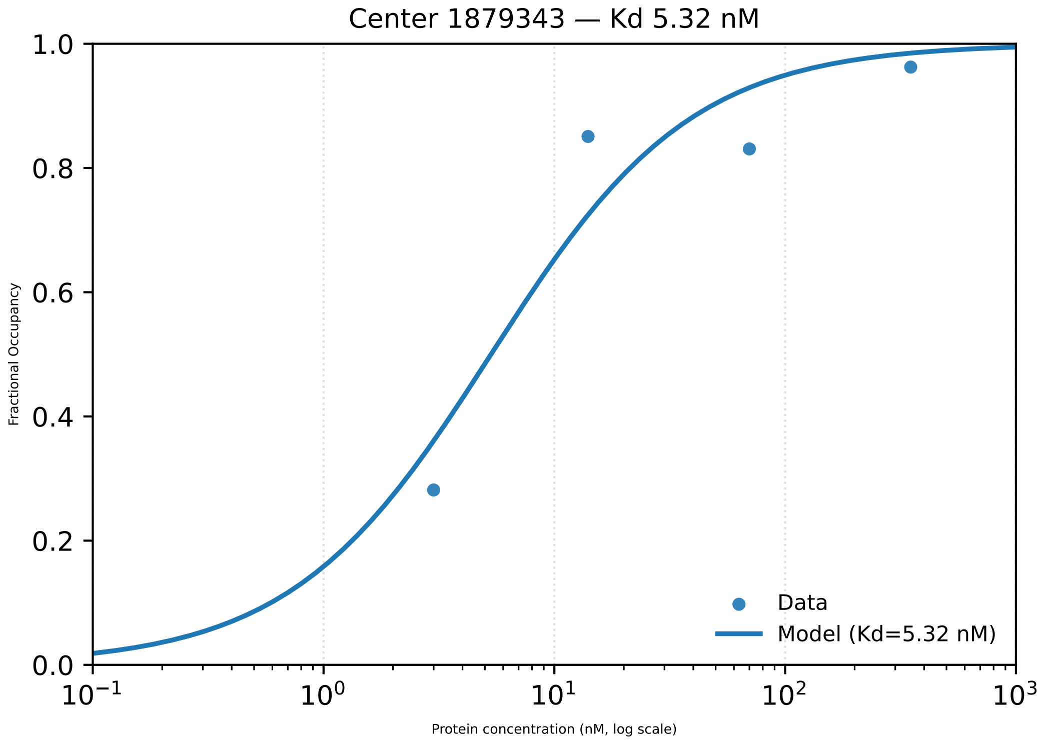

Center 1879343 — Kd 5.32 nM
Fractional Occupancy
1.0
0.8
0.6
0.4
0.2
0.0
$10^{-1}$
$10^{0}$
$10^{1}$
$10^{2}$
$10^{3}$
Protein concentration (nM, log scale)
Data
Model (Kd=5.32 nM)

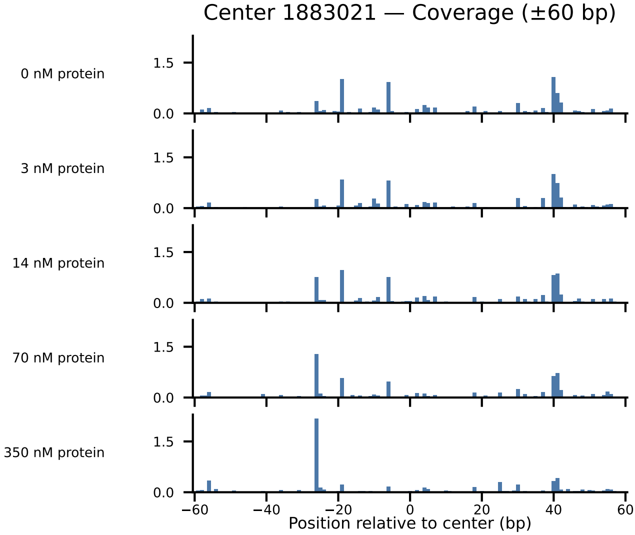
Center 1883021 — Coverage (±60 bp)
0 nM protein
3 nM protein
14 nM protein
70 nM protein
350 nM protein
1.5
0.0
−60
−40
−20
0
20
40
60
Position relative to center (bp)

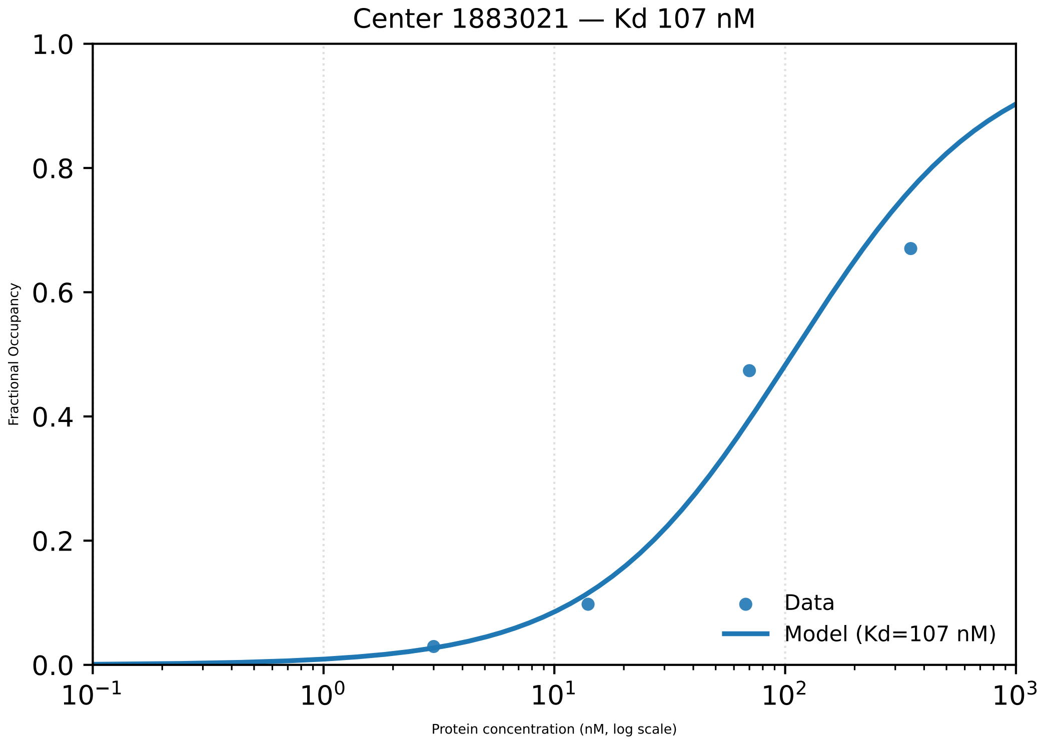

Center 1883021 — Kd 107 nM
1.0
0.8
0.6
0.4
0.2
0.0
Fractional Occupancy
$10^{-1}$
$10^{0}$
$10^{1}$
$10^{2}$
$10^{3}$
Protein concentration (nM, log scale)
Data
Model (Kd=107 nM)

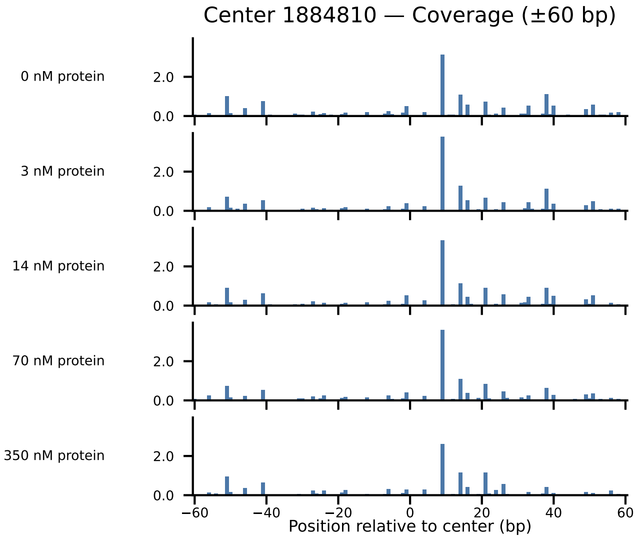

Center 1884810 — Coverage (±60 bp)
0 nM protein
3 nM protein
14 nM protein
70 nM protein
350 nM protein
2.0
0.0
−60
−40
−20
0
20
40
60
Position relative to center (bp)

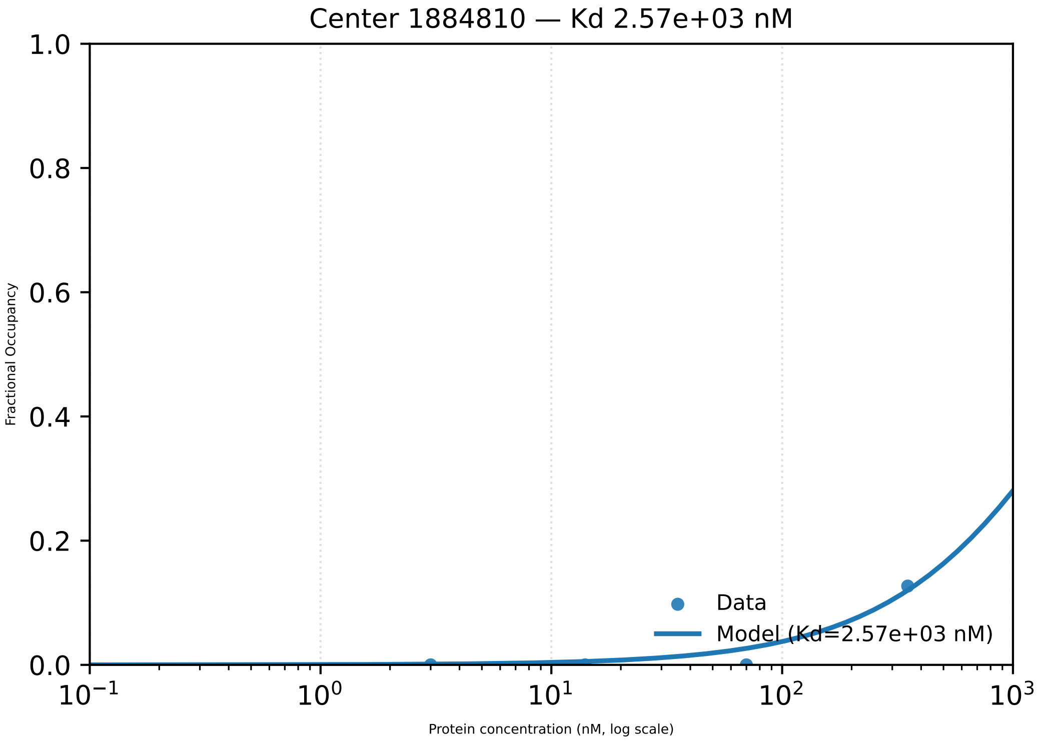
Center 1884810 — Kd 2.57e+03 nM
1.0
0.8
0.6
0.4
0.2
0.0
Fractional Occupancy
$10^{-1}$
$10^{0}$
$10^{1}$
$10^{2}$
$10^{3}$
Protein concentration (nM, log scale)
Data
Model (Kd=2.57e+03 nM)

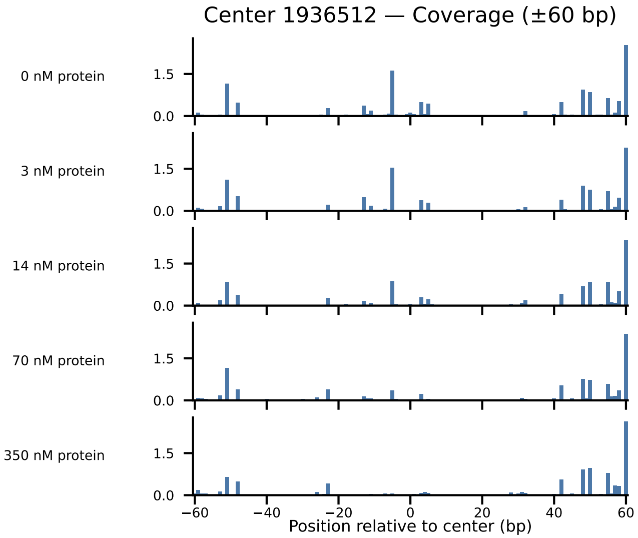

Center 1936512 — Coverage (±60 bp)
0 nM protein
3 nM protein
14 nM protein
70 nM protein
350 nM protein
1.5
0.0
−60
−40
−20
0
20
40
60
Position relative to center (bp)

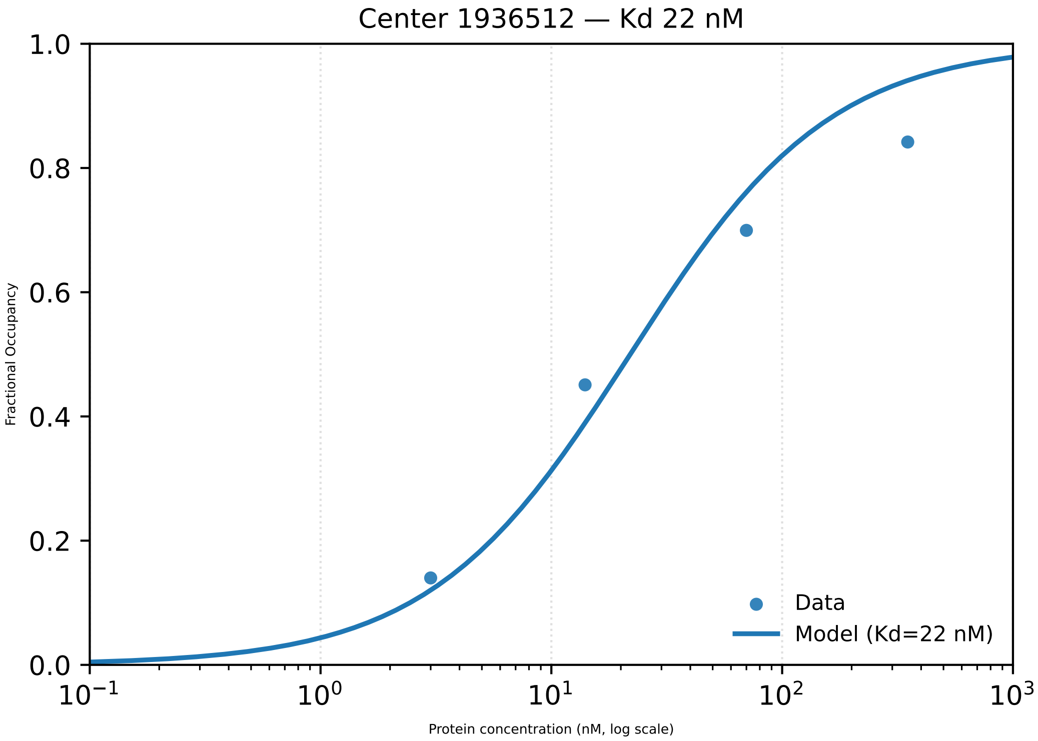

Center 1936512 — Kd 22 nM
Fractional Occupancy
1.0
0.8
0.6
0.4
0.2
0.0
$10^{-1}$
$10^{0}$
$10^{1}$
$10^{2}$
$10^{3}$
Protein concentration (nM, log scale)
Data
Model (Kd=22 nM)

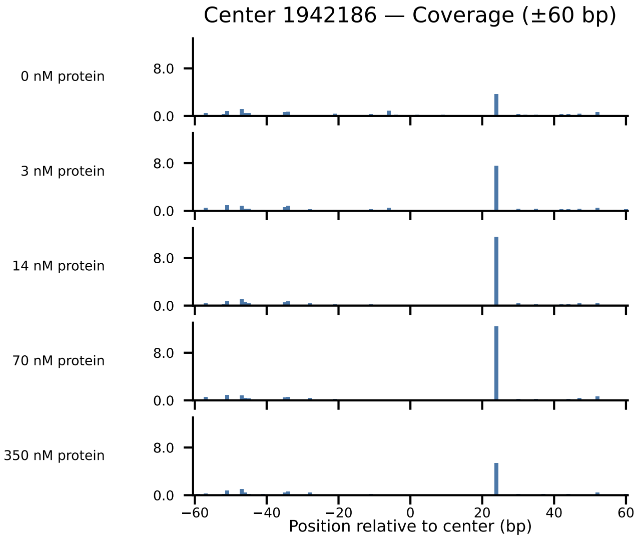

Center 1942186 — Coverage (±60 bp)
0 nM protein
3 nM protein
14 nM protein
70 nM protein
350 nM protein
8.0
0.0
−60
−40
−20
0
20
40
60
Position relative to center (bp)

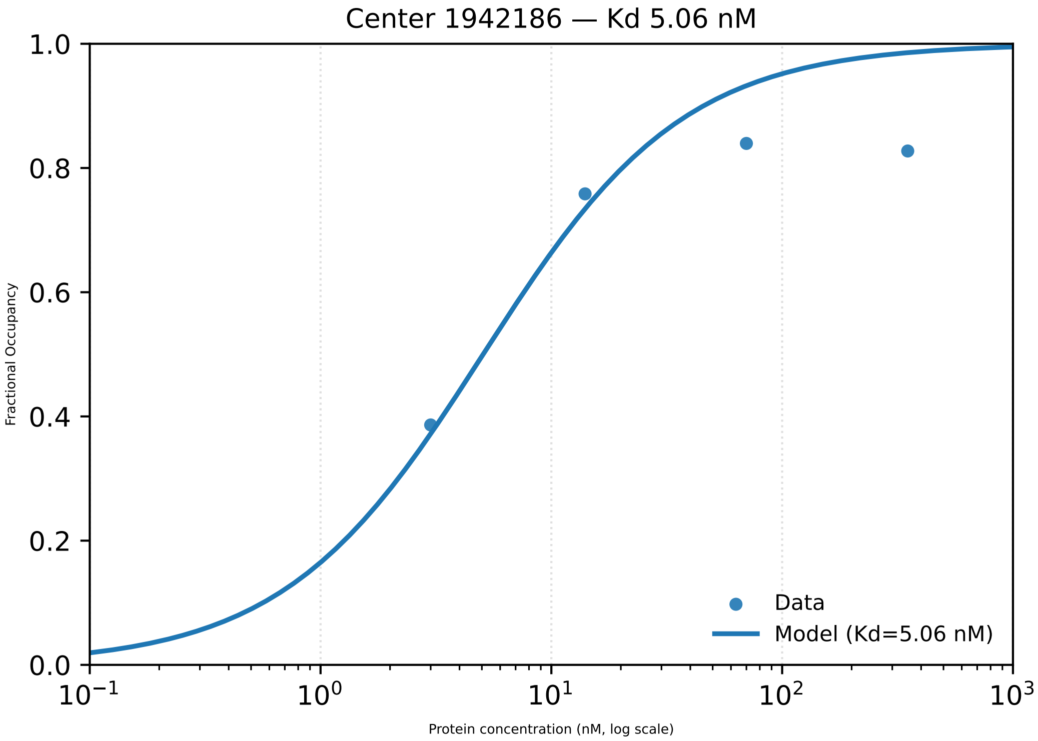

Center 1942186 — Kd 5.06 nM
Fractional Occupancy
1.0
0.8
0.6
0.4
0.2
0.0
$10^{-1}$
$10^{0}$
$10^{1}$
$10^{2}$
$10^{3}$
Protein concentration (nM, log scale)
Data
Model (Kd=5.06 nM)

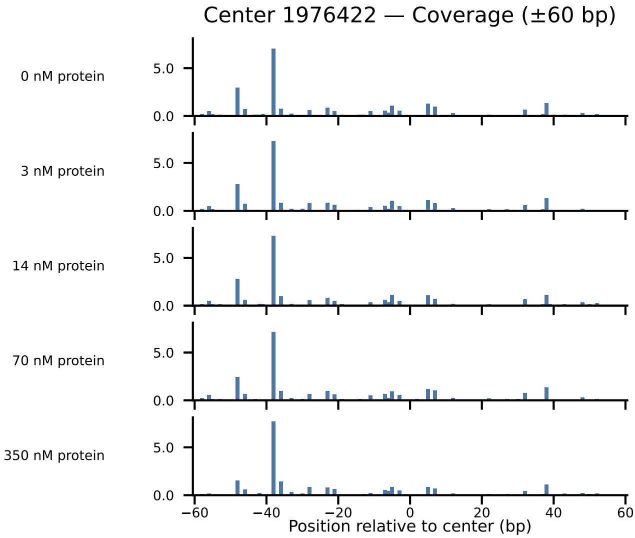

Center 1976422 — Coverage (±60 bp)
0 nM protein
3 nM protein
14 nM protein
70 nM protein
350 nM protein
5.0
0.0
−60
−40
−20
0
20
40
60
Position relative to center (bp)

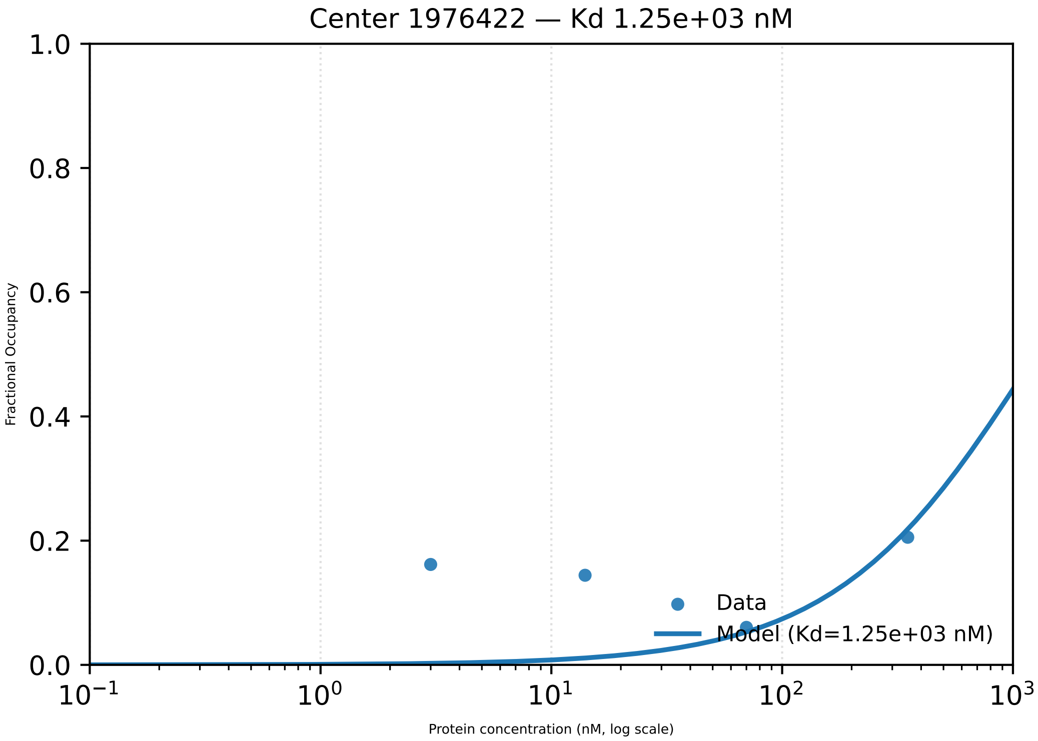

Center 1976422 — Kd 1.25e+03 nM
1.0
0.8
0.6
0.4
0.2
0.0
Fractional Occupancy
$10^{-1}$
$10^{0}$
$10^{1}$
$10^{2}$
$10^{3}$
Protein concentration (nM, log scale)
Data
Model (Kd=1.25e+03 nM)

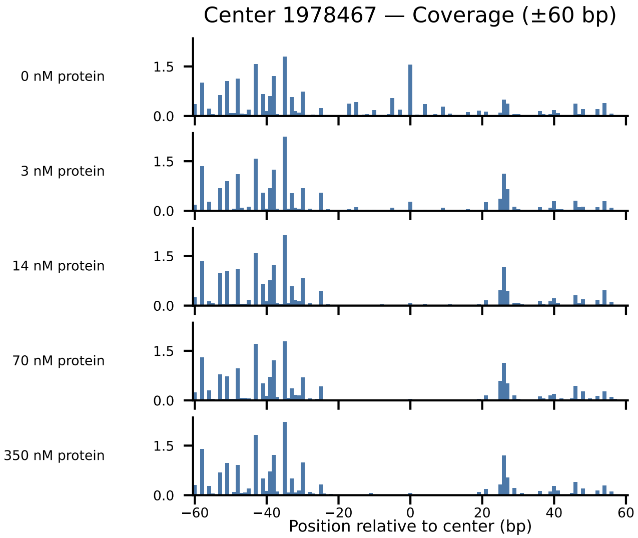

Center 1978467 — Coverage (±60 bp)
0 nM protein
3 nM protein
14 nM protein
70 nM protein
350 nM protein
1.5
0.0
−60
−40
−20
0
20
40
60
Position relative to center (bp)

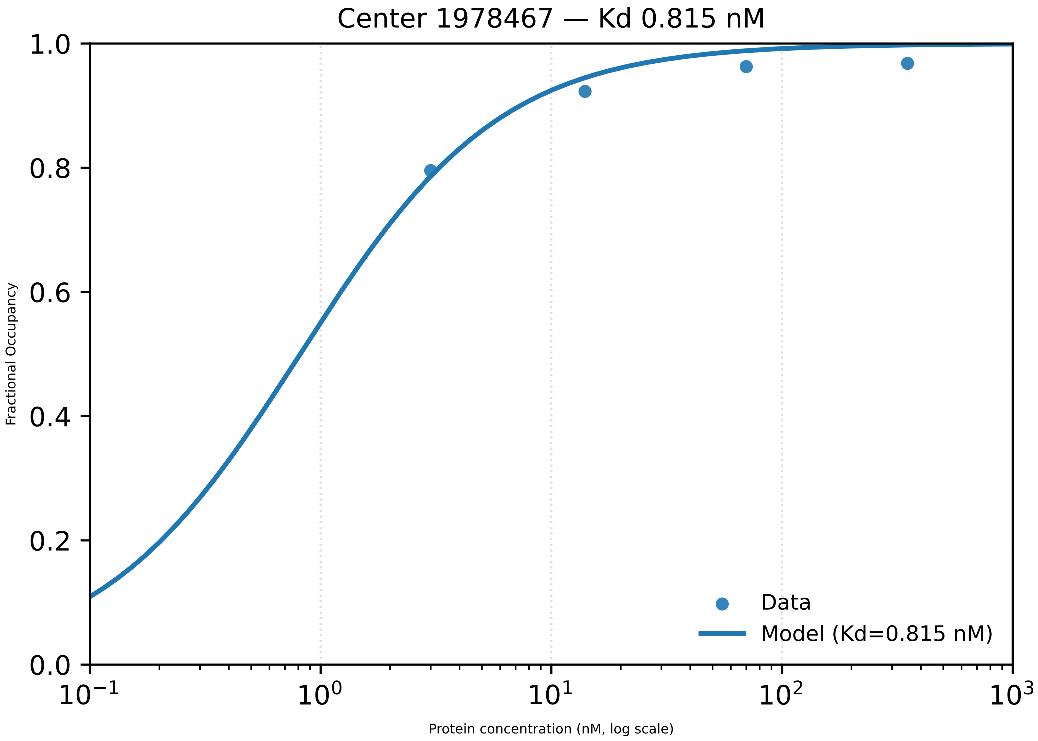

Center 1978467 — Kd 0.815 nM
1.0
0.8
0.6
0.4
0.2
0.0
Fractional Occupancy
$10^{-1}$
$10^{0}$
$10^{1}$
$10^{2}$
$10^{3}$
Protein concentration (nM, log scale)
Data
Model (Kd=0.815 nM)

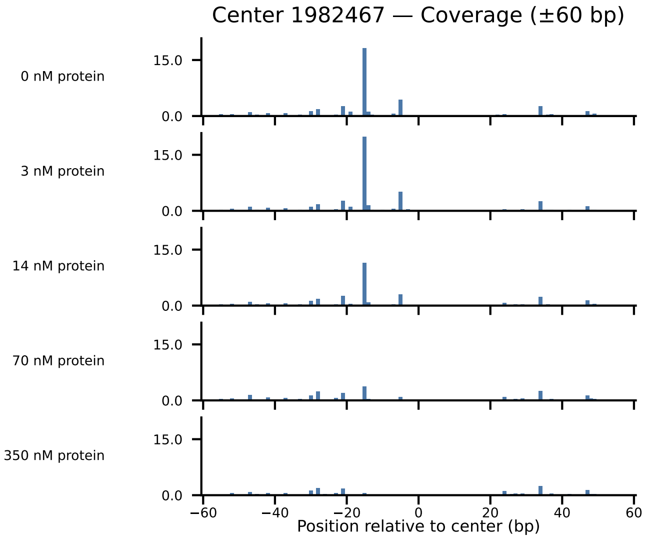

Center 1982467 — Coverage (±60 bp)
0 nM protein
3 nM protein
14 nM protein
70 nM protein
350 nM protein
15.0
0.0
−60
−40
−20
0
20
40
60
Position relative to center (bp)

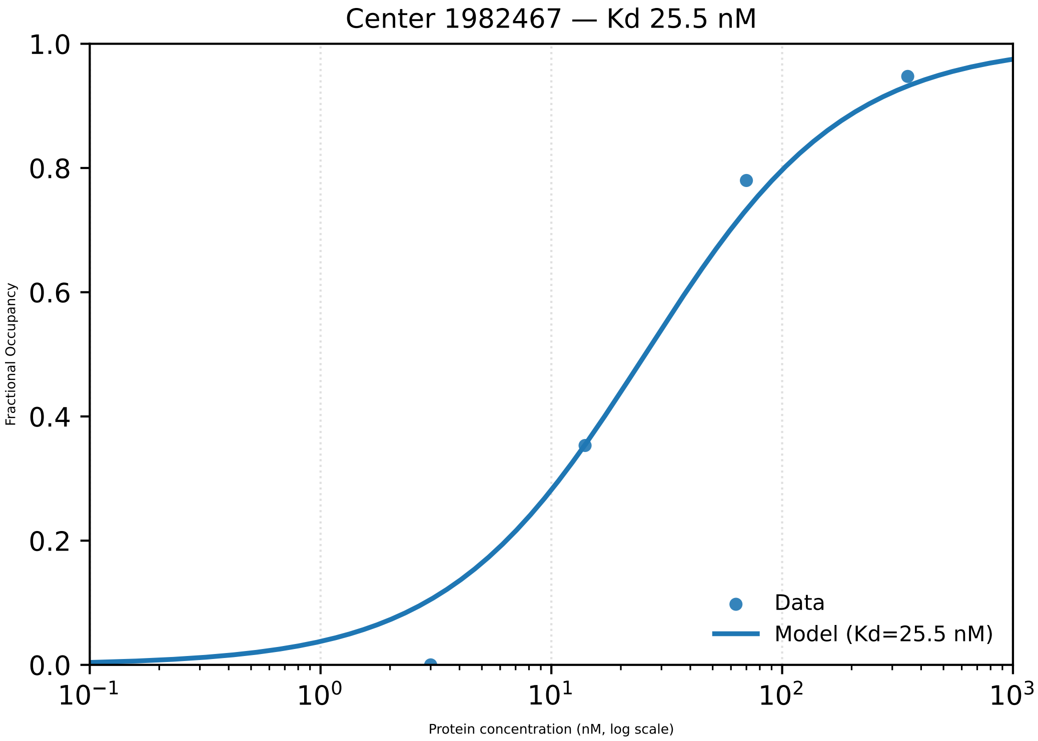

Center 1982467 — Kd 25.5 nM
Fractional Occupancy
1.0
0.8
0.6
0.4
0.2
0.0
$10^{-1}$
$10^{0}$
$10^{1}$
$10^{2}$
$10^{3}$
Protein concentration (nM, log scale)
Data
Model (Kd=25.5 nM)

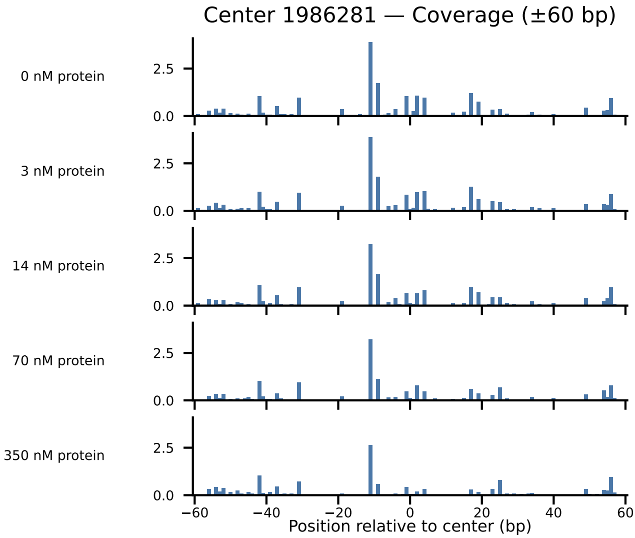

Center 1986281 — Coverage (±60 bp)
0 nM protein
3 nM protein
14 nM protein
70 nM protein
350 nM protein
2.5
0.0
−60
−40
−20
0
20
40
60
Position relative to center (bp)

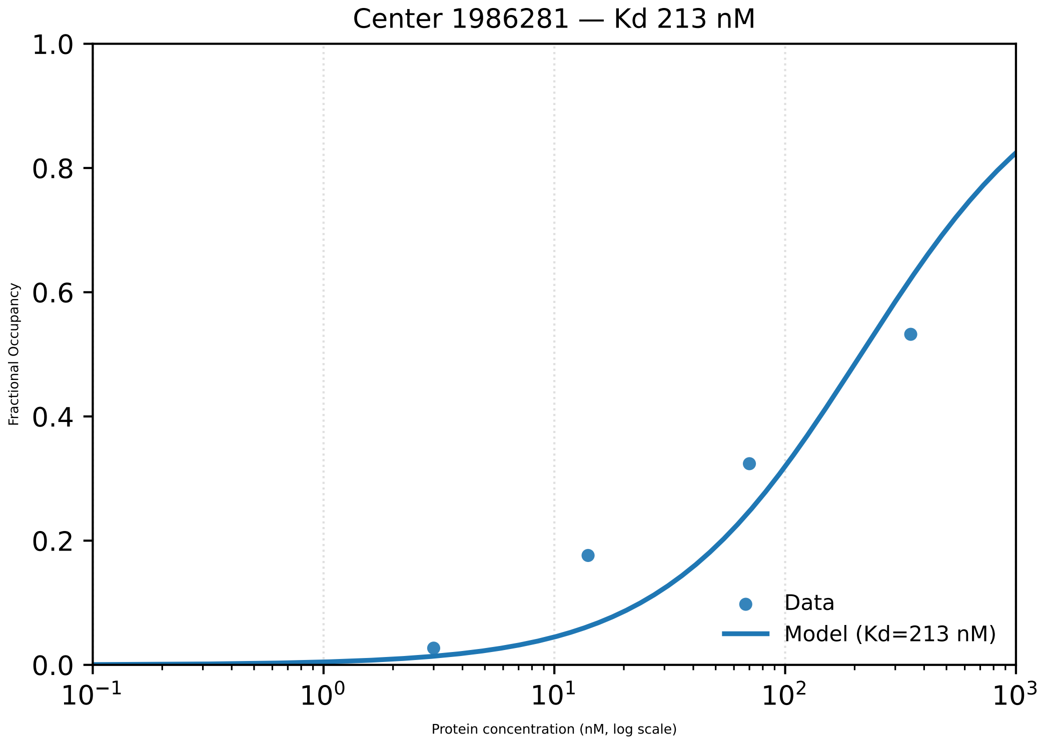

Center 1986281 — Kd 213 nM
Fractional Occupancy
1.0
0.8
0.6
0.4
0.2
0.0
$10^{-1}$
$10^{0}$
$10^{1}$
$10^{2}$
$10^{3}$
Protein concentration (nM, log scale)
Data
Model (Kd=213 nM)

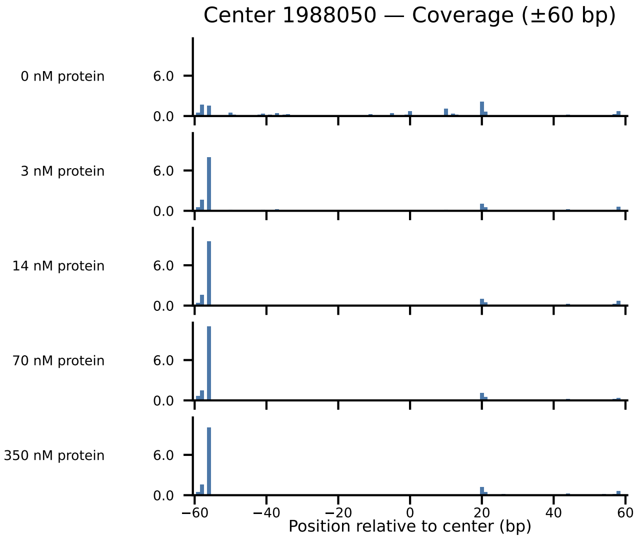

Center 1988050 — Coverage (±60 bp)
0 nM protein
3 nM protein
14 nM protein
70 nM protein
350 nM protein
6.0
0.0
−60
−40
−20
0
20
40
60
Position relative to center (bp)

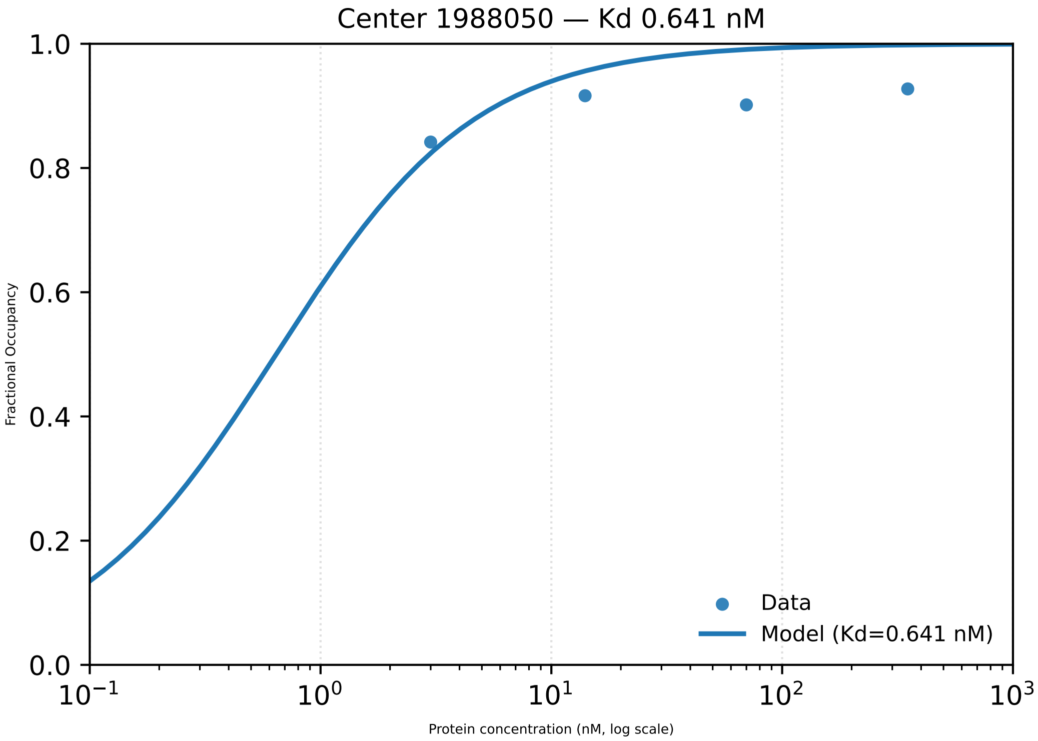
Center 1988050 — Kd 0.641 nM
1.0
0.8
0.6
0.4
0.2
0.0
Fractional Occupancy
$10^{-1}$
$10^{0}$
$10^{1}$
$10^{2}$
$10^{3}$
Protein concentration (nM, log scale)
Data
Model (Kd=0.641 nM)

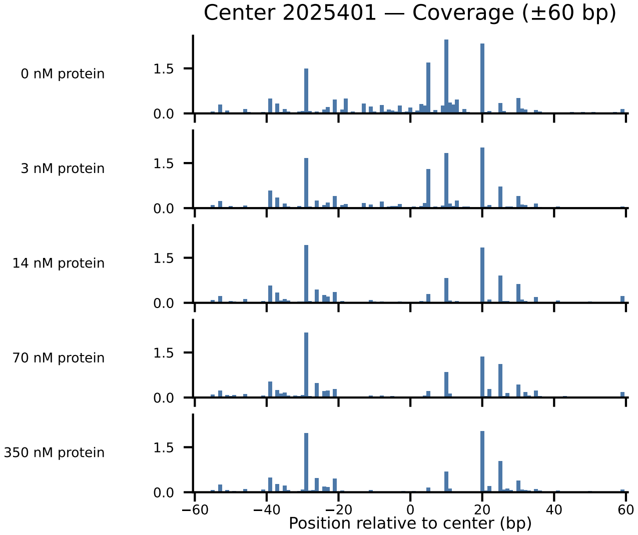
Center 2025401 — Coverage (±60 bp)
0 nM protein
3 nM protein
14 nM protein
70 nM protein
350 nM protein
1.5
0.0
−60
−40
−20
0
20
40
60
Position relative to center (bp)

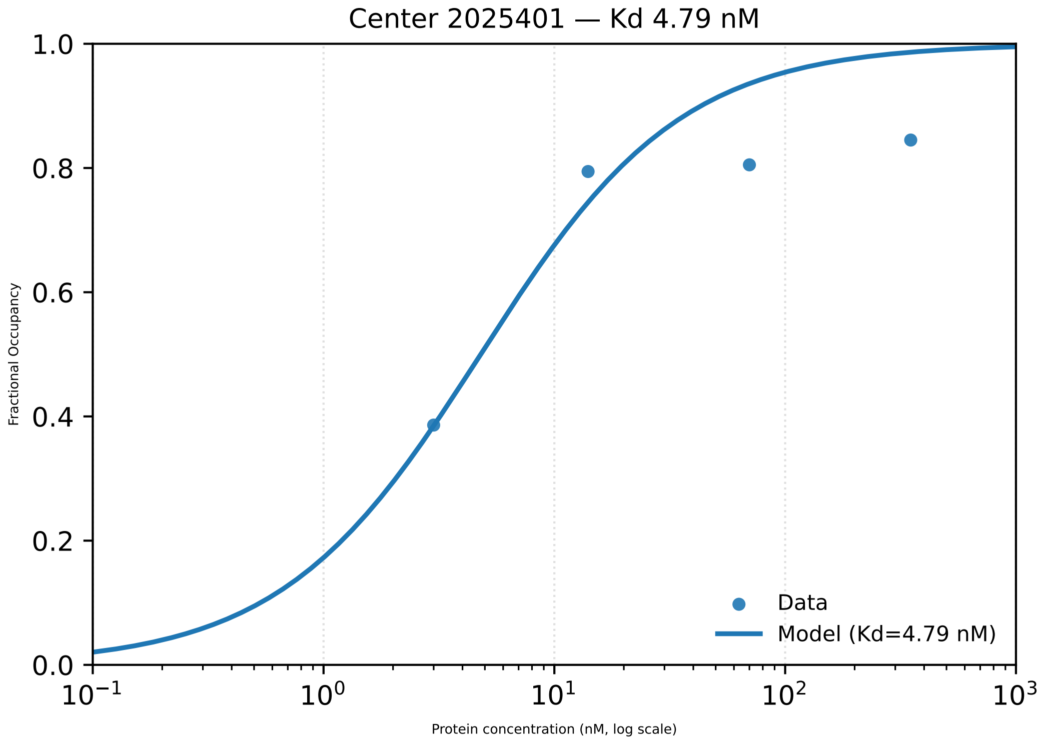

Center 2025401 — Kd 4.79 nM
Fractional Occupancy
1.0
0.8
0.6
0.4
0.2
0.0
$10^{-1}$
$10^{0}$
$10^{1}$
$10^{2}$
$10^{3}$
Protein concentration (nM, log scale)
Data
Model (Kd=4.79 nM)

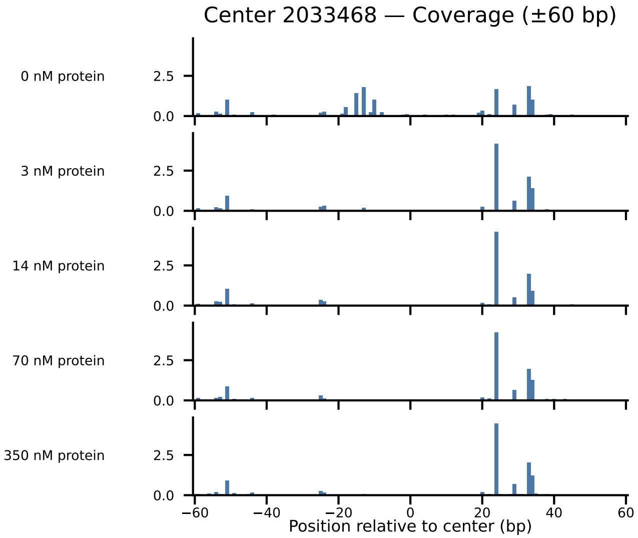

Center 2033468 — Coverage (±60 bp)
0 nM protein
3 nM protein
14 nM protein
70 nM protein
350 nM protein
2.5
0.0
−60
−40
−20
0
20
40
60
Position relative to center (bp)

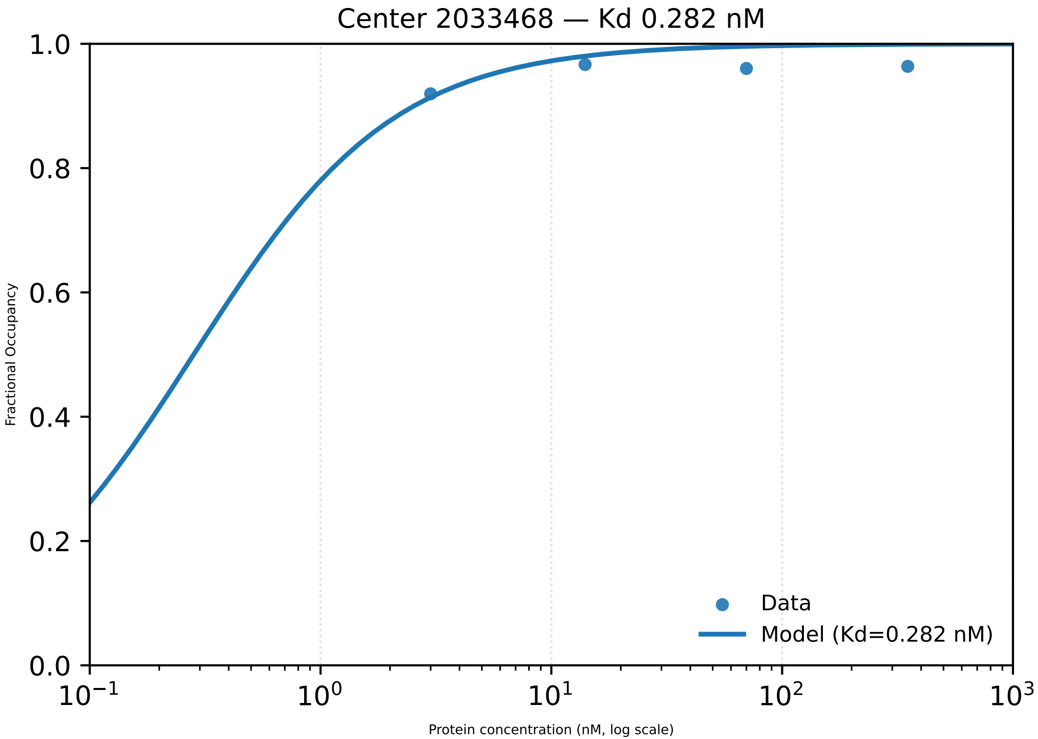

Center 2033468 — Kd 0.282 nM
Fractional Occupancy
0.0
0.2
0.4
0.6
0.8
1.0
Protein concentration (nM, log scale)
$10^{-1}$
$10^{0}$
$10^{1}$
$10^{2}$
$10^{3}$
Data
Model (Kd=0.282 nM)

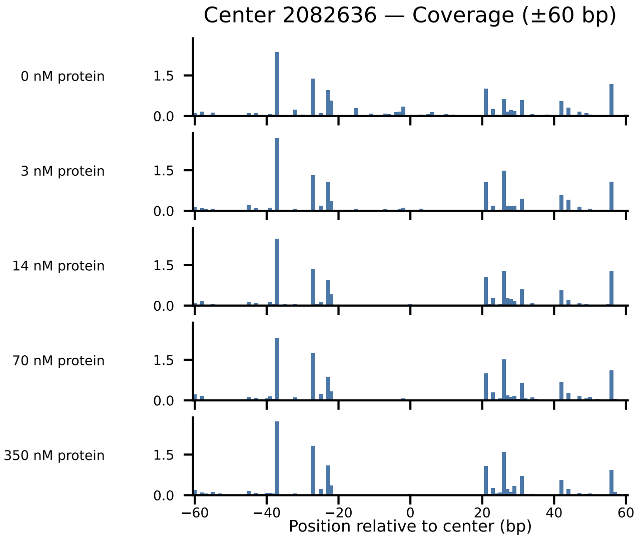

Center 2082636 — Coverage (±60 bp)
0 nM protein
3 nM protein
14 nM protein
70 nM protein
350 nM protein
1.5
0.0
−60
−40
−20
0
20
40
60
Position relative to center (bp)

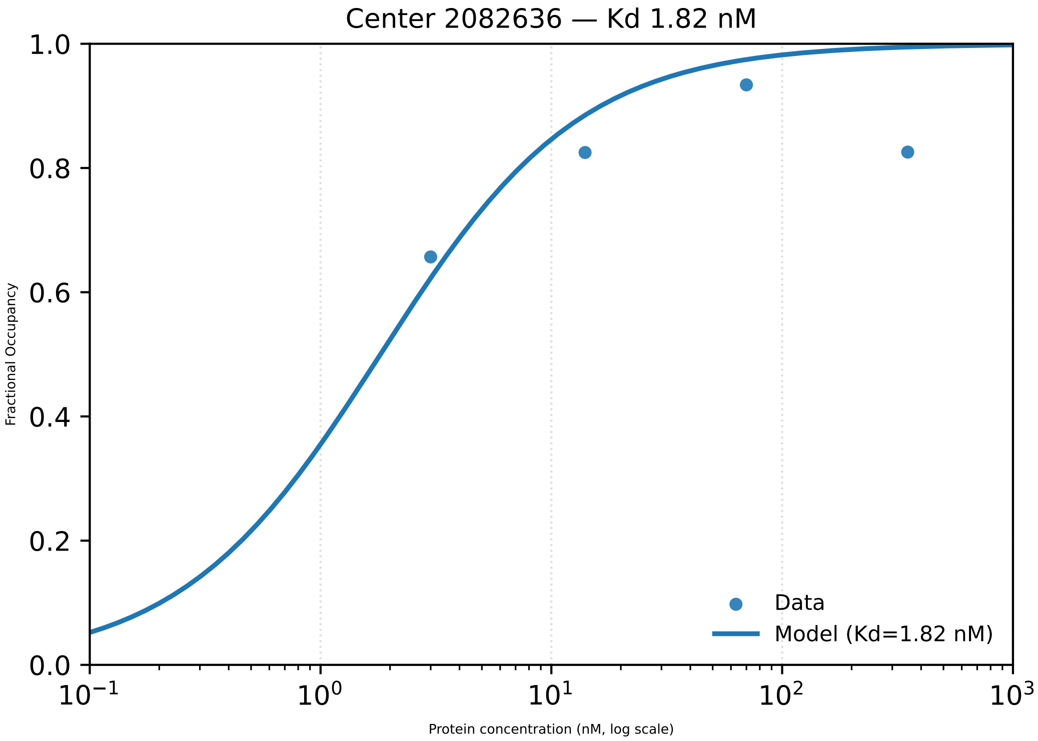
Center 2082636 — Kd 1.82 nM
Fractional Occupancy
Protein concentration (nM, log scale)
Data
Model (Kd=1.82 nM)
0.0
0.2
0.4
0.6
0.8
1.0
$10^{-1}$
$10^{0}$
$10^{1}$
$10^{2}$
$10^{3}$

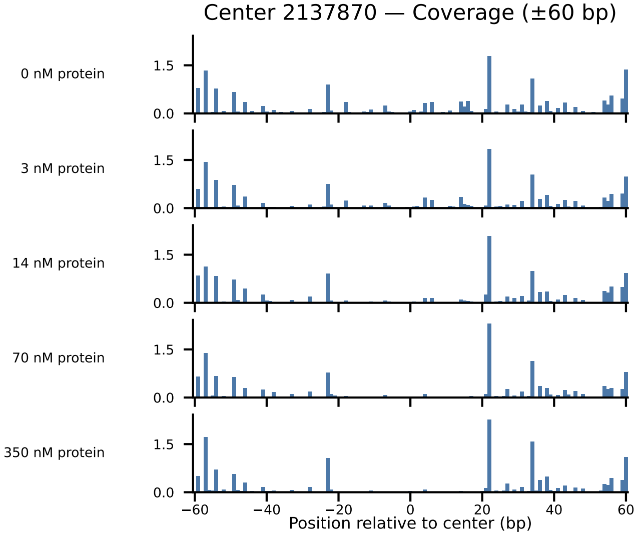

Center 2137870 — Coverage (±60 bp)
0 nM protein
3 nM protein
14 nM protein
70 nM protein
350 nM protein
1.5
0.0
−60
−40
−20
0
20
40
60
Position relative to center (bp)

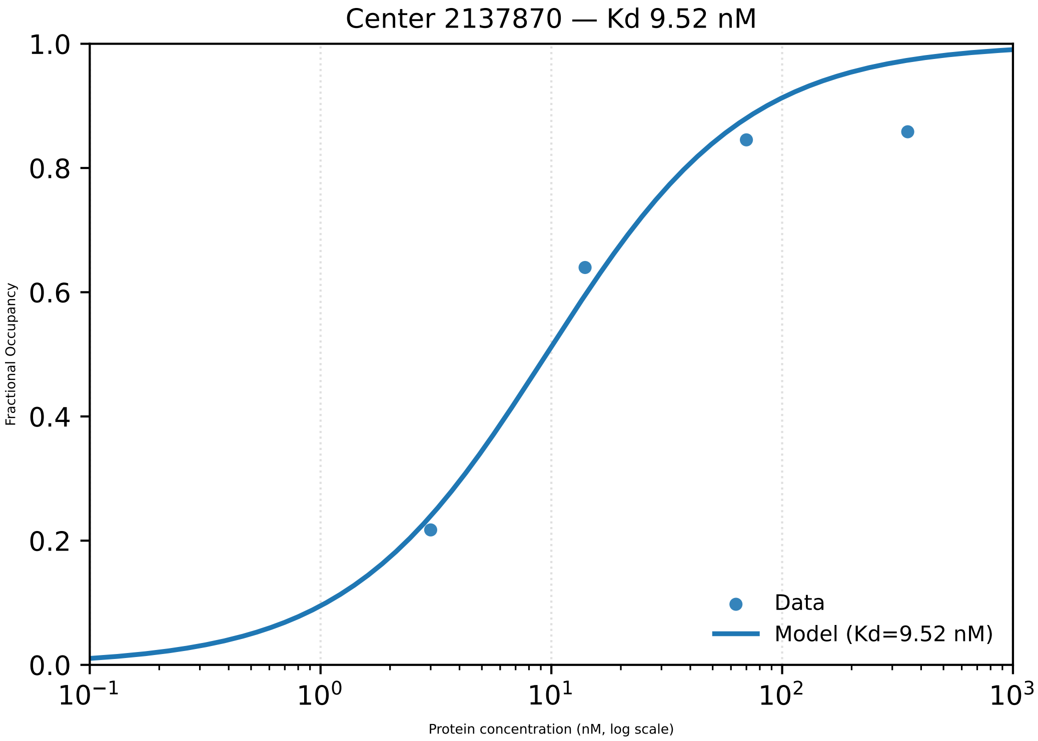

Center 2137870 — Kd 9.52 nM
Fractional Occupancy
1.0
0.8
0.6
0.4
0.2
0.0
$10^{-1}$
$10^{0}$
$10^{1}$
$10^{2}$
$10^{3}$
Protein concentration (nM, log scale)
Data
Model (Kd=9.52 nM)

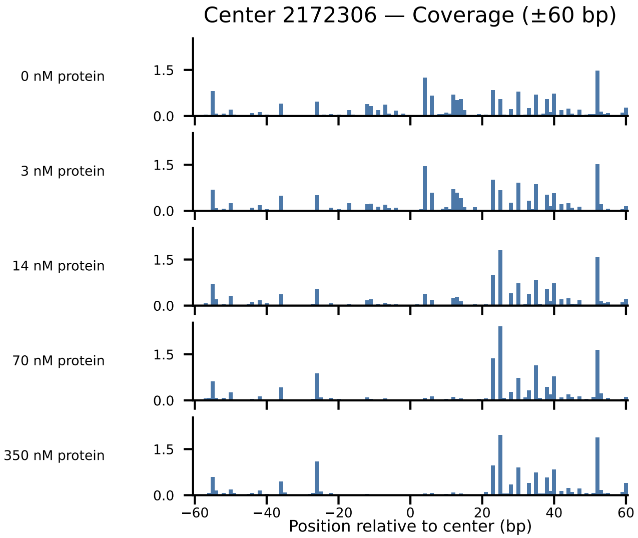

Center 2172306 — Coverage (±60 bp)
0 nM protein
3 nM protein
14 nM protein
70 nM protein
350 nM protein
1.5
0.0
−60
−40
−20
0
20
40
60
Position relative to center (bp)

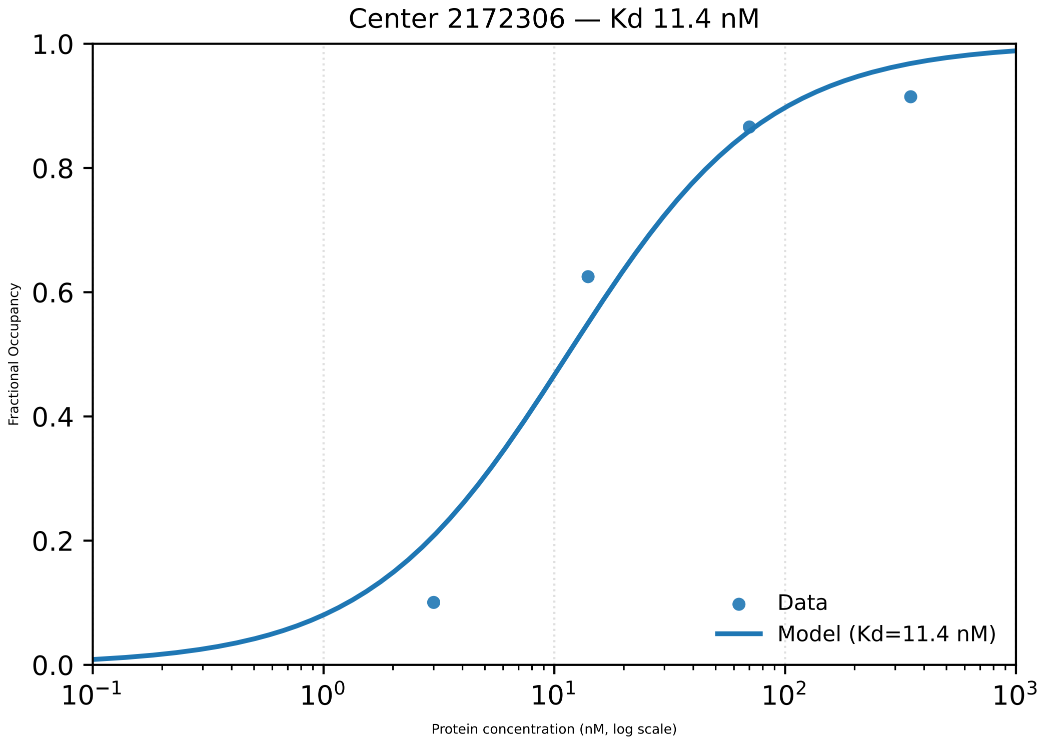
Center 2172306 — Kd 11.4 nM
1.0
0.8
0.6
0.4
0.2
0.0
Fractional Occupancy
$10^{-1}$
$10^{0}$
$10^{1}$
$10^{2}$
$10^{3}$
Protein concentration (nM, log scale)
Data
Model (Kd=11.4 nM)

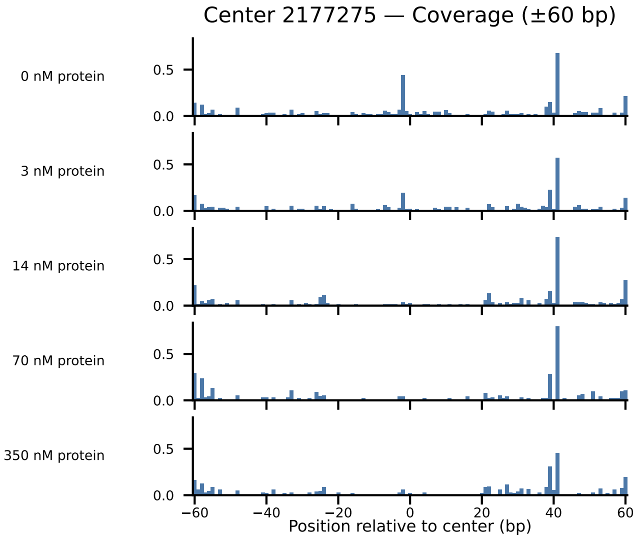

Center 2177275 — Coverage (±60 bp)
0 nM protein
3 nM protein
14 nM protein
70 nM protein
350 nM protein
0.5
0.0
−60
−40
−20
0
20
40
60
Position relative to center (bp)

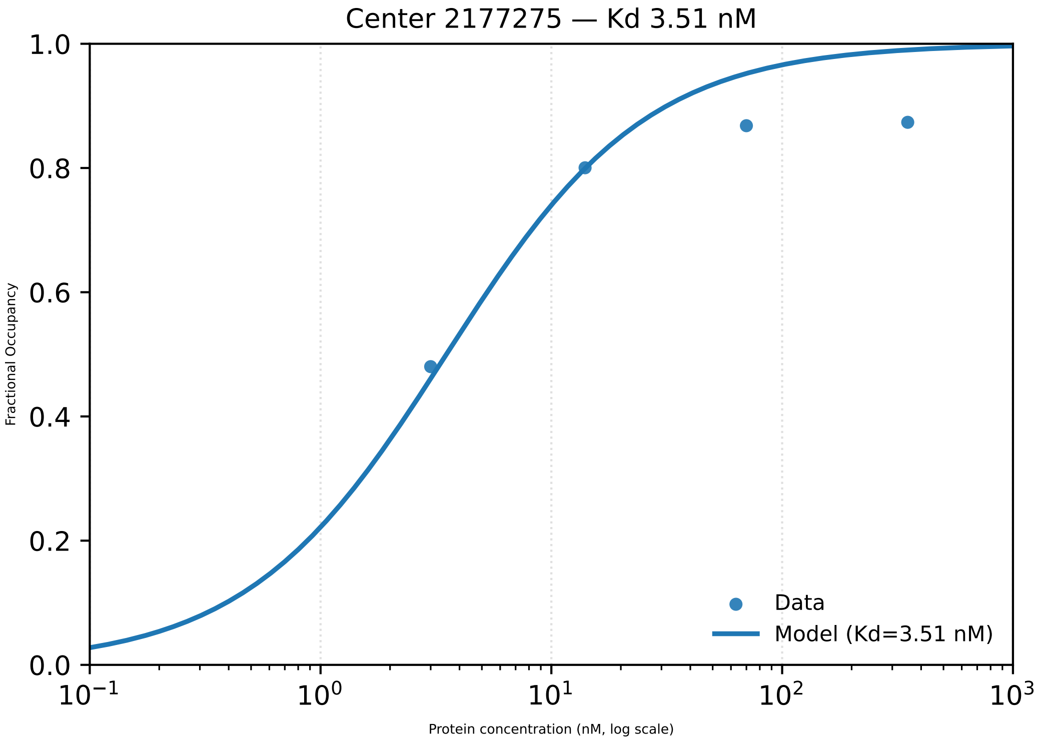

Center 2177275 — Kd 3.51 nM
1.0
0.8
0.6
0.4
0.2
0.0
Fractional Occupancy
$10^{-1}$
$10^{0}$
$10^{1}$
$10^{2}$
$10^{3}$
Protein concentration (nM, log scale)
Data
Model (Kd=3.51 nM)

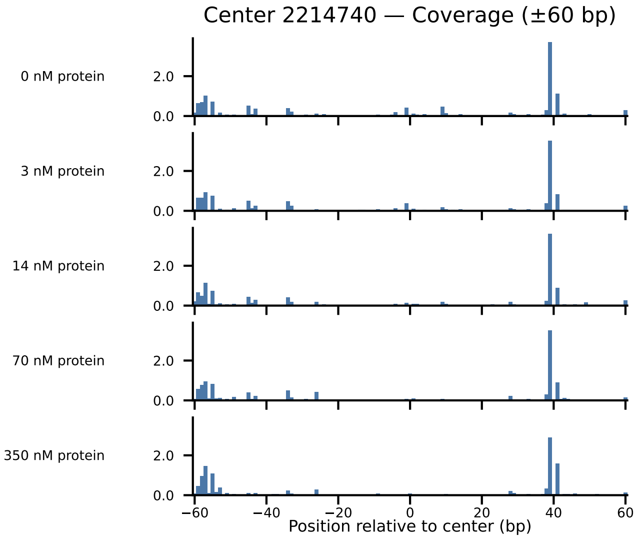

Center 2214740 — Coverage (±60 bp)
0 nM protein
3 nM protein
14 nM protein
70 nM protein
350 nM protein
2.0
0.0
−60
−40
−20
0
20
40
60
Position relative to center (bp)

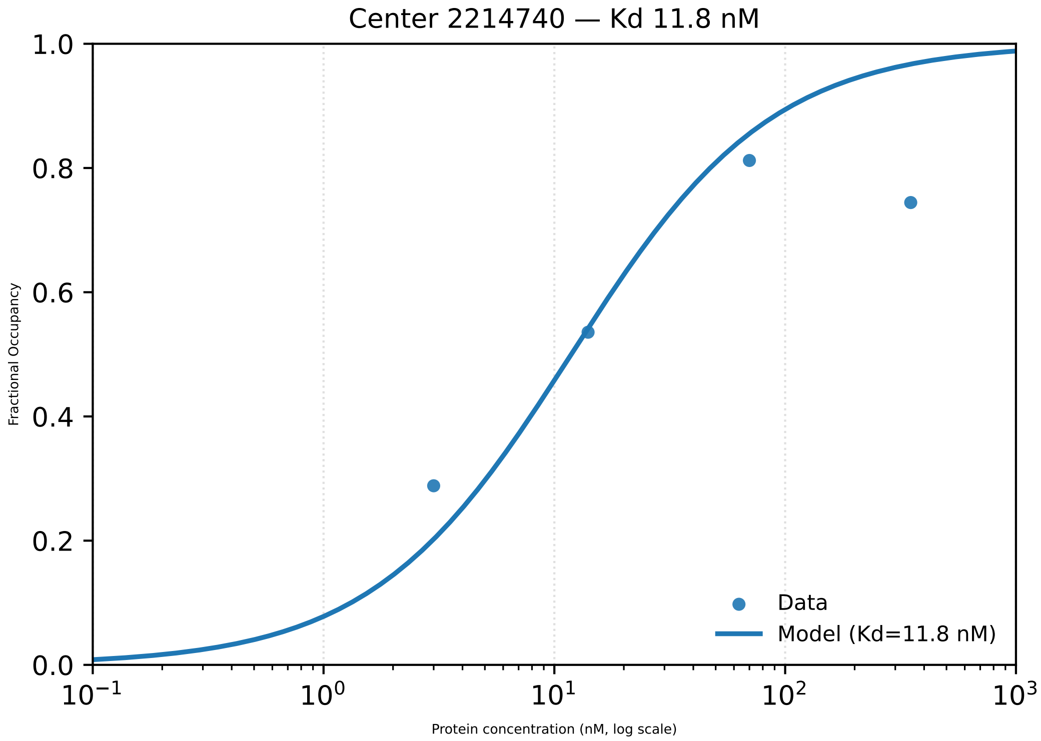

Center 2214740 — Kd 11.8 nM
1.0
0.8
0.6
0.4
0.2
0.0
Fractional Occupancy
$10^{-1}$
$10^{0}$
$10^{1}$
$10^{2}$
$10^{3}$
Protein concentration (nM, log scale)
Data
Model (Kd=11.8 nM)

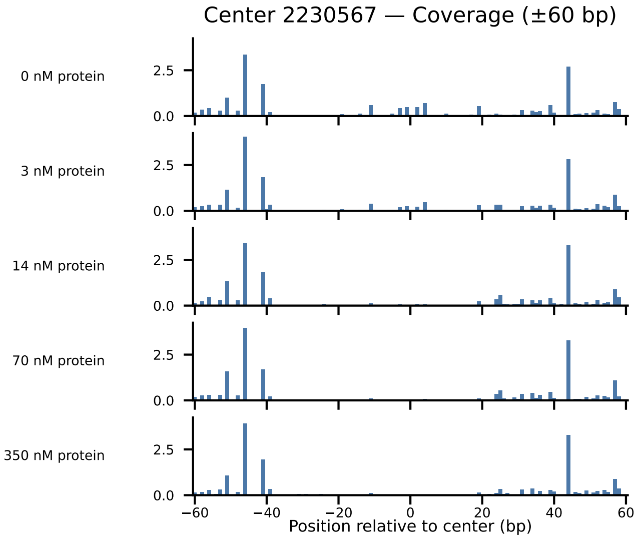

Center 2230567 — Coverage (±60 bp)
0 nM protein
3 nM protein
14 nM protein
70 nM protein
350 nM protein
2.5
0.0
−60
−40
−20
0
20
40
60
Position relative to center (bp)

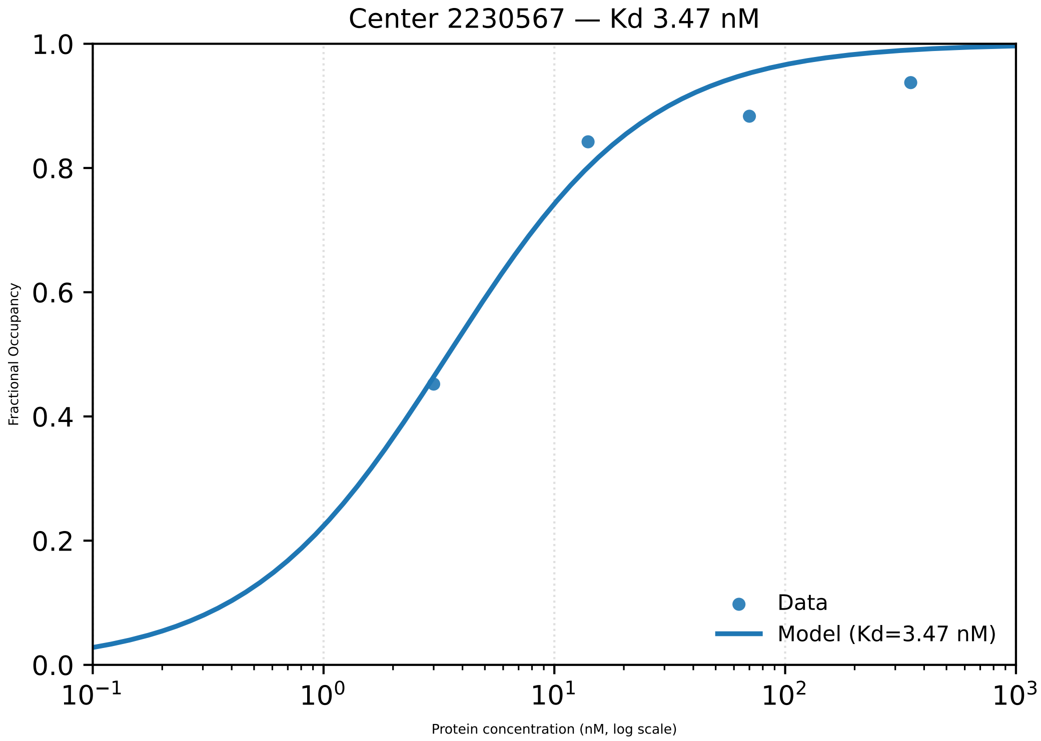
Center 2230567 — Kd 3.47 nM
Fractional Occupancy
1.0
0.8
0.6
0.4
0.2
0.0
$10^{-1}$
$10^{0}$
$10^{1}$
$10^{2}$
$10^{3}$
Protein concentration (nM, log scale)
Data
Model (Kd=3.47 nM)

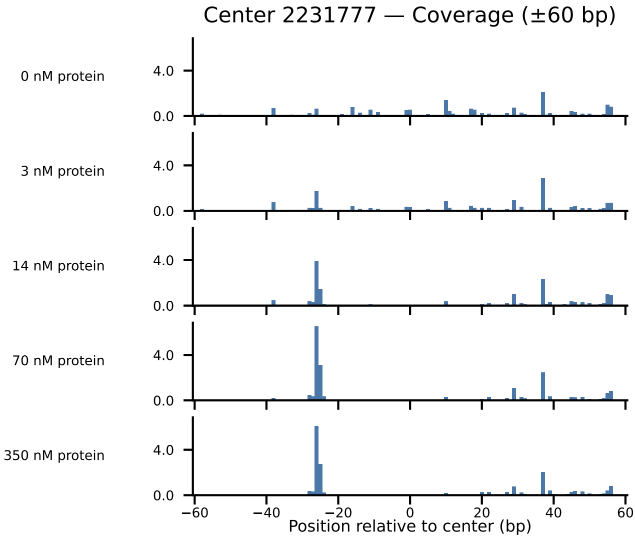
Center 2231777 — Coverage (±60 bp)
0 nM protein
3 nM protein
14 nM protein
70 nM protein
350 nM protein
4.0
0.0
−60
−40
−20
0
20
40
60
Position relative to center (bp)

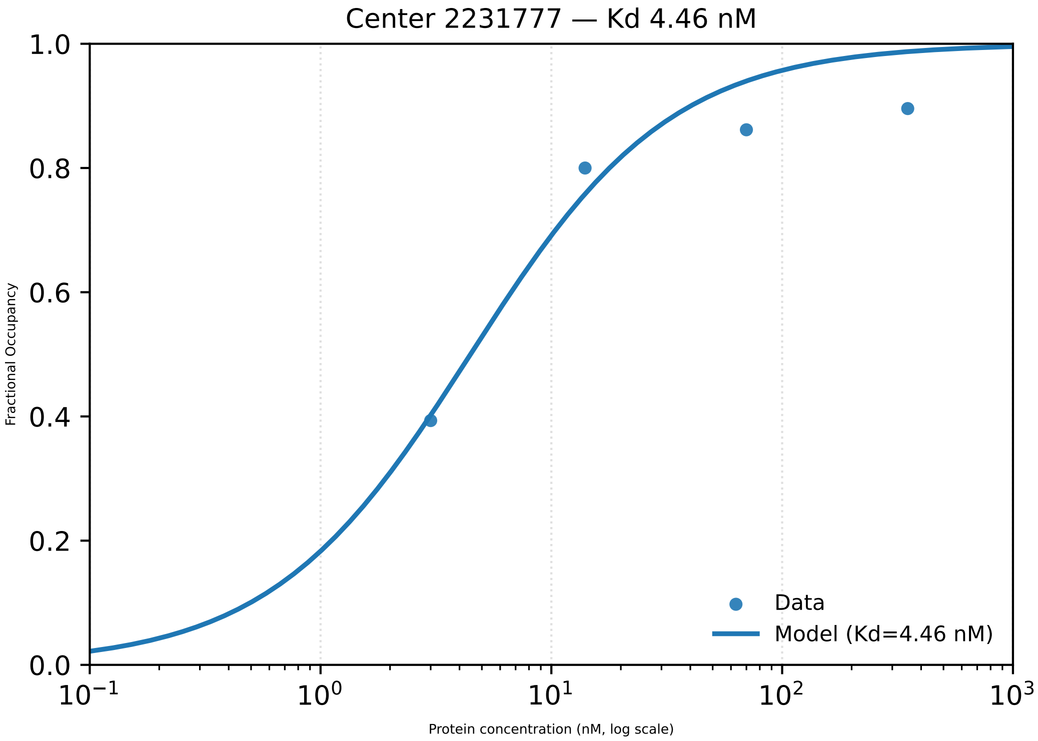

Center 2231777 — Kd 4.46 nM
1.0
0.8
0.6
0.4
0.2
0.0
Fractional Occupancy
$10^{-1}$
$10^{0}$
$10^{1}$
$10^{2}$
$10^{3}$
Protein concentration (nM, log scale)
Data
Model (Kd=4.46 nM)

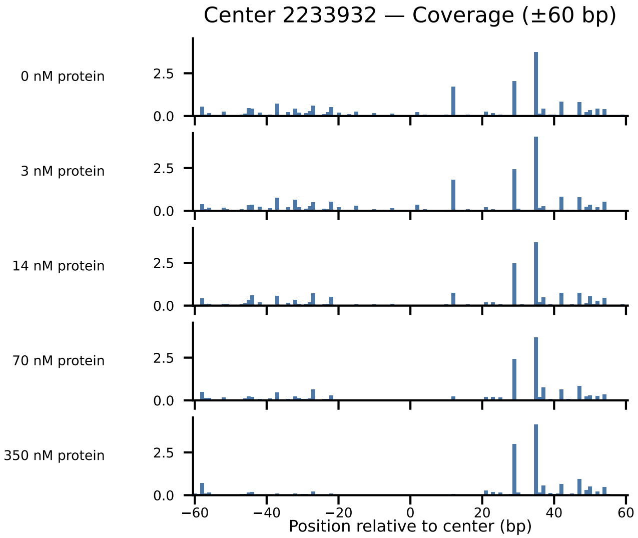

Center 2233932 — Coverage (±60 bp)
0 nM protein
3 nM protein
14 nM protein
70 nM protein
350 nM protein
2.5
0.0
−60
−40
−20
0
20
40
60
Position relative to center (bp)

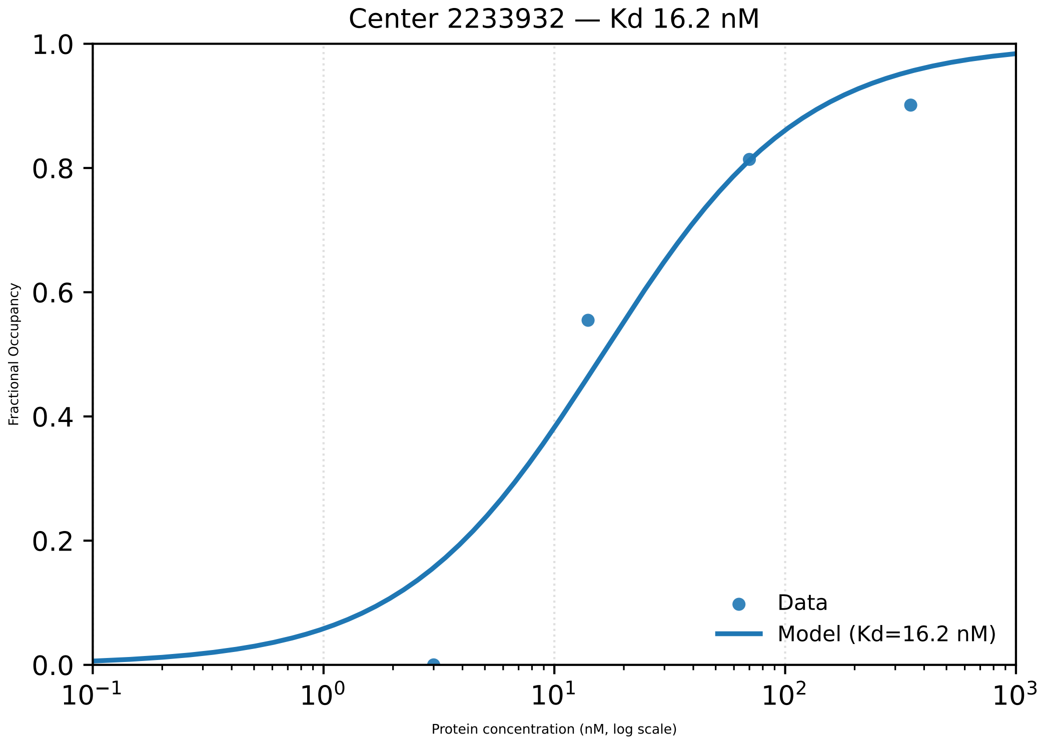

Center 2233932 — Kd 16.2 nM
Fractional Occupancy
Protein concentration (nM, log scale)
Data
Model (Kd=16.2 nM)
0.0
0.2
0.4
0.6
0.8
1.0
$10^{-1}$
$10^{0}$
$10^{1}$
$10^{2}$
$10^{3}$

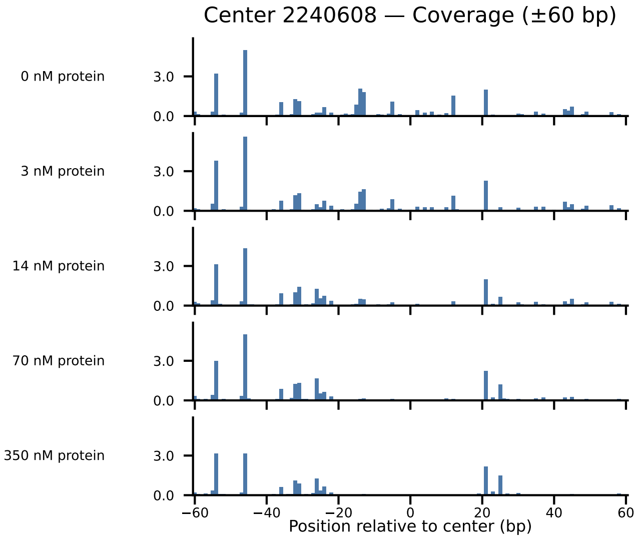
Center 2240608 — Coverage (±60 bp)
0 nM protein
3 nM protein
14 nM protein
70 nM protein
350 nM protein
3.0
0.0
−60
−40
−20
0
20
40
60
Position relative to center (bp)

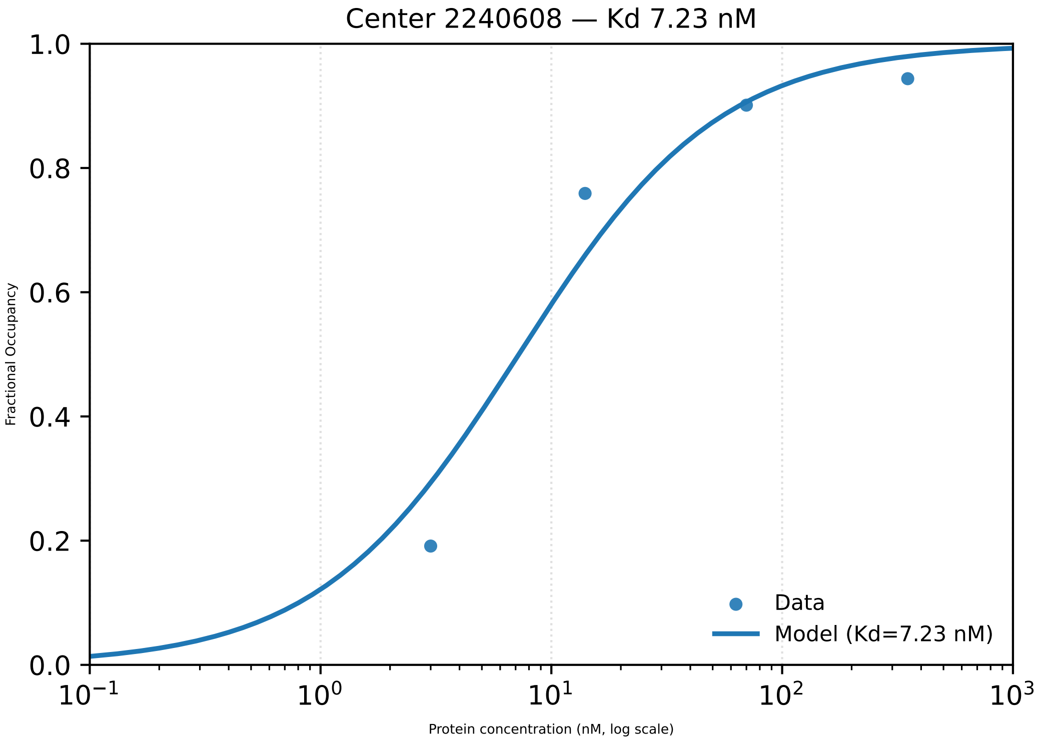

Center 2240608 — Kd 7.23 nM
1.0
0.8
0.6
0.4
0.2
0.0
Fractional Occupancy
$10^{-1}$
$10^{0}$
$10^{1}$
$10^{2}$
$10^{3}$
Protein concentration (nM, log scale)
Data
Model (Kd=7.23 nM)

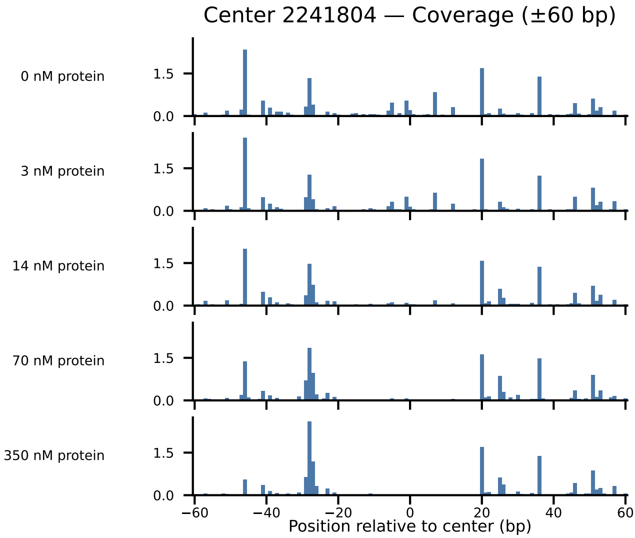
Center 2241804 — Coverage (±60 bp)
0 nM protein
3 nM protein
14 nM protein
70 nM protein
350 nM protein
1.5
0.0
−60
−40
−20
0
20
40
60
Position relative to center (bp)

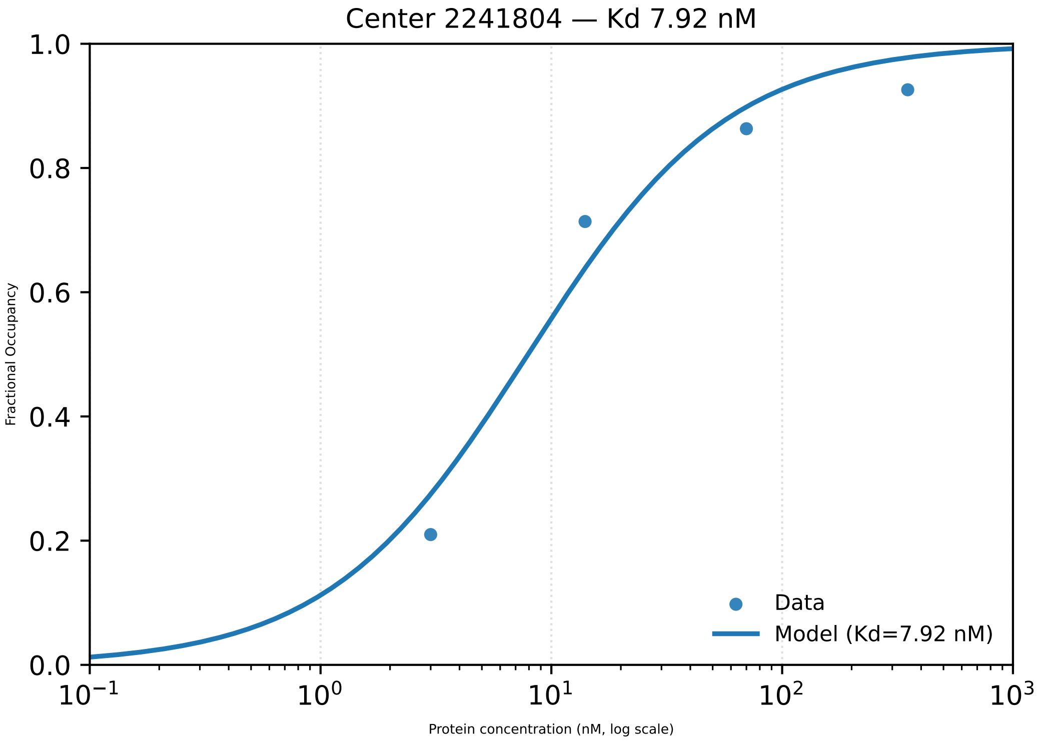

Center 2241804 — Kd 7.92 nM
1.0
0.8
0.6
0.4
0.2
0.0
Fractional Occupancy
$10^{-1}$
$10^{0}$
$10^{1}$
$10^{2}$
$10^{3}$
Protein concentration (nM, log scale)
Data
Model (Kd=7.92 nM)

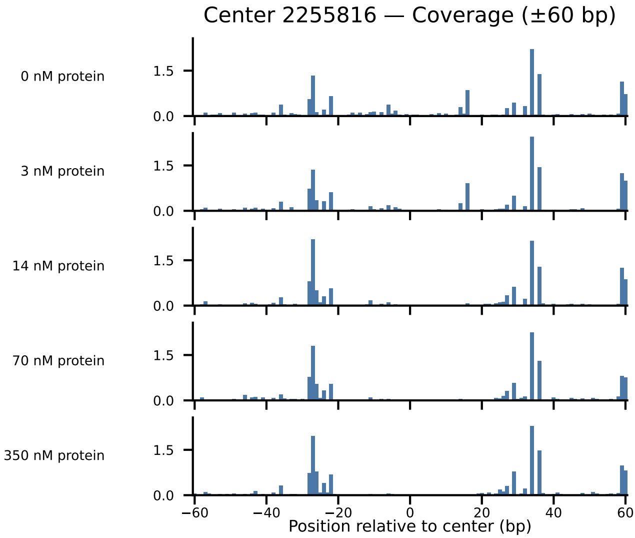
Center 2255816 — Coverage (±60 bp)
0 nM protein
3 nM protein
14 nM protein
70 nM protein
350 nM protein
1.5
0.0
−60
−40
−20
0
20
40
60
Position relative to center (bp)

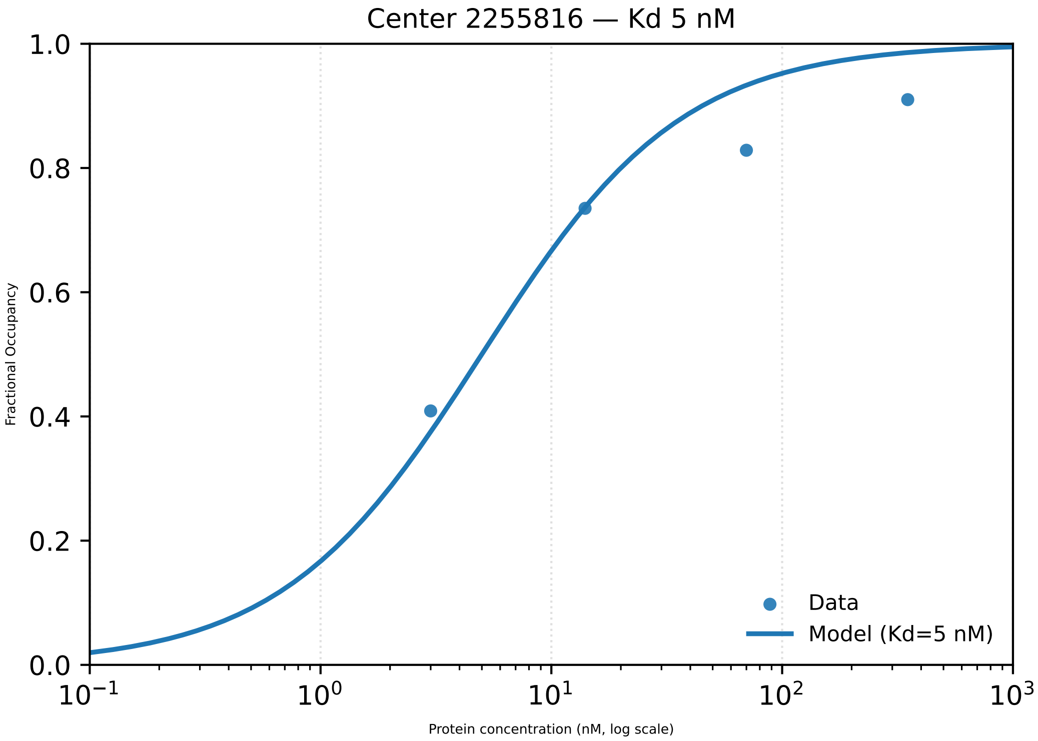

Center 2255816 — Kd 5 nM
1.0
0.8
0.6
0.4
0.2
0.0
Fractional Occupancy
$10^{-1}$
$10^{0}$
$10^{1}$
$10^{2}$
$10^{3}$
Protein concentration (nM, log scale)
Data
Model (Kd=5 nM)

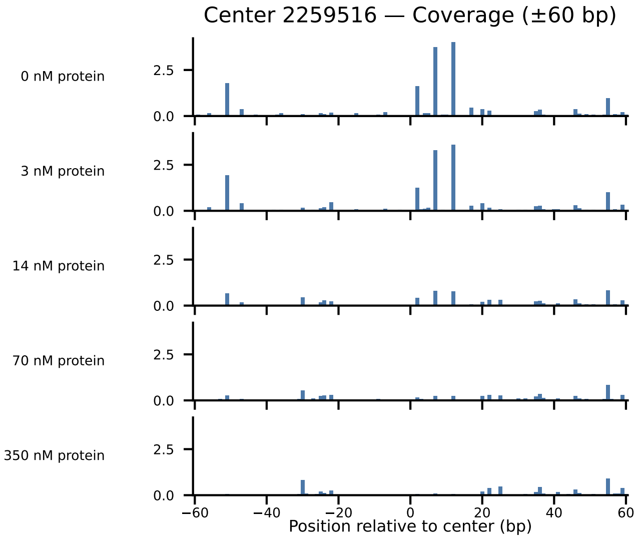

Center 2259516 — Coverage (±60 bp)
0 nM protein
3 nM protein
14 nM protein
70 nM protein
350 nM protein
2.5
0.0
−60
−40
−20
0
20
40
60
Position relative to center (bp)

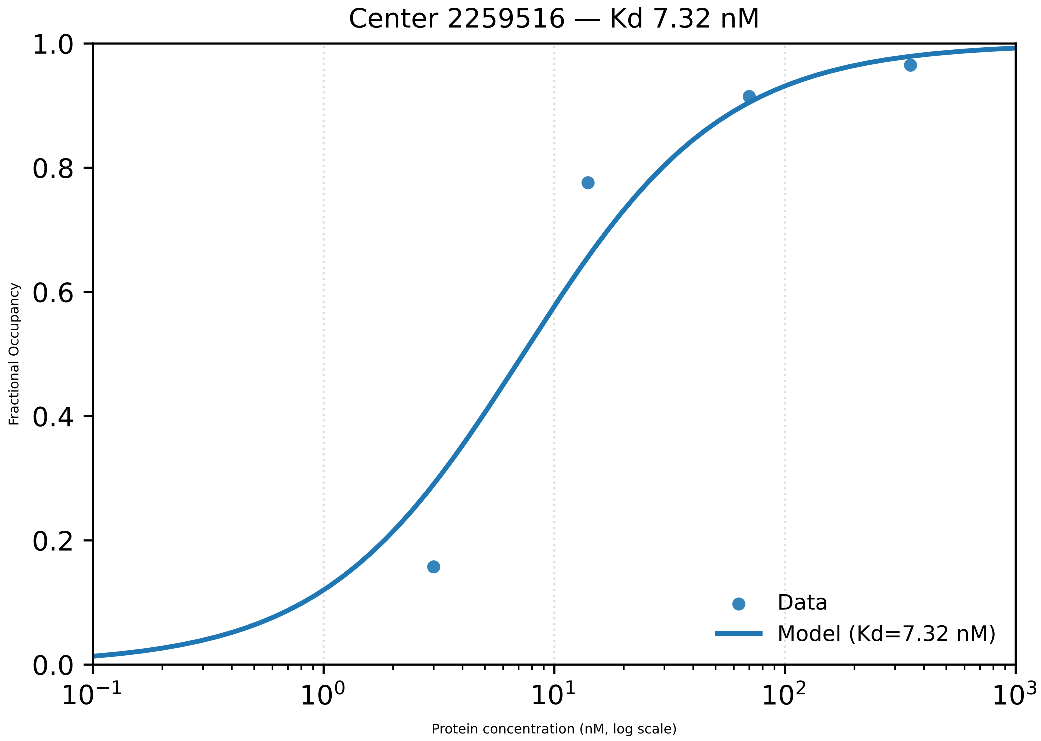

Center 2259516 — Kd 7.32 nM
1.0
0.8
0.6
0.4
0.2
0.0
Fractional Occupancy
$10^{-1}$
$10^{0}$
$10^{1}$
$10^{2}$
$10^{3}$
Protein concentration (nM, log scale)
Data
Model (Kd=7.32 nM)

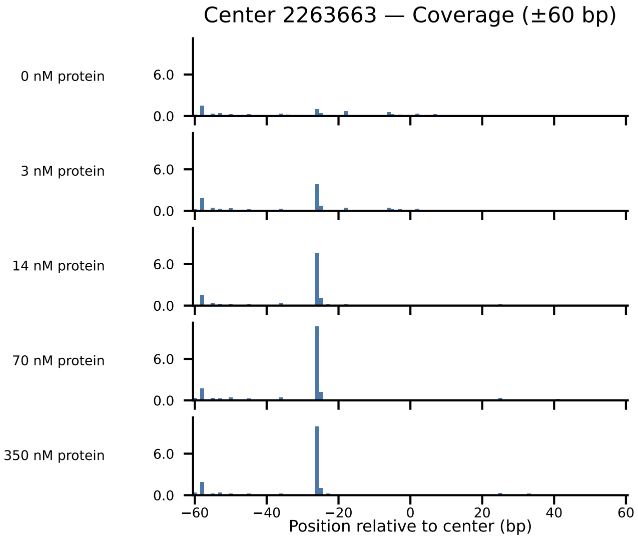

Center 2263663 — Coverage (±60 bp)
0 nM protein
3 nM protein
14 nM protein
70 nM protein
350 nM protein
6.0
0.0
−60
−40
−20
0
20
40
60
Position relative to center (bp)

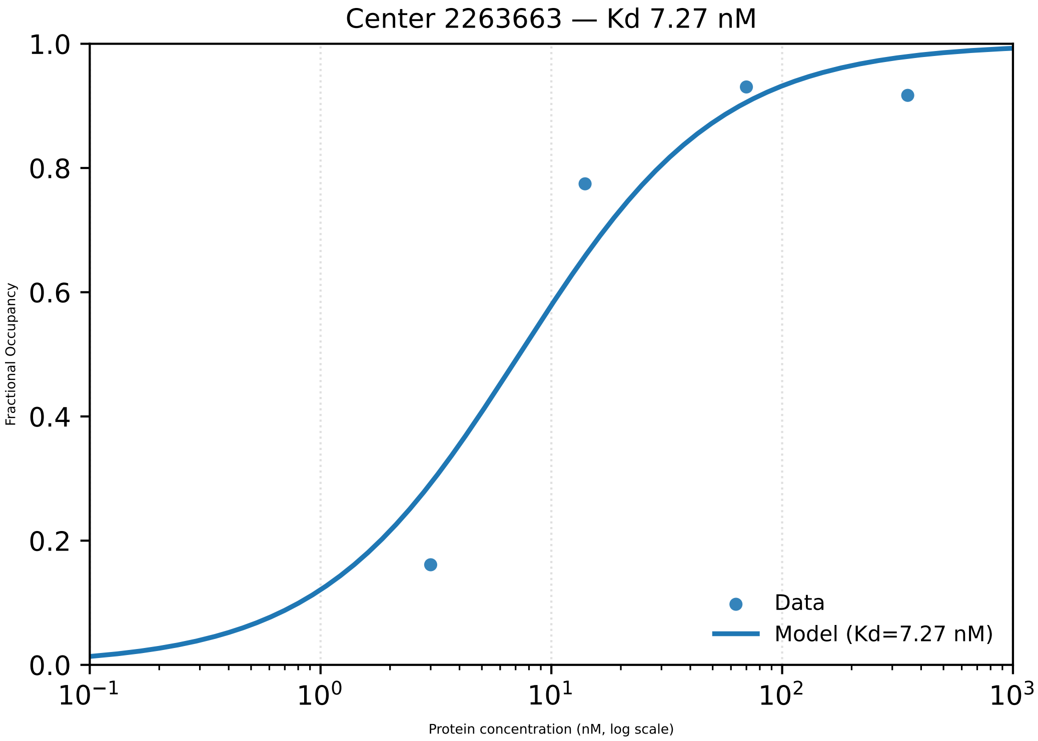

Center 2263663 — Kd 7.27 nM
1.0
0.8
0.6
0.4
0.2
0.0
Fractional Occupancy
$10^{-1}$
$10^{0}$
$10^{1}$
$10^{2}$
$10^{3}$
Protein concentration (nM, log scale)
Data
Model (Kd=7.27 nM)

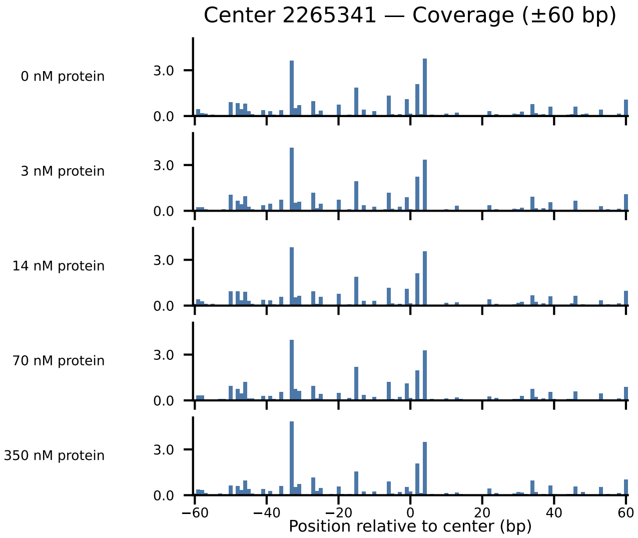

Center 2265341 — Coverage (±60 bp)
0 nM protein
3 nM protein
14 nM protein
70 nM protein
350 nM protein
3.0
0.0
−60
−40
−20
0
20
40
60
Position relative to center (bp)

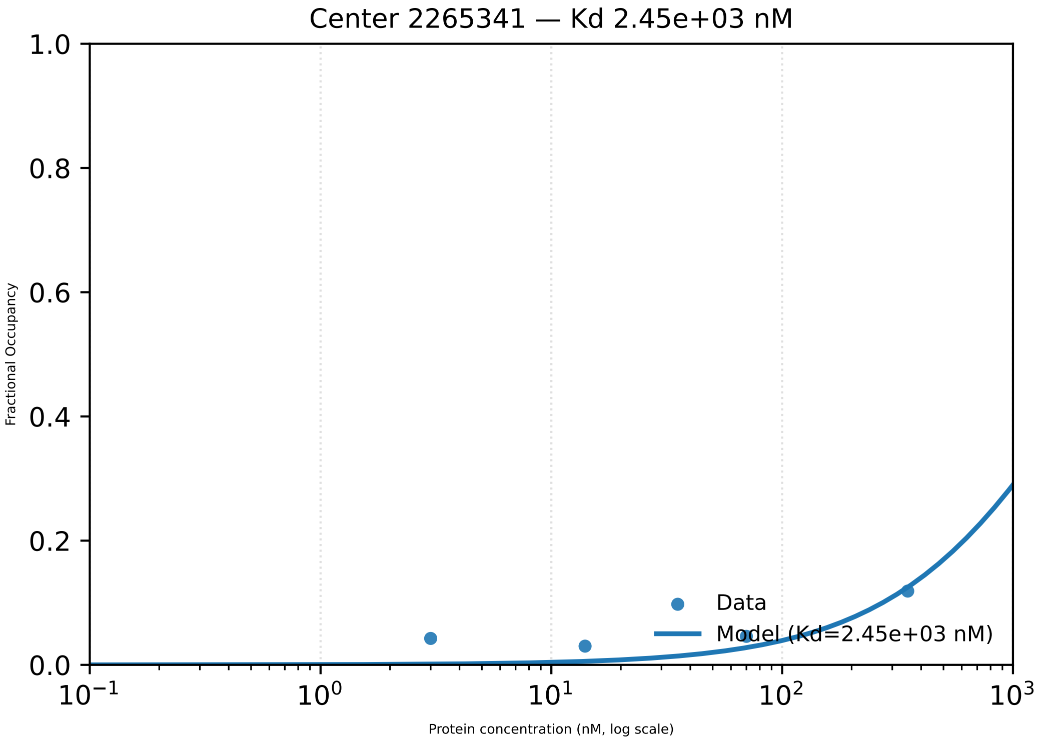

Center 2265341 — Kd 2.45e+03 nM
1.0
0.8
0.6
0.4
0.2
0.0
Fractional Occupancy
$10^{-1}$
$10^{0}$
$10^{1}$
$10^{2}$
$10^{3}$
Protein concentration (nM, log scale)
Data
Model (Kd=2.45e+03 nM)

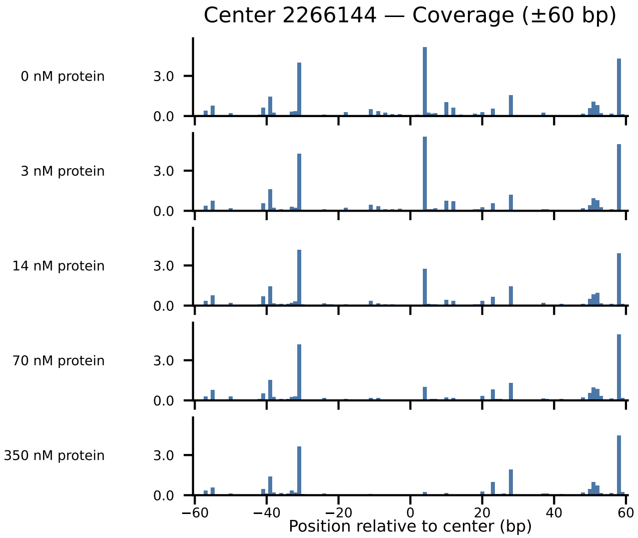

Center 2266144 — Coverage (±60 bp)
0 nM protein
3 nM protein
14 nM protein
70 nM protein
350 nM protein
3.0
0.0
−60
−40
−20
0
20
40
60
Position relative to center (bp)

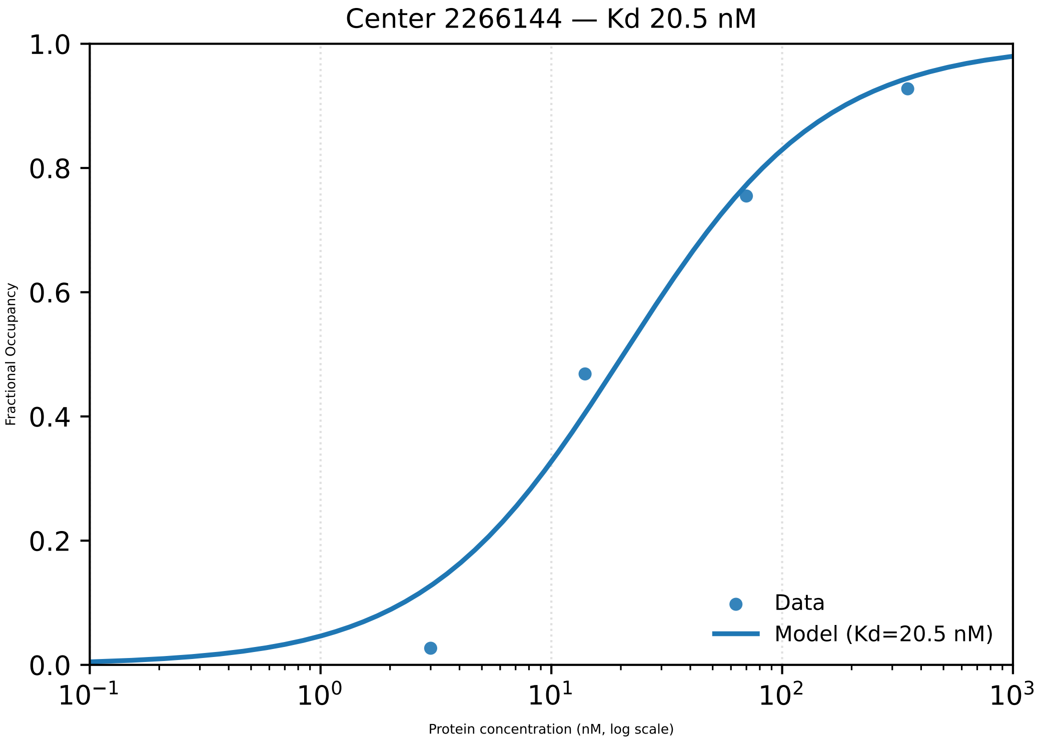

Center 2266144 — Kd 20.5 nM
1.0
0.8
0.6
0.4
0.2
0.0
Fractional Occupancy
$10^{-1}$
$10^{0}$
$10^{1}$
$10^{2}$
$10^{3}$
Protein concentration (nM, log scale)
Data
Model (Kd=20.5 nM)

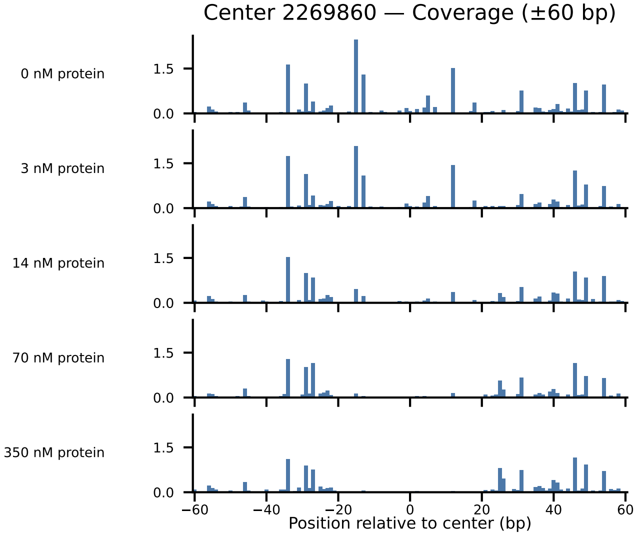

Center 2269860 — Coverage (±60 bp)
0 nM protein
3 nM protein
14 nM protein
70 nM protein
350 nM protein
1.5
0.0
−60
−40
−20
0
20
40
60
Position relative to center (bp)

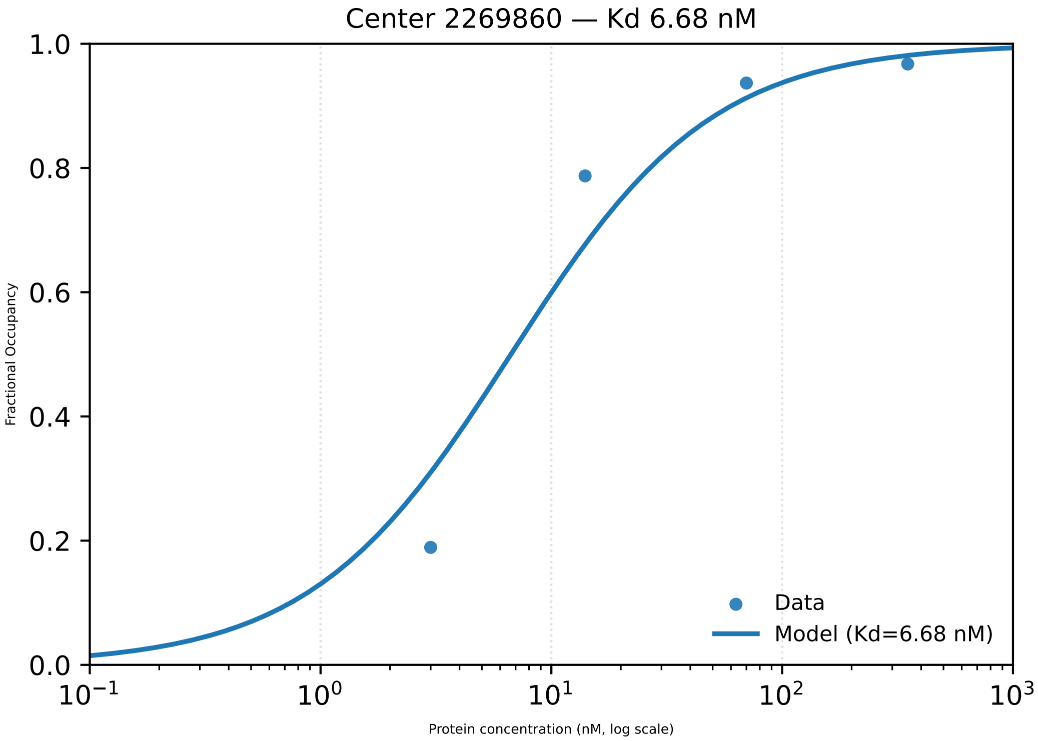

Center 2269860 — Kd 6.68 nM
1.0
0.8
0.6
0.4
0.2
0.0
Fractional Occupancy
$10^{-1}$
$10^{0}$
$10^{1}$
$10^{2}$
$10^{3}$
Protein concentration (nM, log scale)
Data
Model (Kd=6.68 nM)

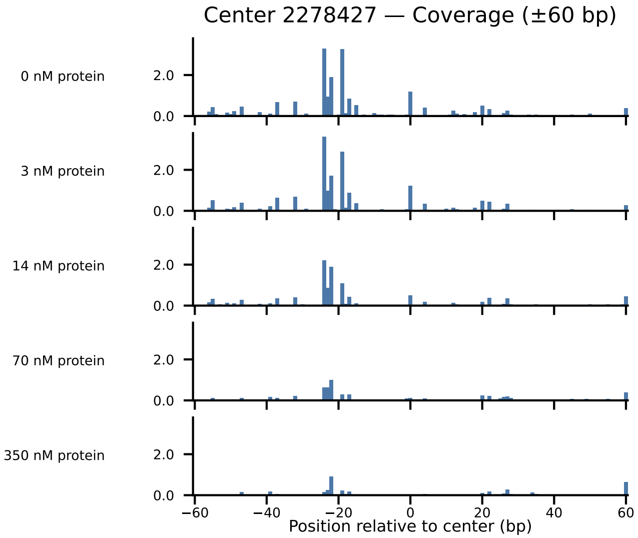

Center 2278427 — Coverage (±60 bp)
0 nM protein
3 nM protein
14 nM protein
70 nM protein
350 nM protein
2.0
0.0
−60
−40
−20
0
20
40
60
Position relative to center (bp)

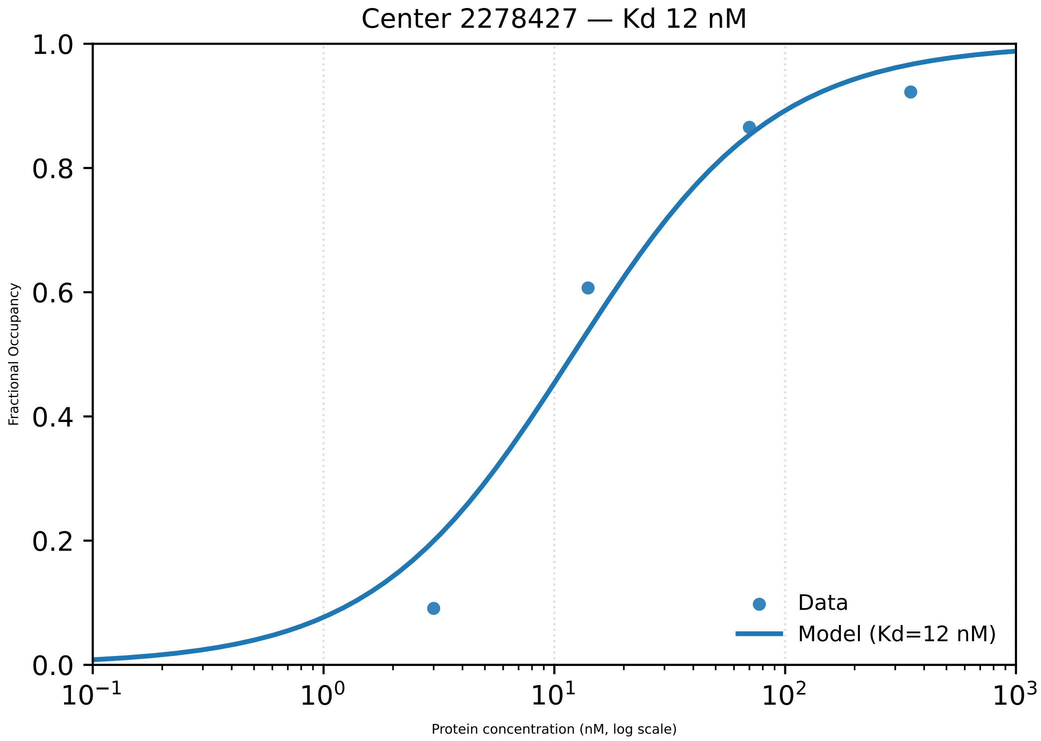

Center 2278427 — Kd 12 nM
Fractional Occupancy
1.0
0.8
0.6
0.4
0.2
0.0
$10^{-1}$
$10^{0}$
$10^{1}$
$10^{2}$
$10^{3}$
Protein concentration (nM, log scale)
Data
Model (Kd=12 nM)

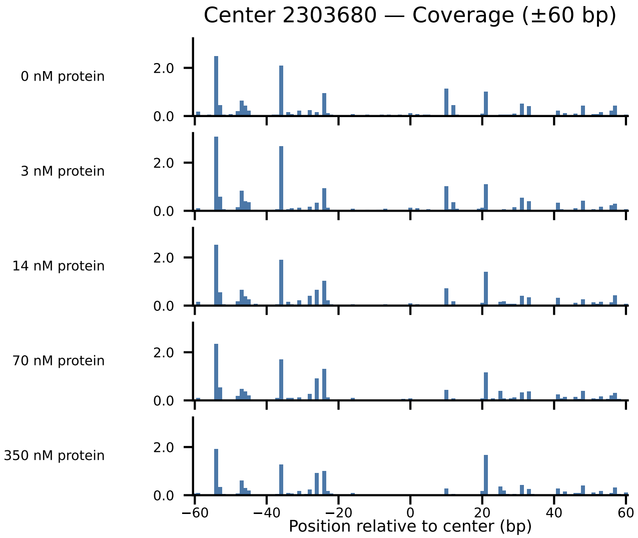

Center 2303680 — Coverage (±60 bp)
0 nM protein
3 nM protein
14 nM protein
70 nM protein
350 nM protein
2.0
0.0
−60
−40
−20
0
20
40
60
Position relative to center (bp)

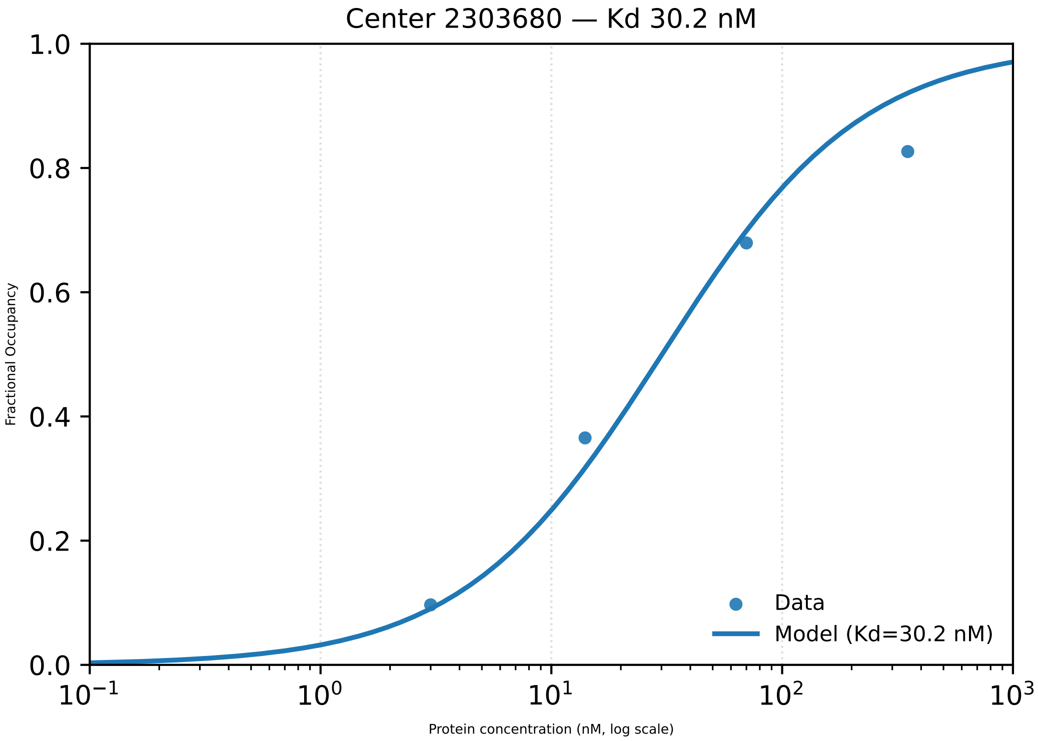
Center 2303680 — Kd 30.2 nM
Fractional Occupancy
Protein concentration (nM, log scale)
Data
Model (Kd=30.2 nM)
1.0
0.8
0.6
0.4
0.2
0.0
$10^{-1}$
$10^{0}$
$10^{1}$
$10^{2}$
$10^{3}$

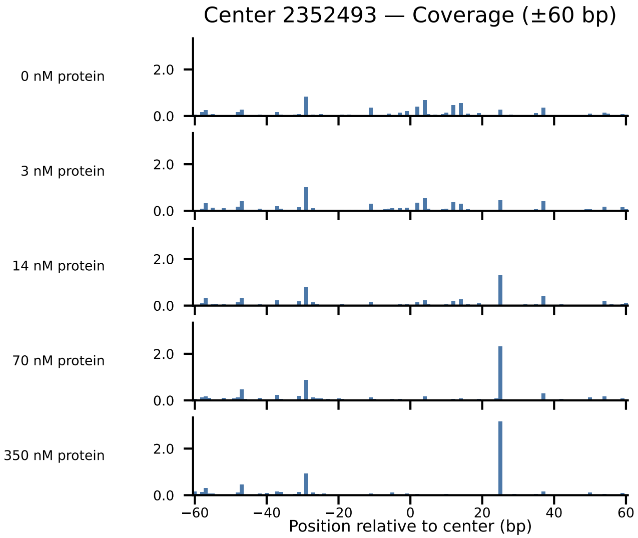

Center 2352493 — Coverage (±60 bp)
0 nM protein
3 nM protein
14 nM protein
70 nM protein
350 nM protein
2.0
0.0
−60
−40
−20
0
20
40
60
Position relative to center (bp)

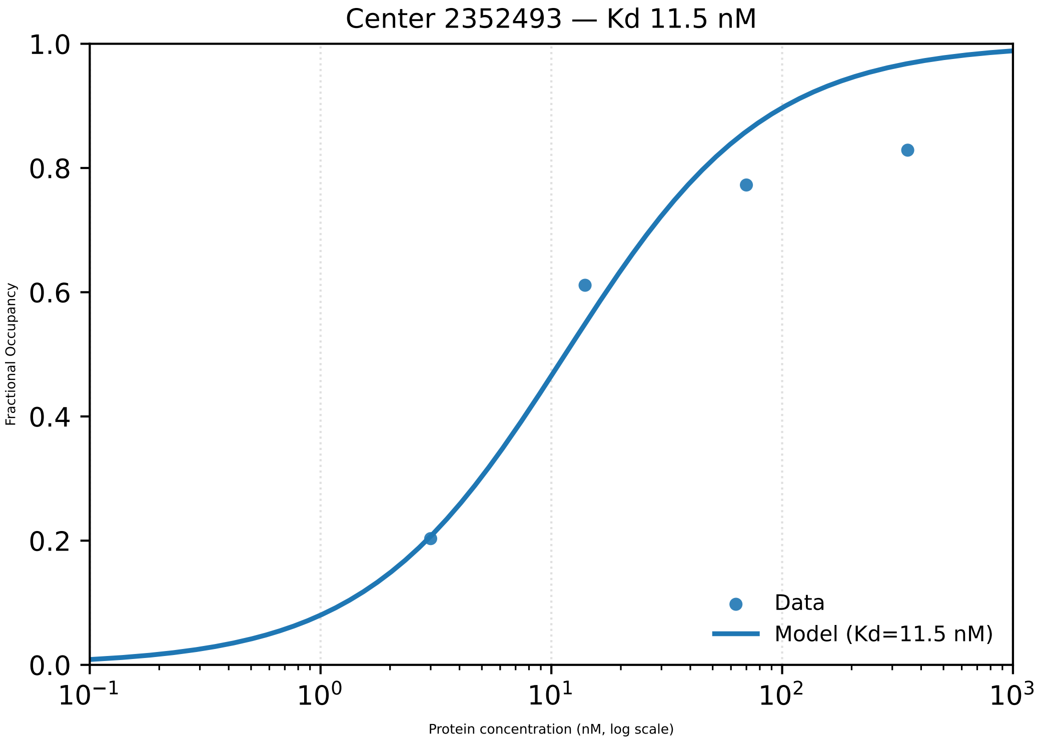

Center 2352493 — Kd 11.5 nM
1.0
0.8
0.6
0.4
0.2
0.0
Fractional Occupancy
$10^{-1}$
$10^{0}$
$10^{1}$
$10^{2}$
$10^{3}$
Protein concentration (nM, log scale)
Data
Model (Kd=11.5 nM)

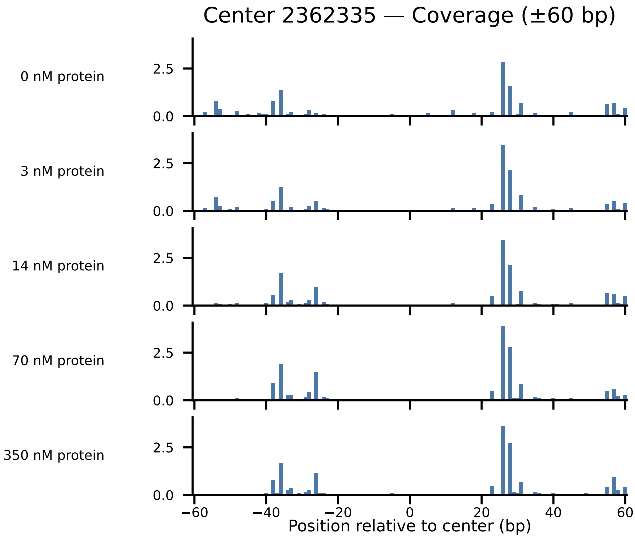

Center 2362335 — Coverage (±60 bp)
0 nM protein
3 nM protein
14 nM protein
70 nM protein
350 nM protein
2.5
0.0
−60
−40
−20
0
20
40
60
Position relative to center (bp)

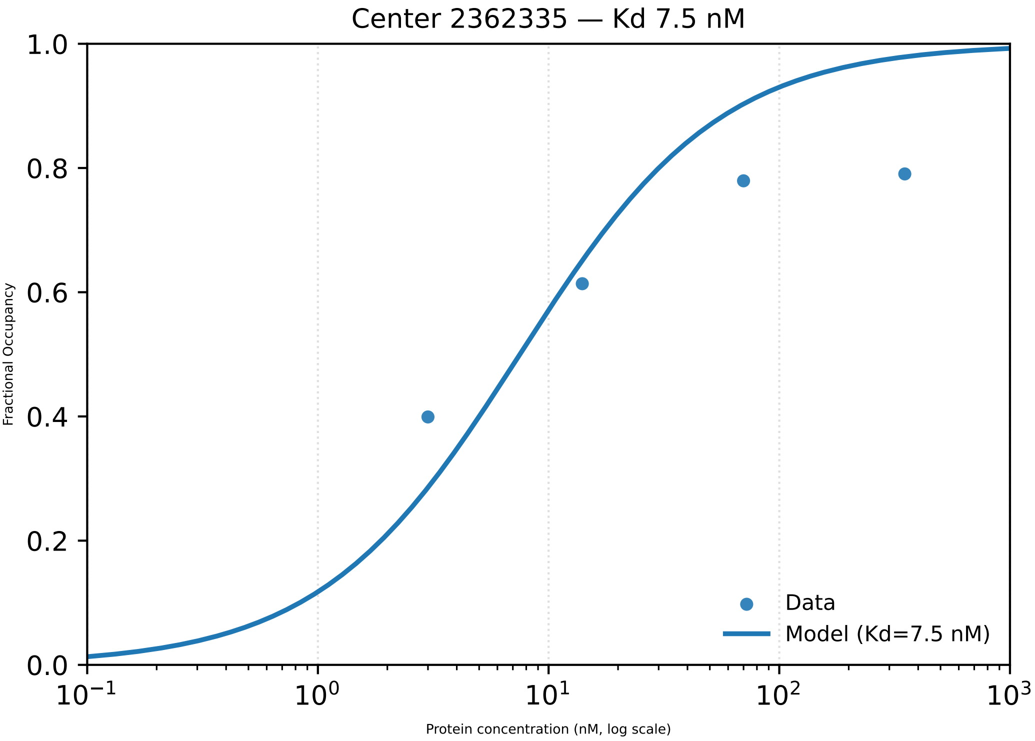

Center 2362335 — Kd 7.5 nM
1.0
0.8
0.6
0.4
0.2
0.0
Fractional Occupancy
$10^{-1}$
$10^{0}$
$10^{1}$
$10^{2}$
$10^{3}$
Protein concentration (nM, log scale)
Data
Model (Kd=7.5 nM)

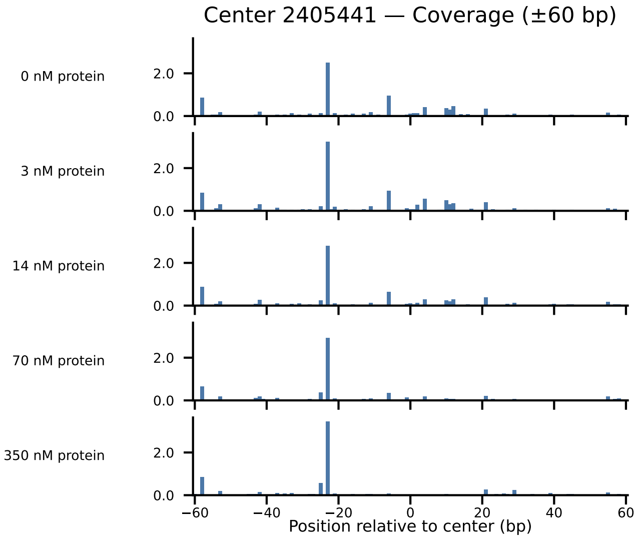

Center 2405441 — Coverage (±60 bp)
0 nM protein
3 nM protein
14 nM protein
70 nM protein
350 nM protein
2.0
0.0
−60
−40
−20
0
20
40
60
Position relative to center (bp)

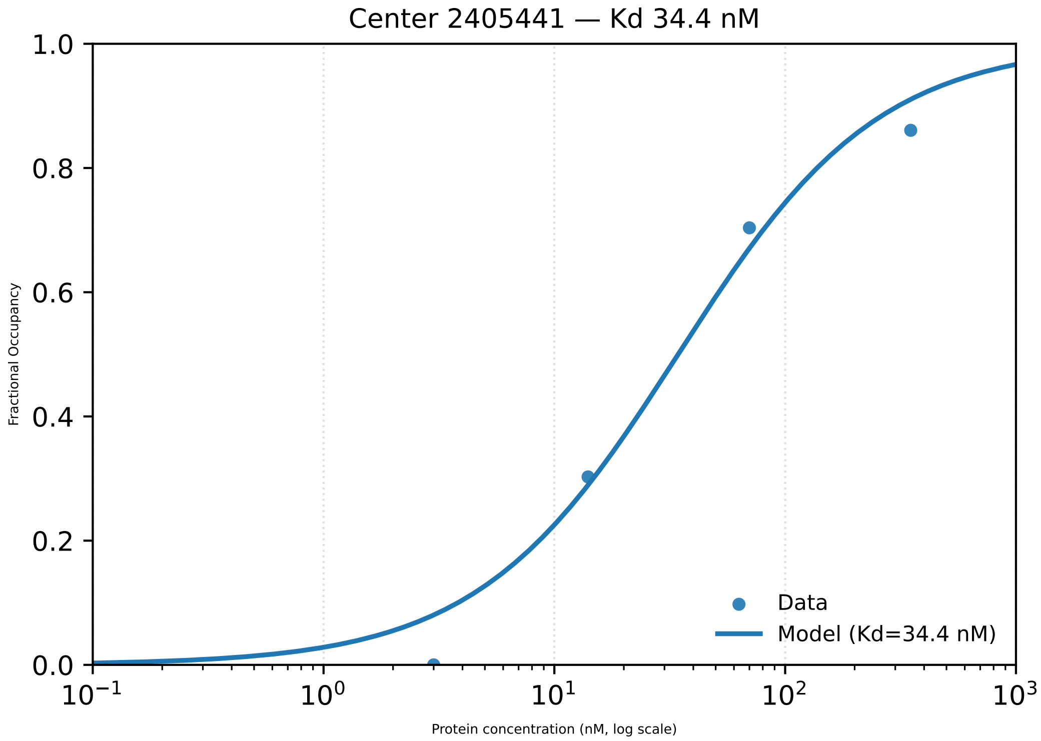

Center 2405441 — Kd 34.4 nM
Fractional Occupancy
1.0
0.8
0.6
0.4
0.2
0.0
$10^{-1}$
$10^{0}$
$10^{1}$
$10^{2}$
$10^{3}$
Protein concentration (nM, log scale)
Data
Model (Kd=34.4 nM)

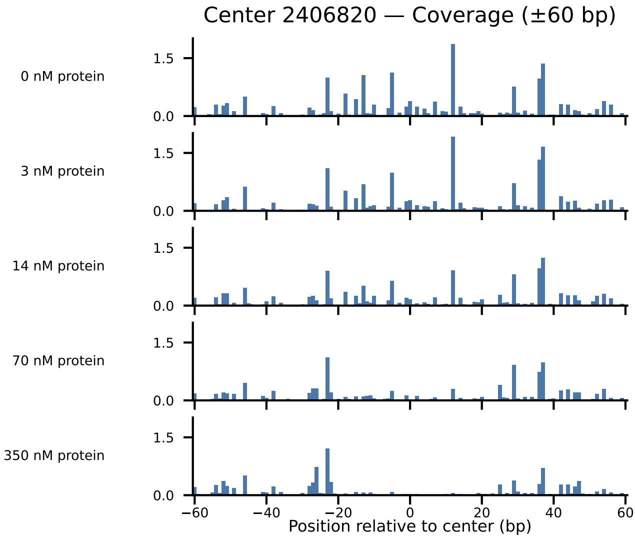
Center 2406820 — Coverage (±60 bp)
0 nM protein
3 nM protein
14 nM protein
70 nM protein
350 nM protein
1.5
0.0
−60
−40
−20
0
20
40
60
Position relative to center (bp)

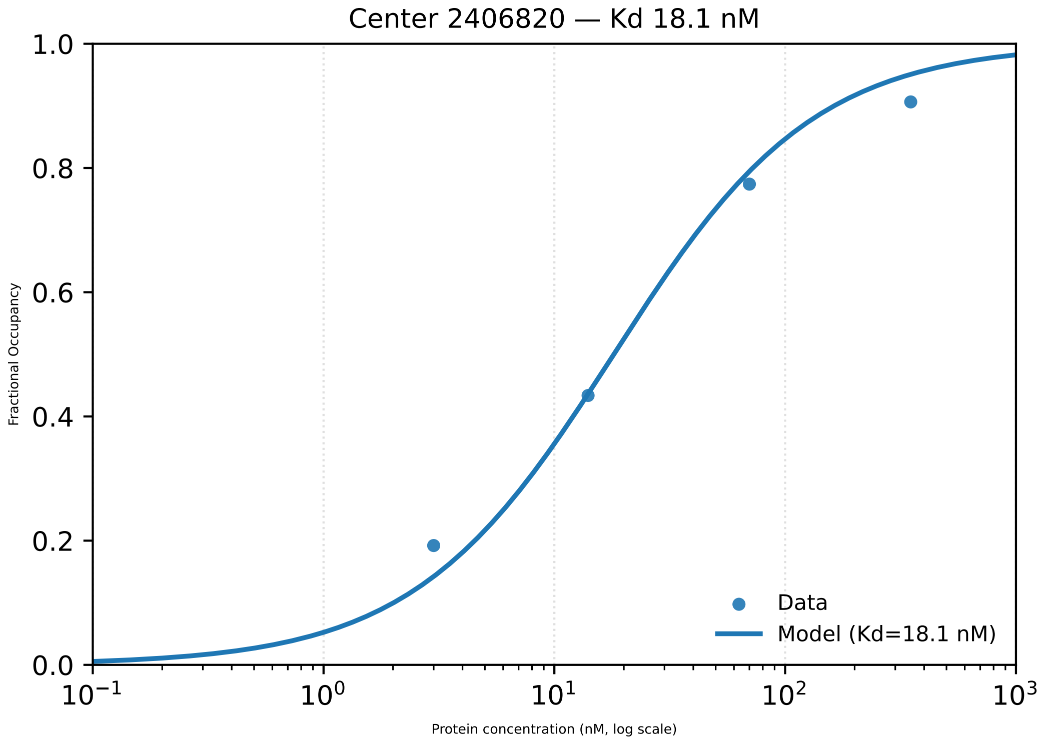
Center 2406820 — Kd 18.1 nM
Fractional Occupancy
Protein concentration (nM, log scale)
Data
Model (Kd=18.1 nM)
0.0
0.2
0.4
0.6
0.8
1.0
$10^{-1}$
$10^{0}$
$10^{1}$
$10^{2}$
$10^{3}$

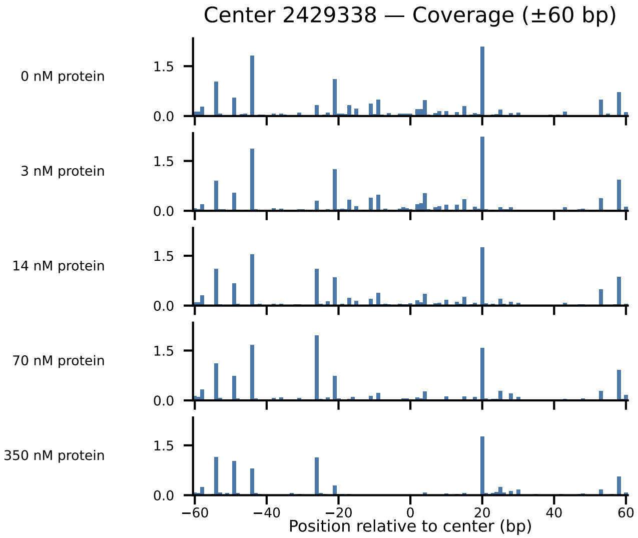
Center 2429338 — Coverage (±60 bp)
0 nM protein
3 nM protein
14 nM protein
70 nM protein
350 nM protein
1.5
0.0
−60
−40
−20
0
20
40
60
Position relative to center (bp)

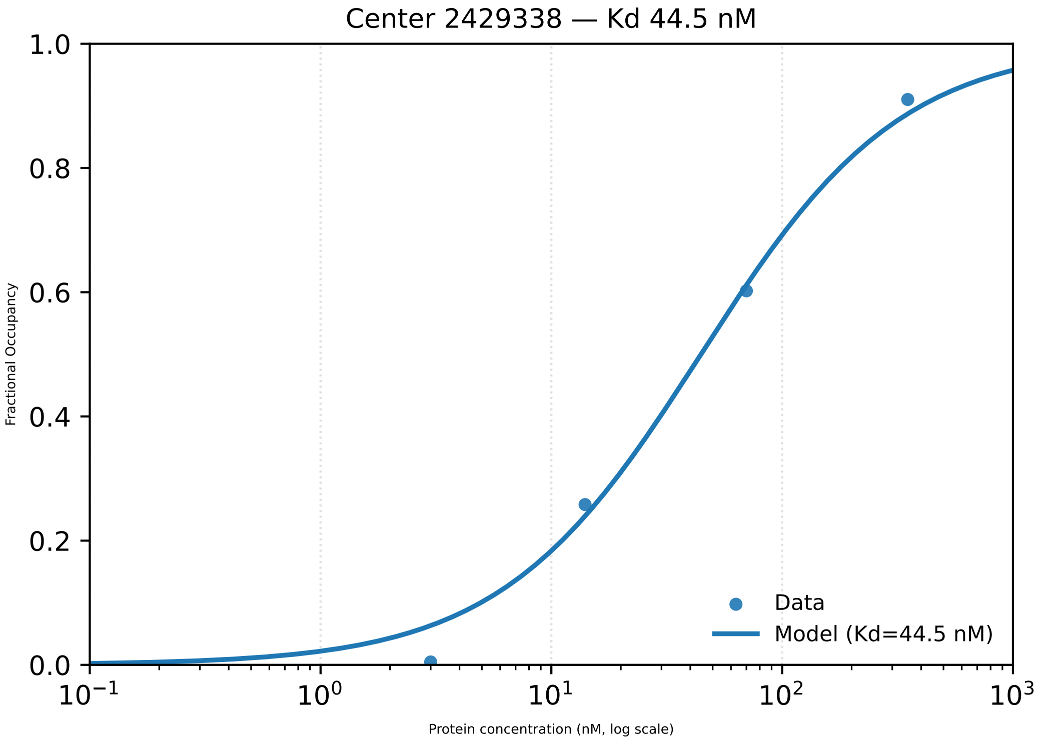
Center 2429338 — Kd 44.5 nM
Fractional Occupancy
1.0
0.8
0.6
0.4
0.2
0.0
$10^{-1}$
$10^{0}$
$10^{1}$
$10^{2}$
$10^{3}$
Protein concentration (nM, log scale)
Data
Model (Kd=44.5 nM)

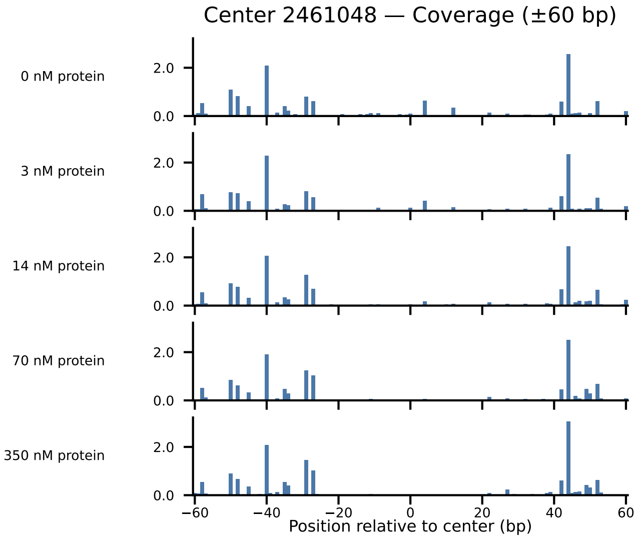
Center 2461048 — Coverage (±60 bp)
0 nM protein
3 nM protein
14 nM protein
70 nM protein
350 nM protein
2.0
0.0
−60
−40
−20
0
20
40
60
Position relative to center (bp)

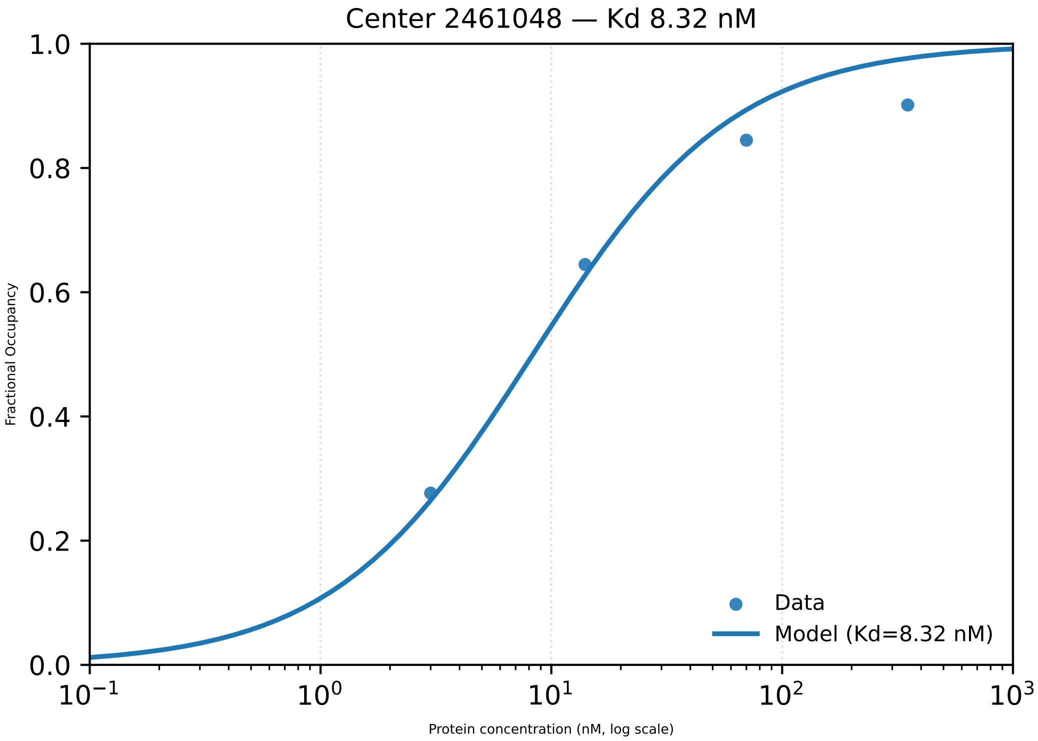

Center 2461048 — Kd 8.32 nM
Fractional Occupancy
1.0
0.8
0.6
0.4
0.2
0.0
$10^{-1}$
$10^{0}$
$10^{1}$
$10^{2}$
$10^{3}$
Protein concentration (nM, log scale)
Data
Model (Kd=8.32 nM)

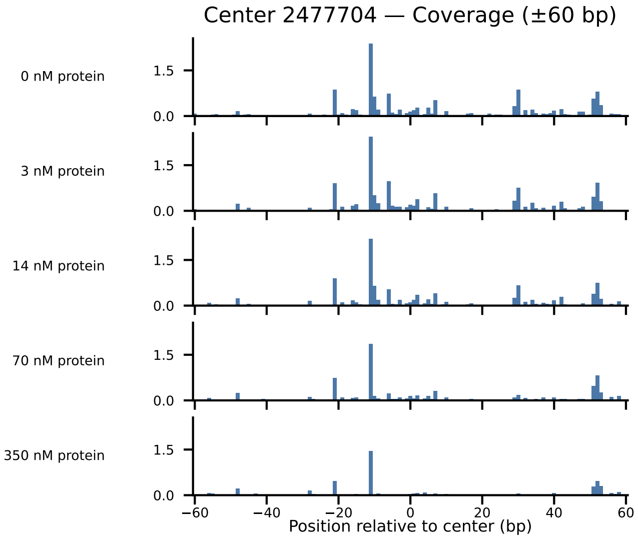

Center 2477704 — Coverage (±60 bp)
0 nM protein
3 nM protein
14 nM protein
70 nM protein
350 nM protein
1.5
0.0
−60
−40
−20
0
20
40
60
Position relative to center (bp)

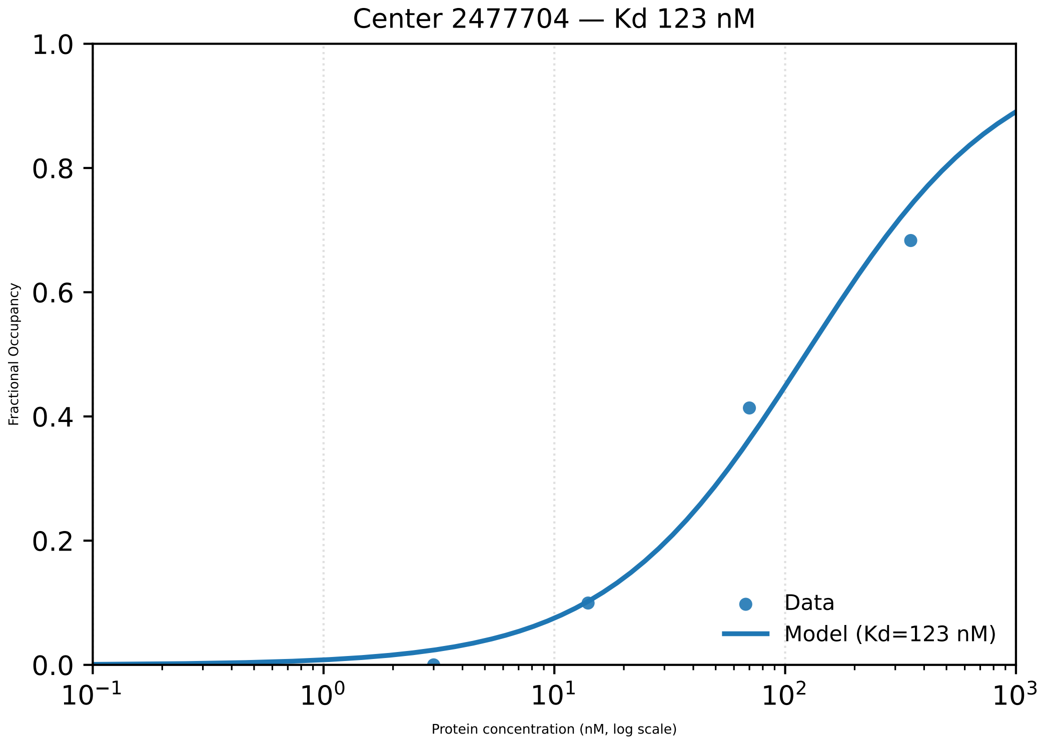

Center 2477704 — Kd 123 nM
1.0
0.8
0.6
0.4
0.2
0.0
Fractional Occupancy
$10^{-1}$
$10^{0}$
$10^{1}$
$10^{2}$
$10^{3}$
Protein concentration (nM, log scale)
Data
Model (Kd=123 nM)

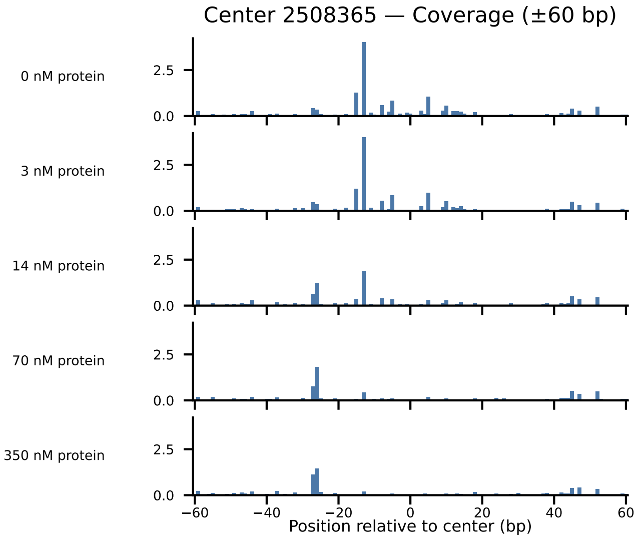

Center 2508365 — Coverage (±60 bp)
0 nM protein
3 nM protein
14 nM protein
70 nM protein
350 nM protein
2.5
0.0
−60
−40
−20
0
20
40
60
Position relative to center (bp)

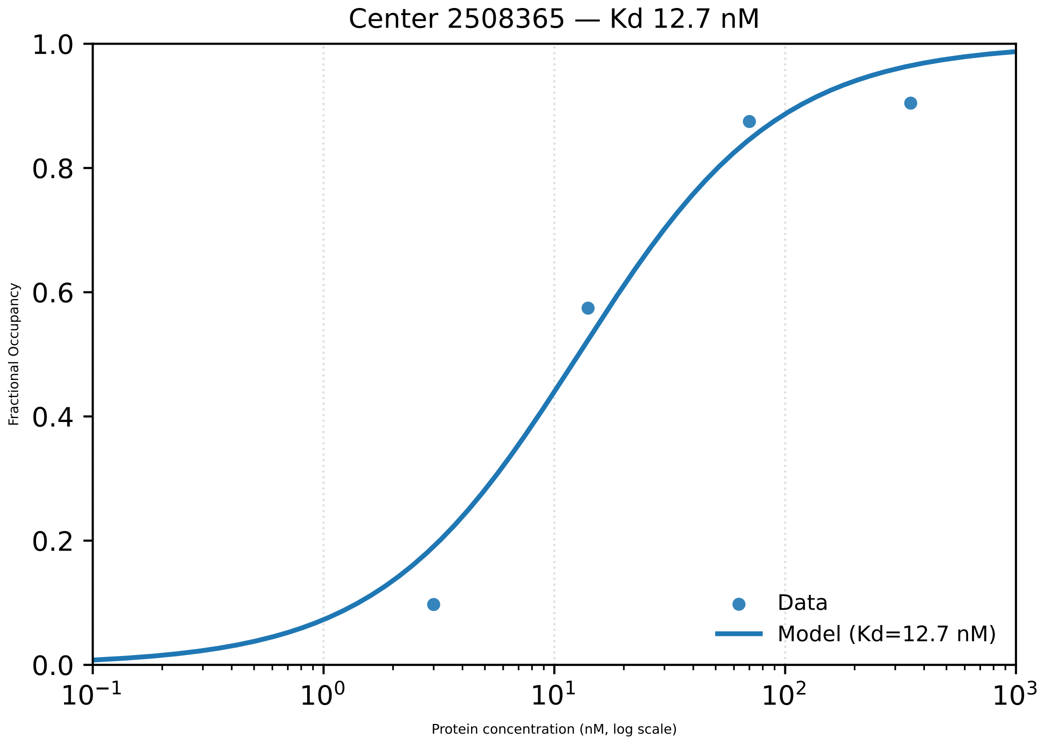

Center 2508365 — Kd 12.7 nM
1.0
0.8
0.6
0.4
0.2
0.0
Fractional Occupancy
$10^{-1}$
$10^{0}$
$10^{1}$
$10^{2}$
$10^{3}$
Protein concentration (nM, log scale)
Data
Model (Kd=12.7 nM)

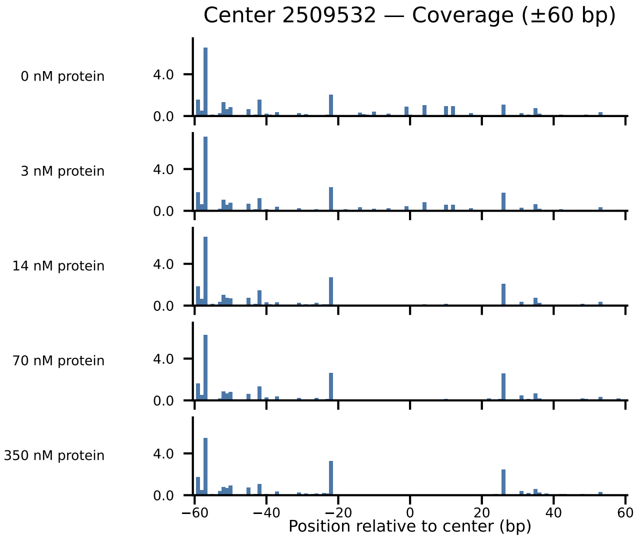

Center 2509532 — Coverage (±60 bp)
0 nM protein
3 nM protein
14 nM protein
70 nM protein
350 nM protein
4.0
0.0
−60
−40
−20
0
20
40
60
Position relative to center (bp)

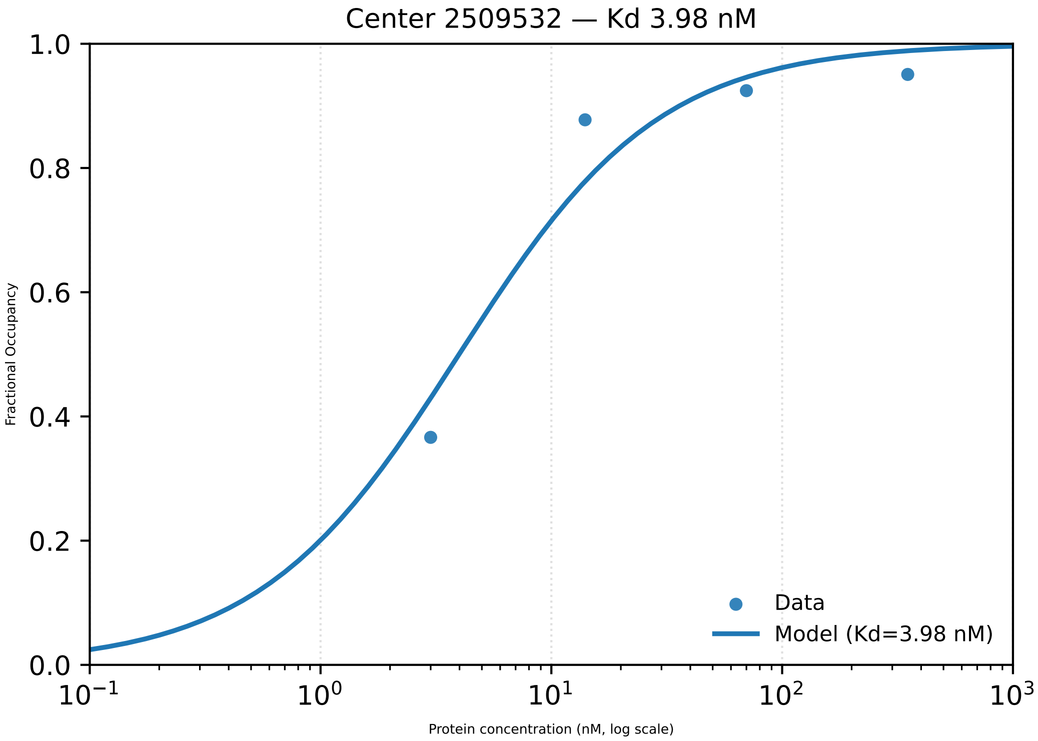

Center 2509532 — Kd 3.98 nM
Fractional Occupancy
1.0
0.8
0.6
0.4
0.2
0.0
$10^{-1}$
$10^{0}$
$10^{1}$
$10^{2}$
$10^{3}$
Protein concentration (nM, log scale)
Data
Model (Kd=3.98 nM)

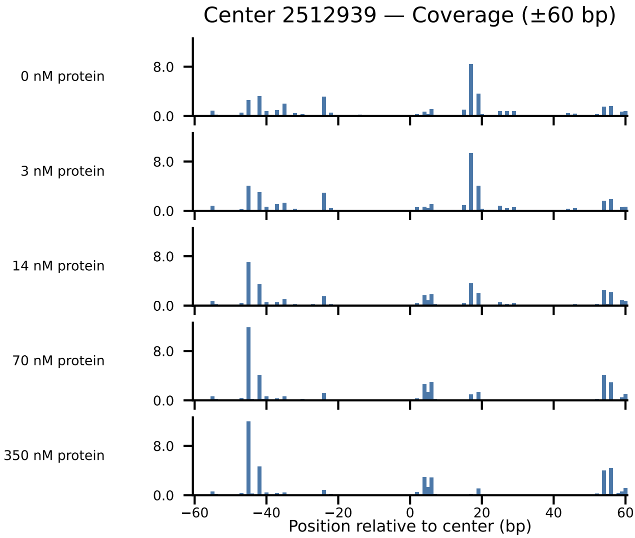

Center 2512939 — Coverage (±60 bp)
0 nM protein
3 nM protein
14 nM protein
70 nM protein
350 nM protein
8.0
0.0
−60
−40
−20
0
20
40
60
Position relative to center (bp)

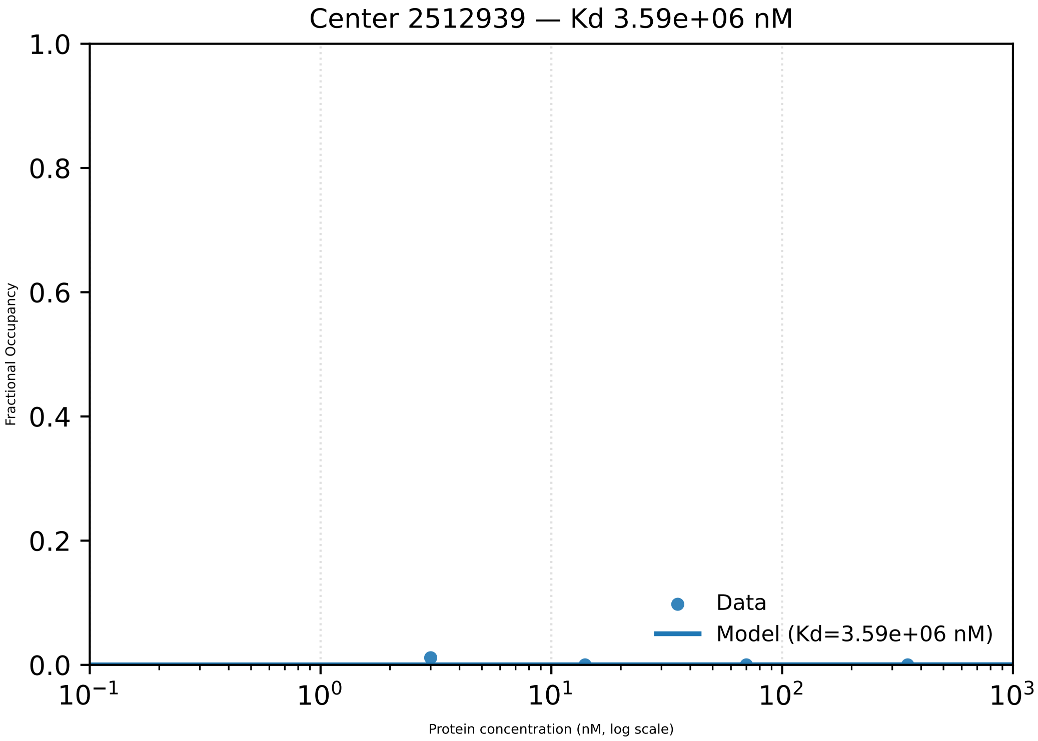
Center 2512939 — Kd 3.59e+06 nM
Fractional Occupancy
1.0
0.8
0.6
0.4
0.2
0.0
$10^{-1}$
$10^{0}$
$10^{1}$
$10^{2}$
$10^{3}$
Protein concentration (nM, log scale)
Data
Model (Kd=3.59e+06 nM)

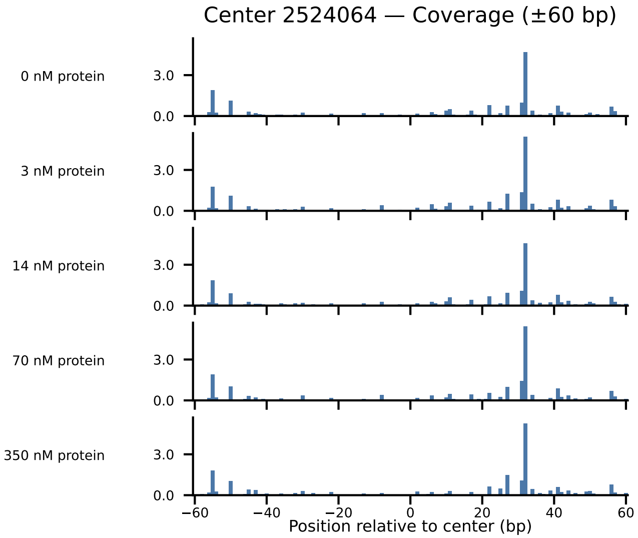

Center 2524064 — Coverage (±60 bp)
0 nM protein
3 nM protein
14 nM protein
70 nM protein
350 nM protein
3.0
0.0
−60
−40
−20
0
20
40
60
Position relative to center (bp)

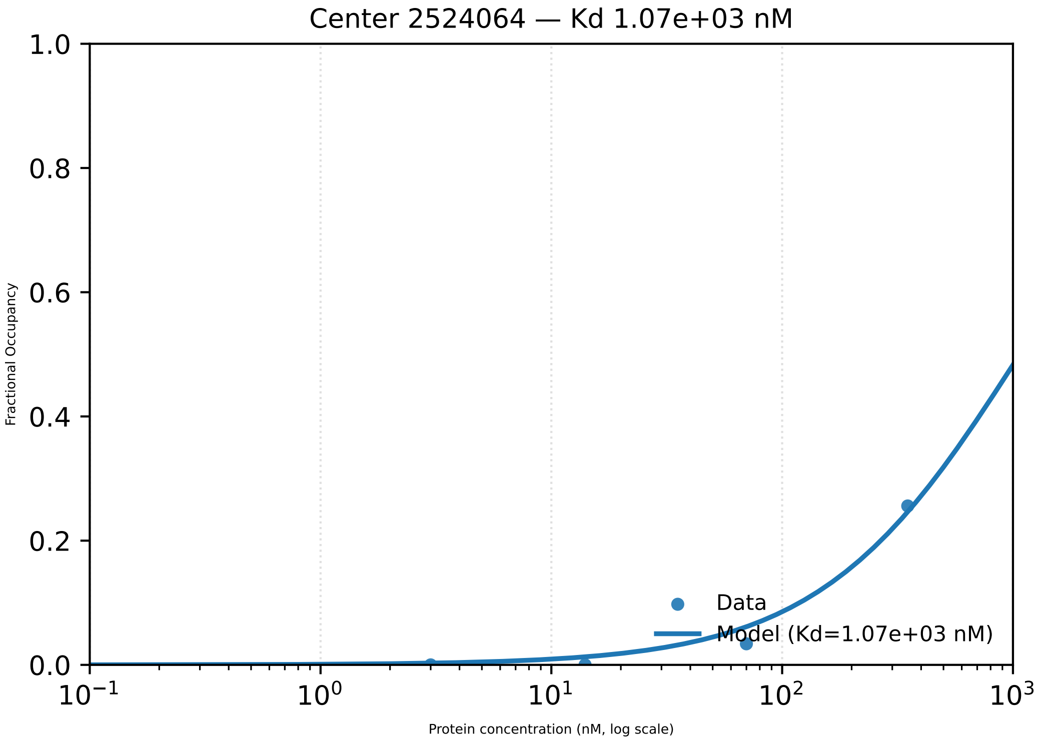

Center 2524064 — Kd 1.07e+03 nM
Fractional Occupancy
1.0
0.8
0.6
0.4
0.2
0.0
$10^{-1}$
$10^{0}$
$10^{1}$
$10^{2}$
$10^{3}$
Protein concentration (nM, log scale)
Data
Model (Kd=1.07e+03 nM)

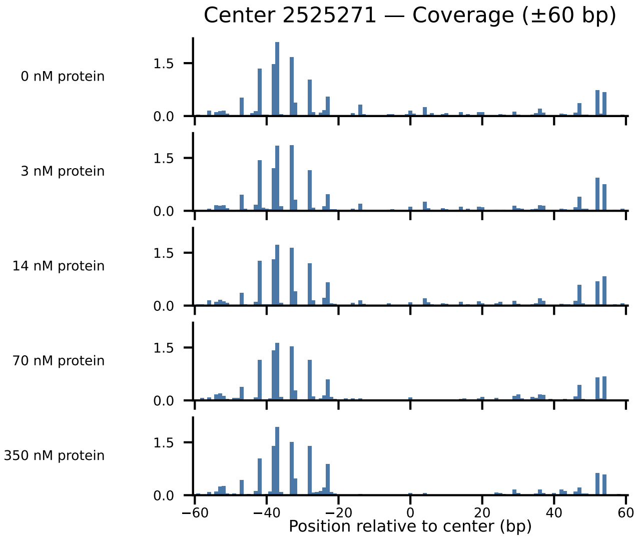
Center 2525271 — Coverage (±60 bp)
0 nM protein
3 nM protein
14 nM protein
70 nM protein
350 nM protein
1.5
0.0
−60
−40
−20
0
20
40
60
Position relative to center (bp)

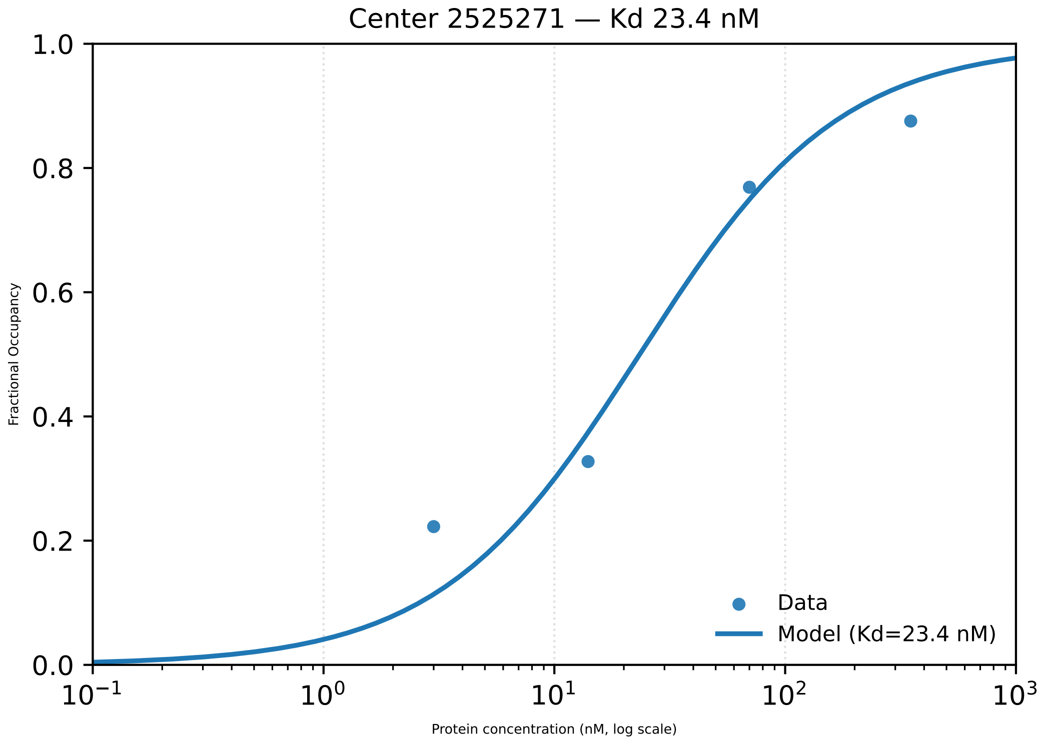

Center 2525271 — Kd 23.4 nM
1.0
0.8
0.6
0.4
0.2
0.0
Fractional Occupancy
$10^{-1}$
$10^{0}$
$10^{1}$
$10^{2}$
$10^{3}$
Protein concentration (nM, log scale)
Data
Model (Kd=23.4 nM)

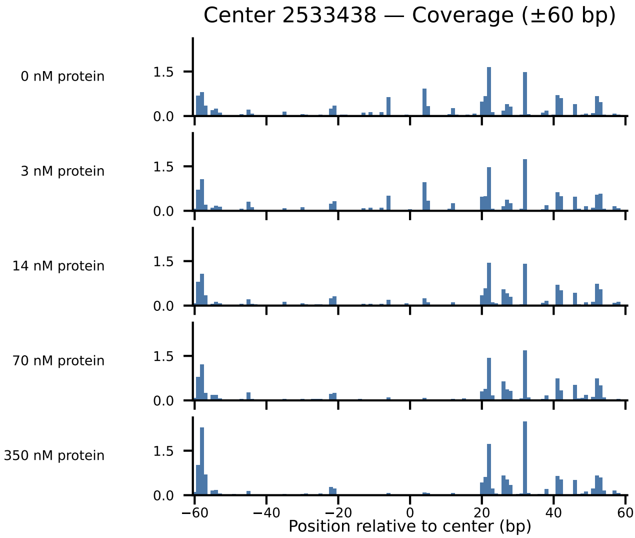

Center 2533438 — Coverage (±60 bp)
0 nM protein
3 nM protein
14 nM protein
70 nM protein
350 nM protein
1.5
0.0
−60
−40
−20
0
20
40
60
Position relative to center (bp)

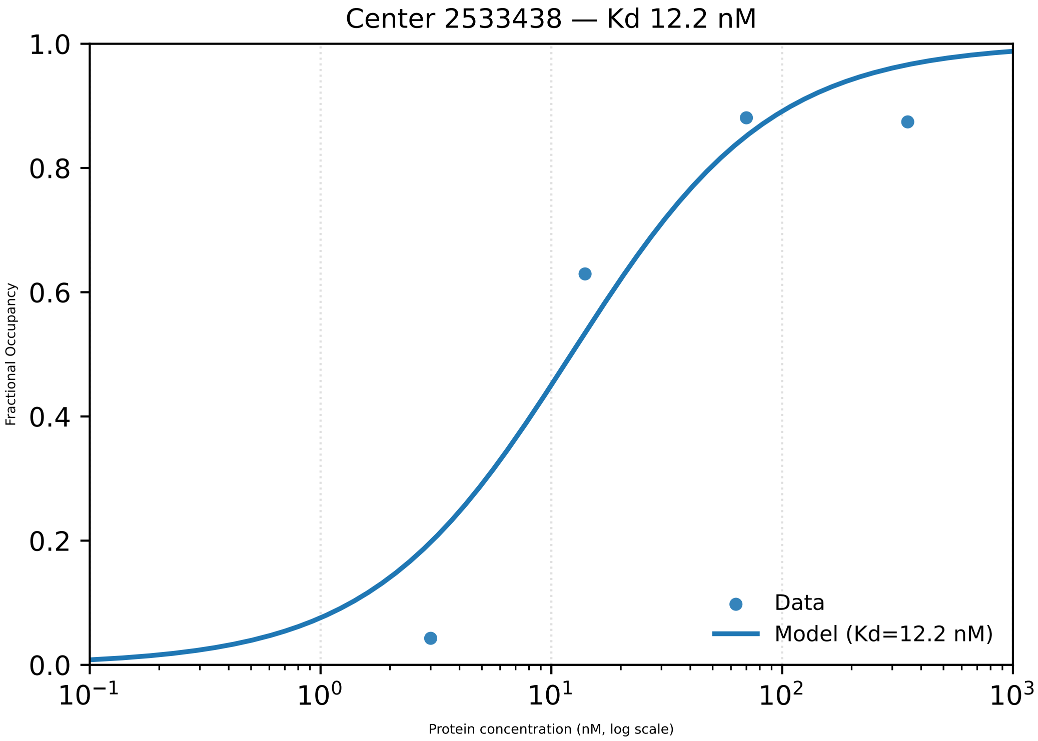

Center 2533438 — Kd 12.2 nM
1.0
0.8
0.6
0.4
0.2
0.0
Fractional Occupancy
$10^{-1}$
$10^{0}$
$10^{1}$
$10^{2}$
$10^{3}$
Protein concentration (nM, log scale)
Data
Model (Kd=12.2 nM)

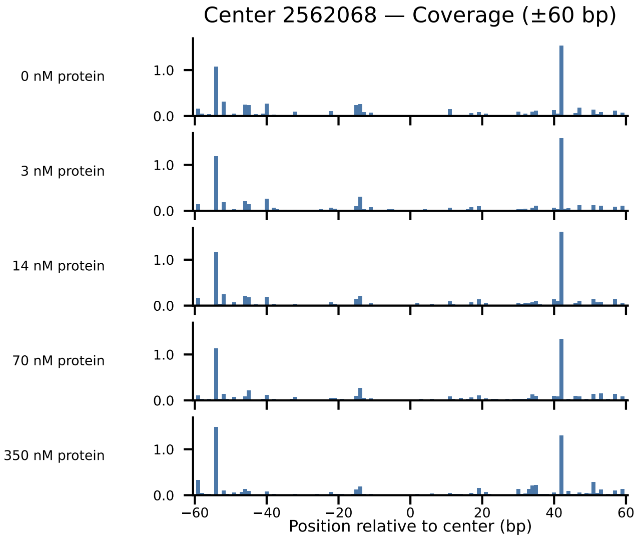
Center 2562068 — Coverage (±60 bp)
0 nM protein
3 nM protein
14 nM protein
70 nM protein
350 nM protein
1.0
0.0
−60
−40
−20
0
20
40
60
Position relative to center (bp)

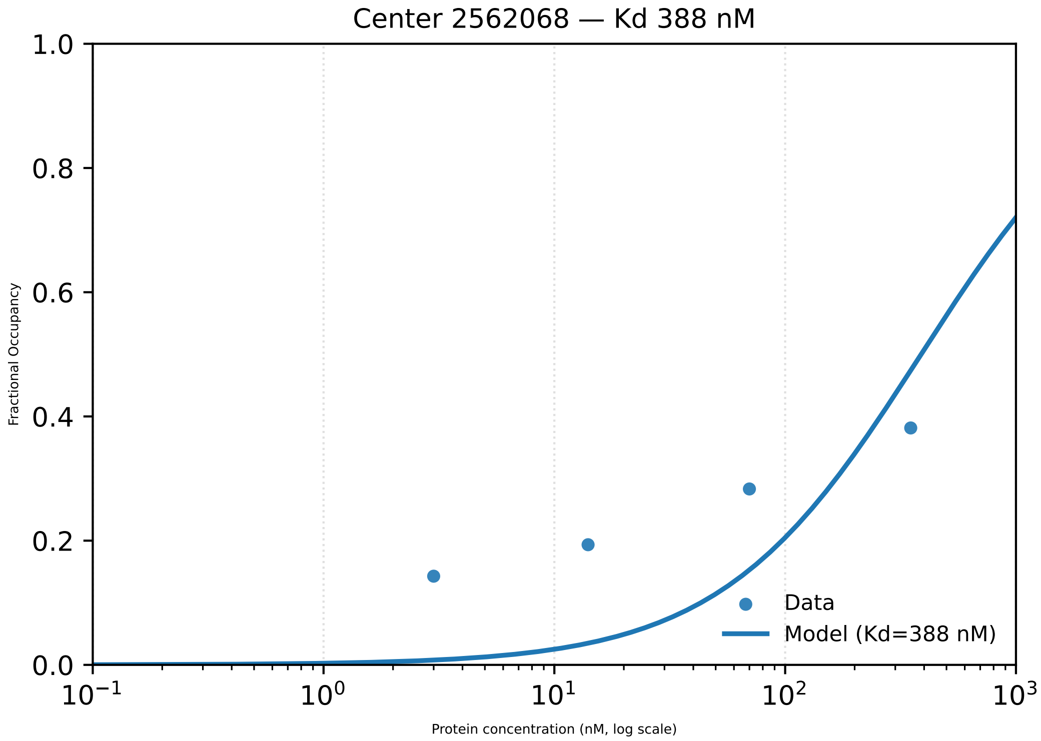

Center 2562068 — Kd 388 nM
1.0
0.8
0.6
0.4
0.2
0.0
Fractional Occupancy
$10^{-1}$
$10^{0}$
$10^{1}$
$10^{2}$
$10^{3}$
Protein concentration (nM, log scale)
Data
Model (Kd=388 nM)

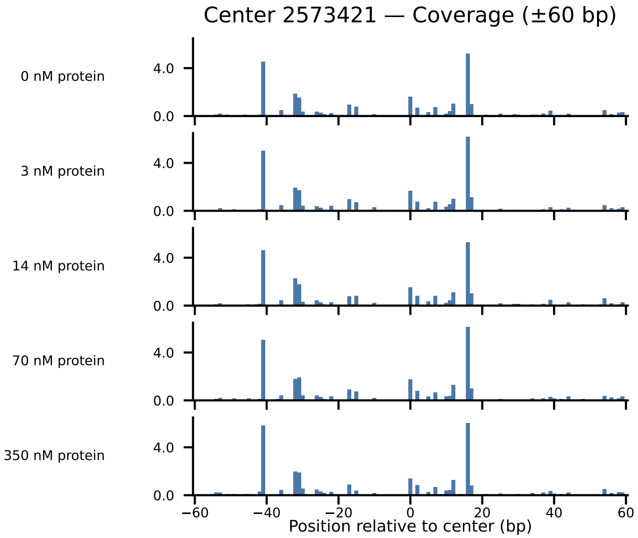
Center 2573421 — Coverage (±60 bp)
0 nM protein
3 nM protein
14 nM protein
70 nM protein
350 nM protein
4.0
0.0
−60
−40
−20
0
20
40
60
Position relative to center (bp)

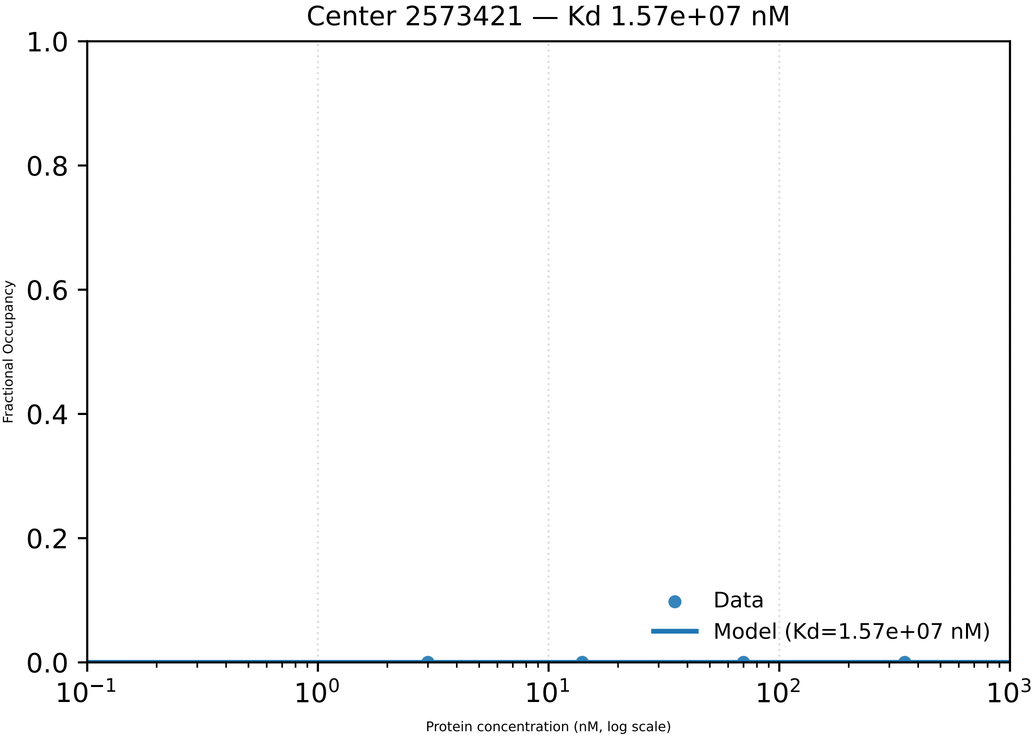

Center 2573421 — Kd 1.57e+07 nM
1.0
0.8
0.6
0.4
0.2
0.0
Fractional Occupancy
Data
Model (Kd=1.57e+07 nM)
$10^{-1}$
$10^{0}$
$10^{1}$
$10^{2}$
$10^{3}$
Protein concentration (nM, log scale)

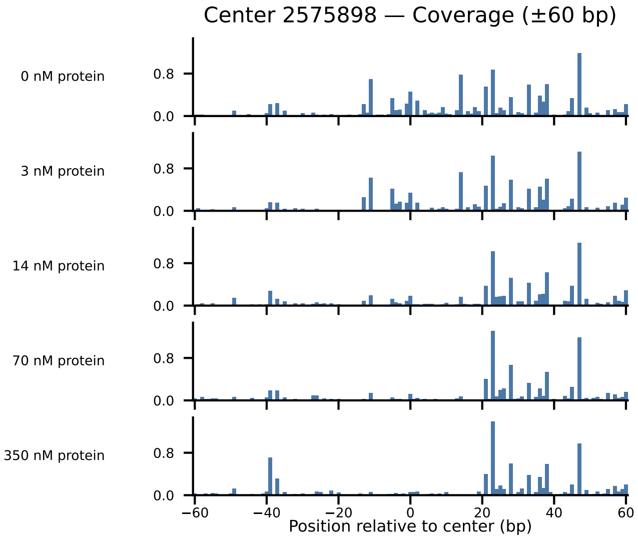
Center 2575898 — Coverage (±60 bp)
0 nM protein
3 nM protein
14 nM protein
70 nM protein
350 nM protein
0.8
0.0
−60
−40
−20
0
20
40
60
Position relative to center (bp)

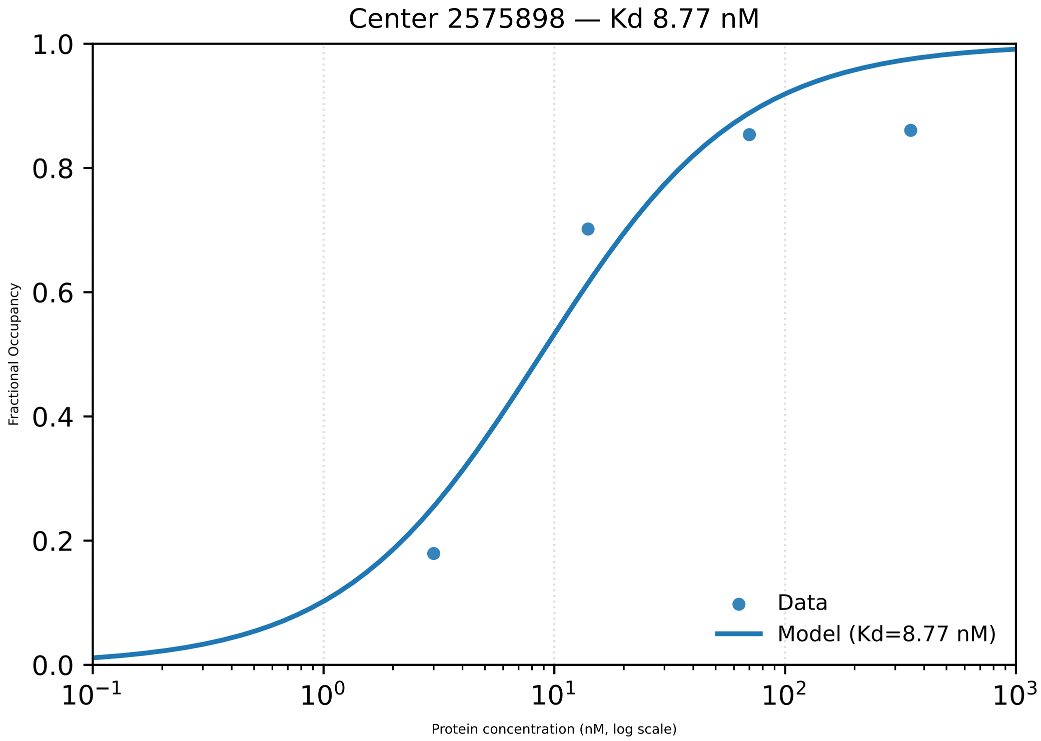

Center 2575898 — Kd 8.77 nM
1.0
0.8
0.6
0.4
0.2
0.0
Fractional Occupancy
$10^{-1}$
$10^{0}$
$10^{1}$
$10^{2}$
$10^{3}$
Protein concentration (nM, log scale)
Data
Model (Kd=8.77 nM)

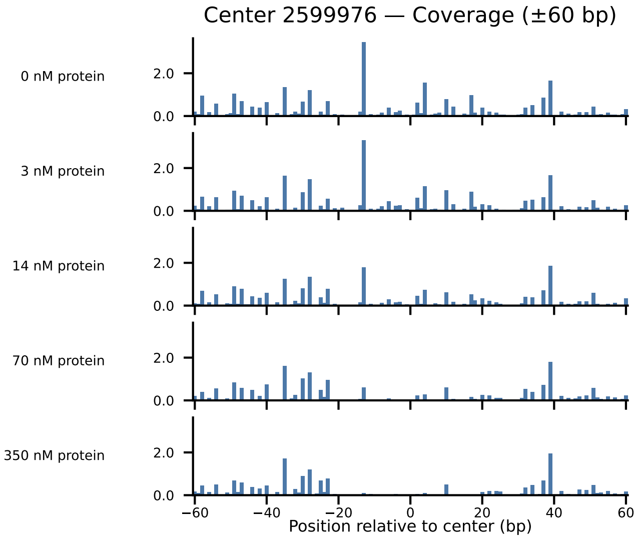

Center 2599976 — Coverage (±60 bp)
0 nM protein
3 nM protein
14 nM protein
70 nM protein
350 nM protein
0.0
2.0
−60
−40
−20
0
20
40
60
Position relative to center (bp)

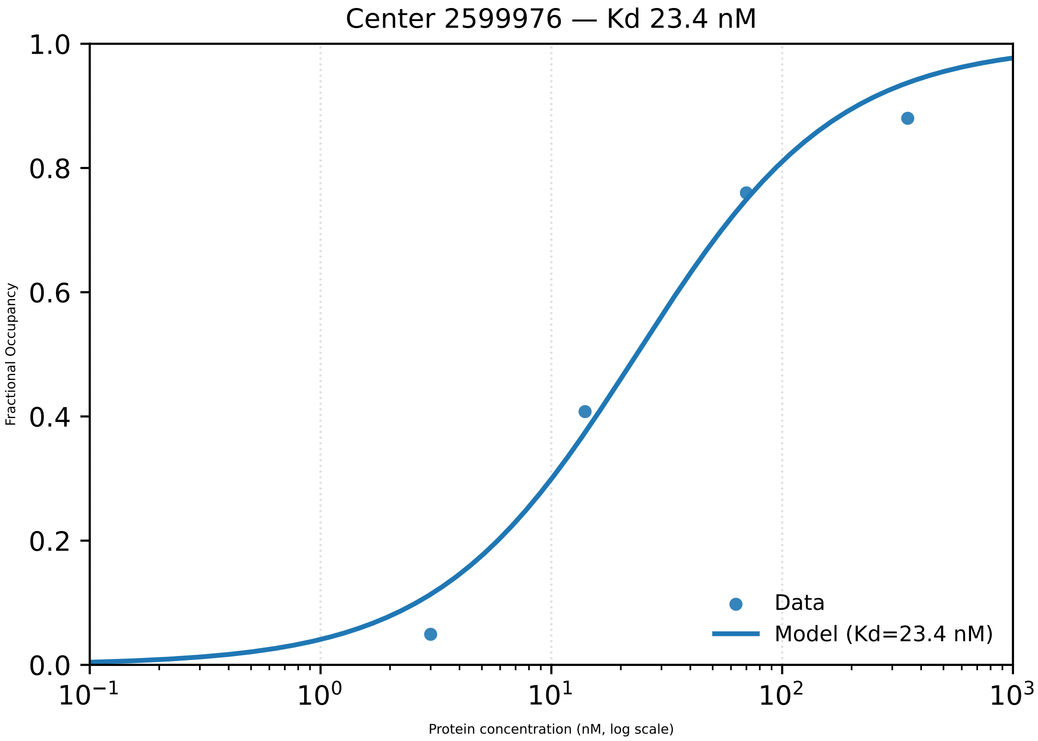

Center 2599976 — Kd 23.4 nM
Fractional Occupancy
1.0
0.8
0.6
0.4
0.2
0.0
$10^{-1}$
$10^{0}$
$10^{1}$
$10^{2}$
$10^{3}$
Protein concentration (nM, log scale)
Data
Model (Kd=23.4 nM)

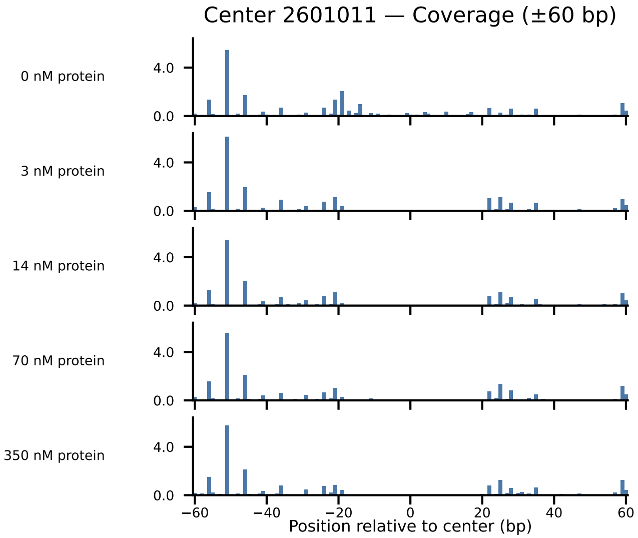
Center 2601011 — Coverage (±60 bp)
0 nM protein
3 nM protein
14 nM protein
70 nM protein
350 nM protein
4.0
0.0
−60
−40
−20
0
20
40
60
Position relative to center (bp)

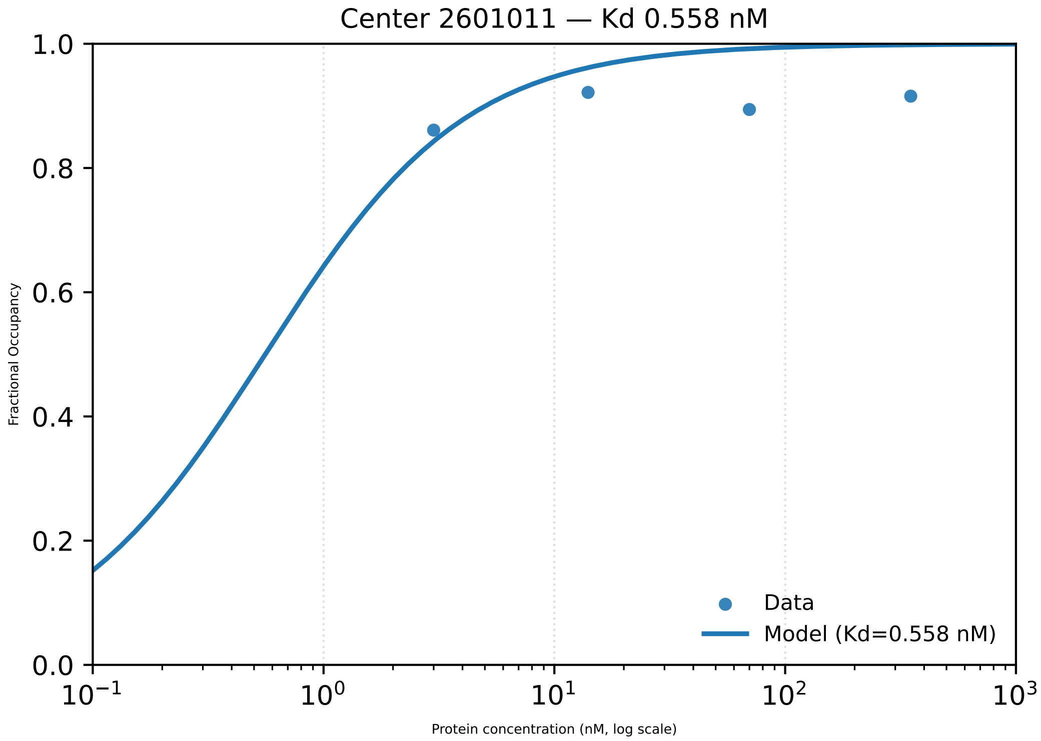

Center 2601011 — Kd 0.558 nM
Fractional Occupancy
1.0
0.8
0.6
0.4
0.2
0.0
$10^{-1}$
$10^{0}$
$10^{1}$
$10^{2}$
$10^{3}$
Protein concentration (nM, log scale)
Data
Model (Kd=0.558 nM)

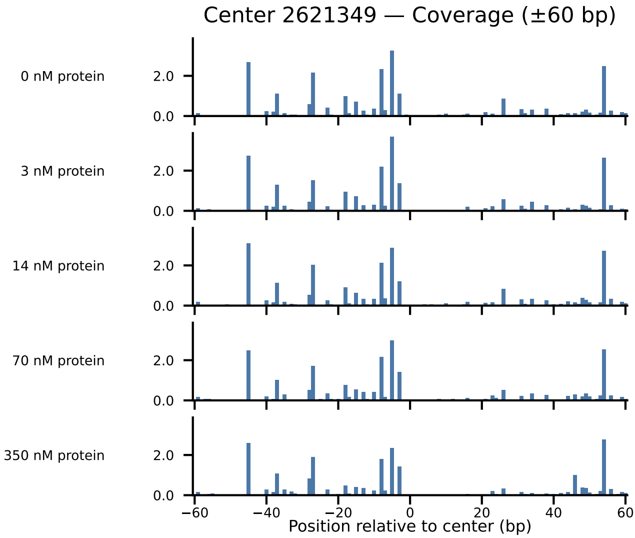

Center 2621349 — Coverage (±60 bp)
0 nM protein
3 nM protein
14 nM protein
70 nM protein
350 nM protein
2.0
0.0
−60
−40
−20
0
20
40
60
Position relative to center (bp)

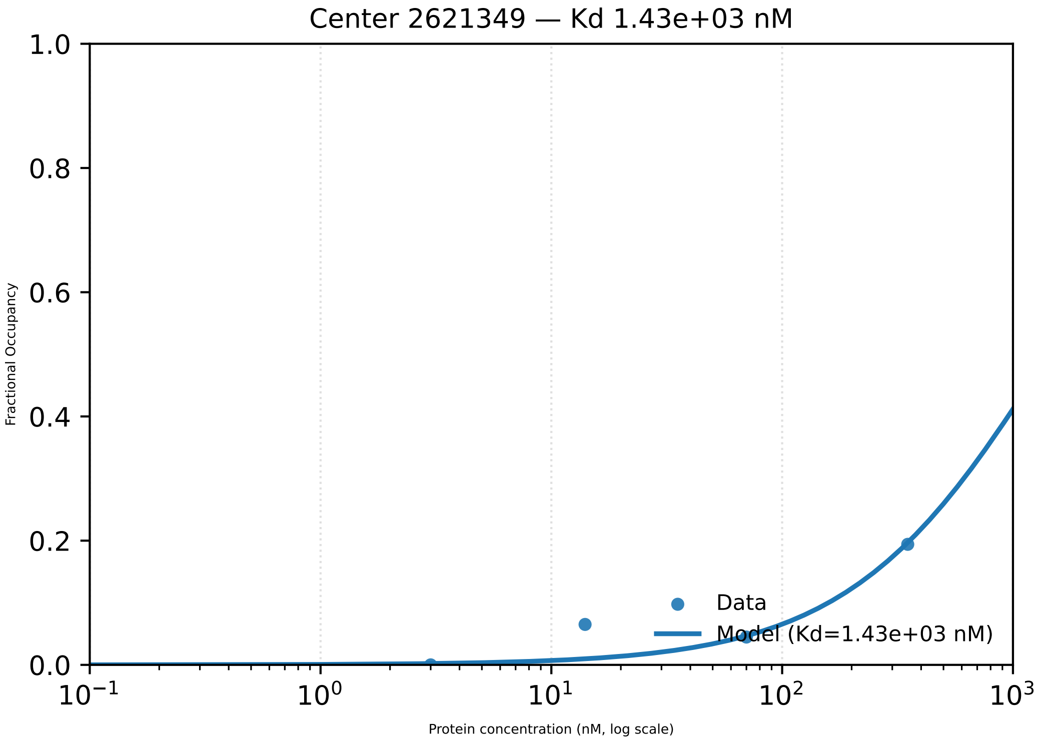

Center 2621349 — Kd 1.43e+03 nM
Fractional Occupancy
Protein concentration (nM, log scale)
Data
Model (Kd=1.43e+03 nM)
1.0
0.8
0.6
0.4
0.2
0.0
$10^{-1}$
$10^{0}$
$10^{1}$
$10^{2}$
$10^{3}$

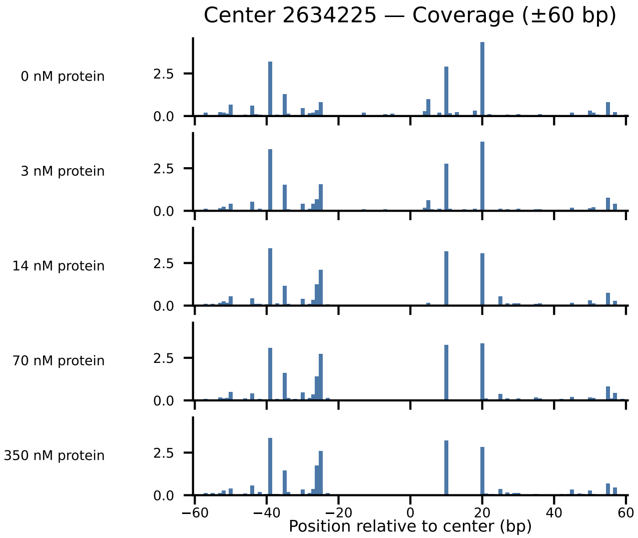

Center 2634225 — Coverage (±60 bp)
0 nM protein
3 nM protein
14 nM protein
70 nM protein
350 nM protein
2.5
0.0
−60
−40
−20
0
20
40
60
Position relative to center (bp)

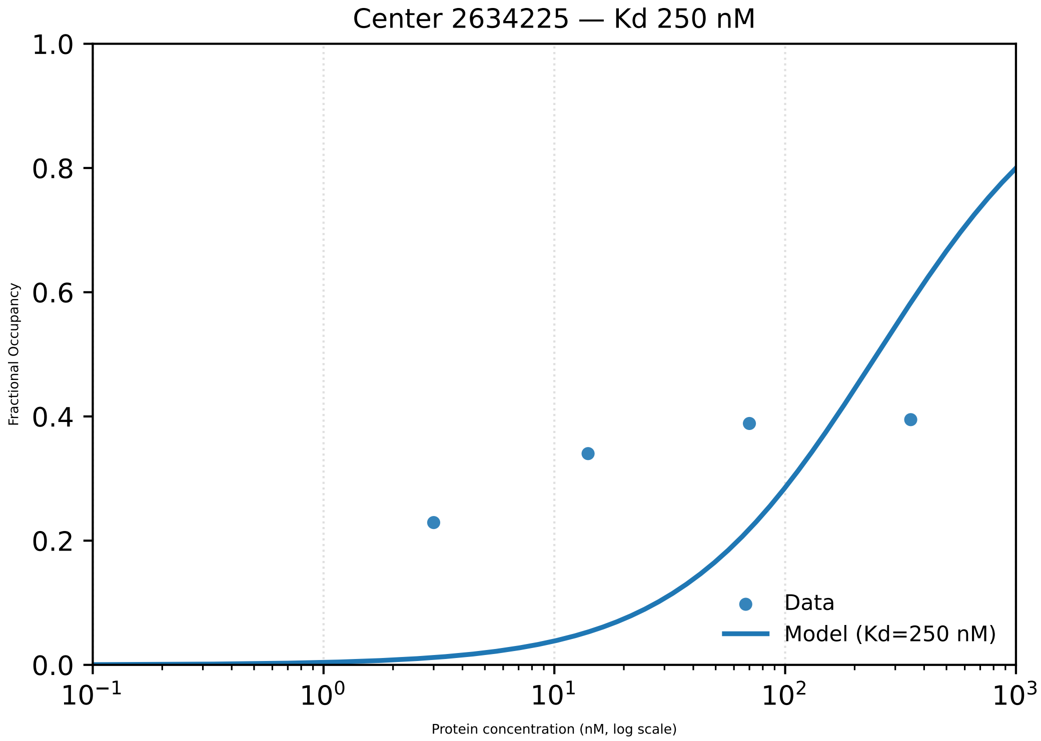

Center 2634225 — Kd 250 nM
Fractional Occupancy
Protein concentration (nM, log scale)
1.0
0.8
0.6
0.4
0.2
0.0
$10^{-1}$
$10^{0}$
$10^{1}$
$10^{2}$
$10^{3}$
Data
Model (Kd=250 nM)

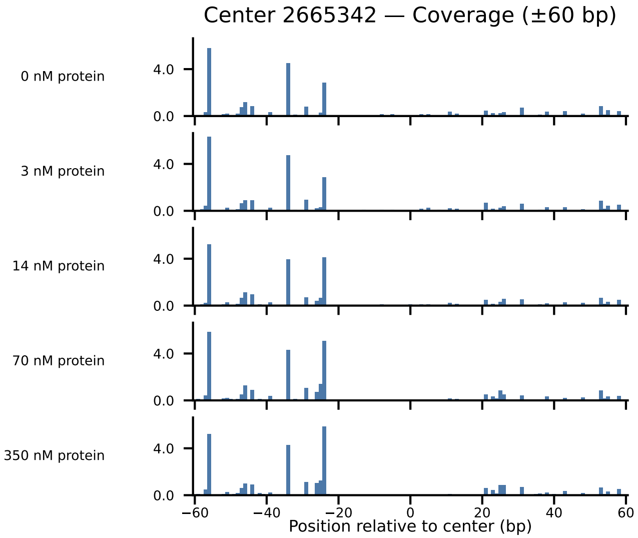

Center 2665342 — Coverage (±60 bp)
0 nM protein
3 nM protein
14 nM protein
70 nM protein
350 nM protein
4.0
0.0
−60
−40
−20
0
20
40
60
Position relative to center (bp)

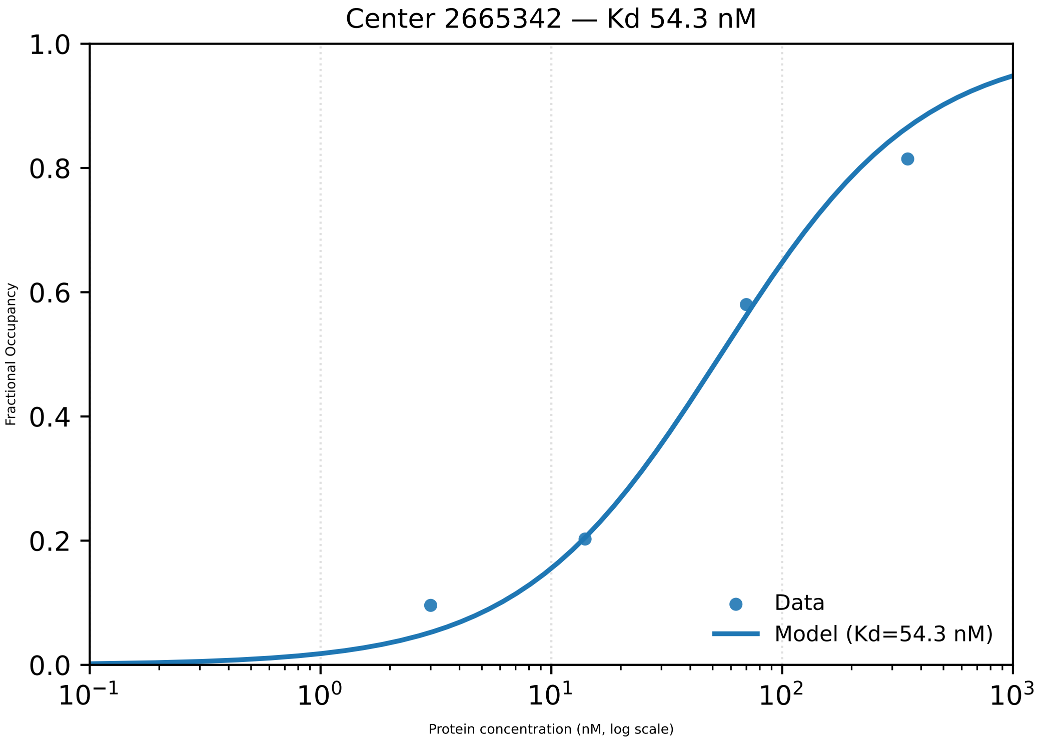

Center 2665342 — Kd 54.3 nM
Fractional Occupancy
Protein concentration (nM, log scale)
1.0
0.8
0.6
0.4
0.2
0.0
$10^{-1}$
$10^{0}$
$10^{1}$
$10^{2}$
$10^{3}$
Data
Model (Kd=54.3 nM)

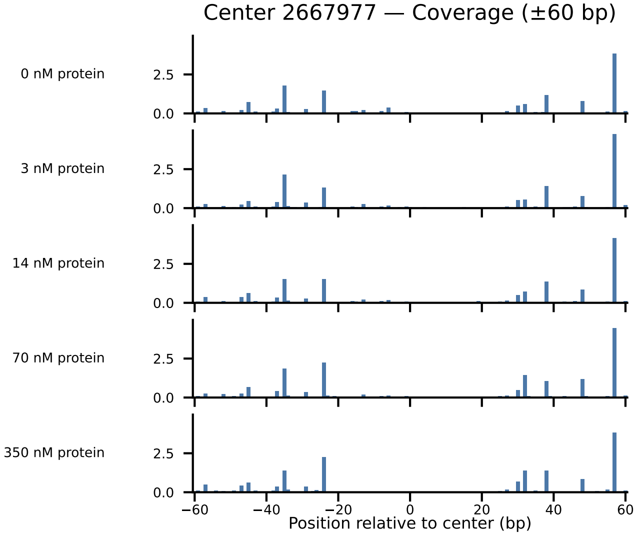

Center 2667977 — Coverage (±60 bp)
0 nM protein
3 nM protein
14 nM protein
70 nM protein
350 nM protein
2.5
0.0
−60
−40
−20
0
20
40
60
Position relative to center (bp)

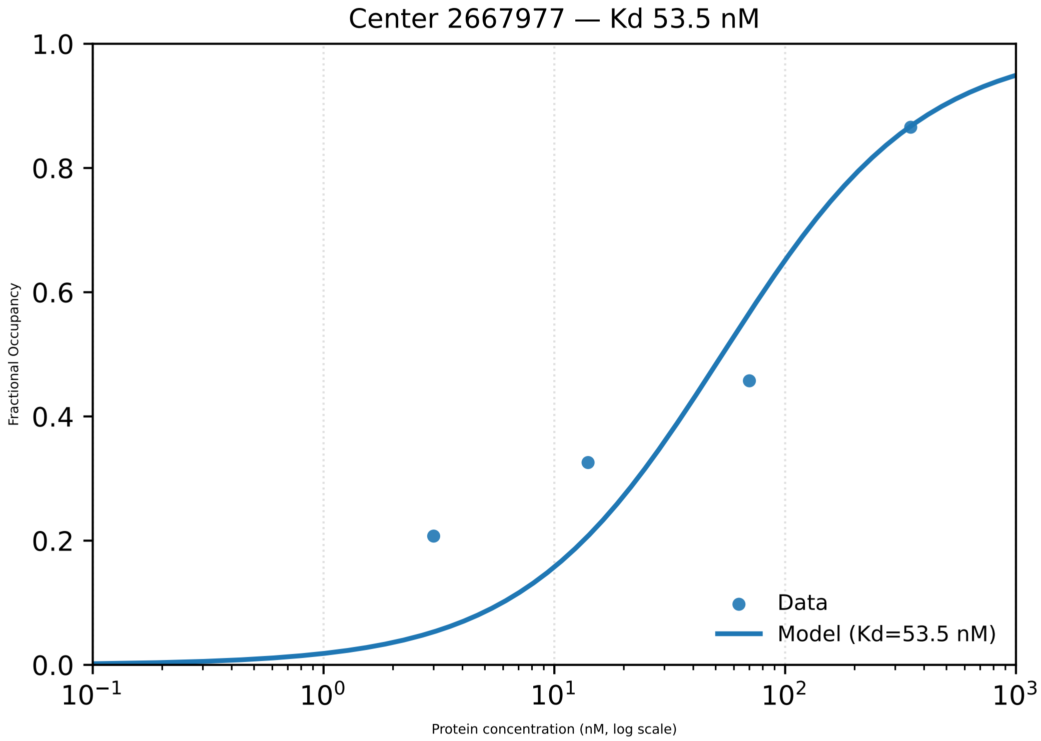

Center 2667977 — Kd 53.5 nM
1.0
0.8
0.6
0.4
0.2
0.0
Fractional Occupancy
$10^{-1}$
$10^{0}$
$10^{1}$
$10^{2}$
$10^{3}$
Protein concentration (nM, log scale)
Data
Model (Kd=53.5 nM)

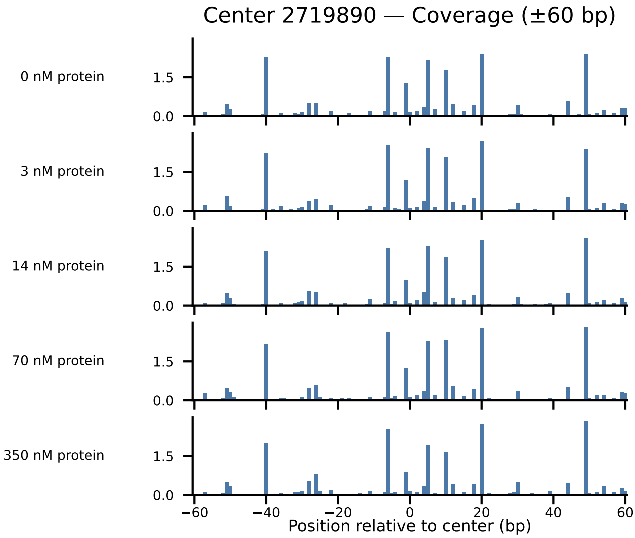

Center 2719890 — Coverage (±60 bp)
0 nM protein
3 nM protein
14 nM protein
70 nM protein
350 nM protein
1.5
0.0
−60
−40
−20
0
20
40
60
Position relative to center (bp)

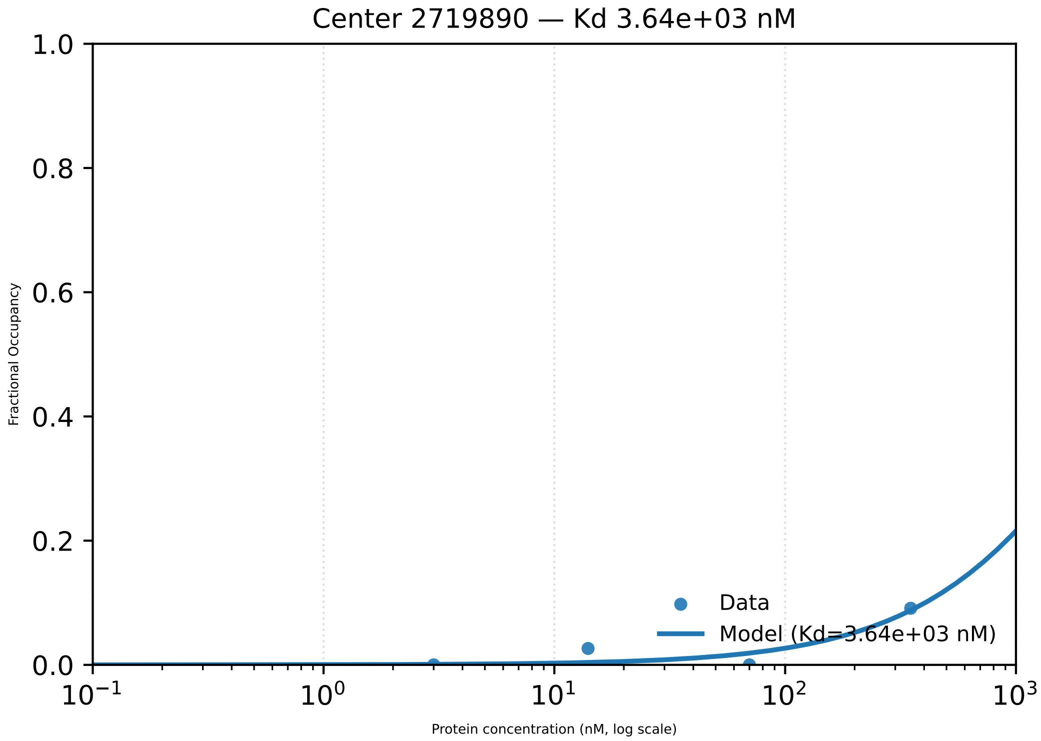

Center 2719890 — Kd 3.64e+03 nM
Fractional Occupancy
1.0
0.8
0.6
0.4
0.2
0.0
$10^{-1}$
$10^{0}$
$10^{1}$
$10^{2}$
$10^{3}$
Protein concentration (nM, log scale)
Data
Model (Kd=3.64e+03 nM)

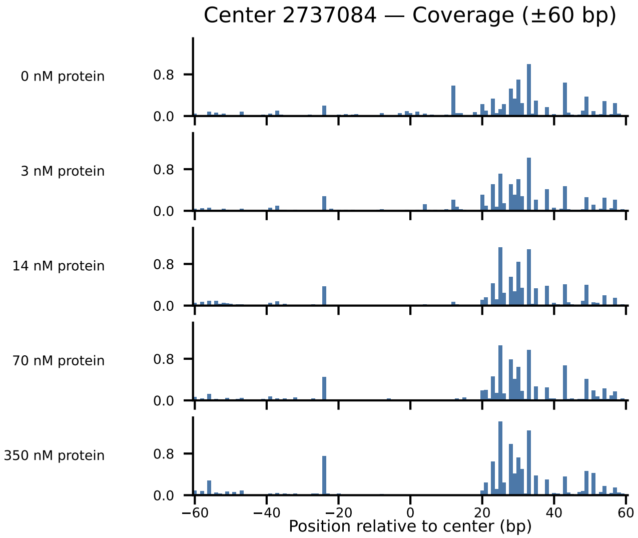

Center 2737084 — Coverage (±60 bp)
0 nM protein
3 nM protein
14 nM protein
70 nM protein
350 nM protein
0.8
0.0
−60
−40
−20
0
20
40
60
Position relative to center (bp)

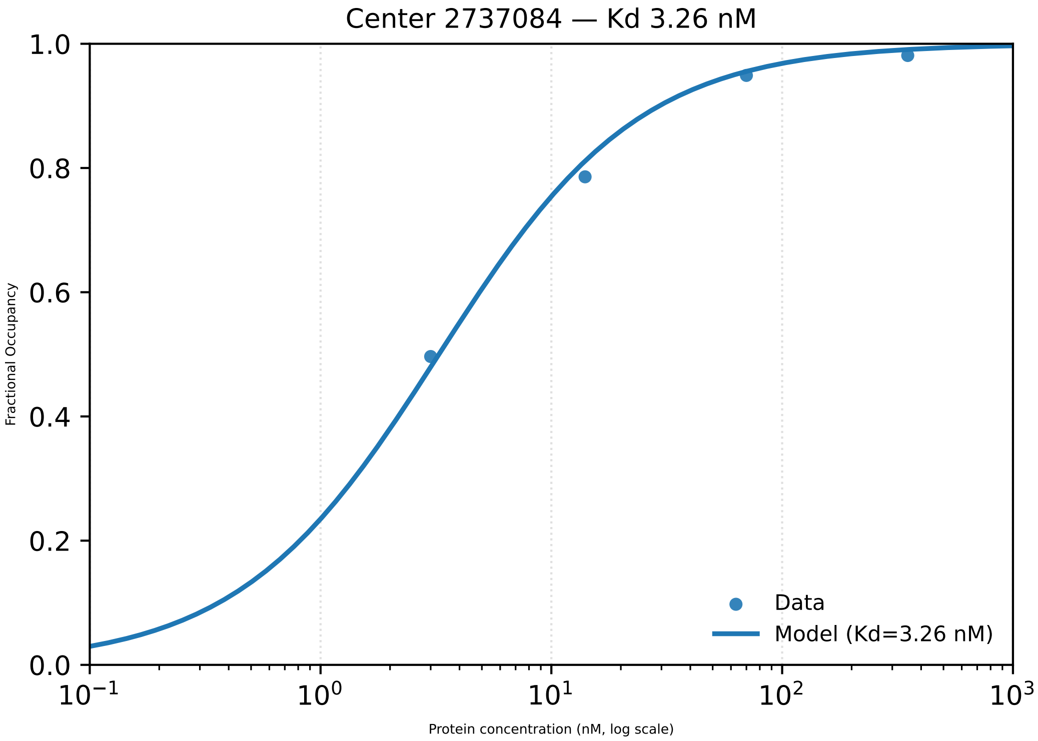

Center 2737084 — Kd 3.26 nM
Fractional Occupancy
0.0
0.2
0.4
0.6
0.8
1.0
$10^{-1}$
$10^{0}$
$10^{1}$
$10^{2}$
$10^{3}$
Protein concentration (nM, log scale)
Data
Model (Kd=3.26 nM)

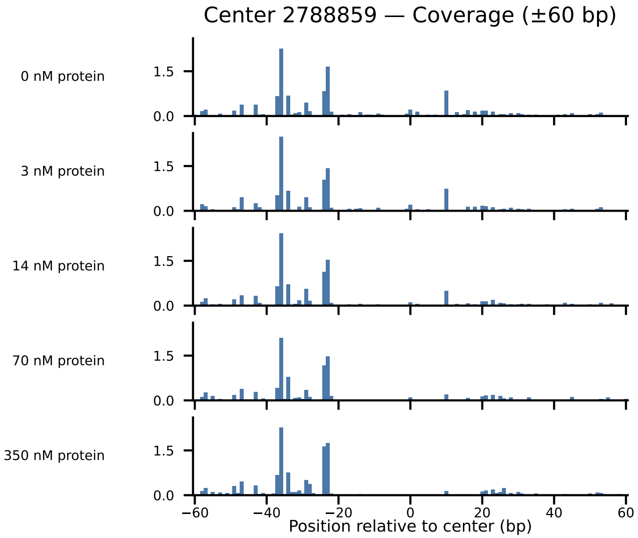

Center 2788859 — Coverage (±60 bp)
0 nM protein
3 nM protein
14 nM protein
70 nM protein
350 nM protein
1.5
0.0
−60
−40
−20
0
20
40
60
Position relative to center (bp)

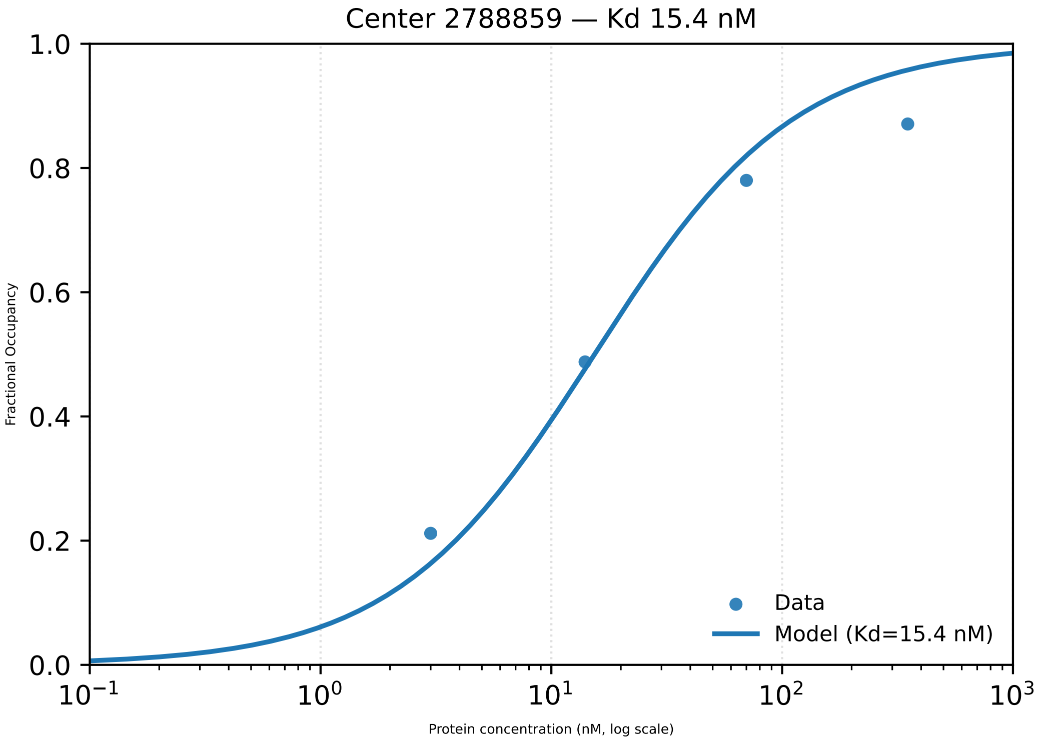

Center 2788859 — Kd 15.4 nM
Fractional Occupancy
1.0
0.8
0.6
0.4
0.2
0.0
$10^{-1}$
$10^{0}$
$10^{1}$
$10^{2}$
$10^{3}$
Protein concentration (nM, log scale)
Data
Model (Kd=15.4 nM)

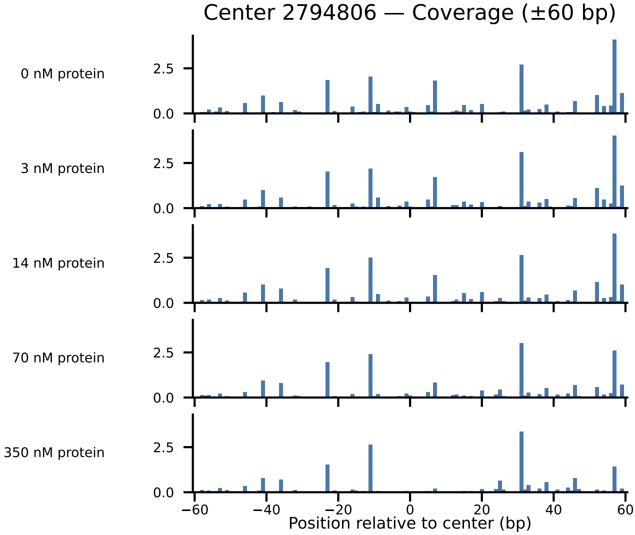
Center 2794806 — Coverage (±60 bp)
0 nM protein
3 nM protein
14 nM protein
70 nM protein
350 nM protein
0.0
2.5
−60
−40
−20
0
20
40
60
Position relative to center (bp)

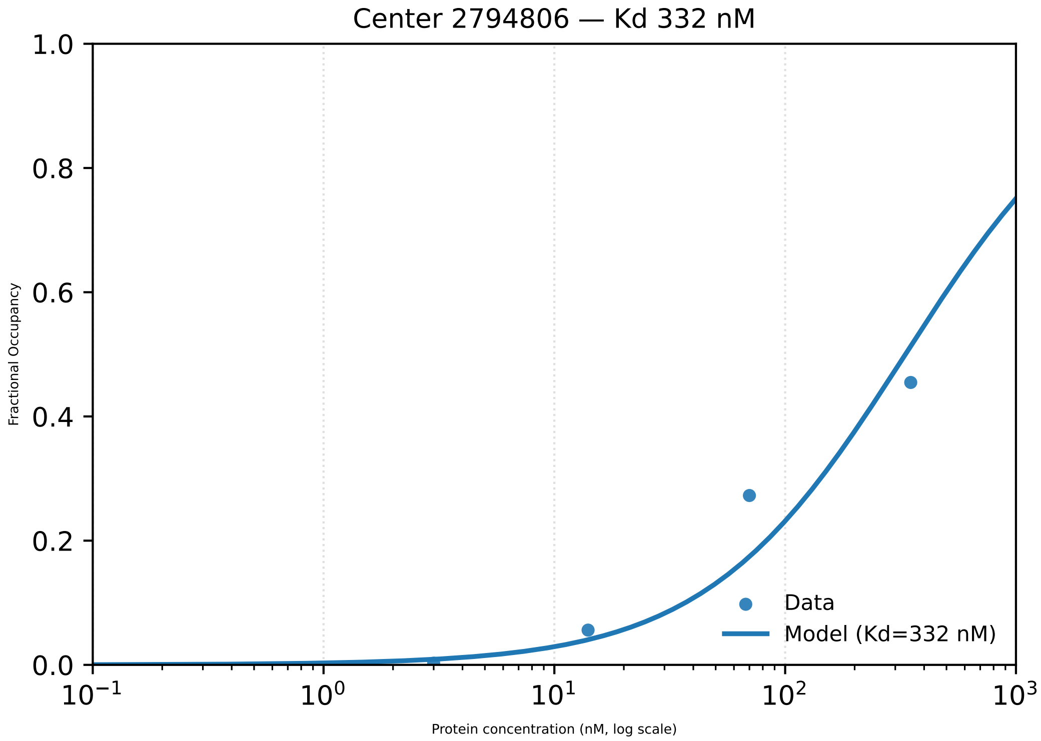

Center 2794806 — Kd 332 nM
Fractional Occupancy
1.0
0.8
0.6
0.4
0.2
0.0
$10^{-1}$
$10^{0}$
$10^{1}$
$10^{2}$
$10^{3}$
Protein concentration (nM, log scale)
Data
Model (Kd=332 nM)

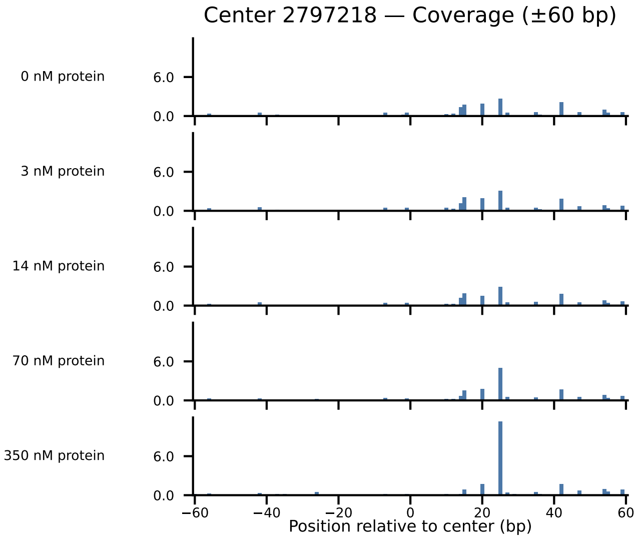

Center 2797218 — Coverage (±60 bp)
0 nM protein
3 nM protein
14 nM protein
70 nM protein
350 nM protein
6.0
0.0
−60
−40
−20
0
20
40
60
Position relative to center (bp)

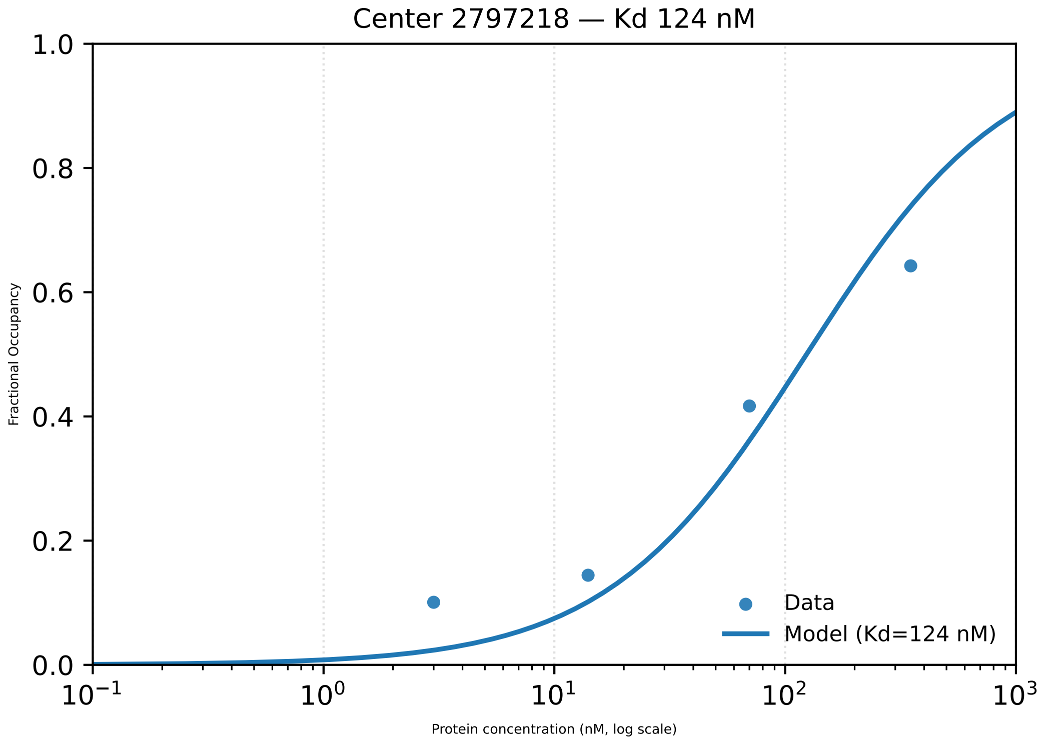

Center 2797218 — Kd 124 nM
1.0
0.8
0.6
0.4
0.2
0.0
Fractional Occupancy
$10^{-1}$
$10^{0}$
$10^{1}$
$10^{2}$
$10^{3}$
Protein concentration (nM, log scale)
Data
Model (Kd=124 nM)

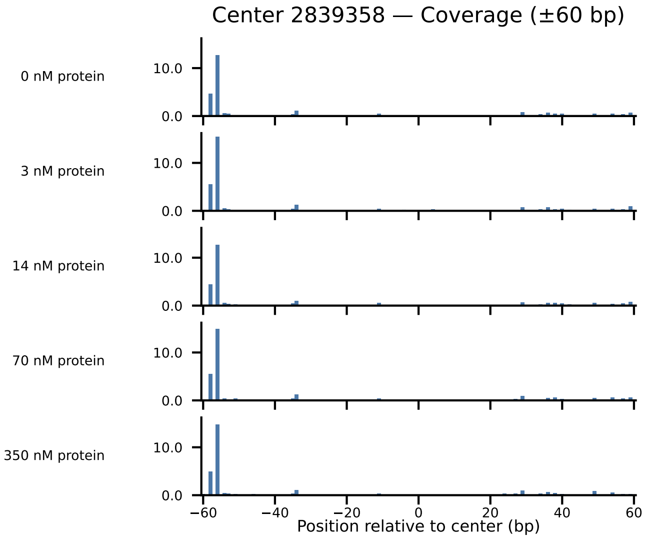
Center 2839358 — Coverage (±60 bp)
0 nM protein
3 nM protein
14 nM protein
70 nM protein
350 nM protein
10.0
0.0
−60
−40
−20
0
20
40
60
Position relative to center (bp)

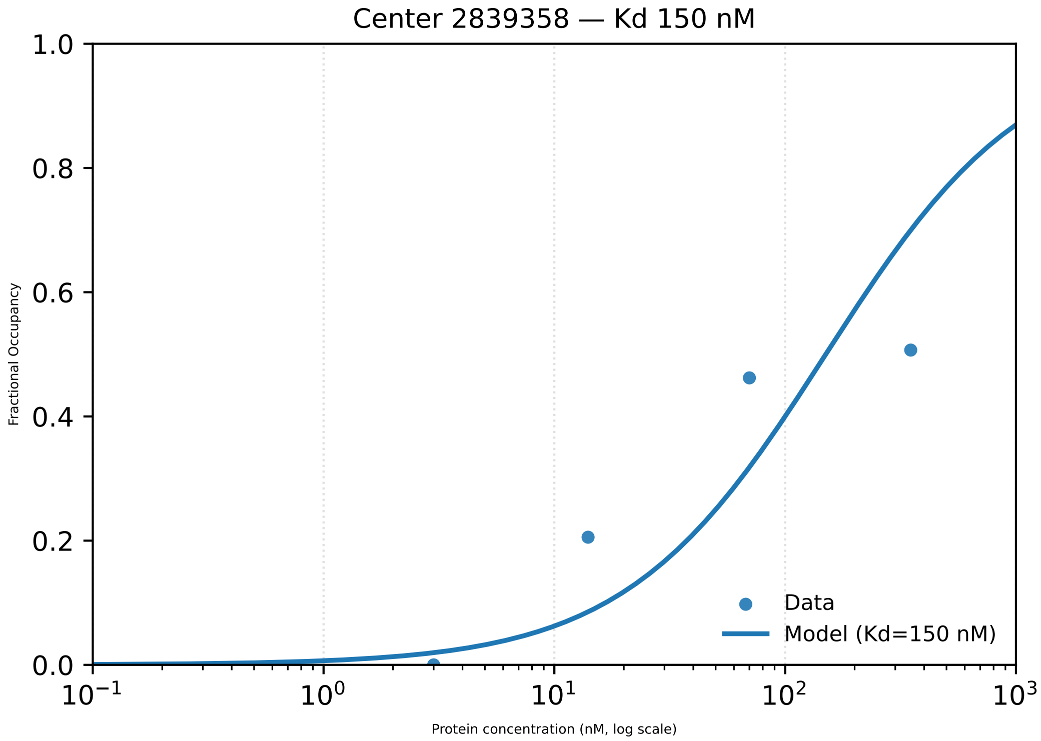

Center 2839358 — Kd 150 nM
Fractional Occupancy
1.0
0.8
0.6
0.4
0.2
0.0
$10^{-1}$
$10^{0}$
$10^{1}$
$10^{2}$
$10^{3}$
Protein concentration (nM, log scale)
Data
Model (Kd=150 nM)

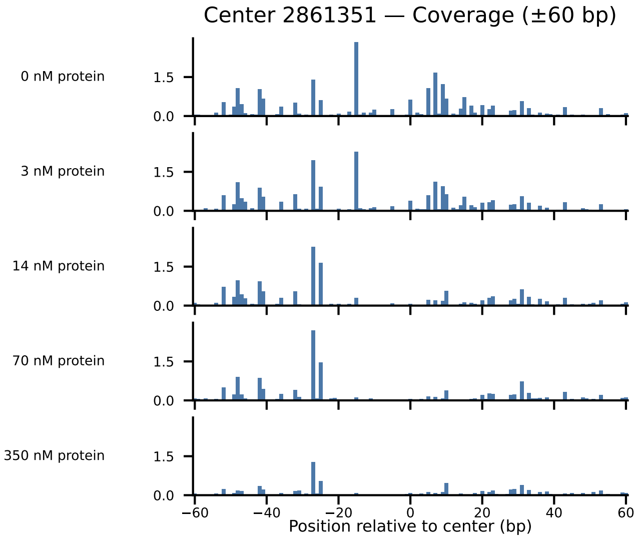
Center 2861351 — Coverage (±60 bp)
0 nM protein
3 nM protein
14 nM protein
70 nM protein
350 nM protein
1.5
0.0
−60
−40
−20
0
20
40
60
Position relative to center (bp)

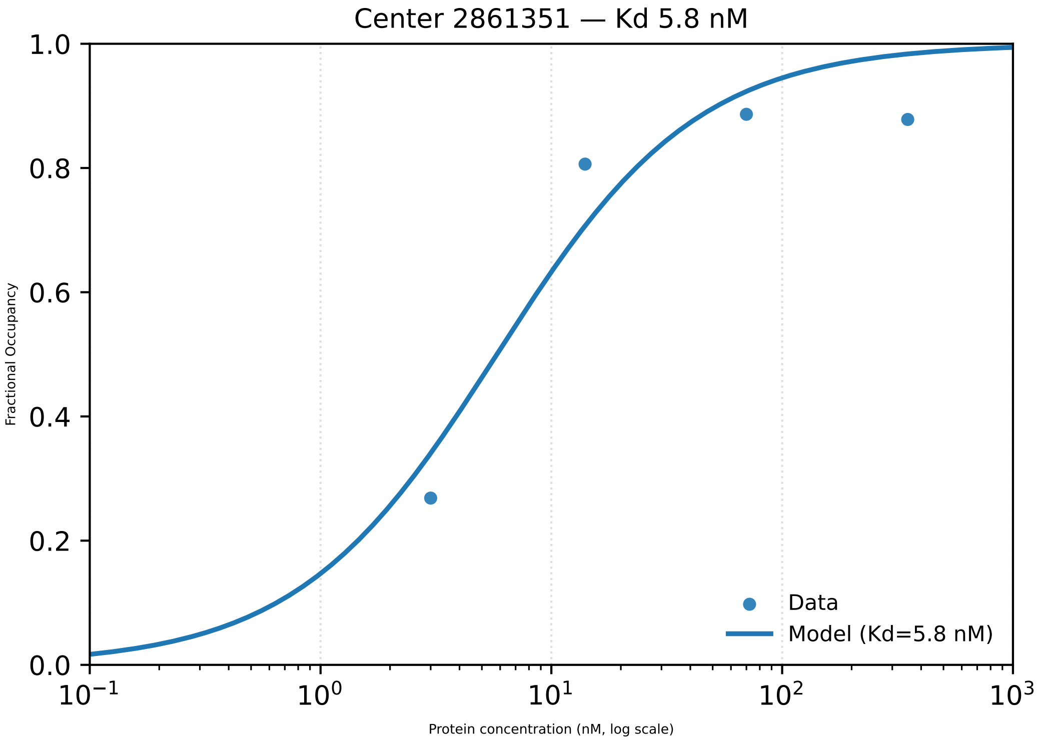

Center 2861351 — Kd 5.8 nM
Fractional Occupancy
Protein concentration (nM, log scale)
Data
Model (Kd=5.8 nM)
0.0
0.2
0.4
0.6
0.8
1.0
$10^{-1}$
$10^{0}$
$10^{1}$
$10^{2}$
$10^{3}$

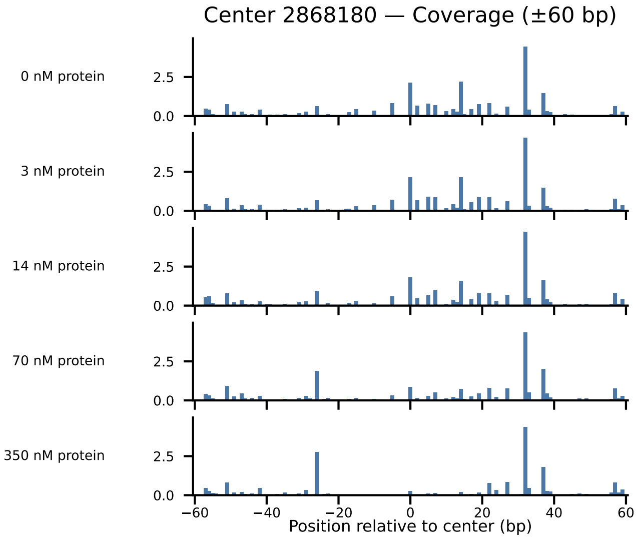

Center 2868180 — Coverage (±60 bp)
0 nM protein
3 nM protein
14 nM protein
70 nM protein
350 nM protein
2.5
0.0
−60
−40
−20
0
20
40
60
Position relative to center (bp)

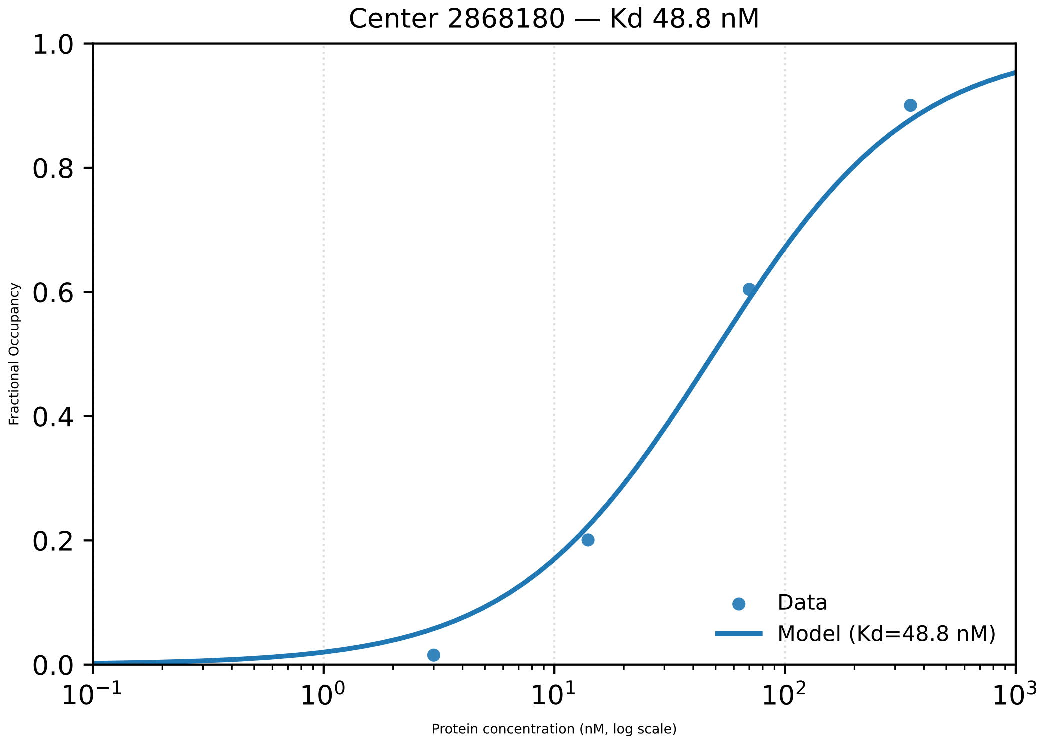

Center 2868180 — Kd 48.8 nM
Fractional Occupancy
1.0
0.8
0.6
0.4
0.2
0.0
$10^{-1}$
$10^{0}$
$10^{1}$
$10^{2}$
$10^{3}$
Protein concentration (nM, log scale)
Data
Model (Kd=48.8 nM)

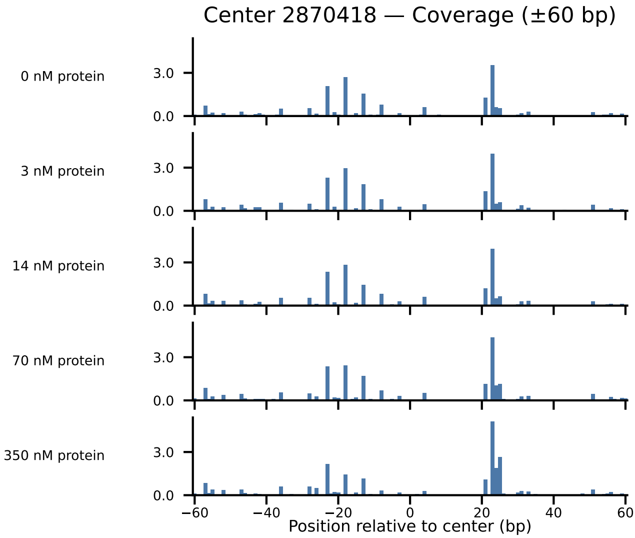

Center 2870418 — Coverage (±60 bp)
0 nM protein
3 nM protein
14 nM protein
70 nM protein
350 nM protein
3.0
0.0
−60
−40
−20
0
20
40
60
Position relative to center (bp)

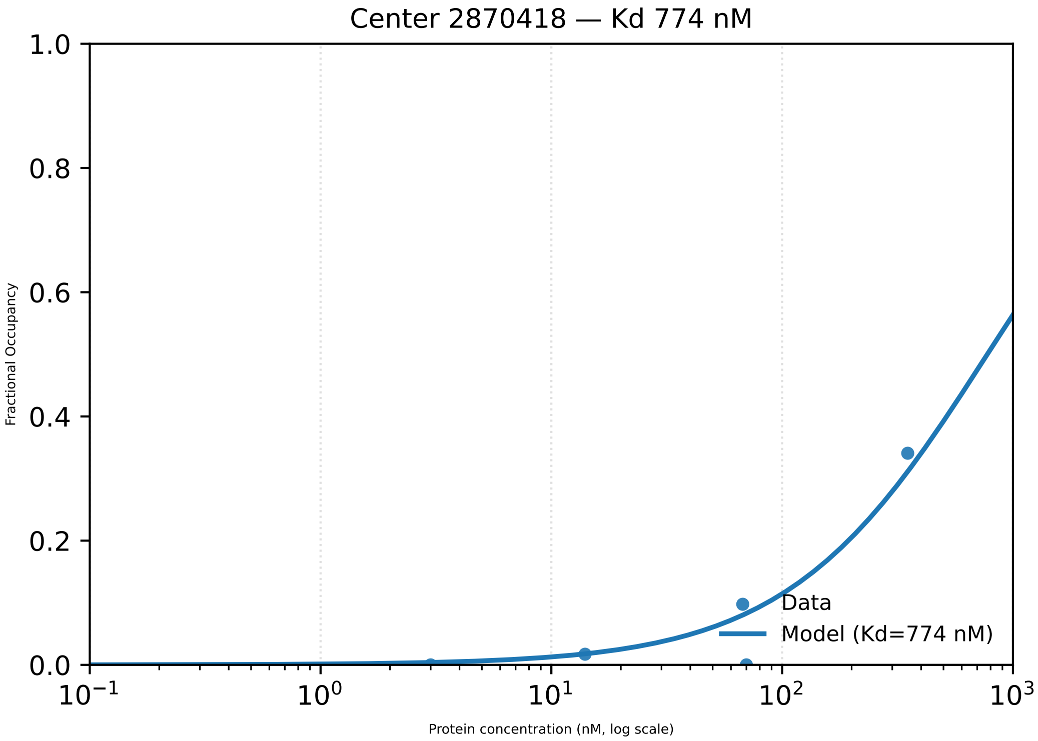

Center 2870418 — Kd 774 nM
1.0
0.8
0.6
0.4
0.2
0.0
Fractional Occupancy
Data
Model (Kd=774 nM)
$10^{-1}$
$10^{0}$
$10^{1}$
$10^{2}$
$10^{3}$
Protein concentration (nM, log scale)

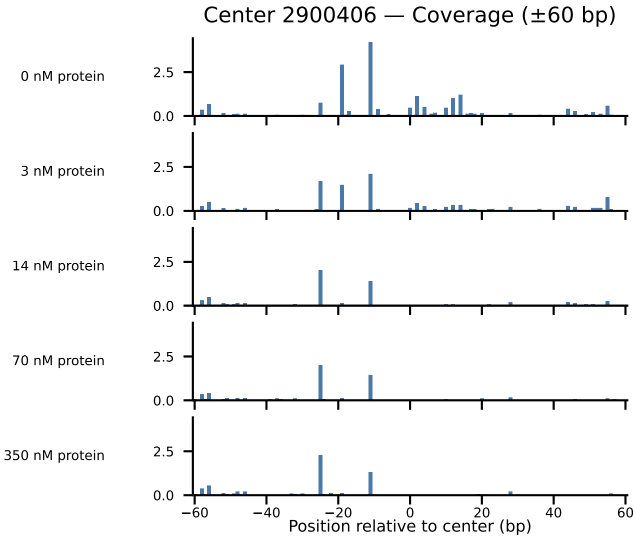
Center 2900406 — Coverage (±60 bp)
0 nM protein
3 nM protein
14 nM protein
70 nM protein
350 nM protein
2.5
0.0
−60
−40
−20
0
20
40
60
Position relative to center (bp)

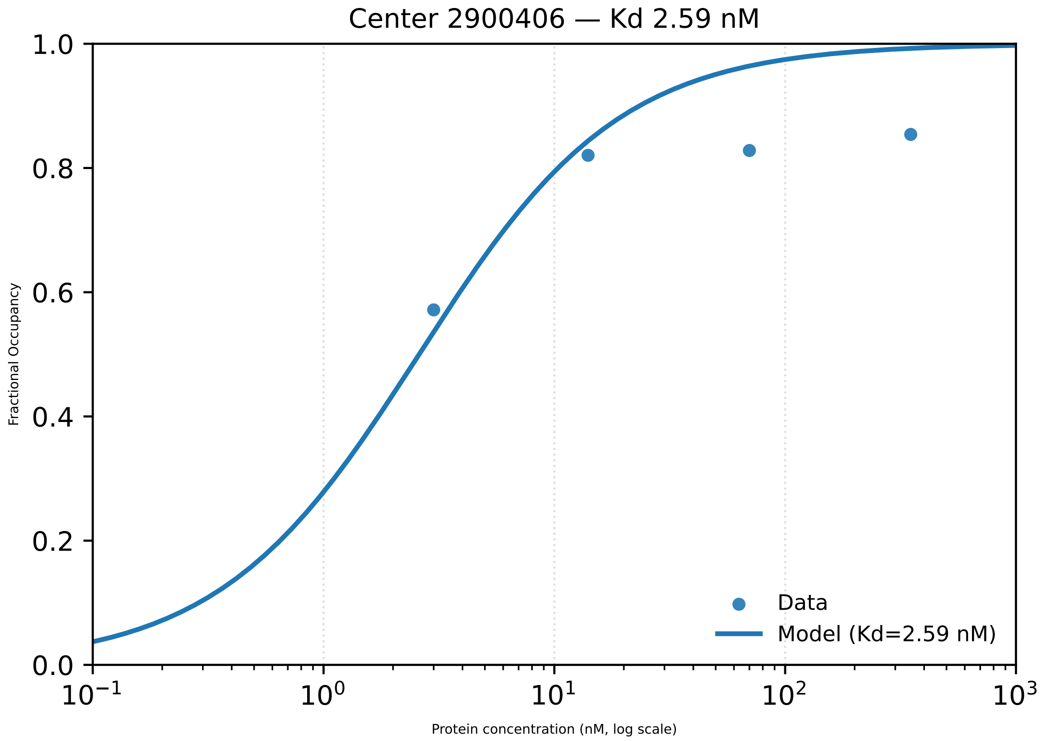

Center 2900406 — Kd 2.59 nM
Fractional Occupancy
Protein concentration (nM, log scale)
0.0
0.2
0.4
0.6
0.8
1.0
$10^{-1}$
$10^{0}$
$10^{1}$
$10^{2}$
$10^{3}$
Data
Model (Kd=2.59 nM)

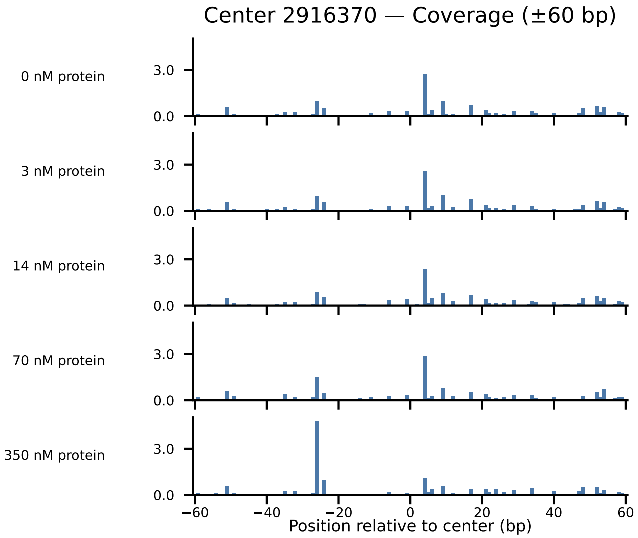

Center 2916370 — Coverage (±60 bp)
0 nM protein
3 nM protein
14 nM protein
70 nM protein
350 nM protein
3.0
0.0
−60
−40
−20
0
20
40
60
Position relative to center (bp)

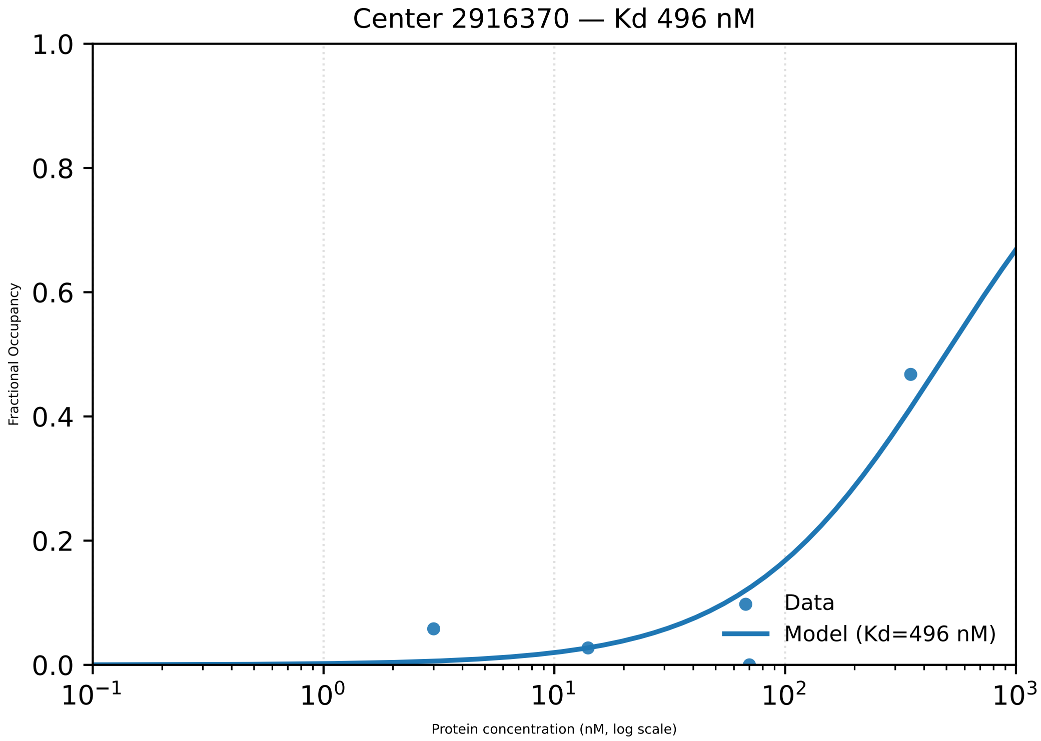

Center 2916370 — Kd 496 nM
1.0
0.8
0.6
0.4
0.2
0.0
Fractional Occupancy
$10^{-1}$
$10^{0}$
$10^{1}$
$10^{2}$
$10^{3}$
Protein concentration (nM, log scale)
Data
Model (Kd=496 nM)

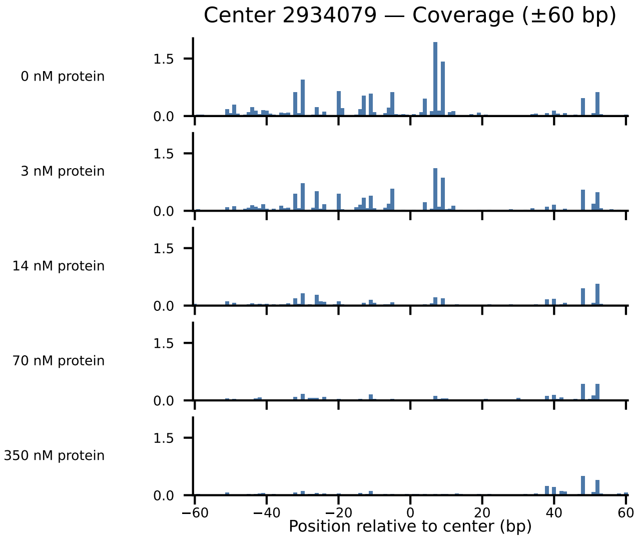

Center 2934079 — Coverage (±60 bp)
0 nM protein
3 nM protein
14 nM protein
70 nM protein
350 nM protein
1.5
0.0
−60
−40
−20
0
20
40
60
Position relative to center (bp)

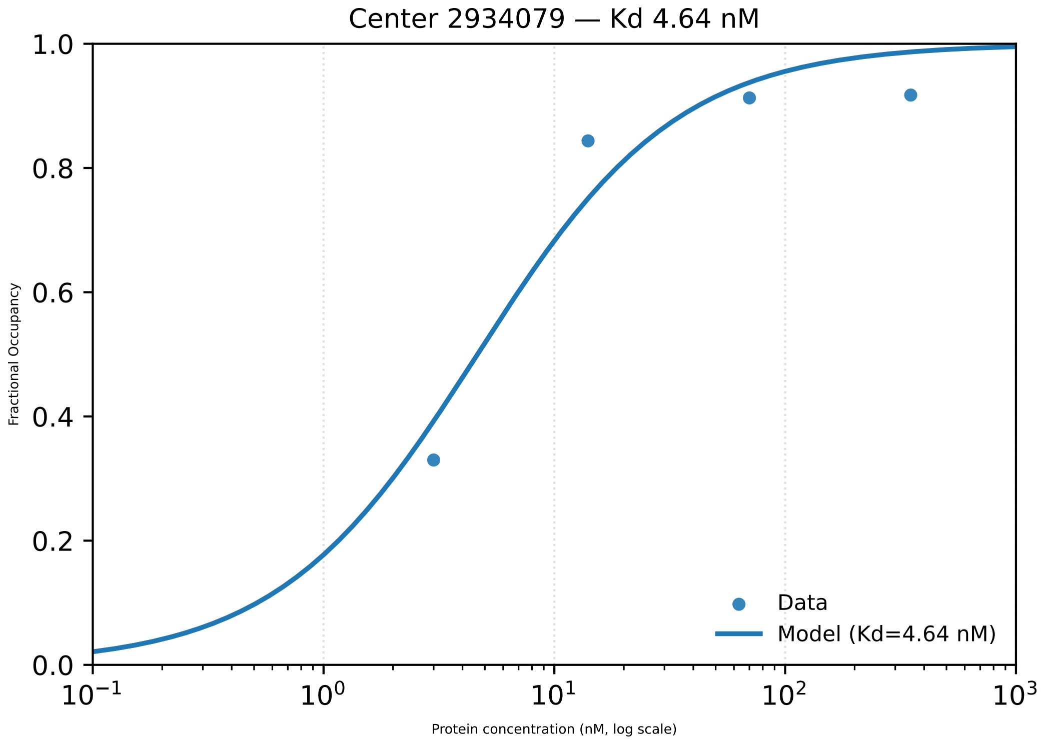
Center 2934079 — Kd 4.64 nM
1.0
0.8
0.6
0.4
0.2
0.0
Fractional Occupancy
$10^{-1}$
$10^{0}$
$10^{1}$
$10^{2}$
$10^{3}$
Protein concentration (nM, log scale)
Data
Model (Kd=4.64 nM)

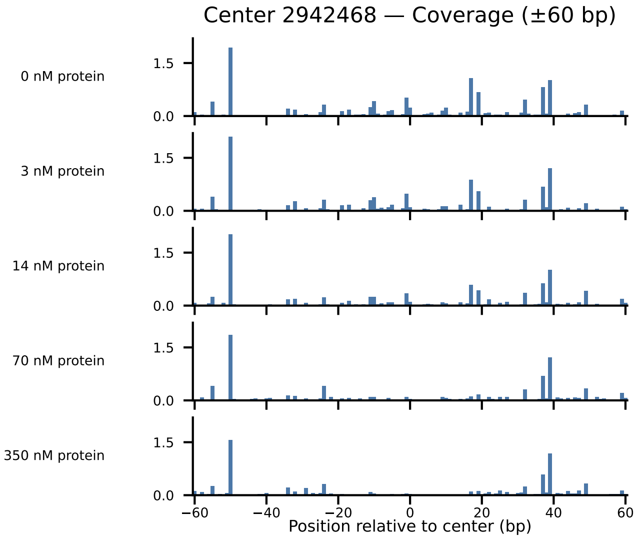

Center 2942468 — Coverage (±60 bp)
0 nM protein
3 nM protein
14 nM protein
70 nM protein
350 nM protein
1.5
0.0
−60
−40
−20
0
20
40
60
Position relative to center (bp)

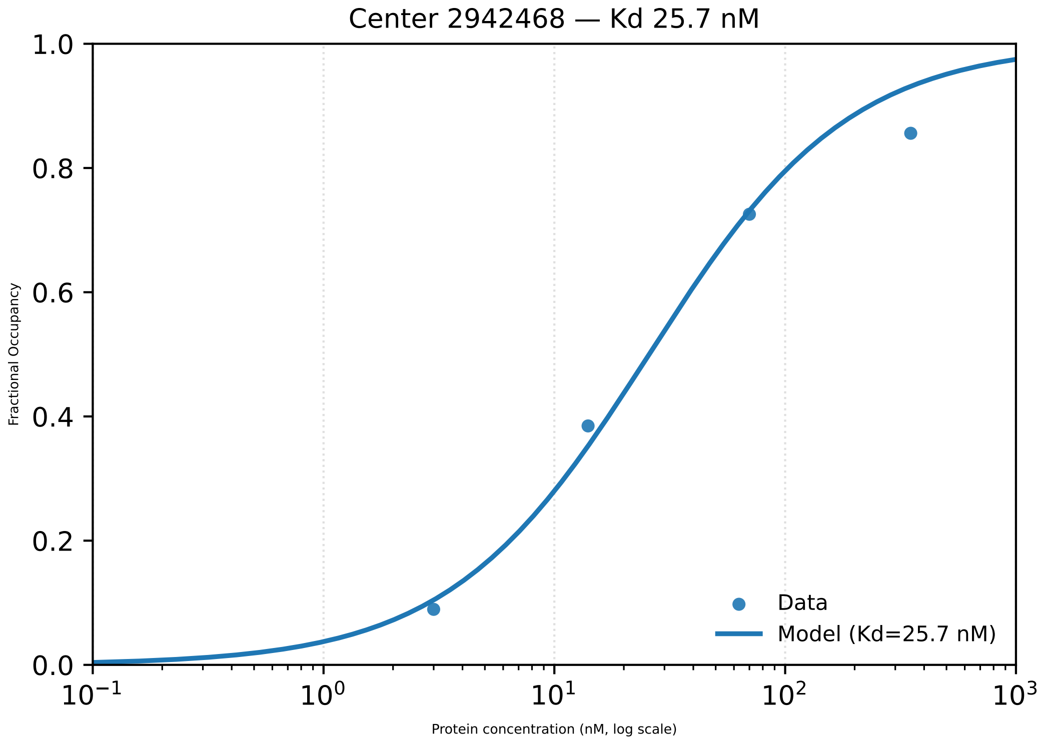

Center 2942468 — Kd 25.7 nM
1.0
0.8
0.6
0.4
0.2
0.0
Fractional Occupancy
$10^{-1}$
$10^{0}$
$10^{1}$
$10^{2}$
$10^{3}$
Protein concentration (nM, log scale)
Data
Model (Kd=25.7 nM)

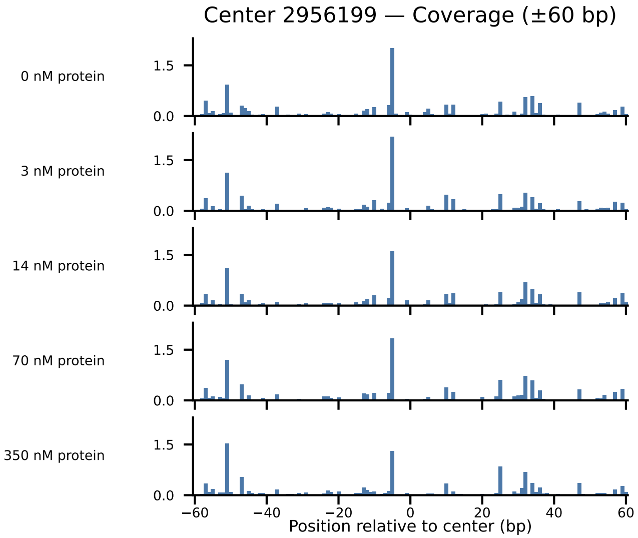
Center 2956199 — Coverage (±60 bp)
0 nM protein
3 nM protein
14 nM protein
70 nM protein
350 nM protein
1.5
0.0
−60
−40
−20
0
20
40
60
Position relative to center (bp)

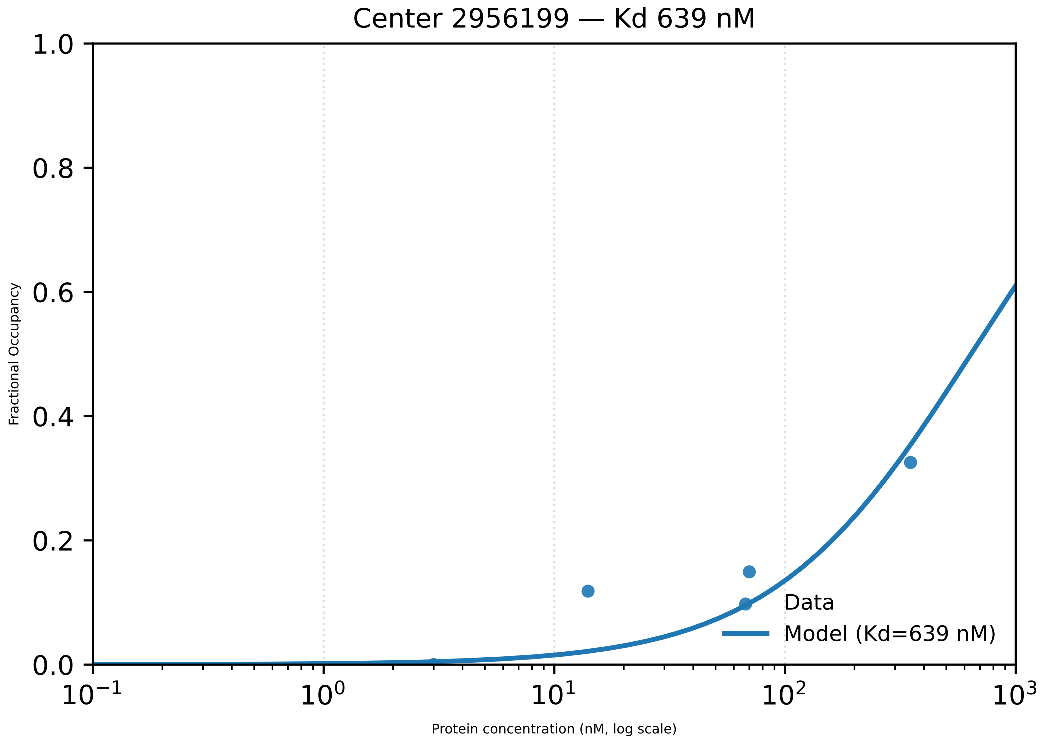

Center 2956199 — Kd 639 nM
Fractional Occupancy
Protein concentration (nM, log scale)
Data
Model (Kd=639 nM)
1.0
0.8
0.6
0.4
0.2
0.0
$10^{-1}$
$10^{0}$
$10^{1}$
$10^{2}$
$10^{3}$

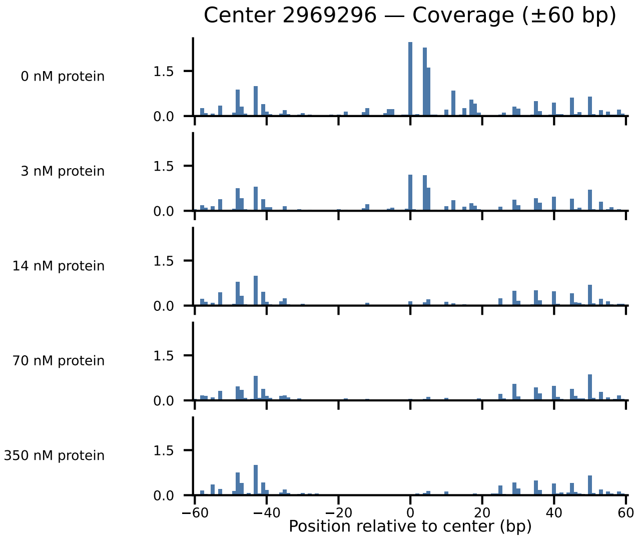

Center 2969296 — Coverage (±60 bp)
0 nM protein
3 nM protein
14 nM protein
70 nM protein
350 nM protein
1.5
0.0
−60
−40
−20
0
20
40
60
Position relative to center (bp)

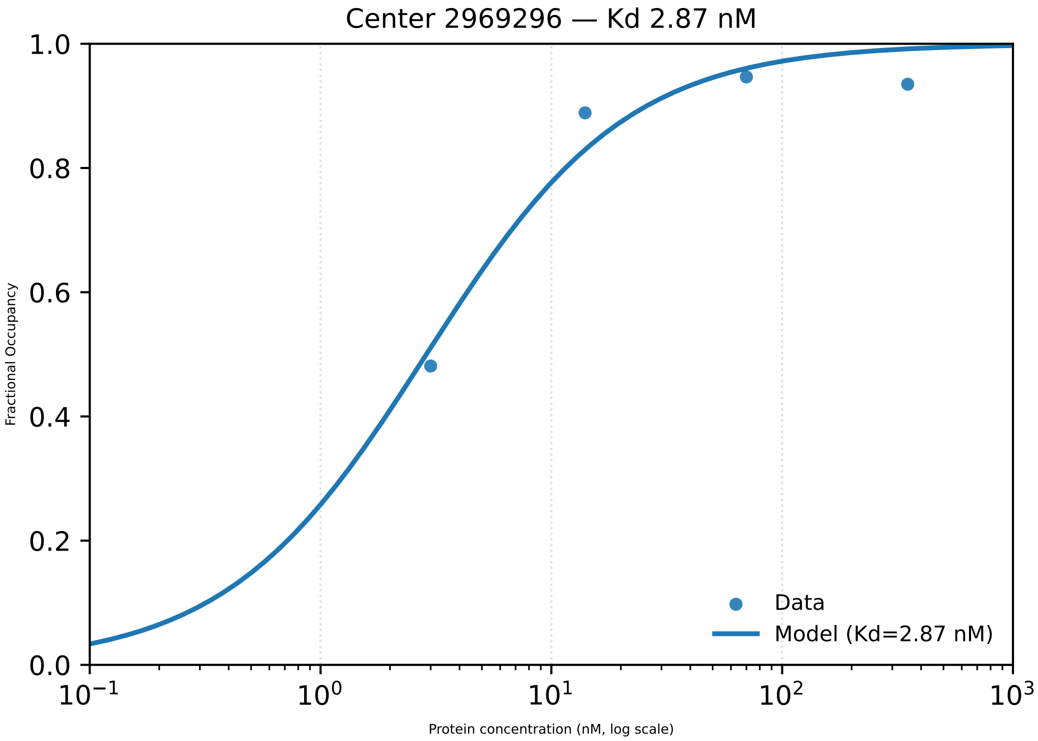

Center 2969296 — Kd 2.87 nM
1.0
0.8
0.6
0.4
0.2
0.0
Fractional Occupancy
$10^{-1}$
$10^{0}$
$10^{1}$
$10^{2}$
$10^{3}$
Protein concentration (nM, log scale)
Data
Model (Kd=2.87 nM)

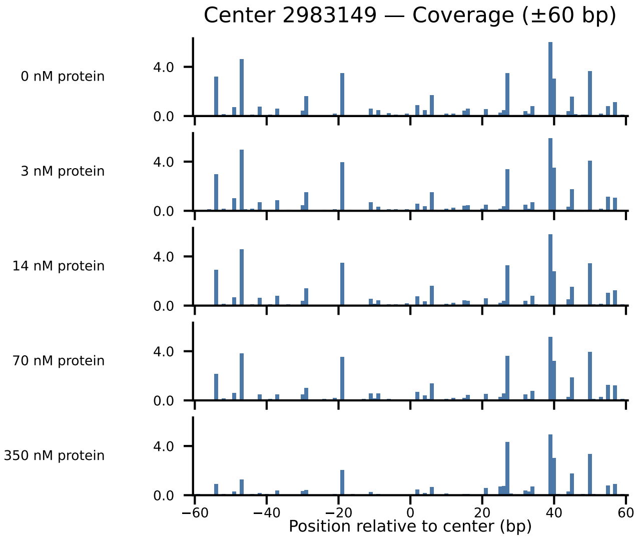

Center 2983149 — Coverage (±60 bp)
0 nM protein
3 nM protein
14 nM protein
70 nM protein
350 nM protein
4.0
0.0
−60
−40
−20
0
20
40
60
Position relative to center (bp)

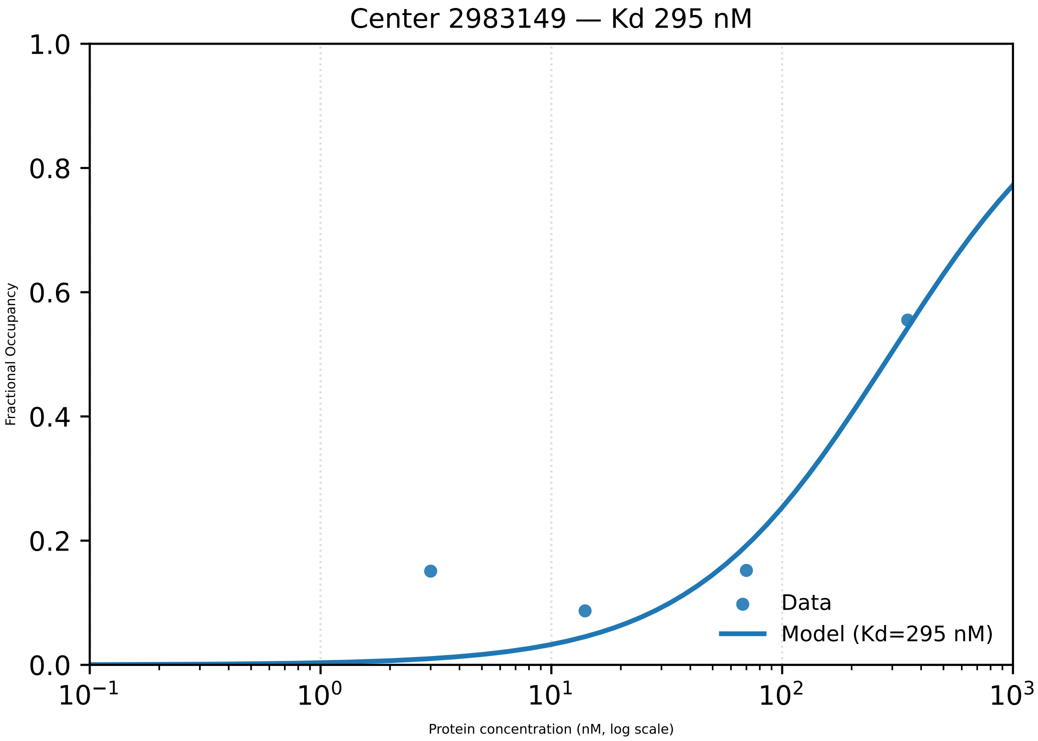

Center 2983149 — Kd 295 nM
1.0
0.8
0.6
0.4
0.2
0.0
Fractional Occupancy
$10^{-1}$
$10^{0}$
$10^{1}$
$10^{2}$
$10^{3}$
Protein concentration (nM, log scale)
Data
Model (Kd=295 nM)

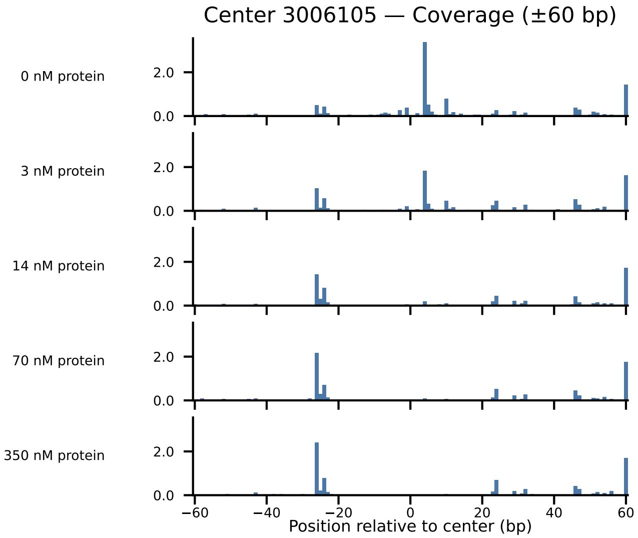

Center 3006105 — Coverage (±60 bp)
0 nM protein
3 nM protein
14 nM protein
70 nM protein
350 nM protein
2.0
0.0
−60
−40
−20
0
20
40
60
Position relative to center (bp)

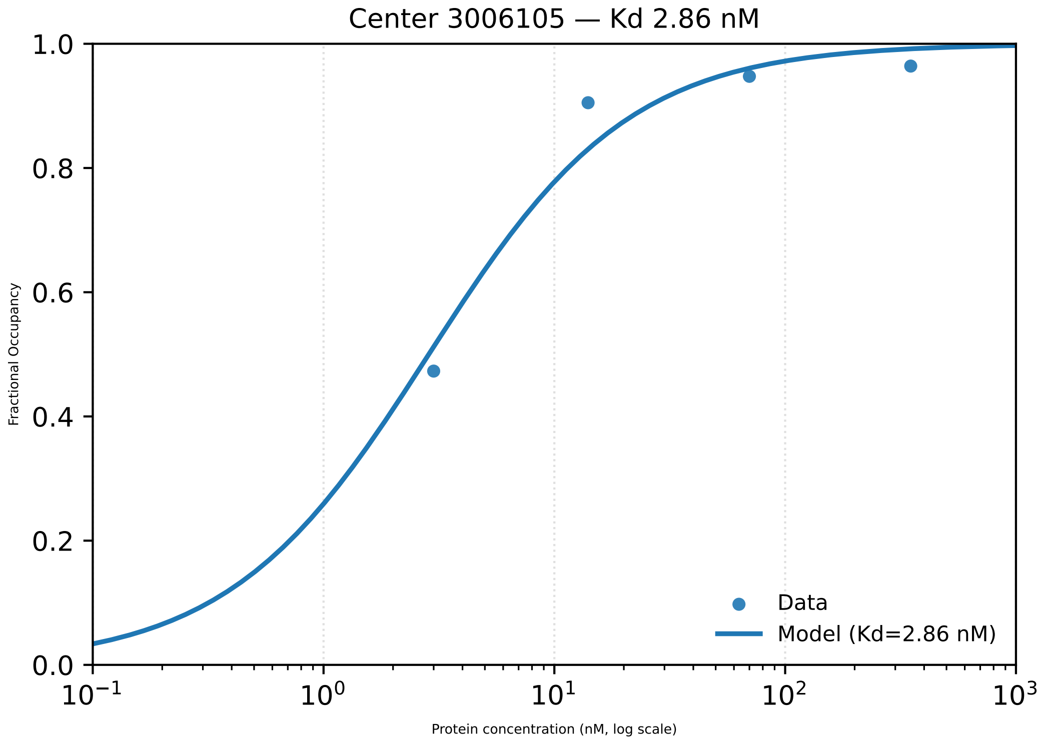

Center 3006105 — Kd 2.86 nM
Fractional Occupancy
1.0
0.8
0.6
0.4
0.2
0.0
$10^{-1}$
$10^{0}$
$10^{1}$
$10^{2}$
$10^{3}$
Protein concentration (nM, log scale)
Data
Model (Kd=2.86 nM)

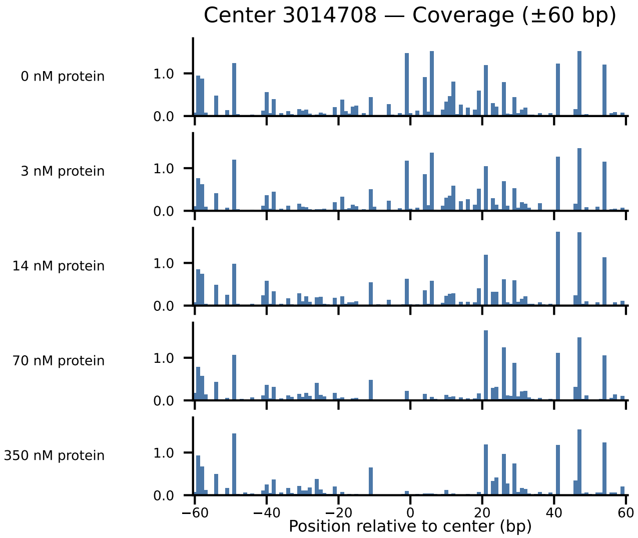

Center 3014708 — Coverage (±60 bp)
0 nM protein
3 nM protein
14 nM protein
70 nM protein
350 nM protein
1.0
0.0
−60
−40
−20
0
20
40
60
Position relative to center (bp)

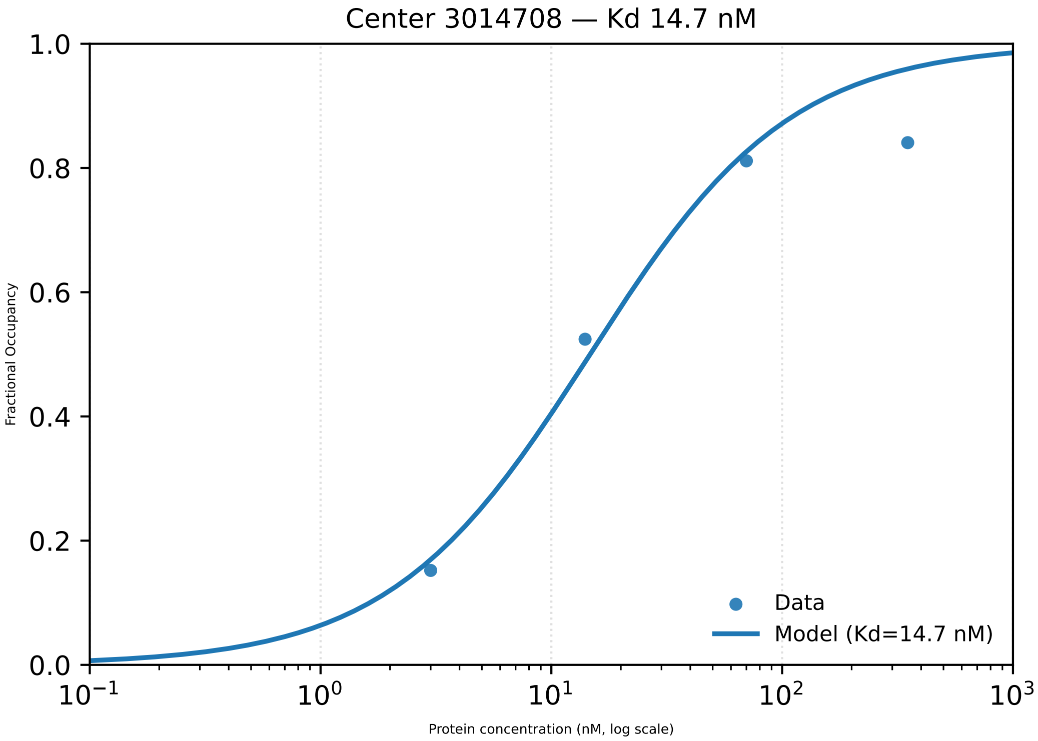

Center 3014708 — Kd 14.7 nM
Fractional Occupancy
1.0
0.8
0.6
0.4
0.2
0.0
$10^{-1}$
$10^{0}$
$10^{1}$
$10^{2}$
$10^{3}$
Protein concentration (nM, log scale)
Data
Model (Kd=14.7 nM)

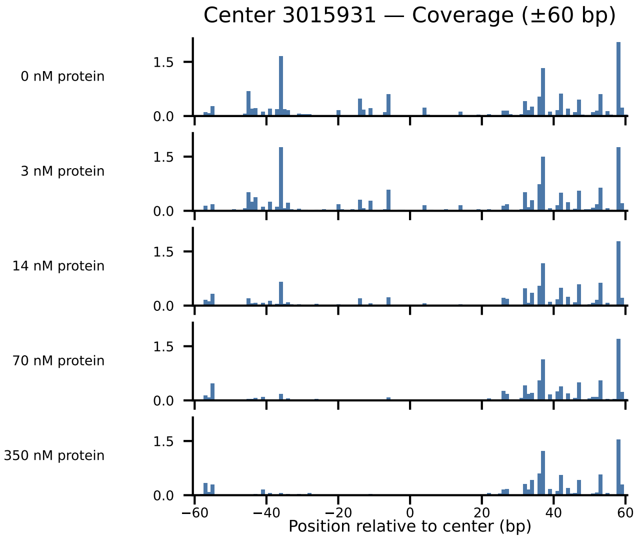

Center 3015931 — Coverage (±60 bp)
0 nM protein
3 nM protein
14 nM protein
70 nM protein
350 nM protein
1.5
0.0
−60
−40
−20
0
20
40
60
Position relative to center (bp)

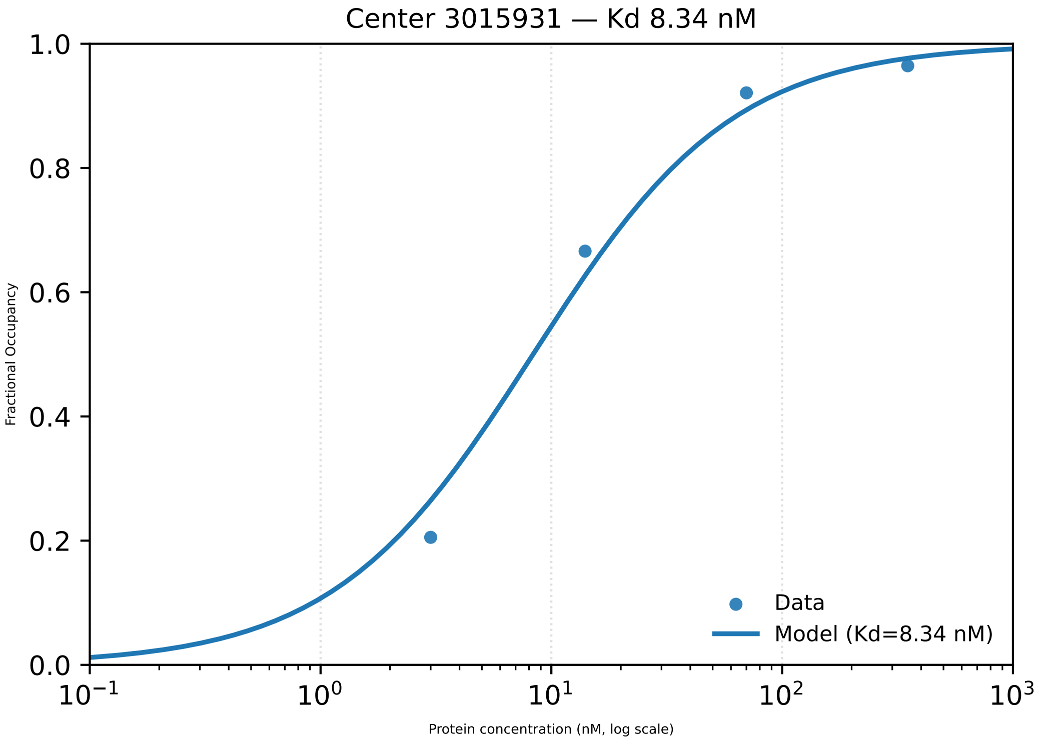

Center 3015931 — Kd 8.34 nM
Fractional Occupancy
1.0
0.8
0.6
0.4
0.2
0.0
$10^{-1}$
$10^{0}$
$10^{1}$
$10^{2}$
$10^{3}$
Protein concentration (nM, log scale)
Data
Model (Kd=8.34 nM)

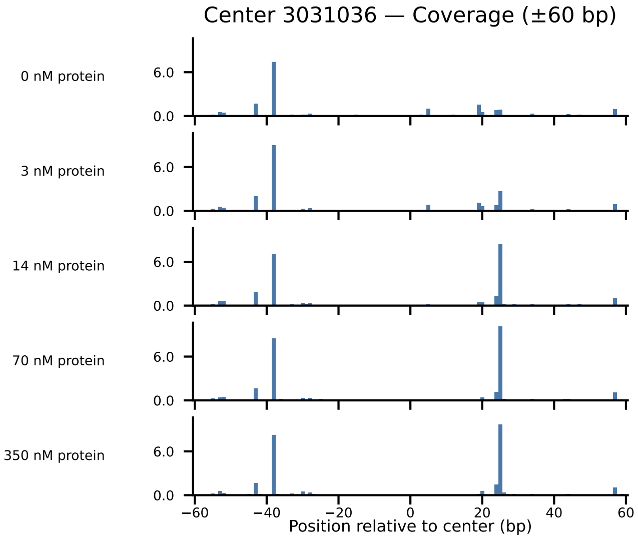

Center 3031036 — Coverage (±60 bp)
0 nM protein
3 nM protein
14 nM protein
70 nM protein
350 nM protein
6.0
0.0
−60
−40
−20
0
20
40
60
Position relative to center (bp)

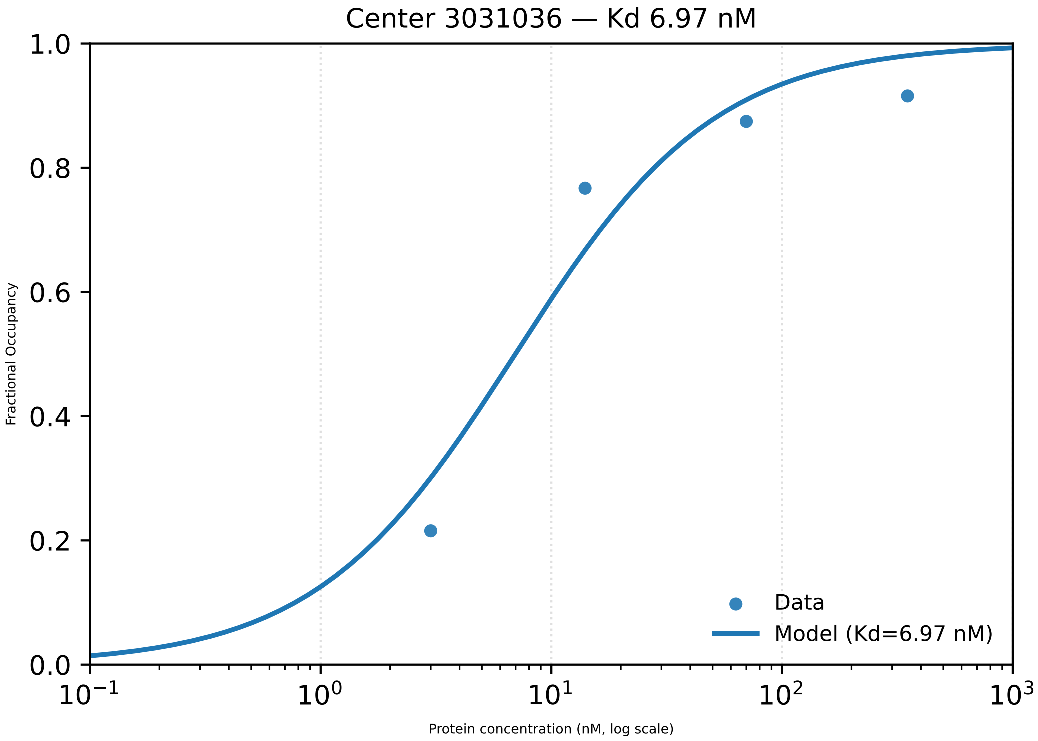

Center 3031036 — Kd 6.97 nM
Fractional Occupancy
1.0
0.8
0.6
0.4
0.2
0.0
$10^{-1}$
$10^{0}$
$10^{1}$
$10^{2}$
$10^{3}$
Protein concentration (nM, log scale)
Data
Model (Kd=6.97 nM)

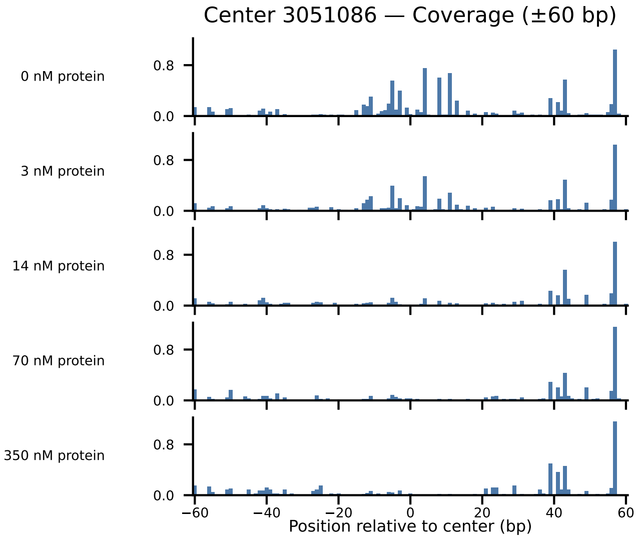

Center 3051086 — Coverage (±60 bp)
0 nM protein
3 nM protein
14 nM protein
70 nM protein
350 nM protein
0.8
0.0
−60
−40
−20
0
20
40
60
Position relative to center (bp)

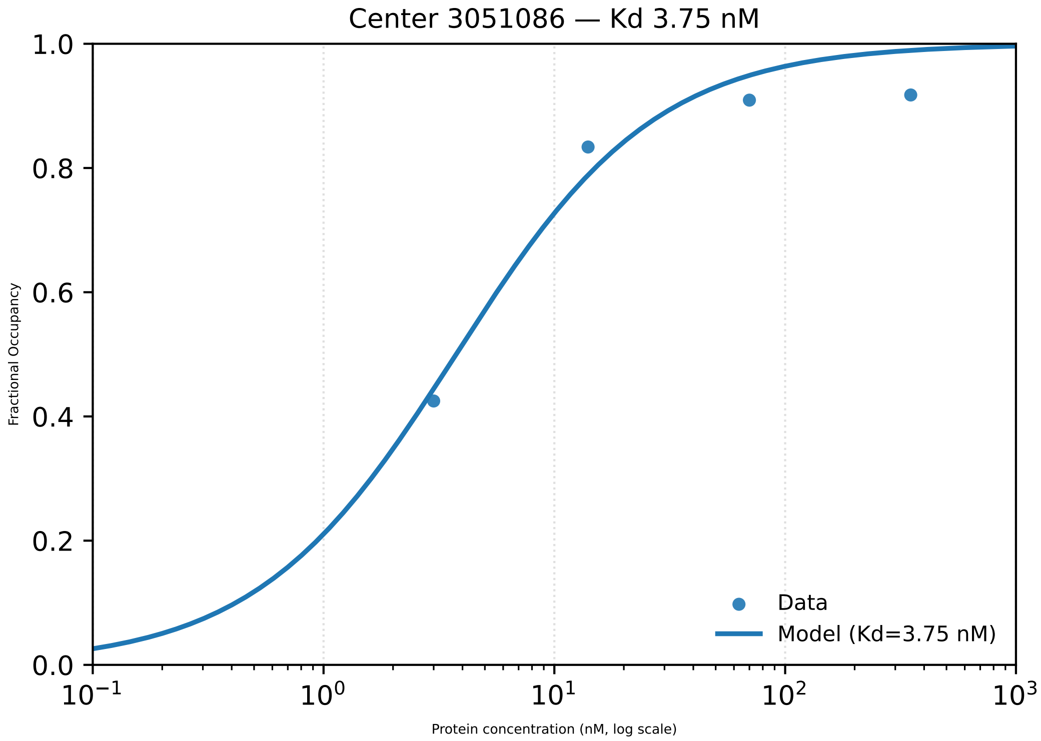

Center 3051086 — Kd 3.75 nM
1.0
0.8
0.6
0.4
0.2
0.0
Fractional Occupancy
$10^{-1}$
$10^{0}$
$10^{1}$
$10^{2}$
$10^{3}$
Protein concentration (nM, log scale)
Data
Model (Kd=3.75 nM)

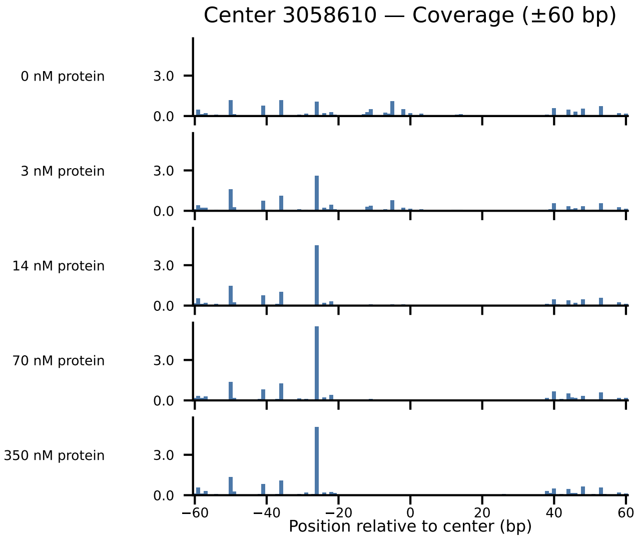
Center 3058610 — Coverage (±60 bp)
0 nM protein
3 nM protein
14 nM protein
70 nM protein
350 nM protein
3.0
0.0
−60
−40
−20
0
20
40
60
Position relative to center (bp)

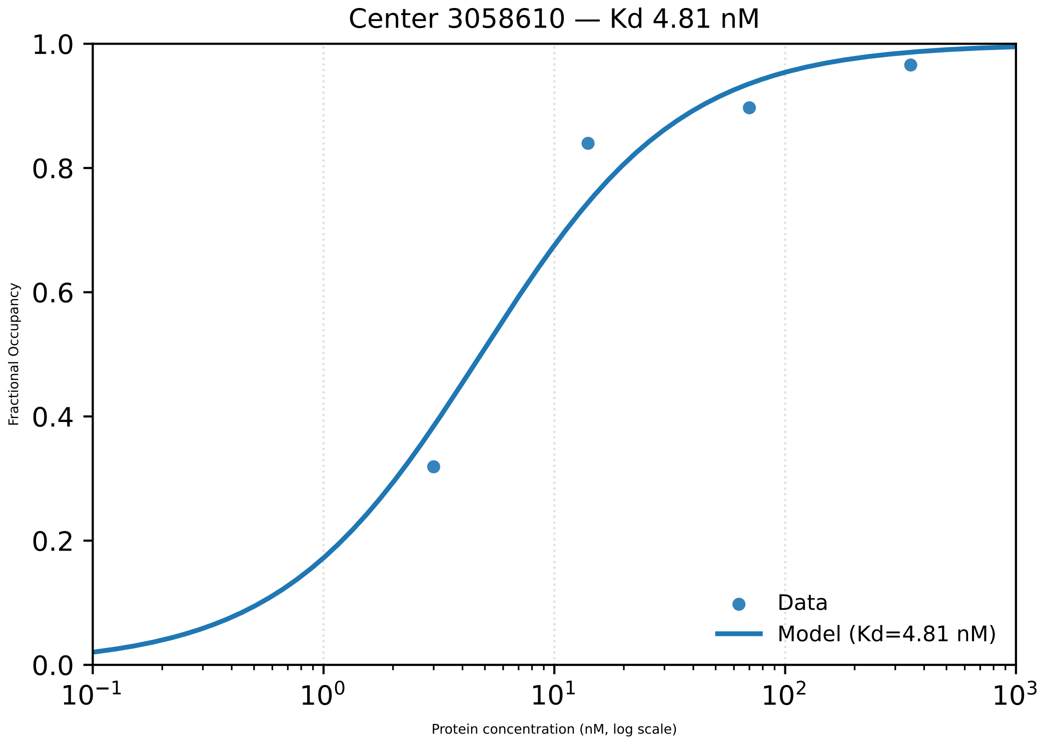
Center 3058610 — Kd 4.81 nM
Fractional Occupancy
Protein concentration (nM, log scale)
Data
Model (Kd=4.81 nM)

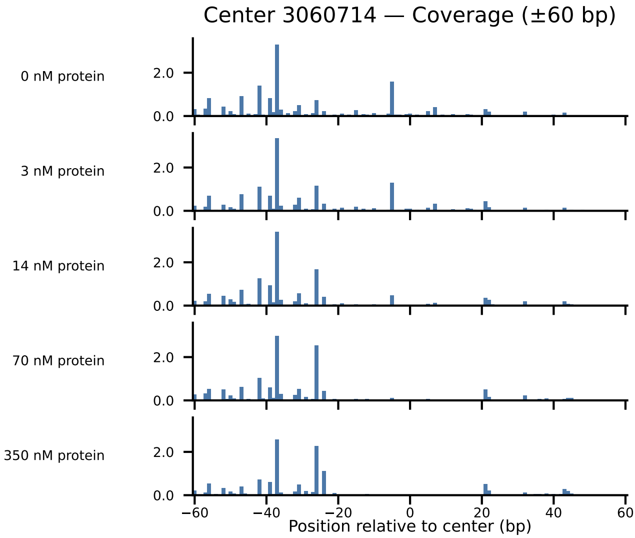

Center 3060714 — Coverage (±60 bp)
0 nM protein
3 nM protein
14 nM protein
70 nM protein
350 nM protein
2.0
0.0
−60
−40
−20
0
20
40
60
Position relative to center (bp)

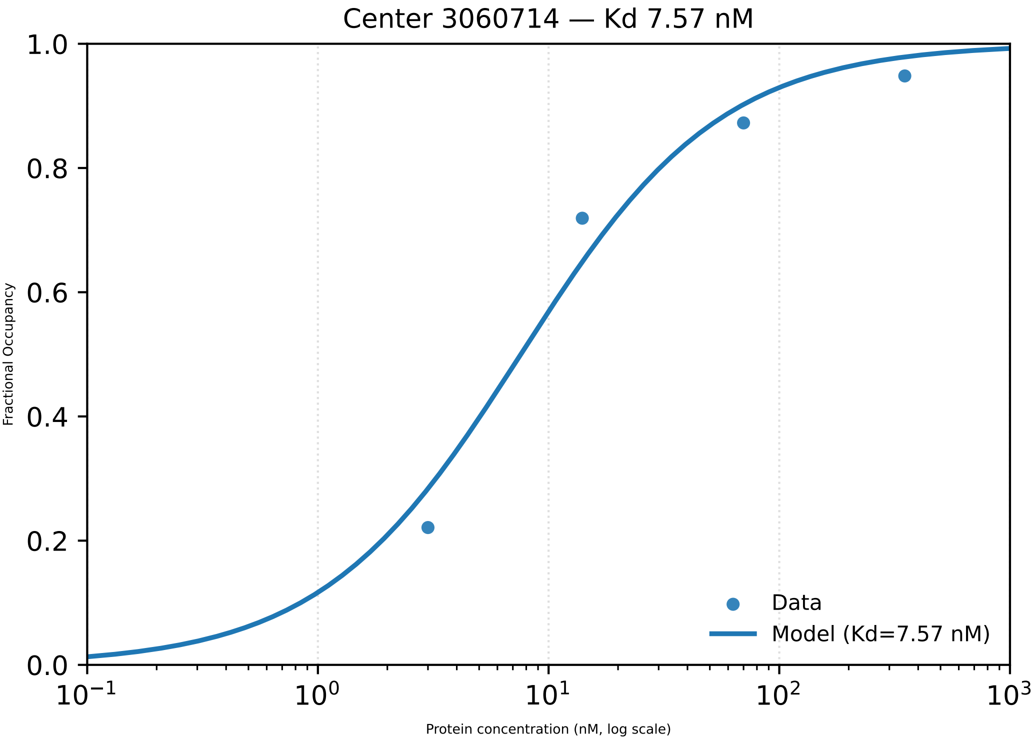

Center 3060714 — Kd 7.57 nM
Fractional Occupancy
1.0
0.8
0.6
0.4
0.2
0.0
$10^{-1}$
$10^{0}$
$10^{1}$
$10^{2}$
$10^{3}$
Protein concentration (nM, log scale)
Data
Model (Kd=7.57 nM)

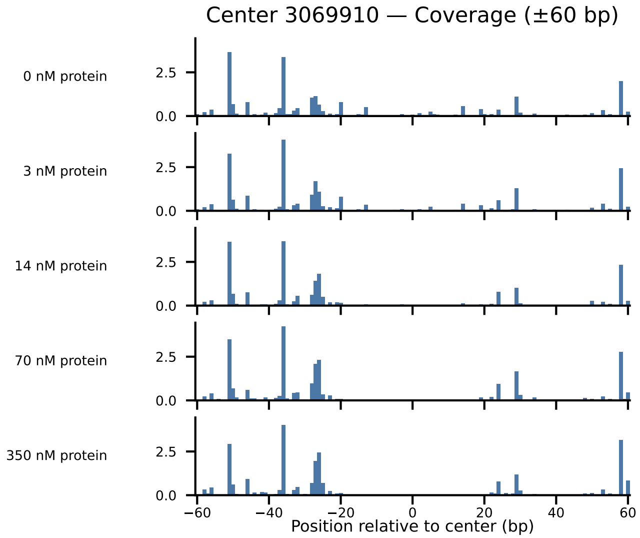

Center 3069910 — Coverage (±60 bp)
0 nM protein
3 nM protein
14 nM protein
70 nM protein
350 nM protein
2.5
0.0
−60
−40
−20
0
20
40
60
Position relative to center (bp)

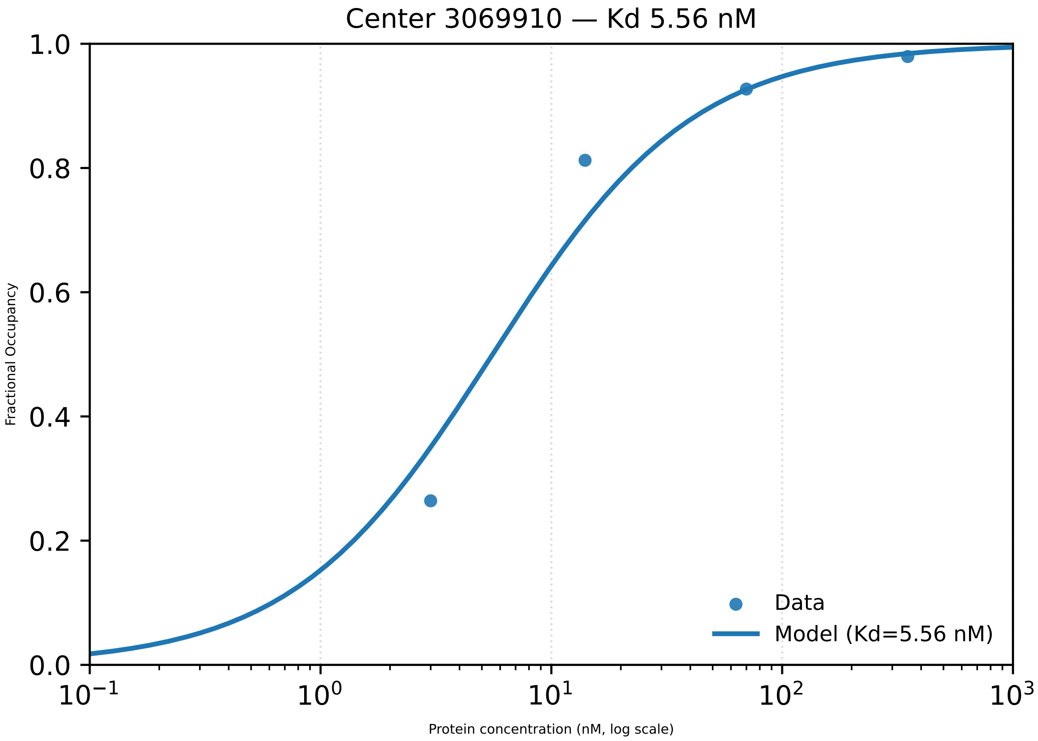

Center 3069910 — Kd 5.56 nM
Fractional Occupancy
1.0
0.8
0.6
0.4
0.2
0.0
$10^{-1}$
$10^{0}$
$10^{1}$
$10^{2}$
$10^{3}$
Protein concentration (nM, log scale)
Data
Model (Kd=5.56 nM)

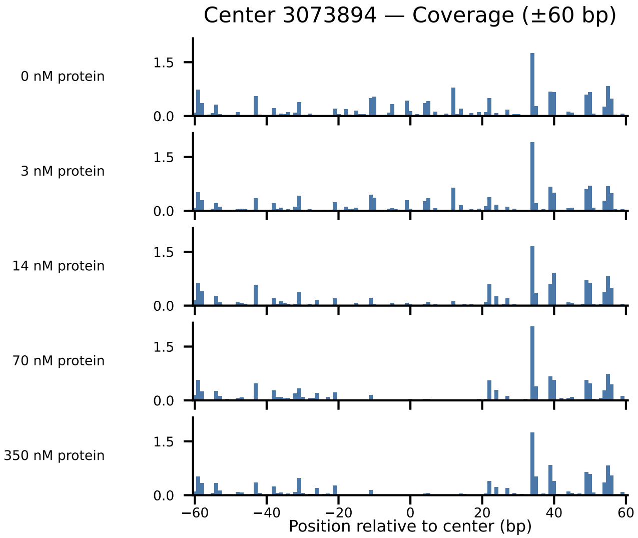

Center 3073894 — Coverage (±60 bp)
0 nM protein
3 nM protein
14 nM protein
70 nM protein
350 nM protein
1.5
0.0
−60
−40
−20
0
20
40
60
Position relative to center (bp)

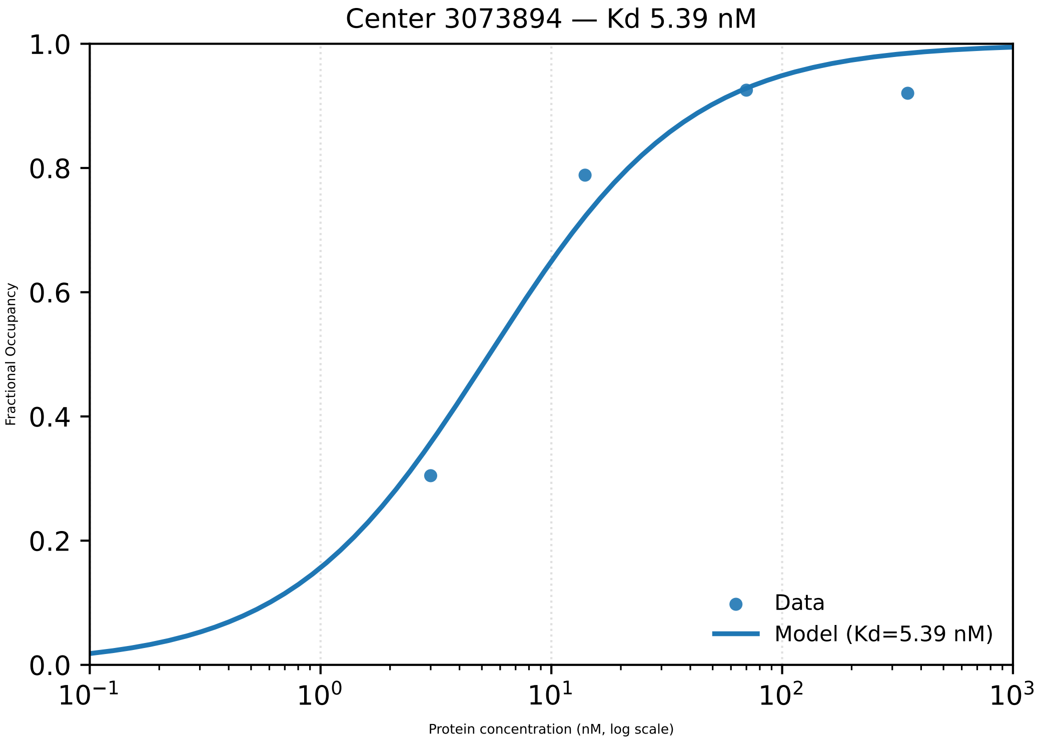

Center 3073894 — Kd 5.39 nM
1.0
0.8
0.6
0.4
0.2
0.0
Fractional Occupancy
10−1
100
101
102
103
Protein concentration (nM, log scale)
Data
Model (Kd=5.39 nM)

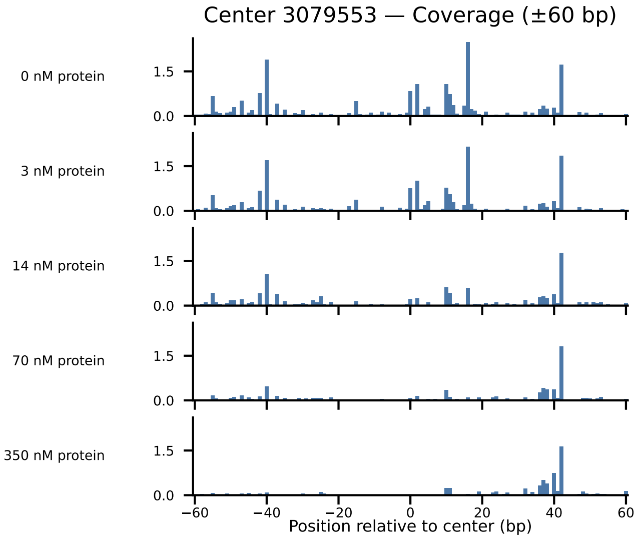

Center 3079553 — Coverage (±60 bp)
0 nM protein
3 nM protein
14 nM protein
70 nM protein
350 nM protein
1.5
0.0
−60
−40
−20
0
20
40
60
Position relative to center (bp)

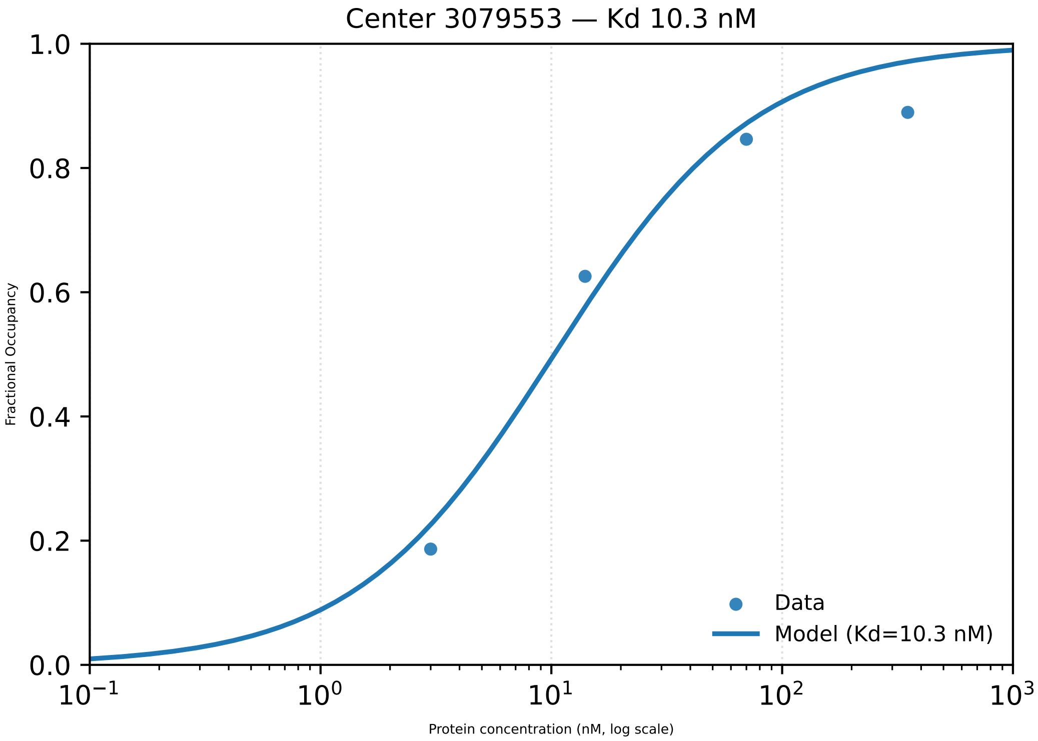

Center 3079553 — Kd 10.3 nM
Fractional Occupancy
1.0
0.8
0.6
0.4
0.2
0.0
$10^{-1}$
$10^{0}$
$10^{1}$
$10^{2}$
$10^{3}$
Protein concentration (nM, log scale)
Data
Model (Kd=10.3 nM)

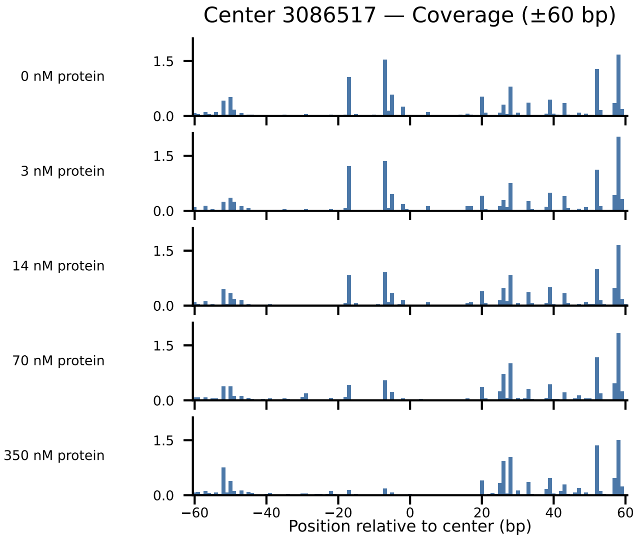

Center 3086517 — Coverage (±60 bp)
0 nM protein
3 nM protein
14 nM protein
70 nM protein
350 nM protein
1.5
0.0
−60
−40
−20
0
20
40
60
Position relative to center (bp)

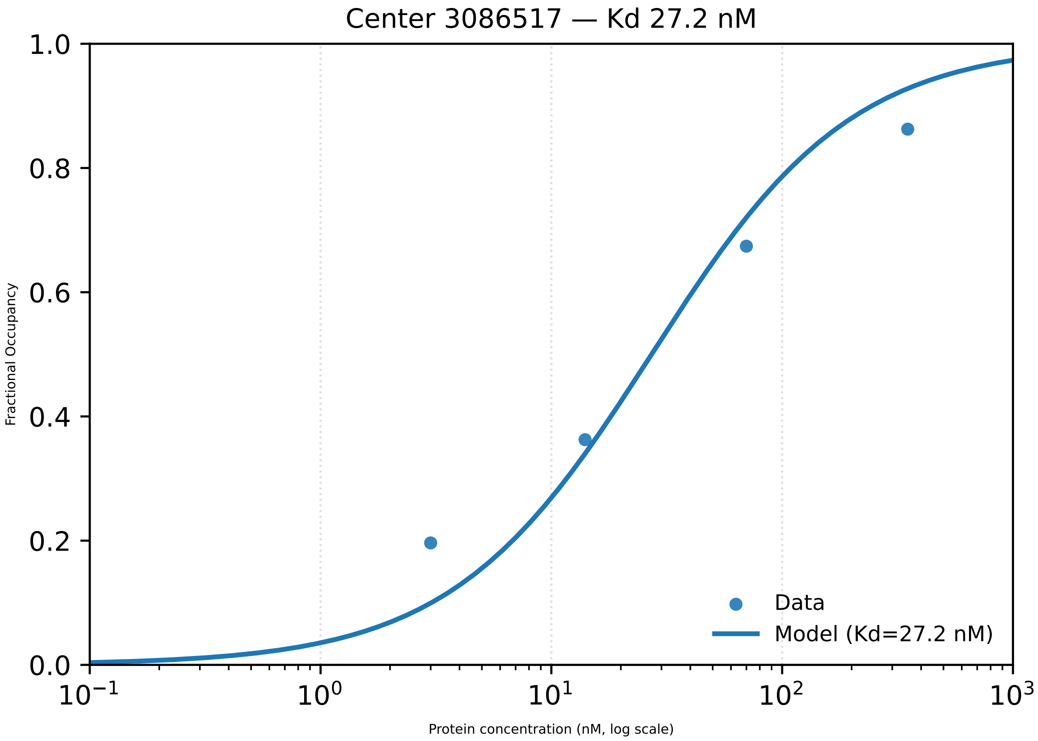
Center 3086517 — Kd 27.2 nM
1.0
0.8
0.6
0.4
0.2
0.0
Fractional Occupancy
$10^{-1}$
$10^{0}$
$10^{1}$
$10^{2}$
$10^{3}$
Protein concentration (nM, log scale)
Data
Model (Kd=27.2 nM)

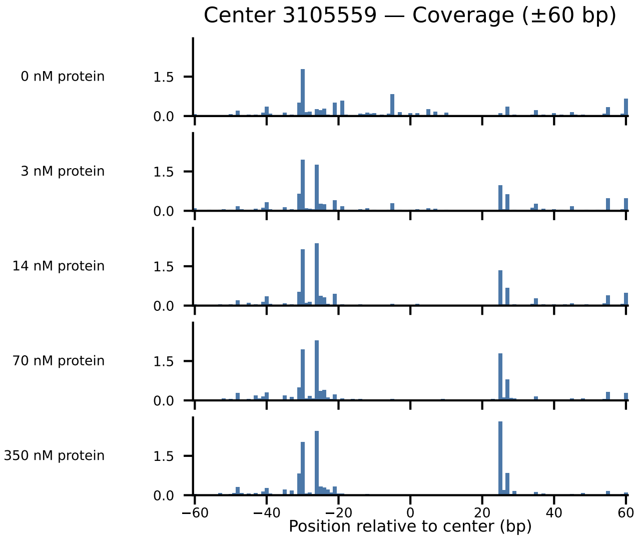
Center 3105559 — Coverage (±60 bp)
0 nM protein
3 nM protein
14 nM protein
70 nM protein
350 nM protein
1.5
0.0
−60
−40
−20
0
20
40
60
Position relative to center (bp)

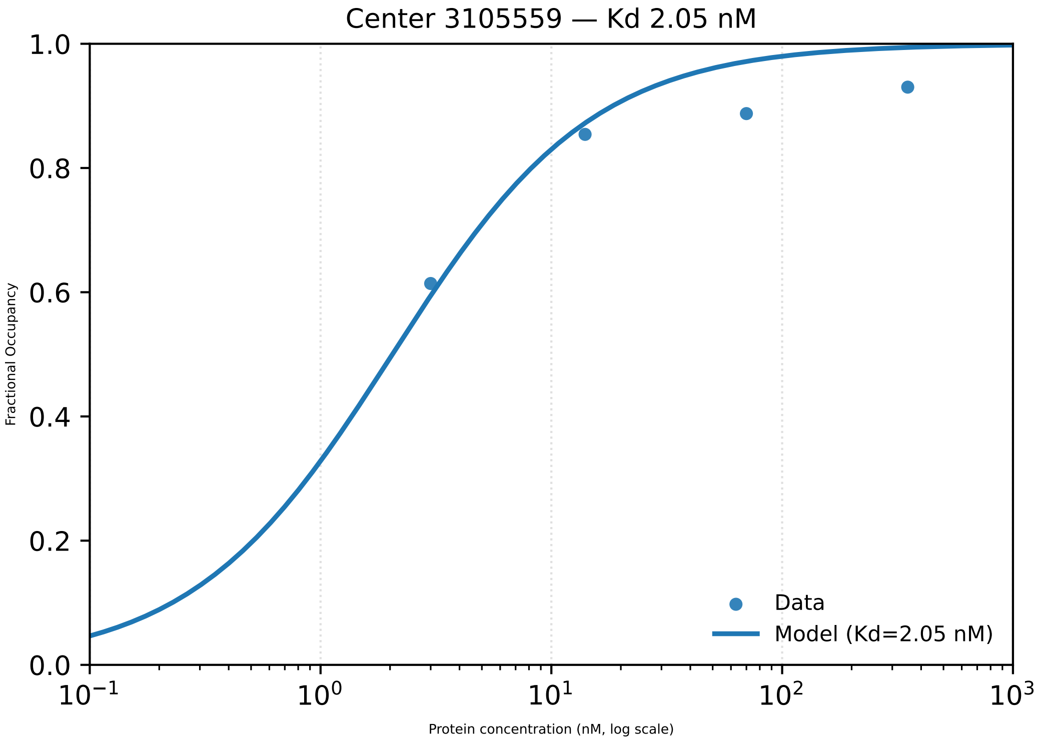

Center 3105559 — Kd 2.05 nM
Fractional Occupancy
1.0
0.8
0.6
0.4
0.2
0.0
$10^{-1}$
$10^{0}$
$10^{1}$
$10^{2}$
$10^{3}$
Protein concentration (nM, log scale)
Data
Model (Kd=2.05 nM)

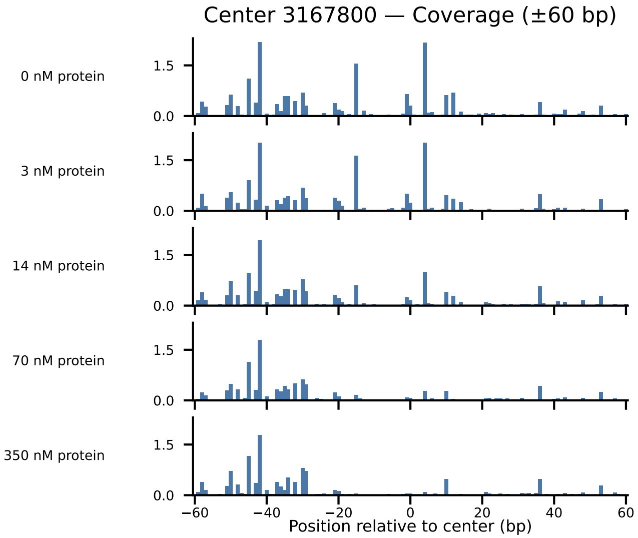

Center 3167800 — Coverage (±60 bp)
0 nM protein
3 nM protein
14 nM protein
70 nM protein
350 nM protein
1.5
0.0
−60
−40
−20
0
20
40
60
Position relative to center (bp)

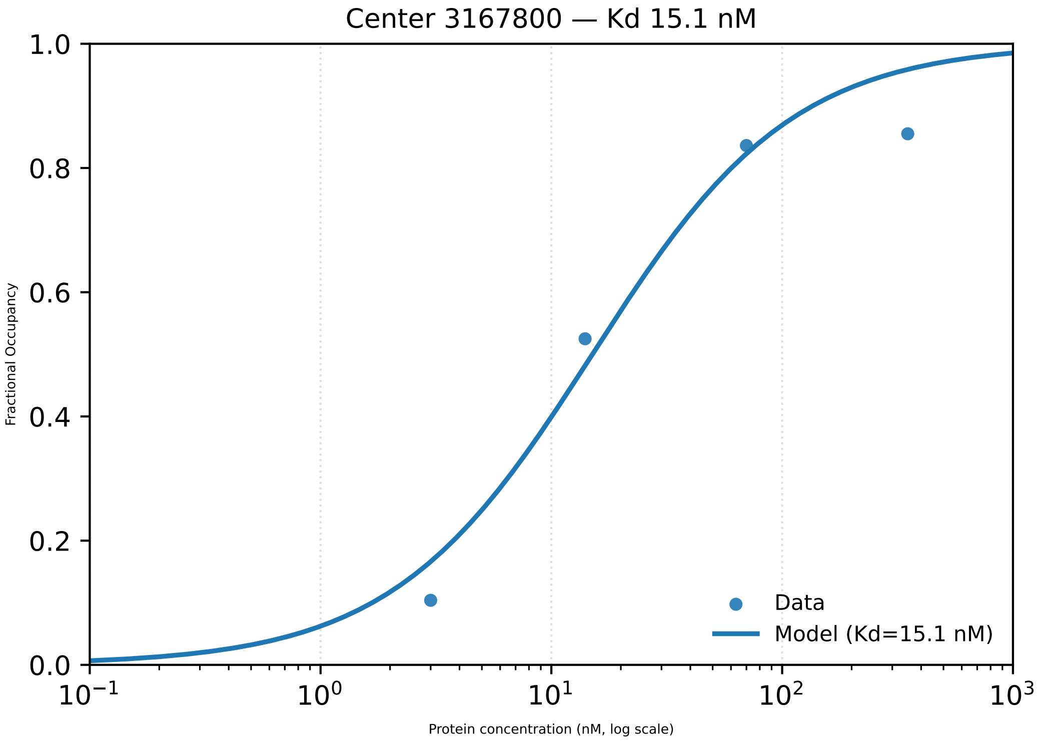
Center 3167800 — Kd 15.1 nM
Fractional Occupancy
1.0
0.8
0.6
0.4
0.2
0.0
$10^{-1}$
$10^{0}$
$10^{1}$
$10^{2}$
$10^{3}$
Protein concentration (nM, log scale)
Data
Model (Kd=15.1 nM)

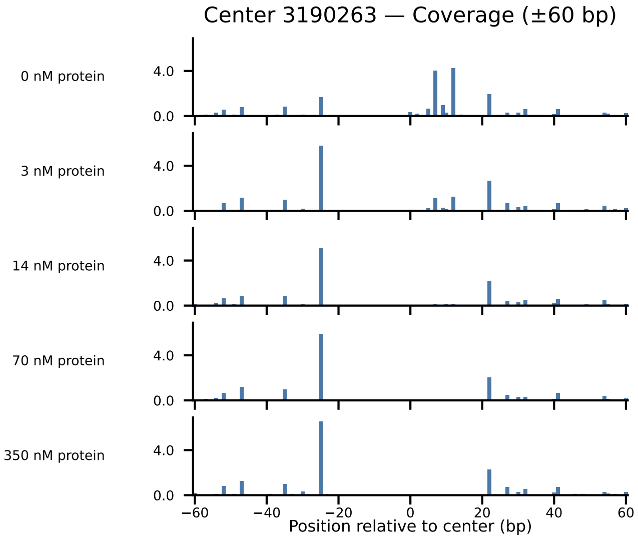

Center 3190263 — Coverage (±60 bp)
0 nM protein
3 nM protein
14 nM protein
70 nM protein
350 nM protein
4.0
0.0
−60
−40
−20
0
20
40
60
Position relative to center (bp)

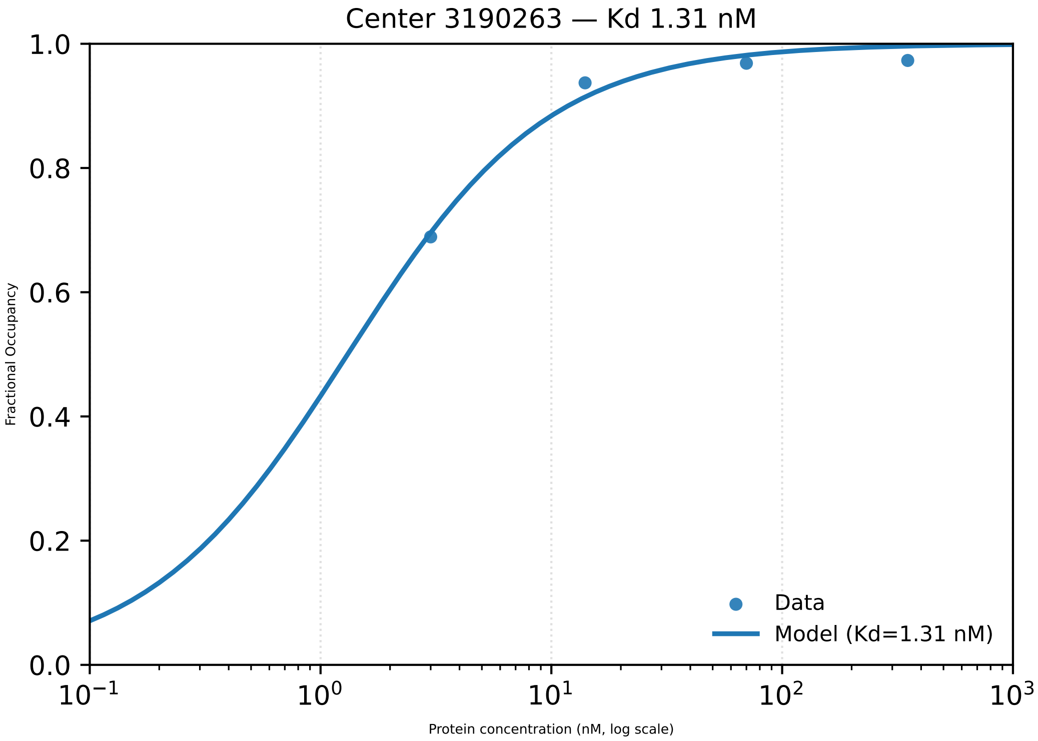

Center 3190263 — Kd 1.31 nM
1.0
0.8
0.6
0.4
0.2
0.0
Fractional Occupancy
$10^{-1}$
$10^{0}$
$10^{1}$
$10^{2}$
$10^{3}$
Protein concentration (nM, log scale)
Data
Model (Kd=1.31 nM)

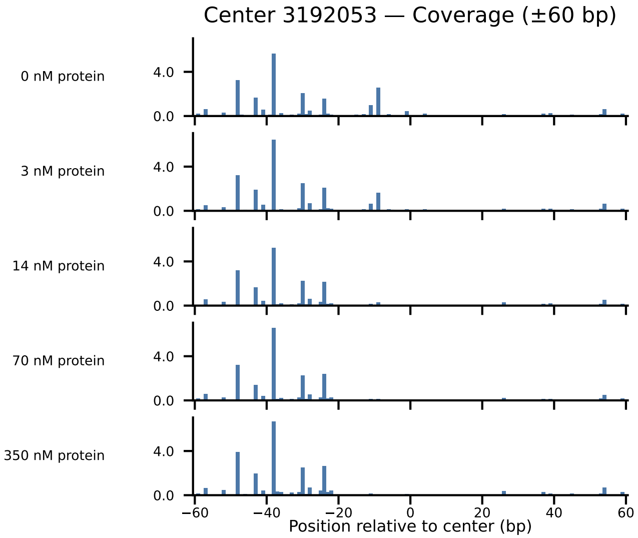

Center 3192053 — Coverage (±60 bp)
0 nM protein
3 nM protein
14 nM protein
70 nM protein
350 nM protein
4.0
0.0
−60
−40
−20
0
20
40
60
Position relative to center (bp)

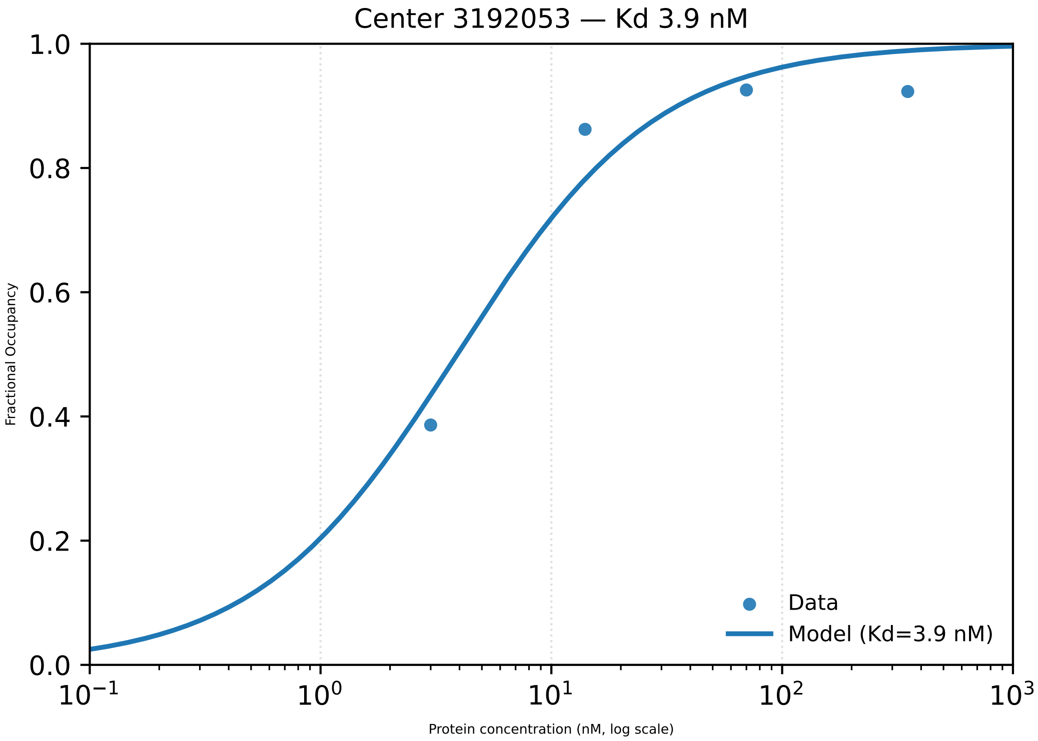

Center 3192053 — Kd 3.9 nM
Fractional Occupancy
1.0
0.8
0.6
0.4
0.2
0.0
$10^{-1}$
$10^{0}$
$10^{1}$
$10^{2}$
$10^{3}$
Protein concentration (nM, log scale)
Data
Model (Kd=3.9 nM)

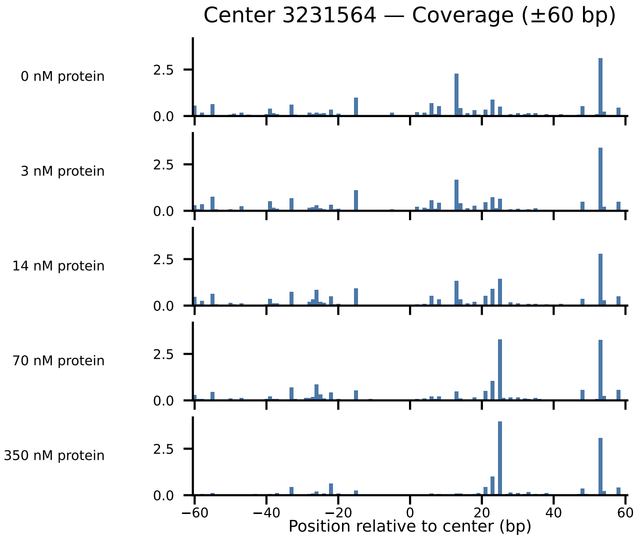

Center 3231564 — Coverage (±60 bp)
0 nM protein
3 nM protein
14 nM protein
70 nM protein
350 nM protein
0.0
2.5
−60
−40
−20
0
20
40
60
Position relative to center (bp)

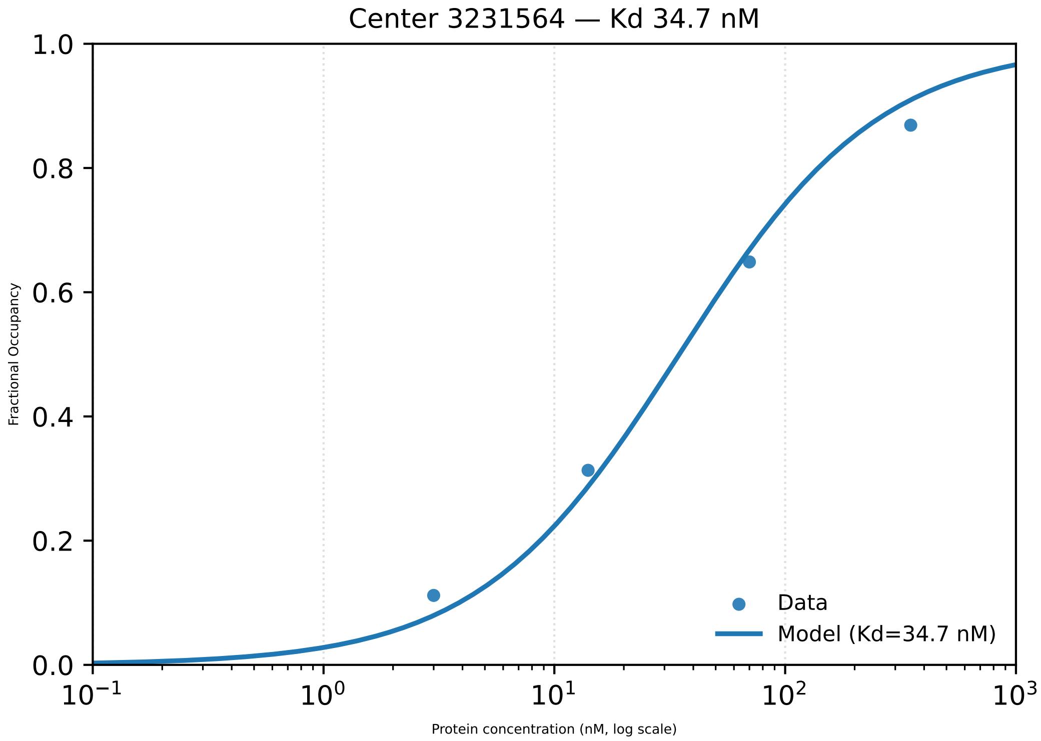

Center 3231564 — Kd 34.7 nM
1.0
0.8
0.6
0.4
0.2
0.0
Fractional Occupancy
$10^{-1}$
$10^{0}$
$10^{1}$
$10^{2}$
$10^{3}$
Protein concentration (nM, log scale)
Data
Model (Kd=34.7 nM)

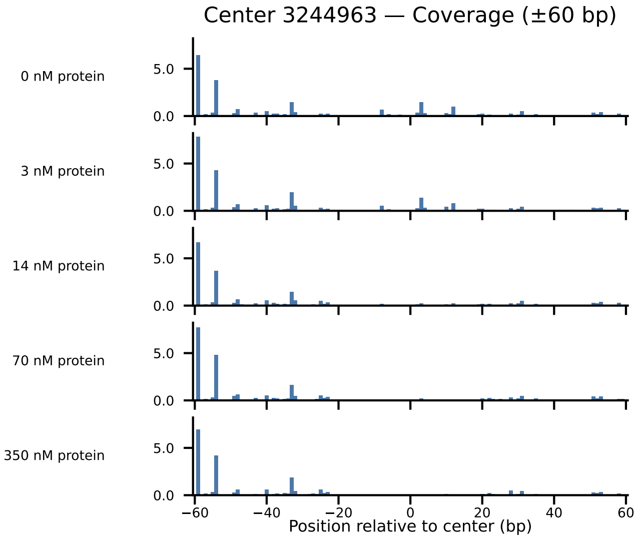
Center 3244963 — Coverage (±60 bp)
0 nM protein
3 nM protein
14 nM protein
70 nM protein
350 nM protein
5.0
0.0
−60
−40
−20
0
20
40
60
Position relative to center (bp)

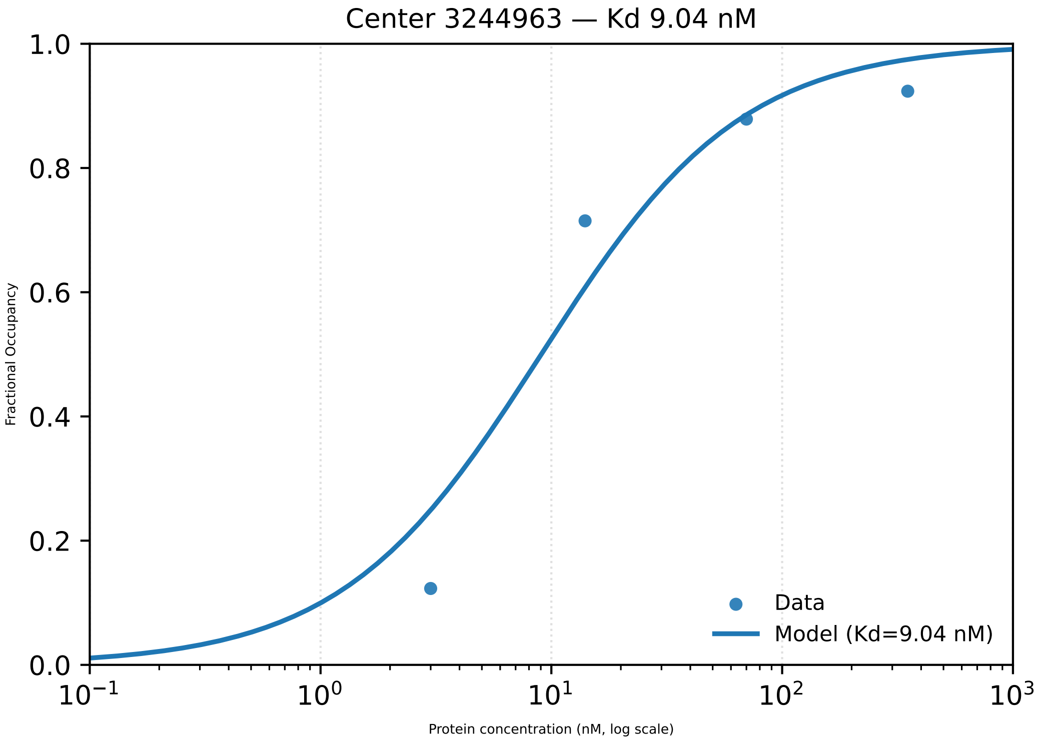

Center 3244963 — Kd 9.04 nM
Fractional Occupancy
0.0
0.2
0.4
0.6
0.8
1.0
$10^{-1}$
$10^{0}$
$10^{1}$
$10^{2}$
$10^{3}$
Protein concentration (nM, log scale)
Data
Model (Kd=9.04 nM)

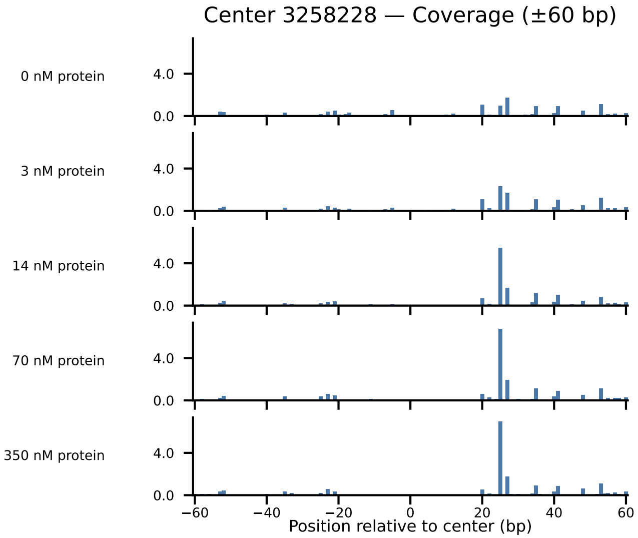

Center 3258228 — Coverage (±60 bp)
0 nM protein
3 nM protein
14 nM protein
70 nM protein
350 nM protein
4.0
0.0
−60
−40
−20
0
20
40
60
Position relative to center (bp)

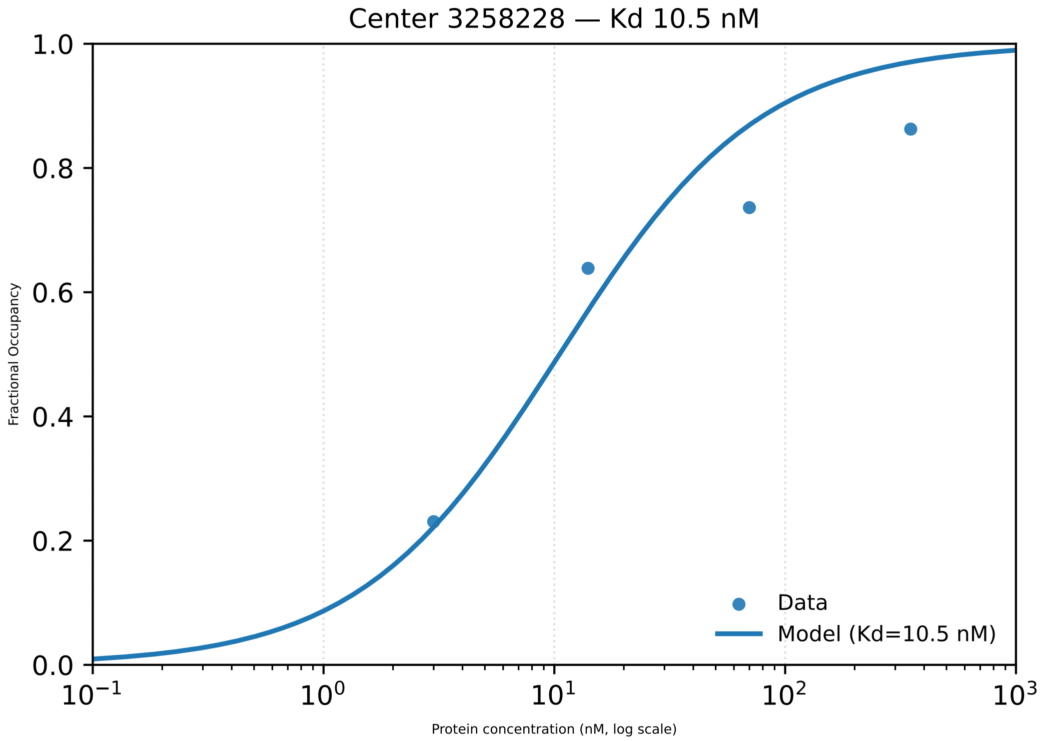

Center 3258228 — Kd 10.5 nM
Fractional Occupancy
1.0
0.8
0.6
0.4
0.2
0.0
$10^{-1}$
$10^{0}$
$10^{1}$
$10^{2}$
$10^{3}$
Protein concentration (nM, log scale)
Data
Model (Kd=10.5 nM)

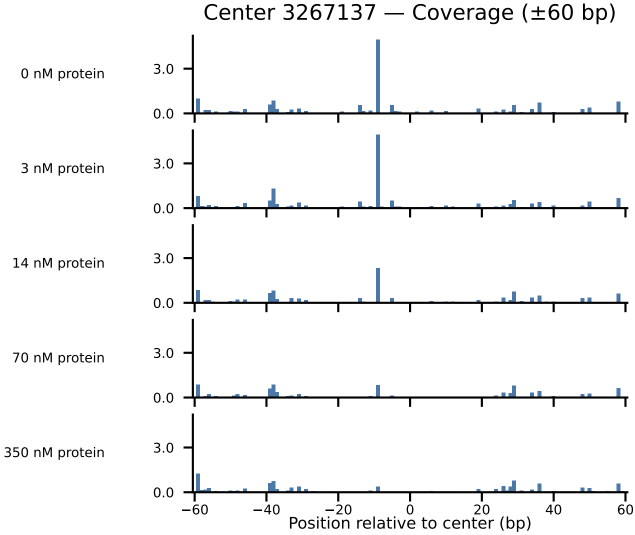

Center 3267137 — Coverage (±60 bp)
0 nM protein
3 nM protein
14 nM protein
70 nM protein
350 nM protein
3.0
0.0
−60
−40
−20
0
20
40
60
Position relative to center (bp)

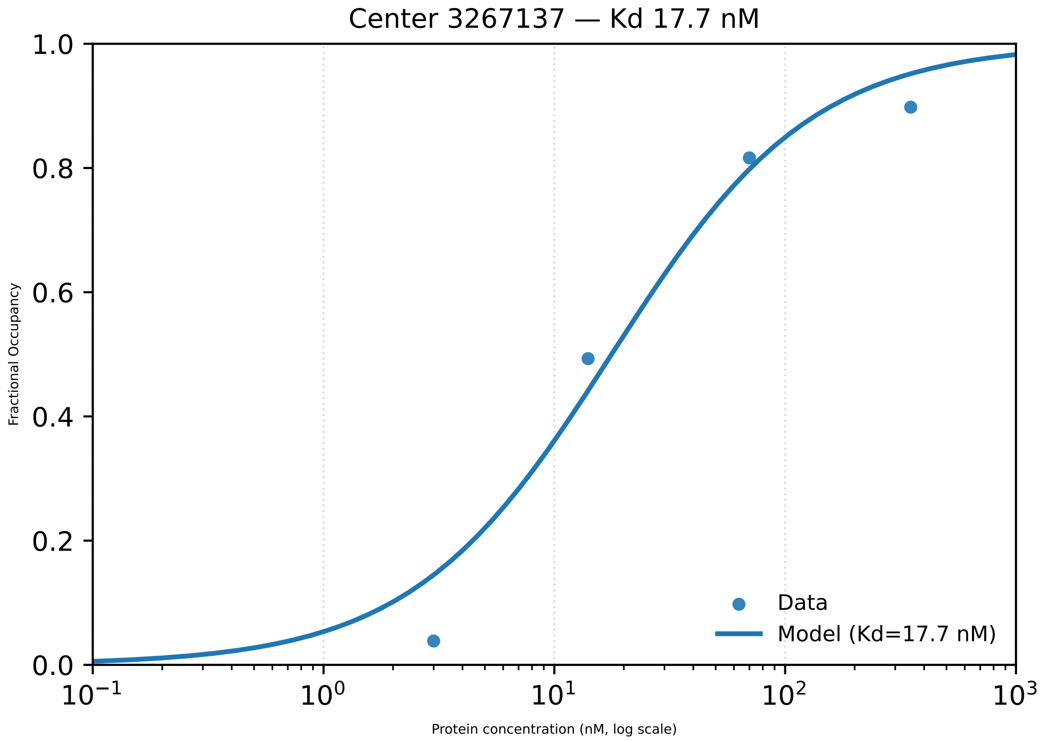

Center 3267137 — Kd 17.7 nM
1.0
0.8
0.6
0.4
0.2
0.0
Fractional Occupancy
$10^{-1}$
$10^{0}$
$10^{1}$
$10^{2}$
$10^{3}$
Protein concentration (nM, log scale)
Data
Model (Kd=17.7 nM)

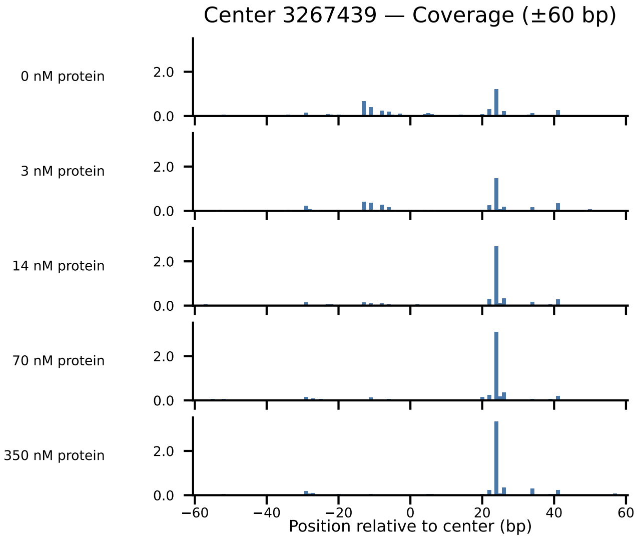

Center 3267439 — Coverage (±60 bp)
0 nM protein
3 nM protein
14 nM protein
70 nM protein
350 nM protein
2.0
0.0
−60
−40
−20
0
20
40
60
Position relative to center (bp)

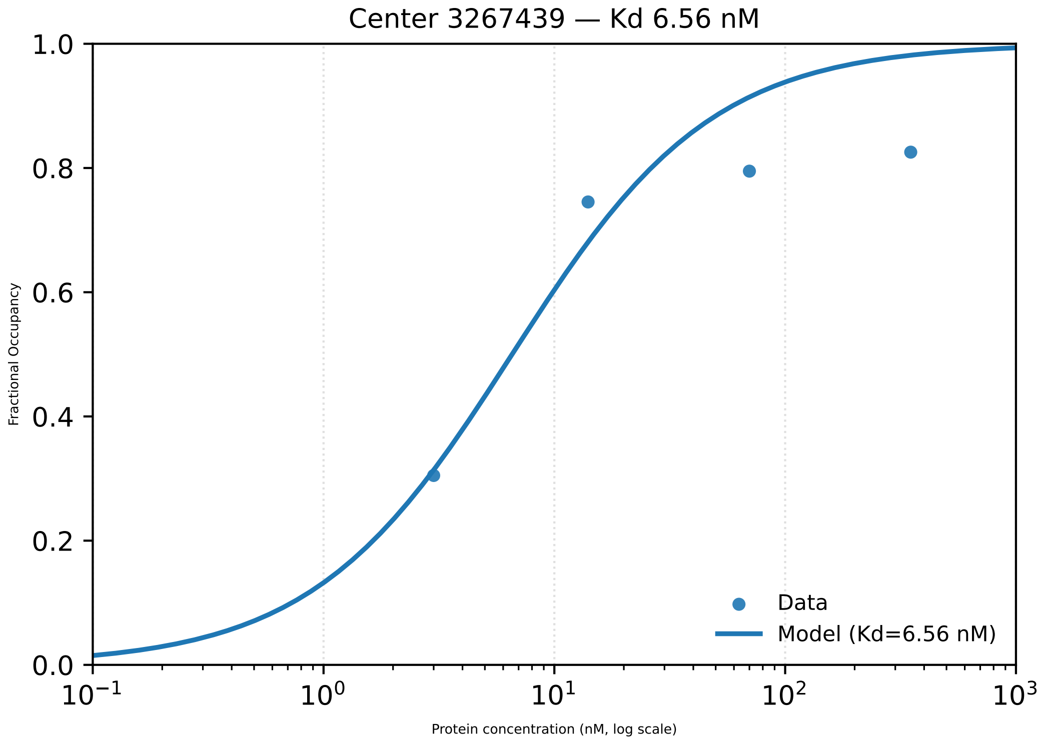

Center 3267439 — Kd 6.56 nM
1.0
0.8
0.6
0.4
0.2
0.0
Fractional Occupancy
$10^{-1}$
$10^{0}$
$10^{1}$
$10^{2}$
$10^{3}$
Protein concentration (nM, log scale)
Data
Model (Kd=6.56 nM)

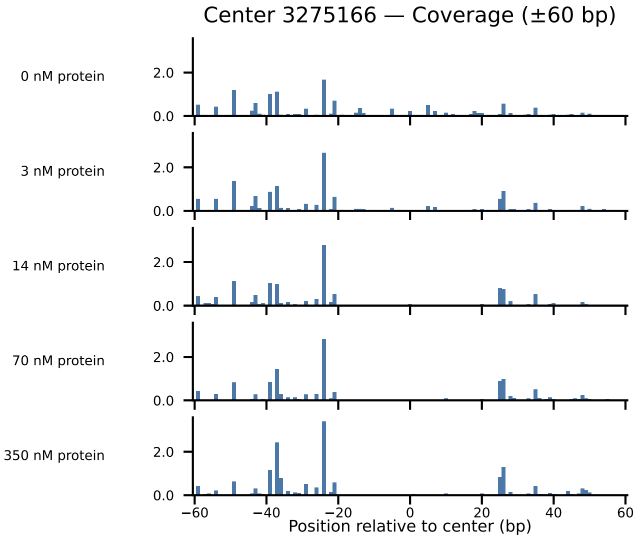

Center 3275166 — Coverage (±60 bp)
0 nM protein
3 nM protein
14 nM protein
70 nM protein
350 nM protein
2.0
0.0
−60
−40
−20
0
20
40
60
Position relative to center (bp)

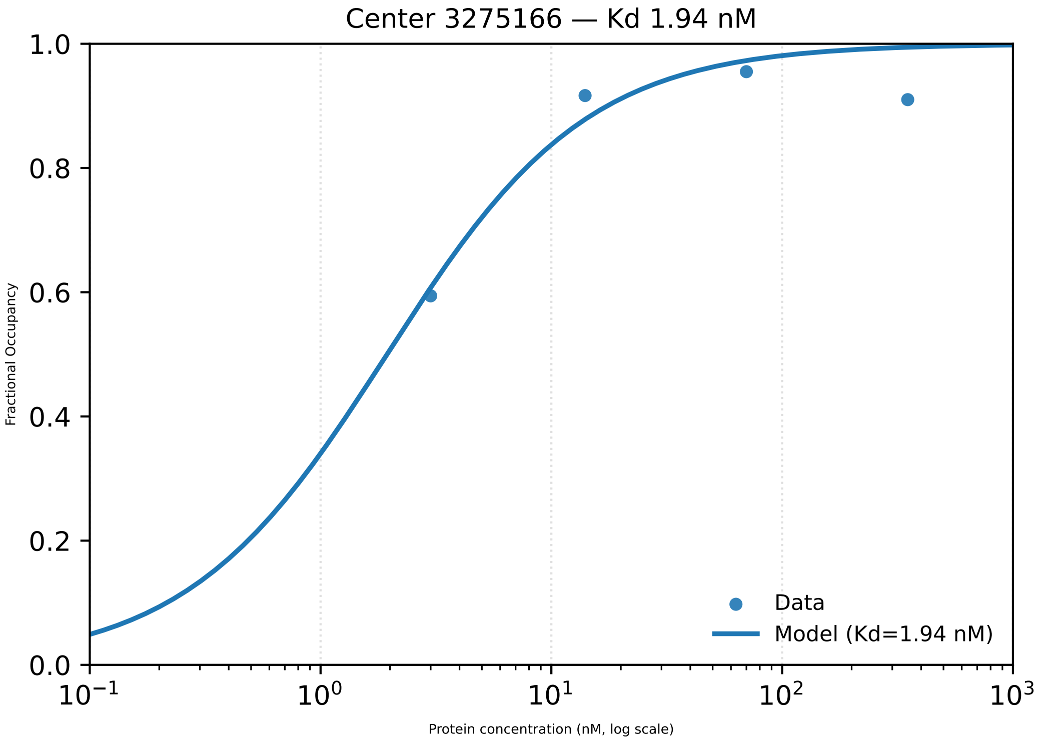

Center 3275166 — Kd 1.94 nM
Fractional Occupancy
1.0
0.8
0.6
0.4
0.2
0.0
$10^{-1}$
$10^{0}$
$10^{1}$
$10^{2}$
$10^{3}$
Protein concentration (nM, log scale)
Data
Model (Kd=1.94 nM)

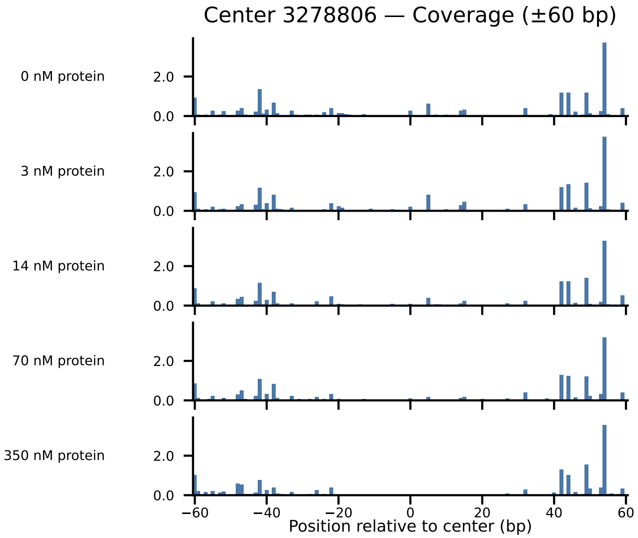

Center 3278806 — Coverage (±60 bp)
0 nM protein
3 nM protein
14 nM protein
70 nM protein
350 nM protein
2.0
0.0
−60
−40
−20
0
20
40
60
Position relative to center (bp)

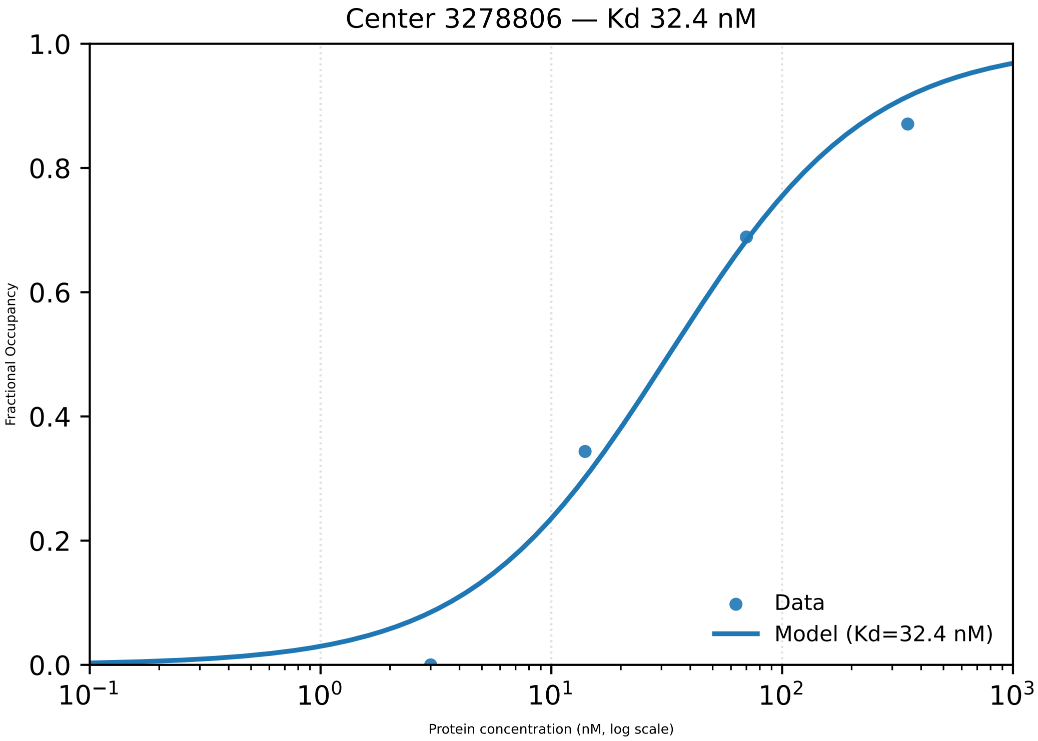

Center 3278806 — Kd 32.4 nM
Fractional Occupancy
1.0
0.8
0.6
0.4
0.2
0.0
$10^{-1}$
$10^{0}$
$10^{1}$
$10^{2}$
$10^{3}$
Protein concentration (nM, log scale)
Data
Model (Kd=32.4 nM)

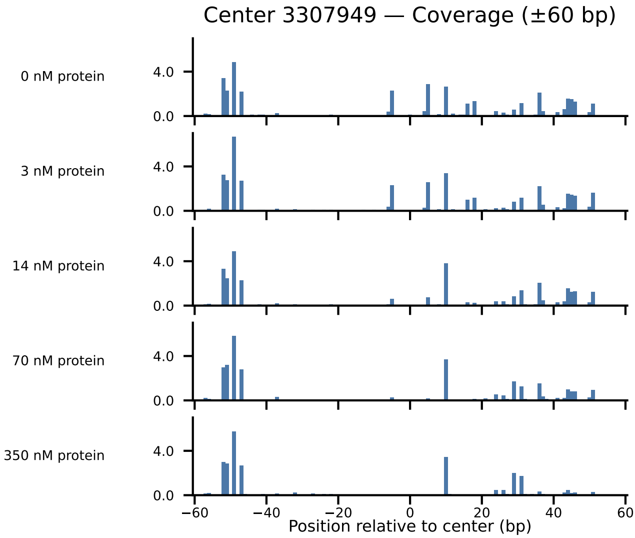
Center 3307949 — Coverage (±60 bp)
0 nM protein
3 nM protein
14 nM protein
70 nM protein
350 nM protein
4.0
0.0
−60
−40
−20
0
20
40
60
Position relative to center (bp)

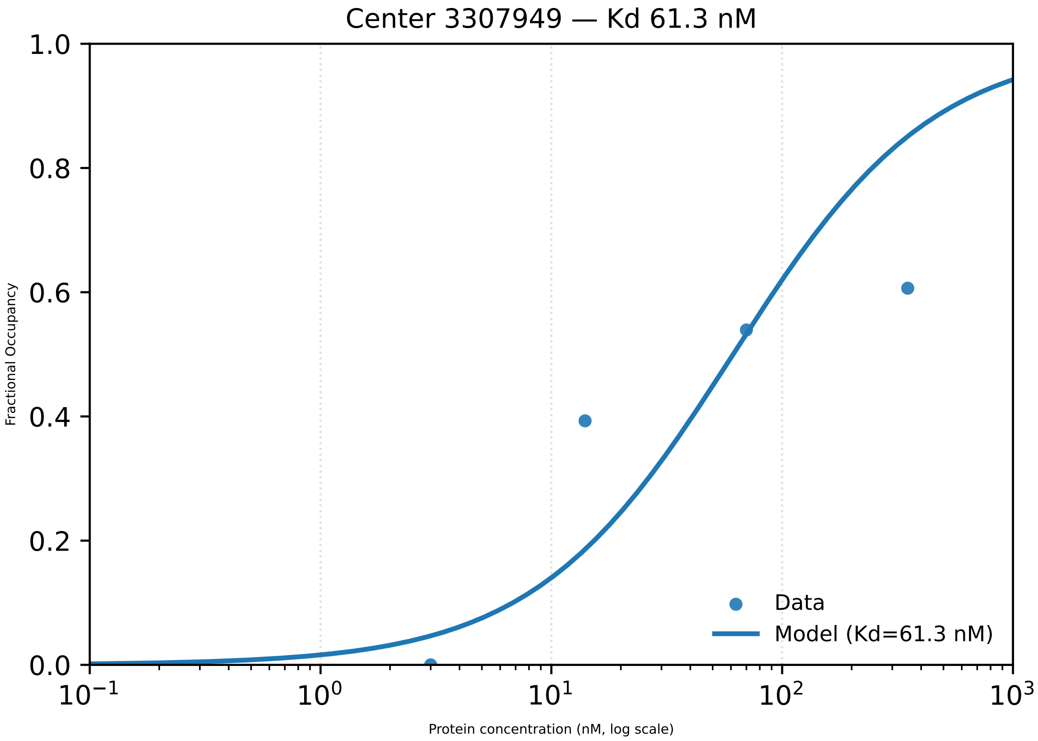

Center 3307949 — Kd 61.3 nM
Fractional Occupancy
1.0
0.8
0.6
0.4
0.2
0.0
$10^{-1}$
$10^{0}$
$10^{1}$
$10^{2}$
$10^{3}$
Protein concentration (nM, log scale)
Data
Model (Kd=61.3 nM)

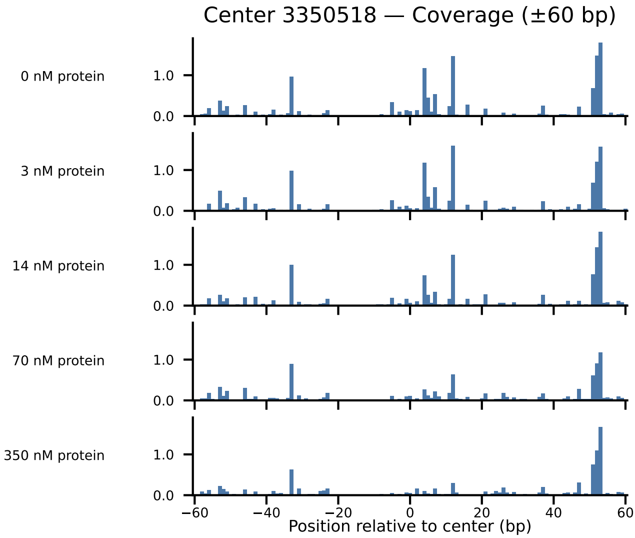
Center 3350518 — Coverage (±60 bp)
0 nM protein
3 nM protein
14 nM protein
70 nM protein
350 nM protein
1.0
0.0
−60
−40
−20
0
20
40
60
Position relative to center (bp)

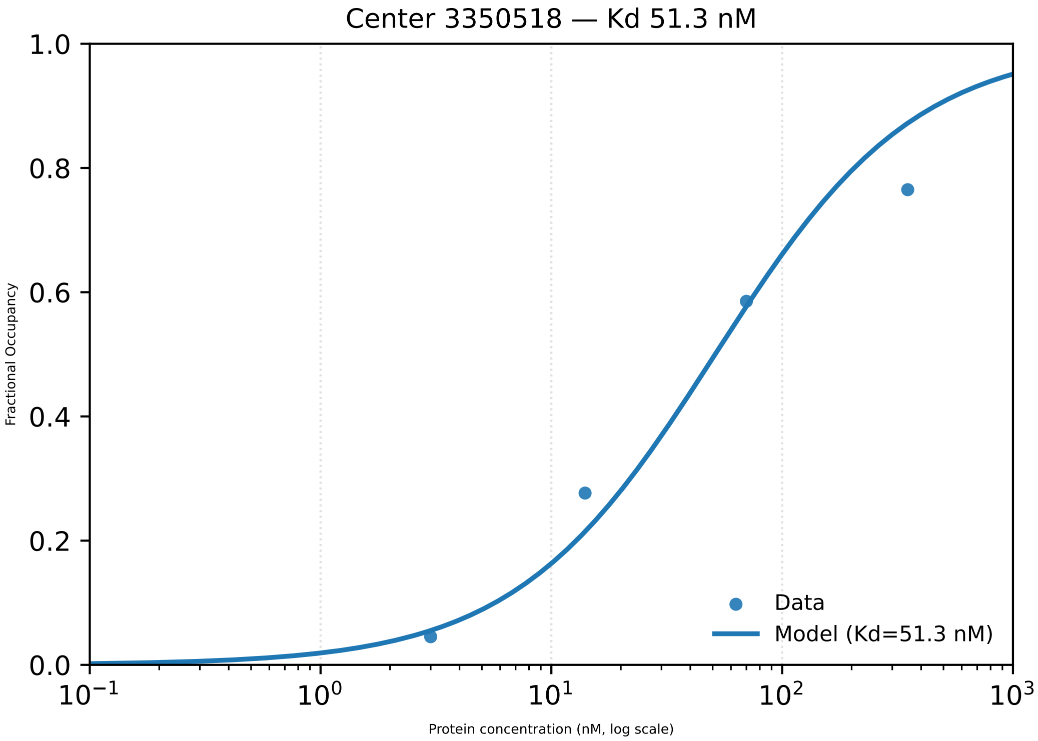

Center 3350518 — Kd 51.3 nM
Fractional Occupancy
1.0
0.8
0.6
0.4
0.2
0.0
$10^{-1}$
$10^{0}$
$10^{1}$
$10^{2}$
$10^{3}$
Protein concentration (nM, log scale)
Data
Model (Kd=51.3 nM)

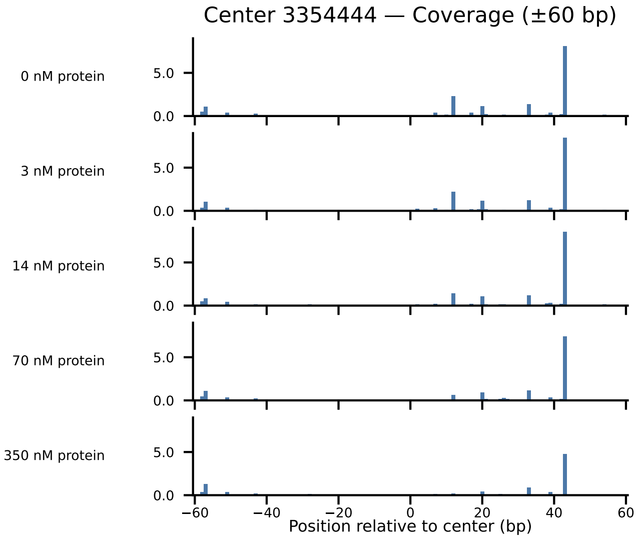

Center 3354444 — Coverage (±60 bp)
0 nM protein
3 nM protein
14 nM protein
70 nM protein
350 nM protein
5.0
0.0
−60
−40
−20
0
20
40
60
Position relative to center (bp)

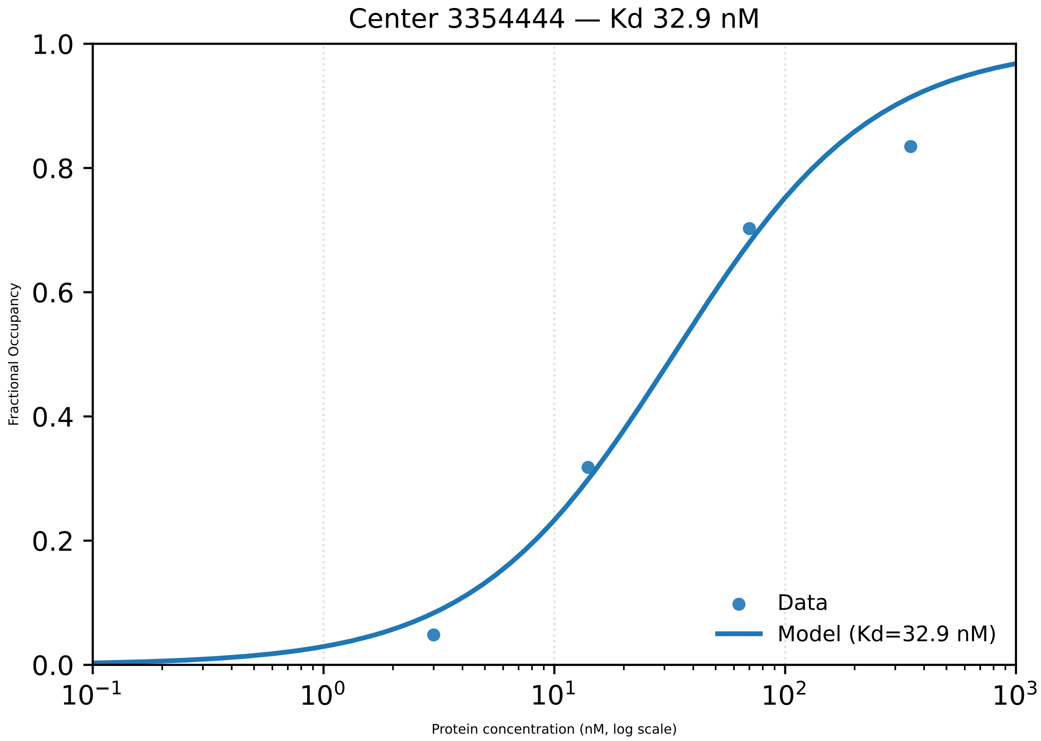

Center 3354444 — Kd 32.9 nM
1.0
0.8
0.6
0.4
0.2
0.0
Fractional Occupancy
$10^{-1}$
$10^{0}$
$10^{1}$
$10^{2}$
$10^{3}$
Protein concentration (nM, log scale)
Data
Model (Kd=32.9 nM)

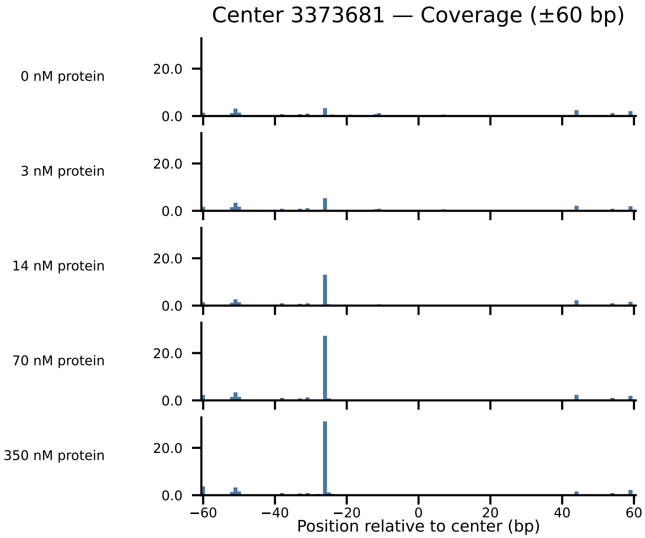

Center 3373681 — Coverage (±60 bp)
0 nM protein
3 nM protein
14 nM protein
70 nM protein
350 nM protein
20.0
0.0
−60
−40
−20
0
20
40
60
Position relative to center (bp)

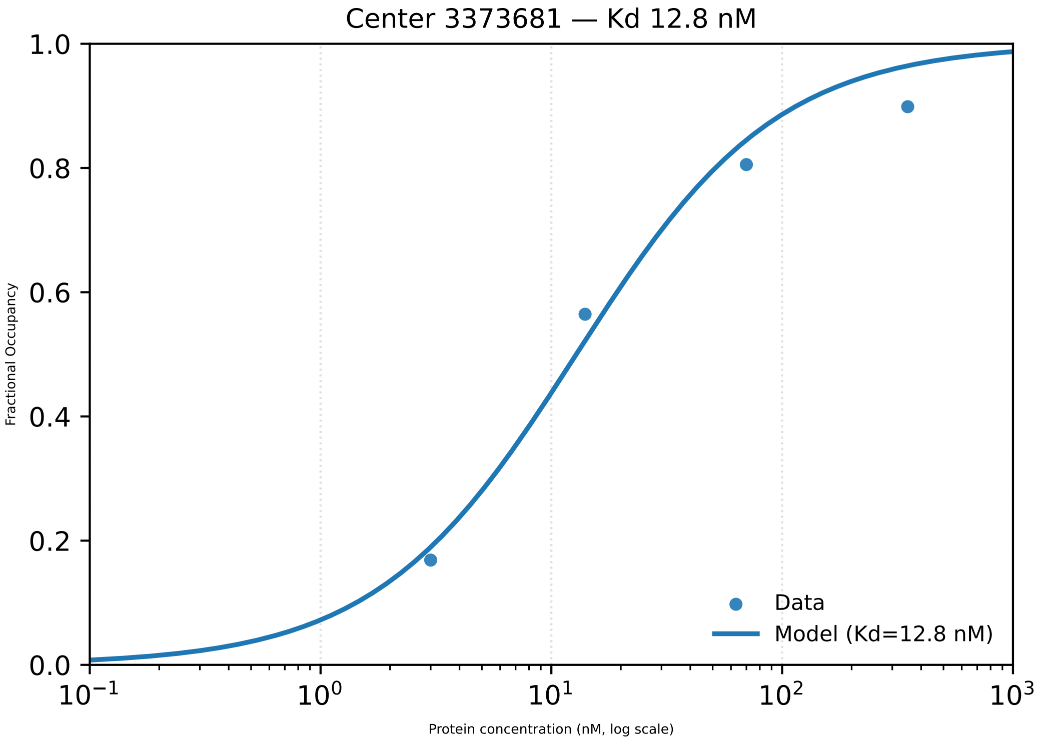

Center 3373681 — Kd 12.8 nM
1.0
0.8
0.6
0.4
0.2
0.0
Fractional Occupancy
Data
Model (Kd=12.8 nM)
$10^{-1}$
$10^{0}$
$10^{1}$
$10^{2}$
$10^{3}$
Protein concentration (nM, log scale)

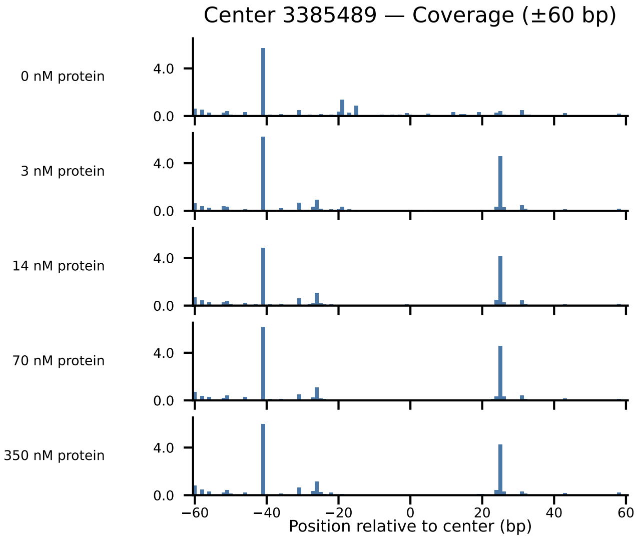

Center 3385489 — Coverage (±60 bp)
0 nM protein
3 nM protein
14 nM protein
70 nM protein
350 nM protein
4.0
0.0
−60
−40
−20
0
20
40
60
Position relative to center (bp)

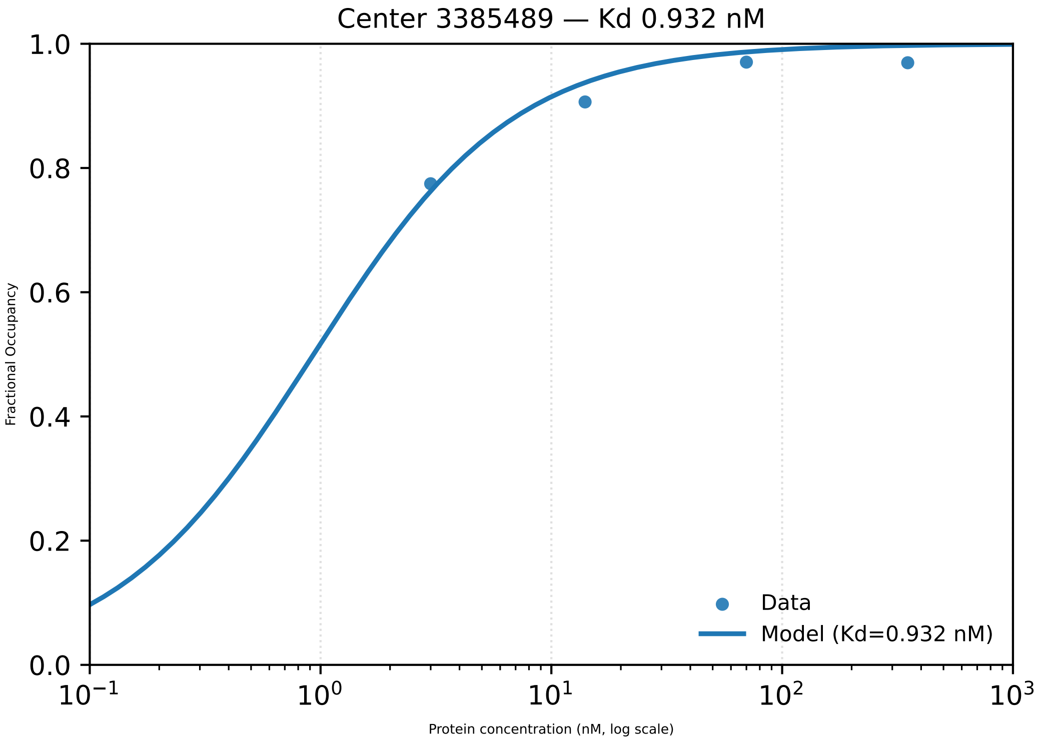

Center 3385489 — Kd 0.932 nM
1.0
0.8
0.6
0.4
0.2
0.0
Fractional Occupancy
$10^{-1}$
$10^{0}$
$10^{1}$
$10^{2}$
$10^{3}$
Protein concentration (nM, log scale)
Data
Model (Kd=0.932 nM)

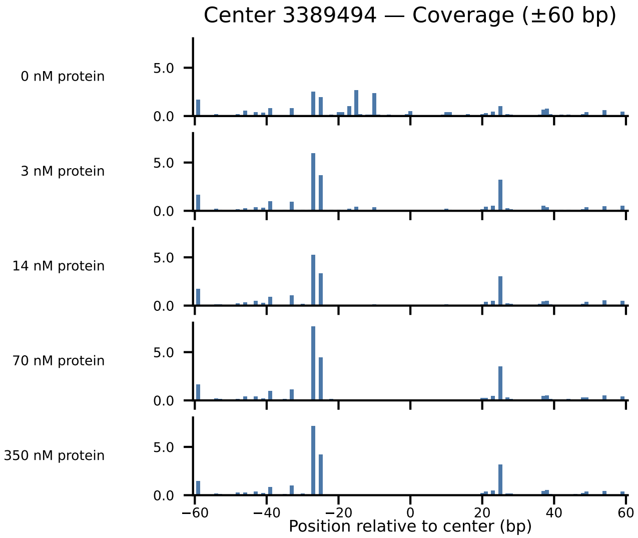

Center 3389494 — Coverage (±60 bp)
0 nM protein
3 nM protein
14 nM protein
70 nM protein
350 nM protein
5.0
0.0
−60
−40
−20
0
20
40
60
Position relative to center (bp)

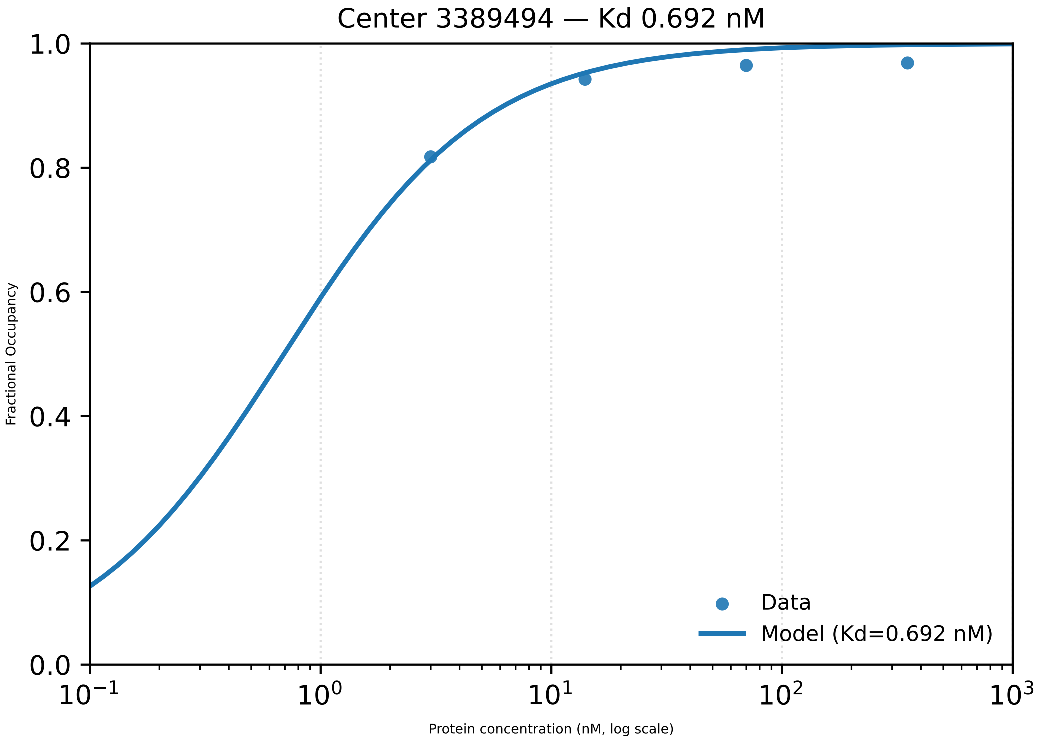

Center 3389494 — Kd 0.692 nM
1.0
0.8
0.6
0.4
0.2
0.0
Fractional Occupancy
$10^{-1}$
$10^{0}$
$10^{1}$
$10^{2}$
$10^{3}$
Protein concentration (nM, log scale)
Data
Model (Kd=0.692 nM)

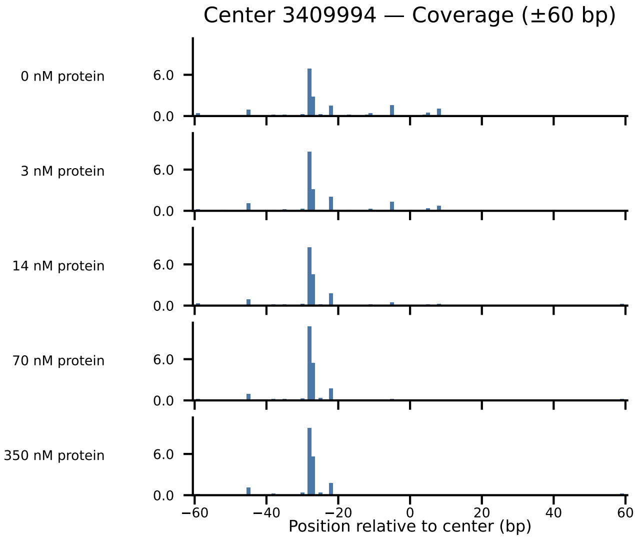
Center 3409994 — Coverage (±60 bp)
0 nM protein
3 nM protein
14 nM protein
70 nM protein
350 nM protein
6.0
0.0
−60
−40
−20
0
20
40
60
Position relative to center (bp)

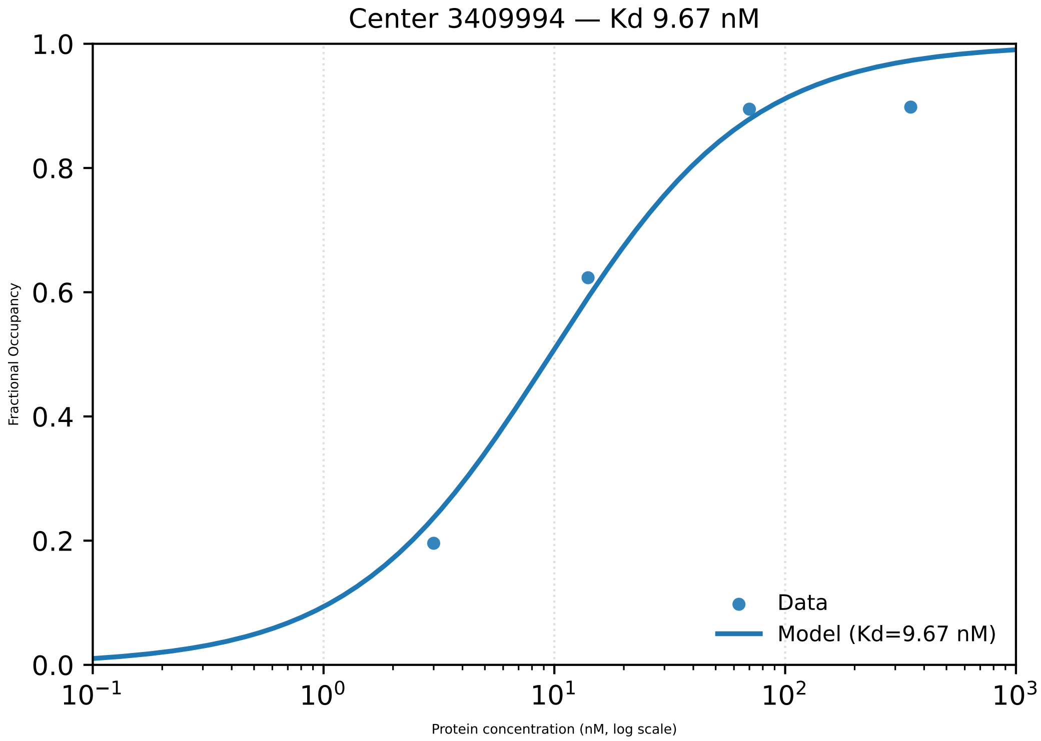

Center 3409994 — Kd 9.67 nM
1.0
0.8
0.6
0.4
0.2
0.0
Fractional Occupancy
$10^{-1}$
$10^{0}$
$10^{1}$
$10^{2}$
$10^{3}$
Protein concentration (nM, log scale)
Data
Model (Kd=9.67 nM)

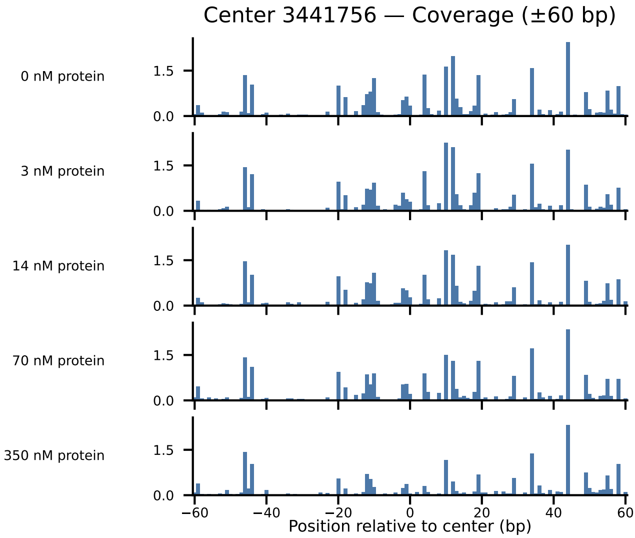

Center 3441756 — Coverage (±60 bp)
0 nM protein
3 nM protein
14 nM protein
70 nM protein
350 nM protein
1.5
0.0
−60
−40
−20
0
20
40
60
Position relative to center (bp)

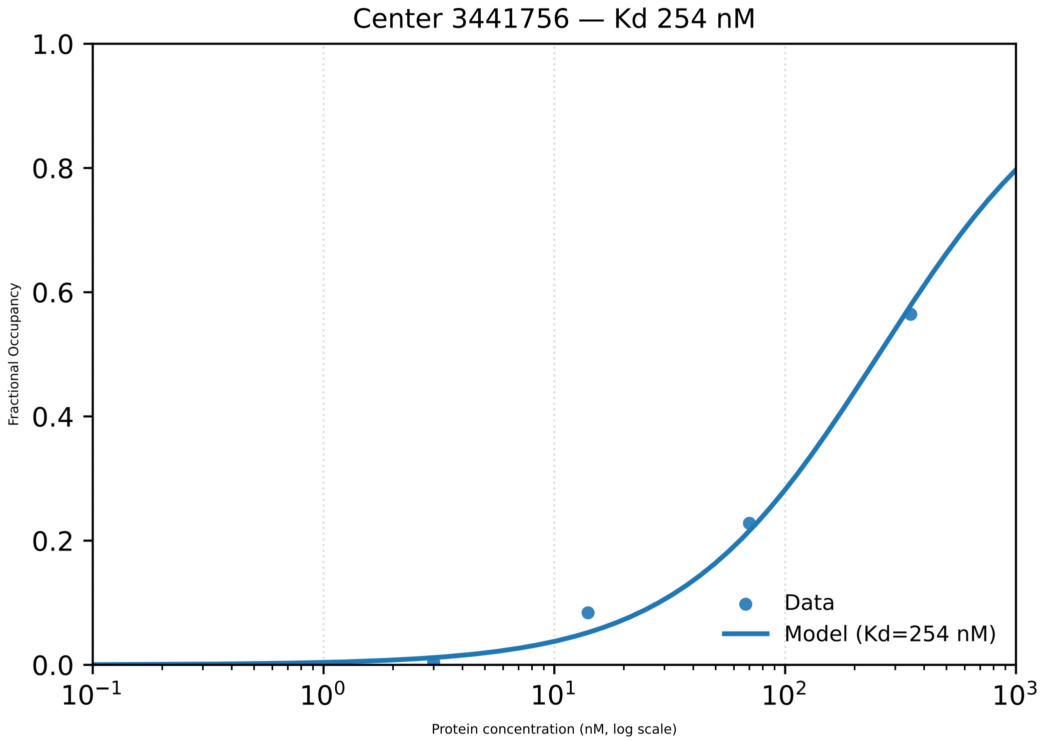

Center 3441756 — Kd 254 nM
1.0
0.8
0.6
0.4
0.2
0.0
Fractional Occupancy
Data
Model (Kd=254 nM)
$10^{-1}$
$10^{0}$
$10^{1}$
$10^{2}$
$10^{3}$
Protein concentration (nM, log scale)

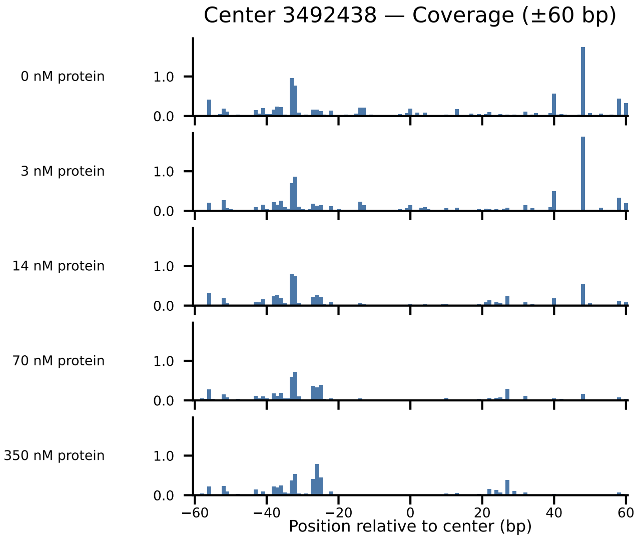

Center 3492438 — Coverage (±60 bp)
0 nM protein
3 nM protein
14 nM protein
70 nM protein
350 nM protein
1.0
0.0
−60
−40
−20
0
20
40
60
Position relative to center (bp)

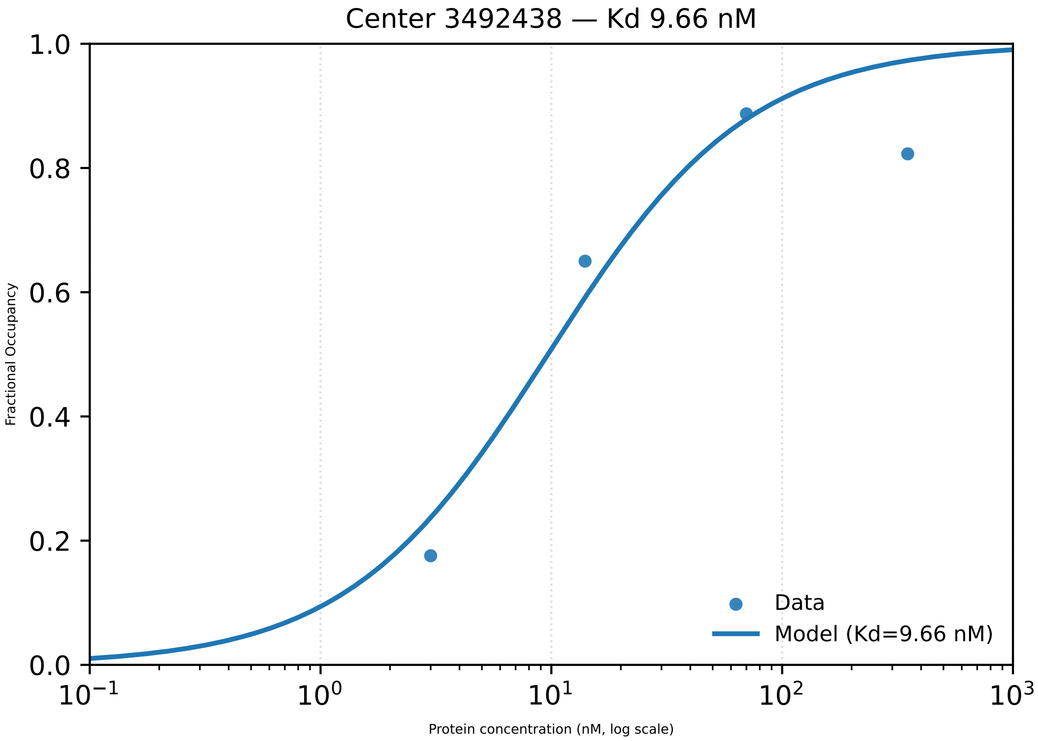

Center 3492438 — Kd 9.66 nM
1.0
0.8
0.6
0.4
0.2
0.0
Fractional Occupancy
$10^{-1}$
$10^{0}$
$10^{1}$
$10^{2}$
$10^{3}$
Protein concentration (nM, log scale)
Data
Model (Kd=9.66 nM)

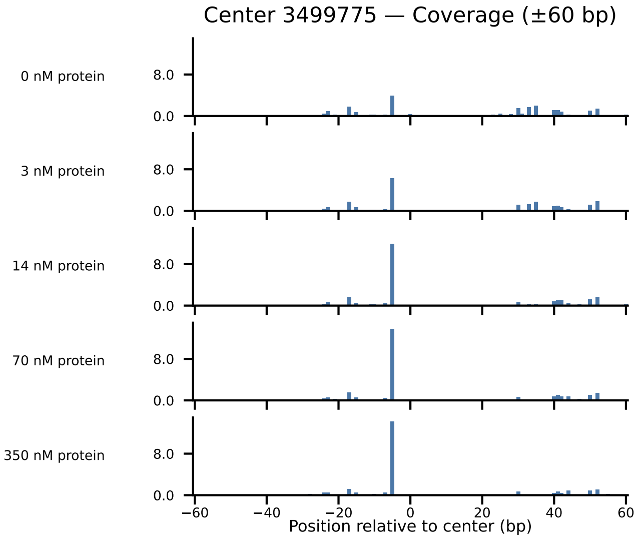

Center 3499775 — Coverage (±60 bp)
0 nM protein
3 nM protein
14 nM protein
70 nM protein
350 nM protein
8.0
0.0
−60
−40
−20
0
20
40
60
Position relative to center (bp)

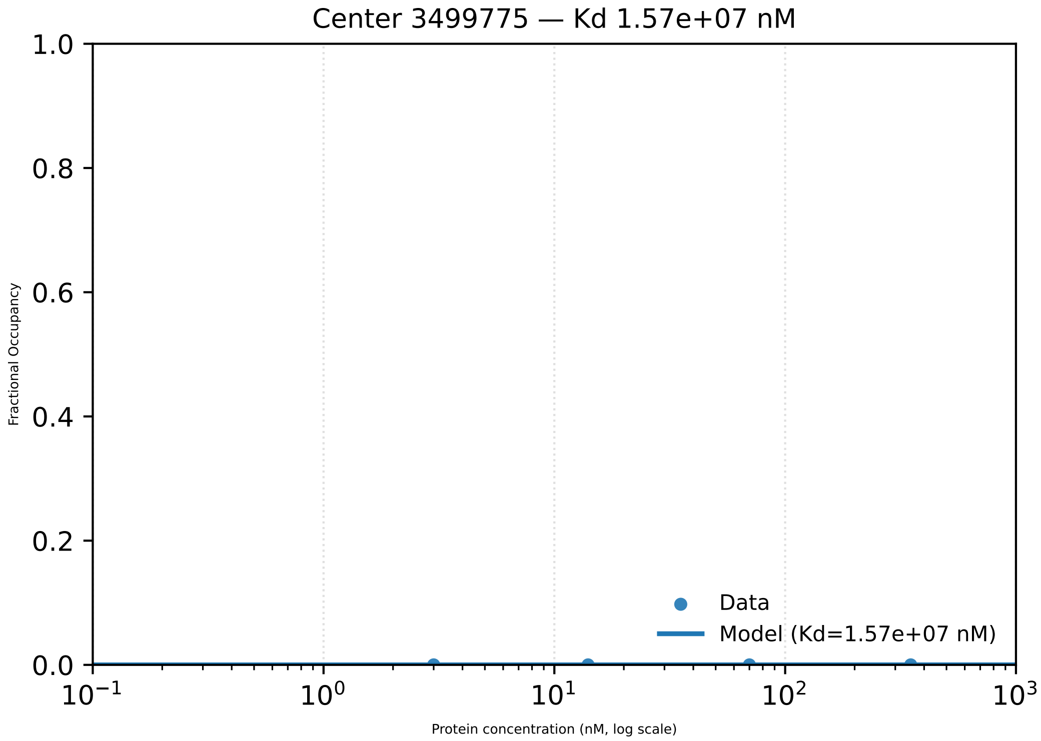

Center 3499775 — Kd 1.57e+07 nM
1.0
0.8
0.6
0.4
0.2
0.0
Fractional Occupancy
$10^{-1}$
$10^{0}$
$10^{1}$
$10^{2}$
$10^{3}$
Protein concentration (nM, log scale)
Data
Model (Kd=1.57e+07 nM)

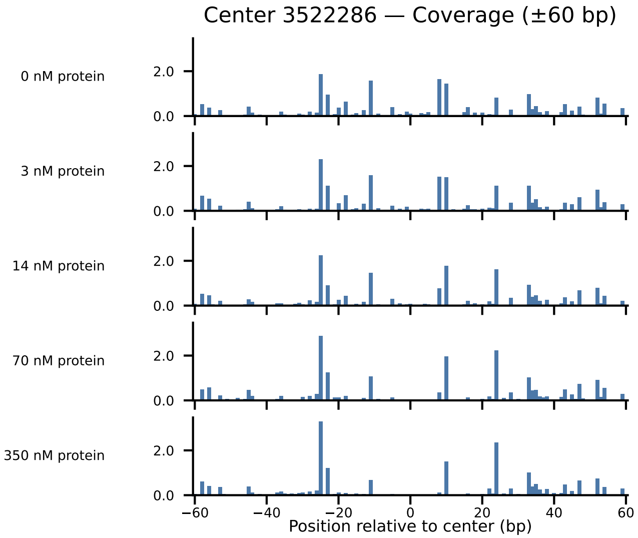

Center 3522286 — Coverage (±60 bp)
0 nM protein
3 nM protein
14 nM protein
70 nM protein
350 nM protein
2.0
0.0
−60
−40
−20
0
20
40
60
Position relative to center (bp)

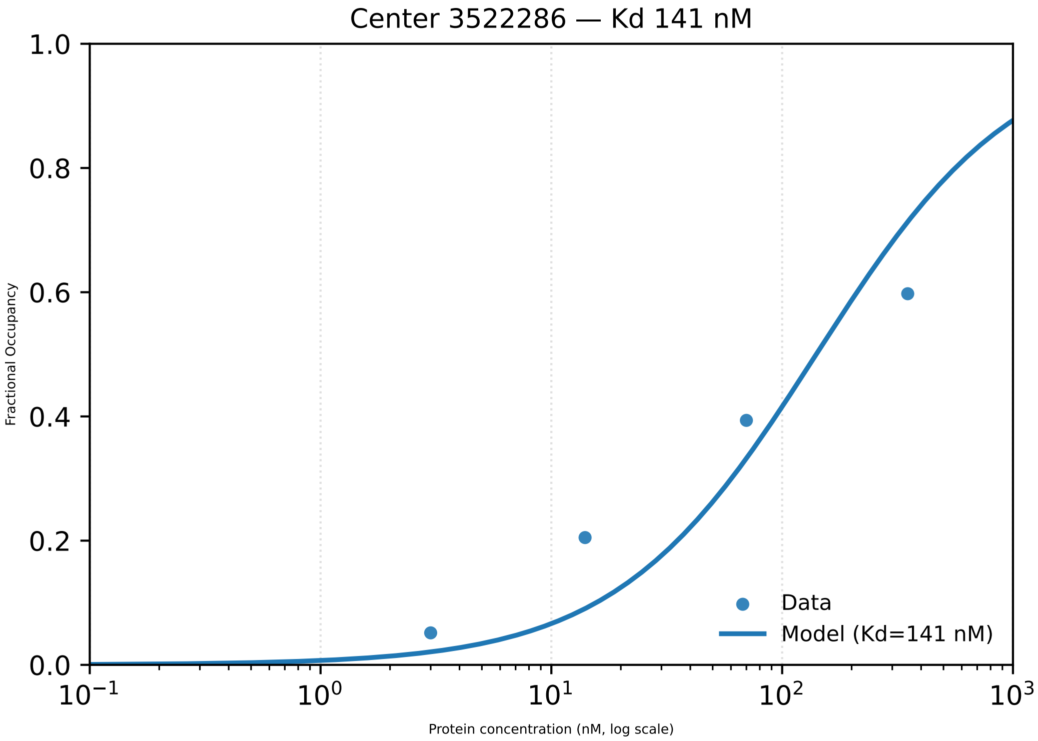

Center 3522286 — Kd 141 nM
1.0
0.8
0.6
0.4
0.2
0.0
Fractional Occupancy
$10^{-1}$
$10^{0}$
$10^{1}$
$10^{2}$
$10^{3}$
Protein concentration (nM, log scale)
Data
Model (Kd=141 nM)

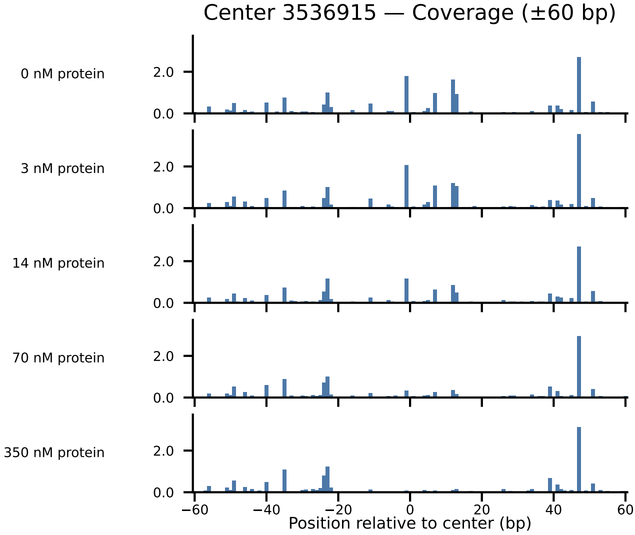

Center 3536915 — Coverage (±60 bp)
0 nM protein
3 nM protein
14 nM protein
70 nM protein
350 nM protein
2.0
0.0
−60
−40
−20
0
20
40
60
Position relative to center (bp)

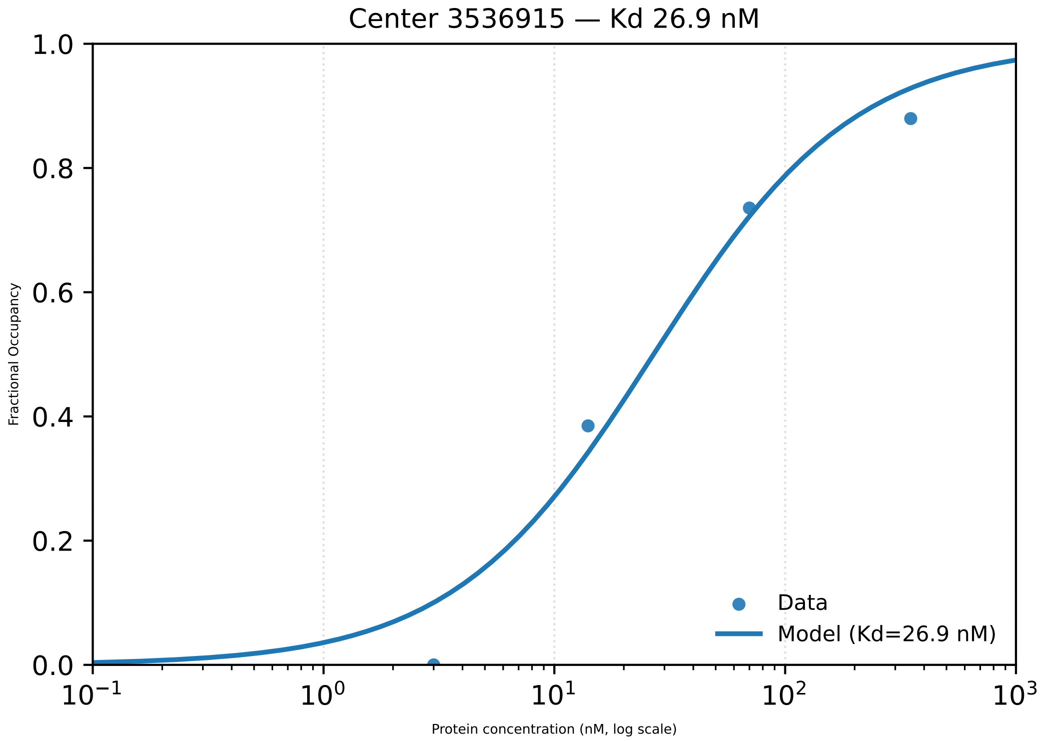
Center 3536915 — Kd 26.9 nM
1.0
0.8
0.6
0.4
0.2
0.0
Fractional Occupancy
$10^{-1}$
$10^{0}$
$10^{1}$
$10^{2}$
$10^{3}$
Protein concentration (nM, log scale)
Data
Model (Kd=26.9 nM)

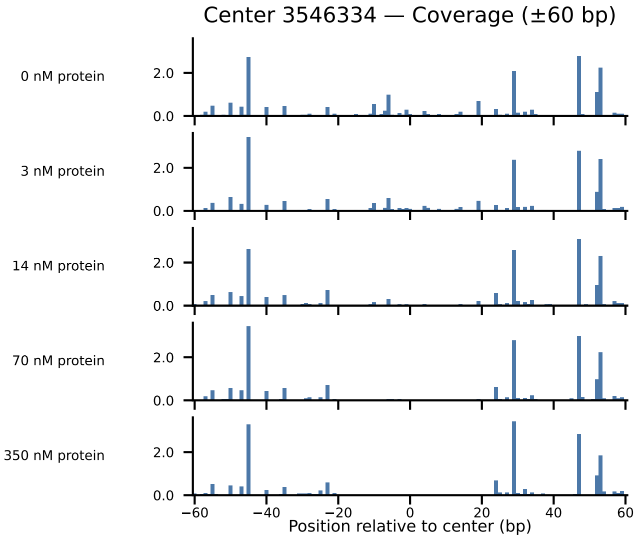

Center 3546334 — Coverage (±60 bp)
0 nM protein
3 nM protein
14 nM protein
70 nM protein
350 nM protein
2.0
0.0
−60
−40
−20
0
20
40
60
Position relative to center (bp)

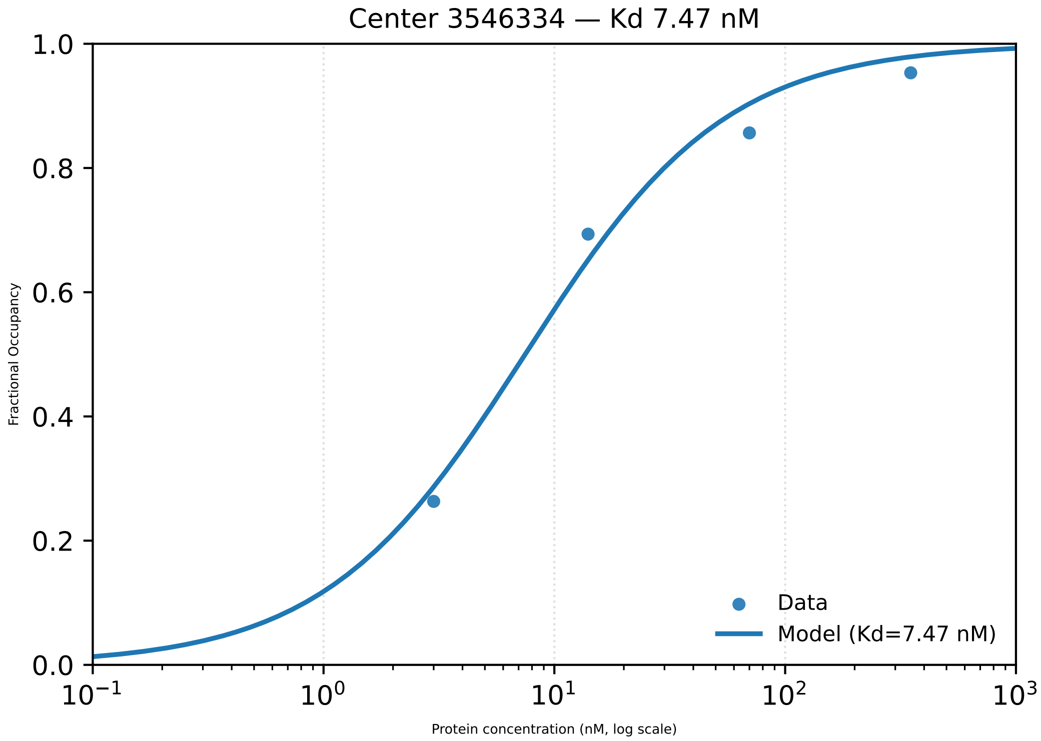

Center 3546334 — Kd 7.47 nM
1.0
0.8
0.6
0.4
0.2
0.0
Fractional Occupancy
$10^{-1}$
$10^{0}$
$10^{1}$
$10^{2}$
$10^{3}$
Protein concentration (nM, log scale)
Data
Model (Kd=7.47 nM)

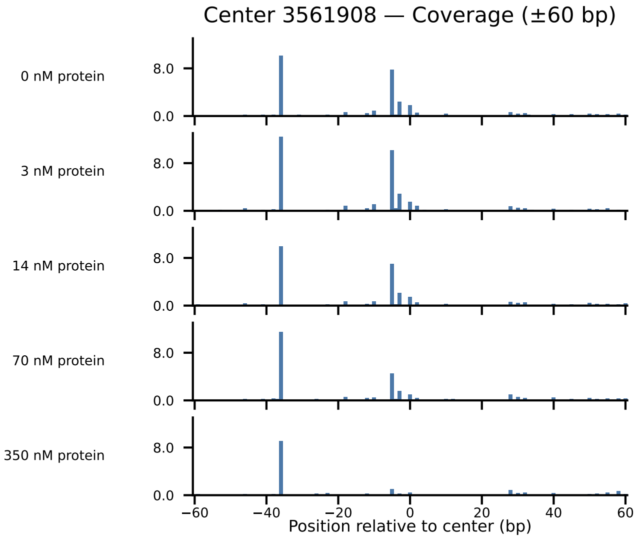

Center 3561908 — Coverage (±60 bp)
0 nM protein
3 nM protein
14 nM protein
70 nM protein
350 nM protein
8.0
0.0
−60
−40
−20
0
20
40
60
Position relative to center (bp)

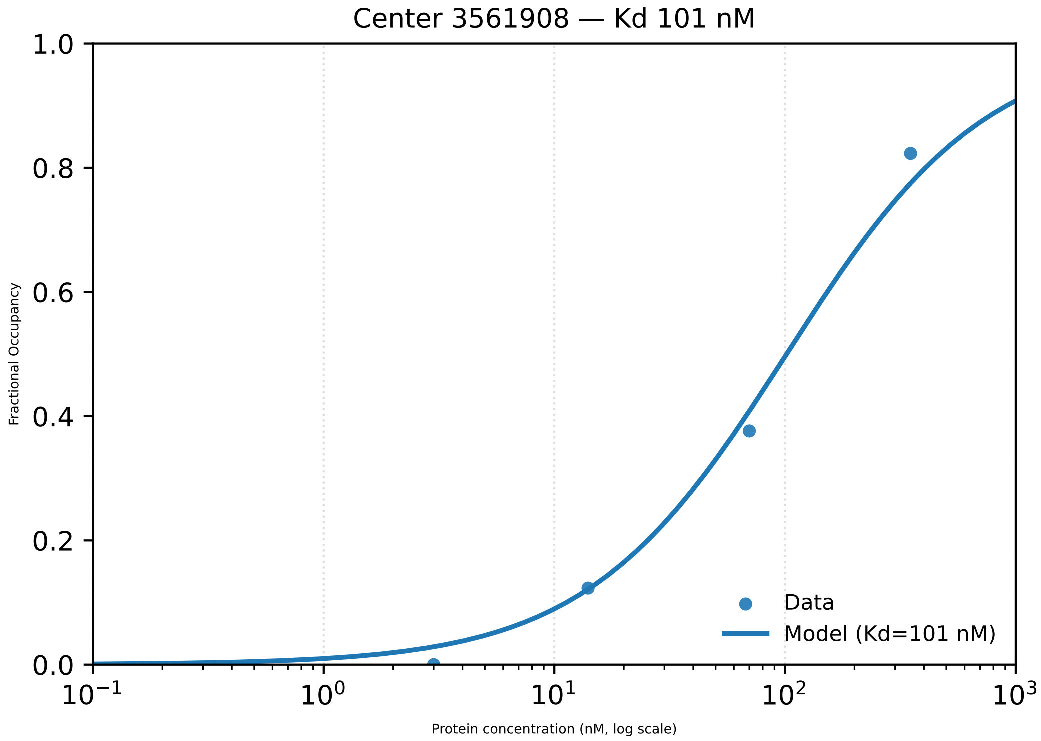
Center 3561908 — Kd 101 nM
1.0
0.8
0.6
0.4
0.2
0.0
Fractional Occupancy
$10^{-1}$
$10^{0}$
$10^{1}$
$10^{2}$
$10^{3}$
Protein concentration (nM, log scale)
Data
Model (Kd=101 nM)

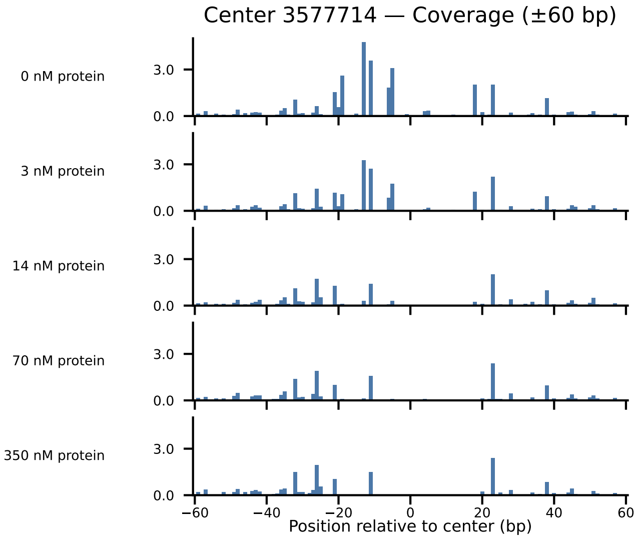

Center 3577714 — Coverage (±60 bp)
0 nM protein
3 nM protein
14 nM protein
70 nM protein
350 nM protein
3.0
0.0
−60
−40
−20
0
20
40
60
Position relative to center (bp)

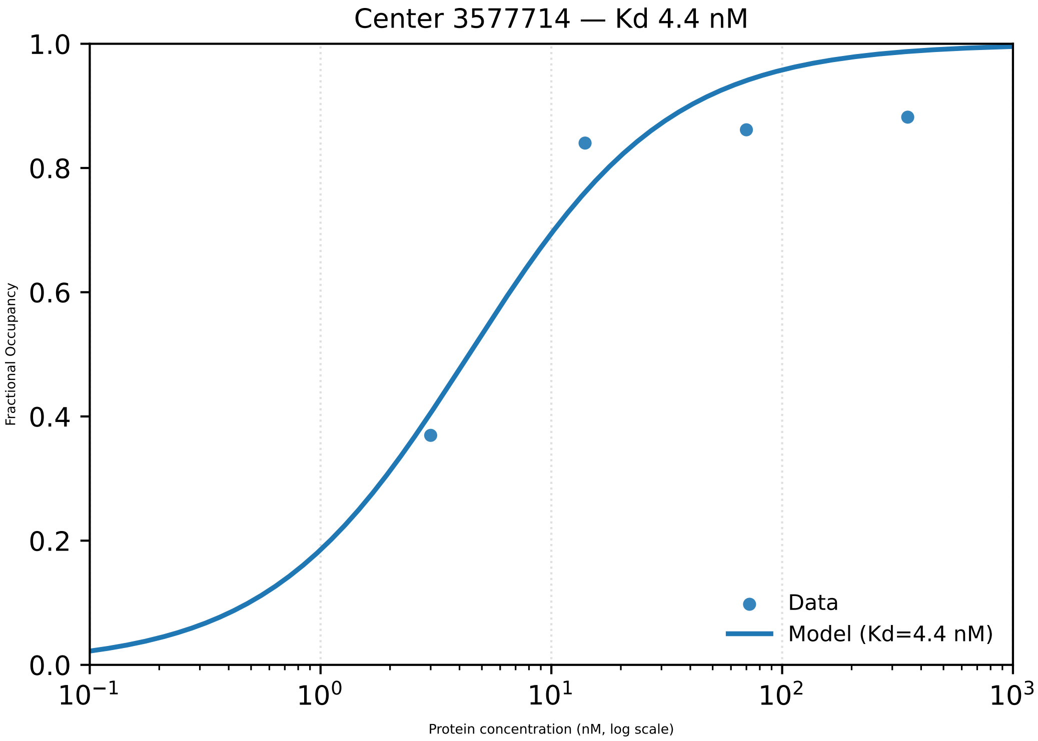

Center 3577714 — Kd 4.4 nM
Fractional Occupancy
1.0
0.8
0.6
0.4
0.2
0.0
$10^{-1}$
$10^{0}$
$10^{1}$
$10^{2}$
$10^{3}$
Protein concentration (nM, log scale)
Data
Model (Kd=4.4 nM)

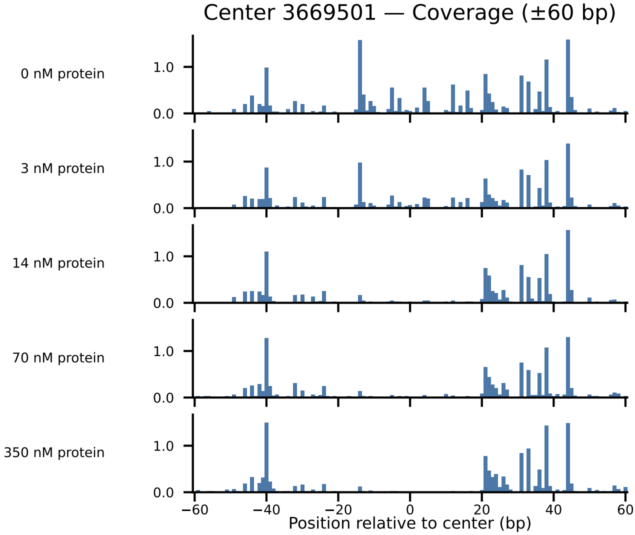

Center 3669501 — Coverage (±60 bp)
0 nM protein
3 nM protein
14 nM protein
70 nM protein
350 nM protein
1.0
0.0
−60
−40
−20
0
20
40
60
Position relative to center (bp)

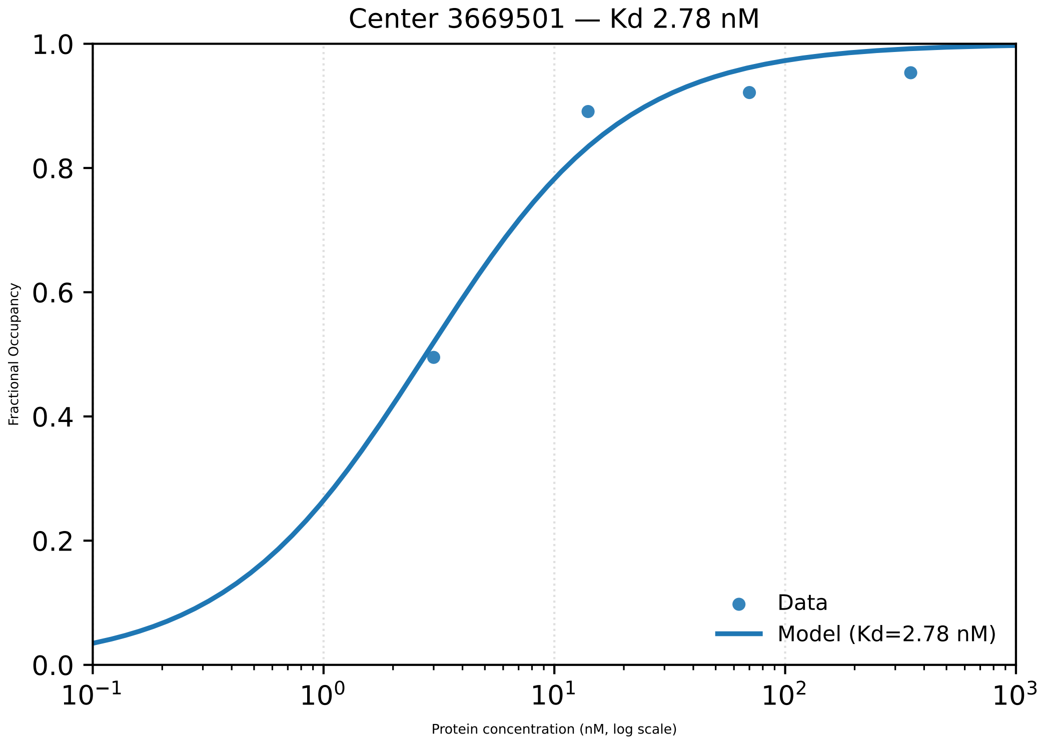

Center 3669501 — Kd 2.78 nM
Fractional Occupancy
1.0
0.8
0.6
0.4
0.2
0.0
$10^{-1}$
$10^{0}$
$10^{1}$
$10^{2}$
$10^{3}$
Protein concentration (nM, log scale)
Data
Model (Kd=2.78 nM)

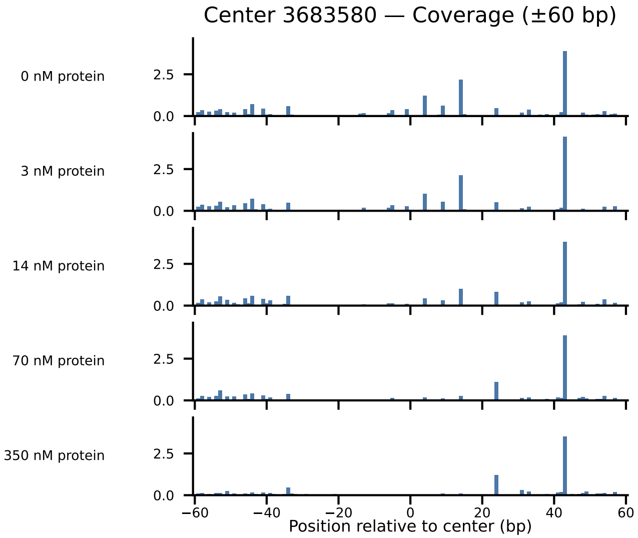

Center 3683580 — Coverage (±60 bp)
0 nM protein
3 nM protein
14 nM protein
70 nM protein
350 nM protein
2.5
0.0
−60
−40
−20
0
20
40
60
Position relative to center (bp)

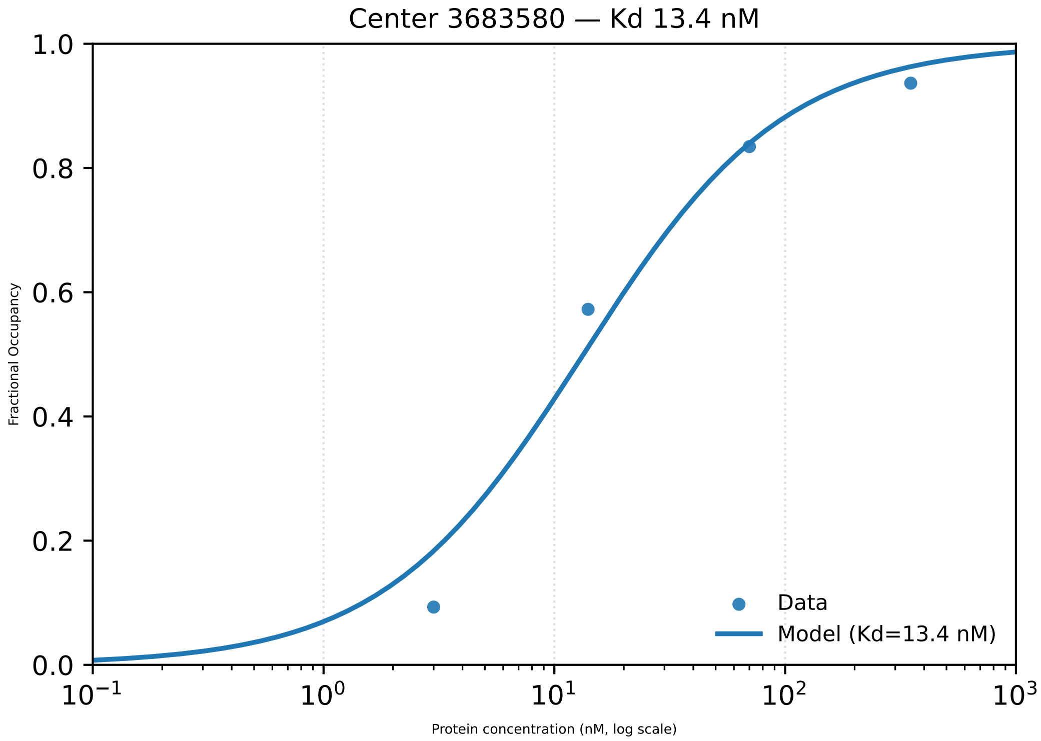

Center 3683580 — Kd 13.4 nM
1.0
0.8
0.6
0.4
0.2
0.0
Fractional Occupancy
$10^{-1}$
$10^{0}$
$10^{1}$
$10^{2}$
$10^{3}$
Protein concentration (nM, log scale)
Data
Model (Kd=13.4 nM)

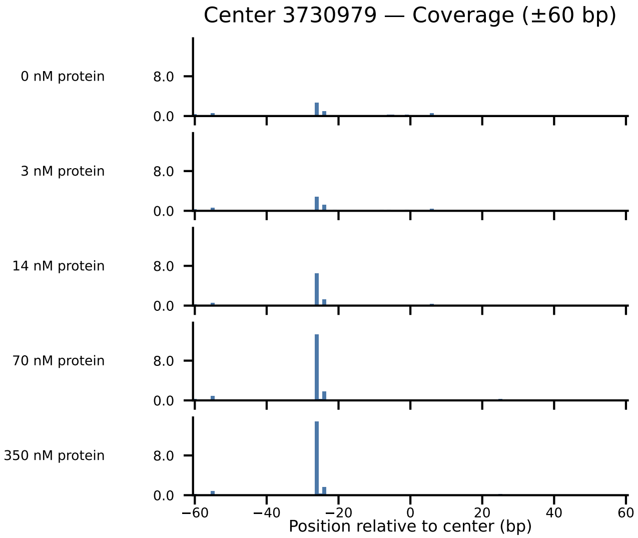

Center 3730979 — Coverage (±60 bp)
0 nM protein
3 nM protein
14 nM protein
70 nM protein
350 nM protein
8.0
0.0
−60
−40
−20
0
20
40
60
Position relative to center (bp)

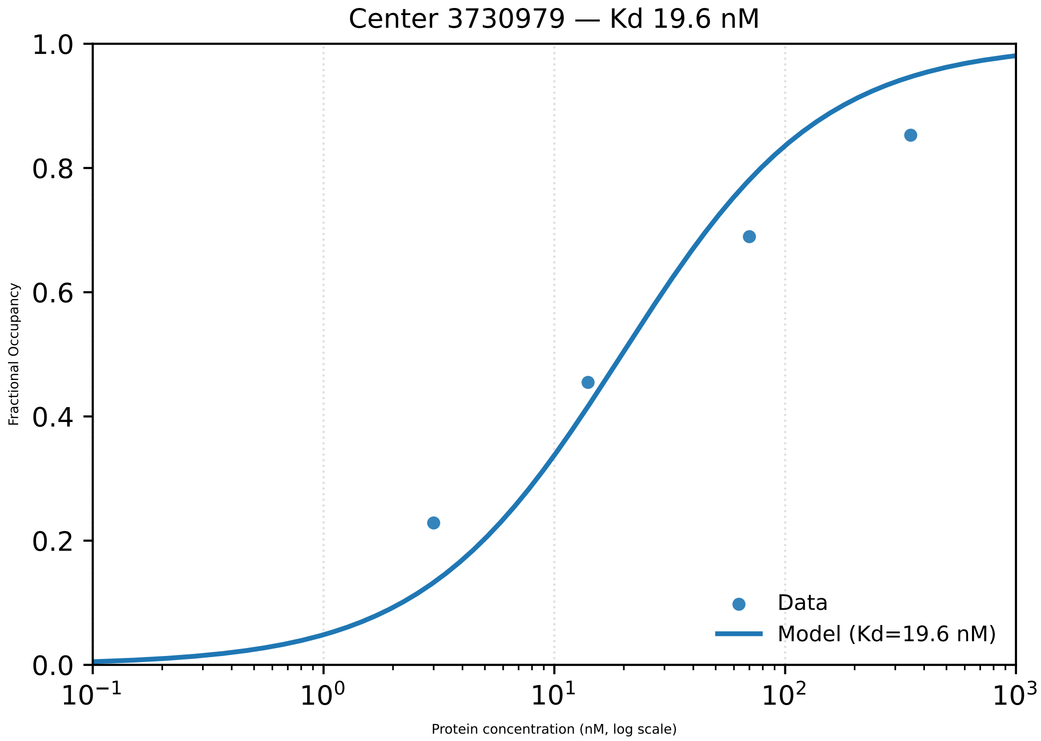
Center 3730979 — Kd 19.6 nM
1.0
0.8
0.6
0.4
0.2
0.0
Fractional Occupancy
$10^{-1}$
$10^{0}$
$10^{1}$
$10^{2}$
$10^{3}$
Protein concentration (nM, log scale)
Data
Model (Kd=19.6 nM)

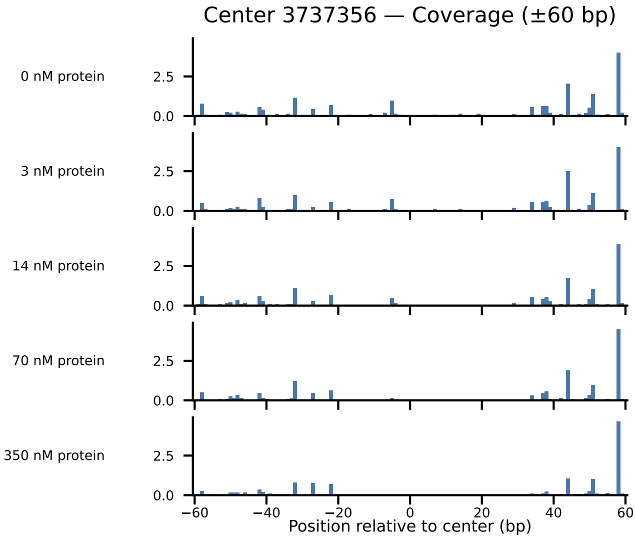

Center 3737356 — Coverage (±60 bp)
0 nM protein
3 nM protein
14 nM protein
70 nM protein
350 nM protein
2.5
0.0
−60
−40
−20
0
20
40
60
Position relative to center (bp)

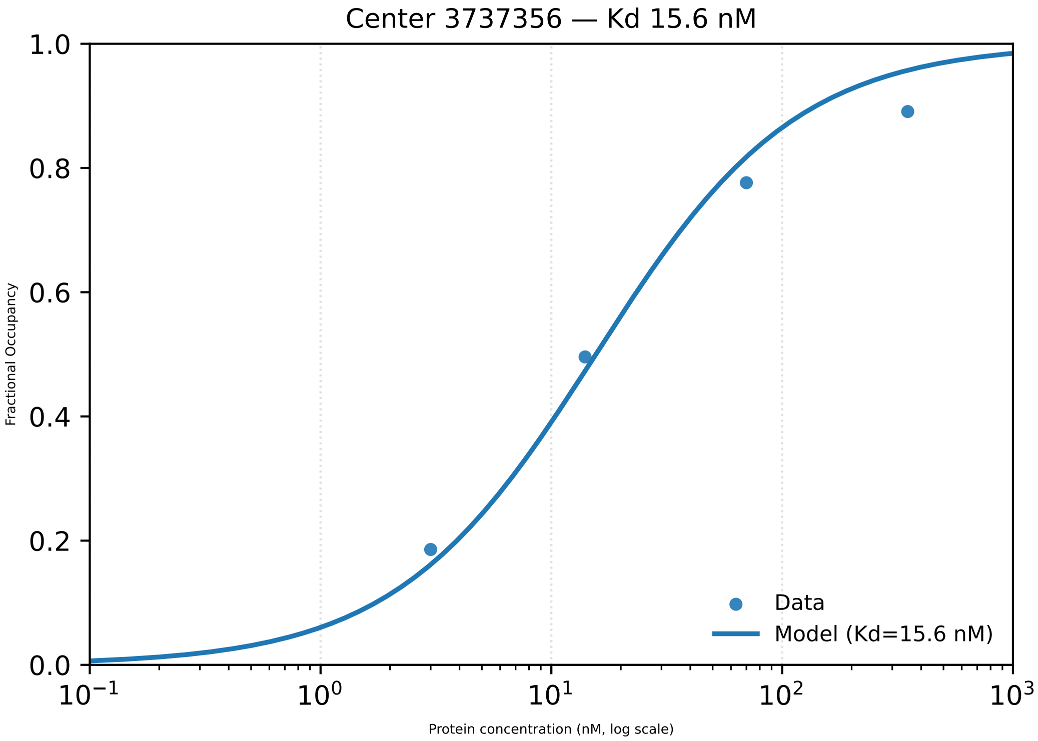

Center 3737356 — Kd 15.6 nM
1.0
0.8
0.6
0.4
0.2
0.0
Fractional Occupancy
$10^{-1}$
$10^{0}$
$10^{1}$
$10^{2}$
$10^{3}$
Protein concentration (nM, log scale)
Data
Model (Kd=15.6 nM)

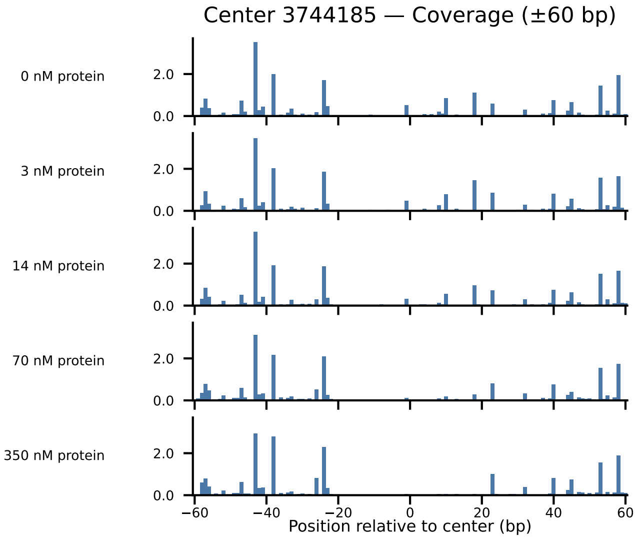

Center 3744185 — Coverage (±60 bp)
0 nM protein
3 nM protein
14 nM protein
70 nM protein
350 nM protein
2.0
0.0
−60
−40
−20
0
20
40
60
Position relative to center (bp)

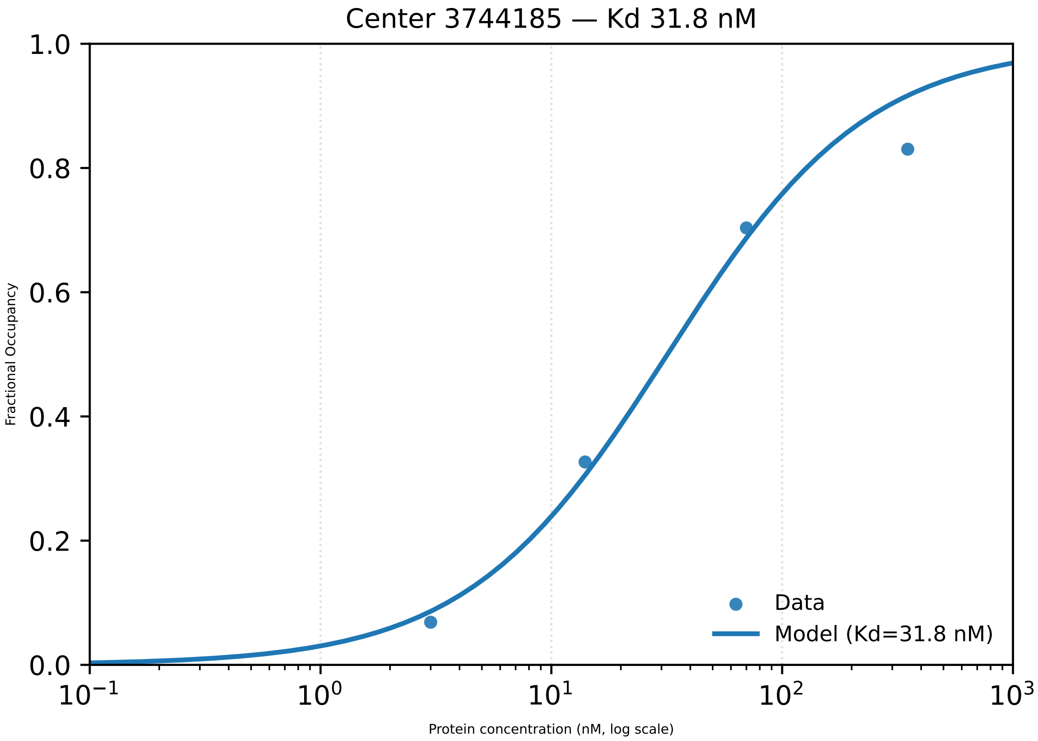
Center 3744185 — Kd 31.8 nM
1.0
0.8
0.6
0.4
0.2
0.0
Fractional Occupancy
$10^{-1}$
$10^{0}$
$10^{1}$
$10^{2}$
$10^{3}$
Protein concentration (nM, log scale)
Data
Model (Kd=31.8 nM)

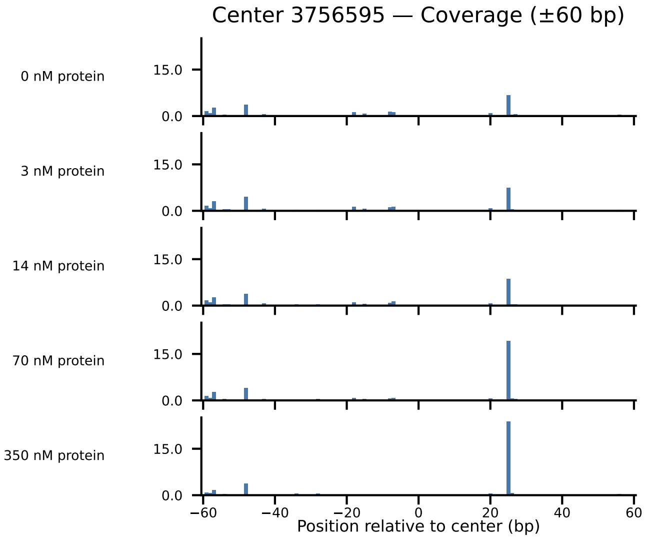

Center 3756595 — Coverage (±60 bp)
0 nM protein
3 nM protein
14 nM protein
70 nM protein
350 nM protein
15.0
0.0
−60
−40
−20
0
20
40
60
Position relative to center (bp)

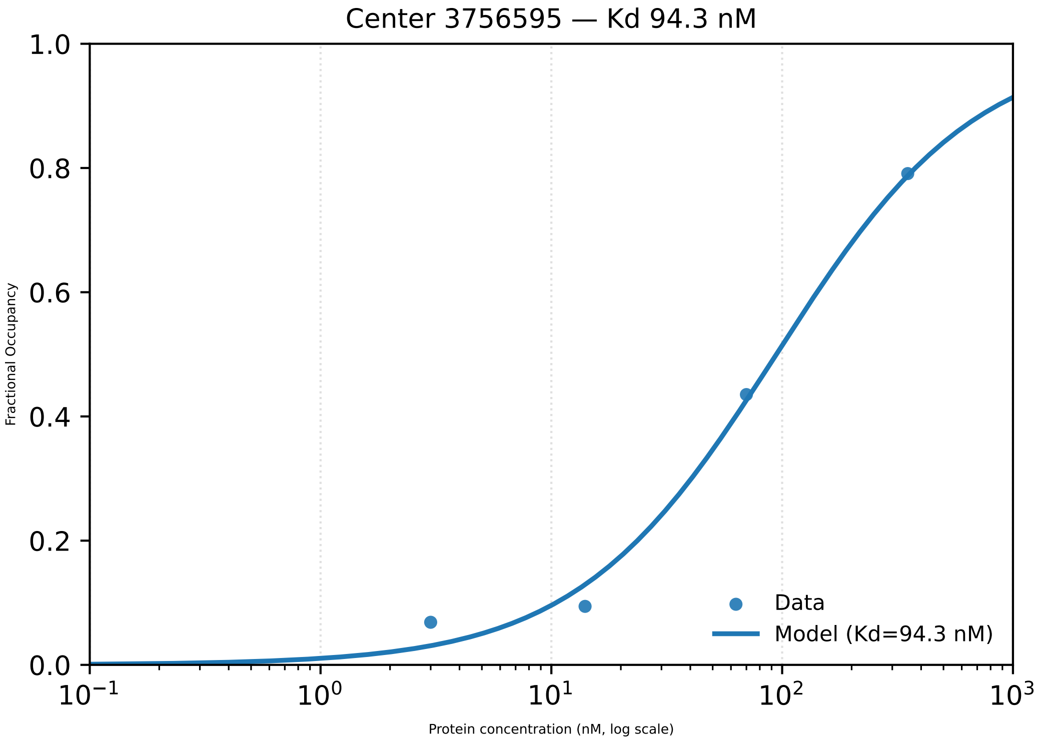

Center 3756595 — Kd 94.3 nM
1.0
0.8
0.6
0.4
0.2
0.0
Fractional Occupancy
$10^{-1}$
$10^{0}$
$10^{1}$
$10^{2}$
$10^{3}$
Protein concentration (nM, log scale)
Data
Model (Kd=94.3 nM)

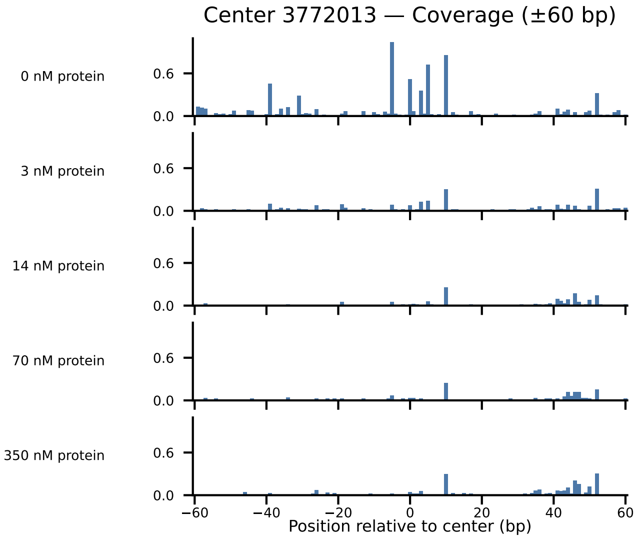
Center 3772013 — Coverage (±60 bp)
0 nM protein
3 nM protein
14 nM protein
70 nM protein
350 nM protein
0.6
0.0
−60
−40
−20
0
20
40
60
Position relative to center (bp)

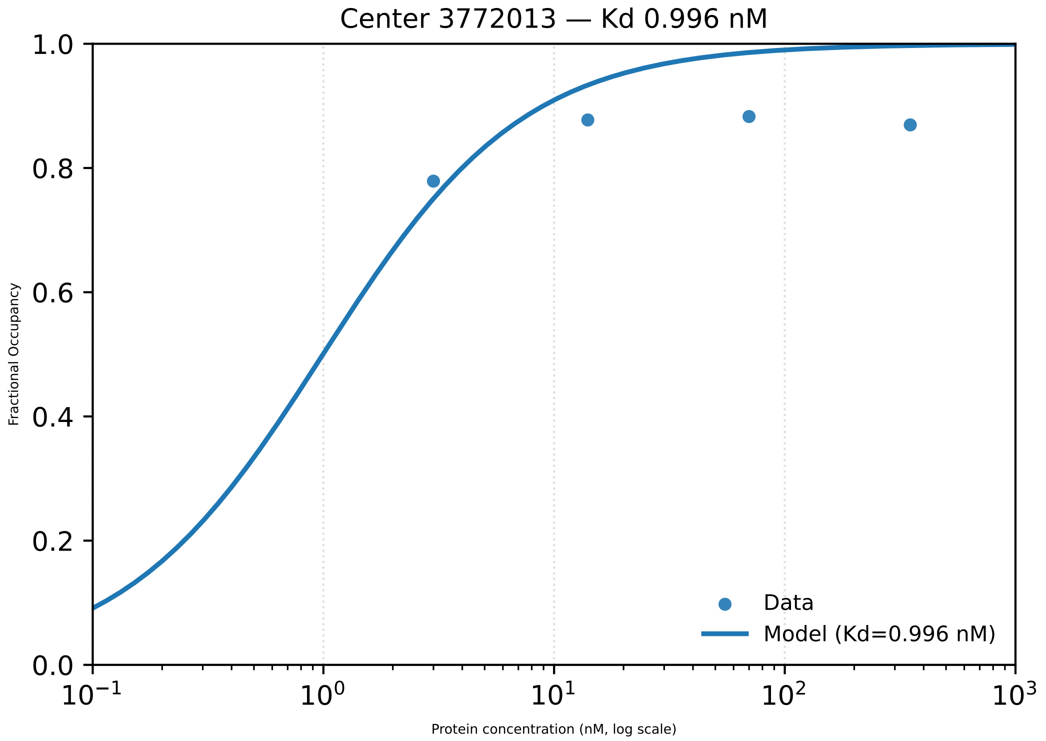
Center 3772013 — Kd 0.996 nM
Fractional Occupancy
1.0
0.8
0.6
0.4
0.2
0.0
$10^{-1}$
$10^{0}$
$10^{1}$
$10^{2}$
$10^{3}$
Protein concentration (nM, log scale)
Data
Model (Kd=0.996 nM)

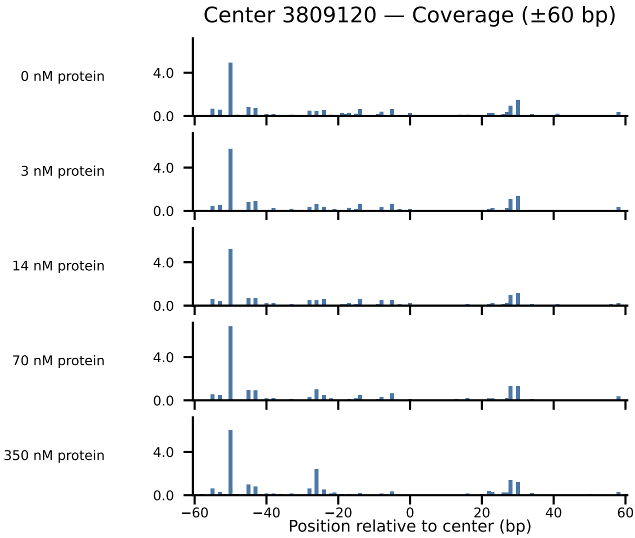

Center 3809120 — Coverage (±60 bp)
0 nM protein
3 nM protein
14 nM protein
70 nM protein
350 nM protein
4.0
0.0
−60
−40
−20
0
20
40
60
Position relative to center (bp)

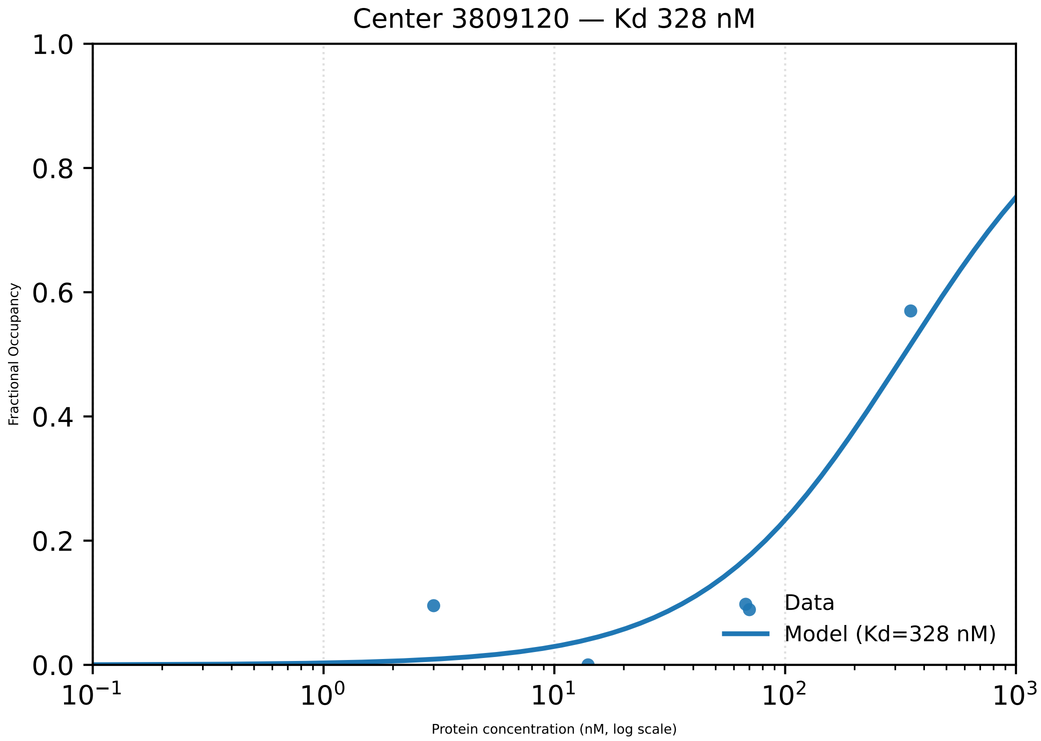

Center 3809120 — Kd 328 nM
Fractional Occupancy
1.0
0.8
0.6
0.4
0.2
0.0
$10^{-1}$
$10^{0}$
$10^{1}$
$10^{2}$
$10^{3}$
Protein concentration (nM, log scale)
Data
Model (Kd=328 nM)

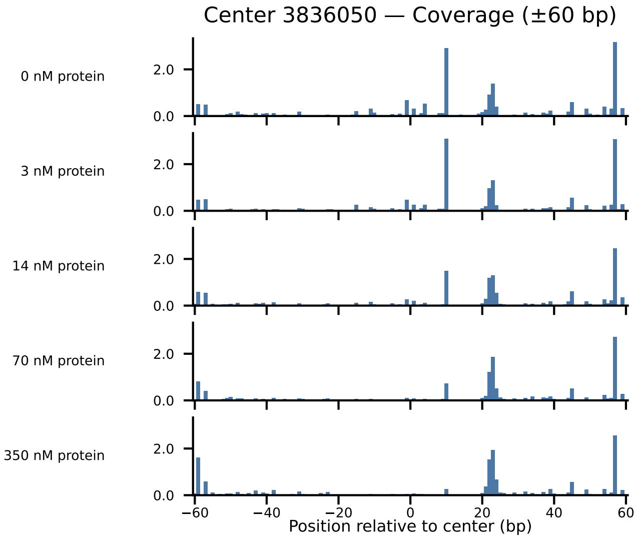

Center 3836050 — Coverage (±60 bp)
0 nM protein
3 nM protein
14 nM protein
70 nM protein
350 nM protein
2.0
0.0
−60
−40
−20
0
20
40
60
Position relative to center (bp)

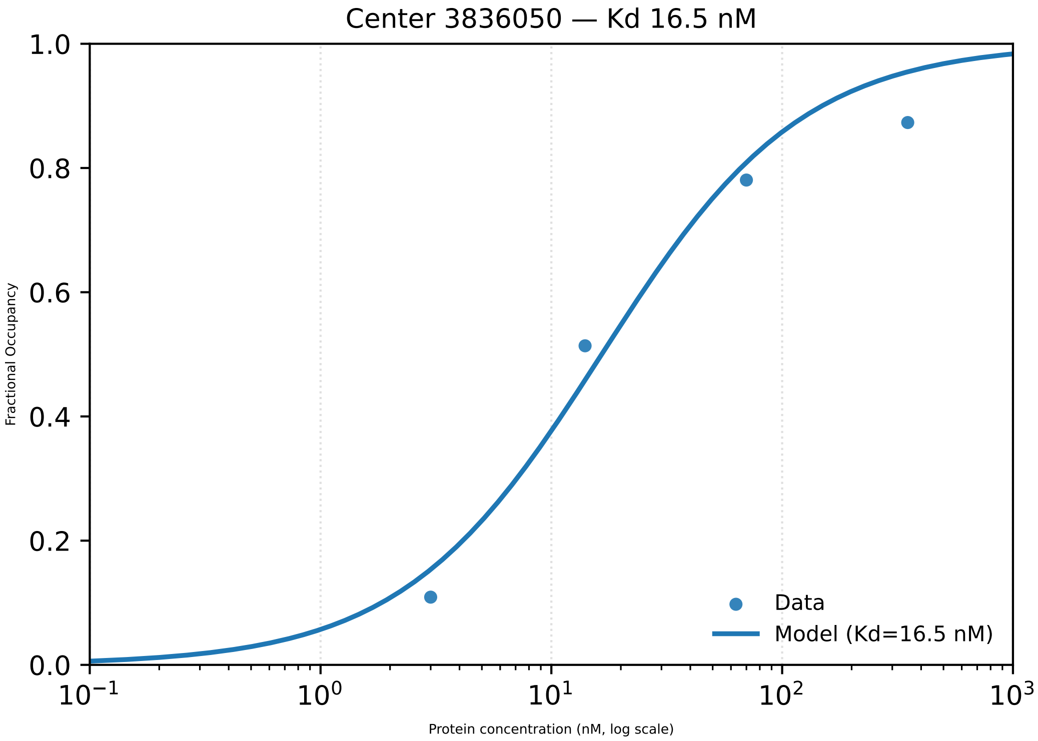

Center 3836050 — Kd 16.5 nM
Fractional Occupancy
Protein concentration (nM, log scale)
0.0
0.2
0.4
0.6
0.8
1.0
$10^{-1}$
$10^{0}$
$10^{1}$
$10^{2}$
$10^{3}$
Data
Model (Kd=16.5 nM)

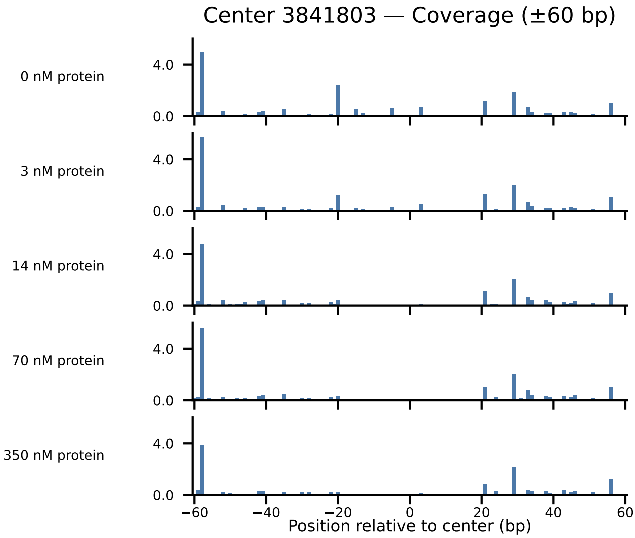

Center 3841803 — Coverage (±60 bp)
0 nM protein
3 nM protein
14 nM protein
70 nM protein
350 nM protein
4.0
0.0
−60
−40
−20
0
20
40
60
Position relative to center (bp)

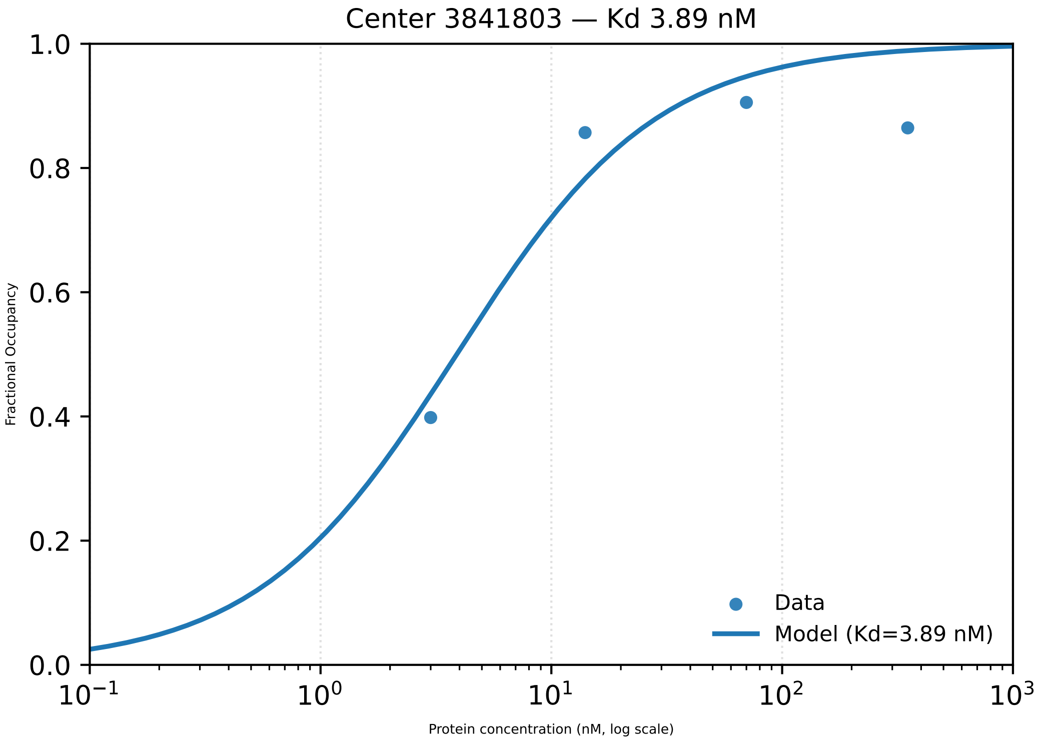

Center 3841803 — Kd 3.89 nM
Fractional Occupancy
1.0
0.8
0.6
0.4
0.2
0.0
$10^{-1}$
$10^{0}$
$10^{1}$
$10^{2}$
$10^{3}$
Protein concentration (nM, log scale)
Data
Model (Kd=3.89 nM)

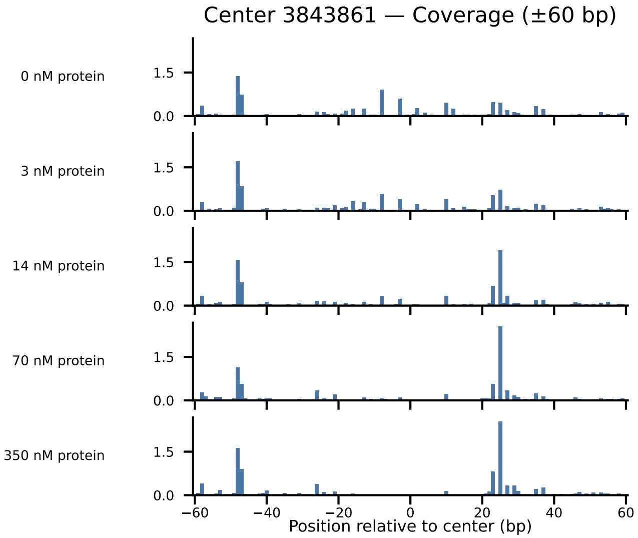

Center 3843861 — Coverage (±60 bp)
0 nM protein
3 nM protein
14 nM protein
70 nM protein
350 nM protein
1.5
0.0
−60
−40
−20
0
20
40
60
Position relative to center (bp)

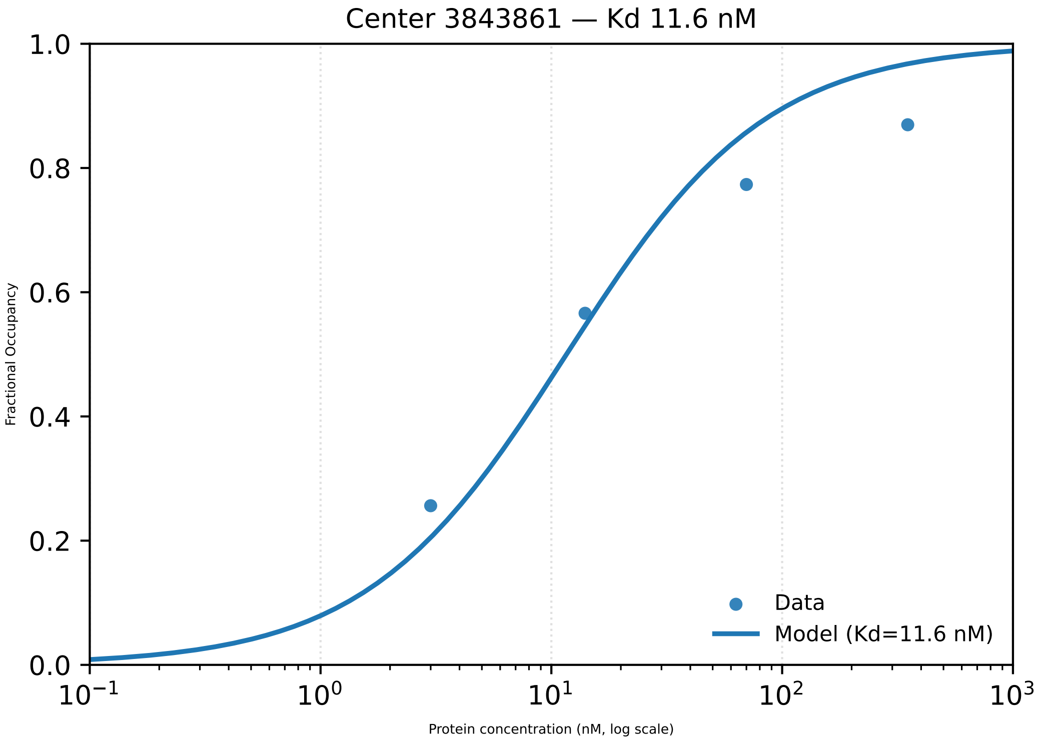

Center 3843861 — Kd 11.6 nM
Fractional Occupancy
1.0
0.8
0.6
0.4
0.2
0.0
$10^{-1}$
$10^{0}$
$10^{1}$
$10^{2}$
$10^{3}$
Protein concentration (nM, log scale)
Data
Model (Kd=11.6 nM)

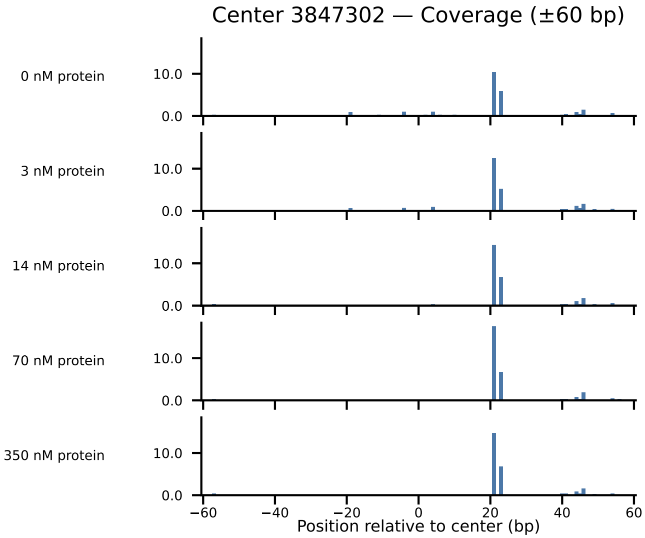

Center 3847302 — Coverage (±60 bp)
0 nM protein
3 nM protein
14 nM protein
70 nM protein
350 nM protein
10.0
0.0
−60
−40
−20
0
20
40
60
Position relative to center (bp)

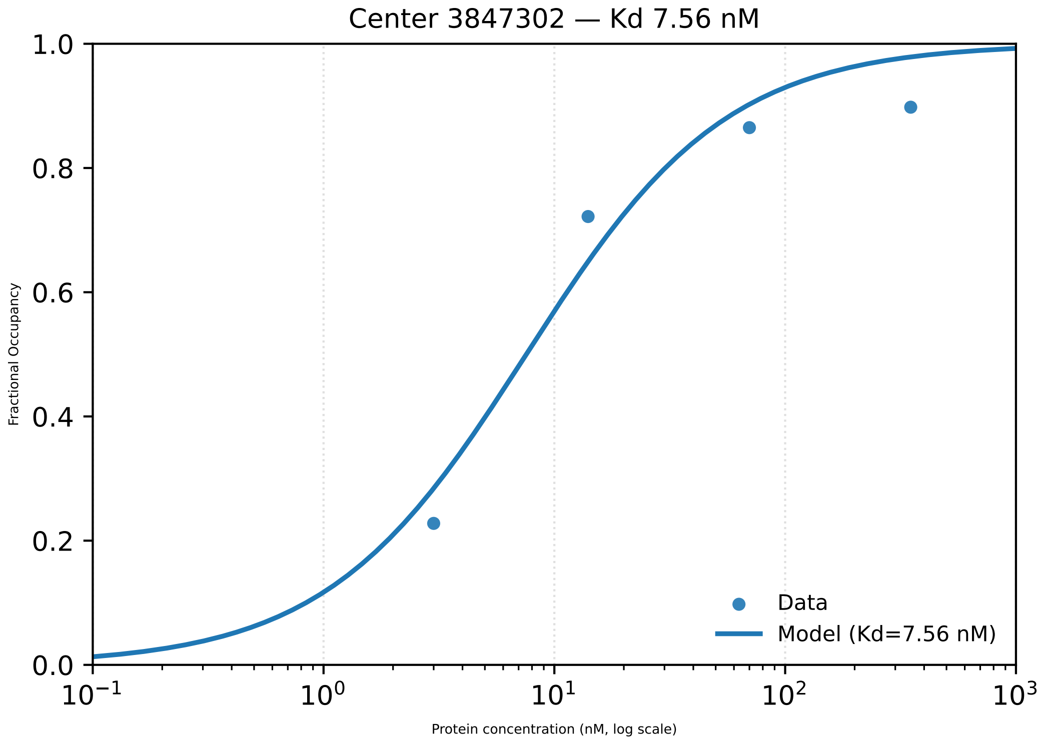

Center 3847302 — Kd 7.56 nM
Fractional Occupancy
1.0
0.8
0.6
0.4
0.2
0.0
$10^{-1}$
$10^{0}$
$10^{1}$
$10^{2}$
$10^{3}$
Protein concentration (nM, log scale)
Data
Model (Kd=7.56 nM)

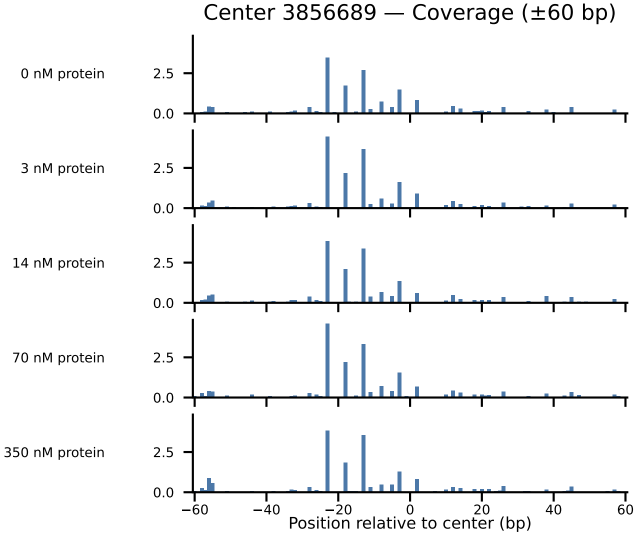

Center 3856689 — Coverage (±60 bp)
0 nM protein
3 nM protein
14 nM protein
70 nM protein
350 nM protein
2.5
0.0
−60
−40
−20
0
20
40
60
Position relative to center (bp)

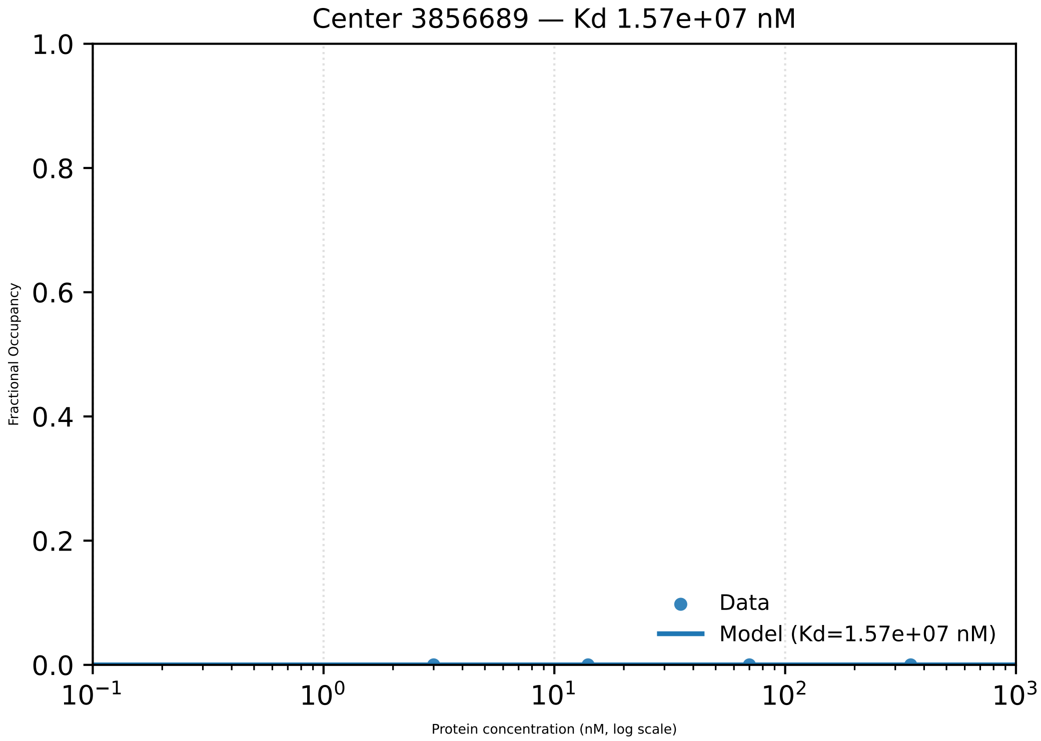
Center 3856689 — Kd 1.57e+07 nM
1.0
0.8
0.6
0.4
0.2
0.0
Fractional Occupancy
$10^{-1}$
$10^{0}$
$10^{1}$
$10^{2}$
$10^{3}$
Protein concentration (nM, log scale)
Data
Model (Kd=1.57e+07 nM)

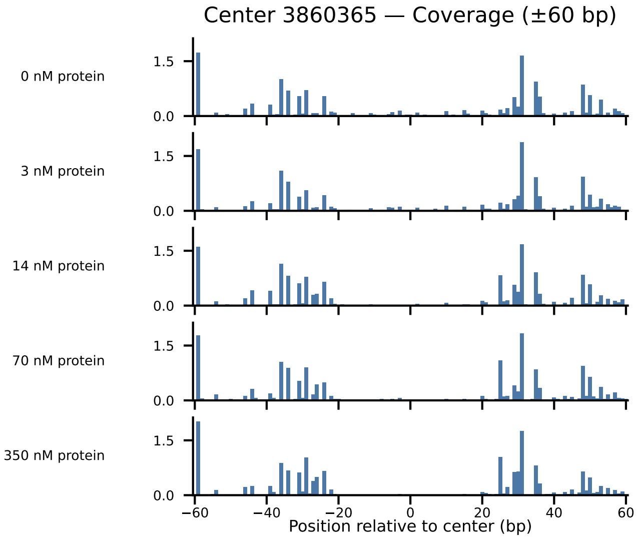

Center 3860365 — Coverage (±60 bp)
0 nM protein
3 nM protein
14 nM protein
70 nM protein
350 nM protein
1.5
0.0
−60
−40
−20
0
20
40
60
Position relative to center (bp)

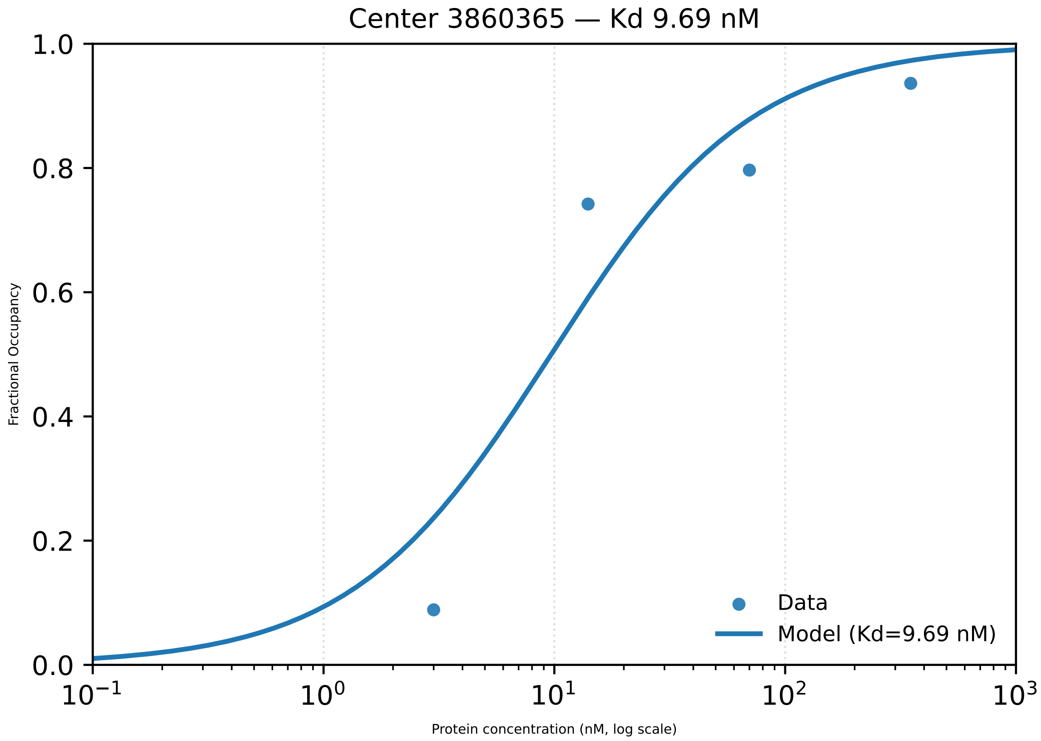

Center 3860365 — Kd 9.69 nM
Fractional Occupancy
1.0
0.8
0.6
0.4
0.2
0.0
$10^{-1}$
$10^{0}$
$10^{1}$
$10^{2}$
$10^{3}$
Protein concentration (nM, log scale)
Data
Model (Kd=9.69 nM)

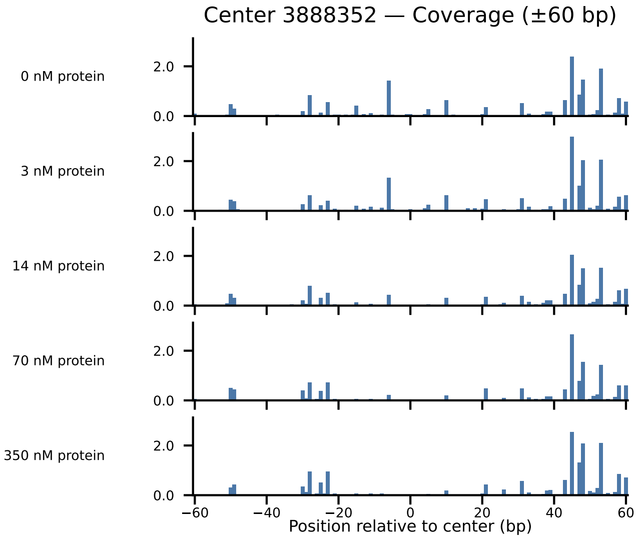

Center 3888352 — Coverage (±60 bp)
0 nM protein
3 nM protein
14 nM protein
70 nM protein
350 nM protein
2.0
0.0
−60
−40
−20
0
20
40
60
Position relative to center (bp)

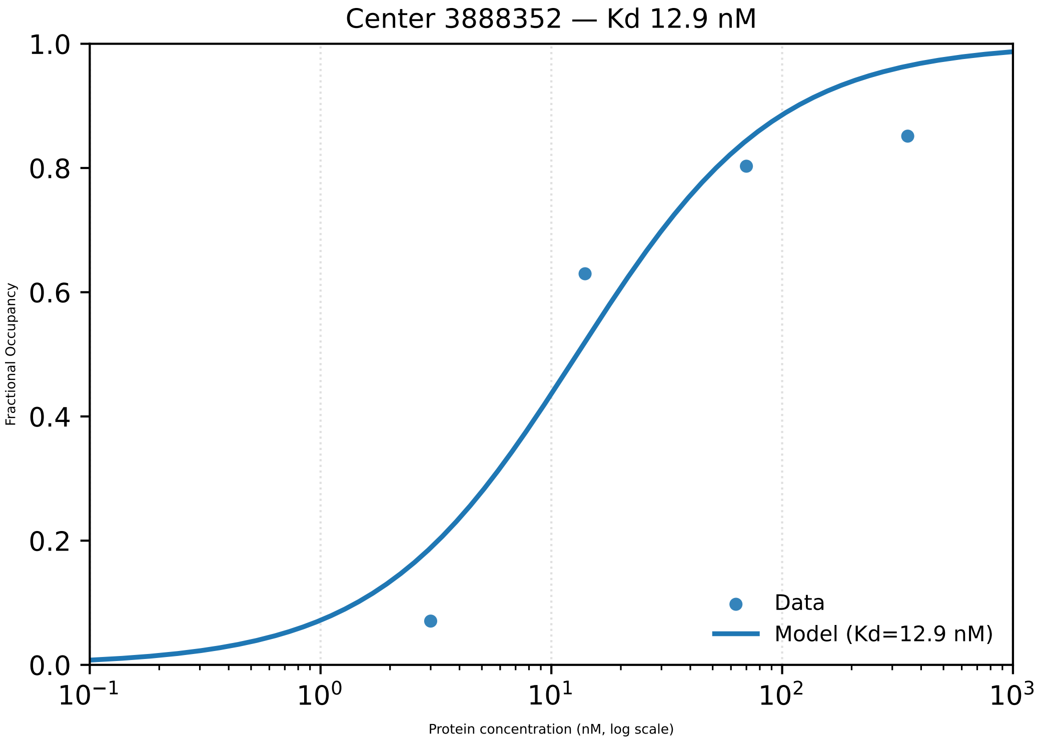
Center 3888352 — Kd 12.9 nM
1.0
0.8
0.6
0.4
0.2
0.0
Fractional Occupancy
$10^{-1}$
$10^{0}$
$10^{1}$
$10^{2}$
$10^{3}$
Protein concentration (nM, log scale)
Data
Model (Kd=12.9 nM)

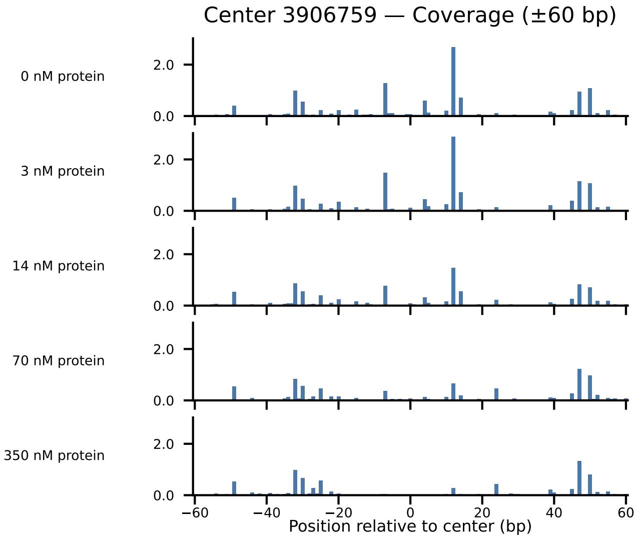

Center 3906759 — Coverage (±60 bp)
0 nM protein
3 nM protein
14 nM protein
70 nM protein
350 nM protein
2.0
0.0
−60
−40
−20
0
20
40
60
Position relative to center (bp)

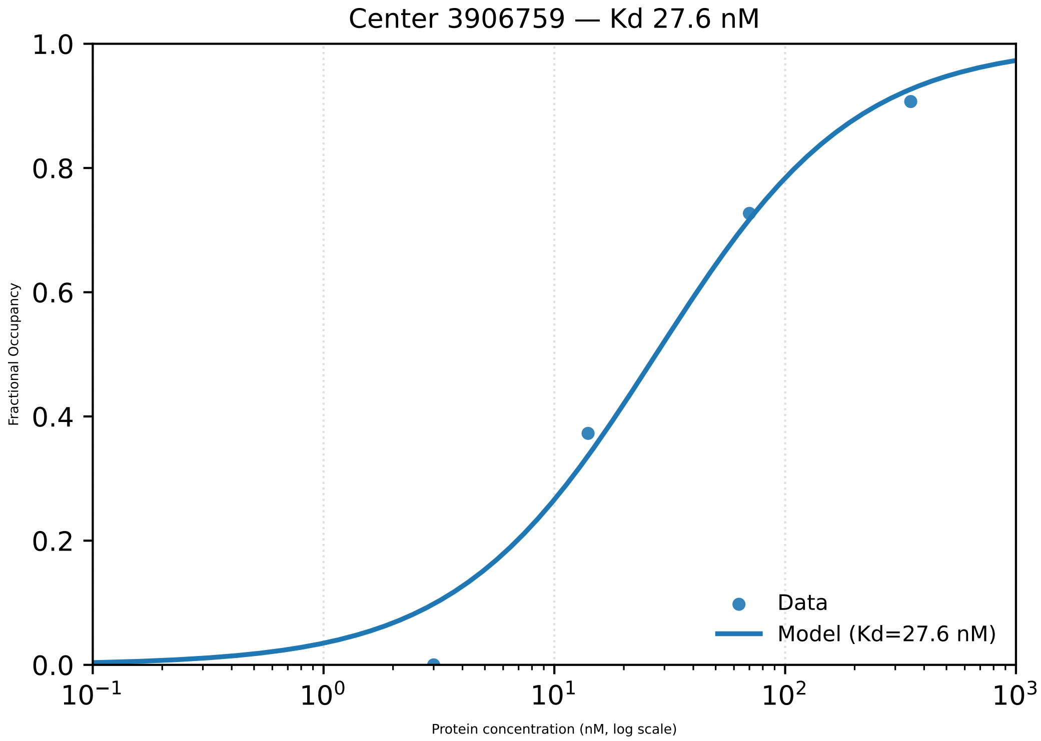

Center 3906759 — Kd 27.6 nM
1.0
0.8
0.6
0.4
0.2
0.0
Fractional Occupancy
$10^{-1}$
$10^{0}$
$10^{1}$
$10^{2}$
$10^{3}$
Protein concentration (nM, log scale)
Data
Model (Kd=27.6 nM)

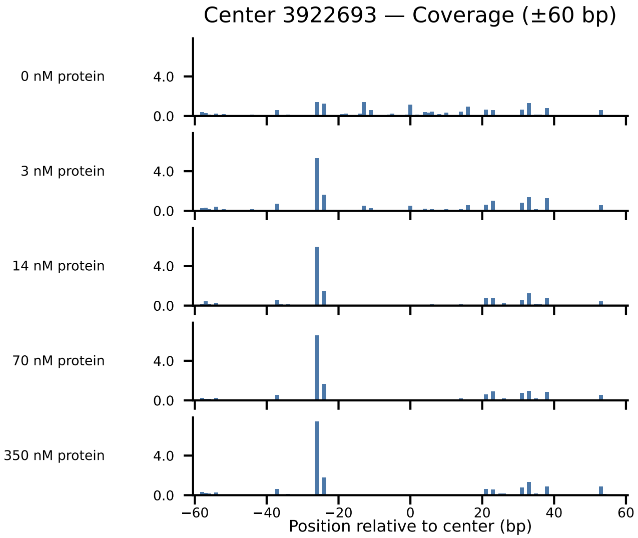

Center 3922693 — Coverage (±60 bp)
0 nM protein
3 nM protein
14 nM protein
70 nM protein
350 nM protein
4.0
0.0
−60
−40
−20
0
20
40
60
Position relative to center (bp)

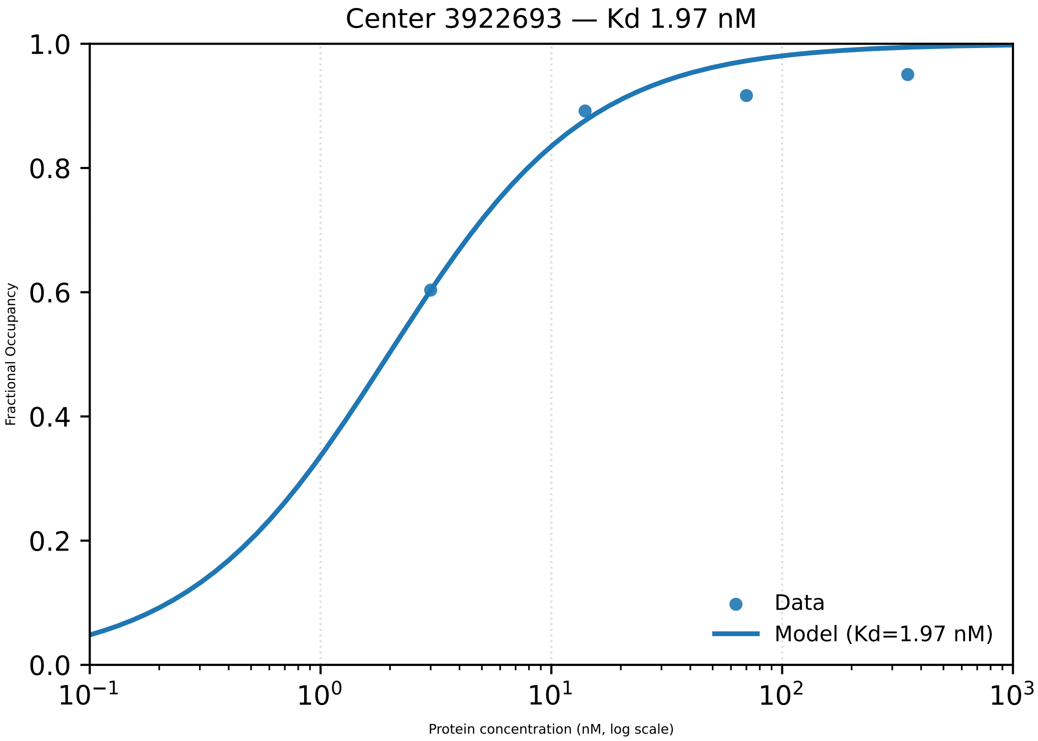

Center 3922693 — Kd 1.97 nM
Fractional Occupancy
1.0
0.8
0.6
0.4
0.2
0.0
$10^{-1}$
$10^{0}$
$10^{1}$
$10^{2}$
$10^{3}$
Protein concentration (nM, log scale)
Data
Model (Kd=1.97 nM)

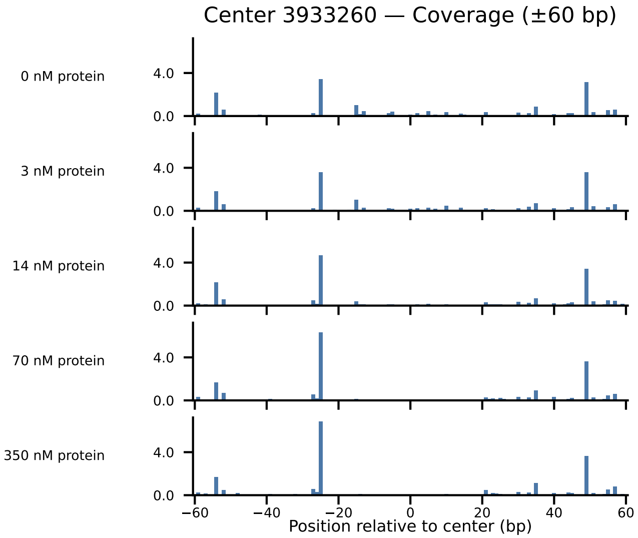
Center 3933260 — Coverage (±60 bp)
0 nM protein
3 nM protein
14 nM protein
70 nM protein
350 nM protein
4.0
0.0
−60
−40
−20
0
20
40
60
Position relative to center (bp)

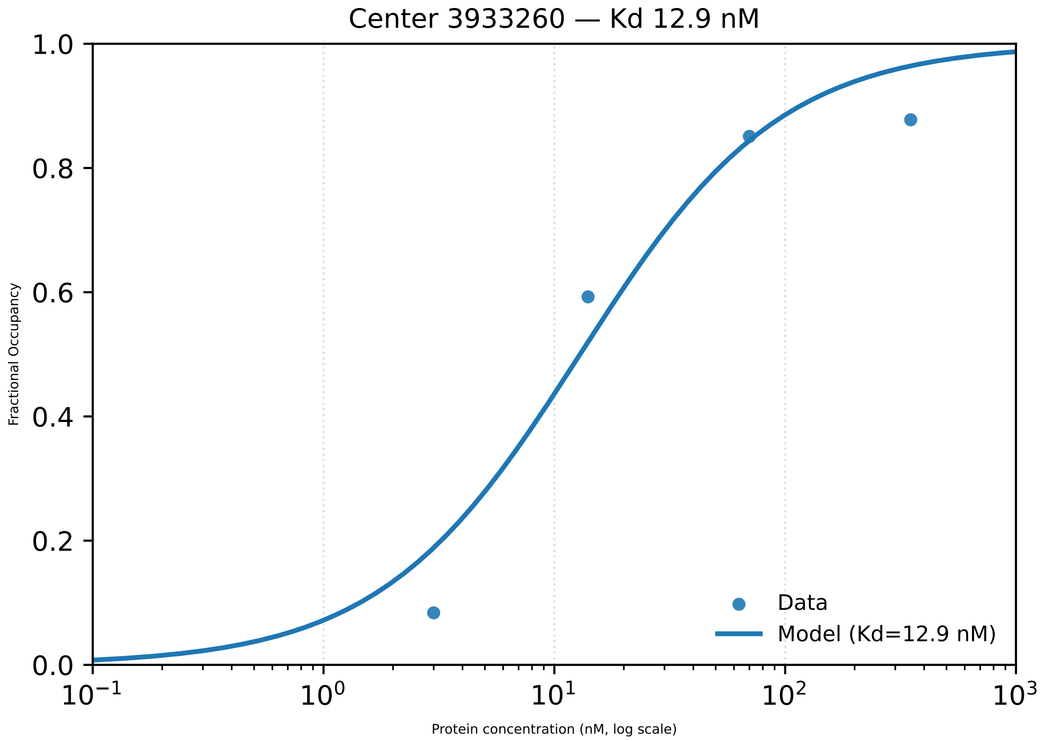
Center 3933260 — Kd 12.9 nM
Fractional Occupancy
1.0
0.8
0.6
0.4
0.2
0.0
$10^{-1}$
$10^{0}$
$10^{1}$
$10^{2}$
$10^{3}$
Protein concentration (nM, log scale)
Data
Model (Kd=12.9 nM)

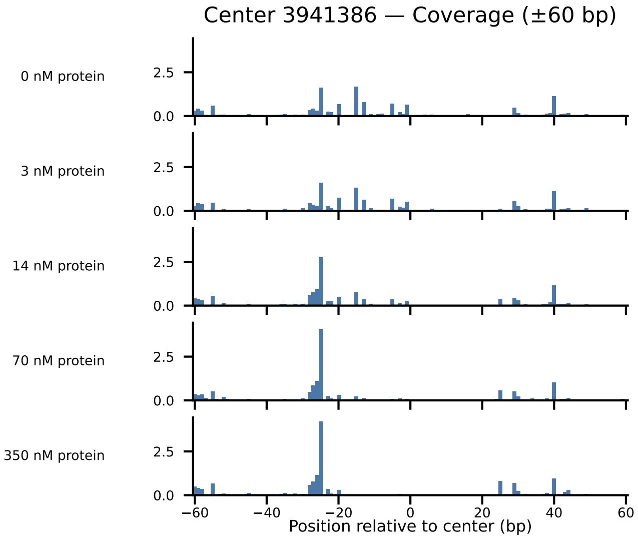

Center 3941386 — Coverage (±60 bp)
0 nM protein
3 nM protein
14 nM protein
70 nM protein
350 nM protein
2.5
0.0
−60
−40
−20
0
20
40
60
Position relative to center (bp)

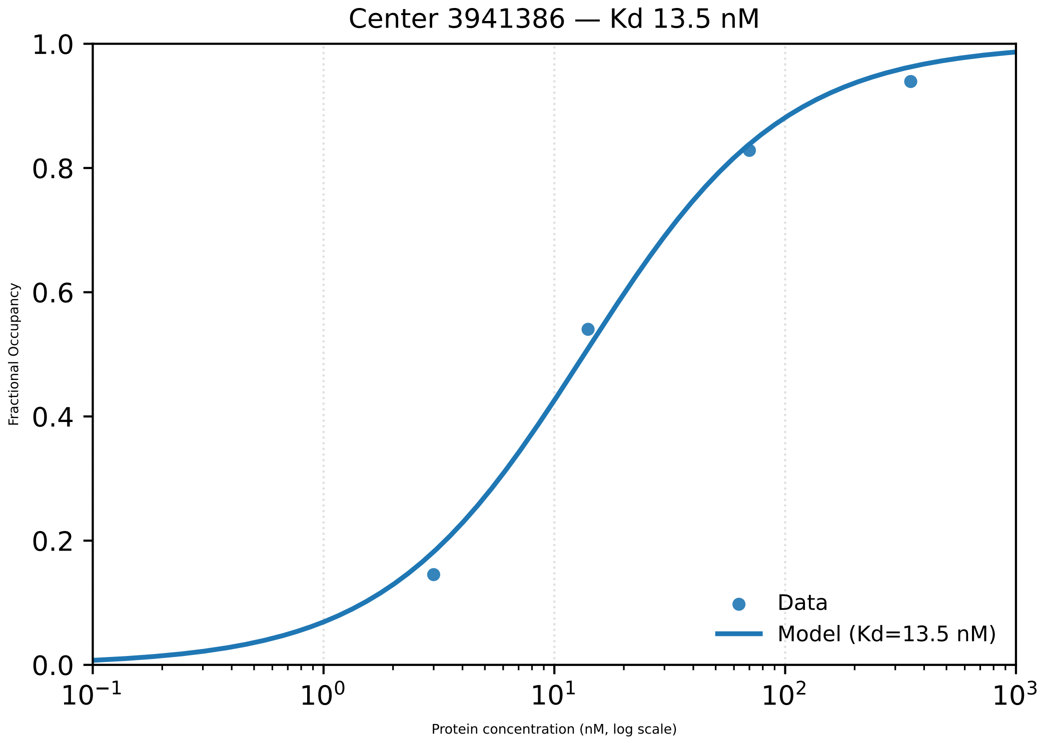

Center 3941386 — Kd 13.5 nM
1.0
0.8
0.6
0.4
0.2
0.0
Fractional Occupancy
Data
Model (Kd=13.5 nM)
$10^{-1}$
$10^{0}$
$10^{1}$
$10^{2}$
$10^{3}$
Protein concentration (nM, log scale)

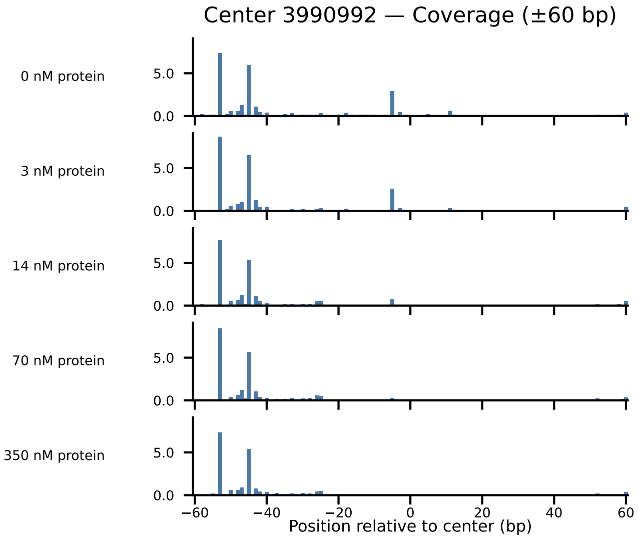
Center 3990992 — Coverage (±60 bp)
0 nM protein
3 nM protein
14 nM protein
70 nM protein
350 nM protein
5.0
0.0
−60
−40
−20
0
20
40
60
Position relative to center (bp)

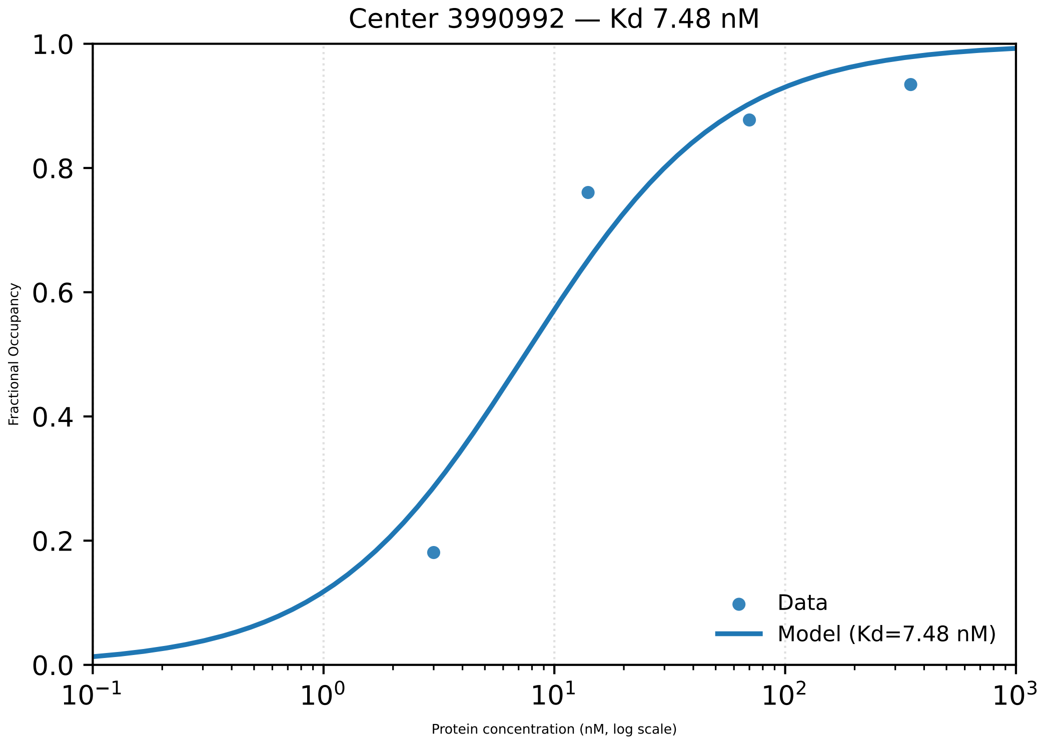

Center 3990992 — Kd 7.48 nM
1.0
0.8
0.6
0.4
0.2
0.0
Fractional Occupancy
$10^{-1}$
$10^{0}$
$10^{1}$
$10^{2}$
$10^{3}$
Protein concentration (nM, log scale)
Data
Model (Kd=7.48 nM)

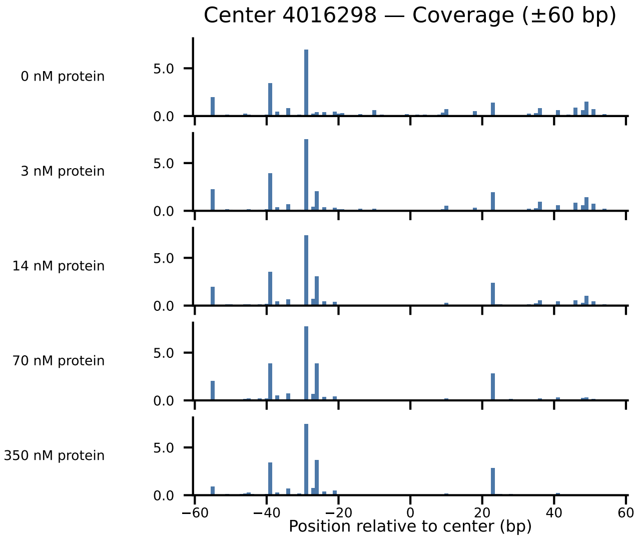
Center 4016298 — Coverage (±60 bp)
0 nM protein
3 nM protein
14 nM protein
70 nM protein
350 nM protein
0.0
5.0
−60
−40
−20
0
20
40
60
Position relative to center (bp)

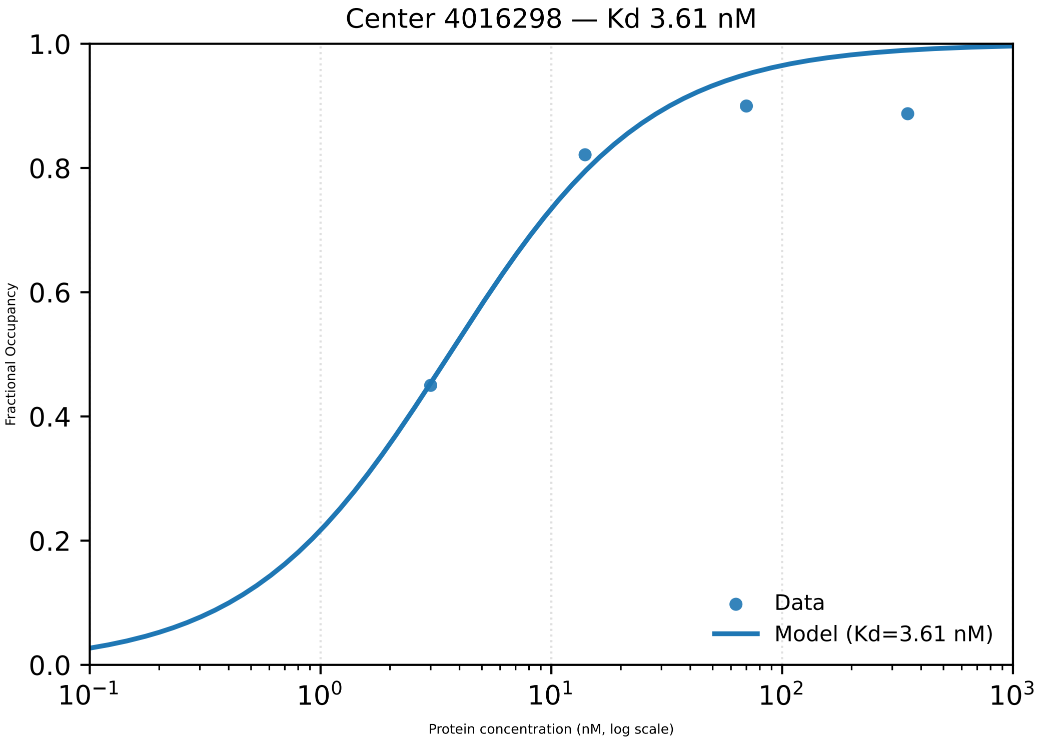

Center 4016298 — Kd 3.61 nM
1.0
0.8
0.6
0.4
0.2
0.0
Fractional Occupancy
$10^{-1}$
$10^{0}$
$10^{1}$
$10^{2}$
$10^{3}$
Protein concentration (nM, log scale)
Data
Model (Kd=3.61 nM)

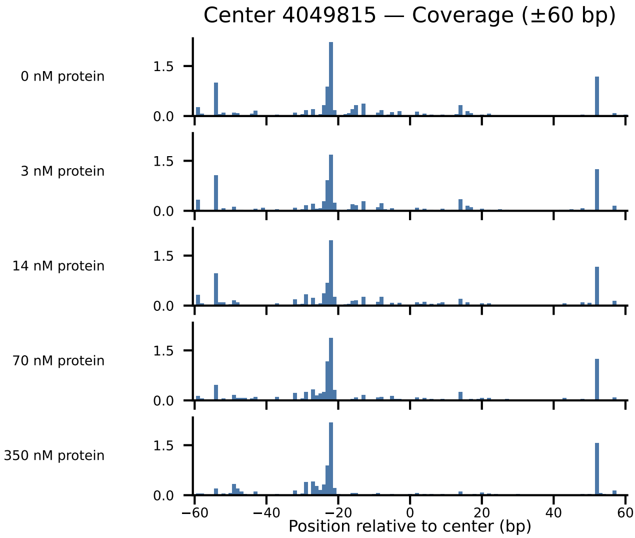

Center 4049815 — Coverage (±60 bp)
0 nM protein
3 nM protein
14 nM protein
70 nM protein
350 nM protein
1.5
0.0
−60
−40
−20
0
20
40
60
Position relative to center (bp)

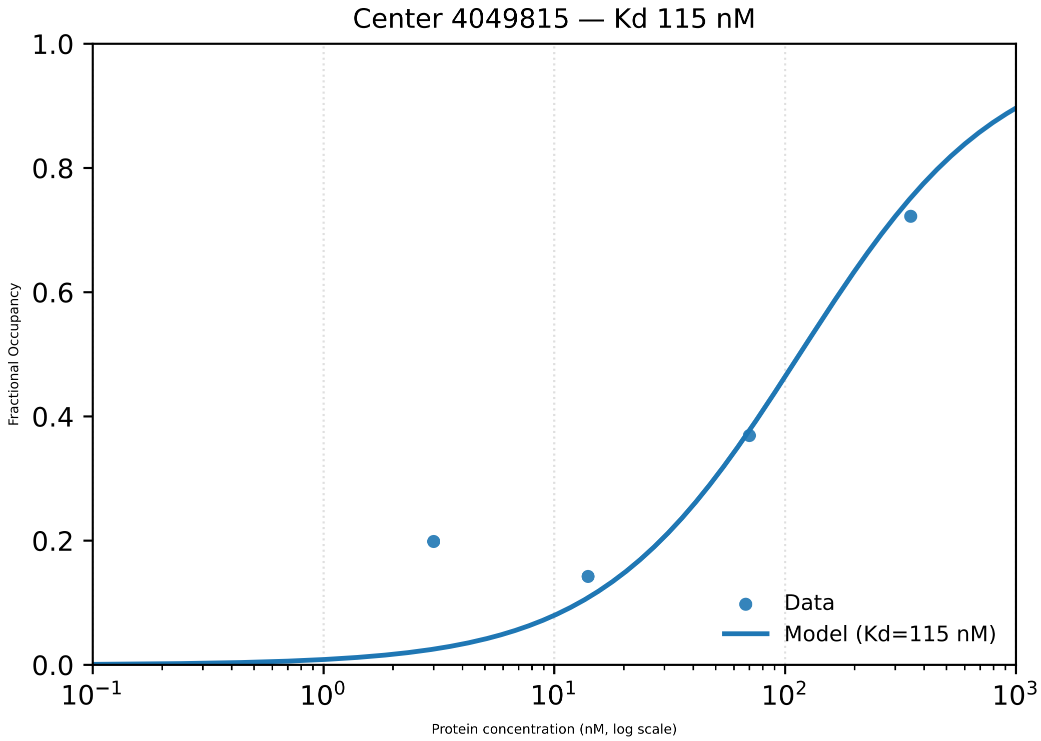

Center 4049815 — Kd 115 nM
Fractional Occupancy
Protein concentration (nM, log scale)
1.0
0.8
0.6
0.4
0.2
0.0
$10^{-1}$
$10^{0}$
$10^{1}$
$10^{2}$
$10^{3}$
Data
Model (Kd=115 nM)

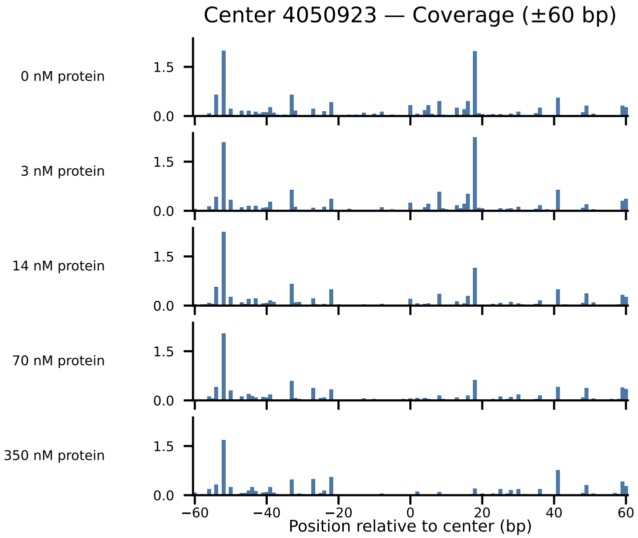

Center 4050923 — Coverage (±60 bp)
0 nM protein
3 nM protein
14 nM protein
70 nM protein
350 nM protein
1.5
0.0
−60
−40
−20
0
20
40
60
Position relative to center (bp)

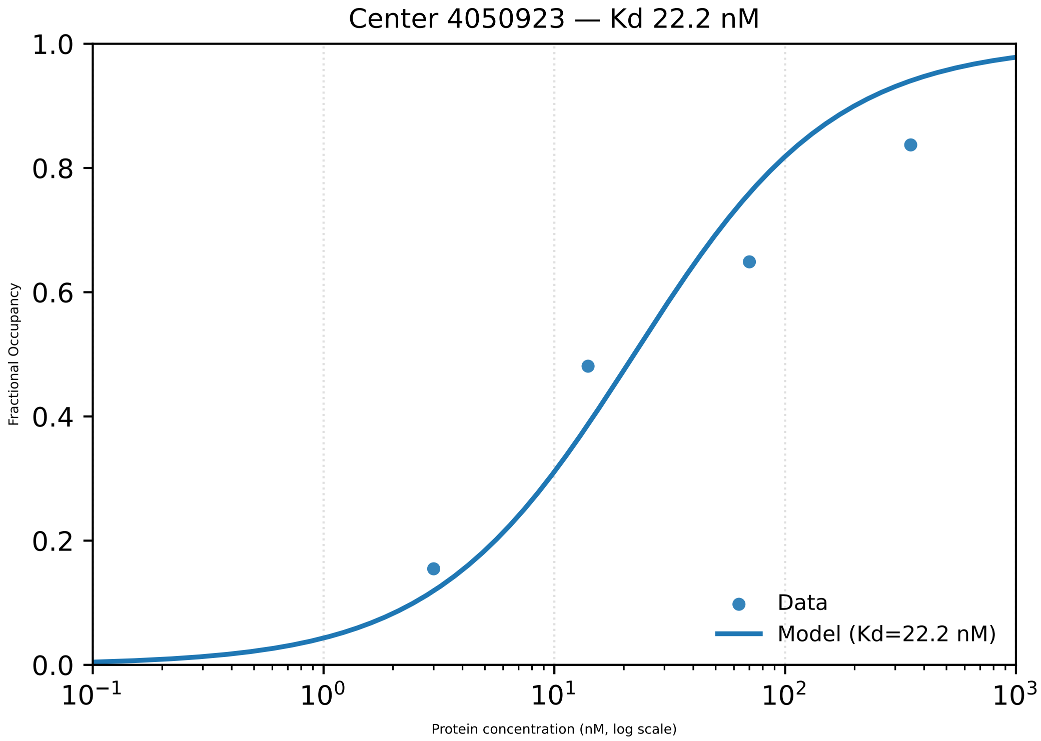

Center 4050923 — Kd 22.2 nM
Fractional Occupancy
1.0
0.8
0.6
0.4
0.2
0.0
$10^{-1}$
$10^{0}$
$10^{1}$
$10^{2}$
$10^{3}$
Protein concentration (nM, log scale)
Data
Model (Kd=22.2 nM)

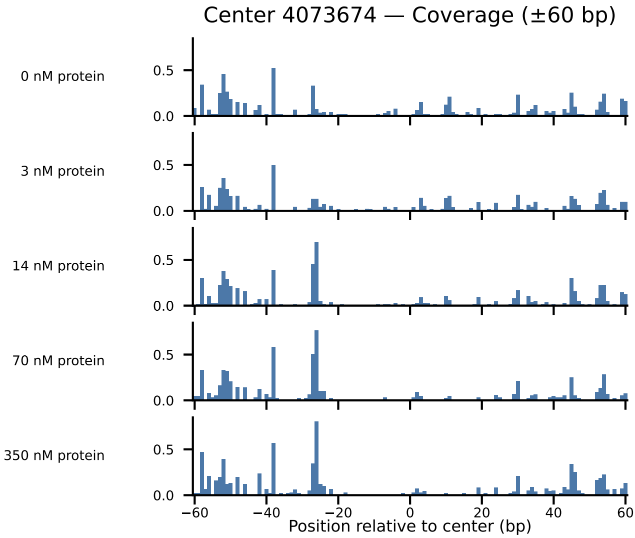
Center 4073674 — Coverage (±60 bp)
0 nM protein
3 nM protein
14 nM protein
70 nM protein
350 nM protein
0.5
0.0
−60
−40
−20
0
20
40
60
Position relative to center (bp)

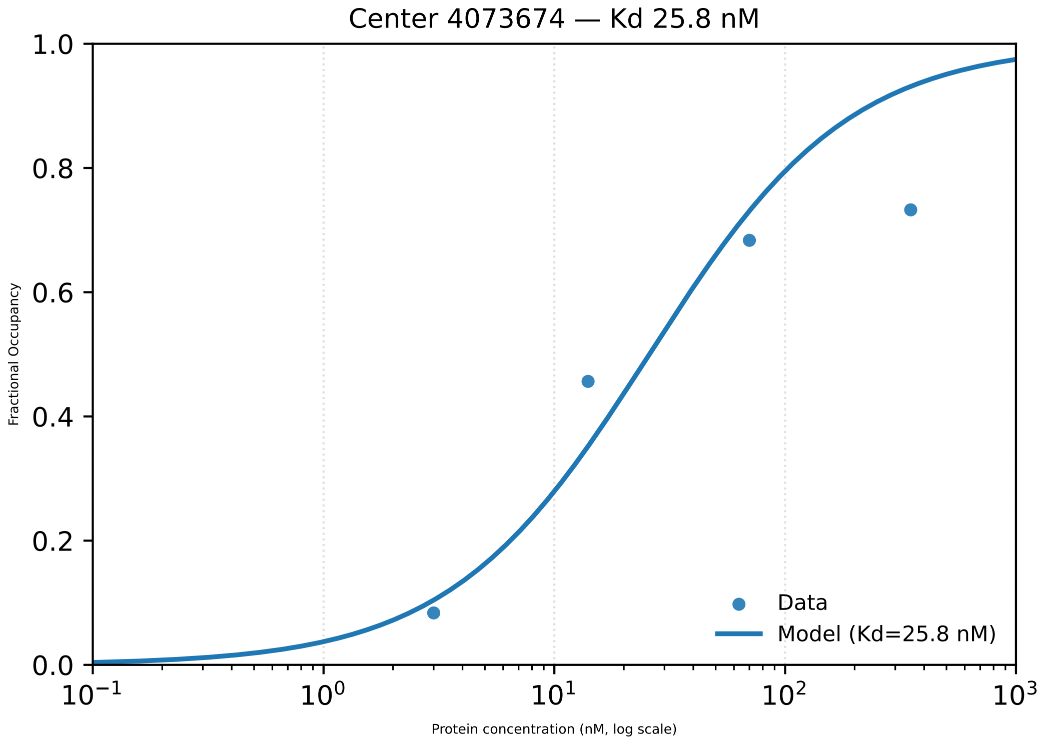

Center 4073674 — Kd 25.8 nM
1.0
0.8
0.6
0.4
0.2
0.0
Fractional Occupancy
10^−1
10^0
10^1
10^2
10^3
Protein concentration (nM, log scale)
Data
Model (Kd=25.8 nM)

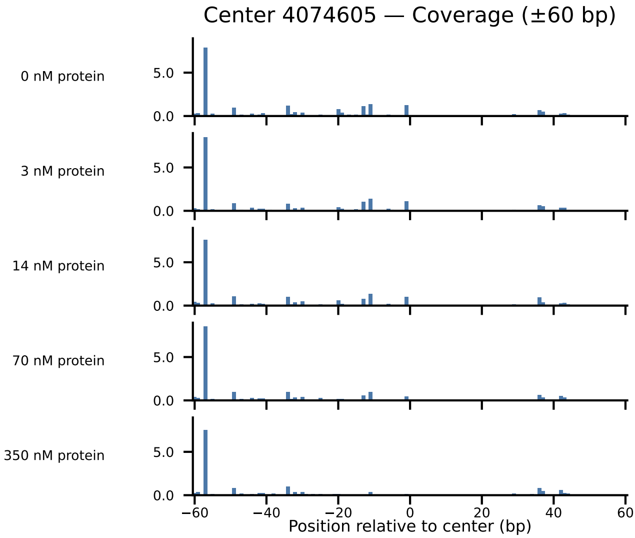

Center 4074605 — Coverage (±60 bp)
0 nM protein
3 nM protein
14 nM protein
70 nM protein
350 nM protein
5.0
0.0
−60
−40
−20
0
20
40
60
Position relative to center (bp)

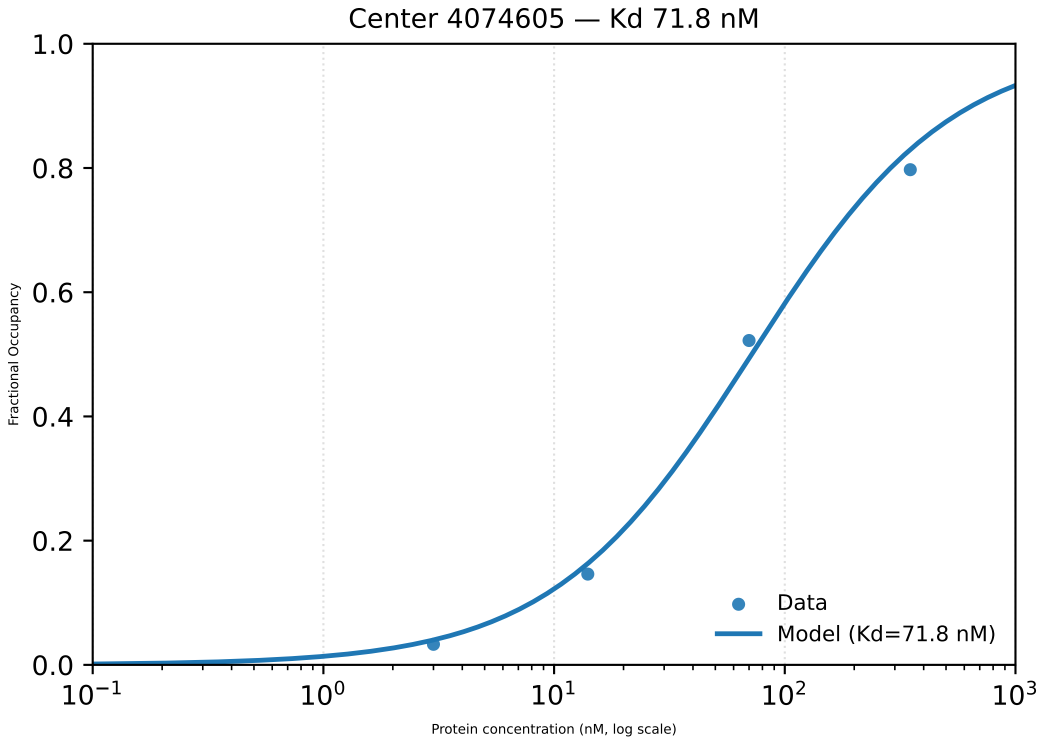

Center 4074605 — Kd 71.8 nM
Fractional Occupancy
Protein concentration (nM, log scale)
0.0
0.2
0.4
0.6
0.8
1.0
$10^{-1}$
$10^{0}$
$10^{1}$
$10^{2}$
$10^{3}$
Data
Model (Kd=71.8 nM)

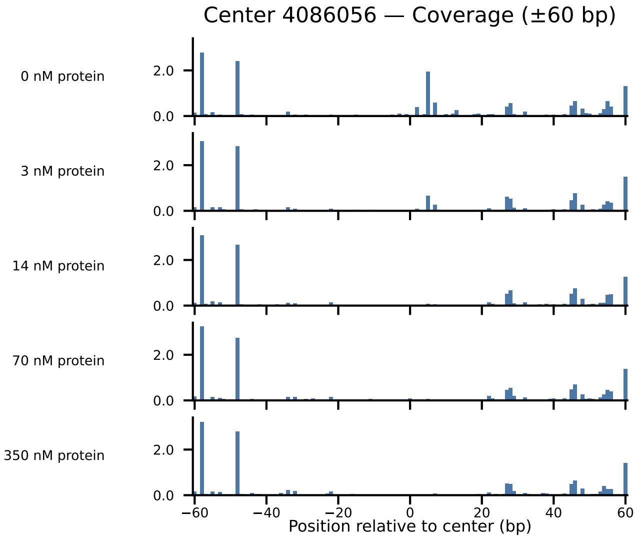

Center 4086056 — Coverage (±60 bp)
0 nM protein
3 nM protein
14 nM protein
70 nM protein
350 nM protein
2.0
0.0
−60
−40
−20
0
20
40
60
Position relative to center (bp)

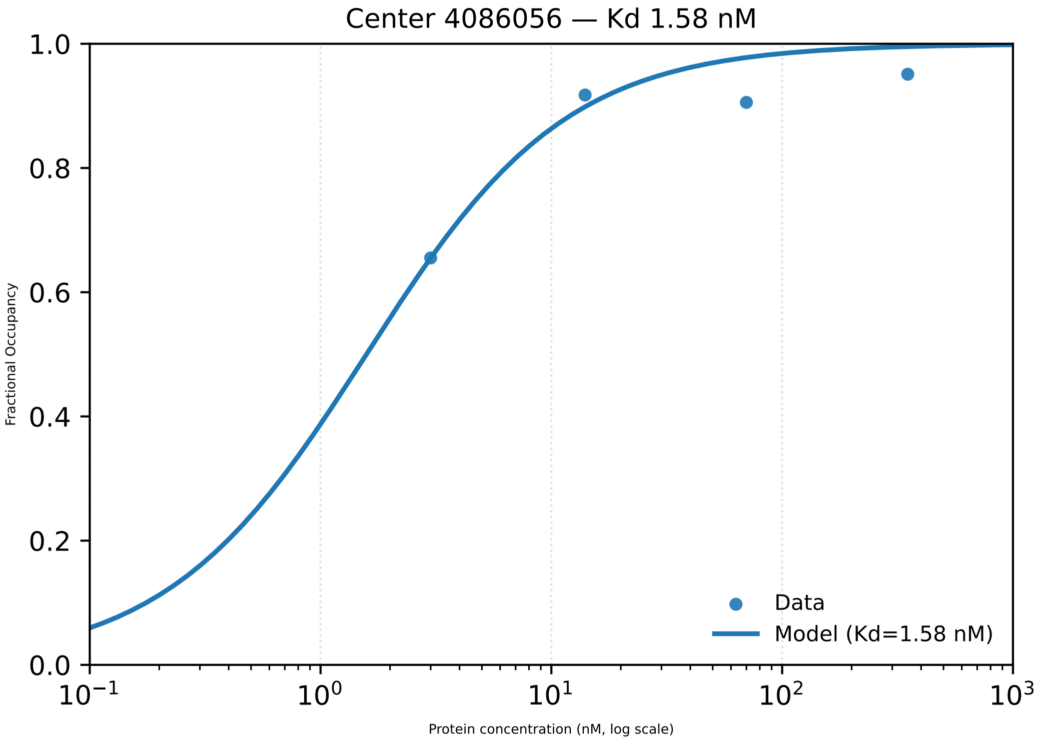

Center 4086056 — Kd 1.58 nM
1.0
0.8
0.6
0.4
0.2
0.0
Fractional Occupancy
$10^{-1}$
$10^{0}$
$10^{1}$
$10^{2}$
$10^{3}$
Protein concentration (nM, log scale)
Data
Model (Kd=1.58 nM)

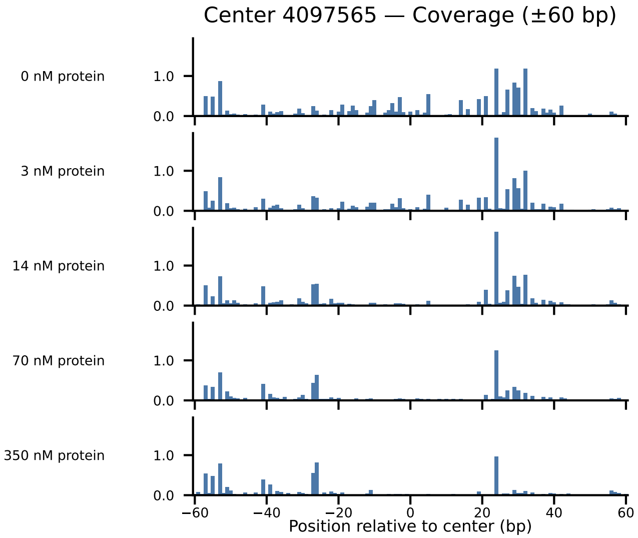

Center 4097565 — Coverage (±60 bp)
0 nM protein
3 nM protein
14 nM protein
70 nM protein
350 nM protein
1.0
0.0
−60
−40
−20
0
20
40
60
Position relative to center (bp)

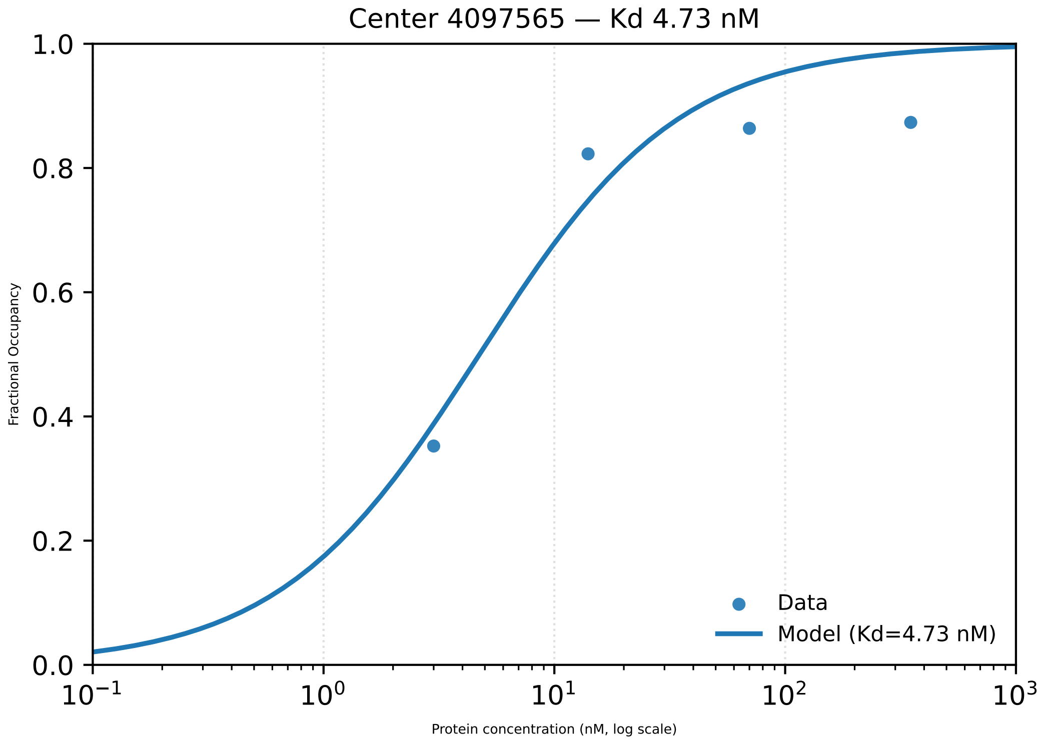

Center 4097565 — Kd 4.73 nM
1.0
0.8
0.6
0.4
0.2
0.0
Fractional Occupancy
$10^{-1}$
$10^{0}$
$10^{1}$
$10^{2}$
$10^{3}$
Protein concentration (nM, log scale)
Data
Model (Kd=4.73 nM)

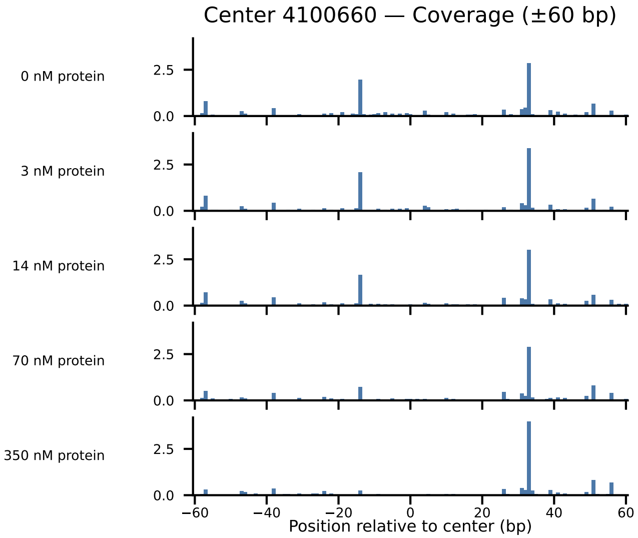

Center 4100660 — Coverage (±60 bp)
0 nM protein
3 nM protein
14 nM protein
70 nM protein
350 nM protein
2.5
0.0
−60
−40
−20
0
20
40
60
Position relative to center (bp)

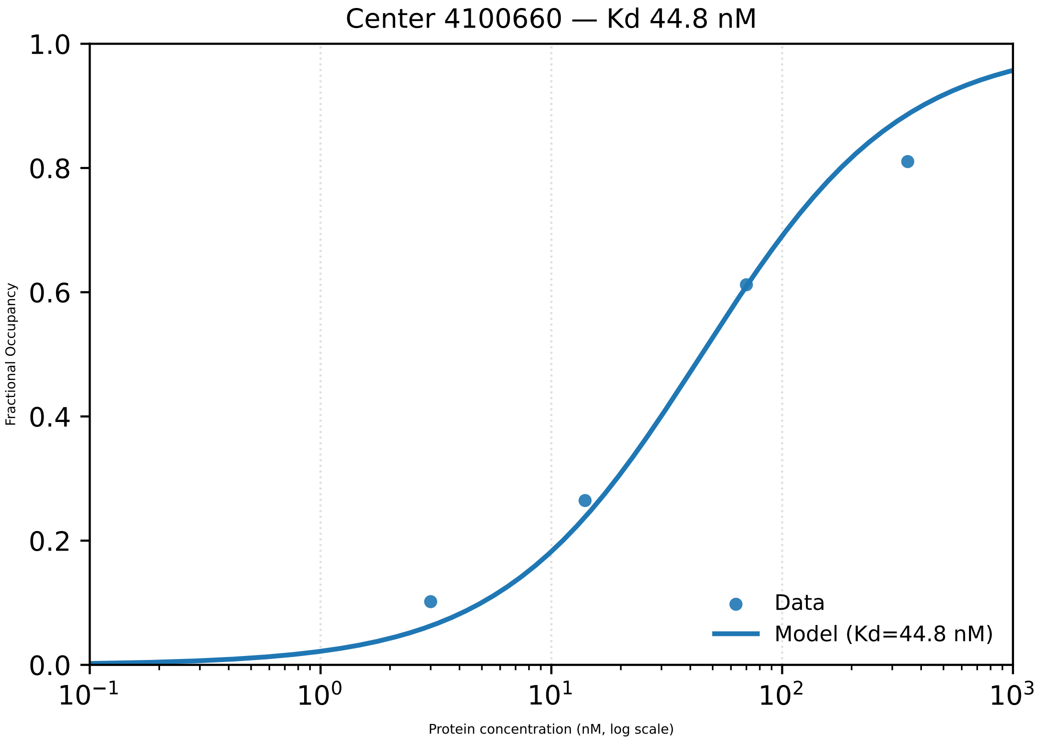

Center 4100660 — Kd 44.8 nM
1.0
0.8
0.6
0.4
0.2
0.0
Fractional Occupancy
$10^{-1}$
$10^{0}$
$10^{1}$
$10^{2}$
$10^{3}$
Protein concentration (nM, log scale)
Data
Model (Kd=44.8 nM)

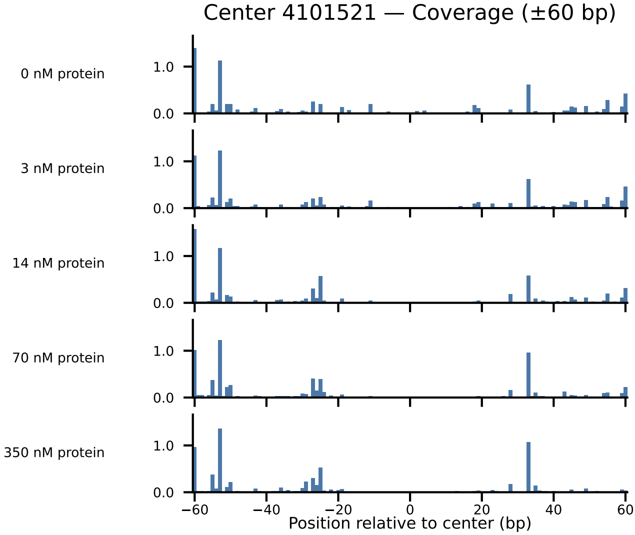

Center 4101521 — Coverage (±60 bp)
0 nM protein
3 nM protein
14 nM protein
70 nM protein
350 nM protein
1.0
0.0
−60
−40
−20
0
20
40
60
Position relative to center (bp)

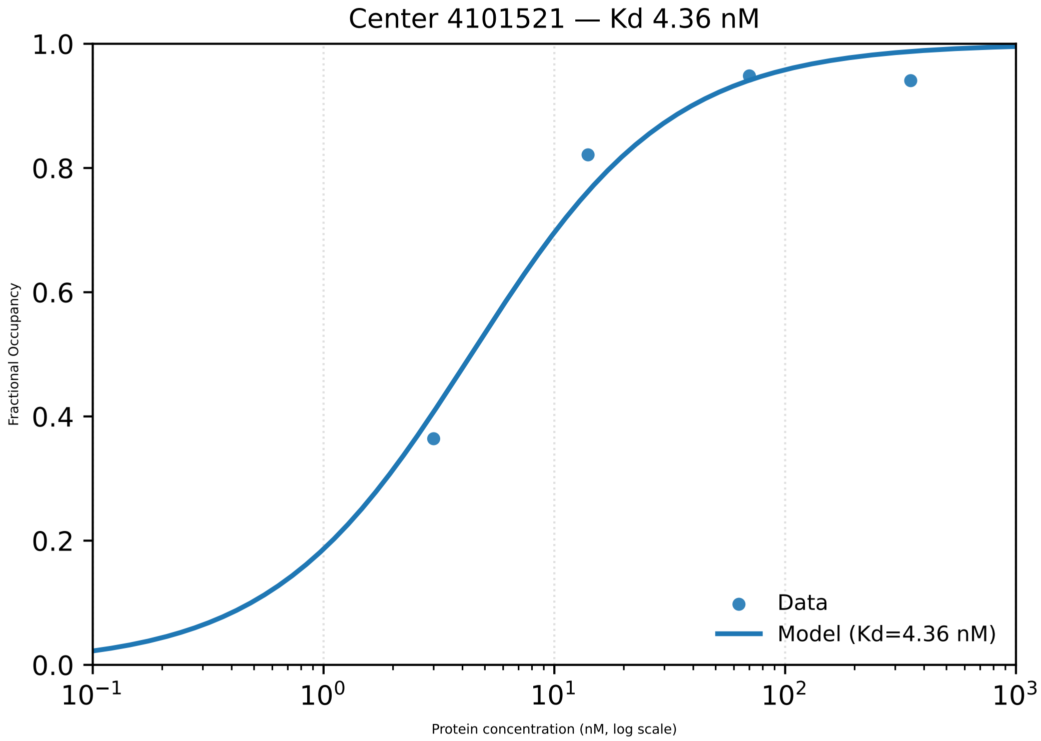

Center 4101521 — Kd 4.36 nM
Fractional Occupancy
0.0
0.2
0.4
0.6
0.8
1.0
$10^{-1}$
$10^{0}$
$10^{1}$
$10^{2}$
$10^{3}$
Protein concentration (nM, log scale)
Data
Model (Kd=4.36 nM)

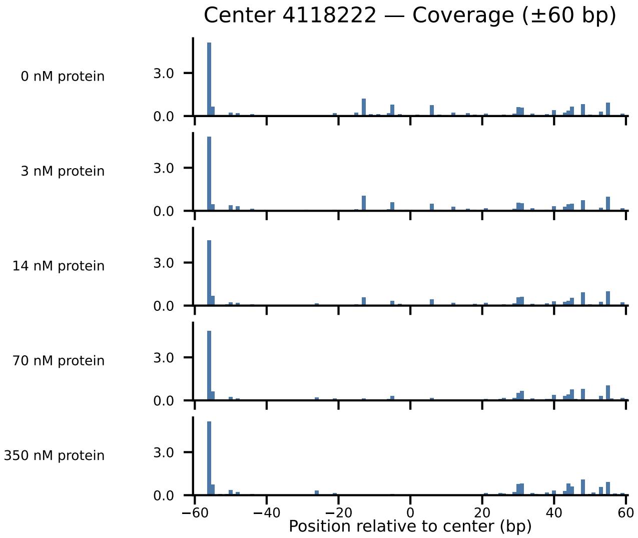

Center 4118222 — Coverage (±60 bp)
0 nM protein
3 nM protein
14 nM protein
70 nM protein
350 nM protein
3.0
0.0
−60
−40
−20
0
20
40
60
Position relative to center (bp)

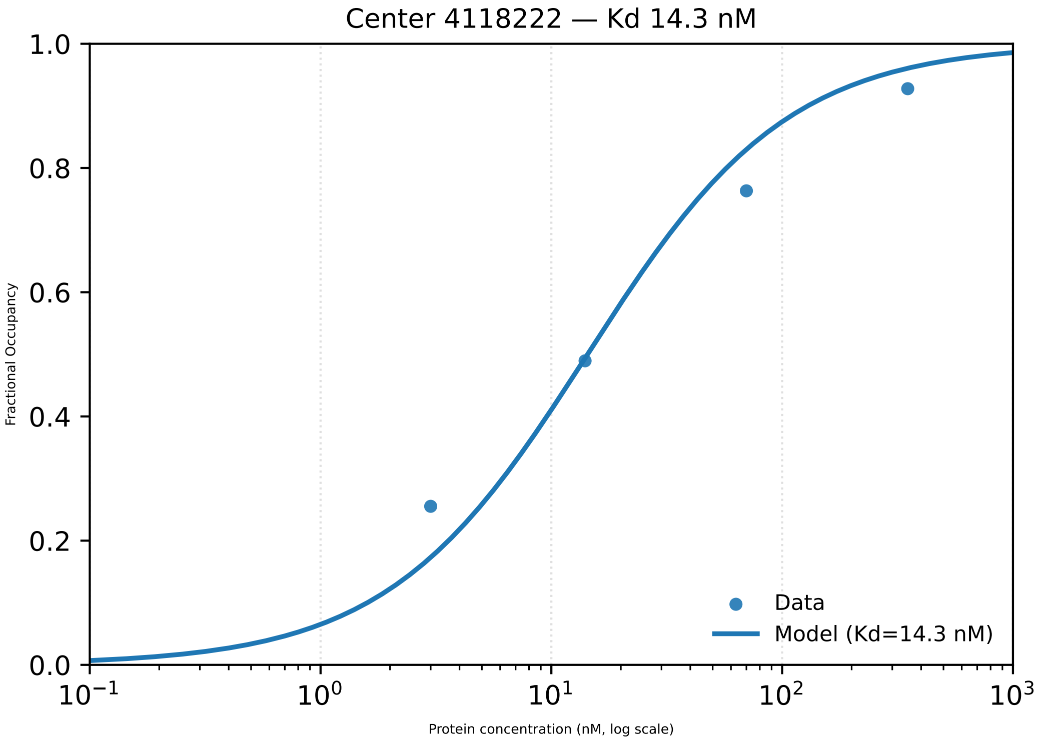

Center 4118222 — Kd 14.3 nM
1.0
0.8
0.6
0.4
0.2
0.0
Fractional Occupancy
$10^{-1}$
$10^{0}$
$10^{1}$
$10^{2}$
$10^{3}$
Protein concentration (nM, log scale)
Data
Model (Kd=14.3 nM)

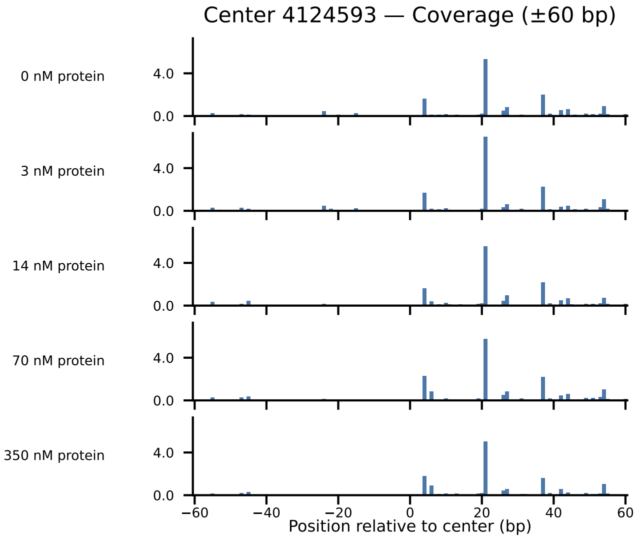
Center 4124593 — Coverage (±60 bp)
0 nM protein
3 nM protein
14 nM protein
70 nM protein
350 nM protein
4.0
0.0
−60
−40
−20
0
20
40
60
Position relative to center (bp)

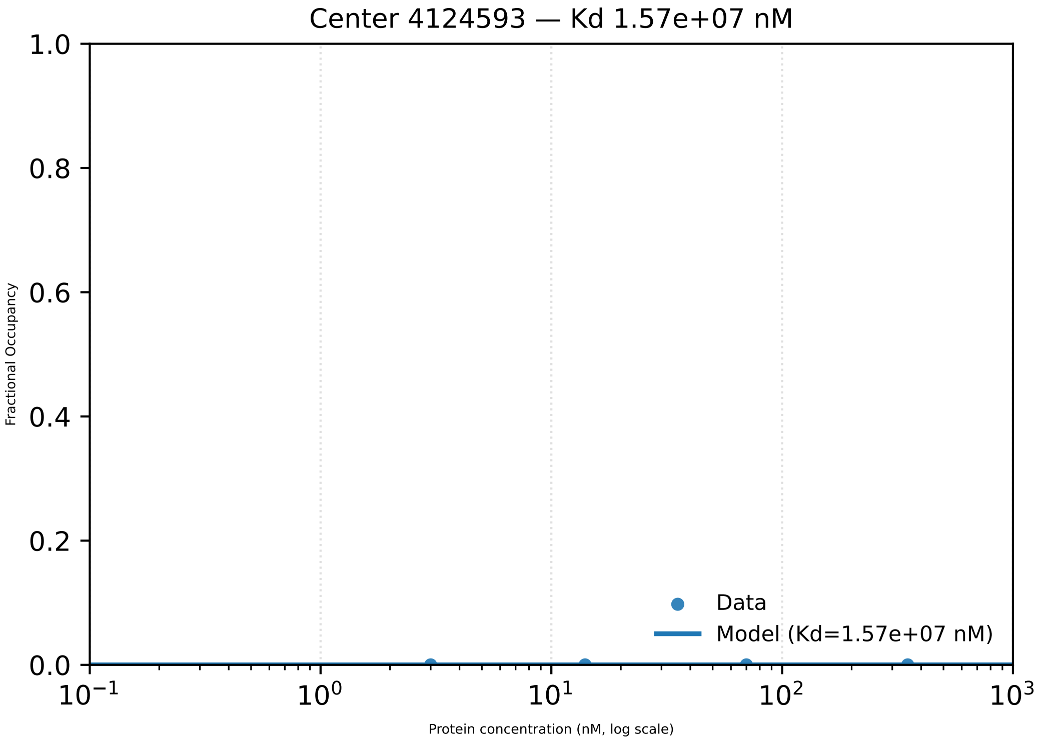
Center 4124593 — Kd 1.57e+07 nM
Fractional Occupancy
1.0
0.8
0.6
0.4
0.2
0.0
Data
Model (Kd=1.57e+07 nM)
$10^{-1}$
$10^{0}$
$10^{1}$
$10^{2}$
$10^{3}$
Protein concentration (nM, log scale)

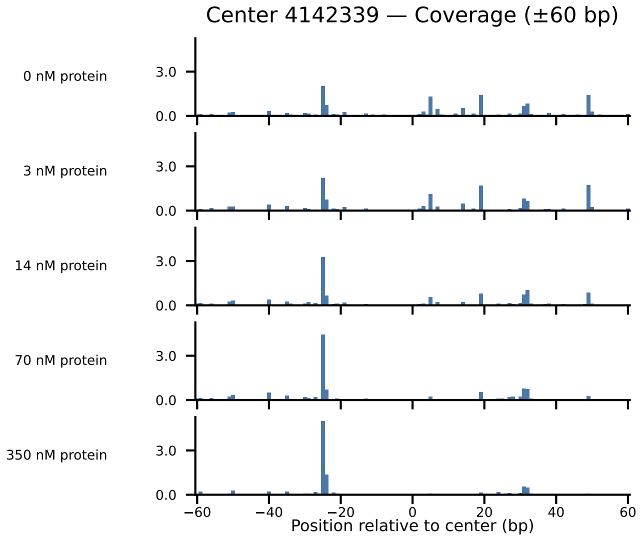

Center 4142339 — Coverage (±60 bp)
0 nM protein
3 nM protein
14 nM protein
70 nM protein
350 nM protein
3.0
0.0
−60
−40
−20
0
20
40
60
Position relative to center (bp)

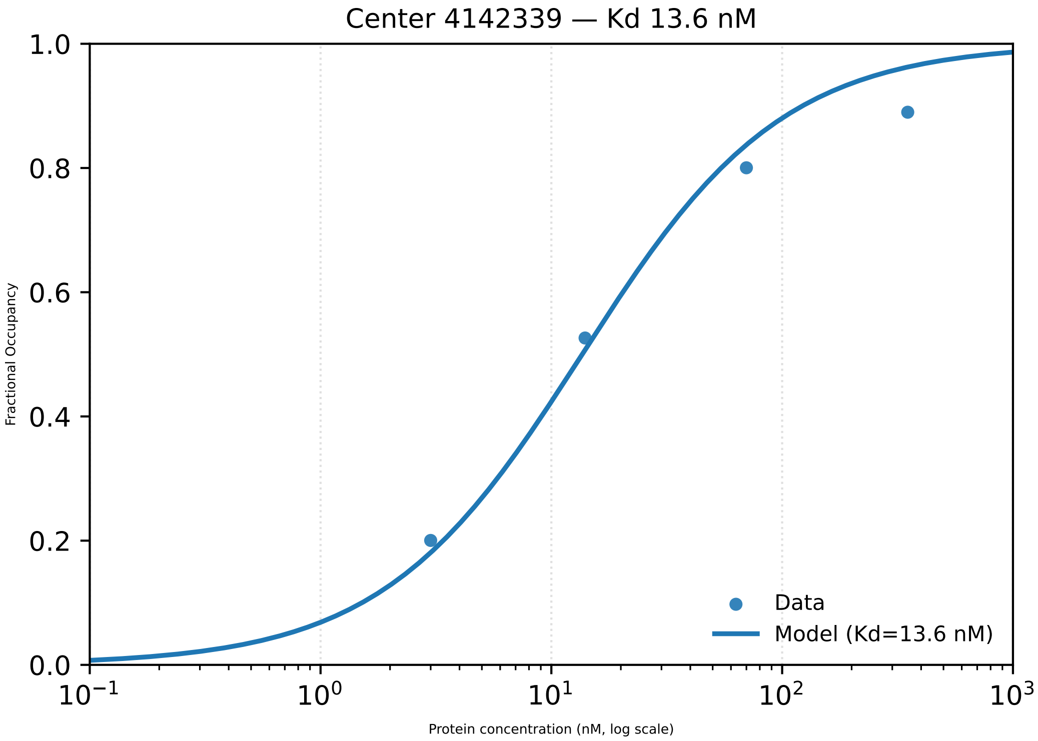

Center 4142339 — Kd 13.6 nM
Fractional Occupancy
Protein concentration (nM, log scale)
Data
Model (Kd=13.6 nM)
0.0
0.2
0.4
0.6
0.8
1.0
$10^{-1}$
$10^{0}$
$10^{1}$
$10^{2}$
$10^{3}$

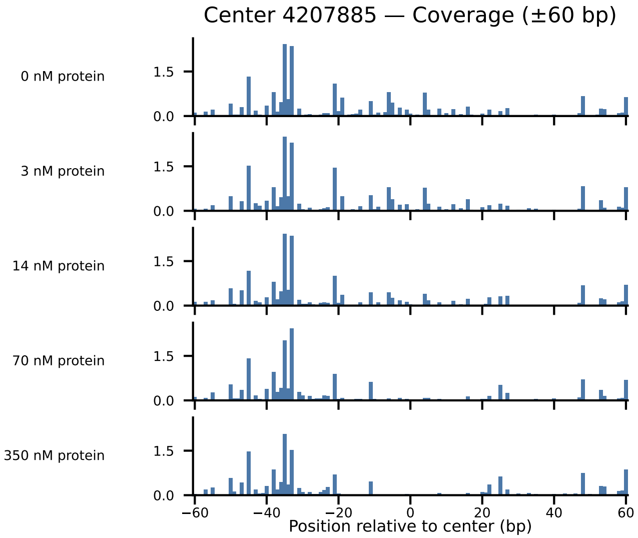

Center 4207885 — Coverage (±60 bp)
0 nM protein
3 nM protein
14 nM protein
70 nM protein
350 nM protein
1.5
0.0
−60
−40
−20
0
20
40
60
Position relative to center (bp)

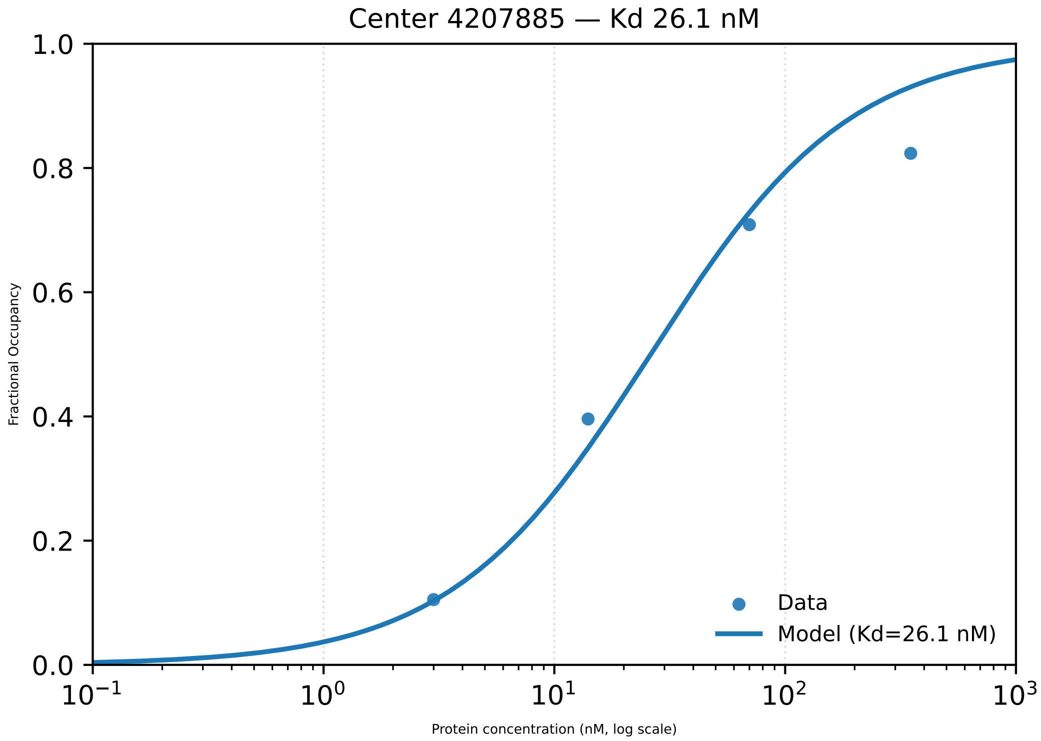
Center 4207885 — Kd 26.1 nM
Fractional Occupancy
1.0
0.8
0.6
0.4
0.2
0.0
$10^{-1}$
$10^{0}$
$10^{1}$
$10^{2}$
$10^{3}$
Protein concentration (nM, log scale)
Data
Model (Kd=26.1 nM)

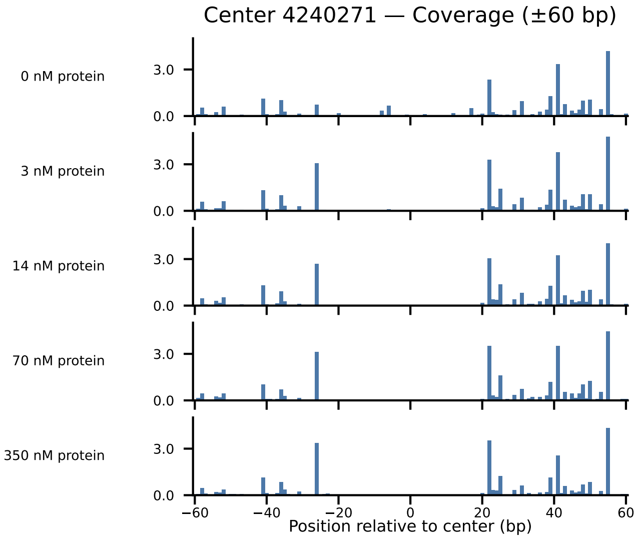

Center 4240271 — Coverage (±60 bp)
0 nM protein
3 nM protein
14 nM protein
70 nM protein
350 nM protein
3.0
0.0
−60
−40
−20
0
20
40
60
Position relative to center (bp)

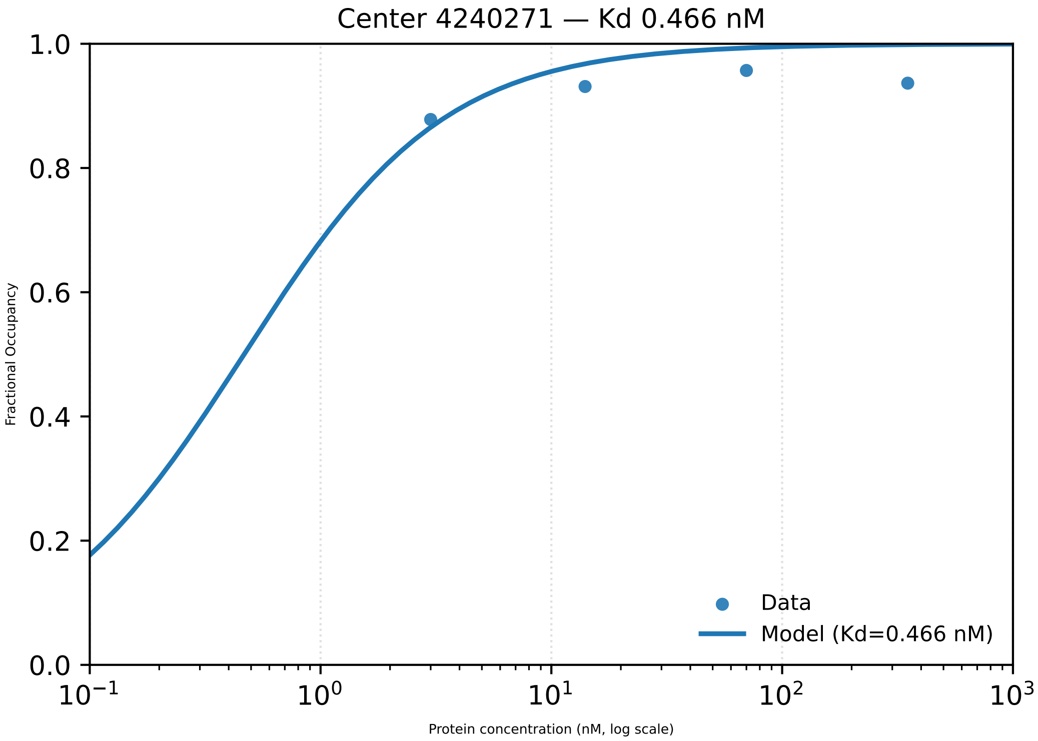

Center 4240271 — Kd 0.466 nM
Fractional Occupancy
1.0
0.8
0.6
0.4
0.2
0.0
$10^{-1}$
$10^{0}$
$10^{1}$
$10^{2}$
$10^{3}$
Protein concentration (nM, log scale)
Data
Model (Kd=0.466 nM)

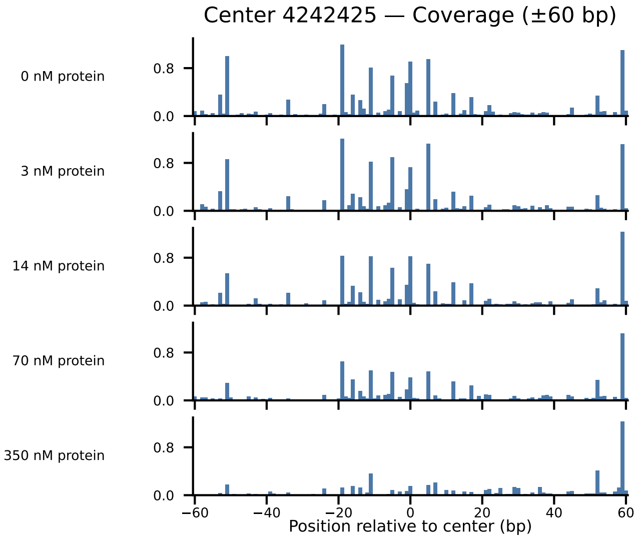
Center 4242425 — Coverage (±60 bp)
0 nM protein
3 nM protein
14 nM protein
70 nM protein
350 nM protein
0.8
0.0
−60
−40
−20
0
20
40
60
Position relative to center (bp)

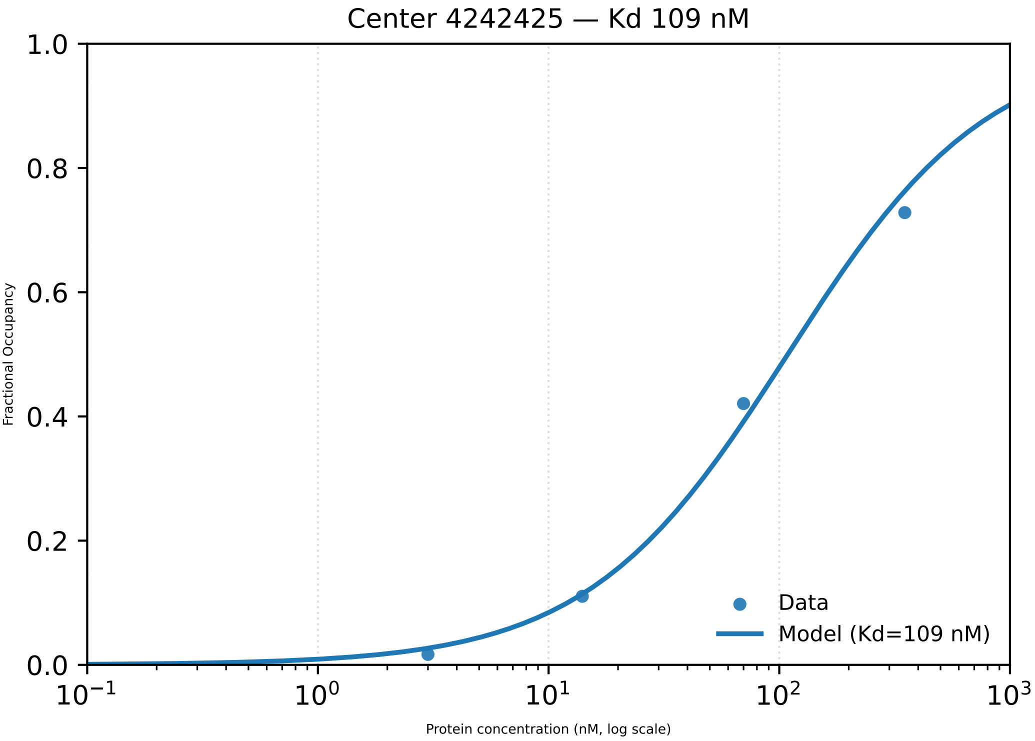

Center 4242425 — Kd 109 nM
1.0
0.8
0.6
0.4
0.2
0.0
Fractional Occupancy
Data
Model (Kd=109 nM)
$10^{-1}$
$10^{0}$
$10^{1}$
$10^{2}$
$10^{3}$
Protein concentration (nM, log scale)

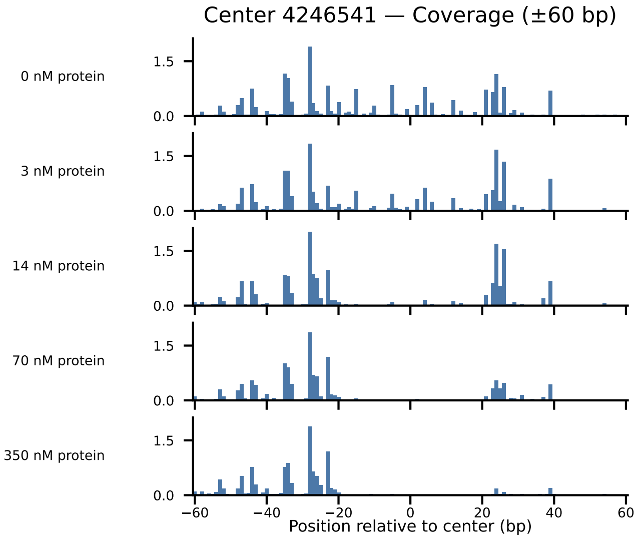
Center 4246541 — Coverage (±60 bp)
0 nM protein
3 nM protein
14 nM protein
70 nM protein
350 nM protein
1.5
0.0
−60
−40
−20
0
20
40
60
Position relative to center (bp)

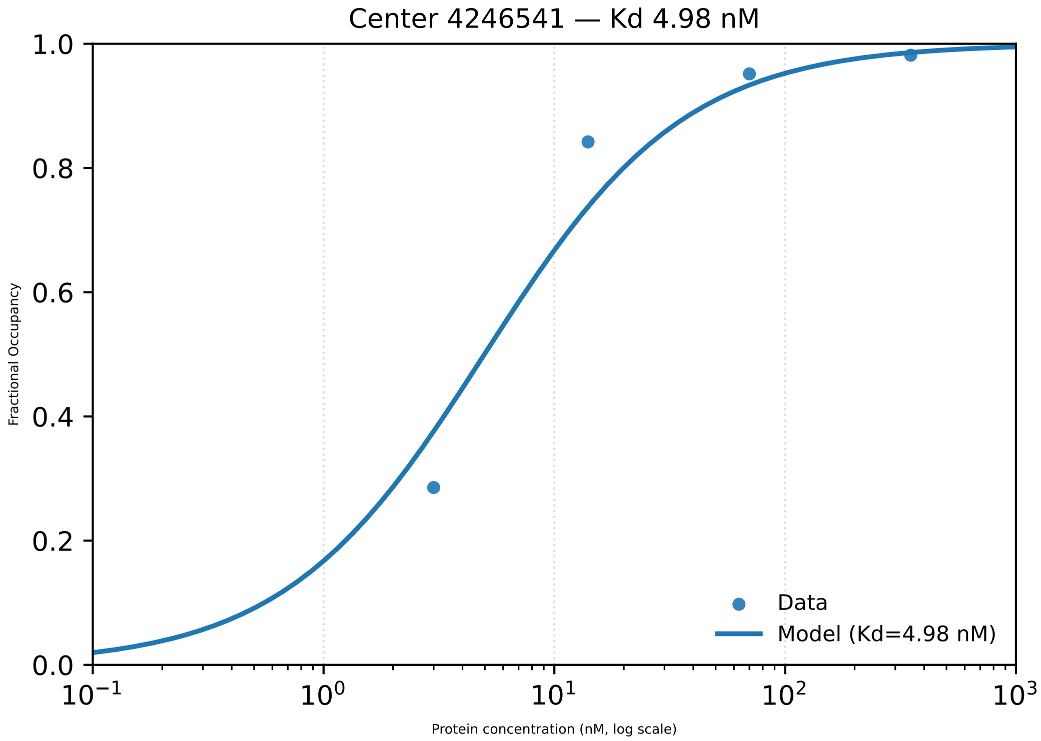

Center 4246541 — Kd 4.98 nM
1.0
0.8
0.6
0.4
0.2
0.0
Fractional Occupancy
$10^{-1}$
$10^{0}$
$10^{1}$
$10^{2}$
$10^{3}$
Protein concentration (nM, log scale)
Data
Model (Kd=4.98 nM)

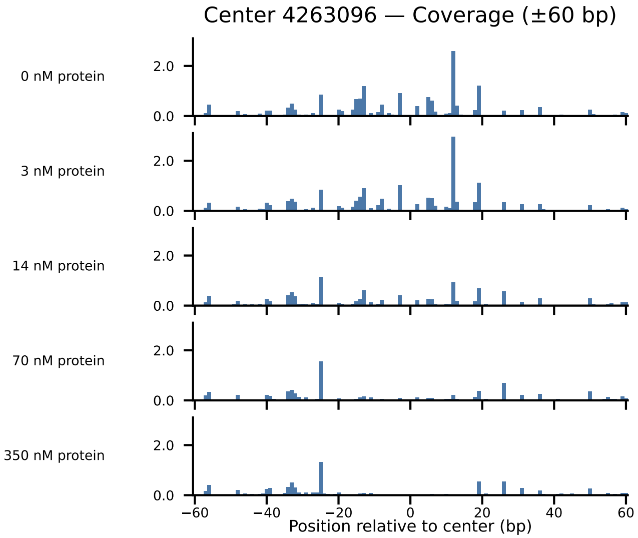
Center 4263096 — Coverage (±60 bp)
0 nM protein
3 nM protein
14 nM protein
70 nM protein
350 nM protein
2.0
0.0
−60
−40
−20
0
20
40
60
Position relative to center (bp)

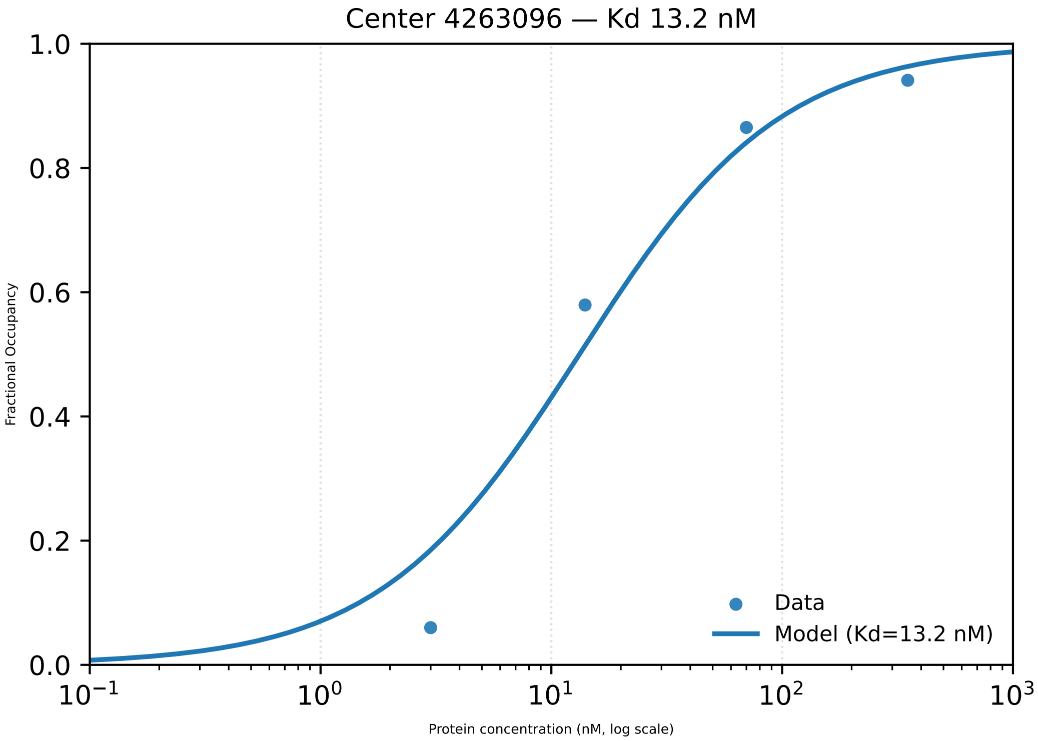

Center 4263096 — Kd 13.2 nM
Fractional Occupancy
Protein concentration (nM, log scale)
0.0
0.2
0.4
0.6
0.8
1.0
$10^{-1}$
$10^{0}$
$10^{1}$
$10^{2}$
$10^{3}$
Data
Model (Kd=13.2 nM)

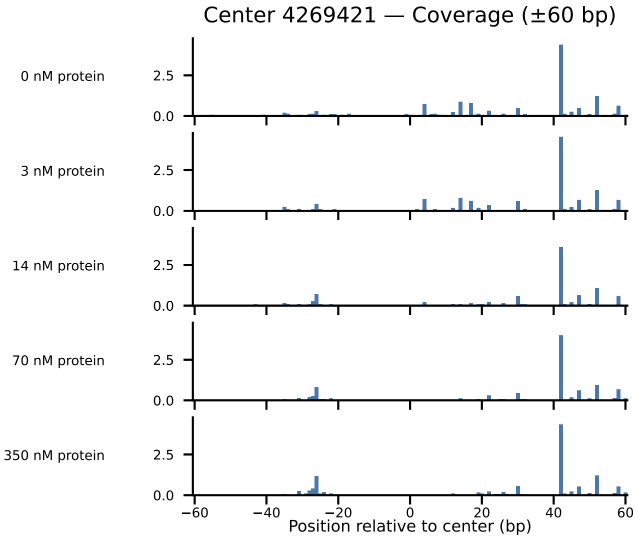

Center 4269421 — Coverage (±60 bp)
0 nM protein
3 nM protein
14 nM protein
70 nM protein
350 nM protein
2.5
0.0
−60
−40
−20
0
20
40
60
Position relative to center (bp)

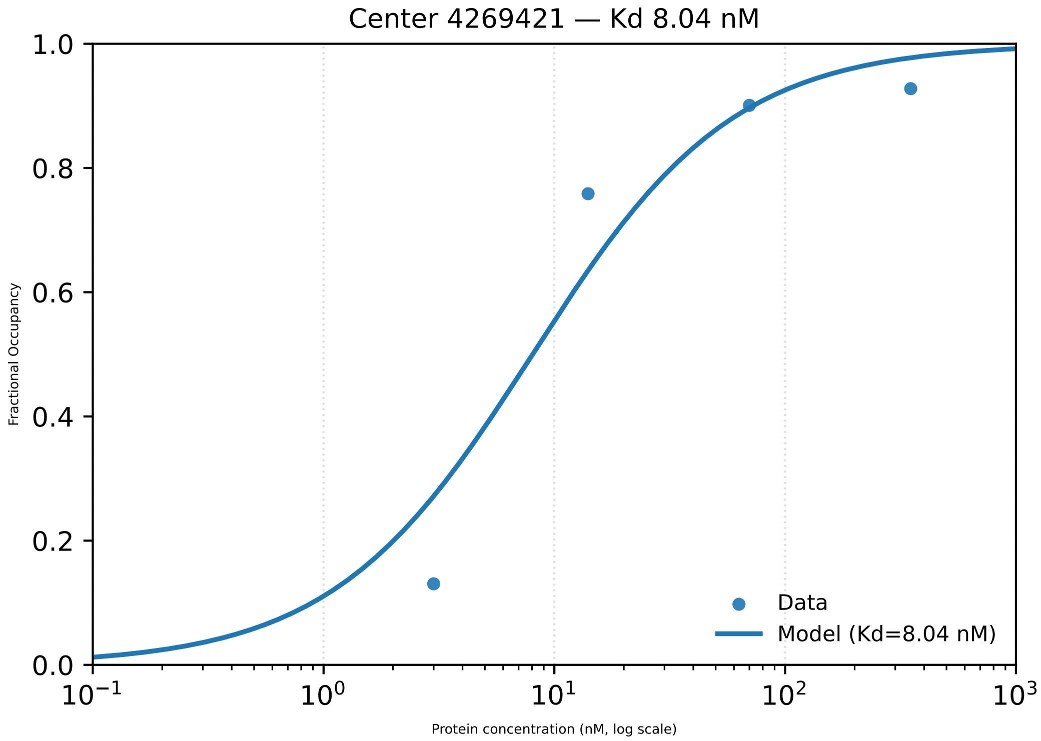

Center 4269421 — Kd 8.04 nM
Fractional Occupancy
Protein concentration (nM, log scale)
Data
Model (Kd=8.04 nM)

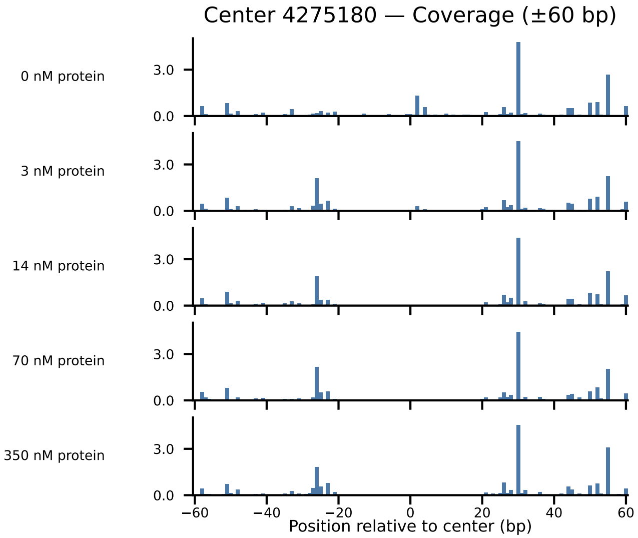

Center 4275180 — Coverage (±60 bp)
0 nM protein
3 nM protein
14 nM protein
70 nM protein
350 nM protein
3.0
0.0
−60
−40
−20
0
20
40
60
Position relative to center (bp)

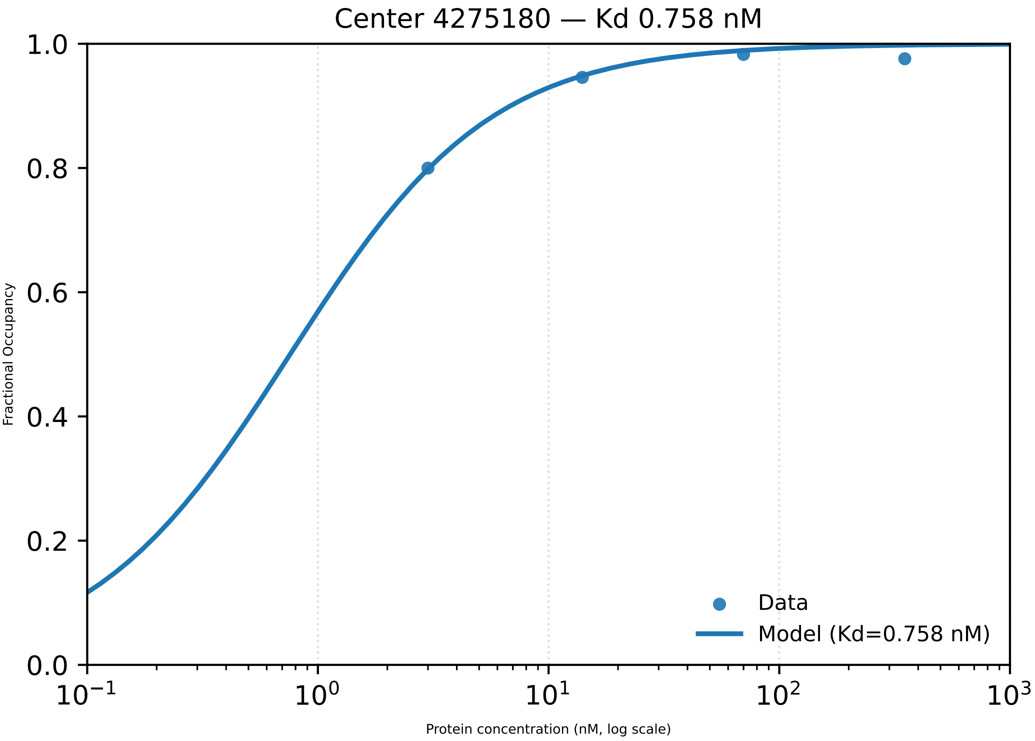

Center 4275180 — Kd 0.758 nM
Fractional Occupancy
1.0
0.8
0.6
0.4
0.2
0.0
$10^{-1}$
$10^{0}$
$10^{1}$
$10^{2}$
$10^{3}$
Protein concentration (nM, log scale)
Data
Model (Kd=0.758 nM)

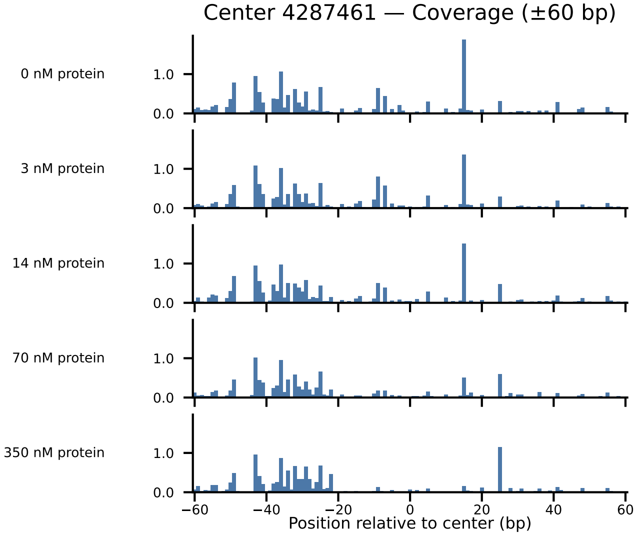

Center 4287461 — Coverage (±60 bp)
0 nM protein
3 nM protein
14 nM protein
70 nM protein
350 nM protein
1.0
0.0
−60
−40
−20
0
20
40
60
Position relative to center (bp)

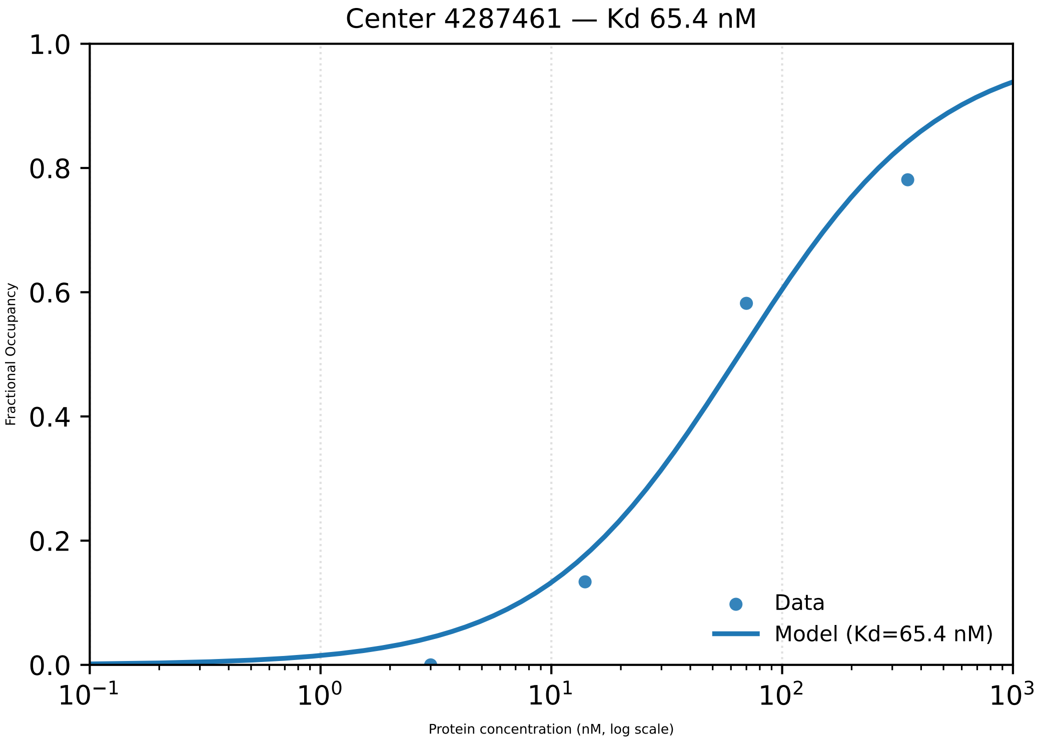

Center 4287461 — Kd 65.4 nM
Fractional Occupancy
1.0
0.8
0.6
0.4
0.2
0.0
$10^{-1}$
$10^{0}$
$10^{1}$
$10^{2}$
$10^{3}$
Protein concentration (nM, log scale)
Data
Model (Kd=65.4 nM)

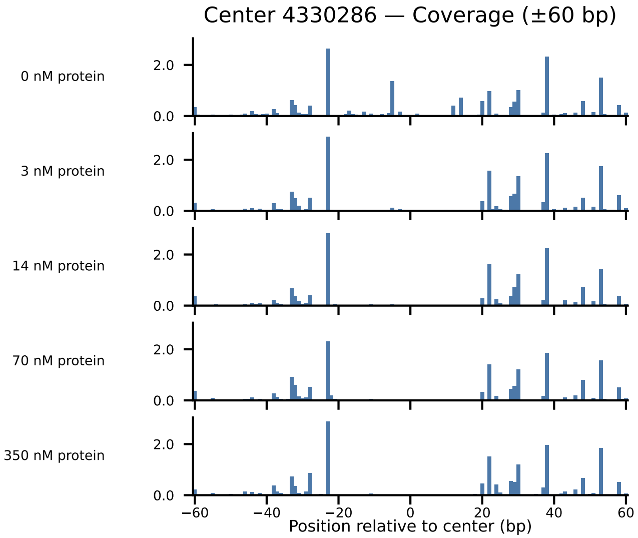

Center 4330286 — Coverage (±60 bp)
0 nM protein
3 nM protein
14 nM protein
70 nM protein
350 nM protein
2.0
0.0
−60
−40
−20
0
20
40
60
Position relative to center (bp)

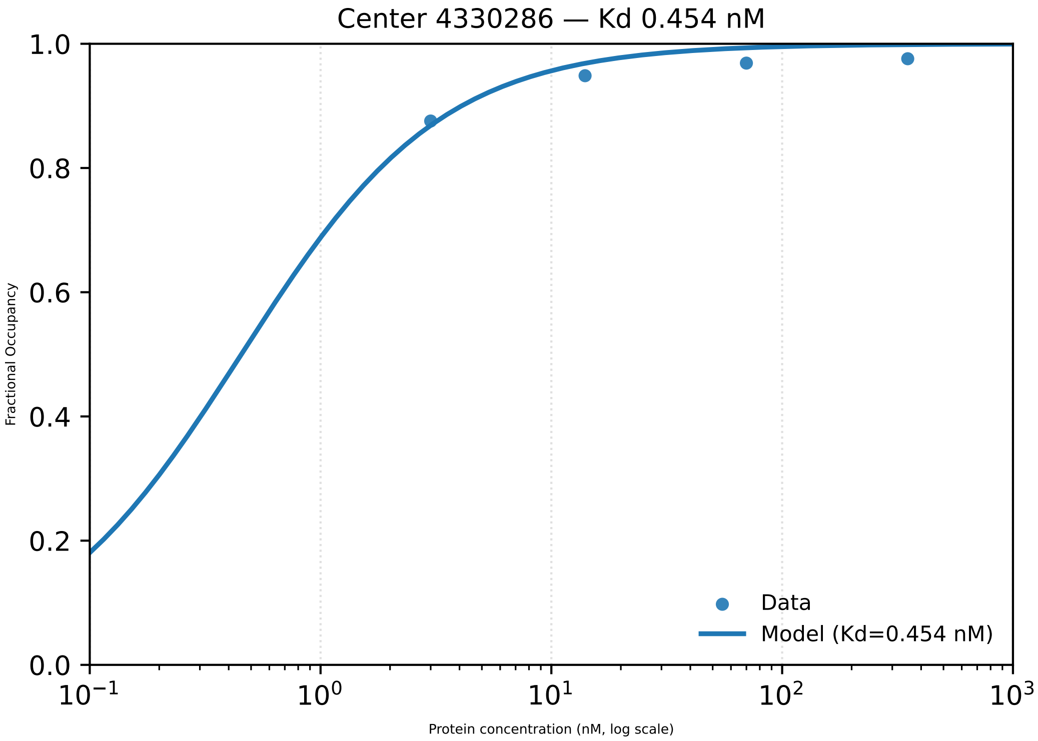

Center 4330286 — Kd 0.454 nM
Fractional Occupancy
Protein concentration (nM, log scale)
1.0
0.8
0.6
0.4
0.2
0.0
$10^{-1}$
$10^{0}$
$10^{1}$
$10^{2}$
$10^{3}$
Data
Model (Kd=0.454 nM)

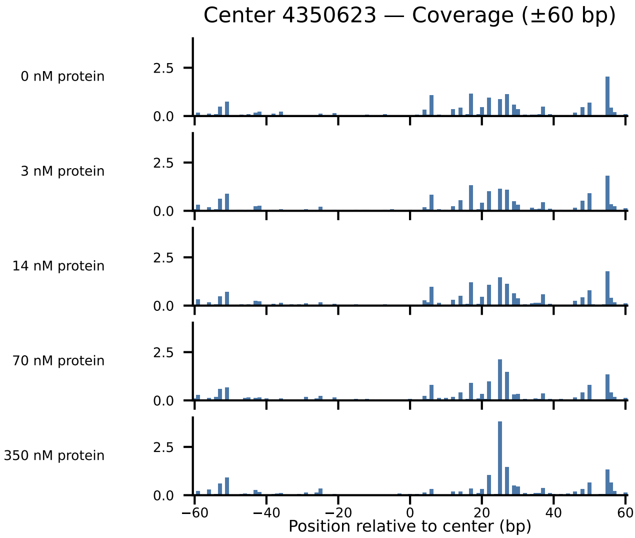

Center 4350623 — Coverage (±60 bp)
0 nM protein
3 nM protein
14 nM protein
70 nM protein
350 nM protein
2.5
0.0
−60
−40
−20
0
20
40
60
Position relative to center (bp)

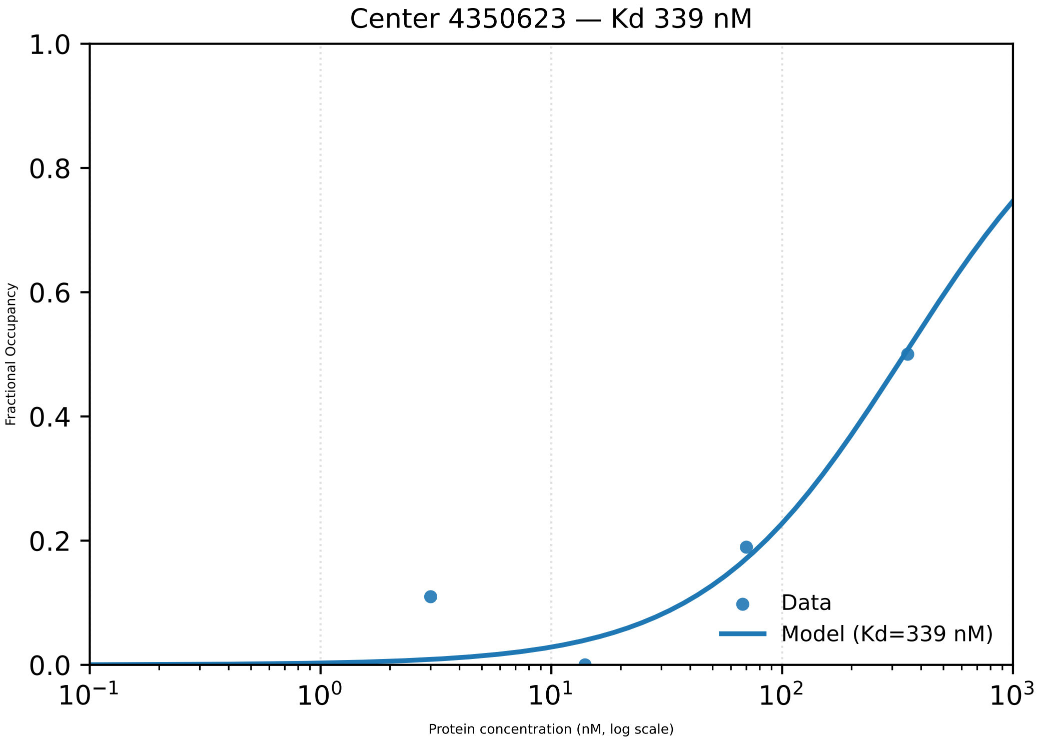
Center 4350623 — Kd 339 nM
1.0
0.8
0.6
0.4
0.2
0.0
Fractional Occupancy
$10^{-1}$
$10^{0}$
$10^{1}$
$10^{2}$
$10^{3}$
Protein concentration (nM, log scale)
Data
Model (Kd=339 nM)

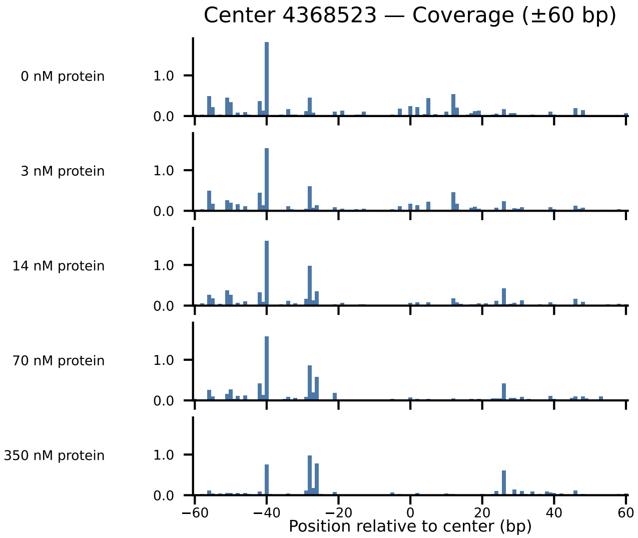

Center 4368523 — Coverage (±60 bp)
0 nM protein
3 nM protein
14 nM protein
70 nM protein
350 nM protein
1.0
0.0
−60
−40
−20
0
20
40
60
Position relative to center (bp)

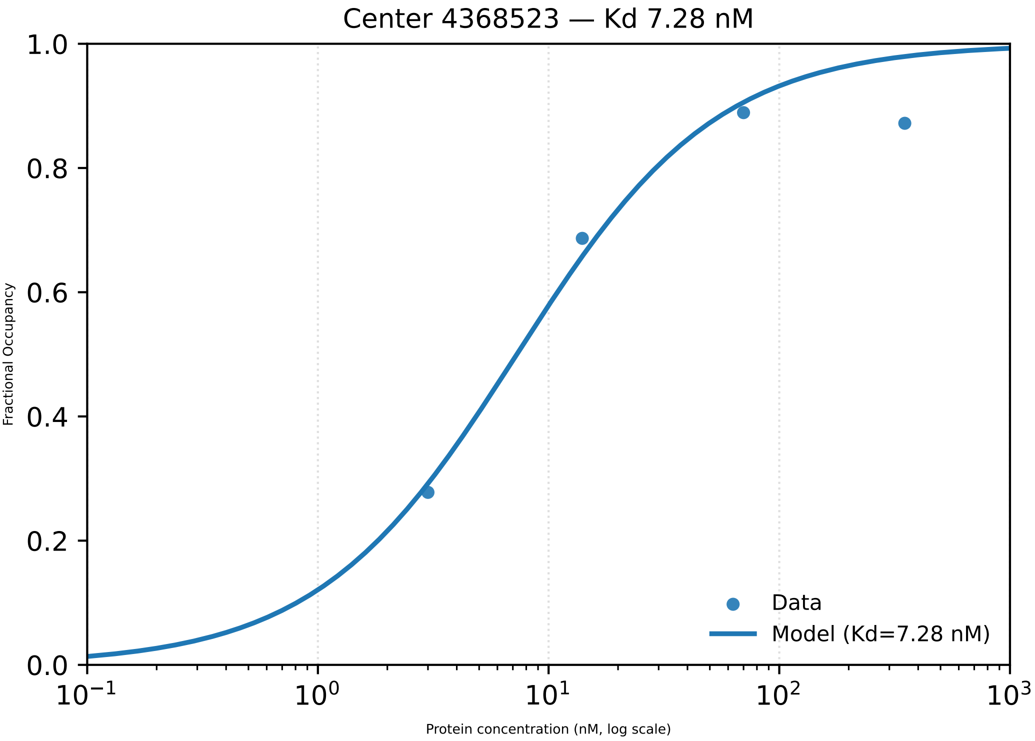

Center 4368523 — Kd 7.28 nM
1.0
0.8
0.6
0.4
0.2
0.0
Fractional Occupancy
10$^{-1}$
10$^{0}$
10$^{1}$
10$^{2}$
10$^{3}$
Protein concentration (nM, log scale)
Data
Model (Kd=7.28 nM)

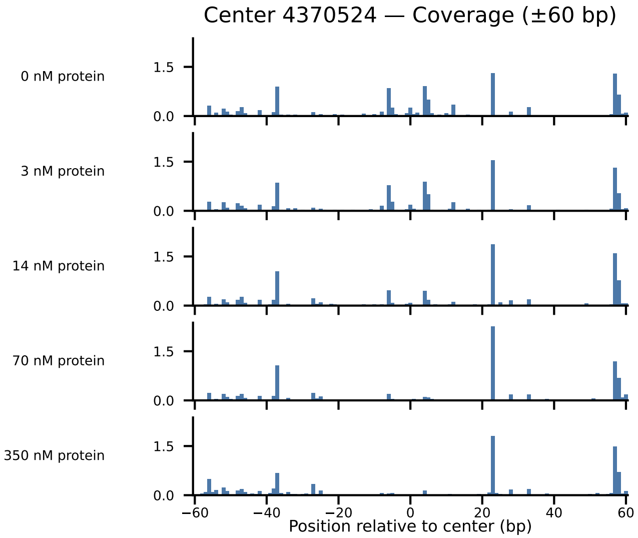
Center 4370524 — Coverage (±60 bp)
0 nM protein
3 nM protein
14 nM protein
70 nM protein
350 nM protein
1.5
0.0
−60
−40
−20
0
20
40
60
Position relative to center (bp)

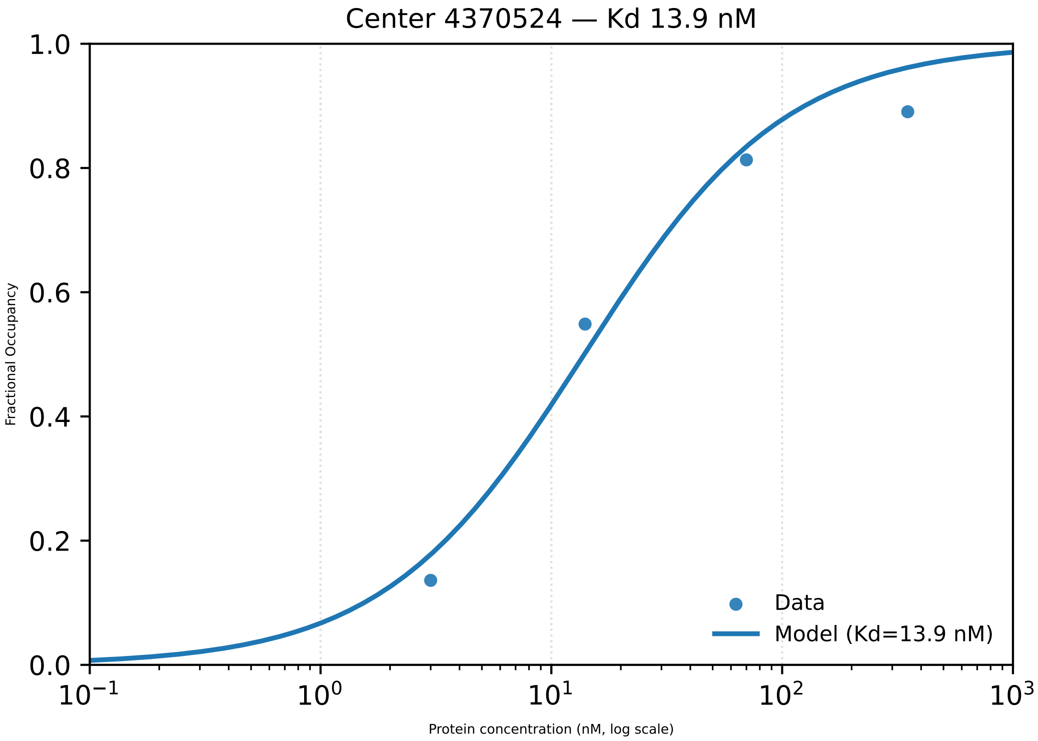

Center 4370524 — Kd 13.9 nM
Fractional Occupancy
Protein concentration (nM, log scale)
Data
Model (Kd=13.9 nM)
1.0
0.8
0.6
0.4
0.2
0.0
$10^{-1}$
$10^{0}$
$10^{1}$
$10^{2}$
$10^{3}$

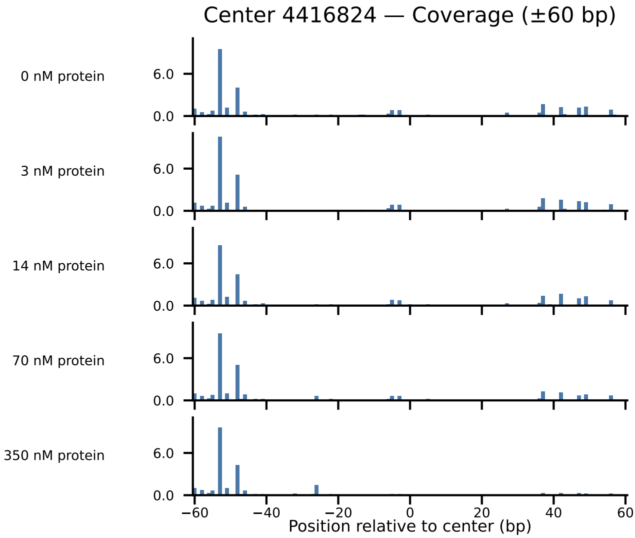

Center 4416824 — Coverage (±60 bp)
0 nM protein
3 nM protein
14 nM protein
70 nM protein
350 nM protein
6.0
0.0
−60
−40
−20
0
20
40
60
Position relative to center (bp)

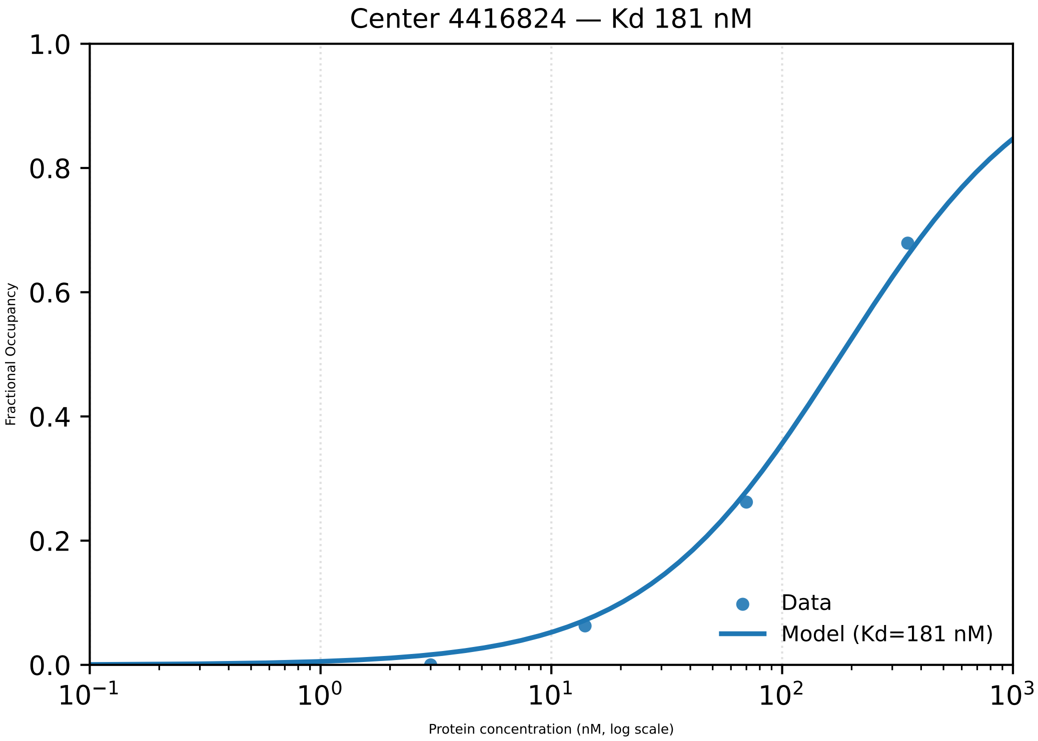

Center 4416824 — Kd 181 nM
1.0
0.8
0.6
0.4
0.2
0.0
Fractional Occupancy
$10^{-1}$
$10^{0}$
$10^{1}$
$10^{2}$
$10^{3}$
Protein concentration (nM, log scale)
Data
Model (Kd=181 nM)

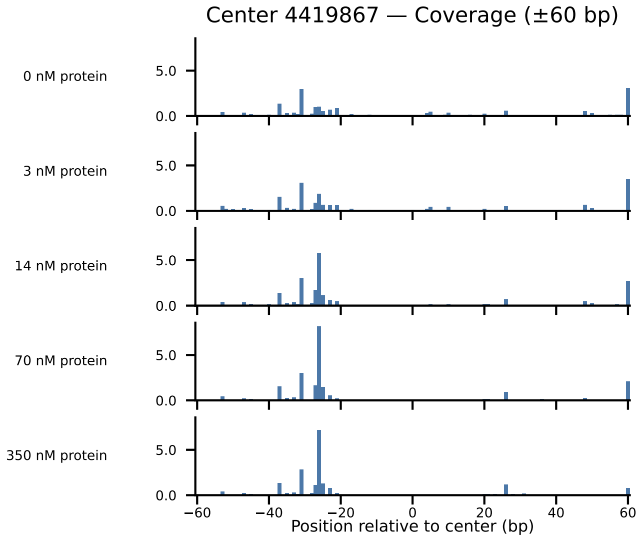
Center 4419867 — Coverage (±60 bp)
0 nM protein
3 nM protein
14 nM protein
70 nM protein
350 nM protein
5.0
0.0
−60
−40
−20
0
20
40
60
Position relative to center (bp)

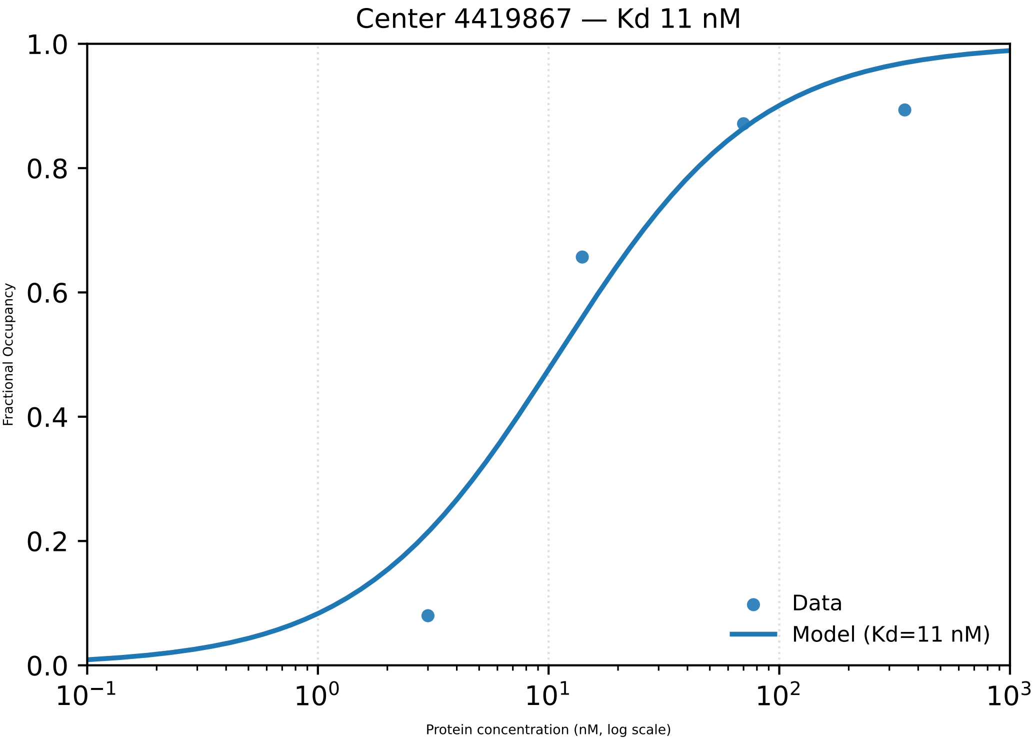

Center 4419867 — Kd 11 nM
1.0
0.8
0.6
0.4
0.2
0.0
Fractional Occupancy
$10^{-1}$
$10^{0}$
$10^{1}$
$10^{2}$
$10^{3}$
Protein concentration (nM, log scale)
Data
Model (Kd=11 nM)

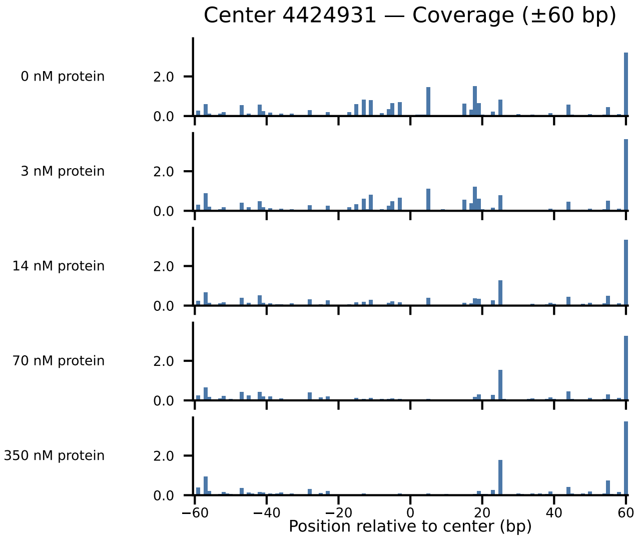
Center 4424931 — Coverage (±60 bp)
0 nM protein
3 nM protein
14 nM protein
70 nM protein
350 nM protein
2.0
0.0
−60
−40
−20
0
20
40
60
Position relative to center (bp)

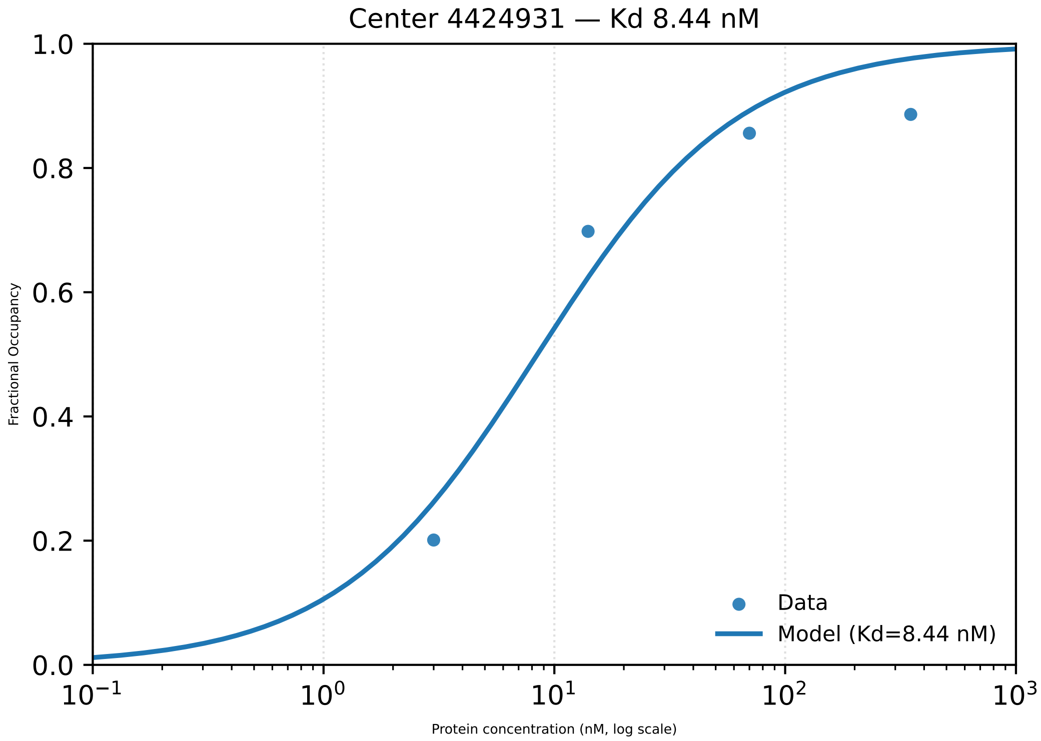

Center 4424931 — Kd 8.44 nM
1.0
0.8
0.6
0.4
0.2
0.0
Fractional Occupancy
Data
Model (Kd=8.44 nM)
$10^{-1}$
$10^{0}$
$10^{1}$
$10^{2}$
$10^{3}$
Protein concentration (nM, log scale)

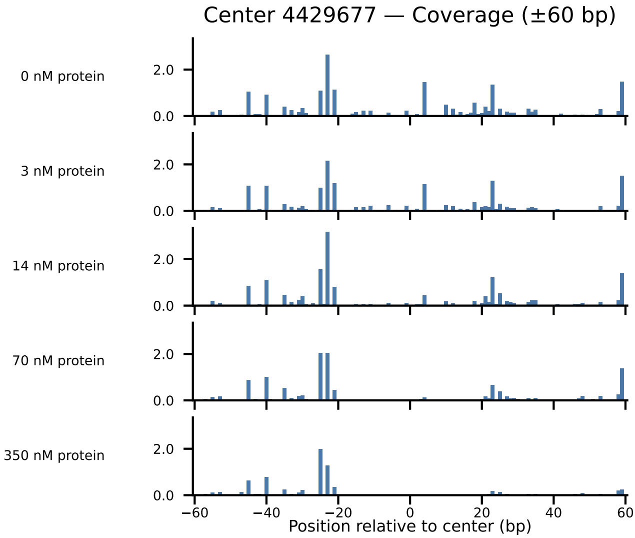

Center 4429677 — Coverage (±60 bp)
0 nM protein
3 nM protein
14 nM protein
70 nM protein
350 nM protein
2.0
0.0
−60
−40
−20
0
20
40
60
Position relative to center (bp)

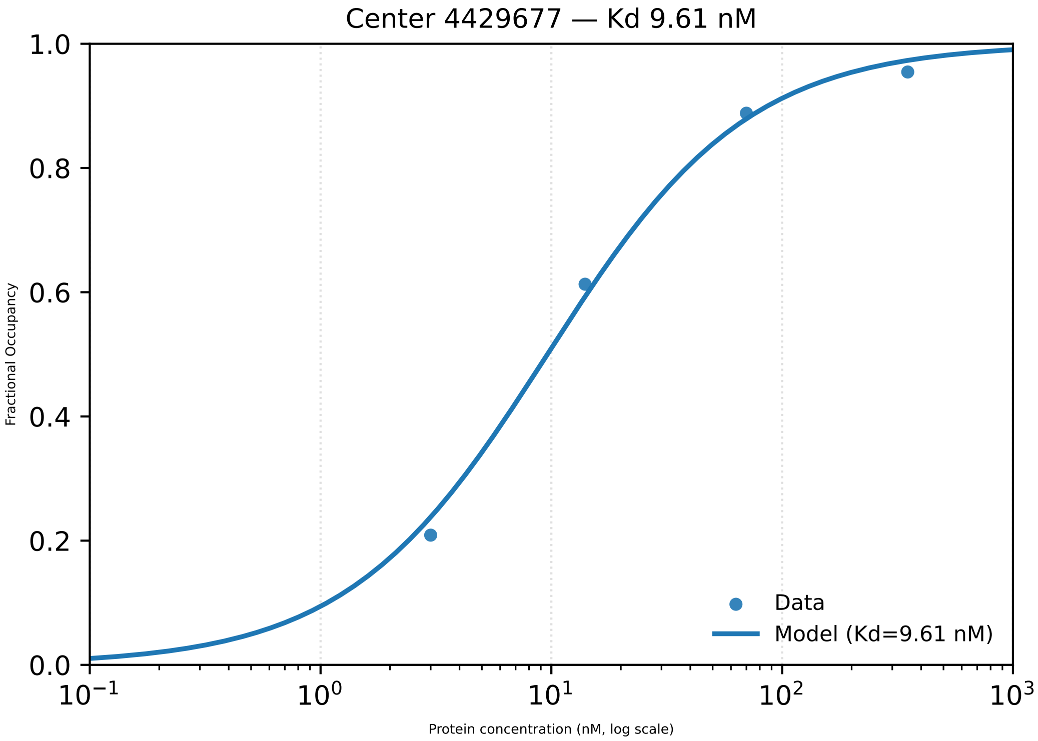

Center 4429677 — Kd 9.61 nM
1.0
0.8
0.6
0.4
0.2
0.0
Fractional Occupancy
Data
Model (Kd=9.61 nM)
$10^{-1}$
$10^{0}$
$10^{1}$
$10^{2}$
$10^{3}$
Protein concentration (nM, log scale)

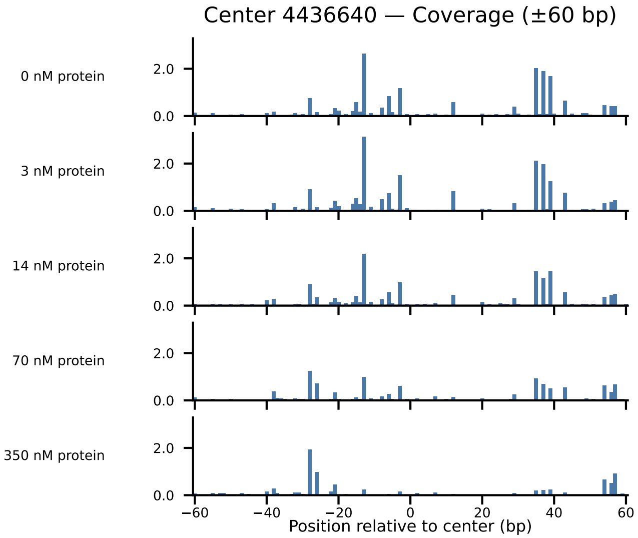

Center 4436640 — Coverage (±60 bp)
0 nM protein
3 nM protein
14 nM protein
70 nM protein
350 nM protein
2.0
0.0
−60
−40
−20
0
20
40
60
Position relative to center (bp)

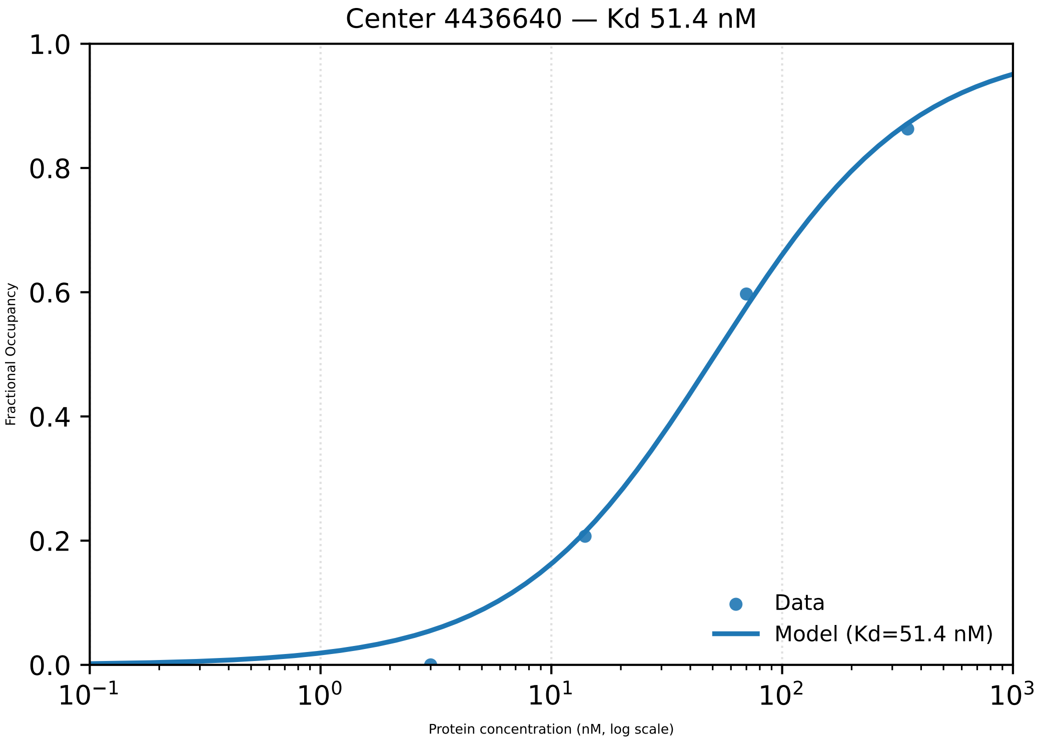

Center 4436640 — Kd 51.4 nM
1.0
0.8
0.6
0.4
0.2
0.0
Fractional Occupancy
$10^{-1}$
$10^{0}$
$10^{1}$
$10^{2}$
$10^{3}$
Protein concentration (nM, log scale)
Data
Model (Kd=51.4 nM)

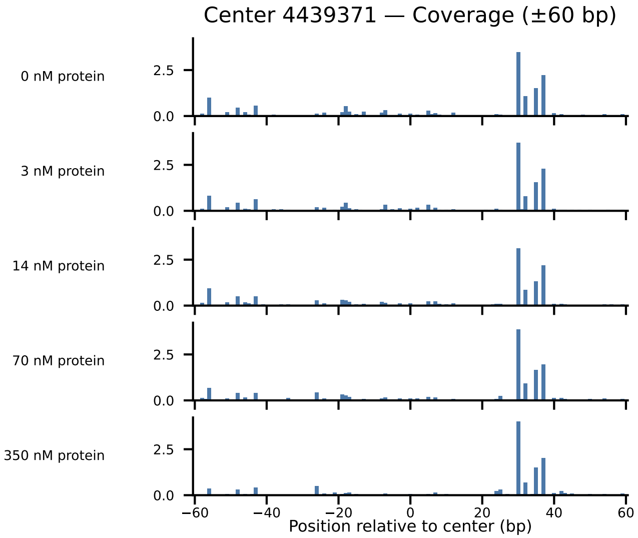
Center 4439371 — Coverage (±60 bp)
0 nM protein
3 nM protein
14 nM protein
70 nM protein
350 nM protein
2.5
0.0
−60
−40
−20
0
20
40
60
Position relative to center (bp)

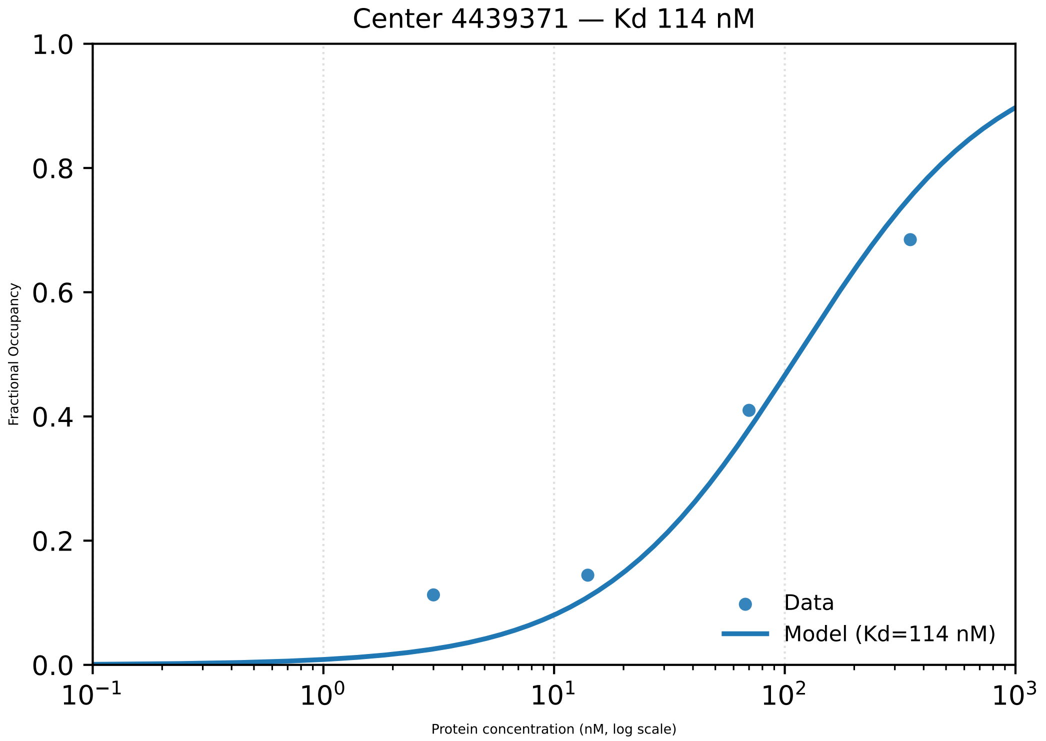

Center 4439371 — Kd 114 nM
1.0
0.8
0.6
0.4
0.2
0.0
Fractional Occupancy
Data
Model (Kd=114 nM)
$10^{-1}$
$10^{0}$
$10^{1}$
$10^{2}$
$10^{3}$
Protein concentration (nM, log scale)

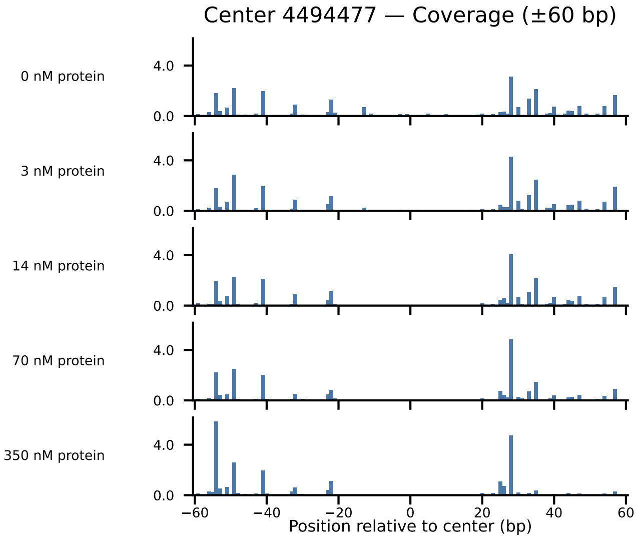
Center 4494477 — Coverage (±60 bp)
0 nM protein
3 nM protein
14 nM protein
70 nM protein
350 nM protein
4.0
0.0
−60
−40
−20
0
20
40
60
Position relative to center (bp)

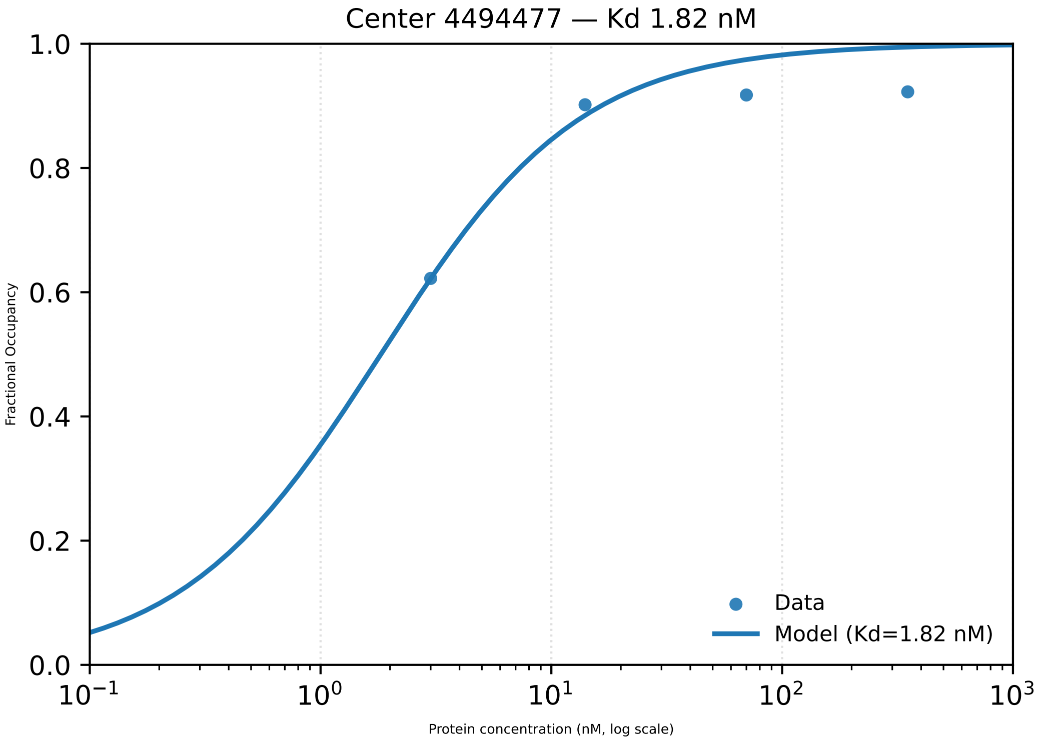
Center 4494477 — Kd 1.82 nM
Fractional Occupancy
Protein concentration (nM, log scale)
Data
Model (Kd=1.82 nM)
1.0
0.8
0.6
0.4
0.2
0.0
$10^{-1}$
$10^{0}$
$10^{1}$
$10^{2}$
$10^{3}$

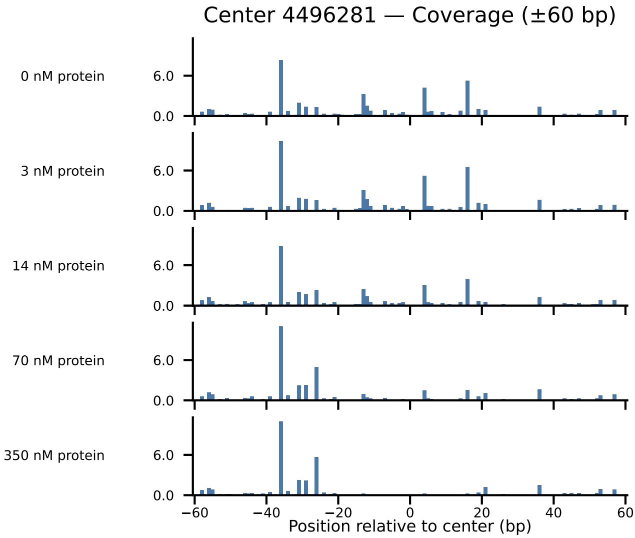
Center 4496281 — Coverage (±60 bp)
0 nM protein
3 nM protein
14 nM protein
70 nM protein
350 nM protein
6.0
0.0
−60
−40
−20
0
20
40
60
Position relative to center (bp)

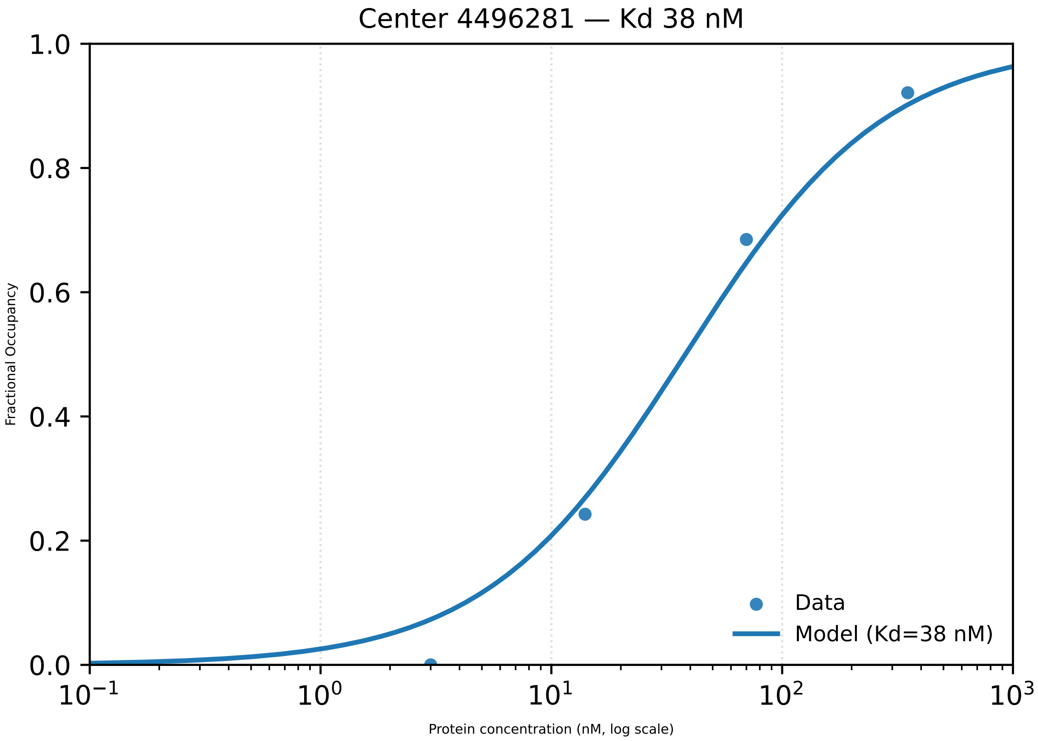

Center 4496281 — Kd 38 nM
Fractional Occupancy
1.0
0.8
0.6
0.4
0.2
0.0
$10^{-1}$
$10^{0}$
$10^{1}$
$10^{2}$
$10^{3}$
Protein concentration (nM, log scale)
Data
Model (Kd=38 nM)

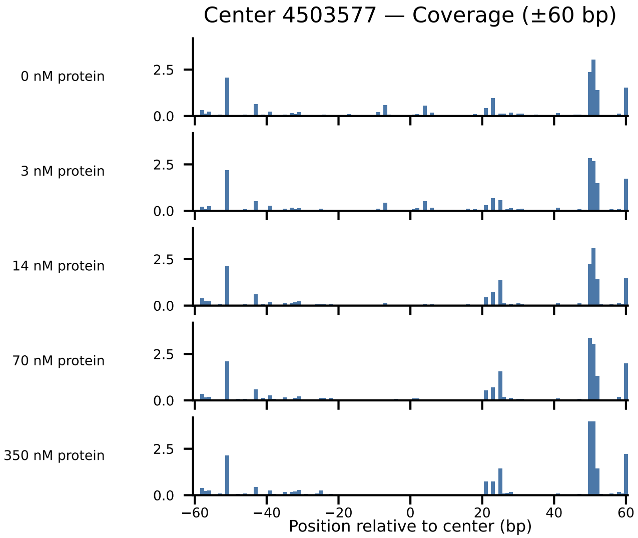

Center 4503577 — Coverage (±60 bp)
0 nM protein
3 nM protein
14 nM protein
70 nM protein
350 nM protein
2.5
0.0
−60
−40
−20
0
20
40
60
Position relative to center (bp)

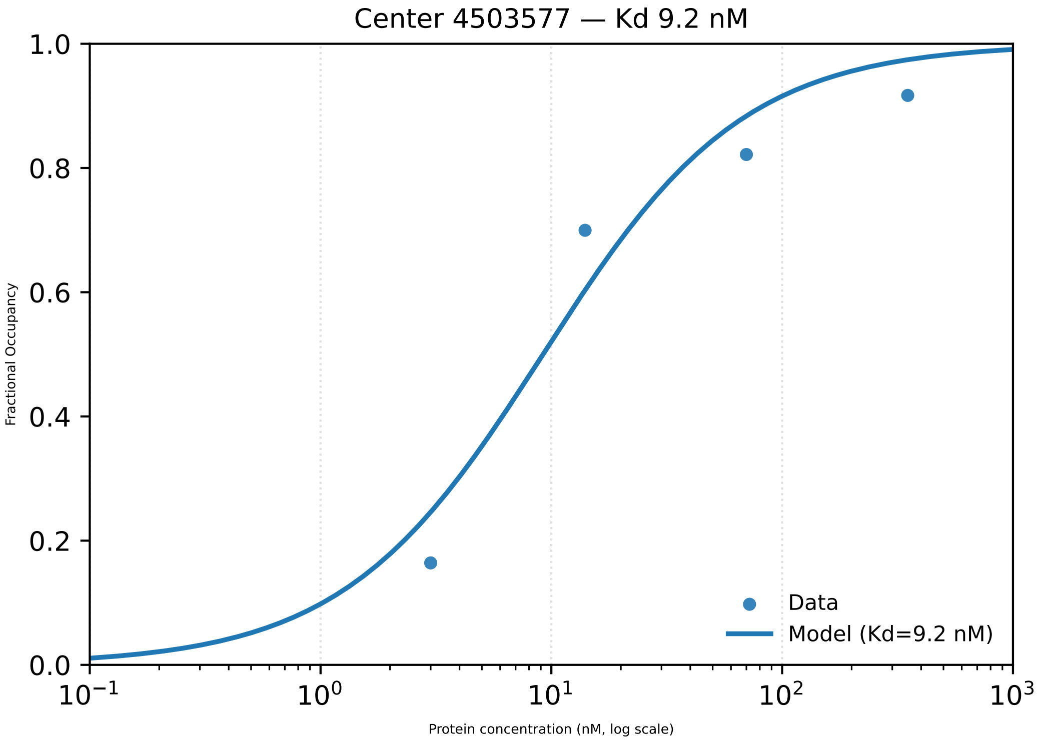

Center 4503577 — Kd 9.2 nM
1.0
0.8
0.6
0.4
0.2
0.0
Fractional Occupancy
$10^{-1}$
$10^{0}$
$10^{1}$
$10^{2}$
$10^{3}$
Protein concentration (nM, log scale)
Data
Model (Kd=9.2 nM)

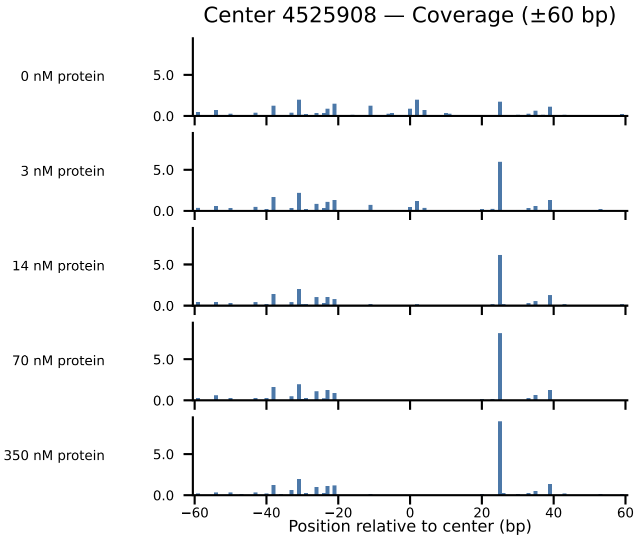

Center 4525908 — Coverage (±60 bp)
0 nM protein
3 nM protein
14 nM protein
70 nM protein
350 nM protein
5.0
0.0
−60
−40
−20
0
20
40
60
Position relative to center (bp)

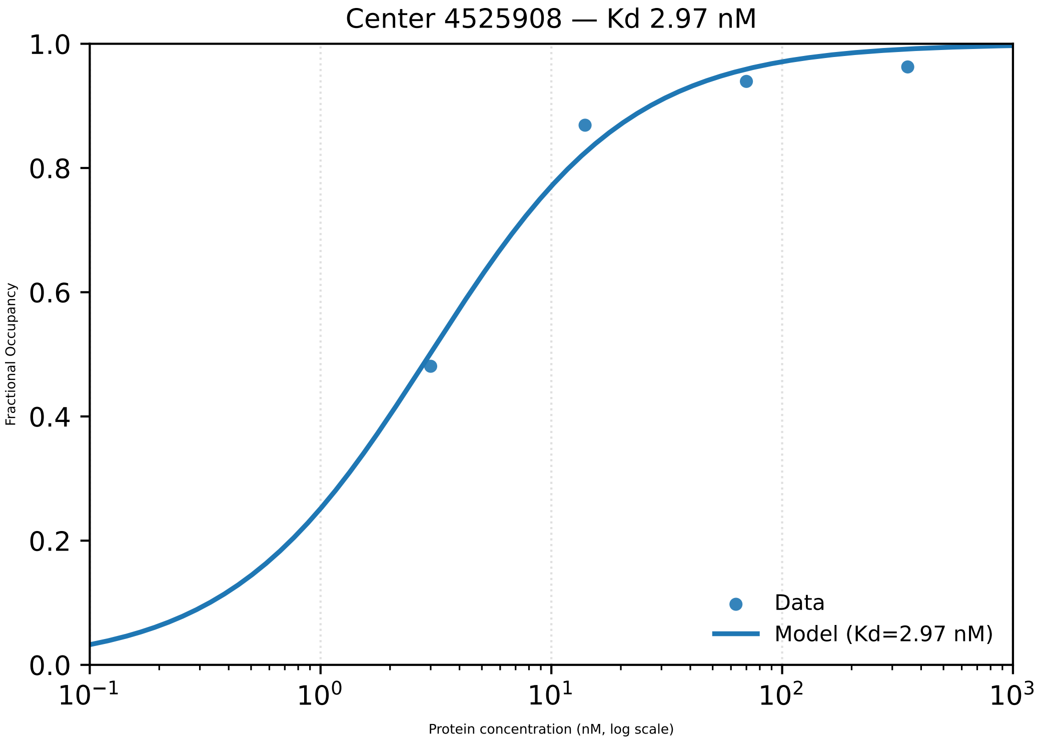

Center 4525908 — Kd 2.97 nM
1.0
0.8
0.6
0.4
0.2
0.0
Fractional Occupancy
$10^{-1}$
$10^{0}$
$10^{1}$
$10^{2}$
$10^{3}$
Protein concentration (nM, log scale)
Data
Model (Kd=2.97 nM)

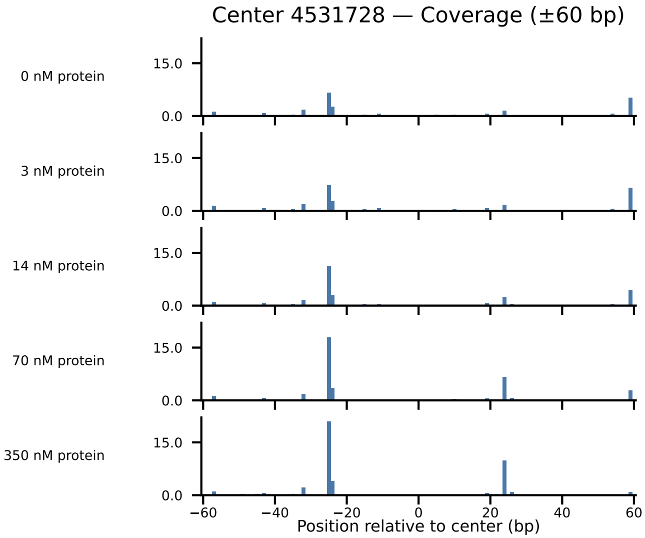

Center 4531728 — Coverage (±60 bp)
0 nM protein
3 nM protein
14 nM protein
70 nM protein
350 nM protein
15.0
0.0
−60
−40
−20
0
20
40
60
Position relative to center (bp)

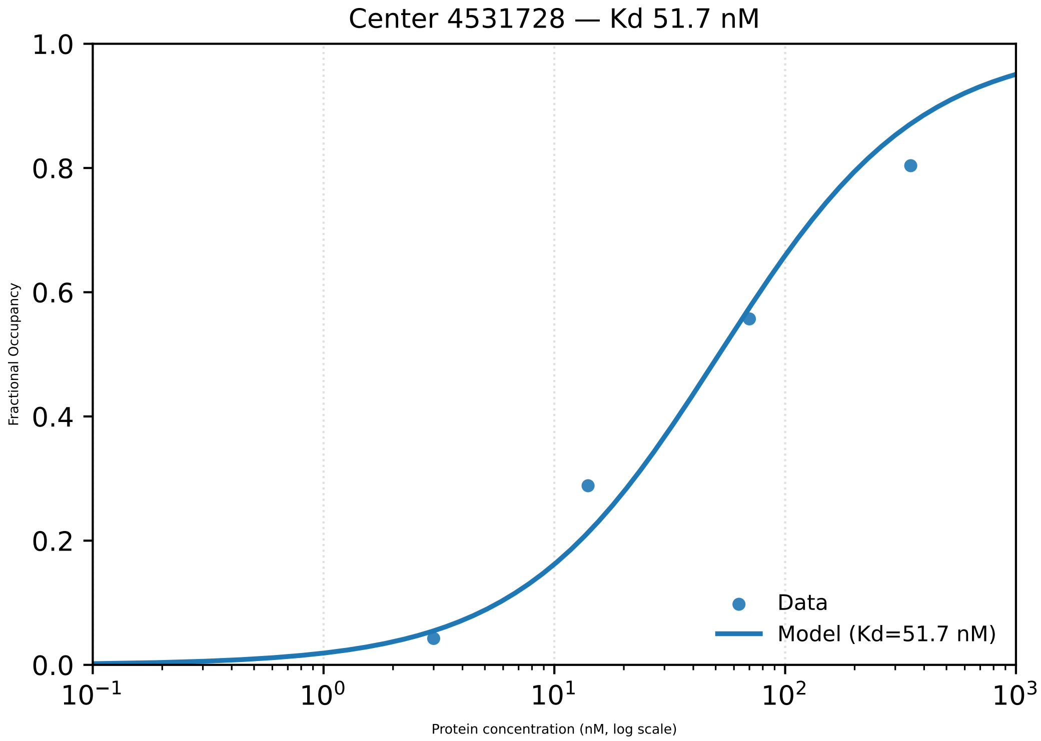

Center 4531728 — Kd 51.7 nM
1.0
0.8
0.6
0.4
0.2
0.0
Fractional Occupancy
10⁻¹
10⁰
10¹
10²
10³
Protein concentration (nM, log scale)
Data
Model (Kd=51.7 nM)

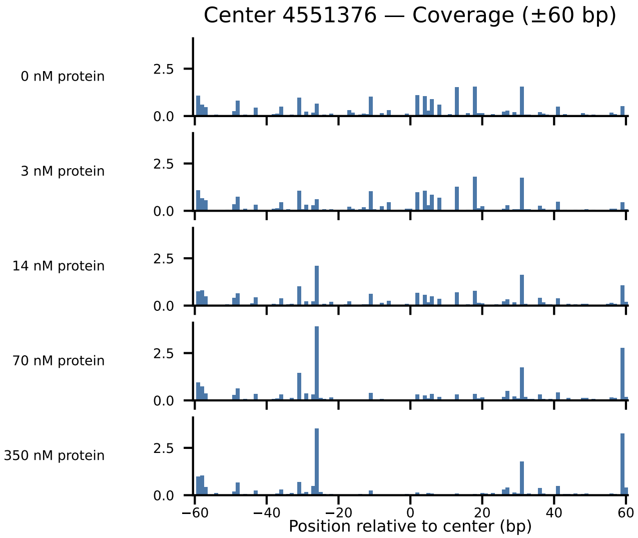

Center 4551376 — Coverage (±60 bp)
0 nM protein
3 nM protein
14 nM protein
70 nM protein
350 nM protein
2.5
0.0
−60
−40
−20
0
20
40
60
Position relative to center (bp)

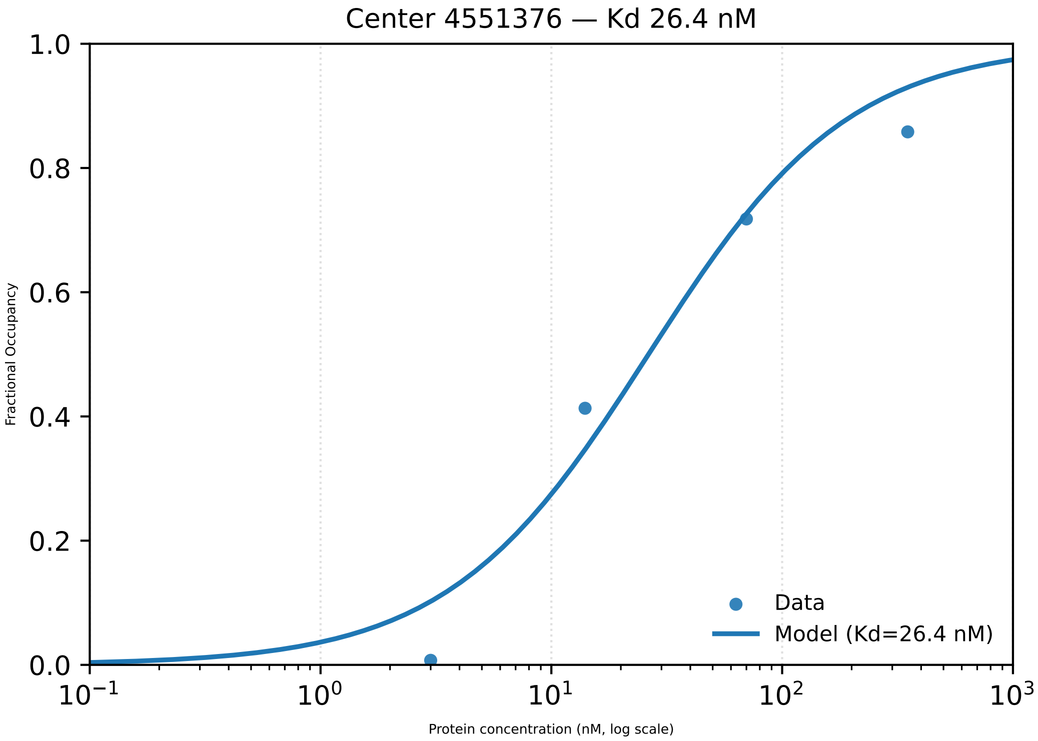

Center 4551376 — Kd 26.4 nM
Fractional Occupancy
1.0
0.8
0.6
0.4
0.2
0.0
$10^{-1}$
$10^{0}$
$10^{1}$
$10^{2}$
$10^{3}$
Protein concentration (nM, log scale)
Data
Model (Kd=26.4 nM)

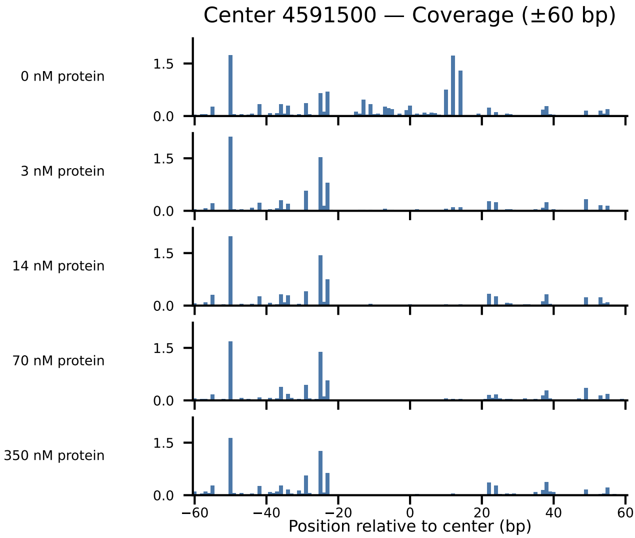

Center 4591500 — Coverage (±60 bp)
0 nM protein
3 nM protein
14 nM protein
70 nM protein
350 nM protein
1.5
0.0
−60
−40
−20
0
20
40
60
Position relative to center (bp)

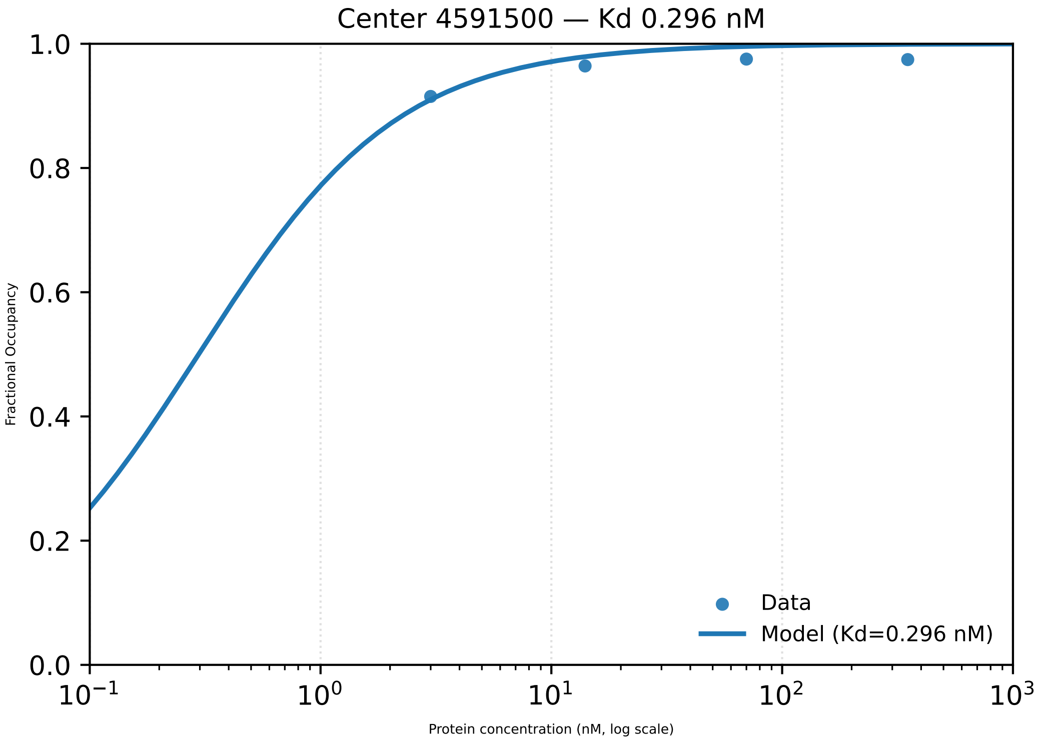

Center 4591500 — Kd 0.296 nM
1.0
0.8
0.6
0.4
0.2
0.0
Fractional Occupancy
$10^{-1}$
$10^{0}$
$10^{1}$
$10^{2}$
$10^{3}$
Protein concentration (nM, log scale)
Data
Model (Kd=0.296 nM)

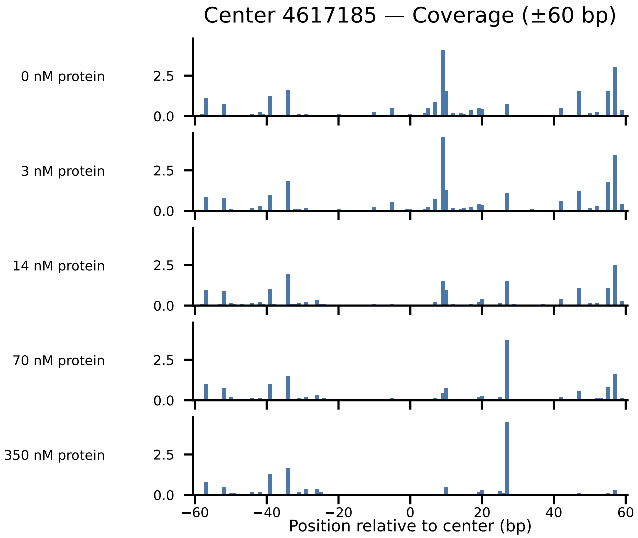

Center 4617185 — Coverage (±60 bp)
0 nM protein
3 nM protein
14 nM protein
70 nM protein
350 nM protein
2.5
0.0
−60
−40
−20
0
20
40
60
Position relative to center (bp)

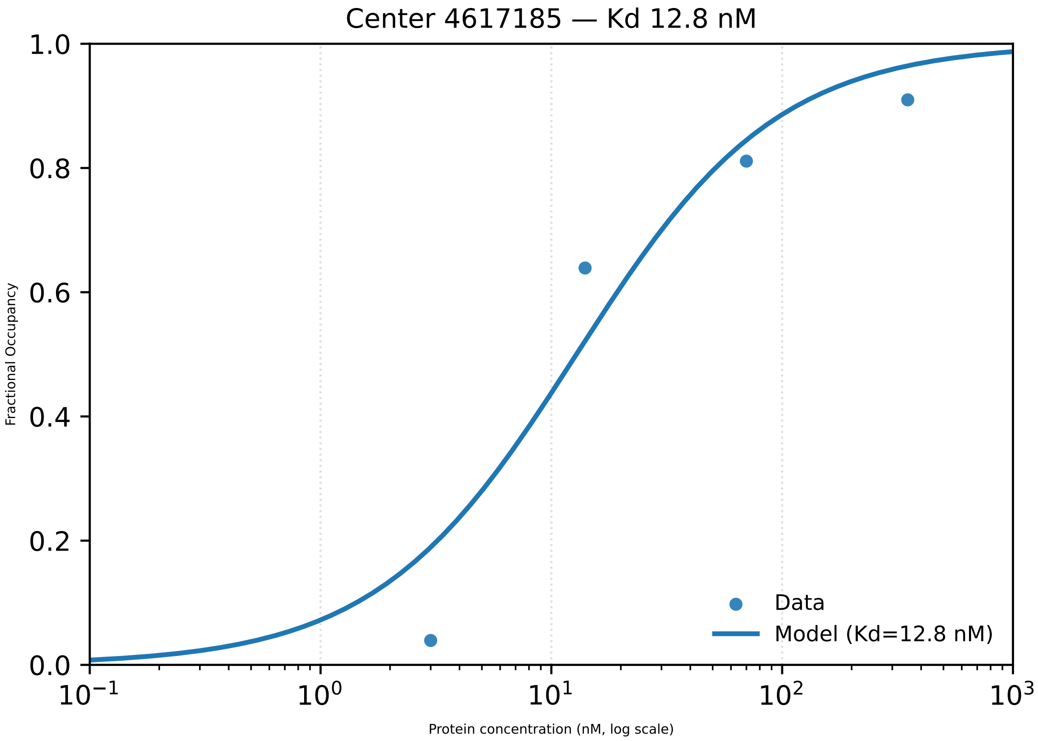
Center 4617185 — Kd 12.8 nM
1.0
0.8
0.6
0.4
0.2
0.0
Fractional Occupancy
$10^{-1}$
$10^{0}$
$10^{1}$
$10^{2}$
$10^{3}$
Protein concentration (nM, log scale)
Data
Model (Kd=12.8 nM)

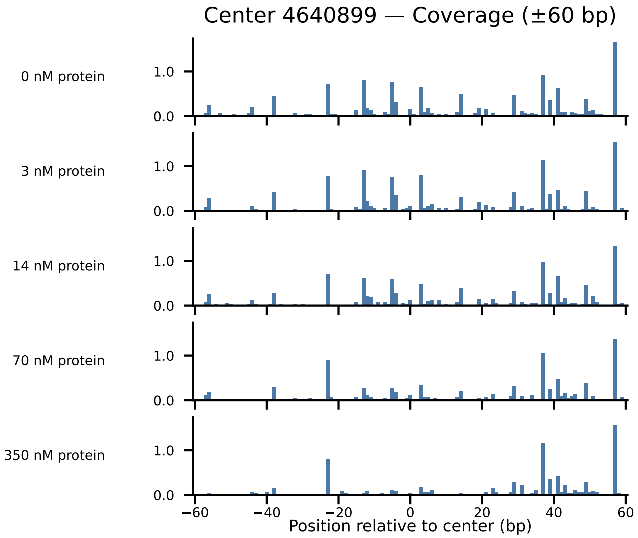

Center 4640899 — Coverage (±60 bp)
0 nM protein
3 nM protein
14 nM protein
70 nM protein
350 nM protein
1.0
0.0
−60
−40
−20
0
20
40
60
Position relative to center (bp)

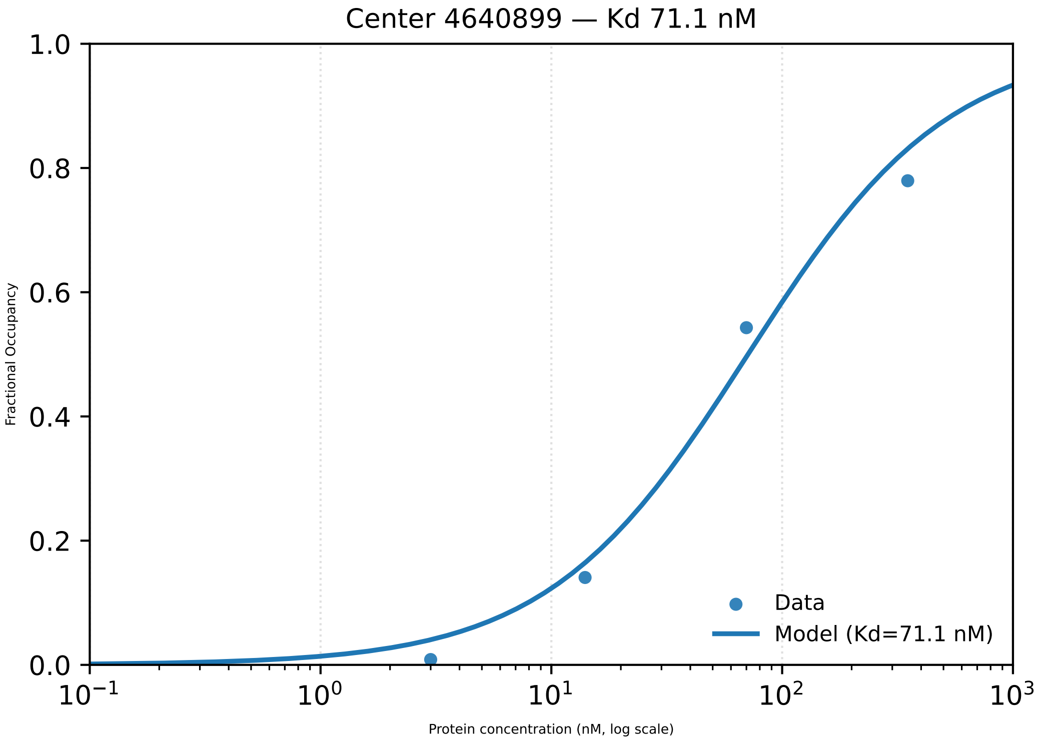
Center 4640899 — Kd 71.1 nM
Fractional Occupancy
Protein concentration (nM, log scale)
Data
Model (Kd=71.1 nM)
1.0
0.8
0.6
0.4
0.2
0.0
$10^{-1}$
$10^{0}$
$10^{1}$
$10^{2}$
$10^{3}$
